# Host association and intracellularity evolved multiple times independently in the *Rickettsiales*

**DOI:** 10.1101/2022.10.13.511287

**Authors:** Michele Castelli, Tiago Nardi, Leandro Gammuto, Greta Bellinzona, Elena Sabaneyeva, Alexey Potekhin, Valentina Serra, Giulio Petroni, Davide Sassera

## Abstract

The order Rickettsiales (*Alphaproteobacteria*) encompasses multiple diverse lineages of host-associated bacteria, including pathogens, reproductive manipulators, and mutualists. In order to understand how intracellularity and host association originated in this order, and whether they are ancestral or convergently evolved characteristics, we built an unprecedentedly large and phylogenetically-balanced dataset that includes *de novo* sequenced genomes and an accurate selection of published genomic and metagenomic assemblies. We performed detailed functional reconstructions that clearly indicated “late” and parallel evolution of obligate host-association and intracellularity in different *Rickettsiales* lineages. According to the depicted scenario, multiple independent series of horizontal acquisitions of transporters led to the progressive loss of biosynthesis of nucleotides, amino acids and other metabolites, producing distinct conditions of host-dependence. Coherently, each clade experienced a different pattern of evolution of the ancestral arsenal of interaction apparatuses, including development of specialised effectors involved in the lineage-specific mechanisms of host cell adhesion/invasion and intracellularity.

## Introduction

*Rickettsiales* are an early diverging^1^ and ancient^2^ alphaproteobacterial order. All the experimentally characterised members of this group engage in obligate associations with eukaryotic host cells^3^. The most long-term and thoroughly studied *Rickettsiales* include vector-borne pathogens, e.g. *Rickettsia* and *Anaplasma*, causing various diseases in humans and vertebrates^4–6^, as well as *Wolbachia*, that can establish complex interactions with arthropod and nematode hosts^7^, chiefly reproductive manipulation and mutualism.

In recent years, our knowledge and understanding of the origin, evolution and diversification of *Rickettsiales* have been remarkably improved. We can identify in particular three main advances. The first is the finding of a plethora of novel lineages (over 30 total genera described, grouped into seven families^8–11^), living in association with a wide variety of hosts^12–14^. Most of those hosts are diverse aquatic unicellular eukaryotes (e.g. ciliates, amoebae, algae)^1,15–21^, which have been deemed as probable ancestral hosts^2,22,23^, even though these associations are still poorly investigated.

Second, “*Candidatus* Deianiraea vastatrix” (from now on, *Candidatus* will be omitted from taxonomic names, e.g. *Deianiraea vastatrix*), a fully extracellular *Rickettsiales* bacterium equipped with an unexpectedly large biosynthetic repertoire for amino acids, was discovered^8^, opening a new perspective on the evolution of *Rickettsiales*. While previous views implied that obligate intracellular association dated back to the last common ancestor of the order (“intracellularity early” hypothesis), this discovery opened a novel alternative scenario. Accordingly, obligate intracellularity could have evolved later and multiple times independently in different sublineages (“intracellularity late” hypothesis), together with a stronger dependence on host cells.

Third, metagenome binning recently allowed the discovery of further *Rickettsiales* sublineages, in particular two early-diverging families^10^. Their genetic repertoire (including nutrient uptake, detoxification, and multiple biosynthetic pathways) led the authors to hypothesise that these bacteria could be free-living, implying that adaptation to the interaction with host cells would have occurred in more “derived” *Rickettsiales* lineages.

However, many open points still exist on the origin and evolution of the interaction between *Rickettsiales* and their hosts, in particular for what concerns the process(es) of transition between facultative/obligate association and between extracellular/intracellular condition, and whether those were “early” or “late” phenomena. In order to address such salient questions, in this study we collected an unprecedentedly large and representative genomic dataset of *Rickettsiales*, thanks to *de novo* sequencing and selection of metagenomic sequences, thus identifying three novel families and remarkably extending the available diversity within previously known lineages. This allowed us to get a view on the diversity and evolution of host adaptation among *Rickettsiales*, finding multiple and convincing lines of evidence supporting the intracellularity late hypothesis.

## Results

### Novel genomes

In this work, we obtained nine complete genome sequences of *Rickettsiales* bacteria (belonging to nine species, eight genera, two families). These represent the first sequences for the respective species, with the exception of *Megaira polyxenophila*^*13*^. Thus, the evolutionary representativity of available *Rickettsiales* genomes results significantly improved, also considering that all the newly sequenced organisms are hosted by ciliates or other protists. All the assemblies were highly curated, resulting in most cases in a very high quality (or even full closed, with five genomes having L50=1, and one L50=2) (Supplementary table 1). In four cases, the quality of the assembly allowed to clearly determine the presence of plasmids (respectively in two *Rickettsiaceae* and one among *Midichloriaceae*, Supplementary table 1).

### Phylogeny

For the successive analyses, we aimed to capture and analyse the widest available diversity of *Rickettsiales* from published sequences, including under-explored (e.g. *Deianiraeaceae*) and possibly yet uncharacterised lineages. To do so, together with a representative set of 36 available *Rickettsiales* genomes, we selected a total of “high-quality” 314 potential *Rickettsiales* metagenome-assembled-genomes (MAGs) from various sources. Their affiliation to *Rickettsiales* was tested by a multi-step phylogeny-based approach, taking advantage of multiple organismal selections tailored on each different MAG lineage to be investigated, as well as of a site-selection approach to counterbalance compositional heterogeneity^1^. After filtering out phylogenetically redundant MAGs, we identified 68 *Rickettsiales* MAGs, ending up with 113 total *Rickettsiales* for the final analysis, including the nine novel high-quality genomes.

The final *Rickettsiales* topology resulted quite robust, with most nodes finding full support, and only eight nodes below the bootstrap threshold of 90% (Fig. 1; Supplementary figure 1). The four families that include at least one characterised organism were highly supported, and mostly with a much increased representativity, thanks to novel genomes sequenced and MAGs identified in this study (novel taxa: 14/29 *Rickettsiaceae*; 10/17 *Midichloriaceae*; 1/14 *Anaplasmataceae*; 5/6 *Deianiraeaceae*). The inner relationships within *Rickettsiaceae* and *Anaplasmataceae* are overall consistent with previous phylogenetic and phylogenomic studies^8,10,12,15–17^. For *Midichloriaceae*, it was possible to infer a novel inner topology with a much higher support with respect to previous studies^14,15,20,21^. Furthermore, it was possible to determine that the *Midichloriaceae* bacterium associated with *Plagiopyla* represents a novel genus and species (Fig 1; from now on, *Vederia obscura*, see Supplementary text 1 for taxonomic description).

**Figure 1.**
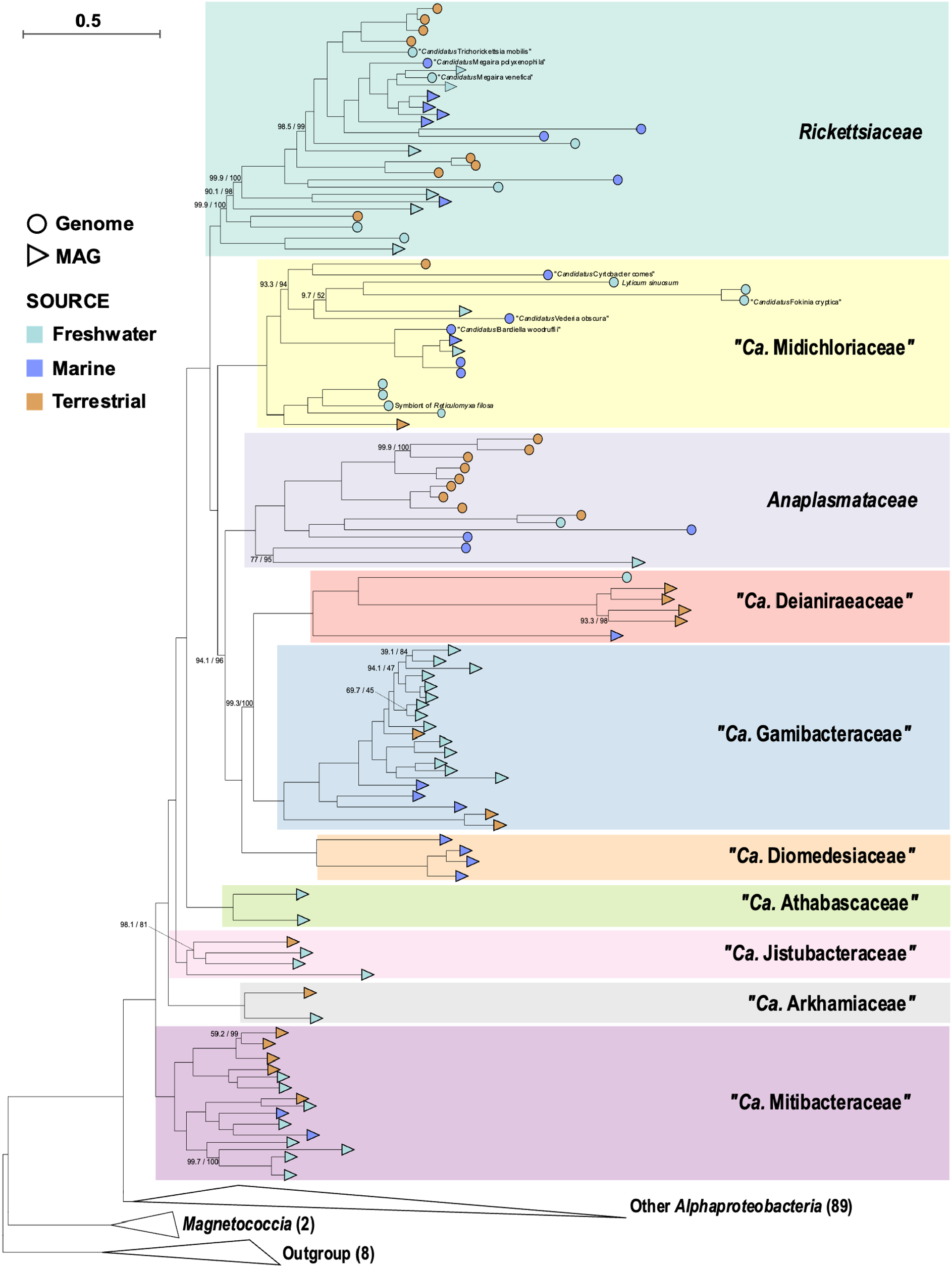
Maximum likelihood phylogenomic tree of 113 *Rickettsiales* inferred on 179 concatenated orthologs. Each *Rickettsiales* family is highlighted by a differently coloured box. At tips, round shapes indicate genome assemblies, while triangular shapes indicate metagenome-assembled-genomes (MAGs). Shape fillings show the sample source, namely light blue for freshwater, dark blue for marine, and orange for terrestrial. Due to space constraints, only the names of the nine newly obtained genome assemblies were reported, and non-*Rickettsiales* lineages (including other *Alphaproteobacteria, Magnetococcia*, and outgroup) are represented by collapsed triangular shapes, with the respective number of organisms reported (full tree is shown in Supplementary figure 1). On each branch, support values by SH-aLRT with 1000 replicates and by 1000 ultrafast bootstraps are reported (full support values were omitted). The tree scale stands for estimated sequence divergence. *Ca*. is an abbreviation for *Candidatus*.

The three recently described families composed only by MAGs ^10^, namely *Gamibacteraceae, Athabascaceae*, and *Mitibacteraceae*, were retrieved with a comparatively higher representativity (Fig. 1). Besides, three further MAG-only families were identified for the first time in this study: i) *Diomedesiaceae*, ii) *Jistubacteraceae*, and iii) *Arkhamiaceae*, the first being sister group of *Deianiraeaceae*+*Gamibacteraceae* (we will name these three families together as the “DDG”-clade), while the latter two forming a sequential branching pattern between *Athabascaceae* and *Mitibacteraceae* (Fig. 1; Supplementary figure 1, see Supplementary text 1 for taxonomic descriptions).

The phyletic relationships among the families are in general consistent with the most recent and comprehensive studies^8,10,19^, with a single partial exception. The present study indicates a closer relationship between the DDG-clade and *Anaplasmataceae* with respect to *Midichloriaceae*, consistent with 16S rRNA gene phylogenies and some phylogenomic analyses^8,19^, while another study placed *Anaplasmataceae* and *Midichloriaceae* as sister groups^10^. It should be considered that the number of available members of the DDG-clade progressively and largely increased in successive studies (in particular, this is the first study considering *Diomedesiaceae*), with a potential significant influence on the results.

In terms of origin, the vast majority of samples derived from aquatic environments (both freshwater and marine), especially in deeper nodes of the tree (Fig 1). The same holds true within each family, with the significant exceptions of *Anaplasmataceae* and *Deianiraeaceae*, mostly derived from terrestrial environments. Many characterised hosts resulted to be protists (most abundant in all families, except for *Anaplasmataceae*, all hosted by metazoans). For all MAGs, including in particular deeply branching lineages, no conclusive information is available on potential hosts, while in few cases there is loose indication of association, e.g. as originating from rumen^24,25^.

### General genome comparisons

Based on phylogeny, we hereby define as “crown *Rickettsiales*”, the smallest monophylum that comprises all characterised organisms (i.e. the one including the six families *Rickettsiaceae, Midichloriaceae, Anaplasmataceae, Deianiraeaceae, Gamibacteraceae, Diomedesiaceae*), thus corresponding to the classical *Rickettsiales* as defined previously^10^. Conversely, the four other early diverging families will be defined as “basal *Rickettsiales*”.

Genome sizes are quite variable between and within *Rickettsiales* families (Supplementary table 2). Crown *Rickettsiales* genomes are mostly in the range 1-1.5 Mb, with some appreciable differences between families, in particular on average *Anaplasmataceae* (1.1 Mb) and *Deianiraeaeaceae* (1.0 Mb), are smaller than others (all averages ≥1.3Mb). Genomes from basal families are much larger (frequently >2 Mb, and on average ≥1.8 Mb, except *Arkhamiaceae*). It should be taken into account that the likely incompleteness of some MAGs (Supplementary table 3) may influence the average size estimates, especially in DDG-clade and basal families. GC content is in general inversely correlated with genome size (32-36% on average in classical families and 40-52% in basal ones, with a maximum of 61%, in *Mitibacteraceae*).

As expected, gene numbers are consistent with respective genome sizes (Supplementary table 2). Our analyses supported the notions from previous studies that *Rickettsiales* have globally experienced genome reduction trends (Supplementary figure 2), as a putative consequence of adaptation and specialisation to host-associated lifestyles^3,8,10,26,27^. In order to investigate on the origin and evolution of such interactions with host cells from a functional and metabolic perspective, we selected a number of relevant traits/functions, in particular biosynthesis and uptake of metabolic precursors (i.e. amino acids and nucleotides), secretion/adhesion/motility apparatuses and putative effector molecules. A detailed overview of gene content variation and evolution in *Rickettsiales*, with a special focus on general “family-level” trends, as well as on the single newly characterised genomes, is presented in (Supplementary text 3).

### Secretion, attachment, and motility

Secretion systems are among the components that may exert a central role in regulating interactions with host cells^28^, potentially enabling specific stages of the bacterial life cycle through the delivery of effectors. In *Rickettsiales*, the hallmark apparatus is type IV secretion system (Supplementary figure 3), representing a probable ancestral horizontal acquisition^29^, and a possible prerequisite for establishing interactions with host cells^10^. Only few genomes among *Rickettsiales* are devoid of this apparatus^17,30^, and few others display an incomplete gene set, possibly indicative of ongoing loss, in particular *Bandiella* and the *Midichloriaceae* symbiont of *Reticulomyxa* (Supplementary figure 3). Type VI secretion system is very rare, being found only in two *Rickettsiaceae* (Supplementary figure 3), in both cases encoded on plasmids, a probable indication of horizontal acquisition. In *Sneabacter* this system has possibly functionally replaced the type IV secretion system^17^, while in *Trichorickettsia* both systems coexist.

Among putative secreted effector proteins (Supplementary figure 4), the “repeat-bearing” ones (ankyrin, tetratricopeptide, leucine-rich or pentapeptide repeats) are overall abundant and enriched in crown families, with many lineage-specific patterns, but are also present in basal families. On the other side, RTX toxins^31,32^ are quite abundant in basal families, and uncommon in crown ones. Interestingly, proteins involved in the intracellular invasion of eukaryotes, such as hemolysins, patatin-like and other phospholipases^33^, are common in some crown families such as *Rickettsiaceae* and *Midichloriacae*, and not uncommon in basal families, while they are rare in DDG clade, especially *Deianiraeaceae* (Supplementary figure 4). Several other putative toxins/effectors, characterised in *Rickettsiales*^*34*^ and/or in other bacteria^13,35–41^, were found more rarely, showing patterns of presence/absence that appear to be quite lineage-specific, and without sharp differences between basal and crown *Rickettsiales*.

Flagellum might be important especially during horizontal transmission in *Rickettsiales*^*21,42*^. Flagellar genes are common in basal *Rickettsiales* (Supplementary figure 5), likely representing ancestral traits^42^. Conversely, they are absent in the DDG-clade, and very rare in *Anaplasmataceae* (found just in the aquatic *Echinorickettsia* and *Xenolissoclinum*). Within families *Rickettsiaceae* and *Midichloriaceae*, they are extremely rare in terrestrial representatives (the only cases being *Midichloria mitochondrii* and the *Rickettsiaceae* symbiont of *Amblyomma* Ac37b), but quite common in aquatic ones, in particular basal *Rickettsiaceae* and *Midichloriaceae* in general, thereby also confirming the few experimental observations of flagella^43–45^. The poor correlation of the presence of flagellar genes with *Rickettsiales* phylogeny (including differential cases within the same genus, such as *Megaira* and *Midichloria*^*30*^) would be indicative of multiple independent reduction/loss events (Supplementary figure 5). Similar considerations may hold for chemotaxis, which possibly works in conjunction with flagella for host targeting^46^, and is present only in basal *Rickettsiales* and in the basal members of the family *Rickettsiaceae* (Supplementary figure 5).

Type 4 pili may be involved in the adhesion/attachment to host cells in *Rickettsiales*^*8*^. Their components are present in basal *Rickettsiales*, in the DDG clade, and only rarely in the other crown families, especially in the respective early-divergent and/or aquatic representatives (Supplementary figure 6). Thus, this apparatus was likely ancestral, experiencing multiple independent losses in crown *Rickettsiales*.

Proteins homologous to the FhaBC two-partner secretion system, which were tentatively linked to the attachment and toxicity towards host cells in *Rickettsiales*^*8*^ and may also participate in bacterial competition^47^, were likely ancestral and lost multiple times among *Rickettsiales*, being present in the basal families, *Deianiraeaceae, Gamibacteraceae*, and very few representatives of the other crown families (Supplementary figure 6).

For what concerns proteins characterised to be involved in adherence/invasion of host cells^48,49,50^ or even immune evasion^51^ in *Rickettsiales*, they are present (almost) exclusively in subgroups of the respective families (Supplementary figure 6).

### Nucleotide and amino acid metabolism

Nucleotide biosynthesis is stably present (both purines and pyrimidines) in basal *Rickettsiales* (Fig 2; Supplementary figure 7), while, differently from other biosynthetic pathways (Supplementary text 3), its presence in crown *Rickettsiales* is “scattered” along the organismal phylogeny. Indeed, it is ubiquitous in the families *Gamibacteraeae, Diomedesiaceae, Anaplasmataceae*, present in the earliest diverging *Deianiraeaceae* MAG and in few basal *Rickettsiaceae* (in particular the endosymbiont of *Amblyomma* Ac37b, able to synthesise both purines and pyrimidines), but absent in *Midichloriaceae* and in the remaining and most numerous *Rickettsiaceae* and *Deianiraeaceae*. Phylogenetic analyses of these pathways indicate that they are likely ancestral in *Rickettsiales*, with quite good correspondence with organismal phylogeny at various levels (up to and even above the families) (Supplementary figure 8).

**Figure 2.**
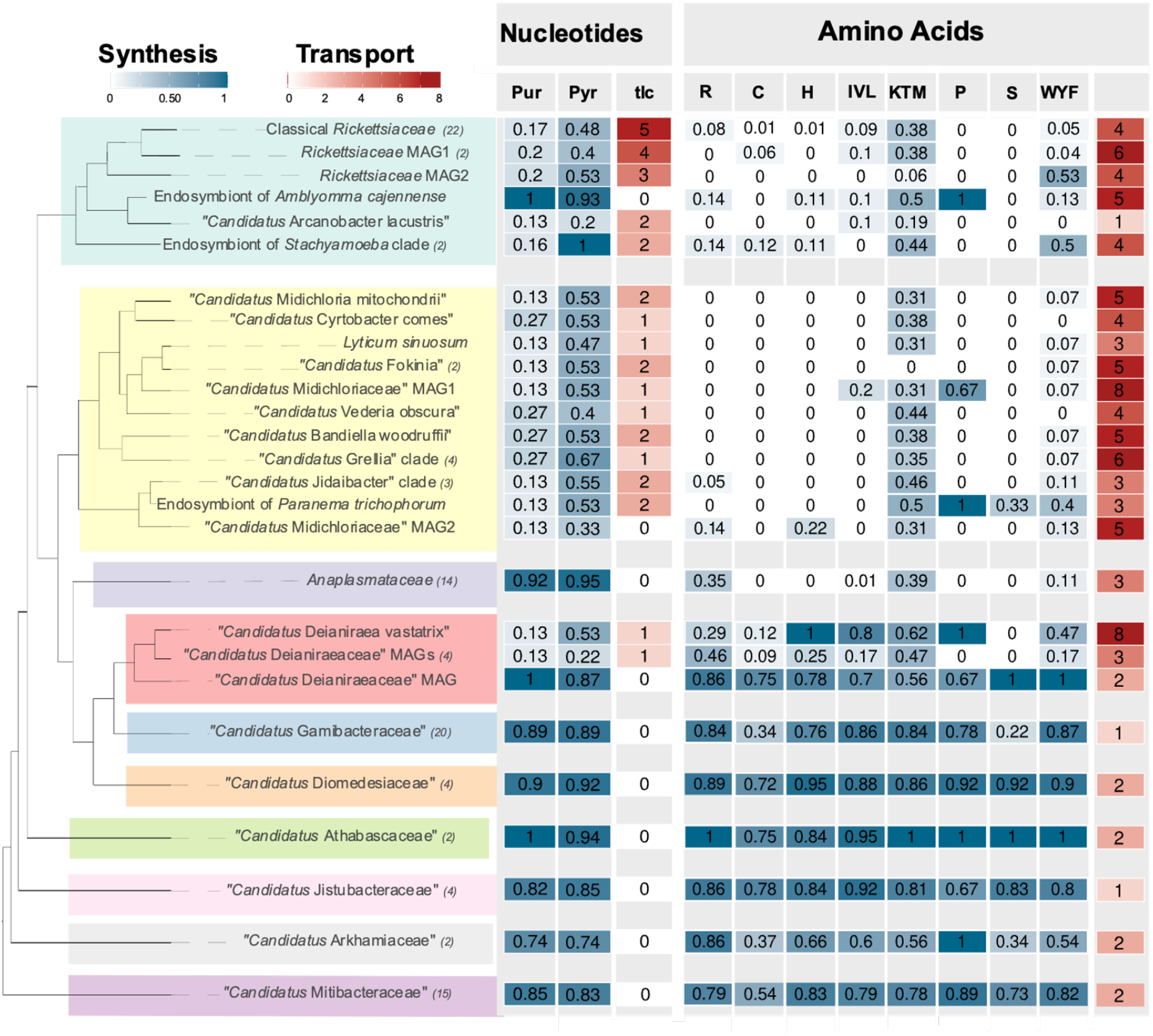
Heat-map showing the presence and abundance of biosynthetic pathways (blue) of nucleotides (purines and pyrimidines) and amino acids (grouped by to their mutually shared enzymatic steps according to BioCyc^94^), as well as their respective transporters (red). For biosynthesis, the proportion of the total genes of the pathway is shown (Supplementary table 6), while for transporters, the number of genes is reported, in particular for amino acid transporters the sum of the “characterised hits” (Supplementary figure 10). A cladogram of the organisms is shown on the left, with each *Rickettsiales* family highlighted by a differently coloured box. Due to space constraints, selected monophyletic clades with homogeneous gene content were collapsed. For each clade, the number of organisms is shown in brackets (if higher than one), and reported values are averaged, while the complete organism sets are shown in (Supplementary figure 7, 10).

In those crown *Rickettsiales* lacking nucleotide biosynthesis, this absence is counterbalanced by the ability to obtain final products (or intermediates) from their hosts. Indeed, the presence/absence pattern of tlc nucleotide translocases almost perfectly inversely correlates with nucleotide biosynthesis (Fig. 2; Supplementary figure 7). These family of transporters include chloroplastic ATP/ADP translocases, as well as a vast array of proteins, able to translocate several different nucleotides^52,53^, and previously reported to have experienced multiple horizontal gene transfer events between phylogenetically unrelated host-associated bacteria^54,55^.

Comparison of tlc and organismal phylogenies (Supplementary figure 9) indicates that these transporters were acquired multiple independent times by different *Rickettsiales* lineages (up to ten, once among *Deianiraeaceae*, three-five among *Midichloriaceae*, three-four among *Rickettsiaceae*, see Fig. 3). We also detected multiple independent events of duplication leading to several paralogs, namely five copies in classical *Rickettsiaceae*, two-four in three distinct basal *Rickettsiaceae* sublineages, and two copies in the *Jidaibacter* lineage among *Midichloriaceae* (Fig. 2,3; Supplementary figure 7, 8).

**Figure 3.**
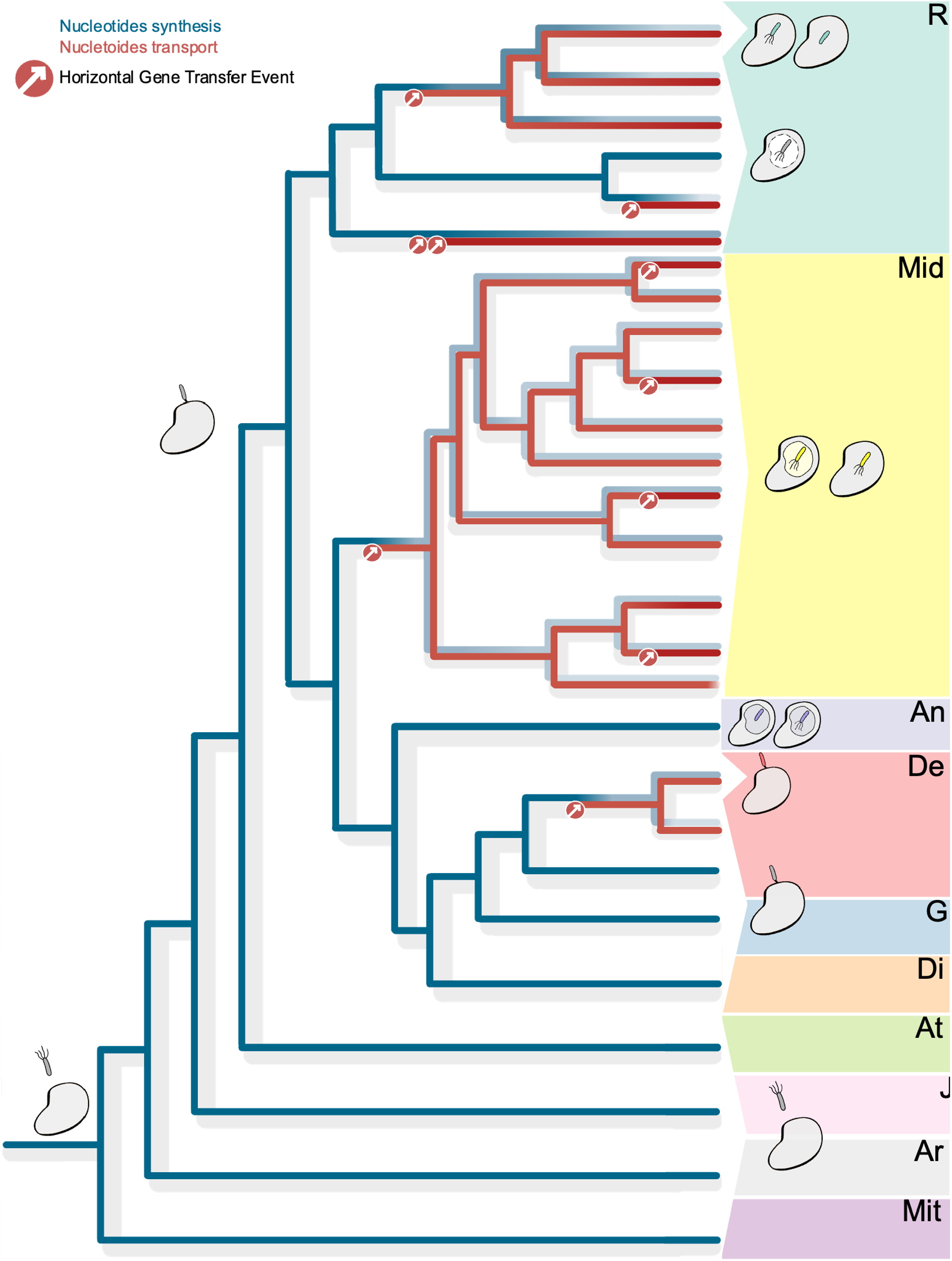
Reconstruction of the main steps of the evolution of *Rickettsiales* with a specific focus on the interactions with and dependence on eukaryotic cells. A cladogram of the main lineages (as represented in Fig. 2) is shown. The case of the biosynthetic pathways (averaged for purines and pyrimidines, blue) and tlc transporters (red) for nucleotides is represented along the tree by a heat-map-like representation, showing the inferred ancestral conditions and hypothesised steps of variation on each branch. In particular, multiple independent acquisitions of transporters by horizontal gene transfer events (red circles with inward arrows) would have led to the progressive reduction/loss of the biosynthesis. *Rickettsiales* families are highlighted by coloured boxes and by abbreviated names (R: *Rickettsiaceae*; Mid: *Midichloriaceae*; An: *Anaplasmataceae*; De: *Deianiraceae*; G: *Gamibacteraceae*; Di: *Diomedesiaceae*; At: *Athabascaceae*; J: *Jistubacteraceae*; Ar: *Arkhamiaceae*; Mit: *Mitibacteraceae*). At tips, groups of tips, and at selected nodes, the drawings represent the known (classical *Rickettsiaceae, Anaplasmataceae, Midichloriaceae*, and *Deianiraea*) or hypothesised (all other lineages) features of the bacteria and their interaction with eukaryotic hosts, in particular, intracellular or extracellular associations, or lack of association, as well as presence/absence of a vacuole and of flagella. Multiple side-by-side pictures represent alternative conditions/reconstructions for the same organisms.

The presence/absence pattern of biosynthetic pathways for amino acids shows significant analogies with what we detected for nucleotides (Fig 2; Supplementary figure 10). Indeed, they are quite uniformly present in basal *Rickettsiales, Diomedesiaceae, Gamibacteraceae*, and partly *Deianiraeceae*. Conversely, they are very rare or fully absent in the non-monophyletic assemblage of *Rickettsiaceae, Midichloriaceae* and *Anaplasmataceae*, with very few exceptions, such as arginine in *Ehrlichia* and *Neoehrlichia* (and partly *Anaplasma*), and proline in endosymbionts of *Amblyomma* Ac37b and *Peranema* (Supplementary figure 10). Our phylogeny indicates an overall vertical descendance of these pathways, with exceptions of possible horizontal transfer events with some *Rickettsiales* as recipients for genes belonging to the pathways of cysteine and histidine (Supplementary figure 8).

The array of putative amino acids transporters is more complex than that for nucleotides, considering the higher numerosity of amino acids and the multiple independent transporter families with variable substrate specificities^56,57^. This impairs precise homology-based prediction of substrate specificity in *Rickettsiales*, making it impossible to define with certainty with amino acid is imported by which transporter. Nevertheless, the total number of transporters is much higher in the three families that are more deprived in biosynthesis (Supplementary figure 10). This is especially true for those with a higher similarity with ascertained amino acid transporters (Fig. 2). Moreover, as seen for nucleotide transporters, the presence of homologs of different amino acid transporters is scattered along the *Rickettsiales* phylogeny (Supplementary figure 10).

We believe that the above presented data clearly indicate a pattern of multiple independent and successive acquisitions of nucleotide and amino acids transporters in different lineages of *Rickettsiales*, events that would have enabled the recipients to efficiently acquire those compounds from their hosts, thus leading to the reduction and eventual loss of the respective biosynthesis, an evolutionary scenario consistent with the intracellularity late hypothesis^8^.

## Discussion

The knowledge and understanding of the evolution of the typically host-associated *Rickettsiales*^*3,15*^, in particular its earlier steps, has been hampered by the limited and phylogenetically unbalanced set of genomes available. Here we present a dataset of over one hundred phylogenetically-diverse *Rickettsiales* assemblies, thanks to *de novo* sequencing of nine high-quality genomes from underexplored protist-associated representatives, and to an accurate selection of published genomes and MAGs. We thus obtained an unprecedented representativity of all known families, including those recently described^10^, and identified three further ones, for a total of ten families in *Rickettsiales*. Leveraging such an extended taxonomic resolution, we investigated whether the obligate association with eukaryotic hosts and specifically intracellular lifestyle were “early” conditions with a single origin, or “late” achievements that evolved multiple times independently in different *Rickettsiales* sublineages.

It is a generally accepted notion that metabolic dependence on the hosts is a key feature in obligate associations such as those involving *Rickettsiales*^*8,10,26,27*^, that evolved as the consequence of the possibility to efficiently acquire metabolites (including precursors and, for energy, ATP). Such ability is likely due to the acquisition of suitable transporters, making the respective biosynthetic and catabolic pathways dispensable, thus leading to their reduction and eventual loss. The pattern of gain of transporters and loss of synthesis pathways can thus be a strong indicator of the state of host dependence through evolution. Based on the herein produced dataset and analyses, the case of nucleotide synthesis and transport is noteworthy among *Rickettsiales*. Indeed, our analysis indicate that the most likely scenario is one of multiple independent horizontal acquisitions (up to ten) of tlc transporters among crown *Rickettsiales*, likely “triggering” independent losses of the ancestral biosynthetic pathways (Fig. 2, 3; Supplementary figure 7, 8, 9). Similar considerations hold for amino acids, even though the impossibility to predict the precise specificity of all transporters impairs a clear reconstruction of single events leading to the multiple independent losses of the ancestral biosynthetic pathways (Fig. 2, 3, Supplementary figure 8, 10).

Besides these more clear-cut cases, detailed analyses of the presence/absence patterns among genes involved in multiple other pathways strongly indicate analogous processes of gradual and independent reduction/losses in different crown *Rickettsiales* sublineages (Supplementary text 3). Sharp differences are present even within single families, such as in the metabolically rich basal *Rickettsiaceae*, as compared to the classical streamlined ones. Thus, in contrast with the more traditional views^10,21,22^, our analyses provide a clear indication that processes of pathway reduction/loss have not taken place just once in *Rickettsiales*, but instead occurred (and are still possibly occurring) multiple independent times in different crown *Rickettsiales* lineages, also in relation with the host features.

Secretion systems and attachment/invasion molecules are other paramount bacterial components for promoting and actively regulating interactions with host cells^28,29,58^. Interestingly, we found that in crown *Rickettsiales* the repertoire for such systems, as well as for flagellar apparatus, is a substantial subset of the one of basal *Rickettsiales*. This result fortifies previous notions that the ancestral *Rickettsiales* already possessed a quite rich set of proteins to interact with (unicellular) eukaryotes^10,29^. It seems reasonable to hypothesise that such arsenal could have represented a prerequisite for the establishment of associations with eukaryotic cells, possibly through a partial process of repurposing^10^ (e.g. delivery of effectors molecules active on eukaryotes, motility and chemotaxis involved in horizontal transmission). In this regard, type IV secretion is a good candidate^10,29^, also considering its almost full conservation among *Rickettsiales* (Supplementary figure 3). Other apparatuses, e.g. flagellum and type IV pilus, are more phylogenetically scattered among crown *Rickettsiales* (Supplementary figure 5, 6), likely as a result of multiple independent losses as well. This indicates unique patterns of specialisations along the *Rickettsiales* evolution, with the concurrent lineage-specific losses of traits that were dispensable for the interaction with respective host cells. Interestingly, the correlation with other traits allows the inference of some functional links, such as flagella and chemotaxis in aquatic environments, likely involved in non host-associated stages such as horizontal transmission^21^, pili for extracellular attachment to host cells in *Deianiraea*^*8*^ and possibly other members of the DDG clade. At the same time, alternative/additional functions for these apparatuses due to their homologies with secretion systems^59,60^ could be possible, and may account for the exceptions to such correlation patterns^30,42^.

Conversely, specialisation in the interaction with host cells has likely implied the expansion of other gene families, in particular those of putative secreted effectors such as the “repeat-containing” ones or acquisition/development of novel ones. In particular, several proteins characterised in pathogenic *Rickettsiales* (e.g. *Rickettsia, Anaplasma*) for their direct involvement in the interaction with the host^34,48–51^, resulted to be lineage-specific (at the family level or even below) rather than conserved in *Rickettsiales* as a whole. Thus, it seems likely that many other still uncharacterised lineage-specific proteins could exist in the other much less investigated *Rickettsiales* (e.g. *Midichloriaceae*, DDG clade). Such a scenario of lineage-specific sets of “interactors” suggests that the mechanisms and conditions of host-association have evolved independently among different (crown) *Rickettsiales* lineages along with their molecular players.

It is more difficult to precisely infer the condition of basal *Rickettsiales* (lacking any experimental data) in terms of potential interaction with eukaryotes. These bacteria are metabolically rich, suggesting independence from possible host cells^10^. At the same time, they bear basically all the putative prerequisite apparatuses for the interaction, and, most significantly, many are also equipped with homologs of effectors typical of crown *Rickettsiales*, such as phospholipases^33^ and the “repeat-containing” effectors, and they even are selectively enriched of additional potential effectors^31,32^. Conversely, free-living-like traits such as inorganic nutrient transport and detoxification, previously found only among basal *Rickettsiales*^*10*^, were retrieved also in some crown representatives (Supplementary text 3). This may be seen as indicative that those crown *Rickettsiales* retain some “primitive” traits, and, more in general, as another hint at more complex evolutionary trajectories than a simple transition towards obligate association at the root of crown *Rickettsiales*.

Taken together, the above presented data clearly indicate that obligate host-association, as well as intracellularity, were most likely “late” conditions in *Rickettsiales*. Under such a scenario, we propose that at some point in the early *Rickettsiales* evolutionary history their presumably aquatic free-living ancestors were engaged in some kind of facultative interaction with eukaryotes. The starting point could have been defence from protist predators through the release of active effectors, as previously hypothesised^10^. It is possible to envision that such defence mechanisms successively paved the way for the (gradual) development of the (at least occasional) ability to gain further advantages by such interactions, such as the capability to acquire metabolites from the damaged/killed eukaryotes, somehow reversing and taking control of the predator-prey interactions. The lifestyle of *Deianiraea* could be reminiscent of this hypothetical condition^8^. Most likely, such transition towards facultative associations would have occurred prior to the last common ancestor of crown *Rickettsiales*, which are all obligatorily host-associated.

Conversely, it is not straightforward to precisely place an upper bound for such transition, given the complete lack of direct information on the lifestyles of extant basal *Rickettsiales*. It could have occurred sharply in the common ancestor of crown *Rickettsiales*, or could have been more nuanced, involving also the ancestors of some basal lineages, possibly up to the ancestor of all *Rickettsiales*. Nevertheless, it cannot be excluded that any potential host-associated representative of basal *Rickettsiales* could be the result of a convergent and independent evolution with respect to crown *Rickettsiales*.

In any case, for what concerns crown *Rickettsiales*, we can envision that a single initial facultative association would have differentiated in the descendants, becoming tighter and tighter (and at some point obligate) through parallel successive steps of acquisition/development of metabolite transporters and interactor molecules. Such further transitions would have occurred separately and independently in different *Rickettsiales* lineages, thus supporting the conclusion of a late origin for the obligate association with hosts. The order and kind of such steps, although somehow similar, would have not been unique in each of the different crown *Rickettsiales* phyletic lines, as reflected in the present-day lineages, which exhibit differential features of metabolic dependence (e.g. for most amino acids but not nucleotides in *Anaplasmataceae*, and vice versa in most *Deianiraeaceae*), as well as differential mechanisms and conditions of interaction (chiefly intracellularity vs extracellarity).

All the above would specifically support the intracellularity late hypothesis, as this trait would have evolved multiple independent times and with differential features in some crown *Rickettsiales* (at least the *Rickettsiaceae, Midichloriaceae*, and *Anaplasmataceae* ancestors, with the exclusion of *Deianiraea* and possibly the whole DDG clade). An alternative reconstruction which cannot be excluded is that the ancestor of crown *Rickettsiales* had already undergone an early transition towards obligate association through some yet unidentified step(s). Even in such a case, analysed data would still robustly support the reconstruction that most further genome reduction/adaptation steps would have occurred later and independently, so that in particular intracellularity would be anyway designated as a late condition.

Our reconstruction may also provide a novel perspective on the origin and evolution of other host-associated bacterial lineages that, similarly to *Rickettsiales*, present the prerogative to “hold the control” of the interaction and to switch hosts by horizontal transmission, and were thus termed “professional symbionts”^61^) (e.g. *Chlamydiae*^*62*^, *Legionellales*^*63*^ and *Holosporales*^*1*^). These lineages could share similarities in the initial establishment and successive stepwise and “late” evolutionary development.

From a more general perspective, it seems worthily a comparison of *Rickettsiales* (and possibly other “professional symbionts”) with the more “canonical” genome evolution models among obligate symbionts, namely nutritional mutualists in insects (with some parallels also in protists^61^). Such symbionts undergo relatively rapid streaminling as a result of an initial host restriction, followed by a more or less prolonged stasis, and are somehow “doomed” to extinction after replacement. Conversely “professional symbionts” would be the controllers of the interaction from its evolutionary beginning, retaining the ability to horizontally change hosts, and undergoing much more gradual and “flexible” streamlining processes, depending also on the external environmental conditions and not just on the features of a single host. Interestingly, the abundance of lineages preferentially associated with marine invertebrates and showing poor cocladogenenesis with their hosts^64^ indicates that there may be more bacteria sharing traits of professional symbionts than currently recognised.

Large-scale comparative genomic analyses such as those presented here and elsewhere^10,62,63^ have huge potential to provide major advances in our understanding of functional traits and the underlying evolutionary processes. However, they also face inherent predictive limits, being quite suitable for deriving metabolic dependencies, and less for the inference of more complex and possibly not-yet-documented traits, such as mechanisms of interaction and subcellular (or extracellular) localisation. For example, it could have been burdensome and highly speculative to infer the so far uniquely observed extracellular condition in *Deianiraea*^*8*^ (representing the first basis to propose the “intracellularity late” hypothesis) only from its genome. Therefore, considering that we still completely lack any experimental data for six out of ten *Rickettsiales* families (including all basal ones), we strongly invoke the need for further experimental investigations. As in other cases^65^, these may provide additional and otherwise unpredictable insights on the lifestyle of present-day organisms, and represent the basis for refining existing hypotheses and inferring novel ones on *Rickettsiales* evolution.

## Methods

### Sample preparation and sequencing

In this work, the nine novel *Rickettsiales* genome sequences were assembled (for a detailed account on sample preparation, sequencing, and genome assembly, see Supplementary text 3; Supplementary table 1, 4), starting from eight protist host samples. Six of those samples were characterised in previous studies^43–45,66–68^, while two others, namely the ciliates *Plagiopyla frontata* IBS-3 and *Euplotes woodruffi* NDG2, were newly isolated. Each sample was differentially processed. Briefly, for Illumina sequencing most samples were subjected to whole-genome amplification (WGA) with the REPLI-g Single Cell Kit (Qiagen), either directly from few ciliate host cells (four samples: *Paramecium biaurelia* US_Bl 11III1, *Paramecium nephridiatum* Sr 2-6, *E. woodruffi* NDG2, *P. frontata* IBS-3) or from a previously obtained DNA extract (one sample: *Euplotes harpa* BOD18^66^; Supplementary table 1). Two additional samples (*P. biaurelia* USBL-36I1 and *Paramecium multimicronucleatum* Kr154-4) were processed for bulk DNA extraction (over 200,000 ciliate host cells each), with CTAB and phenol-chloroform protocols, respectively. Each of those seven DNA samples was processed through a Nextera XT library and sequenced by Admera Health (South Plainfield, NJ, USA) on a Illumina HiSeq X machine, producing 2×150 bp paired-end reads. Read quality was assessed with FastQC^69^. NDG2 sample was also subjected to Nanopore sequencing. For this purpose, a bulk (∼ 300,000 *Euplotes* cells) extract with the NucleoSpin™ Tissue Kit (Macherey-Nagel™) was processed through a SQK-LSK109 ligation-sequencing library, and sequenced in a FLO-MIN106 flow cell. Basecalling was then performed with guppy 5.0.11. Then, reads were processed with Porechop 0.2.4^70^ with default options. Quality of the reads was assessed with NanoPlot 1.23.0^71^. The eighth sample consisted in a *Rickettsiales* bacterium associated with the foraminiferan *Reticulomyxa filosa*, already sequenced together with its host in a previous study^68^. Sequencing reads were kindly provided by the original authors.

### Genome assembly

For each sample, the total Illumina reads were assembled using SPAdes 3.6^72^ with default settings, obtaining a “preliminary assembly”. Then, a multi-step procedure was applied, in order to select only those contigs belonging to the symbiont of interest and discard those belonging to the host or to additional organisms present in the sample (e.g. residual food, additional associated bacteria), as described previously (e.g.^8^). For this purpose, the blobology pipeline was applied^73^, followed by extensive manual curation. Briefly, preliminary contigs were classified according to their length, GC% content, sequencing coverage, and taxonomy. Reads mapping^74^ on selected contigs were reassembled separately with SPAdes 3.6, or, for NDG2, with Unicycler^75^ in a hybrid assembly with the respective Nanopore reads. Two samples (*P. multimicronucleatum* 12 and *P. biaurelia* US_Bl 11III1) were subjected to genome finishing, performing PCR reactions with TaKaRa Ex Taq and reagents (Takara Bio, Japan). Successful results were confirmed by bidirectional Sanger sequencing performed by GATC Biotech (Germany).

### Annotation

The newly obtained genomes were all annotated with Prokka 1.10^76^, using the --rfam option. Afterwards, annotation of the genomes of ciliate symbionts was manually curated by a detailed inspection of blastp hits on NCBI nr and on *Rickettsiales* proteins as described previously^8^.

### Full *Rickettsiales* dataset construction and phylogenomic analyses

Phylogenomic analyses were aimed to collect a representative and comprehensive view on the evolution and diversity of *Rickettsiales*. All sequences were downloaded from NCBI GenBank via ftp (ftp.ncbi.nlm.nih.gov/genomes/all/GCA), and are updated to July 2021. We manually selected a representative set of 36 *Rickettsiales* genomes, including at least one representative per genus. For the phylogeny, other 89 representative non-*Rickettsiales Alphaproteobacteria*, as well as 8 *Gammaproteobacteria* and *Betaproteobacteria* as outgroup were employed, taking inspiration from the selection by Muñoz-Gómez and co-authors^1^. We then aimed to identify all MAGs (metagenome-assembled genomes) which could be assigned to known “core” *Rickettsiales* lineages as of July 2021 (i.e. the four families *Rickettsiaceae, Anaplasmataceae, Midichloriaceae, Deianiraeaceae*), or to any lineage forming a supported monophyletic group with *Rickettsiales* with the exclusion of other (alpha)proteobacterial orders. Identification and representative selection of *Rickettsiales* MAGs was performed by a multi-step procedure (detailed in Supplementary text 4). Briefly, all MAGs assigned to *Rickettsiales* by NCBI taxonomy, those assigned to deep-branching alphaproteobacterial lineages^77^, plus all additional monophyletic relatives from the gtdb tree^78^, were downloaded (394 total MAGs). MAGs were first filtered by assembly quality, retaining only 314 MAGs having ≥50% single-copy and <5% duplicated of 219 proteobacterial orthologs according to BUSCO 5.0.0^79^ (Supplementary table 3).

All phylogenomic analyses were performed on concatenated alignments of 179 orthogroups (Supplementary table 5), which were manually selected on purpose (i.e. presence, after manual polishing of paralogs and poorly aligned sequences, in at least 85% of *Rickettsiales* -MAGs excluded-, 85% *Alphaproteobacteria*, 50% outgroup) from the eggNOG orthogroups^80^ predicted with eggnog-mapper 2.0.6^81^. For each organismal dataset (see below), orthogroups were separately aligned^82^, polished^83^ and concatenated^84^. All phylogenies were performed with IQ-TREE^85^, with 1000 ultrafast bootstraps^86^ and 1000 SH-aLRT replicates, employing the LG+C60+F+R6 model unless specified. A first phylogeny was performed on the full organismal dataset for an initial classification of MAGs, employing ModelFinder^87^ for model selection. In order to avoid artefacts due to compositional heterogeneity in the dataset (in particular potential “erroneous” phylogenetic proximity of MAG lineages to “core” *Rickettsiales* due GC/AT biases)^1,77,88^, the approach by Muñoz-Gómez and co-authors^1^ was applied, thus removing 10%, 20%, 30%, 40% or 50% of most heterogeneous sites from the alignment, and performing a separate phylogeny on each trimmed alignment (Supplementary figure 11). Based on resulting monophyly and Average Amino acid Identity (AAI) >0.85, phylogenetically-redundant MAGs were discarded (Supplementary figure 12). Thirteen clades were identified, grouping all those MAGs which could not be directly assigned to “core” *Rickettsiales* or other orders. In order to minimise potential artefacts (e.g. due to long-branch attraction), the phylogenetic position of the MAGs belonging to each clade was tested separately, with respect to core *Rickettsiales* lineages and other *Alphaproteobacteria* (Supplementary figure 13, 14). Phylogenies were performed accounting for compositional heterogeneity as above, and only MAGs in a monophyletic relationship with *Rickettsiales* were retained. Therefore, 68 total *Rickettsiales* MAGs were selected for all the successive analyses, totalling 210 organisms in the final dataset (113 *Rickettsiales*). For the final dataset, we aimed to “balance” the well-ascertained artefacts in alphaproteobacterial phylogeny due to compositional heterogeneity (e.g.^1,77,88^) and the loss of a considerable amount of phylogenetic information^89^, due to the removal of the most-compositionally heterogeneous sites, and likely important for the reconstruction of the inner relationships within *Rickettsiales*. Therefore, we employed the full alignment, using a guide tree (-g option in IQ-TREE) with a minimal set of just two constrained bipartitions (namely those separating *Rickettsiales* from the AT-rich *Holosporales* and *Pelagibacterales* as per^1^), thus considering the non-compositionally biassed topology among the alphaproteobacterial orders, and leaving at the same time freely unconstrained tree search for what concerns the inner relationships within *Rickettsiales*.

### Creation of a set of orthogroups for gene content comparisons

In order to perform gene content comparisons and reconstruct variations along the inferred species tree, a set of orthogroups was obtained for the 210 total organisms in the final phylogeny, starting from the previously obtained eggNOG orthogroups (see Supplementary text 5). The eggNOG database is hierarchically organised by taxonomy, namely each lineage (at domain, phylum, class, order, family levels) has a dedicated set of orthogroups, linked to those of higher level taxa. We aimed to maximise the advantages of such a system (chiefly lineage-specific annotations), and at the same time minimise potential disadvantages, such as, in particular, potential lack of identification of orthologs due to assignment to eggNOG orthogroups belonging to distinct taxa, which may be especially relevant for organisms displaying high sequence evolutionary rates, such as the *Rickettsiales*^*8*^. For this purpose, we designed a “telescopic” approach, in order to merge genes into large meaningful orthogroups, while still keeping as much as possible lineage-specific refined annotation. Briefly (see Supplementary text 5 for details), we compared taxonomic paths of all the identified eggNOG orthogroups, and grouped together all the orthogroups sharing at least a partial taxonomic path, assigning each group to the orthogroup at lowest possible shared rank in the taxonomic path leading to *Rickettsiales* (root; *Bacteria*; *Proteobacteria*; *Alphaproteobacteria*; *Rickettsiales*). Applying such a “telescopic” approach, a total of 444,226 genes were assigned to 20,041 orthogroups, 4009 of which present in at least one member of *Rickettsiales*, and 2990 of those present in 4 or more organisms of the total dataset, and thus considered for the following analyses.

### Reconstruction of ancestral states of gene copy number

Reconstruction of ancestral states (in terms of gene copy number in each orthogroup) was performed by a tree-reconciliation approach, employing ALE 0.4^90^. Briefly, each of the 2990 orthogroups was aligned^82^, and trimmed^83^. Then, only the 2871 orthogroups ≥30 amino acids after trimming were kept, and, for each of those, a sample of gene trees was obtained running 10000 iterations with PhyloBayes 4.1^91^. Then, amalgamated likelihood estimation was performed with ALEobserve, run with 10% burn-in, followed by ALEml_undated^92^. For each node of the species tree, predicted events and predicted (or observed at tips) copy numbers were rounded using a 0.3 threshold^93^), and then summed considering the whole dataset, as well as separately for each eggNOG functional category.

### Phylogenetic analyses on biosynthetic pathways for amino acids and nucleotides

Reference biosynthetic pathways for amino acids and nucleotides were obtained from the Biocyc database^94^ (Supplementary table 6). The datasets for the respective phylogenies were extensively manually curated (see Supplementary text 5 for details). Briefly, for each gene the corresponding eggNOG orthogroup was identified by a blastp search, and its composition was refined (e.g. excluding paralogs and poorly aligned sequences) by inspection of the respective alignment and of the respective single-gene tree^85^. In order to get more phylogenetically informative datasets for more robust and reliable inferences, we concatenated together the sequences involved in the same pathway, as well as in pathways sharing common reactions. Under similar assumptions, single (or two) gene pathways were not employed in the analyses. We noticed some potential “false positives”, namely organisms (including *Rickettsiales*) displaying only few genes of a given pathway. Alternatively, the apparently missing steps might be “filled” by additional non-specific enzymes, as hypothesised for other pathways in *Rickettsiales*^*26*^. Anyway, due to the “unbalanced” availability of sites, those cases could potentially hamper the accuracy of the phylogenetic inference. Therefore, for each concatenated alignment we opted for two alternative strategies in parallel, namely phylogeny on “full organism dataset”, or on a “selected organism dataset”, keeping only those organisms displaying at least 50% of the included genes, or, alternatively, at least a significant proportion of selected sub-branches of the pathway (Supplementary table 6). Genes were aligned, trimmed and concatenated as described above, and each concatenated alignment was also processed as described above (“Full *Rickettsiales* dataset construction and phylogenomic analyses”) in order to account for compositional heterogeneity^1^. On each resulting alignment, phylogeny was inferred with IQ-TREE and the LG+C60+F+R6 model, as described above.

### Identification of amino acid transporters

All the proteins of the 113 *Rickettsiales* of our final dataset were blasted against the full TCDB database^56^. Then, for each *Rickettsiales* genome the number of proteins having a best significant hit (e-value threshold of 1e-5) on each selected entry were counted (see Supplementary text 5 for details on the selection and refinement of TCDB entries representing putative amino acids transporters and on the rationale for the blast search).

### Identification and phylogenetic analysis of the tlc nucleotide translocases

Analyses were focused on the tlc nucleotide translocase transporters (see Supplementary text 5 for details), common in *Rickettsiales* and in other host-associated lineages^55^. Briefly, the corresponding eggNOG orthogroup was identified by a blastp search. Then, it was joined together with a selection of the phylogenetic dataset of tlc translocases by Major and co-authors^55^, corresponding to the clade of sequences consisting only of the nucleotide transport protein domain. The sequences were then aligned and trimmed as described above, and phylogeny was inferred as described above, employing the LG+C60 model as in^55^.

### Identification of genes involved in the interaction with host cells

For getting information on the presence and multiplicity of genes involved in multiple features of the interaction of *Rickettsiales* with host cells, the VFDB core reference database was employed^95^ (see Supplementary text 5 for details). The VFDB is quite redundant for orthologs identified in different included pathogens. Thus, in order to make it suitable for analyses on our non-model bacteria, for each VFC (Virulence Factor Class) separately, orthologs were identified within the database with OrthoFinder 2.5.4^96^, and manually curated. Then, all proteins of our dataset of *Rickettsiales* were blasted on the full VFDB core database. For each *Rickettsiales* genome, proteins were counted as “assigned” to each curated orthogroup if displaying a significant (evalue 1e-5) best blastp hit on any sequence belonging to the orthogroup.

## Data availability

Sequences obtained in this project were deposited to NCBI under accession number PRJNA831616.

## Acknowledgements

This project was supported by the Human Frontier Science Program (HFSP) Young Investigator Program grant RGY-0075 to DS, by the Italian Ministry of Education, University and Research (MIUR): Dipartimenti di Eccellenza Programme (2018–2022) Department of Biology and Biotechnology ‘L. Spallanzani’ University of Pavia, and by the European Community’s H2020 Programme H2020-MSCA-RISE 2019 under grant agreement n° 872767 to GP. Genomic characterization of *Trichorickettsia* and *Megaira venefica* was performed partly with support of RSF 20-14-00220. The University of Pisa is acknowledged for providing visiting scholarships to ES and AP. The authors would like to thank Venkata Mahesh Nitla for support in culturing *Plagiopyla frontata* IBS-3, Sascha Krenek for DNA preparation of the *Paramecium biaurelia* strain US_Bl 11III1, Umberto Postiglione for assistance in Nanopore assembly, Laura Quattrini and Marco Fagioli for assistance in genome closing by PCR. Gernot Gloeckner and Marco Groth are gratefully acknowledged for sharing genome sequencing data on *Reticulomyxa filosa* and its associated *Rickettsiales* bacterium.

## References

1. Muñoz-Gómez, S. A. et al. An updated phylogeny of the Alphaproteobacteria reveals that the parasitic Rickettsiales and Holosporales have independent origins. eLife vol. 8 Preprint at https://doi.org/10.7554/elife.42535 (2019).

2. Wang, S. & Luo, H. Dating Alphaproteobacteria evolution with eukaryotic fossils. Nat. Commun. 12, 3324 (2021).

3. Salje, J. Cells within cells: Rickettsiales and the obligate intracellular bacterial lifestyle. Nat. Rev. Microbiol. 19, 375–390 (2021).

4. Walker, D. H. & Ismail, N. Emerging and re-emerging rickettsioses: endothelial cell infection and early disease events. Nat. Rev. Microbiol. 6, 375–386 (2008).

5. Rikihisa, Y. Anaplasma phagocytophilum and Ehrlichia chaffeensis: subversive manipulators of host cells. Nat. Rev. Microbiol. 8, 328–339 (2010).

6. Renvoisé, A., Merhej, V., Georgiades, K. & Raoult, D. Intracellular Rickettsiales: Insights into manipulators of eukaryotic cells. Trends Mol. Med. 17, 573–583 (2011).

7. Werren, J. H., Baldo, L. & Clark, M. E. Wolbachia: master manipulators of invertebrate biology. Nat. Rev. Microbiol. 6, 741–751 (2008).

8. Castelli, M. et al. Deianiraea, an extracellular bacterium associated with the ciliate Paramecium, suggests an alternative scenario for the evolution of Rickettsiales. ISME J. 13, 2280–2294 (2019).

9. Montagna, M. et al. ‘Candidatus Midichloriaceae’ fam. nov. (Rickettsiales), an ecologically widespread clade of intracellular alphaproteobacteria. Appl. Environ. Microbiol. 79, 3241–3248 (2013).

10. Schön, M. E., Martijn, J., Vosseberg, J., Köstlbacher, S. & Ettema, T. J. G. The evolutionary origin of host association in the Rickettsiales. Nat Microbiol (2022) doi:10.1038/s41564-022-01169-x.

11. Dumler, J. S., & Walker, D. H. Rickettsiales. Bergey’s Manual of Systematics of Archaea and Bacteria 1–1 Preprint at https://doi.org/10.1002/9781118960608.obm00074 (2015).

12. Carrier, T. J. et al. Microbiome reduction and endosymbiont gain from a switch in sea urchin life history. Proc. Natl. Acad. Sci. U. S. A. 118, (2021).

13. Davison, H. R. et al. Genomic diversity across the Rickettsia and ‘Candidatus Megaira’ genera and proposal of genus status for the Torix group. Nat. Commun. 13, 2630 (2022).

14. Gruber-Vodicka, H. R. et al. Two intracellular and cell type-specific bacterial symbionts in the placozoan Trichoplax H2. Nature Microbiology vol. 4 1465–1474 Preprint at https://doi.org/10.1038/s41564-019-0475-9 (2019).

15. Castelli, M., Sassera, D. & Petroni, G. Biodiversity of ‘Non-model’ Rickettsiales and Their Association with Aquatic Organisms. Rickettsiales 59–91 Preprint at https://doi.org/10.1007/978-3-319-46859-4_3 (2016).

16. Yurchenko, T. et al. A gene transfer event suggests a long-term partnership between eustigmatophyte algae and a novel lineage of endosymbiotic bacteria. ISME J. 12, 2163– 2175 (2018).

17. George, E. E. et al. Highly Reduced Genomes of Protist Endosymbionts Show Evolutionary Convergence. Curr. Biol. 30, 925–933.e3 (2020).

18. Hess, S. Description of Hyalodiscus flabellus sp. nov. (Vampyrellida, Rhizaria) and Identification of its Bacterial Endosymbiont, ‘Candidatus Megaira polyxenophila’ (Rickettsiales, Alphaproteobacteria). Protist 168, 109–133 (2017).

19. Castelli, M. et al. ‘Candidatus Sarmatiella mevalonica’ endosymbiont of the ciliate Paramecium provides insights on evolutionary plasticity among Rickettsiales. Environ. Microbiol. 23, 1684–1701 (2021).

20. Giannotti, D., Boscaro, V., Husnik, F., Vannini, C. & Keeling, P. J. The ‘Other’: an Overview of the Family ‘ Midichloriaceae’. Appl. Environ. Microbiol. 88, e0243221 (2022).

21. Schulz, F. et al. A Rickettsiales symbiont of amoebae with ancient features. Environ. Microbiol. 18, 2326–2342 (2016).

22. Weinert, L. A., Welch, J. J. & Jiggins, F. M. Conjugation genes are common throughout the genus Rickettsia and are transmitted horizontally. Proc. Biol. Sci. 276, 3619–3627 (2009).

23. Vannini, C., Petroni, G., Verni, F. & Rosati, G. A Bacterium Belonging to the Rickettsiaceae Family Inhabits the Cytoplasm of the Marine Ciliate Diophrys appendiculata (Ciliophora, Hypotrichia). Microbial Ecology vol. 49 434–442 Preprint at https://doi.org/10.1007/s00248-004-0055-1 (2005).

24. Xie, F. et al. An integrated gene catalog and over 10,000 metagenome-assembled genomes from the gastrointestinal microbiome of ruminants. Microbiome 9, 137 (2021).

25. Parks, D. H. et al. Recovery of nearly 8,000 metagenome-assembled genomes substantially expands the tree of life. Nat Microbiol 2, 1533–1542 (2017).

26. Driscoll, T. P. et al. Wholly ! Reconstructed Metabolic Profile of the Quintessential Bacterial Parasite of Eukaryotic Cells. MBio 8, (2017).

27. Min, C.-K. et al. Genome-based construction of the metabolic pathways of Orientia tsutsugamushi and comparative analysis within the Rickettsiales order. Comp. Funct. Genomics 623145 (2008).

28. Green, E. R. & Mecsas, J. Bacterial Secretion Systems: An Overview. Microbiol Spectr 4, (2016).

29. Gillespie, J. J. et al. Phylogenomics reveals a diverse Rickettsiales type IV secretion system. Infect. Immun. 78, 1809–1823 (2010).

30. Floriano, A. M. et al. The origin and evolution of mitochondrial tropism in Midichloria bacteria. bioRxiv (2022) doi:10.1101/2022.05.16.490919.

31. Linhartová, I. et al. RTX proteins: a highly diverse family secreted by a common mechanism. FEMS Microbiol. Rev. 34, 1076–1112 (2010).

32. Benz, R. Channel formation by RTX-toxins of pathogenic bacteria: Basis of their biological activity. Biochim. Biophys. Acta 1858, 526–537 (2016).

33. Borgo, G. M. et al. A patatin-like phospholipase mediates Rickettsia parkeri escape from host membranes. Nat. Commun. 13, 3656 (2022).

34. Niu, H., Kozjak-Pavlovic, V., Rudel, T. & Rikihisa, Y. Anaplasma phagocytophilum Ats-1 is imported into host cell mitochondria and interferes with apoptosis induction. PLoS Pathog. 6, e1000774 (2010).

35. Lobato-Márquez, D., Díaz-Orejas, R. & García-Del Portillo, F. Toxin-antitoxins and bacterial virulence. FEMS Microbiol. Rev. 40, 592–609 (2016).

36. Alix, E. et al. The capping domain in RalF regulates effector functions. PLoS Pathog. 8, e1003012 (2012).

37. Nwasike, C., Ewert, S., Jovanovic, S., Haider, S. & Mujtaba, S. SET domain-mediated lysine methylation in lower organisms regulates growth and transcription in hosts. Ann. N. Y. Acad. Sci. 1376, 18–28 (2016).

38. Billington, S. J., Jost, B. H. & Songer, J. G. Thiol-activated cytolysins: structure, function and role in pathogenesis. FEMS Microbiol. Lett. 182, 195–205 (2000).

39. Padmalayam, I., Karem, K., Baumstark, B. & Massung, R. The gene encoding the 17-kDa antigen of Bartonella henselae is located within a cluster of genes homologous to the virB virulence operon. DNA Cell Biol. 19, 377–382 (2000).

40. Swart, A. L., Gomez-Valero, L., Buchrieser, C. & Hilbi, H. Evolution and function of bacterial RCC1 repeat effectors. Cell. Microbiol. 22, e13246 (2020).

41. Veyron, S., Peyroche, G. & Cherfils, J. FIC proteins: from bacteria to humans and back again. Pathog. Dis. 76, (2018).

42. Sassera, D. et al. Phylogenomic evidence for the presence of a flagellum and cbb(3) oxidase in the free-living mitochondrial ancestor. Mol. Biol. Evol. 28, 3285–3296 (2011).

43. Boscaro, V. et al. Rediscovering the genus Lyticum, multiflagellated symbionts of the order Rickettsiales. Sci. Rep. 3, 3305 (2013).

44. Lanzoni, O. et al. Diversity and environmental distribution of the cosmopolitan endosymbiont ‘Candidatus Megaira’. Sci. Rep. 9, 1179 (2019).

45. Mironov, T. & Sabaneyeva, E. A Robust Symbiotic Relationship Between the Ciliate and the Bacterium. Trichorickettsia Mobilis. Front. Microbiol. 11, 603335 (2020).

46. Keegstra, J. M., Carrara, F. & Stocker, R. Publisher Correction: The ecological roles of bacterial chemotaxis. Nat. Rev. Microbiol. 20, 505 (2022).

47. Guérin, J., Bigot, S., Schneider, R., Buchanan, S. K. & Jacob-Dubuisson, F. Two-Partner Secretion: Combining Efficiency and Simplicity in the Secretion of Large Proteins for Bacteria-Host and Bacteria-Bacteria Interactions. Front. Cell. Infect. Microbiol. 7, 148 (2017).

48. Sears, K. T. et al. Surface proteome analysis and characterization of surface cell antigen (Sca) or autotransporter family of Rickettsia typhi. PLoS Pathog. 8, e1002856 (2012).

49. Kahlon, A. et al. Anaplasma phagocytophilum Asp14 is an invasin that interacts with mammalian host cells via its C terminus to facilitate infection. Infect. Immun. 81, 65–79 (2013).

50. Seidman, D. et al. Anaplasma phagocytophilum surface protein AipA mediates invasion of mammalian host cells. Cell. Microbiol. 16, 1133–1145 (2014).

51. Park, J., Kim, K. J., Grab, D. J. & Dumler, J. S. Anaplasma phagocytophilum major surface protein-2 (Msp2) forms multimeric complexes in the bacterial membrane. FEMS Microbiol. Lett. 227, 243–247 (2003).

52. Audia, J. P. & Winkler, H. H. Study of the five Rickettsia prowazekii proteins annotated as ATP/ADP translocases (Tlc): Only Tlc1 transports ATP/ADP, while Tlc4 and Tlc5 transport other ribonucleotides. J. Bacteriol. 188, 6261–6268 (2006).

53. Daugherty, R. M. et al. The nucleotide transporter of Caedibacter caryophilus exhibits an extended substrate spectrum compared to the analogous ATP/ADP translocase of Rickettsia prowazekii. J. Bacteriol. 186, 3262–3265 (2004).

54. Schmitz-Esser, S. et al. ATP/ADP translocases: a common feature of obligate intracellular amoebal symbionts related to Chlamydiae and Rickettsiae. J. Bacteriol. 186, 683–691 (2004).

55. Major, P., Embley, T. M. & Williams, T. A. Phylogenetic Diversity of NTT Nucleotide Transport Proteins in Free-Living and Parasitic Bacteria and Eukaryotes. Genome Biol. Evol. 9, 480–487 (2017).

56. Saier, M. H. et al. The Transporter Classification Database (TCDB): 2021 update. Nucleic Acids Res. 49, D461–D467 (2021).

57. Burkovski, A. & Krämer, R. Bacterial amino acid transport proteins: occurrence, functions, and significance for biotechnological applications. Appl. Microbiol. Biotechnol. 58, 265–274 (2002).

58. Gillespie, J. J. et al. Secretome of obligate intracellular Rickettsia. FEMS Microbiol. Rev. 39, 47–80 (2015).

59. Mattick, J. S. Type IV pili and twitching motility. Annu. Rev. Microbiol. 56, 289–314 (2002).

60. Abby, S. S. & Rocha, E. P. C. The non-flagellar type III secretion system evolved from the bacterial flagellum and diversified into host-cell adapted systems. PLoS Genet. 8, e1002983 (2012).

61. Husnik, F. et al. Bacterial and archaeal symbioses with protists. Current Biology vol. 31 R862–R877 Preprint at https://doi.org/10.1016/j.cub.2021.05.049 (2021).

62. Dharamshi, J. E. et al. Marine Sediments Illuminate Chlamydiae Diversity and Evolution. Curr. Biol. 30, 1032–1048.e7 (2020).

63. Hugoson, E., Guliaev, A., Ammunét, T. & Guy, L. Host Adaptation in Legionellales Is Ga, Coincident with Eukaryogenesis. Mol. Biol. Evol. 39, (2022).

64. Boscaro, V. et al. Microbiomes of microscopic marine invertebrates do not reveal signatures of phylosymbiosis. Nat. Microbiol. 7, 810–819 (2022).

65. Imachi, H. et al. Isolation of an archaeon at the prokaryote-eukaryote interface. Nature 577, 519–525 (2020).

66. Vannini, C. et al. ‘Candidatus anadelfobacter veles’ and ‘Candidatus cyrtobacter comes,’ two new rickettsiales species hosted by the protist ciliate Euplotes harpa (Ciliophora, Spirotrichea). Appl. Environ. Microbiol. 76, 4047–4054 (2010).

67. Szokoli, F. et al. Disentangling the Taxonomy of Rickettsiales and Description of Two Novel Symbionts (‘Candidatus Bealeia paramacronuclearis’ and ‘Candidatus Fokinia cryptica’) Sharing the Cytoplasm of the Ciliate Protist Paramecium biaurelia. Appl. Environ. Microbiol. 82, 7236–7247 (2016).

68. Glöckner, G. et al. The genome of the foraminiferan Reticulomyxa filosa. Curr. Biol. 24, 11–18 (2014).

69. Andrews, S. (2010). FastQC: A Quality Control Tool for High Throughput Sequence Data [https://www.bioinformatics.babraham.ac.uk/projects/fastqc/]

70. Wick, R. R., Judd, L. M., Gorrie, C. L. & Holt, K. E. Completing bacterial genome assemblies with multiplex MinION sequencing. Microb Genom 3, e000132 (2017).

71. De Coster, W., D’Hert, S., Schultz, D. T., Cruts, M. & Van Broeckhoven, C. NanoPack: visualizing and processing long-read sequencing data. Bioinformatics 34, 2666–2669 (2018).

72. Bankevich, A. et al. SPAdes: a new genome assembly algorithm and its applications to single-cell sequencing. J. Comput. Biol. 19, 455–477 (2012).

73. Kumar, S., Jones, M., Koutsovoulos, G., Clarke, M. & Blaxter, M. Blobology: exploring raw genome data for contaminants, symbionts and parasites using taxon-annotated GC-coverage plots. Front. Genet. 4, 237 (2013).

74. Langmead, B. & Salzberg, S. L. Fast gapped-read alignment with Bowtie 2. Nat. Methods 9, 357–359 (2012).

75. Wick, R. R., Judd, L. M., Gorrie, C. L. & Holt, K. E. Unicycler: Resolving bacterial genome assemblies from short and long sequencing reads. PLoS Comput. Biol. 13, e1005595 (2017).

76. Seemann, T. Prokka: rapid prokaryotic genome annotation. Bioinformatics 30, 2068– 2069 (2014).

77. Martijn, J., Vosseberg, J., Guy, L., Offre, P. & Ettema, T. J. G. Deep mitochondrial origin outside the sampled alphaproteobacteria. Nature vol. 557 101–105 Preprint at https://doi.org/10.1038/s41586-018-0059-5 (2018).

78. Parks, D. H. et al. GTDB: an ongoing census of bacterial and archaeal diversity through a phylogenetically consistent, rank normalized and complete genome-based taxonomy. Nucleic Acids Res. 50, D785–D794 (2022).

79. Simão, F. A., Waterhouse, R. M., Ioannidis, P., Kriventseva, E. V. & Zdobnov, E. M. BUSCO: assessing genome assembly and annotation completeness with single-copy orthologs. Bioinformatics 31, 3210–3212 (2015).

80. Huerta-Cepas, J. et al. eggNOG 5.0: a hierarchical, functionally and phylogenetically annotated orthology resource based on 5090 organisms and 2502 viruses. Nucleic Acids Research vol. 47 D309–D314 Preprint at https://doi.org/10.1093/nar/gky1085 (2019).

81. Cantalapiedra, C. P., Hernández-Plaza, A., Letunic, I., Bork, P. & Huerta-Cepas, J. eggNOG-mapper v2: Functional Annotation, Orthology Assignments, and Domain Prediction at the Metagenomic Scale. Mol. Biol. Evol. 38, 5825–5829 (2021).

82. Katoh, K. & Standley, D. M. MAFFT multiple sequence alignment software version 7: improvements in performance and usability. Mol. Biol. Evol. 30, 772–780 (2013).

83. Criscuolo, A. & Gribaldo, S. BMGE (Block Mapping and Gathering with Entropy): a new software for selection of phylogenetic informative regions from multiple sequence alignments. BMC Evolutionary Biology vol. 10 210 Preprint at https://doi.org/10.1186/1471-2148-10-210 (2010).

84. Borowiec, M. L. AMAS: a fast tool for alignment manipulation and computing of summary statistics. PeerJ 4, e1660 (2016).

85. Nguyen, L.-T., Schmidt, H. A., von Haeseler, A. & Minh, B. Q. IQ-TREE: a fast and effective stochastic algorithm for estimating maximum-likelihood phylogenies. Mol. Biol. Evol. 32, 268–274 (2015).

86. Minh, B. Q., Nguyen, M. A. T. & von Haeseler, A. Ultrafast approximation for phylogenetic bootstrap. Mol. Biol. Evol. 30, 1188–1195 (2013).

87. Kalyaanamoorthy, S., Minh, B. Q., Wong, T. K. F., von Haeseler, A. & Jermiin, L. S. ModelFinder: fast model selection for accurate phylogenetic estimates. Nature Methods vol. 14 587–589 Preprint at https://doi.org/10.1038/nmeth.4285 (2017).

88. Viklund, J., Ettema, T. J. G. & Andersson, S. G. E. Independent Genome Reduction and Phylogenetic Reclassification of the Oceanic SAR11 Clade. Molecular Biology and Evolution vol. 29 599–615 Preprint at https://doi.org/10.1093/molbev/msr203 (2012).

89. Fan, L. et al. Phylogenetic analyses with systematic taxon sampling show that mitochondria branch within Alphaproteobacteria. Nature Ecology & Evolution vol. 4 1213–1219 Preprint at https://doi.org/10.1038/s41559-020-1239-x (2020).

90. Szöllõsi, G. J., Rosikiewicz, W., Boussau, B., Tannier, E. & Daubin, V. Efficient exploration of the space of reconciled gene trees. Syst. Biol. 62, 901–912 (2013).

91. Lartillot, N. & Philippe, H. A Bayesian mixture model for across-site heterogeneities in the amino-acid replacement process. Mol. Biol. Evol. 21, 1095–1109 (2004).

92. Szöllősi, G. J., Davín, A. A., Tannier, E., Daubin, V. & Boussau, B. Genome-scale phylogenetic analysis finds extensive gene transfer among fungi. Philos. Trans. R. Soc. Lond. B Biol. Sci. 370, 20140335 (2015).

93. Martijn, J. et al. Hikarchaeia demonstrate an intermediate stage in the methanogen-to-halophile transition. Nature Communications vol. 11 Preprint at https://doi.org/10.1038/s41467-020-19200-2 (2020).

94. Karp, P. D. et al. The BioCyc collection of microbial genomes and metabolic pathways. Brief. Bioinform. 20, 1085–1093 (2019).

95. Liu, B., Zheng, D., Zhou, S., Chen, L. & Yang, J. VFDB 2022: a general classification scheme for bacterial virulence factors. Nucleic Acids Res. 50, D912–D917 (2022).

96. Emms, D. M. & Kelly, S. OrthoFinder: phylogenetic orthology inference for comparative genomics. Genome Biol. 20, 238 (2019).

