## Supplementary material for "Host association and intracellularity evolved multiple times independently in the *Rickettsiales*": figures_and_supplementary_materials: Fig_2.pdf

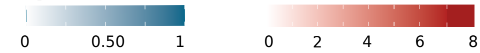


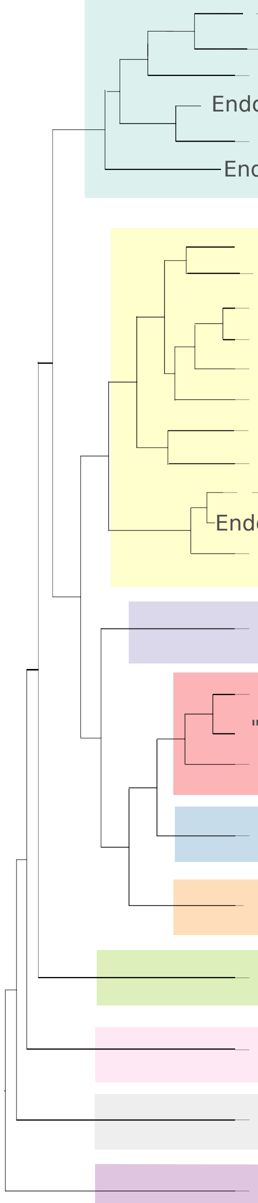

| | Nucleotides | | | Amino Acids | | | | | | | | |
|---|---|---|---|---|---|---|---|---|---|---|---|---|
| | Pur | Pyr | tlc | R | C | H | IVL | KTM | P | S | WYF | |
| Classical *Rickettsiaceae* (22) | 0.17 | 0.48 | 5 | 0.08 | 0.01 | 0.01 | 0.09 | 0.38 | 0 | 0 | 0.05 | 4 |
| *Rickettsiaceae* MAG1 (2) | 0.2 | 0.4 | 4 | 0 | 0.06 | 0 | 0.1 | 0.38 | 0 | 0 | 0.04 | 6 |
| *Rickettsiaceae* MAG2 | 0.2 | 0.53 | 3 | 0 | 0 | 0 | 0 | 0.06 | 0 | 0 | 0.53 | 4 |
| Endosymbiont of *Amblyomma cajennense* | 1 | 0.93 | 0 | 0.14 | 0 | 0.11 | 0.1 | 0.5 | 1 | 0 | 0.13 | 5 |
| "*Candidatus* Arcanobacter lacustris" | 0.13 | 0.2 | 2 | 0 | 0 | 0 | 0.1 | 0.19 | 0 | 0 | 0 | 1 |
| Endosymbiont of *Stachyamoeba* clade (2) | 0.16 | 1 | 2 | 0.14 | 0.12 | 0.11 | 0 | 0.44 | 0 | 0 | 0.5 | 4 |
| "*Candidatus* Midichloria mitochondrii" | 0.13 | 0.53 | 2 | 0 | 0 | 0 | 0 | 0.31 | 0 | 0 | 0.07 | 5 |
| "*Candidatus* Cyrtobacter comes" | 0.27 | 0.53 | 1 | 0 | 0 | 0 | 0 | 0.38 | 0 | 0 | 0 | 4 |
| *Lyticum sinuosum* | 0.13 | 0.47 | 1 | 0 | 0 | 0 | 0 | 0.31 | 0 | 0 | 0.07 | 3 |
| "*Candidatus* Fokinia" (2) | 0.13 | 0.53 | 2 | 0 | 0 | 0 | 0 | 0 | 0 | 0 | 0.07 | 5 |
| "*Candidatus* Midichloriaceae" MAG1 | 0.13 | 0.53 | 1 | 0 | 0 | 0 | 0.2 | 0.31 | 0.67 | 0 | 0.07 | 8 |
| "*Candidatus* Vederia obscura" | 0.27 | 0.4 | 1 | 0 | 0 | 0 | 0 | 0.44 | 0 | 0 | 0 | 4 |
| "*Candidatus* Bandiella woodruffii" | 0.27 | 0.53 | 2 | 0 | 0 | 0 | 0 | 0.38 | 0 | 0 | 0.07 | 5 |
| "*Candidatus* Grellia" clade (4) | 0.27 | 0.67 | 1 | 0 | 0 | 0 | 0 | 0.35 | 0 | 0 | 0.07 | 6 |
| "*Candidatus* Jidaibacter" clade (3) | 0.13 | 0.55 | 2 | 0.05 | 0 | 0 | 0 | 0.46 | 0 | 0 | 0.11 | 3 |
| Endosymbiont of *Paranema trichophorum* | 0.13 | 0.53 | 2 | 0 | 0 | 0 | 0 | 0.5 | 1 | 0.33 | 0.4 | 3 |
| "*Candidatus* Midichloriaceae" MAG2 | 0.13 | 0.33 | 0 | 0.14 | 0 | 0.22 | 0 | 0.31 | 0 | 0 | 0.13 | 5 |
| *Anaplasmataceae* (14) | 0.92 | 0.95 | 0 | 0.35 | 0 | 0 | 0.01 | 0.39 | 0 | 0 | 0.11 | 3 |
| "*Candidatus* Deianiraea vastatrix" | 0.13 | 0.53 | 1 | 0.29 | 0.12 | 1 | 0.8 | 0.62 | 1 | 0 | 0.47 | 8 |
| "*Candidatus* Deianiraeaceae" MAGs (4) | 0.13 | 0.22 | 1 | 0.46 | 0.09 | 0.25 | 0.17 | 0.47 | 0 | 0 | 0.17 | 3 |
| "*Candidatus* Deianiraeaceae" MAG | 1 | 0.87 | 0 | 0.86 | 0.75 | 0.78 | 0.7 | 0.56 | 0.67 | 1 | 1 | 2 |
| "*Candidatus* Gamibacteraceae" (20) | 0.89 | 0.89 | 0 | 0.84 | 0.34 | 0.76 | 0.86 | 0.84 | 0.78 | 0.22 | 0.87 | 1 |
| "*Candidatus* Diomedesiaceae" (4) | 0.9 | 0.92 | 0 | 0.89 | 0.72 | 0.95 | 0.88 | 0.86 | 0.92 | 0.92 | 0.9 | 2 |
| "*Candidatus* Athabascaceae" (2) | 1 | 0.94 | 0 | 1 | 0.75 | 0.84 | 0.95 | 1 | 1 | 1 | 1 | 2 |
| "*Candidatus* Jistubacteraceae" (4) | 0.82 | 0.85 | 0 | 0.86 | 0.78 | 0.84 | 0.92 | 0.81 | 0.67 | 0.83 | 0.8 | 1 |
| "*Candidatus* Arkhamiaceae" (2) | 0.74 | 0.74 | 0 | 0.86 | 0.37 | 0.66 | 0.6 | 0.56 | 1 | 0.34 | 0.54 | 2 |
| "*Candidatus* Mitibacteraceae" (15) | 0.85 | 0.83 | 0 | 0.79 | 0.54 | 0.83 | 0.79 | 0.78 | 0.89 | 0.73 | 0.82 | 2 |
