## Supplementary material for "Host association and intracellularity evolved multiple times independently in the *Rickettsiales*": figures_and_supplementary_materials: Legend_supplementary_materials.docx

**Legends of the supplementary materials**

**Supplementary figure 1.** Full maximum-likelihood phylogenomic tree of the final dataset. *Rickettsiales* families and other groups of organisms are highlighted on the right. On each branch, support values by SH-aLRT with 1000 replicates and by 1000 ultrafast bootstraps are reported (full support values were omitted). The tree scale stands for estimated sequence divergence.

**Supplementary figure 2.** Results of the result of the ancestral states of gene copy number obtained by ALE, including Duplication (D), transfer (T), origin (O), and loss (L) events. The summed results on each node and branch are presented for the total gene dataset, as well as for each functional category separately (after rounding up values the results of single orthologs).

**Supplementary figure 3.** Heat-map showing the numbers of orthologs identified for each component of type IV and type VI secretion systems in the *Rickettsiales* of the final dataset, ordered by their phylogenetic relationships (Supplementary figure 1), with families highlighted by different colours on the right. Empty grey boxes stand for no orthologs found.

**Supplementary figure 4.** Heat-map showing the numbers of orthologs identified for each putative effector protein in the *Rickettsiales* of the final dataset, ordered by their phylogenetic relationships (Supplementary figure 1), with families highlighted by different colours on the right. Empty grey boxes stand for no orthologs found.

**Supplementary figure 5.** Heat-map showing the numbers of orthologs identified for each component of flagellum and chemotaxis apparatuses in the *Rickettsiales* of the final dataset, ordered by their phylogenetic relationships (Supplementary figure 1), with families highlighted by different colours on the right. Empty grey boxes stand for no orthologs found.

**Supplementary figure 6.** Heat-map showing the numbers of orthologs identified for each component of multiple proteins possibly involved in the interaction with host cells in the *Rickettsiales* of the final dataset. Genes are ordered by functional groups (namely, components of type IV pilus/type II secretion, tight adherence pilus, fimbriae, curli, as well as exoproteins homologous to the FhaBC two-partner secretion system, previously characterised effectors in some *Rickettsiales* bacteria, and further other proteins). Organisms are ordered by their phylogenetic relationships (Supplementary figure 1), with families highlighted by different colours on the right. Empty grey boxes stand for no orthologs found.

**Supplementary figure 7.** Heat-map showing the presence and abundance of biosynthetic pathways of nucleotides (purines and pyrimidines; blue) and their tlc transporters (red) in *Rickettsiales*. For biosynthesis, the proportion of the total genes of the pathway is shown (Supplementary table 6), while for transporters, the number of genes is reported. Organisms are ordered by their phylogenetic relationships (Supplementary figure 1), with families highlighted by different colours on the right.

**Supplementary figure 8.** Phylogenetic trees of the concatenated genes of biosynthetic pathways for nucleotides and amino acids. For each pathway, the phylogenies for “all organisms” and “organism selection” sets are shown (Supplementary table 6), in both cases untrimmed and with the 10%, 20%, 30%, 40%, or 50% most compositionally heterogeneous sites removed. On each tree, the members of each *Rickettsiales* family are highlighted by different background colours. On each branch, support values by SH-aLRT with 1000 replicates and by 1000 ultrafast bootstraps are reported. The tree scales stand for estimated sequence divergences.

**Supplementary figure 9.** Phylogenetic tree of the tlc gene family. The members of each *Rickettsiales* family from the dataset of this study are highlighted by different background colours. On each branch, support values by SH-aLRT with 1000 replicates and by 1000 ultrafast bootstraps are reported. The tree scale stands for estimated sequence divergence.

**Supplementary figure 10.** Heat-map showing the presence and abundance of biosynthetic pathways of amino acids (grouped by to their mutually shared enzymatic steps according to BioCyc; blue) and their respective transporters (red) in *Rickettsiales*. Transporters are divided into “characterised” and “uncharacterised” hits, based on whether the corresponding TCDB entry have an experimentally characterised specificity for the given amino acids, or are representatives of a transporter family including amino acid transporters, but their own specificity was not ascertained. For biosynthesis, the proportion of the total genes of the pathway is shown (Supplementary table 6), while for transporters, the number of genes is reported. Organisms are ordered by their phylogenetic relationships (Supplementary figure 1), with families highlighted by different colours on the right.

**Supplementary figure 11.** Phylogenomic trees of *Rickettsiales* and other *Alphaproteobacteria*, for testing the affiliation to *Rickettsiales* of 211 “BUSCO+eggNOG-filtered putative *Rickettsiales* MAGs” (Step 4 in Supplementary text 3). The phylogeny on the untrimmed concatenated gene alignment, as well as on alignments with the 10%, 20%, 30%, 40%, or 50% most compositionally heterogeneous sites removed are presented. In each tree, the ascertained “core” *Rickettsiales* MAGs are highlighted in green, the ascertained non-*Rickettsiales* MAGs in gray and italics, and the members of each of 13 additional groups (labelled from A to M) in a different colour. On each branch, support values by SH-aLRT with 1000 replicates and by 1000 ultrafast bootstraps are reported. The tree scales stand for estimated sequence divergence.

**Supplementary figure 12.** AAI (Average Amino acid Identity) values within each examined clade of MAGs and *Rickettsiales* genomes.

**Supplementary figure 13.** Phylogenomic trees of *Rickettsiales* and other *Alphaproteobacteria*, for testing the affiliation to *Rickettsiales* of the members of each previously identified MAG groups (A-M; Step 5 in Supplementary text 3). For each clade, the phylogenies on the respective untrimmed concatenated gene alignment, as well as on alignments with the 10%, 20%, 30%, 40%, or 50% most compositionally heterogeneous sites removed, are reported in order. In each tree, the investigated MAGs are highlighted in blue. On each branch, support values by SH-aLRT with 1000 replicates and by 1000 ultrafast bootstraps are reported.

**Supplementary figure 14.** Phylogenomic trees of *Rickettsiales* and other *Alphaproteobacteria*, for testing the affiliation to *Rickettsiales* of 95 BUSCO-filtered “new putative *Rickettsiales* MAGs” (Step 6 in Supplementary text 3). The phylogenies on the untrimmed concatenated gene alignment, as well as on alignments with the 10%, 20%, 30%, 40%, or 50% most compositionally heterogeneous sites removed are reported in order. In each tree, the “new putative *Rickettsiales* MAGs” are highlighted in blue. On each branch, support values by SH-aLRT with 1000 replicates and by 1000 ultrafast bootstraps are reported.

**Supplementary table 1.** Summary of the sequencing (A) and assembly (B) procedures and results for the nine novel genomes obtained in this study.

**Supplementary table 2.** General features of the 113 analysed *Rickettsiales* assemblies, including family averages

**Supplementary table 3.** BUSCO scores of the selected published *Rickettsiales* genomes, the novel genomes obtained, and the two different sets of metagenome-assembled-genomes (MAGs) analysed. MAGs with <50% complete orthologs (yellow) or with ≥5% duplicated orthologs (red) were discarded.

**Supplementary table 4.** For each sample separately, list of the contigs of the preliminary assembly in which rRNA genes were identified with barrnap. For each contig, blobology parameters are reported, as well as the positions in which the rRNA gene was inferred, and the best blast hit of this gene sequence. Contigs are sorted by the respective sequencing coverage, and coloured according to the presumed organismal assignment (blue: host; shades of green: *Rickettsiales* symbionts; other colours: additional organisms).

**Supplementary table 5.** List of the 179 eggnog orthogroups employed for the phylogenomic analyses, and their presence/absence pattern in the organisms analysed in this study, including *Rickettsiales*, *Alphaproteobacteria*, outgroup, and, in a separate tab, MAGs (8 MAGs, highlighted in red, were discarded prior to phylogeny due to their identification as non-*Rickettsiales* based on the eggnog assignment).

**Supplementary table 6.** List of the genes analysed for the phylogeny of biosynthetic pathways of nucleotides and amino acids, subdivided by pathway. Each different set of genes taken into account for including organisms in the “selected organismal dataset” (sufficient condition, at least 50% genes in one set, see main text and Supplementary text 4) is highlighted in green on a separate column on the right.

**Supplementary text 1.** Taxonomic descriptions of novel species and families discovered in this study.

**Supplementary text 2.** Detailed description of the genome sequencing and assembly procedures.

**Supplementary text 3.** General comparative description of metabolic and functional features.

**Supplementary text 4.** Detailed description of the phylogenomic analyses.

**Supplementary text 5.** Detailed description of the gene content analyses.
