## Supplementary material for "Host association and intracellularity evolved multiple times independently in the *Rickettsiales*": figures_and_supplementary_materials: Supplementary_figure_2_ALE_results.pdf

### Sum of all COG categories

Total Copies ● 800 ● 1200 ● 1600

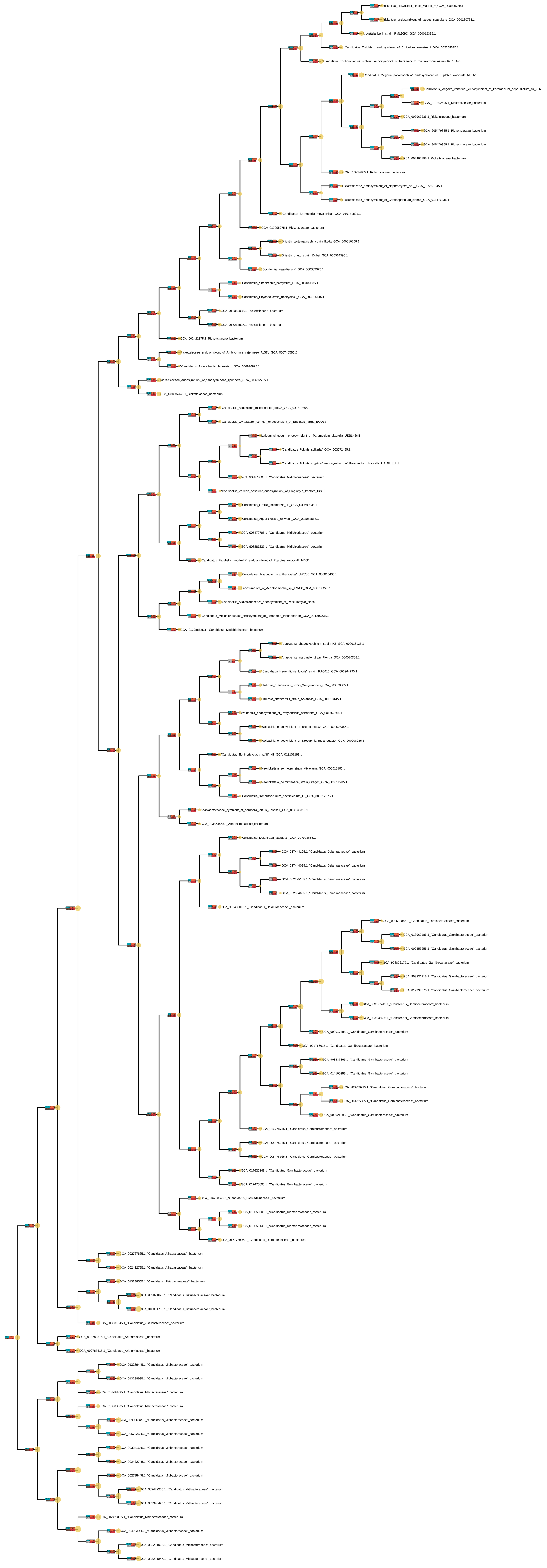

### Category A: RNA processing and modification

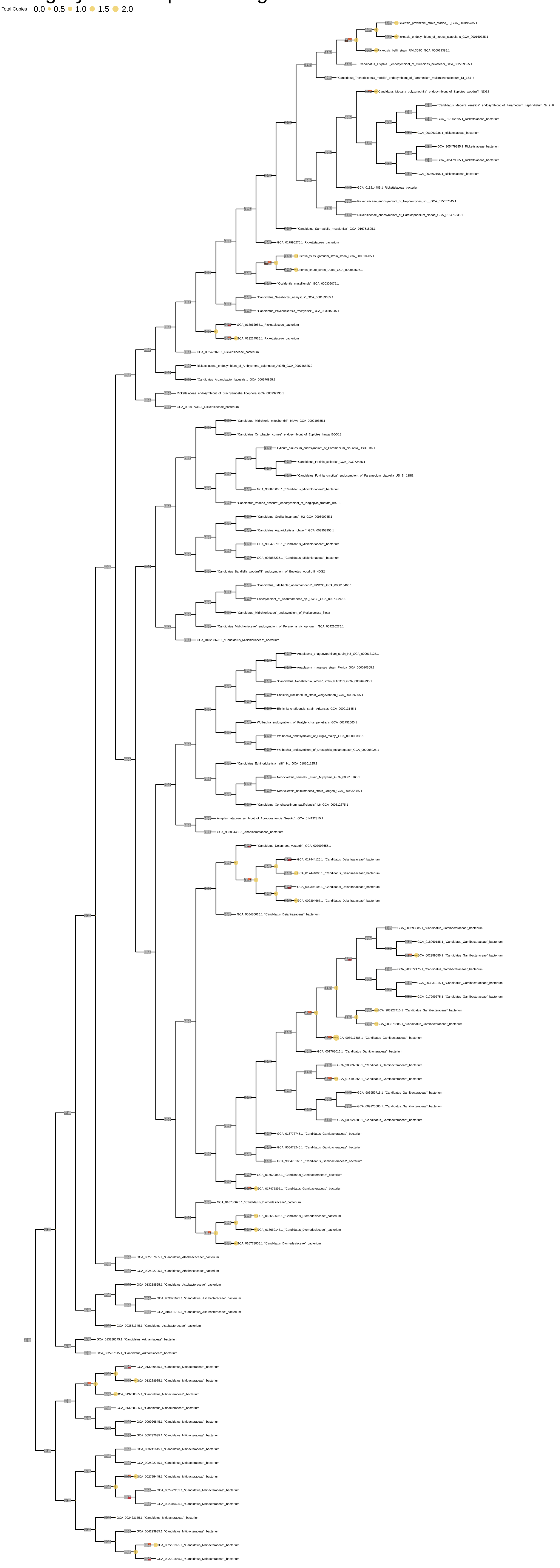

### Category B: Chromatin Structure and dynamics

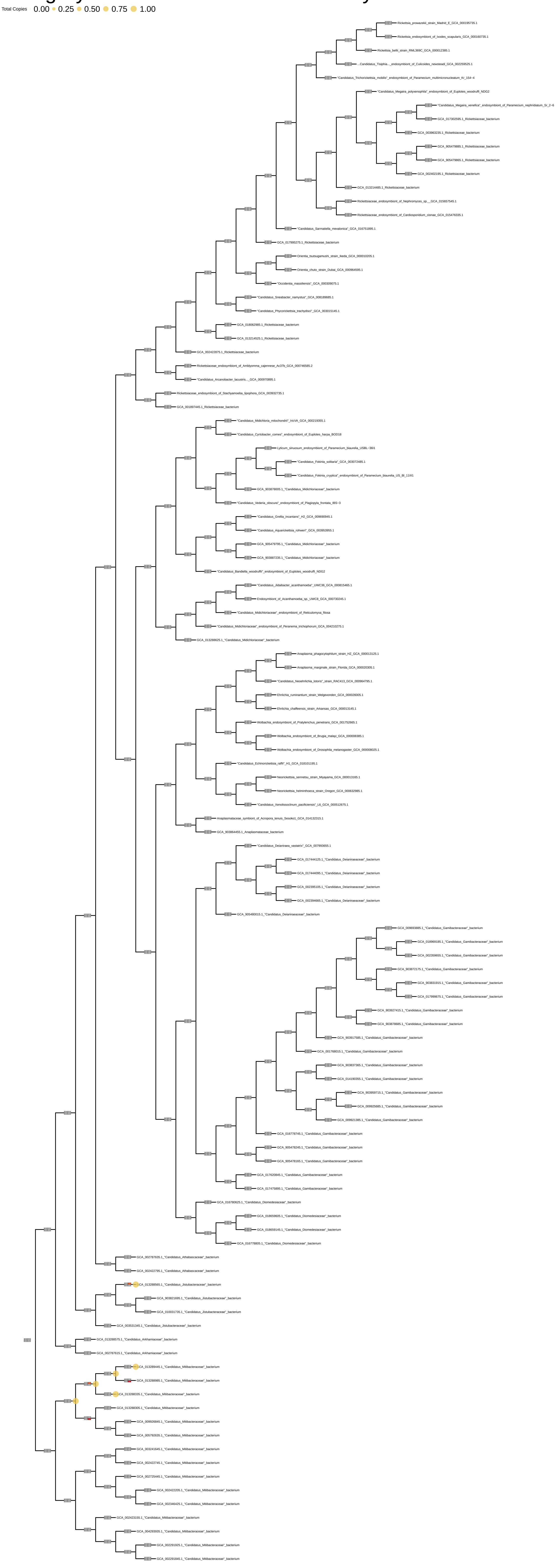

### Category C: Energy production and conversion

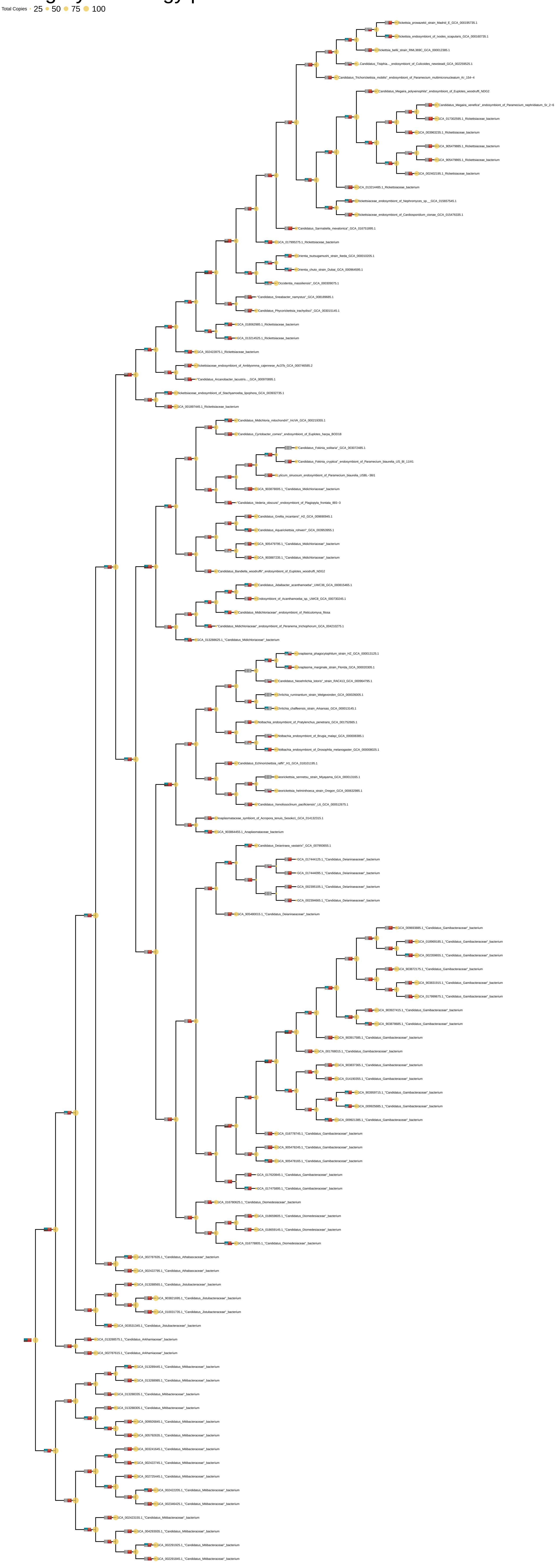

### Category D: Cell cycle control and mitosis

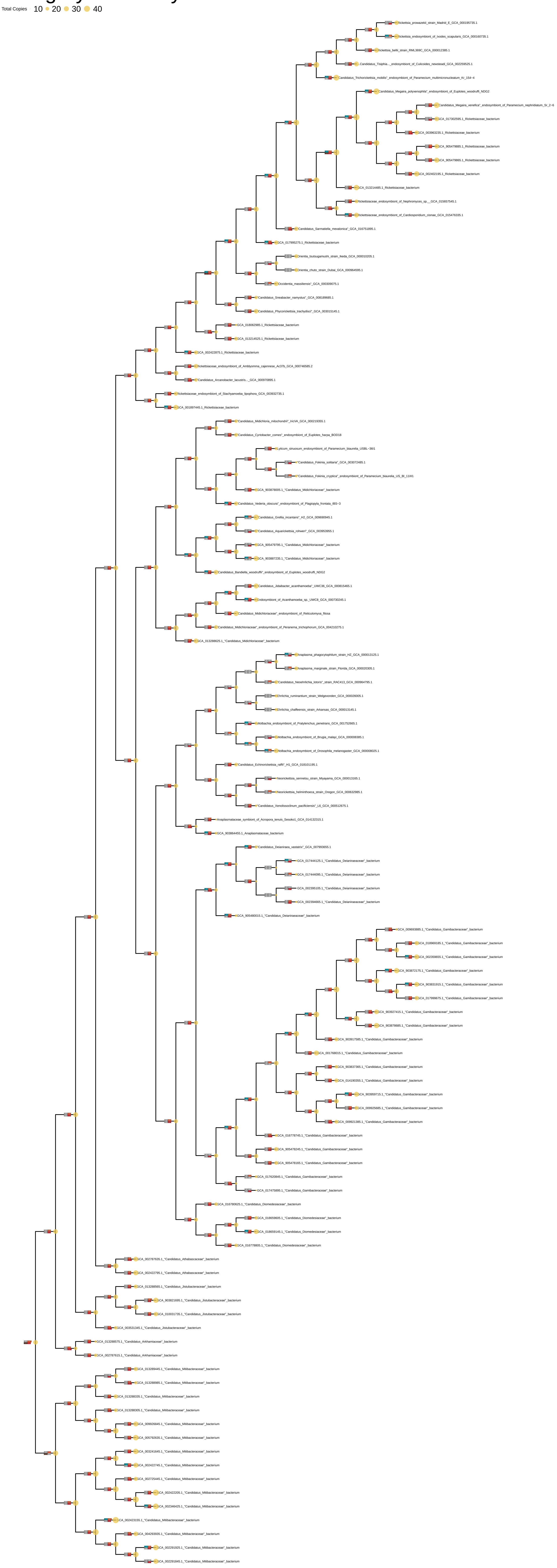

### Category E: Amino Acid metabolism and transport

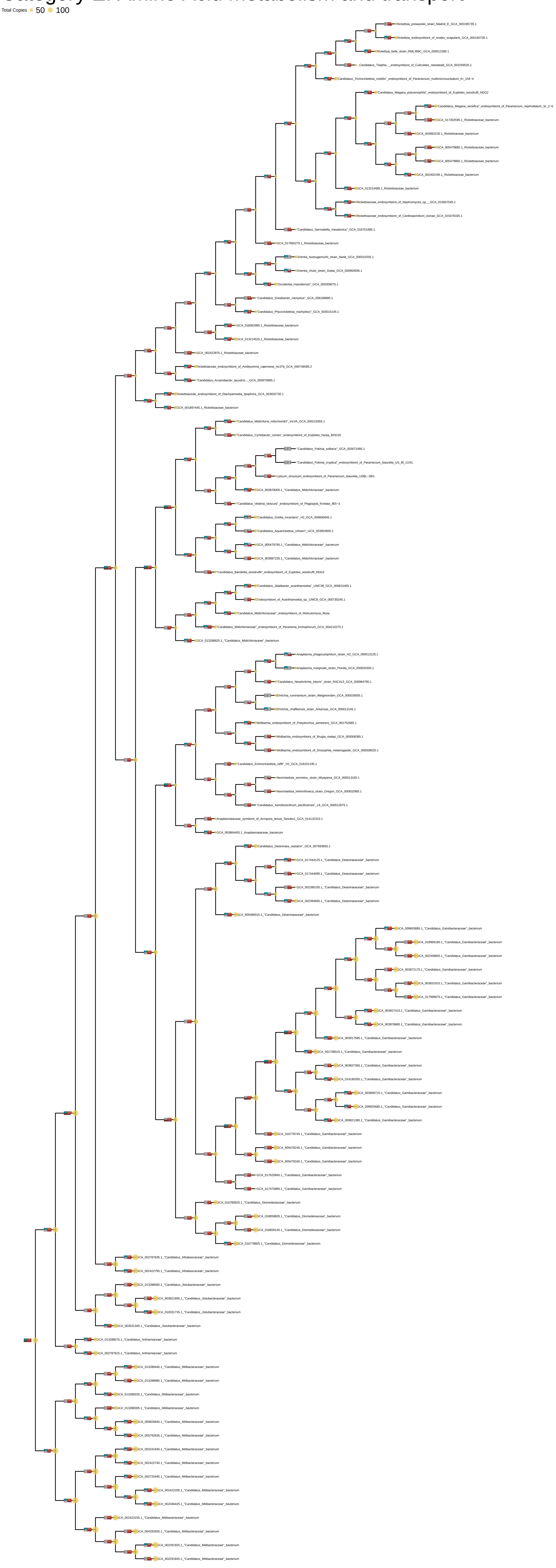

### Category F: Nucleotide metabolism and transport

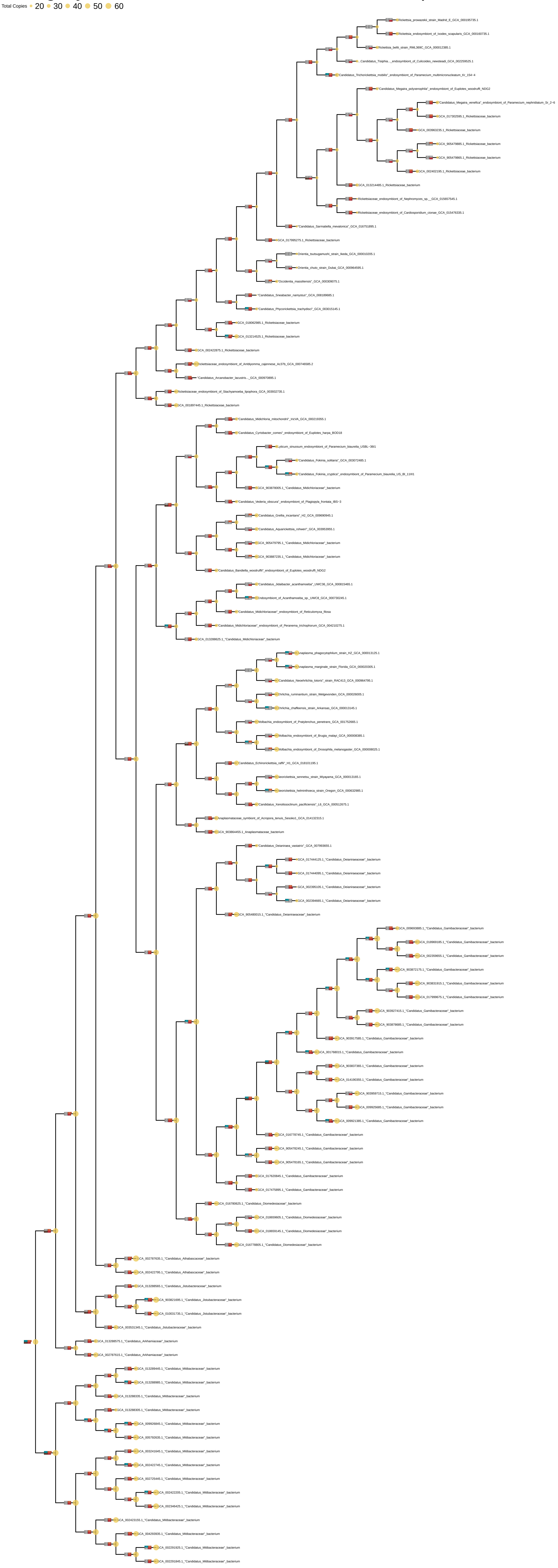

### Category G: Carbohydrate metabolism and transport

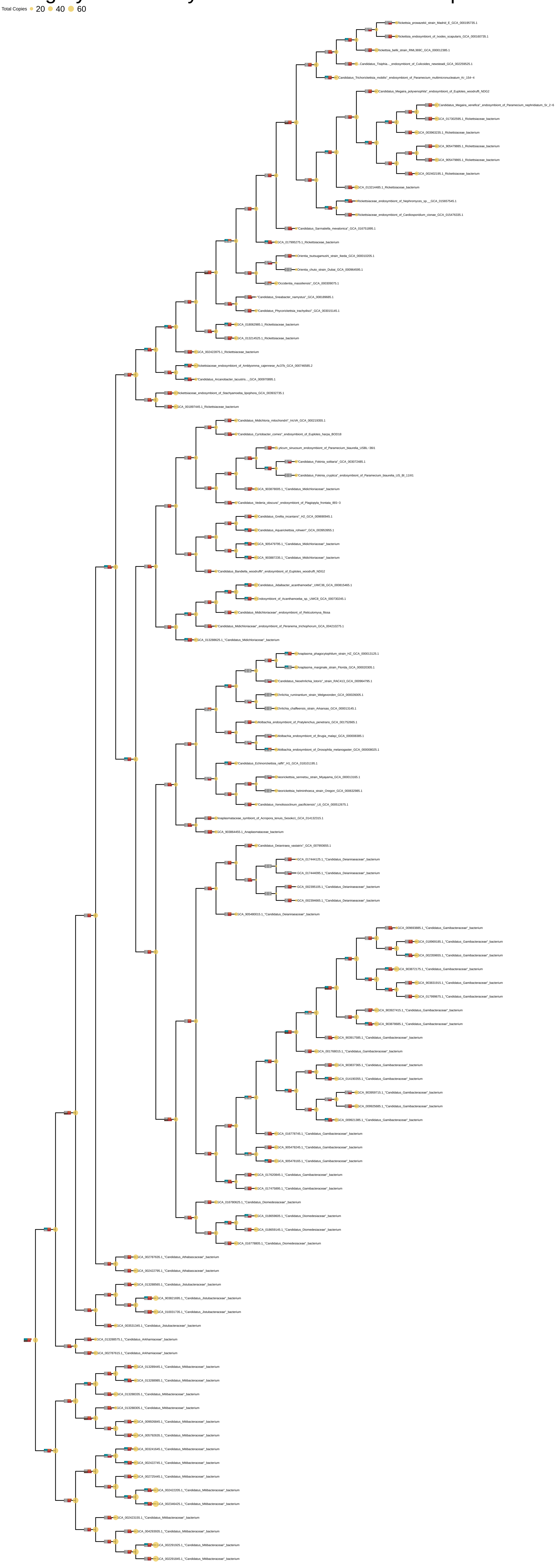

### Category H: Coenzyme transport and metabolism

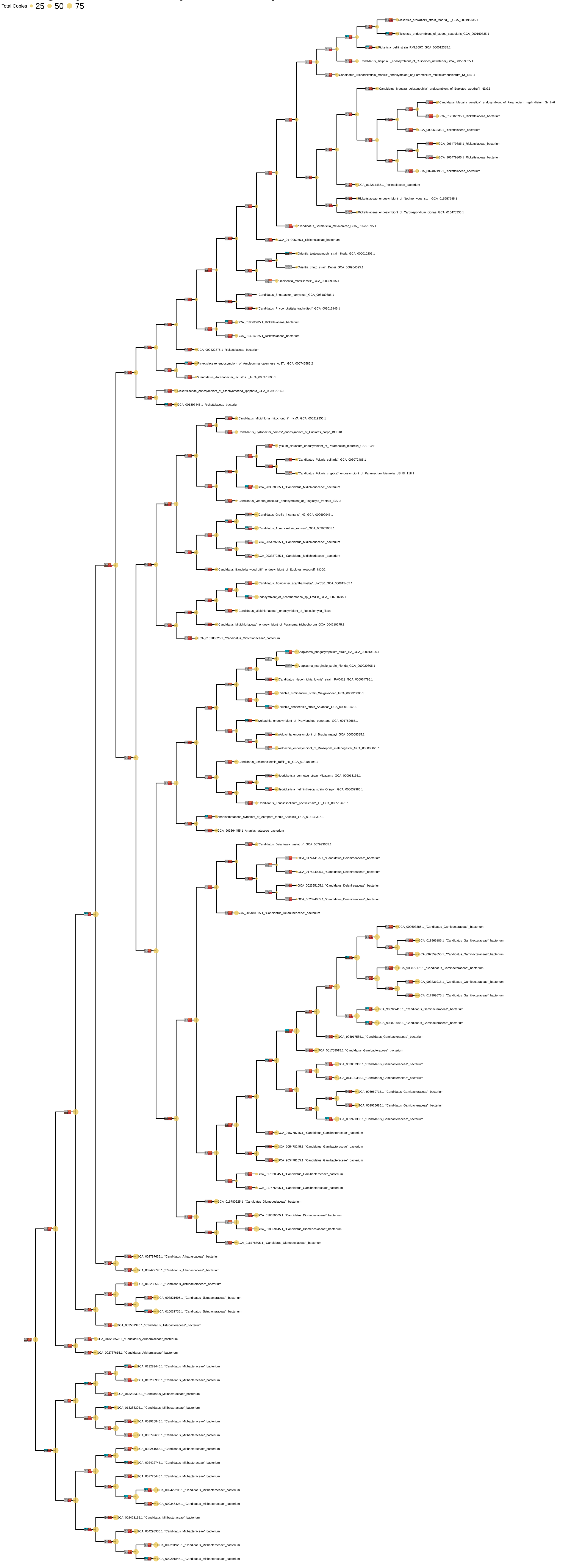

### Category I: Lipid metabolism

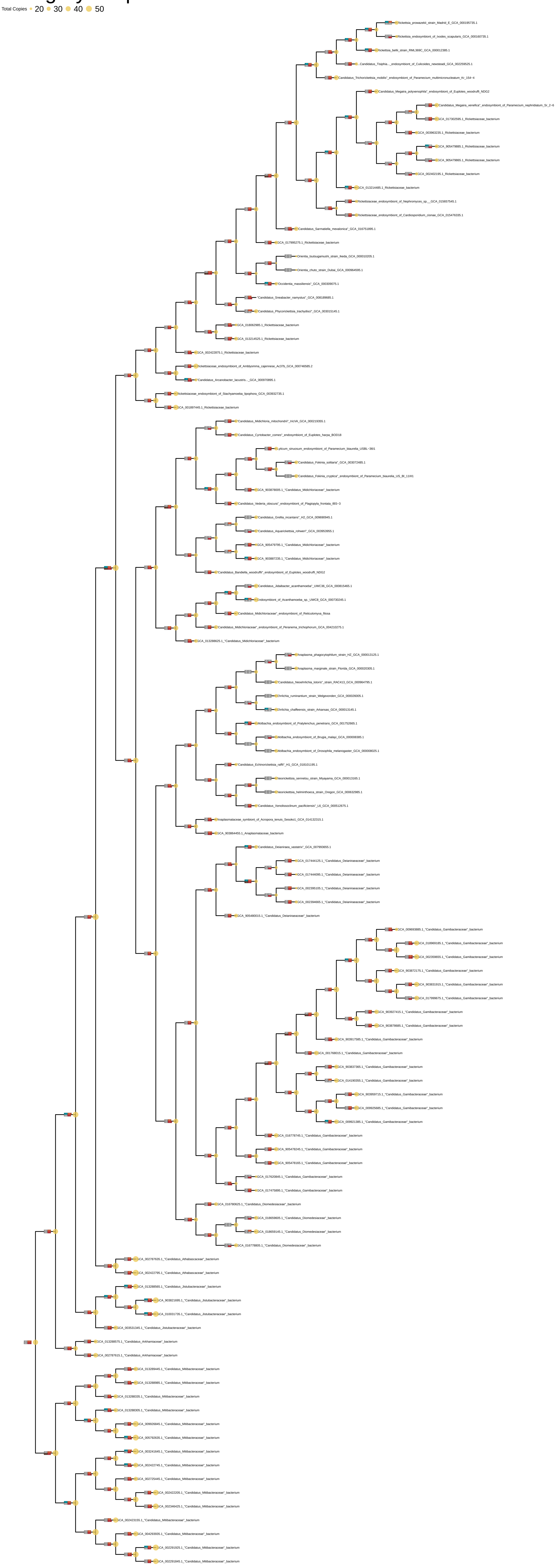

### Category J: Translation

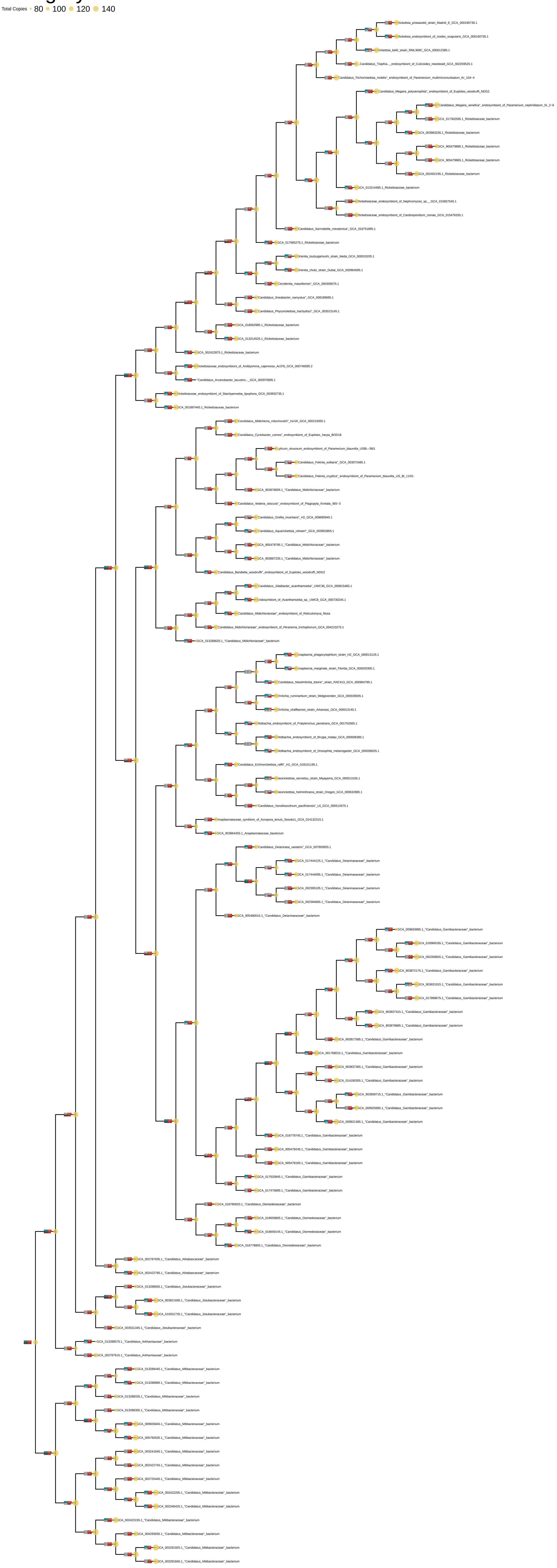

### Category K: Transcription

Total Copies • 20 • 40 • 60

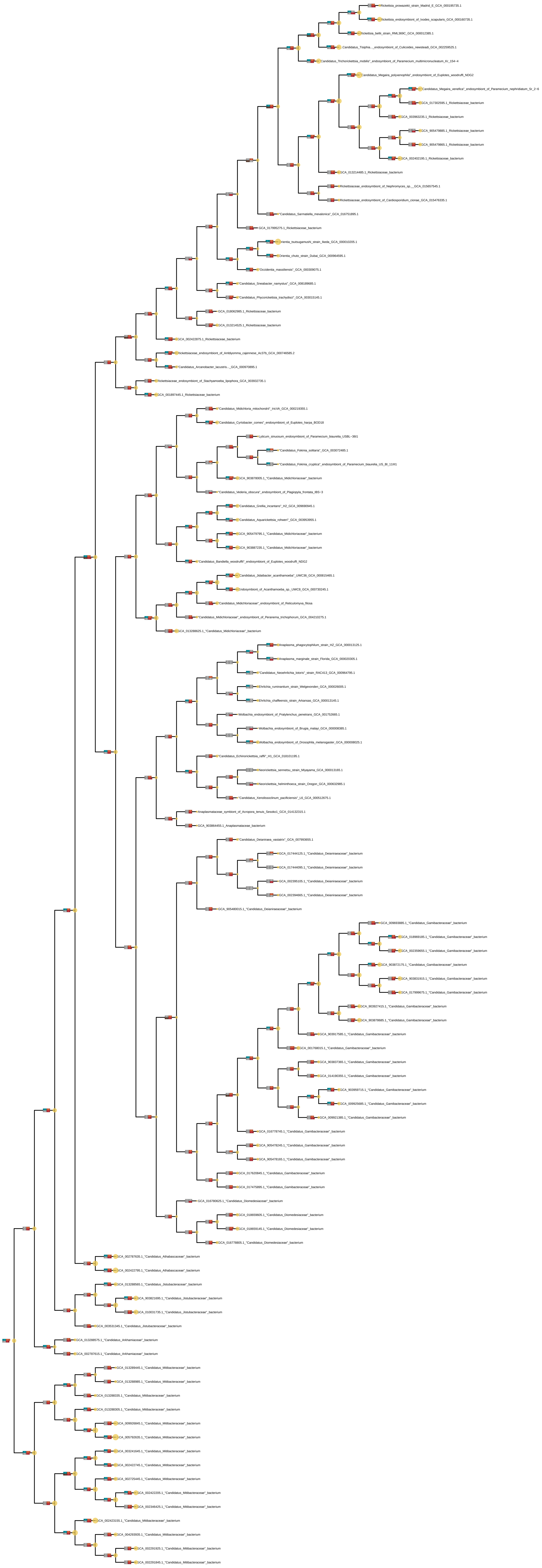

### Category L: Replication and repair

Total Copies • 50 • 100 • 150 • 200 • 250

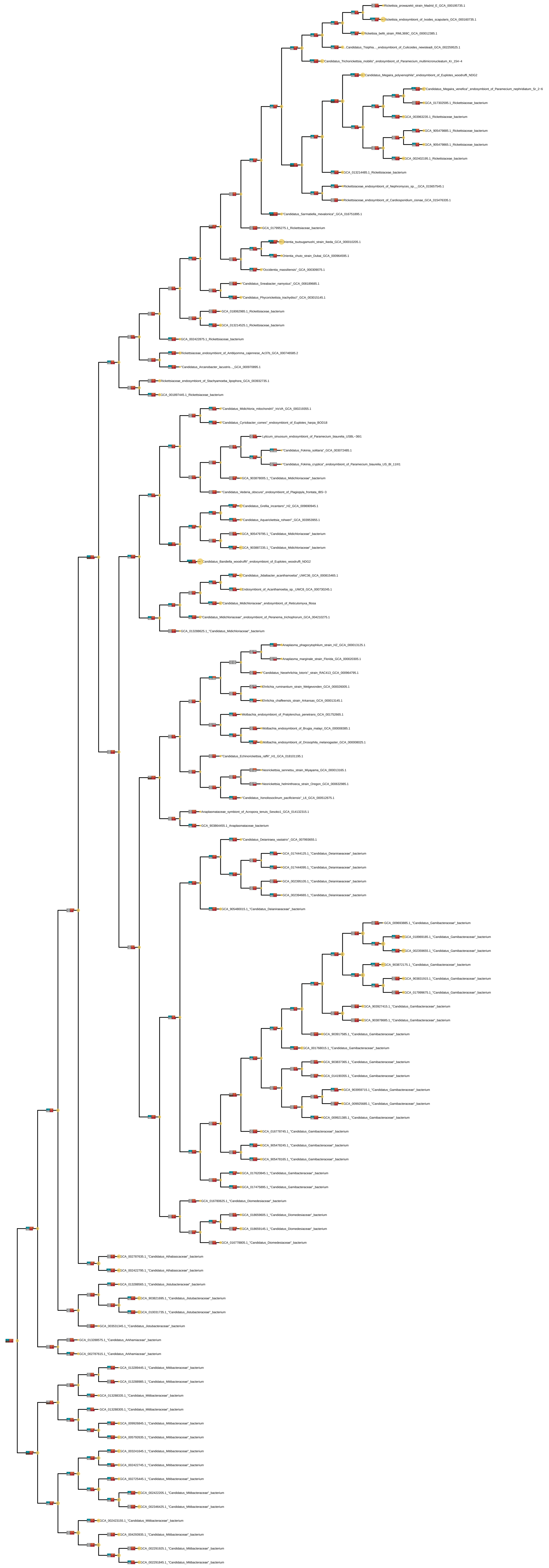

### Category M: Cell wall/membrane/envelope biogenesis

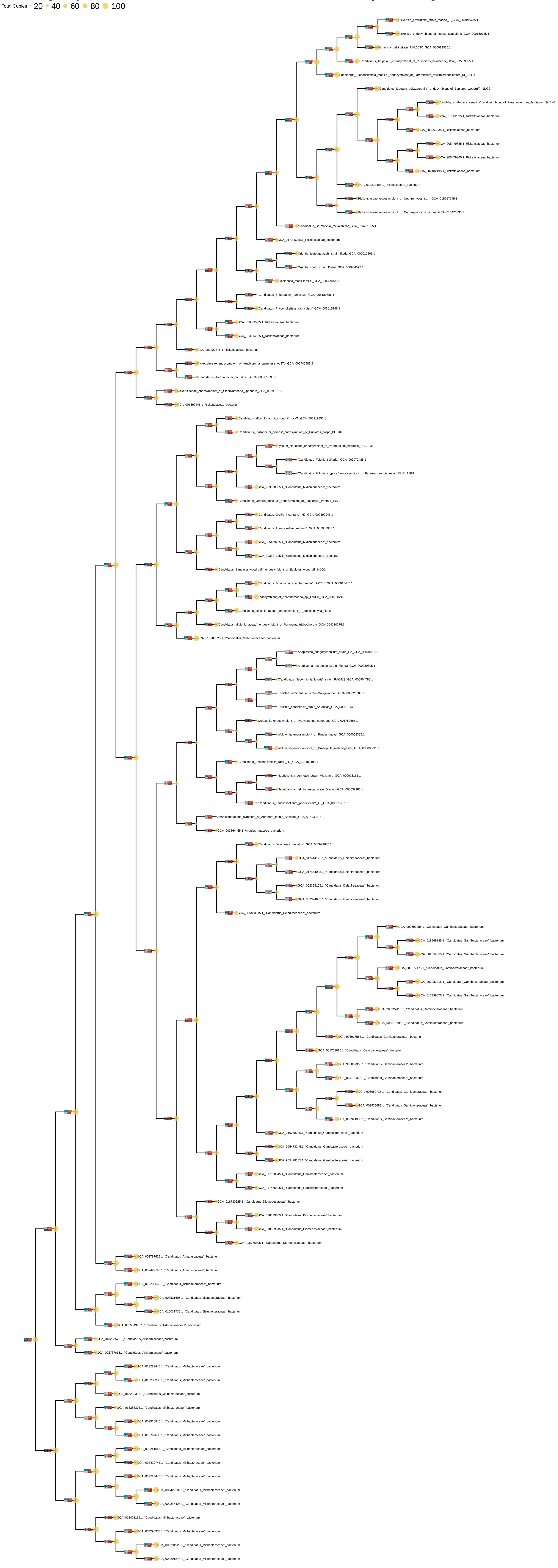

### Category N: Cell motility

Total Copies 25 50 75 100

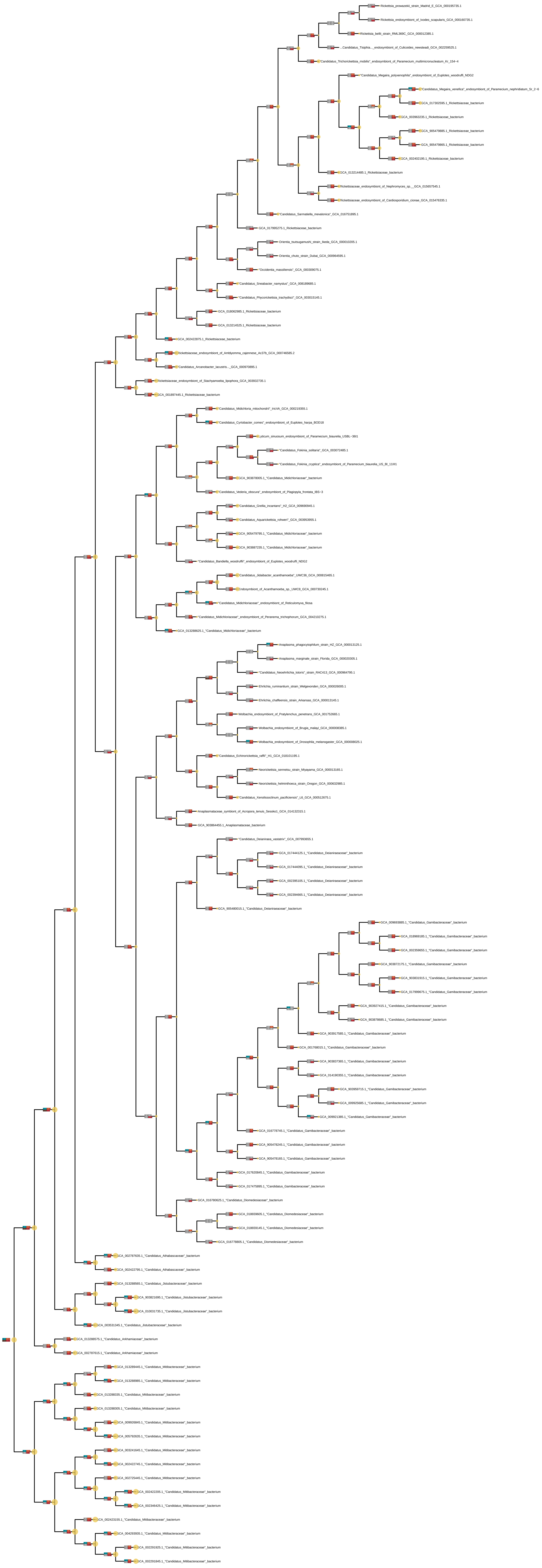

### Category O: Posttranslational modification and chaperones

Total Copies · 20 ● 40 ● 60 ● 80

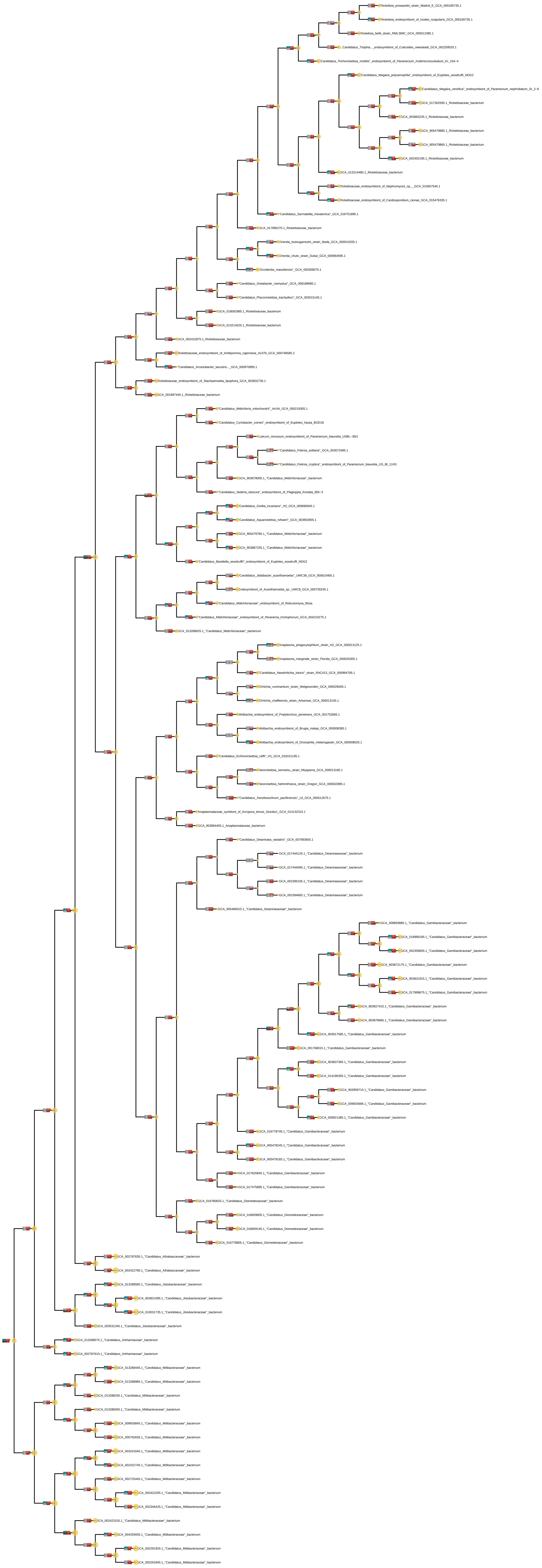

### Category P: Inorganic ion transport and metabolism

Total Copies ● 20 ● 40 ● 60 ● 80

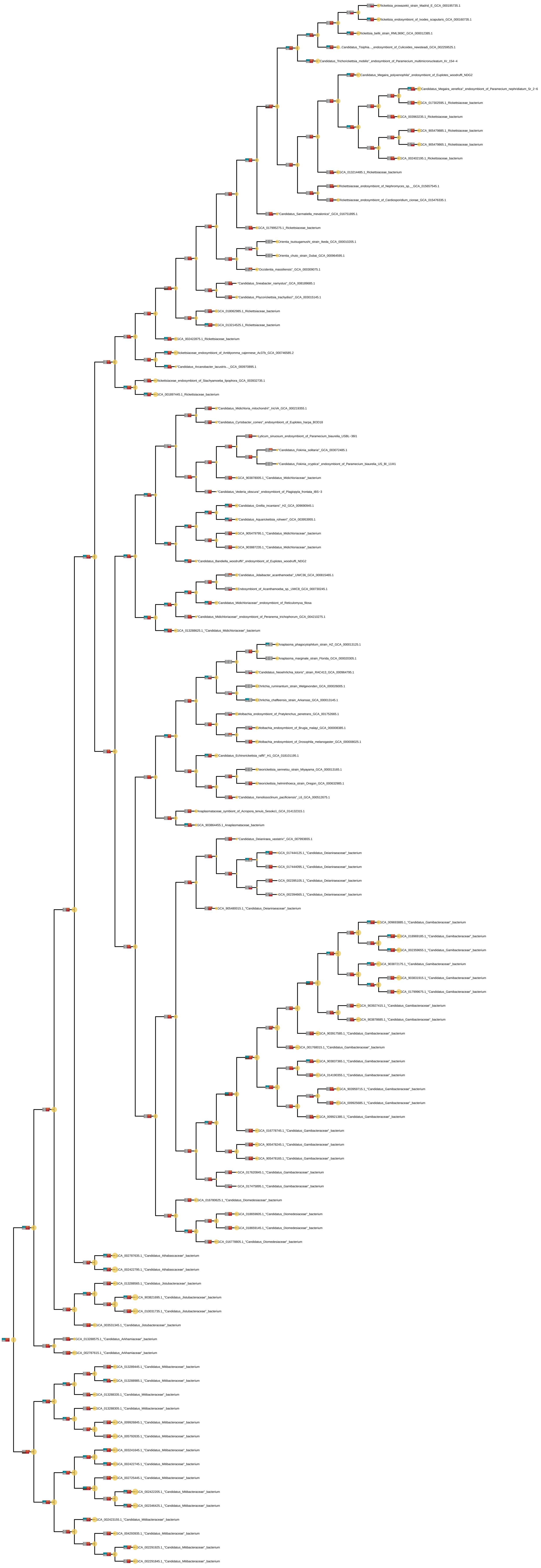

### Category Q: Secondary metabolites biosynthesis, transport and catabolism

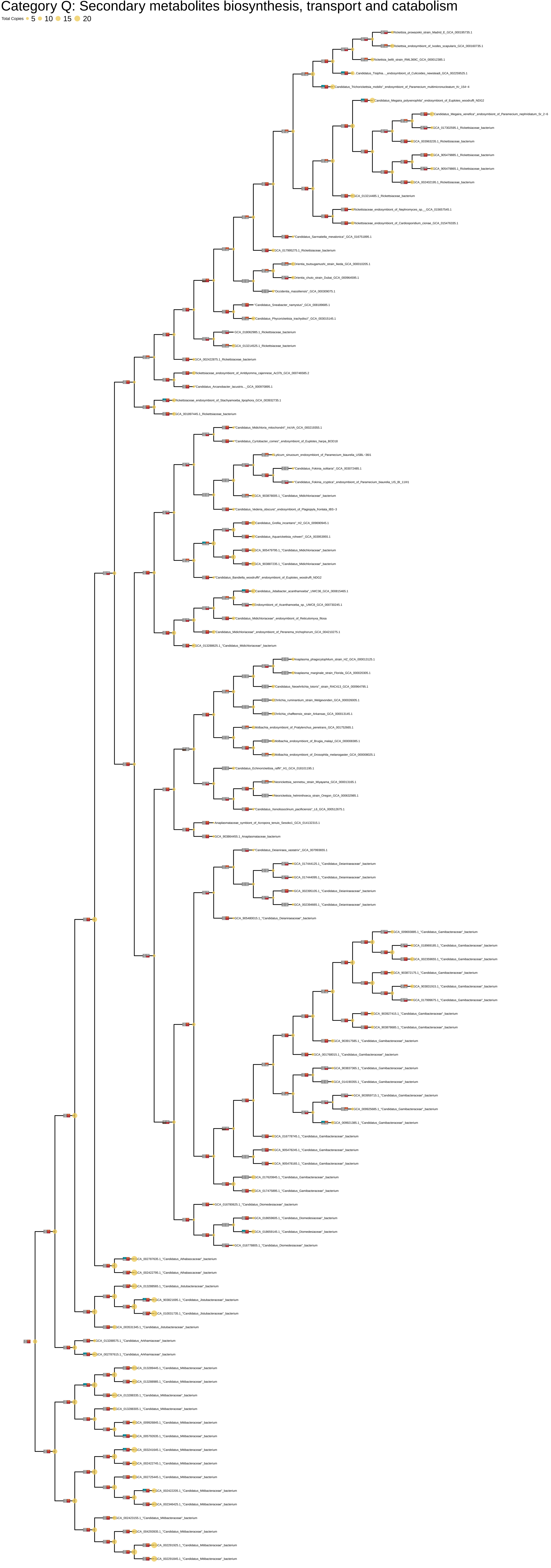

### Category S: Function Unknown

Total Copies ● 50 ● 100 ● 150 ● 200

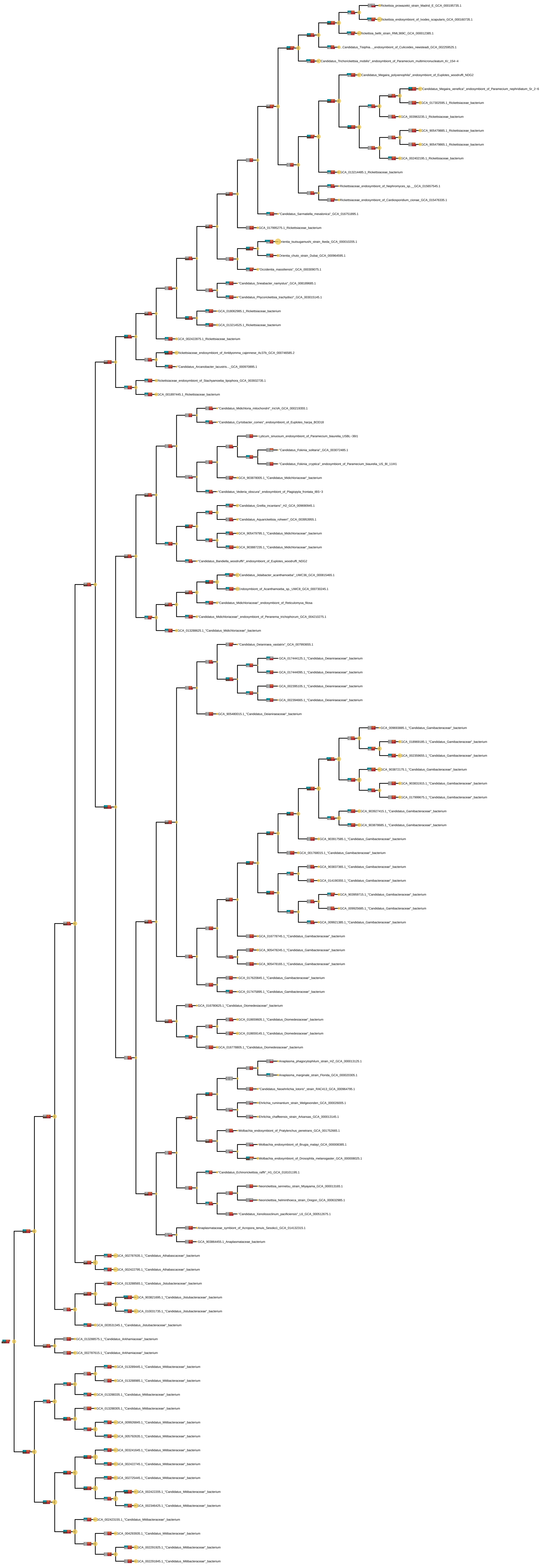

### Category T: Signal Transduction

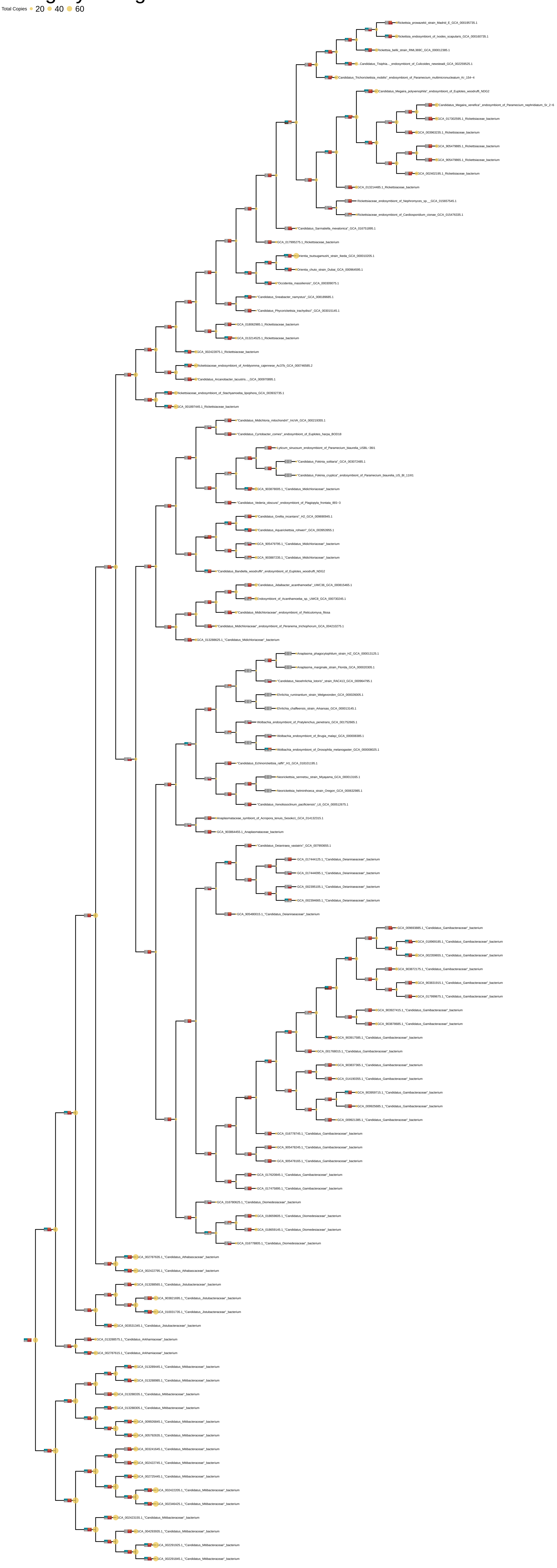

### Category U: Intracellular trafficking and secretion

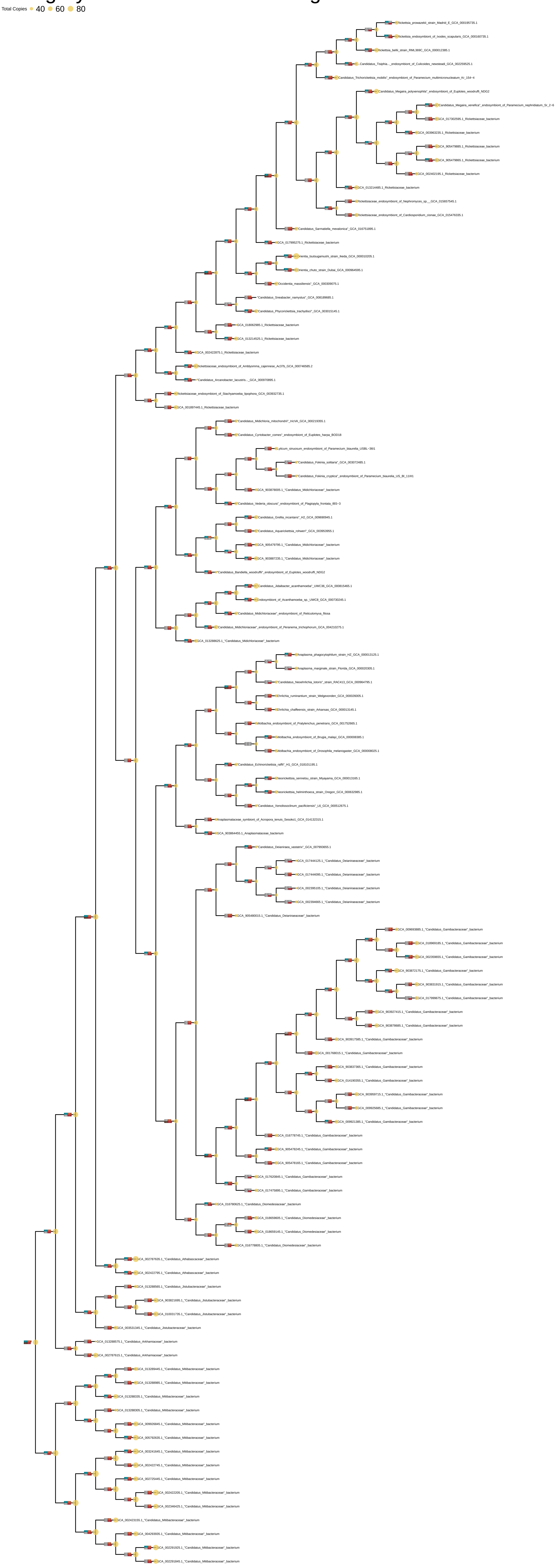

### Category V: Defense mechanisms

Total Copies ● 10 ● 20 ● 30

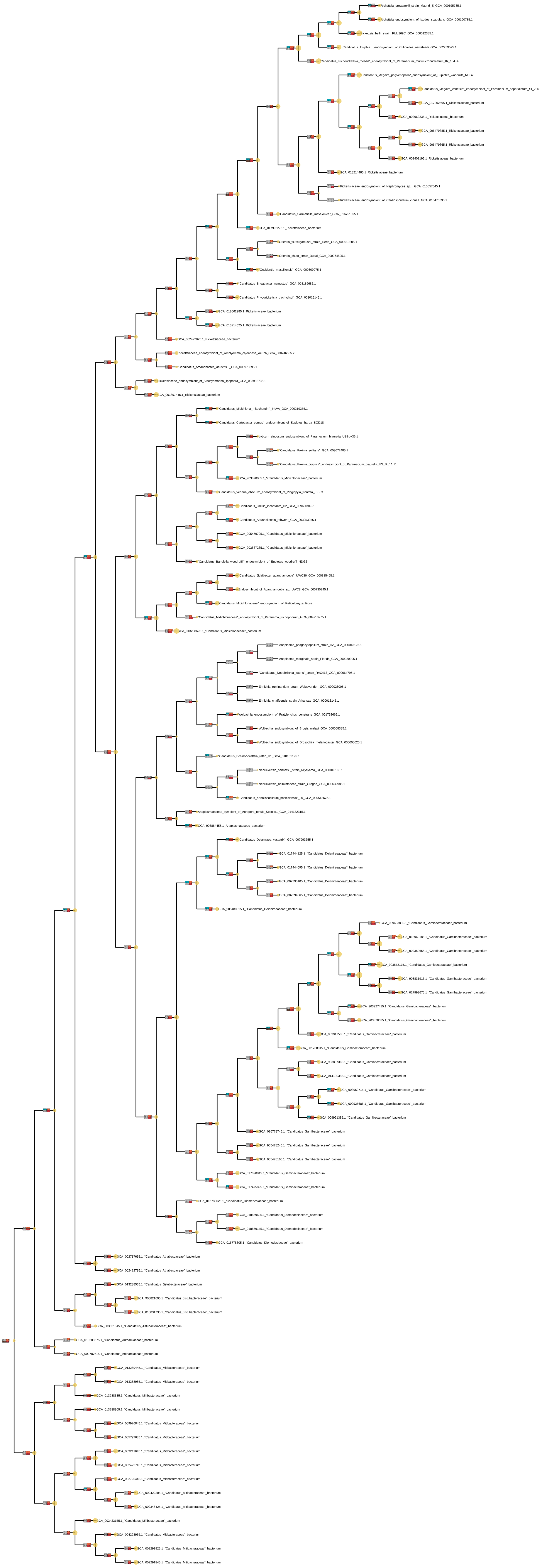

### Category W: Extracellular structures

Total Copies 0 1 2 3 4

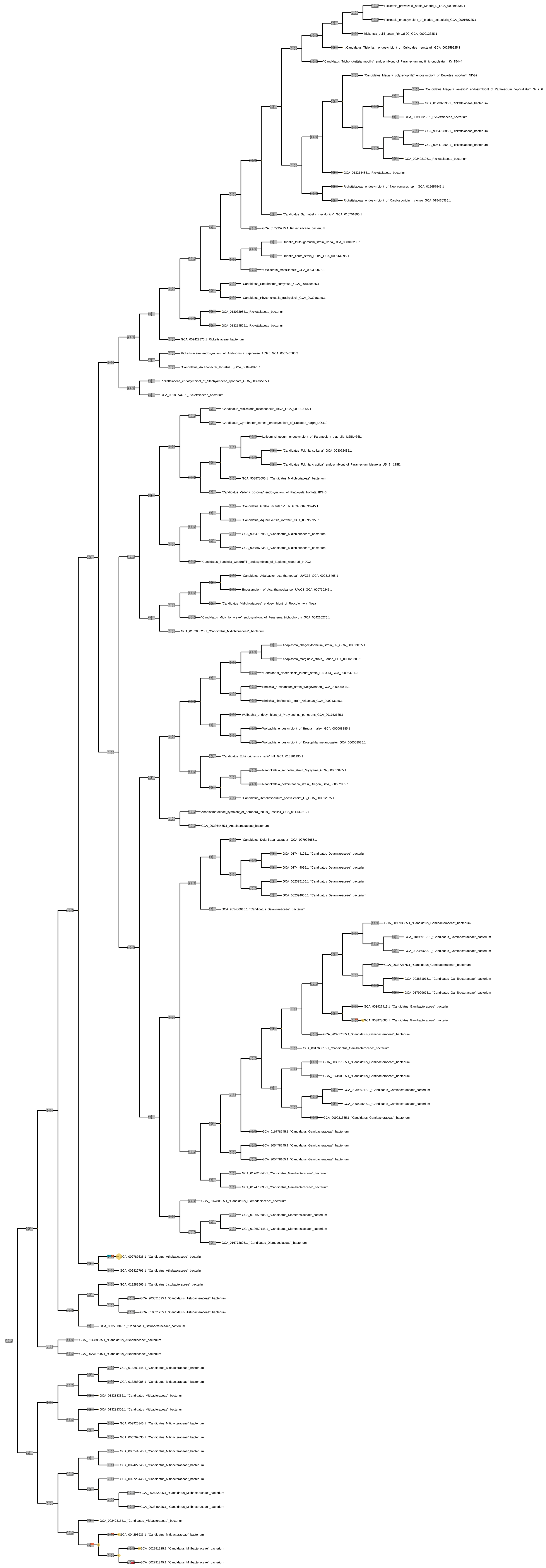

### Category Z: Cytoskeleton

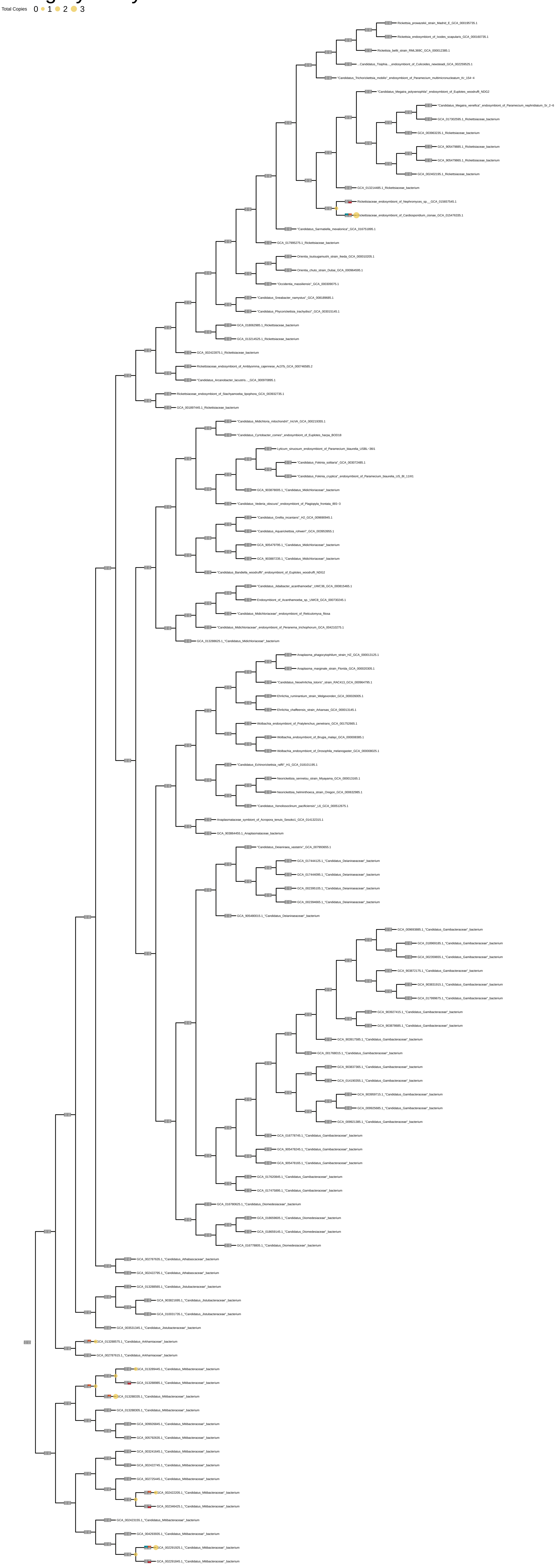
