## Supplementary material for "Host association and intracellularity evolved multiple times independently in the *Rickettsiales*": figures_and_supplementary_materials: Supplementary_figure_8_Biosynthetic_pathways_phylogeny.pdf

### Purines_all_organisms

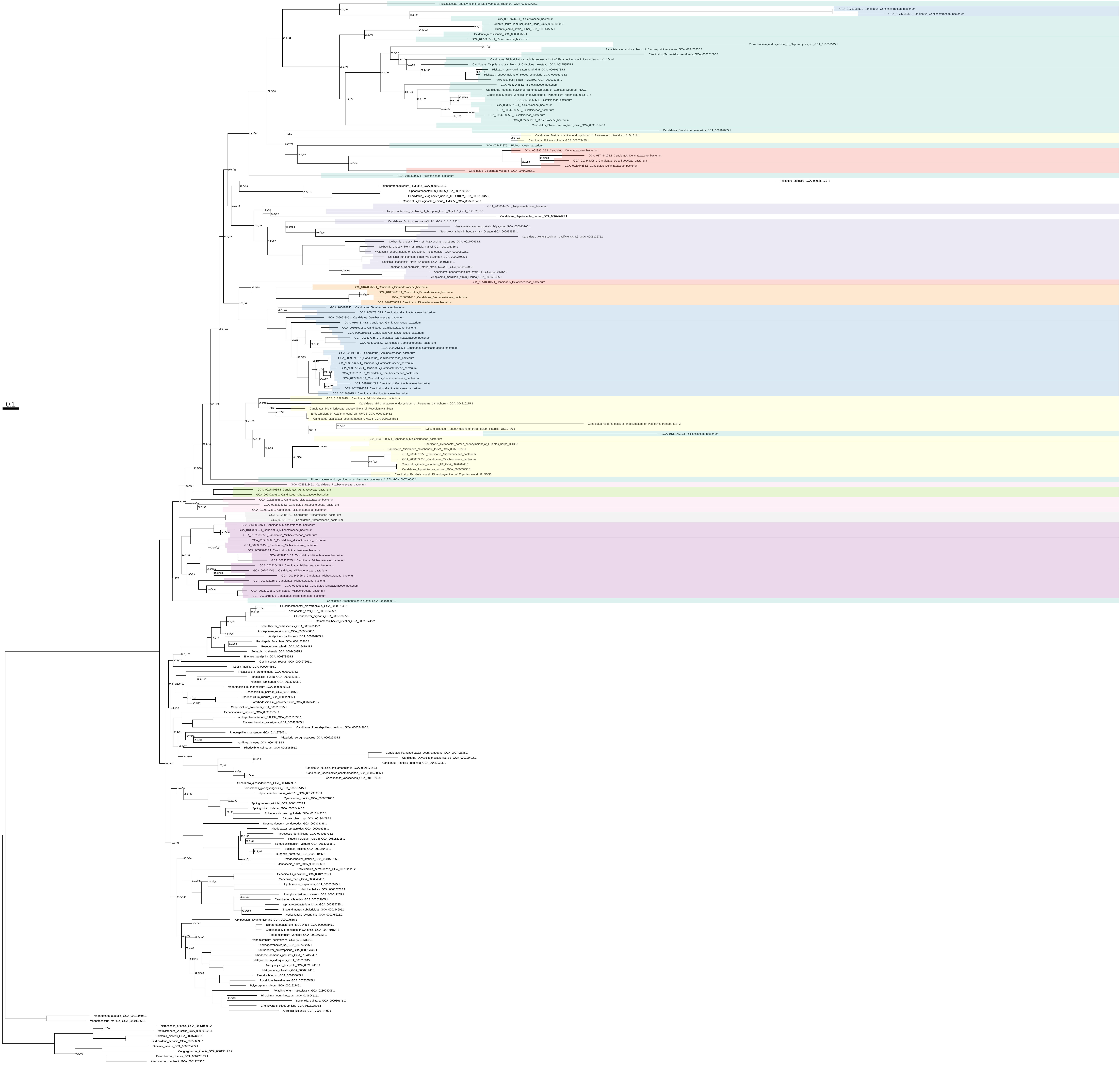

Purines_all_organisms_z_0.10

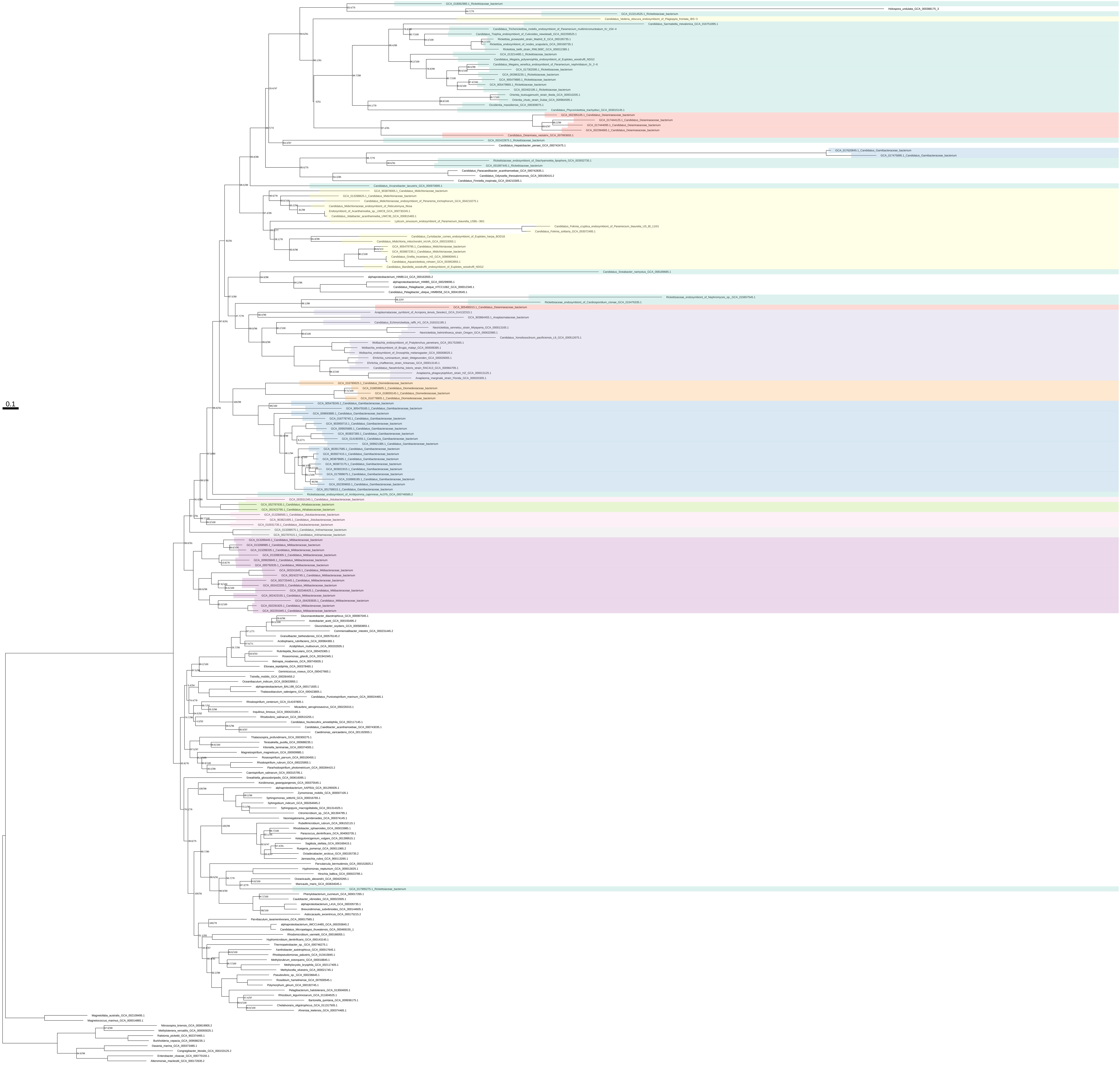

Purines_all_organisms_z_0.20

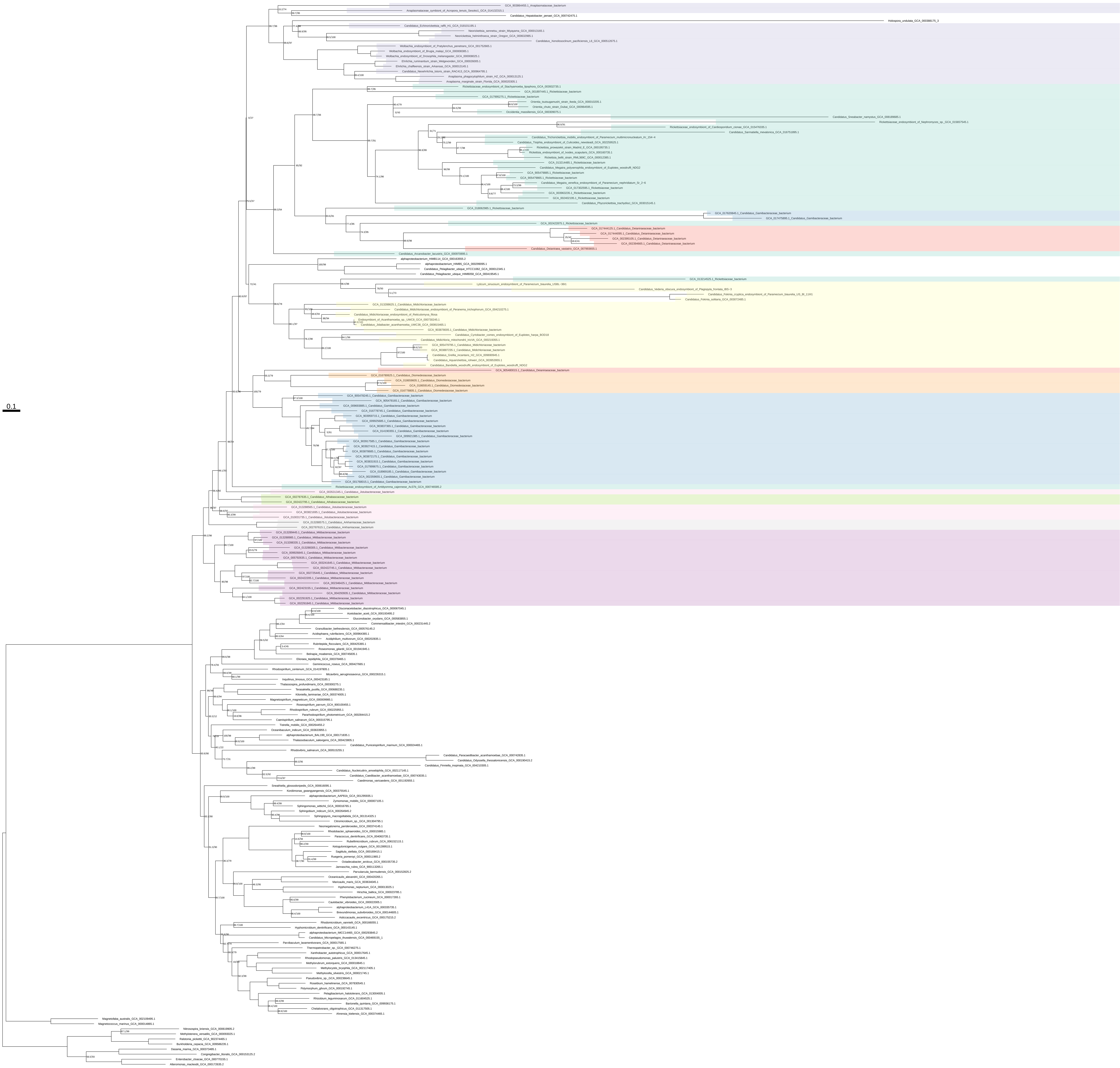

### Purines_all_organisms_z_0.30

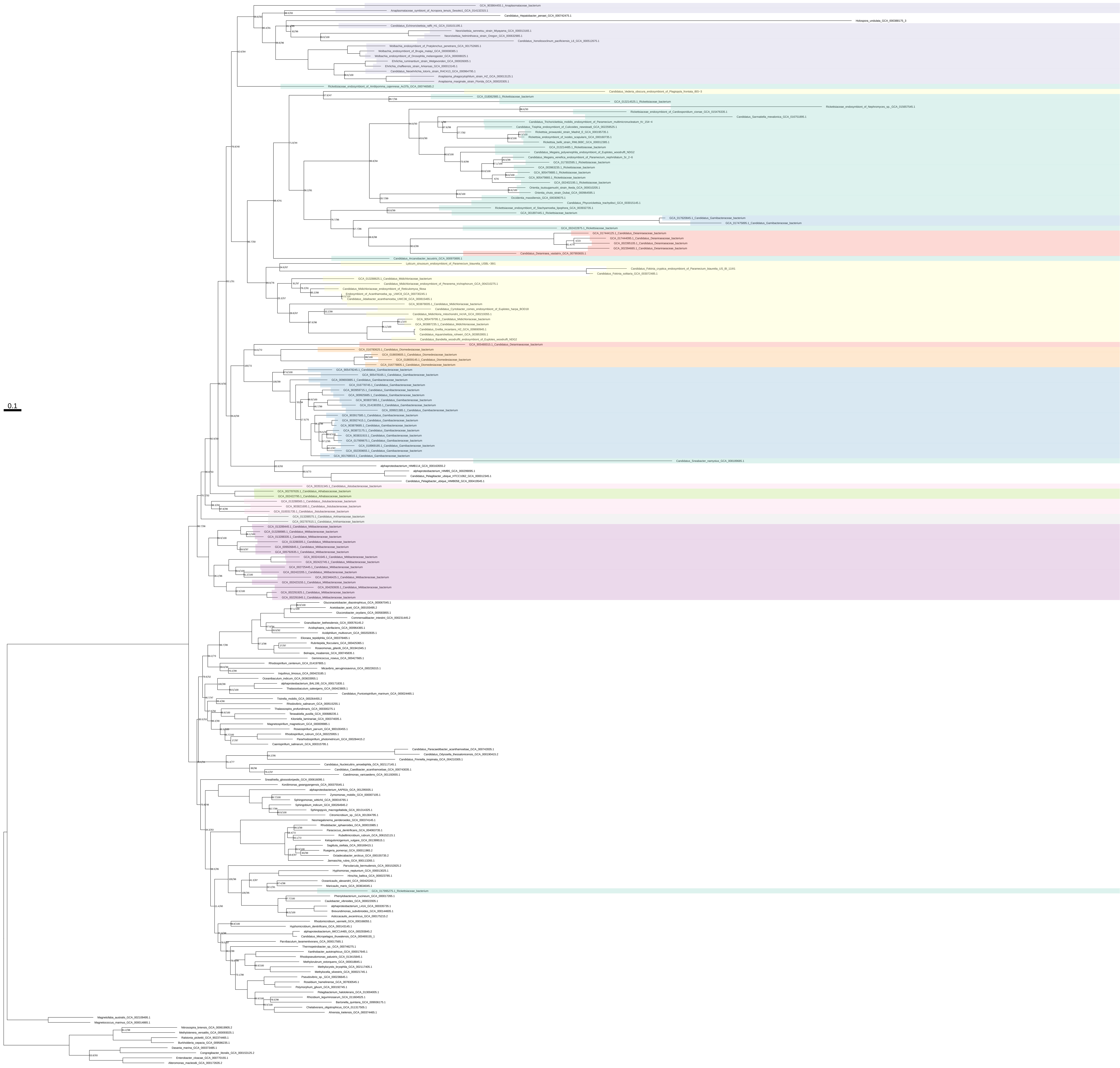

### Purines_all_organisms_z_0.40

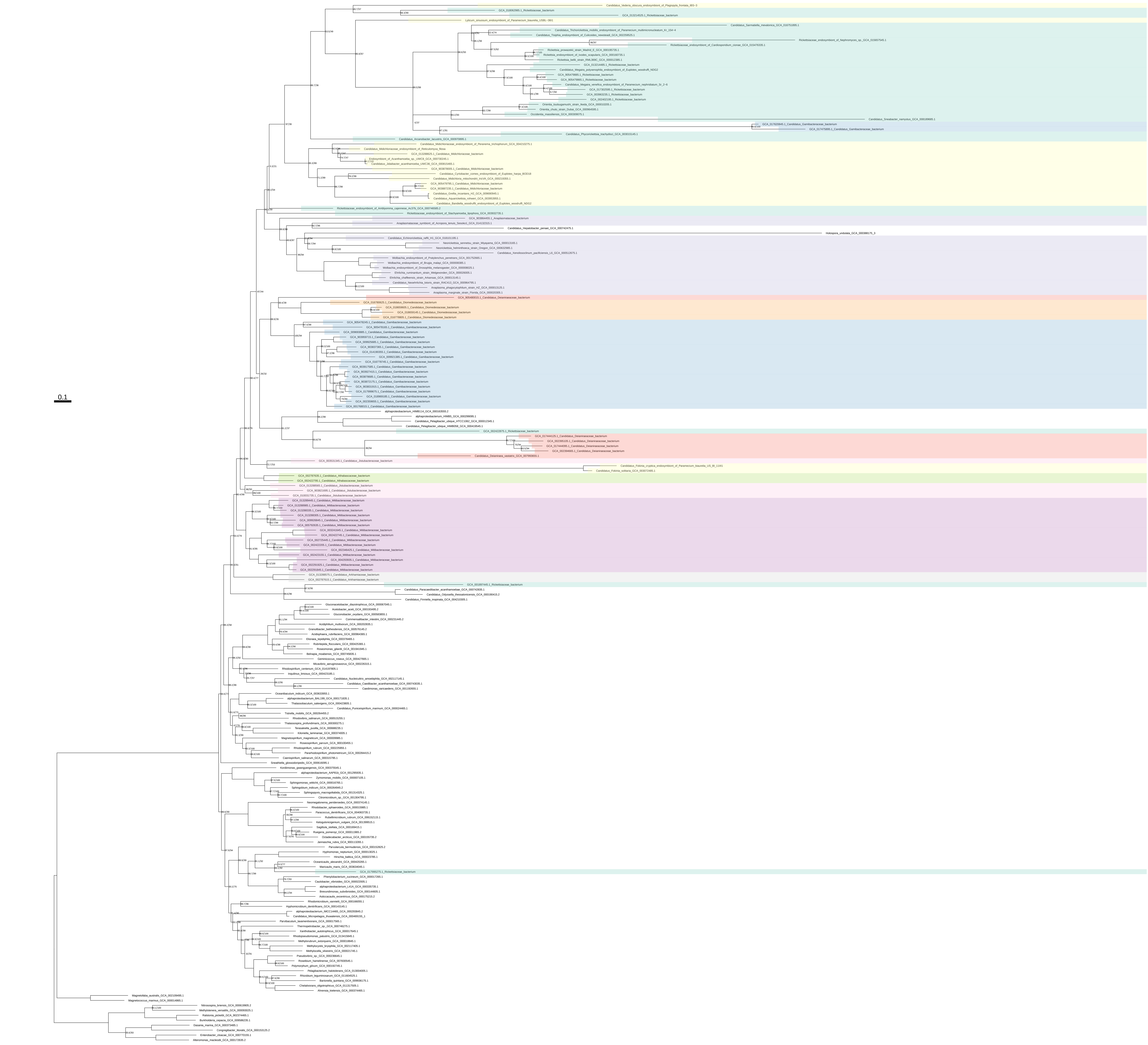

### Purines_all_organisms_z_0.50

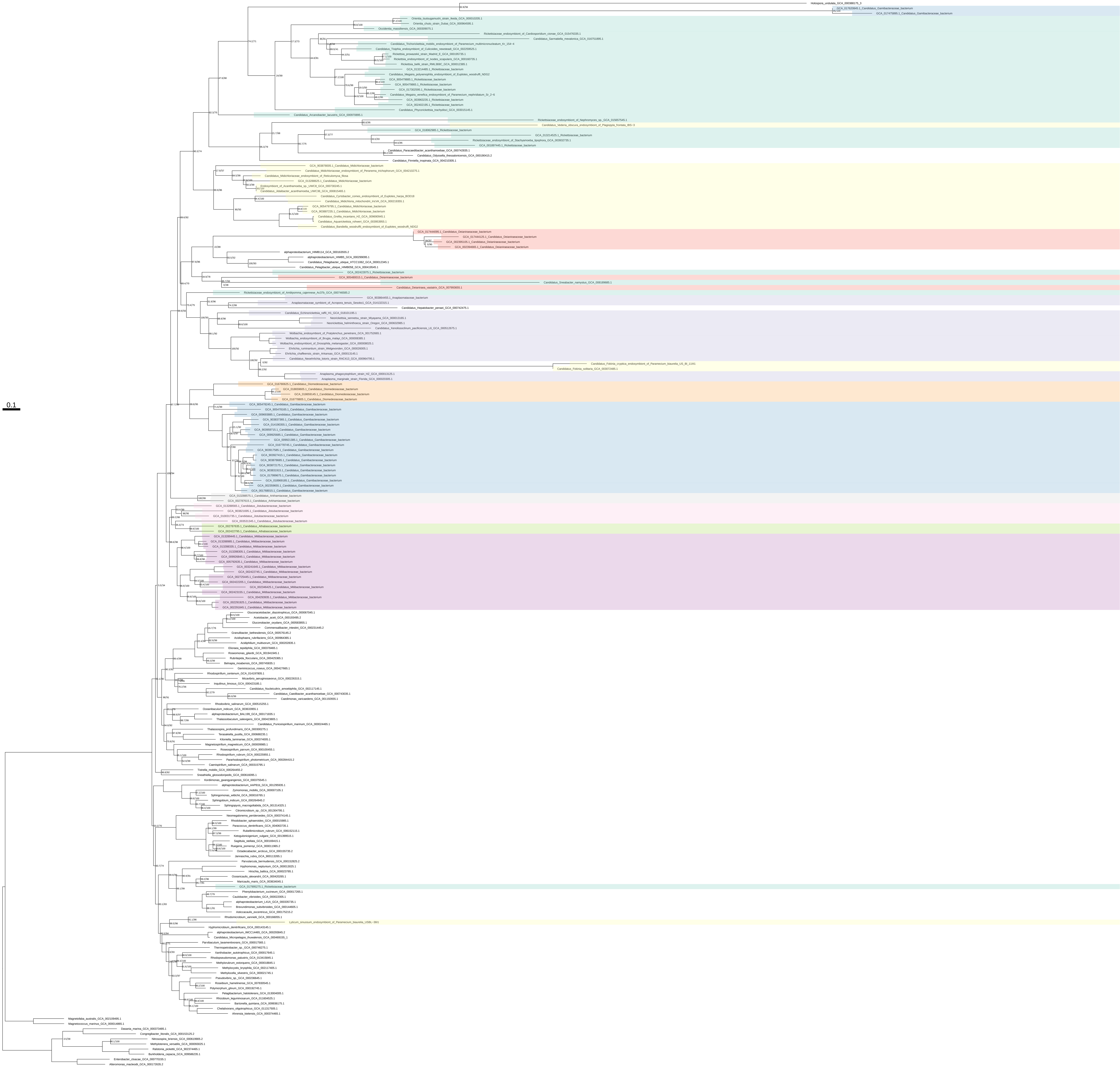

### Purines_organism_selection

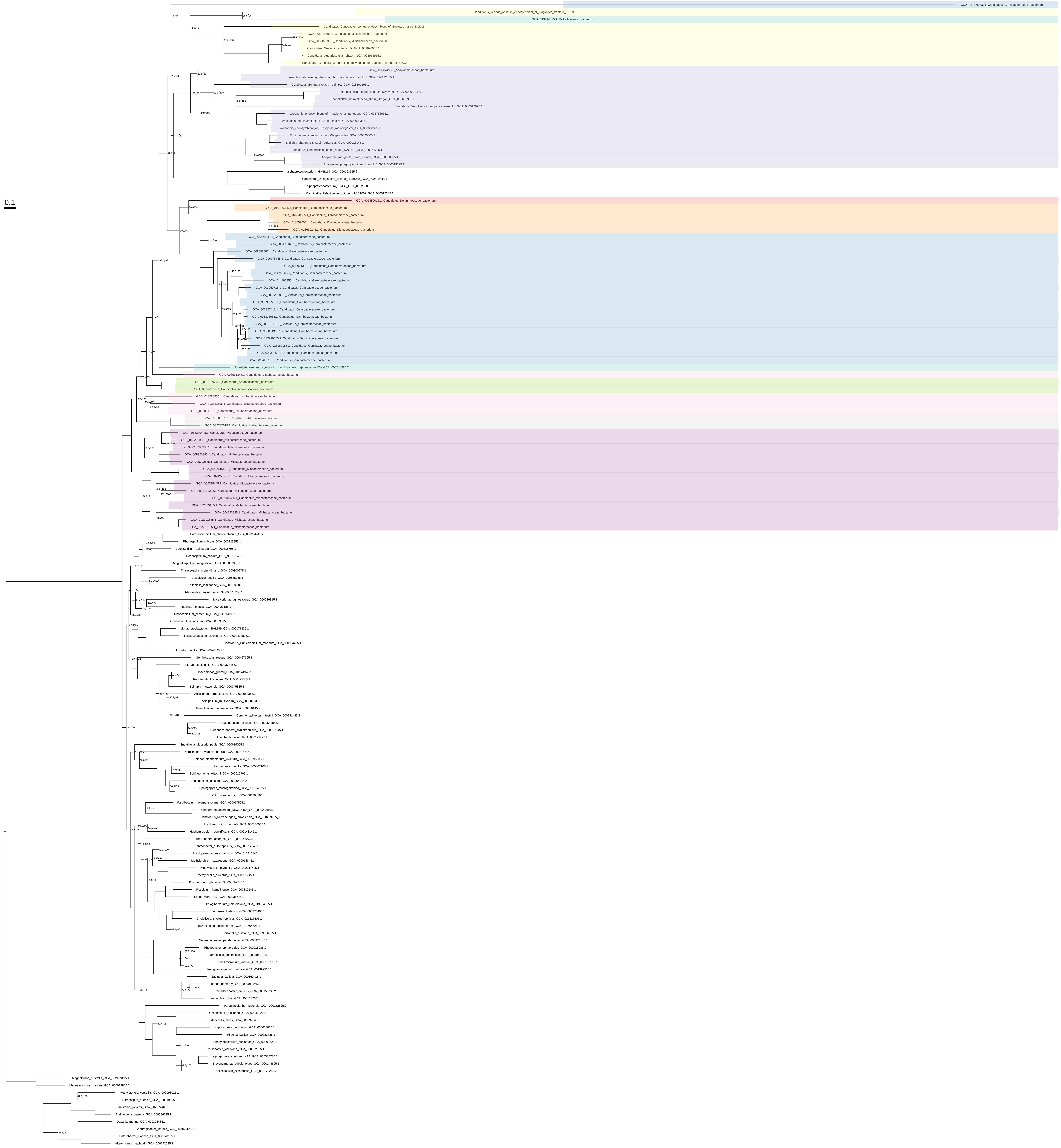

### Purines_organism_selection_z_0.10

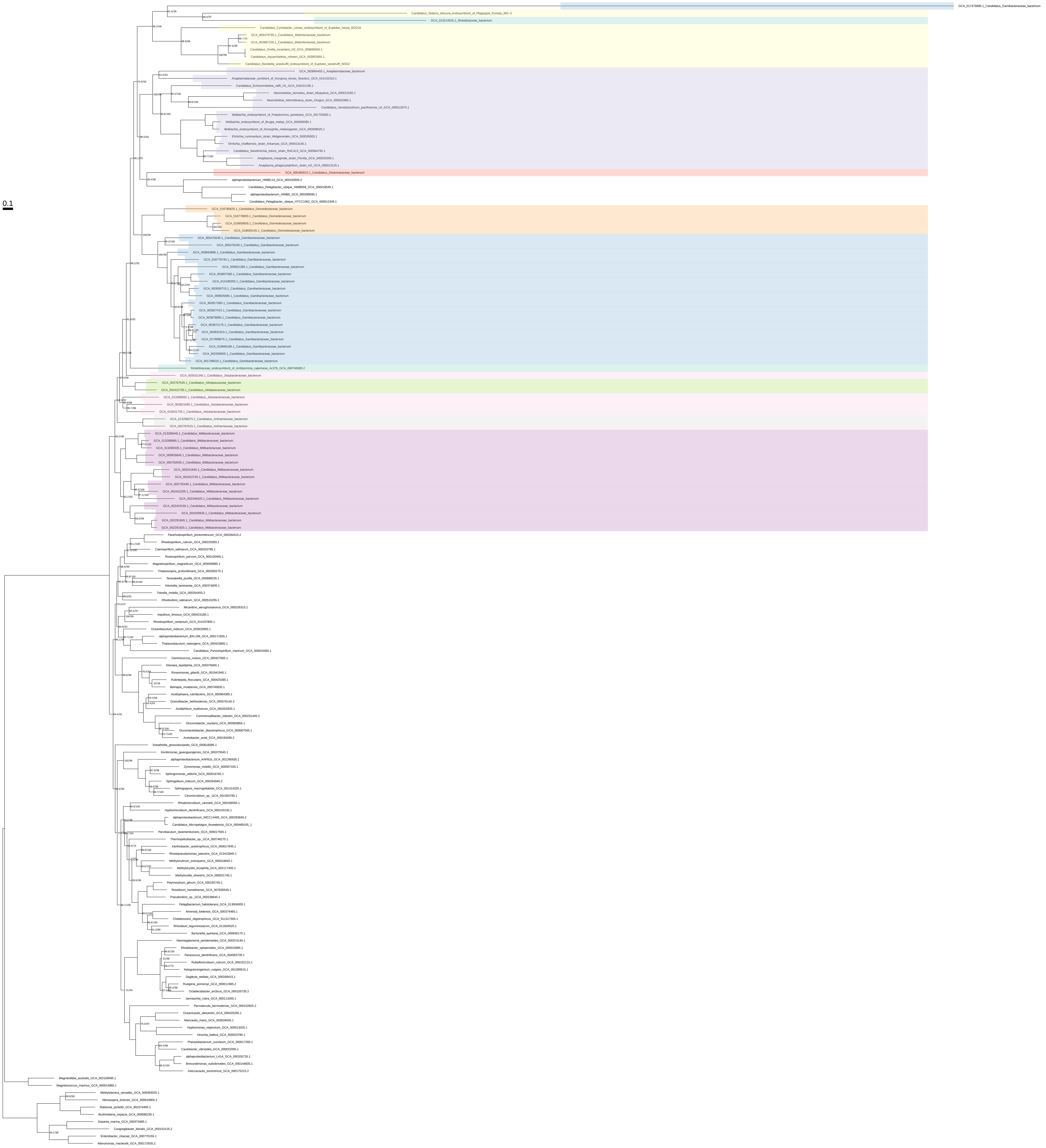

### Purines_organism_selection_z_0.20

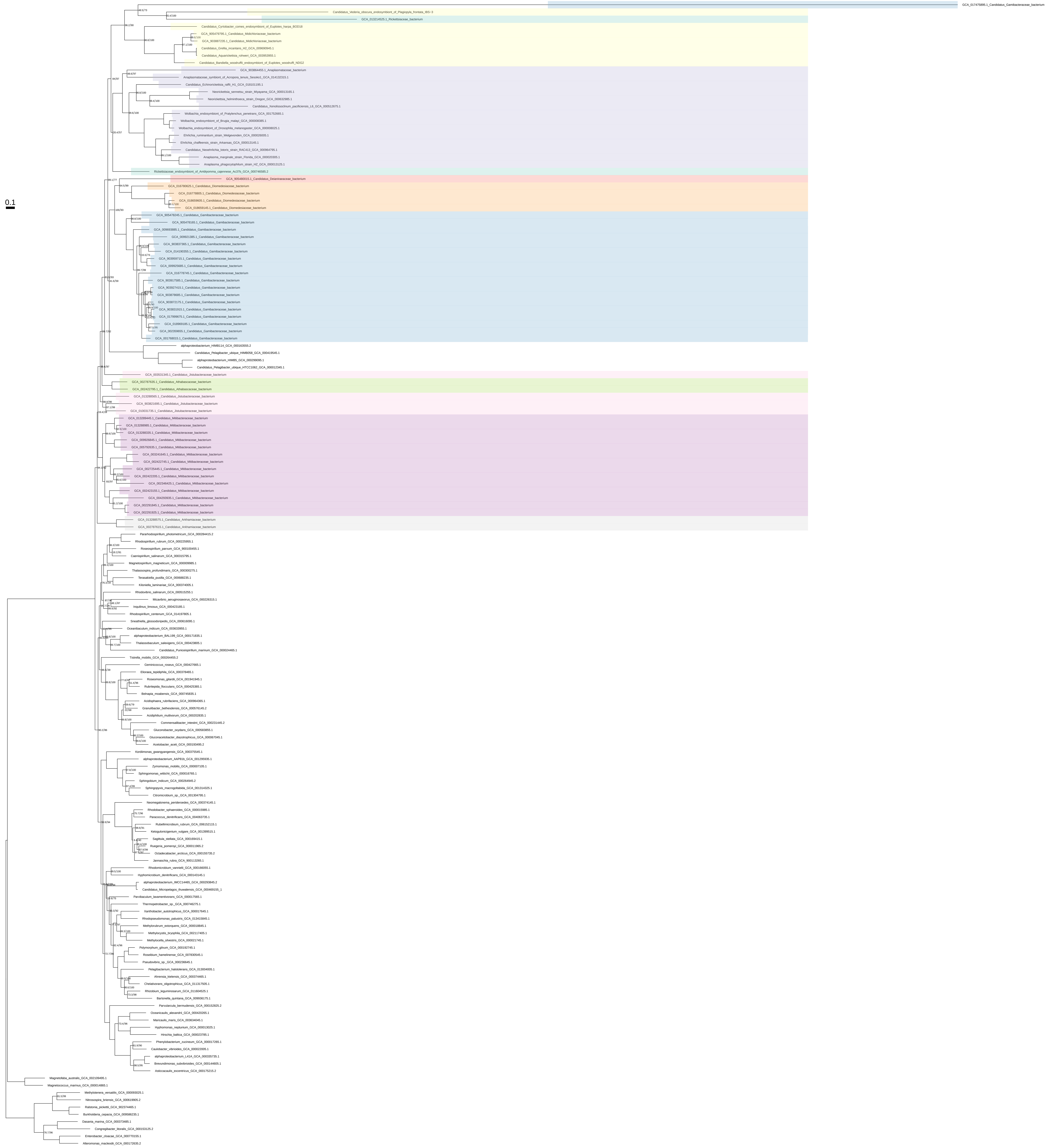

### Purines_organism_selection_z_0.30

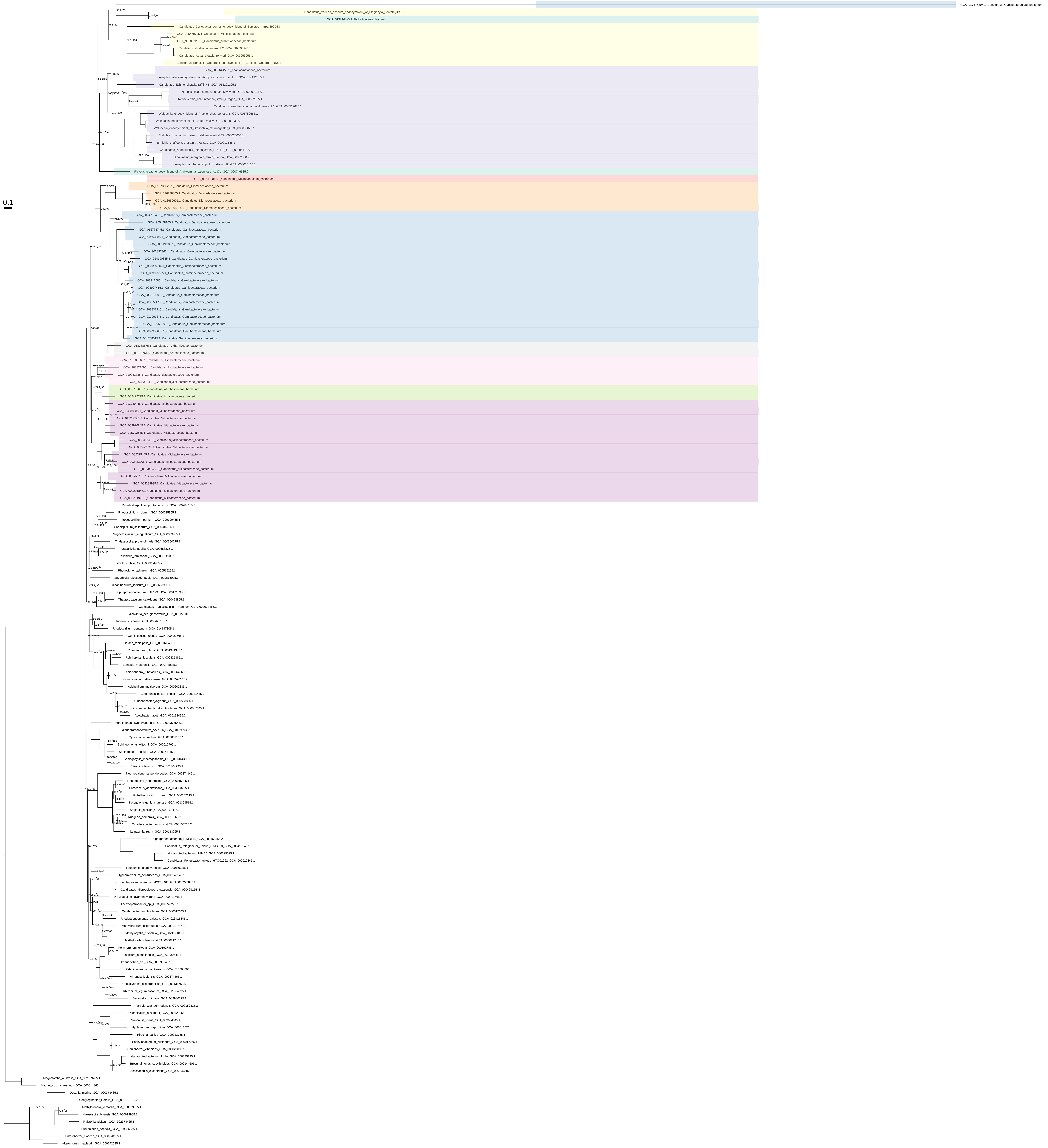

### Purines_organism_selection_z_0.40

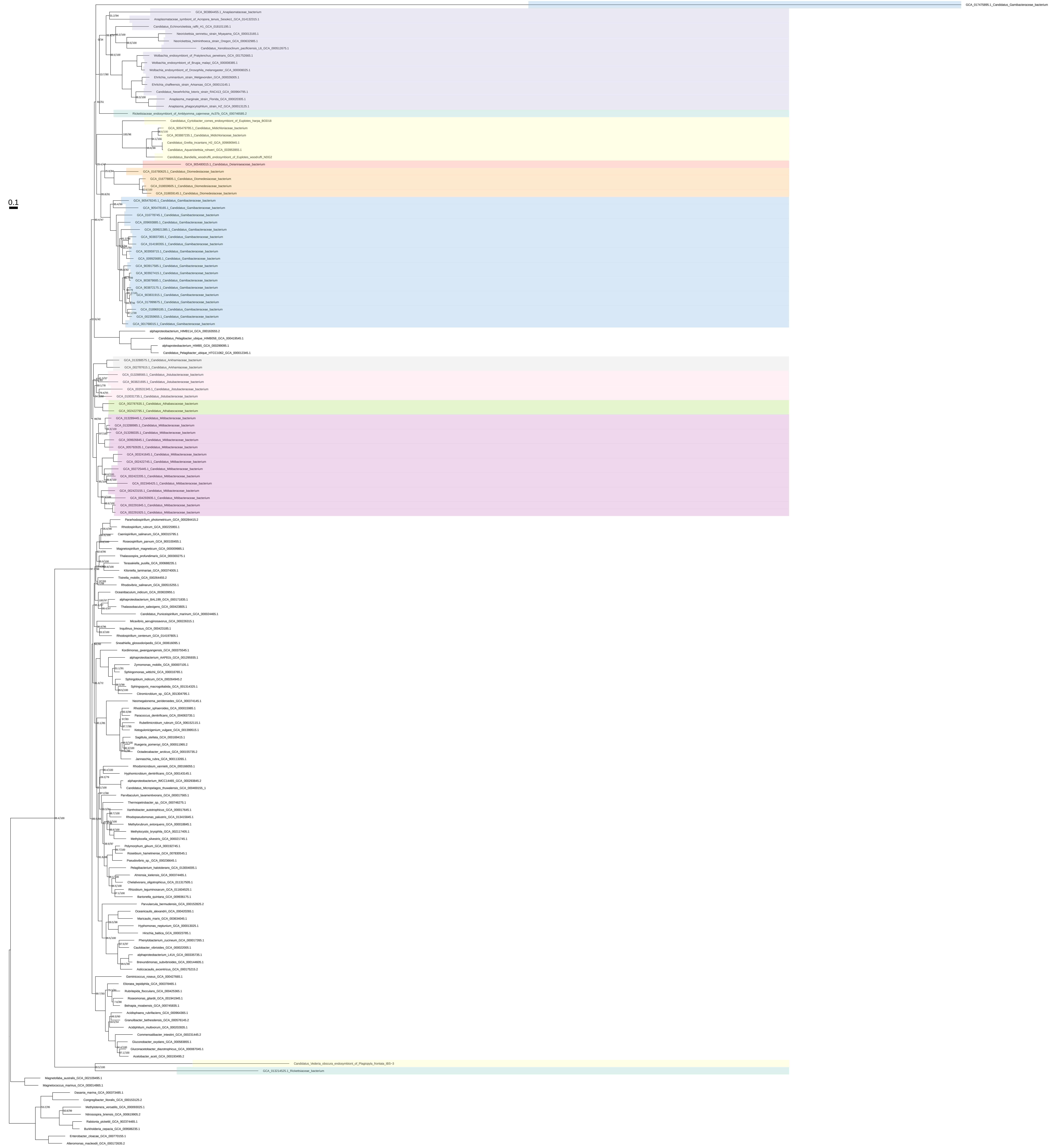

Purines_organism_selection_z_0.50

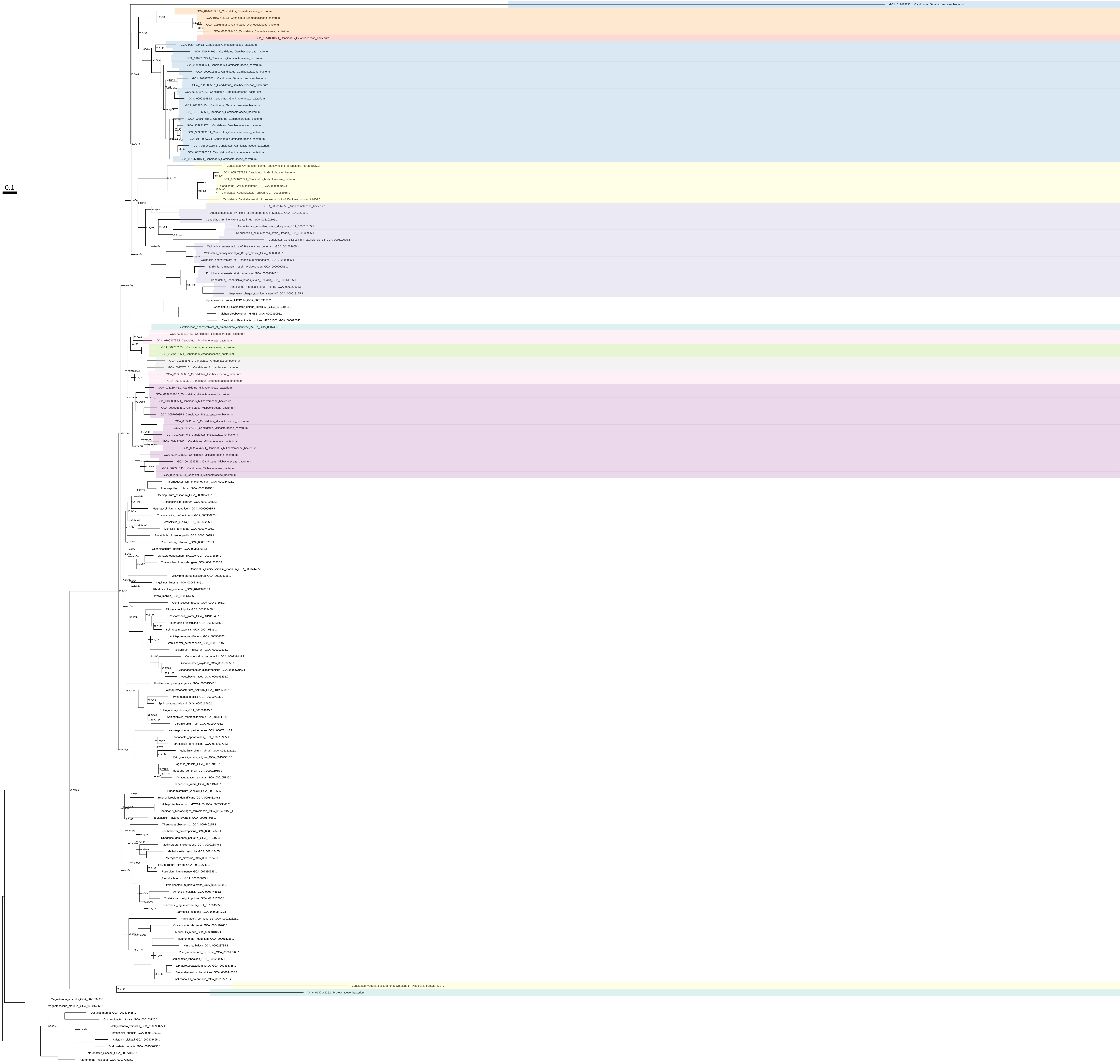

### Pyrimidines_all_organisms

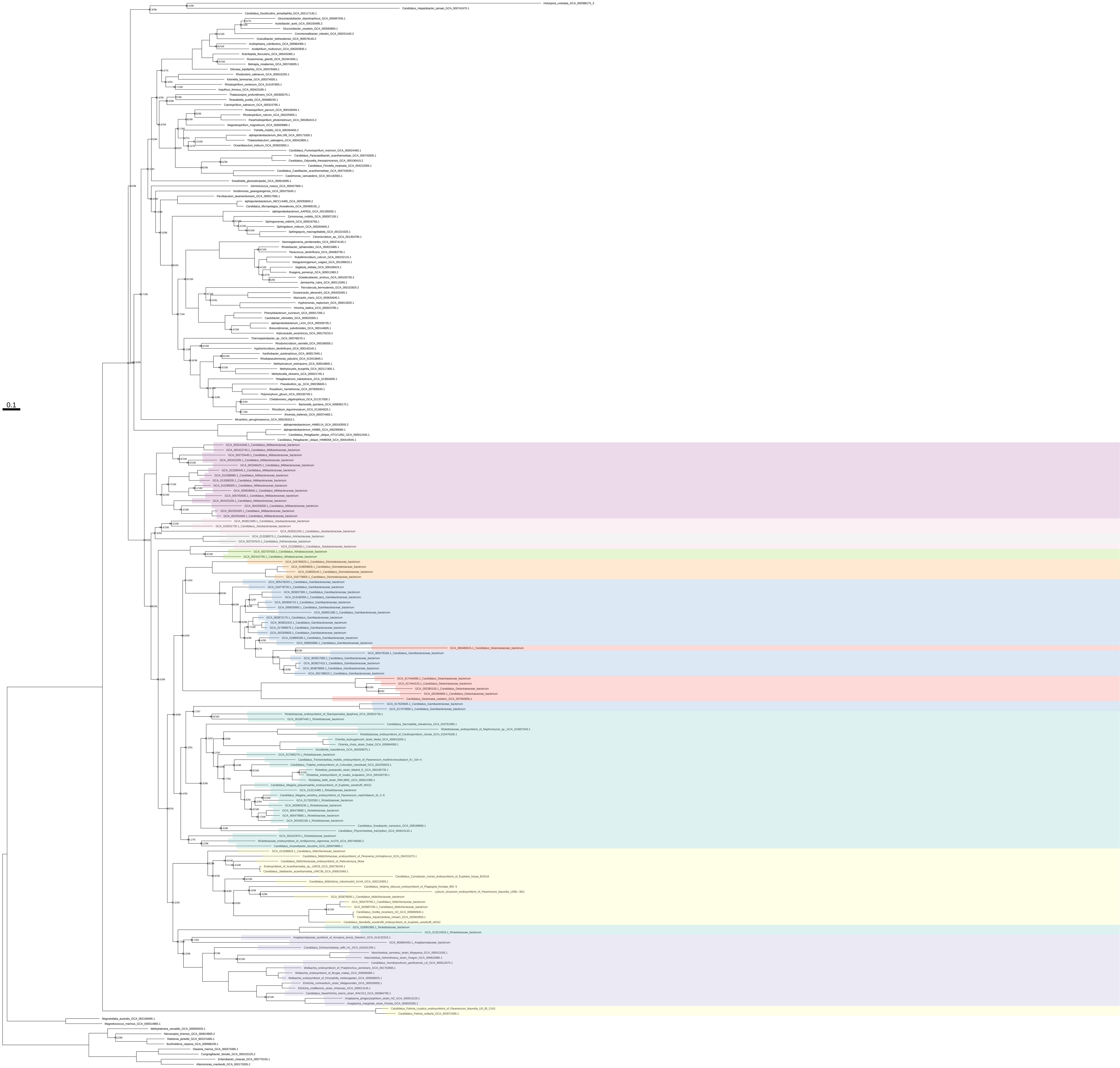

Pyrimidines_all_organisms_z_0.10

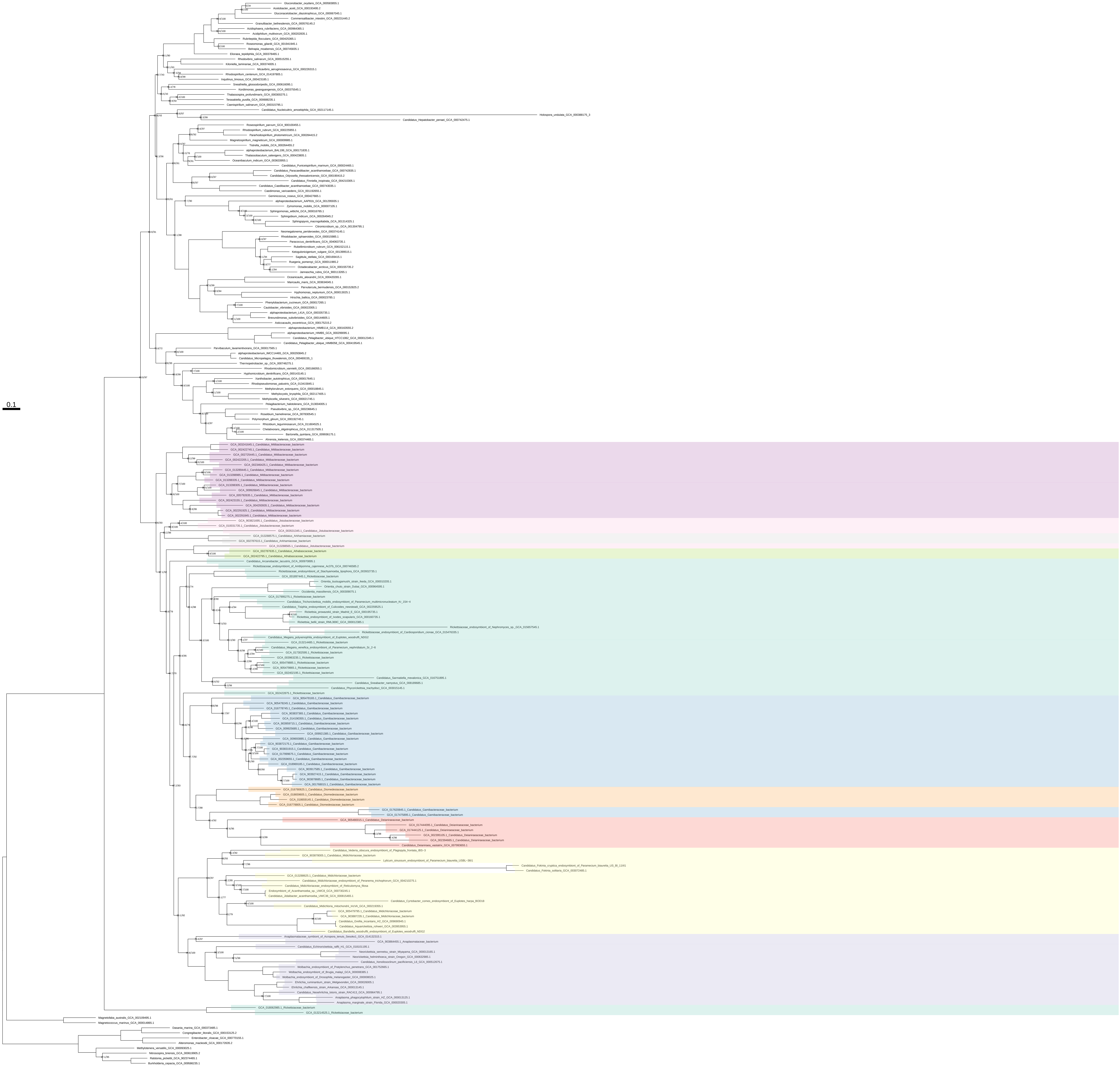

### Pyrimidines_all_organisms_z_0.20

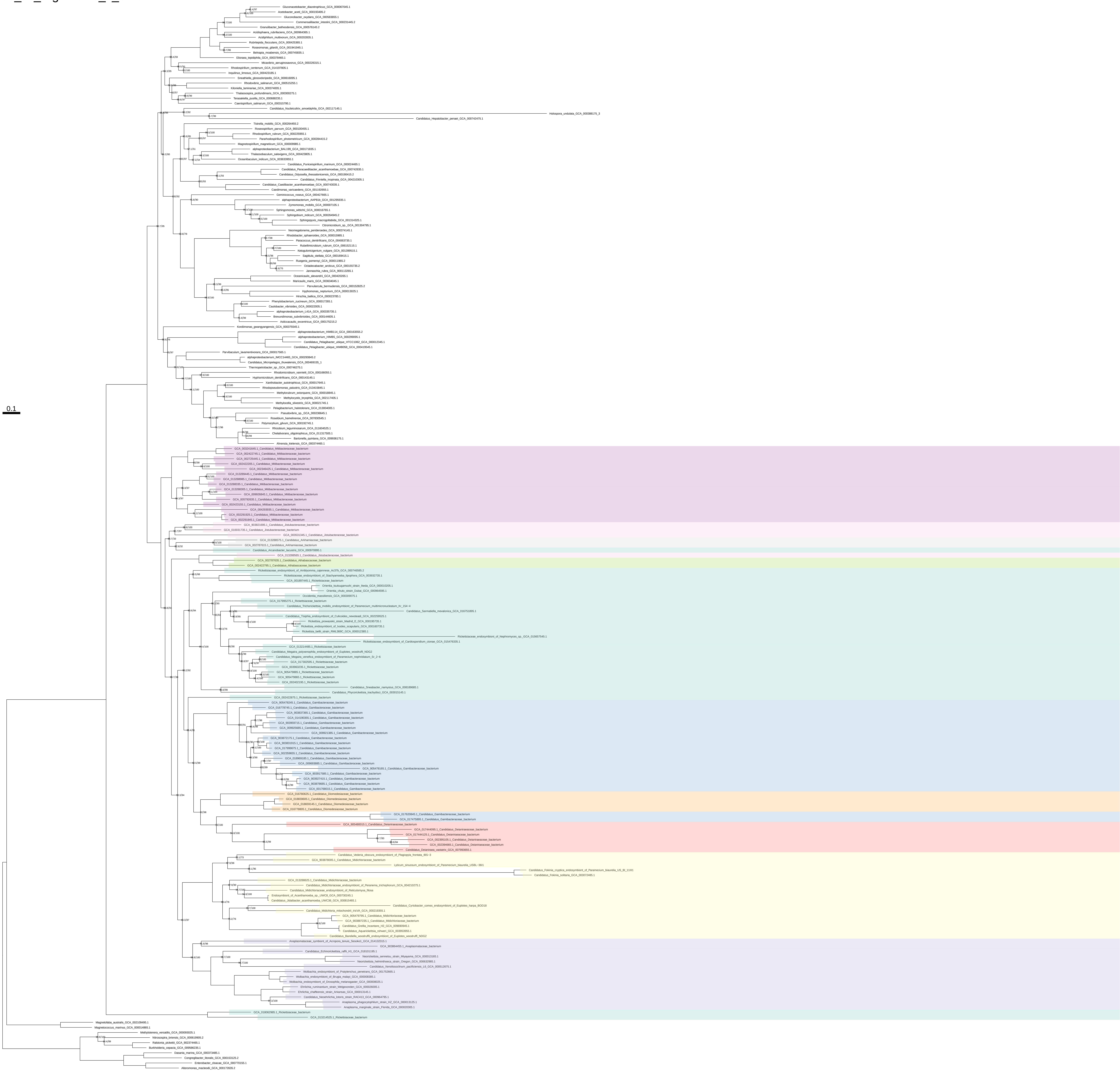

Pyrimidines_all_organisms_z_0.30

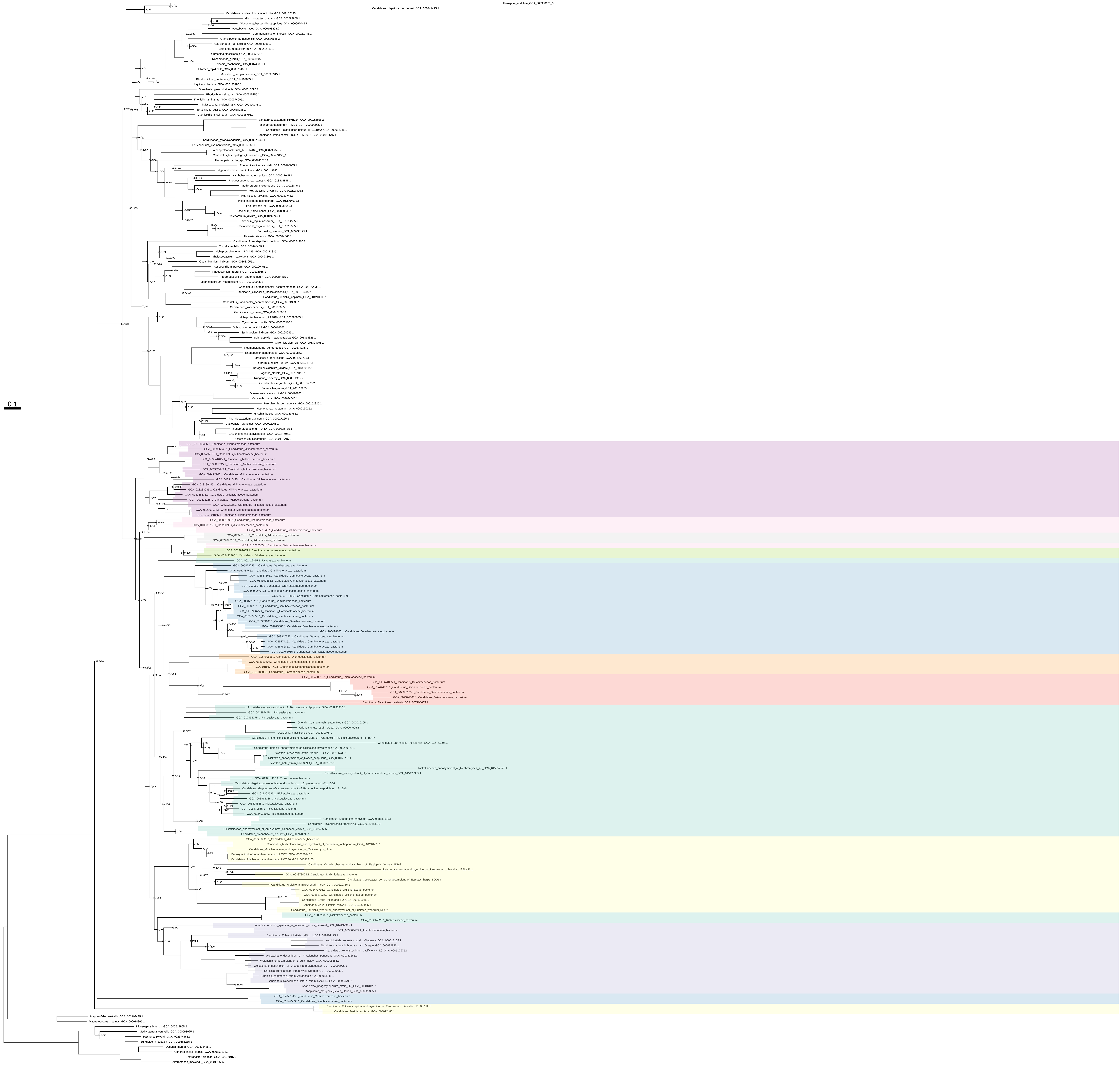

### Pyrimidines_all_organisms_z_0.40

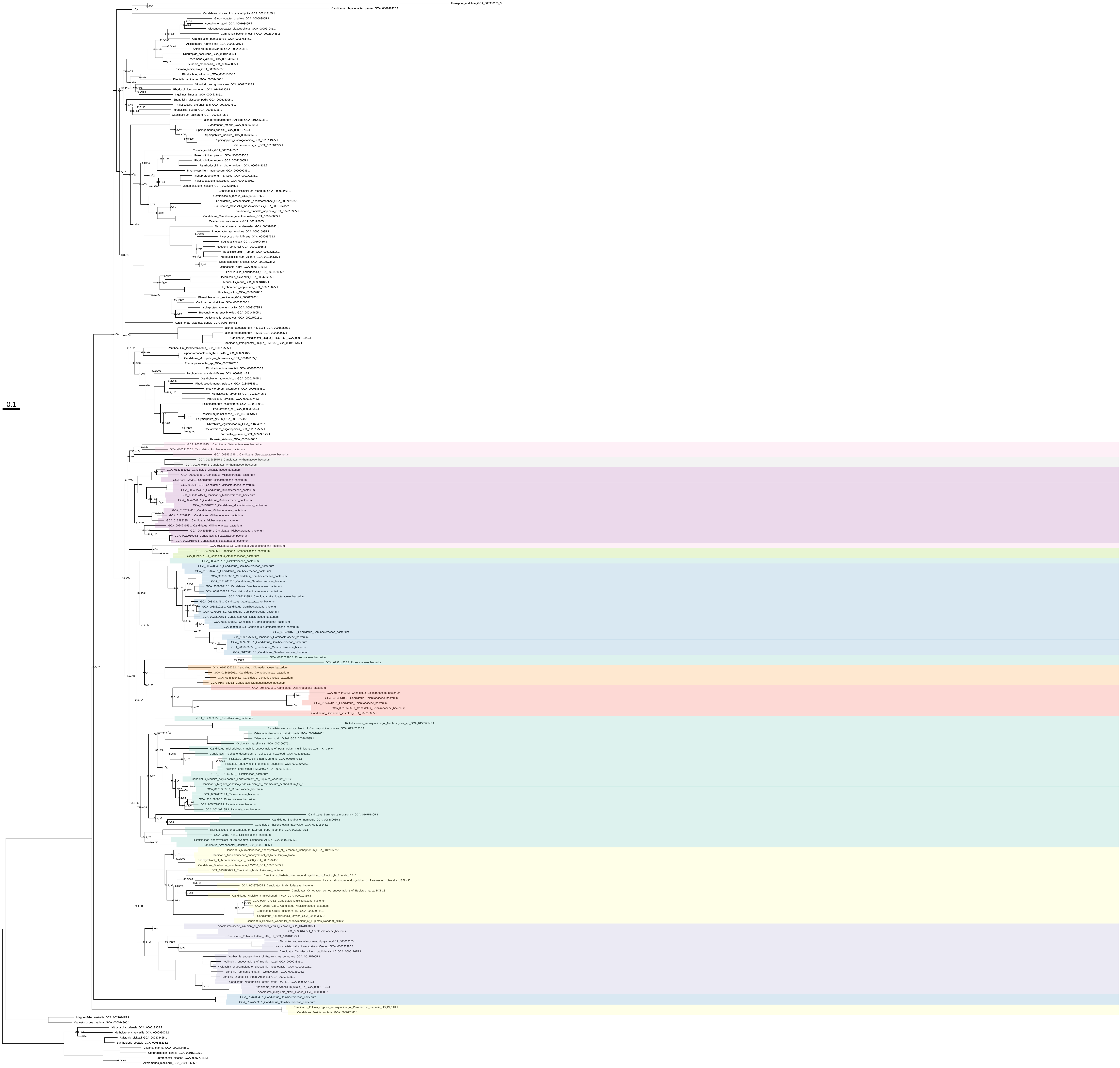

Pyrimidines_all_organisms_z_0.50

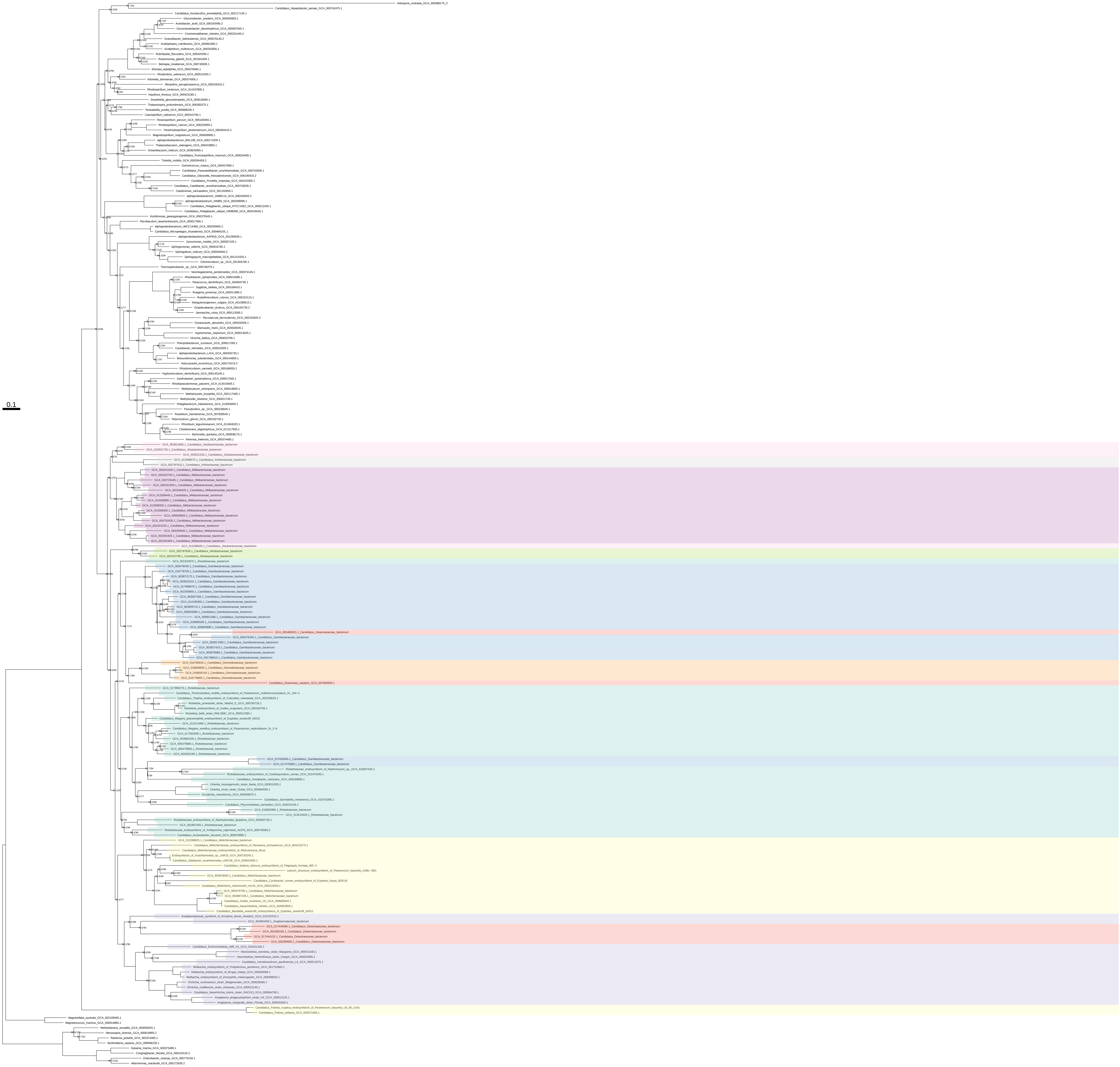

### Pyrimidines_organism_selection

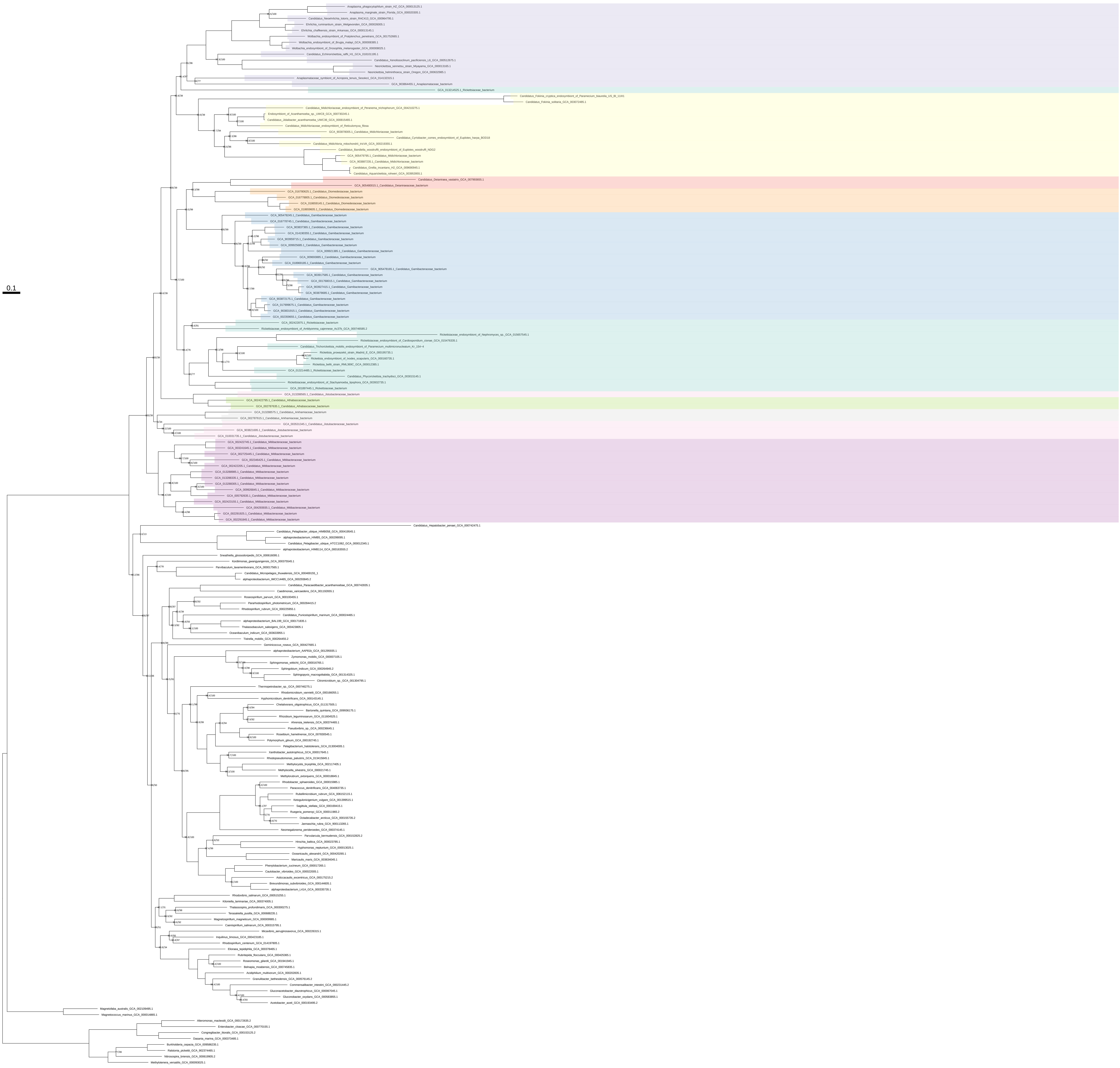

### Pyrimidines_organism_selection_z_0.10

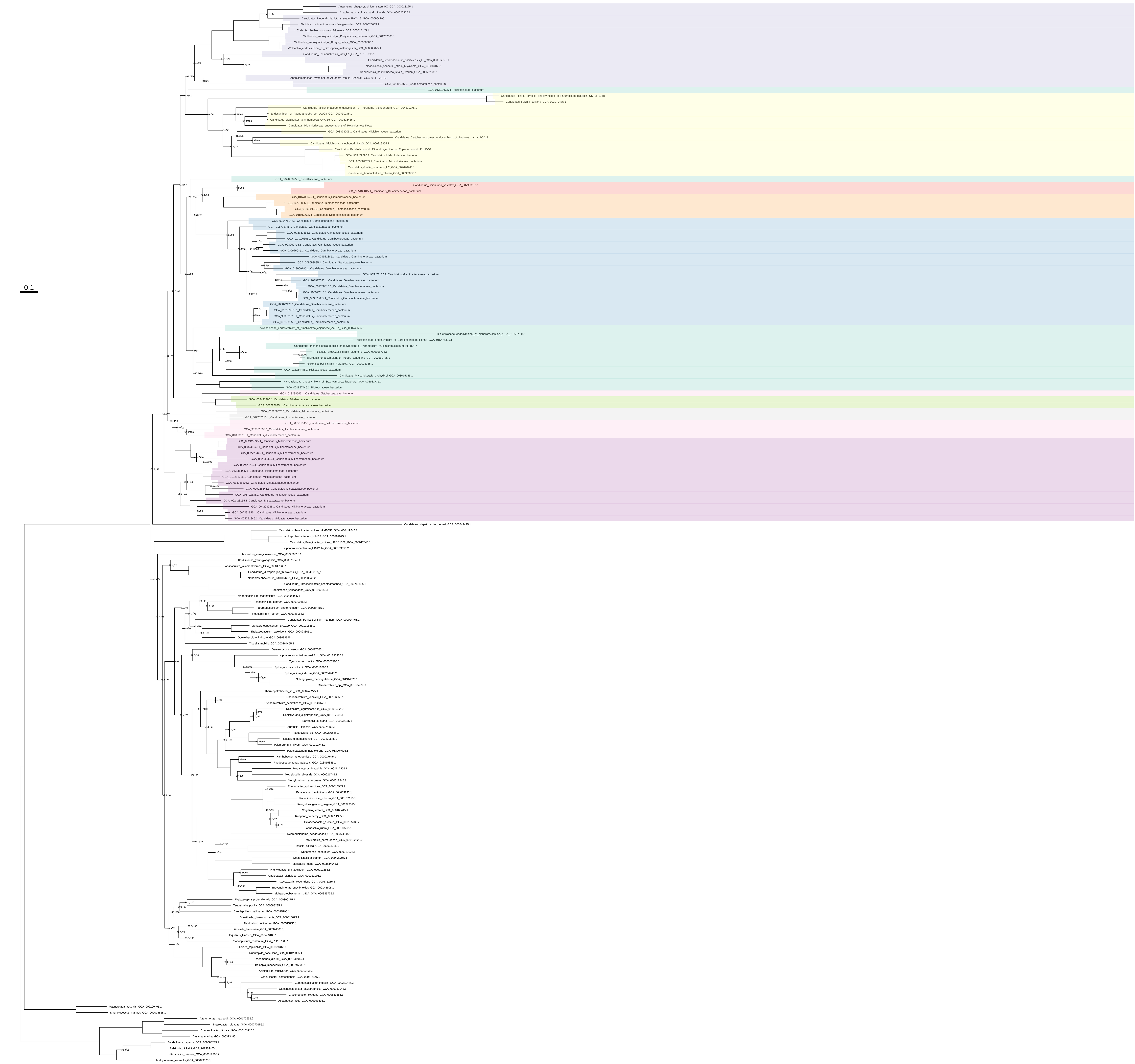

Pyrimidines_organism_selection_z_0.20

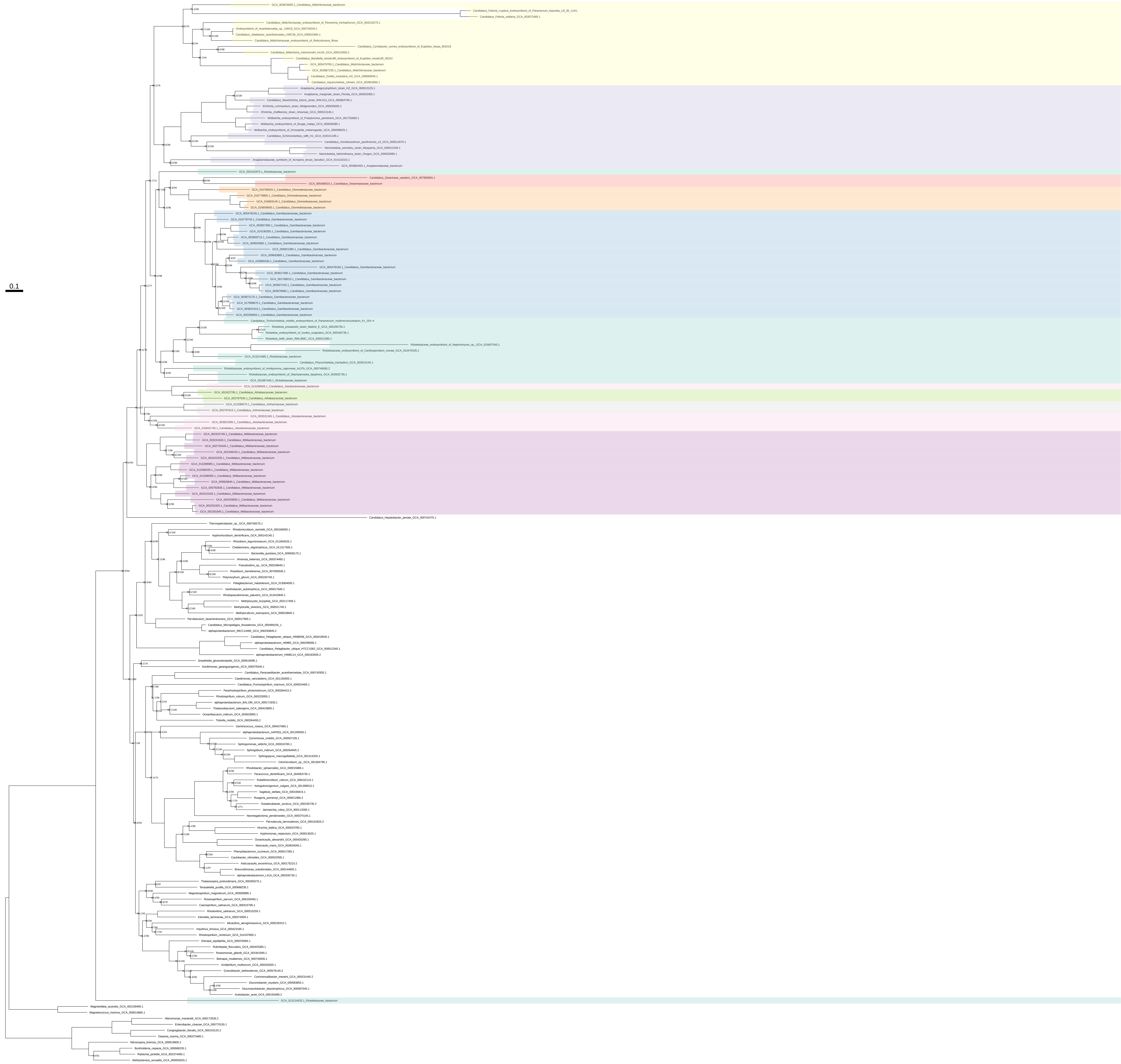

Pyrimidines_organism_selection_z_0.30

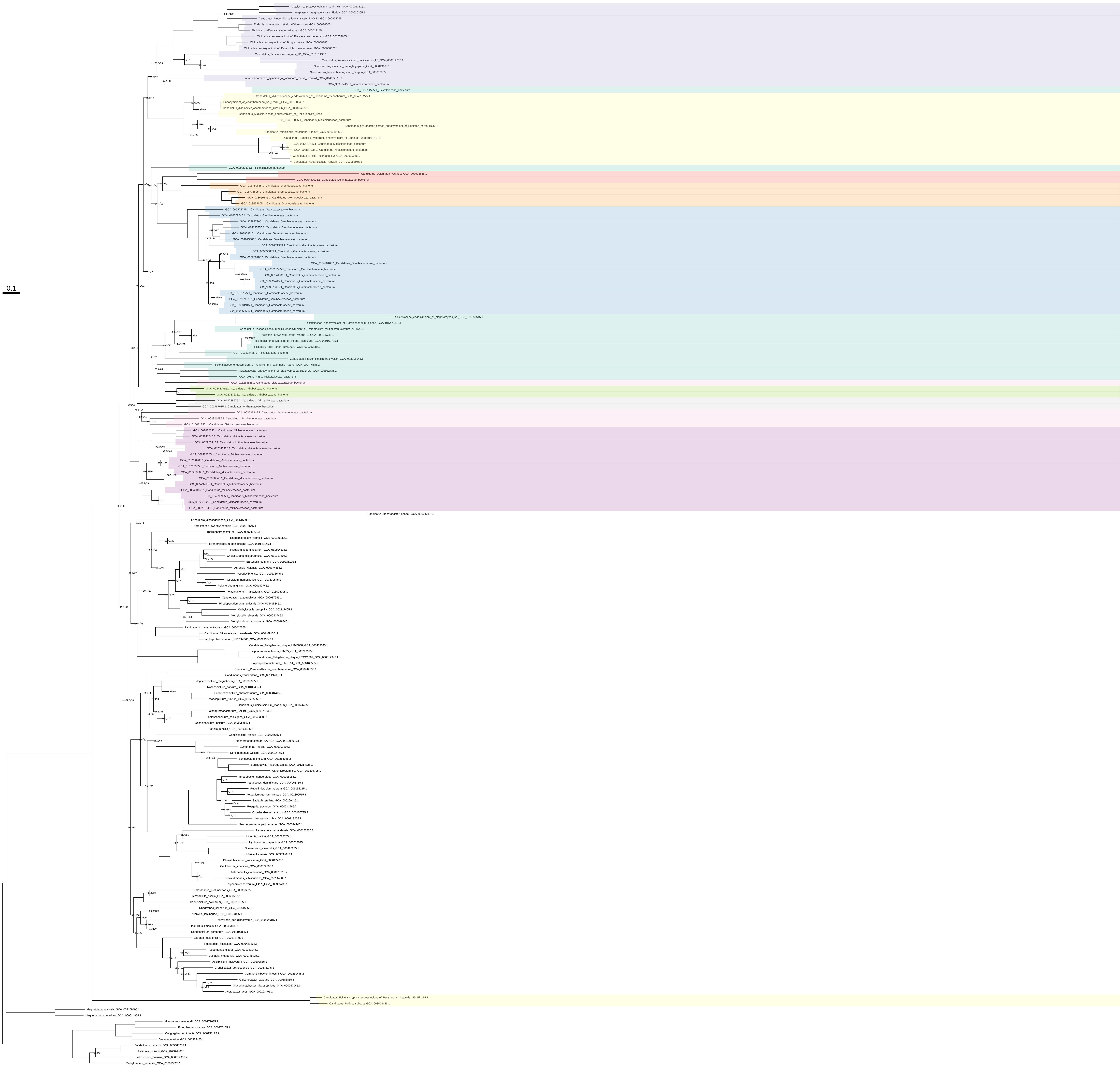

### Pyrimidines_organism_selection_z_0.40

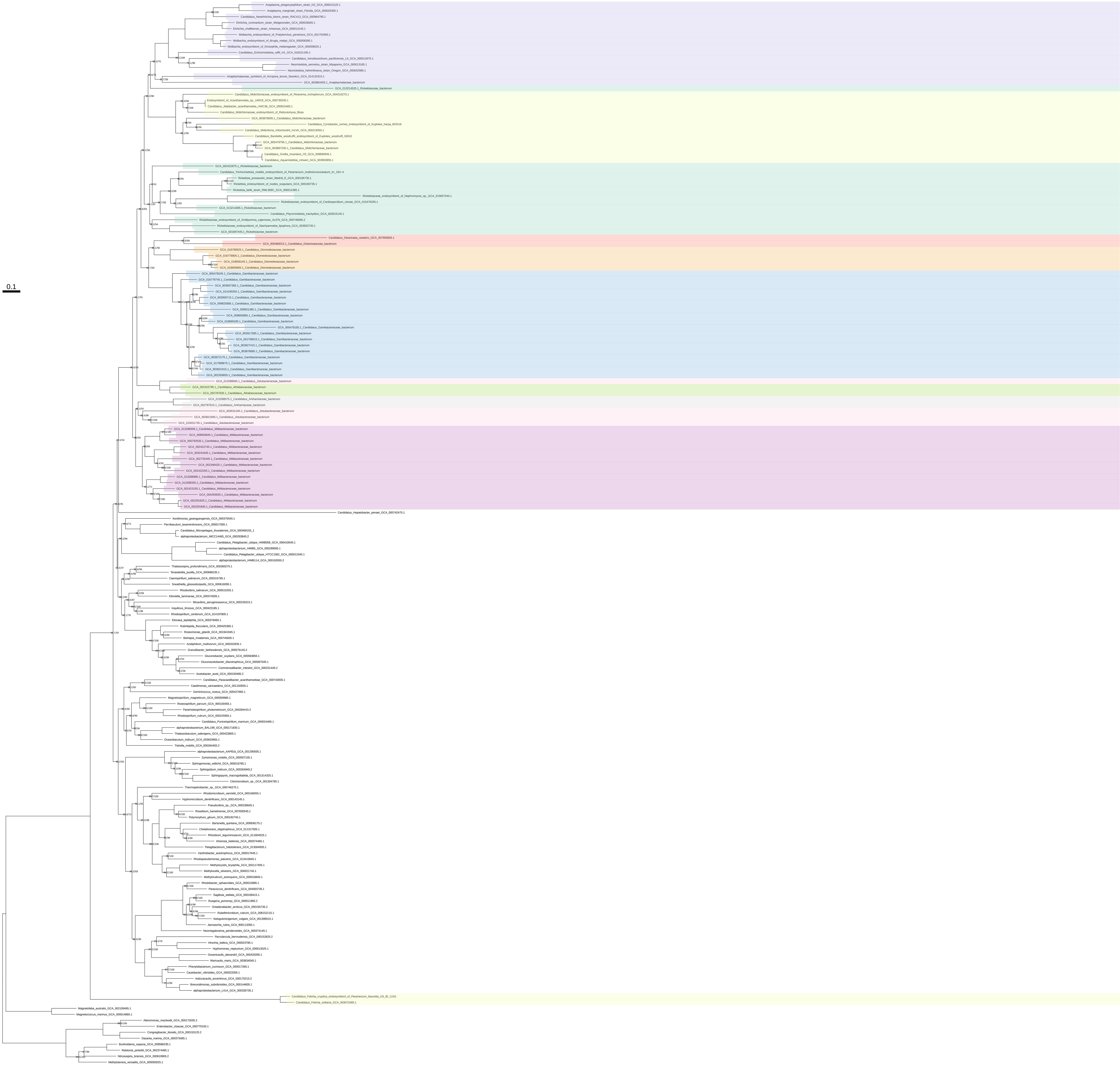

Pyrimidines_organism_selection_z_0.50

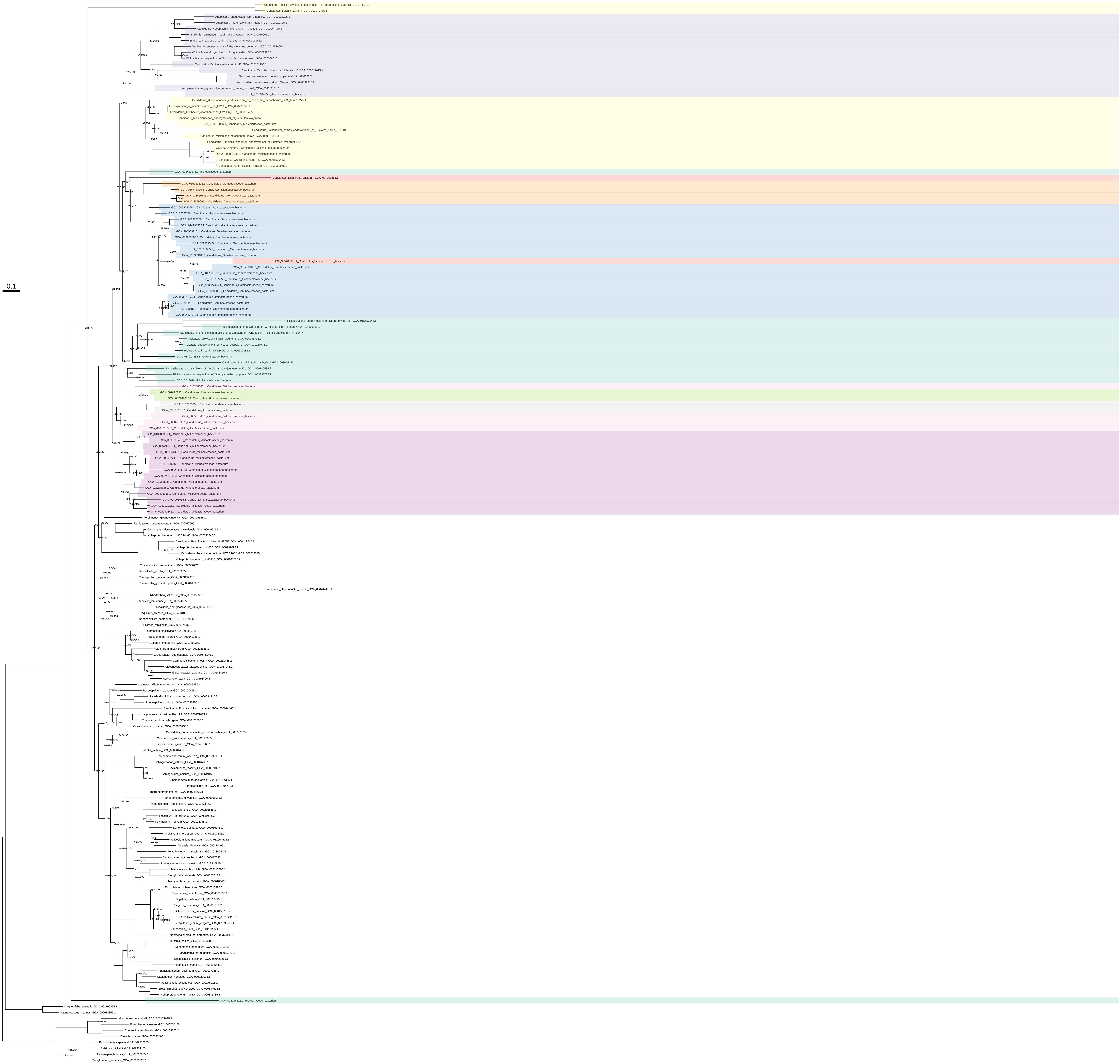

Arg_all_organisms

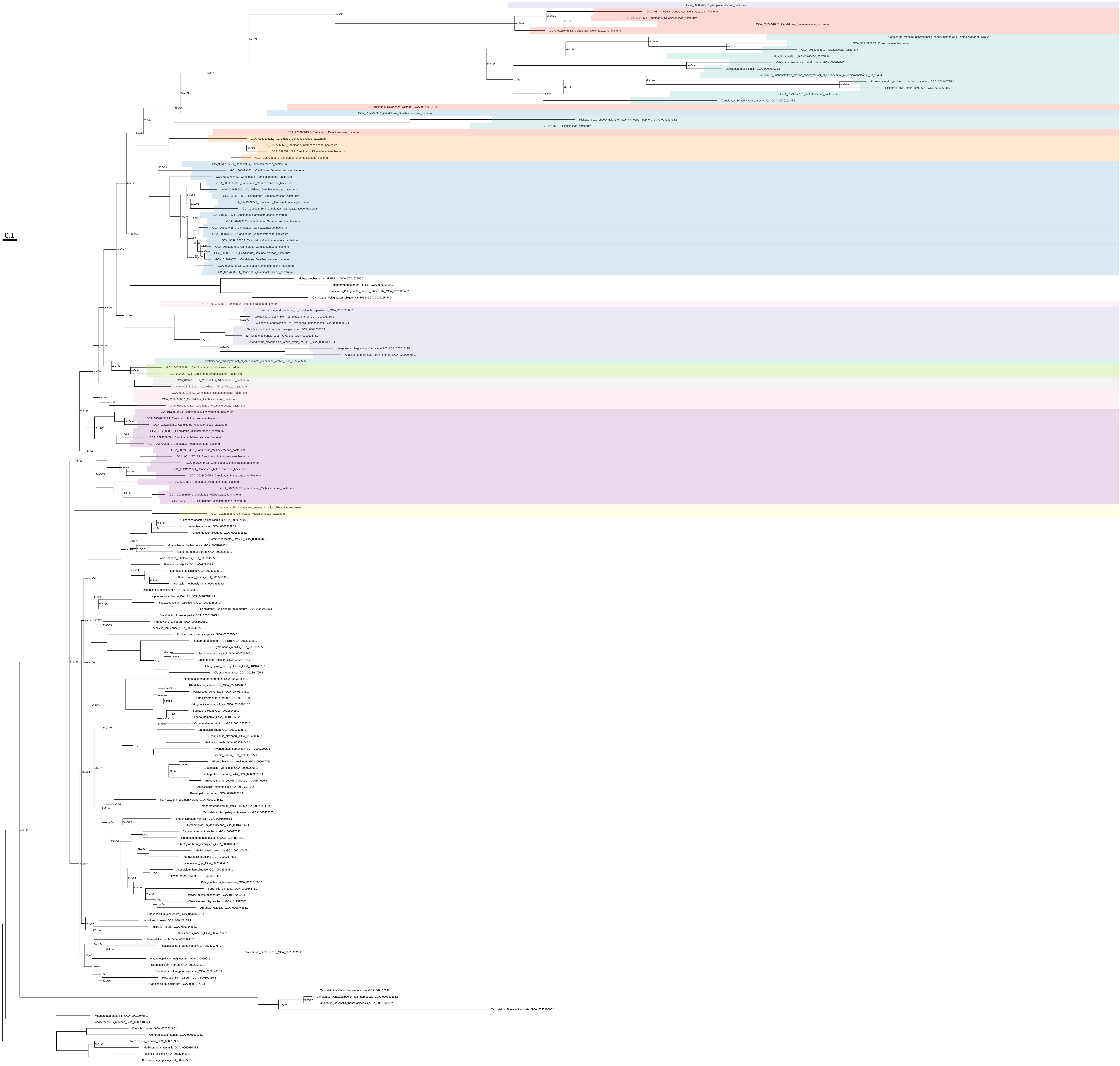

### Arg_all_organisms_z_0.10

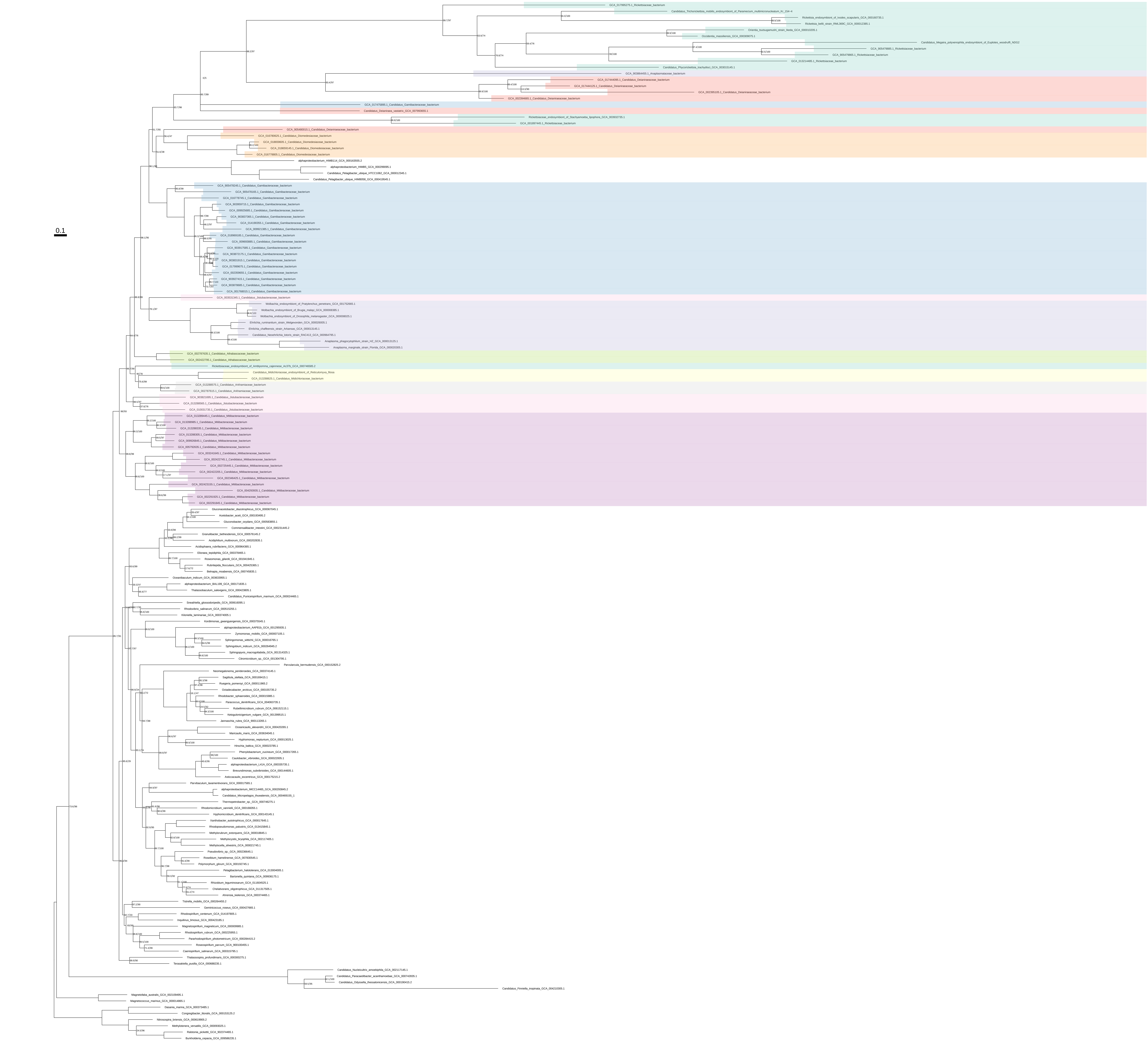

Arg_all_organisms_z_0.20

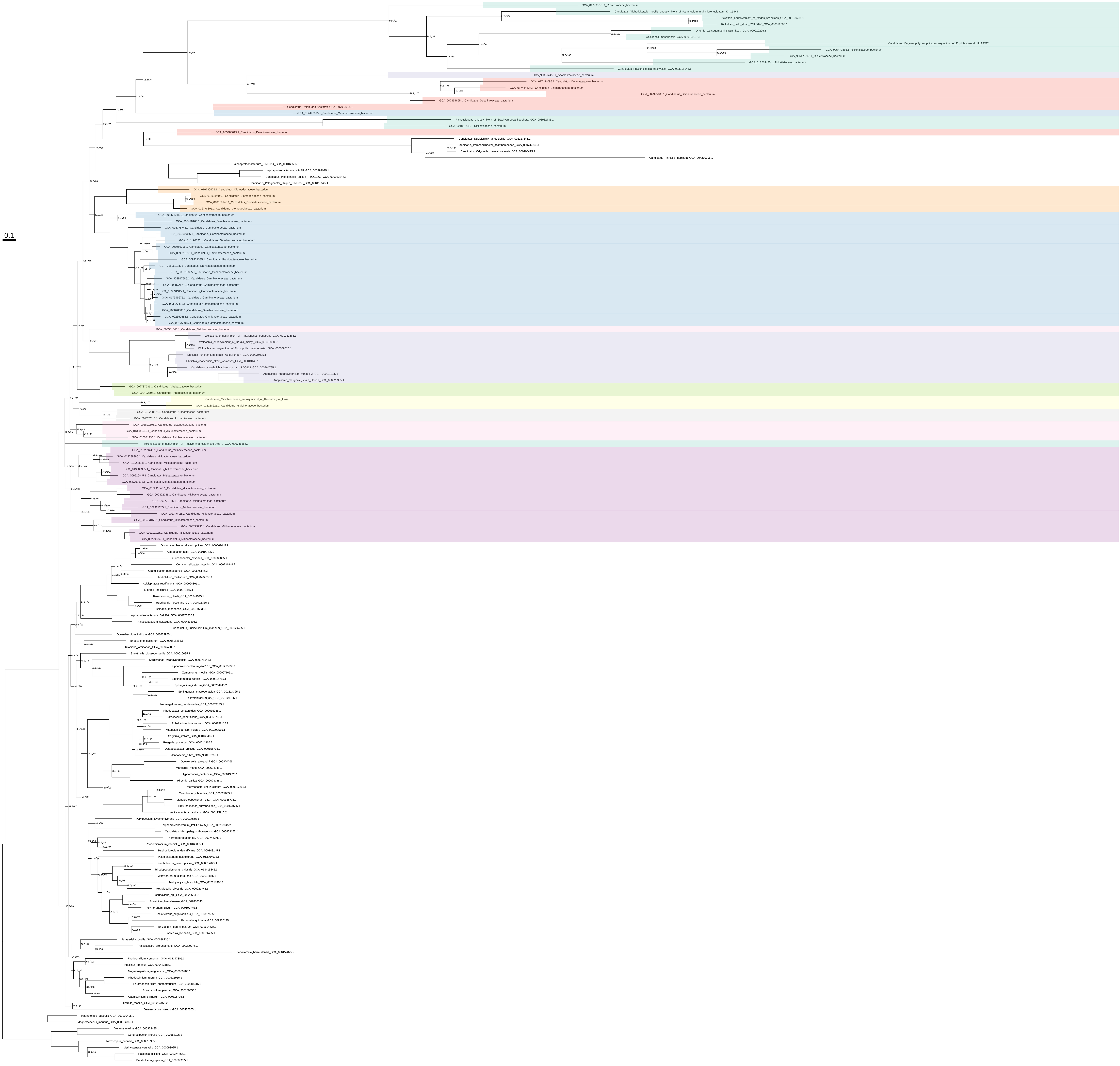

Arg_all_organisms_z_0.30

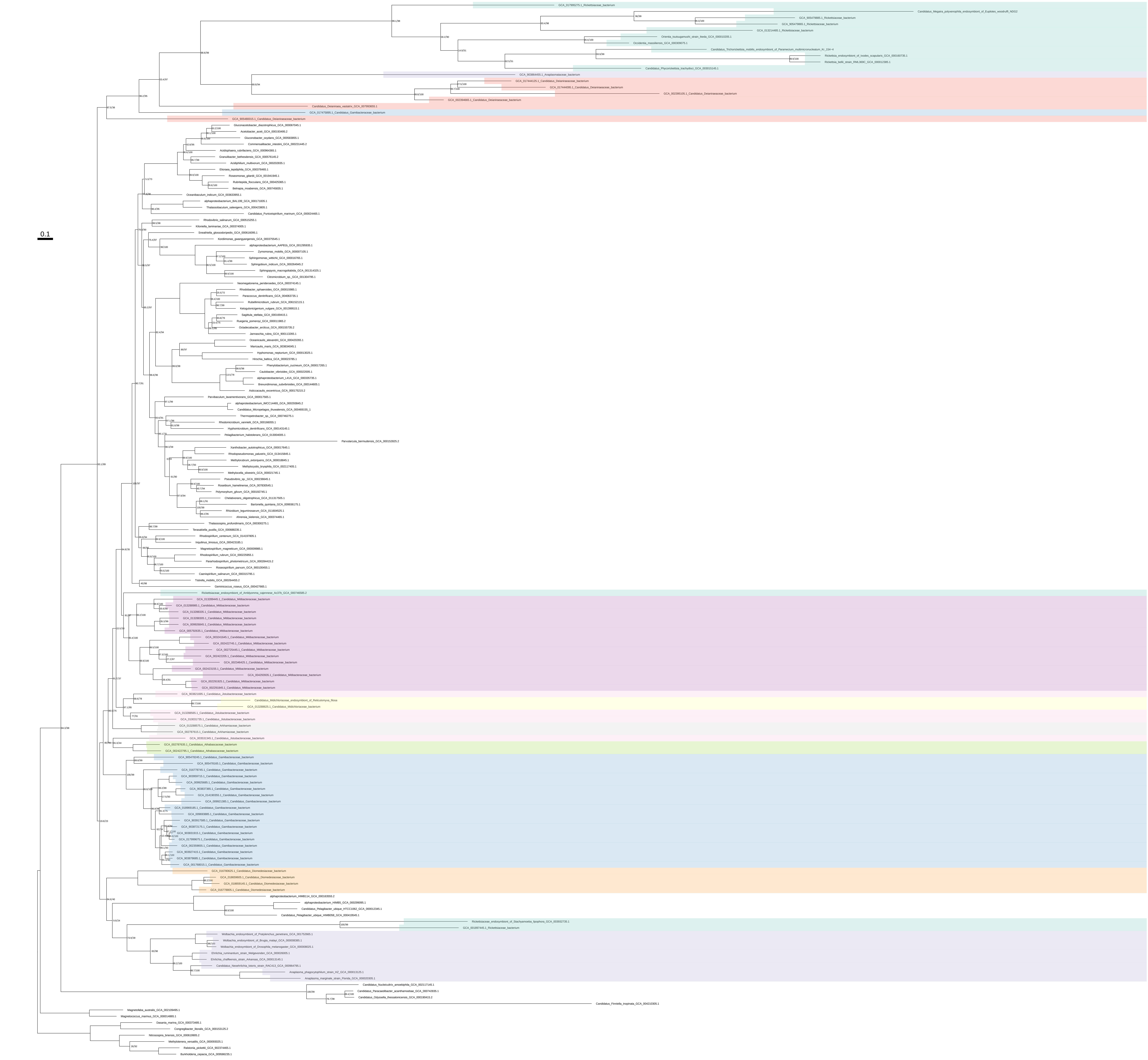

### Arg_all_organisms_z_0.40

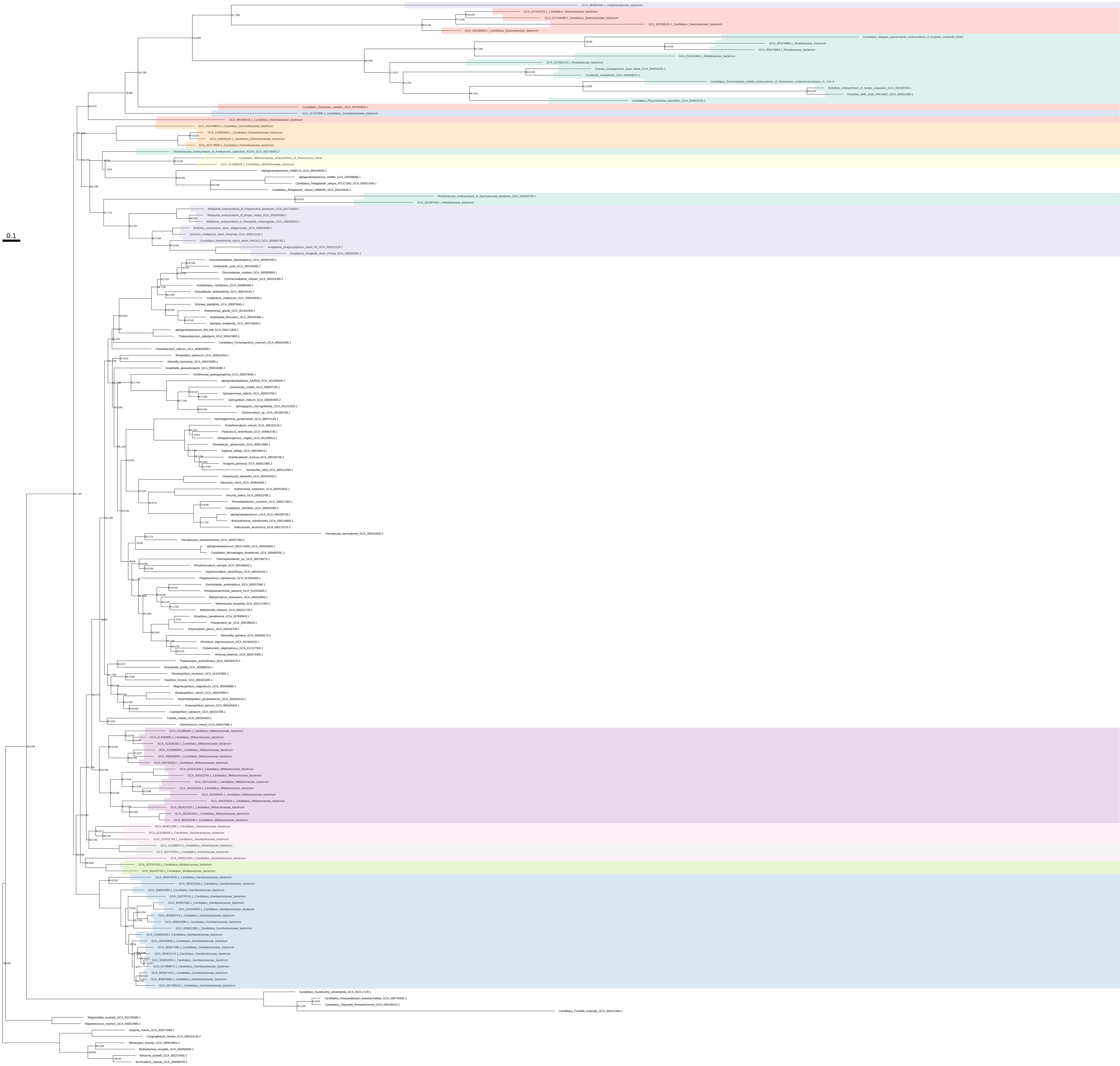

Arg_all_organisms_z_0.50

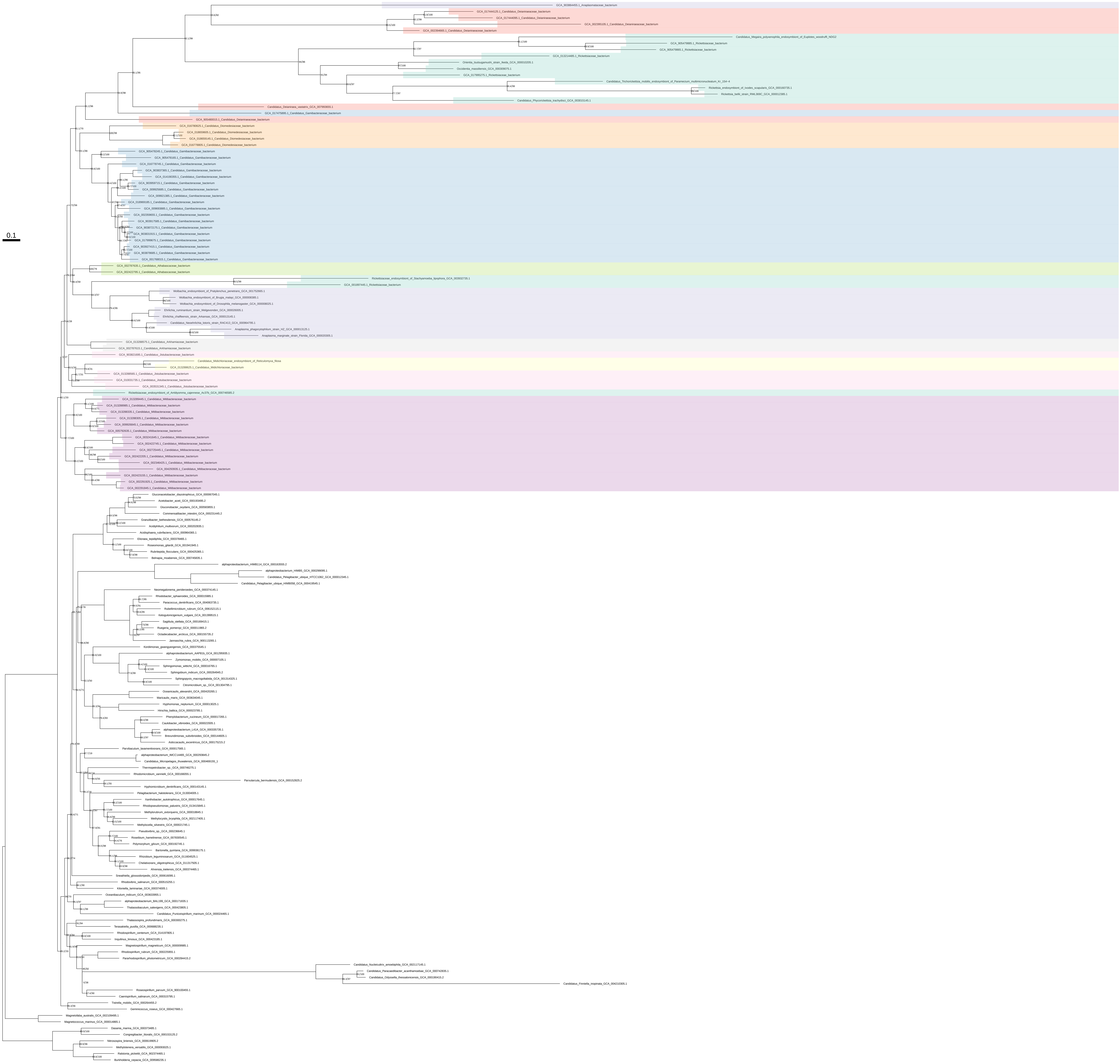

### Arg_organism_selection

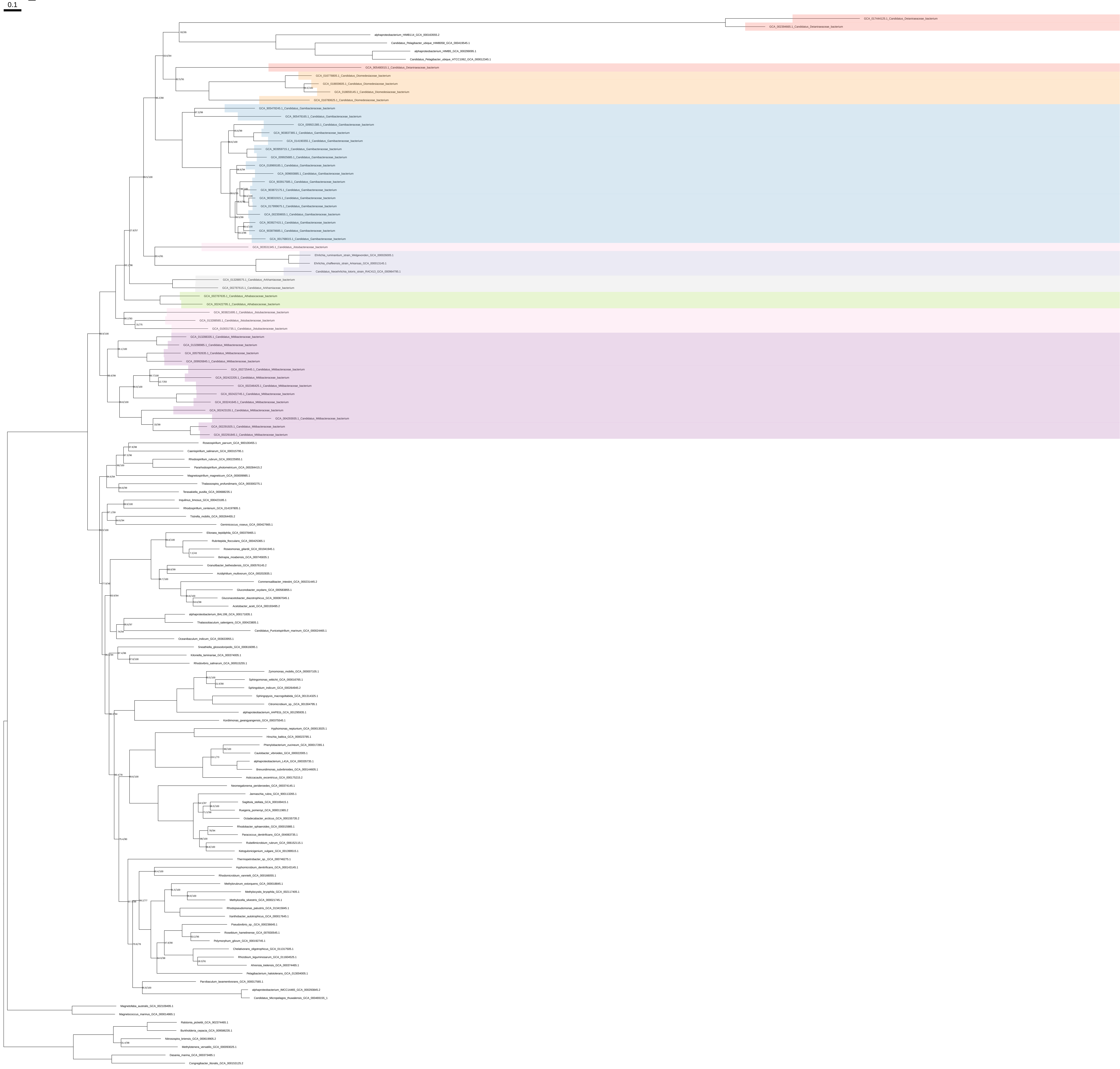

### Arg_organism_selection_z_0.10

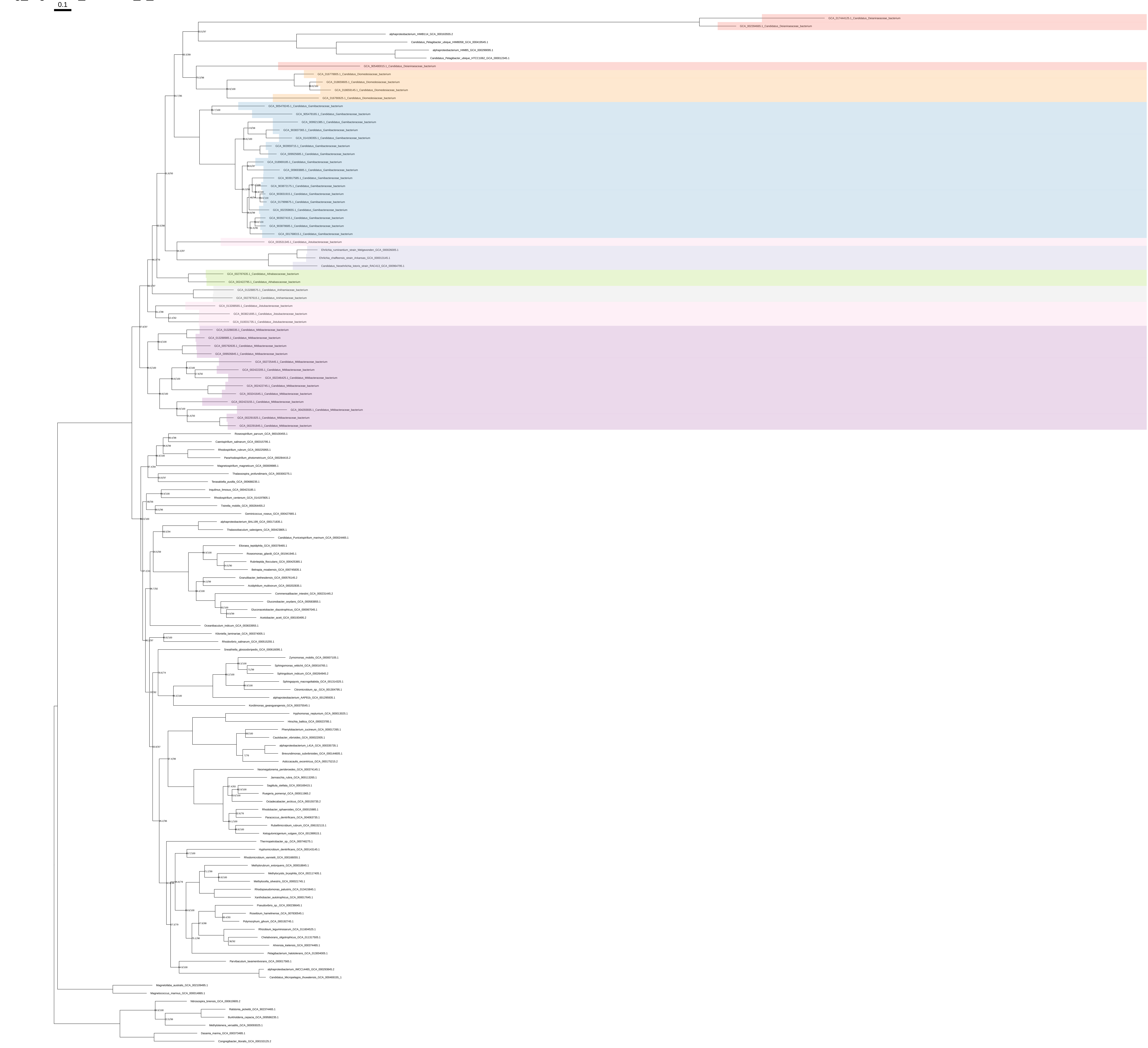

### Arg_organism_selection_z_0.20

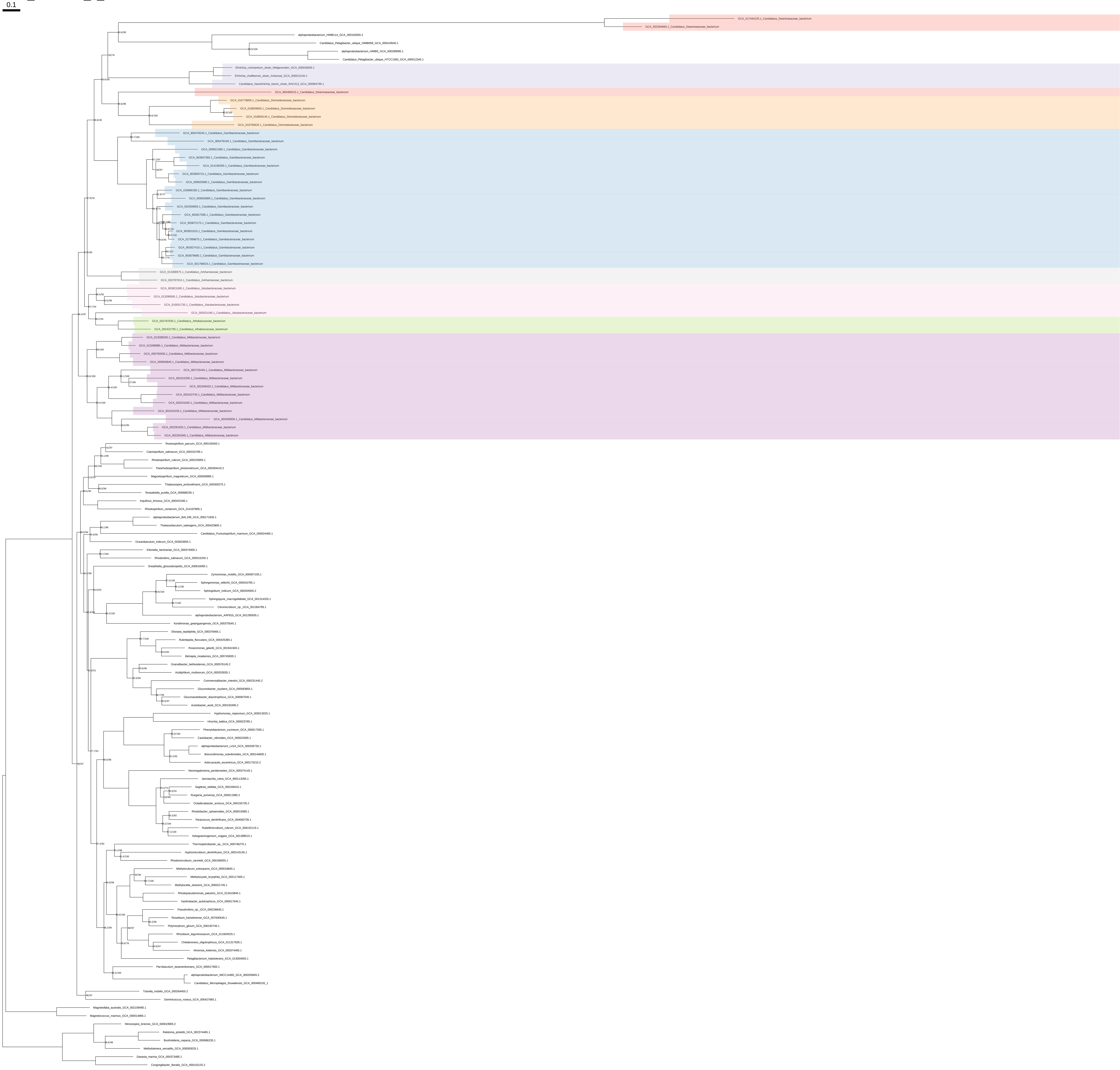

### Arg_organism_selection_z_0.30

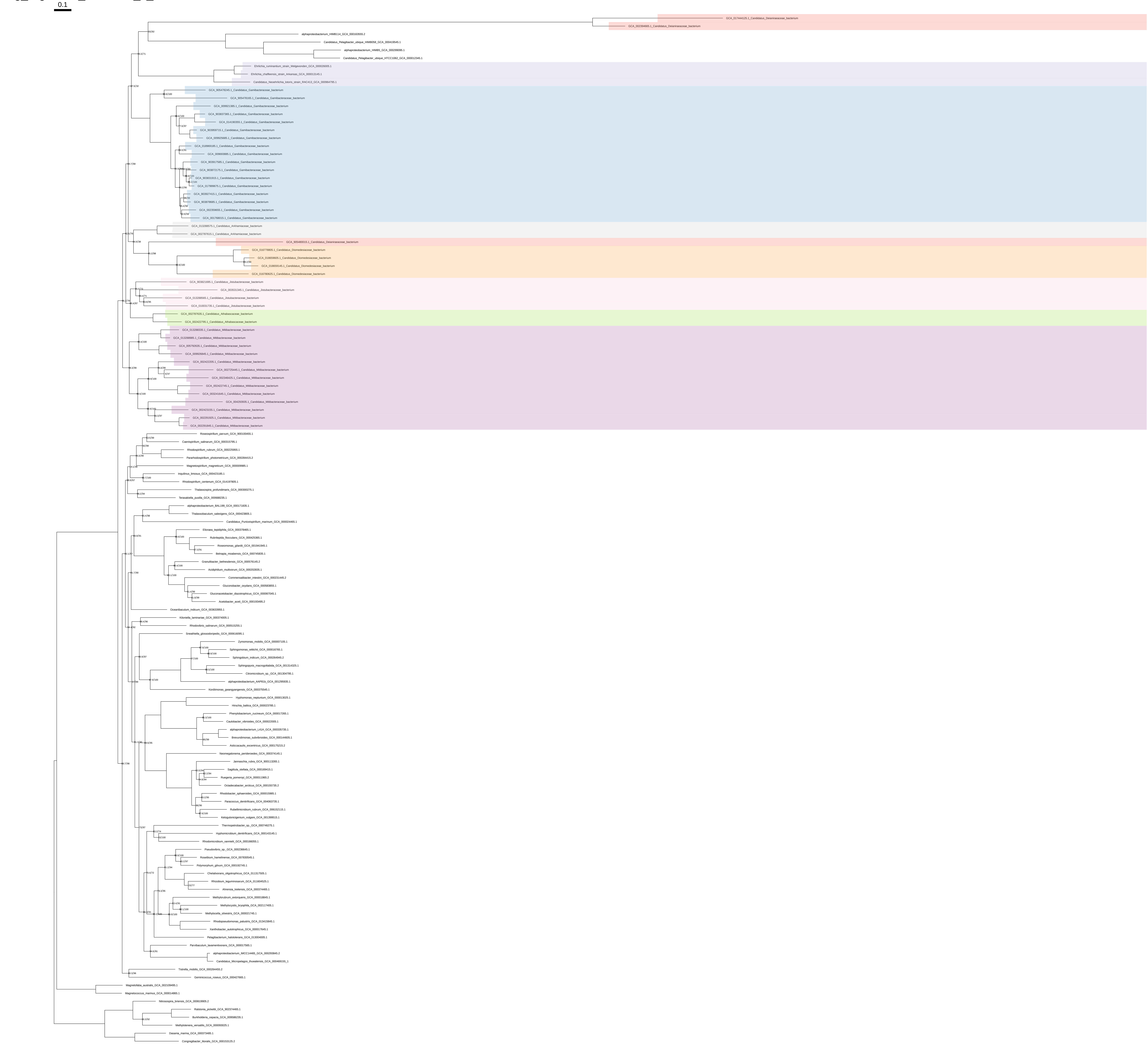

### Arg_organism_selection_z_0.40

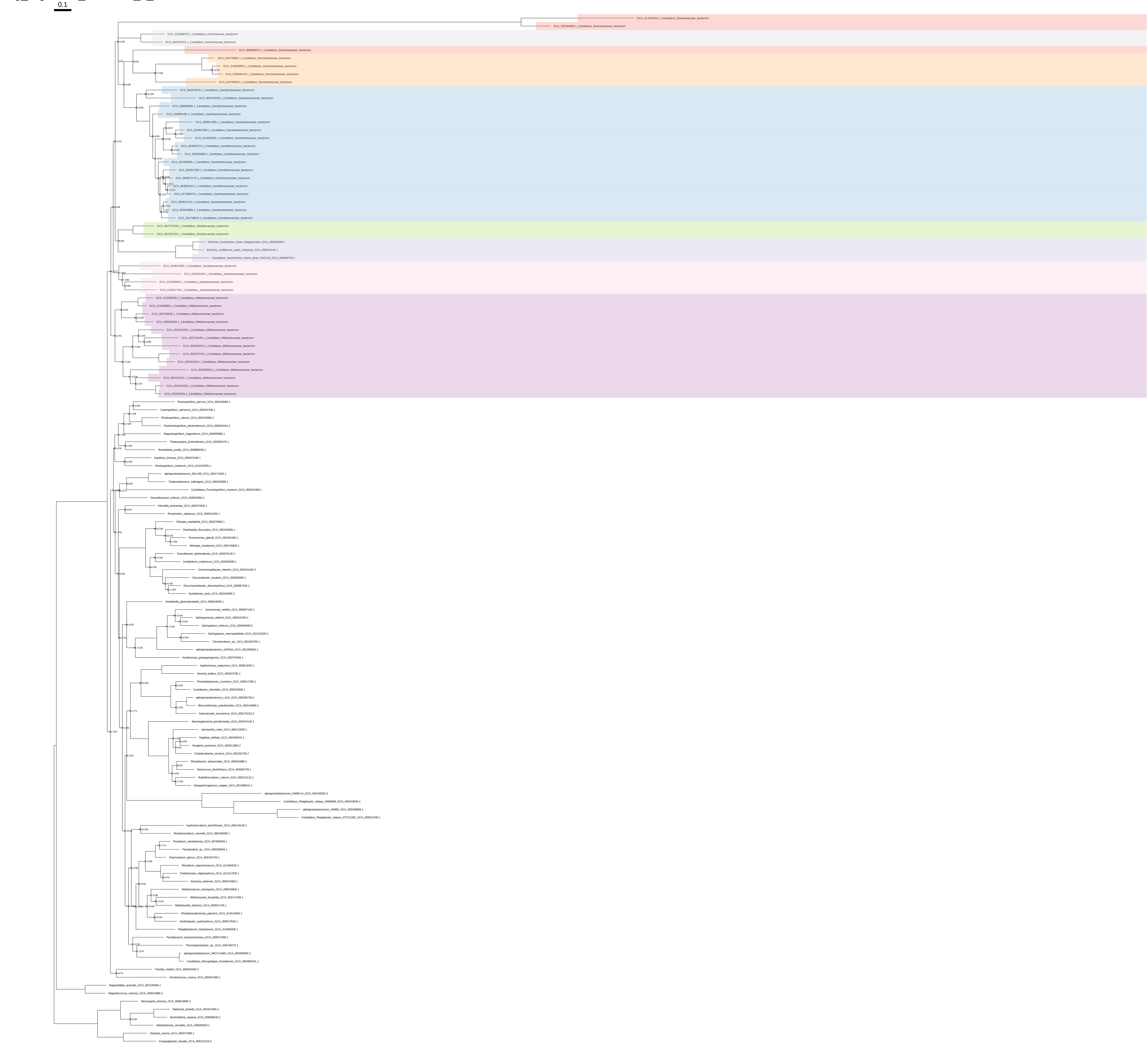

### Arg_organism_selection_z_0.50

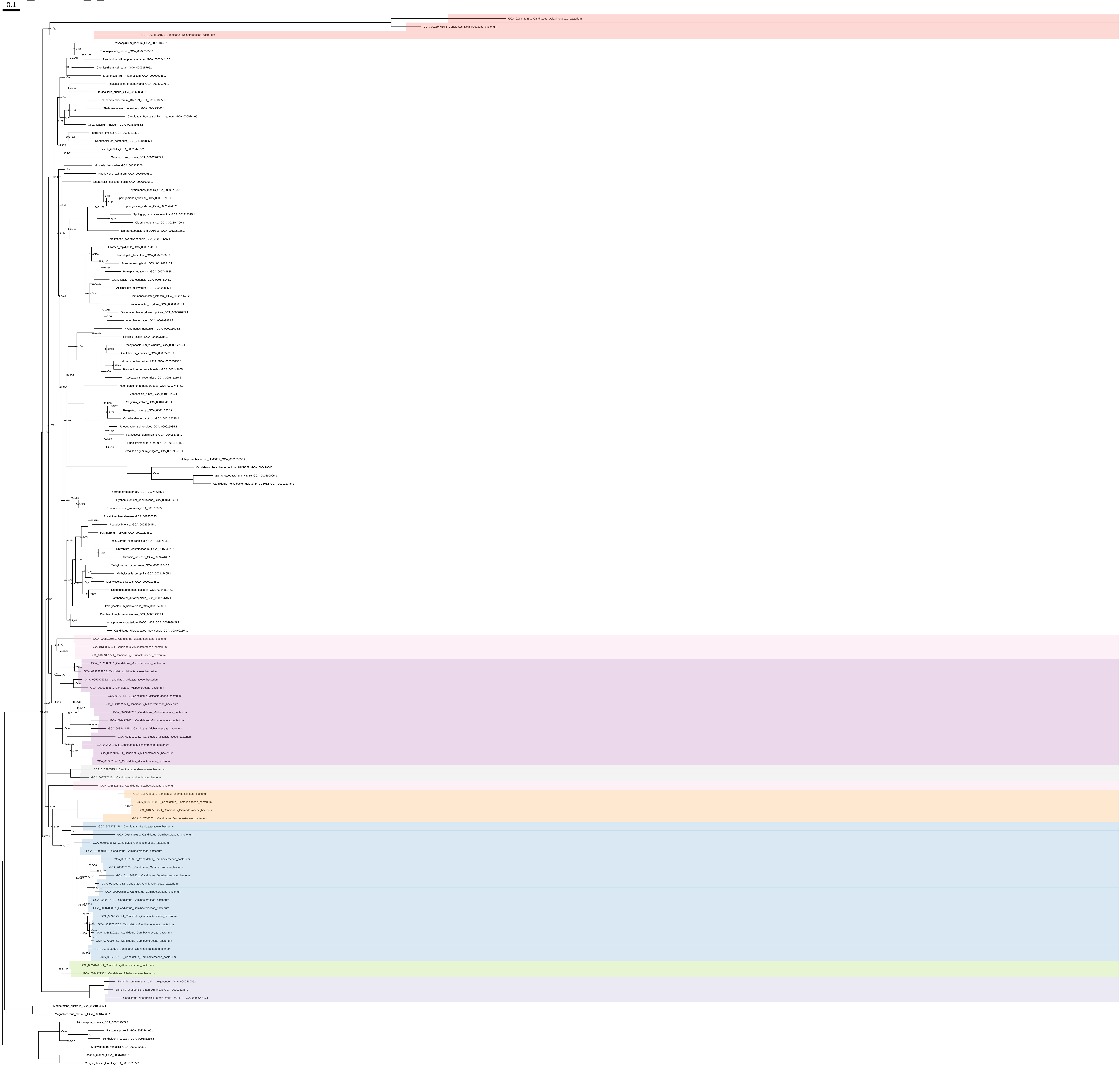

### Cys_all_organisms

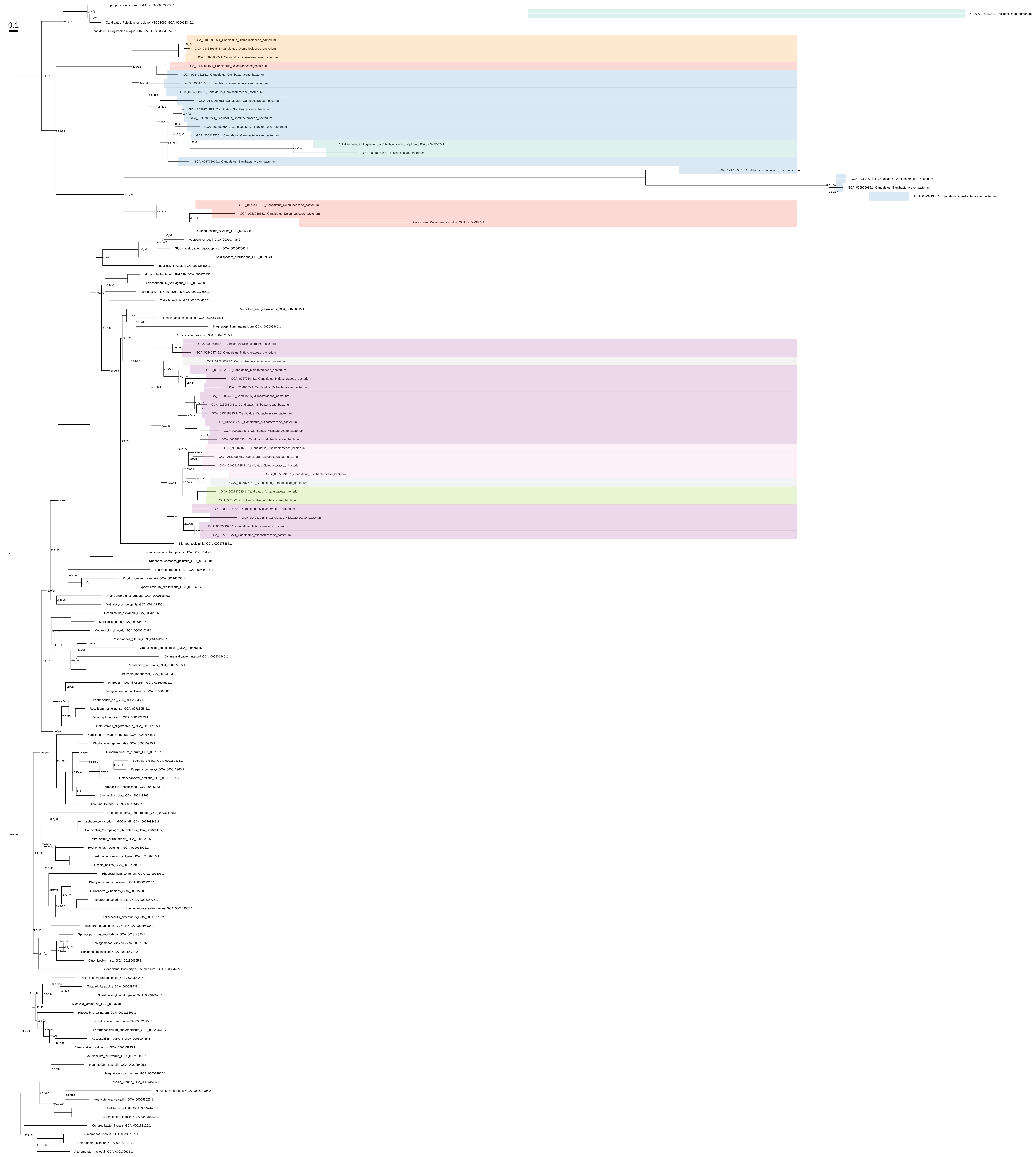

### Cys_all_organisms_z_0.10

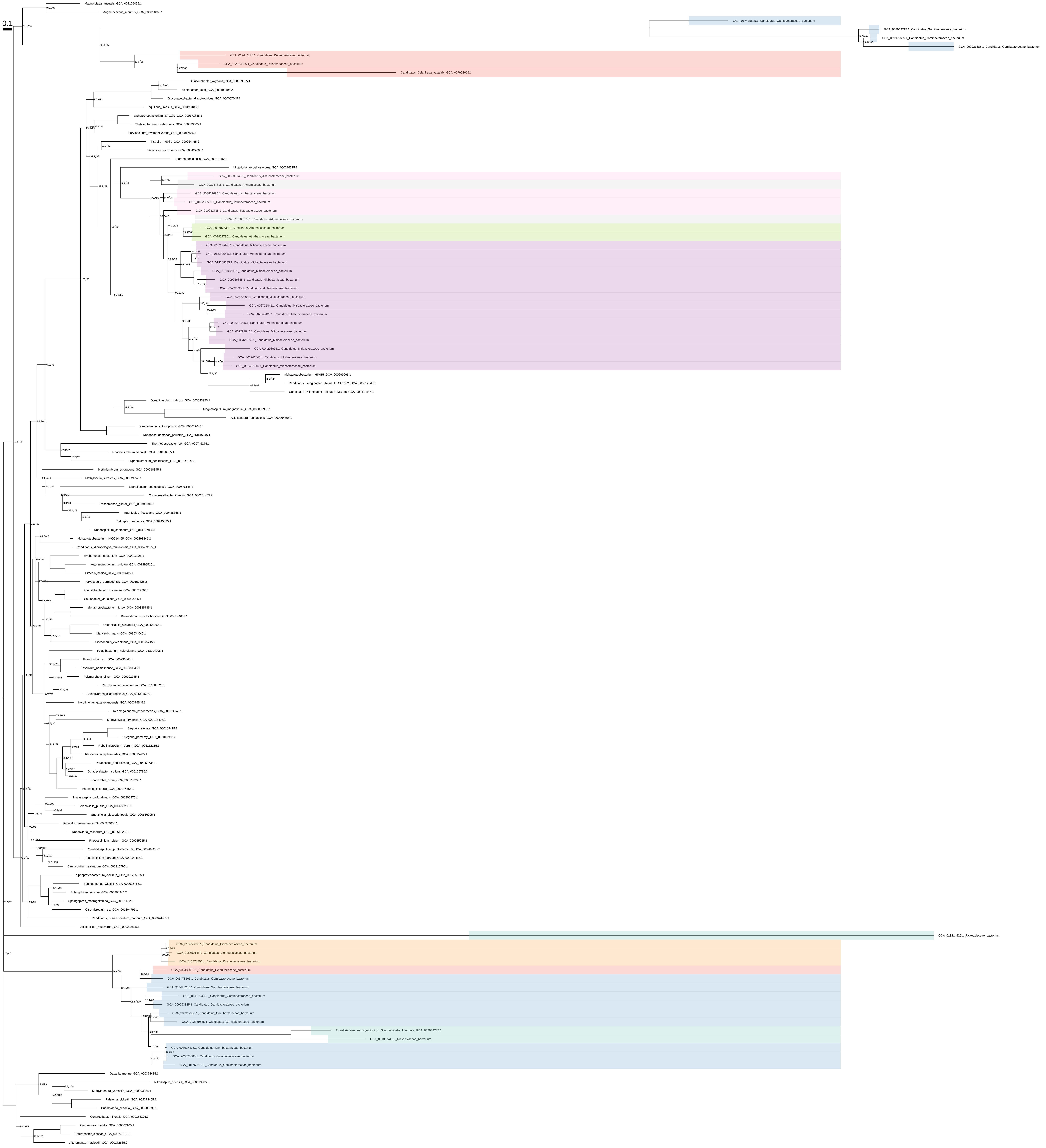

### Cys_all_organisms_z_0.20

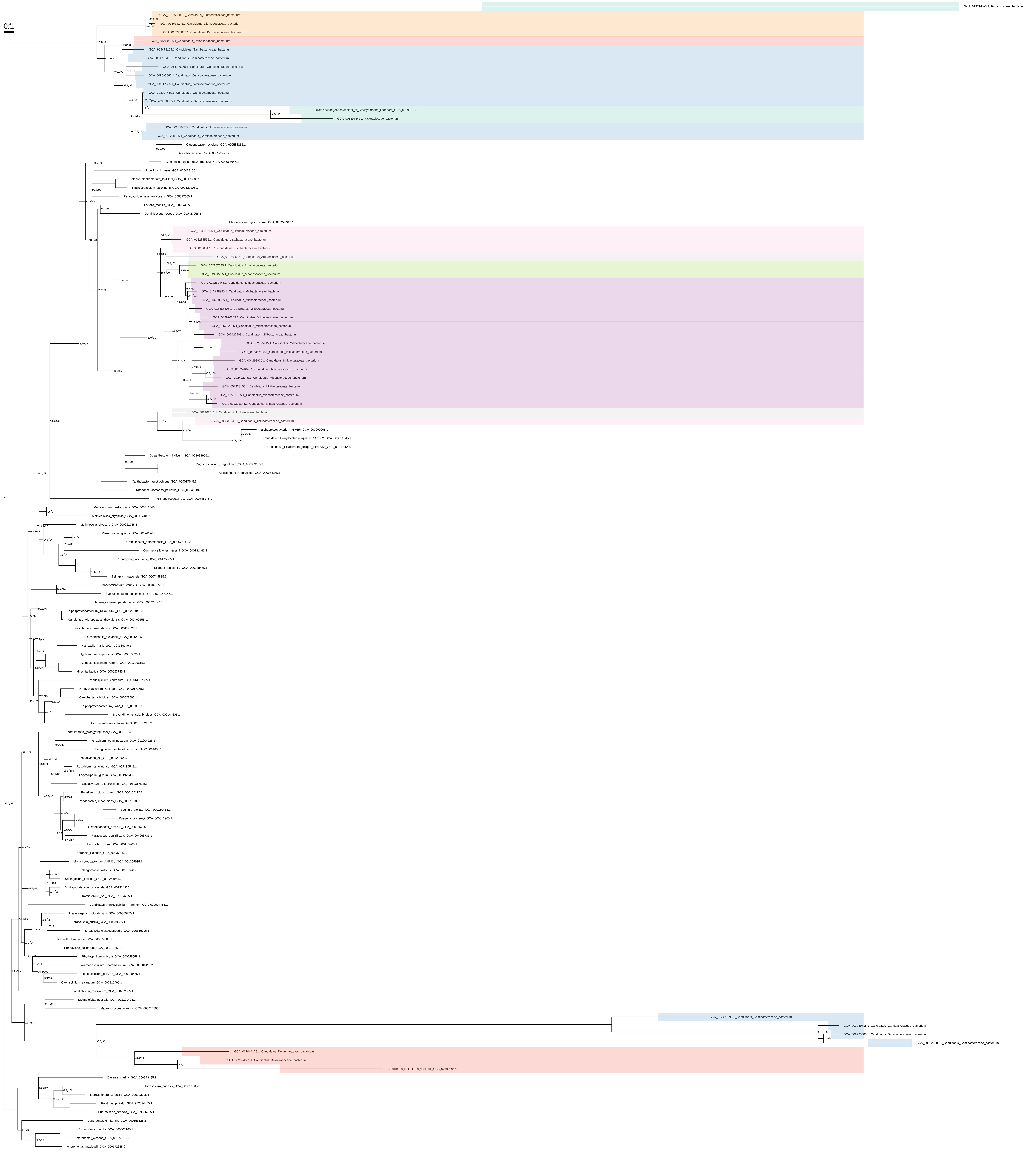

### Cys_all_organisms_z_0.30

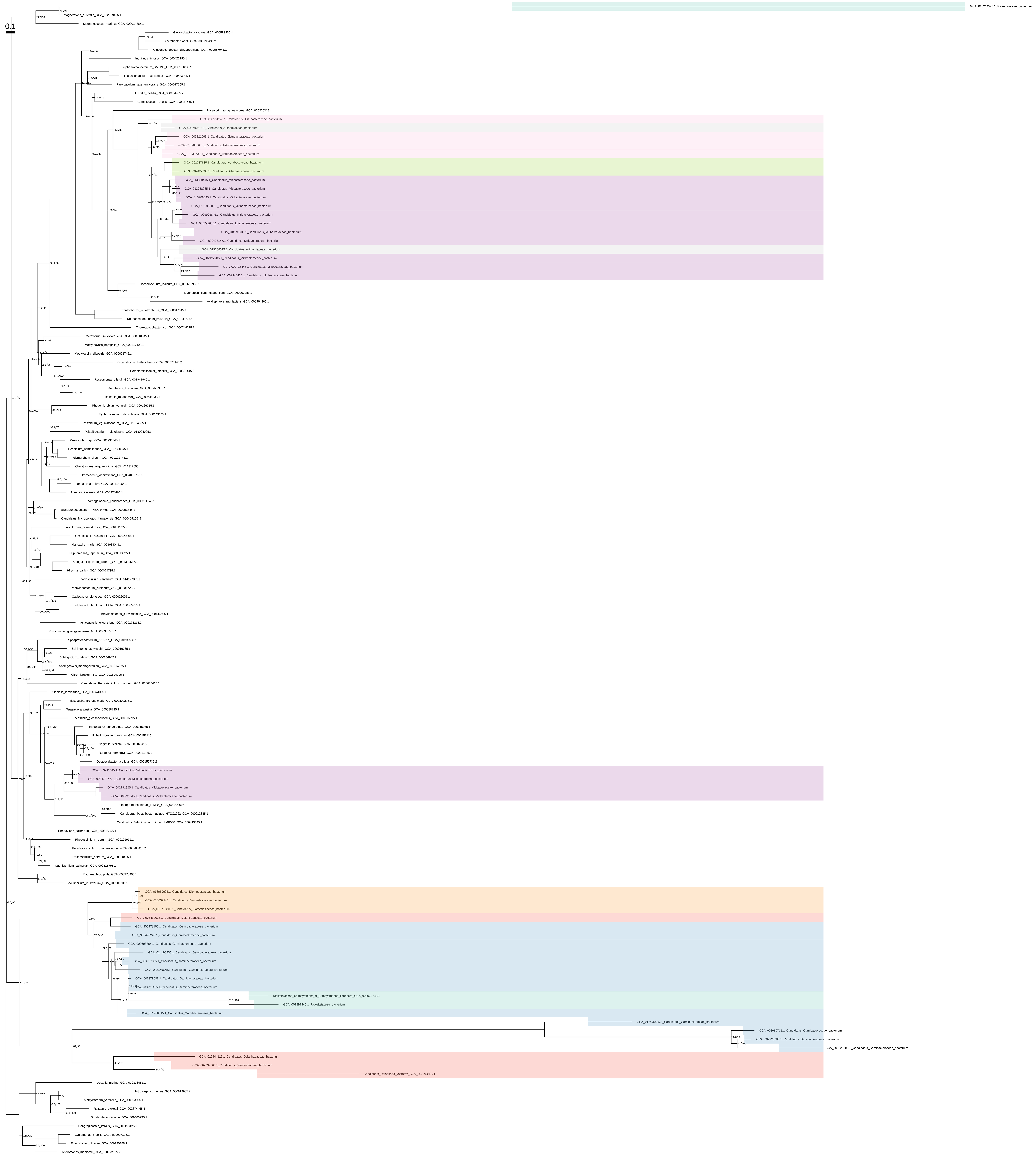

### Cys_all_organisms_z_0.40

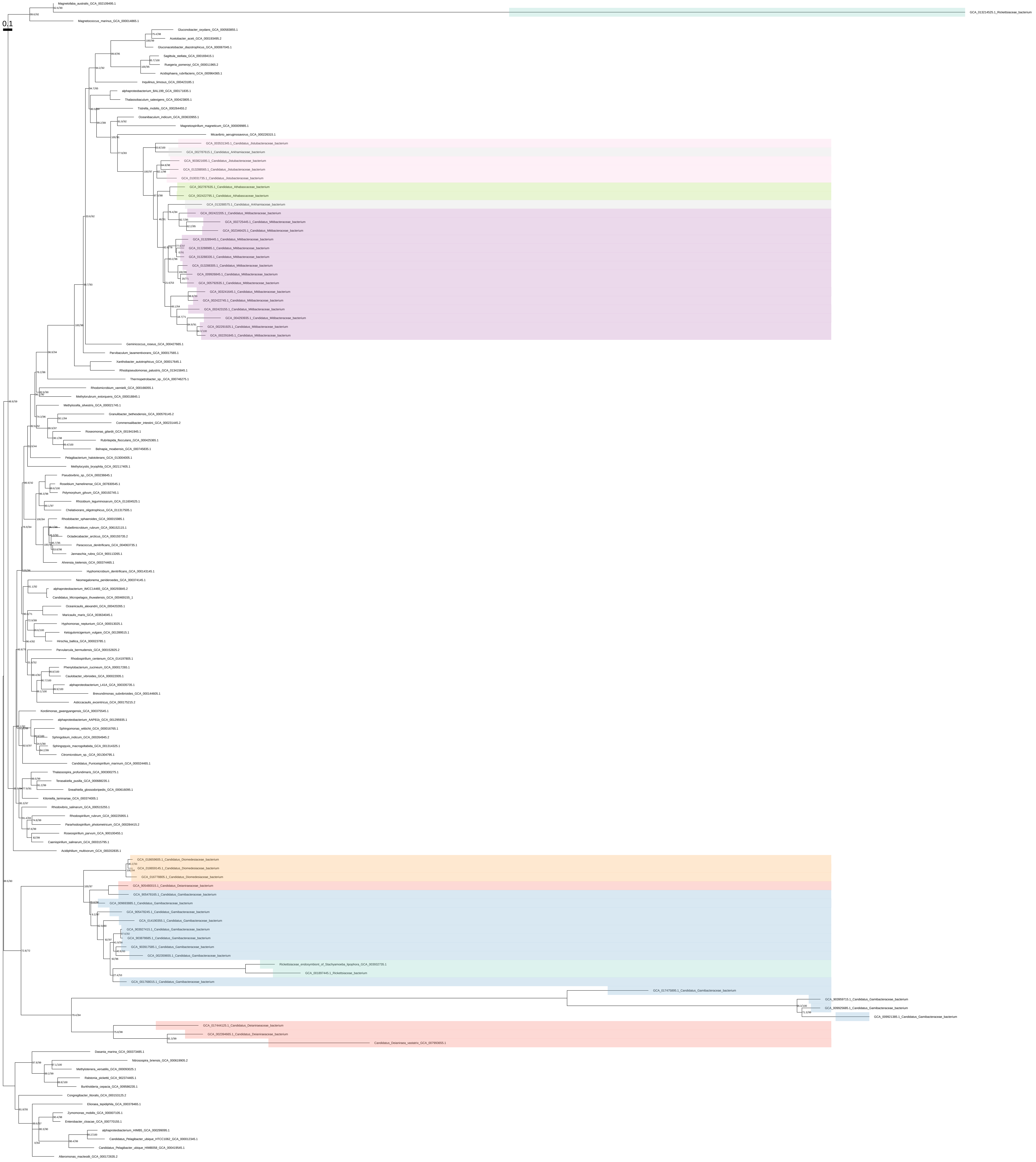

### Cys_all_organisms_z_0.50

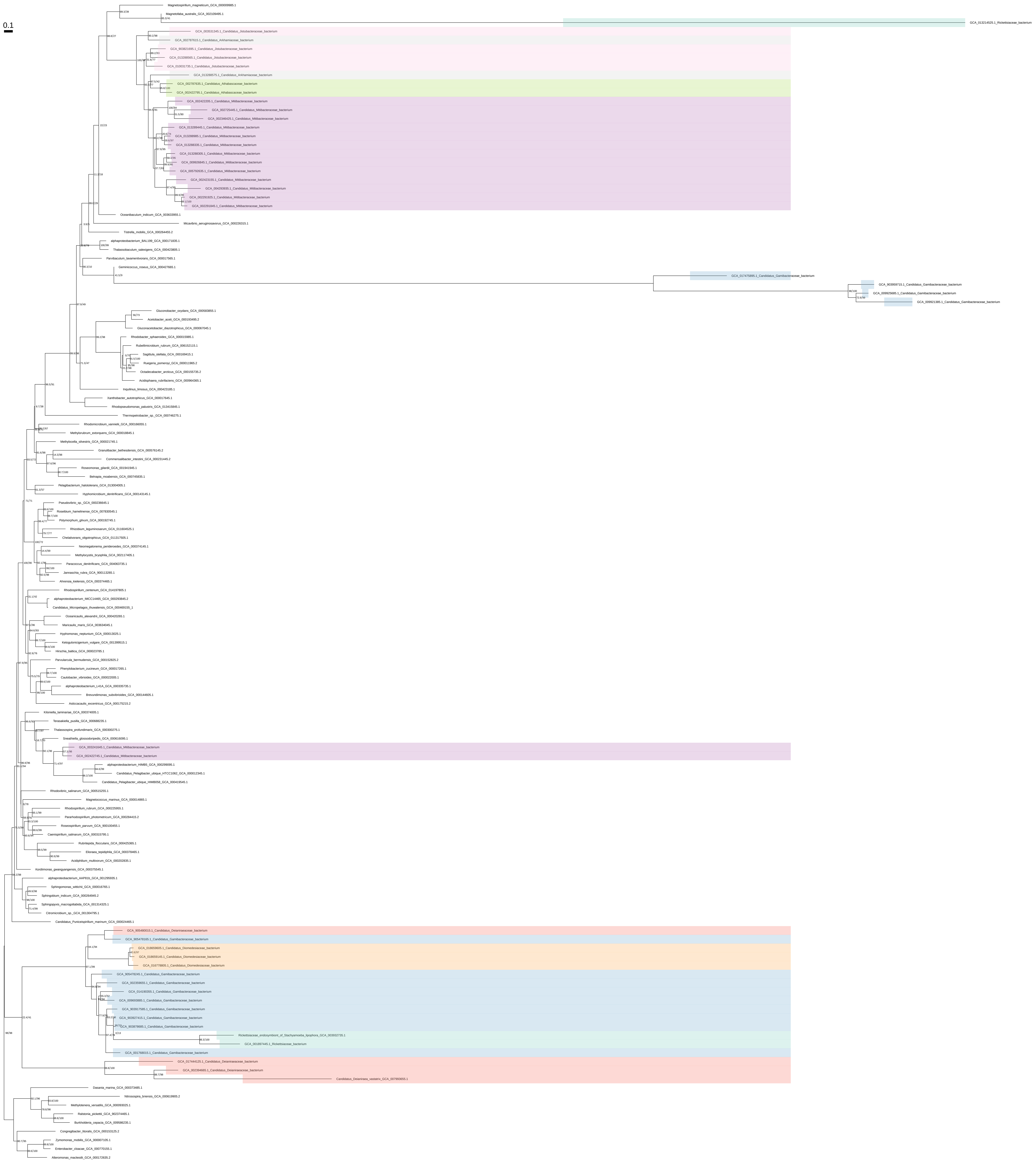

Cys_organism_selection

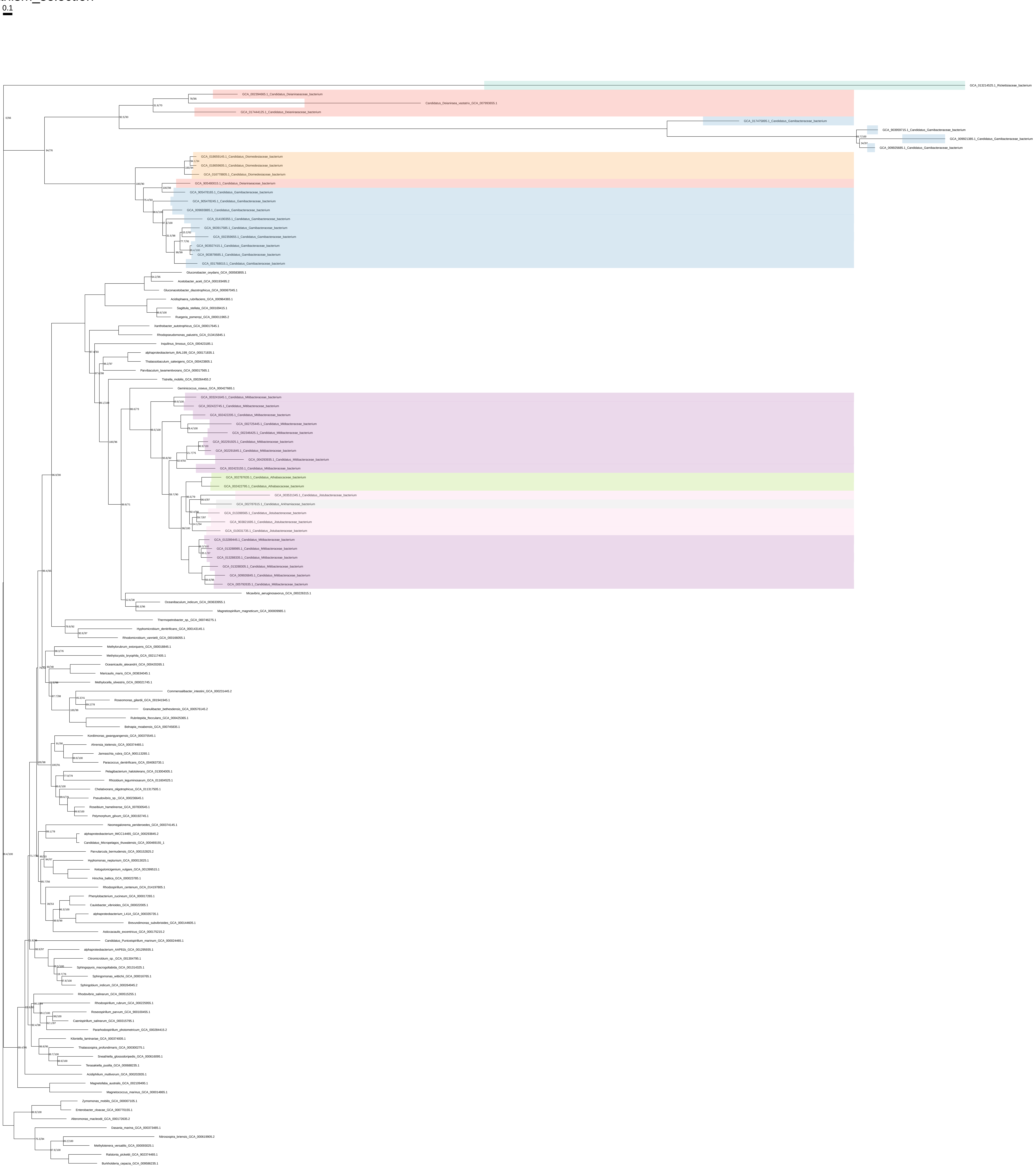

### Cys_organism_selection_z_0.10

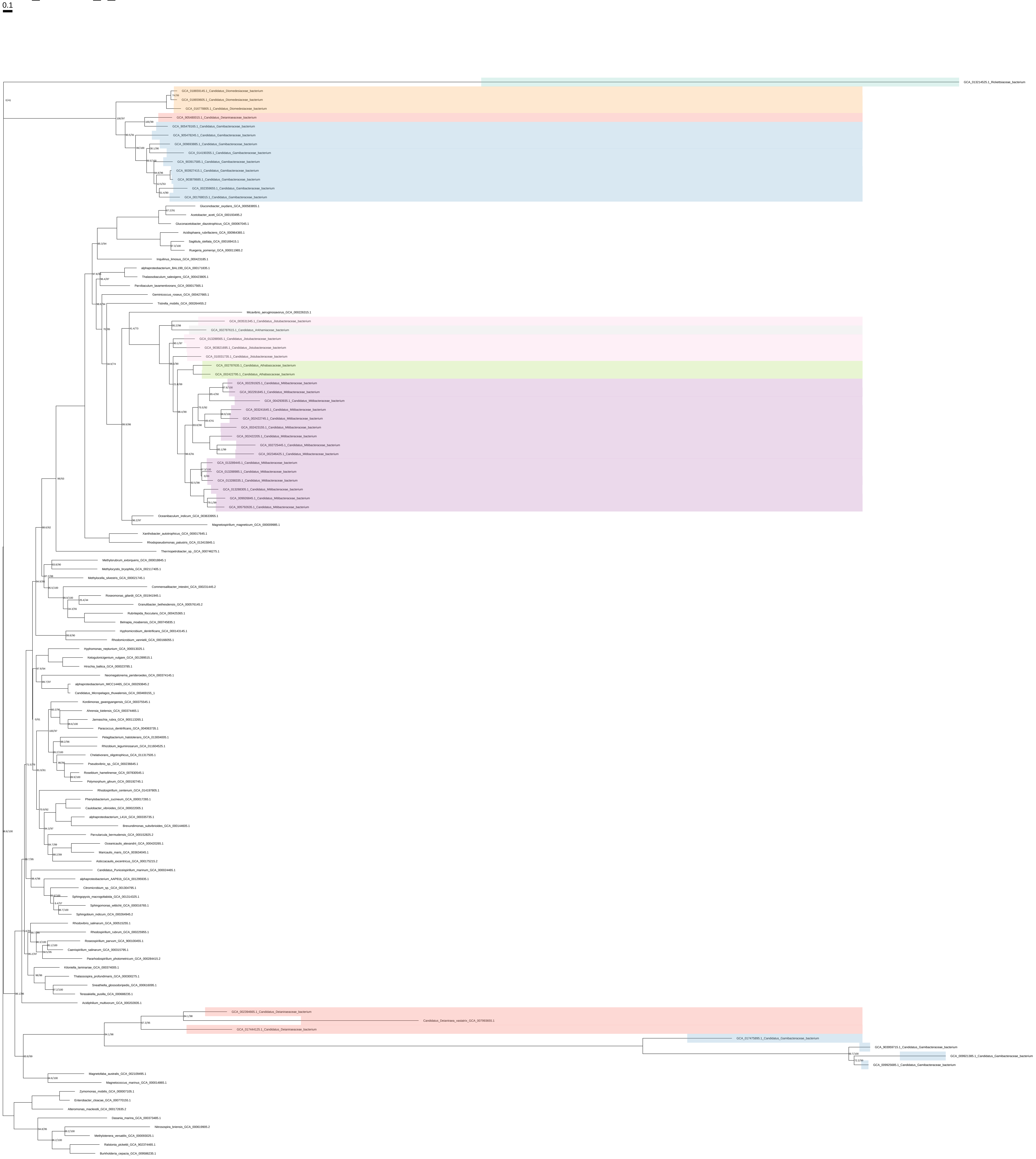

### Cys_organism_selection_z_0.20

0.1

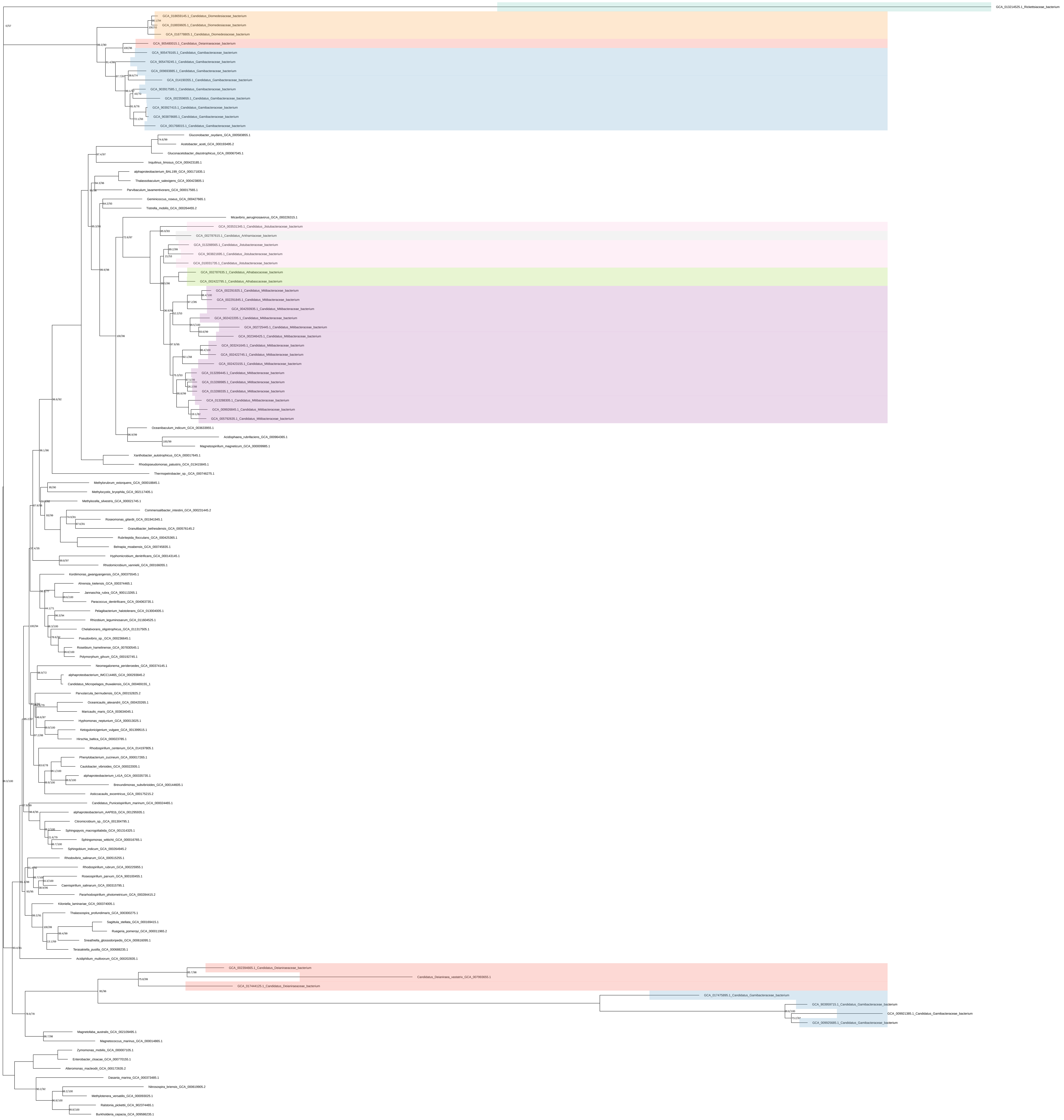

### Cys_organism_selection_z_0.30

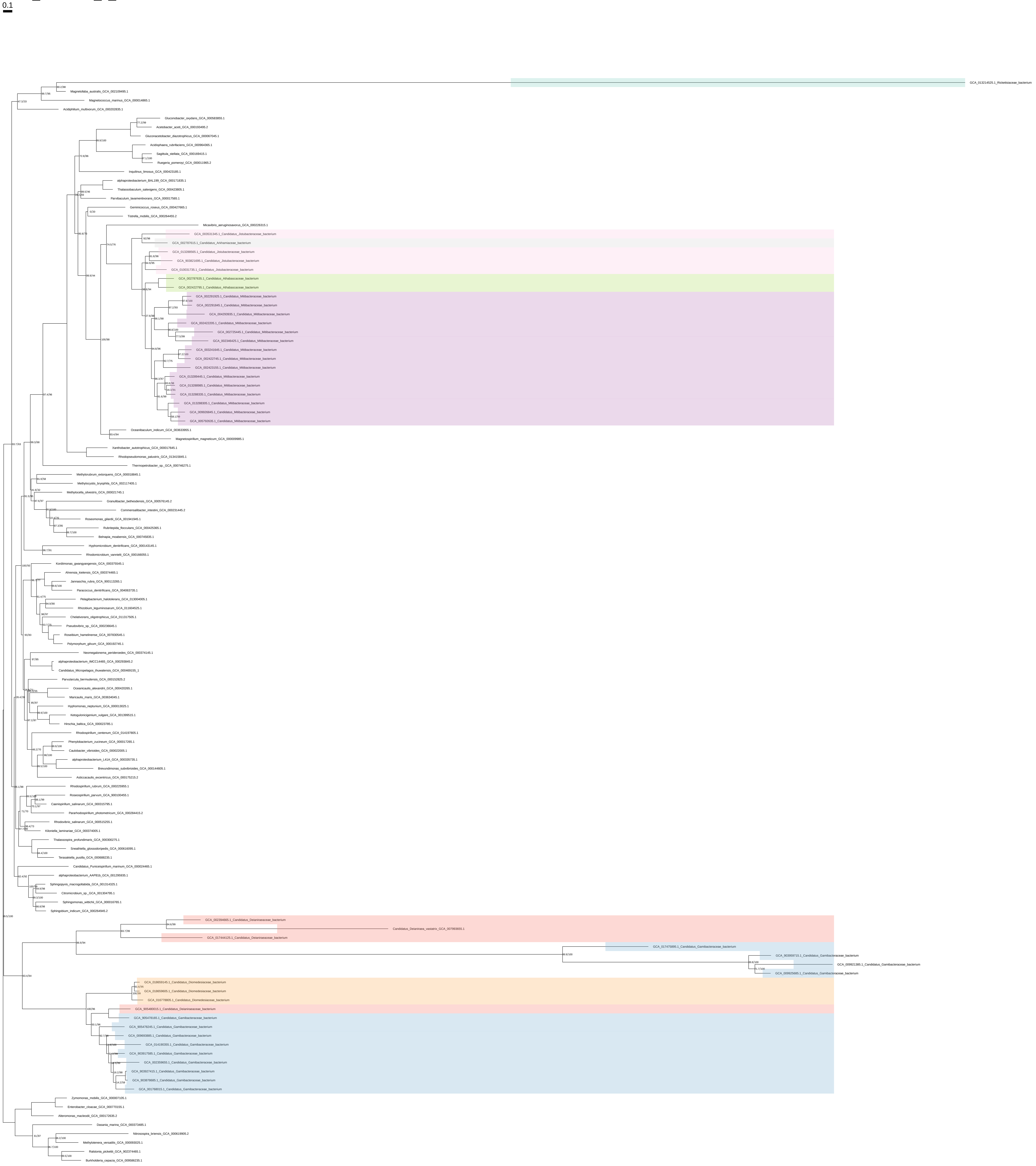

### Cys_organism_selection_z_0.40

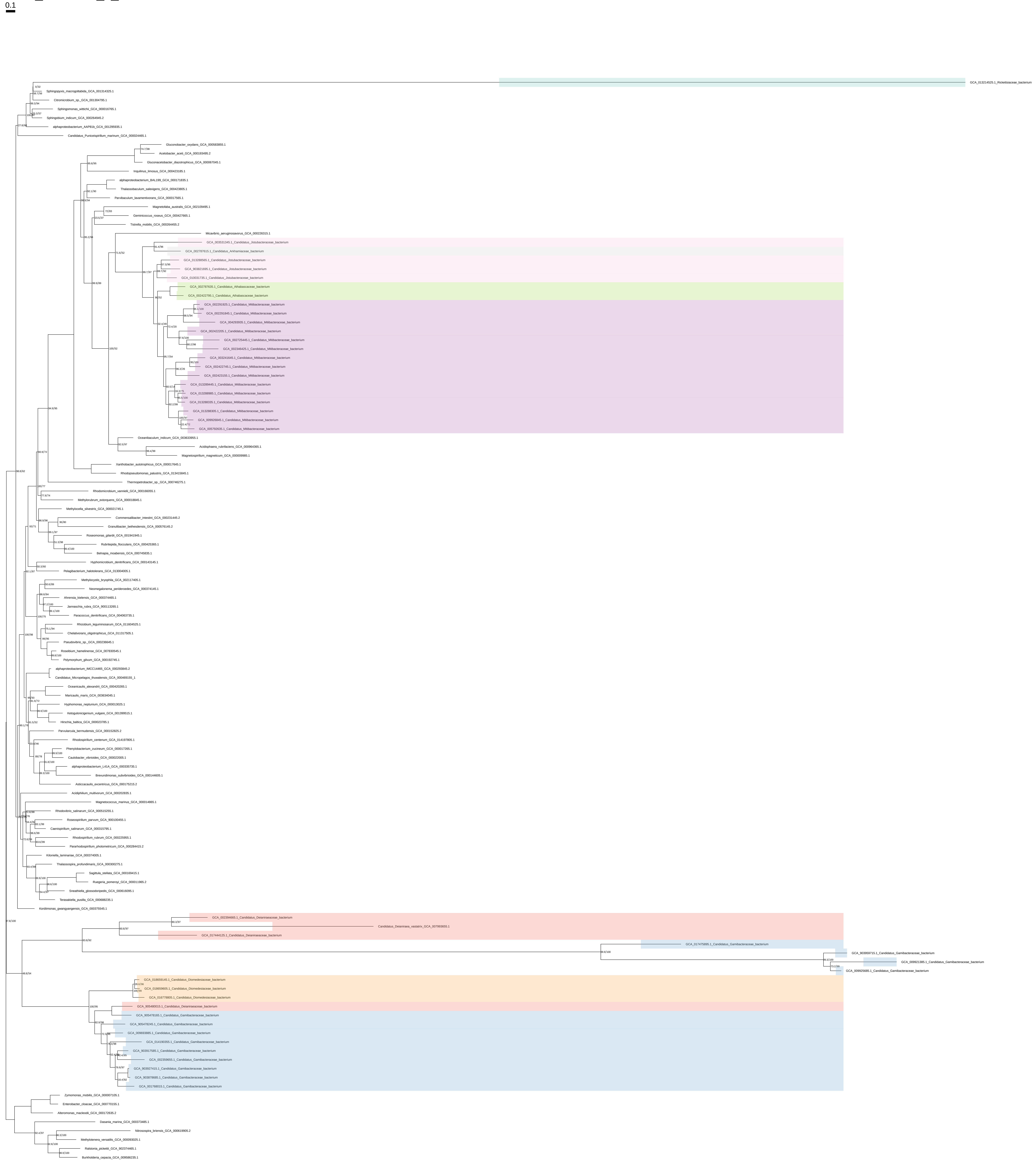

### Cys_organism_selection_z_0.50

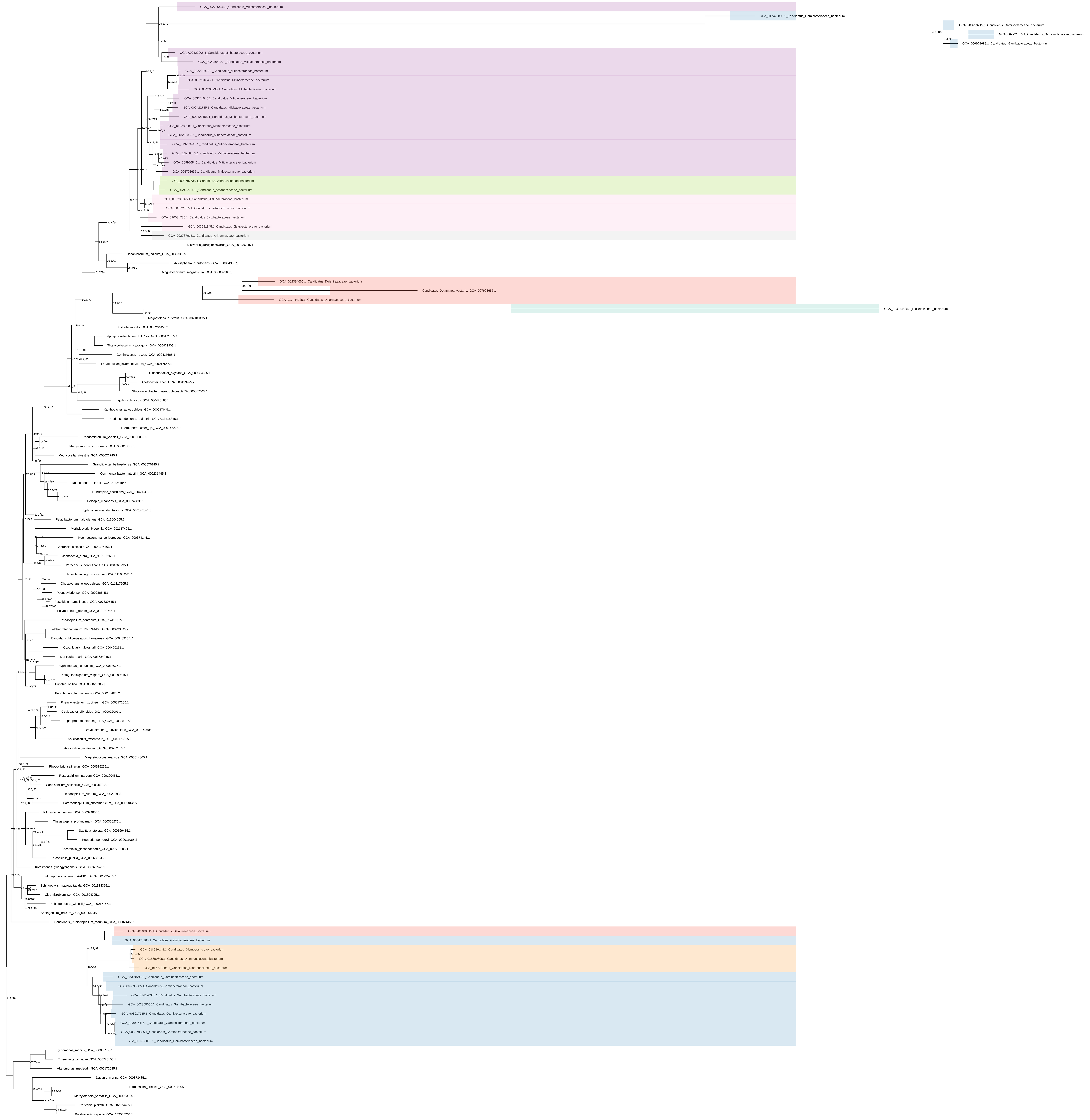

His_all_organisms

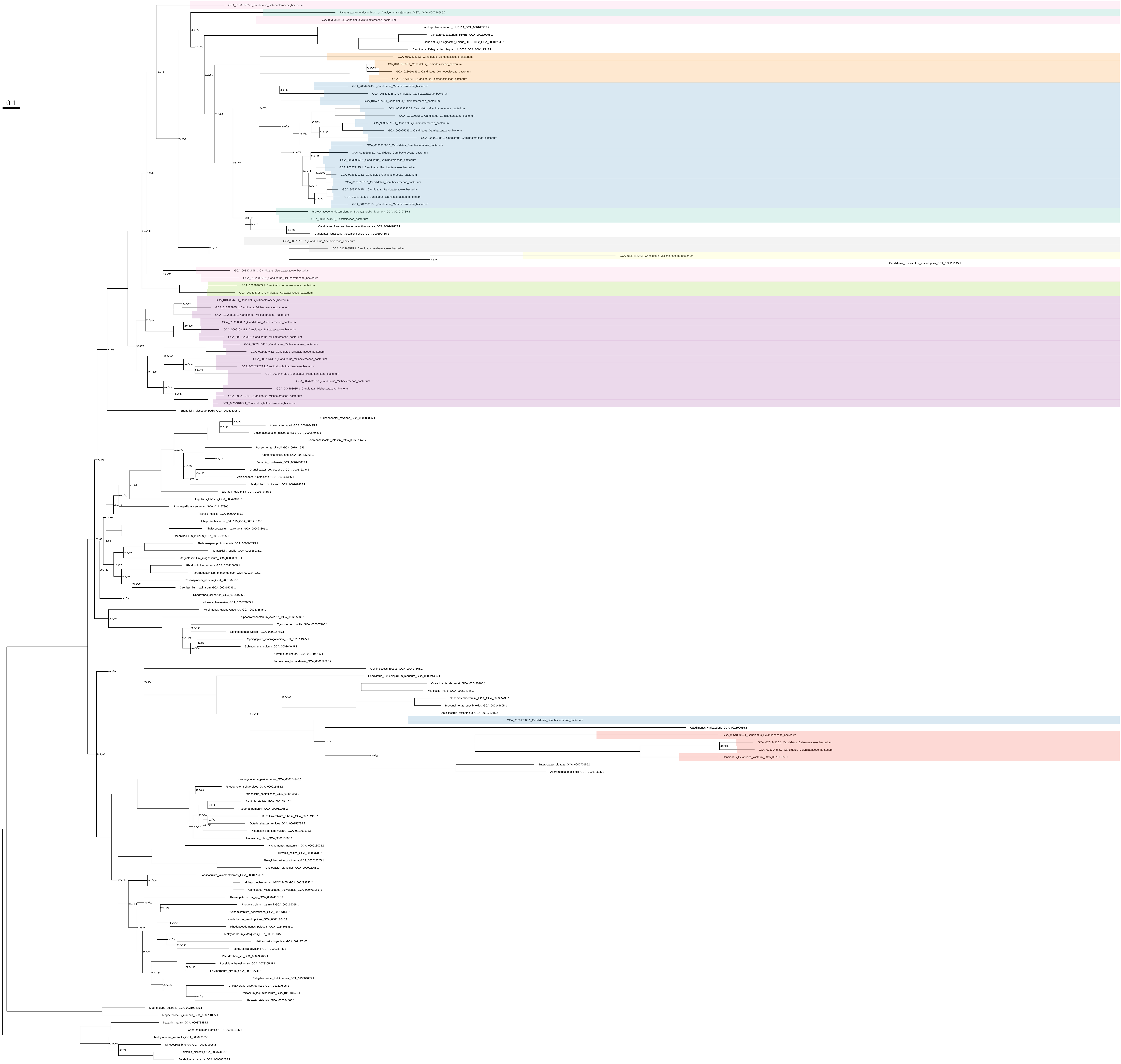

### His_all_organisms_z_0.10

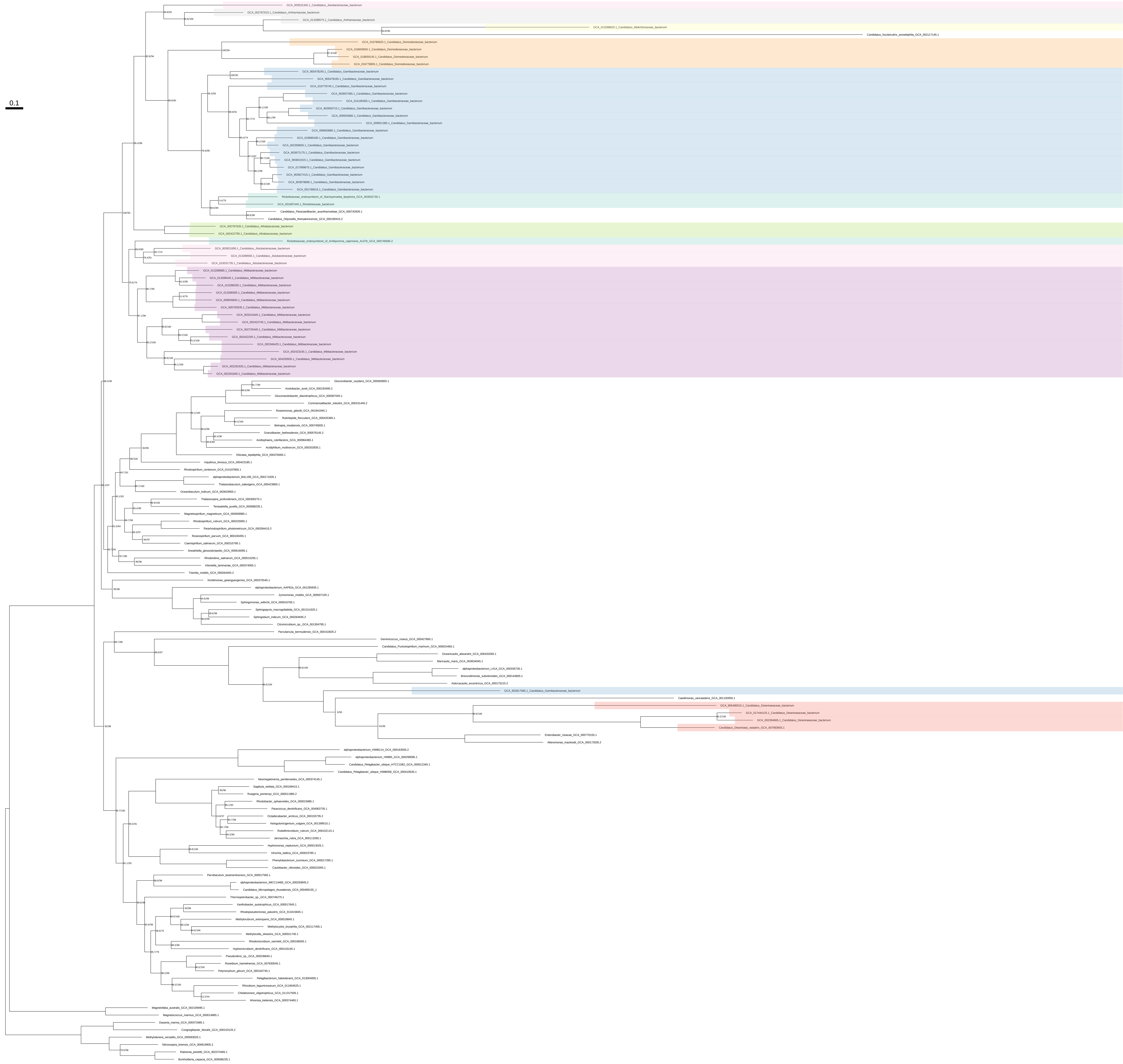

### His_all_organisms_z_0.20

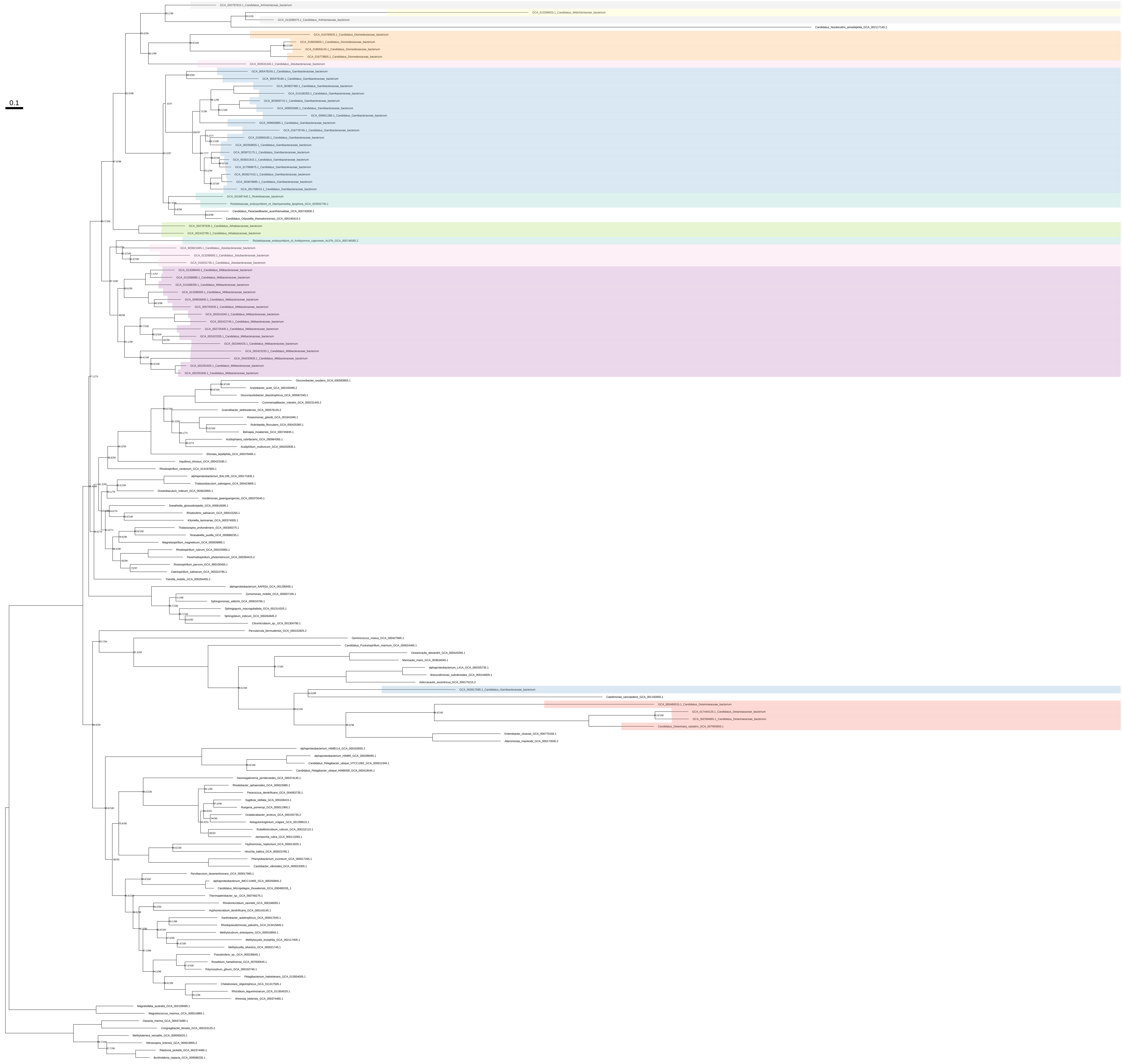

### His_all_organisms_z_0.30

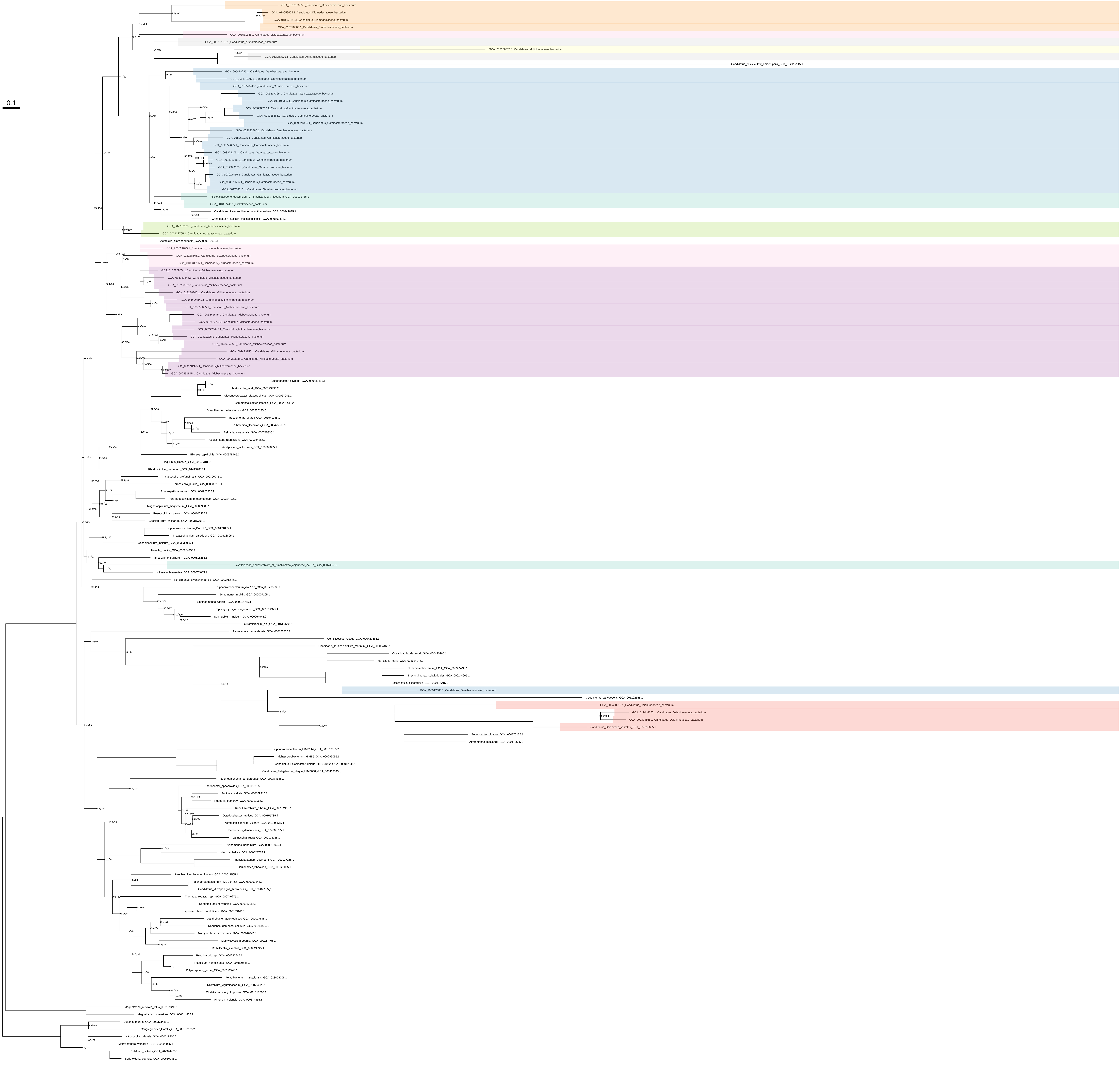

His_all_organisms_z_0.40

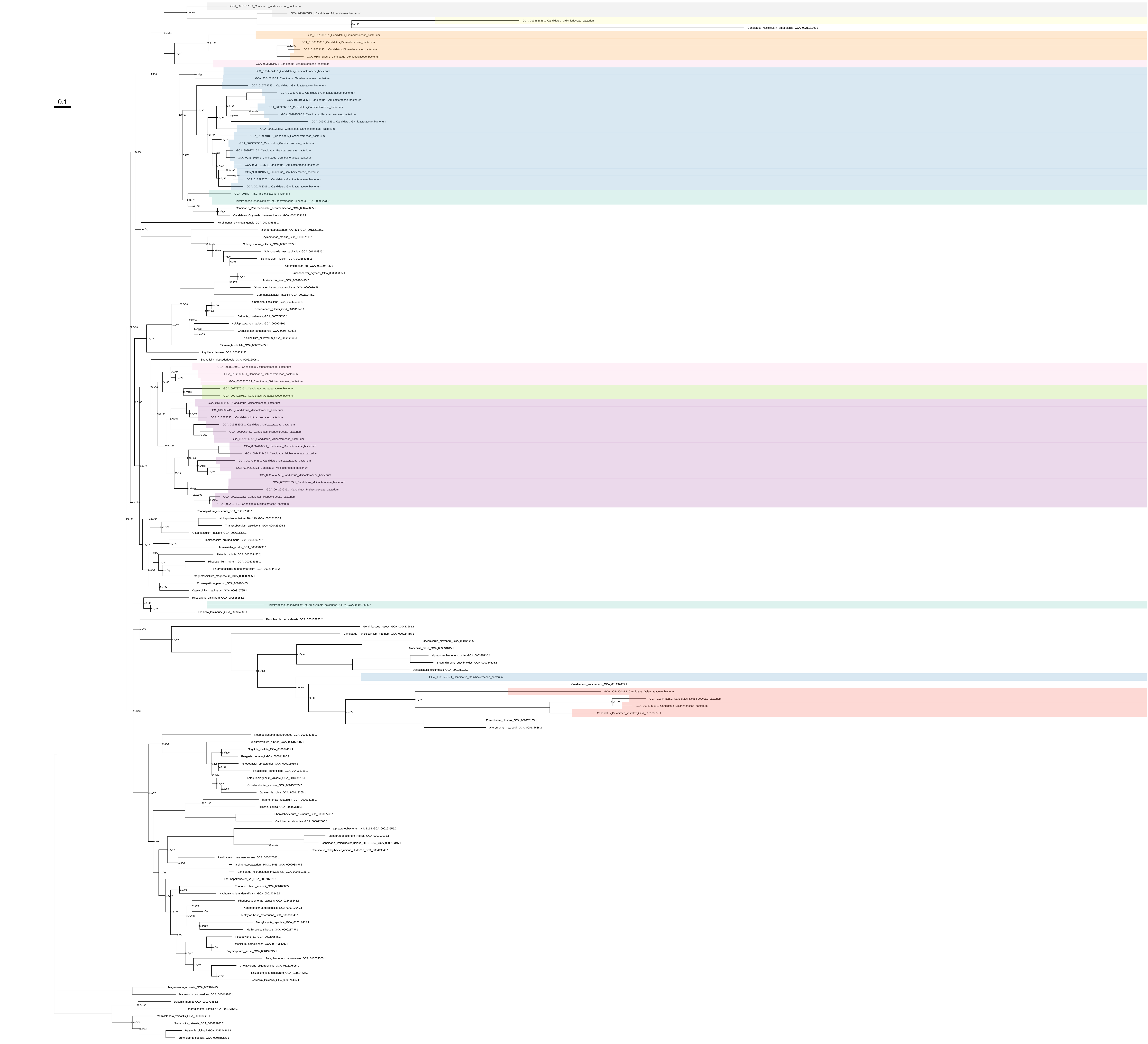

His_all_organisms_z_0.50

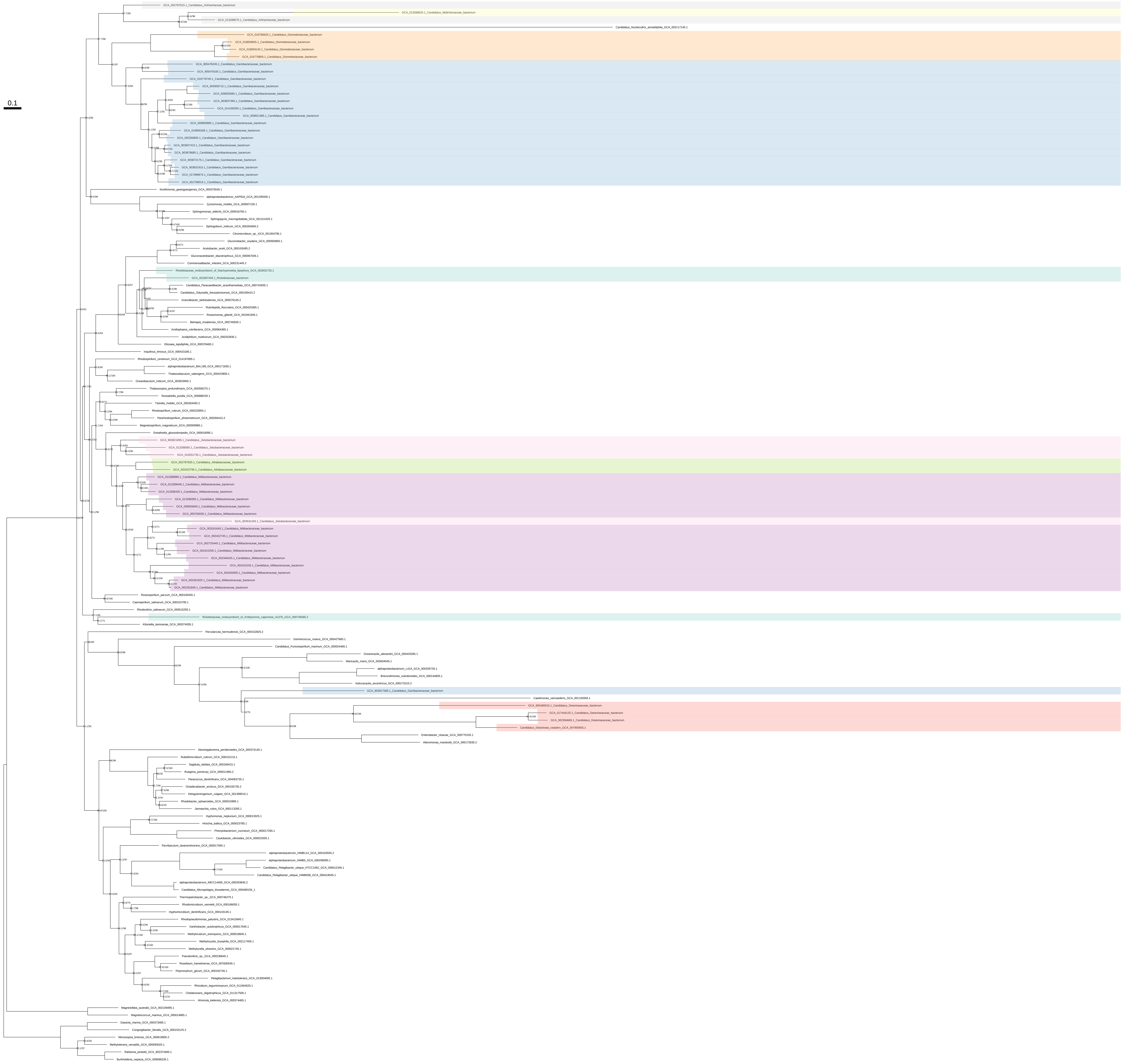

### His_organism_selection

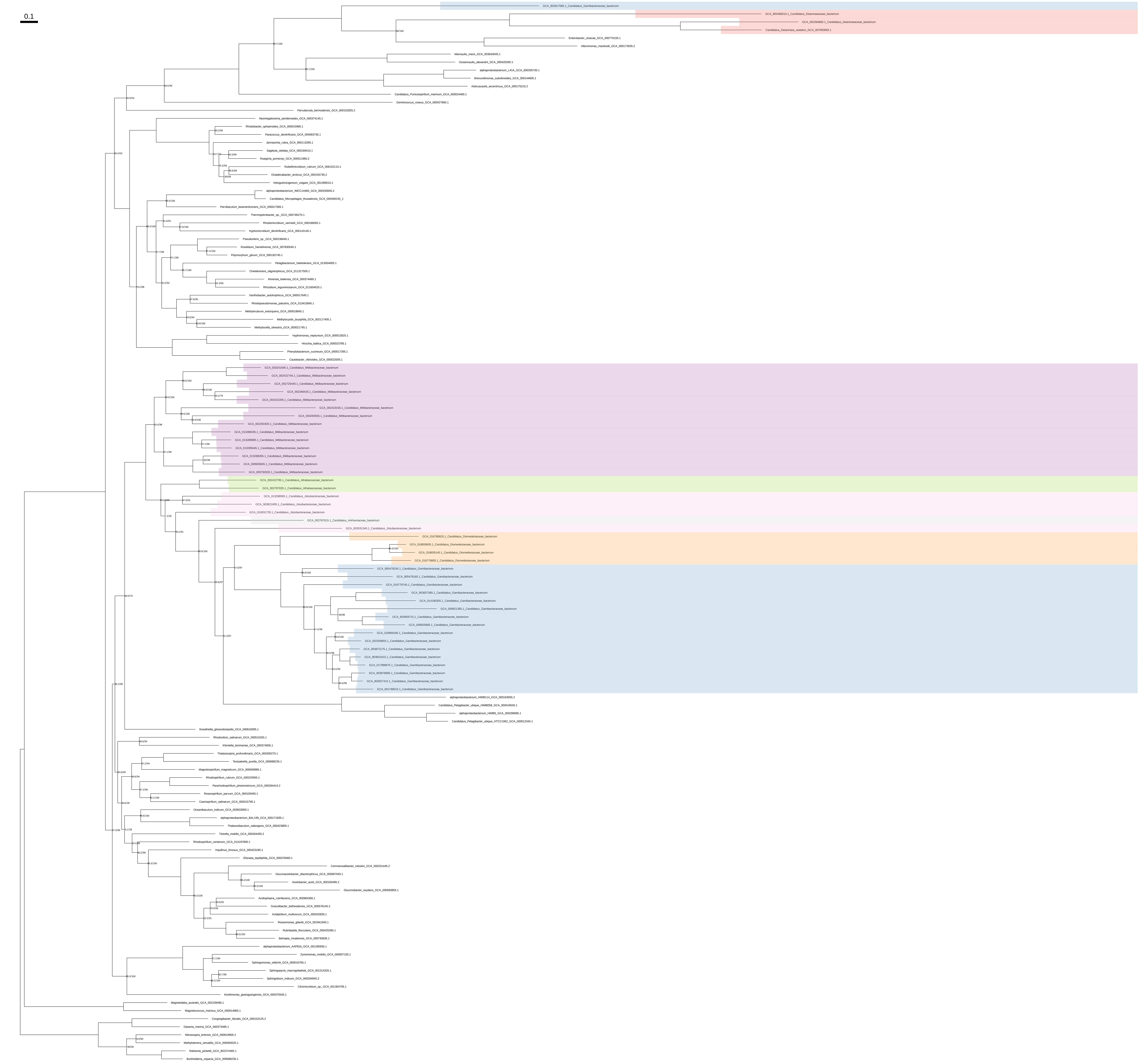

### His_organism_selection_z_0.10

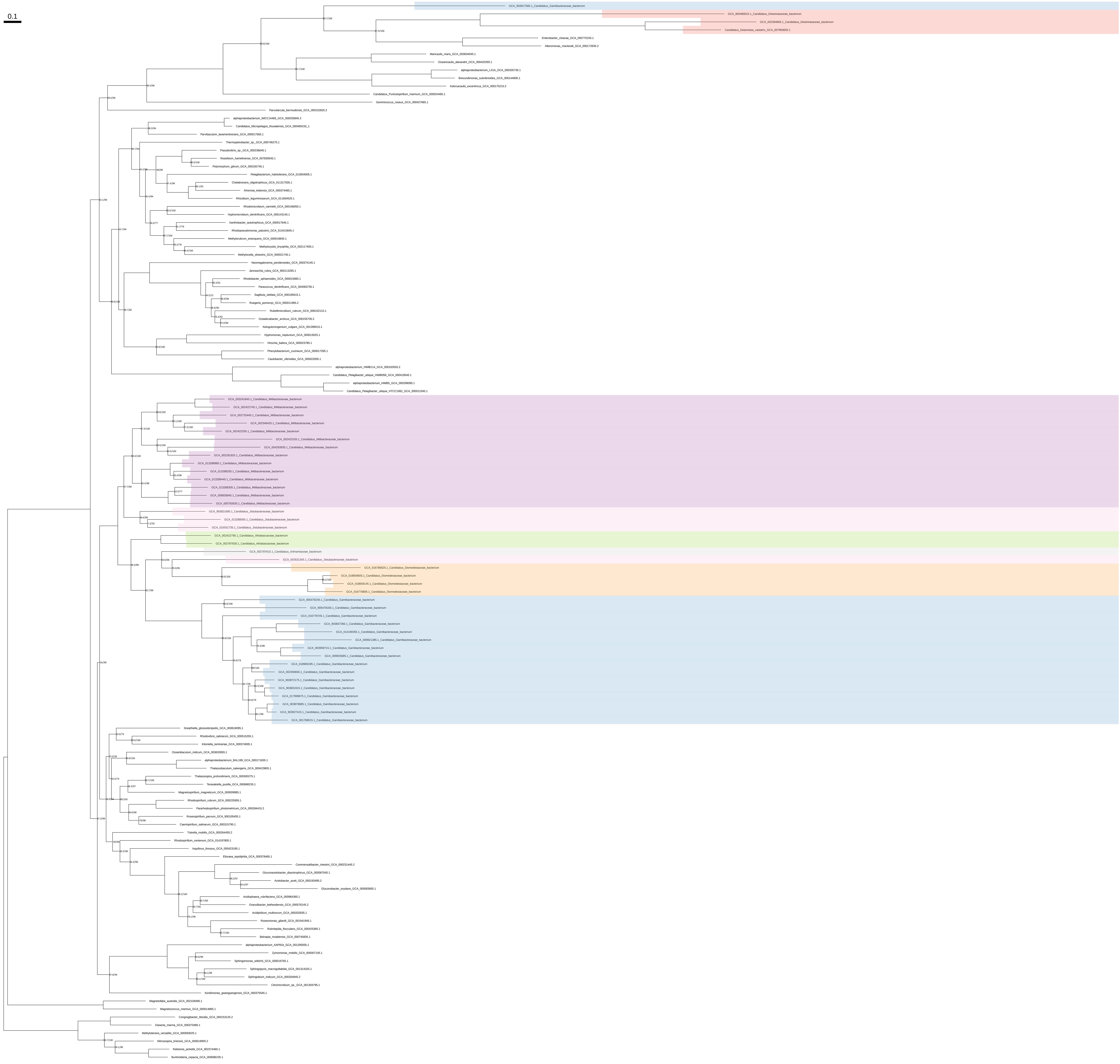

### His_organism_selection_z_0.20

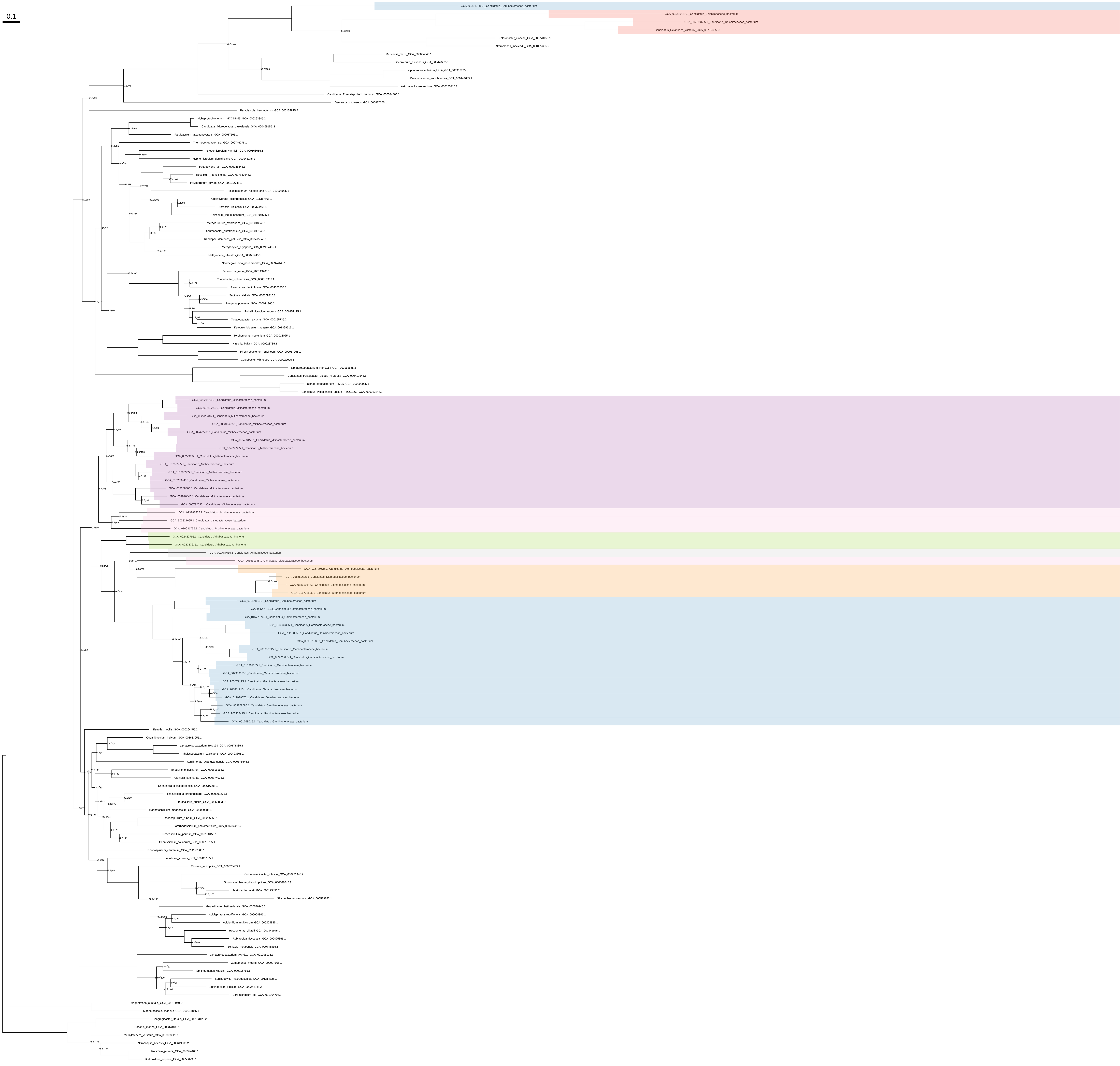

### His_organism_selection_z_0.30

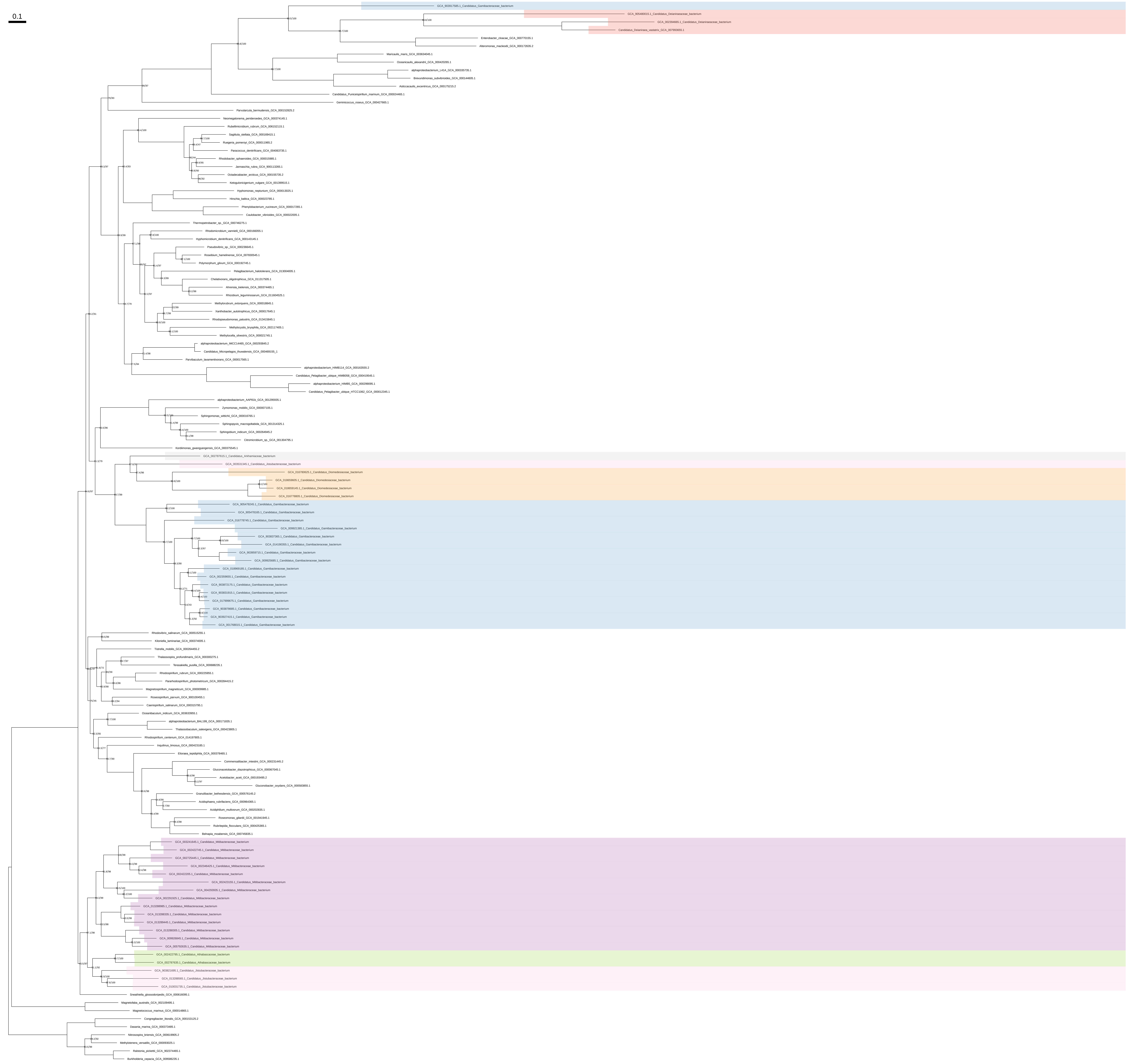

### His_organism_selection_z_0.40

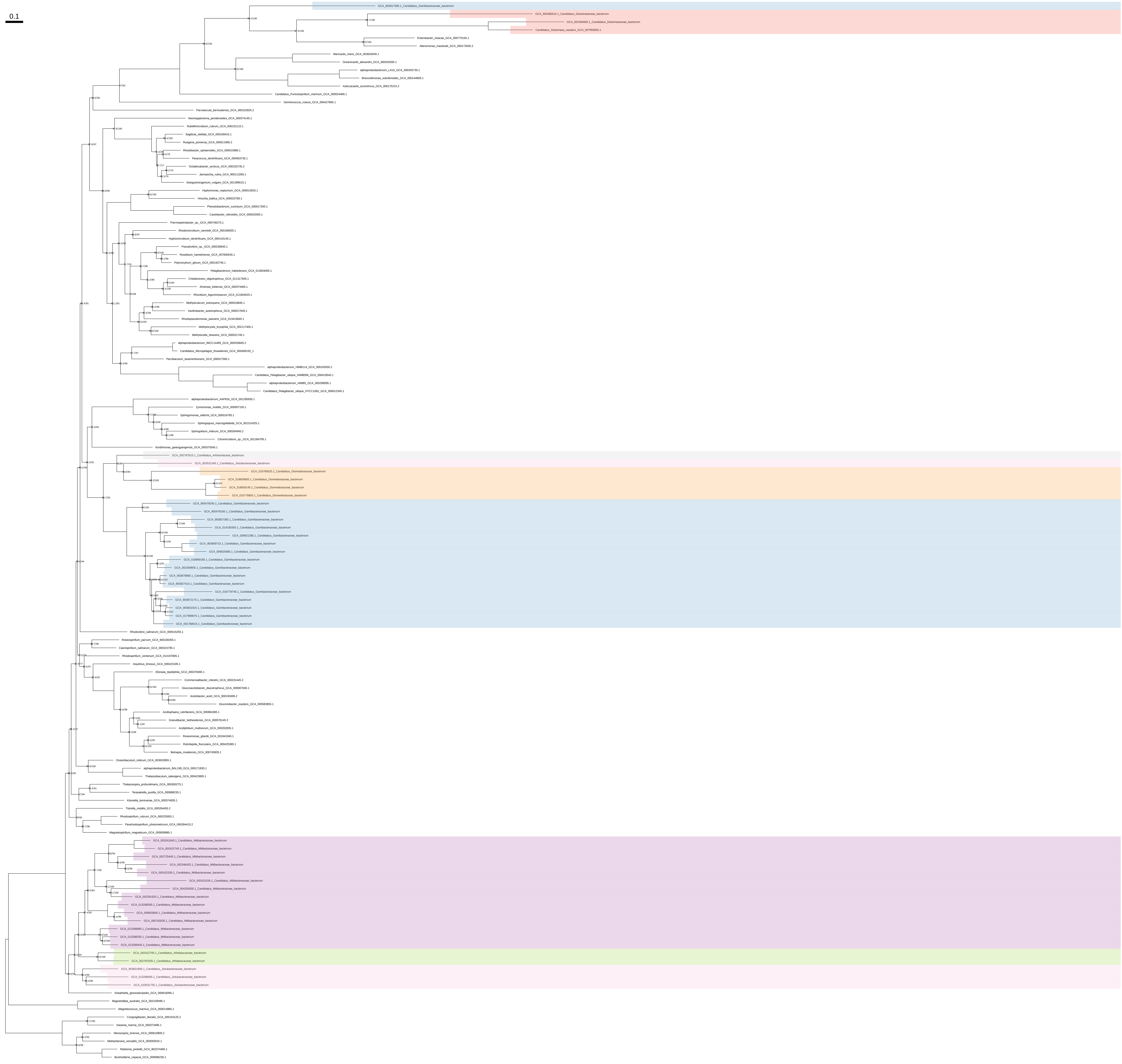

### His_organism_selection_z_0.50

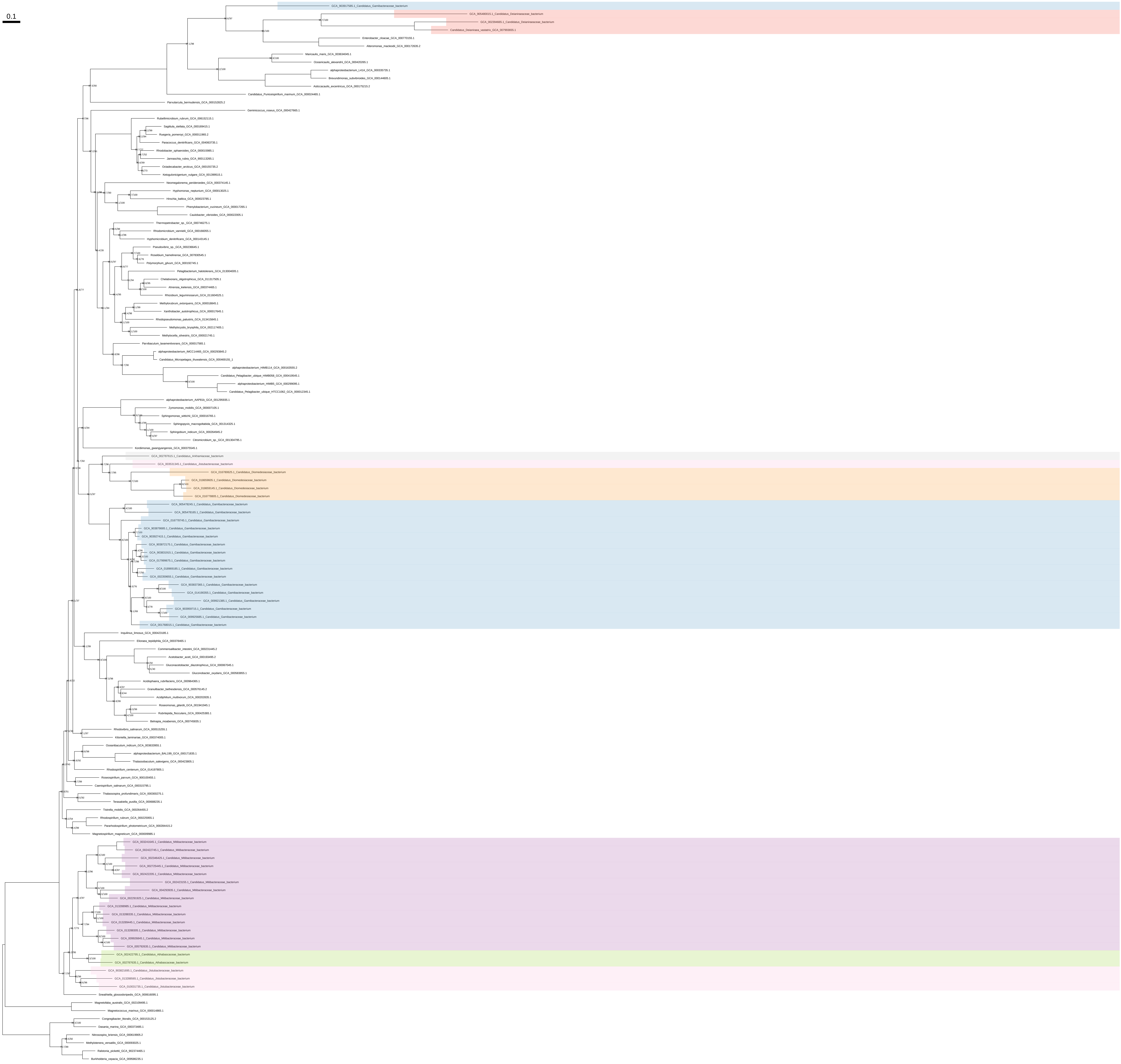

### Ile_Val_Leu_all_organisms

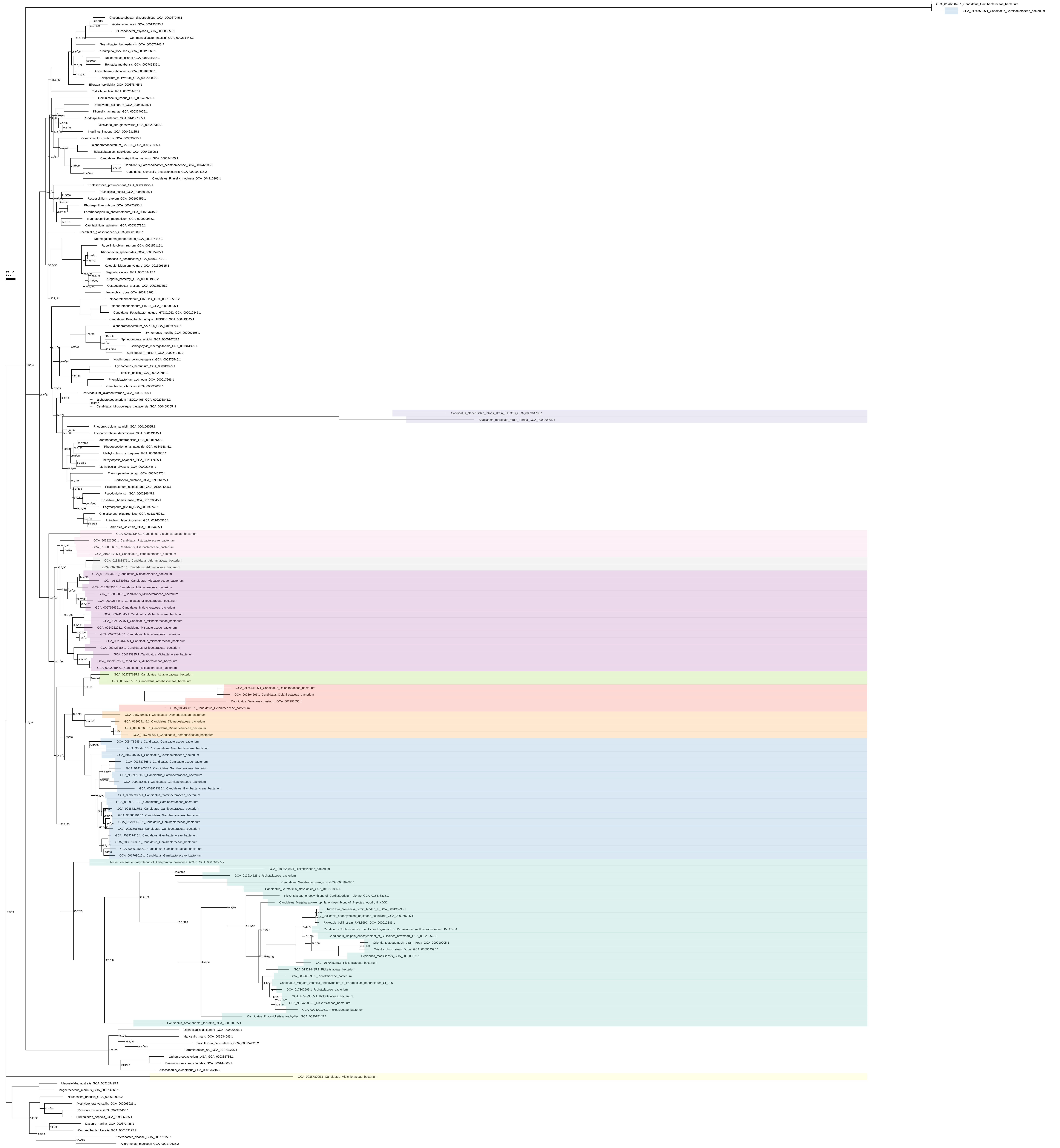

### Ile_Val_Leu_all_organisms_z_0.10

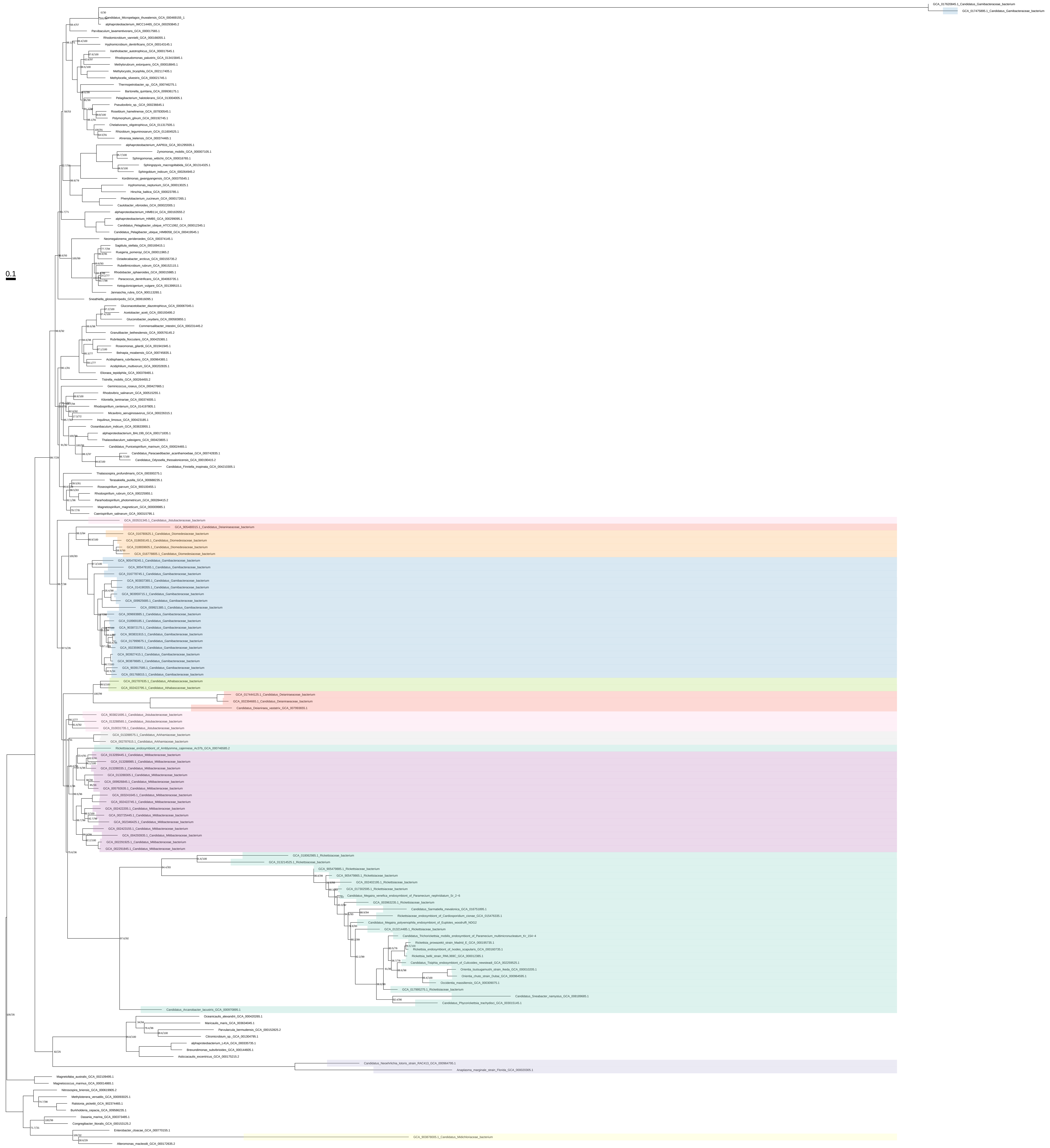

### Ile_Val_Leu_all_organisms_z_0.20

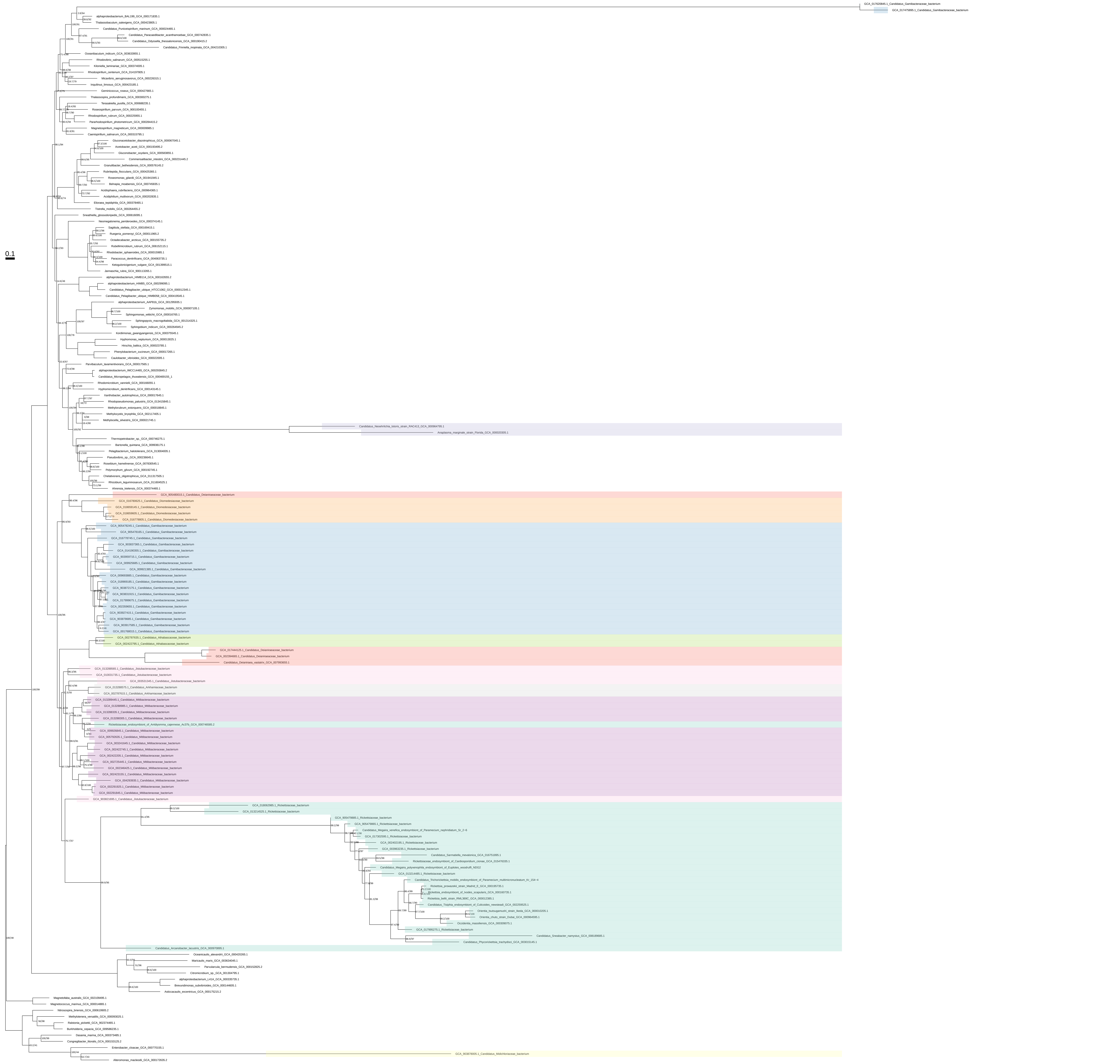

### Ile_Val_Leu_all_organisms_z_0.30

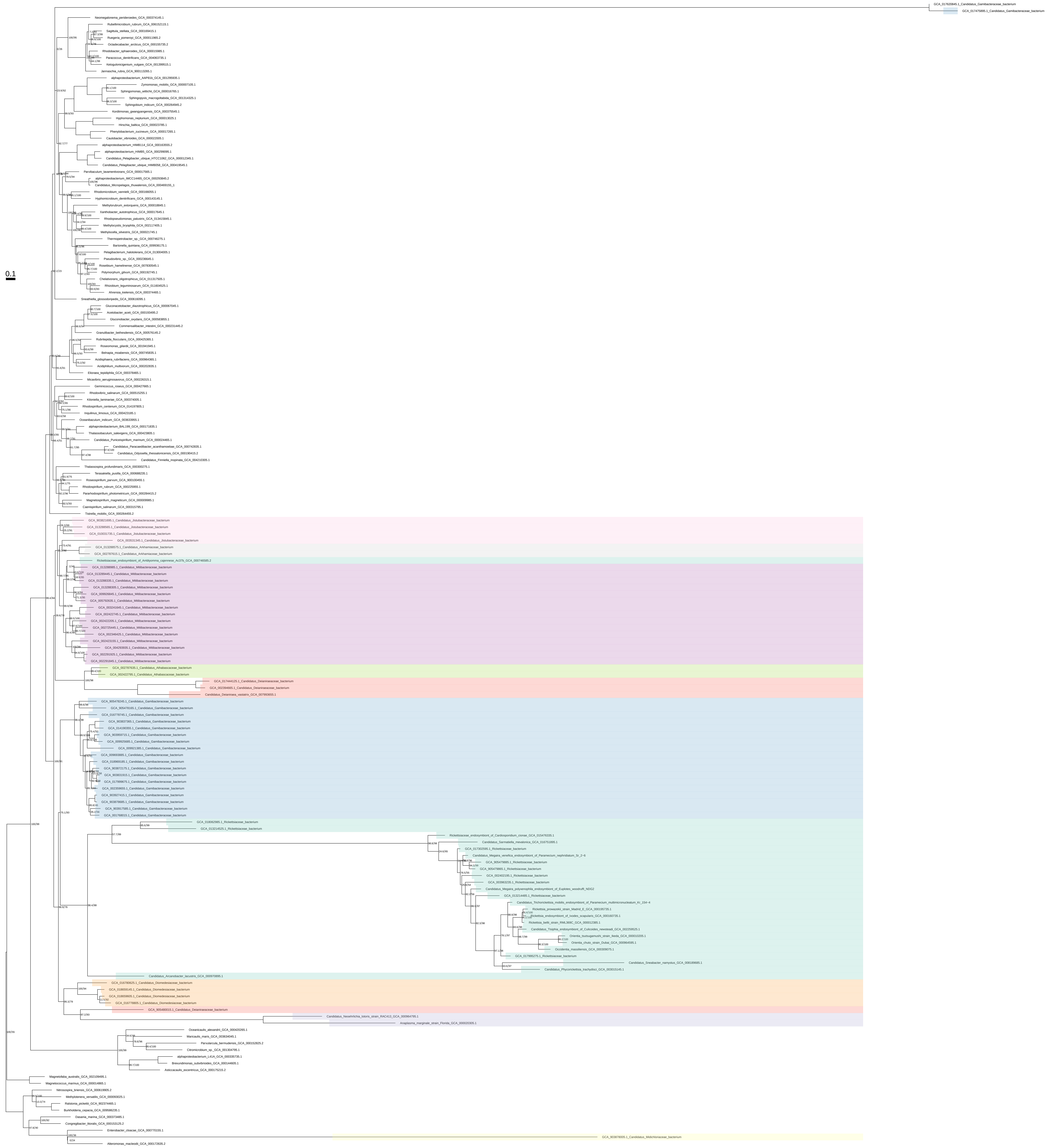

### Ile_Val_Leu_all_organisms_z_0.40

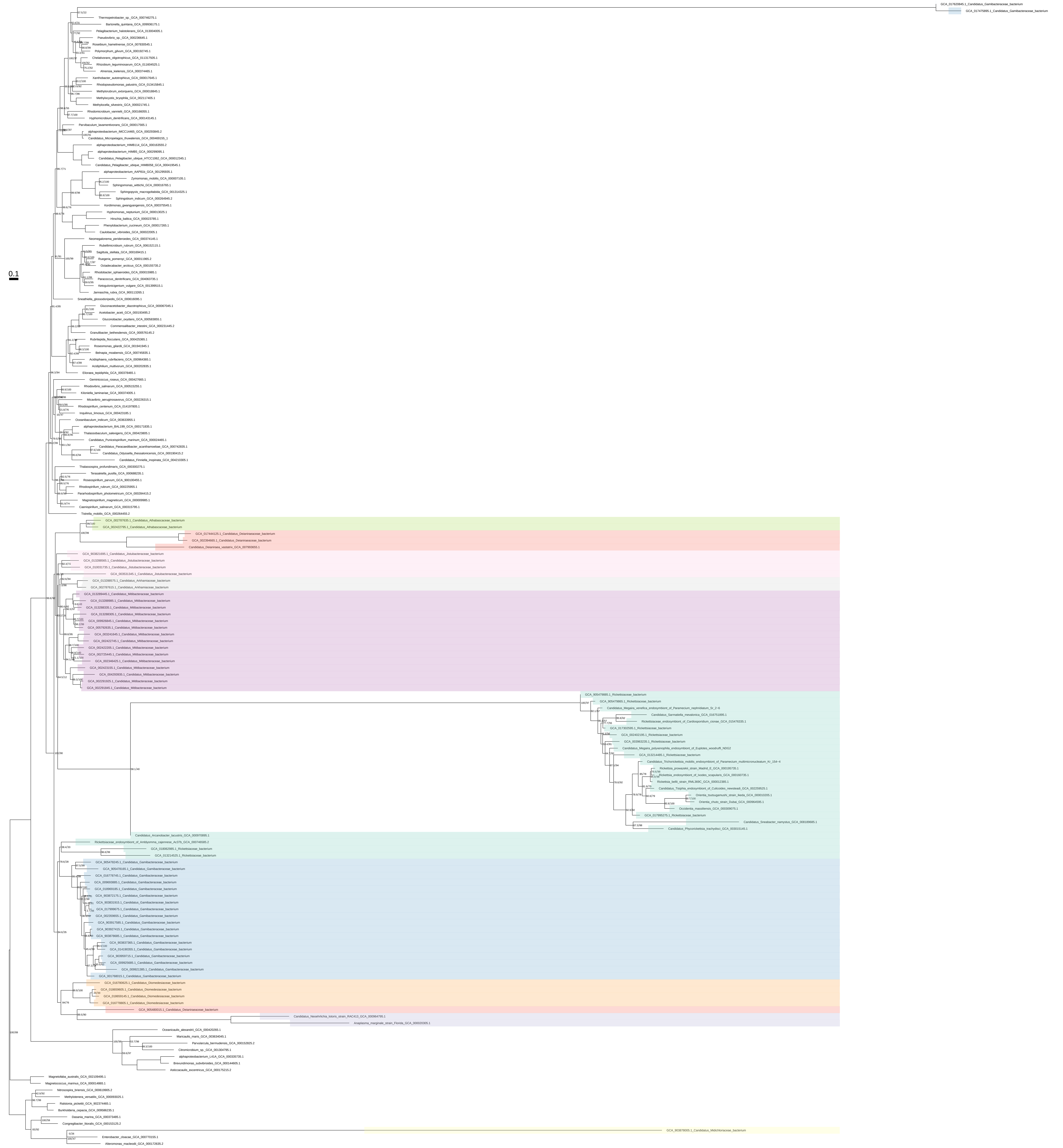

### Ile_Val_Leu_all_organisms_z_0.50

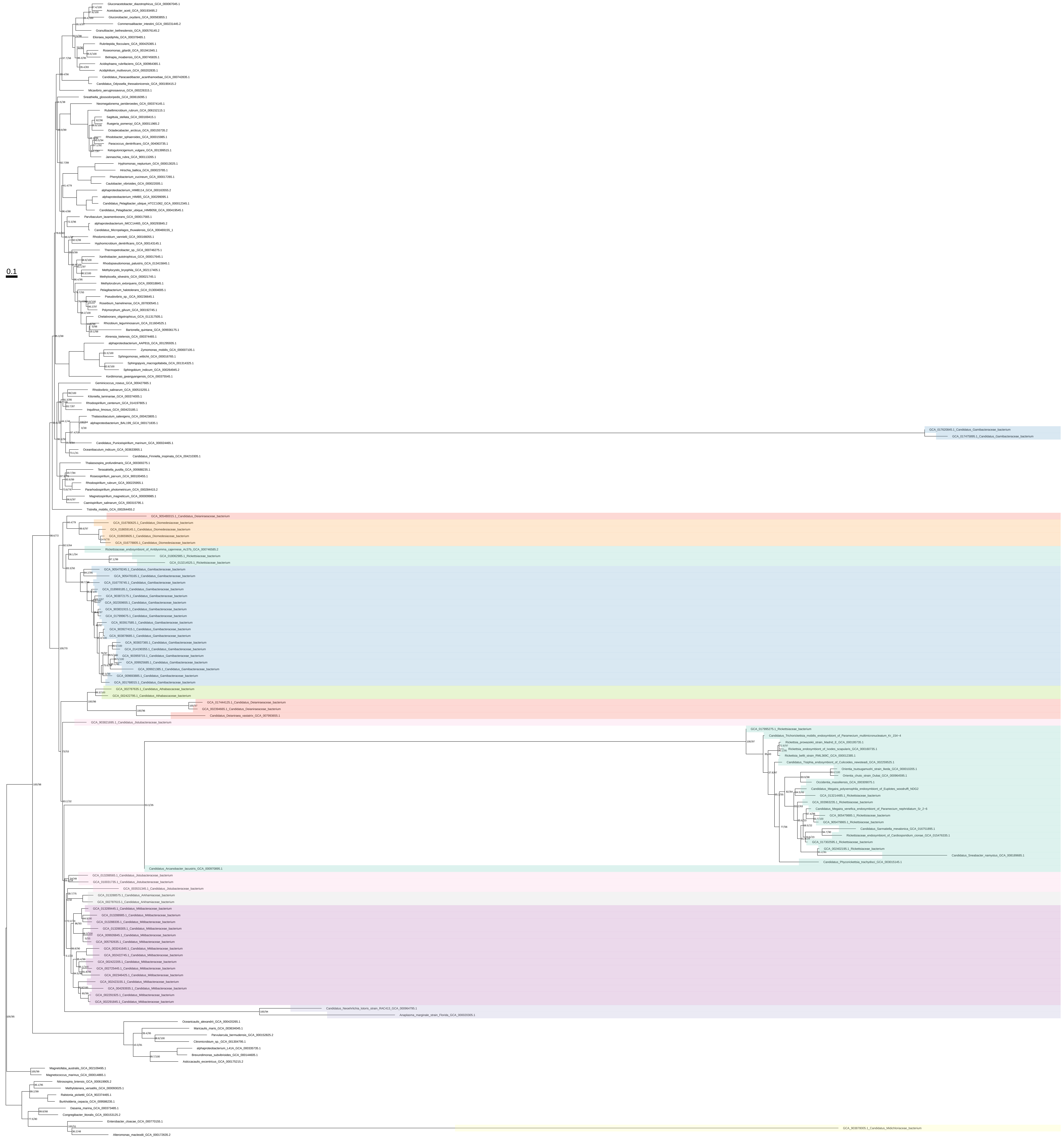

### Ile_Val_Leu_organism_selection

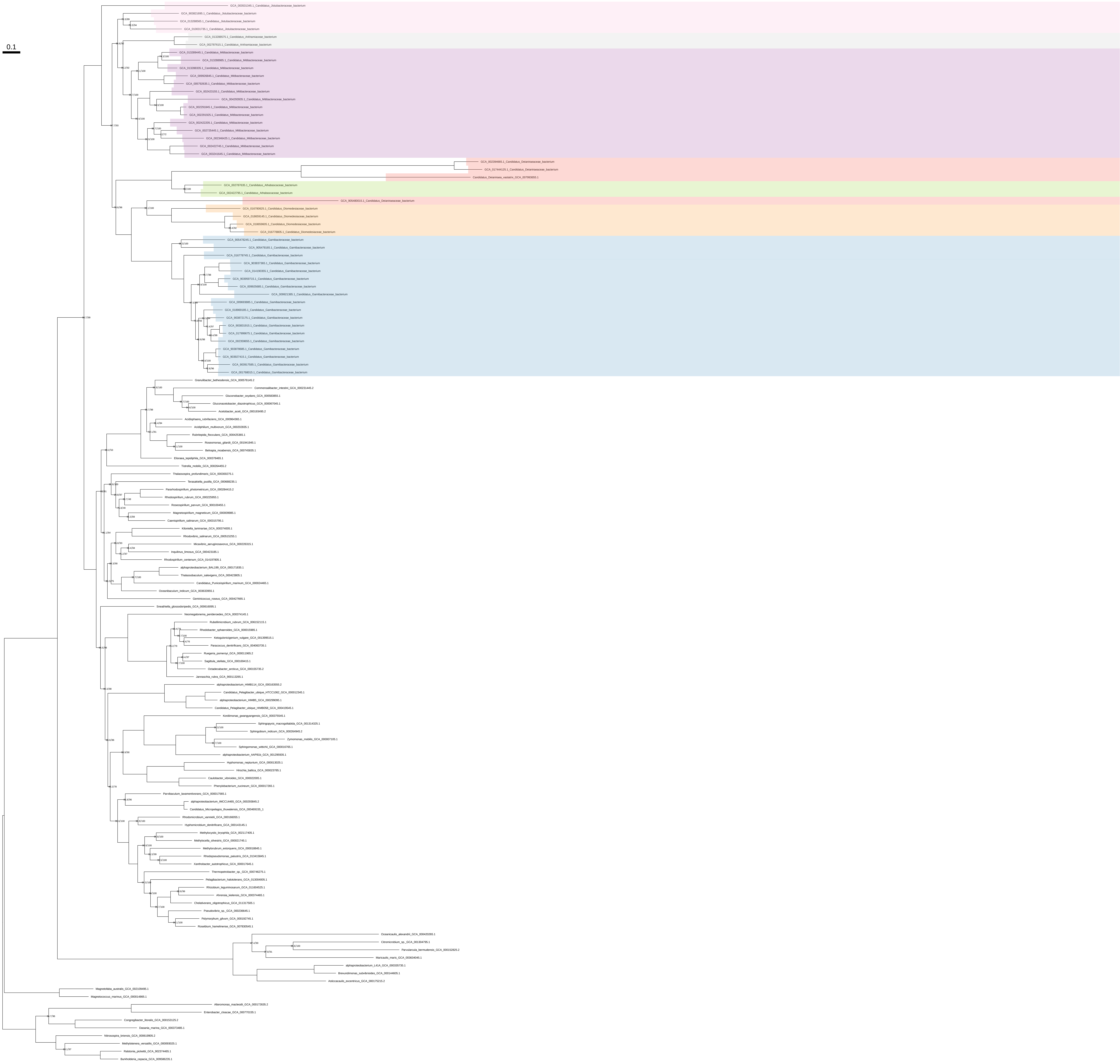

### Ile_Val_Leu_organism_selection_z_0.10

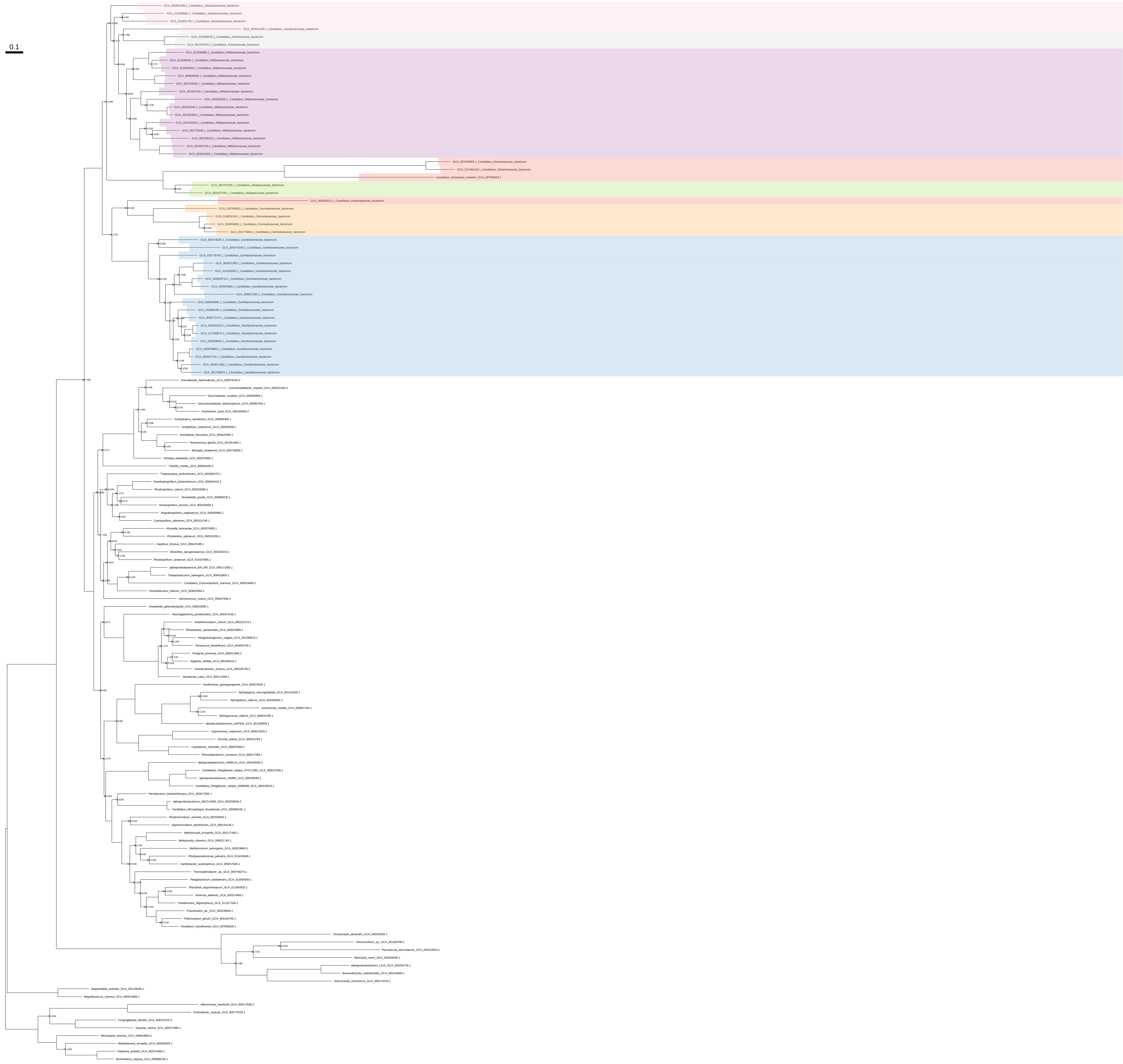

### Ile_Val_Leu_organism_selection_z_0.20

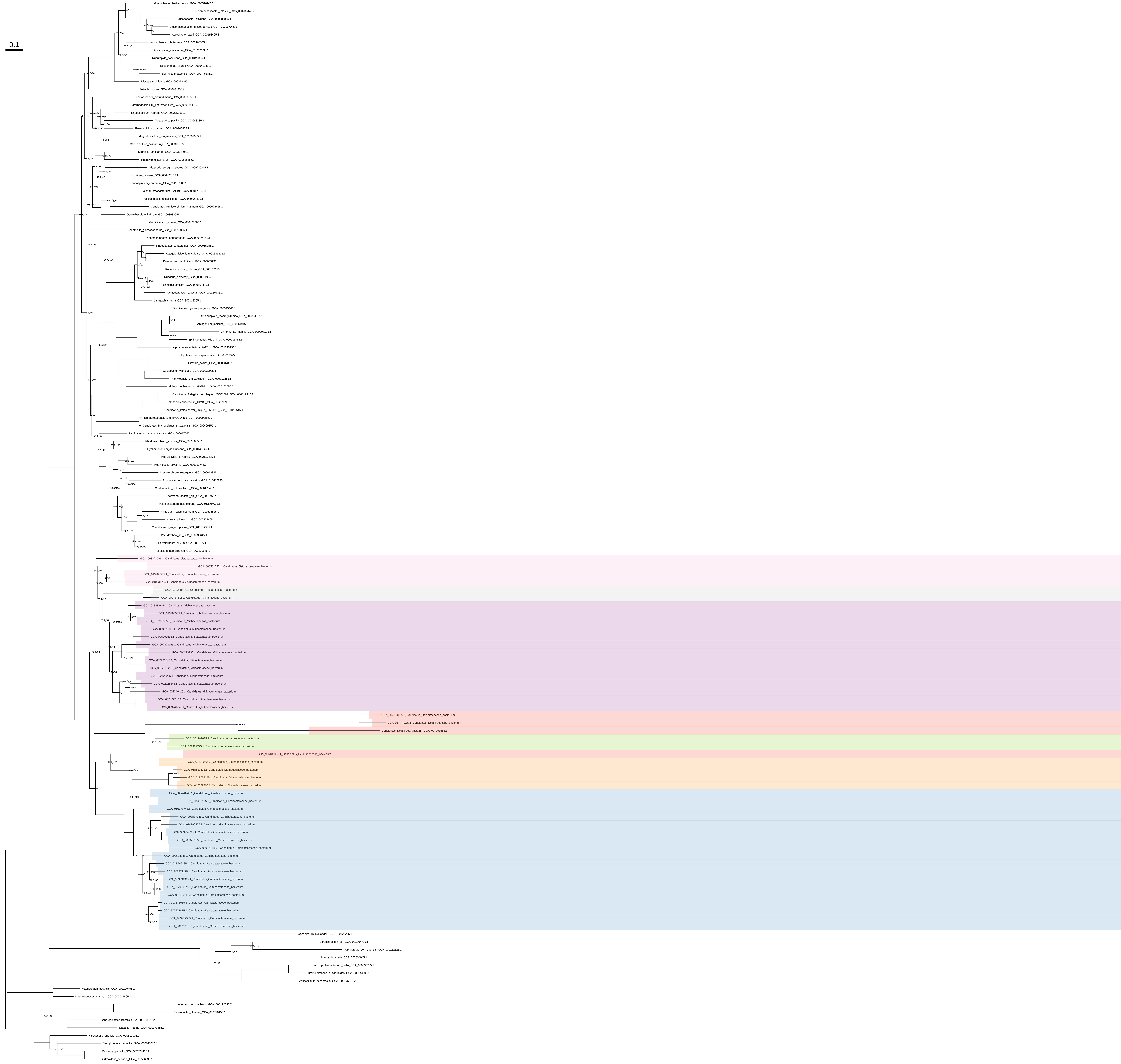

### Ile_Val_Leu_organism_selection_z_0.30

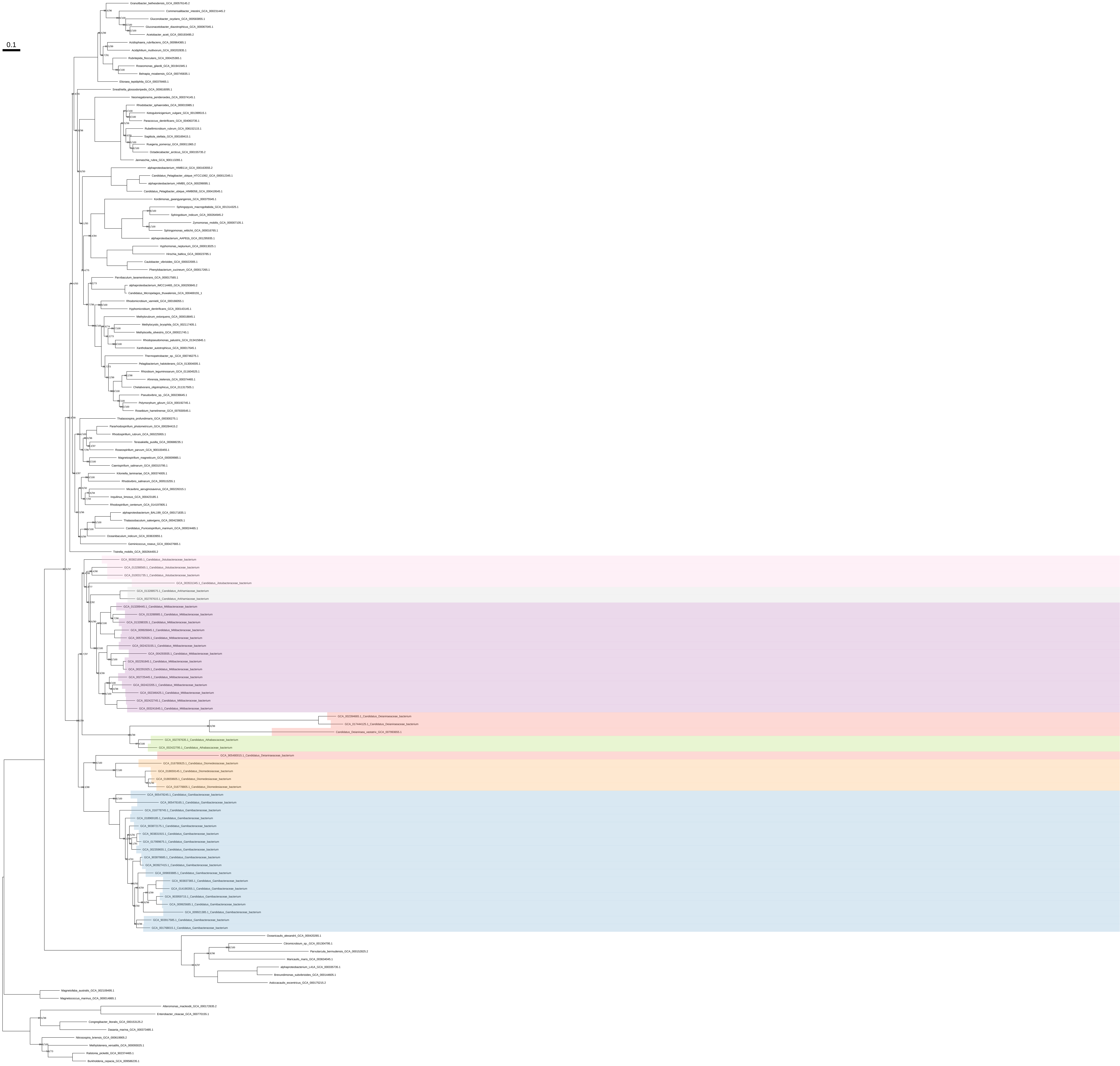

### Ile_Val_Leu_organism_selection_z_0.40

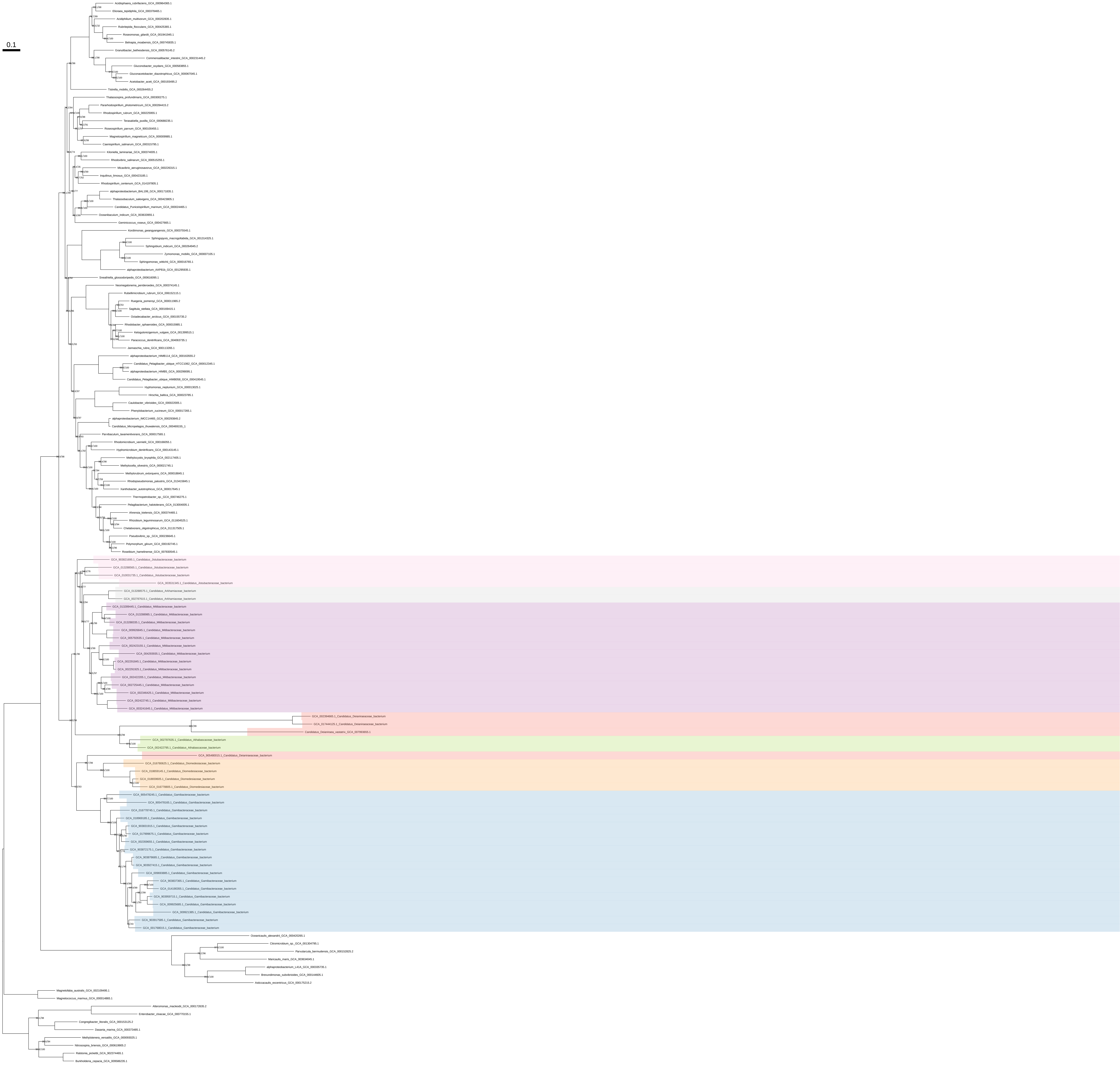

### Ile_Val_Leu_organism_selection_z_0.50

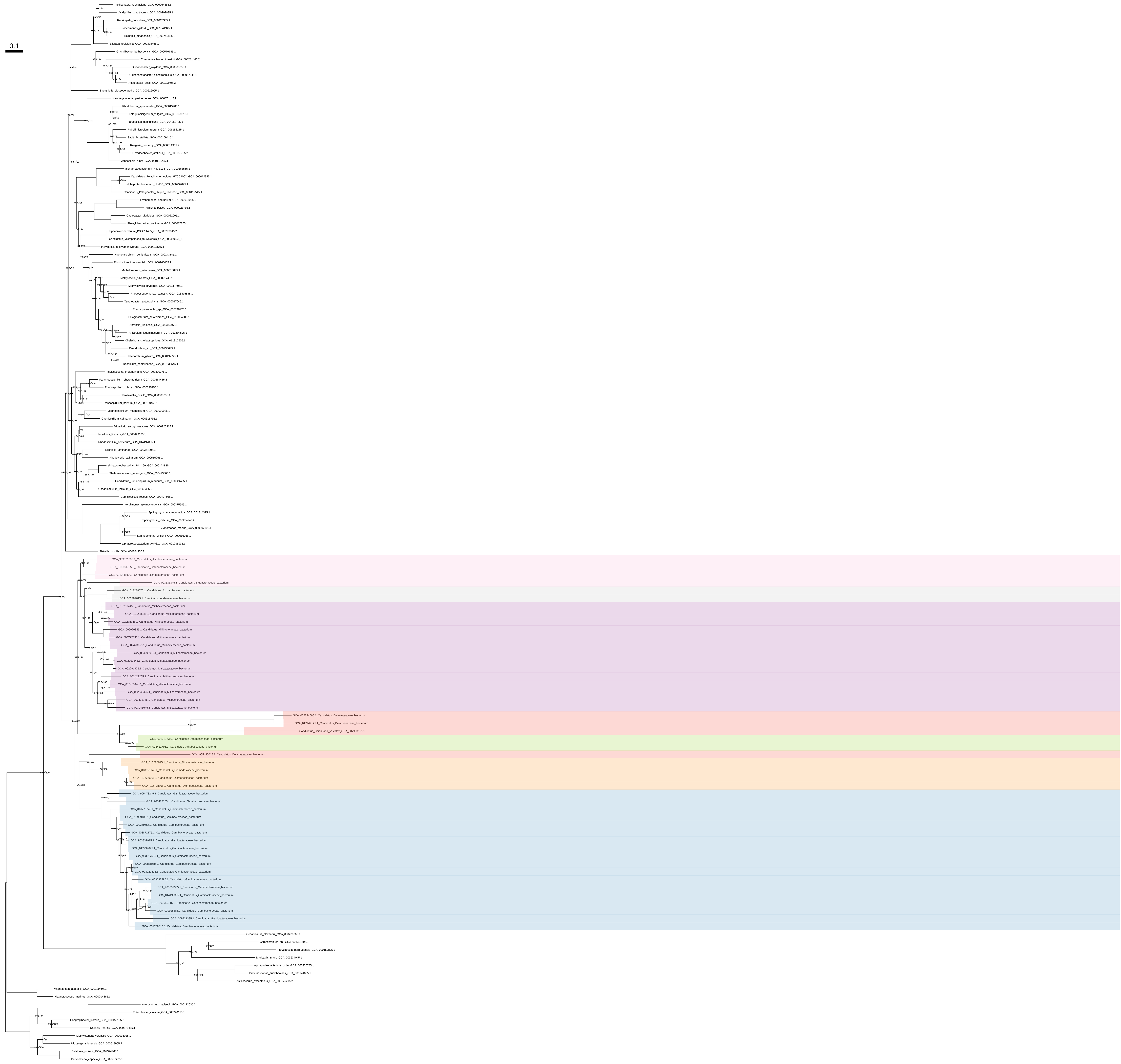

### Lys_Thr_Met_all_organisms

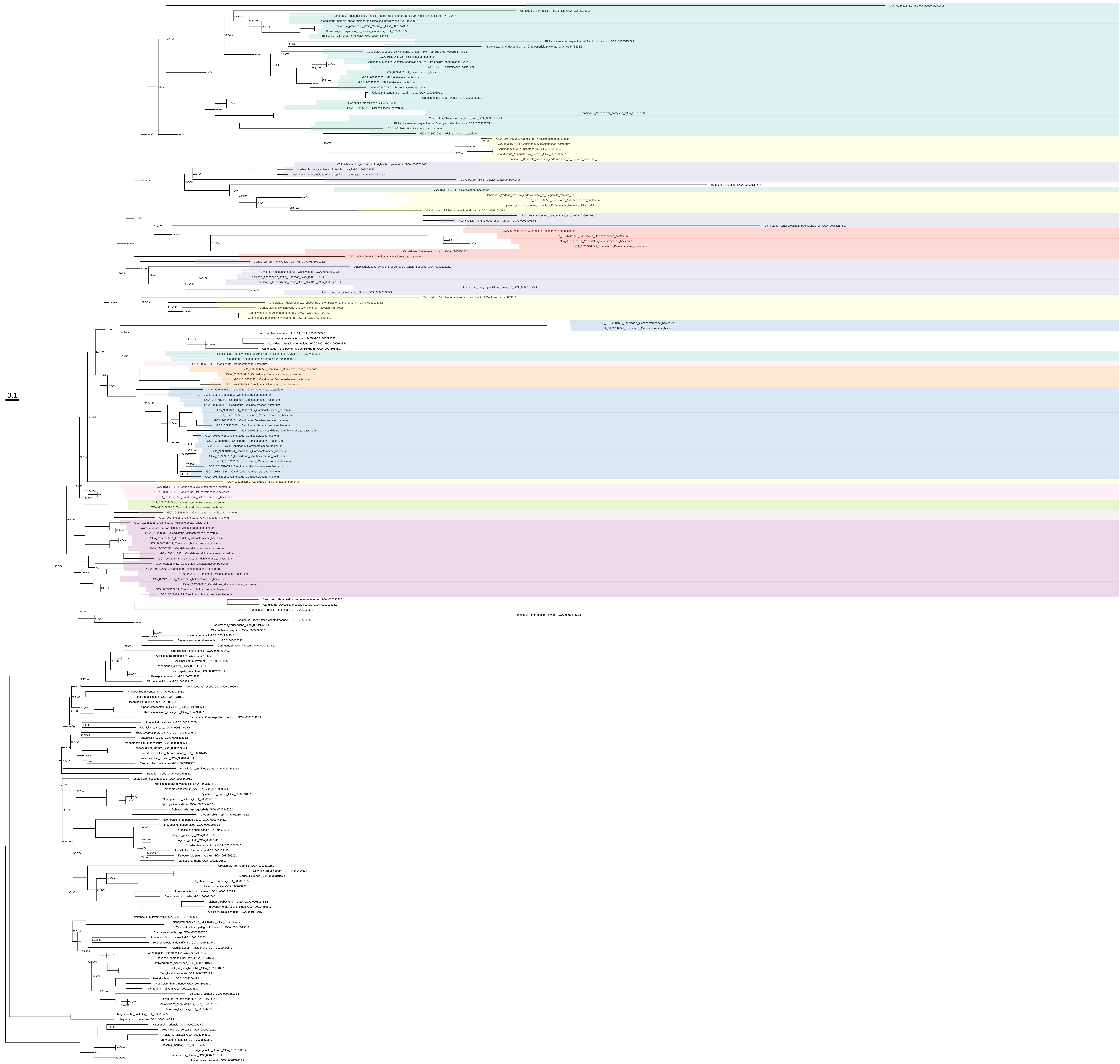

Lys_Thr_Met_all_organisms_z_0.10

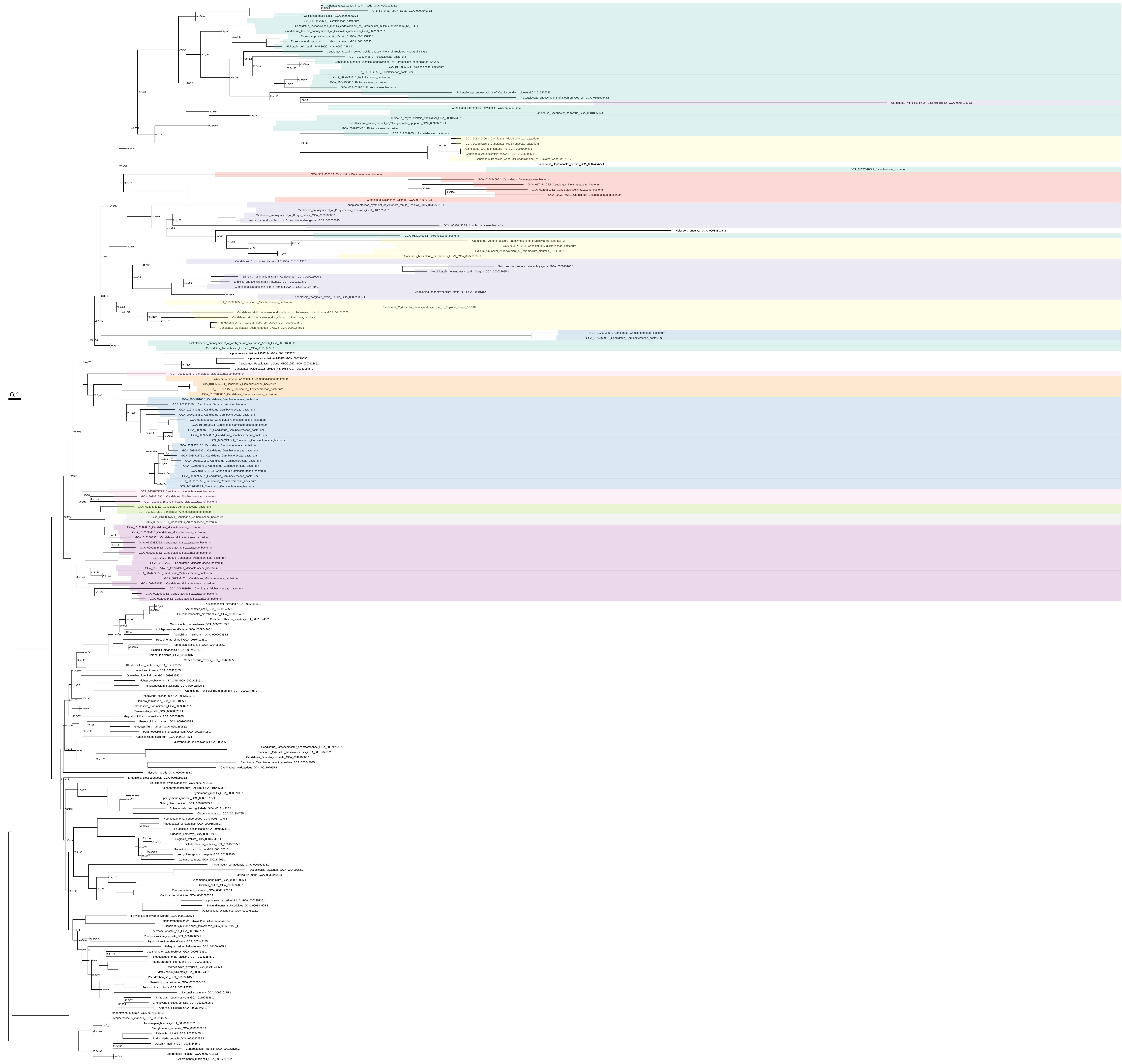

Lys_Thr_Met_all_organisms_z_0.20

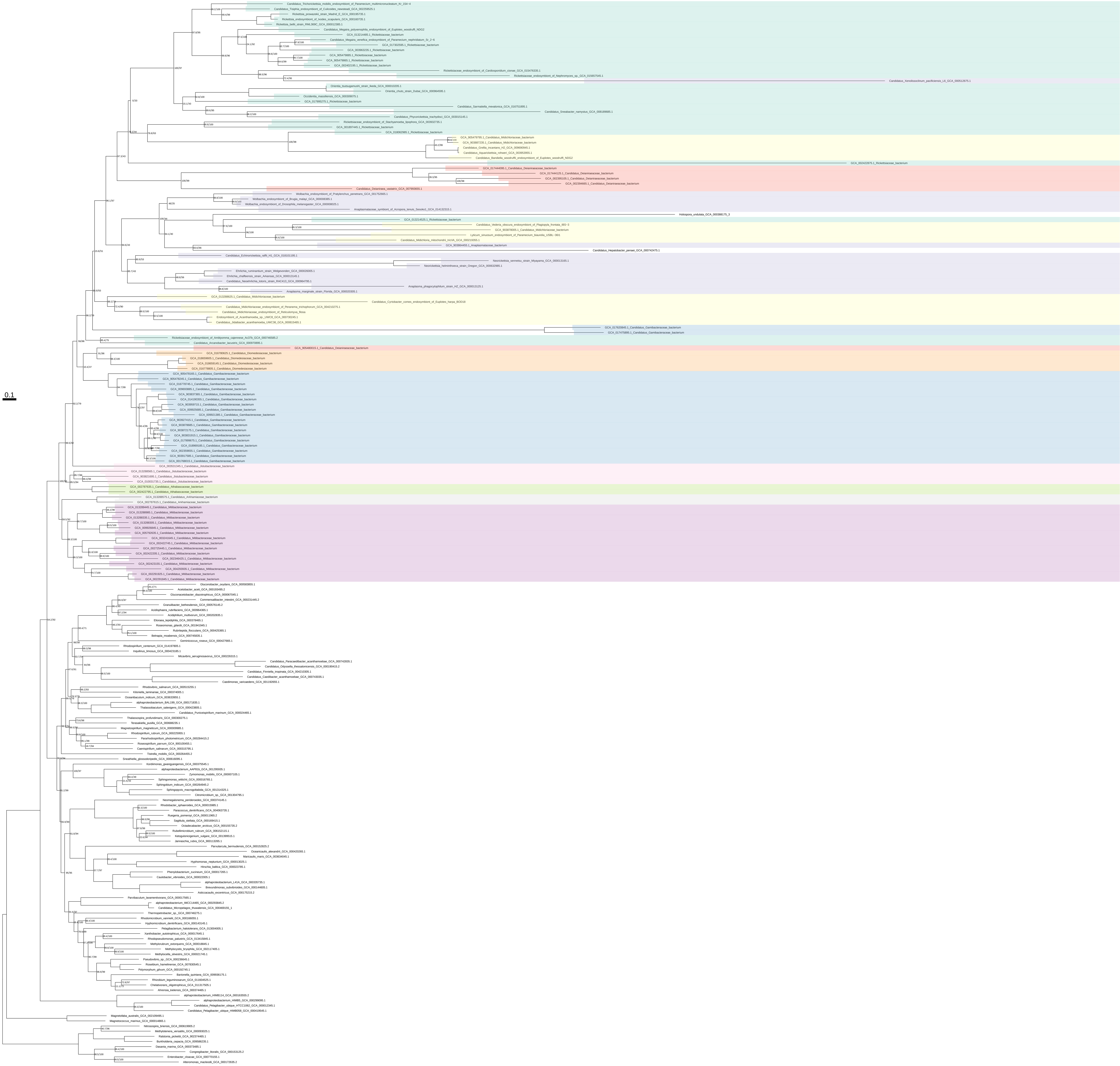

### Lys_Thr_Met_all_organisms_z_0.30

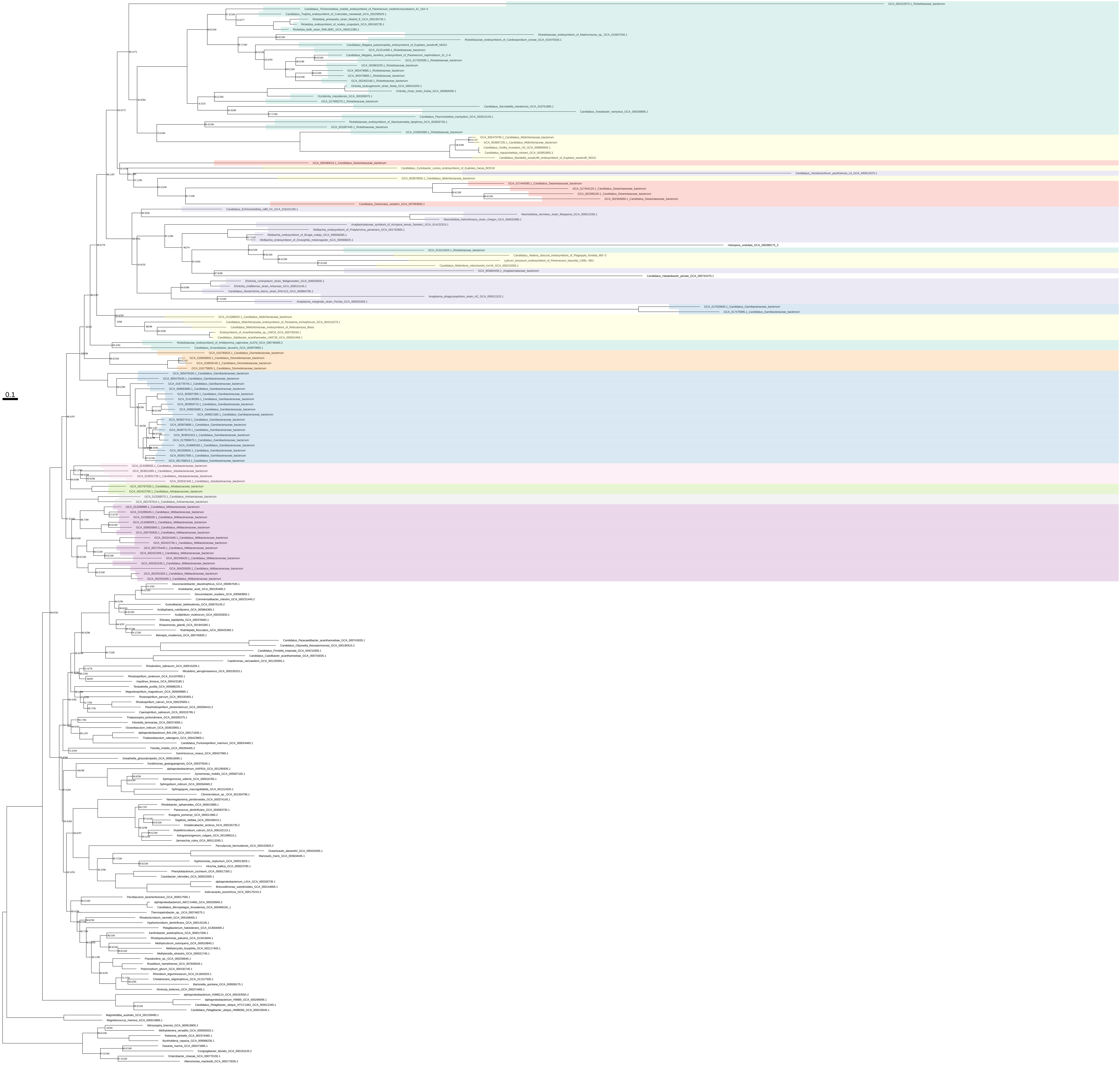

Lys_Thr_Met_all_organisms_z_0.40

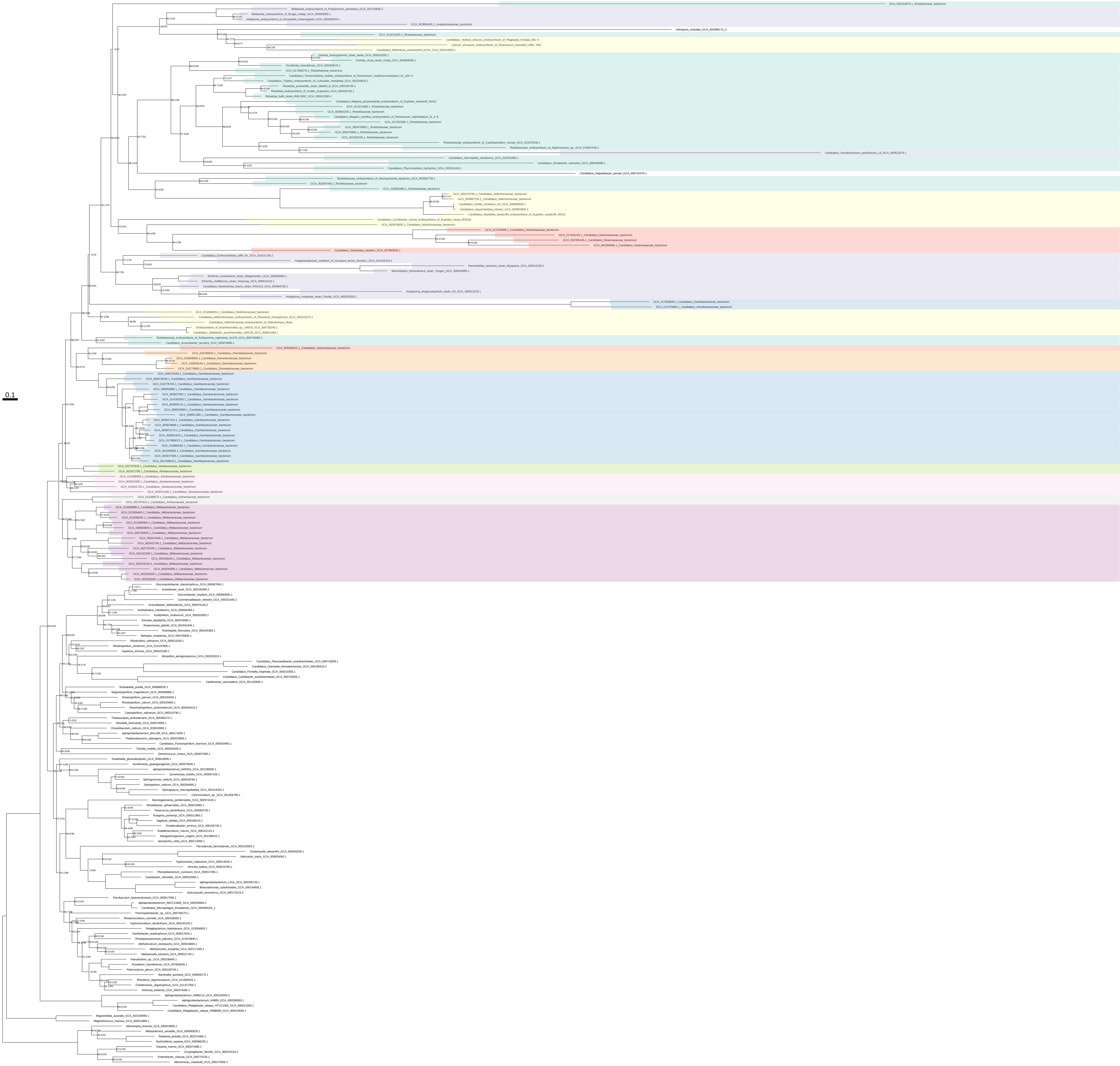

### Lys_Thr_Met_all_organisms_z_0.50

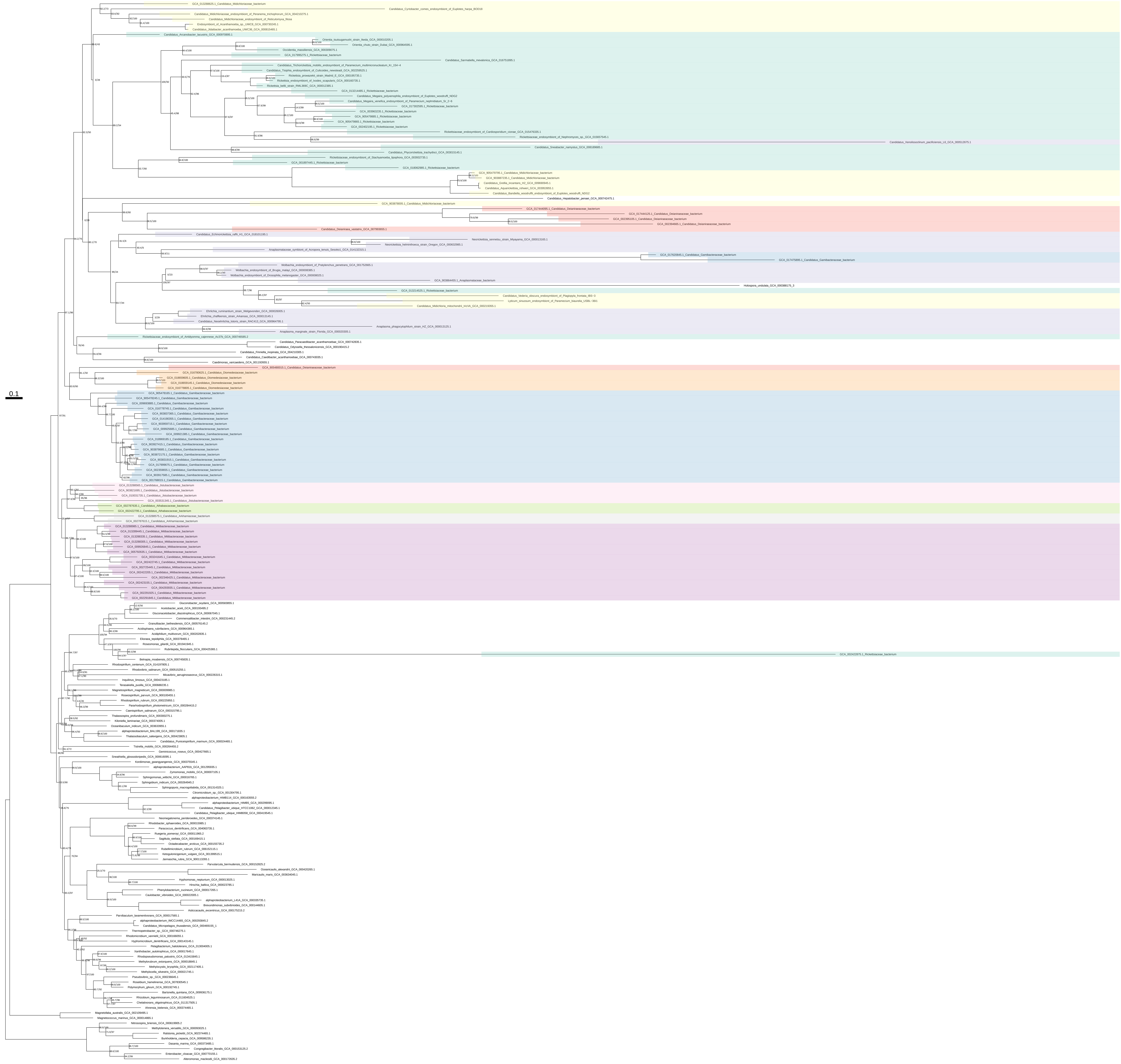

### Lys_Thr_Met_organism_selection

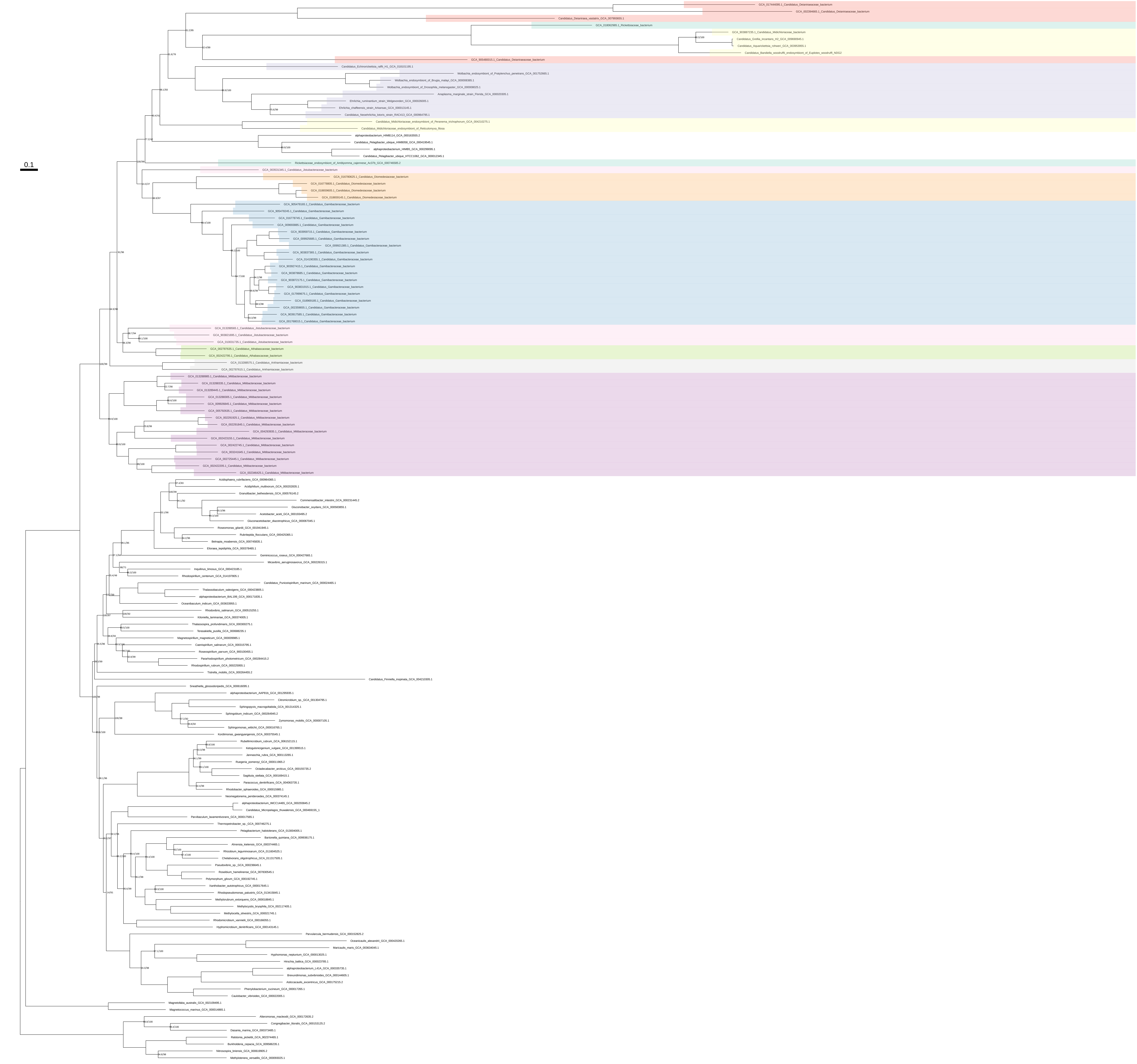

### Lys_Thr_Met_organism_selection_z_0.10

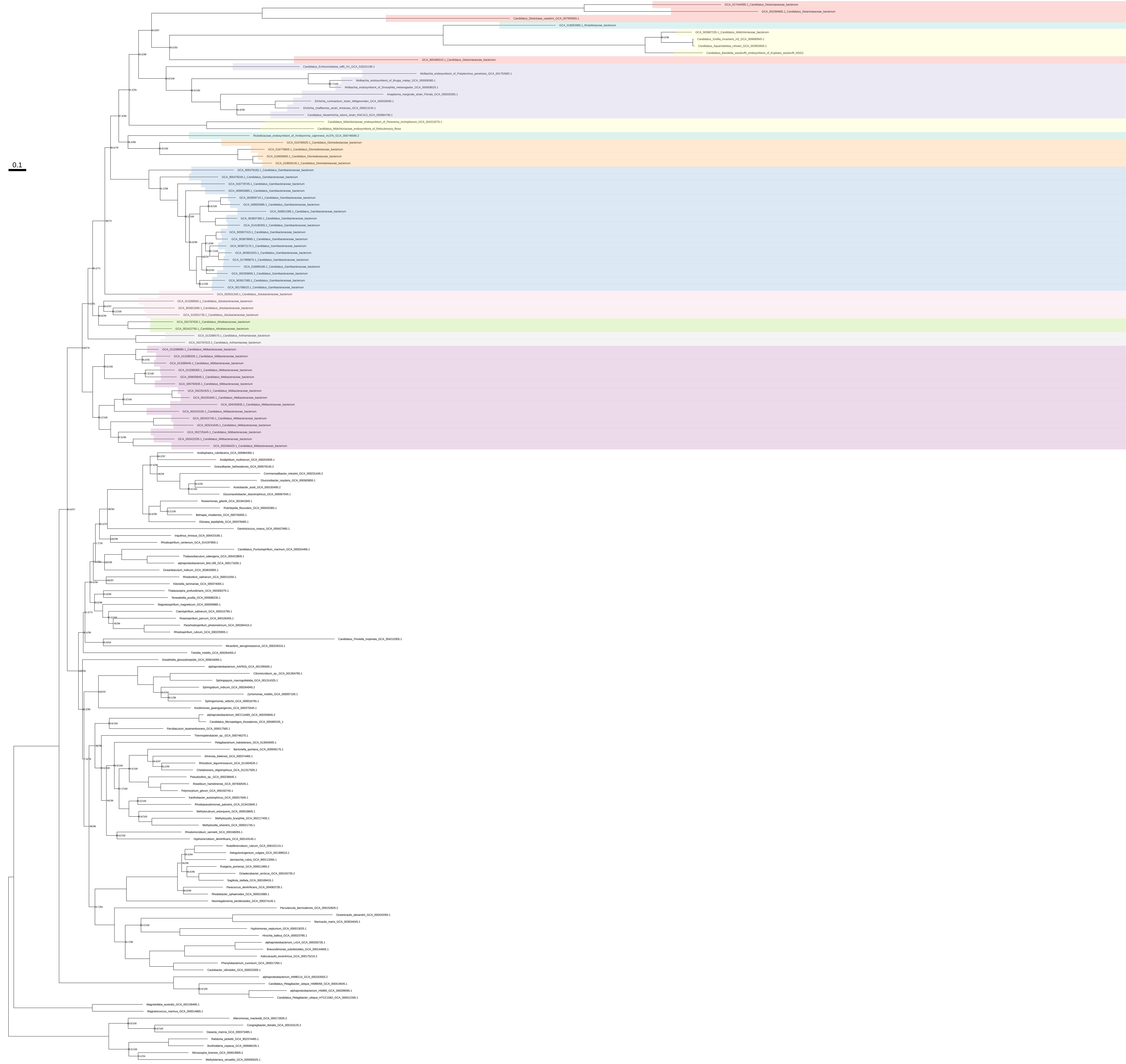

### Lys_Thr_Met_organism_selection_z_0.20

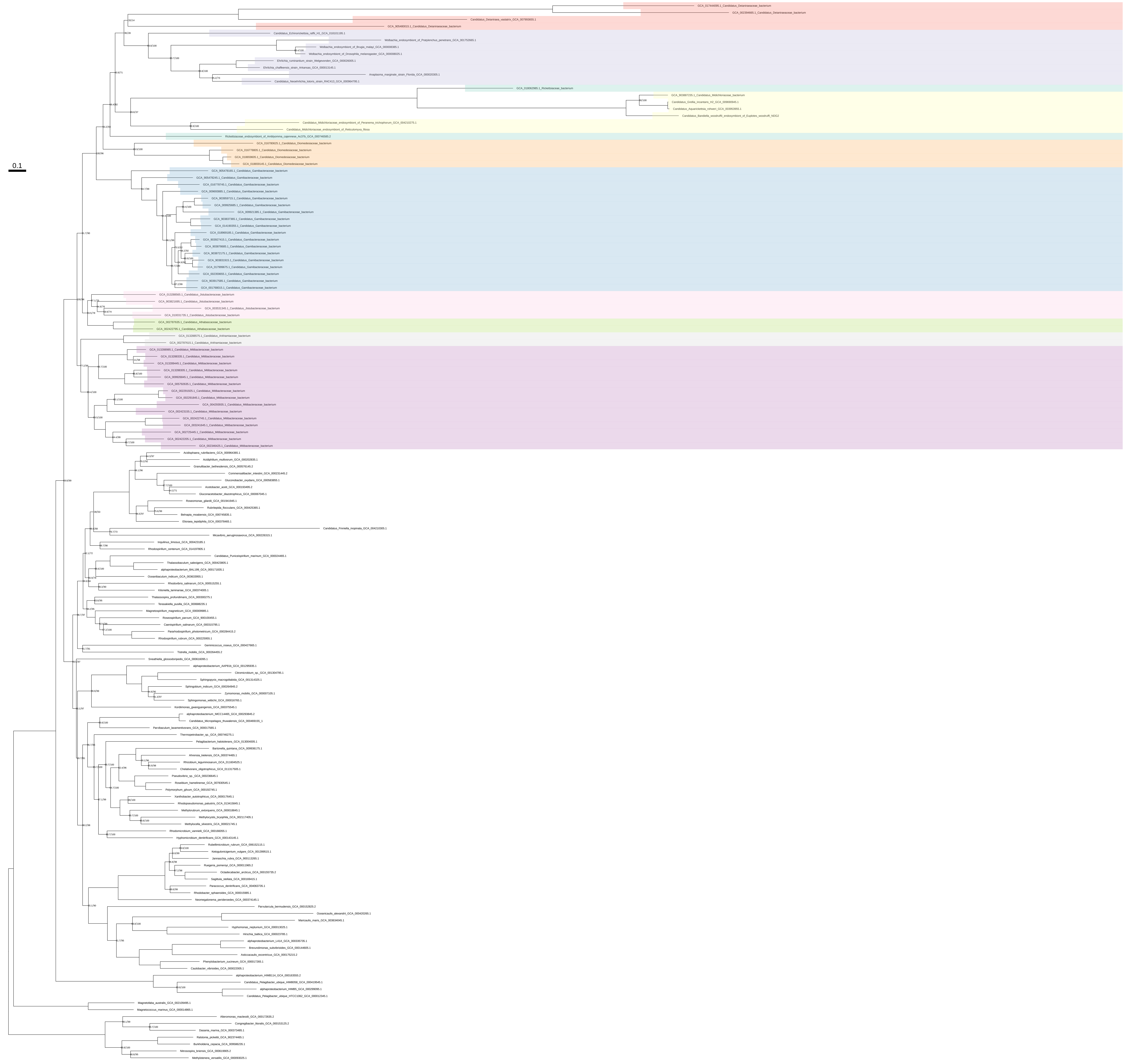

### Lys_Thr_Met_organism_selection_z_0.30

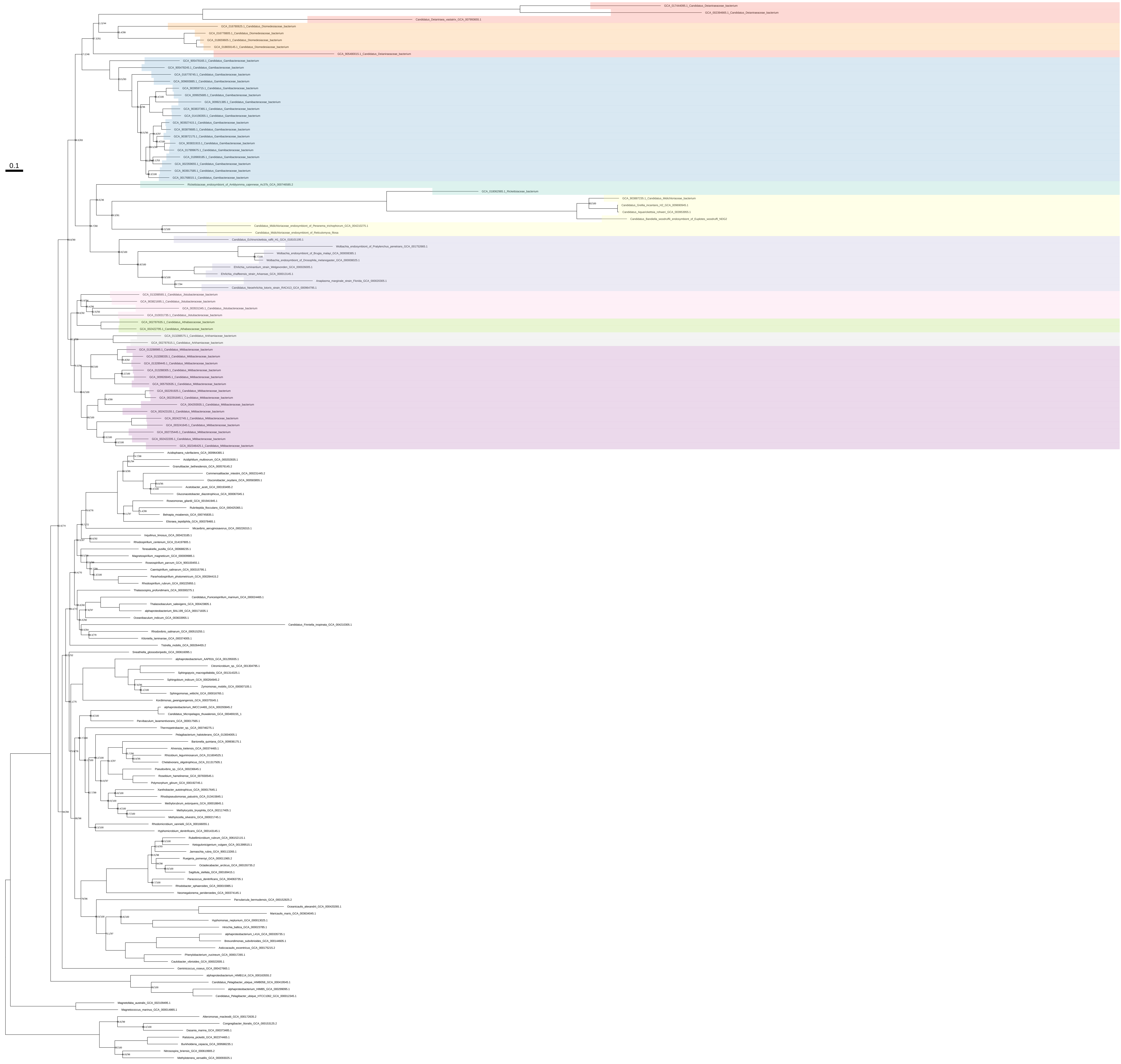

### Lys_Thr_Met_organism_selection_z_0.40

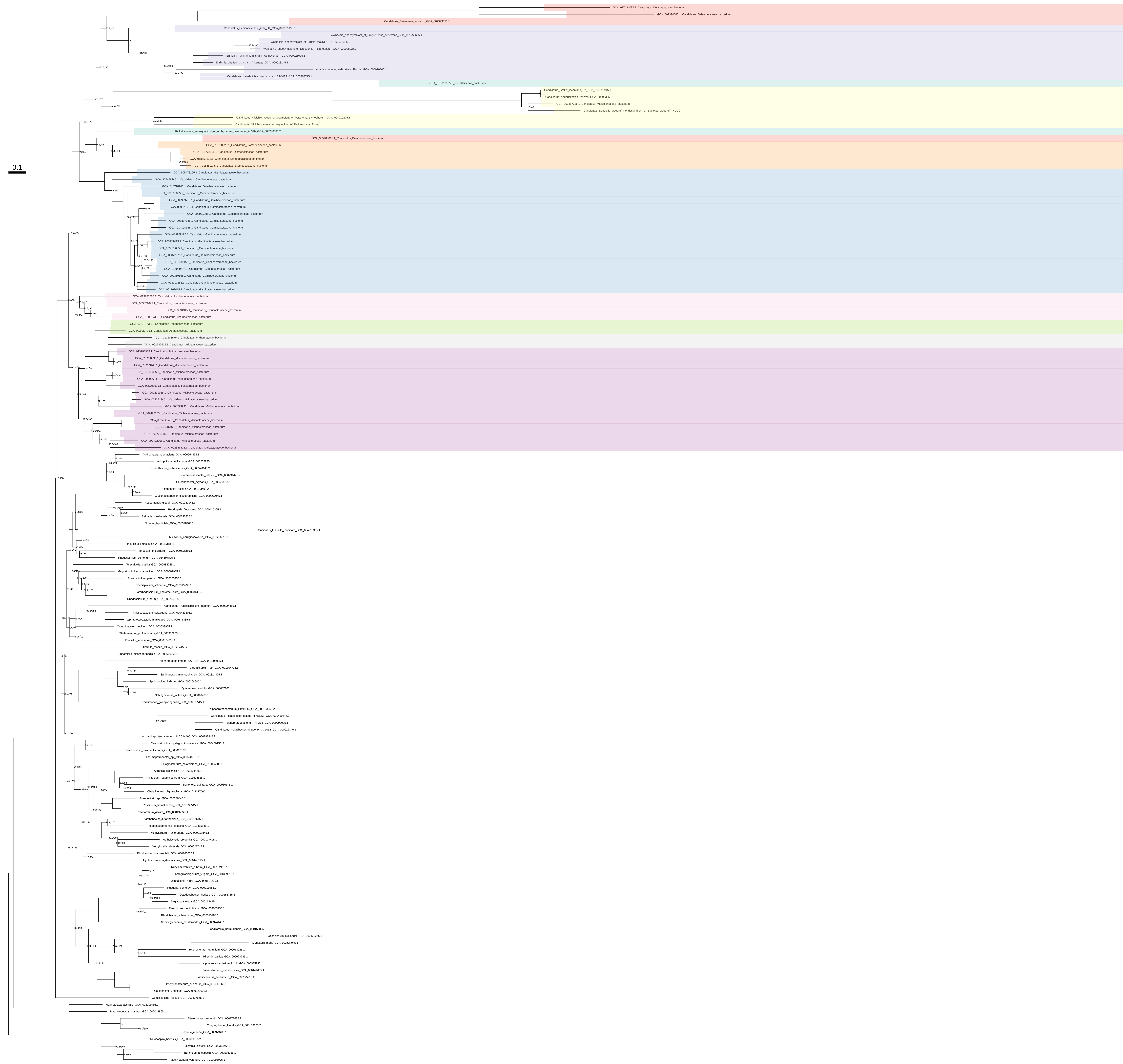

### Lys_Thr_Met_organism_selection_z_0.50

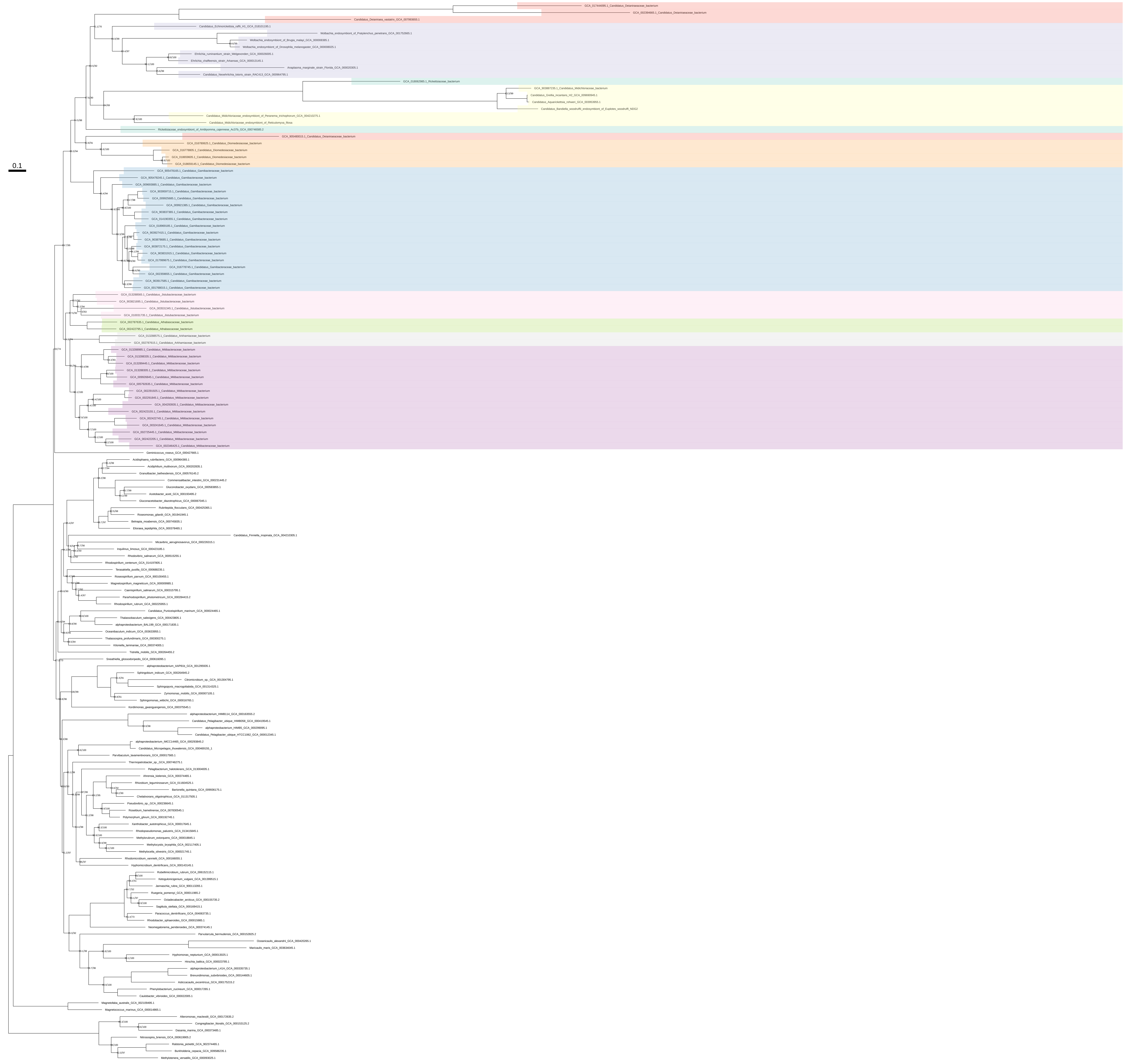

### Pro_all_organisms

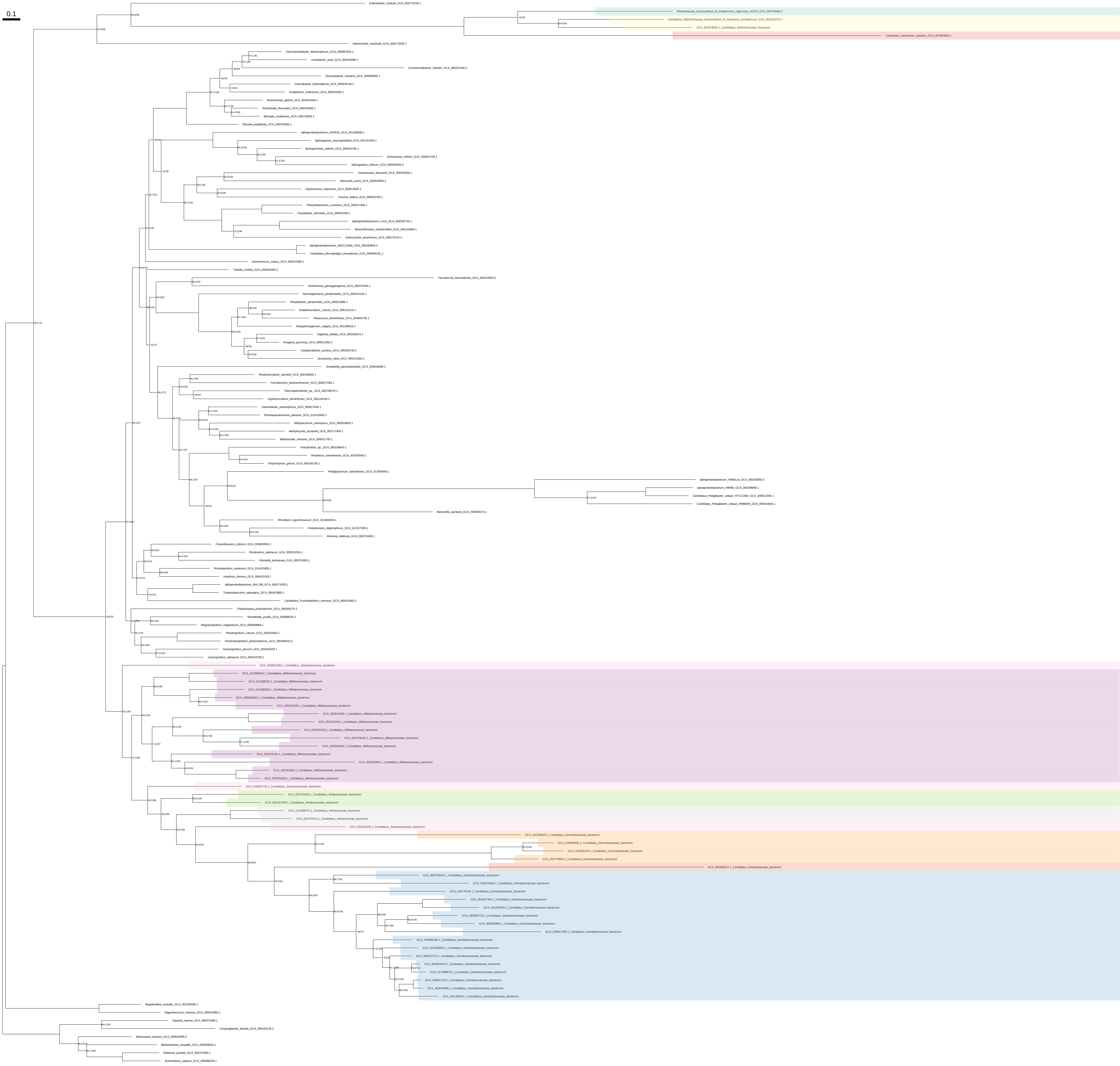

### Pro_all_organisms_z_0.10

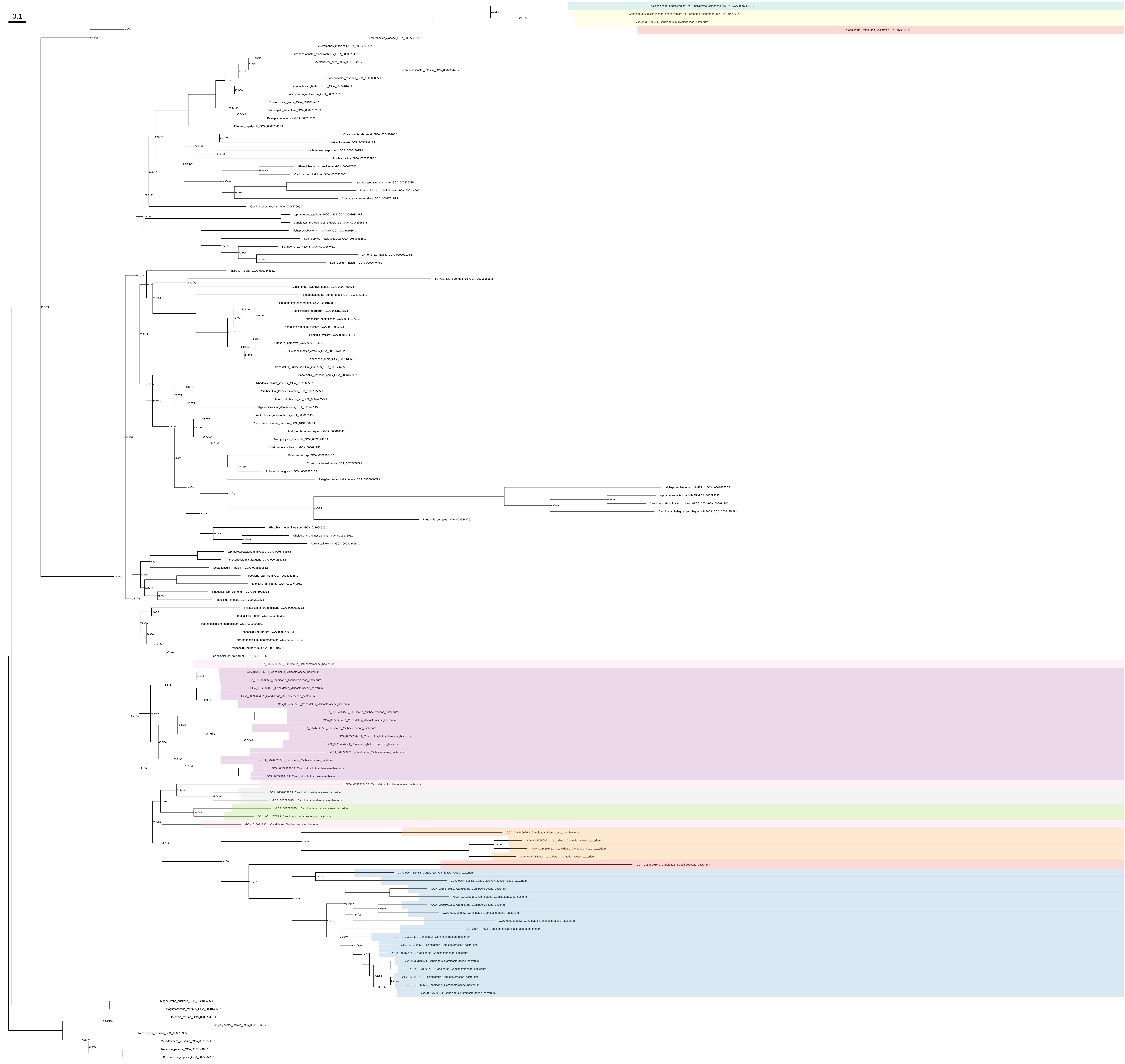

Pro_all_organisms_z_0.20

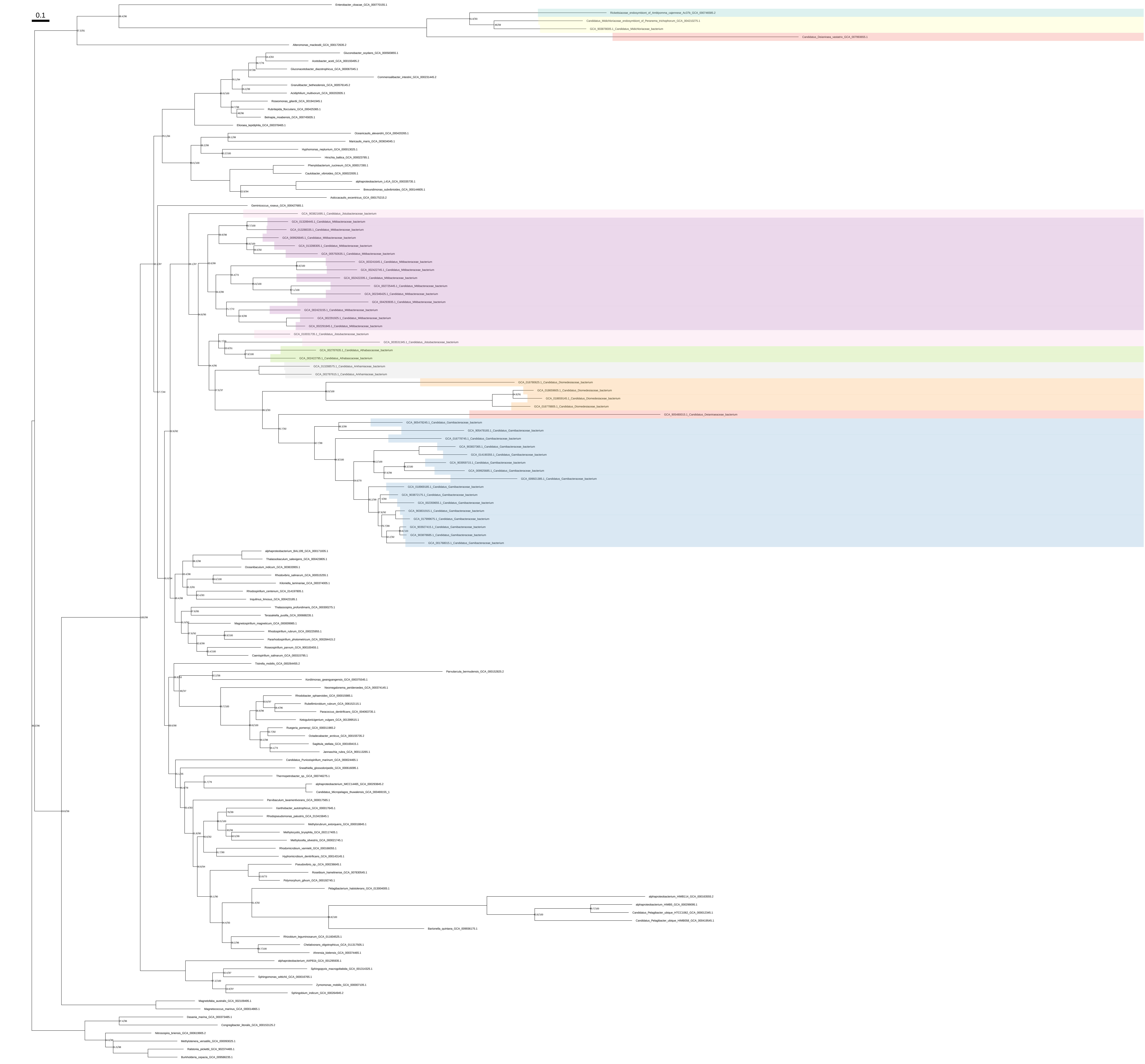

### Pro_all_organisms_z_0.30

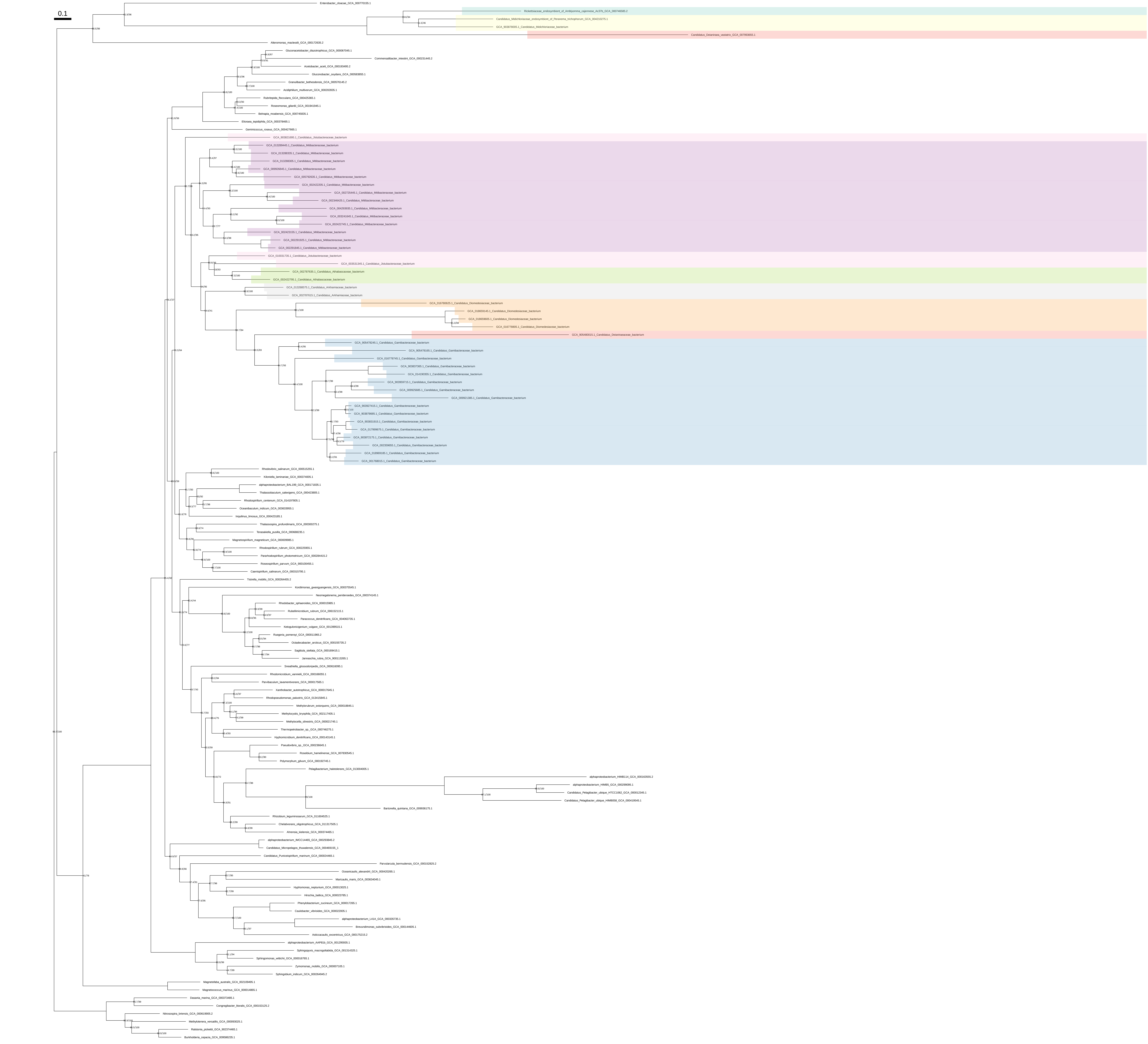

### Pro_all_organisms_z_0.40

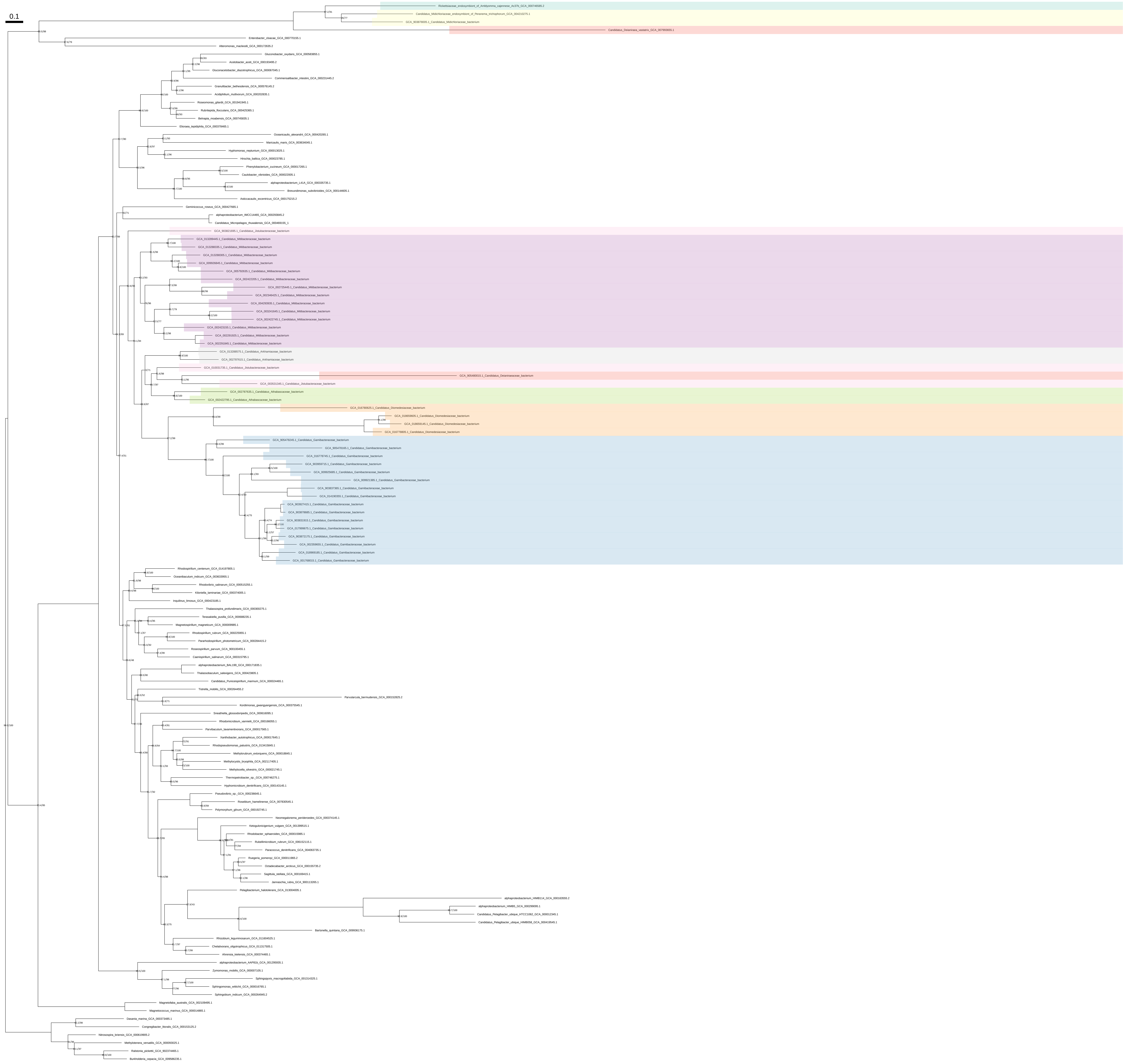

### Pro_all_organisms_z_0.50

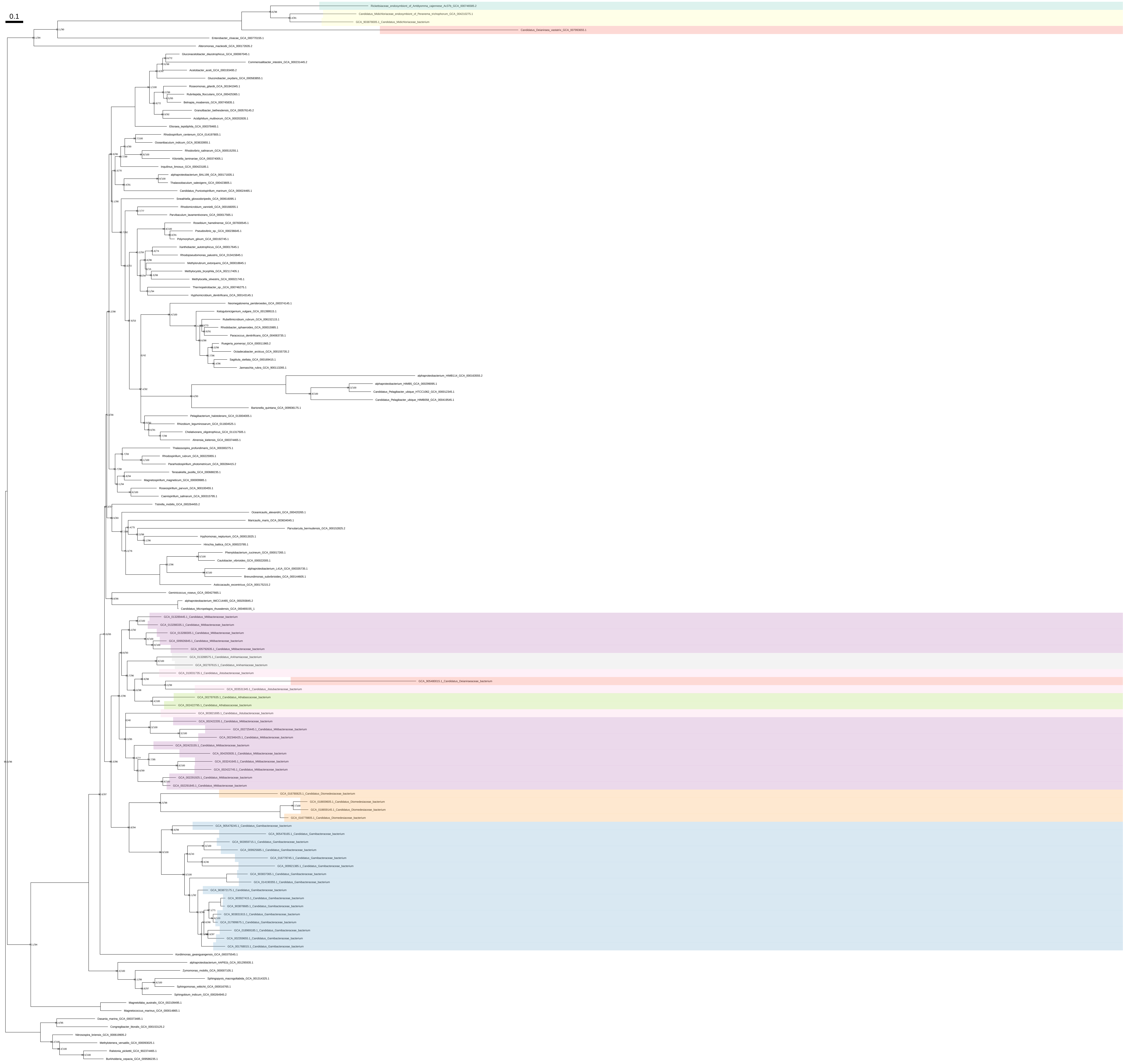

### Ser_all_organisms

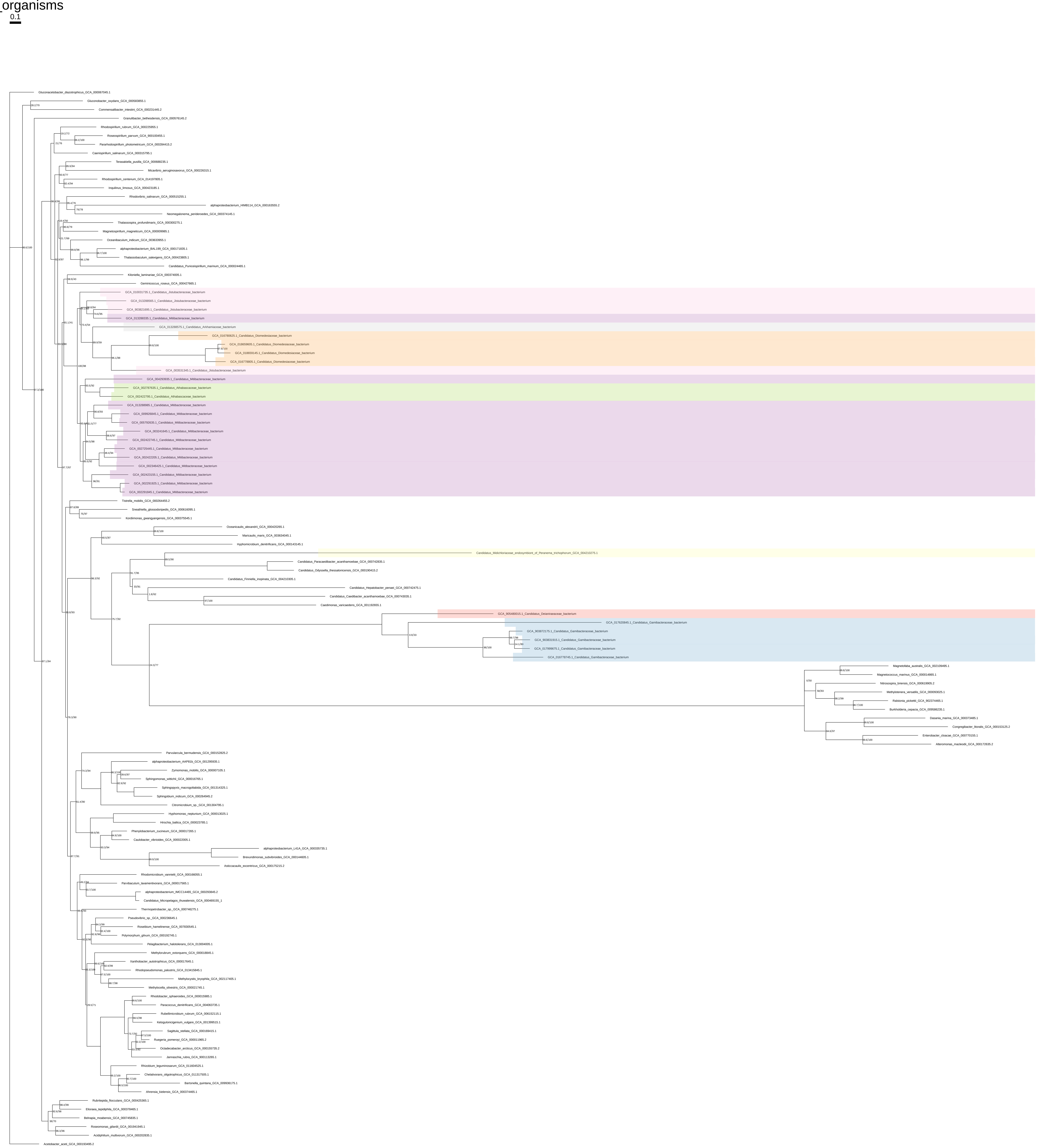

### Ser_all_organisms_z_0.10

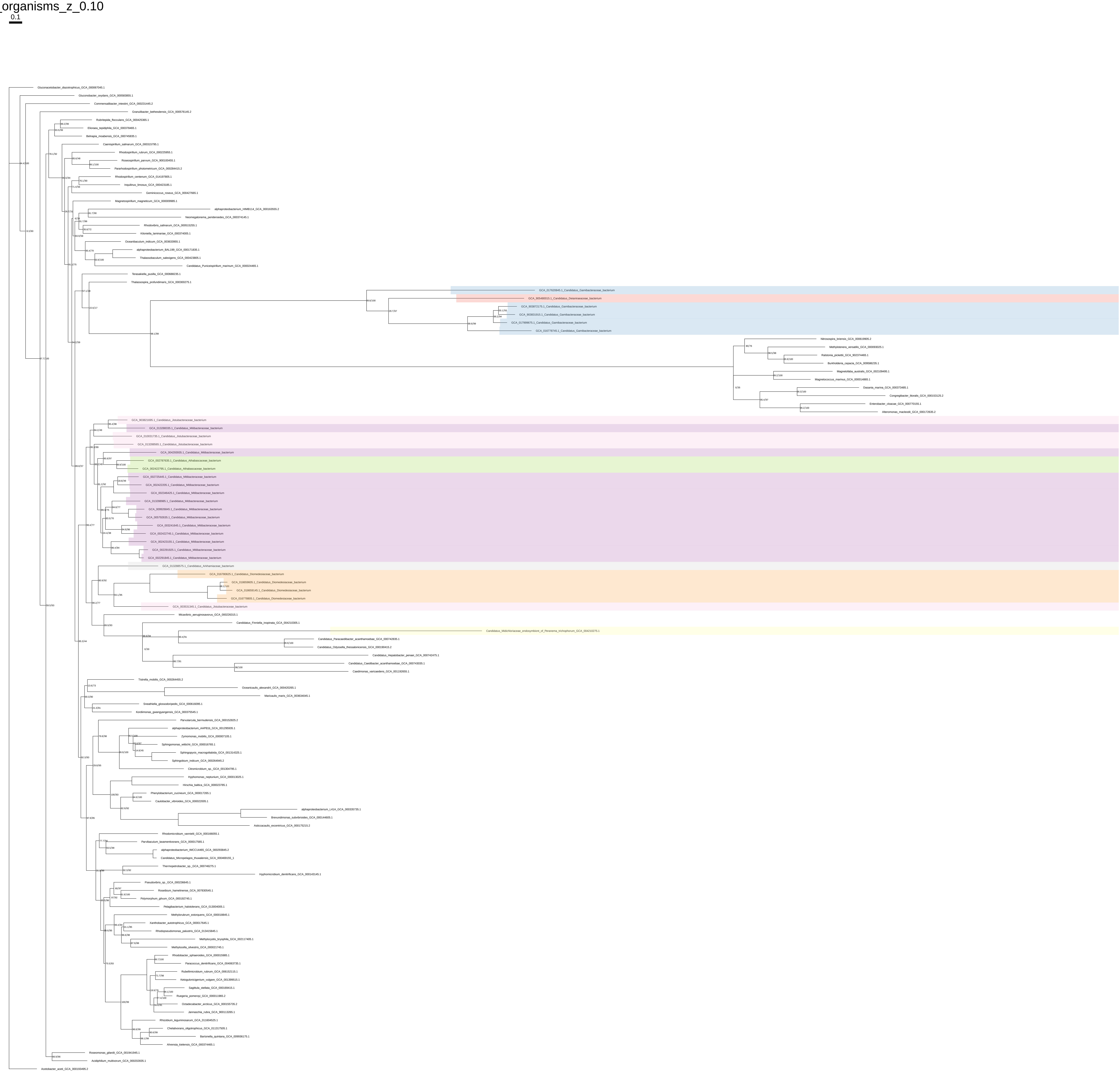

### Ser_all_organisms_z_0.20

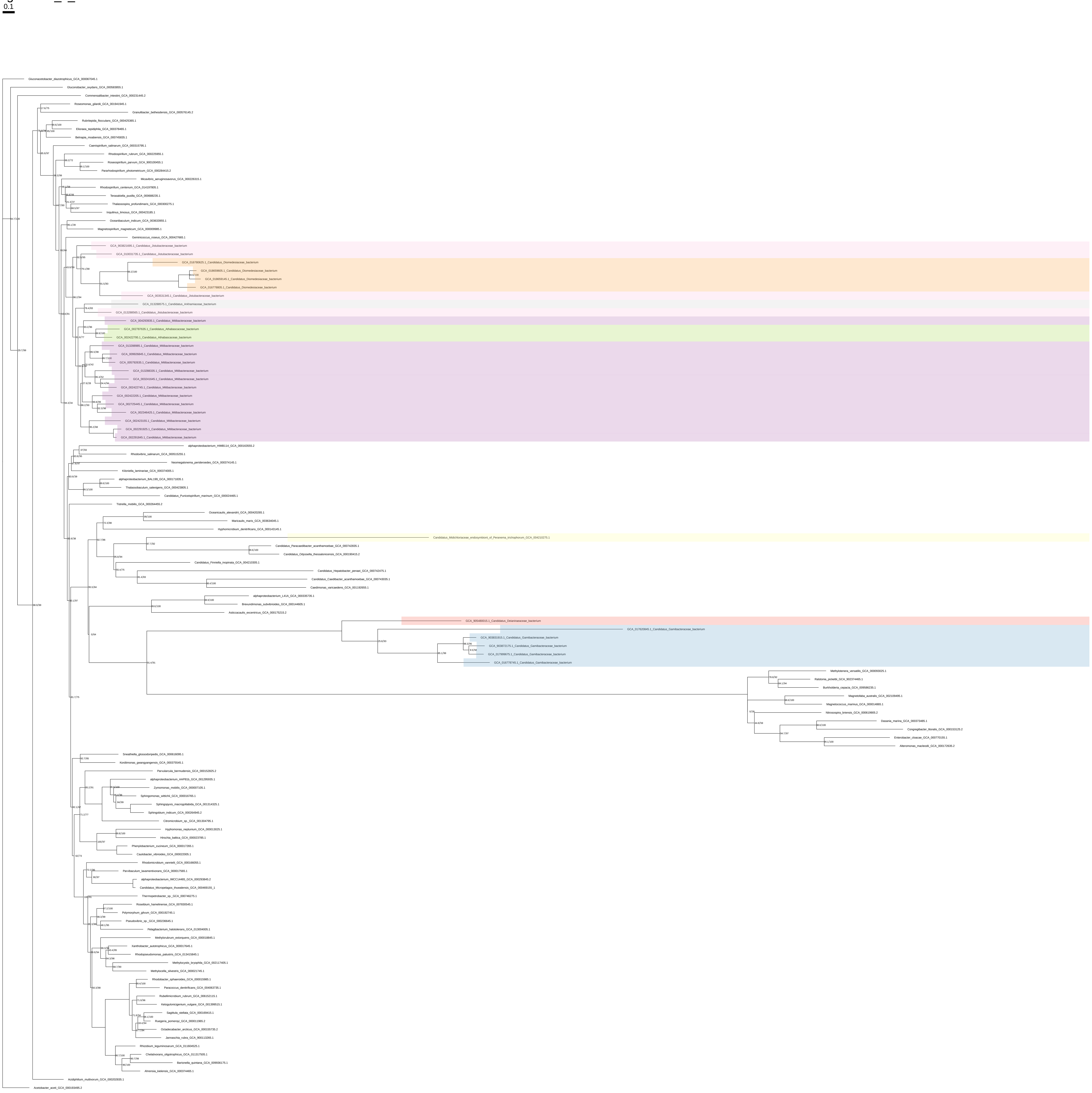

### Ser_all_organisms_z_0.30

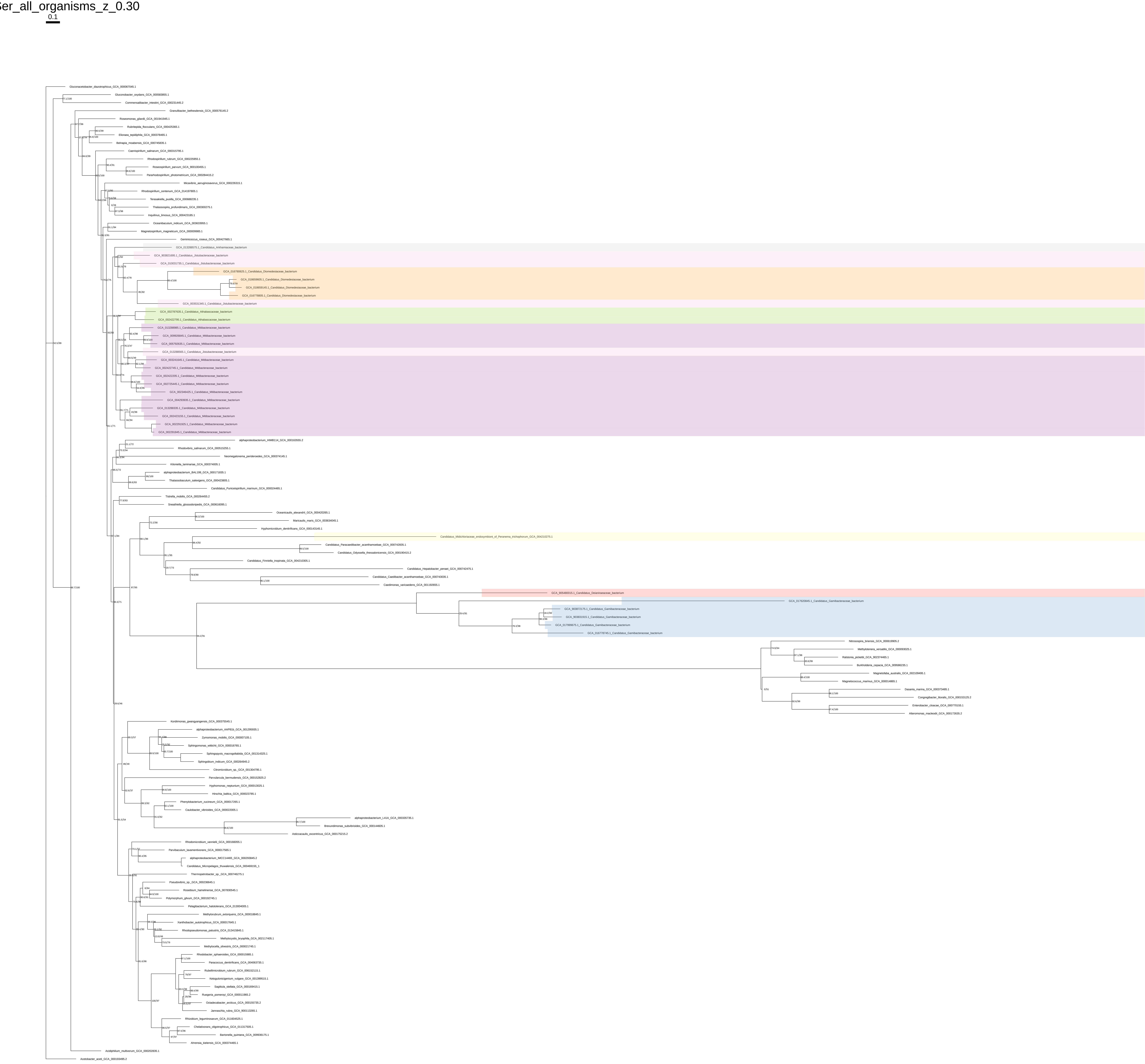

### Ser_all_organisms_z_0.40

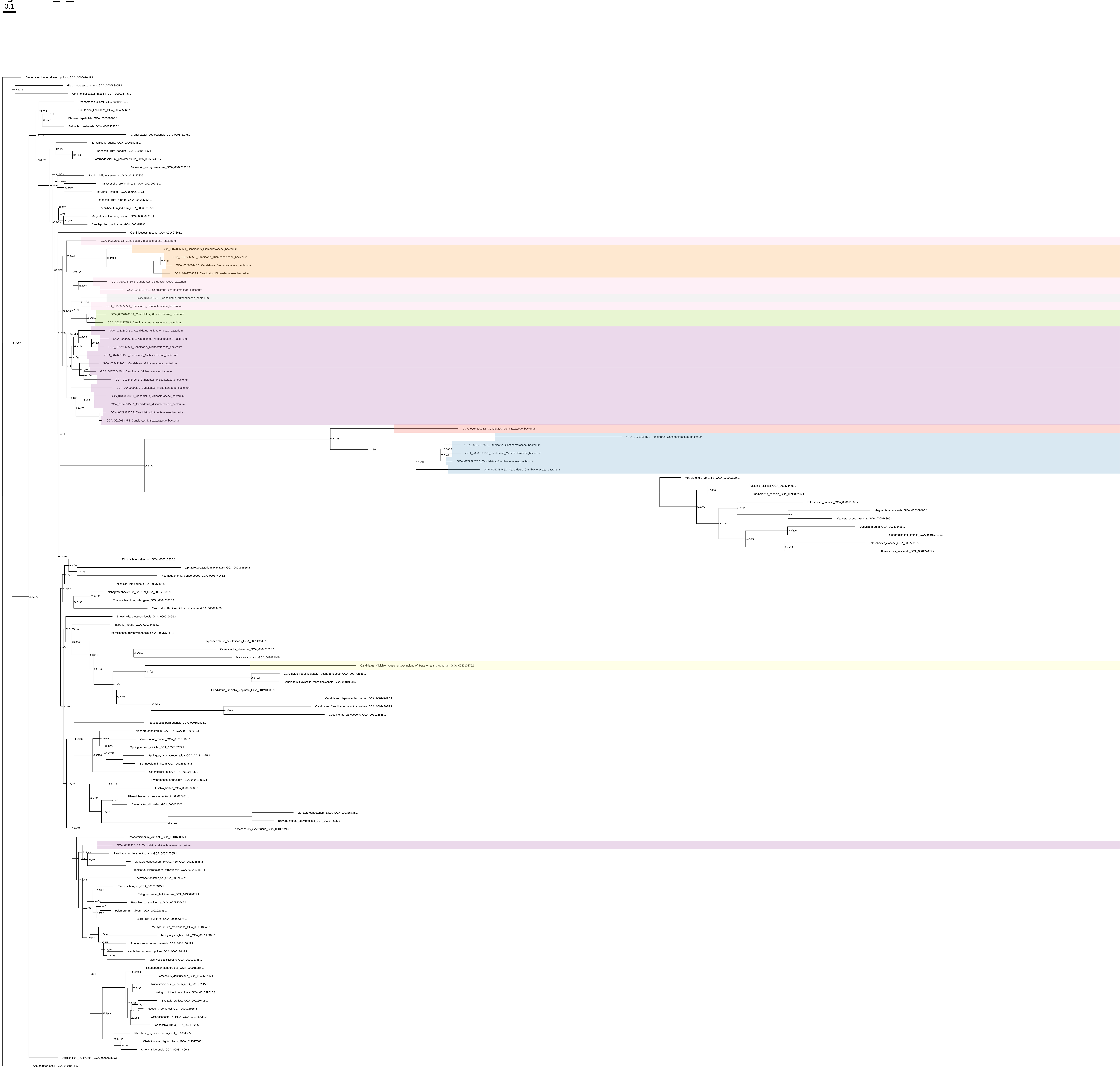

### Ser_all_organisms_z_0.50

0.1

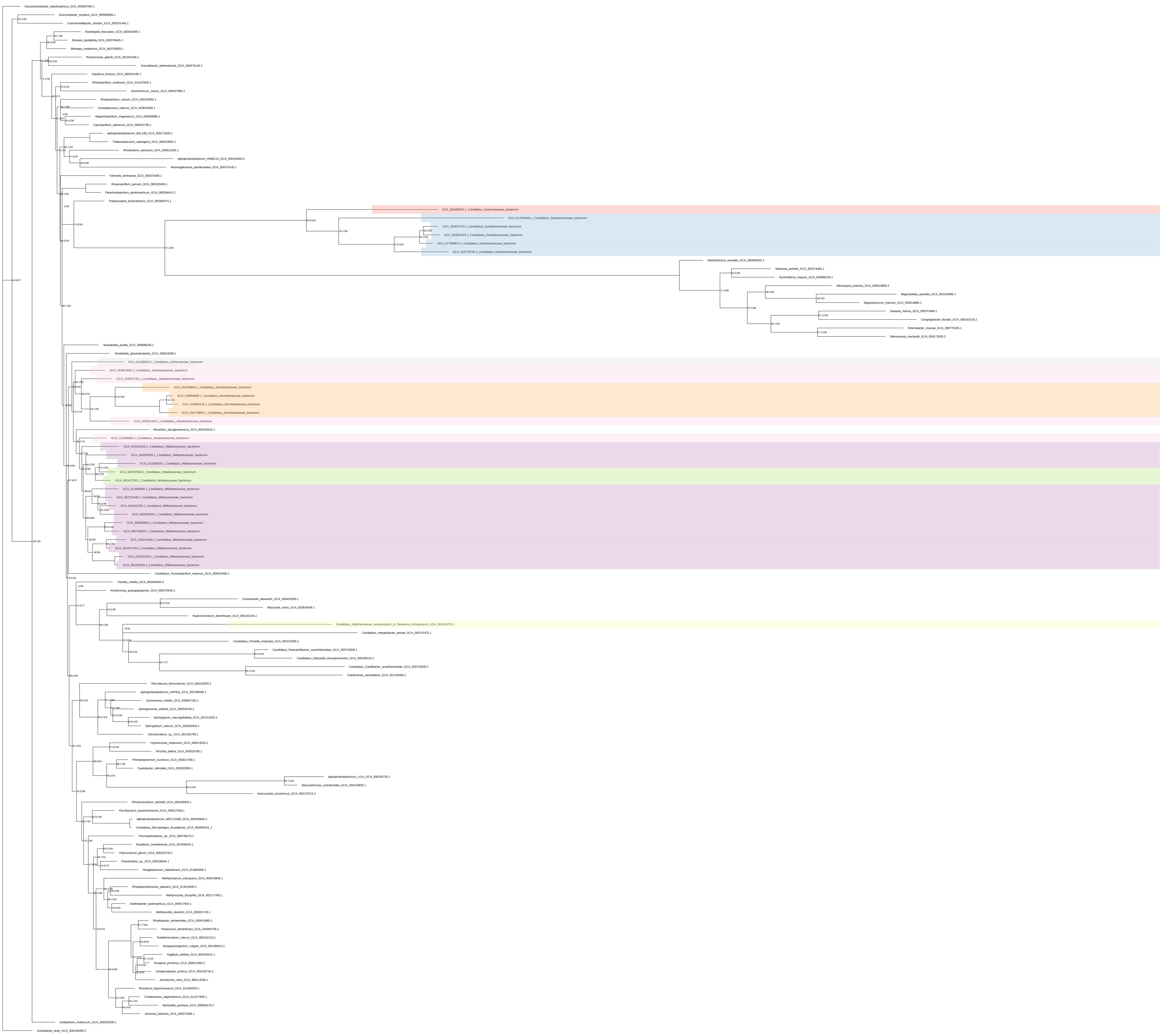

### Ser_organism_selection

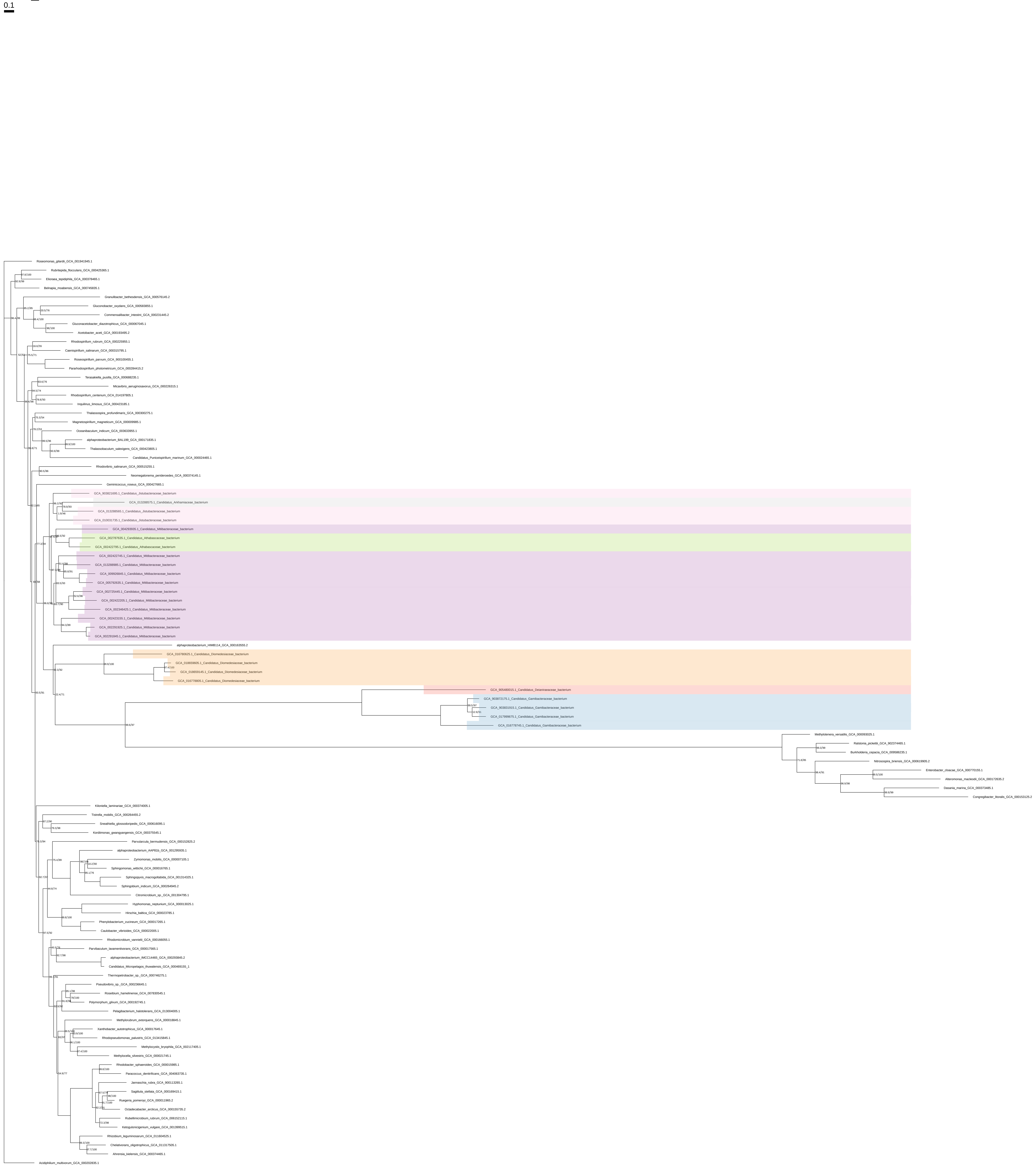

### Ser_organism_selection_z_0.10

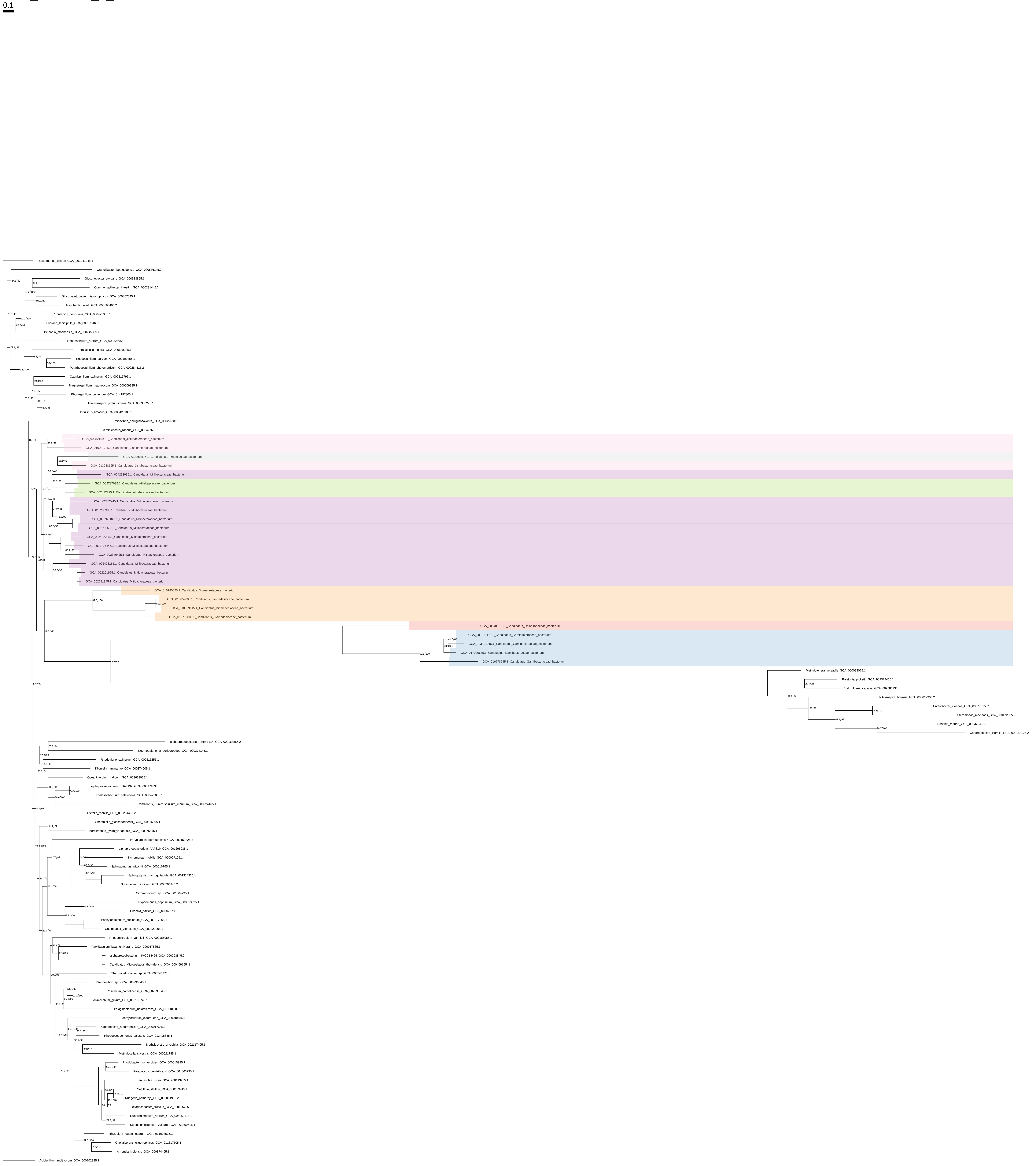

### Ser_organism_selection_z_0.20

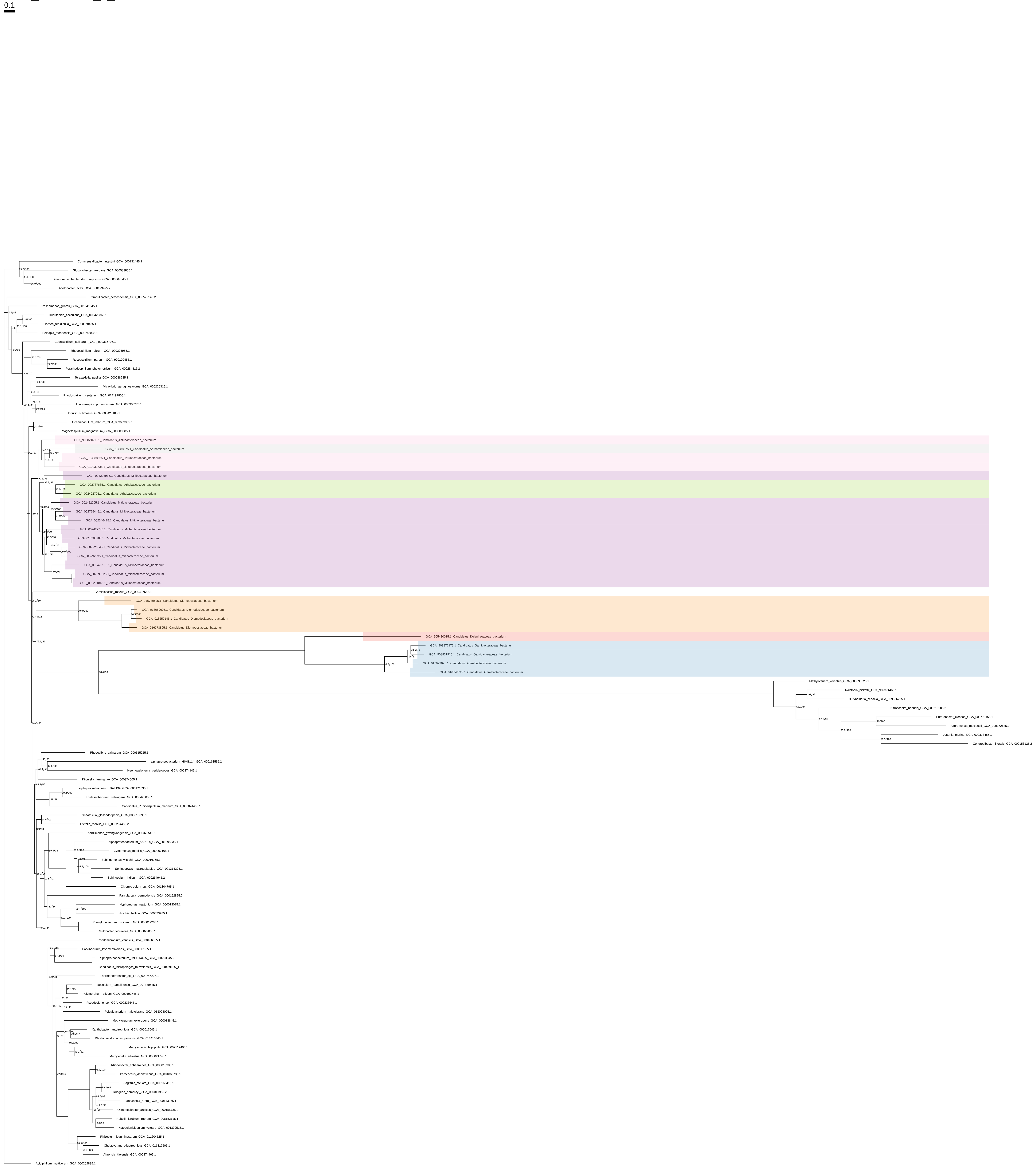

### Ser_organism_selection_z_0.30

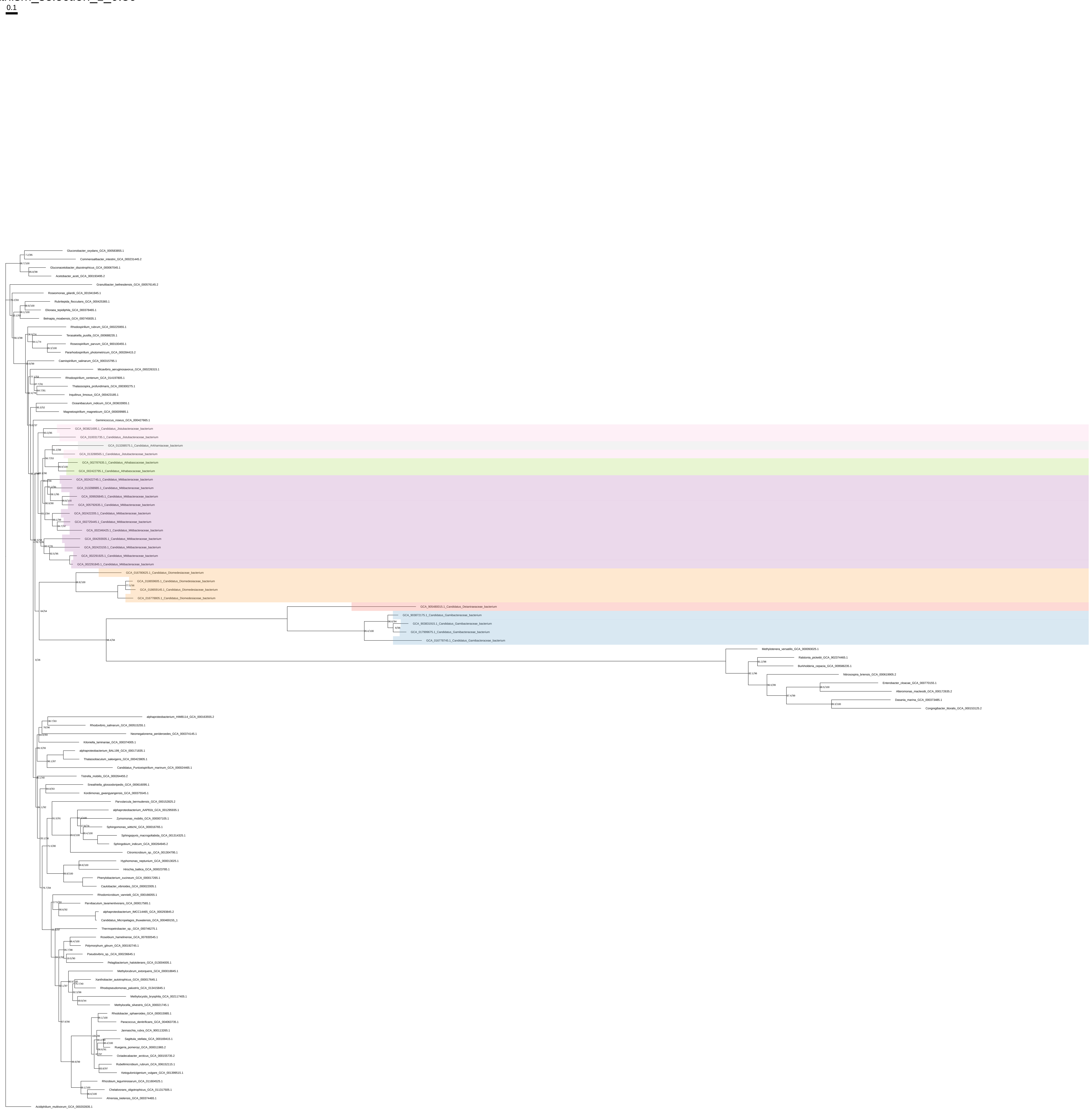

### Ser_organism_selection_z_0.40

0.1

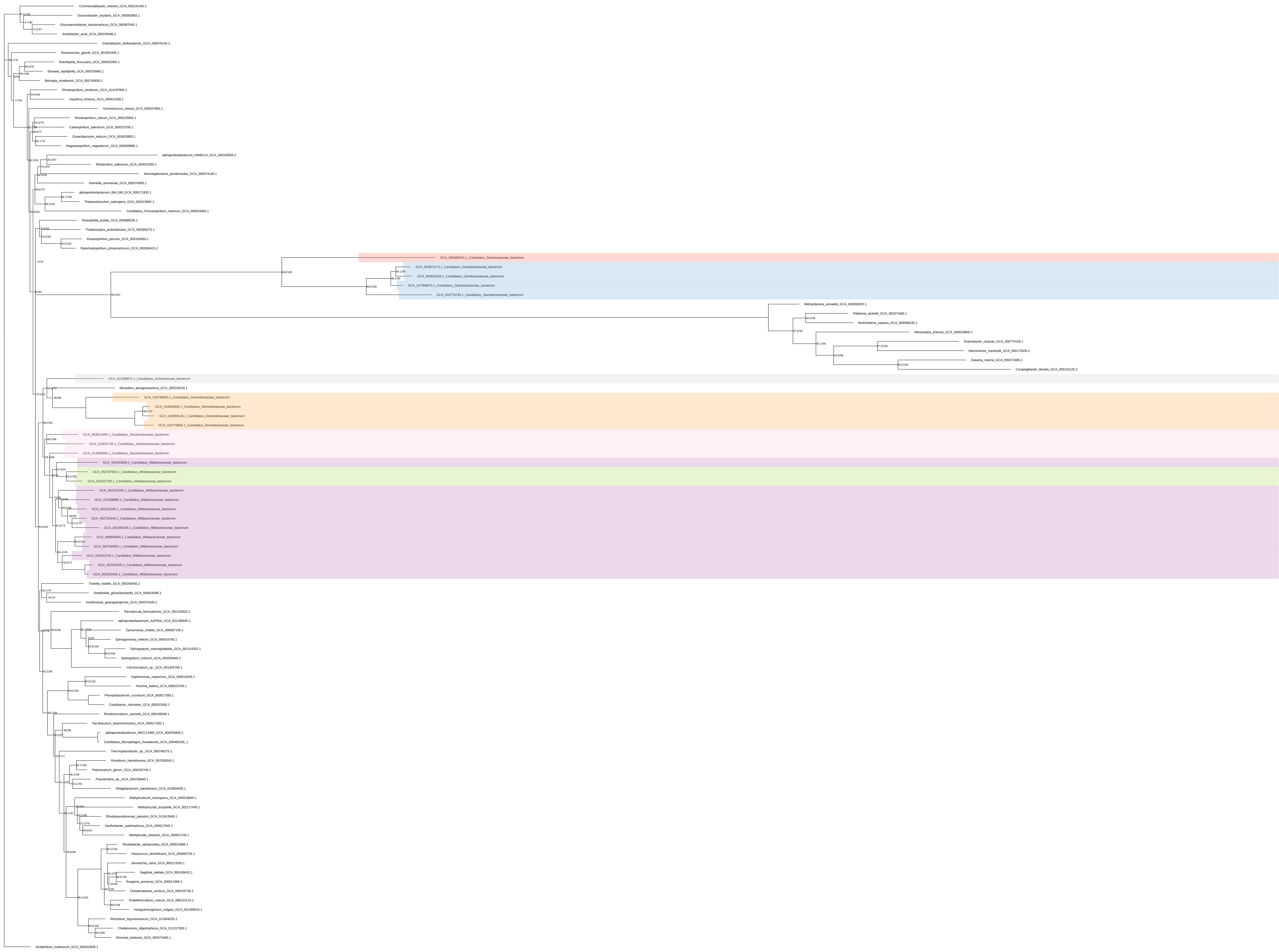

### Ser_organism_selection_z_0.50

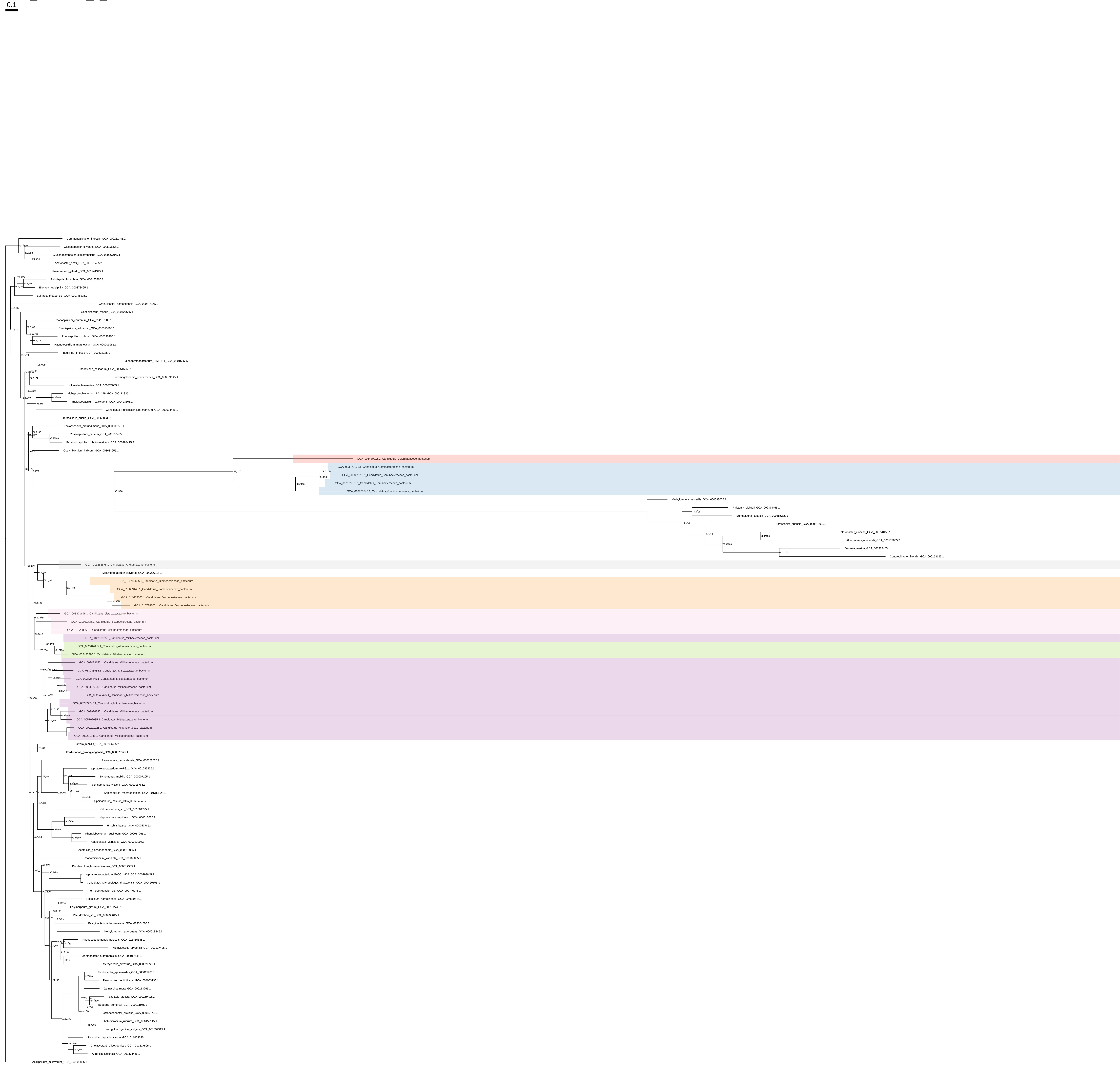

### Trp_Tyr_Phe_all_organisms

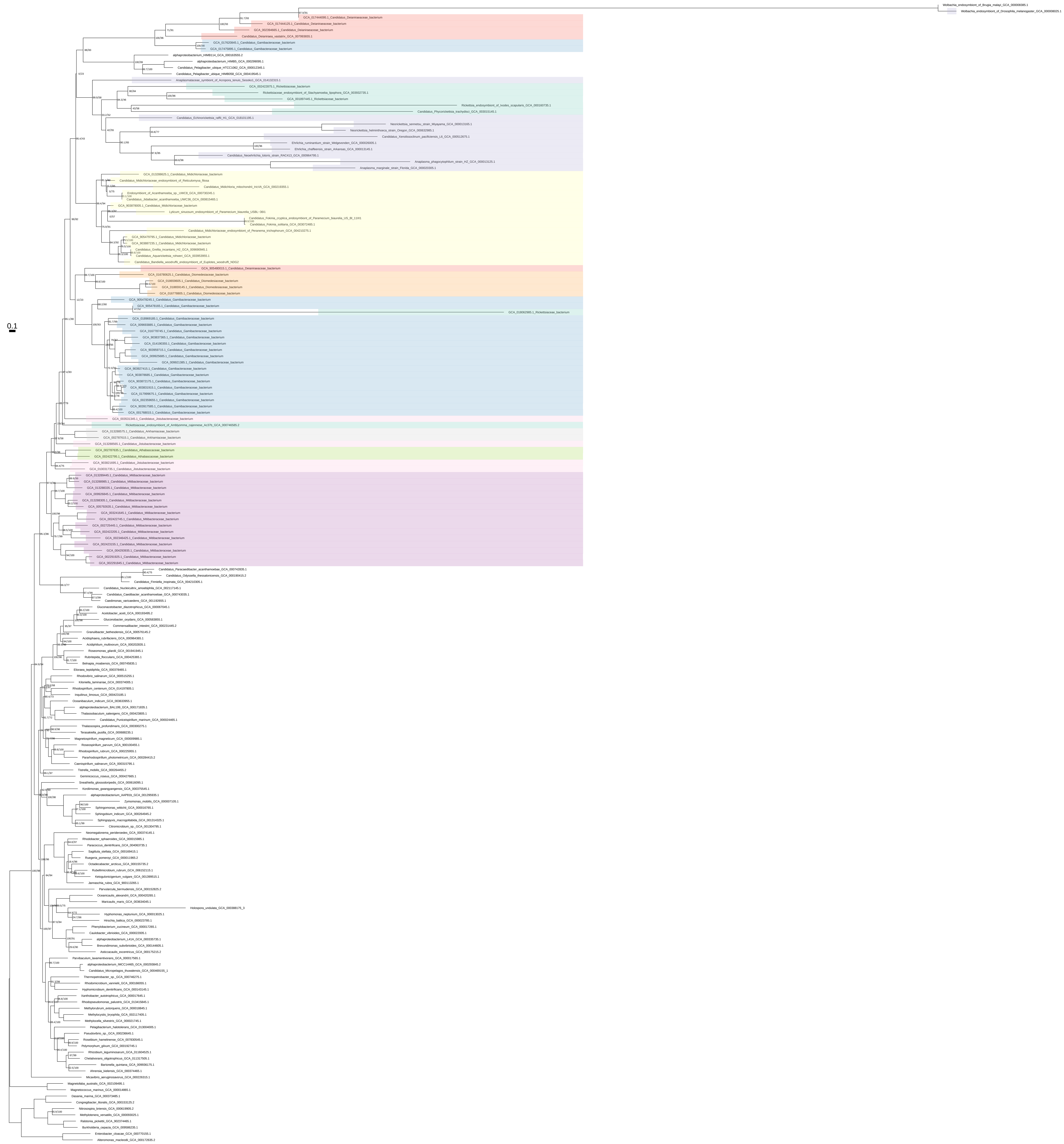

### Trp_Tyr_Phe_all_organisms_z_0.10

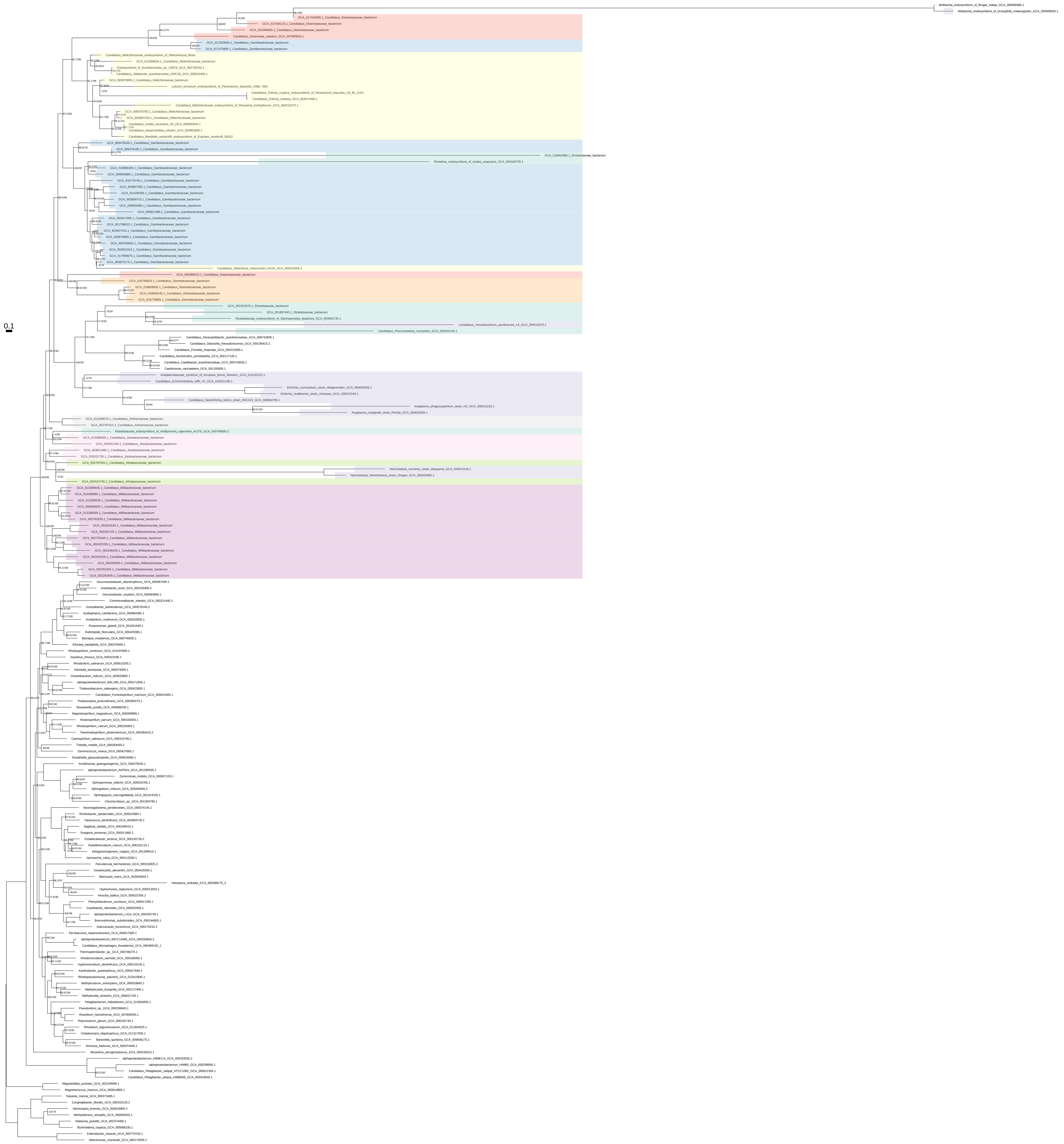

Trp_Tyr_Phe_all_organisms_z_0.20

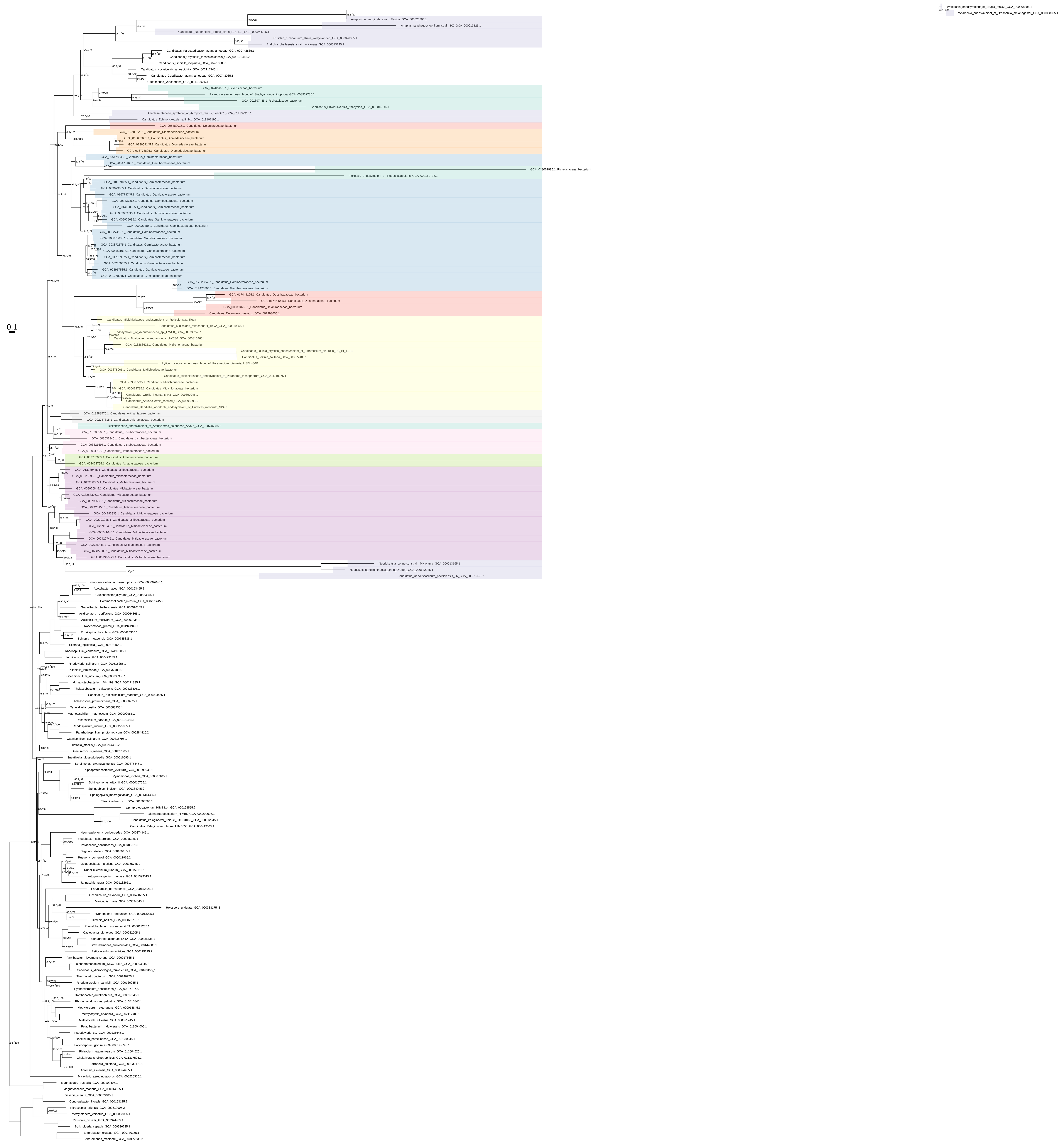

### Trp_Tyr_Phe_all_organisms_z_0.30

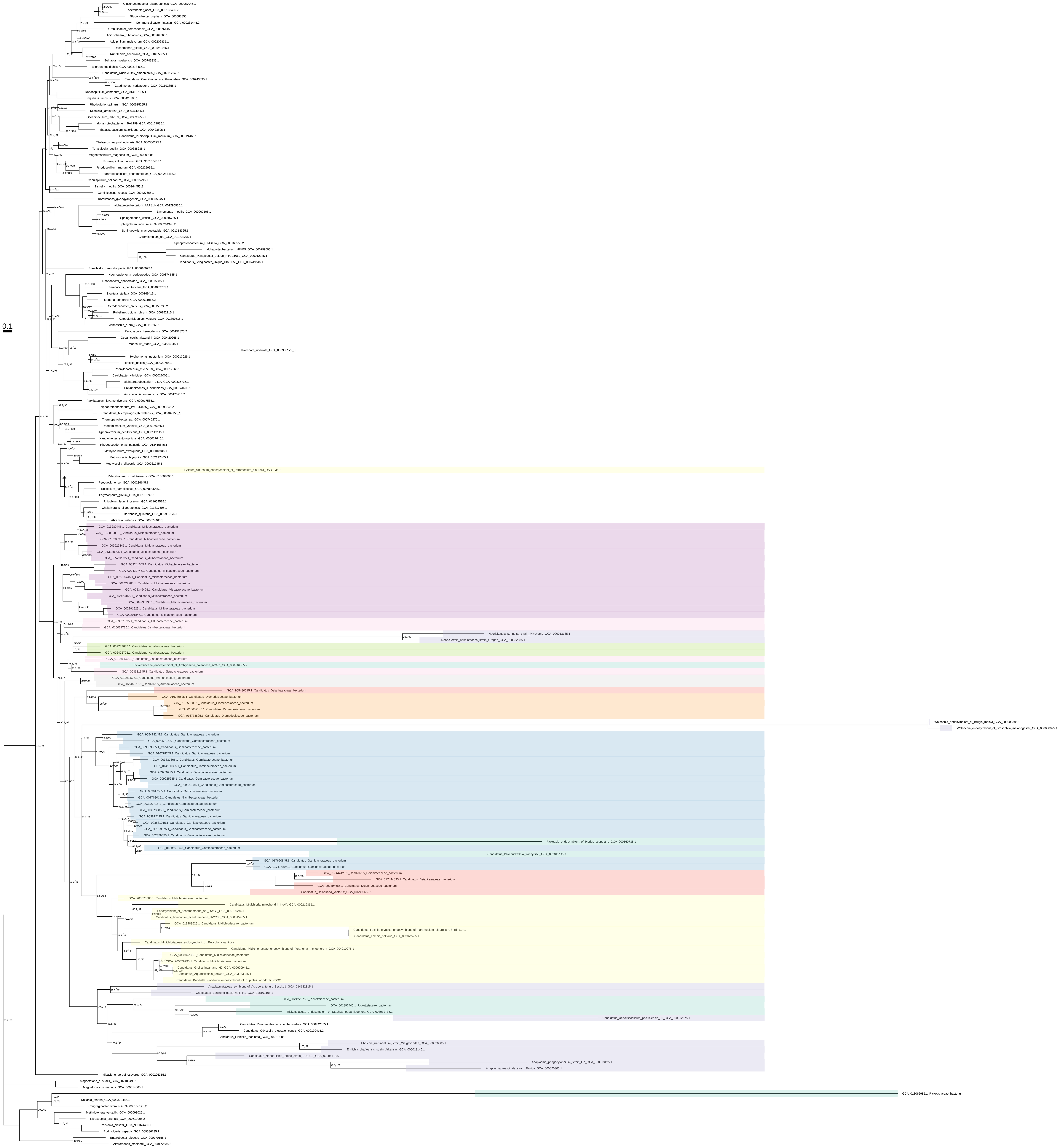

### Trp_Tyr_Phe_all_organisms_z_0.40

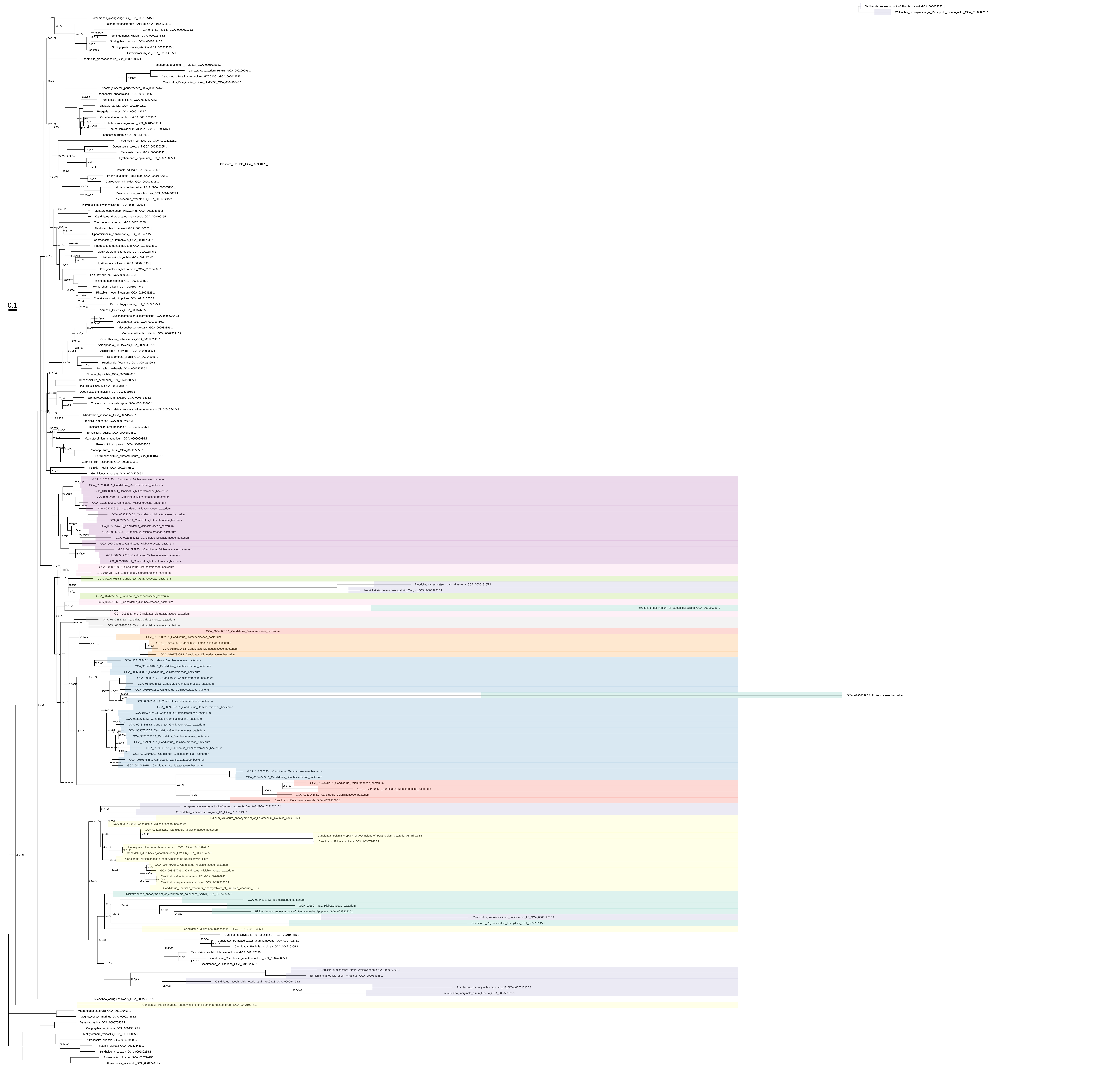

### Trp_Tyr_Phe_all_organisms_z_0.50

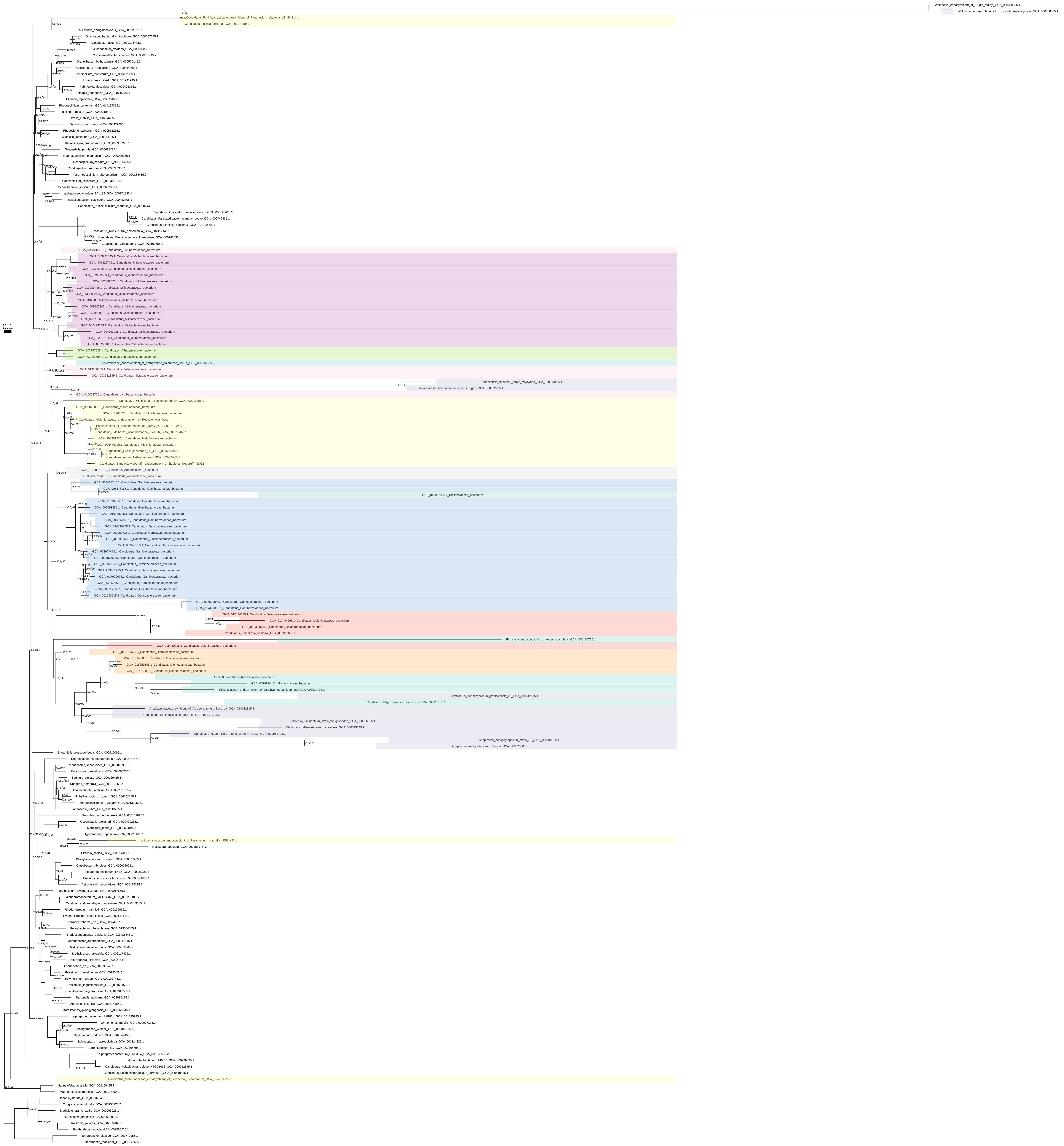

### Trp_Tyr_Phe_organism_selection

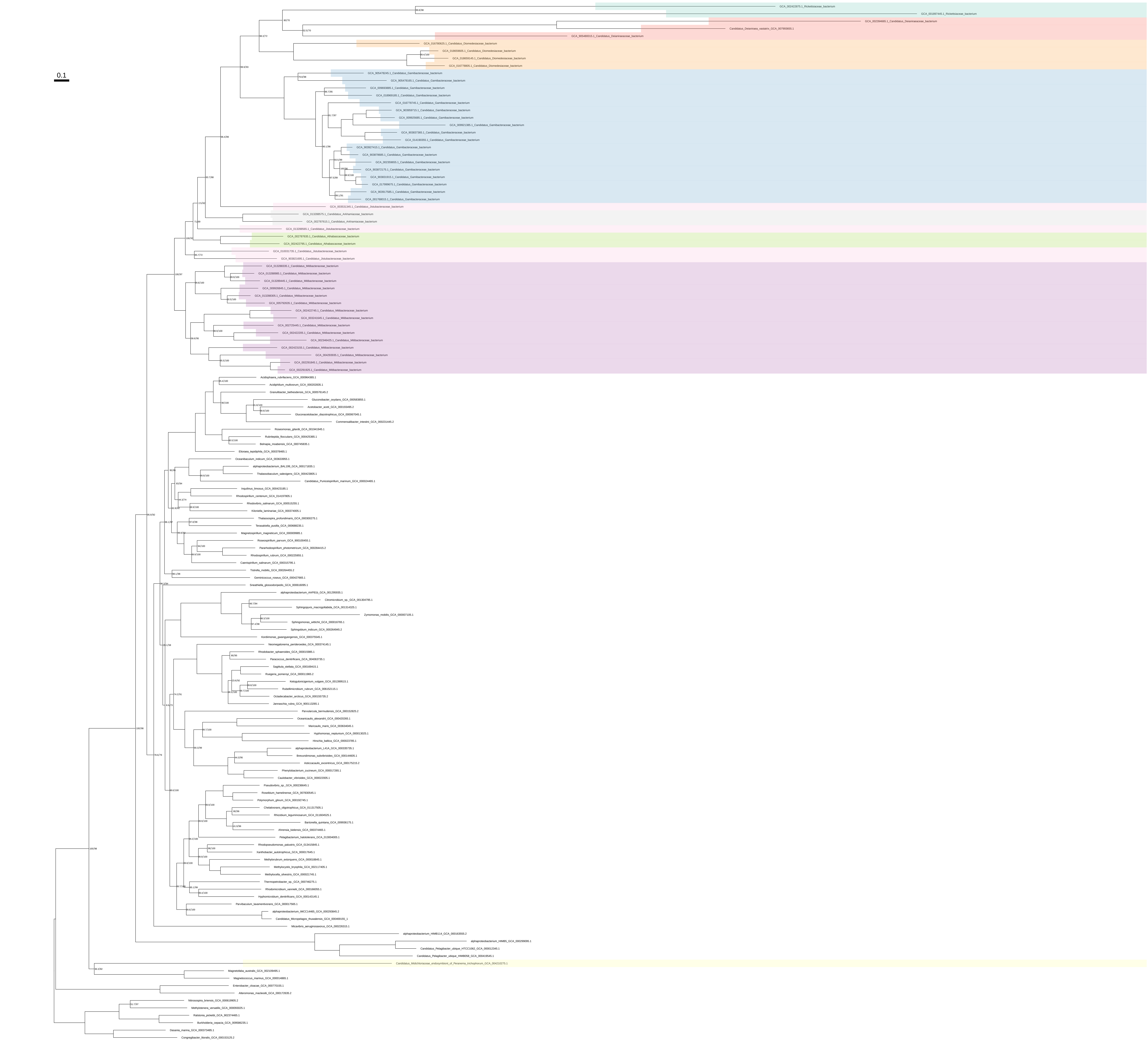

Trp_Tyr_Phe_organism_selection_z_0.10

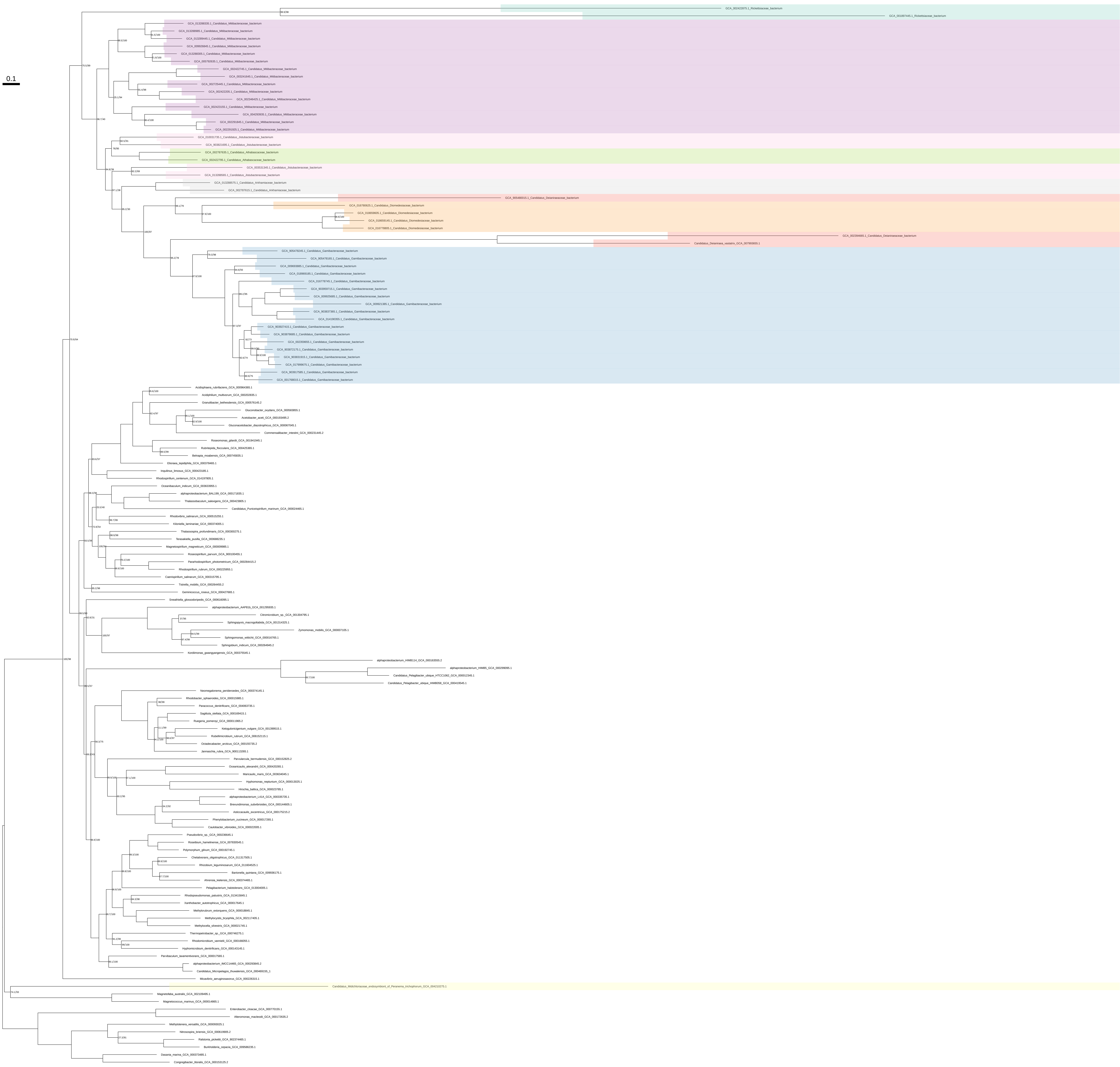

### Trp_Tyr_Phe_organism_selection_z_0.20

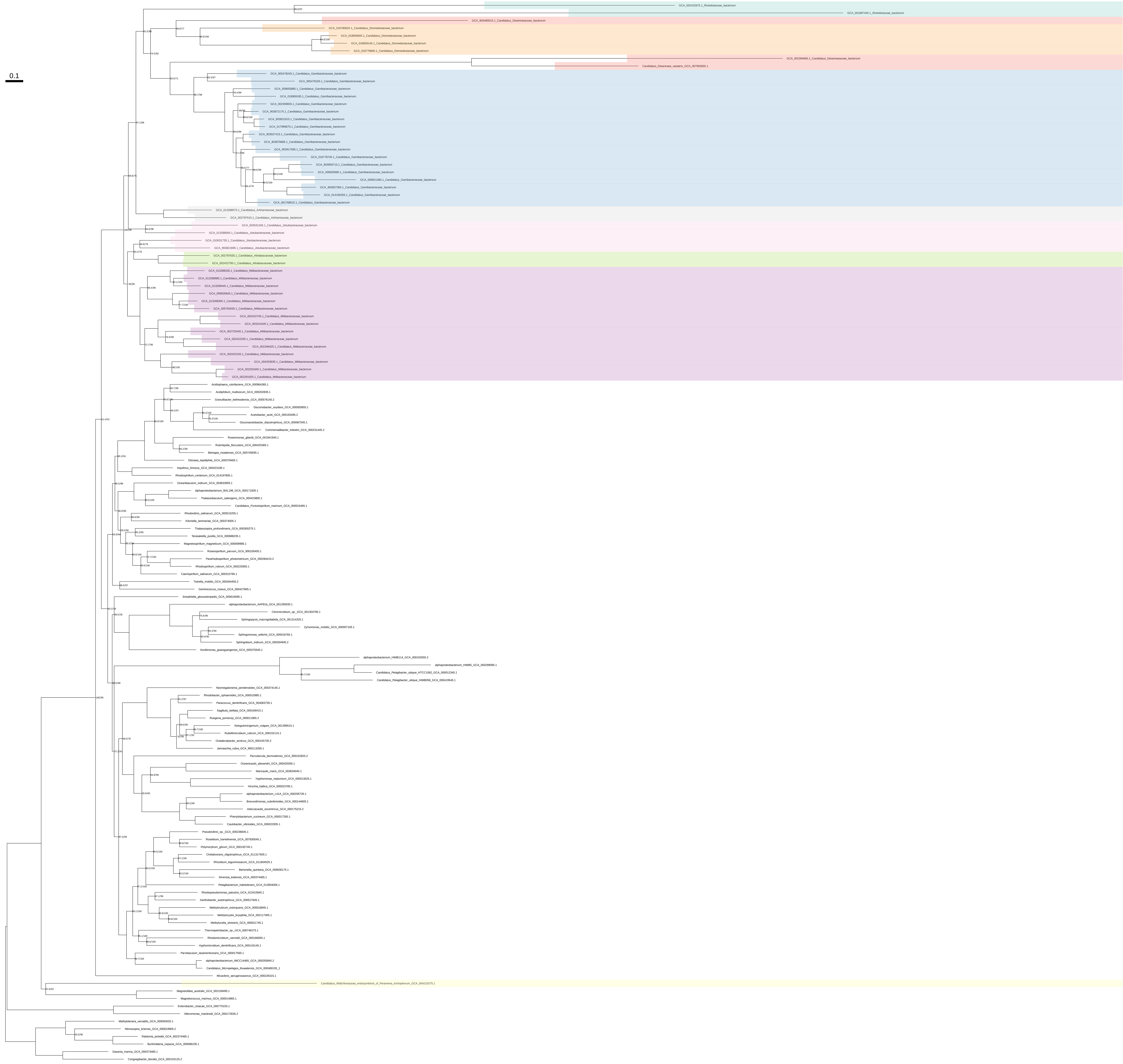

### Trp_Tyr_Phe_organism_selection_z_0.30

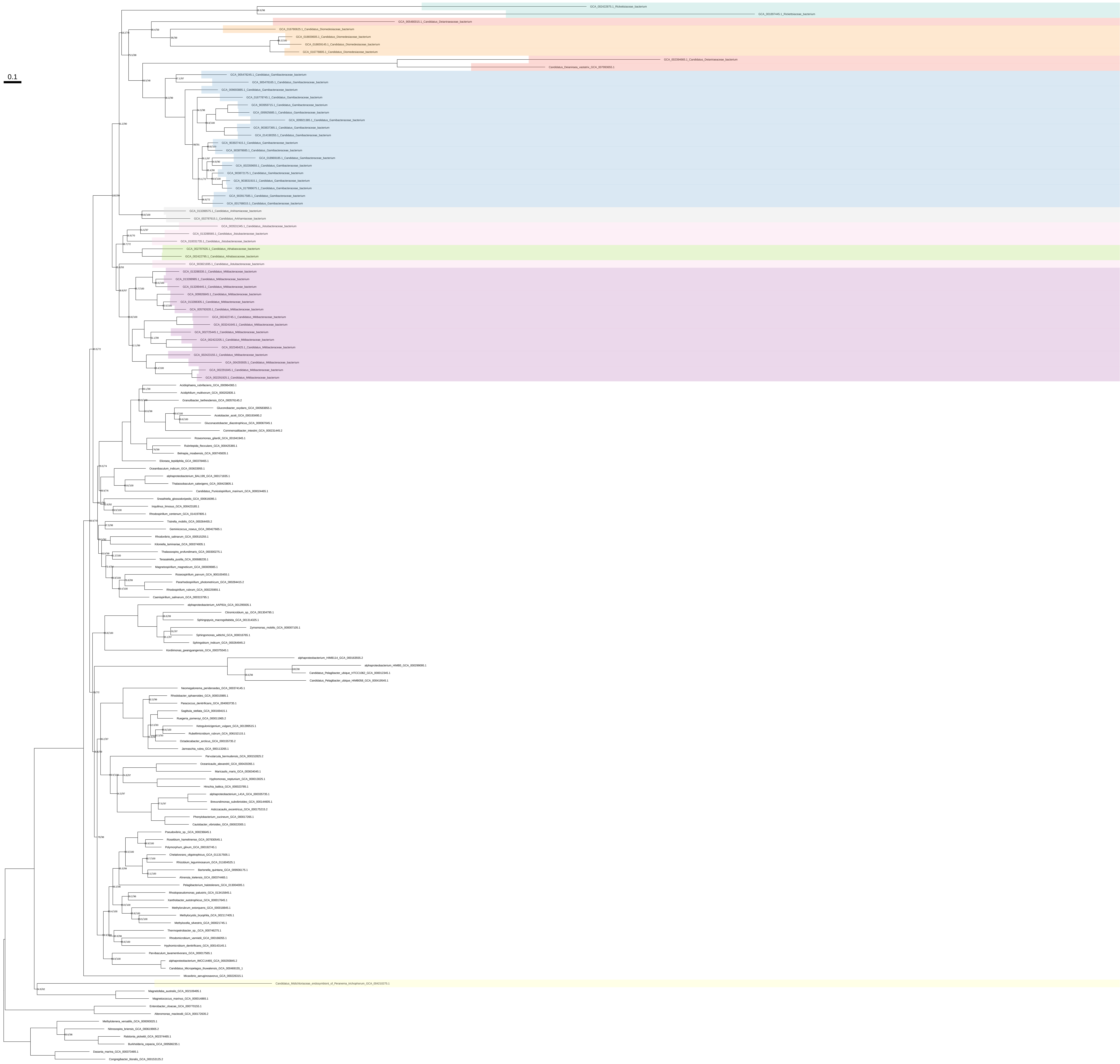

### Trp_Tyr_Phe_organism_selection_z_0.40

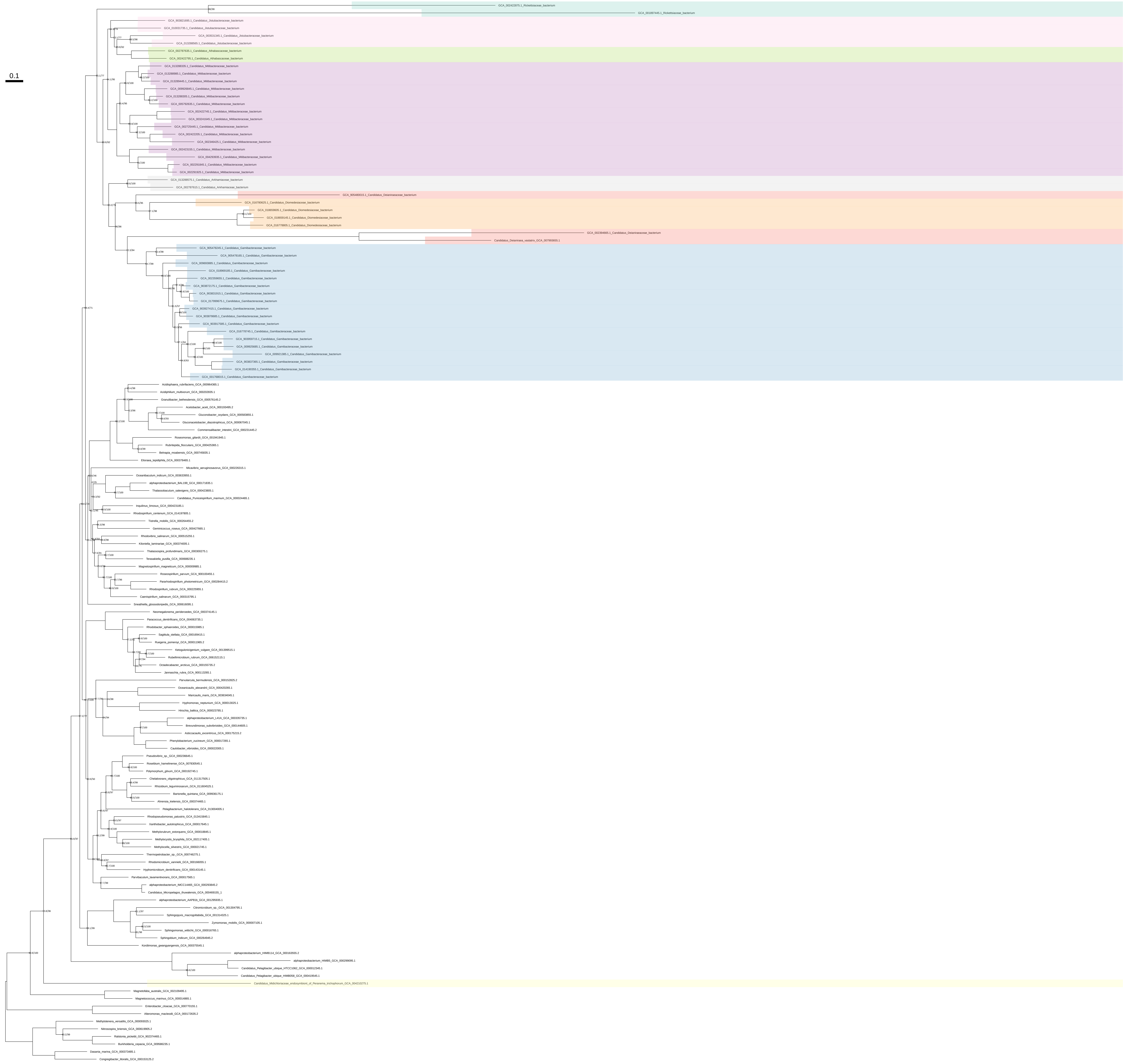

Trp_Tyr_Phe_organism_selection_z_0.50

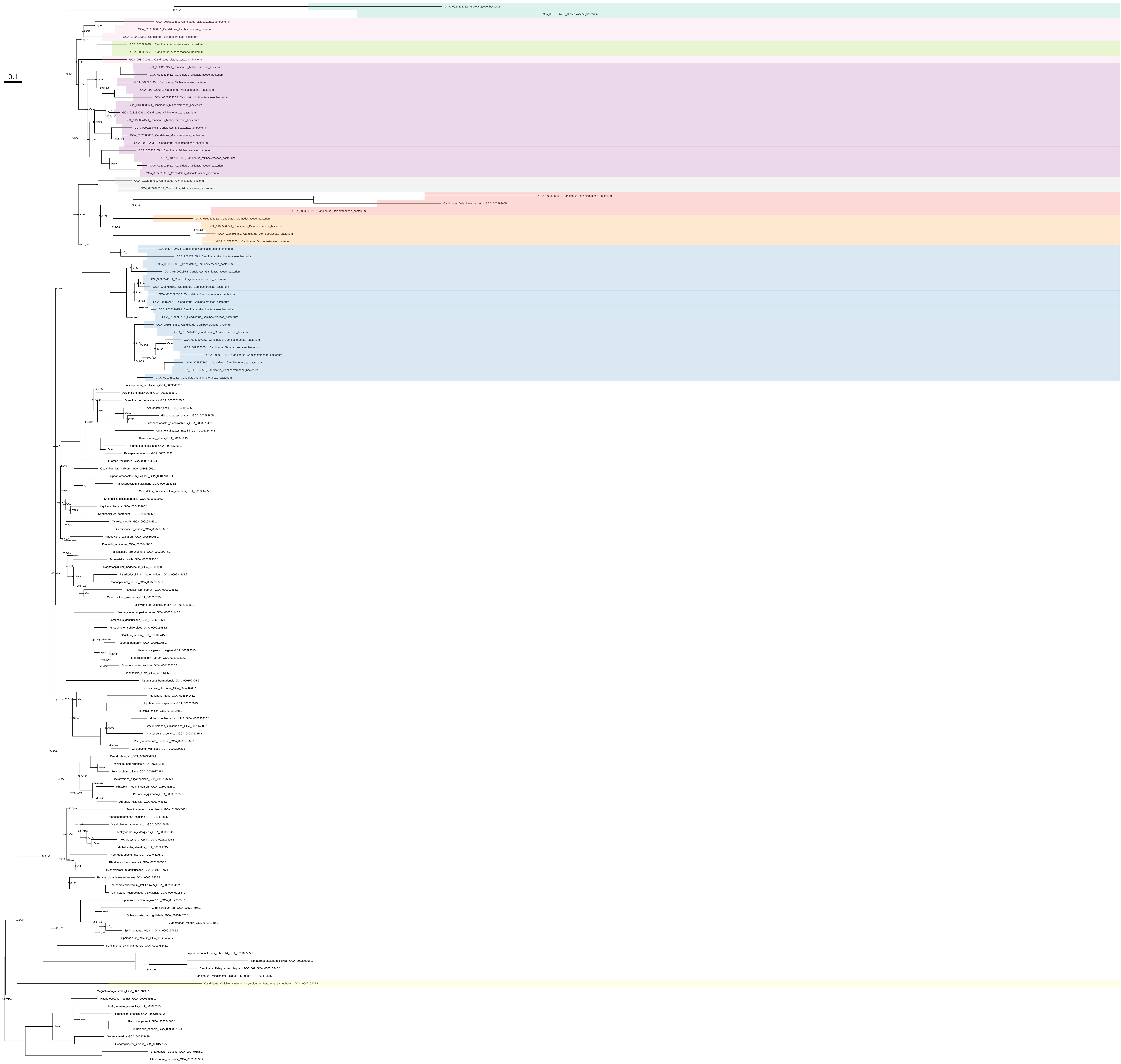
