## Supplementary material for "Host association and intracellularity evolved multiple times independently in the *Rickettsiales*": figures_and_supplementary_materials: Supplementary_figure_11_Phylogenomics_step4_general_trees.pdf

### original concatenated alignment

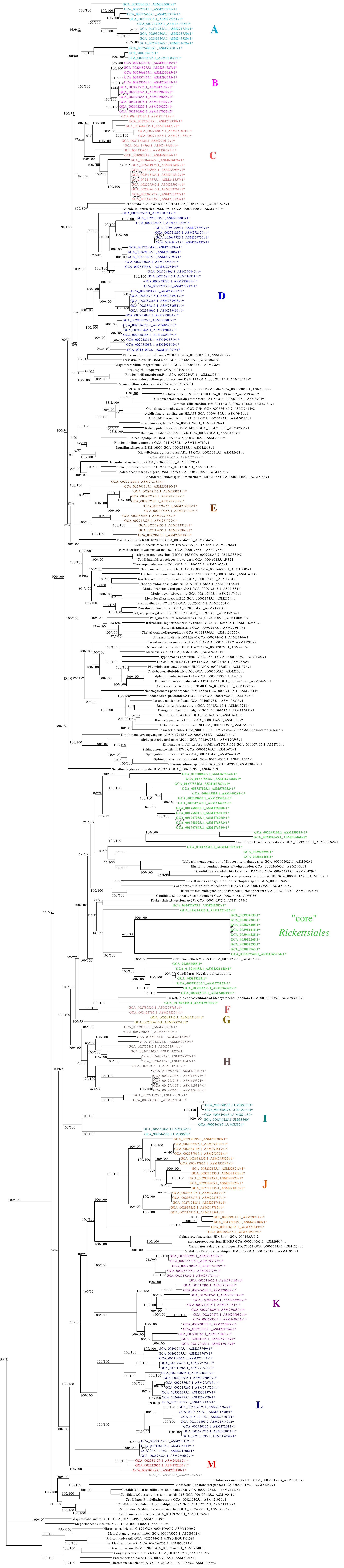

### alignment with 10% most heterogeneous sites removed

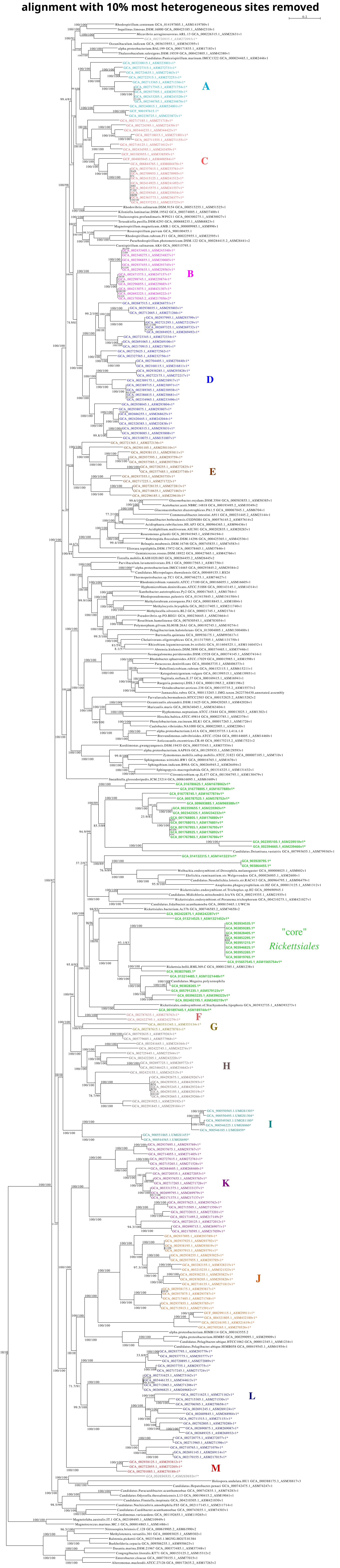

**alignment with 20% most heterogeneous sites removed**

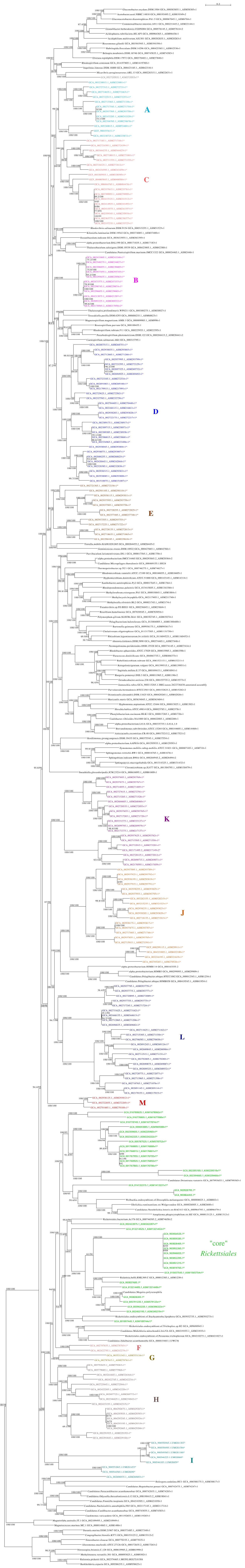

#### alignment with 30% most heterogeneous sites removed

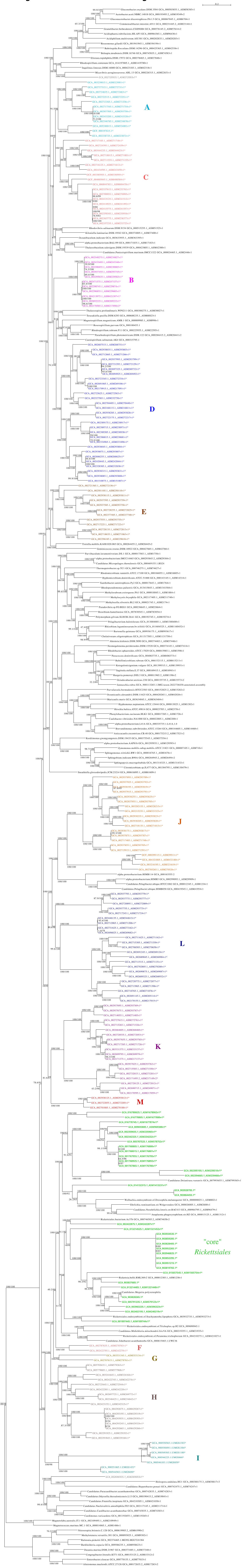

**alignment with 40% most heterogeneous sites removed**

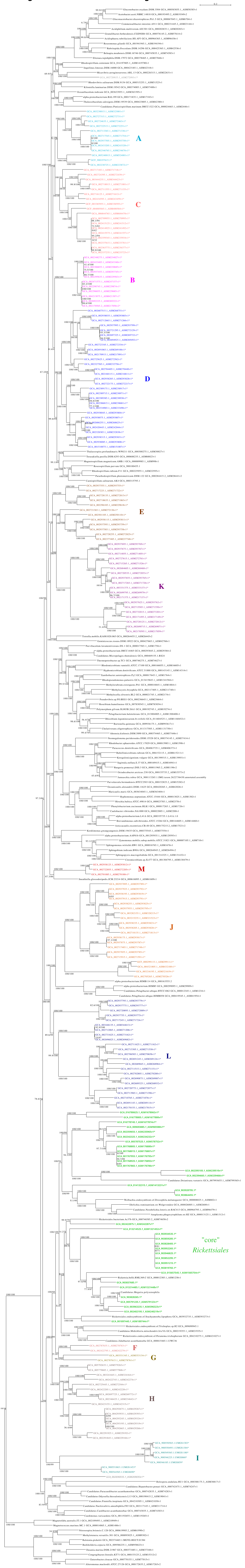

### alignment with 50% most heterogeneous sites removed

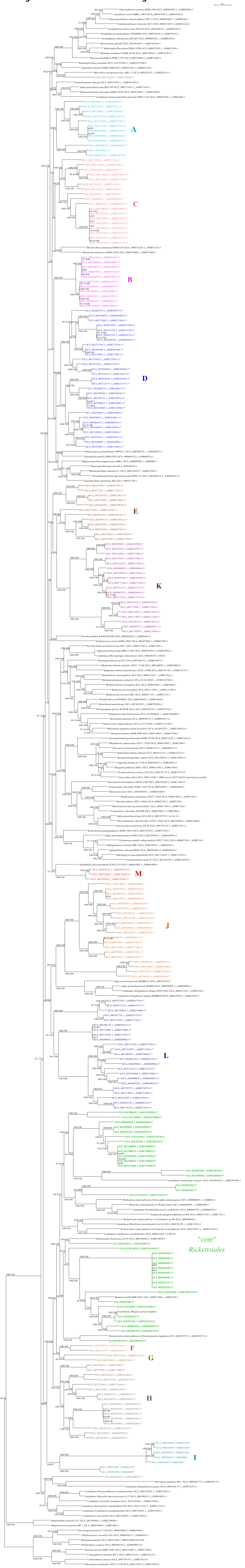
