## Supplementary material for "Host association and intracellularity evolved multiple times independently in the *Rickettsiales*": figures_and_supplementary_materials: Supplementary_figure_12_AAI_values.pdf

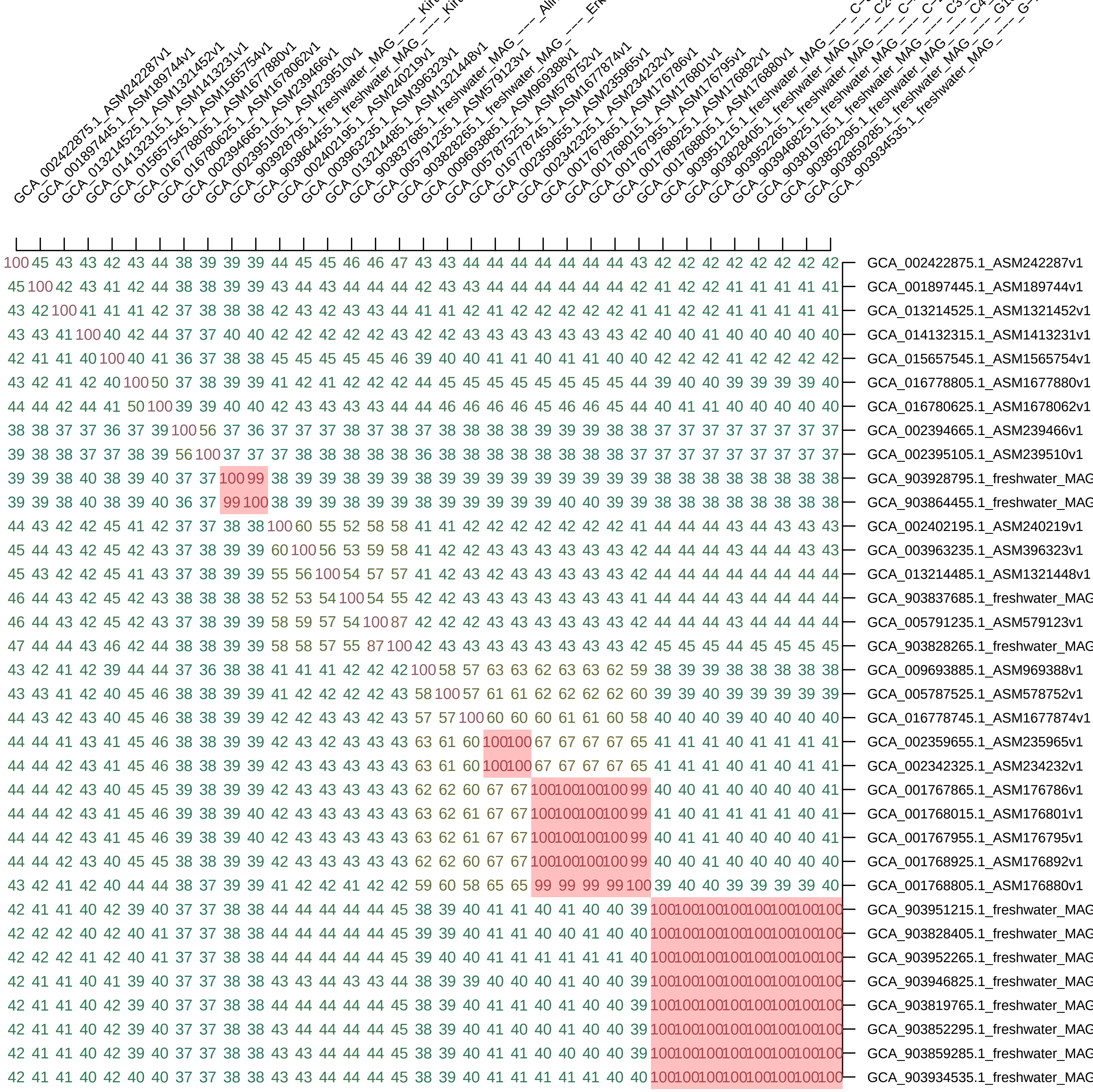

100 45 43 43 42 43 44 38 39 39 39 44 45 45 46 46 47 43 43 44 44 44 44 44 44 44 43 42 42 42 42 42 42 42 42 GCA_002422875.1_ASM242287v1
45 100 42 43 41 42 44 38 38 39 39 43 44 43 44 44 44 42 43 43 44 44 44 44 44 44 42 41 42 42 41 41 41 41 41 GCA_001897445.1_ASM189744v1
43 42 100 41 41 41 42 37 38 38 38 42 43 42 43 43 44 41 41 42 41 42 42 42 42 42 41 41 42 42 41 41 41 41 41 GCA_013214525.1_ASM1321452v1
43 43 41 100 40 42 44 37 37 40 40 42 42 42 42 42 43 42 42 43 43 43 43 43 43 43 42 40 40 41 40 40 40 40 40 GCA_014132315.1_ASM1413231v1
42 41 41 40 100 40 41 36 37 38 38 45 45 45 45 45 46 39 40 40 41 41 40 41 41 40 40 42 42 42 41 42 42 42 42 GCA_015657545.1_ASM1565754v1
43 42 41 42 40 100 50 37 38 39 39 41 42 41 42 42 42 44 45 45 45 45 45 45 45 45 44 39 40 40 39 39 39 39 40 GCA_016778805.1_ASM1677880v1
44 44 42 44 41 50 100 39 39 40 40 42 43 43 43 44 44 46 46 46 46 46 45 46 46 45 44 40 41 41 40 40 40 40 40 GCA_016780625.1_ASM1678062v1
38 38 37 37 36 37 39 100 56 37 36 37 37 37 38 37 38 37 38 38 38 38 39 39 39 38 38 37 37 37 37 37 37 37 37 GCA_002394665.1_ASM239466v1
39 38 38 37 37 38 39 56 100 37 37 37 38 38 38 38 38 36 38 38 38 38 38 38 38 38 37 37 37 37 37 37 37 37 37 GCA_002395105.1_ASM239510v1
39 39 38 40 38 39 40 37 37 100 99 38 39 39 38 39 39 38 39 39 39 39 39 39 39 39 39 38 38 38 38 38 38 38 38 GCA_903928795.1_freshwater_MAG
39 39 38 40 38 39 40 36 37 99 100 38 39 39 38 39 39 38 39 39 39 39 39 40 40 39 39 38 38 38 38 38 38 38 38 GCA_903864455.1_freshwater_MAG
44 43 42 42 45 41 42 37 37 38 38 100 60 55 52 58 58 41 41 42 42 42 42 42 42 42 41 44 44 44 43 44 43 43 43 GCA_002402195.1_ASM240219v1
45 44 43 42 45 42 43 37 38 39 39 60 100 56 53 59 58 41 42 42 43 43 43 43 43 43 42 44 44 44 43 44 44 43 43 GCA_003963235.1_ASM396323v1
45 43 42 42 45 41 43 37 38 39 39 55 56 100 54 57 57 41 42 43 42 43 43 43 43 43 42 44 44 44 44 44 44 44 44 GCA_013214485.1_ASM1321448v1
46 44 43 42 45 42 43 38 38 38 38 52 53 54 100 54 55 42 42 43 43 43 43 43 43 43 41 44 44 44 43 44 44 44 44 GCA_903837685.1_freshwater_MAG
46 44 43 42 45 42 43 37 38 39 39 58 59 57 54 100 87 42 42 42 43 43 43 43 43 43 42 44 44 44 43 44 44 44 44 GCA_005791235.1_ASM579123v1
47 44 44 43 46 42 44 38 38 39 39 58 58 57 55 87 100 42 43 43 43 43 43 43 43 43 42 45 45 45 44 45 45 45 45 GCA_903828265.1_freshwater_MAG
43 42 41 42 39 44 44 37 36 38 38 41 41 41 42 42 42 100 58 57 63 63 62 63 63 62 59 38 39 39 38 38 38 38 38 GCA_009693885.1_ASM969388v1
43 43 41 42 40 45 46 38 38 39 39 41 42 42 42 42 43 58 100 57 61 61 62 62 62 62 60 39 39 40 39 39 39 39 39 GCA_005787525.1_ASM578752v1
44 43 42 43 40 45 46 38 38 39 39 42 42 43 43 42 43 57 57 100 60 60 60 61 61 60 58 40 40 40 39 40 40 40 40 GCA_016778745.1_ASM1677874v1
44 44 41 43 41 45 46 38 38 39 39 42 43 42 43 43 43 63 61 60 100 100 67 67 67 67 65 41 41 41 40 41 41 41 41 GCA_002359655.1_ASM235965v1
44 44 42 43 41 45 46 38 38 39 39 42 43 43 43 43 43 63 61 60 100 100 67 67 67 67 65 41 41 41 40 41 40 41 41 GCA_002342325.1_ASM234232v1
44 44 42 43 40 45 45 39 38 39 39 42 43 43 43 43 43 62 62 60 67 67 100 100 100 100 99 40 40 41 40 40 40 40 41 GCA_001767865.1_ASM176786v1
44 44 42 43 41 45 46 39 38 39 40 42 43 43 43 43 43 63 62 61 67 67 100 100 100 100 99 41 40 41 41 41 41 40 41 GCA_001768015.1_ASM176801v1
44 44 42 43 41 45 46 39 38 39 40 42 43 43 43 43 43 63 62 61 67 67 100 100 100 100 99 40 41 41 40 40 40 40 41 GCA_001767955.1_ASM176795v1
44 44 42 43 40 45 45 38 38 39 39 42 43 43 43 43 43 62 62 60 67 67 100 100 100 100 99 40 40 41 40 40 40 40 40 GCA_001768925.1_ASM176892v1
43 42 41 42 40 44 44 38 37 39 39 41 42 42 41 42 42 59 60 58 65 65 99 99 99 99 100 39 40 40 39 39 39 39 40 GCA_001768805.1_ASM176880v1
42 41 41 40 42 39 40 37 37 38 38 44 44 44 44 44 45 38 39 40 41 41 40 41 40 40 39 100 100 100 100 100 100 100 100 GCA_903951215.1_freshwater_MAG
42 42 42 40 42 40 41 37 37 38 38 44 44 44 44 44 45 39 39 40 41 41 40 40 41 40 40 100 100 100 100 100 100 100 100 GCA_903828405.1_freshwater_MAG
42 42 42 41 42 40 41 37 37 38 38 44 44 44 44 44 45 39 40 40 41 41 41 41 41 41 40 100 100 100 100 100 100 100 100 GCA_903952265.1_freshwater_MAG
42 41 41 40 41 39 40 37 37 38 38 43 43 44 43 43 44 38 39 39 40 40 40 41 40 40 39 100 100 100 100 100 100 100 100 GCA_903946825.1_freshwater_MAG
42 41 41 40 42 39 40 37 37 38 38 44 44 44 44 44 45 38 39 40 41 41 40 41 40 40 39 100 100 100 100 100 100 100 100 GCA_903819765.1_freshwater_MAG
42 41 41 40 42 39 40 37 37 38 38 43 44 44 44 44 45 38 39 40 41 40 40 41 40 40 39 100 100 100 100 100 100 100 100 GCA_903852295.1_freshwater_MAG
42 41 41 40 42 39 40 37 37 38 38 43 43 44 44 44 45 38 39 40 41 41 40 40 40 40 39 100 100 100 100 100 100 100 100 GCA_903859285.1_freshwater_MAG
42 41 41 40 42 40 40 37 37 38 38 43 43 44 44 44 45 38 39 40 41 41 41 41 41 40 40 100 100 100 100 100 100 100 100 GCA_903934535.1_freshwater_MAG

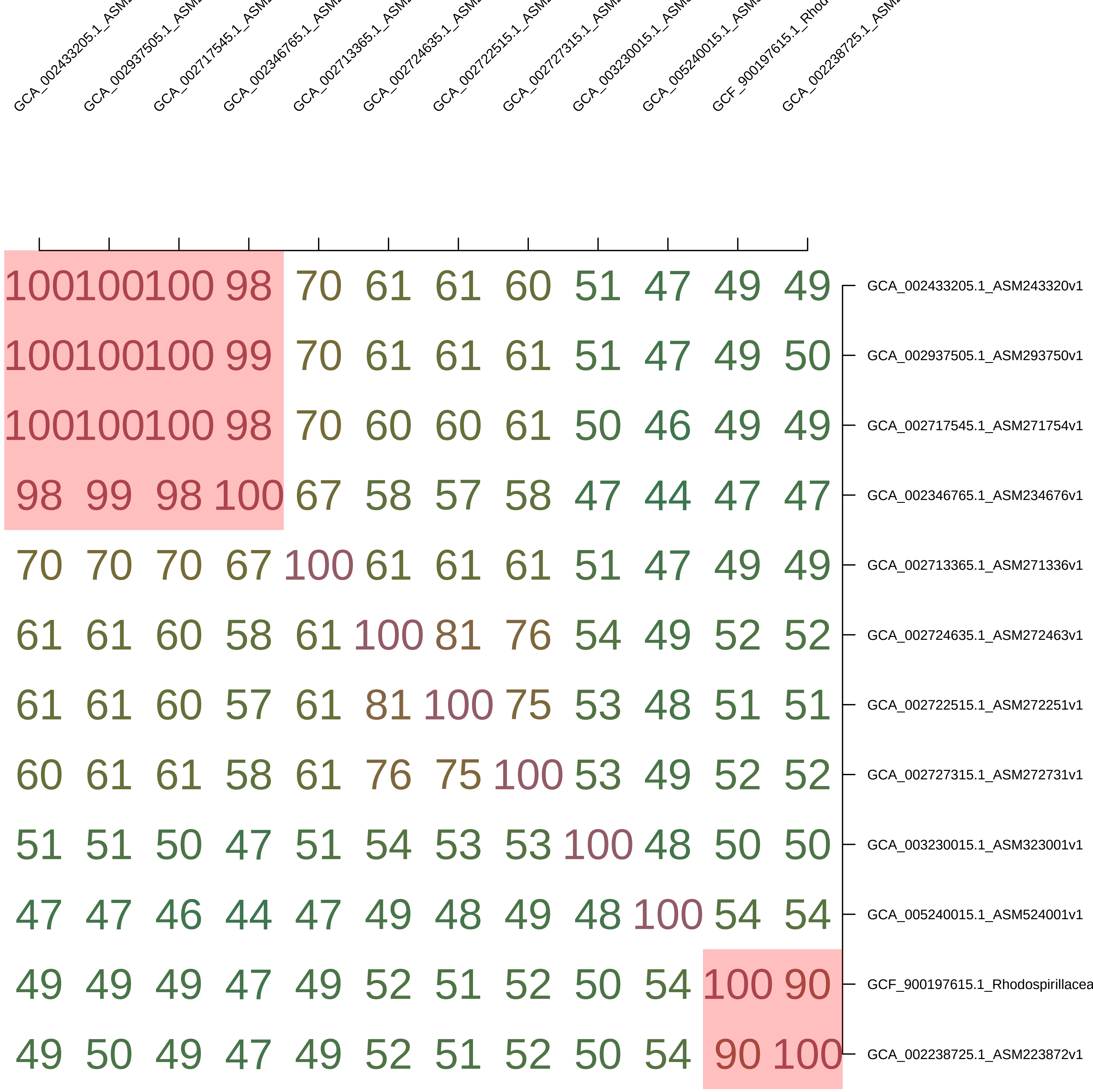
GCA_002433205.1_ASM
GCA_002937505.1_ASM
GCA_002717545.1_ASM
GCA_002346765.1_ASM
GCA_002713365.1_ASM
GCA_002724635.1_ASM
GCA_002722515.1_ASM
GCA_002727315.1_ASM
GCA_003230015.1_ASM
GCA_005240015.1_ASM
GCF_900197615.1_Rhod
GCA_002238725.1_ASM
100 100 100 98 70 61 61 60 51 47 49 49 GCA_002433205.1_ASM243320v1
100 100 100 99 70 61 61 61 51 47 49 50 GCA_002937505.1_ASM293750v1
100 100 100 98 70 60 60 61 50 46 49 49 GCA_002717545.1_ASM271754v1
98 99 98 100 67 58 57 58 47 44 47 47 GCA_002346765.1_ASM234676v1
70 70 70 67 100 61 61 61 51 47 49 49 GCA_002713365.1_ASM271336v1
61 61 60 58 61 100 81 76 54 49 52 52 GCA_002724635.1_ASM272463v1
61 61 60 57 61 81 100 75 53 48 51 51 GCA_002722515.1_ASM272251v1
60 61 61 58 61 76 75 100 53 49 52 52 GCA_002727315.1_ASM272731v1
51 51 50 47 51 54 53 53 100 48 50 50 GCA_003230015.1_ASM323001v1
47 47 46 44 47 49 48 49 48 100 54 54 GCA_005240015.1_ASM524001v1
49 49 49 47 49 52 51 52 50 54 100 90 GCF_900197615.1_Rhodospirillacea
49 50 49 47 49 52 51 52 50 54 90 100 GCA_002238725.1_ASM223872v1

| | GCA_002170565.2_AS | GCA_002692225.1_AS | GCA_004213075.1_AS | GCA_002471575.1_AS | GCA_002298745.1_AS | GCA_002296855.1_AS | GCA_002348275.1_AS | GCA_002306855.1_AS | GCA_002937455.1_AS | GCA_002295635.1_AS | GCA_002433405.1_AS |
|---|---|---|---|---|---|---|---|---|---|---|---|
| GCA_002170565.2_ASM217056v2 | 100 | 100 | 99 | 95 | 95 | 95 | 77 | 78 | 77 | 77 | 76 |
| GCA_002692225.1_ASM269222v1 | 100 | 100 | 99 | 96 | 95 | 95 | 77 | 77 | 77 | 77 | 76 |
| GCA_004213075.1_ASM421307v1 | 99 | 99 | 100 | 96 | 96 | 96 | 79 | 79 | 79 | 79 | 78 |
| GCA_002471575.1_ASM247157v1 | 95 | 96 | 96 | 100 | 100 | 100 | 79 | 79 | 79 | 79 | 78 |
| GCA_002298745.1_ASM229874v1 | 95 | 95 | 96 | 100 | 100 | 100 | 79 | 79 | 79 | 79 | 78 |
| GCA_002296855.1_ASM229685v1 | 95 | 95 | 96 | 100 | 100 | 100 | 79 | 79 | 79 | 79 | 78 |
| GCA_002348275.1_ASM234827v1 | 77 | 77 | 79 | 79 | 79 | 79 | 100 | 100 | 100 | 100 | 100 |
| GCA_002306855.1_ASM230685v1 | 78 | 77 | 79 | 79 | 79 | 79 | 100 | 100 | 100 | 100 | 99 |
| GCA_002937455.1_ASM293745v1 | 77 | 77 | 79 | 79 | 79 | 79 | 100 | 100 | 100 | 100 | 100 |
| GCA_002295635.1_ASM229563v1 | 77 | 77 | 79 | 79 | 79 | 79 | 100 | 100 | 100 | 100 | 99 |
| GCA_002433405.1_ASM243340v1 | 76 | 76 | 78 | 78 | 78 | 78 | 100 | 99 | 100 | 99 | 100 |

| | GCA_006844765.1_ASM684476v1 | GCF_004005845.1_ASM400584v1 | GCF_003385955.1_ASM338595v1 | GCA_002716125.1_ASM271612v1 | GCA_002434595.1_ASM243459v1 | GCA_002717185.1_ASM271718v1 | GCA_003444235.1_ASM344423v1 | GCA_002724395.1_ASM272439v1 | GCA_002718015.1_ASM271801v1 | GCA_002711555.1_ASM271155v1 | GCA_002359345.1_ASM235934v1 | GCA_002414925.1_ASM241492v1 | GCA_002337235.1_ASM233723v1 | GCA_002363775.1_ASM236377v1 | GCA_002337615.1_ASM233761v1 | GCA_002415125.1_ASM241512v1 | GCA_002709955.1_ASM270995v1 | GCA_002415575.1_ASM241557v1 |
|---|---|---|---|---|---|---|---|---|---|---|---|---|---|---|---|---|---|---|
| GCA_006844765.1_ASM684476v1 | 100 | 56 | 54 | 51 | 48 | 50 | 49 | 50 | 47 | 49 | 58 | 58 | 58 | 58 | 57 | 58 | 57 | 57 |
| GCF_004005845.1_ASM400584v1 | 56 | 100 | 55 | 53 | 51 | 52 | 50 | 51 | 50 | 51 | 56 | 56 | 56 | 56 | 56 | 56 | 56 | 56 |
| GCF_003385955.1_ASM338595v1 | 54 | 55 | 100 | 52 | 50 | 51 | 49 | 51 | 49 | 50 | 55 | 55 | 55 | 55 | 55 | 55 | 55 | 54 |
| GCA_002716125.1_ASM271612v1 | 51 | 53 | 52 | 100 | 75 | 60 | 55 | 55 | 52 | 53 | 52 | 51 | 51 | 51 | 51 | 51 | 51 | 51 |
| GCA_002434595.1_ASM243459v1 | 48 | 51 | 50 | 75 | 100 | 58 | 54 | 53 | 49 | 51 | 50 | 50 | 50 | 49 | 49 | 49 | 50 | 49 |
| GCA_002717185.1_ASM271718v1 | 50 | 52 | 51 | 60 | 58 | 100 | 53 | 54 | 51 | 53 | 51 | 51 | 51 | 51 | 51 | 51 | 51 | 50 |
| GCA_003444235.1_ASM344423v1 | 49 | 50 | 49 | 55 | 54 | 53 | 100 | 72 | 66 | 69 | 49 | 49 | 49 | 49 | 49 | 49 | 49 | 49 |
| GCA_002724395.1_ASM272439v1 | 50 | 51 | 51 | 55 | 53 | 54 | 72 | 100 | 67 | 68 | 51 | 51 | 50 | 50 | 51 | 50 | 51 | 50 |
| GCA_002718015.1_ASM271801v1 | 47 | 50 | 49 | 52 | 49 | 51 | 66 | 67 | 100 | 87 | 49 | 49 | 49 | 49 | 49 | 48 | 49 | 48 |
| GCA_002711555.1_ASM271155v1 | 49 | 51 | 50 | 53 | 51 | 53 | 69 | 68 | 87 | 100 | 50 | 50 | 50 | 50 | 50 | 49 | 50 | 49 |
| GCA_002359345.1_ASM235934v1 | 58 | 56 | 55 | 52 | 50 | 51 | 49 | 51 | 49 | 50 | 100 | 100 | 100 | 100 | 100 | 100 | 100 | 100 |
| GCA_002414925.1_ASM241492v1 | 58 | 56 | 55 | 51 | 50 | 51 | 49 | 51 | 49 | 50 | 100 | 100 | 100 | 100 | 100 | 100 | 100 | 100 |
| GCA_002337235.1_ASM233723v1 | 58 | 56 | 55 | 51 | 50 | 51 | 49 | 50 | 49 | 50 | 100 | 100 | 100 | 100 | 100 | 100 | 100 | 100 |
| GCA_002363775.1_ASM236377v1 | 58 | 56 | 55 | 51 | 49 | 51 | 49 | 50 | 49 | 50 | 100 | 100 | 100 | 100 | 100 | 100 | 100 | 100 |
| GCA_002337615.1_ASM233761v1 | 57 | 56 | 55 | 51 | 49 | 51 | 49 | 51 | 49 | 50 | 100 | 100 | 100 | 100 | 100 | 100 | 100 | 99 |
| GCA_002415125.1_ASM241512v1 | 58 | 56 | 55 | 51 | 49 | 51 | 49 | 50 | 48 | 49 | 100 | 100 | 100 | 100 | 100 | 100 | 100 | 100 |
| GCA_002709955.1_ASM270995v1 | 57 | 56 | 55 | 51 | 50 | 51 | 49 | 51 | 49 | 50 | 100 | 100 | 100 | 100 | 100 | 100 | 100 | 100 |
| GCA_002415575.1_ASM241557v1 | 57 | 56 | 54 | 51 | 49 | 50 | 49 | 50 | 48 | 49 | 100 | 100 | 100 | 100 | 99 | 100 | 100 | 100 |

| | GCA_002938315.1_ASM293831v1 | GCA_002938085.1_ASM293808v1 | GCA_001510075.1_ASM151007v1 | GCA_002938045.1_ASM293804v1 | GCA_002686255.1_ASM268625v1 | GCA_002328385.1_ASM232838v1 | GCA_002420445.1_ASM242044v1 | GCA_002938075.1_ASM293807v1 | GCA_002938285.1_ASM293828v1 | GCA_002722175.1_ASM272217v1 | GCA_002334965.1_ASM233496v1 | GCA_002389385.1_ASM238938v1 | GCA_002386815.1_ASM238681v1 | GCA_002389715.1_ASM238971v1 | GCA_002389175.1_ASM238917v1 | GCA_002327565.1_ASM232756v1 | GCA_002725625.1_ASM272562v1 | GCA_002687515.1_ASM268751v1 | GCA_002938035.1_ASM293803v1 | GCA_002712665.1_ASM271266v1 | GCA_002691065.1_ASM269106v1 | GCA_002170915.1_ASM217091v1 | GCA_002723345.1_ASM272334v1 | GCA_002168115.1_ASM216811v1 | GCA_002704405.1_ASM270440v1 | GCA_002697325.1_ASM269732v1 | GCA_002694925.1_ASM269492v1 | GCA_002937995.1_ASM293799v1 | GCA_002721295.1_ASM272129v1 |
|---|---|---|---|---|---|---|---|---|---|---|---|---|---|---|---|---|---|---|---|---|---|---|---|---|---|---|---|---|---|
| GCA_002938315.1_ASM293831v1 | 100 | 99 | 70 | 62 | 71 | 72 | 71 | 71 | 54 | 54 | 57 | 57 | 57 | 57 | 56 | 56 | 55 | 55 | 54 | 52 | 52 | 52 | 54 | 51 | 51 | 49 | 49 | 51 | 50 |
| GCA_002938085.1_ASM293808v1 | 99 | 100 | 68 | 60 | 70 | 71 | 71 | 70 | 53 | 54 | 57 | 57 | 57 | 56 | 56 | 56 | 54 | 55 | 53 | 52 | 51 | 50 | 54 | 51 | 51 | 48 | 49 | 50 | 49 |
| GCA_001510075.1_ASM151007v1 | 70 | 68 | 100 | 60 | 71 | 71 | 71 | 70 | 54 | 54 | 57 | 57 | 57 | 57 | 56 | 55 | 53 | 54 | 52 | 50 | 49 | 49 | 52 | 49 | 49 | 47 | 47 | 49 | 48 |
| GCA_002938045.1_ASM293804v1 | 62 | 60 | 60 | 100 | 63 | 63 | 63 | 62 | 53 | 55 | 56 | 56 | 56 | 56 | 56 | 54 | 53 | 53 | 52 | 49 | 49 | 49 | 52 | 49 | 49 | 45 | 47 | 49 | 47 |
| GCA_002686255.1_ASM268625v1 | 71 | 70 | 71 | 63 | 100 | 99 | 99 | 85 | 54 | 54 | 57 | 57 | 57 | 57 | 56 | 56 | 54 | 55 | 54 | 53 | 51 | 50 | 54 | 50 | 50 | 48 | 49 | 51 | 50 |
| GCA_002328385.1_ASM232838v1 | 72 | 71 | 71 | 63 | 99 | 100 | 99 | 85 | 54 | 54 | 58 | 57 | 58 | 57 | 57 | 56 | 54 | 56 | 54 | 52 | 51 | 51 | 54 | 51 | 51 | 48 | 49 | 51 | 50 |
| GCA_002420445.1_ASM242044v1 | 71 | 71 | 71 | 63 | 99 | 99 | 100 | 85 | 54 | 54 | 57 | 57 | 57 | 57 | 57 | 56 | 54 | 56 | 54 | 52 | 51 | 51 | 54 | 50 | 51 | 48 | 49 | 51 | 50 |
| GCA_002938075.1_ASM293807v1 | 71 | 70 | 70 | 62 | 85 | 85 | 85 | 100 | 54 | 55 | 58 | 57 | 57 | 57 | 57 | 55 | 54 | 54 | 53 | 51 | 51 | 50 | 53 | 50 | 50 | 47 | 48 | 50 | 49 |
| GCA_002938285.1_ASM293828v1 | 54 | 53 | 54 | 53 | 54 | 54 | 54 | 54 | 100 | 99 | 53 | 53 | 53 | 52 | 52 | 49 | 48 | 48 | 49 | 47 | 45 | 45 | 48 | 48 | 47 | 45 | 46 | 47 | 47 |
| GCA_002722175.1_ASM272217v1 | 54 | 54 | 54 | 55 | 54 | 54 | 54 | 55 | 99 | 100 | 53 | 53 | 54 | 52 | 52 | 49 | 48 | 49 | 49 | 48 | 47 | 46 | 49 | 48 | 48 | 46 | 46 | 48 | 47 |
| GCA_002334965.1_ASM233496v1 | 57 | 57 | 57 | 56 | 57 | 58 | 57 | 58 | 53 | 53 | 100 | 99 | 99 | 99 | 69 | 51 | 50 | 50 | 51 | 49 | 49 | 48 | 51 | 48 | 48 | 46 | 47 | 48 | 48 |
| GCA_002389385.1_ASM238938v1 | 57 | 57 | 57 | 56 | 57 | 57 | 57 | 57 | 53 | 53 | 99 | 100 | 99 | 99 | 69 | 51 | 50 | 50 | 51 | 48 | 49 | 48 | 51 | 48 | 48 | 45 | 47 | 48 | 47 |
| GCA_002386815.1_ASM238681v1 | 57 | 57 | 57 | 56 | 57 | 58 | 57 | 57 | 53 | 54 | 99 | 99 | 100 | 99 | 69 | 51 | 50 | 50 | 51 | 49 | 49 | 48 | 51 | 48 | 48 | 46 | 47 | 48 | 47 |
| GCA_002389715.1_ASM238971v1 | 57 | 56 | 57 | 56 | 57 | 57 | 57 | 57 | 52 | 52 | 99 | 99 | 99 | 100 | 68 | 51 | 49 | 50 | 50 | 49 | 49 | 48 | 51 | 48 | 48 | 45 | 46 | 48 | 47 |
| GCA_002389175.1_ASM238917v1 | 56 | 56 | 56 | 56 | 56 | 57 | 57 | 57 | 52 | 52 | 69 | 69 | 69 | 68 | 100 | 51 | 49 | 50 | 50 | 48 | 48 | 48 | 50 | 48 | 48 | 46 | 47 | 47 | 47 |
| GCA_002327565.1_ASM232756v1 | 56 | 56 | 55 | 54 | 56 | 56 | 56 | 55 | 49 | 49 | 51 | 51 | 51 | 51 | 51 | 100 | 67 | 64 | 61 | 60 | 60 | 60 | 62 | 63 | 62 | 56 | 56 | 57 | 56 |
| GCA_002725625.1_ASM272562v1 | 55 | 54 | 53 | 53 | 54 | 54 | 54 | 54 | 48 | 48 | 50 | 50 | 50 | 49 | 49 | 67 | 100 | 62 | 59 | 57 | 58 | 57 | 59 | 57 | 57 | 53 | 54 | 55 | 54 |
| GCA_002687515.1_ASM268751v1 | 55 | 55 | 54 | 53 | 55 | 56 | 56 | 54 | 48 | 49 | 50 | 50 | 50 | 50 | 50 | 64 | 62 | 100 | 59 | 57 | 58 | 57 | 59 | 55 | 54 | 54 | 54 | 54 | 54 |
| GCA_002938035.1_ASM293803v1 | 54 | 53 | 52 | 52 | 54 | 54 | 54 | 53 | 49 | 49 | 51 | 51 | 51 | 50 | 50 | 61 | 59 | 59 | 100 | 93 | 62 | 62 | 65 | 57 | 57 | 60 | 60 | 61 | 61 |
| GCA_002712665.1_ASM271266v1 | 52 | 52 | 50 | 49 | 53 | 52 | 52 | 51 | 47 | 48 | 49 | 48 | 49 | 49 | 48 | 60 | 57 | 57 | 93 | 100 | 61 | 61 | 63 | 55 | 55 | 58 | 59 | 60 | 59 |
| GCA_002691065.1_ASM269106v1 | 52 | 51 | 49 | 49 | 51 | 51 | 51 | 51 | 45 | 47 | 49 | 49 | 49 | 49 | 48 | 60 | 58 | 58 | 62 | 61 | 100 | 100 | 84 | 56 | 56 | 57 | 57 | 57 | 56 |
| GCA_002170915.1_ASM217091v1 | 52 | 50 | 49 | 49 | 50 | 51 | 51 | 50 | 45 | 46 | 48 | 48 | 48 | 48 | 48 | 60 | 57 | 57 | 62 | 61 | 100 | 100 | 83 | 56 | 56 | 57 | 57 | 57 | 56 |
| GCA_002723345.1_ASM272334v1 | 54 | 54 | 52 | 52 | 54 | 54 | 54 | 53 | 48 | 49 | 51 | 51 | 51 | 51 | 50 | 62 | 59 | 59 | 65 | 63 | 84 | 83 | 100 | 57 | 57 | 59 | 59 | 59 | 59 |
| GCA_002168115.1_ASM216811v1 | 51 | 51 | 49 | 49 | 50 | 51 | 50 | 50 | 48 | 48 | 48 | 48 | 48 | 48 | 48 | 63 | 57 | 55 | 57 | 55 | 56 | 56 | 57 | 100 | 100 | 55 | 56 | 56 | 55 |
| GCA_002704405.1_ASM270440v1 | 51 | 51 | 49 | 49 | 50 | 51 | 51 | 50 | 47 | 48 | 48 | 48 | 48 | 48 | 48 | 62 | 57 | 54 | 57 | 55 | 56 | 56 | 57 | 100 | 100 | 55 | 55 | 56 | 55 |
| GCA_002697325.1_ASM269732v1 | 49 | 48 | 47 | 45 | 48 | 48 | 48 | 47 | 45 | 46 | 46 | 45 | 46 | 45 | 46 | 56 | 53 | 54 | 60 | 58 | 57 | 57 | 59 | 55 | 55 | 100 | 95 | 94 | 92 |
| GCA_002694925.1_ASM269492v1 | 49 | 49 | 47 | 47 | 49 | 49 | 49 | 48 | 46 | 46 | 47 | 47 | 47 | 46 | 47 | 56 | 54 | 54 | 60 | 59 | 57 | 57 | 59 | 56 | 55 | 95 | 100 | 95 | 94 |
| GCA_002937995.1_ASM293799v1 | 51 | 50 | 49 | 49 | 51 | 51 | 51 | 50 | 47 | 48 | 48 | 48 | 48 | 48 | 47 | 57 | 55 | 54 | 61 | 60 | 57 | 57 | 59 | 56 | 56 | 94 | 95 | 100 | 96 |
| GCA_002721295.1_ASM272129v1 | 50 | 49 | 48 | 47 | 50 | 50 | 50 | 49 | 47 | 47 | 48 | 47 | 47 | 47 | 47 | 56 | 54 | 54 | 61 | 59 | 56 | 56 | 59 | 55 | 55 | 92 | 94 | 96 | 100 |

| | GCA_002728135.1_ASM | GCA_002718635.1_ASM | GCA_002296185.1_ASM | GCA_002717225.1_ASM | GCA_002937555.1_ASM | GCA_002937585.1_ASM | GCA_002937595.1_ASM | GCA_002938115.1_ASM | GCA_002377485.1_ASM | GCA_002728255.1_ASM | GCA_002501105.1_ASM | GCA_002721365.1_ASM |
|---|---|---|---|---|---|---|---|---|---|---|---|---|
| GCA_002728135.1_ASM272813v1 | 100 | 99 | 66 | 56 | 55 | 49 | 49 | 49 | 47 | 48 | 48 | 49 |
| GCA_002718635.1_ASM271863v1 | 99 | 100 | 64 | 54 | 54 | 47 | 47 | 47 | 46 | 46 | 47 | 48 |
| GCA_002296185.1_ASM229618v1 | 66 | 64 | 100 | 56 | 55 | 49 | 49 | 48 | 47 | 48 | 49 | 49 |
| GCA_002717225.1_ASM271722v1 | 56 | 54 | 56 | 100 | 74 | 51 | 51 | 50 | 48 | 49 | 51 | 51 |
| GCA_002937555.1_ASM293755v1 | 55 | 54 | 55 | 74 | 100 | 52 | 52 | 51 | 48 | 50 | 52 | 53 |
| GCA_002937585.1_ASM293758v1 | 49 | 47 | 49 | 51 | 52 | 100 | 100 | 78 | 62 | 63 | 58 | 55 |
| GCA_002937595.1_ASM293759v1 | 49 | 47 | 49 | 51 | 52 | 100 | 100 | 78 | 62 | 63 | 58 | 56 |
| GCA_002938115.1_ASM293811v1 | 49 | 47 | 48 | 50 | 51 | 78 | 78 | 100 | 61 | 63 | 57 | 55 |
| GCA_002377485.1_ASM237748v1 | 47 | 46 | 47 | 48 | 48 | 62 | 62 | 61 | 100 | 99 | 53 | 51 |
| GCA_002728255.1_ASM272825v1 | 48 | 46 | 48 | 49 | 50 | 63 | 63 | 63 | 99 | 100 | 55 | 52 |
| GCA_002501105.1_ASM250110v1 | 48 | 47 | 49 | 51 | 52 | 58 | 58 | 57 | 53 | 55 | 100 | 56 |
| GCA_002721365.1_ASM272136v1 | 49 | 48 | 49 | 51 | 53 | 55 | 56 | 55 | 51 | 52 | 56 | 100 |

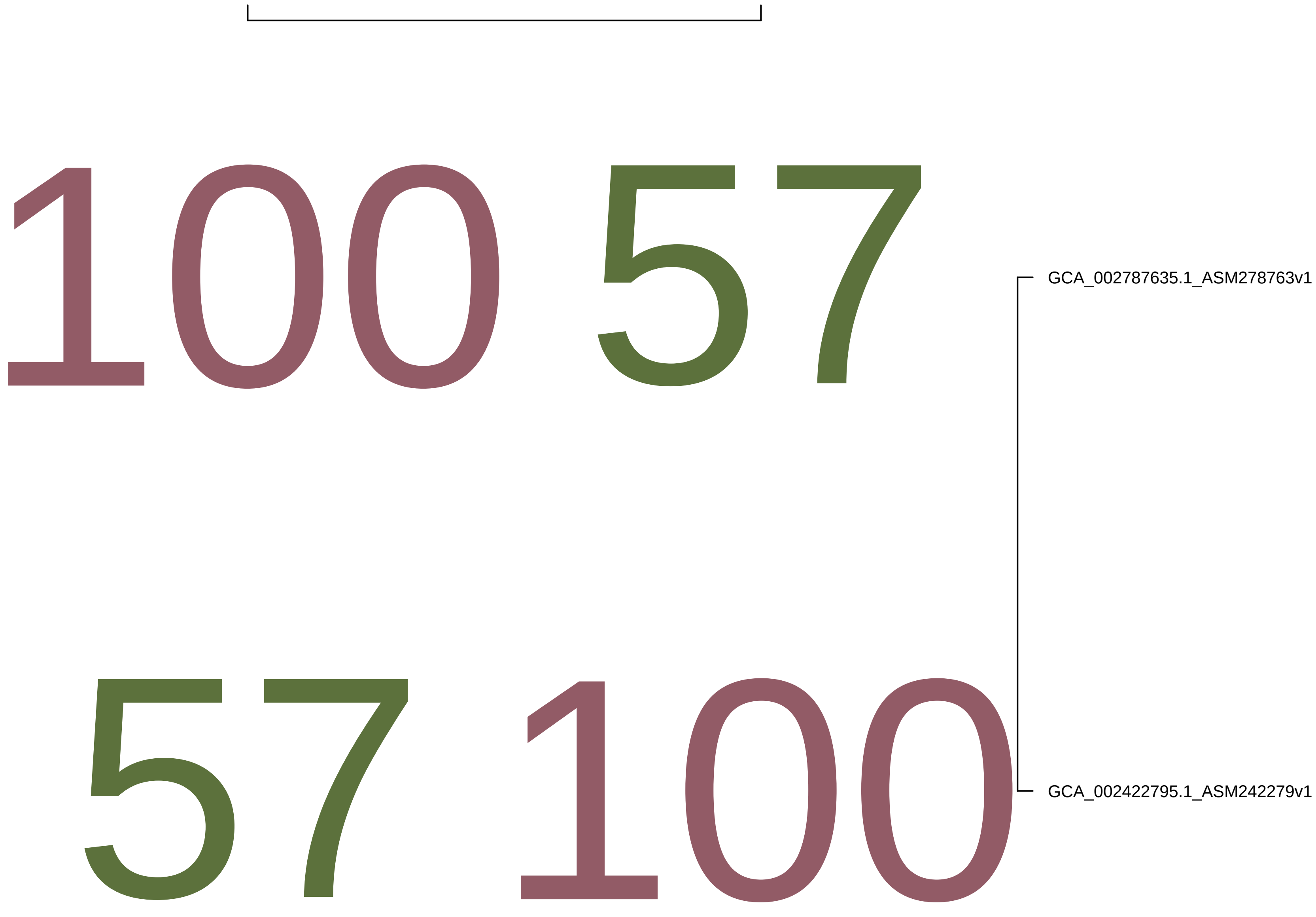
100 57
57 100
GCA_002787635.1_ASM278763v1
GCA_002422795.1_ASM242279v1

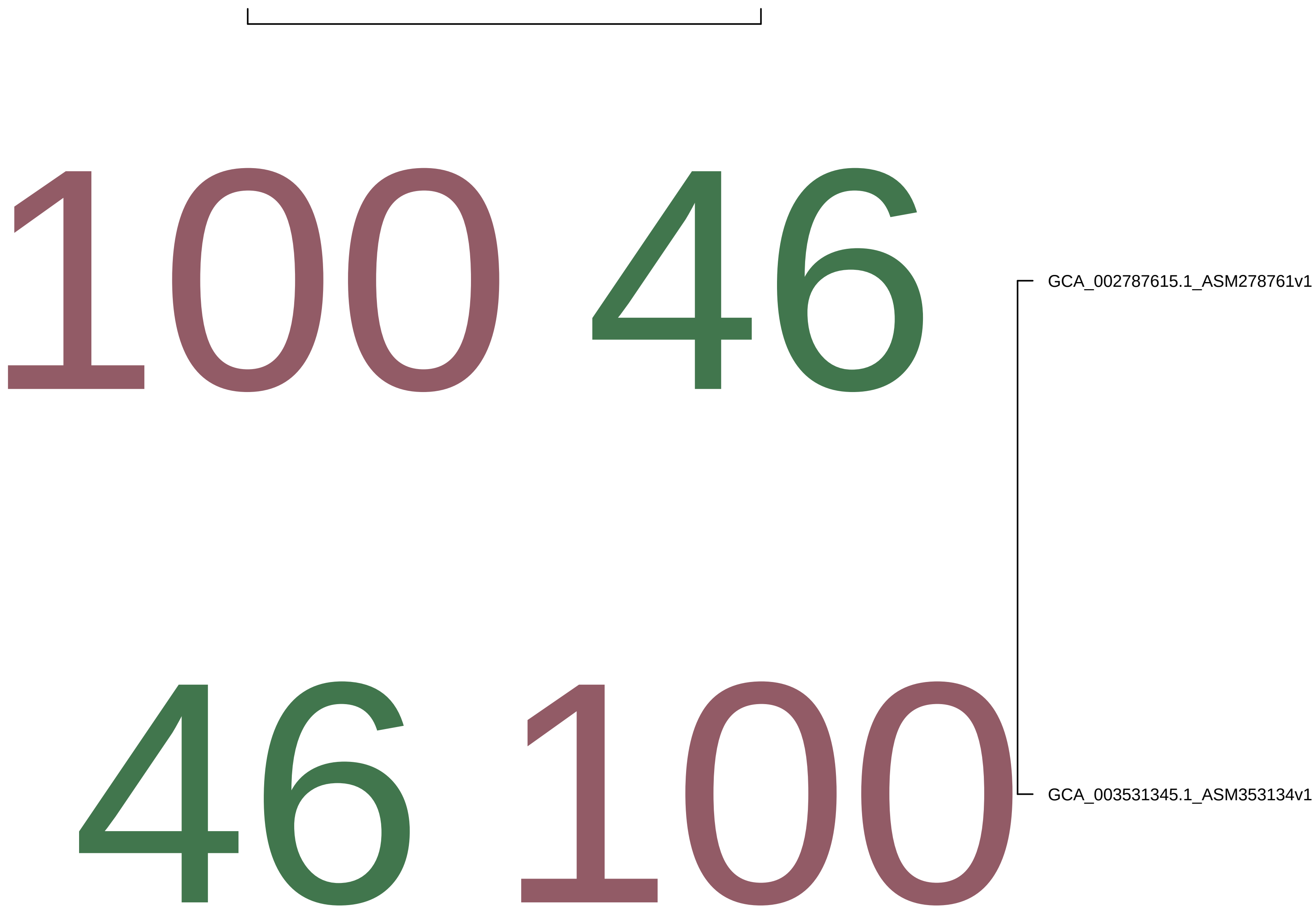
100 46
46 100
GCA_002787615.1_ASM278761v1
GCA_003531345.1_ASM353134v1

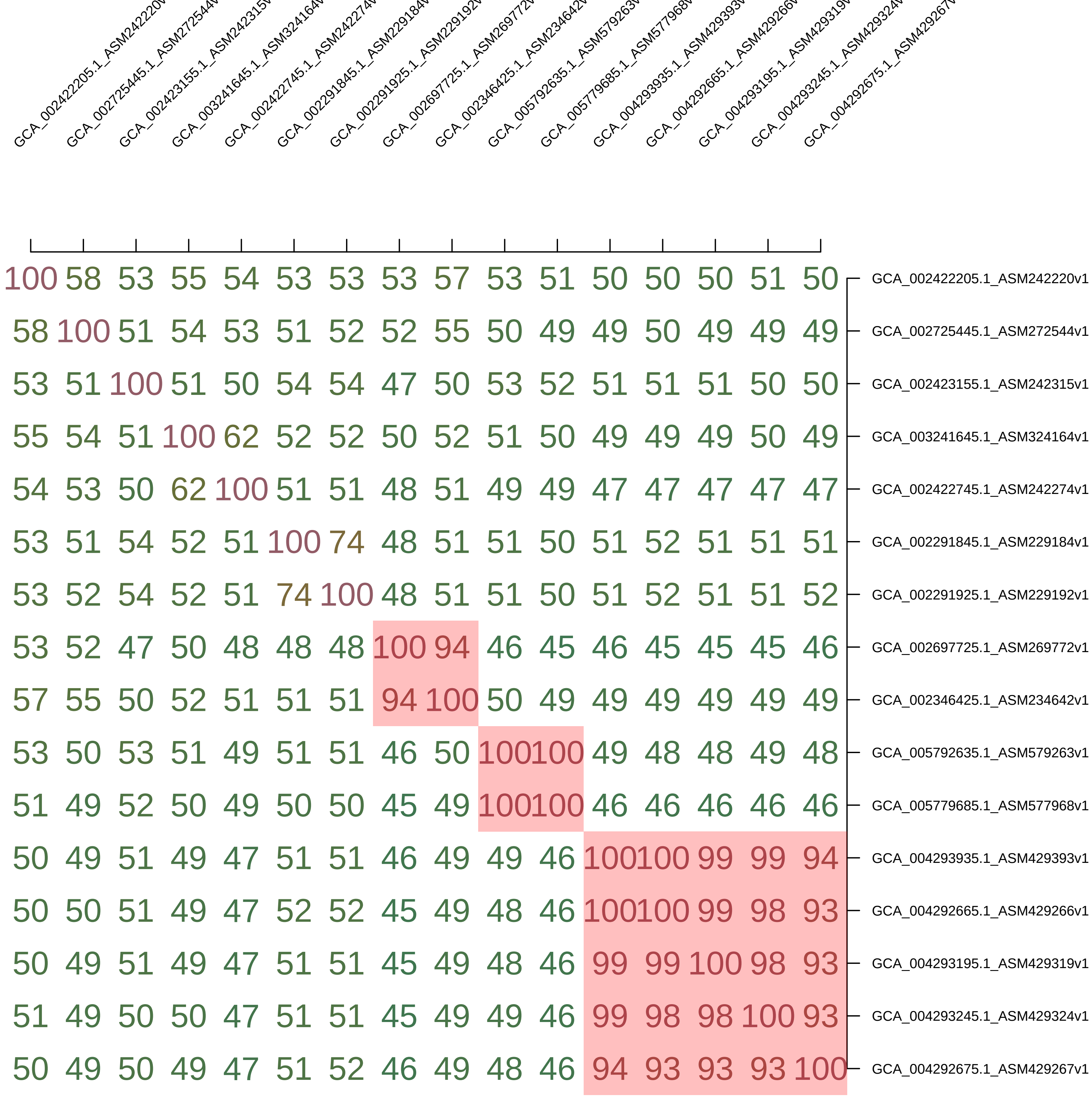
GCA_002422205.1_ASM242220v
GCA_002725445.1_ASM272544v
GCA_002423155.1_ASM242315v
GCA_003241645.1_ASM324164v
GCA_002422745.1_ASM242274v
GCA_002291845.1_ASM229184v
GCA_002291925.1_ASM229192v
GCA_002697725.1_ASM269772v
GCA_002346425.1_ASM234642v
GCA_005792635.1_ASM579263v
GCA_005779685.1_ASM577968v
GCA_004293935.1_ASM429393v
GCA_004292665.1_ASM429266v
GCA_004293195.1_ASM429319v
GCA_004293245.1_ASM429324v
GCA_004292675.1_ASM429267v
100 58 53 55 54 53 53 53 57 53 51 50 50 50 51 50 GCA_002422205.1_ASM242220v1
58 100 51 54 53 51 52 52 55 50 49 49 50 49 49 49 GCA_002725445.1_ASM272544v1
53 51 100 51 50 54 54 47 50 53 52 51 51 51 50 50 GCA_002423155.1_ASM242315v1
55 54 51 100 62 52 52 50 52 51 50 49 49 49 50 49 GCA_003241645.1_ASM324164v1
54 53 50 62 100 51 51 48 51 49 49 47 47 47 47 47 GCA_002422745.1_ASM242274v1
53 51 54 52 51 100 74 48 51 51 50 51 52 51 51 51 GCA_002291845.1_ASM229184v1
53 52 54 52 51 74 100 48 51 51 50 51 52 51 51 52 GCA_002291925.1_ASM229192v1
53 52 47 50 48 48 48 100 94 46 45 46 45 45 45 46 GCA_002697725.1_ASM269772v1
57 55 50 52 51 51 51 94 100 50 49 49 49 49 49 49 GCA_002346425.1_ASM234642v1
53 50 53 51 49 51 51 46 50 100 100 49 48 48 49 48 GCA_005792635.1_ASM579263v1
51 49 52 50 49 50 50 45 49 100 100 46 46 46 46 46 GCA_005779685.1_ASM577968v1
50 49 51 49 47 51 51 46 49 49 46 100 100 99 99 94 GCA_004293935.1_ASM429393v1
50 50 51 49 47 52 52 45 49 48 46 100 100 99 98 93 GCA_004292665.1_ASM429266v1
50 49 51 49 47 51 51 45 49 48 46 99 99 100 98 93 GCA_004293195.1_ASM429319v1
51 49 50 50 47 51 51 45 49 49 46 99 98 98 100 93 GCA_004293245.1_ASM429324v1
50 49 50 49 47 51 52 46 49 48 46 94 93 93 93 100 GCA_004292675.1_ASM429267v1

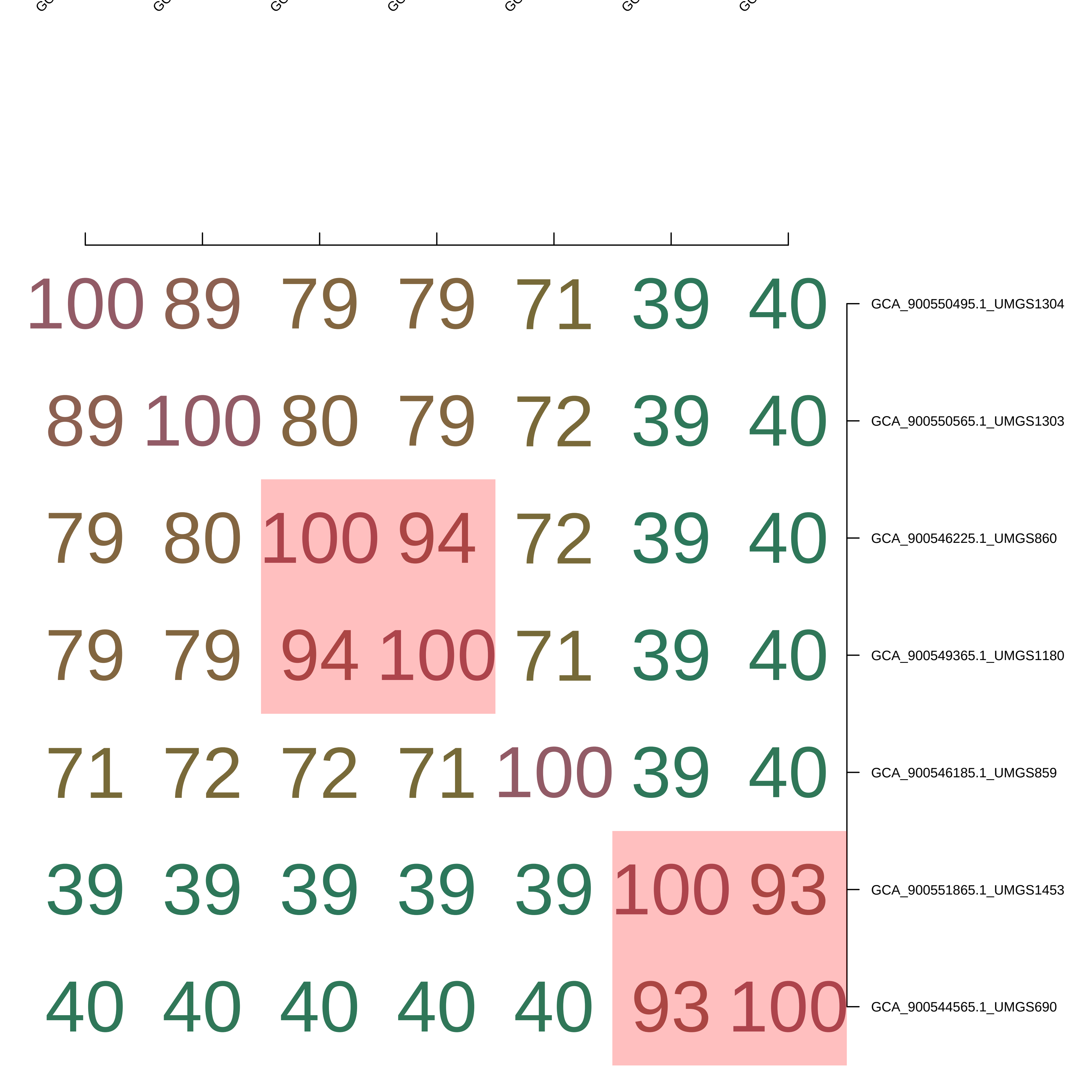

100 89 79 79 71 39 40 GCA_900550495.1_UMGS1304
89 100 80 79 72 39 40 GCA_900550565.1_UMGS1303
79 80 100 94 72 39 40 GCA_900546225.1_UMGS860
79 79 94 100 71 39 40 GCA_900549365.1_UMGS1180
71 72 72 71 100 39 40 GCA_900546185.1_UMGS859
39 39 39 39 39 100 93 GCA_900551865.1_UMGS1453
40 40 40 40 40 93 100 GCA_900544565.1_UMGS690

| | GCA_002938205.1_ASM293820v1 | GCA_002938235.1_ASM293823v1 | GCA_002718135.1_ASM271813v1 | GCA_002937895.1_ASM293789v1 | GCA_003282155.1_ASM328215v1 | GCA_003215235.1_ASM321523v1 | GCA_002938255.1_ASM293825v1 | GCA_002937955.1_ASM293795v1 | GCA_002937925.1_ASM293792v1 | GCA_002938195.1_ASM293819v1 | GCA_002937915.1_ASM293791v1 | GCA_002717485.1_ASM271748v1 | GCA_002938175.1_ASM293817v1 | GCA_002937875.1_ASM293787v1 | GCA_002937855.1_ASM293785v1 | GCA_002715915.1_ASM271591v1 | GCA_004321805.1_ASM432180v1 | GCF_000299115.1_ASM29911v1 | GCA_003216195.1_ASM321619v1 | GCA_002705265.1_ASM270526v1 |
|---|---|---|---|---|---|---|---|---|---|---|---|---|---|---|---|---|---|---|---|---|
| GCA_002938205.1_ASM293820v1 | 100 | 100 | 66 | 62 | 66 | 65 | 64 | 64 | 64 | 64 | 63 | 57 | 57 | 57 | 58 | 58 | 46 | 46 | 46 | 46 |
| GCA_002938235.1_ASM293823v1 | 100 | 100 | 66 | 62 | 65 | 65 | 64 | 64 | 64 | 64 | 63 | 57 | 56 | 56 | 58 | 58 | 45 | 47 | 46 | 46 |
| GCA_002718135.1_ASM271813v1 | 66 | 66 | 100 | 57 | 60 | 60 | 57 | 58 | 58 | 59 | 59 | 52 | 51 | 51 | 54 | 53 | 43 | 44 | 44 | 44 |
| GCA_002937895.1_ASM293789v1 | 62 | 62 | 57 | 100 | 59 | 59 | 61 | 61 | 63 | 63 | 63 | 59 | 58 | 58 | 60 | 60 | 46 | 46 | 46 | 46 |
| GCA_003282155.1_ASM328215v1 | 66 | 65 | 60 | 59 | 100 | 81 | 61 | 61 | 60 | 61 | 60 | 55 | 55 | 54 | 56 | 55 | 46 | 47 | 47 | 46 |
| GCA_003215235.1_ASM321523v1 | 65 | 65 | 60 | 59 | 81 | 100 | 60 | 60 | 60 | 61 | 59 | 55 | 54 | 53 | 55 | 54 | 45 | 46 | 47 | 46 |
| GCA_002938255.1_ASM293825v1 | 64 | 64 | 57 | 61 | 61 | 60 | 100 | 98 | 65 | 65 | 63 | 57 | 55 | 55 | 57 | 56 | 45 | 46 | 45 | 46 |
| GCA_002937955.1_ASM293795v1 | 64 | 64 | 58 | 61 | 61 | 60 | 98 | 100 | 64 | 65 | 64 | 56 | 55 | 55 | 57 | 55 | 45 | 45 | 45 | 45 |
| GCA_002937925.1_ASM293792v1 | 64 | 64 | 58 | 63 | 60 | 60 | 65 | 64 | 100 | 99 | 98 | 57 | 56 | 56 | 58 | 57 | 44 | 45 | 45 | 45 |
| GCA_002938195.1_ASM293819v1 | 64 | 64 | 59 | 63 | 61 | 61 | 65 | 65 | 99 | 100 | 99 | 57 | 57 | 57 | 58 | 57 | 46 | 46 | 45 | 45 |
| GCA_002937915.1_ASM293791v1 | 63 | 63 | 59 | 63 | 60 | 59 | 63 | 64 | 98 | 99 | 100 | 57 | 55 | 55 | 58 | 57 | 44 | 45 | 45 | 45 |
| GCA_002717485.1_ASM271748v1 | 57 | 57 | 52 | 59 | 55 | 55 | 57 | 56 | 57 | 57 | 57 | 100 | 99 | 99 | 66 | 65 | 45 | 46 | 45 | 45 |
| GCA_002938175.1_ASM293817v1 | 57 | 56 | 51 | 58 | 55 | 54 | 55 | 55 | 56 | 57 | 55 | 99 | 100 | 98 | 66 | 65 | 44 | 45 | 44 | 44 |
| GCA_002937875.1_ASM293787v1 | 57 | 56 | 51 | 58 | 54 | 53 | 55 | 55 | 56 | 57 | 55 | 99 | 98 | 100 | 65 | 65 | 44 | 45 | 44 | 44 |
| GCA_002937855.1_ASM293785v1 | 58 | 58 | 54 | 60 | 56 | 55 | 57 | 57 | 58 | 58 | 58 | 66 | 66 | 65 | 100 | 99 | 46 | 47 | 46 | 47 |
| GCA_002715915.1_ASM271591v1 | 58 | 58 | 53 | 60 | 55 | 54 | 56 | 55 | 57 | 57 | 57 | 65 | 65 | 65 | 99 | 100 | 45 | 46 | 46 | 45 |
| GCA_004321805.1_ASM432180v1 | 46 | 45 | 43 | 46 | 46 | 45 | 45 | 45 | 44 | 46 | 44 | 45 | 44 | 44 | 46 | 45 | 100 | 77 | 73 | 62 |
| GCF_000299115.1_ASM29911v1 | 46 | 47 | 44 | 46 | 47 | 46 | 46 | 45 | 45 | 46 | 45 | 46 | 45 | 45 | 47 | 46 | 77 | 100 | 73 | 63 |
| GCA_003216195.1_ASM321619v1 | 46 | 46 | 44 | 46 | 47 | 47 | 45 | 45 | 45 | 45 | 45 | 45 | 44 | 44 | 46 | 46 | 73 | 73 | 100 | 62 |
| GCA_002705265.1_ASM270526v1 | 46 | 46 | 44 | 46 | 46 | 46 | 46 | 45 | 45 | 45 | 45 | 45 | 44 | 44 | 47 | 45 | 62 | 63 | 62 | 100 |

| | GCA_002717265.1_ASM271726v1 | GCA_002937655.1_ASM293765v1 | GCA_002720535.1_ASM272053v1 | GCA_002171375.1_ASM217137v1 | GCA_002699795.1_ASM269979v1 | GCA_003331375.1_ASM333137v1 | GCA_002684605.1_ASM268460v1 | GCA_002715265.1_ASM271526v1 | GCA_002727615.1_ASM272761v1 | GCA_002937695.1_ASM293769v1 | GCA_002937675.1_ASM293767v1 | GCA_002714055.1_ASM271405v1 | GCA_002690715.1_ASM269071v1 | GCA_002170595.1_ASM217059v1 | GCA_002720125.1_ASM272012v1 | GCA_002171495.2_ASM217149v2 | GCA_002732015.1_ASM273201v1 | GCA_002715505.1_ASM271550v1 | GCA_002937625.1_ASM293762v1 |
|---|---|---|---|---|---|---|---|---|---|---|---|---|---|---|---|---|---|---|---|
| GCA_002717265.1_ASM271726v1 | 100 | 99 | 81 | 78 | 80 | 79 | 71 | 66 | 66 | 61 | 61 | 57 | 47 | 48 | 47 | 47 | 48 | 48 | 48 |
| GCA_002937655.1_ASM293765v1 | 99 | 100 | 81 | 80 | 81 | 81 | 71 | 67 | 66 | 61 | 62 | 60 | 48 | 48 | 47 | 48 | 49 | 48 | 49 |
| GCA_002720535.1_ASM272053v1 | 81 | 81 | 100 | 81 | 82 | 81 | 71 | 67 | 67 | 61 | 61 | 59 | 48 | 48 | 47 | 48 | 48 | 48 | 48 |
| GCA_002171375.1_ASM217137v1 | 78 | 80 | 81 | 100 | 100 | 97 | 72 | 67 | 67 | 61 | 62 | 59 | 48 | 48 | 47 | 47 | 48 | 48 | 48 |
| GCA_002699795.1_ASM269979v1 | 80 | 81 | 82 | 100 | 100 | 99 | 72 | 67 | 67 | 62 | 62 | 61 | 49 | 49 | 47 | 48 | 49 | 49 | 49 |
| GCA_003331375.1_ASM333137v1 | 79 | 81 | 81 | 97 | 99 | 100 | 72 | 67 | 67 | 61 | 62 | 58 | 48 | 48 | 47 | 47 | 48 | 48 | 48 |
| GCA_002684605.1_ASM268460v1 | 71 | 71 | 71 | 72 | 72 | 72 | 100 | 65 | 65 | 61 | 61 | 58 | 47 | 47 | 46 | 47 | 47 | 48 | 48 |
| GCA_002715265.1_ASM271526v1 | 66 | 67 | 67 | 67 | 67 | 67 | 65 | 100 | 97 | 61 | 62 | 59 | 48 | 48 | 47 | 48 | 49 | 49 | 49 |
| GCA_002727615.1_ASM272761v1 | 66 | 66 | 67 | 67 | 67 | 67 | 65 | 97 | 100 | 61 | 61 | 58 | 47 | 48 | 47 | 47 | 48 | 48 | 48 |
| GCA_002937695.1_ASM293769v1 | 61 | 61 | 61 | 61 | 62 | 61 | 61 | 61 | 61 | 100 | 99 | 74 | 47 | 48 | 47 | 48 | 48 | 48 | 48 |
| GCA_002937675.1_ASM293767v1 | 61 | 62 | 61 | 62 | 62 | 62 | 61 | 62 | 61 | 99 | 100 | 74 | 48 | 48 | 47 | 48 | 48 | 48 | 49 |
| GCA_002714055.1_ASM271405v1 | 57 | 60 | 59 | 59 | 61 | 58 | 58 | 59 | 58 | 74 | 74 | 100 | 45 | 47 | 44 | 46 | 46 | 47 | 47 |
| GCA_002690715.1_ASM269071v1 | 47 | 48 | 48 | 48 | 49 | 48 | 47 | 48 | 47 | 47 | 48 | 45 | 100 | 99 | 59 | 59 | 60 | 60 | 60 |
| GCA_002170595.1_ASM217059v1 | 48 | 48 | 48 | 48 | 49 | 48 | 47 | 48 | 48 | 48 | 48 | 47 | 99 | 100 | 59 | 60 | 60 | 60 | 60 |
| GCA_002720125.1_ASM272012v1 | 47 | 47 | 47 | 47 | 47 | 47 | 46 | 47 | 47 | 47 | 47 | 44 | 59 | 59 | 100 | 59 | 60 | 60 | 60 |
| GCA_002171495.2_ASM217149v2 | 47 | 48 | 48 | 47 | 48 | 47 | 47 | 48 | 47 | 48 | 48 | 46 | 59 | 60 | 59 | 100 | 82 | 73 | 73 |
| GCA_002732015.1_ASM273201v1 | 48 | 49 | 48 | 48 | 49 | 48 | 47 | 49 | 48 | 48 | 48 | 46 | 60 | 60 | 60 | 82 | 100 | 74 | 74 |
| GCA_002715505.1_ASM271550v1 | 48 | 48 | 48 | 48 | 49 | 48 | 48 | 49 | 48 | 48 | 48 | 47 | 60 | 60 | 60 | 73 | 74 | 100 | 100 |
| GCA_002937625.1_ASM293762v1 | 48 | 49 | 48 | 48 | 49 | 48 | 48 | 49 | 48 | 48 | 49 | 47 | 60 | 60 | 60 | 73 | 74 | 100 | 100 |

| | GCA_002937755.1_ASM293775v1 | GCA_002717245.1_ASM271724v1 | GCA_002720895.1_ASM272089v1 | GCA_002937775.1_ASM293777v1 | GCA_002937795.1_ASM293779v1 | GCA_003446135.1_ASM344613v1 | GCA_002712065.1_ASM271206v1 | GCA_002731625.1_ASM273162v1 | GCA_002696825.1_ASM269682v1 | GCA_002691145.1_ASM269114v1 | GCA_002170155.1_ASM217015v1 | GCA_002710765.1_ASM271076v1 | GCA_002713965.1_ASM271396v1 | GCA_002720775.1_ASM272077v1 | GCA_002702805.1_ASM270280v1 | GCA_002711515.1_ASM271151v1 | GCA_002691245.1_ASM269124v1 | GCA_002689845.1_ASM268984v1 | GCA_002711625.1_ASM271162v1 | GCA_002690875.1_ASM269087v1 | GCA_002689325.1_ASM268932v1 | GCA_002715305.1_ASM271530v1 | GCA_002706585.1_ASM270658v1 |
|---|---|---|---|---|---|---|---|---|---|---|---|---|---|---|---|---|---|---|---|---|---|---|---|
| GCA_002937755.1_ASM293775v1 | 100 | 99 | 75 | 77 | 77 | 47 | 48 | 47 | 48 | 47 | 45 | 44 | 45 | 44 | 45 | 45 | 46 | 45 | 45 | 45 | 46 | 43 | 43 |
| GCA_002717245.1_ASM271724v1 | 99 | 100 | 76 | 77 | 77 | 47 | 47 | 47 | 48 | 46 | 45 | 45 | 46 | 44 | 45 | 45 | 46 | 45 | 45 | 45 | 46 | 43 | 43 |
| GCA_002720895.1_ASM272089v1 | 75 | 76 | 100 | 74 | 71 | 46 | 47 | 46 | 47 | 46 | 44 | 45 | 45 | 43 | 44 | 43 | 45 | 45 | 44 | 44 | 45 | 43 | 42 |
| GCA_002937775.1_ASM293777v1 | 77 | 77 | 74 | 100 | 97 | 47 | 47 | 46 | 48 | 46 | 46 | 45 | 45 | 43 | 44 | 45 | 46 | 46 | 44 | 45 | 46 | 42 | 44 |
| GCA_002937795.1_ASM293779v1 | 77 | 77 | 71 | 97 | 100 | 46 | 47 | 46 | 47 | 46 | 44 | 44 | 45 | 44 | 44 | 44 | 46 | 45 | 44 | 44 | 45 | 42 | 43 |
| GCA_003446135.1_ASM344613v1 | 47 | 47 | 46 | 47 | 46 | 100 | 97 | 95 | 94 | 46 | 44 | 44 | 45 | 43 | 44 | 44 | 45 | 45 | 44 | 44 | 45 | 42 | 42 |
| GCA_002712065.1_ASM271206v1 | 48 | 47 | 47 | 47 | 47 | 97 | 100 | 97 | 95 | 46 | 45 | 45 | 46 | 45 | 45 | 45 | 46 | 46 | 45 | 45 | 46 | 43 | 43 |
| GCA_002731625.1_ASM273162v1 | 47 | 47 | 46 | 46 | 46 | 95 | 97 | 100 | 95 | 46 | 45 | 45 | 45 | 44 | 44 | 44 | 45 | 45 | 44 | 45 | 46 | 42 | 43 |
| GCA_002696825.1_ASM269682v1 | 48 | 48 | 47 | 48 | 47 | 94 | 95 | 95 | 100 | 46 | 45 | 45 | 46 | 44 | 45 | 45 | 45 | 46 | 45 | 45 | 46 | 43 | 43 |
| GCA_002691145.1_ASM269114v1 | 47 | 46 | 46 | 46 | 46 | 46 | 46 | 46 | 46 | 100 | 100 | 59 | 59 | 55 | 56 | 57 | 56 | 56 | 57 | 55 | 57 | 55 | 53 |
| GCA_002170155.1_ASM217015v1 | 45 | 45 | 44 | 46 | 44 | 44 | 45 | 45 | 45 | 100 | 100 | 57 | 57 | 53 | 54 | 55 | 55 | 55 | 55 | 55 | 56 | 53 | 53 |
| GCA_002710765.1_ASM271076v1 | 44 | 45 | 45 | 45 | 44 | 44 | 45 | 45 | 45 | 59 | 57 | 100 | 57 | 54 | 54 | 54 | 55 | 55 | 54 | 54 | 56 | 53 | 51 |
| GCA_002713965.1_ASM271396v1 | 45 | 46 | 45 | 45 | 45 | 45 | 46 | 45 | 46 | 59 | 57 | 57 | 100 | 85 | 55 | 55 | 56 | 55 | 56 | 54 | 56 | 54 | 52 |
| GCA_002720775.1_ASM272077v1 | 44 | 44 | 43 | 43 | 44 | 43 | 45 | 44 | 44 | 55 | 53 | 54 | 85 | 100 | 52 | 53 | 53 | 53 | 53 | 51 | 54 | 49 | 49 |
| GCA_002702805.1_ASM270280v1 | 45 | 45 | 44 | 44 | 44 | 44 | 45 | 44 | 45 | 56 | 54 | 54 | 55 | 52 | 100 | 63 | 60 | 60 | 59 | 64 | 65 | 58 | 55 |
| GCA_002711515.1_ASM271151v1 | 45 | 45 | 43 | 45 | 44 | 44 | 45 | 44 | 45 | 57 | 55 | 54 | 55 | 53 | 63 | 100 | 62 | 60 | 60 | 62 | 65 | 57 | 58 |
| GCA_002691245.1_ASM269124v1 | 46 | 46 | 45 | 46 | 46 | 45 | 46 | 45 | 45 | 56 | 55 | 55 | 56 | 53 | 60 | 62 | 100 | 63 | 60 | 60 | 62 | 60 | 57 |
| GCA_002689845.1_ASM268984v1 | 45 | 45 | 45 | 46 | 45 | 45 | 46 | 45 | 46 | 56 | 55 | 55 | 55 | 53 | 60 | 60 | 63 | 100 | 59 | 60 | 61 | 58 | 57 |
| GCA_002711625.1_ASM271162v1 | 45 | 45 | 44 | 44 | 44 | 44 | 45 | 44 | 45 | 57 | 55 | 54 | 56 | 53 | 59 | 60 | 60 | 59 | 100 | 58 | 60 | 64 | 59 |
| GCA_002690875.1_ASM269087v1 | 45 | 45 | 44 | 45 | 44 | 44 | 45 | 45 | 45 | 55 | 55 | 54 | 54 | 51 | 64 | 62 | 60 | 60 | 58 | 100 | 86 | 58 | 56 |
| GCA_002689325.1_ASM268932v1 | 46 | 46 | 45 | 46 | 45 | 45 | 46 | 46 | 46 | 57 | 56 | 56 | 56 | 54 | 65 | 65 | 62 | 61 | 60 | 86 | 100 | 59 | 59 |
| GCA_002715305.1_ASM271530v1 | 43 | 43 | 43 | 42 | 42 | 42 | 43 | 42 | 43 | 55 | 53 | 53 | 54 | 49 | 58 | 57 | 60 | 58 | 64 | 58 | 59 | 100 | 71 |
| GCA_002706585.1_ASM270658v1 | 43 | 43 | 42 | 44 | 43 | 42 | 43 | 43 | 43 | 53 | 53 | 51 | 52 | 49 | 55 | 58 | 57 | 57 | 59 | 56 | 59 | 71 | 100 |

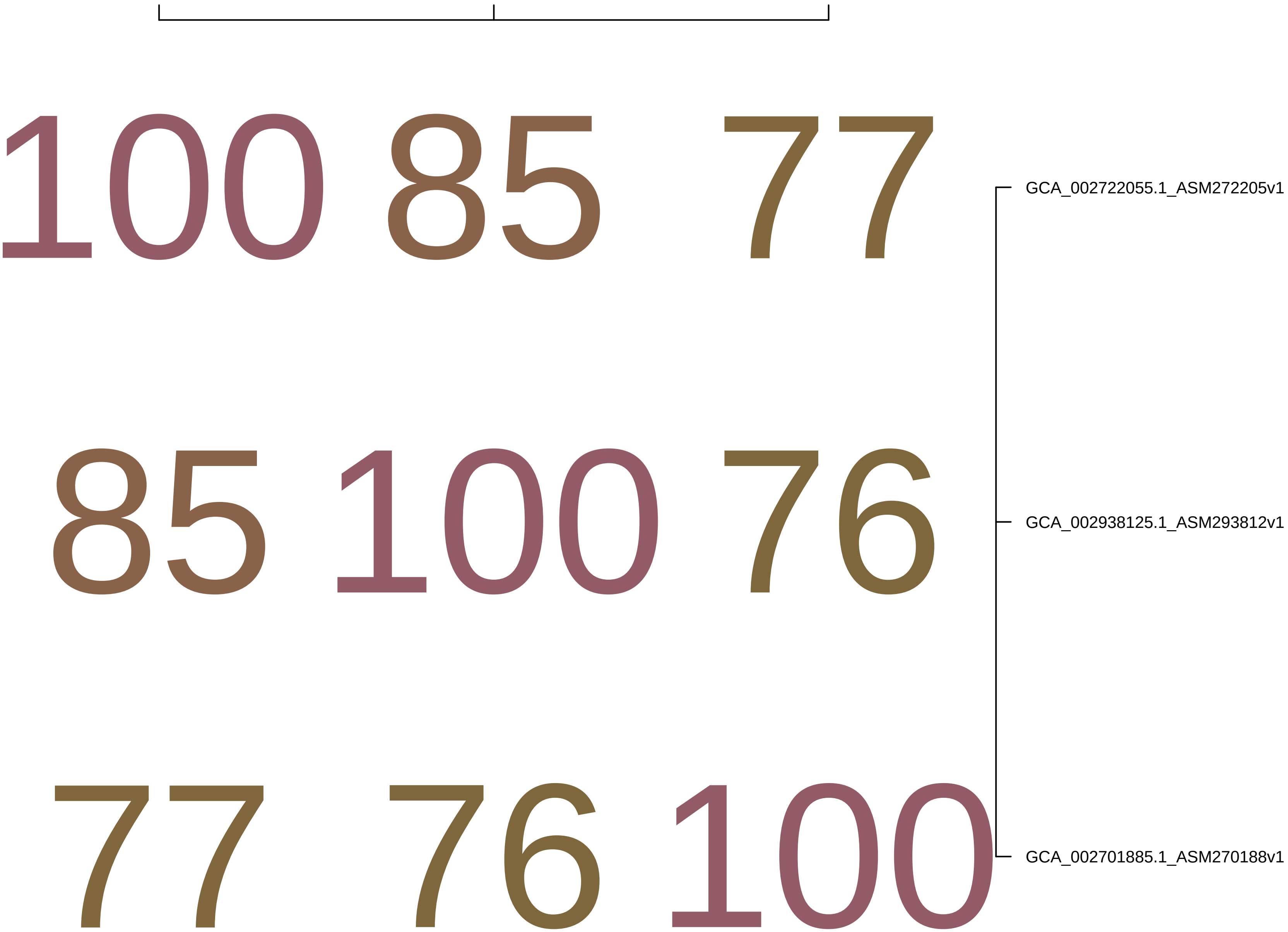
100 85 77
85 100 76
77 76 100
GCA_002722055.1_ASM272205v1
GCA_002938125.1_ASM293812v1
GCA_002701885.1_ASM270188v1

| | GCA_905478245.1_0.2_20180403_Bin_MAX | GCA_905478165.1_0.2_20180403_Bin_25_s | GCA_016780625.1_ASM1678062v1 | GCA_905480015.1_10_20180529_Bin_6_1 | GCA_018659605.1_ASM1865960v1 | GCA_018659145.1_ASM1865914v1 | GCA_016778805.1_ASM1677880v1 | GCA_017620845.1_ASM1762084v1 | GCA_017475895.1_ASM1747589v1 | GCA_017444095.1_ASM1744409v1 | GCA_017444125.1_ASM1744412v1 | GCA_002395105.1_ASM239510v1 | GCA_002394665.1_ASM239466v1 | GCA_014190355.1_ASM1419035v1 | GCA_903837365.1_freshwater_MAG_---_L | GCA_009921385.1_ASM992138v1 | GCA_903959715.1_freshwater_MAG_---_L | GCA_005787525.1_ASM578752v1 | GCA_009925685.1_ASM992568v1 | GCA_009924125.1_ASM992412v1 | GCA_002359655.1_ASM235965v1 | GCA_018969185.1_ASM1896918v1 | GCA_009693885.1_ASM969388v1 | GCA_903917585.1_freshwater_MAG_---_L | GCA_001768015.1_ASM176801v1 | GCA_903927415.1_freshwater_MAG_---_L | GCA_903878685.1_freshwater_MAG_---_L | GCA_017999675.1_ASM1799967v1 | GCA_903831915.1_freshwater_MAG_---_K | GCA_903872175.1_freshwater_MAG_---_L | GCA_016778745.1_ASM1677874v1 |
|---|---|---|---|---|---|---|---|---|---|---|---|---|---|---|---|---|---|---|---|---|---|---|---|---|---|---|---|---|---|---|---|
| GCA_905478245.1_0.2_20180403_B | 100 | 54 | 47 | 42 | 46 | 46 | 46 | 43 | 43 | 38 | 38 | 39 | 38 | 53 | 53 | 51 | 53 | 53 | 53 | 54 | 55 | 55 | 53 | 54 | 55 | 55 | 55 | 54 | 54 | 54 | 54 |
| GCA_905478165.1_0.2_20180403_B | 54 | 100 | 46 | 42 | 45 | 46 | 46 | 42 | 42 | 38 | 38 | 38 | 38 | 50 | 50 | 49 | 50 | 50 | 51 | 51 | 51 | 51 | 50 | 51 | 51 | 52 | 52 | 51 | 51 | 51 | 51 |
| GCA_016780625.1_ASM1678062v1 | 47 | 46 | 100 | 42 | 49 | 50 | 50 | 41 | 40 | 38 | 38 | 39 | 39 | 46 | 45 | 45 | 46 | 46 | 46 | 46 | 46 | 46 | 44 | 46 | 46 | 46 | 46 | 45 | 45 | 45 | 46 |
| GCA_905480015.1_10_20180529_B | 42 | 42 | 42 | 100 | 41 | 41 | 41 | 37 | 38 | 37 | 37 | 37 | 38 | 41 | 41 | 41 | 41 | 41 | 41 | 41 | 41 | 41 | 40 | 42 | 41 | 42 | 41 | 41 | 41 | 41 | 41 |
| GCA_018659605.1_ASM1865960v1 | 46 | 45 | 49 | 41 | 100 | 76 | 67 | 39 | 39 | 37 | 38 | 38 | 38 | 44 | 44 | 44 | 44 | 44 | 44 | 44 | 44 | 45 | 43 | 44 | 45 | 44 | 44 | 44 | 44 | 44 | 44 |
| GCA_018659145.1_ASM1865914v1 | 46 | 46 | 50 | 41 | 76 | 100 | 66 | 39 | 40 | 37 | 38 | 38 | 38 | 44 | 44 | 44 | 44 | 44 | 44 | 44 | 44 | 45 | 44 | 45 | 45 | 45 | 45 | 45 | 44 | 44 | 45 |
| GCA_016778805.1_ASM1677880v1 | 46 | 46 | 50 | 41 | 67 | 66 | 100 | 40 | 40 | 37 | 38 | 38 | 37 | 45 | 44 | 44 | 45 | 45 | 45 | 45 | 45 | 45 | 44 | 45 | 45 | 45 | 45 | 45 | 44 | 44 | 45 |
| GCA_017620845.1_ASM1762084v1 | 43 | 42 | 41 | 37 | 39 | 39 | 40 | 100 | 69 | 38 | 38 | 38 | 38 | 42 | 42 | 42 | 42 | 42 | 42 | 42 | 42 | 43 | 41 | 42 | 42 | 42 | 43 | 42 | 42 | 42 | 42 |
| GCA_017475895.1_ASM1747589v1 | 43 | 42 | 40 | 38 | 39 | 40 | 40 | 69 | 100 | 39 | 38 | 38 | 38 | 42 | 42 | 41 | 42 | 42 | 42 | 42 | 42 | 43 | 41 | 42 | 43 | 42 | 42 | 42 | 42 | 42 | 42 |
| GCA_017444095.1_ASM1744409v1 | 38 | 38 | 38 | 37 | 37 | 37 | 37 | 38 | 39 | 100 | 55 | 54 | 54 | 38 | 38 | 38 | 38 | 37 | 38 | 37 | 37 | 38 | 37 | 38 | 38 | 38 | 38 | 38 | 37 | 38 | 38 |
| GCA_017444125.1_ASM1744412v1 | 38 | 38 | 38 | 37 | 38 | 38 | 38 | 38 | 38 | 55 | 100 | 53 | 55 | 38 | 38 | 38 | 38 | 37 | 38 | 38 | 38 | 38 | 37 | 38 | 38 | 38 | 38 | 38 | 38 | 38 | 38 |
| GCA_002395105.1_ASM239510v1 | 39 | 38 | 39 | 37 | 38 | 38 | 38 | 38 | 38 | 54 | 53 | 100 | 56 | 38 | 38 | 38 | 38 | 38 | 39 | 38 | 38 | 38 | 36 | 38 | 38 | 38 | 38 | 38 | 38 | 38 | 38 |
| GCA_002394665.1_ASM239466v1 | 38 | 38 | 39 | 38 | 38 | 38 | 37 | 38 | 38 | 54 | 55 | 56 | 100 | 39 | 38 | 38 | 38 | 38 | 39 | 38 | 38 | 39 | 37 | 39 | 39 | 38 | 38 | 39 | 39 | 39 | 38 |
| GCA_014190355.1_ASM1419035v1 | 53 | 50 | 46 | 41 | 44 | 44 | 45 | 42 | 42 | 38 | 38 | 38 | 39 | 100 | 70 | 57 | 61 | 61 | 61 | 59 | 60 | 59 | 58 | 60 | 62 | 61 | 61 | 60 | 60 | 60 | 57 |
| GCA_903837365.1_freshwater_MAG | 53 | 50 | 45 | 41 | 44 | 44 | 44 | 42 | 42 | 38 | 38 | 38 | 38 | 70 | 100 | 57 | 62 | 62 | 61 | 60 | 61 | 60 | 58 | 62 | 63 | 62 | 62 | 61 | 61 | 62 | 57 |
| GCA_009921385.1_ASM992138v1 | 51 | 49 | 45 | 41 | 44 | 44 | 44 | 42 | 41 | 38 | 38 | 38 | 38 | 57 | 57 | 100 | 58 | 58 | 58 | 56 | 56 | 56 | 54 | 56 | 57 | 57 | 57 | 56 | 57 | 56 | 54 |
| GCA_903959715.1_freshwater_MAG | 53 | 50 | 46 | 41 | 44 | 44 | 45 | 42 | 42 | 38 | 38 | 38 | 38 | 61 | 62 | 58 | 100 | 86 | 72 | 61 | 61 | 60 | 58 | 62 | 62 | 62 | 62 | 62 | 62 | 62 | 58 |
| GCA_005787525.1_ASM578752v1 | 53 | 50 | 46 | 41 | 44 | 44 | 45 | 42 | 42 | 37 | 37 | 38 | 38 | 61 | 62 | 58 | 86 | 100 | 72 | 61 | 61 | 60 | 58 | 61 | 62 | 62 | 62 | 62 | 62 | 62 | 57 |
| GCA_009925685.1_ASM992568v1 | 53 | 51 | 46 | 41 | 44 | 44 | 45 | 42 | 42 | 38 | 38 | 39 | 39 | 61 | 61 | 58 | 72 | 72 | 100 | 60 | 60 | 60 | 57 | 61 | 62 | 62 | 61 | 61 | 61 | 61 | 57 |
| GCA_009924125.1_ASM992412v1 | 54 | 51 | 46 | 41 | 44 | 44 | 45 | 42 | 42 | 37 | 38 | 38 | 38 | 59 | 60 | 56 | 61 | 61 | 60 | 100 | 100 | 68 | 63 | 67 | 67 | 70 | 70 | 69 | 70 | 71 | 60 |
| GCA_002359655.1_ASM235965v1 | 55 | 51 | 46 | 41 | 44 | 44 | 45 | 42 | 42 | 37 | 38 | 38 | 38 | 60 | 61 | 56 | 61 | 61 | 60 | 100 | 100 | 68 | 63 | 67 | 67 | 70 | 70 | 69 | 70 | 71 | 60 |
| GCA_018969185.1_ASM1896918v1 | 55 | 51 | 46 | 41 | 45 | 45 | 45 | 43 | 43 | 38 | 38 | 38 | 39 | 59 | 60 | 56 | 60 | 60 | 60 | 68 | 68 | 100 | 62 | 65 | 65 | 67 | 67 | 66 | 67 | 67 | 60 |
| GCA_009693885.1_ASM969388v1 | 53 | 50 | 44 | 40 | 43 | 44 | 44 | 41 | 41 | 37 | 37 | 36 | 37 | 58 | 58 | 54 | 58 | 58 | 57 | 63 | 63 | 62 | 100 | 62 | 63 | 64 | 64 | 63 | 64 | 63 | 57 |
| GCA_903917585.1_freshwater_MAG | 54 | 51 | 46 | 42 | 44 | 45 | 45 | 42 | 42 | 38 | 38 | 38 | 39 | 60 | 62 | 56 | 62 | 61 | 61 | 67 | 67 | 65 | 62 | 100 | 69 | 70 | 70 | 68 | 69 | 68 | 60 |
| GCA_001768015.1_ASM176801v1 | 55 | 51 | 46 | 41 | 45 | 45 | 45 | 42 | 43 | 38 | 38 | 38 | 39 | 62 | 63 | 57 | 62 | 62 | 62 | 67 | 67 | 65 | 63 | 69 | 100 | 71 | 71 | 69 | 69 | 69 | 61 |
| GCA_903927415.1_freshwater_MAG | 55 | 52 | 46 | 42 | 44 | 45 | 45 | 42 | 42 | 38 | 38 | 38 | 38 | 61 | 62 | 57 | 62 | 62 | 62 | 70 | 70 | 67 | 64 | 70 | 71 | 100 | 81 | 72 | 72 | 73 | 61 |
| GCA_903878685.1_freshwater_MAG | 55 | 52 | 46 | 41 | 44 | 45 | 45 | 43 | 42 | 38 | 38 | 38 | 38 | 61 | 62 | 57 | 62 | 62 | 61 | 70 | 70 | 67 | 64 | 70 | 71 | 81 | 100 | 71 | 72 | 72 | 61 |
| GCA_017999675.1_ASM1799967v1 | 54 | 51 | 45 | 41 | 44 | 45 | 45 | 42 | 42 | 38 | 38 | 38 | 39 | 60 | 61 | 56 | 62 | 62 | 61 | 69 | 69 | 66 | 63 | 68 | 69 | 72 | 71 | 100 | 82 | 74 | 61 |
| GCA_903831915.1_freshwater_MAG | 54 | 51 | 45 | 41 | 44 | 44 | 44 | 42 | 42 | 37 | 38 | 38 | 39 | 60 | 61 | 57 | 62 | 62 | 61 | 70 | 70 | 67 | 64 | 69 | 69 | 72 | 72 | 82 | 100 | 74 | 61 |
| GCA_903872175.1_freshwater_MAG | 54 | 51 | 45 | 41 | 44 | 44 | 44 | 42 | 42 | 38 | 38 | 38 | 39 | 60 | 62 | 56 | 62 | 62 | 61 | 71 | 71 | 67 | 63 | 68 | 69 | 73 | 72 | 74 | 74 | 100 | 62 |
| GCA_016778745.1_ASM1677874v1 | 54 | 51 | 46 | 41 | 44 | 45 | 45 | 42 | 42 | 38 | 38 | 38 | 38 | 57 | 57 | 54 | 58 | 57 | 57 | 60 | 60 | 60 | 57 | 60 | 61 | 61 | 61 | 61 | 61 | 62 | 100 |

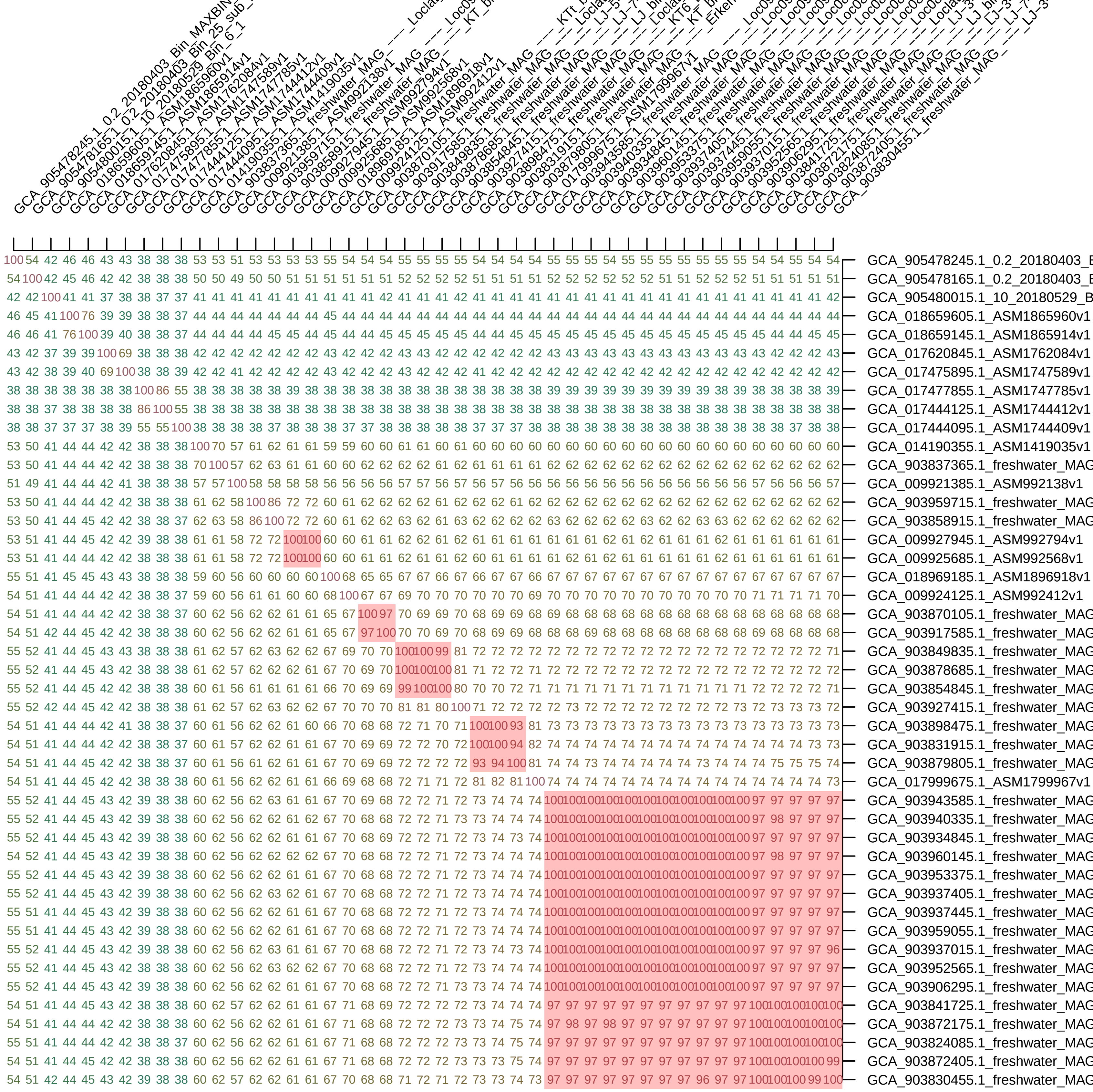
GCA_905478245.1_0.2_20180403_B
GCA_905478165.1_0.2_20180403_B
GCA_905480015.1_10_20180529_B
GCA_018659605.1_ASM1865960v1
GCA_018659145.1_ASM1865914v1
GCA_017620845.1_ASM1762084v1
GCA_017475895.1_ASM1747589v1
GCA_017477855.1_ASM1747785v1
GCA_017444125.1_ASM1744412v1
GCA_017444095.1_ASM1744409v1
GCA_014190355.1_ASM1419035v1
GCA_903837365.1_freshwater_MAG
GCA_009921385.1_ASM992138v1
GCA_903959715.1_freshwater_MAG
GCA_903858915.1_freshwater_MAG
GCA_009927945.1_ASM992794v1
GCA_009925685.1_ASM992568v1
GCA_018969185.1_ASM1896918v1
GCA_009924125.1_ASM992412v1
GCA_903870105.1_freshwater_MAG
GCA_903917585.1_freshwater_MAG
GCA_903849835.1_freshwater_MAG
GCA_903878685.1_freshwater_MAG
GCA_903854845.1_freshwater_MAG
GCA_903927415.1_freshwater_MAG
GCA_903898475.1_freshwater_MAG
GCA_903831915.1_freshwater_MAG
GCA_903879805.1_freshwater_MAG
GCA_017999675.1_ASM1799967v1
GCA_903943585.1_freshwater_MAG
GCA_903940335.1_freshwater_MAG
GCA_903934845.1_freshwater_MAG
GCA_903960145.1_freshwater_MAG
GCA_903953375.1_freshwater_MAG
GCA_903937405.1_freshwater_MAG
GCA_903937445.1_freshwater_MAG
GCA_903959055.1_freshwater_MAG
GCA_903937015.1_freshwater_MAG
GCA_903952565.1_freshwater_MAG
GCA_903906295.1_freshwater_MAG
GCA_903841725.1_freshwater_MAG
GCA_903872175.1_freshwater_MAG
GCA_903824085.1_freshwater_MAG
GCA_903872405.1_freshwater_MAG
GCA_903830455.1_freshwater_MAG

| | | | | | | | | | | | | | | | | | | | | | | | | | | | | | | | | | | | | | | | | | | | | |
|---|---|---|---|---|---|---|---|---|---|---|---|---|---|---|---|---|---|---|---|---|---|---|---|---|---|---|---|---|---|---|---|---|---|---|---|---|---|---|---|---|---|---|---|---|
| 100 | 45 | 43 | 45 | 43 | 44 | 42 | 44 | 42 | 41 | 41 | 39 | 38 | 38 | 41 | 42 | 44 | 44 | 42 | 43 | 43 | 44 | 45 | 43 | 44 | 45 | 44 | 43 | 44 | 44 | 45 | 44 | 43 | 44 | 44 | 43 | 43 | 42 | 42 | 42 | 42 | 43 | 43 | 43 | GCA_001897445.1_ASM189744v1 |
| 45 | 100 | 43 | 46 | 42 | 44 | 43 | 44 | 43 | 42 | 42 | 39 | 39 | 38 | 42 | 41 | 44 | 44 | 43 | 43 | 44 | 44 | 45 | 44 | 45 | 45 | 45 | 45 | 46 | 44 | 48 | 46 | 45 | 47 | 47 | 43 | 43 | 43 | 43 | 43 | 43 | 43 | 43 | 43 | GCA_002422875.1_ASM242287v1 |
| 43 | 43 | 100 | 43 | 42 | 41 | 41 | 44 | 42 | 40 | 40 | 40 | 37 | 37 | 40 | 40 | 43 | 43 | 42 | 42 | 43 | 42 | 43 | 42 | 42 | 42 | 42 | 42 | 42 | 42 | 42 | 43 | 41 | 42 | 43 | 42 | 42 | 42 | 42 | 42 | 42 | 42 | 42 | 42 | GCA_014132315.1_ASM1413231v1 |
| 45 | 46 | 43 | 100 | 48 | 44 | 43 | 44 | 43 | 41 | 41 | 40 | 38 | 38 | 45 | 45 | 44 | 44 | 43 | 44 | 44 | 44 | 44 | 43 | 44 | 44 | 44 | 44 | 44 | 45 | 45 | 45 | 43 | 44 | 45 | 47 | 47 | 47 | 47 | 47 | 47 | 47 | 47 | 47 | GCA_014116815.1_ASM1411681v1 |
| 43 | 42 | 42 | 48 | 100 | 40 | 40 | 41 | 40 | 39 | 39 | 38 | 36 | 36 | 42 | 42 | 42 | 42 | 41 | 41 | 41 | 42 | 42 | 41 | 41 | 42 | 42 | 41 | 42 | 42 | 42 | 42 | 40 | 43 | 42 | 44 | 44 | 44 | 44 | 44 | 45 | 45 | 44 | 44 | GCA_013288625.1_ASM1328862v1 |
| 44 | 44 | 41 | 44 | 40 | 100 | 47 | 42 | 40 | 40 | 40 | 38 | 37 | 37 | 40 | 40 | 42 | 42 | 41 | 41 | 42 | 42 | 43 | 42 | 42 | 44 | 43 | 43 | 43 | 43 | 43 | 43 | 42 | 43 | 44 | 42 | 42 | 42 | 42 | 42 | 42 | 42 | 42 | 42 | GCA_018062985.1_ASM1806298v1 |
| 42 | 43 | 41 | 43 | 40 | 47 | 100 | 42 | 41 | 41 | 41 | 38 | 38 | 37 | 40 | 40 | 41 | 42 | 41 | 41 | 42 | 42 | 42 | 42 | 43 | 43 | 42 | 42 | 43 | 42 | 43 | 43 | 42 | 43 | 44 | 42 | 43 | 42 | 42 | 42 | 42 | 43 | 43 | 43 | GCA_013214525.1_ASM1321452v1 |
| 44 | 44 | 44 | 44 | 41 | 42 | 42 | 100 | 50 | 41 | 40 | 40 | 39 | 39 | 40 | 40 | 46 | 46 | 44 | 46 | 46 | 43 | 44 | 42 | 43 | 43 | 43 | 43 | 43 | 43 | 43 | 44 | 42 | 43 | 44 | 43 | 43 | 42 | 42 | 42 | 42 | 43 | 43 | 43 | GCA_016780625.1_ASM1678062v1 |
| 42 | 43 | 42 | 43 | 40 | 40 | 41 | 50 | 100 | 40 | 39 | 39 | 38 | 37 | 40 | 40 | 45 | 45 | 44 | 45 | 45 | 42 | 42 | 41 | 42 | 42 | 42 | 42 | 42 | 42 | 42 | 42 | 41 | 42 | 42 | 42 | 42 | 42 | 42 | 42 | 42 | 42 | 42 | 42 | GCA_016778805.1_ASM1677880v1 |
| 41 | 42 | 40 | 41 | 39 | 40 | 41 | 41 | 40 | 100 | 41 | 38 | 37 | 36 | 39 | 39 | 41 | 41 | 39 | 40 | 40 | 45 | 46 | 45 | 45 | 45 | 45 | 45 | 45 | 43 | 46 | 46 | 44 | 46 | 46 | 41 | 41 | 41 | 41 | 41 | 41 | 41 | 41 | 41 | GCA_015657545.1_ASM1565754v1 |
| 41 | 42 | 40 | 41 | 39 | 40 | 41 | 40 | 39 | 41 | 100 | 38 | 37 | 37 | 40 | 40 | 40 | 41 | 38 | 39 | 39 | 43 | 44 | 43 | 44 | 44 | 43 | 44 | 43 | 43 | 43 | 43 | 42 | 44 | 44 | 40 | 40 | 40 | 40 | 40 | 40 | 40 | 40 | 40 | GCA_903946825.1_freshwater_MAG |
| 39 | 39 | 40 | 40 | 38 | 38 | 38 | 40 | 39 | 38 | 38 | 100 | 37 | 36 | 38 | 38 | 39 | 40 | 38 | 39 | 39 | 39 | 39 | 38 | 39 | 39 | 38 | 39 | 38 | 39 | 39 | 39 | 38 | 39 | 39 | 39 | 39 | 39 | 39 | 39 | 39 | 39 | 40 | 39 | GCA_903864455.1_freshwater_MAG |
| 38 | 39 | 37 | 38 | 36 | 37 | 38 | 39 | 38 | 37 | 37 | 37 | 100 | 56 | 37 | 37 | 38 | 38 | 36 | 38 | 38 | 38 | 38 | 37 | 38 | 38 | 38 | 38 | 38 | 37 | 38 | 38 | 36 | 38 | 38 | 39 | 39 | 39 | 39 | 39 | 39 | 39 | 39 | 39 | GCA_002395105.1_ASM239510v1 |
| 38 | 38 | 37 | 38 | 36 | 37 | 37 | 39 | 37 | 36 | 37 | 36 | 56 | 100 | 37 | 37 | 38 | 39 | 37 | 38 | 38 | 37 | 37 | 37 | 37 | 38 | 38 | 37 | 38 | 37 | 38 | 38 | 37 | 38 | 38 | 37 | 38 | 37 | 38 | 38 | 38 | 37 | 37 | 37 | GCA_002394665.1_ASM239466v1 |
| 41 | 42 | 40 | 45 | 42 | 40 | 40 | 40 | 40 | 39 | 40 | 38 | 37 | 37 | 100 | 99 | 40 | 41 | 40 | 40 | 40 | 41 | 42 | 40 | 41 | 42 | 41 | 41 | 42 | 41 | 42 | 42 | 41 | 42 | 42 | 44 | 44 | 44 | 44 | 44 | 44 | 44 | 44 | 44 | GCA_903878005.1_freshwater_MAG |
| 42 | 41 | 40 | 45 | 42 | 40 | 40 | 40 | 40 | 39 | 40 | 38 | 37 | 37 | 99 | 100 | 40 | 41 | 40 | 40 | 41 | 41 | 42 | 40 | 41 | 42 | 41 | 41 | 41 | 41 | 42 | 42 | 41 | 42 | 42 | 44 | 44 | 44 | 44 | 44 | 44 | 44 | 44 | 44 | GCA_903860855.1_freshwater_MAG |
| 44 | 44 | 43 | 44 | 42 | 42 | 41 | 46 | 45 | 41 | 40 | 39 | 38 | 38 | 40 | 40 | 100 | 67 | 63 | 61 | 60 | 43 | 43 | 42 | 43 | 43 | 43 | 42 | 43 | 43 | 43 | 43 | 42 | 43 | 43 | 42 | 42 | 42 | 42 | 42 | 42 | 42 | 43 | 43 | GCA_002359655.1_ASM235965v1 |
| 44 | 44 | 43 | 44 | 42 | 42 | 42 | 46 | 45 | 41 | 41 | 40 | 38 | 39 | 41 | 41 | 67 | 100 | 63 | 62 | 61 | 43 | 44 | 42 | 43 | 43 | 43 | 43 | 43 | 43 | 43 | 43 | 42 | 43 | 43 | 43 | 43 | 43 | 43 | 43 | 43 | 43 | 43 | 43 | GCA_001768015.1_ASM176801v1 |
| 42 | 43 | 42 | 43 | 41 | 41 | 41 | 44 | 44 | 39 | 38 | 38 | 36 | 37 | 40 | 40 | 63 | 63 | 100 | 58 | 57 | 41 | 42 | 41 | 41 | 42 | 42 | 41 | 42 | 41 | 42 | 42 | 40 | 42 | 42 | 41 | 41 | 41 | 41 | 41 | 41 | 41 | 41 | 41 | GCA_009693885.1_ASM969388v1 |
| 43 | 43 | 42 | 44 | 41 | 41 | 41 | 46 | 45 | 40 | 39 | 39 | 38 | 38 | 40 | 40 | 61 | 62 | 58 | 100 | 57 | 42 | 42 | 41 | 42 | 43 | 42 | 42 | 42 | 42 | 42 | 43 | 42 | 43 | 43 | 42 | 42 | 42 | 42 | 42 | 42 | 42 | 42 | 42 | GCA_005787525.1_ASM578752v1 |
| 43 | 44 | 43 | 44 | 41 | 42 | 42 | 46 | 45 | 40 | 39 | 39 | 38 | 38 | 40 | 41 | 60 | 61 | 57 | 57 | 100 | 42 | 43 | 42 | 42 | 43 | 42 | 43 | 43 | 42 | 42 | 43 | 41 | 43 | 43 | 42 | 43 | 42 | 42 | 42 | 43 | 42 | 43 | 43 | GCA_016778745.1_ASM1677874v1 |
| 44 | 44 | 42 | 44 | 42 | 42 | 42 | 43 | 42 | 45 | 43 | 39 | 38 | 37 | 41 | 41 | 43 | 43 | 41 | 42 | 42 | 100 | 76 | 65 | 63 | 62 | 62 | 58 | 54 | 50 | 61 | 61 | 58 | 61 | 61 | 42 | 42 | 42 | 42 | 42 | 42 | 42 | 43 | 42 | GCA_905479885.1_10_20180426_B |
| 45 | 45 | 43 | 44 | 42 | 43 | 42 | 44 | 42 | 46 | 44 | 39 | 38 | 37 | 42 | 42 | 43 | 44 | 42 | 42 | 43 | 76 | 100 | 67 | 64 | 63 | 62 | 60 | 55 | 51 | 62 | 62 | 60 | 62 | 62 | 42 | 42 | 42 | 43 | 42 | 42 | 43 | 43 | 43 | GCA_905479865.1_10_20180508_B |
| 43 | 44 | 42 | 43 | 41 | 42 | 42 | 42 | 41 | 45 | 43 | 38 | 37 | 37 | 40 | 40 | 42 | 42 | 41 | 41 | 42 | 65 | 67 | 100 | 60 | 59 | 59 | 56 | 52 | 49 | 58 | 57 | 54 | 58 | 58 | 41 | 41 | 41 | 41 | 41 | 42 | 41 | 42 | 42 | GCA_002402195.1_ASM240219v1 |
| 44 | 45 | 42 | 44 | 41 | 42 | 43 | 43 | 42 | 45 | 44 | 39 | 38 | 37 | 41 | 41 | 43 | 43 | 41 | 42 | 42 | 63 | 64 | 60 | 100 | 62 | 61 | 56 | 53 | 50 | 58 | 58 | 56 | 59 | 58 | 42 | 42 | 42 | 42 | 42 | 42 | 42 | 43 | 42 | GCA_003963235.1_ASM396323v1 |
| 45 | 45 | 42 | 44 | 42 | 44 | 43 | 43 | 42 | 45 | 44 | 39 | 38 | 38 | 42 | 42 | 43 | 43 | 42 | 43 | 43 | 62 | 63 | 59 | 62 | 100 | 90 | 56 | 54 | 50 | 60 | 59 | 58 | 60 | 59 | 43 | 43 | 43 | 43 | 43 | 43 | 43 | 43 | 43 | GCA_017302595.1_ASM1730259v1 |
| 44 | 45 | 42 | 44 | 42 | 43 | 42 | 43 | 42 | 45 | 43 | 38 | 38 | 38 | 41 | 41 | 43 | 43 | 42 | 42 | 42 | 62 | 62 | 59 | 61 | 90 | 100 | 56 | 53 | 50 | 59 | 59 | 57 | 59 | 59 | 42 | 42 | 42 | 42 | 42 | 42 | 42 | 43 | 42 | GCA_013298405.1_ASM1329840v1 |
| 43 | 45 | 42 | 44 | 41 | 43 | 42 | 43 | 42 | 45 | 44 | 39 | 38 | 37 | 41 | 41 | 42 | 43 | 41 | 42 | 43 | 58 | 60 | 56 | 56 | 56 | 56 | 100 | 54 | 49 | 58 | 58 | 55 | 57 | 57 | 42 | 43 | 42 | 42 | 42 | 42 | 43 | 43 | 42 | GCA_013214485.1_ASM1321448v1 |
| 44 | 46 | 42 | 44 | 42 | 43 | 43 | 43 | 42 | 45 | 43 | 38 | 38 | 38 | 42 | 41 | 43 | 43 | 42 | 42 | 43 | 54 | 55 | 52 | 53 | 54 | 53 | 54 | 100 | 50 | 54 | 55 | 52 | 55 | 55 | 43 | 43 | 43 | 43 | 43 | 43 | 43 | 43 | 43 | GCA_903837685.1_freshwater_MAG |
| 44 | 44 | 42 | 45 | 42 | 43 | 42 | 43 | 42 | 43 | 43 | 39 | 37 | 37 | 41 | 41 | 43 | 43 | 41 | 42 | 42 | 50 | 51 | 49 | 50 | 50 | 50 | 49 | 50 | 100 | 50 | 50 | 48 | 51 | 50 | 42 | 42 | 42 | 42 | 42 | 42 | 42 | 43 | 42 | GCA_017995275.1_ASM1799527v1 |
| 45 | 48 | 42 | 45 | 42 | 43 | 43 | 43 | 42 | 46 | 43 | 39 | 38 | 38 | 42 | 42 | 43 | 43 | 42 | 42 | 42 | 61 | 62 | 58 | 58 | 60 | 59 | 58 | 54 | 50 | 100 | 95 | 90 | 88 | 87 | 43 | 43 | 43 | 43 | 43 | 43 | 43 | 43 | 43 | GCA_009927585.1_ASM992758v1 |
| 44 | 46 | 43 | 45 | 42 | 43 | 43 | 44 | 42 | 46 | 43 | 39 | 38 | 38 | 42 | 42 | 43 | 43 | 42 | 43 | 43 | 61 | 62 | 57 | 58 | 59 | 59 | 58 | 55 | 50 | 95 | 100 | 91 | 87 | 86 | 43 | 43 | 43 | 43 | 43 | 44 | 43 | 44 | 43 | GCA_018970295.1_ASM1897029v1 |
| 43 | 45 | 41 | 43 | 40 | 42 | 42 | 42 | 41 | 44 | 42 | 38 | 36 | 37 | 41 | 41 | 42 | 42 | 40 | 42 | 41 | 58 | 60 | 54 | 56 | 58 | 57 | 55 | 52 | 48 | 90 | 91 | 100 | 82 | 82 | 42 | 42 | 42 | 42 | 42 | 42 | 43 | 43 | 42 | GCA_018062005.1_ASM1806200v1 |
| 44 | 47 | 42 | 44 | 43 | 43 | 43 | 43 | 42 | 46 | 44 | 39 | 38 | 38 | 42 | 42 | 43 | 43 | 42 | 43 | 43 | 61 | 62 | 58 | 59 | 60 | 59 | 57 | 55 | 51 | 88 | 87 | 82 | 100 | 94 | 43 | 43 | 43 | 43 | 43 | 43 | 43 | 44 | 43 | GCA_017302665.1_ASM1730266v1 |
| 44 | 47 | 43 | 45 | 42 | 44 | 44 | 44 | 42 | 46 | 44 | 39 | 38 | 38 | 42 | 42 | 43 | 43 | 42 | 43 | 43 | 61 | 62 | 58 | 58 | 59 | 59 | 57 | 55 | 50 | 87 | 86 | 82 | 94 | 100 | 45 | 45 | 45 | 45 | 45 | 46 | 45 | 45 | 43 | GCA_903828265.1_freshwater_MAG |
| 43 | 43 | 42 | 47 | 44 | 42 | 42 | 43 | 42 | 41 | 40 | 39 | 39 | 37 | 44 | 44 | 42 | 43 | 41 | 42 | 42 | 42 | 42 | 41 | 42 | 43 | 42 | 42 | 43 | 42 | 43 | 43 | 42 | 43 | 45 | 100 | 100 | 100 | 100 | 100 | 100 | 100 | 100 | 79 | GCA_903887235.1_freshwater_MAG |
| 43 | 43 | 42 | 47 | 44 | 42 | 43 | 43 | 42 | 41 | 40 | 39 | 39 | 38 | 44 | 44 | 42 | 43 | 41 | 42 | 43 | 42 | 42 | 41 | 42 | 43 | 42 | 43 | 43 | 42 | 43 | 43 | 42 | 43 | 45 | 100 | 100 | 100 | 100 | 100 | 100 | 100 | 100 | 79 | GCA_903953855.1_freshwater_MAG |
| 42 | 43 | 42 | 47 | 44 | 42 | 42 | 42 | 42 | 41 | 40 | 39 | 39 | 37 | 44 | 44 | 42 | 43 | 41 | 42 | 42 | 42 | 42 | 41 | 42 | 43 | 42 | 42 | 43 | 42 | 43 | 43 | 42 | 43 | 45 | 100 | 100 | 100 | 100 | 100 | 100 | 100 | 99 | 79 | GCA_903957775.1_freshwater_MAG |
| 42 | 43 | 42 | 47 | 44 | 42 | 42 | 42 | 42 | 41 | 40 | 39 | 39 | 38 | 44 | 44 | 42 | 43 | 41 | 42 | 42 | 42 | 42 | 41 | 42 | 43 | 42 | 42 | 43 | 42 | 43 | 43 | 42 | 43 | 45 | 100 | 100 | 100 | 100 | 100 | 100 | 100 | 99 | 79 | GCA_903865125.1_freshwater_MAG |
| 42 | 43 | 42 | 47 | 44 | 42 | 42 | 42 | 42 | 41 | 40 | 39 | 39 | 38 | 44 | 44 | 42 | 43 | 41 | 42 | 42 | 42 | 43 | 41 | 42 | 43 | 42 | 42 | 43 | 42 | 43 | 43 | 42 | 43 | 45 | 100 | 100 | 100 | 100 | 100 | 100 | 100 | 99 | 80 | GCA_903903795.1_freshwater_MAG |
| 42 | 43 | 42 | 47 | 45 | 42 | 42 | 42 | 42 | 41 | 40 | 39 | 39 | 38 | 44 | 44 | 42 | 43 | 41 | 42 | 43 | 42 | 42 | 42 | 42 | 43 | 42 | 42 | 43 | 42 | 43 | 44 | 42 | 43 | 46 | 100 | 100 | 100 | 100 | 100 | 100 | 100 | 99 | 79 | GCA_903926815.1_freshwater_MAG |
| 43 | 43 | 42 | 47 | 45 | 42 | 43 | 43 | 42 | 41 | 40 | 39 | 39 | 37 | 44 | 44 | 42 | 43 | 41 | 42 | 42 | 42 | 43 | 41 | 42 | 43 | 42 | 43 | 43 | 42 | 43 | 43 | 43 | 43 | 45 | 100 | 100 | 100 | 100 | 100 | 100 | 100 | 100 | 80 | GCA_903820055.1_freshwater_MAG |
| 43 | 43 | 42 | 47 | 44 | 42 | 43 | 43 | 42 | 41 | 40 | 40 | 39 | 37 | 44 | 44 | 43 | 43 | 41 | 42 | 43 | 43 | 43 | 42 | 43 | 43 | 43 | 43 | 43 | 43 | 43 | 44 | 43 | 44 | 45 | 100 | 100 | 99 | 99 | 99 | 99 | 100 | 100 | 80 | GCA_903916125.1_freshwater_MAG |
| 43 | 43 | 42 | 47 | 44 | 42 | 43 | 43 | 42 | 41 | 40 | 39 | 39 | 37 | 44 | 44 | 43 | 43 | 41 | 42 | 43 | 42 | 43 | 42 | 42 | 43 | 42 | 42 | 43 | 42 | 43 | 43 | 42 | 43 | 43 | 79 | 79 | 79 | 79 | 80 | 79 | 80 | 80 | 100 | GCA_905479795.1_10_20180426_B |

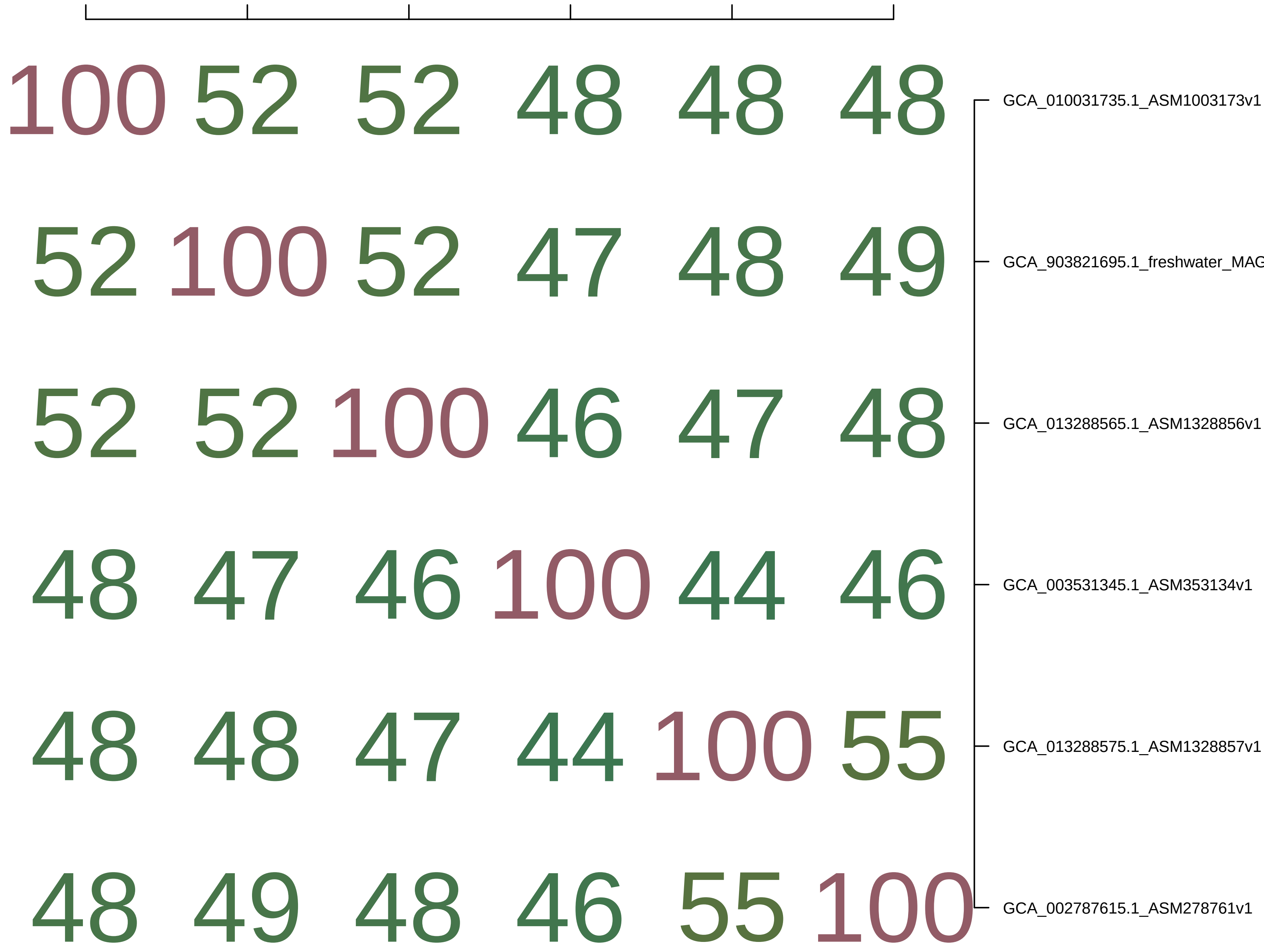

100 52 52 48 48 48
52 100 52 47 48 49
52 52 100 46 47 48
48 47 46 100 44 46
48 48 47 44 100 55
48 49 48 46 55 100
GCA_010031735.1_ASM1003173v1
GCA_903821695.1_freshwater_MAG
GCA_013288565.1_ASM1328856v1
GCA_003531345.1_ASM353134v1
GCA_013288575.1_ASM1328857v1
GCA_002787615.1_ASM278761v1

| GCA_005792635.1_ASM57926 | GCA_009926845.1_ASM99268 | GCA_013288305.1_ASM13288 | GCA_013288335.1_ASM13288 | GCA_013288985.1_ASM13288 | GCA_013289445.1_ASM13289 | GCA_002423155.1_ASM24231 | GCA_004293935.1_ASM42939 | GCA_002422205.1_ASM24222 | GCA_002725445.1_ASM27254 | GCA_002346425.1_ASM23464 | GCA_002422745.1_ASM24227 | GCA_003241645.1_ASM32416 | GCA_002291925.1_ASM22919 | GCA_002291845.1_ASM22918 | |
|---|---|---|---|---|---|---|---|---|---|---|---|---|---|---|---|
| 100 | 62 | 60 | 52 | 52 | 51 | 53 | 49 | 53 | 50 | 50 | 50 | 51 | 51 | 51 | GCA_005792635.1_ASM579263v1 |
| 62 | 100 | 60 | 52 | 52 | 51 | 52 | 49 | 53 | 50 | 49 | 50 | 52 | 52 | 52 | GCA_009926845.1_ASM992684v1 |
| 60 | 60 | 100 | 51 | 52 | 51 | 51 | 48 | 51 | 49 | 50 | 48 | 50 | 51 | 50 | GCA_013288305.1_ASM1328830v1 |
| 52 | 52 | 51 | 100 | 61 | 60 | 51 | 47 | 50 | 49 | 48 | 48 | 49 | 50 | 49 | GCA_013288335.1_ASM1328833v1 |
| 52 | 52 | 52 | 61 | 100 | 60 | 51 | 47 | 51 | 49 | 49 | 49 | 50 | 51 | 50 | GCA_013288985.1_ASM1328898v1 |
| 51 | 51 | 51 | 60 | 60 | 100 | 50 | 47 | 51 | 49 | 48 | 48 | 50 | 50 | 49 | GCA_013289445.1_ASM1328944v1 |
| 53 | 52 | 51 | 51 | 51 | 50 | 100 | 51 | 53 | 51 | 50 | 50 | 51 | 54 | 54 | GCA_002423155.1_ASM242315v1 |
| 49 | 49 | 48 | 47 | 47 | 47 | 51 | 100 | 50 | 49 | 49 | 47 | 49 | 52 | 51 | GCA_004293935.1_ASM429393v1 |
| 53 | 53 | 51 | 50 | 51 | 51 | 53 | 50 | 100 | 58 | 57 | 54 | 55 | 53 | 53 | GCA_002422205.1_ASM242220v1 |
| 50 | 50 | 49 | 49 | 49 | 49 | 51 | 49 | 58 | 100 | 55 | 53 | 54 | 52 | 51 | GCA_002725445.1_ASM272544v1 |
| 50 | 49 | 50 | 48 | 49 | 48 | 50 | 49 | 57 | 55 | 100 | 51 | 52 | 51 | 51 | GCA_002346425.1_ASM234642v1 |
| 50 | 50 | 48 | 48 | 49 | 48 | 50 | 47 | 54 | 53 | 51 | 100 | 62 | 51 | 51 | GCA_002422745.1_ASM242274v1 |
| 51 | 52 | 50 | 49 | 50 | 50 | 51 | 49 | 55 | 54 | 52 | 62 | 100 | 52 | 52 | GCA_003241645.1_ASM324164v1 |
| 51 | 52 | 51 | 50 | 51 | 50 | 54 | 52 | 53 | 52 | 51 | 51 | 52 | 100 | 74 | GCA_002291925.1_ASM229192v1 |
| 51 | 52 | 50 | 49 | 50 | 49 | 54 | 51 | 53 | 51 | 51 | 51 | 52 | 74 | 100 | GCA_002291845.1_ASM229184v1 |

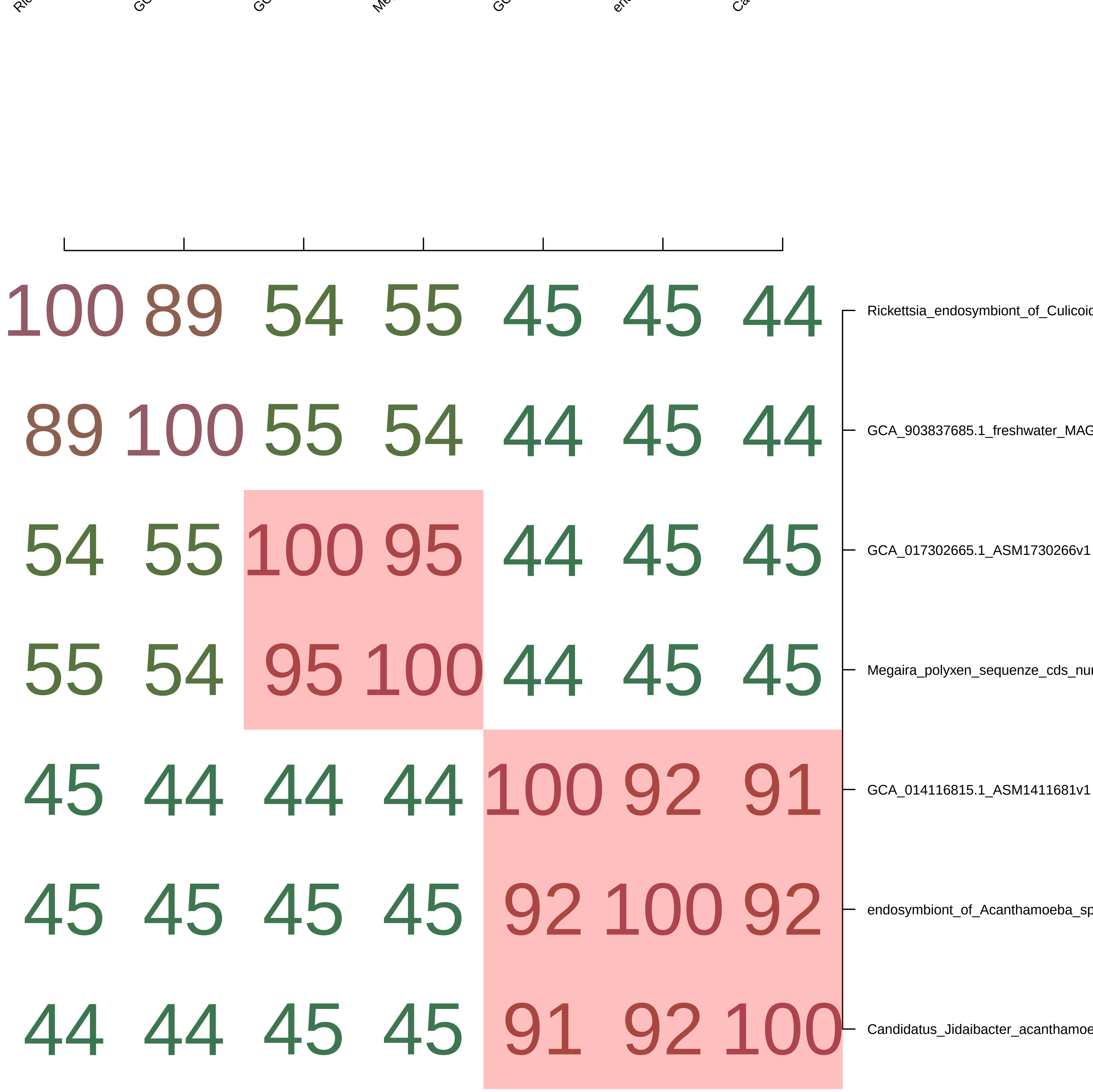

100 89 54 55 45 45 44
89 100 55 54 44 45 44
54 55 100 95 44 45 45
55 54 95 100 44 45 45
45 44 44 44 100 92 91
45 45 45 45 92 100 92
44 44 45 45 91 92 100
Rickettsia_endosymbiont_of_Culicoid
GCA_903837685.1_freshwater_MAG
GCA_017302665.1_ASM1730266v1
Megaira_polyxen_sequenze_cds_nu
GCA_014116815.1_ASM1411681v1
endosymbiont_of_Acanthamoeba_sp
Candidatus_Jidaibacter_acanthamoe
