## Supplementary material for "Host association and intracellularity evolved multiple times independently in the *Rickettsiales*": figures_and_supplementary_materials: Supplementary_figure_13_Phylogenomics_step5_trees_MAG_groups.pdf

### Group A

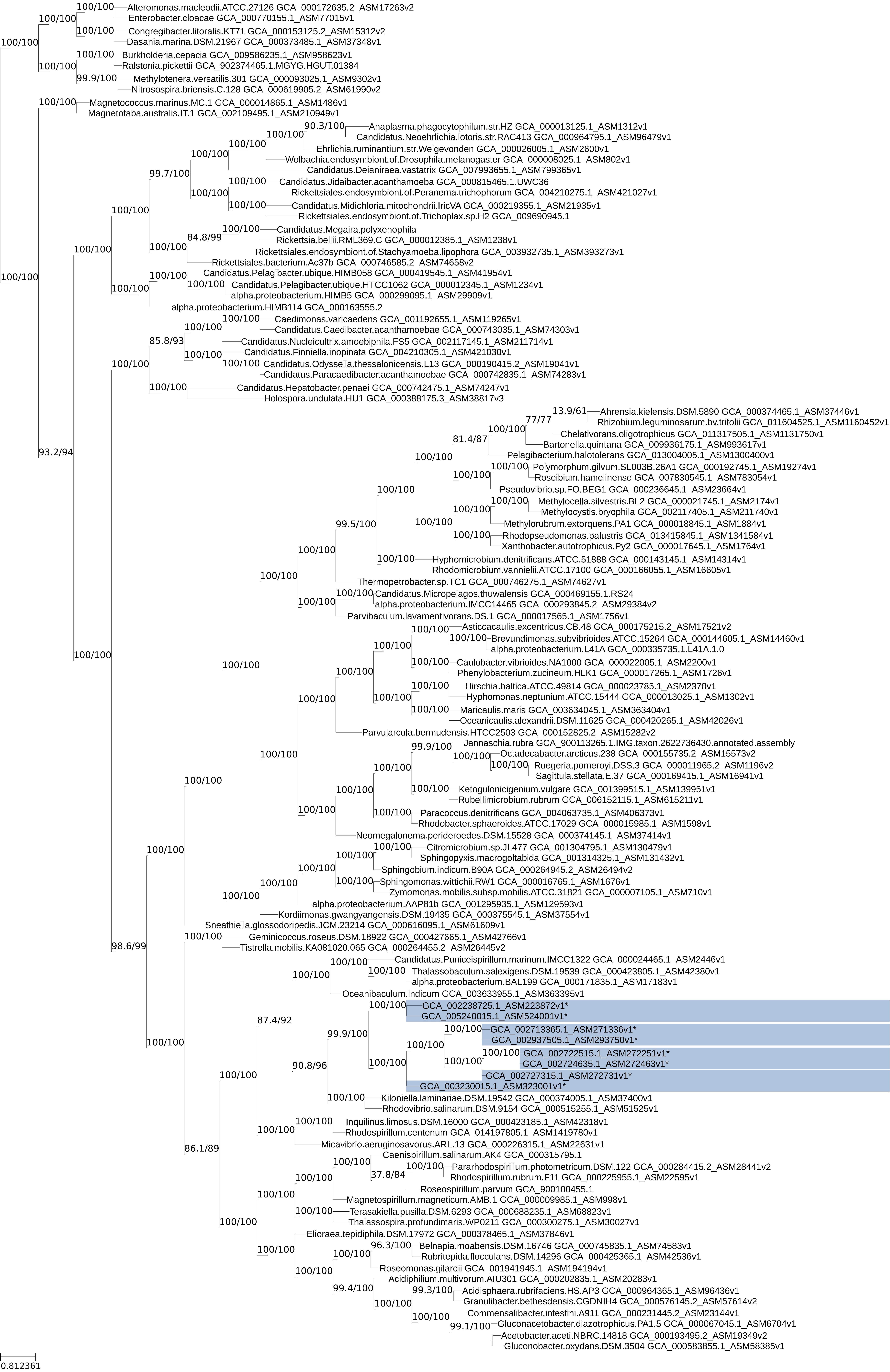

100/100
Alteromonas.macleodii.ATCC.27126 GCA_000172635.2_ASM17263v2
100/100
Enterobacter.cloacae GCA_000770155.1_ASM77015v1
100/100
100/100
Congregibacter.litoralis.KT71 GCA_000153125.2_ASM15312v2
Dasania.marina.DSM.21967 GCA_000373485.1_ASM37348v1
100/100
Burkholderia.cepacia GCA_009586235.1_ASM958623v1
100/100
Ralstonia.pickettii GCA_902374465.1.MGYG.HGUT.01384
99.9/100
Methylotenera.versatilis.301 GCA_000093025.1_ASM9302v1
Nitrosospira.briensis.C.128 GCA_000619905.2_ASM61990v2
100/100
Magnetococcus.marinus.MC.1 GCA_000014865.1_ASM1486v1
Magnetofaba.australis.IT.1 GCA_002109495.1_ASM210949v1
90.3/100
Anaplasma.phagocytophilum.str.HZ GCA_000013125.1_ASM1312v1
100/100
Candidatus.Neoehrlichia.lotoris.str.RAC413 GCA_000964795.1_ASM96479v1
100/100
Ehrlichia.ruminantium.str.Welgevonden GCA_000026005.1_ASM2600v1
100/100
Wolbachia.endosymbiont.of.Drosophila.melanogaster GCA_000008025.1_ASM802v1
99.7/100
Candidatus.Deianiraea.vastatrix GCA_007993655.1_ASM799365v1
100/100
Candidatus.Jidaibacter.acanthamoeba GCA_000815465.1.UWC36
100/100
Rickettsiales.endosymbiont.of.Peranema.trichophorum GCA_004210275.1_ASM421027v1
100/100
Candidatus.Midichloria.mitochondrii.IricVA GCA_000219355.1_ASM21935v1
100/100
Rickettsiales.endosymbiont.of.Trichoplax.sp.H2 GCA_009690945.1
100/100
Candidatus.Megaira.polyxenophila
84.8/99
Rickettsia.bellii.RML369.C GCA_000012385.1_ASM1238v1
100/100
100/100
Rickettsiales.endosymbiont.of.Stachyamoeba.lipophora GCA_003932735.1_ASM393273v1
Rickettsiales.bacterium.Ac37b GCA_000746585.2_ASM74658v2
100/100
100/100
Candidatus.Pelagibacter.ubique.HIMB058 GCA_000419545.1_ASM41954v1
100/100
Candidatus.Pelagibacter.ubique.HTCC1062 GCA_000012345.1_ASM1234v1
100/100
alpha.proteobacterium.HIMB5 GCA_000299095.1_ASM29909v1
alpha.proteobacterium.HIMB114 GCA_000163555.2
100/100
Caedimonas.varicaedens GCA_001192655.1_ASM119265v1
100/100
Candidatus.Caedibacter.acanthamoebae GCA_000743035.1_ASM74303v1
85.8/93
Candidatus.Nucleicultrix.amoebiphila.FS5 GCA_002117145.1_ASM211714v1
100/100
Candidatus.Finniella.inopinata GCA_004210305.1_ASM421030v1
100/100
100/100
Candidatus.Odyssella.thessalonicensis.L13 GCA_000190415.2_ASM19041v1
Candidatus.Paracaedibacter.acanthamoebae GCA_000742835.1_ASM74283v1
100/100
Candidatus.Hepatobacter.penaei GCA_000742475.1_ASM74247v1
Holospora.undulata.HU1 GCA_000388175.3_ASM38817v3
13.9/61
Ahrensia.kielensis.DSM.5890 GCA_000374465.1_ASM37446v1
77/77
Rhizobium.leguminosarum.bv.trifolii GCA_011604525.1_ASM1160452v1
100/100
Chelativorans.oligotrophicus GCA_011317505.1_ASM1131750v1
81.4/87
Bartonella.quintana GCA_009936175.1_ASM993617v1
93.2/94
Pelagibacterium.halotolerans GCA_013004005.1_ASM1300400v1
100/100
100/100
Polymorphum.gilvum.SL003B.26A1 GCA_000192745.1_ASM19274v1
100/100
Roseibium.hamelinense GCA_007830545.1_ASM783054v1
100/100
Pseudovibrio.sp.FO.BEG1 GCA_000236645.1_ASM23664v1
100/100
Methylocella.silvestris.BL2 GCA_000021745.1_ASM2174v1
100/100
Methylocystis.bryophila GCA_002117405.1_ASM211740v1
99.5/100
100/100
Methylorubrum.extorquens.PA1 GCA_000018845.1_ASM1884v1
100/100
Rhodopseudomonas.palustris GCA_013415845.1_ASM1341584v1
Xanthobacter.autotrophicus.Py2 GCA_000017645.1_ASM1764v1
100/100
100/100
Hyphomicrobium.denitrificans.ATCC.51888 GCA_000143145.1_ASM14314v1
Rhodomicrobium.vannielii.ATCC.17100 GCA_000166055.1_ASM16605v1
100/100
Thermopetrobacter.sp.TC1 GCA_000746275.1_ASM74627v1
100/100
Candidatus.Micropelagos.thuwalensis GCA_000469155.1.RS24
100/100
alpha.proteobacterium.IMCC14465 GCA_000293845.2_ASM29384v2
Parvibaculum.lavamentivorans.DS.1 GCA_000017565.1_ASM1756v1
100/100
Asticcacaulis.excentricus.CB.48 GCA_000175215.2_ASM17521v2
100/100
Brevundimonas.subvibrioides.ATCC.15264 GCA_000144605.1_ASM14460v1
100/100
alpha.proteobacterium.L41A GCA_000335735.1.L41A.1.0
100/100
100/100
Caulobacter.vibrioides.NA1000 GCA_000022005.1_ASM2200v1
100/100
Phenylobacterium.zucineum.HLK1 GCA_000017265.1_ASM1726v1
100/100
Hirschia.baltica.ATCC.49814 GCA_000023785.1_ASM2378v1
100/100
100/100
Hyphomonas.neptunium.ATCC.15444 GCA_000013025.1_ASM1302v1
100/100
Maricaulis.maris GCA_003634045.1_ASM363404v1
Oceanicaulis.alexandrii.DSM.11625 GCA_000420265.1_ASM42026v1
Parvularcula.bermudensis.HTCC2503 GCA_000152825.2_ASM15282v2
100/100
Jannaschia.rubra GCA_900113265.1.IMG.taxon.2622736430.annotated.assembly
99.9/100
100/100
Octadecabacter.arcticus.238 GCA_000155735.2_ASM15573v2
100/100
Ruegeria.pomeroyi.DSS.3 GCA_000011965.2_ASM1196v2
100/100
Sagittula.stellata.E.37 GCA_000169415.1_ASM16941v1
100/100
100/100
Ketogulonicigenium.vulgare GCA_001399515.1_ASM139951v1
Rubellimicrobium.rubrum GCA_006152115.1_ASM615211v1
100/100
Paracoccus.denitrificans GCA_004063735.1_ASM406373v1
100/100
Rhodobacter.sphaeroides.ATCC.17029 GCA_000015985.1_ASM1598v1
Neomegalonema.perideroedes.DSM.15528 GCA_000374145.1_ASM37414v1
100/100
100/100
Citromicrobium.sp.JL477 GCA_001304795.1_ASM130479v1
100/100
Sphingopyxis.macrogoltabida GCA_001314325.1_ASM131432v1
100/100
Sphingobium.indicum.B90A GCA_000264945.2_ASM26494v2
100/100
100/100
Sphingomonas.wittichii.RW1 GCA_000016765.1_ASM1676v1
100/100
Zymomonas.mobilis.subsp.mobilis.ATCC.31821 GCA_000007105.1_ASM710v1
alpha.proteobacterium.AAP81b GCA_001295935.1_ASM129593v1
Kordiimonas.gwangyangensis.DSM.19435 GCA_000375545.1_ASM37554v1
Sneathiella.glossodoripedis.JCM.23214 GCA_000616095.1_ASM61609v1
100/100
Geminicoccus.roseus.DSM.18922 GCA_000427665.1_ASM42766v1
98.6/99
Tistrella.mobilis.KA081020.065 GCA_000264455.2_ASM26445v2
100/100
Candidatus.Puniceispirillum.marinum.IMCC1322 GCA_000024465.1_ASM2446v1
100/100
100/100
Thalassobaculum.salexigens.DSM.19539 GCA_000423805.1_ASM42380v1
alpha.proteobacterium.BAL199 GCA_000171835.1_ASM17183v1
Oceanibaculum.indicum GCA_003633955.1_ASM363395v1
100/100
GCA_002238725.1_ASM223872v1*
87.4/92
GCA_005240015.1_ASM524001v1*
99.9/100
100/100
GCA_002713365.1_ASM271336v1*
100/100
GCA_002937505.1_ASM293750v1*
100/100
100/100
GCA_002722515.1_ASM272251v1*
90.8/96
100/100
100/100
GCA_002724635.1_ASM272463v1*
100/100
GCA_002727315.1_ASM272731v1*
GCA_003230015.1_ASM323001v1*
100/100
Kiloniella.laminariae.DSM.19542 GCA_000374005.1_ASM37400v1
Rhodovibrio.salinarum.DSM.9154 GCA_000515255.1_ASM51525v1
100/100
Inquilinus.limosus.DSM.16000 GCA_000423185.1_ASM42318v1
100/100
Rhodospirillum.centenum GCA_014197805.1_ASM1419780v1
86.1/89
Micavibrio.aeruginosavorus.ARL.13 GCA_000226315.1_ASM22631v1
Caenispirillum.salinarum.AK4 GCA_000315795.1
100/100
100/100
Pararhodospirillum.photometricum.DSM.122 GCA_000284415.2_ASM28441v2
37.8/84
100/100
Rhodospirillum.rubrum.F11 GCA_000225955.1_ASM22595v1
Roseospirillum.parvum GCA_900100455.1
100/100
Magnetospirillum.magneticum.AMB.1 GCA_000009985.1_ASM998v1
100/100
Terasakiella.pusilla.DSM.6293 GCA_000688235.1_ASM68823v1
100/100
Thalassospira.profundimaris.WP0211 GCA_000300275.1_ASM30027v1
Elioraea.tepidiphila.DSM.17972 GCA_000378465.1_ASM37846v1
96.3/100
Belnapia.moabensis.DSM.16746 GCA_000745835.1_ASM74583v1
100/100
100/100
Rubritepida.flocculans.DSM.14296 GCA_000425365.1_ASM42536v1
100/100
Roseomonas.gilardii GCA_001941945.1_ASM194194v1
Acidiphilium.multivorum.AIU301 GCA_000202835.1_ASM20283v1
99.4/100
99.3/100
Acidisphaera.rubrifaciens.HS.AP3 GCA_000964365.1_ASM96436v1
100/100
Granulibacter.bethesdensis.CGDNIH4 GCA_000576145.2_ASM57614v2
Commensalibacter.intestini.A911 GCA_000231445.2_ASM23144v1
100/100
99.1/100
Gluconacetobacter.diazotrophicus.PA1.5 GCA_000067045.1_ASM6704v1
Acetobacter.aceti.NBRC.14818 GCA_000193495.2_ASM19349v2
Gluconobacter.oxydans.DSM.3504 GCA_000583855.1_ASM58385v1
0.812361

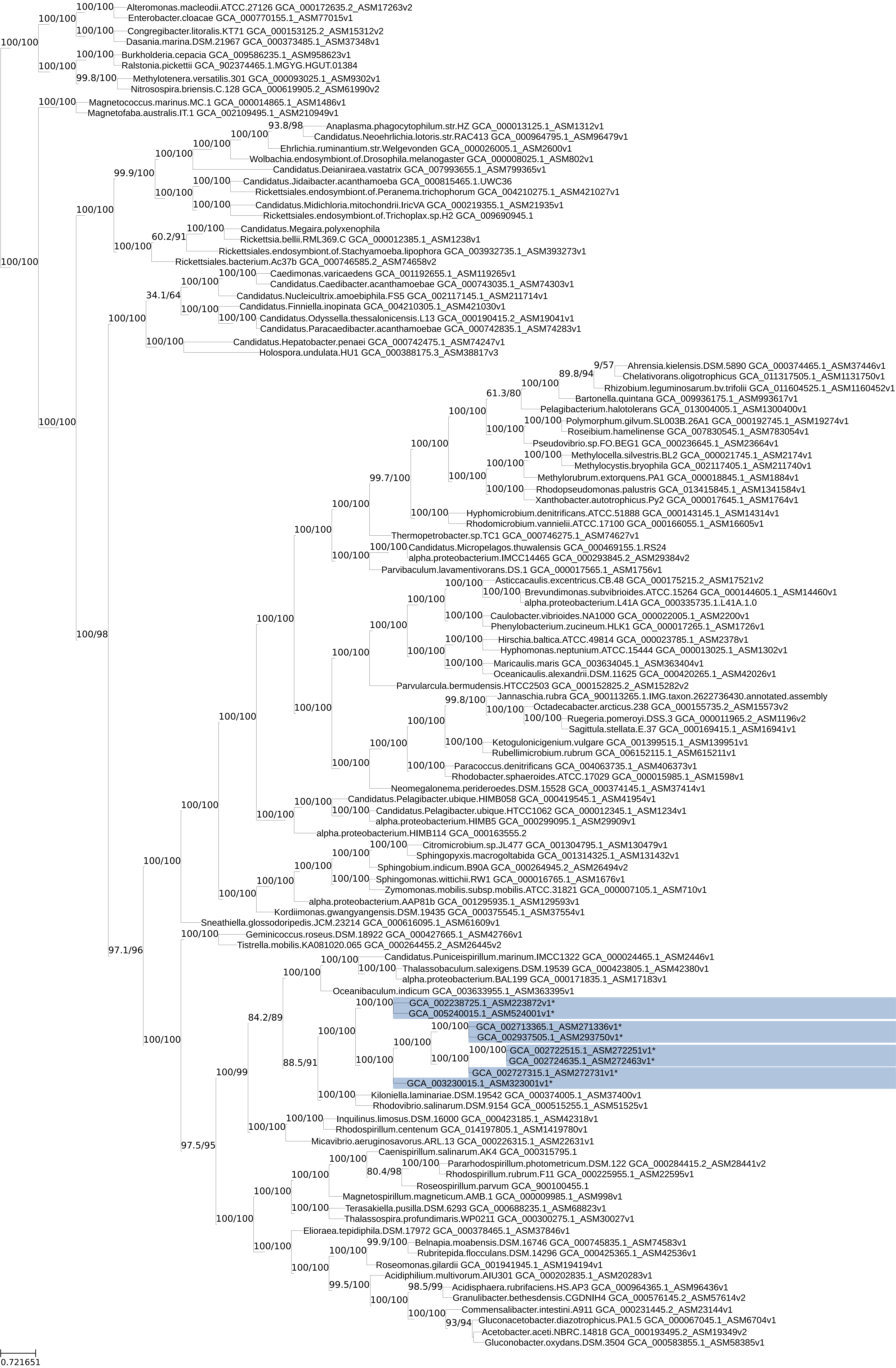

100/100 Alteromonas.macleodii.ATCC.27126 GCA_000172635.2_ASM17263v2
100/100 Enterobacter.cloacae GCA_000770155.1_ASM77015v1
100/100 100/100 Congregibacter.litoralis.KT71 GCA_000153125.2_ASM15312v2
Dasania.marina.DSM.21967 GCA_000373485.1_ASM37348v1
100/100 Burkholderia.cepacia GCA_009586235.1_ASM958623v1
100/100 Ralstonia.pickettii GCA_902374465.1.MGYG.HGUT.01384
99.8/100 Methylotenera.versatilis.301 GCA_000093025.1_ASM9302v1
Nitrosospira.briensis.C.128 GCA_000619905.2_ASM61990v2
100/100 Magnetococcus.marinus.MC.1 GCA_000014865.1_ASM1486v1
Magnetofaba.australis.IT.1 GCA_002109495.1_ASM210949v1
93.8/98 Anaplasma.phagocytophilum.str.HZ GCA_000013125.1_ASM1312v1
100/100 Candidatus.Neoehrlichia.lotoris.str.RAC413 GCA_000964795.1_ASM96479v1
100/100 Ehrlichia.ruminantium.str.Welgevonden GCA_000026005.1_ASM2600v1
100/100 Wolbachia.endosymbiont.of.Drosophila.melanogaster GCA_000008025.1_ASM802v1
99.9/100 Candidatus.Deianiraea.vastatrix GCA_007993655.1_ASM799365v1
100/100 Candidatus.Jidaibacter.acanthamoeba GCA_000815465.1.UWC36
100/100 Rickettsiales.endosymbiont.of.Peranema.trichophorum GCA_004210275.1_ASM421027v1
100/100 Candidatus.Midichloria.mitochondrii.IricVA GCA_000219355.1_ASM21935v1
100/100 Rickettsiales.endosymbiont.of.Trichoplax.sp.H2 GCA_009690945.1
100/100 Candidatus.Megaira.polyxenophila
60.2/91 Rickettsia.bellii.RML369.C GCA_000012385.1_ASM1238v1
100/100 Rickettsiales.endosymbiont.of.Stachyamoeba.lipophora GCA_003932735.1_ASM393273v1
100/100 Rickettsiales.bacterium.Ac37b GCA_000746585.2_ASM74658v2
100/100 Caedimonas.varicaedens GCA_001192655.1_ASM119265v1
100/100 Candidatus.Caedibacter.acanthamoebae GCA_000743035.1_ASM74303v1
34.1/64 Candidatus.Nucleicultrix.amoebiphila.FS5 GCA_002117145.1_ASM211714v1
100/100 Candidatus.Finniella.inopinata GCA_004210305.1_ASM421030v1
100/100 100/100 Candidatus.Odyssella.thessalonicensis.L13 GCA_000190415.2_ASM19041v1
Candidatus.Paracaedibacter.acanthamoebae GCA_000742835.1_ASM74283v1
100/100 Candidatus.Hepatobacter.penaei GCA_000742475.1_ASM74247v1
Holospora.undulata.HU1 GCA_000388175.3_ASM38817v3
9/57 Ahrensia.kielensis.DSM.5890 GCA_000374465.1_ASM37446v1
89.8/94 Chelativorans.oligotrophicus GCA_011317505.1_ASM1131750v1
Rhizobium.leguminosarum.bv.trifolii GCA_011604525.1_ASM1160452v1
100/100 Bartonella.quintana GCA_009936175.1_ASM993617v1
61.3/80 Pelagibacterium.halotolerans GCA_013004005.1_ASM1300400v1
100/100 100/100 Polymorphum.gilvum.SL003B.26A1 GCA_000192745.1_ASM19274v1
100/100 Roseibium.hamelinense GCA_007830545.1_ASM783054v1
100/100 Pseudovibrio.sp.FO.BEG1 GCA_000236645.1_ASM23664v1
100/100 Methylocella.silvestris.BL2 GCA_000021745.1_ASM2174v1
100/100 Methylocystis.bryophila GCA_002117405.1_ASM211740v1
100/100 Methylorubrum.extorquens.PA1 GCA_000018845.1_ASM1884v1
100/100 Rhodopseudomonas.palustris GCA_013415845.1_ASM1341584v1
99.7/100 Xanthobacter.autotrophicus.Py2 GCA_000017645.1_ASM1764v1
100/100 Hyphomicrobium.denitrificans.ATCC.51888 GCA_000143145.1_ASM14314v1
100/100 Rhodomicrobium.vannielii.ATCC.17100 GCA_000166055.1_ASM16605v1
100/100 Thermopetrobacter.sp.TC1 GCA_000746275.1_ASM74627v1
100/100 Candidatus.Micropelagos.thuwalensis GCA_000469155.1.RS24
100/100 alpha.proteobacterium.IMCC14465 GCA_000293845.2_ASM29384v2
Parvibaculum.lavamentivorans.DS.1 GCA_000017565.1_ASM1756v1
100/100 Asticcacaulis.excentricus.CB.48 GCA_000175215.2_ASM17521v2
100/100 Brevundimonas.subvibrioides.ATCC.15264 GCA_000144605.1_ASM14460v1
100/100 alpha.proteobacterium.L41A GCA_000335735.1.L41A.1.0
100/100 Caulobacter.vibrioides.NA1000 GCA_000022005.1_ASM2200v1
100/100 Phenylobacterium.zucineum.HLK1 GCA_000017265.1_ASM1726v1
100/100 Hirschia.baltica.ATCC.49814 GCA_000023785.1_ASM2378v1
100/100 Hyphomonas.neptunium.ATCC.15444 GCA_000013025.1_ASM1302v1
100/100 Maricaulis.maris GCA_003634045.1_ASM363404v1
100/100 Oceanicaulis.alexandrii.DSM.11625 GCA_000420265.1_ASM42026v1
100/100 Parvularcula.bermudensis.HTCC2503 GCA_000152825.2_ASM15282v2
99.8/100 Jannaschia.rubra GCA_900113265.1.IMG.taxon.2622736430.annotated.assembly
100/100 Octadecabacter.arcticus.238 GCA_000155735.2_ASM15573v2
100/100 100/100 Ruegeria.pomeroyi.DSS.3 GCA_000011965.2_ASM1196v2
Sagittula.stellata.E.37 GCA_000169415.1_ASM16941v1
100/100 100/100 Ketogulonicigenium.vulgare GCA_001399515.1_ASM139951v1
Rubellimicrobium.rubrum GCA_006152115.1_ASM615211v1
100/100 100/100 Paracoccus.denitrificans GCA_004063735.1_ASM406373v1
Rhodobacter.sphaeroides.ATCC.17029 GCA_000015985.1_ASM1598v1
100/100 Neomegalonema.perideroedes.DSM.15528 GCA_000374145.1_ASM37414v1
100/100 100/100 Candidatus.Pelagibacter.ubique.HIMB058 GCA_000419545.1_ASM41954v1
100/100 Candidatus.Pelagibacter.ubique.HTCC1062 GCA_000012345.1_ASM1234v1
100/100 alpha.proteobacterium.HIMB5 GCA_000299095.1_ASM29909v1
alpha.proteobacterium.HIMB114 GCA_000163555.2
100/100 Citromicrobium.sp.JL477 GCA_001304795.1_ASM130479v1
100/100 Sphingopyxis.macrogoltabida GCA_001314325.1_ASM131432v1
100/100 Sphingobium.indicum.B90A GCA_000264945.2_ASM26494v2
100/100 100/100 Sphingomonas.wittichii.RW1 GCA_000016765.1_ASM1676v1
100/100 Zymomonas.mobilis.subsp.mobilis.ATCC.31821 GCA_000007105.1_ASM710v1
100/100 alpha.proteobacterium.AAP81b GCA_001295935.1_ASM129593v1
Kordiimonas.gwangyangensis.DSM.19435 GCA_000375545.1_ASM37554v1
Sneathiella.glossodoripedis.JCM.23214 GCA_000616095.1_ASM61609v1
100/100 Geminicoccus.roseus.DSM.18922 GCA_000427665.1_ASM42766v1
Tistrella.mobilis.KA081020.065 GCA_000264455.2_ASM26445v2
100/100 Candidatus.Puniceispirillum.marinum.IMCC1322 GCA_000024445.1_ASM2446v1
100/100 100/100 Thalassobaculum.salexigens.DSM.19539 GCA_000423805.1_ASM42380v1
alpha.proteobacterium.BAL199 GCA_000171835.1_ASM17183v1
Oceanibaculum.indicum GCA_003633955.1_ASM363395v1
100/100 GCA_002238725.1_ASM223872v1*
GCA_005240015.1_ASM524001v1*
100/100 GCA_002713365.1_ASM271336v1*
100/100 GCA_002937505.1_ASM293750v1*
84.2/89 100/100 100/100 GCA_002722515.1_ASM272251v1*
100/100 GCA_002724635.1_ASM272463v1*
88.5/91 GCA_002727315.1_ASM272731v1*
GCA_003230015.1_ASM323001v1*
100/100 Kiloniella.laminariae.DSM.19542 GCA_000374005.1_ASM37400v1
Rhodovibrio.salinarum.DSM.9154 GCA_000515255.1_ASM51525v1
100/99 100/100 Inquilinus.limosus.DSM.16000 GCA_000423185.1_ASM42318v1
100/100 Rhodospirillum.centenum GCA_014197805.1_ASM1419780v1
Micavibrio.aeruginosavorus.ARL.13 GCA_000226315.1_ASM22631v1
97.5/95 100/100 Caenispirillum.salinarum.AK4 GCA_000315795.1
80.4/98 100/100 Pararhodospirillum.photometricum.DSM.122 GCA_000284415.2_ASM28441v2
100/100 Rhodospirillum.rubrum.F11 GCA_000225955.1_ASM22595v1
Roseospirillum.parvum GCA_900100455.1
100/100 Magnetospirillum.magneticum.AMB.1 GCA_000009985.1_ASM998v1
100/100 Terasakiella.pusilla.DSM.6293 GCA_000688235.1_ASM68823v1
Thalassospira.profundimaris.WP0211 GCA_000300275.1_ASM30027v1
100/100 Elioraea.tepidiphila.DSM.17972 GCA_000378465.1_ASM37846v1
100/100 99.9/100 Belnapia.moabensis.DSM.16746 GCA_000745835.1_ASM74583v1
100/100 Rubritepida.flocculans.DSM.14296 GCA_000425365.1_ASM42536v1
100/100 Roseomonas.gilardii GCA_001941945.1_ASM194194v1
99.5/100 Acidiphilium.multivorum.AIU301 GCA_000202835.1_ASM20283v1
98.5/99 Acidisphaera.rubrifaciens.HS.AP3 GCA_000964365.1_ASM96436v1
100/100 Granulibacter.bethesdensis.CGDNIH4 GCA_000576145.2_ASM57614v2
100/100 Commensalibacter.intestini.A911 GCA_000231445.2_ASM23144v1
93/94 Gluconacetobacter.diazotrophicus.PA1.5 GCA_000067045.1_ASM6704v1
Acetobacter.aceti.NBRC.14818 GCA_000193495.2_ASM19349v2
Gluconobacter.oxydans.DSM.3504 GCA_000583855.1_ASM58385v1
100/100
100/100
100/100
100/98
97.1/96
100/100
100/100
0.721651

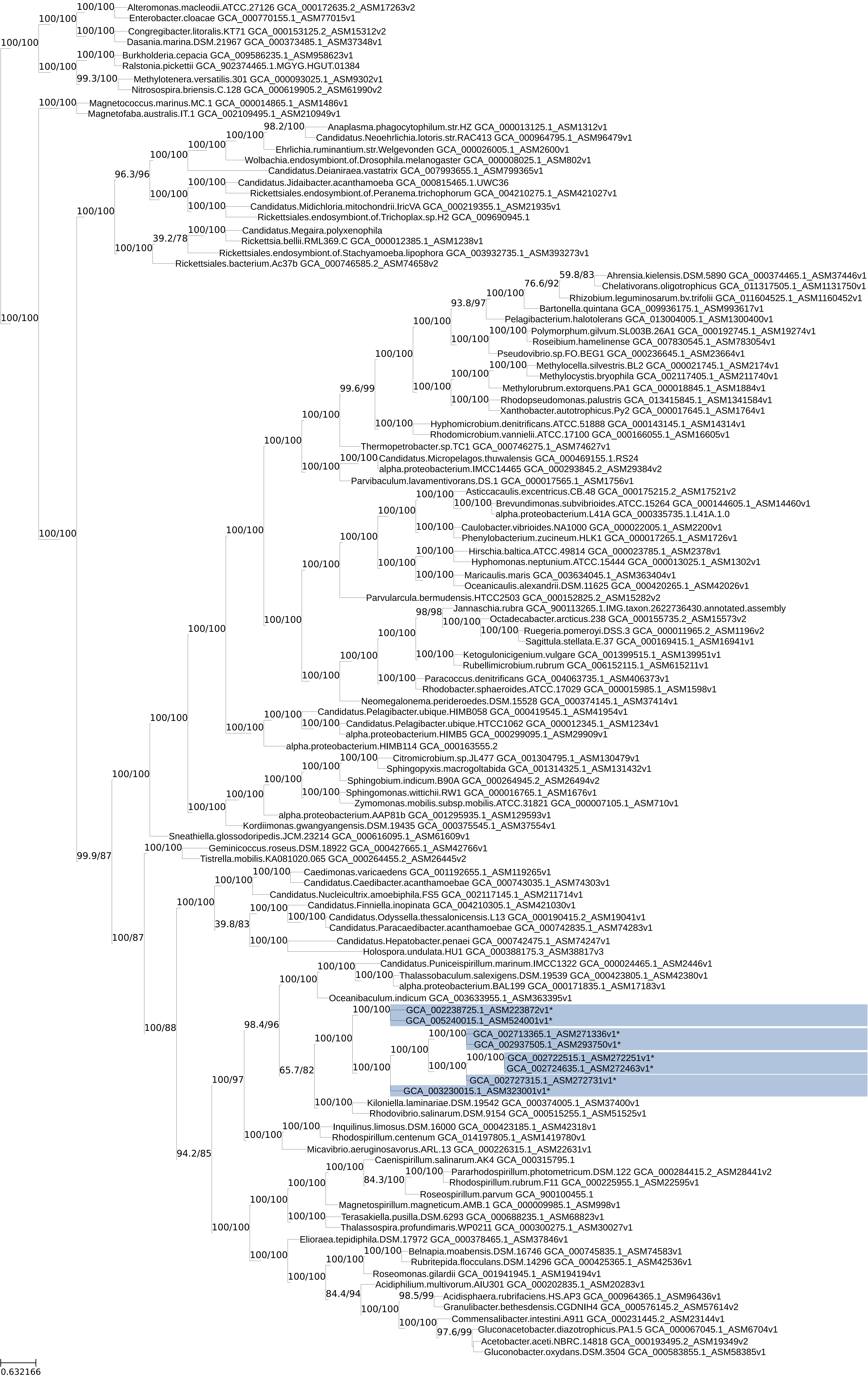

Alteromonas.macleodii.ATCC.27126 GCA_000172635.2_ASM17263v2
Enterobacter.cloacae GCA_000770155.1_ASM77015v1
Congregibacter.litoralis.KT71 GCA_000153125.2_ASM15312v2
Dasania.marina.DSM.21967 GCA_000373485.1_ASM37348v1
Burkholderia.cepacia GCA_009586235.1_ASM958623v1
Ralstonia.pickettii GCA_902374465.1.MGYG.HGUT.01384
Methylotenera.versatilis.301 GCA_000093025.1_ASM9302v1
Nitrosospira.briensis.C.128 GCA_000619905.2_ASM61990v2
Magnetococcus.marinus.MC.1 GCA_000014865.1_ASM1486v1
Magnetofaba.australis.IT.1 GCA_002109495.1_ASM210949v1
Anaplasma.phagocytophilum.str.HZ GCA_000013125.1_ASM1312v1
Candidatus.Neoehrlichia.lotoris.str.RAC413 GCA_000964795.1_ASM96479v1
Ehrlichia.ruminantium.str.Welgevonden GCA_000026005.1_ASM2600v1
Wolbachia.endosymbiont.of.Drosophila.melanogaster GCA_000008025.1_ASM802v1
Candidatus.Deianiraea.vastatrix GCA_007993655.1_ASM799365v1
Candidatus.Jidaibacter.acanthamoeba GCA_000815465.1.UWC36
Rickettsiales.endosymbiont.of.Peranema.trichophorum GCA_004210275.1_ASM421027v1
Candidatus.Midichloria.mitochondrii.IricVA GCA_000219355.1_ASM21935v1
Rickettsiales.endosymbiont.of.Trichoplax.sp.H2 GCA_009690945.1
Candidatus.Megaira.polyxenophila
Rickettsia.bellii.RML369.C GCA_000012385.1_ASM1238v1
Rickettsiales.endosymbiont.of.Stachyamoeba.lipophora GCA_003932735.1_ASM393273v1
Rickettsiales.bacterium.Ac37b GCA_000746585.2_ASM74658v2
Ahrensia.kielensis.DSM.5890 GCA_000374465.1_ASM37446v1
Chelativorans.oligotrophicus GCA_011317505.1_ASM1131750v1
Rhizobium.leguminosarum.bv.trifolii GCA_011604525.1_ASM1160452v1
Bartonella.quintana GCA_009936175.1_ASM993617v1
Pelagibacterium.halotolerans GCA_013004005.1_ASM1300400v1
Polymorphum.gilvum.SL003B.26A1 GCA_000192745.1_ASM19274v1
Roseibium.hamelinense GCA_007830545.1_ASM783054v1
Pseudovibrio.sp.FO.BEG1 GCA_000236645.1_ASM23664v1
Methylocella.silvestris.BL2 GCA_000021745.1_ASM2174v1
Methylocystis.bryophila GCA_002117405.1_ASM211740v1
Methylorubrum.extorquens.PA1 GCA_000018845.1_ASM1884v1
Rhodopseudomonas.palustris GCA_013415845.1_ASM1341584v1
Xanthobacter.autotrophicus.Py2 GCA_000017645.1_ASM1764v1
Hyphomicrobium.denitrificans.ATCC.51888 GCA_000143145.1_ASM14314v1
Rhodomicrobium.vannielii.ATCC.17100 GCA_000166055.1_ASM16605v1
Thermopetrobacter.sp.TC1 GCA_000746275.1_ASM74627v1
Candidatus.Micropelagos.thuwalensis GCA_000469155.1.RS24
alpha.proteobacterium.IMCC14465 GCA_000293845.2_ASM29384v2
Parvibaculum.lavamentivorans.DS.1 GCA_000017565.1_ASM1756v1
Asticcacaulis.excentricus.CB.48 GCA_000175215.2_ASM17521v2
Brevundimonas.subvibrioides.ATCC.15264 GCA_000144605.1_ASM14460v1
alpha.proteobacterium.L41A GCA_000335735.1.L41A.1.0
Caulobacter.vibrioides.NA1000 GCA_000022005.1_ASM2200v1
Phenylobacterium.zucineum.HLK1 GCA_000017265.1_ASM1726v1
Hirschia.baltica.ATCC.49814 GCA_000023785.1_ASM2378v1
Hyphomonas.neptunium.ATCC.15444 GCA_000013025.1_ASM1302v1
Maricaulis.maris GCA_003634045.1_ASM363404v1
Oceanicaulis.alexandrii.DSM.11625 GCA_000420265.1_ASM42026v1
Parvularcula.bermudensis.HTCC2503 GCA_000152825.2_ASM15282v2
Jannaschia.rubra GCA_900113265.1.IMG.taxon.2622736430.annotated.assembly
Octadecabacter.arcticus.238 GCA_000155735.2_ASM15573v2
Ruegeria.pomeroyi.DSS.3 GCA_000011965.2_ASM1196v2
Sagittula.stellata.E.37 GCA_000169415.1_ASM16941v1
Ketogulonicigenium.vulgare GCA_001399515.1_ASM139951v1
Rubellimicrobium.rubrum GCA_006152115.1_ASM615211v1
Paracoccus.denitrificans GCA_004063735.1_ASM406373v1
Rhodobacter.sphaeroides.ATCC.17029 GCA_000015985.1_ASM1598v1
Neomegalonema.perideroedes.DSM.15528 GCA_000374145.1_ASM37414v1
Candidatus.Pelagibacter.ubique.HIMB058 GCA_000419545.1_ASM41954v1
Candidatus.Pelagibacter.ubique.HTCC1062 GCA_000012345.1_ASM1234v1
alpha.proteobacterium.HIMB5 GCA_000299095.1_ASM29909v1
alpha.proteobacterium.HIMB114 GCA_000163555.2
Citromicrobium.sp.JL477 GCA_001304795.1_ASM130479v1
Sphingopyxis.macrogoltabida GCA_001314325.1_ASM131432v1
Sphingobium.indicum.B90A GCA_000264945.2_ASM26494v2
Sphingomonas.wittichii.RW1 GCA_000016765.1_ASM1676v1
Zymomonas.mobilis.subsp.mobilis.ATCC.31821 GCA_000007105.1_ASM710v1
alpha.proteobacterium.AAP81b GCA_001295935.1_ASM129593v1
Kordiimonas.gwangyangensis.DSM.19435 GCA_000375545.1_ASM37554v1
Sneathiella.glossodoripedis.JCM.23214 GCA_000616095.1_ASM61609v1
Geminicoccus.roseus.DSM.18922 GCA_000427665.1_ASM42766v1
Tistrella.mobilis.KA081020.065 GCA_000264455.2_ASM26445v2
Caedimonas.varicaedens GCA_001192655.1_ASM119265v1
Candidatus.Caedibacter.acanthamoebae GCA_000743035.1_ASM74303v1
Candidatus.Nucleicultrix.amoebiphila.FS5 GCA_002117145.1_ASM211714v1
Candidatus.Finniella.inopinata GCA_004210305.1_ASM421030v1
Candidatus.Odyssella.thessalonicensis.L13 GCA_000190415.2_ASM19041v1
Candidatus.Paracaedibacter.acanthamoebae GCA_000742835.1_ASM74283v1
Candidatus.Hepatobacter.penaei GCA_000742475.1_ASM74247v1
Holospora.undulata.HU1 GCA_000388175.3_ASM38817v3
Candidatus.Puniceispirillum.marinum.IMCC1322 GCA_000024465.1_ASM2446v1
Thalassobaculum.salexigens.DSM.19539 GCA_000423805.1_ASM42380v1
alpha.proteobacterium.BAL199 GCA_000171835.1_ASM17183v1
Oceanibaculum.indicum GCA_003633955.1_ASM363395v1
GCA_002238725.1_ASM223872v1*
GCA_005240015.1_ASM524001v1*
GCA_002713365.1_ASM271336v1*
GCA_002937505.1_ASM293750v1*
GCA_002722515.1_ASM272251v1*
GCA_002724635.1_ASM272463v1*
GCA_002727315.1_ASM272731v1*
GCA_003230015.1_ASM323001v1*
Kiloniella.laminariae.DSM.19542 GCA_000374005.1_ASM37400v1
Rhodovibrio.salinarum.DSM.9154 GCA_000515255.1_ASM51525v1
Inquilinus.limosus.DSM.16000 GCA_000423185.1_ASM42318v1
Rhodospirillum.centenum GCA_014197805.1_ASM1419780v1
Micavibrio.aeruginosavorus.ARL.13 GCA_000226315.1_ASM22631v1
Caenispirillum.salinarum.AK4 GCA_000315795.1
Pararhodospirillum.photometricum.DSM.122 GCA_000284415.2_ASM28441v2
Rhodospirillum.rubrum.F11 GCA_000225955.1_ASM22595v1
Roseospirillum.parvum GCA_900100455.1
Magnetospirillum.magneticum.AMB.1 GCA_000009985.1_ASM998v1
Terasakiella.pusilla.DSM.6293 GCA_000688235.1_ASM68823v1
Thalassospira.profundimaris.WP0211 GCA_000300275.1_ASM30027v1
Elioraea.tepidiphila.DSM.17972 GCA_000378465.1_ASM37846v1
Belnapia.moabensis.DSM.16746 GCA_000745835.1_ASM74583v1
Rubritepida.flocculans.DSM.14296 GCA_000425365.1_ASM42536v1
Roseomonas.gilardii GCA_001941945.1_ASM194194v1
Acidiphilium.multivorum.AIU301 GCA_000202835.1_ASM20283v1
Acidisphaera.rubrifaciens.HS.AP3 GCA_000964365.1_ASM96436v1
Granulibacter.bethesdensis.CGDNIH4 GCA_000576145.2_ASM57614v2
Commensalibacter.intestini.A911 GCA_000231445.2_ASM23144v1
Gluconacetobacter.diazotrophicus.PA1.5 GCA_000067045.1_ASM6704v1
Acetobacter.aceti.NBRC.14818 GCA_000193495.2_ASM19349v2
Gluconobacter.oxydans.DSM.3504 GCA_000583855.1_ASM58385v1
0.632166

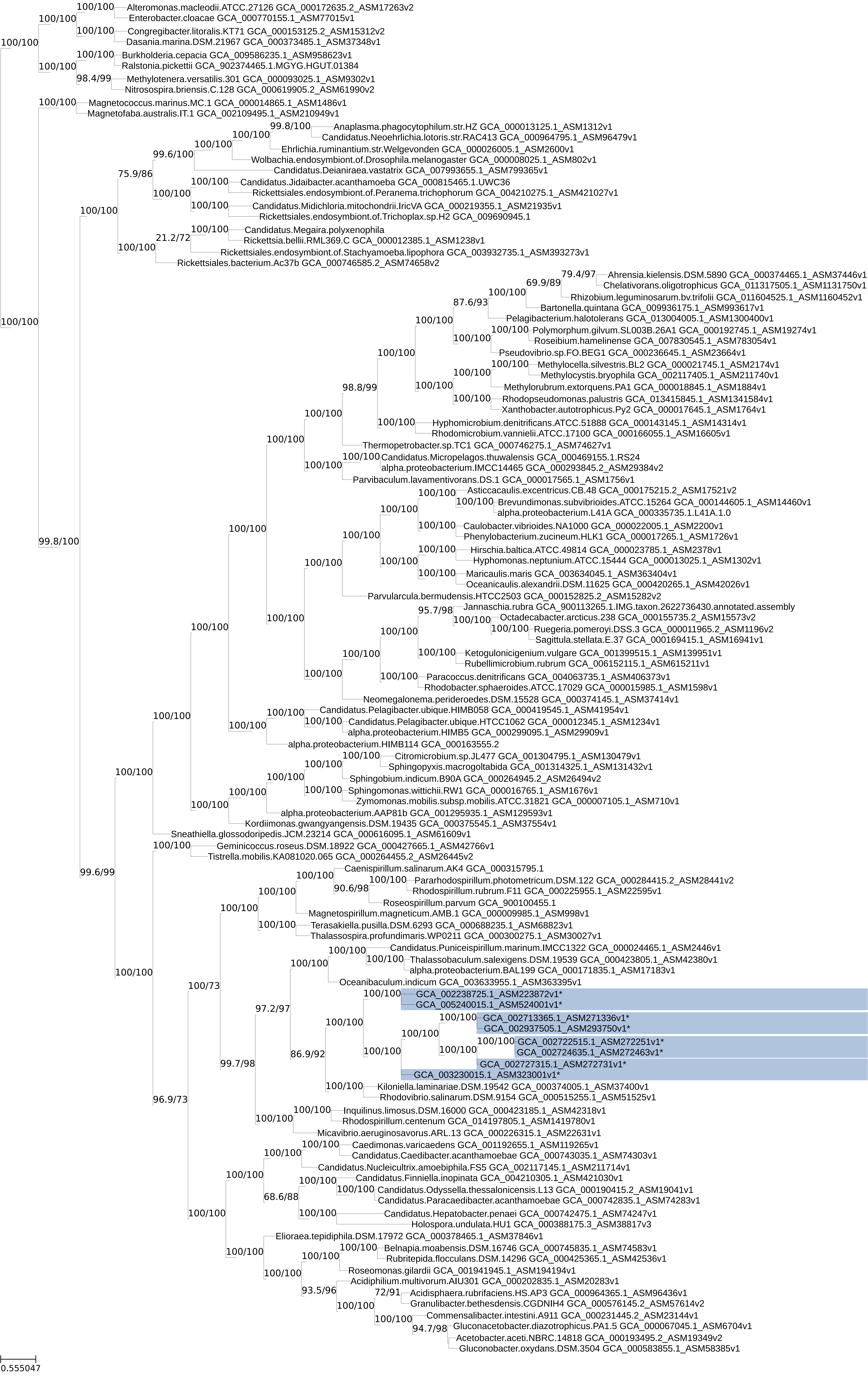
Alteromonas.macleodii.ATCC.27126 GCA_000172635.2_ASM17263v2
Enterobacter.cloacae GCA_000770155.1_ASM77015v1
Congregibacter.litoralis.KT71 GCA_000153125.2_ASM15312v2
Dasania.marina.DSM.21967 GCA_000373485.1_ASM37348v1
Burkholderia.cepacia GCA_009586235.1_ASM958623v1
Ralstonia.pickettii GCA_902374465.1.MGYG.HGUT.01384
Methylotenera.versatilis.301 GCA_000093025.1_ASM9302v1
Nitrosospira.briensis.C.128 GCA_000619905.2_ASM61990v2
Magnetococcus.marinus.MC.1 GCA_000014865.1_ASM1486v1
Magnetofaba.australis.IT.1 GCA_002109495.1_ASM210949v1
Anaplasma.phagocytophilum.str.HZ GCA_000013125.1_ASM1312v1
Candidatus.Neoehrlichia.lotoris.str.RAC413 GCA_000964795.1_ASM96479v1
Ehrlichia.ruminantium.str.Welgevonden GCA_000026005.1_ASM2600v1
Wolbachia.endosymbiont.of.Drosophila.melanogaster GCA_000008025.1_ASM802v1
Candidatus.Deianiraea.vastatrix GCA_007993655.1_ASM799365v1
Candidatus.Jidaibacter.acanthamoeba GCA_000815465.1.UWC36
Rickettsiales.endosymbiont.of.Peranema.trichophorum GCA_004210275.1_ASM421027v1
Candidatus.Midichloria.mitochondrii.IricVA GCA_000219355.1_ASM21935v1
Rickettsiales.endosymbiont.of.Trichoplax.sp.H2 GCA_009690945.1
Candidatus.Megaira.polyxenophila
Rickettsia.bellii.RML369.C GCA_000012385.1_ASM1238v1
Rickettsiales.endosymbiont.of.Stachyamoeba.lipophora GCA_003932735.1_ASM393273v1
Rickettsiales.bacterium.Ac37b GCA_000746585.2_ASM74658v2
Ahrensia.kielensis.DSM.5890 GCA_000374465.1_ASM37446v1
Chelativorans.oligotrophicus GCA_011317505.1_ASM1131750v1
Rhizobium.leguminosarum.bv.trifolii GCA_011604525.1_ASM1160452v1
Bartonella.quintana GCA_009936175.1_ASM993617v1
Pelagibacterium.halotolerans GCA_013004005.1_ASM1300400v1
Polymorphum.gilvum.SL003B.26A1 GCA_000192745.1_ASM19274v1
Roseibium.hamelinense GCA_007830545.1_ASM783054v1
Pseudovibrio.sp.FO.BEG1 GCA_000236645.1_ASM23664v1
Methylocella.silvestris.BL2 GCA_000021745.1_ASM2174v1
Methylocystis.bryophila GCA_002117405.1_ASM211740v1
Methylorubrum.extorquens.PA1 GCA_000018845.1_ASM1884v1
Rhodopseudomonas.palustris GCA_013415845.1_ASM1341584v1
Xanthobacter.autotrophicus.Py2 GCA_000017645.1_ASM1764v1
Hyphomicrobium.denitrificans.ATCC.51888 GCA_000143145.1_ASM14314v1
Rhodomicrobium.vannielii.ATCC.17100 GCA_000166055.1_ASM16605v1
Thermopetrobacter.sp.TC1 GCA_000746275.1_ASM74627v1
Candidatus.Micropelagos.thuwalensis GCA_000469155.1.RS24
alpha.proteobacterium.IMCC14465 GCA_000293845.2_ASM29384v2
Parvibaculum.lavamentivorans.DS.1 GCA_000017565.1_ASM1756v1
Asticcacaulis.excentricus.CB.48 GCA_000175215.2_ASM17521v2
Brevundimonas.subvibrioides.ATCC.15264 GCA_000144605.1_ASM14460v1
alpha.proteobacterium.L41A GCA_000335735.1.L41A.1.0
Caulobacter.vibrioides.NA1000 GCA_000022005.1_ASM2200v1
Phenylobacterium.zucineum.HLK1 GCA_000017265.1_ASM1726v1
Hirschia.baltica.ATCC.49814 GCA_000023785.1_ASM2378v1
Hyphomonas.neptunium.ATCC.15444 GCA_000013025.1_ASM1302v1
Maricaulis.maris GCA_003634045.1_ASM363404v1
Oceanicaulis.alexandrii.DSM.11625 GCA_000420265.1_ASM42026v1
Parvularcula.bermudensis.HTCC2503 GCA_000152825.2_ASM15282v2
Jannaschia.rubra GCA_900113265.1.IMG.taxon.2622736430.annotated.assembly
Octadecabacter.arcticus.238 GCA_000155735.2_ASM15573v2
Ruegeria.pomeroyi.DSS.3 GCA_000011965.2_ASM1196v2
Sagittula.stellata.E.37 GCA_000169415.1_ASM16941v1
Ketogulonicigenium.vulgare GCA_001399515.1_ASM139951v1
Rubellimicrobium.rubrum GCA_006152115.1_ASM615211v1
Paracoccus.denitrificans GCA_004063735.1_ASM406373v1
Rhodobacter.sphaeroides.ATCC.17029 GCA_000015985.1_ASM1598v1
Neomegalonema.perideroedes.DSM.15528 GCA_000374145.1_ASM37414v1
Candidatus.Pelagibacter.ubique.HIMB058 GCA_000419545.1_ASM41954v1
Candidatus.Pelagibacter.ubique.HTCC1062 GCA_000012345.1_ASM1234v1
alpha.proteobacterium.HIMB5 GCA_000299095.1_ASM29909v1
alpha.proteobacterium.HIMB114 GCA_000163555.2
Citromicrobium.sp.JL477 GCA_001304795.1_ASM130479v1
Sphingopyxis.macrogoltabida GCA_001314325.1_ASM131432v1
Sphingobium.indicum.B90A GCA_000264945.2_ASM26494v2
Sphingomonas.wittichii.RW1 GCA_000016765.1_ASM1676v1
Zymomonas.mobilis.subsp.mobilis.ATCC.31821 GCA_000007105.1_ASM710v1
alpha.proteobacterium.AAP81b GCA_001295935.1_ASM129593v1
Kordiimonas.gwangyangensis.DSM.19435 GCA_000375545.1_ASM37554v1
Sneathiella.glossodoripedis.JCM.23214 GCA_000616095.1_ASM61609v1
Geminicoccus.roseus.DSM.18922 GCA_000427665.1_ASM42766v1
Tistrella.mobilis.KA081020.065 GCA_000264455.2_ASM26445v2
Caenispirillum.salinarum.AK4 GCA_000315795.1
Pararhodospirillum.photometricum.DSM.122 GCA_000284415.2_ASM28441v2
Rhodospirillum.rubrum.F11 GCA_000225955.1_ASM22595v1
Roseospirillum.parvum GCA_900100455.1
Magnetospirillum.magneticum.AMB.1 GCA_000009985.1_ASM998v1
Terasakiella.pusilla.DSM.6293 GCA_000688235.1_ASM68823v1
Thalassospira.profundimaris.WP0211 GCA_000300275.1_ASM30027v1
Candidatus.Puniceispirillum.marinum.IMCC1322 GCA_000024465.1_ASM2446v1
Thalassobaculum.salexigens.DSM.19539 GCA_000423805.1_ASM42380v1
alpha.proteobacterium.BAL199 GCA_000171835.1_ASM17183v1
Oceanibaculum.indicum GCA_003633955.1_ASM363395v1
GCA_002238725.1_ASM223872v1*
GCA_005240015.1_ASM524001v1*
GCA_002713365.1_ASM271336v1*
GCA_002937505.1_ASM293750v1*
GCA_002722515.1_ASM272251v1*
GCA_002724635.1_ASM272463v1*
GCA_002727315.1_ASM272731v1*
GCA_003230015.1_ASM323001v1*
Kiloniella.laminariae.DSM.19542 GCA_000374005.1_ASM37400v1
Rhodovibrio.salinarum.DSM.9154 GCA_000515255.1_ASM51525v1
Inquilinus.limosus.DSM.16000 GCA_000423185.1_ASM42318v1
Rhodospirillum.centenum GCA_014197805.1_ASM1419780v1
Micavibrio.aeruginosavorus.ARL.13 GCA_000226315.1_ASM22631v1
Caedimonas.varicaedens GCA_001192655.1_ASM119265v1
Candidatus.Caedibacter.acanthamoebae GCA_000743035.1_ASM74303v1
Candidatus.Nucleicultrix.amoebiphila.FS5 GCA_002117145.1_ASM211714v1
Candidatus.Finniella.inopinata GCA_004210305.1_ASM421030v1
Candidatus.Odyssella.thessalonicensis.L13 GCA_000190415.2_ASM19041v1
Candidatus.Paracaedibacter.acanthamoebae GCA_000742835.1_ASM74283v1
Candidatus.Hepatobacter.penaei GCA_000742475.1_ASM74247v1
Holospora.undulata.HU1 GCA_000388175.3_ASM38817v3
Elioraea.tepidiphila.DSM.17972 GCA_000378465.1_ASM37846v1
Belnapia.moabensis.DSM.16746 GCA_000745835.1_ASM74583v1
Rubritepida.flocculans.DSM.14296 GCA_000425365.1_ASM42536v1
Roseomonas.gilardii GCA_001941945.1_ASM194194v1
Acidiphilium.multivorum.AIU301 GCA_000202835.1_ASM20283v1
Acidisphaera.rubrifaciens.HS.AP3 GCA_000964365.1_ASM96436v1
Granulibacter.bethesdensis.CGDNIH4 GCA_000576145.2_ASM57614v2
Commensalibacter.intestini.A911 GCA_000231445.2_ASM23144v1
Gluconacetobacter.diazotrophicus.PA1.5 GCA_000067045.1_ASM6704v1
Acetobacter.aceti.NBRC.14818 GCA_000193495.2_ASM19349v2
Gluconobacter.oxydans.DSM.3504 GCA_000583855.1_ASM58385v1
0.555047

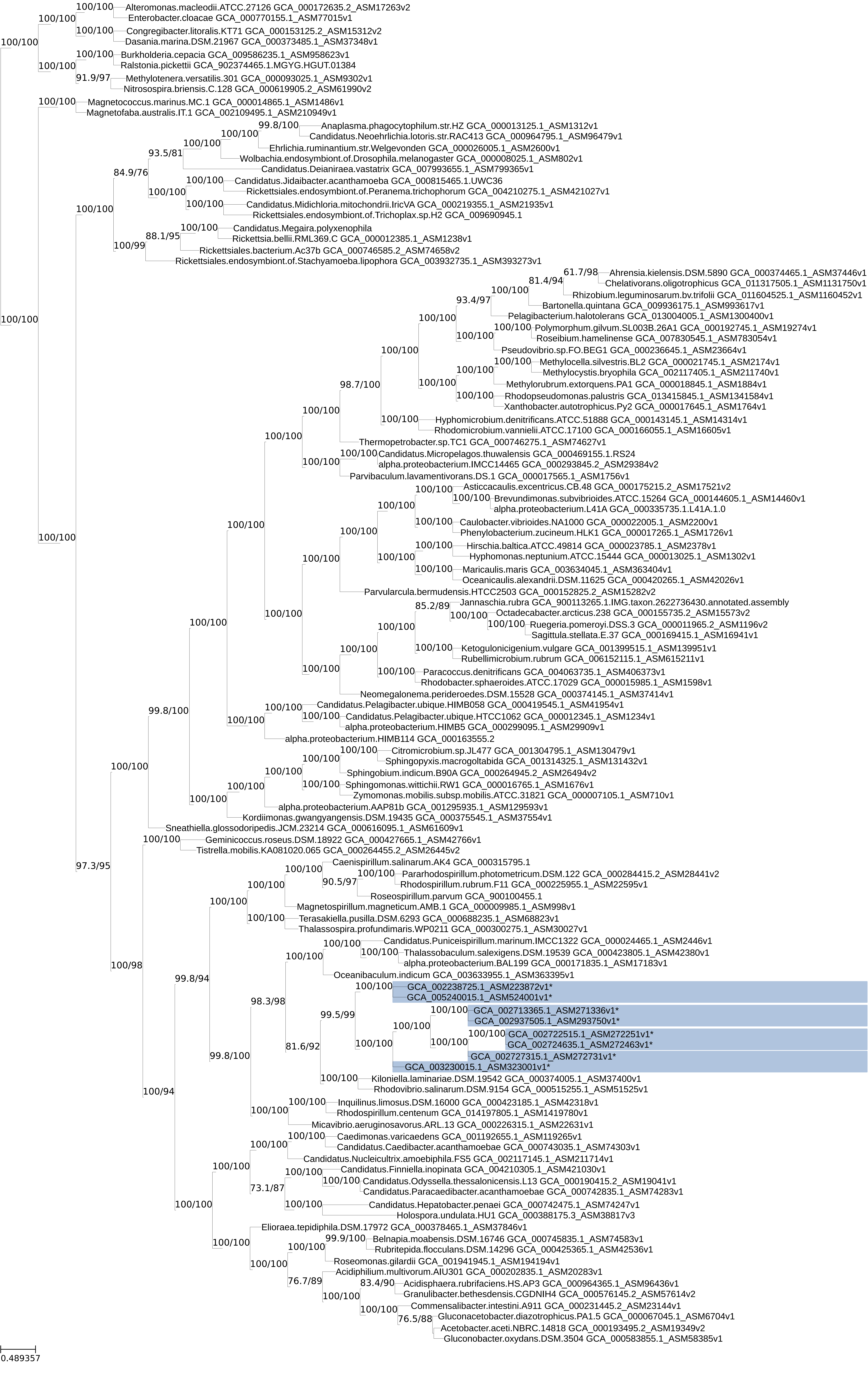
100/100
Alteromonas.macleodii.ATCC.27126 GCA_000172635.2_ASM17263v2
100/100
Enterobacter.cloacae GCA_000770155.1_ASM77015v1
100/100
Congregibacter.litoralis.KT71 GCA_000153125.2_ASM15312v2
100/100
Dasania.marina.DSM.21967 GCA_000373485.1_ASM37348v1
100/100
Burkholderia.cepacia GCA_009586235.1_ASM958623v1
100/100
Ralstonia.pickettii GCA_902374465.1.MGYG.HGUT.01384
91.9/97
Methylotenera.versatilis.301 GCA_000093025.1_ASM9302v1
Nitrosospira.briensis.C.128 GCA_000619905.2_ASM61990v2
100/100
Magnetococcus.marinus.MC.1 GCA_000014865.1_ASM1486v1
Magnetofaba.australis.IT.1 GCA_002109495.1_ASM210949v1
99.8/100
Anaplasma.phagocytophilum.str.HZ GCA_000013125.1_ASM1312v1
100/100
Candidatus.Neoehrlichia.lotoris.str.RAC413 GCA_000964795.1_ASM96479v1
100/100
Ehrlichia.ruminantium.str.Welgevonden GCA_000026005.1_ASM2600v1
93.5/81
Wolbachia.endosymbiont.of.Drosophila.melanogaster GCA_000008025.1_ASM802v1
Candidatus.Deianiraea.vastatrix GCA_007993655.1_ASM799365v1
84.9/76
100/100
Candidatus.Jidaibacter.acanthamoeba GCA_000815465.1.UWC36
100/100
Rickettsiales.endosymbiont.of.Peranema.trichophorum GCA_004210275.1_ASM421027v1
100/100
Candidatus.Midichloria.mitochondrii.IricVA GCA_000219355.1_ASM21935v1
100/100
Rickettsiales.endosymbiont.of.Trichoplax.sp.H2 GCA_009690945.1
100/100
Candidatus.Megaira.polyxenophila
88.1/95
Rickettsia.bellii.RML369.C GCA_000012385.1_ASM1238v1
100/99
Rickettsiales.bacterium.Ac37b GCA_000746585.2_ASM74658v2
Rickettsiales.endosymbiont.of.Stachyamoeba.lipophora GCA_003932735.1_ASM393273v1
100/100
61.7/98
Ahrensia.kielensis.DSM.5890 GCA_000374465.1_ASM37446v1
81.4/94
Chelativorans.oligotrophicus GCA_011317505.1_ASM1131750v1
100/100
Rhizobium.leguminosarum.bv.trifolii GCA_011604525.1_ASM1160452v1
93.4/97
Bartonella.quintana GCA_009936175.1_ASM993617v1
Pelagibacterium.halotolerans GCA_013004005.1_ASM1300400v1
100/100
100/100
Polymorphum.gilvum.SL003B.26A1 GCA_000192745.1_ASM19274v1
100/100
Roseibium.hamelinense GCA_007830545.1_ASM783054v1
Pseudovibrio.sp.FO.BEG1 GCA_000236645.1_ASM23664v1
100/100
100/100
Methylocella.silvestris.BL2 GCA_000021745.1_ASM2174v1
100/100
Methylocystis.bryophila GCA_002117405.1_ASM211740v1
98.7/100
100/100
Methylorubrum.extorquens.PA1 GCA_000018845.1_ASM1884v1
100/100
Rhodopseudomonas.palustris GCA_013415845.1_ASM1341584v1
Xanthobacter.autotrophicus.Py2 GCA_000017645.1_ASM1764v1
100/100
100/100
Hyphomicrobium.denitrificans.ATCC.51888 GCA_000143145.1_ASM14314v1
Rhodomicrobium.vannielii.ATCC.17100 GCA_000166055.1_ASM16605v1
100/100
Thermopetrobacter.sp.TC1 GCA_000746275.1_ASM74627v1
100/100
Candidatus.Micropelagos.thuwalensis GCA_000469155.1.RS24
100/100
alpha.proteobacterium.IMCC14465 GCA_000293845.2_ASM29384v2
Parvibaculum.lavamentivorans.DS.1 GCA_000017565.1_ASM1756v1
Asticcacaulis.excentricus.CB.48 GCA_000175215.2_ASM17521v2
100/100
100/100
Brevundimonas.subvibrioides.ATCC.15264 GCA_000144605.1_ASM14460v1
100/100
alpha.proteobacterium.L41A GCA_000335735.1.L41A.1.0
100/100
Caulobacter.vibrioides.NA1000 GCA_000022005.1_ASM2200v1
100/100
Phenylobacterium.zucineum.HLK1 GCA_000017265.1_ASM1726v1
100/100
Hirschia.baltica.ATCC.49814 GCA_000023785.1_ASM2378v1
100/100
100/100
Hyphomonas.neptunium.ATCC.15444 GCA_000013025.1_ASM1302v1
100/100
Maricaulis.maris GCA_003634045.1_ASM363404v1
Oceanicaulis.alexandrii.DSM.11625 GCA_000420265.1_ASM42026v1
Parvularcula.bermudensis.HTCC2503 GCA_000152825.2_ASM15282v2
100/100
85.2/89
Jannaschia.rubra GCA_900113265.1.IMG.taxon.2622736430.annotated.assembly
100/100
Octadecabacter.arcticus.238 GCA_000155735.2_ASM15573v2
100/100
Ruegeria.pomeroyi.DSS.3 GCA_000011965.2_ASM1196v2
100/100
Sagittula.stellata.E.37 GCA_000169415.1_ASM16941v1
100/100
Ketogulonicigenium.vulgare GCA_001399515.1_ASM139951v1
100/100
Rubellimicrobium.rubrum GCA_006152115.1_ASM615211v1
100/100
100/100
Paracoccus.denitrificans GCA_004063735.1_ASM406373v1
Rhodobacter.sphaeroides.ATCC.17029 GCA_000015985.1_ASM1598v1
Neomegalonema.perideroedes.DSM.15528 GCA_000374145.1_ASM37414v1
99.8/100
100/100
Candidatus.Pelagibacter.ubique.HIMB058 GCA_000419545.1_ASM41954v1
100/100
100/100
Candidatus.Pelagibacter.ubique.HTCC1062 GCA_000012345.1_ASM1234v1
alpha.proteobacterium.HIMB5 GCA_000299095.1_ASM29909v1
alpha.proteobacterium.HIMB114 GCA_000163555.2
100/100
100/100
Citromicrobium.sp.JL477 GCA_001304795.1_ASM130479v1
100/100
Sphingopyxis.macrogoltabida GCA_001314325.1_ASM131432v1
100/100
Sphingobium.indicum.B90A GCA_000264945.2_ASM26494v2
100/100
100/100
Sphingomonas.wittichii.RW1 GCA_000016765.1_ASM1676v1
100/100
Zymomonas.mobilis.subsp.mobilis.ATCC.31821 GCA_000007105.1_ASM710v1
alpha.proteobacterium.AAP81b GCA_001295935.1_ASM129593v1
Kordiimonas.gwangyangensis.DSM.19435 GCA_000375545.1_ASM37554v1
Sneathiella.glossodoripedis.JCM.23214 GCA_000616095.1_ASM61609v1
100/100
Geminicoccus.roseus.DSM.18922 GCA_000427665.1_ASM42766v1
Tistrella.mobilis.KA081020.065 GCA_000264455.2_ASM26445v2
100/100
100/100
Caenispirillum.salinarum.AK4 GCA_000315795.1
97.3/95
100/100
Pararhodospirillum.photometricum.DSM.122 GCA_000284415.2_ASM28441v2
90.5/97
Rhodospirillum.rubrum.F11 GCA_000225955.1_ASM22595v1
100/100
Roseospirillum.parvum GCA_900100455.1
Magnetospirillum.magneticum.AMB.1 GCA_000009985.1_ASM998v1
100/100
Terasakiella.pusilla.DSM.6293 GCA_000688235.1_ASM68823v1
Thalassospira.profundimaris.WP0211 GCA_000300275.1_ASM30027v1
100/100
Candidatus.Puniceispirillum.marinum.IMCC1322 GCA_000024465.1_ASM2446v1
100/100
100/100
Thalassobaculum.salexigens.DSM.19539 GCA_000423805.1_ASM42380v1
alpha.proteobacterium.BAL199 GCA_000171835.1_ASM17183v1
100/98
Oceanibaculum.indicum GCA_003633955.1_ASM363395v1
99.8/94
100/100
GCA_002238725.1_ASM223872v1*
98.3/98
GCA_005240015.1_ASM524001v1*
99.5/99
100/100
GCA_002713365.1_ASM271336v1*
100/100
GCA_002937505.1_ASM293750v1*
100/100
100/100
GCA_002722515.1_ASM272251v1*
100/100
GCA_002724635.1_ASM272463v1*
81.6/92
99.8/100
GCA_002727315.1_ASM272731v1*
GCA_003230015.1_ASM323001v1*
100/100
Kiloniella.laminariae.DSM.19542 GCA_000374005.1_ASM37400v1
Rhodovibrio.salinarum.DSM.9154 GCA_000515255.1_ASM51525v1
100/94
100/100
Inquilinus.limosus.DSM.16000 GCA_000423185.1_ASM42318v1
100/100
Rhodospirillum.centenum GCA_014197805.1_ASM1419780v1
Micavibrio.aeruginosavorus.ARL.13 GCA_000226315.1_ASM22631v1
100/100
Caedimonas.varicaedens GCA_001192655.1_ASM119265v1
100/100
Candidatus.Caedibacter.acanthamoebae GCA_000743035.1_ASM74303v1
100/100
Candidatus.Nucleicultrix.amoebiphila.FS5 GCA_002117145.1_ASM211714v1
100/100
Candidatus.Finniella.inopinata GCA_004210305.1_ASM421030v1
100/100
Candidatus.Odyssella.thessalonicensis.L13 GCA_000190415.2_ASM19041v1
73.1/87
Candidatus.Paracaedibacter.acanthamoebae GCA_000742835.1_ASM74283v1
100/100
100/100
Candidatus.Hepatobacter.penaei GCA_000742475.1_ASM74247v1
Holospora.undulata.HU1 GCA_000388175.3_ASM38817v3
Elioraea.tepidiphila.DSM.17972 GCA_000378465.1_ASM37846v1
99.9/100
Belnapia.moabensis.DSM.16746 GCA_000745835.1_ASM74583v1
100/100
100/100
Rubritepida.flocculans.DSM.14296 GCA_000425365.1_ASM42536v1
100/100
Roseomonas.gilardii GCA_001941945.1_ASM194194v1
Acidiphilium.multivorum.AIU301 GCA_000202835.1_ASM20283v1
76.7/89
83.4/90
Acidisphaera.rubrifaciens.HS.AP3 GCA_000964365.1_ASM96436v1
100/100
Granulibacter.bethesdensis.CGDNIH4 GCA_000576145.2_ASM57614v2
100/100
Commensalibacter.intestini.A911 GCA_000231445.2_ASM23144v1
76.5/88
Gluconacetobacter.diazotrophicus.PA1.5 GCA_000067045.1_ASM6704v1
Acetobacter.aceti.NBRC.14818 GCA_000193495.2_ASM19349v2
Gluconobacter.oxydans.DSM.3504 GCA_000583855.1_ASM58385v1
0.489357

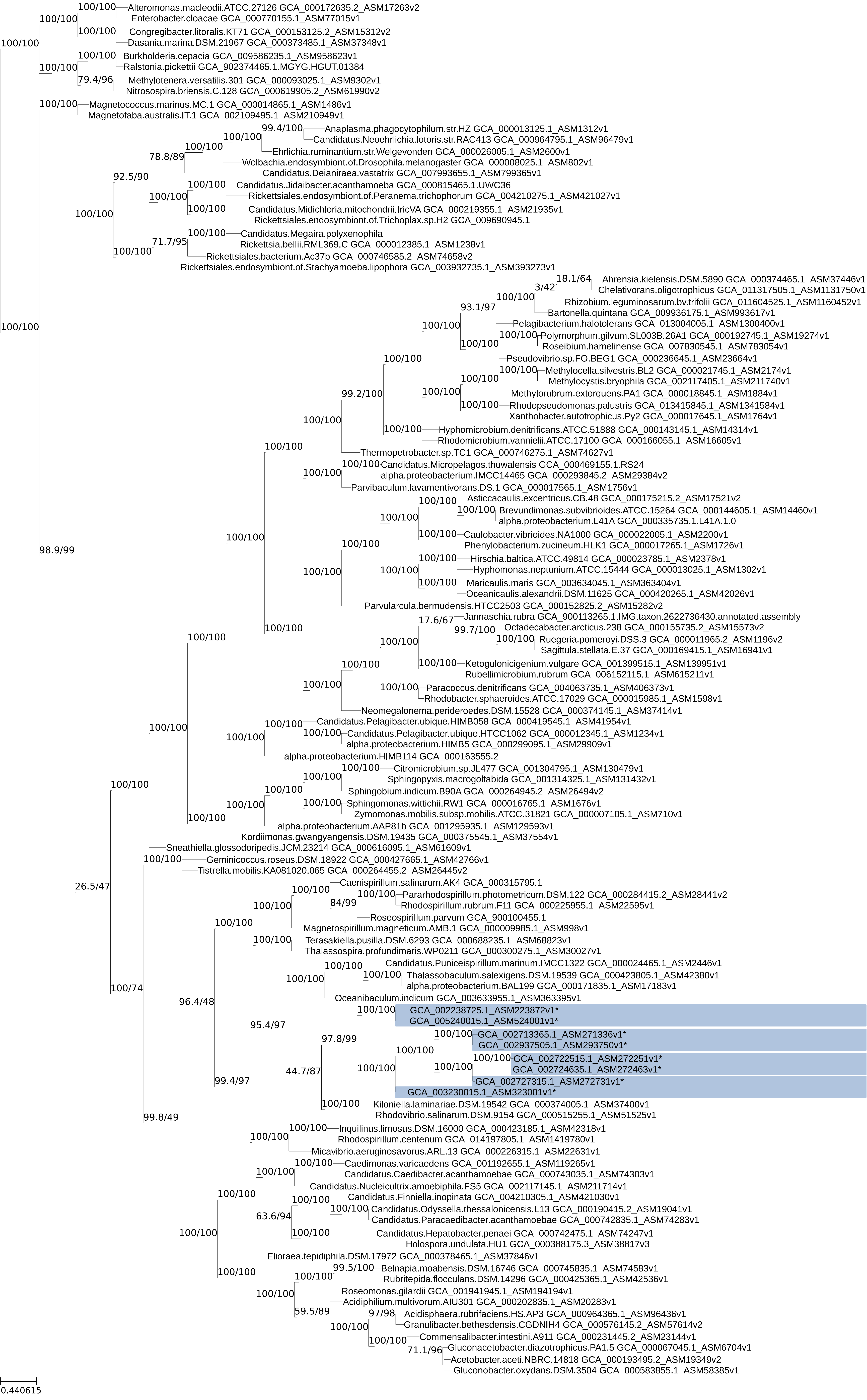
100/100
Alteromonas.macleodii.ATCC.27126 GCA_000172635.2_ASM17263v2
100/100
Enterobacter.cloacae GCA_000770155.1_ASM77015v1
100/100
100/100
Congregibacter.litoralis.KT71 GCA_000153125.2_ASM15312v2
Dasania.marina.DSM.21967 GCA_000373485.1_ASM37348v1
100/100
Burkholderia.cepacia GCA_009586235.1_ASM958623v1
100/100
Ralstonia.pickettii GCA_902374465.1.MGYG.HGUT.01384
79.4/96
Methylotenera.versatilis.301 GCA_000093025.1_ASM9302v1
Nitrosospira.briensis.C.128 GCA_000619905.2_ASM61990v2
100/100
Magnetococcus.marinus.MC.1 GCA_000014865.1_ASM1486v1
Magnetofaba.australis.IT.1 GCA_002109495.1_ASM210949v1
99.4/100
Anaplasma.phagocytophilum.str.HZ GCA_000013125.1_ASM1312v1
100/100
Candidatus.Neoehrlichia.lotoris.str.RAC413 GCA_000964795.1_ASM96479v1
100/100
Ehrlichia.ruminantium.str.Welgevonden GCA_000026005.1_ASM2600v1
78.8/89
Wolbachia.endosymbiont.of.Drosophila.melanogaster GCA_000008025.1_ASM802v1
Candidatus.Deianiraea.vastatrix GCA_007993655.1_ASM799365v1
92.5/90
100/100
Candidatus.Jidaibacter.acanthamoeba GCA_000815465.1.UWC36
100/100
Rickettsiales.endosymbiont.of.Peranema.trichophorum GCA_004210275.1_ASM421027v1
100/100
Candidatus.Midichloria.mitochondrii.IricVA GCA_000219355.1_ASM21935v1
100/100
Rickettsiales.endosymbiont.of.Trichoplax.sp.H2 GCA_009690945.1
100/100
Candidatus.Megaira.polyxenophila
71.7/95
Rickettsia.bellii.RML369.C GCA_000012385.1_ASM1238v1
100/100
Rickettsiales.bacterium.Ac37b GCA_000746585.2_ASM74658v2
Rickettsiales.endosymbiont.of.Stachyamoeba.lipophora GCA_003932735.1_ASM393273v1
18.1/64
Ahrensia.kielensis.DSM.5890 GCA_000374465.1_ASM37446v1
3/42
Chelativorans.oligotrophicus GCA_011317505.1_ASM1131750v1
100/100
Rhizobium.leguminosarum.bv.trifolii GCA_011604525.1_ASM1160452v1
93.1/97
Bartonella.quintana GCA_009936175.1_ASM993617v1
Pelagibacterium.halotolerans GCA_013004005.1_ASM1300400v1
100/100
100/100
Polymorphum.gilvum.SL003B.26A1 GCA_000192745.1_ASM19274v1
100/100
Roseibium.hamelinense GCA_007830545.1_ASM783054v1
Pseudovibrio.sp.FO.BEG1 GCA_000236645.1_ASM23664v1
100/100
100/100
Methylocella.silvestris.BL2 GCA_000021745.1_ASM2174v1
100/100
Methylocystis.bryophila GCA_002117405.1_ASM211740v1
100/100
Methylorubrum.extorquens.PA1 GCA_000018845.1_ASM1884v1
99.2/100
100/100
Rhodopseudomonas.palustris GCA_013415845.1_ASM1341584v1
Xanthobacter.autotrophicus.Py2 GCA_000017645.1_ASM1764v1
100/100
100/100
Hyphomicrobium.denitrificans.ATCC.51888 GCA_000143145.1_ASM14314v1
Rhodomicrobium.vannielii.ATCC.17100 GCA_000166055.1_ASM16605v1
100/100
Thermopetrobacter.sp.TC1 GCA_000746275.1_ASM74627v1
100/100
Candidatus.Micropelagos.thuwalensis GCA_000469155.1.RS24
100/100
alpha.proteobacterium.IMCC14465 GCA_000293845.2_ASM29384v2
Parvibaculum.lavamentivorans.DS.1 GCA_000017565.1_ASM1756v1
100/100
Asticcacaulis.excentricus.CB.48 GCA_000175215.2_ASM17521v2
100/100
Brevundimonas.subvibrioides.ATCC.15264 GCA_000144605.1_ASM14460v1
100/100
alpha.proteobacterium.L41A GCA_000335735.1.L41A.1.0
100/100
Caulobacter.vibrioides.NA1000 GCA_000022005.1_ASM2200v1
100/100
Phenylobacterium.zucineum.HLK1 GCA_000017265.1_ASM1726v1
100/100
Hirschia.baltica.ATCC.49814 GCA_000023785.1_ASM2378v1
100/100
Hyphomonas.neptunium.ATCC.15444 GCA_000013025.1_ASM1302v1
100/100
Maricaulis.maris GCA_003634045.1_ASM363404v1
100/100
Oceanicaulis.alexandrii.DSM.11625 GCA_000420265.1_ASM42026v1
Parvularcula.bermudensis.HTCC2503 GCA_000152825.2_ASM15282v2
17.6/67
Jannaschia.rubra GCA_900113265.1.IMG.taxon.2622736430.annotated.assembly
99.7/100
Octadecabacter.arcticus.238 GCA_000155735.2_ASM15573v2
100/100
Ruegeria.pomeroyi.DSS.3 GCA_000011965.2_ASM1196v2
100/100
100/100
Sagittula.stellata.E.37 GCA_000169415.1_ASM16941v1
100/100
Ketogulonicigenium.vulgare GCA_001399515.1_ASM139951v1
100/100
Rubellimicrobium.rubrum GCA_006152115.1_ASM615211v1
100/100
100/100
Paracoccus.denitrificans GCA_004063735.1_ASM406373v1
Rhodobacter.sphaeroides.ATCC.17029 GCA_000015985.1_ASM1598v1
Neomegalonema.perideroedes.DSM.15528 GCA_000374145.1_ASM37414v1
100/100
Candidatus.Pelagibacter.ubique.HIMB058 GCA_000419545.1_ASM41954v1
100/100
100/100
Candidatus.Pelagibacter.ubique.HTCC1062 GCA_000012345.1_ASM1234v1
alpha.proteobacterium.HIMB5 GCA_000299095.1_ASM29909v1
alpha.proteobacterium.HIMB114 GCA_000163555.2
100/100
Citromicrobium.sp.JL477 GCA_001304795.1_ASM130479v1
100/100
Sphingopyxis.macrogoltabida GCA_001314325.1_ASM131432v1
100/100
Sphingobium.indicum.B90A GCA_000264945.2_ASM26494v2
100/100
100/100
Sphingomonas.wittichii.RW1 GCA_000016765.1_ASM1676v1
100/100
Zymomonas.mobilis.subsp.mobilis.ATCC.31821 GCA_000007105.1_ASM710v1
alpha.proteobacterium.AAP81b GCA_001295935.1_ASM129593v1
Kordiimonas.gwangyangensis.DSM.19435 GCA_000375545.1_ASM37554v1
Sneathiella.glossodoripedis.JCM.23214 GCA_000616095.1_ASM61609v1
100/100
Geminicoccus.roseus.DSM.18922 GCA_000427665.1_ASM42766v1
Tistrella.mobilis.KA081020.065 GCA_000264455.2_ASM26445v2
26.5/47
Caenispirillum.salinarum.AK4 GCA_000315795.1
100/100
100/100
Pararhodospirillum.photometricum.DSM.122 GCA_000284415.2_ASM28441v2
84/99
Rhodospirillum.rubrum.F11 GCA_000225955.1_ASM22595v1
100/100
Roseospirillum.parvum GCA_900100455.1
100/100
Magnetospirillum.magneticum.AMB.1 GCA_000009985.1_ASM998v1
100/100
Terasakiella.pusilla.DSM.6293 GCA_000688235.1_ASM68823v1
Thalassospira.profundimaris.WP0211 GCA_000300275.1_ASM30027v1
100/100
Candidatus.Puniceispirillum.marinum.IMCC1322 GCA_000024465.1_ASM2446v1
100/100
Thalassobaculum.salexigens.DSM.19539 GCA_000423805.1_ASM42380v1
100/100
100/74
alpha.proteobacterium.BAL199 GCA_000171835.1_ASM17183v1
Oceanibaculum.indicum GCA_003633955.1_ASM363395v1
96.4/48
100/100
GCA_002238725.1_ASM223872v1*
GCA_005240015.1_ASM524001v1*
95.4/97
100/100
GCA_002713365.1_ASM271336v1*
97.8/99
GCA_002937505.1_ASM293750v1*
100/100
100/100
GCA_002722515.1_ASM272251v1*
100/100
GCA_002724635.1_ASM272463v1*
44.7/87
100/100
GCA_002727315.1_ASM272731v1*
99.4/97
GCA_003230015.1_ASM323001v1*
100/100
Kiloniella.laminariae.DSM.19542 GCA_000374005.1_ASM37400v1
99.8/49
Rhodovibrio.salinarum.DSM.9154 GCA_000515255.1_ASM51525v1
100/100
Inquilinus.limosus.DSM.16000 GCA_000423185.1_ASM42318v1
100/100
Rhodospirillum.centenum GCA_014197805.1_ASM1419780v1
Micavibrio.aeruginosavorus.ARL.13 GCA_000226315.1_ASM22631v1
100/100
Caedimonas.varicaedens GCA_001192655.1_ASM119265v1
100/100
Candidatus.Caedibacter.acanthamoebae GCA_000743035.1_ASM74303v1
Candidatus.Nucleicultrix.amoebiphila.FS5 GCA_002117145.1_ASM211714v1
100/100
100/100
Candidatus.Finniella.inopinata GCA_004210305.1_ASM421030v1
100/100
Candidatus.Odyssella.thessalonicensis.L13 GCA_000190415.2_ASM19041v1
63.6/94
Candidatus.Paracaedibacter.acanthamoebae GCA_000742835.1_ASM74283v1
100/100
100/100
Candidatus.Hepatobacter.penaei GCA_000742475.1_ASM74247v1
Holospora.undulata.HU1 GCA_000388175.3_ASM38817v3
Elioraea.tepidiphila.DSM.17972 GCA_000378465.1_ASM37846v1
99.5/100
Belnapia.moabensis.DSM.16746 GCA_000745835.1_ASM74583v1
100/100
100/100
Rubritepida.flocculans.DSM.14296 GCA_000425365.1_ASM42536v1
100/100
Roseomonas.gilardii GCA_001941945.1_ASM194194v1
Acidiphilium.multivorum.AIU301 GCA_000202835.1_ASM20283v1
59.5/89
97/98
Acidisphaera.rubrifaciens.HS.AP3 GCA_000964365.1_ASM96436v1
100/100
Granulibacter.bethesdensis.CGDNIH4 GCA_000576145.2_ASM57614v2
100/100
Commensalibacter.intestini.A911 GCA_000231445.2_ASM23144v1
71.1/96
Gluconacetobacter.diazotrophicus.PA1.5 GCA_000067045.1_ASM6704v1
Acetobacter.aceti.NBRC.14818 GCA_000193495.2_ASM19349v2
Gluconobacter.oxydans.DSM.3504 GCA_000583855.1_ASM58385v1
100/100
98.9/99
0.440615

### Group B

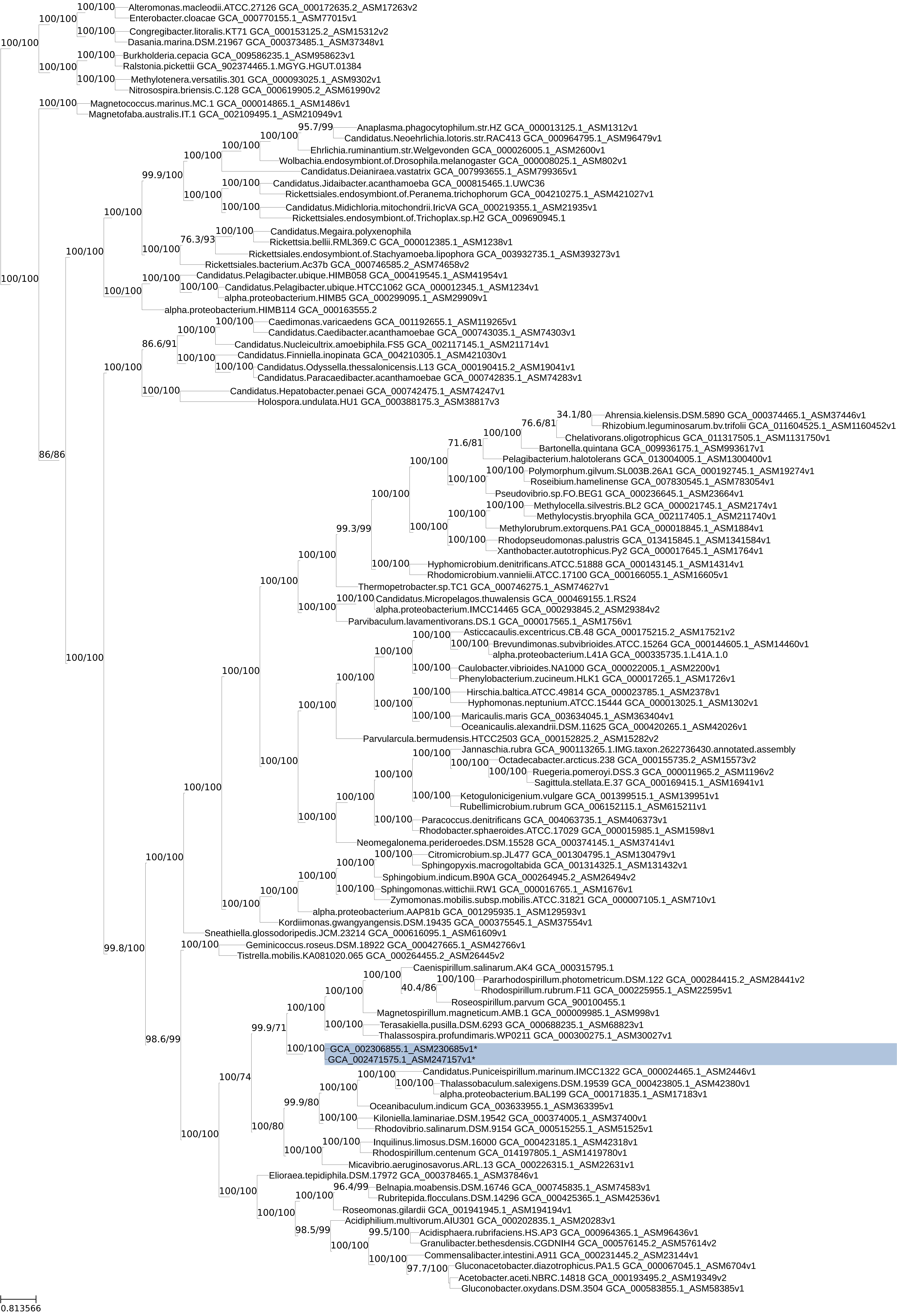
100/100
Alteromonas.macleodii.ATCC.27126 GCA_000172635.2_ASM17263v2
Enterobacter.cloacae GCA_000770155.1_ASM77015v1
100/100
100/100
Congregibacter.litoralis.KT71 GCA_000153125.2_ASM15312v2
Dasania.marina.DSM.21967 GCA_000373485.1_ASM37348v1
100/100
100/100
Burkholderia.cepacia GCA_009586235.1_ASM958623v1
Ralstonia.pickettii GCA_902374465.1.MGYG.HGUT.01384
100/100
100/100
Methylotenera.versatilis.301 GCA_000093025.1_ASM9302v1
Nitrosospira.briensis.C.128 GCA_000619905.2_ASM61990v2
100/100
Magnetococcus.marinus.MC.1 GCA_000014865.1_ASM1486v1
Magnetofaba.australis.IT.1 GCA_002109495.1_ASM210949v1
95.7/99
Anaplasma.phagocytophilum.str.HZ GCA_000013125.1_ASM1312v1
100/100
Candidatus.Neoehrlichia.lotoris.str.RAC413 GCA_000964795.1_ASM96479v1
100/100
Ehrlichia.ruminantium.str.Welgevonden GCA_000026005.1_ASM2600v1
100/100
Wolbachia.endosymbiont.of.Drosophila.melanogaster GCA_000008025.1_ASM802v1
99.9/100
Candidatus.Deianiraea.vastatrix GCA_007993655.1_ASM799365v1
100/100
Candidatus.Jidaibacter.acanthamoeba GCA_000815465.1.UWC36
100/100
Rickettsiales.endosymbiont.of.Peranema.trichophorum GCA_004210275.1_ASM421027v1
100/100
Candidatus.Midichloria.mitochondrii.IricVA GCA_000219355.1_ASM21935v1
100/100
Rickettsiales.endosymbiont.of.Trichoplax.sp.H2 GCA_009690945.1
100/100
Candidatus.Megaira.polyxenophila
76.3/93
Rickettsia.bellii.RML369.C GCA_000012385.1_ASM1238v1
100/100
Rickettsiales.endosymbiont.of.Stachyamoeba.lipophora GCA_003932735.1_ASM393273v1
100/100
Rickettsiales.bacterium.Ac37b GCA_000746585.2_ASM74658v2
100/100
Candidatus.Pelagibacter.ubique.HIMB058 GCA_000419545.1_ASM41954v1
100/100
Candidatus.Pelagibacter.ubique.HTCC1062 GCA_000012345.1_ASM1234v1
100/100
alpha.proteobacterium.HIMB5 GCA_000299095.1_ASM29909v1
alpha.proteobacterium.HIMB114 GCA_000163555.2
100/100
Caedimonas.varicaedens GCA_001192655.1_ASM119265v1
100/100
Candidatus.Caedibacter.acanthamoebae GCA_000743035.1_ASM74303v1
86.6/91
Candidatus.Nucleicultrix.amoebiphila.FS5 GCA_002117145.1_ASM211714v1
Candidatus.Finniella.inopinata GCA_004210305.1_ASM421030v1
100/100
100/100
Candidatus.Odyssella.thessalonicensis.L13 GCA_000190415.2_ASM19041v1
100/100
Candidatus.Paracaedibacter.acanthamoebae GCA_000742835.1_ASM74283v1
100/100
Candidatus.Hepatobacter.penaei GCA_000742475.1_ASM74247v1
Holospora.undulata.HU1 GCA_000388175.3_ASM38817v3
34.1/80
Ahrensia.kielensis.DSM.5890 GCA_000374465.1_ASM37446v1
76.6/81
Rhizobium.leguminosarum.bv.trifolii GCA_011604525.1_ASM1160452v1
100/100
Chelativorans.oligotrophicus GCA_011317505.1_ASM1131750v1
71.6/81
Bartonella.quintana GCA_009936175.1_ASM993617v1
86/86
Pelagibacterium.halotolerans GCA_013004005.1_ASM1300400v1
100/100
100/100
Polymorphum.gilvum.SL003B.26A1 GCA_000192745.1_ASM19274v1
100/100
Roseibium.hamelinense GCA_007830545.1_ASM783054v1
100/100
Pseudovibrio.sp.FO.BEG1 GCA_000236645.1_ASM23664v1
100/100
Methylocella.silvestris.BL2 GCA_000021745.1_ASM2174v1
100/100
Methylocystis.bryophila GCA_002117405.1_ASM211740v1
99.3/99
100/100
Methylorubrum.extorquens.PA1 GCA_000018845.1_ASM1884v1
100/100
Rhodopseudomonas.palustris GCA_013415845.1_ASM1341584v1
100/100
Xanthobacter.autotrophicus.Py2 GCA_000017645.1_ASM1764v1
100/100
Hyphomicrobium.denitrificans.ATCC.51888 GCA_000143145.1_ASM14314v1
100/100
Rhodomicrobium.vannielii.ATCC.17100 GCA_000166055.1_ASM16605v1
Thermopetrobacter.sp.TC1 GCA_000746275.1_ASM74627v1
100/100
Candidatus.Micropelagos.thuwalensis GCA_000469155.1.RS24
100/100
alpha.proteobacterium.IMCC14465 GCA_000293845.2_ASM29384v2
Parvibaculum.lavamentivorans.DS.1 GCA_000017565.1_ASM1756v1
100/100
Asticcacaulis.excentricus.CB.48 GCA_000175215.2_ASM17521v2
100/100
100/100
Brevundimonas.subvibrioides.ATCC.15264 GCA_000144605.1_ASM14460v1
alpha.proteobacterium.L41A GCA_000335735.1.L41A.1.0
100/100
100/100
Caulobacter.vibrioides.NA1000 GCA_000022005.1_ASM2200v1
100/100
Phenylobacterium.zucineum.HLK1 GCA_000017265.1_ASM1726v1
100/100
Hirschia.baltica.ATCC.49814 GCA_000023785.1_ASM2378v1
100/100
Hyphomonas.neptunium.ATCC.15444 GCA_000013025.1_ASM1302v1
100/100
Maricaulis.maris GCA_003634045.1_ASM363404v1
Oceanicaulis.alexandrii.DSM.11625 GCA_000420265.1_ASM42026v1
Parvularcula.bermudensis.HTCC2503 GCA_000152825.2_ASM15282v2
100/100
100/100
Jannaschia.rubra GCA_900113265.1.IMG.taxon.2622736430.annotated.assembly
100/100
Octadecabacter.arcticus.238 GCA_000155735.2_ASM15573v2
100/100
100/100
Ruegeria.pomeroyi.DSS.3 GCA_000011965.2_ASM1196v2
Sagittula.stellata.E.37 GCA_000169415.1_ASM16941v1
100/100
100/100
Ketogulonicigenium.vulgare GCA_001399515.1_ASM139951v1
100/100
Rubellimicrobium.rubrum GCA_006152115.1_ASM615211v1
100/100
Paracoccus.denitrificans GCA_004063735.1_ASM406373v1
100/100
Rhodobacter.sphaeroides.ATCC.17029 GCA_000015985.1_ASM1598v1
Neomegalonema.perideroedes.DSM.15528 GCA_000374145.1_ASM37414v1
100/100
100/100
Citromicrobium.sp.JL477 GCA_001304795.1_ASM130479v1
100/100
Sphingopyxis.macrogoltabida GCA_001314325.1_ASM131432v1
100/100
Sphingobium.indicum.B90A GCA_000264945.2_ASM26494v2
100/100
Sphingomonas.wittichii.RW1 GCA_000016765.1_ASM1676v1
100/100
Zymomonas.mobilis.subsp.mobilis.ATCC.31821 GCA_000007105.1_ASM710v1
100/100
alpha.proteobacterium.AAP81b GCA_001295935.1_ASM129593v1
Kordiimonas.gwangyangensis.DSM.19435 GCA_000375545.1_ASM37554v1
Sneathiella.glossodoripedis.JCM.23214 GCA_000616095.1_ASM61609v1
100/100
Geminicoccus.roseus.DSM.18922 GCA_000427665.1_ASM42766v1
99.8/100
Tistrella.mobilis.KA081020.065 GCA_000264455.2_ASM26445v2
Caenispirillum.salinarum.AK4 GCA_000315795.1
100/100
100/100
Pararhodospirillum.photometricum.DSM.122 GCA_000284415.2_ASM28441v2
100/100
40.4/86
Rhodospirillum.rubrum.F11 GCA_000225955.1_ASM22595v1
100/100
Roseospirillum.parvum GCA_900100455.1
Magnetospirillum.magneticum.AMB.1 GCA_000009985.1_ASM998v1
98.6/99
100/100
Terasakiella.pusilla.DSM.6293 GCA_000688235.1_ASM68823v1
99.9/71
Thalassospira.profundimaris.WP0211 GCA_000300275.1_ASM30027v1
100/100
GCA_002306855.1_ASM230685v1*
GCA_002471575.1_ASM247157v1*
100/100
Candidatus.Puniceispirillum.marinum.IMCC1322 GCA_000024465.1_ASM2446v1
100/74
100/100
100/100
Thalassobaculum.salexigens.DSM.19539 GCA_000423805.1_ASM42380v1
alpha.proteobacterium.BAL199 GCA_000171835.1_ASM17183v1
99.9/80
Oceanibaculum.indicum GCA_003633955.1_ASM363395v1
100/100
Kiloniella.laminariae.DSM.19542 GCA_000374005.1_ASM37400v1
100/100
100/80
Rhodovibrio.salinarum.DSM.9154 GCA_000515255.1_ASM51525v1
100/100
Inquilinus.limosus.DSM.16000 GCA_000423185.1_ASM42318v1
100/100
Rhodospirillum.centenum GCA_014197805.1_ASM1419780v1
Micavibrio.aeruginosavorus.ARL.13 GCA_000226315.1_ASM22631v1
Elioraea.tepidiphila.DSM.17972 GCA_000378465.1_ASM37846v1
96.4/99
100/100
Belnapia.moabensis.DSM.16746 GCA_000745835.1_ASM74583v1
100/100
Rubritepida.flocculans.DSM.14296 GCA_000425365.1_ASM42536v1
100/100
Roseomonas.gilardii GCA_001941945.1_ASM194194v1
Acidiphilium.multivorum.AIU301 GCA_000202835.1_ASM20283v1
98.5/99
99.5/100
Acidisphaera.rubrifaciens.HS.AP3 GCA_000964365.1_ASM96436v1
100/100
Granulibacter.bethesdensis.CGDNIH4 GCA_000576145.2_ASM57614v2
100/100
Commensalibacter.intestini.A911 GCA_000231445.2_ASM23144v1
97.7/100
Gluconacetobacter.diazotrophicus.PA1.5 GCA_000067045.1_ASM6704v1
Acetobacter.aceti.NBRC.14818 GCA_000193495.2_ASM19349v2
Gluconobacter.oxydans.DSM.3504 GCA_000583855.1_ASM58385v1
0.813566

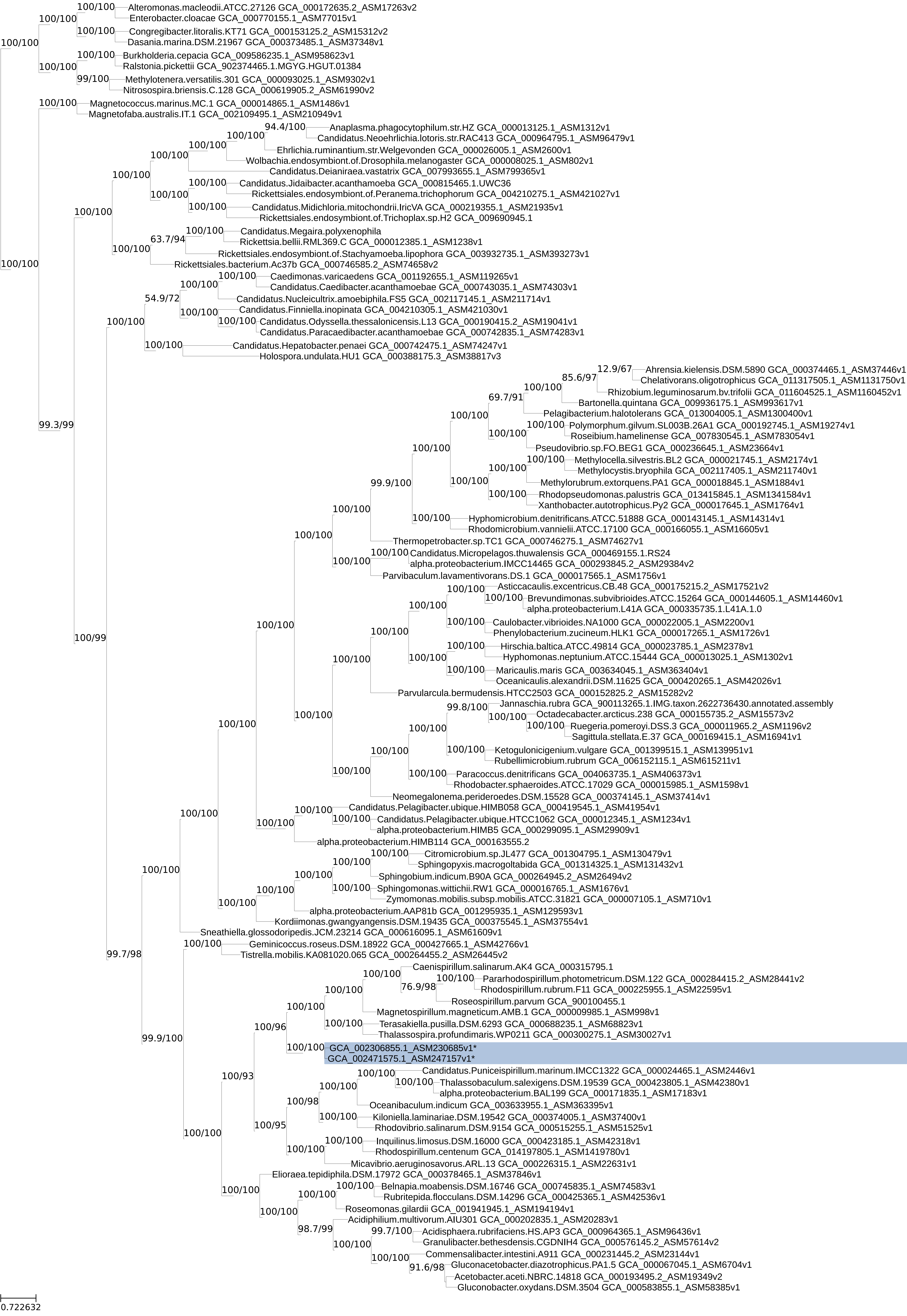

100/100 Alteromonas.macleodii.ATCC.27126 GCA_000172635.2_ASM17263v2
100/100 Enterobacter.cloacae GCA_000770155.1_ASM77015v1
100/100 100/100 Congregibacter.litoralis.KT71 GCA_000153125.2_ASM15312v2
Dasania.marina.DSM.21967 GCA_000373485.1_ASM37348v1
100/100 Burkholderia.cepacia GCA_009586235.1_ASM958623v1
100/100 Ralstonia.pickettii GCA_902374465.1.MGYG.HGUT.01384
99/100 Methylotenera.versatilis.301 GCA_000093025.1_ASM9302v1
Nitrosospira.briensis.C.128 GCA_000619905.2_ASM61990v2
100/100 Magnetococcus.marinus.MC.1 GCA_000014865.1_ASM1486v1
Magnetofaba.australis.IT.1 GCA_002109495.1_ASM210949v1
94.4/100 Anaplasma.phagocytophilum.str.HZ GCA_000013125.1_ASM1312v1
100/100 Candidatus.Neoehrlichia.lotoris.str.RAC413 GCA_000964795.1_ASM96479v1
100/100 Ehrlichia.ruminantium.str.Welgevonden GCA_000026005.1_ASM2600v1
100/100 Wolbachia.endosymbiont.of.Drosophila.melanogaster GCA_000008025.1_ASM802v1
100/100 Candidatus.Deianiraea.vastatrix GCA_007993655.1_ASM799365v1
100/100 Candidatus.Jidaibacter.acanthamoeba GCA_000815465.1.UWC36
100/100 Rickettsiales.endosymbiont.of.Peranema.trichophorum GCA_004210275.1_ASM421027v1
100/100 Candidatus.Midichloria.mitochondrii.IricVA GCA_000219355.1_ASM21935v1
100/100 Rickettsiales.endosymbiont.of.Trichoplax.sp.H2 GCA_009690945.1
100/100 Candidatus.Megaira.polyxenophila
63.7/94 Rickettsia.bellii.RML369.C GCA_000012385.1_ASM1238v1
Rickettsiales.endosymbiont.of.Stachyamoeba.lipophora GCA_003932735.1_ASM393273v1
100/100 Rickettsiales.bacterium.Ac37b GCA_000746585.2_ASM74658v2
100/100 100/100 Caedimonas.varicaedens GCA_001192655.1_ASM119265v1
Candidatus.Caedibacter.acanthamoebae GCA_000743035.1_ASM74303v1
54.9/72 Candidatus.Nucleicultrix.amoebiphila.FS5 GCA_002117145.1_ASM211714v1
100/100 Candidatus.Finniella.inopinata GCA_004210305.1_ASM421030v1
100/100 Candidatus.Odyssella.thessalonicensis.L13 GCA_000190415.2_ASM19041v1
100/100 Candidatus.Paracaedibacter.acanthamoebae GCA_000742835.1_ASM74283v1
100/100 Candidatus.Hepatobacter.penaei GCA_000742475.1_ASM74247v1
Holospora.undulata.HU1 GCA_000388175.3_ASM38817v3
12.9/67 Ahrensia.kielensis.DSM.5890 GCA_000374465.1_ASM37446v1
85.6/97 Chelativorans.oligotrophicus GCA_011317505.1_ASM1131750v1
100/100 Rhizobium.leguminosarum.bv.trifolii GCA_011604525.1_ASM1160452v1
69.7/91 Bartonella.quintana GCA_009936175.1_ASM993617v1
Pelagibacterium.halotolerans GCA_013004005.1_ASM1300400v1
100/100 100/100 Polymorphum.gilvum.SL003B.26A1 GCA_000192745.1_ASM19274v1
100/100 Roseibium.hamelinense GCA_007830545.1_ASM783054v1
99.3/99 Pseudovibrio.sp.FO.BEG1 GCA_000236645.1_ASM23664v1
100/100 100/100 Methylocella.silvestris.BL2 GCA_000021745.1_ASM2174v1
100/100 Methylocystis.bryophila GCA_002117405.1_ASM211740v1
100/100 Methylorubrum.extorquens.PA1 GCA_000018845.1_ASM1884v1
99.9/100 100/100 Rhodopseudomonas.palustris GCA_013415845.1_ASM1341584v1
Xanthobacter.autotrophicus.Py2 GCA_000017645.1_ASM1764v1
100/100 100/100 Hyphomicrobium.denitrificans.ATCC.51888 GCA_000143145.1_ASM14314v1
Rhodomicrobium.vannielii.ATCC.17100 GCA_000166055.1_ASM16605v1
100/100 Thermopetrobacter.sp.TC1 GCA_000746275.1_ASM74627v1
100/100 Candidatus.Micropelagos.thuwalensis GCA_000469155.1.RS24
100/100 alpha.proteobacterium.IMCC14465 GCA_000293845.2_ASM29384v2
Parvibaculum.lavamentivorans.DS.1 GCA_000017565.1_ASM1756v1
100/100 Asticcacaulis.excentricus.CB.48 GCA_000175215.2_ASM17521v2
100/100 100/100 Brevundimonas.subvibrioides.ATCC.15264 GCA_000144605.1_ASM14460v1
alpha.proteobacterium.L41A GCA_000335735.1.L41A.1.0
100/100 Caulobacter.vibrioides.NA1000 GCA_000022005.1_ASM2200v1
100/100 Phenylobacterium.zucineum.HLK1 GCA_000017265.1_ASM1726v1
100/100 100/100 Hirschia.baltica.ATCC.49814 GCA_000023785.1_ASM2378v1
100/100 Hyphomonas.neptunium.ATCC.15444 GCA_000013025.1_ASM1302v1
100/100 100/100 Maricaulis.maris GCA_003634045.1_ASM363404v1
Oceanicaulis.alexandrii.DSM.11625 GCA_000420265.1_ASM42026v1
100/99 Parvularcula.bermudensis.HTCC2503 GCA_000152825.2_ASM15282v2
100/100 99.8/100 Jannaschia.rubra GCA_900113265.1.IMG.taxon.2622736430.annotated.assembly
100/100 100/100 Octadecabacter.arcticus.238 GCA_000155735.2_ASM15573v2
100/100 100/100 Ruegeria.pomeroyi.DSS.3 GCA_000011965.2_ASM1196v2
Sagittula.stellata.E.37 GCA_000169415.1_ASM16941v1
100/100 100/100 Ketogulonicigenium.vulgare GCA_001399515.1_ASM139951v1
Rubellimicrobium.rubrum GCA_006152115.1_ASM615211v1
100/100 100/100 Paracoccus.denitrificans GCA_004063735.1_ASM406373v1
Rhodobacter.sphaeroides.ATCC.17029 GCA_000015985.1_ASM1598v1
Neomegalonema.perideroedes.DSM.15528 GCA_000374145.1_ASM37414v1
100/100 100/100 Candidatus.Pelagibacter.ubique.HIMB058 GCA_000419545.1_ASM41954v1
100/100 Candidatus.Pelagibacter.ubique.HTCC1062 GCA_000012345.1_ASM1234v1
100/100 alpha.proteobacterium.HIMB5 GCA_000299095.1_ASM29909v1
alpha.proteobacterium.HIMB114 GCA_000163555.2
100/100 Citromicrobium.sp.JL477 GCA_001304795.1_ASM130479v1
100/100 100/100 Sphingopyxis.macrogoltabida GCA_001314325.1_ASM131432v1
100/100 Sphingobium.indicum.B90A GCA_000264945.2_ASM26494v2
100/100 100/100 Sphingomonas.wittichii.RW1 GCA_000016765.1_ASM1676v1
100/100 Zymomonas.mobilis.subsp.mobilis.ATCC.31821 GCA_000007105.1_ASM710v1
alpha.proteobacterium.AAP81b GCA_001295935.1_ASM129593v1
Kordiimonas.gwangyangensis.DSM.19435 GCA_000375545.1_ASM37554v1
Sneathiella.glossodoripedis.JCM.23214 GCA_000616095.1_ASM61609v1
100/100 Geminicoccus.roseus.DSM.18922 GCA_000427665.1_ASM42766v1
99.7/98 Tistrella.mobilis.KA081020.065 GCA_000264455.2_ASM26445v2
Caenispirillum.salinarum.AK4 GCA_000315795.1
100/100 100/100 Pararhodospirillum.photometricum.DSM.122 GCA_000284415.2_ASM28441v2
100/100 76.9/98 Rhodospirillum.rubrum.F11 GCA_000225955.1_ASM22595v1
100/100 Roseospirillum.parvum GCA_900100455.1
Magnetospirillum.magneticum.AMB.1 GCA_000009985.1_ASM998v1
99.9/100 100/96 100/100 Terasakiella.pusilla.DSM.6293 GCA_000688235.1_ASM68823v1
Thalassospira.profundimaris.WP0211 GCA_000300275.1_ASM30027v1
100/100 GCA_002306855.1_ASM230685v1*
GCA_002471575.1_ASM247157v1*
100/100 Candidatus.Puniceispirillum.marinum.IMCC1322 GCA_000024465.1_ASM2446v1
100/93 100/100 Thalassobaculum.salexigens.DSM.19539 GCA_000423805.1_ASM42380v1
100/100 alpha.proteobacterium.BAL199 GCA_000171835.1_ASM17183v1
100/98 Oceanibaculum.indicum GCA_003633955.1_ASM363395v1
100/100 100/100 Kiloniella.laminariae.DSM.19542 GCA_000374005.1_ASM37400v1
100/95 Rhodovibrio.salinarum.DSM.9154 GCA_000515255.1_ASM51525v1
100/100 100/100 Inquilinus.limosus.DSM.16000 GCA_000423185.1_ASM42318v1
Rhodospirillum.centenum GCA_014197805.1_ASM1419780v1
Micavibrio.aeruginosavorus.ARL.13 GCA_000226315.1_ASM22631v1
Elioraea.tepidiphila.DSM.17972 GCA_000378465.1_ASM37846v1
100/100 Belnapia.moabensis.DSM.16746 GCA_000745835.1_ASM74583v1
100/100 100/100 Rubritepida.flocculans.DSM.14296 GCA_000425365.1_ASM42536v1
100/100 Roseomonas.gilardii GCA_001941945.1_ASM194194v1
Acidiphilium.multivorum.AIU301 GCA_000202835.1_ASM20283v1
98.7/99 99.7/100 Acidisphaera.rubrifaciens.HS.AP3 GCA_000964365.1_ASM96436v1
100/100 Granulibacter.bethesdensis.CGDNIH4 GCA_000576145.2_ASM57614v2
Commensalibacter.intestini.A911 GCA_000231445.2_ASM23144v1
100/100 Gluconacetobacter.diazotrophicus.PA1.5 GCA_000067045.1_ASM6704v1
91.6/98 Acetobacter.aceti.NBRC.14818 GCA_000193495.2_ASM19349v2
Gluconobacter.oxydans.DSM.3504 GCA_000583855.1_ASM58385v1
0.722632

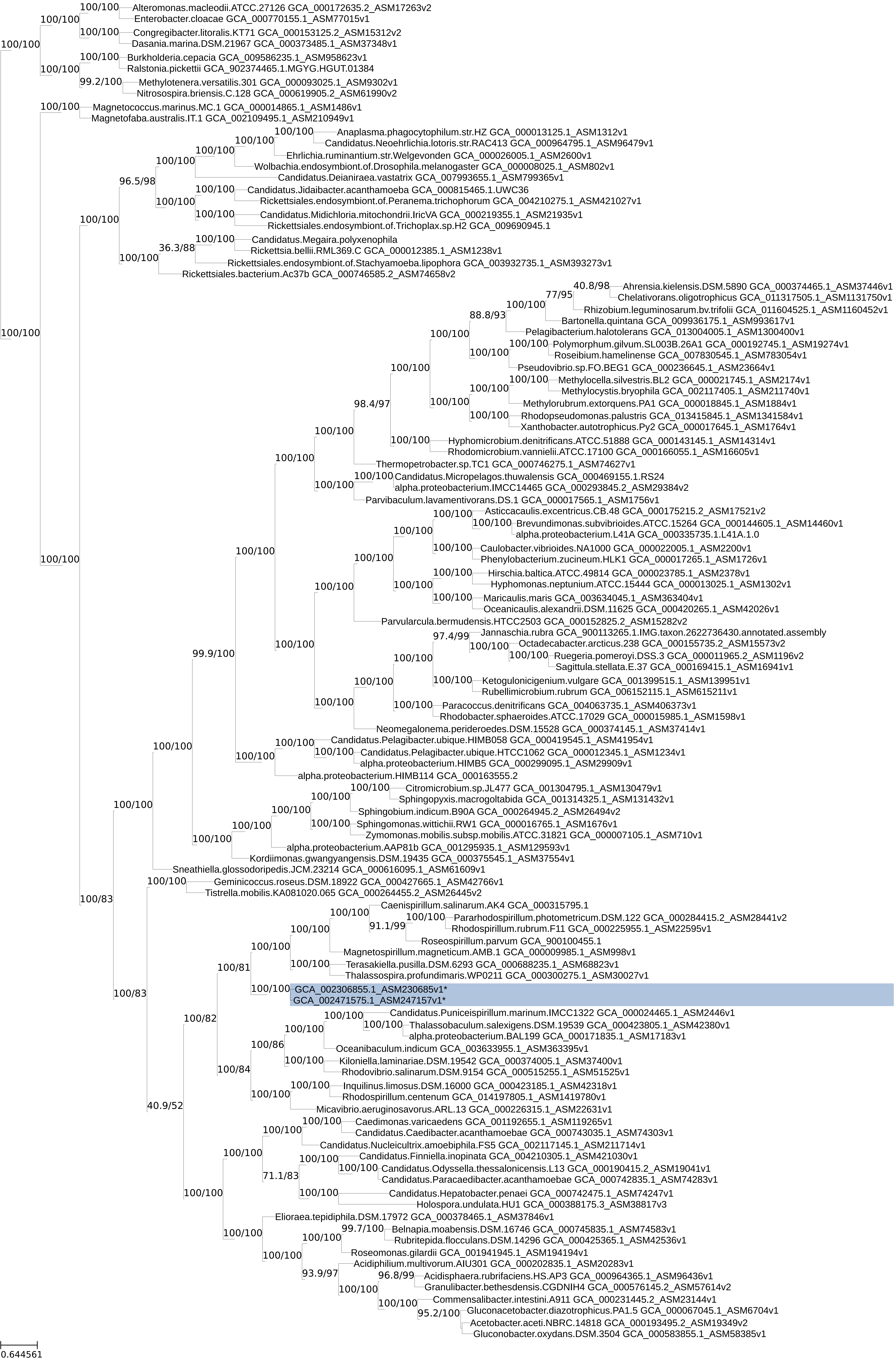
Alteromonas.macleodii.ATCC.27126 GCA_000172635.2_ASM17263v2
Enterobacter.cloacae GCA_000770155.1_ASM77015v1
Congregibacter.litoralis.KT71 GCA_000153125.2_ASM15312v2
Dasania.marina.DSM.21967 GCA_000373485.1_ASM37348v1
Burkholderia.cepacia GCA_009586235.1_ASM958623v1
Ralstonia.pickettii GCA_902374465.1.MGYG.HGUT.01384
Methylotenera.versatilis.301 GCA_000093025.1_ASM9302v1
Nitrosospira.briensis.C.128 GCA_000619905.2_ASM61990v2
Magnetococcus.marinus.MC.1 GCA_000014865.1_ASM1486v1
Magnetofaba.australis.IT.1 GCA_002109495.1_ASM210949v1
Anaplasma.phagocytophilum.str.HZ GCA_000013125.1_ASM1312v1
Candidatus.Neoehrlichia.lotoris.str.RAC413 GCA_000964795.1_ASM96479v1
Ehrlichia.ruminantium.str.Welgevonden GCA_000026005.1_ASM2600v1
Wolbachia.endosymbiont.of.Drosophila.melanogaster GCA_000008025.1_ASM802v1
Candidatus.Deianiraea.vastatrix GCA_007993655.1_ASM799365v1
Candidatus.Jidaibacter.acanthamoeba GCA_000815465.1.UWC36
Rickettsiales.endosymbiont.of.Peranema.trichophorum GCA_004210275.1_ASM421027v1
Candidatus.Midichloria.mitochondrii.IricVA GCA_000219355.1_ASM21935v1
Rickettsiales.endosymbiont.of.Trichoplax.sp.H2 GCA_009690945.1
Candidatus.Megaira.polyxenophila
Rickettsia.bellii.RML369.C GCA_000012385.1_ASM1238v1
Rickettsiales.endosymbiont.of.Stachyamoeba.lipophora GCA_003932735.1_ASM393273v1
Rickettsiales.bacterium.Ac37b GCA_000746585.2_ASM74658v2
Ahrensia.kielensis.DSM.5890 GCA_000374465.1_ASM37446v1
Chelativorans.oligotrophicus GCA_011317505.1_ASM1131750v1
Rhizobium.leguminosarum.bv.trifolii GCA_011604525.1_ASM1160452v1
Bartonella.quintana GCA_009936175.1_ASM993617v1
Pelagibacterium.halotolerans GCA_013004005.1_ASM1300400v1
Polymorphum.gilvum.SL003B.26A1 GCA_000192745.1_ASM19274v1
Roseibium.hamelinense GCA_007830545.1_ASM783054v1
Pseudovibrio.sp.FO.BEG1 GCA_000236645.1_ASM23664v1
Methylocella.silvestris.BL2 GCA_000021745.1_ASM2174v1
Methylocystis.bryophila GCA_002117405.1_ASM211740v1
Methylorubrum.extorquens.PA1 GCA_000018845.1_ASM1884v1
Rhodopseudomonas.palustris GCA_013415845.1_ASM1341584v1
Xanthobacter.autotrophicus.Py2 GCA_000017645.1_ASM1764v1
Hyphomicrobium.denitrificans.ATCC.51888 GCA_000143145.1_ASM14314v1
Rhodomicrobium.vannielii.ATCC.17100 GCA_000166055.1_ASM16605v1
Thermopetrobacter.sp.TC1 GCA_000746275.1_ASM74627v1
Candidatus.Micropelagos.thuwalensis GCA_000469155.1.RS24
alpha.proteobacterium.IMCC14465 GCA_000293845.2_ASM29384v2
Parvibaculum.lavamentivorans.DS.1 GCA_000017565.1_ASM1756v1
Asticcacaulis.excentricus.CB.48 GCA_000175215.2_ASM17521v2
Brevundimonas.subvibrioides.ATCC.15264 GCA_000144605.1_ASM14460v1
alpha.proteobacterium.L41A GCA_000335735.1.L41A.1.0
Caulobacter.vibrioides.NA1000 GCA_000022005.1_ASM2200v1
Phenylobacterium.zucineum.HLK1 GCA_000017265.1_ASM1726v1
Hirschia.baltica.ATCC.49814 GCA_000023785.1_ASM2378v1
Hyphomonas.neptunium.ATCC.15444 GCA_000013025.1_ASM1302v1
Maricaulis.maris GCA_003634045.1_ASM363404v1
Oceanicaulis.alexandrii.DSM.11625 GCA_000420265.1_ASM42026v1
Parvularcula.bermudensis.HTCC2503 GCA_000152825.2_ASM15282v2
Jannaschia.rubra GCA_900113265.1.IMG.taxon.2622736430.annotated.assembly
Octadecabacter.arcticus.238 GCA_000155735.2_ASM15573v2
Ruegeria.pomeroyi.DSS.3 GCA_000011965.2_ASM1196v2
Sagittula.stellata.E.37 GCA_000169415.1_ASM16941v1
Ketogulonicigenium.vulgare GCA_001399515.1_ASM139951v1
Rubellimicrobium.rubrum GCA_006152115.1_ASM615211v1
Paracoccus.denitrificans GCA_004063735.1_ASM406373v1
Rhodobacter.sphaeroides.ATCC.17029 GCA_000015985.1_ASM1598v1
Neomegalonema.perideroedes.DSM.15528 GCA_000374145.1_ASM37414v1
Candidatus.Pelagibacter.ubique.HIMB058 GCA_000419545.1_ASM41954v1
Candidatus.Pelagibacter.ubique.HTCC1062 GCA_000012345.1_ASM1234v1
alpha.proteobacterium.HIMB5 GCA_000299095.1_ASM29909v1
alpha.proteobacterium.HIMB114 GCA_000163555.2
Citromicrobium.sp.JL477 GCA_001304795.1_ASM130479v1
Sphingopyxis.macrogoltabida GCA_001314325.1_ASM131432v1
Sphingobium.indicum.B90A GCA_000264945.2_ASM26494v2
Sphingomonas.wittichii.RW1 GCA_000016765.1_ASM1676v1
Zymomonas.mobilis.subsp.mobilis.ATCC.31821 GCA_000007105.1_ASM710v1
alpha.proteobacterium.AAP81b GCA_001295935.1_ASM129593v1
Kordiimonas.gwangyangensis.DSM.19435 GCA_000375545.1_ASM37554v1
Sneathiella.glossodoripedis.JCM.23214 GCA_000616095.1_ASM61609v1
Geminicoccus.roseus.DSM.18922 GCA_000427665.1_ASM42766v1
Tistrella.mobilis.KA081020.065 GCA_000264455.2_ASM26445v2
Caenispirillum.salinarum.AK4 GCA_000315795.1
Pararhodospirillum.photometricum.DSM.122 GCA_000284415.2_ASM28441v2
Rhodospirillum.rubrum.F11 GCA_000225955.1_ASM22595v1
Roseospirillum.parvum GCA_900100455.1
Magnetospirillum.magneticum.AMB.1 GCA_000009985.1_ASM998v1
Terasakiella.pusilla.DSM.6293 GCA_000688235.1_ASM68823v1
Thalassospira.profundimaris.WP0211 GCA_000300275.1_ASM30027v1
GCA_002306855.1_ASM230685v1*
GCA_002471575.1_ASM247157v1*
Candidatus.Puniceispirillum.marinum.IMCC1322 GCA_000024465.1_ASM2446v1
Thalassobaculum.salexigens.DSM.19539 GCA_000423805.1_ASM42380v1
alpha.proteobacterium.BAL199 GCA_000171835.1_ASM17183v1
Oceanibaculum.indicum GCA_003633955.1_ASM363395v1
Kiloniella.laminariae.DSM.19542 GCA_000374005.1_ASM37400v1
Rhodovibrio.salinarum.DSM.9154 GCA_000515255.1_ASM51525v1
Inquilinus.limosus.DSM.16000 GCA_000423185.1_ASM42318v1
Rhodospirillum.centenum GCA_014197805.1_ASM1419780v1
Micavibrio.aeruginosavorus.ARL.13 GCA_000226315.1_ASM22631v1
Caedimonas.varicaedens GCA_001192655.1_ASM119265v1
Candidatus.Caedibacter.acanthamoebae GCA_000743035.1_ASM74303v1
Candidatus.Nucleicultrix.amoebiphila.FS5 GCA_002117145.1_ASM211714v1
Candidatus.Finniella.inopinata GCA_004210305.1_ASM421030v1
Candidatus.Odyssella.thessalonicensis.L13 GCA_000190415.2_ASM19041v1
Candidatus.Paracaedibacter.acanthamoebae GCA_000742835.1_ASM74283v1
Candidatus.Hepatobacter.penaei GCA_000742475.1_ASM74247v1
Holospora.undulata.HU1 GCA_000388175.3_ASM38817v3
Elioraea.tepidiphila.DSM.17972 GCA_000378465.1_ASM37846v1
Belnapia.moabensis.DSM.16746 GCA_000745835.1_ASM74583v1
Rubritepida.flocculans.DSM.14296 GCA_000425365.1_ASM42536v1
Roseomonas.gilardii GCA_001941945.1_ASM194194v1
Acidiphilium.multivorum.AIU301 GCA_000202835.1_ASM20283v1
Acidisphaera.rubrifaciens.HS.AP3 GCA_000964365.1_ASM96436v1
Granulibacter.bethesdensis.CGDNIH4 GCA_000576145.2_ASM57614v2
Commensalibacter.intestini.A911 GCA_000231445.2_ASM23144v1
Gluconacetobacter.diazotrophicus.PA1.5 GCA_000067045.1_ASM6704v1
Acetobacter.aceti.NBRC.14818 GCA_000193495.2_ASM19349v2
Gluconobacter.oxydans.DSM.3504 GCA_000583855.1_ASM58385v1
0.644561

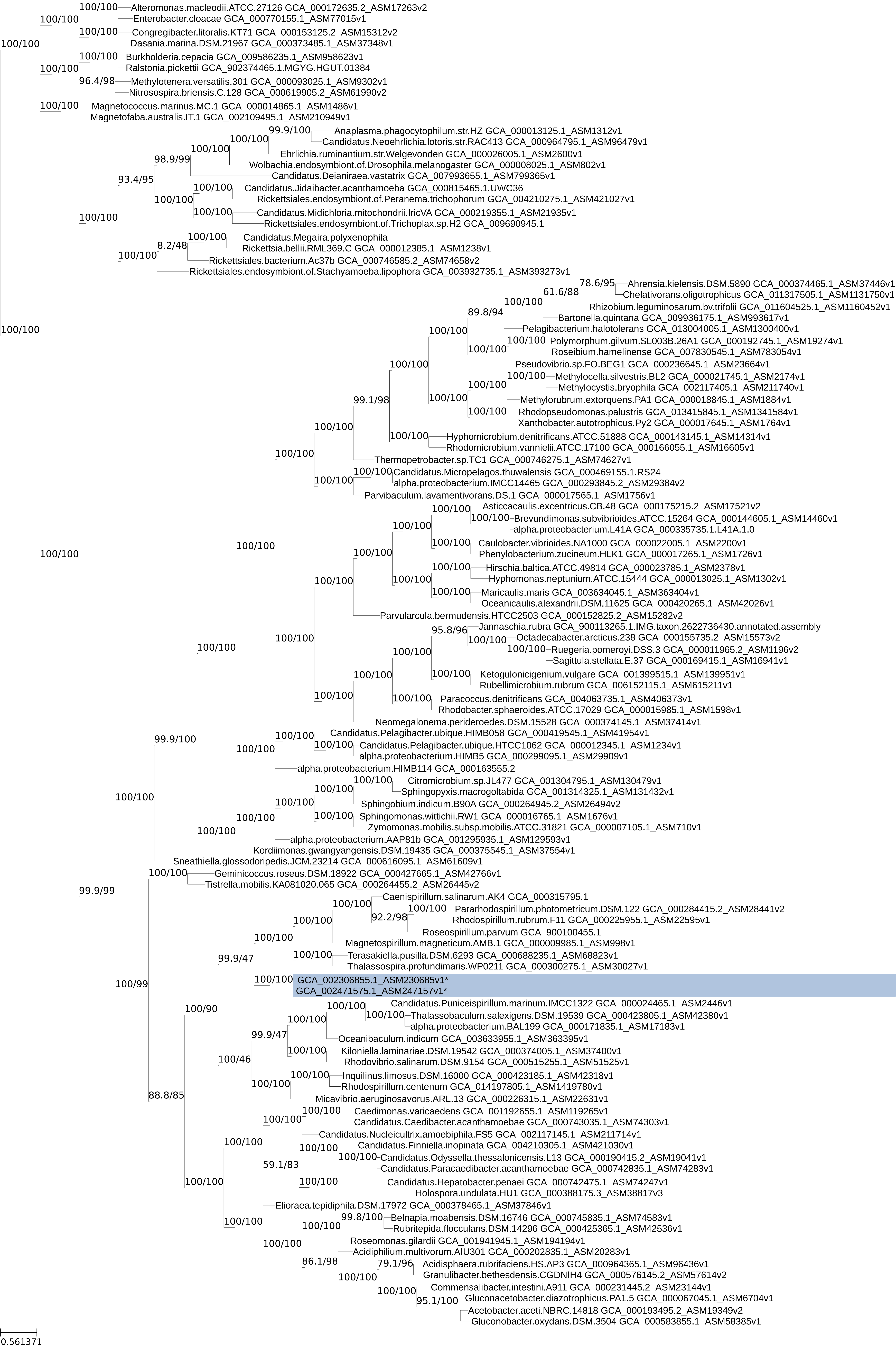

100/100
100/100
100/100
Alteromonas.macleodii.ATCC.27126 GCA_000172635.2_ASM17263v2
Enterobacter.cloacae GCA_000770155.1_ASM77015v1
100/100
Congregibacter.litoralis.KT71 GCA_000153125.2_ASM15312v2
Dasania.marina.DSM.21967 GCA_000373485.1_ASM37348v1
100/100
100/100
Burkholderia.cepacia GCA_009586235.1_ASM958623v1
Ralstonia.pickettii GCA_902374465.1.MGYG.HGUT.01384
96.4/98
Methylotenera.versatilis.301 GCA_000093025.1_ASM9302v1
Nitrosospira.briensis.C.128 GCA_000619905.2_ASM61990v2
100/100
Magnetococcus.marinus.MC.1 GCA_000014865.1_ASM1486v1
Magnetofaba.australis.IT.1 GCA_002109495.1_ASM210949v1
99.9/100
Anaplasma.phagocytophilum.str.HZ GCA_000013125.1_ASM1312v1
Candidatus.Neoehrlichia.lotoris.str.RAC413 GCA_000964795.1_ASM96479v1
100/100
Ehrlichia.ruminantium.str.Welgevonden GCA_000026005.1_ASM2600v1
100/100
Wolbachia.endosymbiont.of.Drosophila.melanogaster GCA_000008025.1_ASM802v1
98.9/99
Candidatus.Deianiraea.vastatrix GCA_007993655.1_ASM799365v1
93.4/95
100/100
Candidatus.Jidaibacter.acanthamoeba GCA_000815465.1.UWC36
Rickettsiales.endosymbiont.of.Peranema.trichophorum GCA_004210275.1_ASM421027v1
100/100
100/100
Candidatus.Midichloria.mitochondrii.IricVA GCA_000219355.1_ASM21935v1
Rickettsiales.endosymbiont.of.Trichoplax.sp.H2 GCA_009690945.1
100/100
100/100
Candidatus.Megaira.polyxenophila
8.2/48
Rickettsia.bellii.RML369.C GCA_000012385.1_ASM1238v1
Rickettsiales.bacterium.Ac37b GCA_000746585.2_ASM74658v2
100/100
Rickettsiales.endosymbiont.of.Stachyamoeba.lipophora GCA_003932735.1_ASM393273v1
78.6/95
Ahrensia.kielensis.DSM.5890 GCA_000374465.1_ASM37446v1
61.6/88
Chelativorans.oligotrophicus GCA_011317505.1_ASM1131750v1
100/100
Rhizobium.leguminosarum.bv.trifolii GCA_011604525.1_ASM1160452v1
89.8/94
Bartonella.quintana GCA_009936175.1_ASM993617v1
Pelagibacterium.halotolerans GCA_013004005.1_ASM1300400v1
100/100
100/100
100/100
Polymorphum.gilvum.SL003B.26A1 GCA_000192745.1_ASM19274v1
Roseibium.hamelinense GCA_007830545.1_ASM783054v1
Pseudovibrio.sp.FO.BEG1 GCA_000236645.1_ASM23664v1
100/100
100/100
100/100
Methylocella.silvestris.BL2 GCA_000021745.1_ASM2174v1
Methylocystis.bryophila GCA_002117405.1_ASM211740v1
99.1/98
100/100
Methylorubrum.extorquens.PA1 GCA_000018845.1_ASM1884v1
100/100
Rhodopseudomonas.palustris GCA_013415845.1_ASM1341584v1
Xanthobacter.autotrophicus.Py2 GCA_000017645.1_ASM1764v1
100/100
100/100
Hyphomicrobium.denitrificans.ATCC.51888 GCA_000143145.1_ASM14314v1
Rhodomicrobium.vannielii.ATCC.17100 GCA_000166055.1_ASM16605v1
100/100
Thermopetrobacter.sp.TC1 GCA_000746275.1_ASM74627v1
100/100
Candidatus.Micropelagos.thuwalensis GCA_000469155.1.RS24
100/100
alpha.proteobacterium.IMCC14465 GCA_000293845.2_ASM29384v2
Parvibaculum.lavamentivorans.DS.1 GCA_000017565.1_ASM1756v1
100/100
Asticcacaulis.excentricus.CB.48 GCA_000175215.2_ASM17521v2
100/100
100/100
Brevundimonas.subvibrioides.ATCC.15264 GCA_000144605.1_ASM14460v1
alpha.proteobacterium.L41A GCA_000335735.1.L41A.1.0
100/100
100/100
Caulobacter.vibrioides.NA1000 GCA_000022005.1_ASM2200v1
Phenylobacterium.zucineum.HLK1 GCA_000017265.1_ASM1726v1
100/100
100/100
Hirschia.baltica.ATCC.49814 GCA_000023785.1_ASM2378v1
Hyphomonas.neptunium.ATCC.15444 GCA_000013025.1_ASM1302v1
100/100
100/100
Maricaulis.maris GCA_003634045.1_ASM363404v1
Oceanicaulis.alexandrii.DSM.11625 GCA_000420265.1_ASM42026v1
Parvularcula.bermudensis.HTCC2503 GCA_000152825.2_ASM15282v2
95.8/96
Jannaschia.rubra GCA_900113265.1.IMG.taxon.2622736430.annotated.assembly
100/100
100/100
Octadecabacter.arcticus.238 GCA_000155735.2_ASM15573v2
100/100
Ruegeria.pomeroyi.DSS.3 GCA_000011965.2_ASM1196v2
Sagittula.stellata.E.37 GCA_000169415.1_ASM16941v1
100/100
100/100
Ketogulonicigenium.vulgare GCA_001399515.1_ASM139951v1
Rubellimicrobium.rubrum GCA_006152115.1_ASM615211v1
100/100
100/100
Paracoccus.denitrificans GCA_004063735.1_ASM406373v1
Rhodobacter.sphaeroides.ATCC.17029 GCA_000015985.1_ASM1598v1
Neomegalonema.perideroedes.DSM.15528 GCA_000374145.1_ASM37414v1
100/100
Candidatus.Pelagibacter.ubique.HIMB058 GCA_000419545.1_ASM41954v1
100/100
100/100
Candidatus.Pelagibacter.ubique.HTCC1062 GCA_000012345.1_ASM1234v1
alpha.proteobacterium.HIMB5 GCA_000299095.1_ASM29909v1
99.9/100
alpha.proteobacterium.HIMB114 GCA_000163555.2
100/100
Citromicrobium.sp.JL477 GCA_001304795.1_ASM130479v1
100/100
Sphingopyxis.macrogoltabida GCA_001314325.1_ASM131432v1
100/100
Sphingobium.indicum.B90A GCA_000264945.2_ASM26494v2
100/100
100/100
Sphingomonas.wittichii.RW1 GCA_000016765.1_ASM1676v1
100/100
Zymomonas.mobilis.subsp.mobilis.ATCC.31821 GCA_000007105.1_ASM710v1
100/100
alpha.proteobacterium.AAP81b GCA_001295935.1_ASM129593v1
Kordiimonas.gwangyangensis.DSM.19435 GCA_000375545.1_ASM37554v1
Sneathiella.glossodoripedis.JCM.23214 GCA_000616095.1_ASM61609v1
100/100
Geminicoccus.roseus.DSM.18922 GCA_000427665.1_ASM42766v1
Tistrella.mobilis.KA081020.065 GCA_000264455.2_ASM26445v2
99.9/99
Caenispirillum.salinarum.AK4 GCA_000315795.1
100/100
100/100
Pararhodospirillum.photometricum.DSM.122 GCA_000284415.2_ASM28441v2
92.2/98
Rhodospirillum.rubrum.F11 GCA_000225955.1_ASM22595v1
100/100
Roseospirillum.parvum GCA_900100455.1
100/100
Magnetospirillum.magneticum.AMB.1 GCA_000009985.1_ASM998v1
99.9/47
100/100
Terasakiella.pusilla.DSM.6293 GCA_000688235.1_ASM68823v1
Thalassospira.profundimaris.WP0211 GCA_000300275.1_ASM30027v1
100/100
GCA_002306855.1_ASM230685v1*
GCA_002471575.1_ASM247157v1*
100/99
100/90
100/100
Candidatus.Puniceispirillum.marinum.IMCC1322 GCA_000024465.1_ASM2446v1
100/100
100/100
Thalassobaculum.salexigens.DSM.19539 GCA_000423805.1_ASM42380v1
alpha.proteobacterium.BAL199 GCA_000171835.1_ASM17183v1
99.9/47
Oceanibaculum.indicum GCA_003633955.1_ASM363395v1
100/100
Kiloniella.laminariae.DSM.19542 GCA_000374005.1_ASM37400v1
100/46
Rhodovibrio.salinarum.DSM.9154 GCA_000515255.1_ASM51525v1
100/100
Inquilinus.limosus.DSM.16000 GCA_000423185.1_ASM42318v1
100/100
Rhodospirillum.centenum GCA_014197805.1_ASM1419780v1
88.8/85
Micavibrio.aeruginosavorus.ARL.13 GCA_000226315.1_ASM22631v1
100/100
Caedimonas.varicaedens GCA_001192655.1_ASM119265v1
100/100
Candidatus.Caedibacter.acanthamoebae GCA_000743035.1_ASM74303v1
Candidatus.Nucleicultrix.amoebiphila.FS5 GCA_002117145.1_ASM211714v1
100/100
100/100
Candidatus.Finniella.inopinata GCA_004210305.1_ASM421030v1
100/100
Candidatus.Odyssella.thessalonicensis.L13 GCA_000190415.2_ASM19041v1
59.1/83
Candidatus.Paracaedibacter.acanthamoebae GCA_000742835.1_ASM74283v1
100/100
100/100
Candidatus.Hepatobacter.penaei GCA_000742475.1_ASM74247v1
Holospora.undulata.HU1 GCA_000388175.3_ASM38817v3
Elioraea.tepidiphila.DSM.17972 GCA_000378465.1_ASM37846v1
100/100
99.8/100
Belnapia.moabensis.DSM.16746 GCA_000745835.1_ASM74583v1
100/100
Rubritepida.flocculans.DSM.14296 GCA_000425365.1_ASM42536v1
100/100
Roseomonas.gilardii GCA_001941945.1_ASM194194v1
Acidiphilium.multivorum.AIU301 GCA_000202835.1_ASM20283v1
86.1/98
79.1/96
Acidisphaera.rubrifaciens.HS.AP3 GCA_000964365.1_ASM96436v1
100/100
Granulibacter.bethesdensis.CGDNIH4 GCA_000576145.2_ASM57614v2
Commensalibacter.intestini.A911 GCA_000231445.2_ASM23144v1
100/100
95.1/100
Gluconacetobacter.diazotrophicus.PA1.5 GCA_000067045.1_ASM6704v1
Acetobacter.aceti.NBRC.14818 GCA_000193495.2_ASM19349v2
Gluconobacter.oxydans.DSM.3504 GCA_000583855.1_ASM58385v1
0.561371

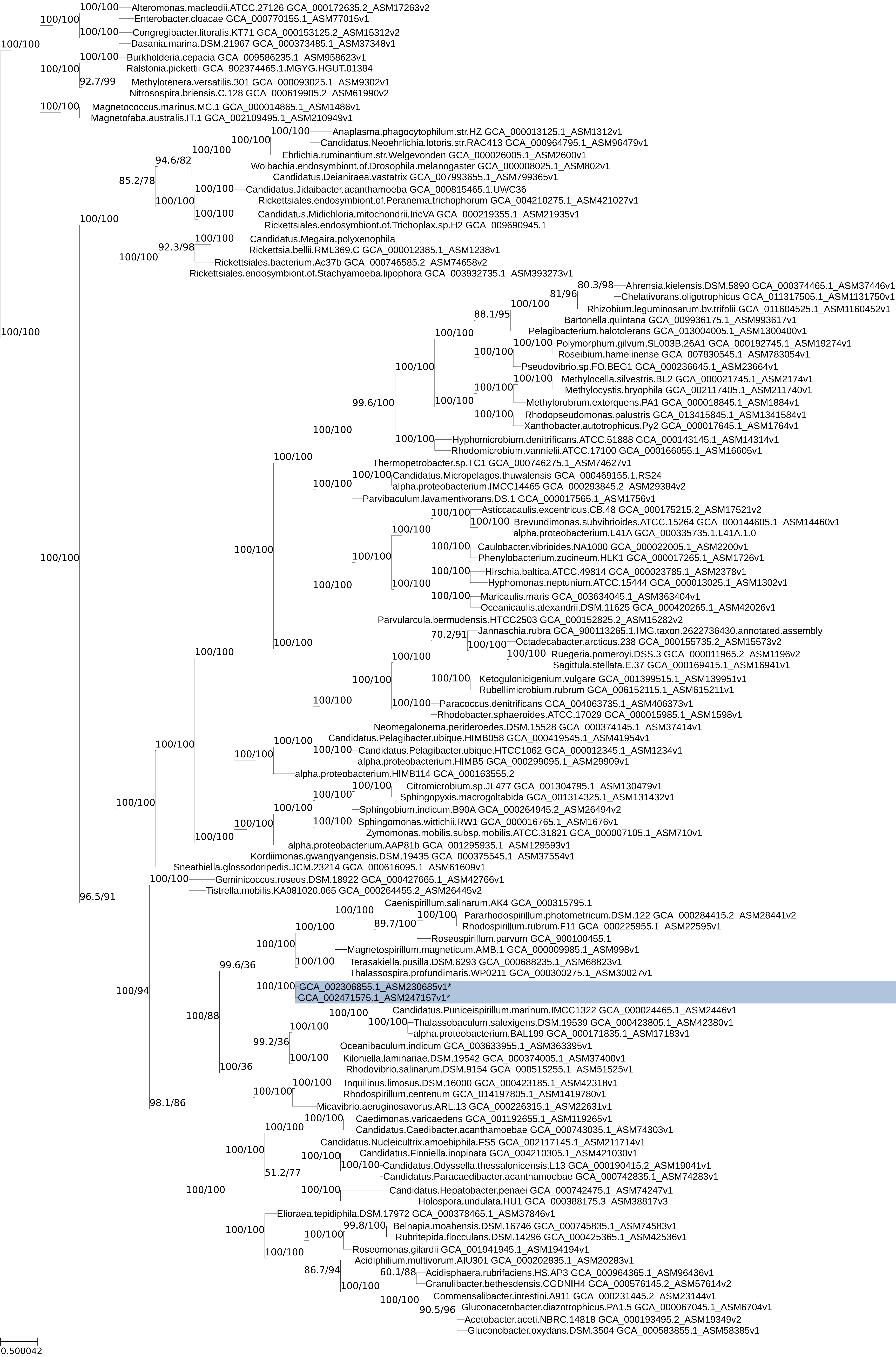
Alteromonas.macleodii.ATCC.27126 GCA_000172635.2_ASM17263v2
Enterobacter.cloacae GCA_000770155.1_ASM77015v1
Congregibacter.litoralis.KT71 GCA_000153125.2_ASM15312v2
Dasania.marina.DSM.21967 GCA_000373485.1_ASM37348v1
Burkholderia.cepacia GCA_009586235.1_ASM958623v1
Ralstonia.pickettii GCA_902374465.1.MGYG.HGUT.01384
Methylotenera.versatilis.301 GCA_000093025.1_ASM9302v1
Nitrosospira.briensis.C.128 GCA_000619905.2_ASM61990v2
Magnetococcus.marinus.MC.1 GCA_000014865.1_ASM1486v1
Magnetofaba.australis.IT.1 GCA_002109495.1_ASM210949v1
Anaplasma.phagocytophilum.str.HZ GCA_000013125.1_ASM1312v1
Candidatus.Neoehrlichia.lotoris.str.RAC413 GCA_000964795.1_ASM96479v1
Ehrlichia.ruminantium.str.Welgevonden GCA_000026005.1_ASM2600v1
Wolbachia.endosymbiont.of.Drosophila.melanogaster GCA_000008025.1_ASM802v1
Candidatus.Deianiraea.vastatrix GCA_007993655.1_ASM799365v1
Candidatus.Jidaibacter.acanthamoeba GCA_000815465.1.UWC36
Rickettsiales.endosymbiont.of.Peranema.trichophorum GCA_004210275.1_ASM421027v1
Candidatus.Midichloria.mitochondrii.IricVA GCA_000219355.1_ASM21935v1
Rickettsiales.endosymbiont.of.Trichoplax.sp.H2 GCA_009690945.1
Candidatus.Megaira.polyxenophila
Rickettsia.bellii.RML369.C GCA_000012385.1_ASM1238v1
Rickettsiales.bacterium.Ac37b GCA_000746585.2_ASM74658v2
Rickettsiales.endosymbiont.of.Stachyamoeba.lipophora GCA_003932735.1_ASM393273v1
Ahrensia.kielensis.DSM.5890 GCA_000374465.1_ASM37446v1
Chelativorans.oligotrophicus GCA_011317505.1_ASM1131750v1
Rhizobium.leguminosarum.bv.trifolii GCA_011604525.1_ASM1160452v1
Bartonella.quintana GCA_009936175.1_ASM993617v1
Pelagibacterium.halotolerans GCA_013004005.1_ASM1300400v1
Polymorphum.gilvum.SL003B.26A1 GCA_000192745.1_ASM19274v1
Roseibium.hamelinense GCA_007830545.1_ASM783054v1
Pseudovibrio.sp.FO.BEG1 GCA_000236645.1_ASM23664v1
Methylocella.silvestris.BL2 GCA_000021745.1_ASM2174v1
Methylocystis.bryophila GCA_002117405.1_ASM211740v1
Methylorubrum.extorquens.PA1 GCA_000018845.1_ASM1884v1
Rhodopseudomonas.palustris GCA_013415845.1_ASM1341584v1
Xanthobacter.autotrophicus.Py2 GCA_000017645.1_ASM1764v1
Hyphomicrobium.denitrificans.ATCC.51888 GCA_000143145.1_ASM14314v1
Rhodomicrobium.vannielii.ATCC.17100 GCA_000166055.1_ASM16605v1
Thermopetrobacter.sp.TC1 GCA_000746275.1_ASM74627v1
Candidatus.Micropelagos.thuwalensis GCA_000469155.1.RS24
alpha.proteobacterium.IMCC14465 GCA_000293845.2_ASM29384v2
Parvibaculum.lavamentivorans.DS.1 GCA_000017565.1_ASM1756v1
Asticcacaulis.excentricus.CB.48 GCA_000175215.2_ASM17521v2
Brevundimonas.subvibrioides.ATCC.15264 GCA_000144605.1_ASM14460v1
alpha.proteobacterium.L41A GCA_000335735.1.L41A.1.0
Caulobacter.vibrioides.NA1000 GCA_000022005.1_ASM2200v1
Phenylobacterium.zucineum.HLK1 GCA_000017265.1_ASM1726v1
Hirschia.baltica.ATCC.49814 GCA_000023785.1_ASM2378v1
Hyphomonas.neptunium.ATCC.15444 GCA_000013025.1_ASM1302v1
Maricaulis.maris GCA_003634045.1_ASM363404v1
Oceanicaulis.alexandrii.DSM.11625 GCA_000420265.1_ASM42026v1
Parvularcula.bermudensis.HTCC2503 GCA_000152825.2_ASM15282v2
Jannaschia.rubra GCA_900113265.1.IMG.taxon.2622736430.annotated.assembly
Octadecabacter.arcticus.238 GCA_000155735.2_ASM15573v2
Ruegeria.pomeroyi.DSS.3 GCA_000011965.2_ASM1196v2
Sagittula.stellata.E.37 GCA_000169415.1_ASM16941v1
Ketogulonicigenium.vulgare GCA_001399515.1_ASM139951v1
Rubellimicrobium.rubrum GCA_006152115.1_ASM615211v1
Paracoccus.denitrificans GCA_004063735.1_ASM406373v1
Rhodobacter.sphaeroides.ATCC.17029 GCA_000015985.1_ASM1598v1
Neomegalonema.perideroedes.DSM.15528 GCA_000374145.1_ASM37414v1
Candidatus.Pelagibacter.ubique.HIMB058 GCA_000419545.1_ASM41954v1
Candidatus.Pelagibacter.ubique.HTCC1062 GCA_000012345.1_ASM1234v1
alpha.proteobacterium.HIMB5 GCA_000299095.1_ASM29909v1
alpha.proteobacterium.HIMB114 GCA_000163555.2
Citromicrobium.sp.JL477 GCA_001304795.1_ASM130479v1
Sphingopyxis.macrogoltabida GCA_001314325.1_ASM131432v1
Sphingobium.indicum.B90A GCA_000264945.2_ASM26494v2
Sphingomonas.wittichii.RW1 GCA_000016765.1_ASM1676v1
Zymomonas.mobilis.subsp.mobilis.ATCC.31821 GCA_000007105.1_ASM710v1
alpha.proteobacterium.AAP81b GCA_001295935.1_ASM129593v1
Kordiimonas.gwangyangensis.DSM.19435 GCA_000375545.1_ASM37554v1
Sneathiella.glossodoripedis.JCM.23214 GCA_000616095.1_ASM61609v1
Geminicoccus.roseus.DSM.18922 GCA_000427665.1_ASM42766v1
Tistrella.mobilis.KA081020.065 GCA_000264455.2_ASM26445v2
Caenispirillum.salinarum.AK4 GCA_000315795.1
Pararhodospirillum.photometricum.DSM.122 GCA_000284415.2_ASM28441v2
Rhodospirillum.rubrum.F11 GCA_000225955.1_ASM22595v1
Roseospirillum.parvum GCA_900100455.1
Magnetospirillum.magneticum.AMB.1 GCA_000009985.1_ASM998v1
Terasakiella.pusilla.DSM.6293 GCA_000688235.1_ASM68823v1
Thalassospira.profundimaris.WP0211 GCA_000300275.1_ASM30027v1
GCA_002306855.1_ASM230685v1*
GCA_002471575.1_ASM247157v1*
Candidatus.Puniceispirillum.marinum.IMCC1322 GCA_000024465.1_ASM2446v1
Thalassobaculum.salexigens.DSM.19539 GCA_000423805.1_ASM42380v1
alpha.proteobacterium.BAL199 GCA_000171835.1_ASM17183v1
Oceanibaculum.indicum GCA_003633955.1_ASM363395v1
Kiloniella.laminariae.DSM.19542 GCA_000374005.1_ASM37400v1
Rhodovibrio.salinarum.DSM.9154 GCA_000515255.1_ASM51525v1
Inquilinus.limosus.DSM.16000 GCA_000423185.1_ASM42318v1
Rhodospirillum.centenum GCA_014197805.1_ASM1419780v1
Micavibrio.aeruginosavorus.ARL.13 GCA_000226315.1_ASM22631v1
Caedimonas.varicaedens GCA_001192655.1_ASM119265v1
Candidatus.Caedibacter.acanthamoebae GCA_000743035.1_ASM74303v1
Candidatus.Nucleicultrix.amoebiphila.FS5 GCA_002117145.1_ASM211714v1
Candidatus.Finniella.inopinata GCA_004210305.1_ASM421030v1
Candidatus.Odyssella.thessalonicensis.L13 GCA_000190415.2_ASM19041v1
Candidatus.Paracaedibacter.acanthamoebae GCA_000742835.1_ASM74283v1
Candidatus.Hepatobacter.penaei GCA_000742475.1_ASM74247v1
Holospora.undulata.HU1 GCA_000388175.3_ASM38817v3
Elioraea.tepidiphila.DSM.17972 GCA_000378465.1_ASM37846v1
Belnapia.moabensis.DSM.16746 GCA_000745835.1_ASM74583v1
Rubritepida.flocculans.DSM.14296 GCA_000425365.1_ASM42536v1
Roseomonas.gilardii GCA_001941945.1_ASM194194v1
Acidiphilium.multivorum.AIU301 GCA_000202835.1_ASM20283v1
Acidisphaera.rubrifaciens.HS.AP3 GCA_000964365.1_ASM96436v1
Granulibacter.bethesdensis.CGDNIH4 GCA_000576145.2_ASM57614v2
Commensalibacter.intestini.A911 GCA_000231445.2_ASM23144v1
Gluconacetobacter.diazotrophicus.PA1.5 GCA_000067045.1_ASM6704v1
Acetobacter.aceti.NBRC.14818 GCA_000193495.2_ASM19349v2
Gluconobacter.oxydans.DSM.3504 GCA_000583855.1_ASM58385v1
0.500042

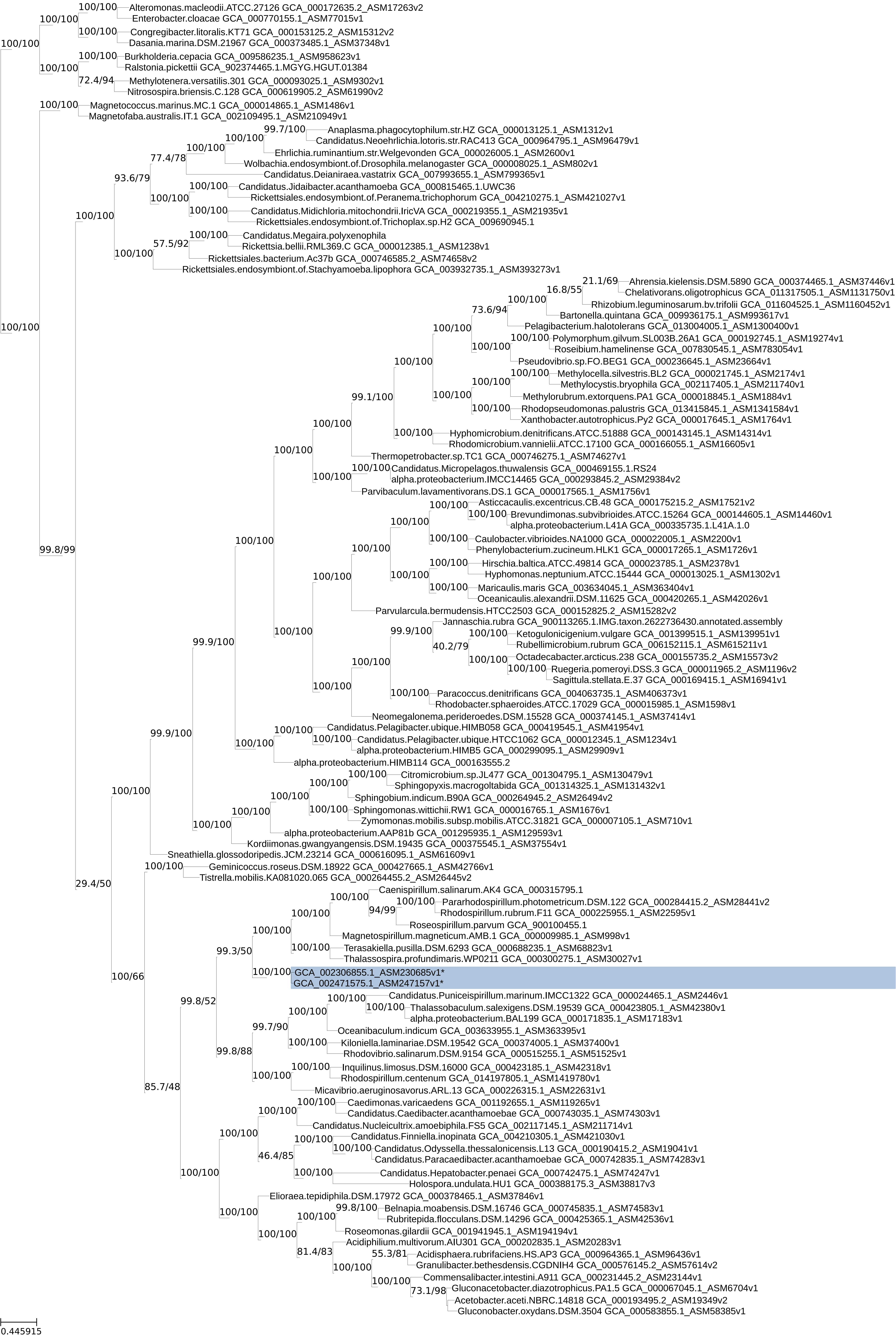
100/100
Alteromonas.macleodii.ATCC.27126 GCA_000172635.2_ASM17263v2
100/100
Enterobacter.cloacae GCA_000770155.1_ASM77015v1
100/100
Congregibacter.litoralis.KT71 GCA_000153125.2_ASM15312v2
100/100
Dasania.marina.DSM.21967 GCA_000373485.1_ASM37348v1
100/100
Burkholderia.cepacia GCA_009586235.1_ASM958623v1
100/100
Ralstonia.pickettii GCA_902374465.1.MGYG.HGUT.01384
72.4/94
Methylotenera.versatilis.301 GCA_000093025.1_ASM9302v1
Nitrosospira.briensis.C.128 GCA_000619905.2_ASM61990v2
100/100
Magnetococcus.marinus.MC.1 GCA_000014865.1_ASM1486v1
Magnetofaba.australis.IT.1 GCA_002109495.1_ASM210949v1
99.7/100
Anaplasma.phagocytophilum.str.HZ GCA_000013125.1_ASM1312v1
100/100
Candidatus.Neoehrlichia.lotoris.str.RAC413 GCA_000964795.1_ASM96479v1
100/100
Ehrlichia.ruminantium.str.Welgevonden GCA_000026005.1_ASM2600v1
77.4/78
Wolbachia.endosymbiont.of.Drosophila.melanogaster GCA_000008025.1_ASM802v1
Candidatus.Deianiraea.vastatrix GCA_007993655.1_ASM799365v1
93.6/79
100/100
Candidatus.Jidaibacter.acanthamoeba GCA_000815465.1.UWC36
100/100
Rickettsiales.endosymbiont.of.Peranema.trichophorum GCA_004210275.1_ASM421027v1
100/100
Candidatus.Midichloria.mitochondrii.IricVA GCA_000219355.1_ASM21935v1
100/100
Rickettsiales.endosymbiont.of.Trichoplax.sp.H2 GCA_009690945.1
100/100
Candidatus.Megaira.polyxenophila
57.5/92
Rickettsia.bellii.RML369.C GCA_000012385.1_ASM1238v1
100/100
Rickettsiales.bacterium.Ac37b GCA_000746585.2_ASM74658v2
Rickettsiales.endosymbiont.of.Stachyamoeba.lipophora GCA_003932735.1_ASM393273v1
21.1/69
Ahrensia.kielensis.DSM.5890 GCA_000374465.1_ASM37446v1
16.8/55
Chelativorans.oligotrophicus GCA_011317505.1_ASM1131750v1
100/100
Rhizobium.leguminosarum.bv.trifolii GCA_011604525.1_ASM1160452v1
73.6/94
Bartonella.quintana GCA_009936175.1_ASM993617v1
100/100
Pelagibacterium.halotolerans GCA_013004005.1_ASM1300400v1
100/100
Polymorphum.gilvum.SL003B.26A1 GCA_000192745.1_ASM19274v1
100/100
Roseibium.hamelinense GCA_007830545.1_ASM783054v1
100/100
Pseudovibrio.sp.FO.BEG1 GCA_000236645.1_ASM23664v1
100/100
Methylocella.silvestris.BL2 GCA_000021745.1_ASM2174v1
100/100
Methylocystis.bryophila GCA_002117405.1_ASM211740v1
99.1/100
100/100
Methylorubrum.extorquens.PA1 GCA_000018845.1_ASM1884v1
100/100
Rhodopseudomonas.palustris GCA_013415845.1_ASM1341584v1
Xanthobacter.autotrophicus.Py2 GCA_000017645.1_ASM1764v1
100/100
100/100
Hyphomicrobium.denitrificans.ATCC.51888 GCA_000143145.1_ASM14314v1
Rhodomicrobium.vannielii.ATCC.17100 GCA_000166055.1_ASM16605v1
100/100
Thermopetrobacter.sp.TC1 GCA_000746275.1_ASM74627v1
100/100
Candidatus.Micropelagos.thuwalensis GCA_000469155.1.RS24
100/100
alpha.proteobacterium.IMCC14465 GCA_000293845.2_ASM29384v2
Parvibaculum.lavamentivorans.DS.1 GCA_000017565.1_ASM1756v1
100/100
Asticcacaulis.excentricus.CB.48 GCA_000175215.2_ASM17521v2
100/100
100/100
Brevundimonas.subvibrioides.ATCC.15264 GCA_000144605.1_ASM14460v1
alpha.proteobacterium.L41A GCA_000335735.1_L41A.1.0
100/100
100/100
Caulobacter.vibrioides.NA1000 GCA_000022005.1_ASM2200v1
100/100
Phenylobacterium.zucineum.HLK1 GCA_000017265.1_ASM1726v1
100/100
Hirschia.baltica.ATCC.49814 GCA_000023785.1_ASM2378v1
100/100
100/100
Hyphomonas.neptunium.ATCC.15444 GCA_000013025.1_ASM1302v1
100/100
Maricaulis.maris GCA_003634045.1_ASM363404v1
Oceanicaulis.alexandrii.DSM.11625 GCA_000420265.1_ASM42026v1
Parvularcula.bermudensis.HTCC2503 GCA_000152825.2_ASM15282v2
Jannaschia.rubra GCA_900113265.1.IMG.taxon.2622736430.annotated.assembly
99.9/100
100/100
Ketogulonicigenium.vulgare GCA_001399515.1_ASM139951v1
100/100
40.2/79
Rubellimicrobium.rubrum GCA_006152115.1_ASM615211v1
99.9/100
Octadecabacter.arcticus.238 GCA_000155735.2_ASM15573v2
100/100
100/100
Ruegeria.pomeroyi.DSS.3 GCA_000011965.2_ASM1196v2
Sagittula.stellata.E.37 GCA_000169415.1_ASM16941v1
100/100
100/100
Paracoccus.denitrificans GCA_004063735.1_ASM406373v1
Rhodobacter.sphaeroides.ATCC.17029 GCA_000015985.1_ASM1598v1
Neomegalonema.perideroedes.DSM.15528 GCA_000374145.1_ASM37414v1
99.9/100
100/100
Candidatus.Pelagibacter.ubique.HIMB058 GCA_000419545.1_ASM41954v1
100/100
Candidatus.Pelagibacter.ubique.HTCC1062 GCA_000012345.1_ASM1234v1
100/100
alpha.proteobacterium.HIMB5 GCA_000299095.1_ASM29909v1
alpha.proteobacterium.HIMB114 GCA_000163555.2
100/100
Citromicrobium.sp.JL477 GCA_001304795.1_ASM130479v1
100/100
Sphingopyxis.macrogoltabida GCA_001314325.1_ASM131432v1
100/100
100/100
Sphingobium.indicum.B90A GCA_000264945.2_ASM26494v2
100/100
Sphingomonas.wittichii.RW1 GCA_000016765.1_ASM1676v1
100/100
Zymomonas.mobilis.subsp.mobilis.ATCC.31821 GCA_000007105.1_ASM710v1
99.8/99
alpha.proteobacterium.AAP81b GCA_001295935.1_ASM129593v1
Kordiimonas.gwangyangensis.DSM.19435 GCA_000375545.1_ASM37554v1
Sneathiella.glossodoripedis.JCM.23214 GCA_000616095.1_ASM61609v1
100/100
Geminicoccus.roseus.DSM.18922 GCA_000427665.1_ASM42766v1
Tistrella.mobilis.KA081020.065 GCA_000264455.2_ASM26445v2
29.4/50
Caenispirillum.salinarum.AK4 GCA_000315795.1
100/100
100/100
Pararhodospirillum.photometricum.DSM.122 GCA_000284415.2_ASM28441v2
94/99
Rhodospirillum.rubrum.F11 GCA_000225955.1_ASM22595v1
100/100
Roseospirillum.parvum GCA_900100455.1
100/100
Magnetospirillum.magneticum.AMB.1 GCA_000009985.1_ASM998v1
100/100
Terasakiella.pusilla.DSM.6293 GCA_000688235.1_ASM68823v1
99.3/50
Thalassospira.profundimaris.WP0211 GCA_000300275.1_ASM30027v1
100/100
GCA_002306855.1_ASM230685v1*
100/66
GCA_002471575.1_ASM247157v1*
Candidatus.Puniceispirillum.marinum.IMCC1322 GCA_000024465.1_ASM2446v1
100/100
99.8/52
100/100
Thalassobaculum.salexigens.DSM.19539 GCA_000423805.1_ASM42380v1
100/100
alpha.proteobacterium.BAL199 GCA_000171835.1_ASM17183v1
99.7/90
Oceanibaculum.indicum GCA_003633955.1_ASM363395v1
100/100
Kiloniella.laminariae.DSM.19542 GCA_000374005.1_ASM37400v1
99.8/88
Rhodovibrio.salinarum.DSM.9154 GCA_000515255.1_ASM51525v1
100/100
Inquilinus.limosus.DSM.16000 GCA_000423185.1_ASM42318v1
100/100
Rhodospirillum.centenum GCA_014197805.1_ASM1419780v1
85.7/48
Micavibrio.aeruginosavorus.ARL.13 GCA_000226315.1_ASM22631v1
100/100
Caedimonas.varicaedens GCA_001192655.1_ASM119265v1
100/100
Candidatus.Caedibacter.acanthamoebae GCA_000743035.1_ASM74303v1
100/100
Candidatus.Nucleicultrix.amoebiphila.FS5 GCA_002117145.1_ASM211714v1
100/100
Candidatus.Finniella.inopinata GCA_004210305.1_ASM421030v1
100/100
Candidatus.Odyssella.thessalonicensis.L13 GCA_000190415.2_ASM19041v1
46.4/85
Candidatus.Paracaedibacter.acanthamoebae GCA_000742835.1_ASM74283v1
100/100
100/100
Candidatus.Hepatobacter.penaei GCA_000742475.1_ASM74247v1
Holospora.undulata.HU1 GCA_000388175.3_ASM38817v3
Elioraea.tepidiphila.DSM.17972 GCA_000378465.1_ASM37846v1
99.8/100
Belnapia.moabensis.DSM.16746 GCA_000745835.1_ASM74583v1
100/100
100/100
Rubritepida.flocculans.DSM.14296 GCA_000425365.1_ASM42536v1
Roseomonas.gilardii GCA_001941945.1_ASM194194v1
100/100
Acidiphilium.multivorum.AIU301 GCA_000202835.1_ASM20283v1
81.4/83
55.3/81
Acidisphaera.rubrifaciens.HS.AP3 GCA_000964365.1_ASM96436v1
100/100
Granulibacter.bethesdensis.CGDNIH4 GCA_000576145.2_ASM57614v2
Commensalibacter.intestini.A911 GCA_000231445.2_ASM23144v1
100/100
73.1/98
Gluconacetobacter.diazotrophicus.PA1.5 GCA_000067045.1_ASM6704v1
Acetobacter.aceti.NBRC.14818 GCA_000193495.2_ASM19349v2
Gluconobacter.oxydans.DSM.3504 GCA_000583855.1_ASM58385v1
0.445915

### Group C

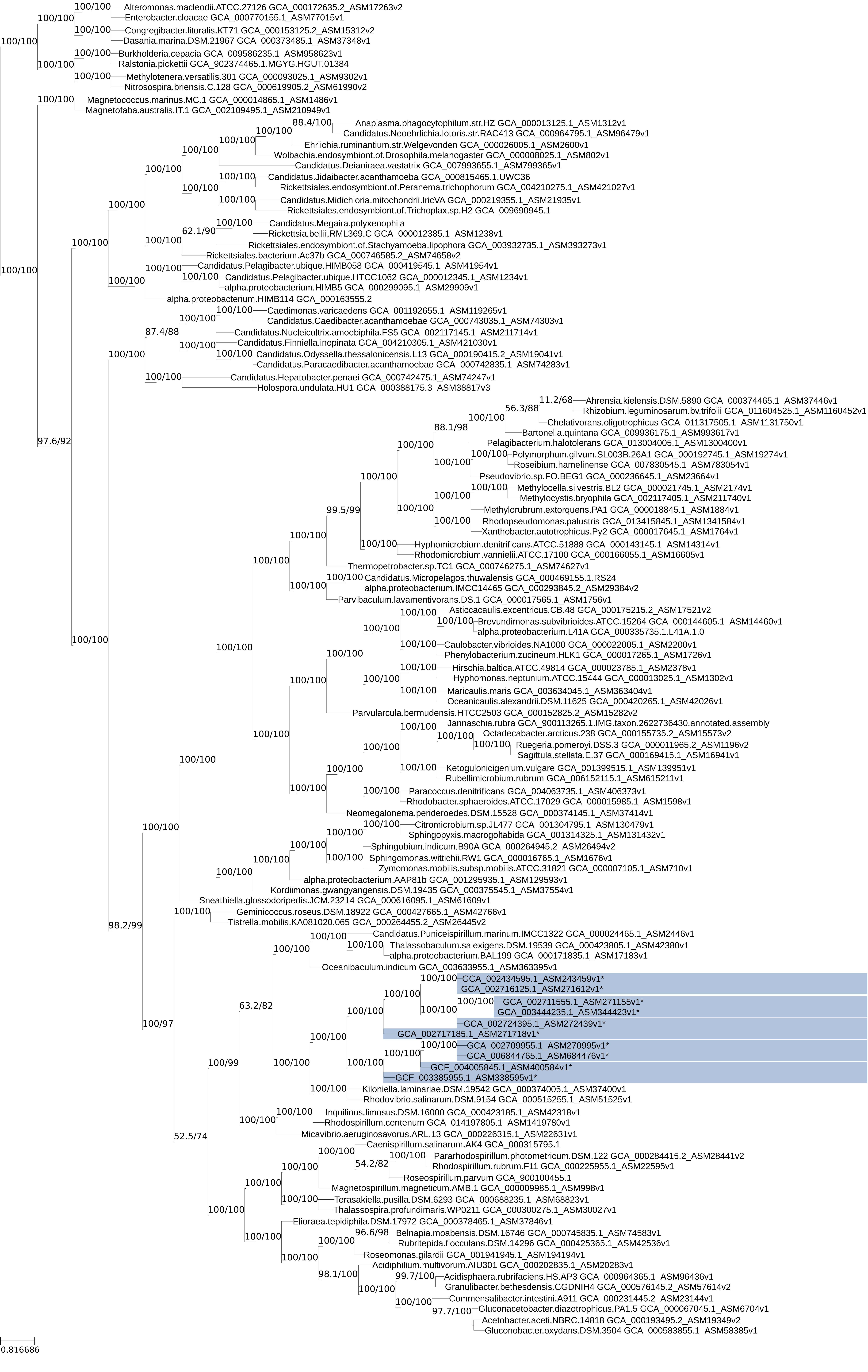
Alteromonas.macleodii.ATCC.27126 GCA_000172635.2_ASM17263v2
Enterobacter.cloacae GCA_000770155.1_ASM77015v1
Congregibacter.litoralis.KT71 GCA_000153125.2_ASM15312v2
Dasania.marina.DSM.21967 GCA_000373485.1_ASM37348v1
Burkholderia.cepacia GCA_009586235.1_ASM958623v1
Ralstonia.pickettii GCA_902374465.1.MGYG.HGUT.01384
Methylotenera.versatilis.301 GCA_000093025.1_ASM9302v1
Nitrosospira.briensis.C.128 GCA_000619905.2_ASM61990v2
Magnetococcus.marinus.MC.1 GCA_000014865.1_ASM1486v1
Magnetofaba.australis.IT.1 GCA_002109495.1_ASM210949v1
Anaplasma.phagocytophilum.str.HZ GCA_000013125.1_ASM1312v1
Candidatus.Neoehrlichia.lotoris.str.RAC413 GCA_000964795.1_ASM96479v1
Ehrlichia.ruminantium.str.Welgevonden GCA_000026005.1_ASM2600v1
Wolbachia.endosymbiont.of.Drosophila.melanogaster GCA_000008025.1_ASM802v1
Candidatus.Deianiraea.vastatrix GCA_007993655.1_ASM799365v1
Candidatus.Jidaibacter.acanthamoeba GCA_000815465.1.UWC36
Rickettsiales.endosymbiont.of.Peranema.trichophorum GCA_004210275.1_ASM421027v1
Candidatus.Midichloria.mitochondrii.IricVA GCA_000219355.1_ASM21935v1
Rickettsiales.endosymbiont.of.Trichoplax.sp.H2 GCA_009690945.1
Candidatus.Megaira.polyxenophila
Rickettsia.bellii.RML369.C GCA_000012385.1_ASM1238v1
Rickettsiales.endosymbiont.of.Stachyamoeba.lipophora GCA_003932735.1_ASM393273v1
Rickettsiales.bacterium.Ac37b GCA_000746585.2_ASM74658v2
Candidatus.Pelagibacter.ubique.HIMB058 GCA_000419545.1_ASM41954v1
Candidatus.Pelagibacter.ubique.HTCC1062 GCA_000012345.1_ASM1234v1
alpha.proteobacterium.HIMB5 GCA_000299095.1_ASM29909v1
alpha.proteobacterium.HIMB114 GCA_000163555.2
Caedimonas.varicaedens GCA_001192655.1_ASM119265v1
Candidatus.Caedibacter.acanthamoebae GCA_000743035.1_ASM74303v1
Candidatus.Nucleicultrix.amoebiphila.FS5 GCA_002117145.1_ASM211714v1
Candidatus.Finniella.inopinata GCA_004210305.1_ASM421030v1
Candidatus.Odyssella.thessalonicensis.L13 GCA_000190415.2_ASM19041v1
Candidatus.Paracaedibacter.acanthamoebae GCA_000742835.1_ASM74283v1
Candidatus.Hepatobacter.penaei GCA_000742475.1_ASM74247v1
Holospora.undulata.HU1 GCA_000388175.3_ASM38817v3
Ahrensia.kielensis.DSM.5890 GCA_000374465.1_ASM37446v1
Rhizobium.leguminosarum.bv.trifolii GCA_011604525.1_ASM1160452v1
Chelativorans.oligotrophicus GCA_011317505.1_ASM1131750v1
Bartonella.quintana GCA_009936175.1_ASM993617v1
Pelagibacterium.halotolerans GCA_013004005.1_ASM1300400v1
Polymorphum.gilvum.SL003B.26A1 GCA_000192745.1_ASM19274v1
Roseibium.hamelinense GCA_007830545.1_ASM783054v1
Pseudovibrio.sp.FO.BEG1 GCA_000236645.1_ASM23664v1
Methylocella.silvestris.BL2 GCA_000021745.1_ASM2174v1
Methylocystis.bryophila GCA_002117405.1_ASM211740v1
Methylorubrum.extorquens.PA1 GCA_000018845.1_ASM1884v1
Rhodopseudomonas.palustris GCA_013415845.1_ASM1341584v1
Xanthobacter.autotrophicus.Py2 GCA_000017645.1_ASM1764v1
Hyphomicrobium.denitrificans.ATCC.51888 GCA_000143145.1_ASM14314v1
Rhodomicrobium.vannielii.ATCC.17100 GCA_000166055.1_ASM16605v1
Thermopetrobacter.sp.TC1 GCA_000746275.1_ASM74627v1
Candidatus.Micropelagos.thuwalensis GCA_004469155.1.RS24
alpha.proteobacterium.IMCC14465 GCA_000293845.2_ASM29384v2
Parvibaculum.lavamentivorans.DS.1 GCA_000017565.1_ASM1756v1
Asticcacaulis.excentricus.CB.48 GCA_000175215.2_ASM17521v2
Brevundimonas.subvibrioides.ATCC.15264 GCA_000144605.1_ASM14460v1
alpha.proteobacterium.L41A GCA_000335735.1.L41A.1.0
Caulobacter.vibrioides.NA1000 GCA_000022005.1_ASM2200v1
Phenylobacterium.zucineum.HLK1 GCA_000017265.1_ASM1726v1
Hirschia.baltica.ATCC.49814 GCA_000023785.1_ASM2378v1
Hyphomonas.neptunium.ATCC.15444 GCA_000013025.1_ASM1302v1
Maricaulis.maris GCA_003634045.1_ASM363404v1
Oceanicaulis.alexandrii.DSM.11625 GCA_000420265.1_ASM42026v1
Parvularcula.bermudensis.HTCC2503 GCA_000152825.2_ASM15282v2
Jannaschia.rubra GCA_900113265.1.IMG.taxon.2622736430.annotated.assembly
Octadecabacter.arcticus.238 GCA_000155735.2_ASM15573v2
Ruegeria.pomeroyi.DSS.3 GCA_000011965.2_ASM1196v2
Sagittula.stellata.E.37 GCA_000169415.1_ASM16941v1
Ketogulonicigenium.vulgare GCA_001399515.1_ASM139951v1
Rubellimicrobium.rubrum GCA_006152115.1_ASM615211v1
Paracoccus.denitrificans GCA_004063735.1_ASM406373v1
Rhodobacter.sphaeroides.ATCC.17029 GCA_000015985.1_ASM1598v1
Neomegalonema.perideroedes.DSM.15528 GCA_000374145.1_ASM37414v1
Citromicrobium.sp.JL477 GCA_001304795.1_ASM130479v1
Sphingopyxis.macrogoltabida GCA_001314325.1_ASM131432v1
Sphingobium.indicum.B90A GCA_000264945.2_ASM26494v2
Sphingomonas.wittichii.RW1 GCA_000016765.1_ASM1676v1
Zymomonas.mobilis.subsp.mobilis.ATCC.31821 GCA_000007105.1_ASM710v1
alpha.proteobacterium.AAP81b GCA_001295935.1_ASM129593v1
Kordiimonas.gwangyangensis.DSM.19435 GCA_000375545.1_ASM37554v1
Sneathiella.glossodoripedis.JCM.23214 GCA_000616095.1_ASM61609v1
Geminicoccus.roseus.DSM.18922 GCA_000427665.1_ASM42766v1
Tistrella.mobilis.KA081020.065 GCA_000264455.2_ASM26445v2
Candidatus.Puniceispirillum.marinum.IMCC1322 GCA_000024465.1_ASM2446v1
Thalassobaculum.salexigens.DSM.19539 GCA_000423805.1_ASM42380v1
alpha.proteobacterium.BAL199 GCA_000171835.1_ASM17183v1
Oceanibaculum.indicum GCA_003633955.1_ASM363395v1
GCA_002434595.1_ASM243459v1*
GCA_002716125.1_ASM271612v1*
GCA_002711555.1_ASM271155v1*
GCA_003444235.1_ASM344423v1*
GCA_002724395.1_ASM272439v1*
GCA_002717185.1_ASM271718v1*
GCA_002709955.1_ASM270995v1*
GCA_006844765.1_ASM684476v1*
GCF_004005845.1_ASM400584v1*
GCF_003385955.1_ASM338595v1*
Kiloniella.laminariae.DSM.19542 GCA_000374005.1_ASM37400v1
Rhodovibrio.salinarum.DSM.9154 GCA_000515255.1_ASM51525v1
Inquilinus.limosus.DSM.16000 GCA_000423185.1_ASM42318v1
Rhodospirillum.centenum GCA_014197805.1_ASM1419780v1
Micavibrio.aeruginosavorus.ARL.13 GCA_000226315.1_ASM22631v1
Caenispirillum.salinarum.AK4 GCA_000315795.1
Pararhodospirillum.photometricum.DSM.122 GCA_000284415.2_ASM28441v2
Rhodospirillum.rubrum.F11 GCA_000225955.1_ASM22595v1
Roseospirillum.parvum GCA_900100455.1
Magnetospirillum.magneticum.AMB.1 GCA_000009985.1_ASM998v1
Terasakiella.pusilla.DSM.6293 GCA_000688235.1_ASM68823v1
Thalassospira.profundimaris.WP0211 GCA_000300275.1_ASM30027v1
Elioraea.tepidiphila.DSM.17972 GCA_000378465.1_ASM37846v1
Belnapia.moabensis.DSM.16746 GCA_000745835.1_ASM74583v1
Rubritepida.flocculans.DSM.14296 GCA_000425365.1_ASM42536v1
Roseomonas.gilardii GCA_001941945.1_ASM194194v1
Acidiphilium.multivorum.AIU301 GCA_000202835.1_ASM20283v1
Acidisphaera.rubrifaciens.HS.AP3 GCA_000964365.1_ASM96436v1
Granulibacter.bethesdensis.CGDNIH4 GCA_000576145.2_ASM57614v2
Commensalibacter.intestini.A911 GCA_000231445.2_ASM23144v1
Gluconacetobacter.diazotrophicus.PA1.5 GCA_000067045.1_ASM6704v1
Acetobacter.aceti.NBRC.14818 GCA_000193495.2_ASM19349v2
Gluconobacter.oxydans.DSM.3504 GCA_000583855.1_ASM58385v1
0.816686

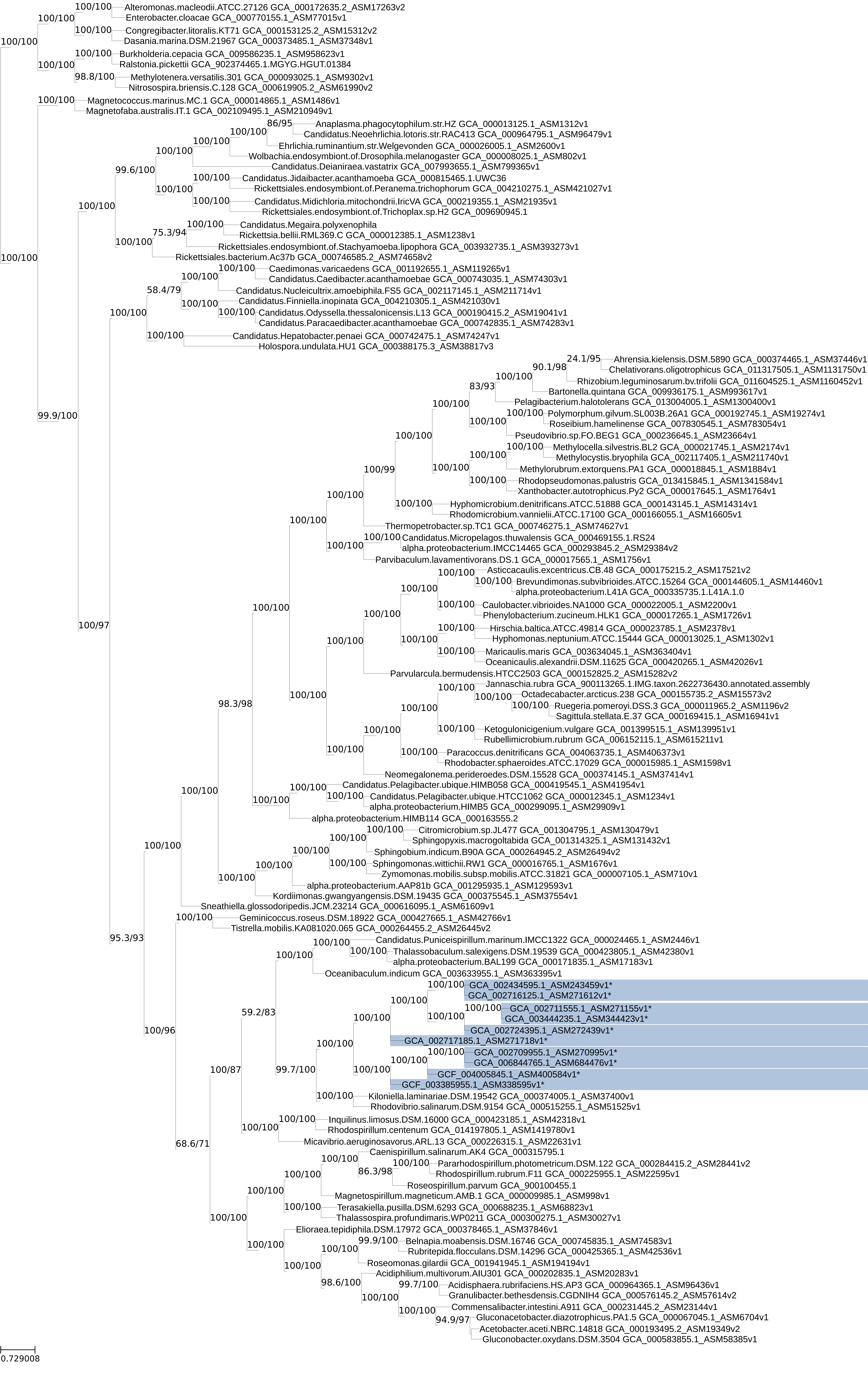
Alteromonas.macleodii.ATCC.27126 GCA_000172635.2_ASM17263v2
Enterobacter.cloacae GCA_000770155.1_ASM77015v1
Congregibacter.litoralis.KT71 GCA_000153125.2_ASM15312v2
Dasania.marina.DSM.21967 GCA_000373485.1_ASM37348v1
Burkholderia.cepacia GCA_009586235.1_ASM958623v1
Ralstonia.pickettii GCA_902374465.1.MGYG.HGUT.01384
Methylotenera.versatilis.301 GCA_000093025.1_ASM9302v1
Nitrosospira.briensis.C.128 GCA_000619905.2_ASM61990v2
Magnetococcus.marinus.MC.1 GCA_000014865.1_ASM1486v1
Magnetofaba.australis.IT.1 GCA_002109495.1_ASM210949v1
Anaplasma.phagocytophilum.str.HZ GCA_000013125.1_ASM1312v1
Candidatus.Neoehrlichia.lotoris.str.RAC413 GCA_000964795.1_ASM96479v1
Ehrlichia.ruminantium.str.Welgevonden GCA_000026005.1_ASM2600v1
Wolbachia.endosymbiont.of.Drosophila.melanogaster GCA_000008025.1_ASM802v1
Candidatus.Deianiraea.vastatrix GCA_007993655.1_ASM799365v1
Candidatus.Jidaibacter.acanthamoeba GCA_000815465.1.UWC36
Rickettsiales.endosymbiont.of.Peranema.trichophorum GCA_004210275.1_ASM421027v1
Candidatus.Midichloria.mitochondrii.IricVA GCA_000219355.1_ASM21935v1
Rickettsiales.endosymbiont.of.Trichoplax.sp.H2 GCA_009690945.1
Candidatus.Megaira.polyxenophila
Rickettsia.bellii.RML369.C GCA_000012385.1_ASM1238v1
Rickettsiales.endosymbiont.of.Stachyamoeba.lipophora GCA_003932735.1_ASM393273v1
Rickettsiales.bacterium.Ac37b GCA_000746585.2_ASM74658v2
Caedimonas.varicaedens GCA_001192655.1_ASM119265v1
Candidatus.Caedibacter.acanthamoebae GCA_000743035.1_ASM74303v1
Candidatus.Nucleicultrix.amoebiphila.FS5 GCA_002117145.1_ASM211714v1
Candidatus.Finniella.inopinata GCA_004210305.1_ASM421030v1
Candidatus.Odyssella.thessalonicensis.L13 GCA_000190415.2_ASM19041v1
Candidatus.Paracaedibacter.acanthamoebae GCA_000742835.1_ASM74283v1
Candidatus.Hepatobacter.penaei GCA_000742475.1_ASM74247v1
Holospora.undulata.HU1 GCA_000388175.3_ASM38817v3
Ahrensia.kielensis.DSM.5890 GCA_000374465.1_ASM37446v1
Chelativorans.oligotrophicus GCA_011317505.1_ASM1131750v1
Rhizobium.leguminosarum.bv.trifolii GCA_011604525.1_ASM1160452v1
Bartonella.quintana GCA_009936175.1_ASM993617v1
Pelagibacterium.halotolerans GCA_013004005.1_ASM1300400v1
Polymorphum.gilvum.SL003B.26A1 GCA_000192745.1_ASM19274v1
Roseibium.hamelinense GCA_007830545.1_ASM783054v1
Pseudovibrio.sp.FO.BEG1 GCA_000236645.1_ASM23664v1
Methylocella.silvestris.BL2 GCA_000021745.1_ASM2174v1
Methylocystis.bryophila GCA_002117405.1_ASM211740v1
Methylorubrum.extorquens.PA1 GCA_000018845.1_ASM1884v1
Rhodopseudomonas.palustris GCA_013415845.1_ASM1341584v1
Xanthobacter.autotrophicus.Py2 GCA_000017645.1_ASM1764v1
Hyphomicrobium.denitrificans.ATCC.51888 GCA_000143145.1_ASM14314v1
Rhodomicrobium.vannielii.ATCC.17100 GCA_000166055.1_ASM16605v1
Thermopetrobacter.sp.TC1 GCA_000746275.1_ASM74627v1
Candidatus.Micropelagos.thuwalensis GCA_000469155.1.RS24
alpha.proteobacterium.IMCC14465 GCA_000293845.2_ASM29384v2
Parvibaculum.lavamentivorans.DS.1 GCA_000017565.1_ASM1756v1
Asticcacaulis.excentricus.CB.48 GCA_000175215.2_ASM17521v2
Brevundimonas.subvibrioides.ATCC.15264 GCA_000144605.1_ASM14460v1
alpha.proteobacterium.L41A GCA_000335735.1.L41A.1.0
Caulobacter.vibrioides.NA1000 GCA_000022005.1_ASM2200v1
Phenylobacterium.zucineum.HLK1 GCA_000017265.1_ASM1726v1
Hirschia.baltica.ATCC.49814 GCA_000023785.1_ASM2378v1
Hyphomonas.neptunium.ATCC.15444 GCA_000013025.1_ASM1302v1
Maricaulis.maris GCA_003634045.1_ASM363404v1
Oceanicaulis.alexandrii.DSM.11625 GCA_000420265.1_ASM42026v1
Parvularcula.bermudensis.HTCC2503 GCA_000152825.2_ASM15282v2
Jannaschia.rubra GCA_900113265.1.IMG.taxon.2622736430.annotated.assembly
Octadecabacter.arcticus.238 GCA_000155735.2_ASM15573v2
Ruegeria.pomeroyi.DSS.3 GCA_000011965.2_ASM1196v2
Sagittula.stellata.E.37 GCA_000169415.1_ASM16941v1
Ketogulonicigenium.vulgare GCA_001399515.1_ASM139951v1
Rubellimicrobium.rubrum GCA_006152115.1_ASM615211v1
Paracoccus.denitrificans GCA_004063735.1_ASM406373v1
Rhodobacter.sphaeroides.ATCC.17029 GCA_000015985.1_ASM1598v1
Neomegalonema.perideroedes.DSM.15528 GCA_000374145.1_ASM37414v1
Candidatus.Pelagibacter.ubique.HIMB058 GCA_000419545.1_ASM41954v1
Candidatus.Pelagibacter.ubique.HTCC1062 GCA_000012345.1_ASM1234v1
alpha.proteobacterium.HIMB5 GCA_000299095.1_ASM29909v1
alpha.proteobacterium.HIMB114 GCA_000163555.2
Citromicrobium.sp.JL477 GCA_001304795.1_ASM130479v1
Sphingopyxis.macrogoltabida GCA_001314325.1_ASM131432v1
Sphingobium.indicum.B90A GCA_000264945.2_ASM26494v2
Sphingomonas.wittichii.RW1 GCA_000016765.1_ASM1676v1
Zymomonas.mobilis.subsp.mobilis.ATCC.31821 GCA_000007105.1_ASM710v1
alpha.proteobacterium.AAP81b GCA_001295935.1_ASM129593v1
Kordiimonas.gwangyangensis.DSM.19435 GCA_000375545.1_ASM37554v1
Sneathiella.glossodoripedis.JCM.23214 GCA_000616095.1_ASM61609v1
Geminicoccus.roseus.DSM.18922 GCA_000427665.1_ASM42766v1
Tistrella.mobilis.KA081020.065 GCA_000264455.2_ASM26445v2
Candidatus.Puniceispirillum.marinum.IMCC1322 GCA_000024465.1_ASM2446v1
Thalassobaculum.salexigens.DSM.19539 GCA_000423805.1_ASM42380v1
alpha.proteobacterium.BAL199 GCA_000171835.1_ASM17183v1
Oceanibaculum.indicum GCA_003633955.1_ASM363395v1
GCA_002434595.1_ASM243459v1*
GCA_002716125.1_ASM271612v1*
GCA_002711555.1_ASM271155v1*
GCA_003444235.1_ASM344423v1*
GCA_002724395.1_ASM272439v1*
GCA_002717185.1_ASM271718v1*
GCA_002709955.1_ASM270995v1*
GCA_006844765.1_ASM684476v1*
GCF_004005845.1_ASM400584v1*
GCF_003385955.1_ASM338595v1*
Kiloniella.laminariae.DSM.19542 GCA_000374005.1_ASM37400v1
Rhodovibrio.salinarum.DSM.9154 GCA_000515255.1_ASM51525v1
Inquilinus.limosus.DSM.16000 GCA_000423185.1_ASM42318v1
Rhodospirillum.centenum GCA_014197805.1_ASM1419780v1
Micavibrio.aeruginosavorus.ARL.13 GCA_000226315.1_ASM22631v1
Caenispirillum.salinarum.AK4 GCA_000315795.1
Pararhodospirillum.photometricum.DSM.122 GCA_000284415.2_ASM28441v2
Rhodospirillum.rubrum.F11 GCA_000225955.1_ASM22595v1
Roseospirillum.parvum GCA_900100455.1
Magnetospirillum.magneticum.AMB.1 GCA_000009985.1_ASM998v1
Terasakiella.pusilla.DSM.6293 GCA_000688235.1_ASM68823v1
Thalassospira.profundimaris.WP0211 GCA_000300275.1_ASM30027v1
Elioraea.tepidiphila.DSM.17972 GCA_000378465.1_ASM37846v1
Belnapia.moabensis.DSM.16746 GCA_000745835.1_ASM74583v1
Rubritepida.flocculans.DSM.14296 GCA_000425365.1_ASM42536v1
Roseomonas.gilardii GCA_001941945.1_ASM194194v1
Acidiphilium.multivorum.AIU301 GCA_000202835.1_ASM20283v1
Acidisphaera.rubrifaciens.HS.AP3 GCA_000964365.1_ASM96436v1
Granulibacter.bethesdensis.CGDNIH4 GCA_000576145.2_ASM57614v2
Commensalibacter.intestini.A911 GCA_000231445.2_ASM23144v1
Gluconacetobacter.diazotrophicus.PA1.5 GCA_000067045.1_ASM6704v1
Acetobacter.aceti.NBRC.14818 GCA_000193495.2_ASM19349v2
Gluconobacter.oxydans.DSM.3504 GCA_000583855.1_ASM58385v1
0.729008

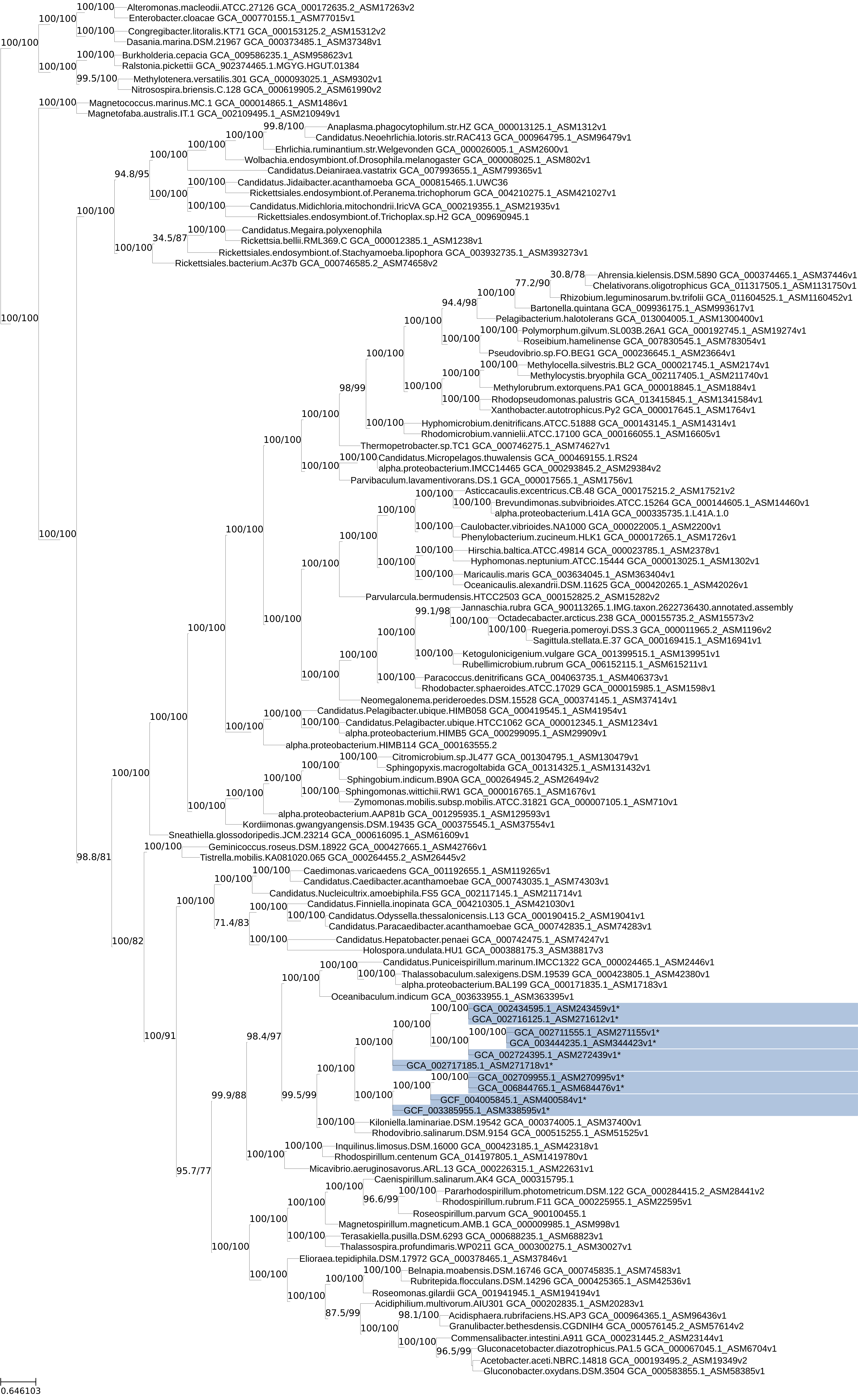

100/100
Alteromonas.macleodii.ATCC.27126 GCA_000172635.2_ASM17263v2
100/100
Enterobacter.cloacae GCA_000770155.1_ASM77015v1
100/100
100/100
Congregibacter.litoralis.KT71 GCA_000153125.2_ASM15312v2
Dasania.marina.DSM.21967 GCA_000373485.1_ASM37348v1
100/100
Burkholderia.cepacia GCA_009586235.1_ASM958623v1
100/100
Ralstonia.pickettii GCA_902374465.1.MGYG.HGUT.01384
99.5/100
Methylotenera.versatilis.301 GCA_000093025.1_ASM9302v1
Nitrosospira.briensis.C.128 GCA_000619905.2_ASM61990v2
100/100
Magnetococcus.marinus.MC.1 GCA_000014865.1_ASM1486v1
Magnetofaba.australis.IT.1 GCA_002109495.1_ASM210949v1
99.8/100
Anaplasma.phagocytophilum.str.HZ GCA_000013125.1_ASM1312v1
100/100
Candidatus.Neoehrlichia.lotoris.str.RAC413 GCA_000964795.1_ASM96479v1
100/100
Ehrlichia.ruminantium.str.Welgevonden GCA_000026005.1_ASM2600v1
100/100
Wolbachia.endosymbiont.of.Drosophila.melanogaster GCA_000008025.1_ASM802v1
Candidatus.Deianiraea.vastatrix GCA_007993655.1_ASM799365v1
94.8/95
100/100
Candidatus.Jidaibacter.acanthamoeba GCA_000815465.1.UWC36
100/100
Rickettsiales.endosymbiont.of.Peranema.trichophorum GCA_004210275.1_ASM421027v1
100/100
Candidatus.Midichloria.mitochondrii.IricVA GCA_000219355.1_ASM21935v1
100/100
Rickettsiales.endosymbiont.of.Trichoplax.sp.H2 GCA_009690945.1
100/100
Candidatus.Megaira.polyxenophila
34.5/87
Rickettsia.bellii.RML369.C GCA_000012385.1_ASM1238v1
Rickettsiales.endosymbiont.of.Stachyamoeba.lipophora GCA_003932735.1_ASM393273v1
100/100
Rickettsiales.bacterium.Ac37b GCA_000746585.2_ASM74658v2
30.8/78
Ahrensia.kielensis.DSM.5890 GCA_000374465.1_ASM37446v1
77.2/90
Chelativorans.oligotrophicus GCA_011317505.1_ASM1131750v1
100/100
Rhizobium.leguminosarum.bv.trifolii GCA_011604525.1_ASM1160452v1
100/100
Bartonella.quintana GCA_009936175.1_ASM993617v1
94.4/98
Pelagibacterium.halotolerans GCA_013004005.1_ASM1300400v1
100/100
100/100
Polymorphum.gilvum.SL003B.26A1 GCA_000192745.1_ASM19274v1
100/100
Roseibium.hamelinense GCA_007830545.1_ASM783054v1
Pseudovibrio.sp.FO.BEG1 GCA_000236645.1_ASM23664v1
100/100
100/100
Methylocella.silvestris.BL2 GCA_000021745.1_ASM2174v1
100/100
Methylocystis.bryophila GCA_002117405.1_ASM211740v1
100/100
Methylorubrum.extorquens.PA1 GCA_000018845.1_ASM1884v1
98/99
100/100
Rhodopseudomonas.palustris GCA_013415845.1_ASM1341584v1
Xanthobacter.autotrophicus.Py2 GCA_000017645.1_ASM1764v1
100/100
100/100
Hyphomicrobium.denitrificans.ATCC.51888 GCA_000143145.1_ASM14314v1
Rhodomicrobium.vannielii.ATCC.17100 GCA_000166055.1_ASM16605v1
100/100
Thermopetrobacter.sp.TC1 GCA_000746275.1_ASM74627v1
100/100
Candidatus.Micropelagos.thuwalensis GCA_000469155.1.RS24
100/100
alpha.proteobacterium.IMCC14465 GCA_000293845.2_ASM29384v2
Parvibaculum.lavamentivorans.DS.1 GCA_000017565.1_ASM1756v1
100/100
Asticcacaulis.excentricus.CB.48 GCA_000175215.2_ASM17521v2
100/100
Brevundimonas.subvibrioides.ATCC.15264 GCA_000144605.1_ASM14460v1
100/100
alpha.proteobacterium.L41A GCA_000335735.1.L41A.1.0
100/100
Caulobacter.vibrioides.NA1000 GCA_000022005.1_ASM2200v1
100/100
Phenylobacterium.zucineum.HLK1 GCA_000017265.1_ASM1726v1
100/100
Hirschia.baltica.ATCC.49814 GCA_000023785.1_ASM2378v1
100/100
100/100
Hyphomonas.neptunium.ATCC.15444 GCA_000013025.1_ASM1302v1
100/100
Maricaulis.maris GCA_003634045.1_ASM363404v1
Oceanicaulis.alexandrii.DSM.11625 GCA_000420265.1_ASM42026v1
Parvularcula.bermudensis.HTCC2503 GCA_000152825.2_ASM15282v2
99.1/98
Jannaschia.rubra GCA_900113265.1.IMG.taxon.2622736430.annotated.assembly
100/100
Octadecabacter.arcticus.238 GCA_000155735.2_ASM15573v2
100/100
Ruegeria.pomeroyi.DSS.3 GCA_000011965.2_ASM1196v2
100/100
Sagittula.stellata.E.37 GCA_000169415.1_ASM16941v1
100/100
100/100
Ketogulonicigenium.vulgare GCA_001399515.1_ASM139951v1
100/100
Rubellimicrobium.rubrum GCA_006152115.1_ASM615211v1
100/100
Paracoccus.denitrificans GCA_004063735.1_ASM406373v1
100/100
Rhodobacter.sphaeroides.ATCC.17029 GCA_000015985.1_ASM1598v1
Neomegalonema.perideroedes.DSM.15528 GCA_000374145.1_ASM37414v1
100/100
100/100
Candidatus.Pelagibacter.ubique.HIMB058 GCA_000419545.1_ASM41954v1
100/100
Candidatus.Pelagibacter.ubique.HTCC1062 GCA_000012345.1_ASM1234v1
100/100
alpha.proteobacterium.HIMB5 GCA_000299095.1_ASM29909v1
alpha.proteobacterium.HIMB114 GCA_000163555.2
100/100
Citromicrobium.sp.JL477 GCA_001304795.1_ASM130479v1
100/100
Sphingopyxis.macrogoltabida GCA_001314325.1_ASM131432v1
100/100
Sphingobium.indicum.B90A GCA_000264945.2_ASM26494v2
100/100
Sphingomonas.wittichii.RW1 GCA_000016765.1_ASM1676v1
100/100
Zymomonas.mobilis.subsp.mobilis.ATCC.31821 GCA_000007105.1_ASM710v1
100/100
alpha.proteobacterium.AAP81b GCA_001295935.1_ASM129593v1
Kordiimonas.gwangyangensis.DSM.19435 GCA_000375545.1_ASM37554v1
Sneathiella.glossodoripedis.JCM.23214 GCA_000616095.1_ASM61609v1
100/100
Geminicoccus.roseus.DSM.18922 GCA_000427665.1_ASM42766v1
98.8/81
Tistrella.mobilis.KA081020.065 GCA_000264455.2_ASM26445v2
100/100
Caedimonas.varicaedens GCA_001192655.1_ASM119265v1
100/100
Candidatus.Caedibacter.acanthamoebae GCA_000743035.1_ASM74303v1
100/100
Candidatus.Nucleicultrix.amoebiphila.FS5 GCA_002117145.1_ASM211714v1
100/100
Candidatus.Finniella.inopinata GCA_004210305.1_ASM421030v1
100/100
Candidatus.Odyssella.thessalonicensis.L13 GCA_000190415.2_ASM19041v1
71.4/83
Candidatus.Paracaedibacter.acanthamoebae GCA_000742835.1_ASM74283v1
100/100
Candidatus.Hepatobacter.penaei GCA_000742475.1_ASM74247v1
100/82
Holospora.undulata.HU1 GCA_000388175.3_ASM38817v3
100/100
Candidatus.Puniceispirillum.marinum.IMCC1322 GCA_000024465.1_ASM2446v1
100/100
Thalassobaculum.salexigens.DSM.19539 GCA_000423805.1_ASM42380v1
100/100
alpha.proteobacterium.BAL199 GCA_000171835.1_ASM17183v1
Oceanibaculum.indicum GCA_003633955.1_ASM363395v1
100/100
GCA_002434595.1_ASM243459v1*
100/100
GCA_002716125.1_ASM271612v1*
100/100
GCA_002711555.1_ASM271155v1*
100/100
GCA_003444235.1_ASM344423v1*
100/91
98.4/97
100/100
GCA_002724395.1_ASM272439v1*
GCA_002717185.1_ASM271718v1*
100/100
100/100
GCA_002709955.1_ASM270995v1*
100/100
GCA_006844765.1_ASM684476v1*
99.9/88
99.5/99
100/100
GCF_004005845.1_ASM400584v1*
GCF_003385955.1_ASM338595v1*
100/100
Kiloniella.laminariae.DSM.19542 GCA_000374005.1_ASM37400v1
Rhodovibrio.salinarum.DSM.9154 GCA_000515255.1_ASM51525v1
100/100
Inquilinus.limosus.DSM.16000 GCA_000423185.1_ASM42318v1
100/100
Rhodospirillum.centenum GCA_014197805.1_ASM1419780v1
Micavibrio.aeruginosavorus.ARL.13 GCA_000226315.1_ASM22631v1
95.7/77
100/100
Caenispirillum.salinarum.AK4 GCA_000315795.1
100/100
Pararhodospirillum.photometricum.DSM.122 GCA_000284415.2_ASM28441v2
96.6/99
Rhodospirillum.rubrum.F11 GCA_000225955.1_ASM22595v1
100/100
100/100
Roseospirillum.parvum GCA_900100455.1
Magnetospirillum.magneticum.AMB.1 GCA_000009985.1_ASM998v1
100/100
Terasakiella.pusilla.DSM.6293 GCA_000688235.1_ASM68823v1
100/100
Thalassospira.profundimaris.WP0211 GCA_000300275.1_ASM30027v1
Elioraea.tepidiphila.DSM.17972 GCA_000378465.1_ASM37846v1
100/100
Belnapia.moabensis.DSM.16746 GCA_000745835.1_ASM74583v1
100/100
100/100
Rubritepida.flocculans.DSM.14296 GCA_000425365.1_ASM42536v1
100/100
Roseomonas.gilardii GCA_001941945.1_ASM194194v1
Acidiphilium.multivorum.AIU301 GCA_000202835.1_ASM20283v1
87.5/99
98.1/100
Acidisphaera.rubrifaciens.HS.AP3 GCA_000964365.1_ASM96436v1
100/100
Granulibacter.bethesdensis.CGDNIH4 GCA_000576145.2_ASM57614v2
Commensalibacter.intestini.A911 GCA_000231445.2_ASM23144v1
100/100
Gluconacetobacter.diazotrophicus.PA1.5 GCA_000067045.1_ASM6704v1
96.5/99
Acetobacter.aceti.NBRC.14818 GCA_000193495.2_ASM19349v2
Gluconobacter.oxydans.DSM.3504 GCA_000583855.1_ASM58385v1
0.646103

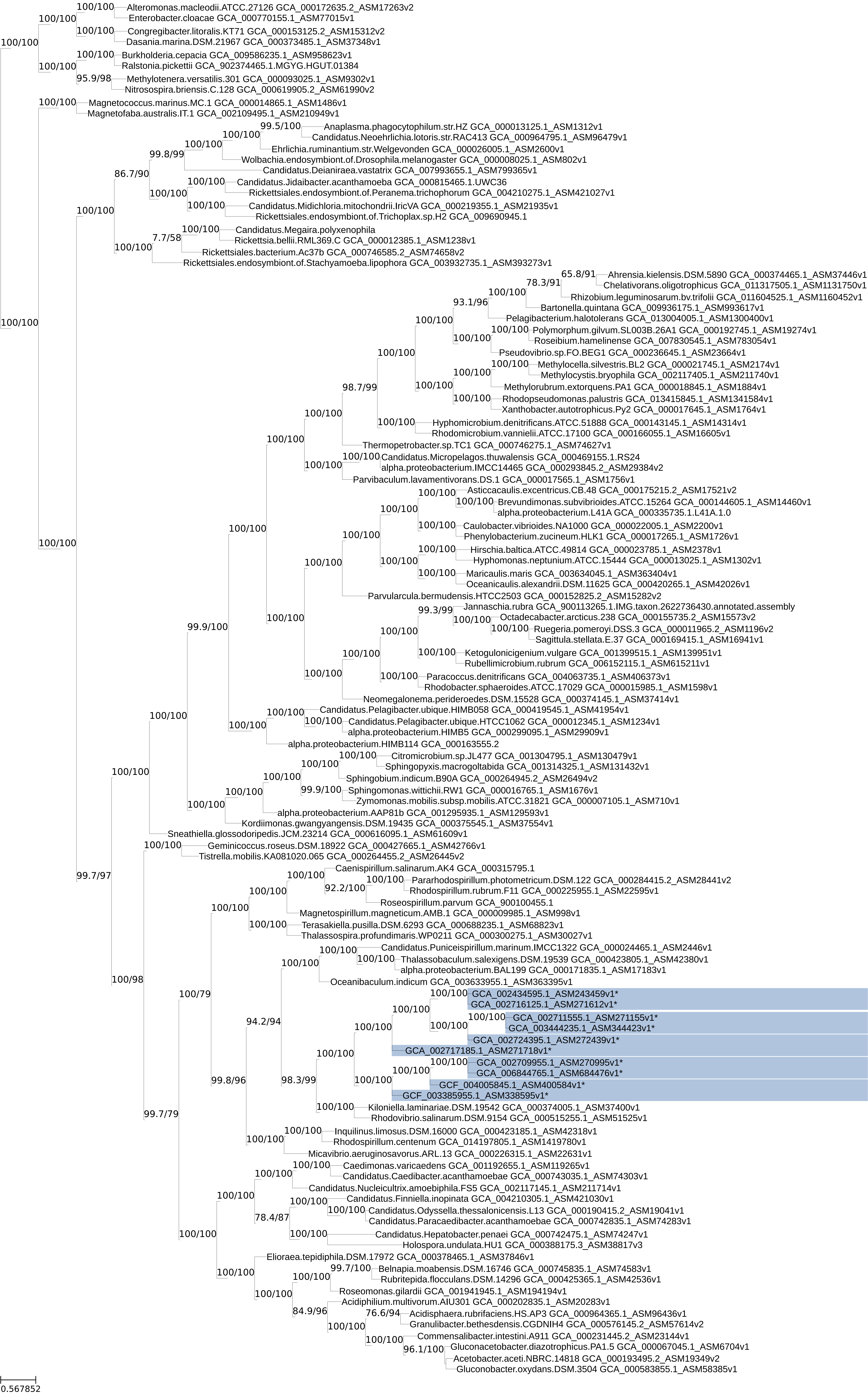
100/100
Alteromonas.macleodii.ATCC.27126 GCA_000172635.2_ASM17263v2
100/100
Enterobacter.cloacae GCA_000770155.1_ASM77015v1
100/100
Congregibacter.litoralis.KT71 GCA_000153125.2_ASM15312v2
100/100
Dasania.marina.DSM.21967 GCA_000373485.1_ASM37348v1
100/100
Burkholderia.cepacia GCA_009586235.1_ASM958623v1
100/100
Ralstonia.pickettii GCA_902374465.1.MGYG.HGUT.01384
95.9/98
Methylotenera.versatilis.301 GCA_000093025.1_ASM9302v1
Nitrosospira.briensis.C.128 GCA_000619905.2_ASM61990v2
100/100
Magnetococcus.marinus.MC.1 GCA_000014865.1_ASM1486v1
Magnetofaba.australis.IT.1 GCA_002109495.1_ASM210949v1
99.5/100
Anaplasma.phagocytophilum.str.HZ GCA_000013125.1_ASM1312v1
100/100
Candidatus.Neoehrlichia.lotoris.str.RAC413 GCA_000964795.1_ASM96479v1
100/100
Ehrlichia.ruminantium.str.Welgevonden GCA_000026005.1_ASM2600v1
99.8/99
Wolbachia.endosymbiont.of.Drosophila.melanogaster GCA_000008025.1_ASM802v1
86.7/90
Candidatus.Deianiraea.vastatrix GCA_007993655.1_ASM799365v1
100/100
Candidatus.Jidaibacter.acanthamoeba GCA_000815465.1.UWC36
100/100
Rickettsiales.endosymbiont.of.Peranema.trichophorum GCA_004210275.1_ASM421027v1
100/100
Candidatus.Midichloria.mitochondrii.IricVA GCA_000219355.1_ASM21935v1
100/100
Rickettsiales.endosymbiont.of.Trichoplax.sp.H2 GCA_009690945.1
100/100
Candidatus.Megaira.polyxenophila
7.7/58
Rickettsia.bellii.RML369.C GCA_000012385.1_ASM1238v1
100/100
Rickettsiales.bacterium.Ac37b GCA_000746585.2_ASM74658v2
Rickettsiales.endosymbiont.of.Stachyamoeba.lipophora GCA_003932735.1_ASM393273v1
65.8/91
Ahrensia.kielensis.DSM.5890 GCA_000374465.1_ASM37446v1
78.3/91
Chelativorans.oligotrophicus GCA_011317505.1_ASM1131750v1
100/100
Rhizobium.leguminosarum.bv.trifolii GCA_011604525.1_ASM1160452v1
93.1/96
Bartonella.quintana GCA_009936175.1_ASM993617v1
100/100
Pelagibacterium.halotolerans GCA_013004005.1_ASM1300400v1
100/100
Polymorphum.gilvum.SL003B.26A1 GCA_000192745.1_ASM19274v1
100/100
Roseibium.hamelinense GCA_007830545.1_ASM783054v1
100/100
Pseudovibrio.sp.FO.BEG1 GCA_000236645.1_ASM23664v1
100/100
Methylocella.silvestris.BL2 GCA_000021745.1_ASM2174v1
100/100
Methylocystis.bryophila GCA_002117405.1_ASM211740v1
100/100
Methylorubrum.extorquens.PA1 GCA_000018845.1_ASM1884v1
98.7/99
100/100
Rhodopseudomonas.palustris GCA_013415845.1_ASM1341584v1
Xanthobacter.autotrophicus.Py2 GCA_000017645.1_ASM1764v1
100/100
100/100
Hyphomicrobium.denitrificans.ATCC.51888 GCA_000143145.1_ASM14314v1
100/100
Rhodomicrobium.vannielii.ATCC.17100 GCA_000166055.1_ASM16605v1
Thermopetrobacter.sp.TC1 GCA_000746275.1_ASM74627v1
100/100
Candidatus.Micropelagos.thuwalensis GCA_000469155.1.RS24
100/100
alpha.proteobacterium.IMCC14465 GCA_000293845.2_ASM29384v2
Parvibaculum.lavamentivorans.DS.1 GCA_000017565.1_ASM1756v1
100/100
Asticcacaulis.excentricus.CB.48 GCA_000175215.2_ASM17521v2
100/100
Brevundimonas.subvibrioides.ATCC.15264 GCA_000144605.1_ASM14460v1
100/100
alpha.proteobacterium.L41A GCA_000335735.1.L41A.1.0
100/100
Caulobacter.vibrioides.NA1000 GCA_000022005.1_ASM2200v1
100/100
Phenylobacterium.zucineum.HLK1 GCA_000017265.1_ASM1726v1
100/100
Hirschia.baltica.ATCC.49814 GCA_000023785.1_ASM2378v1
100/100
Hyphomonas.neptunium.ATCC.15444 GCA_000013025.1_ASM1302v1
100/100
Maricaulis.maris GCA_003634045.1_ASM363404v1
100/100
Oceanicaulis.alexandrii.DSM.11625 GCA_000420265.1_ASM42026v1
Parvularcula.bermudensis.HTCC2503 GCA_000152825.2_ASM15282v2
99.3/99
Jannaschia.rubra GCA_900113265.1.IMG.taxon.2622736430.annotated.assembly
100/100
Octadecabacter.arcticus.238 GCA_000155735.2_ASM15573v2
100/100
100/100
Ruegeria.pomeroyi.DSS.3 GCA_000011965.2_ASM1196v2
99.9/100
Sagittula.stellata.E.37 GCA_000169415.1_ASM16941v1
100/100
Ketogulonicigenium.vulgare GCA_001399515.1_ASM139951v1
100/100
Rubellimicrobium.rubrum GCA_006152115.1_ASM615211v1
100/100
100/100
Paracoccus.denitrificans GCA_004063735.1_ASM406373v1
Rhodobacter.sphaeroides.ATCC.17029 GCA_000015985.1_ASM1598v1
Neomegalonema.perideroedes.DSM.15528 GCA_000374145.1_ASM37414v1
100/100
Candidatus.Pelagibacter.ubique.HIMB058 GCA_000419545.1_ASM41954v1
100/100
Candidatus.Pelagibacter.ubique.HTCC1062 GCA_000012345.1_ASM1234v1
100/100
100/100
alpha.proteobacterium.HIMB5 GCA_000299095.1_ASM29909v1
alpha.proteobacterium.HIMB114 GCA_000163555.2
100/100
Citromicrobium.sp.JL477 GCA_001304795.1_ASM130479v1
100/100
100/100
Sphingopyxis.macrogoltabida GCA_001314325.1_ASM131432v1
100/100
Sphingobium.indicum.B90A GCA_000264945.2_ASM26494v2
100/100
99.9/100
Sphingomonas.wittichii.RW1 GCA_000016765.1_ASM1676v1
Zymomonas.mobilis.subsp.mobilis.ATCC.31821 GCA_000007105.1_ASM710v1
100/100
alpha.proteobacterium.AAP81b GCA_001295935.1_ASM129593v1
Kordiimonas.gwangyangensis.DSM.19435 GCA_000375545.1_ASM37554v1
Sneathiella.glossodoripedis.JCM.23214 GCA_000616095.1_ASM61609v1
100/100
Geminicoccus.roseus.DSM.18922 GCA_000427665.1_ASM42766v1
Tistrella.mobilis.KA081020.065 GCA_000264455.2_ASM26445v2
Caenispirillum.salinarum.AK4 GCA_000315795.1
100/100
100/100
Pararhodospirillum.photometricum.DSM.122 GCA_000284415.2_ASM28441v2
99.7/97
100/100
92.2/100
Rhodospirillum.rubrum.F11 GCA_000225955.1_ASM22595v1
Roseospirillum.parvum GCA_900100455.1
100/100
Magnetospirillum.magneticum.AMB.1 GCA_000009985.1_ASM998v1
100/100
Terasakiella.pusilla.DSM.6293 GCA_000688235.1_ASM68823v1
Thalassospira.profundimaris.WP0211 GCA_000300275.1_ASM30027v1
100/100
Candidatus.Puniceispirillum.marinum.IMCC1322 GCA_000024465.1_ASM2446v1
100/100
100/100
Thalassobaculum.salexigens.DSM.19539 GCA_000423805.1_ASM42380v1
alpha.proteobacterium.BAL199 GCA_000171835.1_ASM17183v1
100/98
Oceanibaculum.indicum GCA_003633955.1_ASM363395v1
100/79
100/100
GCA_002434595.1_ASM243459v1*
GCA_002716125.1_ASM271612v1*
100/100
94.2/94
100/100
100/100
GCA_002711555.1_ASM271155v1*
GCA_003444235.1_ASM344423v1*
100/100
GCA_002724395.1_ASM272439v1*
GCA_002717185.1_ASM271718v1*
100/100
100/100
GCA_002709955.1_ASM270995v1*
100/100
GCA_006844765.1_ASM684476v1*
99.8/96
98.3/99
100/100
GCF_004005845.1_ASM400584v1*
GCF_003385955.1_ASM338595v1*
100/100
Kiloniella.laminariae.DSM.19542 GCA_000374005.1_ASM37400v1
99.7/79
Rhodovibrio.salinarum.DSM.9154 GCA_000515255.1_ASM51525v1
100/100
Inquilinus.limosus.DSM.16000 GCA_000423185.1_ASM42318v1
100/100
Rhodospirillum.centenum GCA_014197805.1_ASM1419780v1
Micavibrio.aeruginosavorus.ARL.13 GCA_000226315.1_ASM22631v1
100/100
Caedimonas.varicaedens GCA_001192655.1_ASM119265v1
100/100
Candidatus.Caedibacter.acanthamoebae GCA_000743035.1_ASM74303v1
100/100
Candidatus.Nucleicultrix.amoebiphila.FS5 GCA_002117145.1_ASM211714v1
100/100
Candidatus.Finniella.inopinata GCA_004210305.1_ASM421030v1
100/100
Candidatus.Odyssella.thessalonicensis.L13 GCA_000190415.2_ASM19041v1
78.4/87
Candidatus.Paracaedibacter.acanthamoebae GCA_000742835.1_ASM74283v1
100/100
100/100
Candidatus.Hepatobacter.penaei GCA_000742475.1_ASM74247v1
Holospora.undulata.HU1 GCA_000388175.3_ASM38817v3
Elioraea.tepidiphila.DSM.17972 GCA_000378465.1_ASM37846v1
99.7/100
Belnapia.moabensis.DSM.16746 GCA_000745835.1_ASM74583v1
100/100
100/100
Rubritepida.flocculans.DSM.14296 GCA_000425365.1_ASM42536v1
100/100
Roseomonas.gilardii GCA_001941945.1_ASM194194v1
Acidiphilium.multivorum.AIU301 GCA_000202835.1_ASM20283v1
76.6/94
84.9/96
Acidisphaera.rubrifaciens.HS.AP3 GCA_000964365.1_ASM96436v1
100/100
Granulibacter.bethesdensis.CGDNIH4 GCA_000576145.2_ASM57614v2
100/100
Commensalibacter.intestini.A911 GCA_000231445.2_ASM23144v1
96.1/100
Gluconacetobacter.diazotrophicus.PA1.5 GCA_000067045.1_ASM6704v1
Acetobacter.aceti.NBRC.14818 GCA_000193495.2_ASM19349v2
Gluconobacter.oxydans.DSM.3504 GCA_000583855.1_ASM58385v1
0.567852

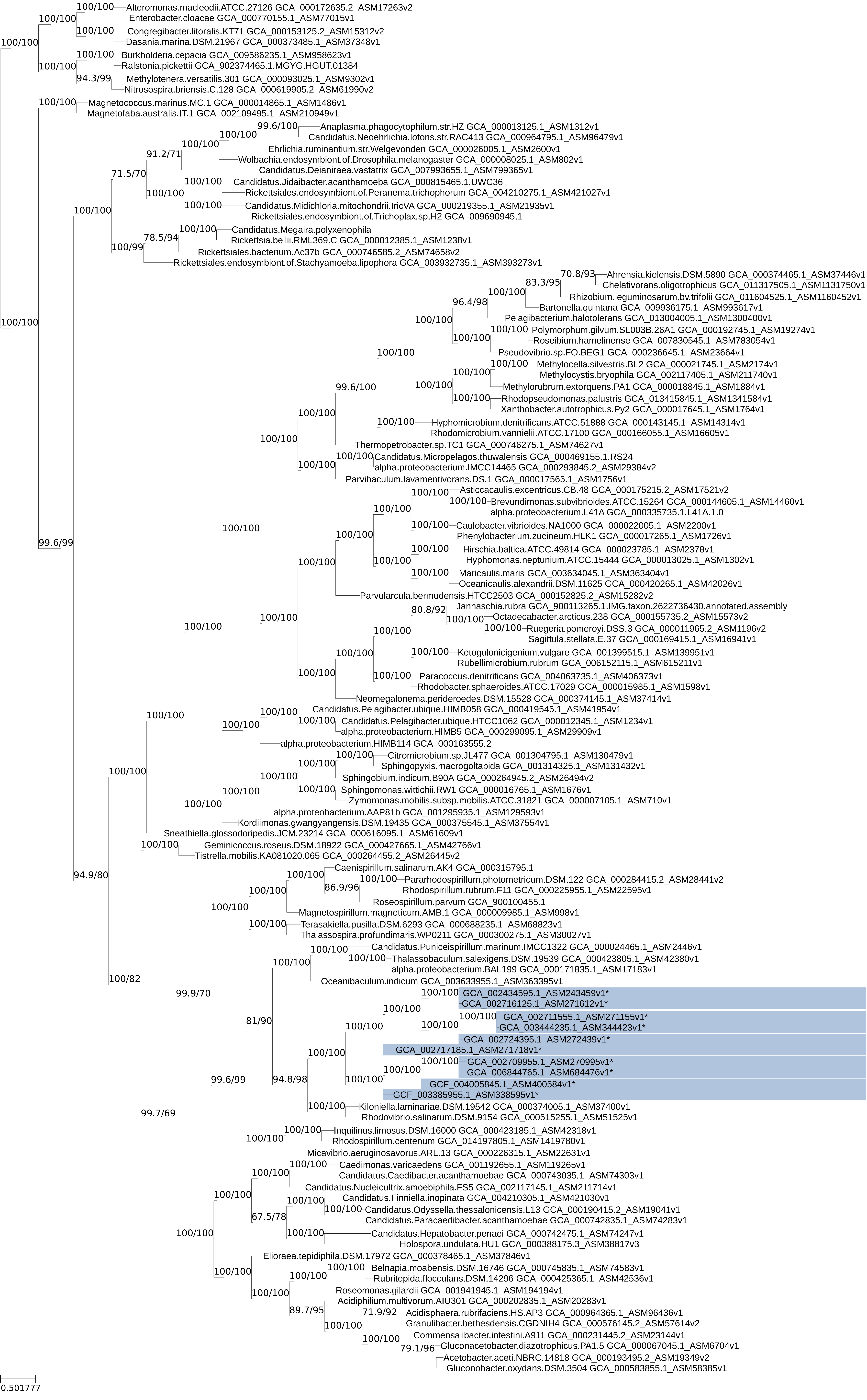
Alteromonas.macleodii.ATCC.27126 GCA_000172635.2_ASM17263v2
Enterobacter.cloacae GCA_000770155.1_ASM77015v1
Congregibacter.litoralis.KT71 GCA_000153125.2_ASM15312v2
Dasania.marina.DSM.21967 GCA_000373485.1_ASM37348v1
Burkholderia.cepacia GCA_009586235.1_ASM958623v1
Ralstonia.pickettii GCA_902374465.1.MGYG.HGUT.01384
Methylotenera.versatilis.301 GCA_000093025.1_ASM9302v1
Nitrosospira.briensis.C.128 GCA_000619905.2_ASM61990v2
Magnetococcus.marinus.MC.1 GCA_000014865.1_ASM1486v1
Magnetofaba.australis.IT.1 GCA_002109495.1_ASM210949v1
Anaplasma.phagocytophilum.str.HZ GCA_000013125.1_ASM1312v1
Candidatus.Neoehrlichia.lotoris.str.RAC413 GCA_000964795.1_ASM96479v1
Ehrlichia.ruminantium.str.Welgevonden GCA_000026005.1_ASM2600v1
Wolbachia.endosymbiont.of.Drosophila.melanogaster GCA_000008025.1_ASM802v1
Candidatus.Deianiraea.vastatrix GCA_007993655.1_ASM799365v1
Candidatus.Jidaibacter.acanthamoeba GCA_000815465.1.UWC36
Rickettsiales.endosymbiont.of.Peranema.trichophorum GCA_004210275.1_ASM421027v1
Candidatus.Midichloria.mitochondrii.IricVA GCA_000219355.1_ASM21935v1
Rickettsiales.endosymbiont.of.Trichoplax.sp.H2 GCA_009690945.1
Candidatus.Megaira.polyxenophila
Rickettsia.bellii.RML369.C GCA_000012385.1_ASM1238v1
Rickettsiales.bacterium.Ac37b GCA_000746585.2_ASM74658v2
Rickettsiales.endosymbiont.of.Stachyamoeba.lipophora GCA_003932735.1_ASM393273v1
Ahrensia.kielensis.DSM.5890 GCA_000374465.1_ASM37446v1
Chelativorans.oligotrophicus GCA_011317505.1_ASM1131750v1
Rhizobium.leguminosarum.bv.trifolii GCA_011604525.1_ASM1160452v1
Bartonella.quintana GCA_009936175.1_ASM993617v1
Pelagibacterium.halotolerans GCA_013004005.1_ASM1300400v1
Polymorphum.gilvum.SL003B.26A1 GCA_000192745.1_ASM19274v1
Roseibium.hamelinense GCA_007830545.1_ASM783054v1
Pseudovibrio.sp.FO.BEG1 GCA_000236645.1_ASM23664v1
Methylocella.silvestris.BL2 GCA_000021745.1_ASM2174v1
Methylocystis.bryophila GCA_002117405.1_ASM211740v1
Methylorubrum.extorquens.PA1 GCA_000018845.1_ASM1884v1
Rhodopseudomonas.palustris GCA_013415845.1_ASM1341584v1
Xanthobacter.autotrophicus.Py2 GCA_000017645.1_ASM1764v1
Hyphomicrobium.denitrificans.ATCC.51888 GCA_000143145.1_ASM14314v1
Rhodomicrobium.vannielii.ATCC.17100 GCA_000166055.1_ASM16605v1
Thermopetrobacter.sp.TC1 GCA_000746275.1_ASM74627v1
Candidatus.Micropelagos.thuwalensis GCA_000469155.1.RS24
alpha.proteobacterium.IMCC14465 GCA_000293845.2_ASM29384v2
Parvibaculum.lavamentivorans.DS.1 GCA_000017565.1_ASM1756v1
Asticcacaulis.excentricus.CB.48 GCA_000175215.2_ASM17521v2
Brevundimonas.subvibrioides.ATCC.15264 GCA_000144605.1_ASM14460v1
alpha.proteobacterium.L41A GCA_000335735.1.L41A.1.0
Caulobacter.vibrioides.NA1000 GCA_000022005.1_ASM2200v1
Phenylobacterium.zucineum.HLK1 GCA_000017265.1_ASM1726v1
Hirschia.baltica.ATCC.49814 GCA_000023785.1_ASM2378v1
Hyphomonas.neptunium.ATCC.15444 GCA_000013025.1_ASM1302v1
Maricaulis.maris GCA_003634045.1_ASM363404v1
Oceanicaulis.alexandrii.DSM.11625 GCA_000420265.1_ASM42026v1
Parvularcula.bermudensis.HTCC2503 GCA_000152825.2_ASM15282v2
Jannaschia.rubra GCA_900113265.1.IMG.taxon.2622736430.annotated.assembly
Octadecabacter.arcticus.238 GCA_000155735.2_ASM15573v2
Ruegeria.pomeroyi.DSS.3 GCA_000011965.2_ASM1196v2
Sagittula.stellata.E.37 GCA_000169415.1_ASM16941v1
Ketogulonicigenium.vulgare GCA_001399515.1_ASM139951v1
Rubellimicrobium.rubrum GCA_006152115.1_ASM615211v1
Paracoccus.denitrificans GCA_004063735.1_ASM406373v1
Rhodobacter.sphaeroides.ATCC.17029 GCA_000015985.1_ASM1598v1
Neomegalonema.perideroedes.DSM.15528 GCA_000374145.1_ASM37414v1
Candidatus.Pelagibacter.ubique.HIMB058 GCA_000419545.1_ASM41954v1
Candidatus.Pelagibacter.ubique.HTCC1062 GCA_000012345.1_ASM1234v1
alpha.proteobacterium.HIMB5 GCA_000299095.1_ASM29909v1
alpha.proteobacterium.HIMB114 GCA_000163555.2
Citromicrobium.sp.JL477 GCA_001304795.1_ASM130479v1
Sphingopyxis.macrogoltabida GCA_001314325.1_ASM131432v1
Sphingobium.indicum.B90A GCA_000264945.2_ASM26494v2
Sphingomonas.wittichii.RW1 GCA_000016765.1_ASM1676v1
Zymomonas.mobilis.subsp.mobilis.ATCC.31821 GCA_000007105.1_ASM710v1
alpha.proteobacterium.AAP81b GCA_001295935.1_ASM129593v1
Kordiimonas.gwangyangensis.DSM.19435 GCA_000375545.1_ASM37554v1
Sneathiella.glossodoripedis.JCM.23214 GCA_000616095.1_ASM61609v1
Geminicoccus.roseus.DSM.18922 GCA_000427665.1_ASM42766v1
Tistrella.mobilis.KA081020.065 GCA_000264455.2_ASM26445v2
Caenispirillum.salinarum.AK4 GCA_000315795.1
Pararhodospirillum.photometricum.DSM.122 GCA_000284415.2_ASM28441v2
Rhodospirillum.rubrum.F11 GCA_000225955.1_ASM22595v1
Roseospirillum.parvum GCA_900100455.1
Magnetospirillum.magneticum.AMB.1 GCA_000009985.1_ASM998v1
Terasakiella.pusilla.DSM.6293 GCA_000688235.1_ASM68823v1
Thalassospira.profundimaris.WP0211 GCA_000300275.1_ASM30027v1
Candidatus.Puniceispirillum.marinum.IMCC1322 GCA_000024465.1_ASM2446v1
Thalassobaculum.salexigens.DSM.19539 GCA_000423805.1_ASM42380v1
alpha.proteobacterium.BAL199 GCA_000171835.1_ASM17183v1
Oceanibaculum.indicum GCA_003633955.1_ASM363395v1
GCA_002434595.1_ASM243459v1*
GCA_002716125.1_ASM271612v1*
GCA_002711555.1_ASM271155v1*
GCA_003444235.1_ASM344423v1*
GCA_002724395.1_ASM272439v1*
GCA_002717185.1_ASM271718v1*
GCA_002709955.1_ASM270995v1*
GCA_006844765.1_ASM684476v1*
GCF_004005845.1_ASM400584v1*
GCF_003385955.1_ASM338595v1*
Kiloniella.laminariae.DSM.19542 GCA_000374005.1_ASM37400v1
Rhodovibrio.salinarum.DSM.9154 GCA_000515255.1_ASM51525v1
Inquilinus.limosus.DSM.16000 GCA_000423185.1_ASM42318v1
Rhodospirillum.centenum GCA_014197805.1_ASM1419780v1
Micavibrio.aeruginosavorus.ARL.13 GCA_000226315.1_ASM22631v1
Caedimonas.varicaedens GCA_001192655.1_ASM119265v1
Candidatus.Caedibacter.acanthamoebae GCA_000743035.1_ASM74303v1
Candidatus.Nucleicultrix.amoebiphila.FS5 GCA_002117145.1_ASM211714v1
Candidatus.Finniella.inopinata GCA_004210305.1_ASM421030v1
Candidatus.Odyssella.thessalonicensis.L13 GCA_000190415.2_ASM19041v1
Candidatus.Paracaedibacter.acanthamoebae GCA_000742835.1_ASM74283v1
Candidatus.Hepatobacter.penaei GCA_000742475.1_ASM74247v1
Holospora.undulata.HU1 GCA_000388175.3_ASM38817v3
Elioraea.tepidiphila.DSM.17972 GCA_000378465.1_ASM37846v1
Belnapia.moabensis.DSM.16746 GCA_000745835.1_ASM74583v1
Rubritepida.flocculans.DSM.14296 GCA_000425365.1_ASM42536v1
Roseomonas.gilardii GCA_001941945.1_ASM194194v1
Acidiphilium.multivorum.AIU301 GCA_000202835.1_ASM20283v1
Acidisphaera.rubrifaciens.HS.AP3 GCA_000964365.1_ASM96436v1
Granulibacter.bethesdensis.CGDNIH4 GCA_000576145.2_ASM57614v2
Commensalibacter.intestini.A911 GCA_000231445.2_ASM23144v1
Gluconacetobacter.diazotrophicus.PA1.5 GCA_000067045.1_ASM6704v1
Acetobacter.aceti.NBRC.14818 GCA_000193495.2_ASM19349v2
Gluconobacter.oxydans.DSM.3504 GCA_000583855.1_ASM58385v1
0.501777

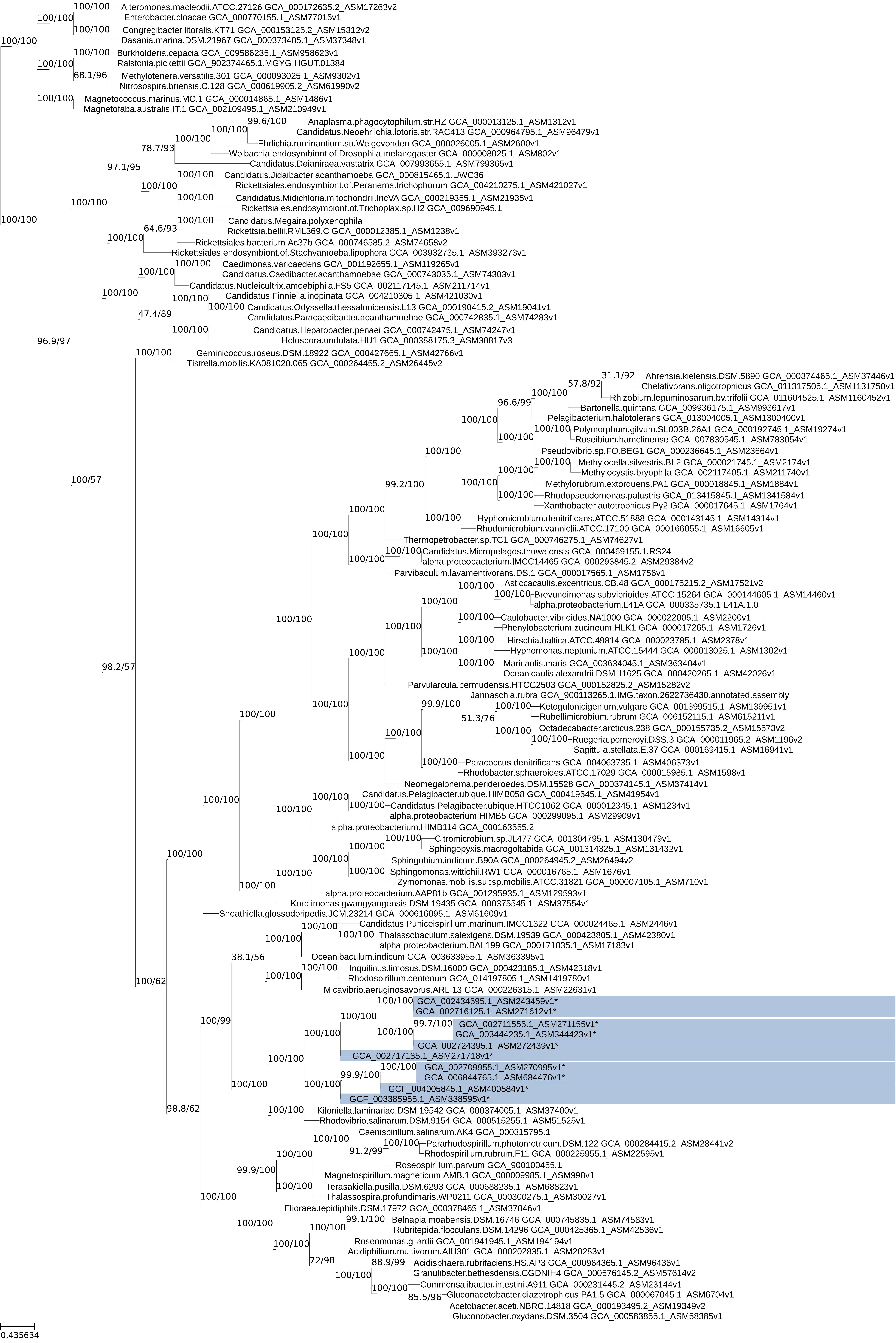

Alteromonas.macleodii.ATCC.27126 GCA_000172635.2_ASM17263v2
Enterobacter.cloacae GCA_000770155.1_ASM77015v1
Congregibacter.litoralis.KT71 GCA_000153125.2_ASM15312v2
Dasania.marina.DSM.21967 GCA_000373485.1_ASM37348v1
Burkholderia.cepacia GCA_009586235.1_ASM958623v1
Ralstonia.pickettii GCA_902374465.1.MGYG.HGUT.01384
Methylotenera.versatilis.301 GCA_000093025.1_ASM9302v1
Nitrosospira.briensis.C.128 GCA_000619905.2_ASM61990v2
Magnetococcus.marinus.MC.1 GCA_000014865.1_ASM1486v1
Magnetofaba.australis.IT.1 GCA_002109495.1_ASM210949v1
Anaplasma.phagocytophilum.str.HZ GCA_000013125.1_ASM1312v1
Candidatus.Neoehrlichia.lotoris.str.RAC413 GCA_000964795.1_ASM96479v1
Ehrlichia.ruminantium.str.Welgevonden GCA_000026005.1_ASM2600v1
Wolbachia.endosymbiont.of.Drosophila.melanogaster GCA_000008025.1_ASM802v1
Candidatus.Deianiraea.vastatrix GCA_007993655.1_ASM799365v1
Candidatus.Jidaibacter.acanthamoeba GCA_000815465.1.UWC36
Rickettsiales.endosymbiont.of.Peranema.trichophorum GCA_004210275.1_ASM421027v1
Candidatus.Midichloria.mitochondrii.IricVA GCA_000219355.1_ASM21935v1
Rickettsiales.endosymbiont.of.Trichoplax.sp.H2 GCA_009690945.1
Candidatus.Megaira.polyxenophila
Rickettsia.bellii.RML369.C GCA_000012385.1_ASM1238v1
Rickettsiales.bacterium.Ac37b GCA_000746585.2_ASM74658v2
Rickettsiales.endosymbiont.of.Stachyamoeba.lipophora GCA_003932735.1_ASM393273v1
Caedimonas.varicaedens GCA_001192655.1_ASM119265v1
Candidatus.Caedibacter.acanthamoebae GCA_000743035.1_ASM74303v1
Candidatus.Nucleicultrix.amoebiphila.FS5 GCA_002117145.1_ASM211714v1
Candidatus.Finniella.inopinata GCA_004210305.1_ASM421030v1
Candidatus.Odyssella.thessalonicensis.L13 GCA_000190415.2_ASM19041v1
Candidatus.Paracaedibacter.acanthamoebae GCA_000742835.1_ASM74283v1
Candidatus.Hepatobacter.penaei GCA_000742475.1_ASM74247v1
Holospora.undulata.HU1 GCA_000388175.3_ASM38817v3
Geminicoccus.roseus.DSM.18922 GCA_000427665.1_ASM42766v1
Tistrella.mobilis.KA081020.065 GCA_000264455.2_ASM26445v2
Ahrensia.kielensis.DSM.5890 GCA_000374465.1_ASM37446v1
Chelativorans.oligotrophicus GCA_011317505.1_ASM1131750v1
Rhizobium.leguminosarum.bv.trifolii GCA_011604525.1_ASM1160452v1
Bartonella.quintana GCA_009936175.1_ASM993617v1
Pelagibacterium.halotolerans GCA_013004005.1_ASM1300400v1
Polymorphum.gilvum.SL003B.26A1 GCA_000192745.1_ASM19274v1
Roseibium.hamelinense GCA_007830545.1_ASM783054v1
Pseudovibrio.sp.FO.BEG1 GCA_000236645.1_ASM23664v1
Methylocella.silvestris.BL2 GCA_000021745.1_ASM2174v1
Methylocystis.bryophila GCA_002117405.1_ASM211740v1
Methylorubrum.extorquens.PA1 GCA_000018845.1_ASM1884v1
Rhodopseudomonas.palustris GCA_013415845.1_ASM1341584v1
Xanthobacter.autotrophicus.Py2 GCA_000017645.1_ASM1764v1
Hyphomicrobium.denitrificans.ATCC.51888 GCA_000143145.1_ASM14314v1
Rhodomicrobium.vannielii.ATCC.17100 GCA_000166055.1_ASM16605v1
Thermopetrobacter.sp.TC1 GCA_000746275.1_ASM74627v1
Candidatus.Micropelagos.thuwalensis GCA_000469155.1.RS24
alpha.proteobacterium.IMCC14465 GCA_000293845.2_ASM29384v2
Parvibaculum.lavamentivorans.DS.1 GCA_000017565.1_ASM1756v1
Asticcacaulis.excentricus.CB.48 GCA_000175215.2_ASM17521v2
Brevundimonas.subvibrioides.ATCC.15264 GCA_000144605.1_ASM14460v1
alpha.proteobacterium.L41A GCA_000335735.1.L41A.1.0
Caulobacter.vibrioides.NA1000 GCA_000022005.1_ASM2200v1
Phenylobacterium.zucineum.HLK1 GCA_000017265.1_ASM1726v1
Hirschia.baltica.ATCC.49814 GCA_000023785.1_ASM2378v1
Hyphomonas.neptunium.ATCC.15444 GCA_000013025.1_ASM1302v1
Maricaulis.maris GCA_003634045.1_ASM363404v1
Oceanicaulis.alexandrii.DSM.11625 GCA_000420265.1_ASM42026v1
Parvularcula.bermudensis.HTCC2503 GCA_000152825.2_ASM15282v2
Jannaschia.rubra GCA_900113265.1.IMG.taxon.2622736430.annotated.assembly
Ketogulonicigenium.vulgare GCA_001399515.1_ASM139951v1
Rubellimicrobium.rubrum GCA_006152115.1_ASM615211v1
Octadecabacter.arcticus.238 GCA_000155735.2_ASM15573v2
Ruegeria.pomeroyi.DSS.3 GCA_000011965.2_ASM1196v2
Sagittula.stellata.E.37 GCA_000169415.1_ASM16941v1
Paracoccus.denitrificans GCA_004063735.1_ASM406373v1
Rhodobacter.sphaeroides.ATCC.17029 GCA_000015985.1_ASM1598v1
Neomegalonema.perideroedes.DSM.15528 GCA_000374145.1_ASM37414v1
Candidatus.Pelagibacter.ubique.HIMB058 GCA_000419545.1_ASM41954v1
Candidatus.Pelagibacter.ubique.HTCC1062 GCA_000012345.1_ASM1234v1
alpha.proteobacterium.HIMB5 GCA_000299095.1_ASM29909v1
alpha.proteobacterium.HIMB114 GCA_000163555.2
Citromicrobium.sp.JL477 GCA_001304795.1_ASM130479v1
Sphingopyxis.macrogoltabida GCA_001314325.1_ASM131432v1
Sphingobium.indicum.B90A GCA_000264945.2_ASM26494v2
Sphingomonas.wittichii.RW1 GCA_000016765.1_ASM1676v1
Zymomonas.mobilis.subsp.mobilis.ATCC.31821 GCA_000007105.1_ASM710v1
alpha.proteobacterium.AAP81b GCA_001295935.1_ASM129593v1
Kordiimonas.gwangyangensis.DSM.19435 GCA_000375545.1_ASM37554v1
Sneathiella.glossodoripedis.JCM.23214 GCA_000616095.1_ASM61609v1
Candidatus.Puniceispirillum.marinum.IMCC1322 GCA_000024465.1_ASM2446v1
Thalassobaculum.salexigens.DSM.19539 GCA_000423805.1_ASM42380v1
alpha.proteobacterium.BAL199 GCA_000171835.1_ASM17183v1
Oceanibaculum.indicum GCA_003633955.1_ASM363395v1
Inquilinus.limosus.DSM.16000 GCA_000423185.1_ASM42318v1
Rhodospirillum.centenum GCA_014197805.1_ASM1419780v1
Micavibrio.aeruginosavorus.ARL.13 GCA_000226315.1_ASM22631v1
GCA_002434595.1_ASM243459v1*
GCA_002716125.1_ASM271612v1*
GCA_002711555.1_ASM271155v1*
GCA_003444235.1_ASM344423v1*
GCA_002724395.1_ASM272439v1*
GCA_002717185.1_ASM271718v1*
GCA_002709955.1_ASM270995v1*
GCA_006844765.1_ASM684476v1*
GCF_004005845.1_ASM400584v1*
GCF_003385955.1_ASM338595v1*
Kiloniella.laminariae.DSM.19542 GCA_000374005.1_ASM37400v1
Rhodovibrio.salinarum.DSM.9154 GCA_000515255.1_ASM51525v1
Caenispirillum.salinarum.AK4 GCA_000315795.1
Pararhodospirillum.photometricum.DSM.122 GCA_000284415.2_ASM28441v2
Rhodospirillum.rubrum.F11 GCA_000225955.1_ASM22595v1
Roseospirillum.parvum GCA_900100455.1
Magnetospirillum.magneticum.AMB.1 GCA_000009985.1_ASM998v1
Terasakiella.pusilla.DSM.6293 GCA_000688235.1_ASM68823v1
Thalassospira.profundimaris.WP0211 GCA_000300275.1_ASM30027v1
Elioraea.tepidiphila.DSM.17972 GCA_000378465.1_ASM37846v1
Belnapia.moabensis.DSM.16746 GCA_000745835.1_ASM74583v1
Rubritepida.flocculans.DSM.14296 GCA_000425365.1_ASM42536v1
Roseomonas.gilardii GCA_001941945.1_ASM194194v1
Acidiphilium.multivorum.AIU301 GCA_000202835.1_ASM20283v1
Acidisphaera.rubrifaciens.HS.AP3 GCA_000964365.1_ASM96436v1
Granulibacter.bethesdensis.CGDNIH4 GCA_000576145.2_ASM57614v2
Commensalibacter.intestini.A911 GCA_000231445.2_ASM23144v1
Gluconacetobacter.diazotrophicus.PA1.5 GCA_000067045.1_ASM6704v1
Acetobacter.aceti.NBRC.14818 GCA_000193495.2_ASM19349v2
Gluconobacter.oxydans.DSM.3504 GCA_000583855.1_ASM58385v1
0.435634

### Group D

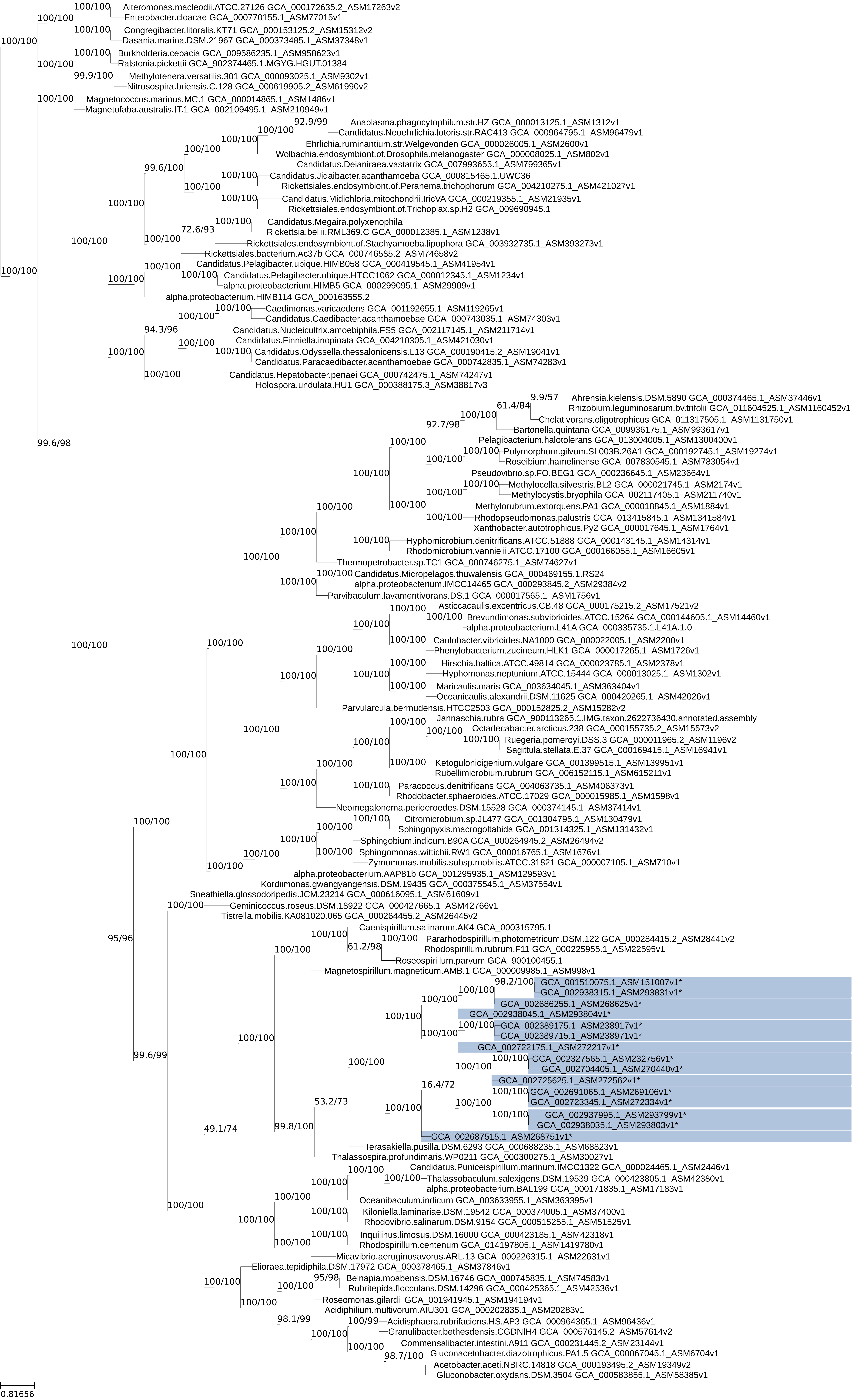
Alteromonas.macleodii.ATCC.27126 GCA_000172635.2_ASM17263v2
Enterobacter.cloacae GCA_000770155.1_ASM77015v1
Congregibacter.litoralis.KT71 GCA_000153125.2_ASM15312v2
Dasania.marina.DSM.21967 GCA_000373485.1_ASM37348v1
Burkholderia.cepacia GCA_009586235.1_ASM958623v1
Ralstonia.pickettii GCA_902374465.1.MGYG.HGUT.01384
Methylotenera.versatilis.301 GCA_000093025.1_ASM9302v1
Nitrosospira.briensis.C.128 GCA_000619905.2_ASM61990v2
Magnetococcus.marinus.MC.1 GCA_000014865.1_ASM1486v1
Magnetofaba.australis.IT.1 GCA_002109495.1_ASM210949v1
Anaplasma.phagocytophilum.str.HZ GCA_000013125.1_ASM1312v1
Candidatus.Neoehrlichia.lotoris.str.RAC413 GCA_000964795.1_ASM96479v1
Ehrlichia.ruminantium.str.Welgevonden GCA_000026005.1_ASM2600v1
Wolbachia.endosymbiont.of.Drosophila.melanogaster GCA_000008025.1_ASM802v1
Candidatus.Deianiraea.vastatrix GCA_007993655.1_ASM799365v1
Candidatus.Jidaibacter.acanthamoeba GCA_000815465.1.UWC36
Rickettsiales.endosymbiont.of.Peranema.trichophorum GCA_004210275.1_ASM421027v1
Candidatus.Midichloria.mitochondrii.IricVA GCA_000219355.1_ASM21935v1
Rickettsiales.endosymbiont.of.Trichoplax.sp.H2 GCA_009690945.1
Candidatus.Megaira.polyxenophila
Rickettsia.bellii.RML369.C GCA_000012385.1_ASM1238v1
Rickettsiales.endosymbiont.of.Stachyamoeba.lipophora GCA_003932735.1_ASM393273v1
Rickettsiales.bacterium.Ac37b GCA_000746585.2_ASM74658v2
Candidatus.Pelagibacter.ubique.HIMB058 GCA_000419545.1_ASM41954v1
Candidatus.Pelagibacter.ubique.HTCC1062 GCA_000012345.1_ASM1234v1
alpha.proteobacterium.HIMB5 GCA_000299095.1_ASM29909v1
alpha.proteobacterium.HIMB114 GCA_000163555.2
Caedimonas.varicaedens GCA_001192655.1_ASM119265v1
Candidatus.Caedibacter.acanthamoebae GCA_000743035.1_ASM74303v1
Candidatus.Nucleicultrix.amoebiphila.FS5 GCA_002117145.1_ASM211714v1
Candidatus.Finniella.inopinata GCA_004210305.1_ASM421030v1
Candidatus.Odyssella.thessalonicensis.L13 GCA_000190415.2_ASM19041v1
Candidatus.Paracaedibacter.acanthamoebae GCA_000742835.1_ASM74283v1
Candidatus.Hepatobacter.penaei GCA_000742475.1_ASM74247v1
Holospora.undulata.HU1 GCA_000388175.3_ASM38817v3
Ahrensia.kielensis.DSM.5890 GCA_000374465.1_ASM37446v1
Rhizobium.leguminosarum.bv.trifolii GCA_011604525.1_ASM1160452v1
Chelativorans.oligotrophicus GCA_011317505.1_ASM1131750v1
Bartonella.quintana GCA_009936175.1_ASM993617v1
Pelagibacterium.halotolerans GCA_013004005.1_ASM1300400v1
Polymorphum.gilvum.SL003B.26A1 GCA_000192745.1_ASM19274v1
Roseibium.hamelinense GCA_007830545.1_ASM783054v1
Pseudovibrio.sp.FO.BEG1 GCA_000236645.1_ASM23664v1
Methylocella.silvestris.BL2 GCA_000021745.1_ASM2174v1
Methylocystis.bryophila GCA_002117405.1_ASM211740v1
Methylorubrum.extorquens.PA1 GCA_000018845.1_ASM1884v1
Rhodopseudomonas.palustris GCA_013415845.1_ASM1341584v1
Xanthobacter.autotrophicus.Py2 GCA_000017645.1_ASM1764v1
Hyphomicrobium.denitrificans.ATCC.51888 GCA_000143145.1_ASM14314v1
Rhodomicrobium.vannielii.ATCC.17100 GCA_000166055.1_ASM16605v1
Thermopetrobacter.sp.TC1 GCA_000746275.1_ASM74627v1
Candidatus.Micropelagos.thuwalensis GCA_000469155.1.RS24
alpha.proteobacterium.IMCC14465 GCA_000293845.2_ASM29384v2
Parvibaculum.lavamentivorans.DS.1 GCA_000017565.1_ASM1756v1
Asticcacaulis.excentricus.CB.48 GCA_000175215.2_ASM17521v2
Brevundimonas.subvibrioides.ATCC.15264 GCA_000144605.1_ASM14460v1
alpha.proteobacterium.L41A GCA_000335735.1.L41A.1.0
Caulobacter.vibrioides.NA1000 GCA_000022005.1_ASM2200v1
Phenylobacterium.zucineum.HLK1 GCA_000017265.1_ASM1726v1
Hirschia.baltica.ATCC.49814 GCA_000023785.1_ASM2378v1
Hyphomonas.neptunium.ATCC.15444 GCA_000013025.1_ASM1302v1
Maricaulis.maris GCA_003634045.1_ASM363404v1
Oceanicaulis.alexandrii.DSM.11625 GCA_000420265.1_ASM42026v1
Parvularcula.bermudensis.HTCC2503 GCA_000152825.2_ASM15282v2
Jannaschia.rubra GCA_900113265.1.IMG.taxon.2622736430.annotated.assembly
Octadecabacter.arcticus.238 GCA_000155735.2_ASM15573v2
Ruegeria.pomeroyi.DSS.3 GCA_000011965.2_ASM1196v2
Sagittula.stellata.E.37 GCA_000169415.1_ASM16941v1
Ketogulonicigenium.vulgare GCA_001399515.1_ASM139951v1
Rubellimicrobium.rubrum GCA_006152115.1_ASM615211v1
Paracoccus.denitrificans GCA_004063735.1_ASM406373v1
Rhodobacter.sphaeroides.ATCC.17029 GCA_000015985.1_ASM1598v1
Neomegalonema.perideroedes.DSM.15528 GCA_000374145.1_ASM37414v1
Citromicrobium.sp.JL477 GCA_001304795.1_ASM130479v1
Sphingopyxis.macrogoltabida GCA_001314325.1_ASM131432v1
Sphingobium.indicum.B90A GCA_000264945.2_ASM26494v2
Sphingomonas.wittichii.RW1 GCA_000016765.1_ASM1676v1
Zymomonas.mobilis.subsp.mobilis.ATCC.31821 GCA_000007105.1_ASM710v1
alpha.proteobacterium.AAP81b GCA_001295935.1_ASM129593v1
Kordiimonas.gwangyangensis.DSM.19435 GCA_000375545.1_ASM37554v1
Sneathiella.glossodoripedis.JCM.23214 GCA_000616095.1_ASM61609v1
Geminicoccus.roseus.DSM.18922 GCA_000427665.1_ASM42766v1
Tistrella.mobilis.KA081020.065 GCA_000264455.2_ASM26445v2
Caenispirillum.salinarum.AK4 GCA_000315795.1
Pararhodospirillum.photometricum.DSM.122 GCA_000284415.2_ASM28441v2
Rhodospirillum.rubrum.F11 GCA_000225955.1_ASM22595v1
Roseospirillum.parvum GCA_900100455.1
Magnetospirillum.magneticum.AMB.1 GCA_000009985.1_ASM998v1
GCA_001510075.1_ASM151007v1*
GCA_002938315.1_ASM293831v1*
GCA_002686255.1_ASM268625v1*
GCA_002938045.1_ASM293804v1*
GCA_002389175.1_ASM238917v1*
GCA_002389715.1_ASM238971v1*
GCA_002722175.1_ASM272217v1*
GCA_002327565.1_ASM232756v1*
GCA_002704405.1_ASM270440v1*
GCA_002725625.1_ASM272562v1*
GCA_002691065.1_ASM269106v1*
GCA_002723345.1_ASM272334v1*
GCA_002937995.1_ASM293799v1*
GCA_002938035.1_ASM293803v1*
GCA_002687515.1_ASM268751v1*
Terasakiella.pusilla.DSM.6293 GCA_000688235.1_ASM68823v1
Thalassospira.profundimaris.WP0211 GCA_000300275.1_ASM30027v1
Candidatus.Puniceispirillum.marinum.IMCC1322 GCA_000024465.1_ASM2446v1
Thalassobaculum.salexigens.DSM.19539 GCA_000423805.1_ASM42380v1
alpha.proteobacterium.BAL199 GCA_000171835.1_ASM17183v1
Oceanibaculum.indicum GCA_003633955.1_ASM363395v1
Kiloniella.laminariae.DSM.19542 GCA_000374005.1_ASM37400v1
Rhodovibrio.salinarum.DSM.9154 GCA_000515255.1_ASM51525v1
Inquilinus.limosus.DSM.16000 GCA_000423185.1_ASM42318v1
Rhodospirillum.centenum GCA_014197805.1_ASM1419780v1
Micavibrio.aeruginosavorus.ARL.13 GCA_000226315.1_ASM22631v1
Elioraea.tepidiphila.DSM.17972 GCA_000378465.1_ASM37846v1
Belnapia.moabensis.DSM.16746 GCA_000745835.1_ASM74583v1
Rubritepida.flocculans.DSM.14296 GCA_000425365.1_ASM42536v1
Roseomonas.gilardii GCA_001941945.1_ASM194194v1
Acidiphilium.multivorum.AIU301 GCA_000202835.1_ASM20283v1
Acidisphaera.rubrifaciens.HS.AP3 GCA_000964365.1_ASM96436v1
Granulibacter.bethesdensis.CGDNIH4 GCA_000576145.2_ASM57614v2
Commensalibacter.intestini.A911 GCA_000231445.2_ASM23144v1
Gluconacetobacter.diazotrophicus.PA1.5 GCA_000067045.1_ASM6704v1
Acetobacter.aceti.NBRC.14818 GCA_000193495.2_ASM19349v2
Gluconobacter.oxydans.DSM.3504 GCA_000583855.1_ASM58385v1
0.81656

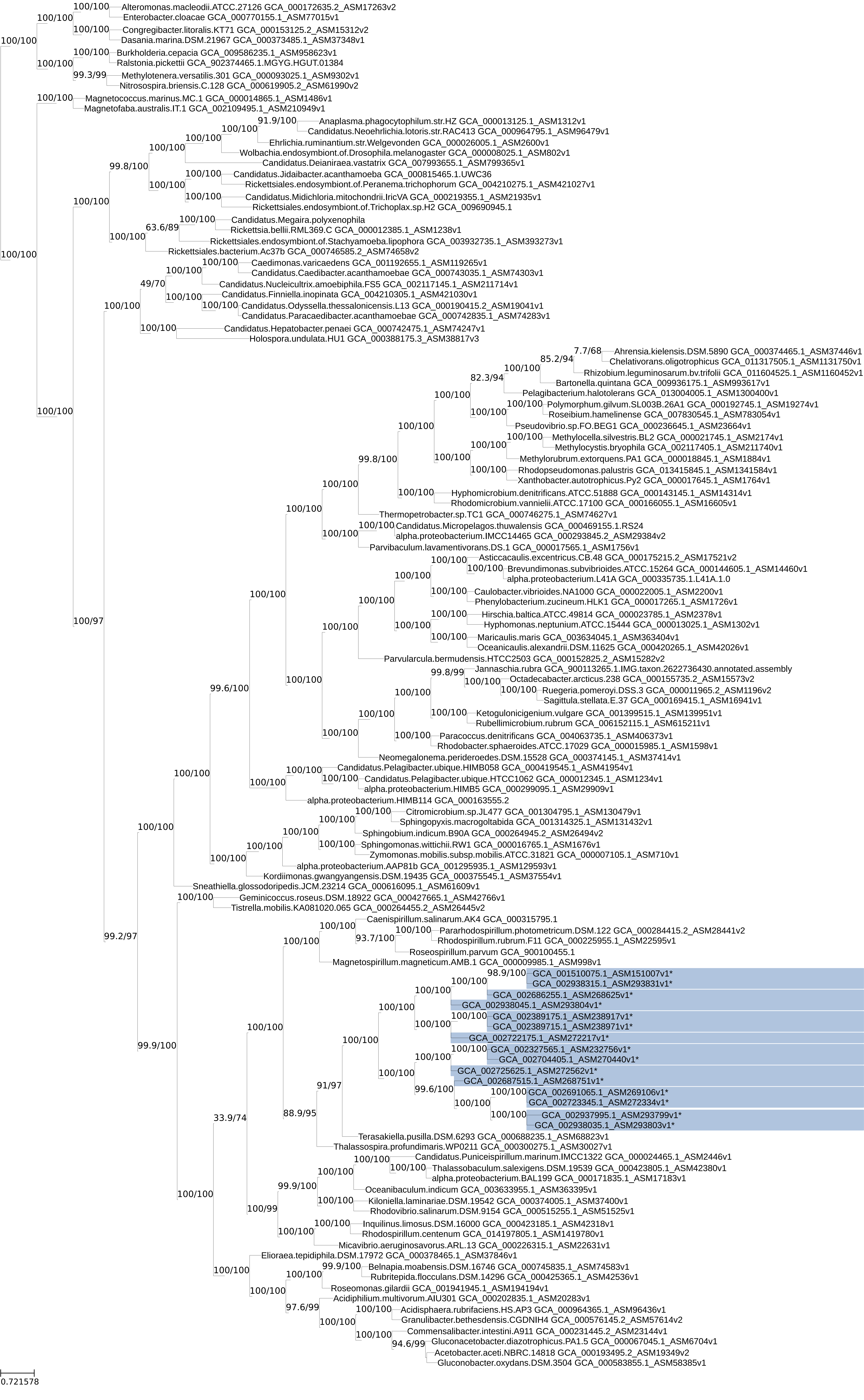
Alteromonas.macleodii.ATCC.27126 GCA_000172635.2_ASM17263v2
Enterobacter.cloacae GCA_000770155.1_ASM77015v1
Congregibacter.litoralis.KT71 GCA_000153125.2_ASM15312v2
Dasania.marina.DSM.21967 GCA_000373485.1_ASM37348v1
Burkholderia.cepacia GCA_009586235.1_ASM958623v1
Ralstonia.pickettii GCA_902374465.1.MGYG.HGUT.01384
Methylotenera.versatilis.301 GCA_000093025.1_ASM9302v1
Nitrosospira.briensis.C.128 GCA_000619905.2_ASM61990v2
Magnetococcus.marinus.MC.1 GCA_000014865.1_ASM1486v1
Magnetofaba.australis.IT.1 GCA_002109495.1_ASM210949v1
Anaplasma.phagocytophilum.str.HZ GCA_000013125.1_ASM1312v1
Candidatus.Neoehrlichia.lotoris.str.RAC413 GCA_000964795.1_ASM96479v1
Ehrlichia.ruminantium.str.Welgevonden GCA_000026005.1_ASM2600v1
Wolbachia.endosymbiont.of.Drosophila.melanogaster GCA_000008025.1_ASM802v1
Candidatus.Deianiraea.vastatrix GCA_007993655.1_ASM799365v1
Candidatus.Jidaibacter.acanthamoeba GCA_000815465.1.UWC36
Rickettsiales.endosymbiont.of.Peranema.trichophorum GCA_004210275.1_ASM421027v1
Candidatus.Midichloria.mitochondrii.IricVA GCA_000219355.1_ASM21935v1
Rickettsiales.endosymbiont.of.Trichoplax.sp.H2 GCA_009690945.1
Candidatus.Megaira.polyxenophila
Rickettsia.bellii.RML369.C GCA_000012385.1_ASM1238v1
Rickettsiales.endosymbiont.of.Stachyamoeba.lipophora GCA_003932735.1_ASM393273v1
Rickettsiales.bacterium.Ac37b GCA_000746585.2_ASM74658v2
Caedimonas.varicaedens GCA_001192655.1_ASM119265v1
Candidatus.Caedibacter.acanthamoebae GCA_000743035.1_ASM74303v1
Candidatus.Nucleicultrix.amoebiphila.FS5 GCA_002117145.1_ASM211714v1
Candidatus.Finniella.inopinata GCA_004210305.1_ASM421030v1
Candidatus.Odyssella.thessalonicensis.L13 GCA_000190415.2_ASM19041v1
Candidatus.Paracaedibacter.acanthamoebae GCA_000742835.1_ASM74283v1
Candidatus.Hepatobacter.penaei GCA_000742475.1_ASM74247v1
Holospora.undulata.HU1 GCA_000388175.3_ASM38817v3
Ahrensia.kielensis.DSM.5890 GCA_000374465.1_ASM37446v1
Chelativorans.oligotrophicus GCA_011317505.1_ASM1131750v1
Rhizobium.leguminosarum.bv.trifolii GCA_011604525.1_ASM1160452v1
Bartonella.quintana GCA_009936175.1_ASM993617v1
Pelagibacterium.halotolerans GCA_013004005.1_ASM1300400v1
Polymorphum.gilvum.SL003B.26A1 GCA_000192745.1_ASM19274v1
Roseibium.hamelinense GCA_007830545.1_ASM783054v1
Pseudovibrio.sp.FO.BEG1 GCA_000236645.1_ASM23664v1
Methylocella.silvestris.BL2 GCA_000021745.1_ASM2174v1
Methylocystis.bryophila GCA_002117405.1_ASM211740v1
Methylorubrum.extorquens.PA1 GCA_000018845.1_ASM1884v1
Rhodopseudomonas.palustris GCA_013415845.1_ASM1341584v1
Xanthobacter.autotrophicus.Py2 GCA_000017645.1_ASM1764v1
Hyphomicrobium.denitrificans.ATCC.51888 GCA_000143145.1_ASM14314v1
Rhodomicrobium.vannielii.ATCC.17100 GCA_000166055.1_ASM16605v1
Thermopetrobacter.sp.TC1 GCA_000746275.1_ASM74627v1
Candidatus.Micropelagos.thuwalensis GCA_000469155.1.RS24
alpha.proteobacterium.IMCC14465 GCA_000293845.2_ASM29384v2
Parvibaculum.lavamentivorans.DS.1 GCA_000017565.1_ASM1756v1
Asticcacaulis.excentricus.CB.48 GCA_000175215.2_ASM17521v2
Brevundimonas.subvibrioides.ATCC.15264 GCA_000144605.1_ASM14460v1
alpha.proteobacterium.L41A GCA_000335735.1.L41A.1.0
Caulobacter.vibrioides.NA1000 GCA_000022005.1_ASM2200v1
Phenylobacterium.zucineum.HLK1 GCA_000017265.1_ASM1726v1
Hirschia.baltica.ATCC.49814 GCA_000023785.1_ASM2378v1
Hyphomonas.neptunium.ATCC.15444 GCA_000013025.1_ASM1302v1
Maricaulis.maris GCA_003634045.1_ASM363404v1
Oceanicaulis.alexandrii.DSM.11625 GCA_000420265.1_ASM42026v1
Parvularcula.bermudensis.HTCC2503 GCA_000152825.2_ASM15282v2
Jannaschia.rubra GCA_900113265.1.IMG.taxon.2622736430.annotated.assembly
Octadecabacter.arcticus.238 GCA_000155735.2_ASM15573v2
Ruegeria.pomeroyi.DSS.3 GCA_000011965.2_ASM1196v2
Sagittula.stellata.E.37 GCA_000169415.1_ASM16941v1
Ketogulonicigenium.vulgare GCA_001399515.1_ASM139951v1
Rubellimicrobium.rubrum GCA_006152115.1_ASM615211v1
Paracoccus.denitrificans GCA_004063735.1_ASM406373v1
Rhodobacter.sphaeroides.ATCC.17029 GCA_000015985.1_ASM1598v1
Neomegalonema.perideroedes.DSM.15528 GCA_000374145.1_ASM37414v1
Candidatus.Pelagibacter.ubique.HIMB058 GCA_000419545.1_ASM41954v1
Candidatus.Pelagibacter.ubique.HTCC1062 GCA_000012345.1_ASM1234v1
alpha.proteobacterium.HIMB5 GCA_000299095.1_ASM29909v1
alpha.proteobacterium.HIMB114 GCA_000163555.2
Citromicrobium.sp.JL477 GCA_001304795.1_ASM130479v1
Sphingopyxis.macrogoltabida GCA_001314325.1_ASM131432v1
Sphingobium.indicum.B90A GCA_000264945.2_ASM26494v2
Sphingomonas.wittichii.RW1 GCA_000016765.1_ASM1676v1
Zymomonas.mobilis.subsp.mobilis.ATCC.31821 GCA_000007105.1_ASM710v1
alpha.proteobacterium.AAP81b GCA_001295935.1_ASM129593v1
Kordiimonas.gwangyangensis.DSM.19435 GCA_000375545.1_ASM37554v1
Sneathiella.glossodoripedis.JCM.23214 GCA_000616095.1_ASM61609v1
Geminicoccus.roseus.DSM.18922 GCA_000427665.1_ASM42766v1
Tistrella.mobilis.KA081020.065 GCA_000264455.2_ASM26445v2
Caenispirillum.salinarum.AK4 GCA_000315795.1
Pararhodospirillum.photometricum.DSM.122 GCA_000284415.2_ASM28441v2
Rhodospirillum.rubrum.F11 GCA_000225955.1_ASM22595v1
Roseospirillum.parvum GCA_900100455.1
Magnetospirillum.magneticum.AMB.1 GCA_000009985.1_ASM998v1
GCA_001510075.1_ASM151007v1*
GCA_002938315.1_ASM293831v1*
GCA_002686255.1_ASM268625v1*
GCA_002938045.1_ASM293804v1*
GCA_002389175.1_ASM238917v1*
GCA_002389715.1_ASM238971v1*
GCA_002722175.1_ASM272217v1*
GCA_002327565.1_ASM232756v1*
GCA_002704405.1_ASM270440v1*
GCA_002725625.1_ASM272562v1*
GCA_002687515.1_ASM268751v1*
GCA_002691065.1_ASM269106v1*
GCA_002723345.1_ASM272334v1*
GCA_002937995.1_ASM293799v1*
GCA_002938035.1_ASM293803v1*
Terasakiella.pusilla.DSM.6293 GCA_000688235.1_ASM68823v1
Thalassospira.profundimaris.WP0211 GCA_000300275.1_ASM30027v1
Candidatus.Puniceispirillum.marinum.IMCC1322 GCA_000024465.1_ASM2446v1
Thalassobaculum.salexigens.DSM.19539 GCA_000423805.1_ASM42380v1
alpha.proteobacterium.BAL199 GCA_000171835.1_ASM17183v1
Oceanibaculum.indicum GCA_003633955.1_ASM363395v1
Kiloniella.laminariae.DSM.19542 GCA_000374005.1_ASM37400v1
Rhodovibrio.salinarum.DSM.9154 GCA_000515255.1_ASM51525v1
Inquilinus.limosus.DSM.16000 GCA_000423185.1_ASM42318v1
Rhodospirillum.centenum GCA_014197805.1_ASM1419780v1
Micavibrio.aeruginosavorus.ARL.13 GCA_000226315.1_ASM22631v1
Elioraea.tepidiphila.DSM.17972 GCA_000378465.1_ASM37846v1
Belnapia.moabensis.DSM.16746 GCA_000745835.1_ASM74583v1
Rubritepida.flocculans.DSM.14296 GCA_000425365.1_ASM42536v1
Roseomonas.gilardii GCA_001941945.1_ASM194194v1
Acidiphilium.multivorum.AIU301 GCA_000202835.1_ASM20283v1
Acidisphaera.rubrifaciens.HS.AP3 GCA_000964365.1_ASM96436v1
Granulibacter.bethesdensis.CGDNIH4 GCA_000576145.2_ASM57614v2
Commensalibacter.intestini.A911 GCA_000231445.2_ASM23144v1
Gluconacetobacter.diazotrophicus.PA1.5 GCA_000067045.1_ASM6704v1
Acetobacter.aceti.NBRC.14818 GCA_000193495.2_ASM19349v2
Gluconobacter.oxydans.DSM.3504 GCA_000583855.1_ASM58385v1
0.721578

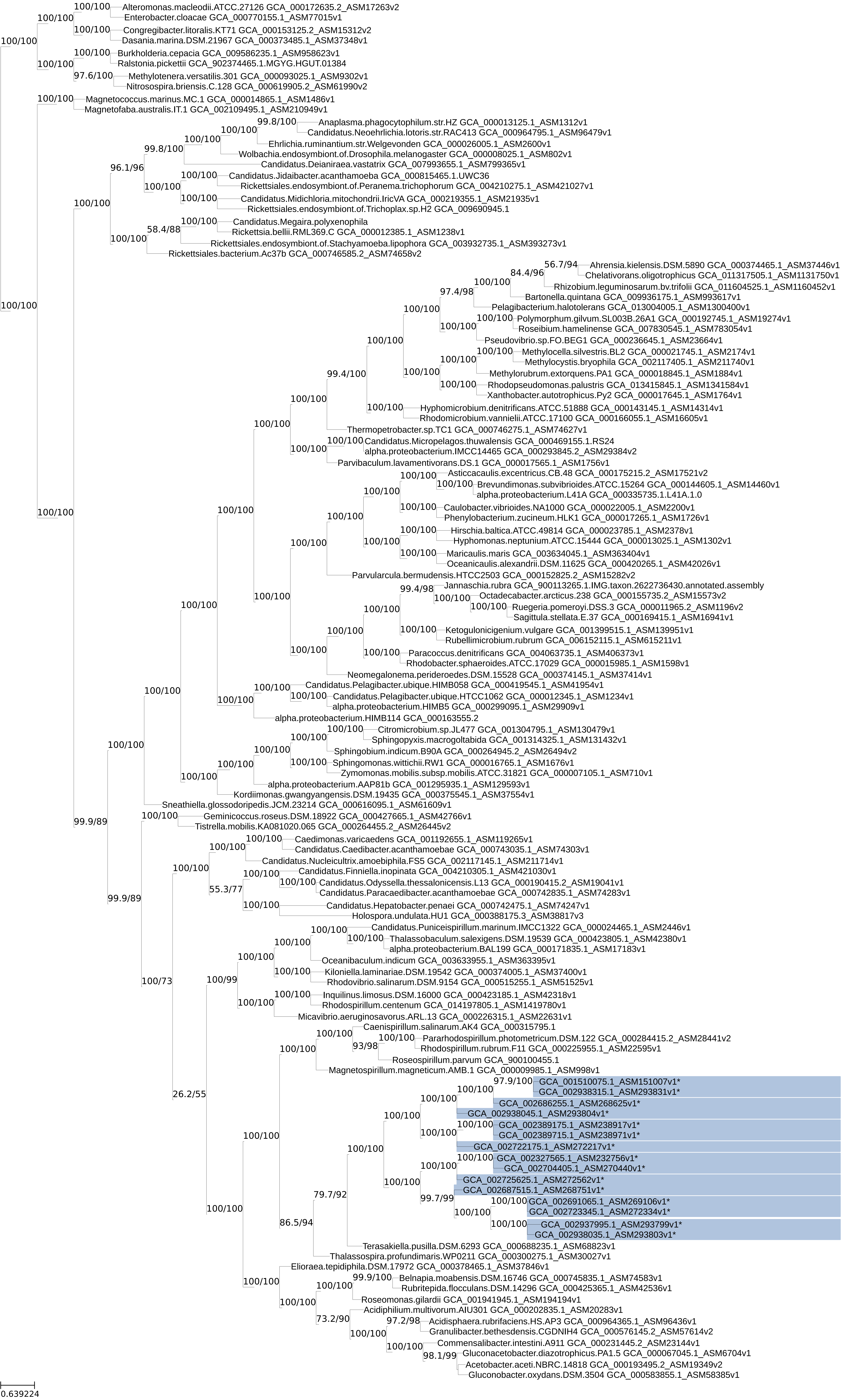
100/100
Alteromonas.macleodii.ATCC.27126 GCA_000172635.2_ASM17263v2
100/100
Enterobacter.cloacae GCA_000770155.1_ASM77015v1
100/100
Congregibacter.litoralis.KT71 GCA_000153125.2_ASM15312v2
100/100
Dasania.marina.DSM.21967 GCA_000373485.1_ASM37348v1
100/100
Burkholderia.cepacia GCA_009586235.1_ASM958623v1
100/100
Ralstonia.pickettii GCA_902374465.1.MGYG.HGUT.01384
97.6/100
Methylotenera.versatilis.301 GCA_000093025.1_ASM9302v1
Nitrosospira.briensis.C.128 GCA_000619905.2_ASM61990v2
100/100
Magnetococcus.marinus.MC.1 GCA_000014865.1_ASM1486v1
Magnetofaba.australis.IT.1 GCA_002109495.1_ASM210949v1
99.8/100
Anaplasma.phagocytophilum.str.HZ GCA_000013125.1_ASM1312v1
100/100
Candidatus.Neoehrlichia.lotoris.str.RAC413 GCA_000964795.1_ASM96479v1
100/100
Ehrlichia.ruminantium.str.Welgevonden GCA_000026005.1_ASM2600v1
99.8/100
Wolbachia.endosymbiont.of.Drosophila.melanogaster GCA_000008025.1_ASM802v1
Candidatus.Deianiraea.vastatrix GCA_007993655.1_ASM799365v1
96.1/96
100/100
Candidatus.Jidaibacter.acanthamoeba GCA_000815465.1.UWC36
100/100
Rickettsiales.endosymbiont.of.Peranema.trichophorum GCA_004210275.1_ASM421027v1
100/100
Candidatus.Midichloria.mitochondrii.IricVA GCA_000219355.1_ASM21935v1
100/100
Rickettsiales.endosymbiont.of.Trichoplax.sp.H2 GCA_009690945.1
100/100
Candidatus.Megaira.polyxenophila
58.4/88
Rickettsia.bellii.RML369.C GCA_000012385.1_ASM1238v1
100/100
Rickettsiales.endosymbiont.of.Stachyamoeba.lipophora GCA_003932735.1_ASM393273v1
Rickettsiales.bacterium.Ac37b GCA_000746585.2_ASM74658v2
56.7/94
Ahrensia.kielensis.DSM.5890 GCA_000374465.1_ASM37446v1
84.4/96
Chelativorans.oligotrophicus GCA_011317505.1_ASM1131750v1
100/100
Rhizobium.leguminosarum.bv.trifolii GCA_011604525.1_ASM1160452v1
97.4/98
Bartonella.quintana GCA_009936175.1_ASM993617v1
Pelagibacterium.halotolerans GCA_013004005.1_ASM1300400v1
100/100
100/100
Polymorphum.gilvum.SL003B.26A1 GCA_000192745.1_ASM19274v1
100/100
Roseibium.hamelinense GCA_007830545.1_ASM783054v1
100/100
Pseudovibrio.sp.FO.BEG1 GCA_000236645.1_ASM23664v1
100/100
Methylocella.silvestris.BL2 GCA_000021745.1_ASM2174v1
100/100
Methylocystis.bryophila GCA_002117405.1_ASM211740v1
99.4/100
100/100
Methylorubrum.extorquens.PA1 GCA_000018845.1_ASM1884v1
100/100
Rhodopseudomonas.palustris GCA_013415845.1_ASM1341584v1
Xanthobacter.autotrophicus.Py2 GCA_000017645.1_ASM1764v1
100/100
100/100
Hyphomicrobium.denitrificans.ATCC.51888 GCA_000143145.1_ASM14314v1
Rhodomicrobium.vannielii.ATCC.17100 GCA_000166055.1_ASM16605v1
100/100
Thermopetrobacter.sp.TC1 GCA_000746275.1_ASM74627v1
100/100
Candidatus.Micropelagos.thuwalensis GCA_000469155.1.RS24
100/100
alpha.proteobacterium.IMCC14465 GCA_000293845.2_ASM29384v2
Parvibaculum.lavamentivorans.DS.1 GCA_000017565.1_ASM1756v1
100/100
Asticcacaulis.excentricus.CB.48 GCA_000175215.2_ASM17521v2
100/100
Brevundimonas.subvibrioides.ATCC.15264 GCA_000144605.1_ASM14460v1
100/100
alpha.proteobacterium.L41A GCA_000335735.1.L41A.1.0
100/100
Caulobacter.vibrioides.NA1000 GCA_000022005.1_ASM2200v1
100/100
Phenylobacterium.zucineum.HLK1 GCA_000017265.1_ASM1726v1
100/100
Hirschia.baltica.ATCC.49814 GCA_000023785.1_ASM2378v1
100/100
Hyphomonas.neptunium.ATCC.15444 GCA_000013025.1_ASM1302v1
100/100
Maricaulis.maris GCA_003634045.1_ASM363404v1
100/100
Oceanicaulis.alexandrii.DSM.11625 GCA_000420265.1_ASM42026v1
Parvularcula.bermudensis.HTCC2503 GCA_000152825.2_ASM15282v2
99.4/98
Jannaschia.rubra GCA_900113265.1.IMG.taxon.2622736430.annotated.assembly
100/100
Octadecabacter.arcticus.238 GCA_000155735.2_ASM15573v2
100/100
100/100
Ruegeria.pomeroyi.DSS.3 GCA_000011965.2_ASM1196v2
Sagittula.stellata.E.37 GCA_000169415.1_ASM16941v1
100/100
Ketogulonicigenium.vulgare GCA_001399515.1_ASM139951v1
100/100
Rubellimicrobium.rubrum GCA_006152115.1_ASM615211v1
100/100
Paracoccus.denitrificans GCA_004063735.1_ASM406373v1
100/100
Rhodobacter.sphaeroides.ATCC.17029 GCA_000015985.1_ASM1598v1
Neomegalonema.perideroedes.DSM.15528 GCA_000374145.1_ASM37414v1
100/100
Candidatus.Pelagibacter.ubique.HIMB058 GCA_000419545.1_ASM41954v1
100/100
100/100
Candidatus.Pelagibacter.ubique.HTCC1062 GCA_000012345.1_ASM1234v1
100/100
alpha.proteobacterium.HIMB5 GCA_000299095.1_ASM29909v1
alpha.proteobacterium.HIMB114 GCA_000163555.2
100/100
Citromicrobium.sp.JL477 GCA_001304795.1_ASM130479v1
100/100
Sphingopyxis.macrogoltabida GCA_001314325.1_ASM131432v1
100/100
Sphingobium.indicum.B90A GCA_000264945.2_ASM26494v2
100/100
100/100
Sphingomonas.wittichii.RW1 GCA_000016765.1_ASM1676v1
100/100
Zymomonas.mobilis.subsp.mobilis.ATCC.31821 GCA_000007105.1_ASM710v1
100/100
alpha.proteobacterium.AAP81b GCA_001295935.1_ASM129593v1
Kordiimonas.gwangyangensis.DSM.19435 GCA_000375545.1_ASM37554v1
Sneathiella.glossodoripedis.JCM.23214 GCA_000616095.1_ASM61609v1
100/100
Geminicoccus.roseus.DSM.18922 GCA_000427665.1_ASM42766v1
Tistrella.mobilis.KA081020.065 GCA_000264455.2_ASM26445v2
99.9/89
100/100
Caedimonas.varicaedens GCA_001192655.1_ASM119265v1
100/100
Candidatus.Caedibacter.acanthamoebae GCA_000743035.1_ASM74303v1
100/100
Candidatus.Nucleicultrix.amoebiphila.FS5 GCA_002117145.1_ASM211714v1
100/100
Candidatus.Finniella.inopinata GCA_004210305.1_ASM421030v1
100/100
Candidatus.Odyssella.thessalonicensis.L13 GCA_000190415.2_ASM19041v1
55.3/77
Candidatus.Paracaedibacter.acanthamoebae GCA_000742835.1_ASM74283v1
99.9/89
100/100
Candidatus.Hepatobacter.penaei GCA_000742475.1_ASM74247v1
Holospora.undulata.HU1 GCA_000388175.3_ASM38817v3
100/100
Candidatus.Puniceispirillum.marinum.IMCC1322 GCA_000024465.1_ASM2446v1
100/100
Thalassobaculum.salexigens.DSM.19539 GCA_000423805.1_ASM42380v1
100/100
alpha.proteobacterium.BAL199 GCA_000171835.1_ASM17183v1
100/100
Oceanibaculum.indicum GCA_003633955.1_ASM363395v1
100/100
Kiloniella.laminariae.DSM.19542 GCA_000374005.1_ASM37400v1
100/73
100/99
Rhodovibrio.salinarum.DSM.9154 GCA_000515255.1_ASM51525v1
100/100
Inquilinus.limosus.DSM.16000 GCA_000423185.1_ASM42318v1
100/100
Rhodospirillum.centenum GCA_014197805.1_ASM1419780v1
Micavibrio.aeruginosavorus.ARL.13 GCA_000226315.1_ASM22631v1
100/100
Caenispirillum.salinarum.AK4 GCA_000315795.1
100/100
100/100
Pararhodospirillum.photometricum.DSM.122 GCA_000284415.2_ASM28441v2
93/98
Rhodospirillum.rubrum.F11 GCA_000225955.1_ASM22595v1
Roseospirillum.parvum GCA_900100455.1
Magnetospirillum.magneticum.AMB.1 GCA_000009985.1_ASM998v1
97.9/100
GCA_001510075.1_ASM151007v1*
100/100
GCA_002938315.1_ASM293831v1*
100/100
GCA_002686255.1_ASM268625v1*
26.2/55
GCA_002938045.1_ASM293804v1*
100/100
100/100
GCA_002389175.1_ASM238917v1*
100/100
GCA_002389715.1_ASM238971v1*
100/100
GCA_002722175.1_ASM272217v1*
100/100
GCA_002327565.1_ASM232756v1*
100/100
100/100
GCA_002704405.1_ASM270440v1*
100/100
GCA_002725625.1_ASM272562v1*
GCA_002687515.1_ASM268751v1*
79.7/92
99.7/99
100/100
GCA_002691065.1_ASM269106v1*
100/100
GCA_002723345.1_ASM272334v1*
100/100
86.5/94
100/100
GCA_002937995.1_ASM293799v1*
GCA_002938035.1_ASM293803v1*
Terasakiella.pusilla.DSM.6293 GCA_000688235.1_ASM68823v1
Thalassospira.profundimaris.WP0211 GCA_000300275.1_ASM30027v1
Elioraea.tepidiphila.DSM.17972 GCA_000378465.1_ASM37846v1
99.9/100
Belnapia.moabensis.DSM.16746 GCA_000745835.1_ASM74583v1
100/100
100/100
Rubritepida.flocculans.DSM.14296 GCA_000425365.1_ASM42536v1
Roseomonas.gilardii GCA_001941945.1_ASM194194v1
100/100
Acidiphilium.multivorum.AIU301 GCA_000202835.1_ASM20283v1
97.2/98
73.2/90
Acidisphaera.rubrifaciens.HS.AP3 GCA_000964365.1_ASM96436v1
100/100
Granulibacter.bethesdensis.CGDNIH4 GCA_000576145.2_ASM57614v2
100/100
Commensalibacter.intestini.A911 GCA_000231445.2_ASM23144v1
98.1/99
Gluconacetobacter.diazotrophicus.PA1.5 GCA_000067045.1_ASM6704v1
Acetobacter.aceti.NBRC.14818 GCA_000193495.2_ASM19349v2
Gluconobacter.oxydans.DSM.3504 GCA_000583855.1_ASM58385v1
0.639224

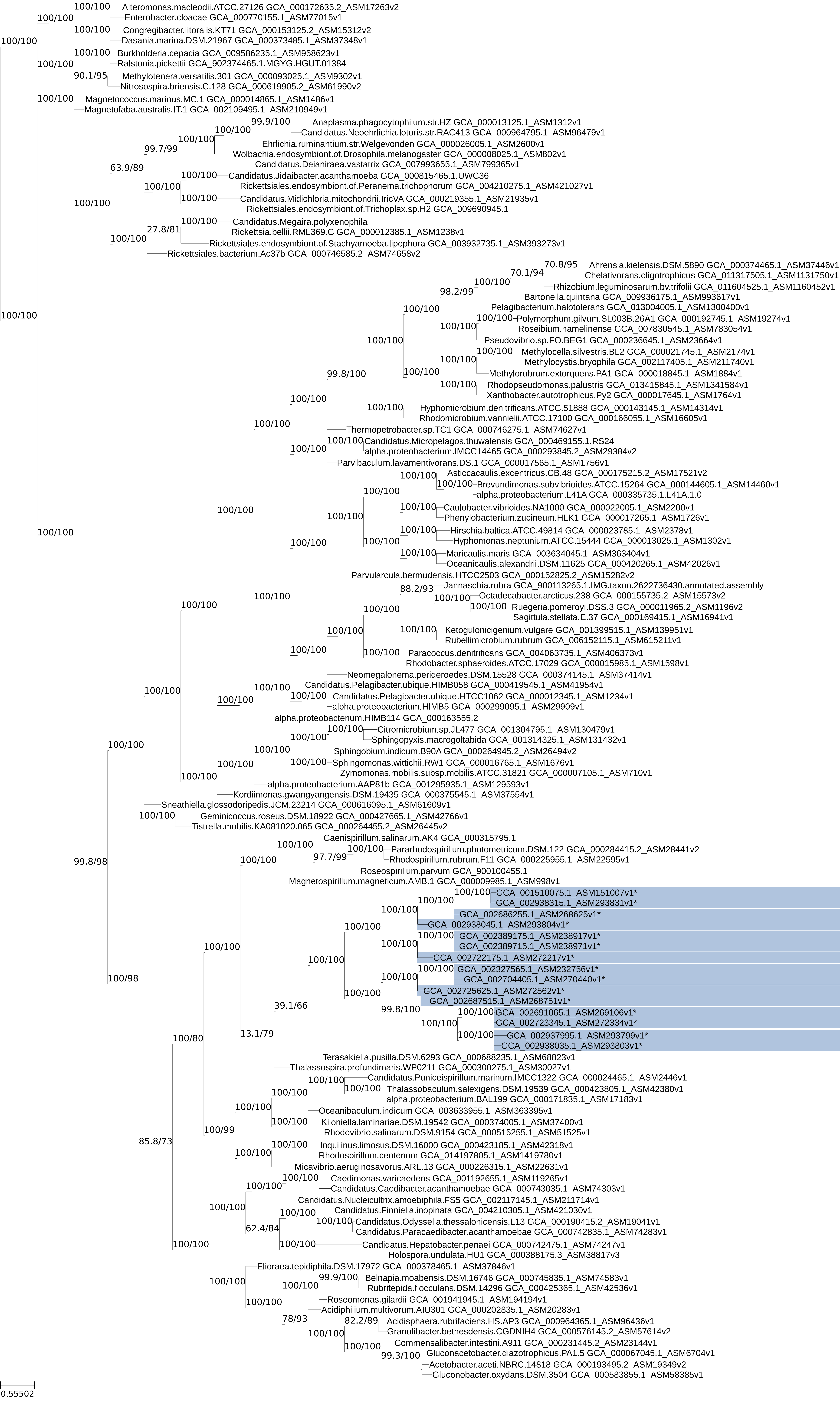
Alteromonas.macleodii.ATCC.27126 GCA_000172635.2_ASM17263v2
Enterobacter.cloacae GCA_000770155.1_ASM77015v1
Congregibacter.litoralis.KT71 GCA_000153125.2_ASM15312v2
Dasania.marina.DSM.21967 GCA_000373485.1_ASM37348v1
Burkholderia.cepacia GCA_009586235.1_ASM958623v1
Ralstonia.pickettii GCA_902374465.1.MGYG.HGUT.01384
Methylotenera.versatilis.301 GCA_000093025.1_ASM9302v1
Nitrosospira.briensis.C.128 GCA_000619905.2_ASM61990v2
Magnetococcus.marinus.MC.1 GCA_000014865.1_ASM1486v1
Magnetofaba.australis.IT.1 GCA_002109495.1_ASM210949v1
Anaplasma.phagocytophilum.str.HZ GCA_000013125.1_ASM1312v1
Candidatus.Neoehrlichia.lotoris.str.RAC413 GCA_000964795.1_ASM96479v1
Ehrlichia.ruminantium.str.Welgevonden GCA_000026005.1_ASM2600v1
Wolbachia.endosymbiont.of.Drosophila.melanogaster GCA_000008025.1_ASM802v1
Candidatus.Deianiraea.vastatrix GCA_007993655.1_ASM799365v1
Candidatus.Jidaibacter.acanthamoeba GCA_000815465.1.UWC36
Rickettsiales.endosymbiont.of.Peranema.trichophorum GCA_004210275.1_ASM421027v1
Candidatus.Midichloria.mitochondrii.IricVA GCA_000219355.1_ASM21935v1
Rickettsiales.endosymbiont.of.Trichoplax.sp.H2 GCA_009690945.1
Candidatus.Megaira.polyxenophila
Rickettsia.bellii.RML369.C GCA_000012385.1_ASM1238v1
Rickettsiales.endosymbiont.of.Stachyamoeba.lipophora GCA_003932735.1_ASM393273v1
Rickettsiales.bacterium.Ac37b GCA_000746585.2_ASM74658v2
Ahrensia.kielensis.DSM.5890 GCA_000374465.1_ASM37446v1
Chelativorans.oligotrophicus GCA_011317505.1_ASM1131750v1
Rhizobium.leguminosarum.bv.trifolii GCA_011604525.1_ASM1160452v1
Bartonella.quintana GCA_009936175.1_ASM993617v1
Pelagibacterium.halotolerans GCA_013004005.1_ASM1300400v1
Polymorphum.gilvum.SL003B.26A1 GCA_000192745.1_ASM19274v1
Roseibium.hamelinense GCA_007830545.1_ASM783054v1
Pseudovibrio.sp.FO.BEG1 GCA_000236645.1_ASM23664v1
Methylocella.silvestris.BL2 GCA_000021745.1_ASM2174v1
Methylocystis.bryophila GCA_002117405.1_ASM211740v1
Methylorubrum.extorquens.PA1 GCA_000018845.1_ASM1884v1
Rhodopseudomonas.palustris GCA_013415845.1_ASM1341584v1
Xanthobacter.autotrophicus.Py2 GCA_000017645.1_ASM1764v1
Hyphomicrobium.denitrificans.ATCC.51888 GCA_000143145.1_ASM14314v1
Rhodomicrobium.vannielii.ATCC.17100 GCA_000166055.1_ASM16605v1
Thermopetrobacter.sp.TC1 GCA_000746275.1_ASM74627v1
Candidatus.Micropelagos.thuwalensis GCA_000469155.1.RS24
alpha.proteobacterium.IMCC14465 GCA_000293845.2_ASM29384v2
Parvibaculum.lavamentivorans.DS.1 GCA_000017565.1_ASM1756v1
Asticcacaulis.excentricus.CB.48 GCA_000175215.2_ASM17521v2
Brevundimonas.subvibrioides.ATCC.15264 GCA_000144605.1_ASM14460v1
alpha.proteobacterium.L41A GCA_000335735.1.L41A.1.0
Caulobacter.vibrioides.NA1000 GCA_000022005.1_ASM2200v1
Phenylobacterium.zucineum.HLK1 GCA_000017265.1_ASM1726v1
Hirschia.baltica.ATCC.49814 GCA_000023785.1_ASM2378v1
Hyphomonas.neptunium.ATCC.15444 GCA_000013025.1_ASM1302v1
Maricaulis.maris GCA_003634045.1_ASM363404v1
Oceanicaulis.alexandrii.DSM.11625 GCA_000420265.1_ASM42026v1
Parvularcula.bermudensis.HTCC2503 GCA_000152825.2_ASM15282v2
Jannaschia.rubra GCA_900113265.1.IMG.taxon.2622736430.annotated.assembly
Octadecabacter.arcticus.238 GCA_000155735.2_ASM15573v2
Ruegeria.pomeroyi.DSS.3 GCA_000011965.2_ASM1196v2
Sagittula.stellata.E.37 GCA_000169415.1_ASM16941v1
Ketogulonicigenium.vulgare GCA_001399515.1_ASM139951v1
Rubellimicrobium.rubrum GCA_006152115.1_ASM615211v1
Paracoccus.denitrificans GCA_004063735.1_ASM406373v1
Rhodobacter.sphaeroides.ATCC.17029 GCA_000015985.1_ASM1598v1
Neomegalonema.perideroedes.DSM.15528 GCA_000374145.1_ASM37414v1
Candidatus.Pelagibacter.ubique.HIMB058 GCA_000419545.1_ASM41954v1
Candidatus.Pelagibacter.ubique.HTCC1062 GCA_000012345.1_ASM1234v1
alpha.proteobacterium.HIMB5 GCA_000299095.1_ASM29909v1
alpha.proteobacterium.HIMB114 GCA_000163555.2
Citromicrobium.sp.JL477 GCA_001304795.1_ASM130479v1
Sphingopyxis.macrogoltabida GCA_001314325.1_ASM131432v1
Sphingobium.indicum.B90A GCA_000264945.2_ASM26494v2
Sphingomonas.wittichii.RW1 GCA_000016765.1_ASM1676v1
Zymomonas.mobilis.subsp.mobilis.ATCC.31821 GCA_000007105.1_ASM710v1
alpha.proteobacterium.AAP81b GCA_001295935.1_ASM129593v1
Kordiimonas.gwangyangensis.DSM.19435 GCA_000375545.1_ASM37554v1
Sneathiella.glossodoripedis.JCM.23214 GCA_000616095.1_ASM61609v1
Geminicoccus.roseus.DSM.18922 GCA_000427665.1_ASM42766v1
Tistrella.mobilis.KA081020.065 GCA_000264455.2_ASM26445v2
Caenispirillum.salinarum.AK4 GCA_000315795.1
Pararhodospirillum.photometricum.DSM.122 GCA_000284415.2_ASM28441v2
Rhodospirillum.rubrum.F11 GCA_000225955.1_ASM22595v1
Roseospirillum.parvum GCA_900100455.1
Magnetospirillum.magneticum.AMB.1 GCA_000009985.1_ASM998v1
GCA_001510075.1_ASM151007v1*
GCA_002938315.1_ASM293831v1*
GCA_002686255.1_ASM268625v1*
GCA_002938045.1_ASM293804v1*
GCA_002389175.1_ASM238917v1*
GCA_002389715.1_ASM238971v1*
GCA_002722175.1_ASM272217v1*
GCA_002327565.1_ASM232756v1*
GCA_002704405.1_ASM270440v1*
GCA_002725625.1_ASM272562v1*
GCA_002687515.1_ASM268751v1*
GCA_002691065.1_ASM269106v1*
GCA_002723345.1_ASM272334v1*
GCA_002937995.1_ASM293799v1*
GCA_002938035.1_ASM293803v1*
Terasakiella.pusilla.DSM.6293 GCA_000688235.1_ASM68823v1
Thalassospira.profundimaris.WP0211 GCA_000300275.1_ASM30027v1
Candidatus.Puniceispirillum.marinum.IMCC1322 GCA_000024465.1_ASM2446v1
Thalassobaculum.salexigens.DSM.19539 GCA_000423805.1_ASM42380v1
alpha.proteobacterium.BAL199 GCA_000171835.1_ASM17183v1
Oceanibaculum.indicum GCA_003633955.1_ASM363395v1
Kiloniella.laminariae.DSM.19542 GCA_000374005.1_ASM37400v1
Rhodovibrio.salinarum.DSM.9154 GCA_000515255.1_ASM51525v1
Inquilinus.limosus.DSM.16000 GCA_000423185.1_ASM42318v1
Rhodospirillum.centenum GCA_014197805.1_ASM1419780v1
Micavibrio.aeruginosavorus.ARL.13 GCA_000226315.1_ASM22631v1
Caedimonas.varicaedens GCA_001192655.1_ASM119265v1
Candidatus.Caedibacter.acanthamoebae GCA_000743035.1_ASM74303v1
Candidatus.Nucleicultrix.amoebiphila.FS5 GCA_002117145.1_ASM211714v1
Candidatus.Finniella.inopinata GCA_004210305.1_ASM421030v1
Candidatus.Odyssella.thessalonicensis.L13 GCA_000190415.2_ASM19041v1
Candidatus.Paracaedibacter.acanthamoebae GCA_000742835.1_ASM74283v1
Candidatus.Hepatobacter.penaei GCA_000742475.1_ASM74247v1
Holospora.undulata.HU1 GCA_000388175.3_ASM38817v3
Elioraea.tepidiphila.DSM.17972 GCA_000378465.1_ASM37846v1
Belnapia.moabensis.DSM.16746 GCA_000745835.1_ASM74583v1
Rubritepida.flocculans.DSM.14296 GCA_000425365.1_ASM42536v1
Roseomonas.gilardii GCA_001941945.1_ASM194194v1
Acidiphilium.multivorum.AIU301 GCA_000202835.1_ASM20283v1
Acidisphaera.rubrifaciens.HS.AP3 GCA_000964365.1_ASM96436v1
Granulibacter.bethesdensis.CGDNIH4 GCA_000576145.2_ASM57614v2
Commensalibacter.intestini.A911 GCA_000231445.2_ASM23144v1
Gluconacetobacter.diazotrophicus.PA1.5 GCA_000067045.1_ASM6704v1
Acetobacter.aceti.NBRC.14818 GCA_000193495.2_ASM19349v2
Gluconobacter.oxydans.DSM.3504 GCA_000583855.1_ASM58385v1
0.55502

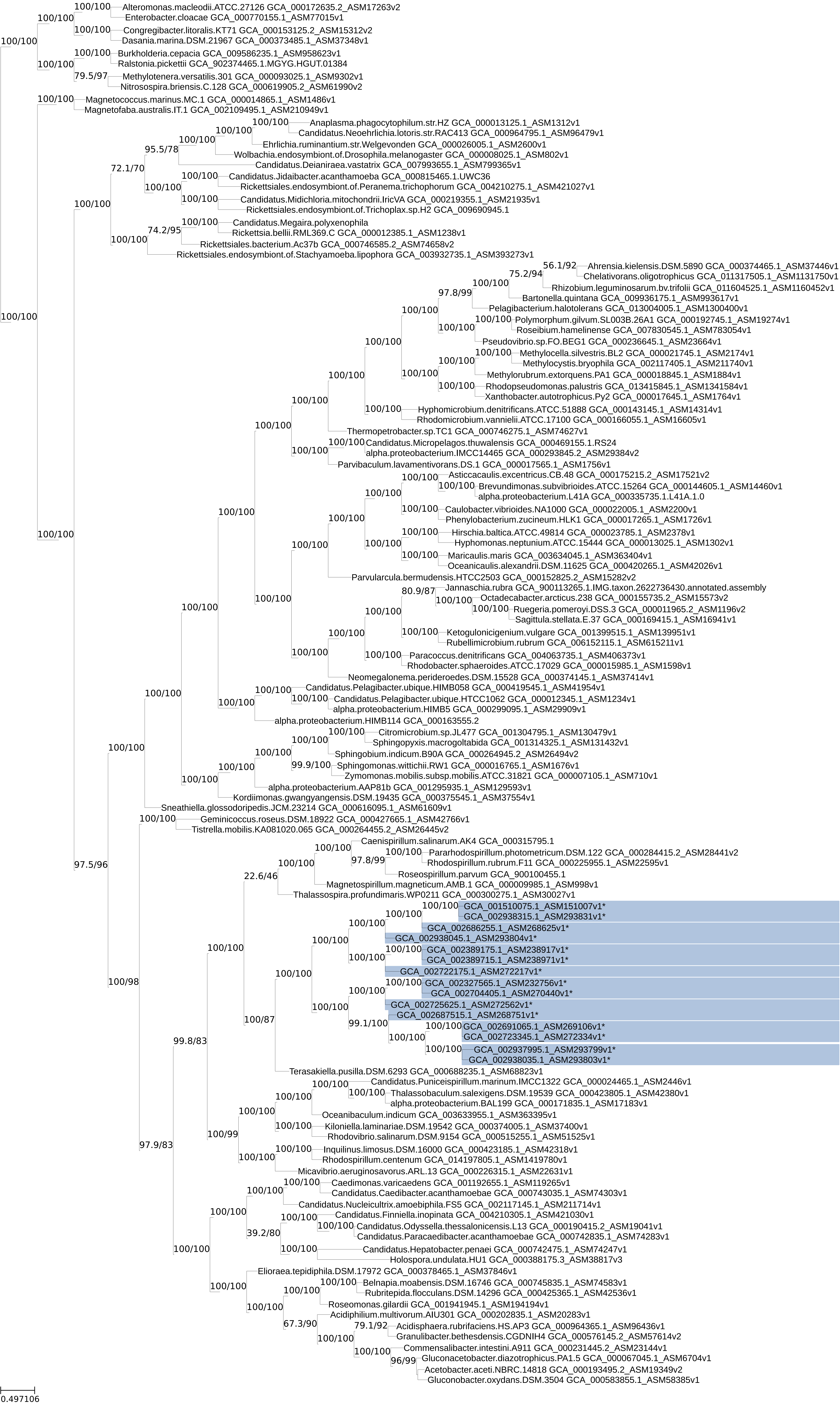
100/100
Alteromonas.macleodii.ATCC.27126 GCA_000172635.2_ASM17263v2
100/100
Enterobacter.cloacae GCA_000770155.1_ASM77015v1
100/100
100/100
Congregibacter.litoralis.KT71 GCA_000153125.2_ASM15312v2
Dasania.marina.DSM.21967 GCA_000373485.1_ASM37348v1
100/100
Burkholderia.cepacia GCA_009586235.1_ASM958623v1
100/100
Ralstonia.pickettii GCA_902374465.1.MGYG.HGUT.01384
79.5/97
Methylotenera.versatilis.301 GCA_000093025.1_ASM9302v1
Nitrosospira.briensis.C.128 GCA_000619905.2_ASM61990v2
100/100
Magnetococcus.marinus.MC.1 GCA_000014865.1_ASM1486v1
Magnetofaba.australis.IT.1 GCA_002109495.1_ASM210949v1
100/100
Anaplasma.phagocytophilum.str.HZ GCA_000013125.1_ASM1312v1
100/100
Candidatus.Neoehrlichia.lotoris.str.RAC413 GCA_000964795.1_ASM96479v1
100/100
Ehrlichia.ruminantium.str.Welgevonden GCA_000026005.1_ASM2600v1
95.5/78
Wolbachia.endosymbiont.of.Drosophila.melanogaster GCA_000008025.1_ASM802v1
Candidatus.Deianiraea.vastatrix GCA_007993655.1_ASM799365v1
72.1/70
100/100
Candidatus.Jidaibacter.acanthamoeba GCA_000815465.1.UWC36
100/100
Rickettsiales.endosymbiont.of.Peranema.trichophorum GCA_004210275.1_ASM421027v1
100/100
100/100
Candidatus.Midichloria.mitochondrii.IricVA GCA_000219355.1_ASM21935v1
Rickettsiales.endosymbiont.of.Trichoplax.sp.H2 GCA_009690945.1
100/100
Candidatus.Megaira.polyxenophila
74.2/95
Rickettsia.bellii.RML369.C GCA_000012385.1_ASM1238v1
100/100
Rickettsiales.bacterium.Ac37b GCA_000746585.2_ASM74658v2
Rickettsiales.endosymbiont.of.Stachyamoeba.lipophora GCA_003932735.1_ASM393273v1
56.1/92
Ahrensia.kielensis.DSM.5890 GCA_000374465.1_ASM37446v1
75.2/94
Chelativorans.oligotrophicus GCA_011317505.1_ASM1131750v1
100/100
Rhizobium.leguminosarum.bv.trifolii GCA_011604525.1_ASM1160452v1
97.8/99
Bartonella.quintana GCA_009936175.1_ASM993617v1
100/100
Pelagibacterium.halotolerans GCA_013004005.1_ASM1300400v1
100/100
Polymorphum.gilvum.SL003B.26A1 GCA_000192745.1_ASM19274v1
100/100
Roseibium.hamelinense GCA_007830545.1_ASM783054v1
100/100
Pseudovibrio.sp.FO.BEG1 GCA_000236645.1_ASM23664v1
100/100
Methylocella.silvestris.BL2 GCA_000021745.1_ASM2174v1
100/100
Methylocystis.bryophila GCA_002117405.1_ASM211740v1
100/100
Methylorubrum.extorquens.PA1 GCA_000018845.1_ASM1884v1
100/100
Rhodopseudomonas.palustris GCA_013415845.1_ASM1341584v1
100/100
Xanthobacter.autotrophicus.Py2 GCA_000017645.1_ASM1764v1
100/100
Hyphomicrobium.denitrificans.ATCC.51888 GCA_000143145.1_ASM14314v1
100/100
Rhodomicrobium.vannielii.ATCC.17100 GCA_000166055.1_ASM16605v1
Thermopetrobacter.sp.TC1 GCA_000746275.1_ASM74627v1
100/100
Candidatus.Micropelagos.thuwalensis GCA_000469155.1.RS24
100/100
alpha.proteobacterium.IMCC14465 GCA_000293845.2_ASM29384v2
Parvibaculum.lavamentivorans.DS.1 GCA_000017565.1_ASM1756v1
100/100
Asticcacaulis.excentricus.CB.48 GCA_000175215.2_ASM17521v2
100/100
Brevundimonas.subvibrioides.ATCC.15264 GCA_000144605.1_ASM14460v1
100/100
alpha.proteobacterium.L41A GCA_000335735.1.L41A.1.0
100/100
Caulobacter.vibrioides.NA1000 GCA_000022005.1_ASM2200v1
100/100
Phenylobacterium.zucineum.HLK1 GCA_000017265.1_ASM1726v1
100/100
Hirschia.baltica.ATCC.49814 GCA_000023785.1_ASM2378v1
100/100
100/100
Hyphomonas.neptunium.ATCC.15444 GCA_000013025.1_ASM1302v1
100/100
Maricaulis.maris GCA_003634045.1_ASM363404v1
100/100
Oceanicaulis.alexandrii.DSM.11625 GCA_000420265.1_ASM42026v1
Parvularcula.bermudensis.HTCC2503 GCA_000152825.2_ASM15282v2
100/100
Jannaschia.rubra GCA_900113265.1.IMG.taxon.2622736430.annotated.assembly
80.9/87
100/100
Octadecabacter.arcticus.238 GCA_000155735.2_ASM15573v2
100/100
Ruegeria.pomeroyi.DSS.3 GCA_000011965.2_ASM1196v2
100/100
Sagittula.stellata.E.37 GCA_000169415.1_ASM16941v1
100/100
Ketogulonicigenium.vulgare GCA_001399515.1_ASM139951v1
100/100
Rubellimicrobium.rubrum GCA_006152115.1_ASM615211v1
100/100
Paracoccus.denitrificans GCA_004063735.1_ASM406373v1
100/100
Rhodobacter.sphaeroides.ATCC.17029 GCA_000015985.1_ASM1598v1
Neomegalonema.perideroedes.DSM.15528 GCA_000374145.1_ASM37414v1
100/100
Candidatus.Pelagibacter.ubique.HIMB058 GCA_000419545.1_ASM41954v1
100/100
Candidatus.Pelagibacter.ubique.HTCC1062 GCA_000012345.1_ASM1234v1
100/100
alpha.proteobacterium.HIMB5 GCA_000299095.1_ASM29909v1
100/100
alpha.proteobacterium.HIMB114 GCA_000163555.2
100/100
100/100
Citromicrobium.sp.JL477 GCA_001304795.1_ASM130479v1
100/100
Sphingopyxis.macrogoltabida GCA_001314325.1_ASM131432v1
100/100
Sphingobium.indicum.B90A GCA_000264945.2_ASM26494v2
100/100
99.9/100
Sphingomonas.wittichii.RW1 GCA_000016765.1_ASM1676v1
100/100
Zymomonas.mobilis.subsp.mobilis.ATCC.31821 GCA_000007105.1_ASM710v1
100/100
alpha.proteobacterium.AAP81b GCA_001295935.1_ASM129593v1
Kordiimonas.gwangyangensis.DSM.19435 GCA_000375545.1_ASM37554v1
Sneathiella.glossodoripedis.JCM.23214 GCA_000616095.1_ASM61609v1
100/100
Geminicoccus.roseus.DSM.18922 GCA_000427665.1_ASM42766v1
100/100
Tistrella.mobilis.KA081020.065 GCA_000264455.2_ASM26445v2
Caenispirillum.salinarum.AK4 GCA_000315795.1
100/100
100/100
Pararhodospirillum.photometricum.DSM.122 GCA_000284415.2_ASM28441v2
97.8/99
Rhodospirillum.rubrum.F11 GCA_000225955.1_ASM22595v1
100/100
Roseospirillum.parvum GCA_900100455.1
22.6/46
Magnetospirillum.magneticum.AMB.1 GCA_000009985.1_ASM998v1
Thalassospira.profundimaris.WP0211 GCA_000300275.1_ASM30027v1
100/100
GCA_001510075.1_ASM151007v1*
100/100
GCA_002938315.1_ASM293831v1*
100/100
GCA_002686255.1_ASM268625v1*
GCA_002938045.1_ASM293804v1*
100/100
100/100
GCA_002389175.1_ASM238917v1*
100/100
GCA_002389715.1_ASM238971v1*
100/100
GCA_002722175.1_ASM272217v1*
97.5/96
100/100
GCA_002327565.1_ASM232756v1*
100/100
GCA_002704405.1_ASM270440v1*
100/100
GCA_002725625.1_ASM272562v1*
100/98
GCA_002687515.1_ASM268751v1*
100/87
99.1/100
100/100
GCA_002691065.1_ASM269106v1*
100/100
GCA_002723345.1_ASM272334v1*
100/100
99.8/83
GCA_002937995.1_ASM293799v1*
GCA_002938035.1_ASM293803v1*
Terasakiella.pusilla.DSM.6293 GCA_000688235.1_ASM68823v1
100/100
Candidatus.Puniceispirillum.marinum.IMCC1322 GCA_000024465.1_ASM2446v1
100/100
Thalassobaculum.salexigens.DSM.19539 GCA_000423805.1_ASM42380v1
100/100
alpha.proteobacterium.BAL199 GCA_000171835.1_ASM17183v1
100/100
Oceanibaculum.indicum GCA_003633955.1_ASM363395v1
100/100
Kiloniella.laminariae.DSM.19542 GCA_000374005.1_ASM37400v1
100/99
Rhodovibrio.salinarum.DSM.9154 GCA_000515255.1_ASM51525v1
97.9/83
100/100
Inquilinus.limosus.DSM.16000 GCA_000423185.1_ASM42318v1
100/100
Rhodospirillum.centenum GCA_014197805.1_ASM1419780v1
Micavibrio.aeruginosavorus.ARL.13 GCA_000226315.1_ASM22631v1
100/100
Caedimonas.varicaedens GCA_001192655.1_ASM119265v1
100/100
Candidatus.Caedibacter.acanthamoebae GCA_000743035.1_ASM74303v1
100/100
Candidatus.Nucleicultrix.amoebiphila.FS5 GCA_002117145.1_ASM211714v1
100/100
Candidatus.Finniella.inopinata GCA_004210305.1_ASM421030v1
100/100
Candidatus.Odyssella.thessalonicensis.L13 GCA_000190415.2_ASM19041v1
39.2/80
Candidatus.Paracaedibacter.acanthamoebae GCA_000742835.1_ASM74283v1
100/100
100/100
Candidatus.Hepatobacter.penaei GCA_000742475.1_ASM74247v1
Holospora.undulata.HU1 GCA_000388175.3_ASM38817v3
Elioraea.tepidiphila.DSM.17972 GCA_000378465.1_ASM37846v1
100/100
Belnapia.moabensis.DSM.16746 GCA_000745835.1_ASM74583v1
100/100
100/100
Rubritepida.flocculans.DSM.14296 GCA_000425365.1_ASM42536v1
100/100
Roseomonas.gilardii GCA_001941945.1_ASM194194v1
Acidiphilium.multivorum.AIU301 GCA_000202835.1_ASM20283v1
67.3/90
79.1/92
Acidisphaera.rubrifaciens.HS.AP3 GCA_000964365.1_ASM96436v1
100/100
Granulibacter.bethesdensis.CGDNIH4 GCA_000576145.2_ASM57614v2
100/100
Commensalibacter.intestini.A911 GCA_000231445.2_ASM23144v1
96/99
Gluconacetobacter.diazotrophicus.PA1.5 GCA_000067045.1_ASM6704v1
Acetobacter.aceti.NBRC.14818 GCA_000193495.2_ASM19349v2
Gluconobacter.oxydans.DSM.3504 GCA_000583855.1_ASM58385v1
0.497106

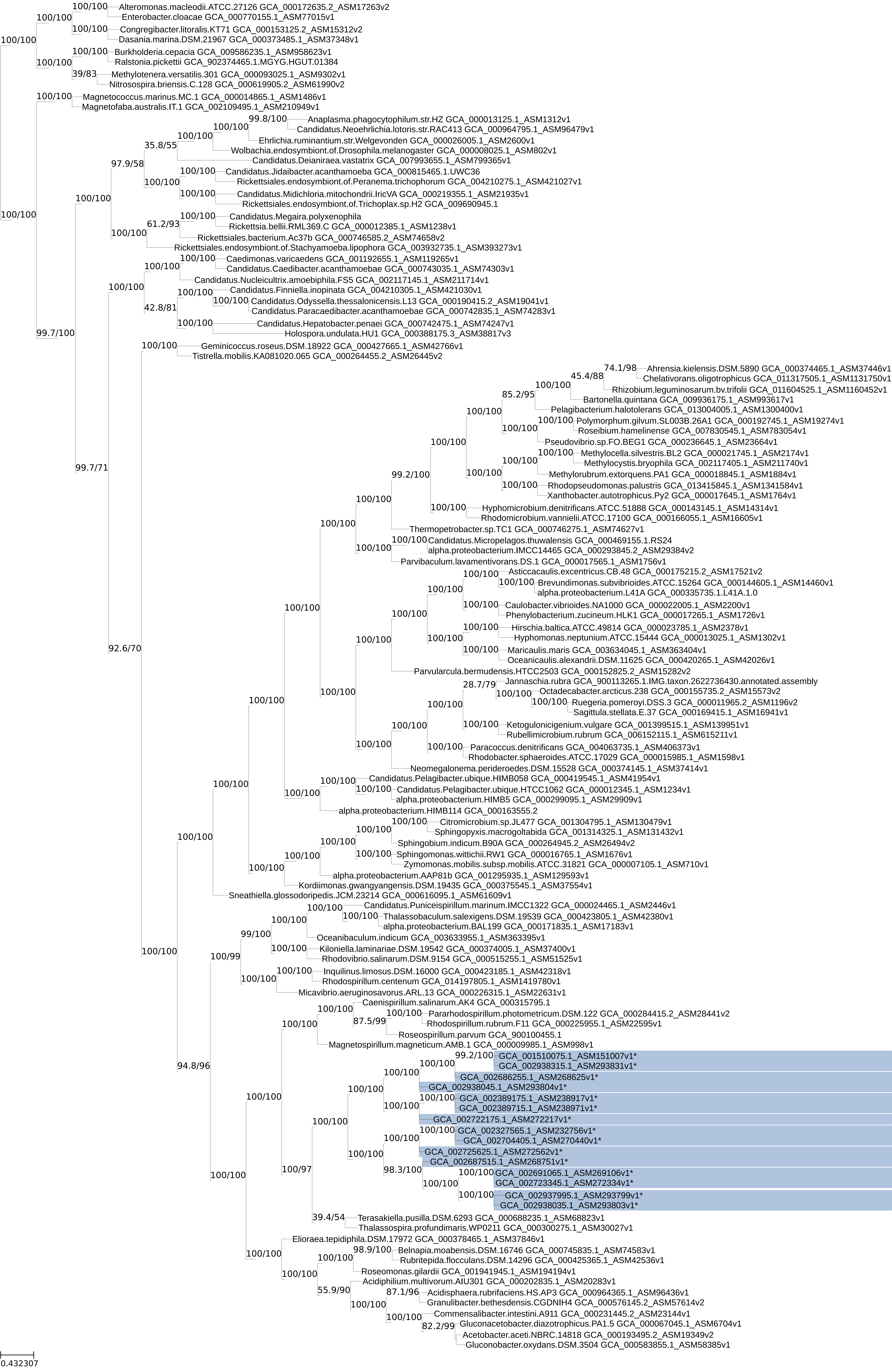

Alteromonas.macleodii.ATCC.27126 GCA_000172635.2_ASM17263v2
Enterobacter.cloacae GCA_000770155.1_ASM77015v1
Congregibacter.litoralis.KT71 GCA_000153125.2_ASM15312v2
Dasania.marina.DSM.21967 GCA_000373485.1_ASM37348v1
Burkholderia.cepacia GCA_009586235.1_ASM958623v1
Ralstonia.pickettii GCA_902374465.1.MGYG.HGUT.01384
Methylotenera.versatilis.301 GCA_000093025.1_ASM9302v1
Nitrosospira.briensis.C.128 GCA_000619905.2_ASM61990v2
Magnetococcus.marinus.MC.1 GCA_000014865.1_ASM1486v1
Magnetofaba.australis.IT.1 GCA_002109495.1_ASM210949v1
Anaplasma.phagocytophilum.str.HZ GCA_000013125.1_ASM1312v1
Candidatus.Neoehrlichia.lotoris.str.RAC413 GCA_000964795.1_ASM96479v1
Ehrlichia.ruminantium.str.Welgevonden GCA_000026005.1_ASM2600v1
Wolbachia.endosymbiont.of.Drosophila.melanogaster GCA_000008025.1_ASM802v1
Candidatus.Deianiraea.vastatrix GCA_007993655.1_ASM799365v1
Candidatus.Jidaibacter.acanthamoeba GCA_000815465.1.UWC36
Rickettsiales.endosymbiont.of.Peranema.trichophorum GCA_004210275.1_ASM421027v1
Candidatus.Midichloria.mitochondrii.IricVA GCA_000219355.1_ASM21935v1
Rickettsiales.endosymbiont.of.Trichoplax.sp.H2 GCA_009690945.1
Candidatus.Megaira.polyxenophila
Rickettsia.bellii.RML369.C GCA_000012385.1_ASM1238v1
Rickettsiales.bacterium.Ac37b GCA_000746585.2_ASM74658v2
Rickettsiales.endosymbiont.of.Stachyamoeba.lipophora GCA_003932735.1_ASM393273v1
Caedimonas.varicaedens GCA_001192655.1_ASM119265v1
Candidatus.Caedibacter.acanthamoebae GCA_000743035.1_ASM74303v1
Candidatus.Nucleicultrix.amoebiphila.FS5 GCA_002117145.1_ASM211714v1
Candidatus.Finniella.inopinata GCA_004210305.1_ASM421030v1
Candidatus.Odyssella.thessalonicensis.L13 GCA_000190415.2_ASM19041v1
Candidatus.Paracaedibacter.acanthamoebae GCA_000742835.1_ASM74283v1
Candidatus.Hepatobacter.penaei GCA_000742475.1_ASM74247v1
Holospora.undulata.HU1 GCA_000388175.3_ASM38817v3
Geminicoccus.roseus.DSM.18922 GCA_000427665.1_ASM42766v1
Tistrella.mobilis.KA081020.065 GCA_000264455.2_ASM26445v2
Ahrensia.kielensis.DSM.5890 GCA_000374465.1_ASM37446v1
Chelativorans.oligotrophicus GCA_011317505.1_ASM1131750v1
Rhizobium.leguminosarum.bv.trifolii GCA_011604525.1_ASM1160452v1
Bartonella.quintana GCA_009936175.1_ASM993617v1
Pelagibacterium.halotolerans GCA_013004005.1_ASM1300400v1
Polymorphum.gilvum.SL003B.26A1 GCA_000192745.1_ASM19274v1
Roseibium.hamelinense GCA_007830545.1_ASM783054v1
Pseudovibrio.sp.FO.BEG1 GCA_000236645.1_ASM23664v1
Methylocella.silvestris.BL2 GCA_000021745.1_ASM2174v1
Methylocystis.bryophila GCA_002117405.1_ASM211740v1
Methylorubrum.extorquens.PA1 GCA_000018845.1_ASM1884v1
Rhodopseudomonas.palustris GCA_013415845.1_ASM1341584v1
Xanthobacter.autotrophicus.Py2 GCA_000017645.1_ASM1764v1
Hyphomicrobium.denitrificans.ATCC.51888 GCA_000143145.1_ASM14314v1
Rhodomicrobium.vannielii.ATCC.17100 GCA_000166055.1_ASM16605v1
Thermopetrobacter.sp.TC1 GCA_000746275.1_ASM74627v1
Candidatus.Micropelagos.thuwalensis GCA_000469155.1.RS24
alpha.proteobacterium.IMCC14465 GCA_000293845.2_ASM29384v2
Parvibaculum.lavamentivorans.DS.1 GCA_000017565.1_ASM1756v1
Asticcacaulis.excentricus.CB.48 GCA_000175215.2_ASM17521v2
Brevundimonas.subvibrioides.ATCC.15264 GCA_000144605.1_ASM14460v1
alpha.proteobacterium.L41A GCA_000335735.1.L41A.1.0
Caulobacter.vibrioides.NA1000 GCA_000022005.1_ASM2200v1
Phenylobacterium.zucineum.HLK1 GCA_000017265.1_ASM1726v1
Hirschia.baltica.ATCC.49814 GCA_000023785.1_ASM2378v1
Hyphomonas.neptunium.ATCC.15444 GCA_000013025.1_ASM1302v1
Maricaulis.maris GCA_003634045.1_ASM363404v1
Oceanicaulis.alexandrii.DSM.11625 GCA_000420265.1_ASM42026v1
Parvularcula.bermudensis.HTCC2503 GCA_000152825.2_ASM15282v2
Jannaschia.rubra GCA_900113265.1.IMG.taxon.2622736430.annotated.assembly
Octadecabacter.arcticus.238 GCA_000155735.2_ASM15573v2
Ruegeria.pomeroyi.DSS.3 GCA_000011965.2_ASM1196v2
Sagittula.stellata.E.37 GCA_000169415.1_ASM16941v1
Ketogulonicigenium.vulgare GCA_001399515.1_ASM139951v1
Rubellimicrobium.rubrum GCA_006152115.1_ASM615211v1
Paracoccus.denitrificans GCA_004063735.1_ASM406373v1
Rhodobacter.sphaeroides.ATCC.17029 GCA_000015985.1_ASM1598v1
Neomegalonema.perideroedes.DSM.15528 GCA_000374145.1_ASM37414v1
Candidatus.Pelagibacter.ubique.HIMB058 GCA_000419545.1_ASM41954v1
Candidatus.Pelagibacter.ubique.HTCC1062 GCA_000012345.1_ASM1234v1
alpha.proteobacterium.HIMB5 GCA_000299095.1_ASM29909v1
alpha.proteobacterium.HIMB114 GCA_000163555.2
Citromicrobium.sp.JL477 GCA_001304795.1_ASM130479v1
Sphingopyxis.macrogoltabida GCA_001314325.1_ASM131432v1
Sphingobium.indicum.B90A GCA_000264945.2_ASM26494v2
Sphingomonas.wittichii.RW1 GCA_000016765.1_ASM1676v1
Zymomonas.mobilis.subsp.mobilis.ATCC.31821 GCA_000007105.1_ASM710v1
alpha.proteobacterium.AAP81b GCA_001295935.1_ASM129593v1
Kordiimonas.gwangyangensis.DSM.19435 GCA_000375545.1_ASM37554v1
Sneathiella.glossodoripedis.JCM.23214 GCA_000616095.1_ASM61609v1
Candidatus.Puniceispirillum.marinum.IMCC1322 GCA_000024465.1_ASM2446v1
Thalassobaculum.salexigens.DSM.19539 GCA_000423805.1_ASM42380v1
alpha.proteobacterium.BAL199 GCA_000171835.1_ASM17183v1
Oceanibaculum.indicum GCA_003633955.1_ASM363395v1
Kiloniella.laminariae.DSM.19542 GCA_000374005.1_ASM37400v1
Rhodovibrio.salinarum.DSM.9154 GCA_000515255.1_ASM51525v1
Inquilinus.limosus.DSM.16000 GCA_000423185.1_ASM42318v1
Rhodospirillum.centenum GCA_014197805.1_ASM1419780v1
Micavibrio.aeruginosavorus.ARL.13 GCA_000226315.1_ASM22631v1
Caenispirillum.salinarum.AK4 GCA_000315795.1
Pararhodospirillum.photometricum.DSM.122 GCA_000284415.2_ASM28441v2
Rhodospirillum.rubrum.F11 GCA_000225955.1_ASM22595v1
Roseospirillum.parvum GCA_900100455.1
Magnetospirillum.magneticum.AMB.1 GCA_000009985.1_ASM998v1
GCA_001510075.1_ASM151007v1*
GCA_002938315.1_ASM293831v1*
GCA_002686255.1_ASM268625v1*
GCA_002938045.1_ASM293804v1*
GCA_002389175.1_ASM238917v1*
GCA_002389715.1_ASM238971v1*
GCA_002722175.1_ASM272217v1*
GCA_002327565.1_ASM232756v1*
GCA_002704405.1_ASM270440v1*
GCA_002725625.1_ASM272562v1*
GCA_002687515.1_ASM268751v1*
GCA_002691065.1_ASM269106v1*
GCA_002723345.1_ASM272334v1*
GCA_002937995.1_ASM293799v1*
GCA_002938035.1_ASM293803v1*
Terasakiella.pusilla.DSM.6293 GCA_000688235.1_ASM68823v1
Thalassospira.profundimaris.WP0211 GCA_000300275.1_ASM30027v1
Elioraea.tepidiphila.DSM.17972 GCA_000378465.1_ASM37846v1
Belnapia.moabensis.DSM.16746 GCA_000745835.1_ASM74583v1
Rubritepida.flocculans.DSM.14296 GCA_000425365.1_ASM42536v1
Roseomonas.gilardii GCA_001941945.1_ASM194194v1
Acidiphilium.multivorum.AIU301 GCA_000202835.1_ASM20283v1
Acidisphaera.rubrifaciens.HS.AP3 GCA_000964365.1_ASM96436v1
Granulibacter.bethesdensis.CGDNIH4 GCA_000576145.2_ASM57614v2
Commensalibacter.intestini.A911 GCA_000231445.2_ASM23144v1
Gluconacetobacter.diazotrophicus.PA1.5 GCA_000067045.1_ASM6704v1
Acetobacter.aceti.NBRC.14818 GCA_000193495.2_ASM19349v2
Gluconobacter.oxydans.DSM.3504 GCA_000583855.1_ASM58385v1
0.432307

### Group E

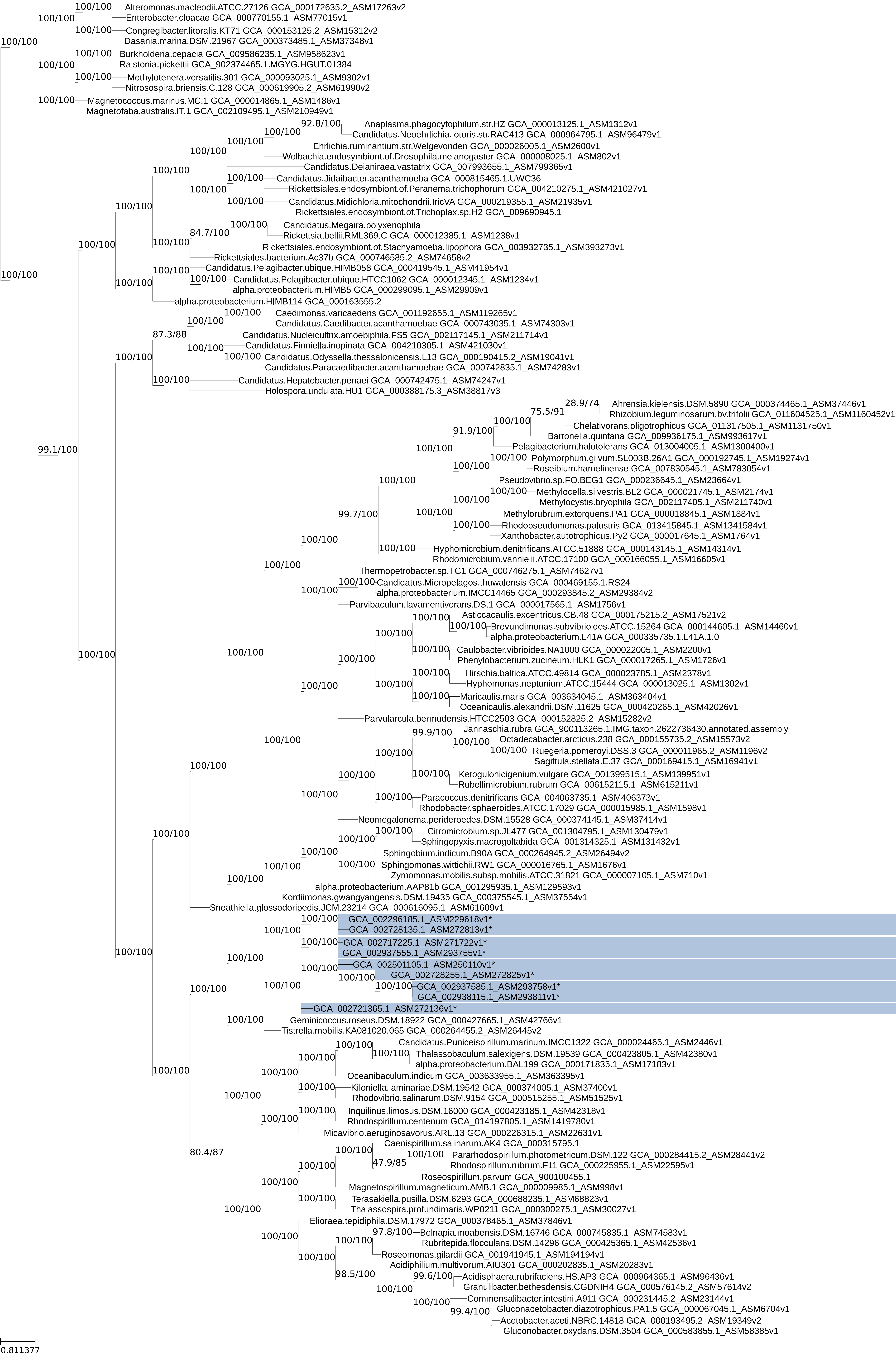
Alteromonas.macleodii.ATCC.27126 GCA_000172635.2_ASM17263v2
Enterobacter.cloacae GCA_000770155.1_ASM77015v1
Congregibacter.litoralis.KT71 GCA_000153125.2_ASM15312v2
Dasania.marina.DSM.21967 GCA_000373485.1_ASM37348v1
Burkholderia.cepacia GCA_009586235.1_ASM958623v1
Ralstonia.pickettii GCA_902374465.1.MGYG.HGUT.01384
Methylotenera.versatilis.301 GCA_000093025.1_ASM9302v1
Nitrosospira.briensis.C.128 GCA_000619905.2_ASM61990v2
Magnetococcus.marinus.MC.1 GCA_000014865.1_ASM1486v1
Magnetofaba.australis.IT.1 GCA_002109495.1_ASM210949v1
Anaplasma.phagocytophilum.str.HZ GCA_000013125.1_ASM1312v1
Candidatus.Neoehrlichia.lotoris.str.RAC413 GCA_000964795.1_ASM96479v1
Ehrlichia.ruminantium.str.Welgevonden GCA_000026005.1_ASM2600v1
Wolbachia.endosymbiont.of.Drosophila.melanogaster GCA_000008025.1_ASM802v1
Candidatus.Deianiraea.vastatrix GCA_007993655.1_ASM799365v1
Candidatus.Jidaibacter.acanthamoeba GCA_000815465.1.UWC36
Rickettsiales.endosymbiont.of.Peranema.trichophorum GCA_004210275.1_ASM421027v1
Candidatus.Midichloria.mitochondrii.IricVA GCA_000219355.1_ASM21935v1
Rickettsiales.endosymbiont.of.Trichoplax.sp.H2 GCA_009690945.1
Candidatus.Megaira.polyxenophila
Rickettsia.bellii.RML369.C GCA_000012385.1_ASM1238v1
Rickettsiales.endosymbiont.of.Stachyamoeba.lipophora GCA_003932735.1_ASM393273v1
Rickettsiales.bacterium.Ac37b GCA_000746585.2_ASM74658v2
Candidatus.Pelagibacter.ubique.HIMB058 GCA_000419545.1_ASM41954v1
Candidatus.Pelagibacter.ubique.HTCC1062 GCA_000012345.1_ASM1234v1
alpha.proteobacterium.HIMB5 GCA_000299095.1_ASM29909v1
alpha.proteobacterium.HIMB114 GCA_000163555.2
Caedimonas.varicaedens GCA_001192655.1_ASM119265v1
Candidatus.Caedibacter.acanthamoebae GCA_000743035.1_ASM74303v1
Candidatus.Nucleicultrix.amoebiphila.FS5 GCA_002117145.1_ASM211714v1
Candidatus.Finniella.inopinata GCA_004210305.1_ASM421030v1
Candidatus.Odyssella.thessalonicensis.L13 GCA_000190415.2_ASM19041v1
Candidatus.Paracaedibacter.acanthamoebae GCA_000742835.1_ASM74283v1
Candidatus.Hepatobacter.penaei GCA_000742475.1_ASM74247v1
Holospora.undulata.HU1 GCA_000388175.3_ASM38817v3
Ahrensia.kielensis.DSM.5890 GCA_000374465.1_ASM37446v1
Rhizobium.leguminosarum.bv.trifolii GCA_011604525.1_ASM1160452v1
Chelativorans.oligotrophicus GCA_011317505.1_ASM1131750v1
Bartonella.quintana GCA_009936175.1_ASM993617v1
Pelagibacterium.halotolerans GCA_013004005.1_ASM1300400v1
Polymorphum.gilvum.SL003B.26A1 GCA_000192745.1_ASM19274v1
Roseibium.hamelinense GCA_007830545.1_ASM783054v1
Pseudovibrio.sp.FO.BEG1 GCA_000236645.1_ASM23664v1
Methylocella.silvestris.BL2 GCA_000021745.1_ASM2174v1
Methylocystis.bryophila GCA_002117405.1_ASM211740v1
Methylorubrum.extorquens.PA1 GCA_000018845.1_ASM1884v1
Rhodopseudomonas.palustris GCA_013415845.1_ASM1341584v1
Xanthobacter.autotrophicus.Py2 GCA_000017645.1_ASM1764v1
Hyphomicrobium.denitrificans.ATCC.51888 GCA_000143145.1_ASM14314v1
Rhodomicrobium.vannielii.ATCC.17100 GCA_000166055.1_ASM16605v1
Thermopetrobacter.sp.TC1 GCA_000746275.1_ASM74627v1
Candidatus.Micropelagos.thuwalensis GCA_000469155.1.RS24
alpha.proteobacterium.IMCC14465 GCA_000293845.2_ASM29384v2
Parvibaculum.lavamentivorans.DS.1 GCA_000017565.1_ASM1756v1
Asticcacaulis.excentricus.CB.48 GCA_000175215.2_ASM17521v2
Brevundimonas.subvibrioides.ATCC.15264 GCA_000144605.1_ASM14460v1
alpha.proteobacterium.L41A GCA_000335735.1.L41A.1.0
Caulobacter.vibrioides.NA1000 GCA_000022005.1_ASM2200v1
Phenylobacterium.zucineum.HLK1 GCA_000017265.1_ASM1726v1
Hirschia.baltica.ATCC.49814 GCA_000023785.1_ASM2378v1
Hyphomonas.neptunium.ATCC.15444 GCA_000013025.1_ASM1302v1
Maricaulis.maris GCA_003634045.1_ASM363404v1
Oceanicaulis.alexandrii.DSM.11625 GCA_000420265.1_ASM42026v1
Parvularcula.bermudensis.HTCC2503 GCA_000152825.2_ASM15282v2
Jannaschia.rubra GCA_900113265.1.IMG.taxon.2622736430.annotated.assembly
Octadecabacter.arcticus.238 GCA_000155735.2_ASM15573v2
Ruegeria.pomeroyi.DSS.3 GCA_000011965.2_ASM1196v2
Sagittula.stellata.E.37 GCA_000169415.1_ASM16941v1
Ketogulonicigenium.vulgare GCA_001399515.1_ASM139951v1
Rubellimicrobium.rubrum GCA_006152115.1_ASM615211v1
Paracoccus.denitrificans GCA_004063735.1_ASM406373v1
Rhodobacter.sphaeroides.ATCC.17029 GCA_000015985.1_ASM1598v1
Neomegalonema.perideroedes.DSM.15528 GCA_000374145.1_ASM37414v1
Citromicrobium.sp.JL477 GCA_001304795.1_ASM130479v1
Sphingopyxis.macrogoltabida GCA_001314325.1_ASM131432v1
Sphingobium.indicum.B90A GCA_000264945.2_ASM26494v2
Sphingomonas.wittichii.RW1 GCA_000016765.1_ASM1676v1
Zymomonas.mobilis.subsp.mobilis.ATCC.31821 GCA_000007105.1_ASM710v1
alpha.proteobacterium.AAP81b GCA_001295935.1_ASM129593v1
Kordiimonas.gwangyangensis.DSM.19435 GCA_000375545.1_ASM37554v1
Sneathiella.glossodoripedis.JCM.23214 GCA_000616095.1_ASM61609v1
GCA_002296185.1_ASM229618v1*
GCA_002728135.1_ASM272813v1*
GCA_002717225.1_ASM271722v1*
GCA_002937555.1_ASM293755v1*
GCA_002501105.1_ASM250110v1*
GCA_002728255.1_ASM272825v1*
GCA_002937585.1_ASM293758v1*
GCA_002938115.1_ASM293811v1*
GCA_002721365.1_ASM272136v1*
Geminicoccus.roseus.DSM.18922 GCA_000427665.1_ASM42766v1
Tistrella.mobilis.KA081020.065 GCA_000264455.2_ASM26445v2
Candidatus.Puniceispirillum.marinum.IMCC1322 GCA_000024465.1_ASM2446v1
Thalassobaculum.salexigens.DSM.19539 GCA_000423805.1_ASM42380v1
alpha.proteobacterium.BAL199 GCA_000171835.1_ASM17183v1
Oceanibaculum.indicum GCA_003633955.1_ASM363395v1
Kiloniella.laminariae.DSM.19542 GCA_000374005.1_ASM37400v1
Rhodovibrio.salinarum.DSM.9154 GCA_000515255.1_ASM51525v1
Inquilinus.limosus.DSM.16000 GCA_000423185.1_ASM42318v1
Rhodospirillum.centenum GCA_014197805.1_ASM1419780v1
Micavibrio.aeruginosavorus.ARL.13 GCA_000226315.1_ASM22631v1
Caenispirillum.salinarum.AK4 GCA_000315795.1
Pararhodospirillum.photometricum.DSM.122 GCA_000284415.2_ASM28441v2
Rhodospirillum.rubrum.F11 GCA_000225955.1_ASM22595v1
Roseospirillum.parvum GCA_900100455.1
Magnetospirillum.magneticum.AMB.1 GCA_000009985.1_ASM998v1
Terasakiella.pusilla.DSM.6293 GCA_000688235.1_ASM68823v1
Thalassospira.profundimaris.WP0211 GCA_000300275.1_ASM30027v1
Elioraea.tepidiphila.DSM.17972 GCA_000378465.1_ASM37846v1
Belnapia.moabensis.DSM.16746 GCA_000745835.1_ASM74583v1
Rubritepida.flocculans.DSM.14296 GCA_000425365.1_ASM42536v1
Roseomonas.gilardii GCA_001941945.1_ASM194194v1
Acidiphilium.multivorum.AIU301 GCA_000202835.1_ASM20283v1
Acidisphaera.rubrifaciens.HS.AP3 GCA_000964365.1_ASM96436v1
Granulibacter.bethesdensis.CGDNIH4 GCA_000576145.2_ASM57614v2
Commensalibacter.intestini.A911 GCA_000231445.2_ASM23144v1
Gluconacetobacter.diazotrophicus.PA1.5 GCA_000067045.1_ASM6704v1
Acetobacter.aceti.NBRC.14818 GCA_000193495.2_ASM19349v2
Gluconobacter.oxydans.DSM.3504 GCA_000583855.1_ASM58385v1
0.811377

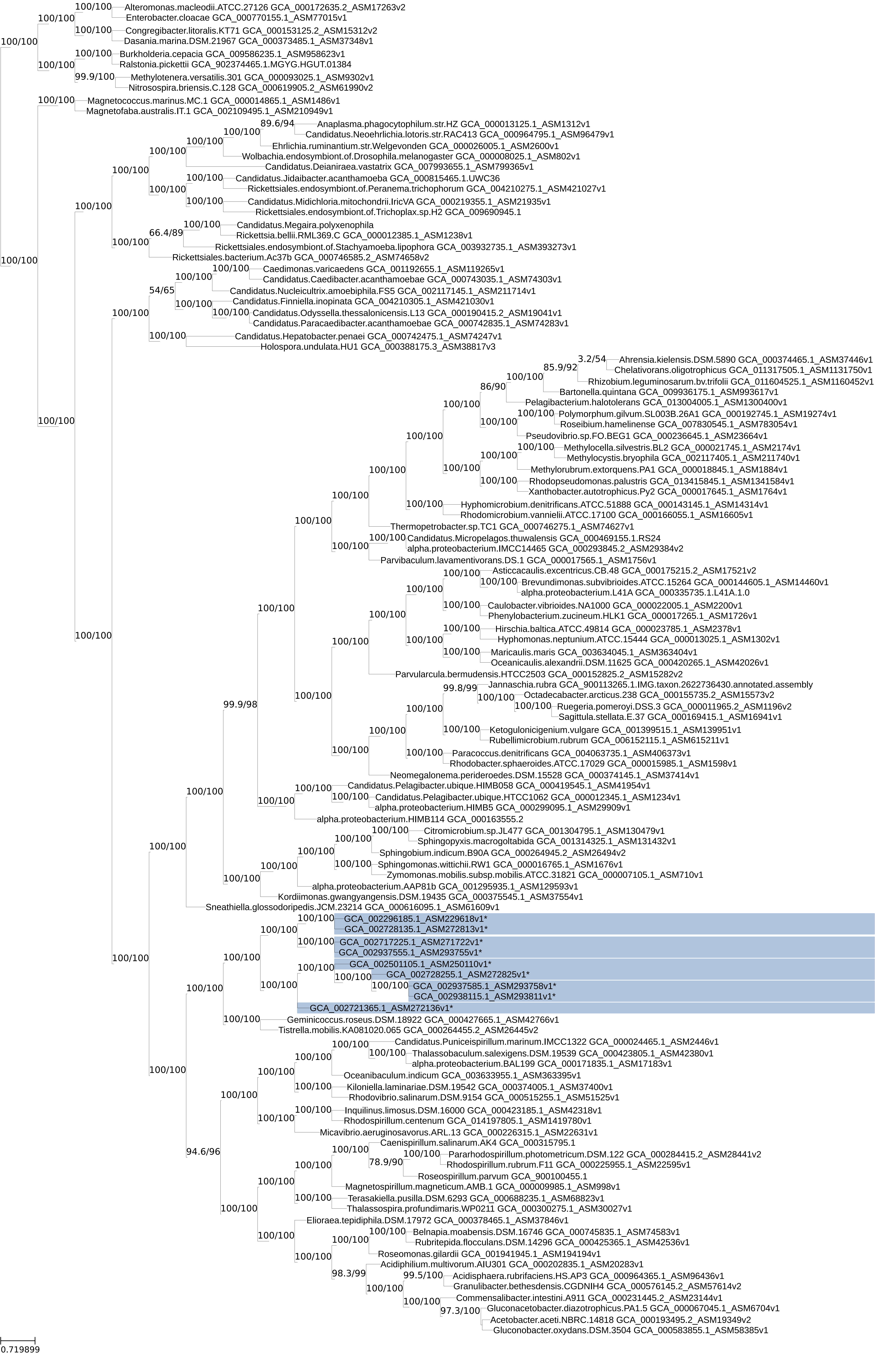
100/100
Alteromonas.macleodii.ATCC.27126 GCA_000172635.2_ASM17263v2
100/100
Enterobacter.cloacae GCA_000770155.1_ASM77015v1
100/100
100/100
Congregibacter.litoralis.KT71 GCA_000153125.2_ASM15312v2
Dasania.marina.DSM.21967 GCA_000373485.1_ASM37348v1
100/100
Burkholderia.cepacia GCA_009586235.1_ASM958623v1
100/100
Ralstonia.pickettii GCA_902374465.1.MGYG.HGUT.01384
99.9/100
Methylotenera.versatilis.301 GCA_000093025.1_ASM9302v1
Nitrosospira.briensis.C.128 GCA_000619905.2_ASM61990v2
100/100
100/100
Magnetococcus.marinus.MC.1 GCA_000014865.1_ASM1486v1
Magnetofaba.australis.IT.1 GCA_002109495.1_ASM210949v1
89.6/94
Anaplasma.phagocytophilum.str.HZ GCA_000013125.1_ASM1312v1
100/100
Candidatus.Neoehrlichia.lotoris.str.RAC413 GCA_000964795.1_ASM96479v1
100/100
Ehrlichia.ruminantium.str.Welgevonden GCA_000026005.1_ASM2600v1
100/100
Wolbachia.endosymbiont.of.Drosophila.melanogaster GCA_000008025.1_ASM802v1
100/100
Candidatus.Deianiraea.vastatrix GCA_007993655.1_ASM799365v1
100/100
Candidatus.Jidaibacter.acanthamoeba GCA_000815465.1.UWC36
100/100
Rickettsiales.endosymbiont.of.Peranema.trichophorum GCA_004210275.1_ASM421027v1
100/100
100/100
Candidatus.Midichloria.mitochondrii.IricVA GCA_000219355.1_ASM21935v1
Rickettsiales.endosymbiont.of.Trichoplax.sp.H2 GCA_009690945.1
100/100
Candidatus.Megaira.polyxenophila
66.4/89
Rickettsia.bellii.RML369.C GCA_000012385.1_ASM1238v1
100/100
Rickettsiales.endosymbiont.of.Stachyamoeba.lipophora GCA_003932735.1_ASM393273v1
100/100
Rickettsiales.bacterium.Ac37b GCA_000746585.2_ASM74658v2
100/100
Caedimonas.varicaedens GCA_001192655.1_ASM119265v1
100/100
Candidatus.Caedibacter.acanthamoebae GCA_000743035.1_ASM74303v1
54/65
Candidatus.Nucleicultrix.amoebiphila.FS5 GCA_002117145.1_ASM211714v1
100/100
Candidatus.Finniella.inopinata GCA_004210305.1_ASM421030v1
100/100
100/100
Candidatus.Odyssella.thessalonicensis.L13 GCA_000190415.2_ASM19041v1
Candidatus.Paracaedibacter.acanthamoebae GCA_000742835.1_ASM74283v1
100/100
Candidatus.Hepatobacter.penaei GCA_000742475.1_ASM74247v1
Holospora.undulata.HU1 GCA_000388175.3_ASM38817v3
3.2/54
Ahrensia.kielensis.DSM.5890 GCA_000374465.1_ASM37446v1
85.9/92
Chelativorans.oligotrophicus GCA_011317505.1_ASM1131750v1
100/100
Rhizobium.leguminosarum.bv.trifolii GCA_011604525.1_ASM1160452v1
86/90
Bartonella.quintana GCA_009936175.1_ASM993617v1
100/100
Pelagibacterium.halotolerans GCA_013004005.1_ASM1300400v1
100/100
Polymorphum.gilvum.SL003B.26A1 GCA_000192745.1_ASM19274v1
100/100
Roseibium.hamelinense GCA_007830545.1_ASM783054v1
100/100
Pseudovibrio.sp.FO.BEG1 GCA_000236645.1_ASM23664v1
100/100
Methylocella.silvestris.BL2 GCA_000021745.1_ASM2174v1
100/100
Methylocystis.bryophila GCA_002117405.1_ASM211740v1
100/100
Methylorubrum.extorquens.PA1 GCA_000018845.1_ASM1884v1
100/100
100/100
Rhodopseudomonas.palustris GCA_013415845.1_ASM1341584v1
Xanthobacter.autotrophicus.Py2 GCA_000017645.1_ASM1764v1
100/100
100/100
Hyphomicrobium.denitrificans.ATCC.51888 GCA_000143145.1_ASM14314v1
Rhodomicrobium.vannielii.ATCC.17100 GCA_000166055.1_ASM16605v1
100/100
Thermopetrobacter.sp.TC1 GCA_000746275.1_ASM74627v1
100/100
Candidatus.Micropelagos.thuwalensis GCA_000469155.1.RS24
100/100
alpha.proteobacterium.IMCC14465 GCA_000293845.2_ASM29384v2
Parvibaculum.lavamentivorans.DS.1 GCA_000017565.1_ASM1756v1
100/100
Asticcacaulis.excentricus.CB.48 GCA_000175215.2_ASM17521v2
100/100
Brevundimonas.subvibrioides.ATCC.15264 GCA_000144605.1_ASM14460v1
100/100
alpha.proteobacterium.L41A GCA_000335735.1.L41A.1.0
100/100
Caulobacter.vibrioides.NA1000 GCA_000022005.1_ASM2200v1
100/100
Phenylobacterium.zucineum.HLK1 GCA_000017265.1_ASM1726v1
100/100
Hirschia.baltica.ATCC.49814 GCA_000023785.1_ASM2378v1
100/100
Hyphomonas.neptunium.ATCC.15444 GCA_000013025.1_ASM1302v1
100/100
100/100
Maricaulis.maris GCA_003634045.1_ASM363404v1
Oceanicaulis.alexandrii.DSM.11625 GCA_000420265.1_ASM42026v1
100/100
Parvularcula.bermudensis.HTCC2503 GCA_000152825.2_ASM15282v2
99.8/99
Jannaschia.rubra GCA_900113265.1.IMG.taxon.2622736430.annotated.assembly
100/100
Octadecabacter.arcticus.238 GCA_000155735.2_ASM15573v2
100/100
Ruegeria.pomeroyi.DSS.3 GCA_000011965.2_ASM1196v2
100/100
Sagittula.stellata.E.37 GCA_000169415.1_ASM16941v1
99.9/98
100/100
Ketogulonicigenium.vulgare GCA_001399515.1_ASM139951v1
100/100
Rubellimicrobium.rubrum GCA_006152115.1_ASM615211v1
100/100
100/100
Paracoccus.denitrificans GCA_004063735.1_ASM406373v1
Rhodobacter.sphaeroides.ATCC.17029 GCA_000015985.1_ASM1598v1
Neomegalonema.perideroedes.DSM.15528 GCA_000374145.1_ASM37414v1
100/100
Candidatus.Pelagibacter.ubique.HIMB058 GCA_000419545.1_ASM41954v1
100/100
100/100
Candidatus.Pelagibacter.ubique.HTCC1062 GCA_000012345.1_ASM1234v1
100/100
alpha.proteobacterium.HIMB5 GCA_000299095.1_ASM29909v1
alpha.proteobacterium.HIMB114 GCA_000163555.2
100/100
Citromicrobium.sp.JL477 GCA_001304795.1_ASM130479v1
100/100
Sphingopyxis.macrogoltabida GCA_001314325.1_ASM131432v1
100/100
100/100
Sphingobium.indicum.B90A GCA_000264945.2_ASM26494v2
100/100
Sphingomonas.wittichii.RW1 GCA_000016765.1_ASM1676v1
100/100
Zymomonas.mobilis.subsp.mobilis.ATCC.31821 GCA_000007105.1_ASM710v1
100/100
alpha.proteobacterium.AAP81b GCA_001295935.1_ASM129593v1
Kordiimonas.gwangyangensis.DSM.19435 GCA_000375545.1_ASM37554v1
Sneathiella.glossodoripedis.JCM.23214 GCA_000616095.1_ASM61609v1
100/100
GCA_002296185.1_ASM229618v1*
100/100
GCA_002728135.1_ASM272813v1*
100/100
GCA_002717225.1_ASM271722v1*
100/100
GCA_002937555.1_ASM293755v1*
100/100
GCA_002501105.1_ASM250110v1*
100/100
100/100
GCA_002728255.1_ASM272825v1*
100/100
100/100
GCA_002937585.1_ASM293758v1*
GCA_002938115.1_ASM293811v1*
100/100
GCA_002721365.1_ASM272136v1*
100/100
Geminicoccus.roseus.DSM.18922 GCA_000427665.1_ASM42766v1
Tistrella.mobilis.KA081020.065 GCA_000264455.2_ASM26445v2
100/100
Candidatus.Puniceispirillum.marinum.IMCC1322 GCA_000024465.1_ASM2446v1
100/100
Thalassobaculum.salexigens.DSM.19539 GCA_000423805.1_ASM42380v1
100/100
alpha.proteobacterium.BAL199 GCA_000171835.1_ASM17183v1
100/100
Oceanibaculum.indicum GCA_003633955.1_ASM363395v1
100/100
Kiloniella.laminariae.DSM.19542 GCA_000374005.1_ASM37400v1
100/100
Rhodovibrio.salinarum.DSM.9154 GCA_000515255.1_ASM51525v1
100/100
Inquilinus.limosus.DSM.16000 GCA_000423185.1_ASM42318v1
100/100
Rhodospirillum.centenum GCA_014197805.1_ASM1419780v1
Micavibrio.aeruginosavorus.ARL.13 GCA_000226315.1_ASM22631v1
100/100
Caenispirillum.salinarum.AK4 GCA_000315795.1
100/100
Pararhodospirillum.photometricum.DSM.122 GCA_000284415.2_ASM28441v2
94.6/96
78.9/90
Rhodospirillum.rubrum.F11 GCA_000225955.1_ASM22595v1
100/100
Roseospirillum.parvum GCA_900100455.1
100/100
Magnetospirillum.magneticum.AMB.1 GCA_000009985.1_ASM998v1
100/100
Terasakiella.pusilla.DSM.6293 GCA_000688235.1_ASM68823v1
100/100
Thalassospira.profundimaris.WP0211 GCA_000300275.1_ASM30027v1
Elioraea.tepidiphila.DSM.17972 GCA_000378465.1_ASM37846v1
100/100
Belnapia.moabensis.DSM.16746 GCA_000745835.1_ASM74583v1
100/100
Rubritepida.flocculans.DSM.14296 GCA_000425365.1_ASM42536v1
100/100
100/100
Roseomonas.gilardii GCA_001941945.1_ASM194194v1
Acidiphilium.multivorum.AIU301 GCA_000202835.1_ASM20283v1
99.5/100
Acidisphaera.rubrifaciens.HS.AP3 GCA_000964365.1_ASM96436v1
98.3/99
100/100
Granulibacter.bethesdensis.CGDNIH4 GCA_000576145.2_ASM57614v2
Commensalibacter.intestini.A911 GCA_000231445.2_ASM23144v1
100/100
Gluconacetobacter.diazotrophicus.PA1.5 GCA_000067045.1_ASM6704v1
97.3/100
Acetobacter.aceti.NBRC.14818 GCA_000193495.2_ASM19349v2
Gluconobacter.oxydans.DSM.3504 GCA_000583855.1_ASM58385v1
0.719899

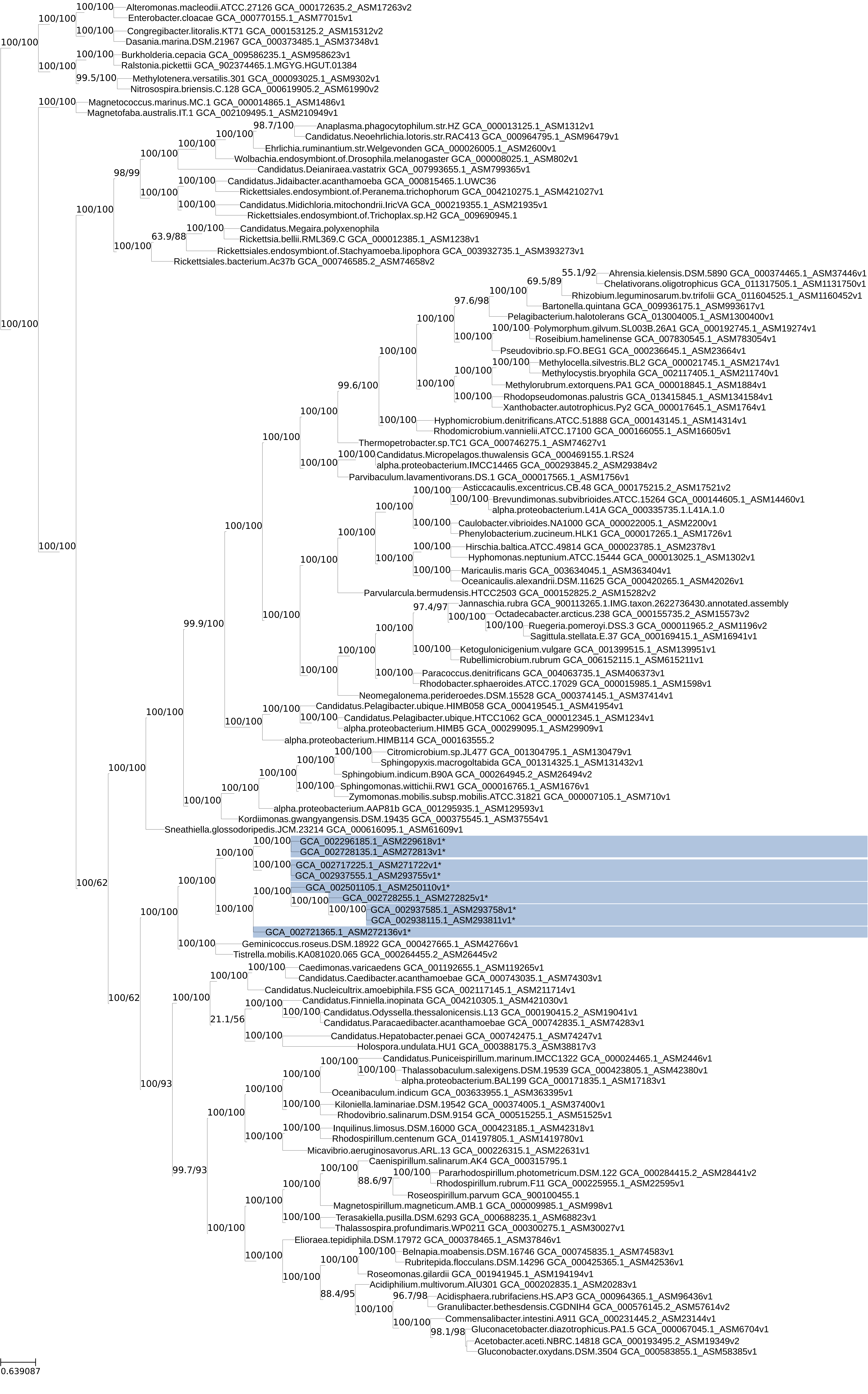

Alteromonas.macleodii.ATCC.27126 GCA_000172635.2_ASM17263v2
Enterobacter.cloacae GCA_000770155.1_ASM77015v1
Congregibacter.litoralis.KT71 GCA_000153125.2_ASM15312v2
Dasania.marina.DSM.21967 GCA_000373485.1_ASM37348v1
Burkholderia.cepacia GCA_009586235.1_ASM958623v1
Ralstonia.pickettii GCA_902374465.1.MGYG.HGUT.01384
Methylotenera.versatilis.301 GCA_000093025.1_ASM9302v1
Nitrosospira.briensis.C.128 GCA_000619905.2_ASM61990v2
Magnetococcus.marinus.MC.1 GCA_000014865.1_ASM1486v1
Magnetofaba.australis.IT.1 GCA_002109495.1_ASM210949v1
Anaplasma.phagocytophilum.str.HZ GCA_000013125.1_ASM1312v1
Candidatus.Neoehrlichia.lotoris.str.RAC413 GCA_000964795.1_ASM96479v1
Ehrlichia.ruminantium.str.Welgevonden GCA_000026005.1_ASM2600v1
Wolbachia.endosymbiont.of.Drosophila.melanogaster GCA_000008025.1_ASM802v1
Candidatus.Deianiraea.vastatrix GCA_007993655.1_ASM799365v1
Candidatus.Jidaibacter.acanthamoeba GCA_000815465.1.UWC36
Rickettsiales.endosymbiont.of.Peranema.trichophorum GCA_004210275.1_ASM421027v1
Candidatus.Midichloria.mitochondrii.IricVA GCA_000219355.1_ASM21935v1
Rickettsiales.endosymbiont.of.Trichoplax.sp.H2 GCA_009690945.1
Candidatus.Megaira.polyxenophila
Rickettsia.bellii.RML369.C GCA_000012385.1_ASM1238v1
Rickettsiales.endosymbiont.of.Stachyamoeba.lipophora GCA_003932735.1_ASM393273v1
Rickettsiales.bacterium.Ac37b GCA_000746585.2_ASM74658v2
Ahrensia.kielensis.DSM.5890 GCA_000374465.1_ASM37446v1
Chelativorans.oligotrophicus GCA_011317505.1_ASM1131750v1
Rhizobium.leguminosarum.bv.trifolii GCA_011604525.1_ASM1160452v1
Bartonella.quintana GCA_009936175.1_ASM993617v1
Pelagibacterium.halotolerans GCA_013004005.1_ASM1300400v1
Polymorphum.gilvum.SL003B.26A1 GCA_000192745.1_ASM19274v1
Roseibium.hamelinense GCA_007830545.1_ASM783054v1
Pseudovibrio.sp.FO.BEG1 GCA_000236645.1_ASM23664v1
Methylocella.silvestris.BL2 GCA_000021745.1_ASM2174v1
Methylocystis.bryophila GCA_002117405.1_ASM211740v1
Methylorubrum.extorquens.PA1 GCA_000018845.1_ASM1884v1
Rhodopseudomonas.palustris GCA_013415845.1_ASM1341584v1
Xanthobacter.autotrophicus.Py2 GCA_000017645.1_ASM1764v1
Hyphomicrobium.denitrificans.ATCC.51888 GCA_000143145.1_ASM14314v1
Rhodomicrobium.vannielii.ATCC.17100 GCA_000166055.1_ASM16605v1
Thermopetrobacter.sp.TC1 GCA_000746275.1_ASM74627v1
Candidatus.Micropelagos.thuwalensis GCA_000469155.1.RS24
alpha.proteobacterium.IMCC14465 GCA_000293845.2_ASM29384v2
Parvibaculum.lavamentivorans.DS.1 GCA_000017565.1_ASM1756v1
Asticcacaulis.excentricus.CB.48 GCA_000175215.2_ASM17521v2
Brevundimonas.subvibrioides.ATCC.15264 GCA_000144605.1_ASM14460v1
alpha.proteobacterium.L41A GCA_000335735.1.L41A.1.0
Caulobacter.vibrioides.NA1000 GCA_000022005.1_ASM2200v1
Phenylobacterium.zucineum.HLK1 GCA_000017265.1_ASM1726v1
Hirschia.baltica.ATCC.49814 GCA_000023785.1_ASM2378v1
Hyphomonas.neptunium.ATCC.15444 GCA_000013025.1_ASM1302v1
Maricaulis.maris GCA_003634045.1_ASM363404v1
Oceanicaulis.alexandrii.DSM.11625 GCA_000420265.1_ASM42026v1
Parvularcula.bermudensis.HTCC2503 GCA_000152825.2_ASM15282v2
Jannaschia.rubra GCA_900113265.1.IMG.taxon.2622736430.annotated.assembly
Octadecabacter.arcticus.238 GCA_000155735.2_ASM15573v2
Ruegeria.pomeroyi.DSS.3 GCA_000011965.2_ASM1196v2
Sagittula.stellata.E.37 GCA_000169415.1_ASM16941v1
Ketogulonicigenium.vulgare GCA_001399515.1_ASM139951v1
Rubellimicrobium.rubrum GCA_006152115.1_ASM615211v1
Paracoccus.denitrificans GCA_004063735.1_ASM406373v1
Rhodobacter.sphaeroides.ATCC.17029 GCA_000015985.1_ASM1598v1
Neomegalonema.perideroedes.DSM.15528 GCA_000374145.1_ASM37414v1
Candidatus.Pelagibacter.ubique.HIMB058 GCA_000419545.1_ASM41954v1
Candidatus.Pelagibacter.ubique.HTCC1062 GCA_000012345.1_ASM1234v1
alpha.proteobacterium.HIMB5 GCA_000299095.1_ASM29909v1
alpha.proteobacterium.HIMB114 GCA_000163555.2
Citromicrobium.sp.JL477 GCA_001304795.1_ASM130479v1
Sphingopyxis.macrogoltabida GCA_001314325.1_ASM131432v1
Sphingobium.indicum.B90A GCA_000264945.2_ASM26494v2
Sphingomonas.wittichii.RW1 GCA_000016765.1_ASM1676v1
Zymomonas.mobilis.subsp.mobilis.ATCC.31821 GCA_000007105.1_ASM710v1
alpha.proteobacterium.AAP81b GCA_001295935.1_ASM129593v1
Kordiimonas.gwangyangensis.DSM.19435 GCA_000375545.1_ASM37554v1
Sneathiella.glossodoripedis.JCM.23214 GCA_000616095.1_ASM61609v1
GCA_002296185.1_ASM229618v1*
GCA_002728135.1_ASM272813v1*
GCA_002717225.1_ASM271722v1*
GCA_002937555.1_ASM293755v1*
GCA_002501105.1_ASM250110v1*
GCA_002728255.1_ASM272825v1*
GCA_002937585.1_ASM293758v1*
GCA_002938115.1_ASM293811v1*
GCA_002721365.1_ASM272136v1*
Geminicoccus.roseus.DSM.18922 GCA_000427665.1_ASM42766v1
Tistrella.mobilis.KA081020.065 GCA_000264455.2_ASM26445v2
Caedimonas.varicaedens GCA_001192655.1_ASM119265v1
Candidatus.Caedibacter.acanthamoebae GCA_000743035.1_ASM74303v1
Candidatus.Nucleicultrix.amoebiphila.FS5 GCA_002117145.1_ASM211714v1
Candidatus.Finniella.inopinata GCA_004210305.1_ASM421030v1
Candidatus.Odyssella.thessalonicensis.L13 GCA_000190415.2_ASM19041v1
Candidatus.Paracaedibacter.acanthamoebae GCA_000742835.1_ASM74283v1
Candidatus.Hepatobacter.penaei GCA_000742475.1_ASM74247v1
Holospora.undulata.HU1 GCA_000388175.3_ASM38817v3
Candidatus.Puniceispirillum.marinum.IMCC1322 GCA_000024465.1_ASM2446v1
Thalassobaculum.salexigens.DSM.19539 GCA_000423805.1_ASM42380v1
alpha.proteobacterium.BAL199 GCA_000171835.1_ASM17183v1
Oceanibaculum.indicum GCA_003633955.1_ASM363395v1
Kiloniella.laminariae.DSM.19542 GCA_000374005.1_ASM37400v1
Rhodovibrio.salinarum.DSM.9154 GCA_000515255.1_ASM51525v1
Inquilinus.limosus.DSM.16000 GCA_000423185.1_ASM42318v1
Rhodospirillum.centenum GCA_014197805.1_ASM1419780v1
Micavibrio.aeruginosavorus.ARL.13 GCA_000226315.1_ASM22631v1
Caenispirillum.salinarum.AK4 GCA_000315795.1
Pararhodospirillum.photometricum.DSM.122 GCA_000284415.2_ASM28441v2
Rhodospirillum.rubrum.F11 GCA_000225955.1_ASM22595v1
Roseospirillum.parvum GCA_900100455.1
Magnetospirillum.magneticum.AMB.1 GCA_000009985.1_ASM998v1
Terasakiella.pusilla.DSM.6293 GCA_000688235.1_ASM68823v1
Thalassospira.profundimaris.WP0211 GCA_000300275.1_ASM30027v1
Elioraea.tepidiphila.DSM.17972 GCA_000378465.1_ASM37846v1
Belnapia.moabensis.DSM.16746 GCA_000745835.1_ASM74583v1
Rubritepida.flocculans.DSM.14296 GCA_000425365.1_ASM42536v1
Roseomonas.gilardii GCA_001941945.1_ASM194194v1
Acidiphilium.multivorum.AIU301 GCA_000202835.1_ASM20283v1
Acidisphaera.rubrifaciens.HS.AP3 GCA_000964365.1_ASM96436v1
Granulibacter.bethesdensis.CGDNIH4 GCA_000576145.2_ASM57614v2
Commensalibacter.intestini.A911 GCA_000231445.2_ASM23144v1
Gluconacetobacter.diazotrophicus.PA1.5 GCA_000067045.1_ASM6704v1
Acetobacter.aceti.NBRC.14818 GCA_000193495.2_ASM19349v2
Gluconobacter.oxydans.DSM.3504 GCA_000583855.1_ASM58385v1
0.639087

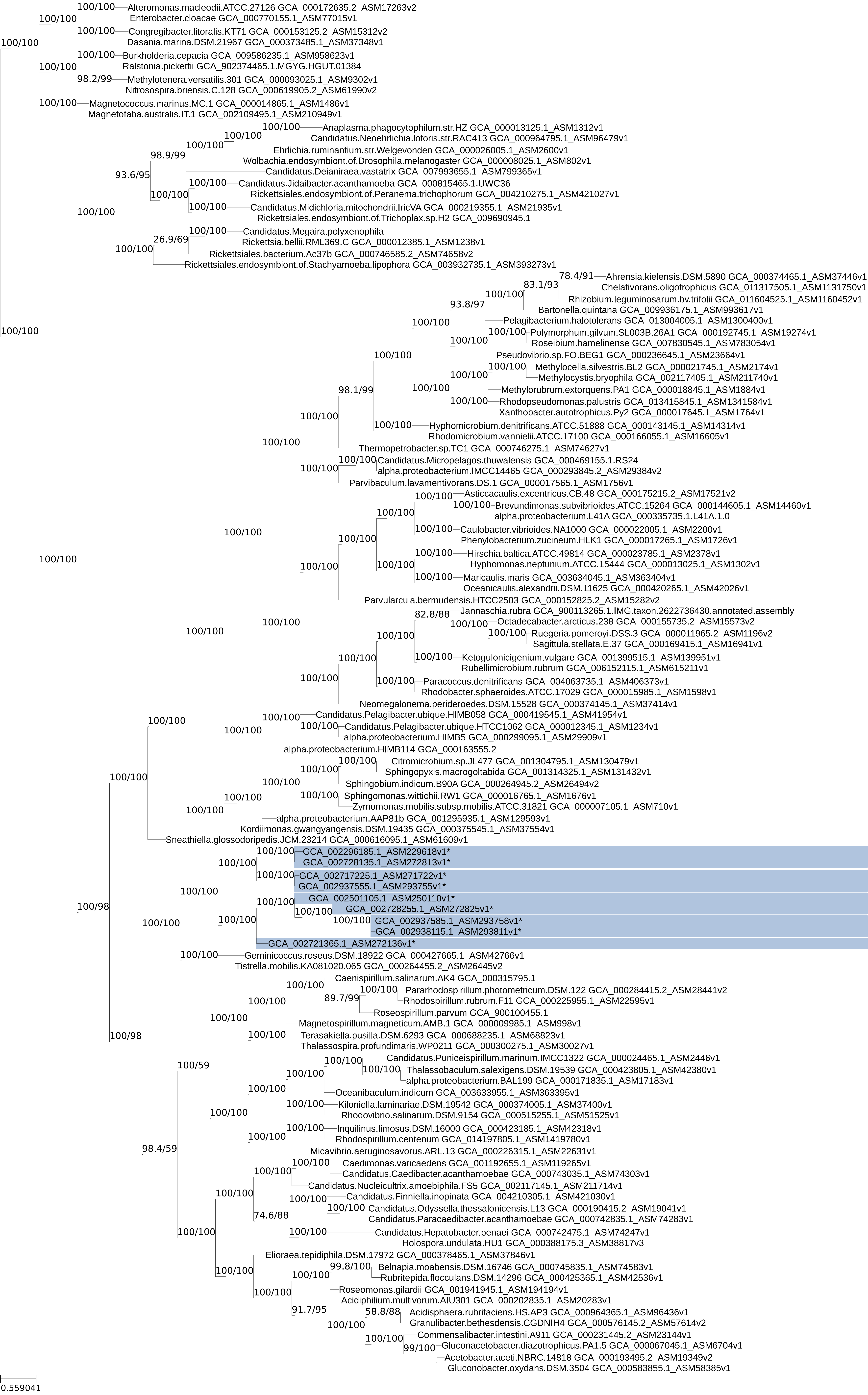

100/100
Alteromonas.macleodii.ATCC.27126 GCA_000172635.2_ASM17263v2
100/100
Enterobacter.cloacae GCA_000770155.1_ASM77015v1
100/100
100/100
Congregibacter.litoralis.KT71 GCA_000153125.2_ASM15312v2
Dasania.marina.DSM.21967 GCA_000373485.1_ASM37348v1
100/100
Burkholderia.cepacia GCA_009586235.1_ASM958623v1
100/100
Ralstonia.pickettii GCA_902374465.1.MGYG.HGUT.01384
98.2/99
Methylotenera.versatilis.301 GCA_000093025.1_ASM9302v1
Nitrosospira.briensis.C.128 GCA_000619905.2_ASM61990v2
100/100
Magnetococcus.marinus.MC.1 GCA_000014865.1_ASM1486v1
Magnetofaba.australis.IT.1 GCA_002109495.1_ASM210949v1
100/100
Anaplasma.phagocytophilum.str.HZ GCA_000013125.1_ASM1312v1
100/100
Candidatus.Neoehrlichia.lotoris.str.RAC413 GCA_000964795.1_ASM96479v1
100/100
Ehrlichia.ruminantium.str.Welgevonden GCA_000026005.1_ASM2600v1
98.9/99
Wolbachia.endosymbiont.of.Drosophila.melanogaster GCA_000008025.1_ASM802v1
Candidatus.Deianiraea.vastatrix GCA_007993655.1_ASM799365v1
93.6/95
100/100
Candidatus.Jidaibacter.acanthamoeba GCA_000815465.1.UWC36
100/100
Rickettsiales.endosymbiont.of.Peranema.trichophorum GCA_004210275.1_ASM421027v1
100/100
Candidatus.Midichloria.mitochondrii.IricVA GCA_000219355.1_ASM21935v1
100/100
Rickettsiales.endosymbiont.of.Trichoplax.sp.H2 GCA_009690945.1
100/100
Candidatus.Megaira.polyxenophila
26.9/69
Rickettsia.bellii.RML369.C GCA_000012385.1_ASM1238v1
100/100
Rickettsiales.bacterium.Ac37b GCA_000746585.2_ASM74658v2
Rickettsiales.endosymbiont.of.Stachyamoeba.lipophora GCA_003932735.1_ASM393273v1
78.4/91
Ahrensia.kielensis.DSM.5890 GCA_000374465.1_ASM37446v1
83.1/93
Chelativorans.oligotrophicus GCA_011317505.1_ASM1131750v1
100/100
Rhizobium.leguminosarum.bv.trifolii GCA_011604525.1_ASM1160452v1
93.8/97
Bartonella.quintana GCA_009936175.1_ASM993617v1
Pelagibacterium.halotolerans GCA_013004005.1_ASM1300400v1
100/100
100/100
Polymorphum.gilvum.SL003B.26A1 GCA_000192745.1_ASM19274v1
100/100
Roseibium.hamelinense GCA_007830545.1_ASM783054v1
100/100
Pseudovibrio.sp.FO.BEG1 GCA_000236645.1_ASM23664v1
100/100
100/100
Methylocella.silvestris.BL2 GCA_000021745.1_ASM2174v1
Methylocystis.bryophila GCA_002117405.1_ASM211740v1
100/100
Methylorubrum.extorquens.PA1 GCA_000018845.1_ASM1884v1
98.1/99
100/100
Rhodopseudomonas.palustris GCA_013415845.1_ASM1341584v1
Xanthobacter.autotrophicus.Py2 GCA_000017645.1_ASM1764v1
100/100
100/100
Hyphomicrobium.denitrificans.ATCC.51888 GCA_000143145.1_ASM14314v1
Rhodomicrobium.vannielii.ATCC.17100 GCA_000166055.1_ASM16605v1
100/100
Thermopetrobacter.sp.TC1 GCA_000746275.1_ASM74627v1
100/100
Candidatus.Micropelagos.thuwalensis GCA_000469155.1.RS24
100/100
alpha.proteobacterium.IMCC14465 GCA_000293845.2_ASM29384v2
Parvibaculum.lavamentivorans.DS.1 GCA_000017565.1_ASM1756v1
100/100
Asticcacaulis.excentricus.CB.48 GCA_000175215.2_ASM17521v2
100/100
100/100
Brevundimonas.subvibrioides.ATCC.15264 GCA_000144605.1_ASM14460v1
alpha.proteobacterium.L41A GCA_000335735.1_L41A.1.0
100/100
100/100
Caulobacter.vibrioides.NA1000 GCA_000022005.1_ASM2200v1
100/100
Phenylobacterium.zucineum.HLK1 GCA_000017265.1_ASM1726v1
100/100
Hirschia.baltica.ATCC.49814 GCA_000023785.1_ASM2378v1
100/100
Hyphomonas.neptunium.ATCC.15444 GCA_000013025.1_ASM1302v1
100/100
100/100
Maricaulis.maris GCA_003634045.1_ASM363404v1
Oceanicaulis.alexandrii.DSM.11625 GCA_000420265.1_ASM42026v1
Parvularcula.bermudensis.HTCC2503 GCA_000152825.2_ASM15282v2
100/100
82.8/88
Jannaschia.rubra GCA_900113265.1.IMG.taxon.2622736430.annotated.assembly
100/100
Octadecabacter.arcticus.238 GCA_000155735.2_ASM15573v2
100/100
Ruegeria.pomeroyi.DSS.3 GCA_000011965.2_ASM1196v2
100/100
Sagittula.stellata.E.37 GCA_000169415.1_ASM16941v1
100/100
100/100
Ketogulonicigenium.vulgare GCA_001399515.1_ASM139951v1
Rubellimicrobium.rubrum GCA_006152115.1_ASM615211v1
100/100
100/100
Paracoccus.denitrificans GCA_004063735.1_ASM406373v1
Rhodobacter.sphaeroides.ATCC.17029 GCA_000015985.1_ASM1598v1
Neomegalonema.perideroedes.DSM.15528 GCA_000374145.1_ASM37414v1
100/100
100/100
Candidatus.Pelagibacter.ubique.HIMB058 GCA_000419545.1_ASM41954v1
100/100
Candidatus.Pelagibacter.ubique.HTCC1062 GCA_000012345.1_ASM1234v1
100/100
alpha.proteobacterium.HIMB5 GCA_000299095.1_ASM29909v1
alpha.proteobacterium.HIMB114 GCA_000163555.2
100/100
100/100
Citromicrobium.sp.JL477 GCA_001304795.1_ASM130479v1
100/100
Sphingopyxis.macrogoltabida GCA_001314325.1_ASM131432v1
100/100
Sphingobium.indicum.B90A GCA_000264945.2_ASM26494v2
100/100
100/100
Sphingomonas.wittichii.RW1 GCA_000016765.1_ASM1676v1
100/100
Zymomonas.mobilis.subsp.mobilis.ATCC.31821 GCA_000007105.1_ASM710v1
alpha.proteobacterium.AAP81b GCA_001295935.1_ASM129593v1
Kordiimonas.gwangyangensis.DSM.19435 GCA_000375545.1_ASM37554v1
Sneathiella.glossodoripedis.JCM.23214 GCA_000616095.1_ASM61609v1
100/100
GCA_002296185.1_ASM229618v1*
100/100
GCA_002728135.1_ASM272813v1*
100/100
GCA_002717225.1_ASM271722v1*
GCA_002937555.1_ASM293755v1*
100/100
100/100
GCA_002501105.1_ASM250110v1*
100/100
GCA_002728255.1_ASM272825v1*
100/98
100/100
100/100
GCA_002937585.1_ASM293758v1*
GCA_002938115.1_ASM293811v1*
GCA_002721365.1_ASM272136v1*
100/100
Geminicoccus.roseus.DSM.18922 GCA_000427665.1_ASM42766v1
Tistrella.mobilis.KA081020.065 GCA_000264455.2_ASM26445v2
Caenispirillum.salinarum.AK4 GCA_000315795.1
100/100
100/100
Pararhodospirillum.photometricum.DSM.122 GCA_000284415.2_ASM28441v2
89.7/99
Rhodospirillum.rubrum.F11 GCA_000225955.1_ASM22595v1
100/100
Roseospirillum.parvum GCA_900100455.1
100/100
Magnetospirillum.magneticum.AMB.1 GCA_000009985.1_ASM998v1
100/100
Terasakiella.pusilla.DSM.6293 GCA_000688235.1_ASM68823v1
100/98
Thalassospira.profundimaris.WP0211 GCA_000300275.1_ASM30027v1
100/100
Candidatus.Puniceispirillum.marinum.IMCC1322 GCA_000024465.1_ASM2446v1
100/59
100/100
100/100
Thalassobaculum.salexigens.DSM.19539 GCA_000423805.1_ASM42380v1
alpha.proteobacterium.BAL199 GCA_000171835.1_ASM17183v1
100/100
Oceanibaculum.indicum GCA_003633955.1_ASM363395v1
100/100
Kiloniella.laminariae.DSM.19542 GCA_000374005.1_ASM37400v1
100/100
Rhodovibrio.salinarum.DSM.9154 GCA_000515255.1_ASM51525v1
100/100
Inquilinus.limosus.DSM.16000 GCA_000423185.1_ASM42318v1
100/100
Rhodospirillum.centenum GCA_014197805.1_ASM1419780v1
98.4/59
Micavibrio.aeruginosavorus.ARL.13 GCA_000226315.1_ASM22631v1
100/100
Caedimonas.varicaedens GCA_001192655.1_ASM119265v1
100/100
Candidatus.Caedibacter.acanthamoebae GCA_000743035.1_ASM74303v1
Candidatus.Nucleicultrix.amoebiphila.FS5 GCA_002117145.1_ASM211714v1
100/100
100/100
Candidatus.Finniella.inopinata GCA_004210305.1_ASM421030v1
100/100
Candidatus.Odyssella.thessalonicensis.L13 GCA_000190415.2_ASM19041v1
74.6/88
Candidatus.Paracaedibacter.acanthamoebae GCA_000742835.1_ASM74283v1
100/100
100/100
Candidatus.Hepatobacter.penaei GCA_000742475.1_ASM74247v1
Holospora.undulata.HU1 GCA_000388175.3_ASM38817v3
Elioraea.tepidiphila.DSM.17972 GCA_000378465.1_ASM37846v1
99.8/100
Belnapia.moabensis.DSM.16746 GCA_000745835.1_ASM74583v1
100/100
100/100
Rubritepida.flocculans.DSM.14296 GCA_000425365.1_ASM42536v1
100/100
Roseomonas.gilardii GCA_001941945.1_ASM194194v1
Acidiphilium.multivorum.AIU301 GCA_000202835.1_ASM20283v1
91.7/95
58.8/88
Acidisphaera.rubrifaciens.HS.AP3 GCA_000964365.1_ASM96436v1
100/100
Granulibacter.bethesdensis.CGDNIH4 GCA_000576145.2_ASM57614v2
100/100
Commensalibacter.intestini.A911 GCA_000231445.2_ASM23144v1
99/100
Gluconacetobacter.diazotrophicus.PA1.5 GCA_000067045.1_ASM6704v1
Acetobacter.aceti.NBRC.14818 GCA_000193495.2_ASM19349v2
Gluconobacter.oxydans.DSM.3504 GCA_000583855.1_ASM58385v1
0.559041

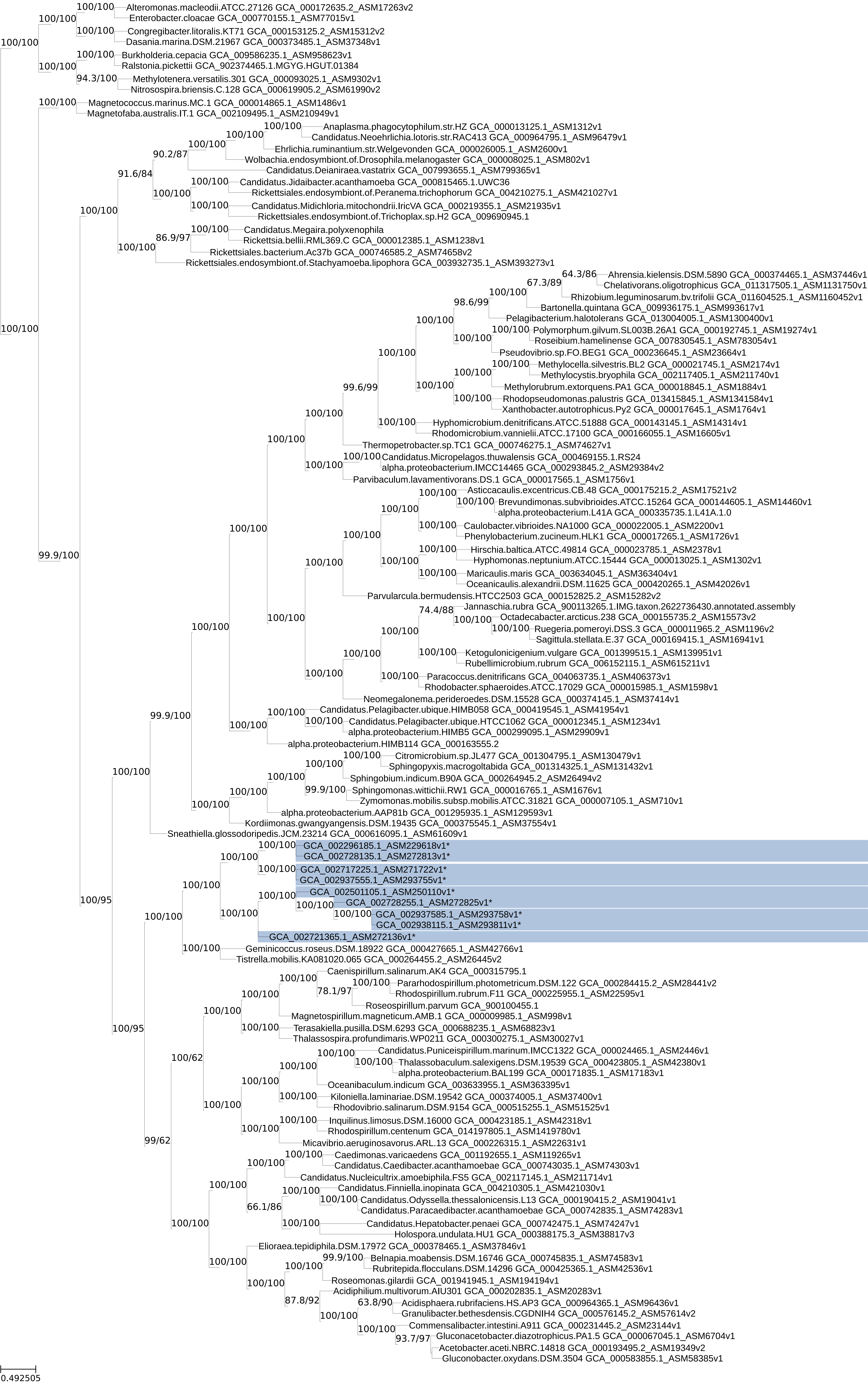

100/100
100/100
100/100
Alteromonas.macleodii.ATCC.27126 GCA_000172635.2_ASM17263v2
Enterobacter.cloacae GCA_000770155.1_ASM77015v1
100/100
Congregibacter.litoralis.KT71 GCA_000153125.2_ASM15312v2
Dasania.marina.DSM.21967 GCA_000373485.1_ASM37348v1
100/100
Burkholderia.cepacia GCA_009586235.1_ASM958623v1
100/100
Ralstonia.pickettii GCA_902374465.1.MGYG.HGUT.01384
94.3/100
Methylotenera.versatilis.301 GCA_000093025.1_ASM9302v1
Nitrosospira.briensis.C.128 GCA_000619905.2_ASM61990v2
100/100
Magnetococcus.marinus.MC.1 GCA_000014865.1_ASM1486v1
Magnetofaba.australis.IT.1 GCA_002109495.1_ASM210949v1
100/100
Anaplasma.phagocytophilum.str.HZ GCA_000013125.1_ASM1312v1
100/100
Candidatus.Neoehrlichia.lotoris.str.RAC413 GCA_000964795.1_ASM96479v1
100/100
Ehrlichia.ruminantium.str.Welgevonden GCA_000026005.1_ASM2600v1
90.2/87
Wolbachia.endosymbiont.of.Drosophila.melanogaster GCA_000008025.1_ASM802v1
Candidatus.Deianiraea.vastatrix GCA_007993655.1_ASM799365v1
91.6/84
100/100
Candidatus.Jidaibacter.acanthamoeba GCA_000815465.1.UWC36
100/100
Rickettsiales.endosymbiont.of.Peranema.trichophorum GCA_004210275.1_ASM421027v1
100/100
Candidatus.Midichloria.mitochondrii.IricVA GCA_000219355.1_ASM21935v1
Rickettsiales.endosymbiont.of.Trichoplax.sp.H2 GCA_009690945.1
100/100
Candidatus.Megaira.polyxenophila
86.9/97
Rickettsia.bellii.RML369.C GCA_000012385.1_ASM1238v1
100/100
Rickettsiales.bacterium.Ac37b GCA_000746585.2_ASM74658v2
Rickettsiales.endosymbiont.of.Stachyamoeba.lipophora GCA_003932735.1_ASM393273v1
100/100
64.3/86
Ahrensia.kielensis.DSM.5890 GCA_000374465.1_ASM37446v1
67.3/89
Chelativorans.oligotrophicus GCA_011317505.1_ASM1131750v1
100/100
Rhizobium.leguminosarum.bv.trifolii GCA_011604525.1_ASM1160452v1
98.6/99
Bartonella.quintana GCA_009936175.1_ASM993617v1
Pelagibacterium.halotolerans GCA_013004005.1_ASM1300400v1
100/100
100/100
Polymorphum.gilvum.SL003B.26A1 GCA_000192745.1_ASM19274v1
100/100
Roseibium.hamelinense GCA_007830545.1_ASM783054v1
Pseudovibrio.sp.FO.BEG1 GCA_000236645.1_ASM23664v1
100/100
100/100
Methylocella.silvestris.BL2 GCA_000021745.1_ASM2174v1
100/100
Methylocystis.bryophila GCA_002117405.1_ASM211740v1
99.6/99
100/100
Methylorubrum.extorquens.PA1 GCA_000018845.1_ASM1884v1
100/100
Rhodopseudomonas.palustris GCA_013415845.1_ASM1341584v1
Xanthobacter.autotrophicus.Py2 GCA_000017645.1_ASM1764v1
100/100
100/100
Hyphomicrobium.denitrificans.ATCC.51888 GCA_000143145.1_ASM14314v1
Rhodomicrobium.vannielii.ATCC.17100 GCA_000166055.1_ASM16605v1
100/100
Thermopetrobacter.sp.TC1 GCA_000746275.1_ASM74627v1
100/100
Candidatus.Micropelagos.thuwalensis GCA_000469155.1.RS24
100/100
alpha.proteobacterium.IMCC14465 GCA_000293845.2_ASM29384v2
Parvibaculum.lavamentivorans.DS.1 GCA_000017565.1_ASM1756v1
100/100
Asticcacaulis.excentricus.CB.48 GCA_000175215.2_ASM17521v2
100/100
Brevundimonas.subvibrioides.ATCC.15264 GCA_000144605.1_ASM14460v1
100/100
alpha.proteobacterium.L41A GCA_000335735.1.L41A.1.0
100/100
Caulobacter.vibrioides.NA1000 GCA_000022005.1_ASM2200v1
100/100
Phenylobacterium.zucineum.HLK1 GCA_000017265.1_ASM1726v1
100/100
Hirschia.baltica.ATCC.49814 GCA_000023785.1_ASM2378v1
100/100
Hyphomonas.neptunium.ATCC.15444 GCA_000013025.1_ASM1302v1
99.9/100
100/100
Maricaulis.maris GCA_003634045.1_ASM363404v1
100/100
Oceanicaulis.alexandrii.DSM.11625 GCA_000420265.1_ASM42026v1
Parvularcula.bermudensis.HTCC2503 GCA_000152825.2_ASM15282v2
74.4/88
Jannaschia.rubra GCA_900113265.1.IMG.taxon.2622736430.annotated.assembly
100/100
100/100
Octadecabacter.arcticus.238 GCA_000155735.2_ASM15573v2
100/100
Ruegeria.pomeroyi.DSS.3 GCA_000011965.2_ASM1196v2
100/100
Sagittula.stellata.E.37 GCA_000169415.1_ASM16941v1
100/100
Ketogulonicigenium.vulgare GCA_001399515.1_ASM139951v1
100/100
Rubellimicrobium.rubrum GCA_006152115.1_ASM615211v1
100/100
Paracoccus.denitrificans GCA_004063735.1_ASM406373v1
Rhodobacter.sphaeroides.ATCC.17029 GCA_000015985.1_ASM1598v1
Neomegalonema.perideroedes.DSM.15528 GCA_000374145.1_ASM37414v1
100/100
Candidatus.Pelagibacter.ubique.HIMB058 GCA_000419545.1_ASM41954v1
100/100
100/100
Candidatus.Pelagibacter.ubique.HTCC1062 GCA_000012345.1_ASM1234v1
alpha.proteobacterium.HIMB5 GCA_000299095.1_ASM29909v1
alpha.proteobacterium.HIMB114 GCA_000163555.2
100/100
100/100
Citromicrobium.sp.JL477 GCA_001304795.1_ASM130479v1
100/100
Sphingopyxis.macrogoltabida GCA_001314325.1_ASM131432v1
100/100
Sphingobium.indicum.B90A GCA_000264945.2_ASM26494v2
100/100
99.9/100
Sphingomonas.wittichii.RW1 GCA_000016765.1_ASM1676v1
100/100
Zymomonas.mobilis.subsp.mobilis.ATCC.31821 GCA_000007105.1_ASM710v1
alpha.proteobacterium.AAP81b GCA_001295935.1_ASM129593v1
Kordiimonas.gwangyangensis.DSM.19435 GCA_000375545.1_ASM37554v1
Sneathiella.glossodoripedis.JCM.23214 GCA_000616095.1_ASM61609v1
100/100
GCA_002296185.1_ASM229618v1*
100/100
GCA_002728135.1_ASM272813v1*
100/100
GCA_002717225.1_ASM271722v1*
GCA_002937555.1_ASM293755v1*
100/100
GCA_002501105.1_ASM250110v1*
100/100
100/100
GCA_002728255.1_ASM272825v1*
100/100
GCA_002937585.1_ASM293758v1*
GCA_002938115.1_ASM293811v1*
100/95
GCA_002721365.1_ASM272136v1*
100/100
Geminicoccus.roseus.DSM.18922 GCA_000427665.1_ASM42766v1
Tistrella.mobilis.KA081020.065 GCA_000264455.2_ASM26445v2
100/100
Caenispirillum.salinarum.AK4 GCA_000315795.1
100/100
Pararhodospirillum.photometricum.DSM.122 GCA_000284415.2_ASM28441v2
78.1/97
Rhodospirillum.rubrum.F11 GCA_000225955.1_ASM22595v1
100/100
Roseospirillum.parvum GCA_900100455.1
Magnetospirillum.magneticum.AMB.1 GCA_000009985.1_ASM998v1
100/95
100/100
Terasakiella.pusilla.DSM.6293 GCA_000688235.1_ASM68823v1
Thalassospira.profundimaris.WP0211 GCA_000300275.1_ASM30027v1
100/62
100/100
Candidatus.Puniceispirillum.marinum.IMCC1322 GCA_000024465.1_ASM2446v1
100/100
Thalassobaculum.salexigens.DSM.19539 GCA_000423805.1_ASM42380v1
100/100
alpha.proteobacterium.BAL199 GCA_000171835.1_ASM17183v1
Oceanibaculum.indicum GCA_003633955.1_ASM363395v1
100/100
100/100
Kiloniella.laminariae.DSM.19542 GCA_000374005.1_ASM37400v1
Rhodovibrio.salinarum.DSM.9154 GCA_000515255.1_ASM51525v1
100/100
Inquilinus.limosus.DSM.16000 GCA_000423185.1_ASM42318v1
100/100
Rhodospirillum.centenum GCA_014197805.1_ASM1419780v1
Micavibrio.aeruginosavorus.ARL.13 GCA_000226315.1_ASM22631v1
99/62
100/100
Caedimonas.varicaedens GCA_001192655.1_ASM119265v1
100/100
Candidatus.Caedibacter.acanthamoebae GCA_000743035.1_ASM74303v1
Candidatus.Nucleicultrix.amoebiphila.FS5 GCA_002117145.1_ASM211714v1
100/100
100/100
Candidatus.Finniella.inopinata GCA_004210305.1_ASM421030v1
100/100
Candidatus.Odyssella.thessalonicensis.L13 GCA_000190415.2_ASM19041v1
66.1/86
Candidatus.Paracaedibacter.acanthamoebae GCA_000742835.1_ASM74283v1
100/100
100/100
Candidatus.Hepatobacter.penaei GCA_000742475.1_ASM74247v1
Holospora.undulata.HU1 GCA_000388175.3_ASM38817v3
Elioraea.tepidiphila.DSM.17972 GCA_000378465.1_ASM37846v1
99.9/100
Belnapia.moabensis.DSM.16746 GCA_000745835.1_ASM74583v1
100/100
Rubritepida.flocculans.DSM.14296 GCA_000425365.1_ASM42536v1
100/100
100/100
Roseomonas.gilardii GCA_001941945.1_ASM194194v1
Acidiphilium.multivorum.AIU301 GCA_000202835.1_ASM20283v1
63.8/90
Acidisphaera.rubrifaciens.HS.AP3 GCA_000964365.1_ASM96436v1
87.8/92
Granulibacter.bethesdensis.CGDNIH4 GCA_000576145.2_ASM57614v2
100/100
Commensalibacter.intestini.A911 GCA_000231445.2_ASM23144v1
100/100
Gluconacetobacter.diazotrophicus.PA1.5 GCA_000067045.1_ASM6704v1
93.7/97
Acetobacter.aceti.NBRC.14818 GCA_000193495.2_ASM19349v2
Gluconobacter.oxydans.DSM.3504 GCA_000583855.1_ASM58385v1
0.492505

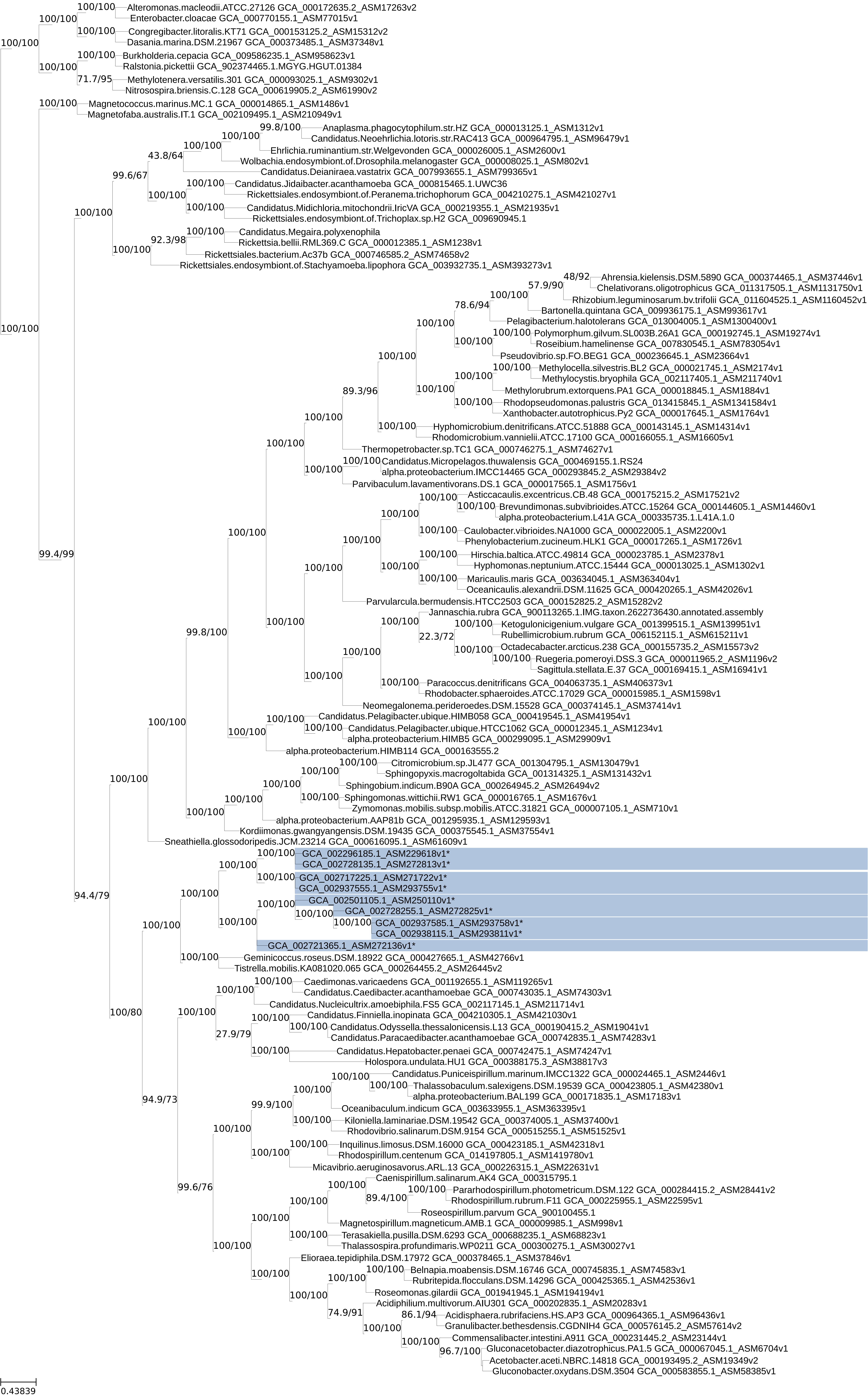
Alteromonas.macleodii.ATCC.27126 GCA_000172635.2_ASM17263v2
Enterobacter.cloacae GCA_000770155.1_ASM77015v1
Congregibacter.litoralis.KT71 GCA_000153125.2_ASM15312v2
Dasania.marina.DSM.21967 GCA_000373485.1_ASM37348v1
Burkholderia.cepacia GCA_009586235.1_ASM958623v1
Ralstonia.pickettii GCA_902374465.1.MGYG.HGUT.01384
Methylotenera.versatilis.301 GCA_000093025.1_ASM9302v1
Nitrosospira.briensis.C.128 GCA_000619905.2_ASM61990v2
Magnetococcus.marinus.MC.1 GCA_000014865.1_ASM1486v1
Magnetofaba.australis.IT.1 GCA_002109495.1_ASM210949v1
Anaplasma.phagocytophilum.str.HZ GCA_000013125.1_ASM1312v1
Candidatus.Neoehrlichia.lotoris.str.RAC413 GCA_000964795.1_ASM96479v1
Ehrlichia.ruminantium.str.Welgevonden GCA_000026005.1_ASM2600v1
Wolbachia.endosymbiont.of.Drosophila.melanogaster GCA_000008025.1_ASM802v1
Candidatus.Deianiraea.vastatrix GCA_007993655.1_ASM799365v1
Candidatus.Jidaibacter.acanthamoeba GCA_000815465.1.UWC36
Rickettsiales.endosymbiont.of.Peranema.trichophorum GCA_004210275.1_ASM421027v1
Candidatus.Midichloria.mitochondrii.IricVA GCA_000219355.1_ASM21935v1
Rickettsiales.endosymbiont.of.Trichoplax.sp.H2 GCA_009690945.1
Candidatus.Megaira.polyxenophila
Rickettsia.bellii.RML369.C GCA_000012385.1_ASM1238v1
Rickettsiales.bacterium.Ac37b GCA_000746585.2_ASM74658v2
Rickettsiales.endosymbiont.of.Stachyamoeba.lipophora GCA_003932735.1_ASM393273v1
Ahrensia.kielensis.DSM.5890 GCA_000374465.1_ASM37446v1
Chelativorans.oligotrophicus GCA_011317505.1_ASM1131750v1
Rhizobium.leguminosarum.bv.trifolii GCA_011604525.1_ASM1160452v1
Bartonella.quintana GCA_009936175.1_ASM993617v1
Pelagibacterium.halotolerans GCA_013004005.1_ASM1300400v1
Polymorphum.gilvum.SL003B.26A1 GCA_000192745.1_ASM19274v1
Roseibium.hamelinense GCA_007830545.1_ASM783054v1
Pseudovibrio.sp.FO.BEG1 GCA_000236645.1_ASM23664v1
Methylocella.silvestris.BL2 GCA_000021745.1_ASM2174v1
Methylocystis.bryophila GCA_002117405.1_ASM211740v1
Methylorubrum.extorquens.PA1 GCA_000018845.1_ASM1884v1
Rhodopseudomonas.palustris GCA_013415845.1_ASM1341584v1
Xanthobacter.autotrophicus.Py2 GCA_000017645.1_ASM1764v1
Hyphomicrobium.denitrificans.ATCC.51888 GCA_000143145.1_ASM14314v1
Rhodomicrobium.vannielii.ATCC.17100 GCA_000166055.1_ASM16605v1
Thermopetrobacter.sp.TC1 GCA_000746275.1_ASM74627v1
Candidatus.Micropelagos.thuwalensis GCA_000469155.1.RS24
alpha.proteobacterium.IMCC14465 GCA_000293845.2_ASM29384v2
Parvibaculum.lavamentivorans.DS.1 GCA_000017565.1_ASM1756v1
Asticcacaulis.excentricus.CB.48 GCA_000175215.2_ASM17521v2
Brevundimonas.subvibrioides.ATCC.15264 GCA_000144605.1_ASM14460v1
alpha.proteobacterium.L41A GCA_000335735.1.L41A.1.0
Caulobacter.vibrioides.NA1000 GCA_000022005.1_ASM2200v1
Phenylobacterium.zucineum.HLK1 GCA_000017265.1_ASM1726v1
Hirschia.baltica.ATCC.49814 GCA_000023785.1_ASM2378v1
Hyphomonas.neptunium.ATCC.15444 GCA_000013025.1_ASM1302v1
Maricaulis.maris GCA_003634045.1_ASM363404v1
Oceanicaulis.alexandrii.DSM.11625 GCA_000420265.1_ASM42026v1
Parvularcula.bermudensis.HTCC2503 GCA_000152825.2_ASM15282v2
Jannaschia.rubra GCA_900113265.1.IMG.taxon.2622736430.annotated.assembly
Ketogulonicigenium.vulgare GCA_001399515.1_ASM139951v1
Rubellimicrobium.rubrum GCA_006152115.1_ASM615211v1
Octadecabacter.arcticus.238 GCA_000155735.2_ASM15573v2
Ruegeria.pomeroyi.DSS.3 GCA_000011965.2_ASM1196v2
Sagittula.stellata.E.37 GCA_000169415.1_ASM16941v1
Paracoccus.denitrificans GCA_004063735.1_ASM406373v1
Rhodobacter.sphaeroides.ATCC.17029 GCA_000015985.1_ASM1598v1
Neomegalonema.perideroedes.DSM.15528 GCA_000374145.1_ASM37414v1
Candidatus.Pelagibacter.ubique.HIMB058 GCA_000419545.1_ASM41954v1
Candidatus.Pelagibacter.ubique.HTCC1062 GCA_000012345.1_ASM1234v1
alpha.proteobacterium.HIMB5 GCA_000299095.1_ASM29909v1
alpha.proteobacterium.HIMB114 GCA_000163555.2
Citromicrobium.sp.JL477 GCA_001304795.1_ASM130479v1
Sphingopyxis.macrogoltabida GCA_001314325.1_ASM131432v1
Sphingobium.indicum.B90A GCA_000264945.2_ASM26494v2
Sphingomonas.wittichii.RW1 GCA_000016765.1_ASM1676v1
Zymomonas.mobilis.subsp.mobilis.ATCC.31821 GCA_000007105.1_ASM710v1
alpha.proteobacterium.AAP81b GCA_001295935.1_ASM129593v1
Kordiimonas.gwangyangensis.DSM.19435 GCA_000375545.1_ASM37554v1
Sneathiella.glossodoripedis.JCM.23214 GCA_000616095.1_ASM61609v1
GCA_002296185.1_ASM229618v1*
GCA_002728135.1_ASM272813v1*
GCA_002717225.1_ASM271722v1*
GCA_002937555.1_ASM293755v1*
GCA_002501105.1_ASM250110v1*
GCA_002728255.1_ASM272825v1*
GCA_002937585.1_ASM293758v1*
GCA_002938115.1_ASM293811v1*
GCA_002721365.1_ASM272136v1*
Geminicoccus.roseus.DSM.18922 GCA_000427665.1_ASM42766v1
Tistrella.mobilis.KA081020.065 GCA_000264455.2_ASM26445v2
Caedimonas.varicaedens GCA_001192655.1_ASM119265v1
Candidatus.Caedibacter.acanthamoebae GCA_000743035.1_ASM74303v1
Candidatus.Nucleicultrix.amoebiphila.FS5 GCA_002117145.1_ASM211714v1
Candidatus.Finniella.inopinata GCA_004210305.1_ASM421030v1
Candidatus.Odyssella.thessalonicensis.L13 GCA_000190415.2_ASM19041v1
Candidatus.Paracaedibacter.acanthamoebae GCA_000742835.1_ASM74283v1
Candidatus.Hepatobacter.penaei GCA_000742475.1_ASM74247v1
Holospora.undulata.HU1 GCA_000388175.3_ASM38817v3
Candidatus.Puniceispirillum.marinum.IMCC1322 GCA_000024465.1_ASM2446v1
Thalassobaculum.salexigens.DSM.19539 GCA_000423805.1_ASM42380v1
alpha.proteobacterium.BAL199 GCA_000171835.1_ASM17183v1
Oceanibaculum.indicum GCA_003633955.1_ASM363395v1
Kiloniella.laminariae.DSM.19542 GCA_000374005.1_ASM37400v1
Rhodovibrio.salinarum.DSM.9154 GCA_000515255.1_ASM51525v1
Inquilinus.limosus.DSM.16000 GCA_000423185.1_ASM42318v1
Rhodospirillum.centenum GCA_014197805.1_ASM1419780v1
Micavibrio.aeruginosavorus.ARL.13 GCA_000226315.1_ASM22631v1
Caenispirillum.salinarum.AK4 GCA_000315795.1
Pararhodospirillum.photometricum.DSM.122 GCA_000284415.2_ASM28441v2
Rhodospirillum.rubrum.F11 GCA_000225955.1_ASM22595v1
Roseospirillum.parvum GCA_900100455.1
Magnetospirillum.magneticum.AMB.1 GCA_000009985.1_ASM998v1
Terasakiella.pusilla.DSM.6293 GCA_000688235.1_ASM68823v1
Thalassospira.profundimaris.WP0211 GCA_000300275.1_ASM30027v1
Elioraea.tepidiphila.DSM.17972 GCA_000378465.1_ASM37846v1
Belnapia.moabensis.DSM.16746 GCA_000745835.1_ASM74583v1
Rubritepida.flocculans.DSM.14296 GCA_000425365.1_ASM42536v1
Roseomonas.gilardii GCA_001941945.1_ASM194194v1
Acidiphilium.multivorum.AIU301 GCA_000202835.1_ASM20283v1
Acidisphaera.rubrifaciens.HS.AP3 GCA_000964365.1_ASM96436v1
Granulibacter.bethesdensis.CGDNIH4 GCA_000576145.2_ASM57614v2
Commensalibacter.intestini.A911 GCA_000231445.2_ASM23144v1
Gluconacetobacter.diazotrophicus.PA1.5 GCA_000067045.1_ASM6704v1
Acetobacter.aceti.NBRC.14818 GCA_000193495.2_ASM19349v2
Gluconobacter.oxydans.DSM.3504 GCA_000583855.1_ASM58385v1
0.43839

### Group F

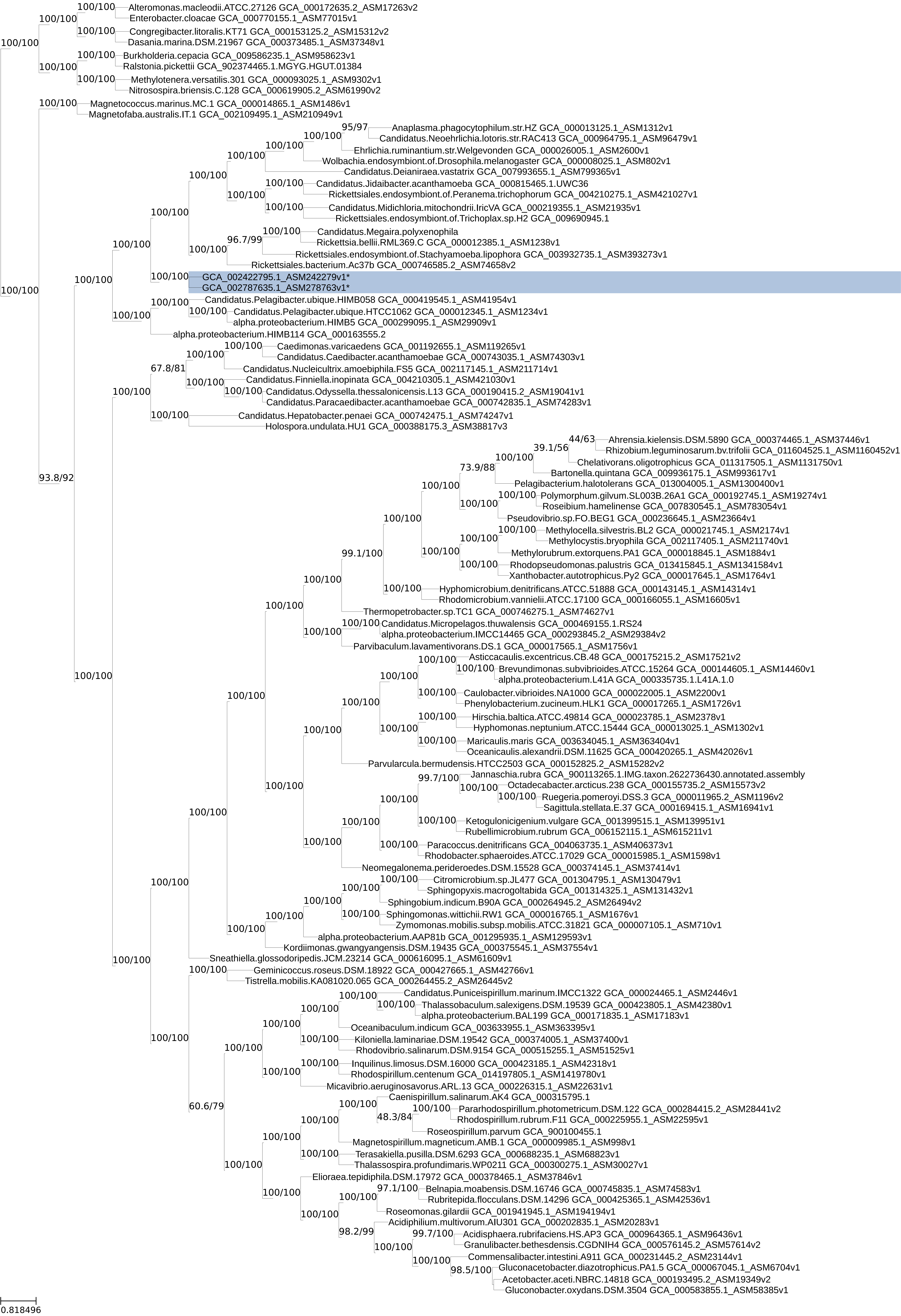
Alteromonas.macleodii.ATCC.27126 GCA_000172635.2_ASM17263v2
Enterobacter.cloacae GCA_000770155.1_ASM77015v1
Congregibacter.litoralis.KT71 GCA_000153125.2_ASM15312v2
Dasania.marina.DSM.21967 GCA_000373485.1_ASM37348v1
Burkholderia.cepacia GCA_009586235.1_ASM958623v1
Ralstonia.pickettii GCA_902374465.1.MGYG.HGUT.01384
Methylotenera.versatilis.301 GCA_000093025.1_ASM9302v1
Nitrosospira.briensis.C.128 GCA_000619905.2_ASM61990v2
Magnetococcus.marinus.MC.1 GCA_000014865.1_ASM1486v1
Magnetofaba.australis.IT.1 GCA_002109495.1_ASM210949v1
Anaplasma.phagocytophilum.str.HZ GCA_000013125.1_ASM1312v1
Candidatus.Neoehrlichia.lotoris.str.RAC413 GCA_000964795.1_ASM96479v1
Ehrlichia.ruminantium.str.Welgevonden GCA_000026005.1_ASM2600v1
Wolbachia.endosymbiont.of.Drosophila.melanogaster GCA_000008025.1_ASM802v1
Candidatus.Deianiraea.vastatrix GCA_007993655.1_ASM799365v1
Candidatus.Jidaibacter.acanthamoeba GCA_000815465.1.UWC36
Rickettsiales.endosymbiont.of.Peranema.trichophorum GCA_004210275.1_ASM421027v1
Candidatus.Midichloria.mitochondrii.IricVA GCA_000219355.1_ASM21935v1
Rickettsiales.endosymbiont.of.Trichoplax.sp.H2 GCA_009690945.1
Candidatus.Megaira.polyxenophila
Rickettsia.bellii.RML369.C GCA_000012385.1_ASM1238v1
Rickettsiales.endosymbiont.of.Stachyamoeba.lipophora GCA_003932735.1_ASM393273v1
Rickettsiales.bacterium.Ac37b GCA_000746585.2_ASM74658v2
GCA_002422795.1_ASM242279v1*
GCA_002787635.1_ASM278763v1*
Candidatus.Pelagibacter.ubique.HIMB058 GCA_000419545.1_ASM41954v1
Candidatus.Pelagibacter.ubique.HTCC1062 GCA_000012345.1_ASM1234v1
alpha.proteobacterium.HIMB5 GCA_000299095.1_ASM29909v1
alpha.proteobacterium.HIMB114 GCA_000163555.2
Caedimonas.varicaedens GCA_001192655.1_ASM119265v1
Candidatus.Caedibacter.acanthamoebae GCA_000743035.1_ASM74303v1
Candidatus.Nucleicultrix.amoebiphila.FS5 GCA_002117145.1_ASM211714v1
Candidatus.Finniella.inopinata GCA_004210305.1_ASM421030v1
Candidatus.Odyssella.thessalonicensis.L13 GCA_000190415.2_ASM19041v1
Candidatus.Paracaedibacter.acanthamoebae GCA_000742835.1_ASM74283v1
Candidatus.Hepatobacter.penaei GCA_000742475.1_ASM74247v1
Holospora.undulata.HU1 GCA_000388175.3_ASM38817v3
Ahrensia.kielensis.DSM.5890 GCA_000374465.1_ASM37446v1
Rhizobium.leguminosarum.bv.trifolii GCA_011604525.1_ASM1160452v1
Chelativorans.oligotrophicus GCA_011317505.1_ASM1131750v1
Bartonella.quintana GCA_009936175.1_ASM993617v1
Pelagibacterium.halotolerans GCA_013004005.1_ASM1300400v1
Polymorphum.gilvum.SL003B.26A1 GCA_000192745.1_ASM19274v1
Roseibium.hamelinense GCA_007830545.1_ASM783054v1
Pseudovibrio.sp.FO.BEG1 GCA_000236645.1_ASM23664v1
Methylocella.silvestris.BL2 GCA_000021745.1_ASM2174v1
Methylocystis.bryophila GCA_002117405.1_ASM211740v1
Methylorubrum.extorquens.PA1 GCA_000018845.1_ASM1884v1
Rhodopseudomonas.palustris GCA_013415845.1_ASM1341584v1
Xanthobacter.autotrophicus.Py2 GCA_000017645.1_ASM1764v1
Hyphomicrobium.denitrificans.ATCC.51888 GCA_000143145.1_ASM14314v1
Rhodomicrobium.vannielii.ATCC.17100 GCA_000166055.1_ASM16605v1
Thermopetrobacter.sp.TC1 GCA_000746275.1_ASM74627v1
Candidatus.Micropelagos.thuwalensis GCA_000469155.1.RS24
alpha.proteobacterium.IMCC14465 GCA_000293845.2_ASM29384v2
Parvibaculum.lavamentivorans.DS.1 GCA_000017565.1_ASM1756v1
Asticcacaulis.excentricus.CB.48 GCA_000175215.2_ASM17521v2
Brevundimonas.subvibrioides.ATCC.15264 GCA_000144605.1_ASM14460v1
alpha.proteobacterium.L41A GCA_000335735.1.L41A.1.0
Caulobacter.vibrioides.NA1000 GCA_000022005.1_ASM2200v1
Phenylobacterium.zucineum.HLK1 GCA_000017265.1_ASM1726v1
Hirschia.baltica.ATCC.49814 GCA_000023785.1_ASM2378v1
Hyphomonas.neptunium.ATCC.15444 GCA_000013025.1_ASM1302v1
Maricaulis.maris GCA_003634045.1_ASM363404v1
Oceanicaulis.alexandrii.DSM.11625 GCA_000420265.1_ASM42026v1
Parvularcula.bermudensis.HTCC2503 GCA_000152825.2_ASM15282v2
Jannaschia.rubra GCA_900113265.1.IMG.taxon.2622736430.annotated.assembly
Octadecabacter.arcticus.238 GCA_000155735.2_ASM15573v2
Ruegeria.pomeroyi.DSS.3 GCA_000011965.2_ASM1196v2
Sagittula.stellata.E.37 GCA_000169415.1_ASM16941v1
Ketogulonicigenium.vulgare GCA_001399515.1_ASM139951v1
Rubellimicrobium.rubrum GCA_006152115.1_ASM615211v1
Paracoccus.denitrificans GCA_004063735.1_ASM406373v1
Rhodobacter.sphaeroides.ATCC.17029 GCA_000015985.1_ASM1598v1
Neomegalonema.perideroedes.DSM.15528 GCA_000374145.1_ASM37414v1
Citromicrobium.sp.JL477 GCA_001304795.1_ASM130479v1
Sphingopyxis.macrogoltabida GCA_001314325.1_ASM131432v1
Sphingobium.indicum.B90A GCA_000264945.2_ASM26494v2
Sphingomonas.wittichii.RW1 GCA_000016765.1_ASM1676v1
Zymomonas.mobilis.subsp.mobilis.ATCC.31821 GCA_000007105.1_ASM710v1
alpha.proteobacterium.AAP81b GCA_001295935.1_ASM129593v1
Kordiimonas.gwangyangensis.DSM.19435 GCA_000375545.1_ASM37554v1
Sneathiella.glossodoripedis.JCM.23214 GCA_000616095.1_ASM61609v1
Geminicoccus.roseus.DSM.18922 GCA_000427665.1_ASM42766v1
Tistrella.mobilis.KA081020.065 GCA_000264455.2_ASM26445v2
Candidatus.Puniceispirillum.marinum.IMCC1322 GCA_000024465.1_ASM2446v1
Thalassobaculum.salexigens.DSM.19539 GCA_000423805.1_ASM42380v1
alpha.proteobacterium.BAL199 GCA_000171835.1_ASM17183v1
Oceanibaculum.indicum GCA_003633955.1_ASM363395v1
Kiloniella.laminariae.DSM.19542 GCA_000374005.1_ASM37400v1
Rhodovibrio.salinarum.DSM.9154 GCA_000515255.1_ASM51525v1
Inquilinus.limosus.DSM.16000 GCA_000423185.1_ASM42318v1
Rhodospirillum.centenum GCA_014197805.1_ASM1419780v1
Micavibrio.aeruginosavorus.ARL.13 GCA_000226315.1_ASM22631v1
Caenispirillum.salinarum.AK4 GCA_000315795.1
Pararhodospirillum.photometricum.DSM.122 GCA_000284415.2_ASM28441v2
Rhodospirillum.rubrum.F11 GCA_000225955.1_ASM22595v1
Roseospirillum.parvum GCA_900100455.1
Magnetospirillum.magneticum.AMB.1 GCA_000009985.1_ASM998v1
Terasakiella.pusilla.DSM.6293 GCA_000688235.1_ASM68823v1
Thalassospira.profundimaris.WP0211 GCA_000300275.1_ASM30027v1
Elioraea.tepidiphila.DSM.17972 GCA_000378465.1_ASM37846v1
Belnapia.moabensis.DSM.16746 GCA_000745835.1_ASM74583v1
Rubritepida.flocculans.DSM.14296 GCA_000425365.1_ASM42536v1
Roseomonas.gilardii GCA_001941945.1_ASM194194v1
Acidiphilium.multivorum.AIU301 GCA_000202835.1_ASM20283v1
Acidisphaera.rubrifaciens.HS.AP3 GCA_000964365.1_ASM96436v1
Granulibacter.bethesdensis.CGDNIH4 GCA_000576145.2_ASM57614v2
Commensalibacter.intestini.A911 GCA_000231445.2_ASM23144v1
Gluconacetobacter.diazotrophicus.PA1.5 GCA_000067045.1_ASM6704v1
Acetobacter.aceti.NBRC.14818 GCA_000193495.2_ASM19349v2
Gluconobacter.oxydans.DSM.3504 GCA_000583855.1_ASM58385v1
0.818496

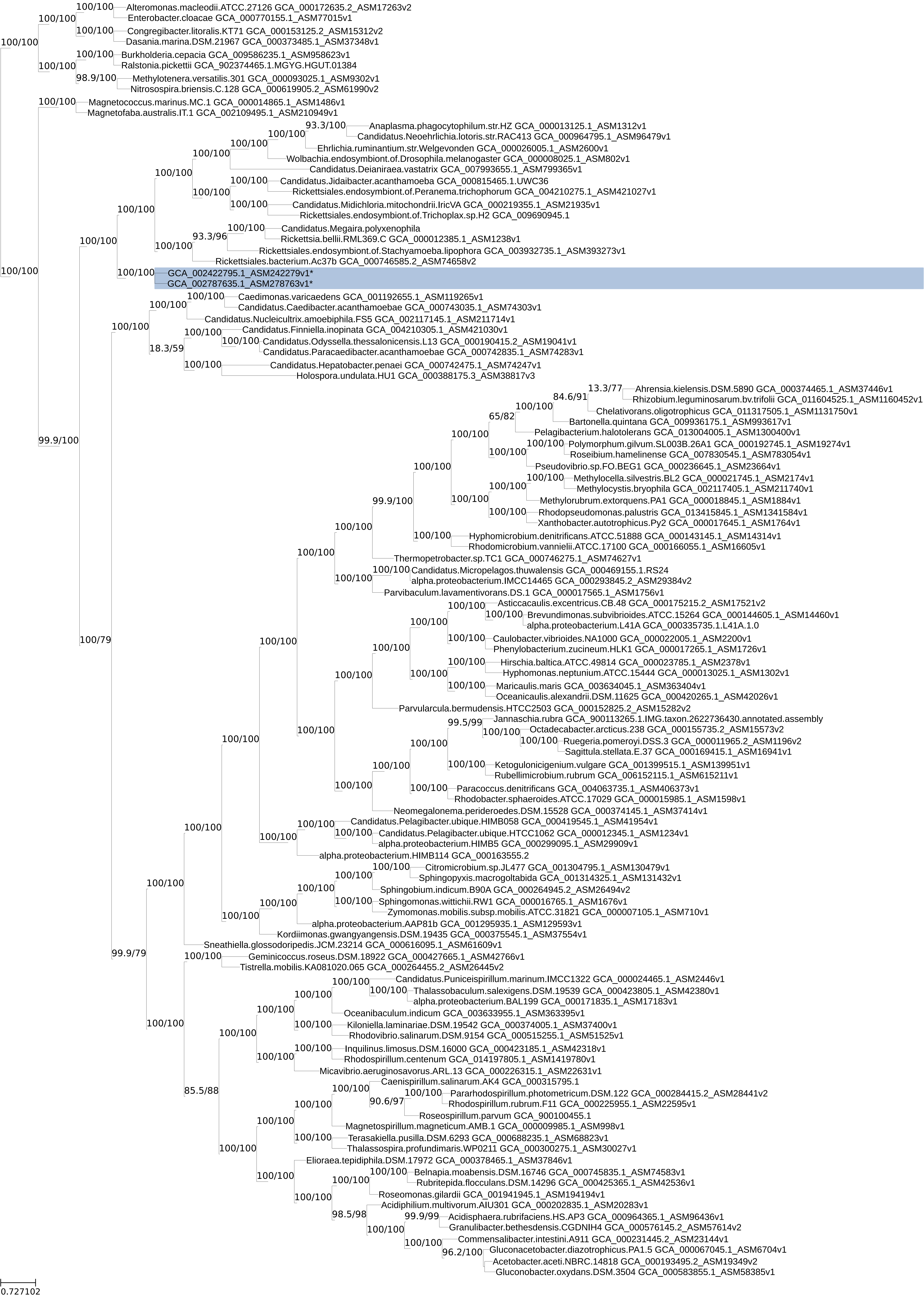
100/100
Alteromonas.macleodii.ATCC.27126 GCA_000172635.2_ASM17263v2
100/100
Enterobacter.cloacae GCA_000770155.1_ASM77015v1
100/100
100/100
Congregibacter.litoralis.KT71 GCA_000153125.2_ASM15312v2
Dasania.marina.DSM.21967 GCA_000373485.1_ASM37348v1
100/100
Burkholderia.cepacia GCA_009586235.1_ASM958623v1
100/100
Ralstonia.pickettii GCA_902374465.1.MGYG.HGUT.01384
98.9/100
Methylotenera.versatilis.301 GCA_000093025.1_ASM9302v1
Nitrosospira.briensis.C.128 GCA_000619905.2_ASM61990v2
100/100
Magnetococcus.marinus.MC.1 GCA_000014865.1_ASM1486v1
Magnetofaba.australis.IT.1 GCA_002109495.1_ASM210949v1
93.3/100
Anaplasma.phagocytophilum.str.HZ GCA_000013125.1_ASM1312v1
100/100
Candidatus.Neoehrlichia.lotoris.str.RAC413 GCA_000964795.1_ASM96479v1
100/100
Ehrlichia.ruminantium.str.Welgevonden GCA_000026005.1_ASM2600v1
100/100
Wolbachia.endosymbiont.of.Drosophila.melanogaster GCA_000008025.1_ASM802v1
100/100
Candidatus.Deianiraea.vastatrix GCA_007993655.1_ASM799365v1
100/100
Candidatus.Jidaibacter.acanthamoeba GCA_000815465.1.UWC36
100/100
Rickettsiales.endosymbiont.of.Peranema.trichophorum GCA_004210275.1_ASM421027v1
100/100
Candidatus.Midichloria.mitochondrii.IricVA GCA_000219355.1_ASM21935v1
100/100
Rickettsiales.endosymbiont.of.Trichoplax.sp.H2 GCA_009690945.1
100/100
Candidatus.Megaira.polyxenophila
93.3/96
Rickettsia.bellii.RML369.C GCA_000012385.1_ASM1238v1
100/100
Rickettsiales.endosymbiont.of.Stachyamoeba.lipophora GCA_003932735.1_ASM393273v1
100/100
Rickettsiales.bacterium.Ac37b GCA_000746585.2_ASM74658v2
100/100
100/100
GCA_002422795.1_ASM242279v1*
GCA_002787635.1_ASM278763v1*
100/100
Caedimonas.varicaedens GCA_001192655.1_ASM119265v1
100/100
Candidatus.Caedibacter.acanthamoebae GCA_000743035.1_ASM74303v1
Candidatus.Nucleicultrix.amoebiphila.FS5 GCA_002117145.1_ASM211714v1
100/100
100/100
Candidatus.Finniella.inopinata GCA_004210305.1_ASM421030v1
100/100
Candidatus.Odyssella.thessalonicensis.L13 GCA_000190415.2_ASM19041v1
18.3/59
Candidatus.Paracaedibacter.acanthamoebae GCA_000742835.1_ASM74283v1
100/100
Candidatus.Hepatobacter.penaei GCA_000742475.1_ASM74247v1
Holospora.undulata.HU1 GCA_000388175.3_ASM38817v3
13.3/77
Ahrensia.kielensis.DSM.5890 GCA_000374465.1_ASM37446v1
84.6/91
Rhizobium.leguminosarum.bv.trifolii GCA_011604525.1_ASM1160452v1
100/100
Chelativorans.oligotrophicus GCA_011317505.1_ASM1131750v1
65/82
Bartonella.quintana GCA_009936175.1_ASM993617v1
Pelagibacterium.halotolerans GCA_013004005.1_ASM1300400v1
100/100
100/100
Polymorphum.gilvum.SL003B.26A1 GCA_000192745.1_ASM19274v1
99.9/100
100/100
Roseibium.hamelinense GCA_007830545.1_ASM783054v1
Pseudovibrio.sp.FO.BEG1 GCA_000236645.1_ASM23664v1
100/100
100/100
Methylocella.silvestris.BL2 GCA_000021745.1_ASM2174v1
100/100
Methylocystis.bryophila GCA_002117405.1_ASM211740v1
100/100
Methylorubrum.extorquens.PA1 GCA_000018845.1_ASM1884v1
100/100
Rhodopseudomonas.palustris GCA_013415845.1_ASM1341584v1
Xanthobacter.autotrophicus.Py2 GCA_000017645.1_ASM1764v1
100/100
Hyphomicrobium.denitrificans.ATCC.51888 GCA_000143145.1_ASM14314v1
100/100
Rhodomicrobium.vannielii.ATCC.17100 GCA_000166055.1_ASM16605v1
Thermopetrobacter.sp.TC1 GCA_000746275.1_ASM74627v1
100/100
Candidatus.Micropelagos.thuwalensis GCA_000469155.1.RS24
100/100
alpha.proteobacterium.IMCC14465 GCA_000293845.2_ASM29384v2
Parvibaculum.lavamentivorans.DS.1 GCA_000017565.1_ASM1756v1
100/100
Asticcacaulis.excentricus.CB.48 GCA_000175215.2_ASM17521v2
100/100
100/100
Brevundimonas.subvibrioides.ATCC.15264 GCA_000144605.1_ASM14460v1
alpha.proteobacterium.L41A GCA_000335735.1.L41A.1.0
100/100
100/100
Caulobacter.vibrioides.NA1000 GCA_000022005.1_ASM2200v1
100/100
Phenylobacterium.zucineum.HLK1 GCA_000017265.1_ASM1726v1
100/100
Hirschia.baltica.ATCC.49814 GCA_000023785.1_ASM2378v1
100/100
Hyphomonas.neptunium.ATCC.15444 GCA_000013025.1_ASM1302v1
100/100
Maricaulis.maris GCA_003634045.1_ASM363404v1
Oceanicaulis.alexandrii.DSM.11625 GCA_000420265.1_ASM42026v1
100/100
Parvularcula.bermudensis.HTCC2503 GCA_000152825.2_ASM15282v2
99.5/99
Jannaschia.rubra GCA_900113265.1.IMG.taxon.2622736430.annotated.assembly
100/100
Octadecabacter.arcticus.238 GCA_000155735.2_ASM15573v2
100/100
Ruegeria.pomeroyi.DSS.3 GCA_000011965.2_ASM1196v2
100/100
Sagittula.stellata.E.37 GCA_000169415.1_ASM16941v1
100/100
100/100
Ketogulonicigenium.vulgare GCA_001399515.1_ASM139951v1
Rubellimicrobium.rubrum GCA_006152115.1_ASM615211v1
100/100
100/100
Paracoccus.denitrificans GCA_004063735.1_ASM406373v1
Rhodobacter.sphaeroides.ATCC.17029 GCA_000015985.1_ASM1598v1
Neomegalonema.perideroedes.DSM.15528 GCA_000374145.1_ASM37414v1
100/100
100/100
Candidatus.Pelagibacter.ubique.HIMB058 GCA_000419545.1_ASM41954v1
100/100
Candidatus.Pelagibacter.ubique.HTCC1062 GCA_000012345.1_ASM1234v1
100/100
alpha.proteobacterium.HIMB5 GCA_000299095.1_ASM29909v1
alpha.proteobacterium.HIMB114 GCA_000163555.2
100/100
Citromicrobium.sp.JL477 GCA_001304795.1_ASM130479v1
100/100
Sphingopyxis.macrogoltabida GCA_001314325.1_ASM131432v1
100/100
100/100
Sphingobium.indicum.B90A GCA_000264945.2_ASM26494v2
100/100
100/100
Sphingomonas.wittichii.RW1 GCA_000016765.1_ASM1676v1
Zymomonas.mobilis.subsp.mobilis.ATCC.31821 GCA_000007105.1_ASM710v1
100/100
alpha.proteobacterium.AAP81b GCA_001295935.1_ASM129593v1
Kordiimonas.gwangyangensis.DSM.19435 GCA_000375545.1_ASM37554v1
Sneathiella.glossodoripedis.JCM.23214 GCA_000616095.1_ASM61609v1
100/100
Geminicoccus.roseus.DSM.18922 GCA_000427665.1_ASM42766v1
99.9/79
Tistrella.mobilis.KA081020.065 GCA_000264455.2_ASM26445v2
100/100
Candidatus.Puniceispirillum.marinum.IMCC1322 GCA_000024465.1_ASM2446v1
100/100
Thalassobaculum.salexigens.DSM.19539 GCA_000423805.1_ASM42380v1
100/100
alpha.proteobacterium.BAL199 GCA_000171835.1_ASM17183v1
100/100
Oceanibaculum.indicum GCA_003633955.1_ASM363395v1
100/100
Kiloniella.laminariae.DSM.19542 GCA_000374005.1_ASM37400v1
100/100
100/100
Rhodovibrio.salinarum.DSM.9154 GCA_000515255.1_ASM51525v1
100/100
Inquilinus.limosus.DSM.16000 GCA_000423185.1_ASM42318v1
100/100
Rhodospirillum.centenum GCA_014197805.1_ASM1419780v1
Micavibrio.aeruginosavorus.ARL.13 GCA_000226315.1_ASM22631v1
85.5/88
Caenispirillum.salinarum.AK4 GCA_000315795.1
100/100
100/100
Pararhodospirillum.photometricum.DSM.122 GCA_000284415.2_ASM28441v2
90.6/97
Rhodospirillum.rubrum.F11 GCA_000225955.1_ASM22595v1
100/100
Roseospirillum.parvum GCA_900100455.1
100/100
Magnetospirillum.magneticum.AMB.1 GCA_000009985.1_ASM998v1
100/100
Terasakiella.pusilla.DSM.6293 GCA_000688235.1_ASM68823v1
100/100
Thalassospira.profundimaris.WP0211 GCA_000300275.1_ASM30027v1
Elioraea.tepidiphila.DSM.17972 GCA_000378465.1_ASM37846v1
100/100
100/100
Belnapia.moabensis.DSM.16746 GCA_000745835.1_ASM74583v1
100/100
Rubritepida.flocculans.DSM.14296 GCA_000425365.1_ASM42536v1
100/100
Roseomonas.gilardii GCA_001941945.1_ASM194194v1
Acidiphilium.multivorum.AIU301 GCA_000202835.1_ASM20283v1
98.5/98
99.9/99
Acidisphaera.rubrifaciens.HS.AP3 GCA_000964365.1_ASM96436v1
100/100
Granulibacter.bethesdensis.CGDNIH4 GCA_000576145.2_ASM57614v2
100/100
Commensalibacter.intestini.A911 GCA_000231445.2_ASM23144v1
96.2/100
Gluconacetobacter.diazotrophicus.PA1.5 GCA_000067045.1_ASM6704v1
Acetobacter.aceti.NBRC.14818 GCA_000193495.2_ASM19349v2
Gluconobacter.oxydans.DSM.3504 GCA_000583855.1_ASM58385v1
100/100
100/100
100/100
99.9/100
100/79
0.727102

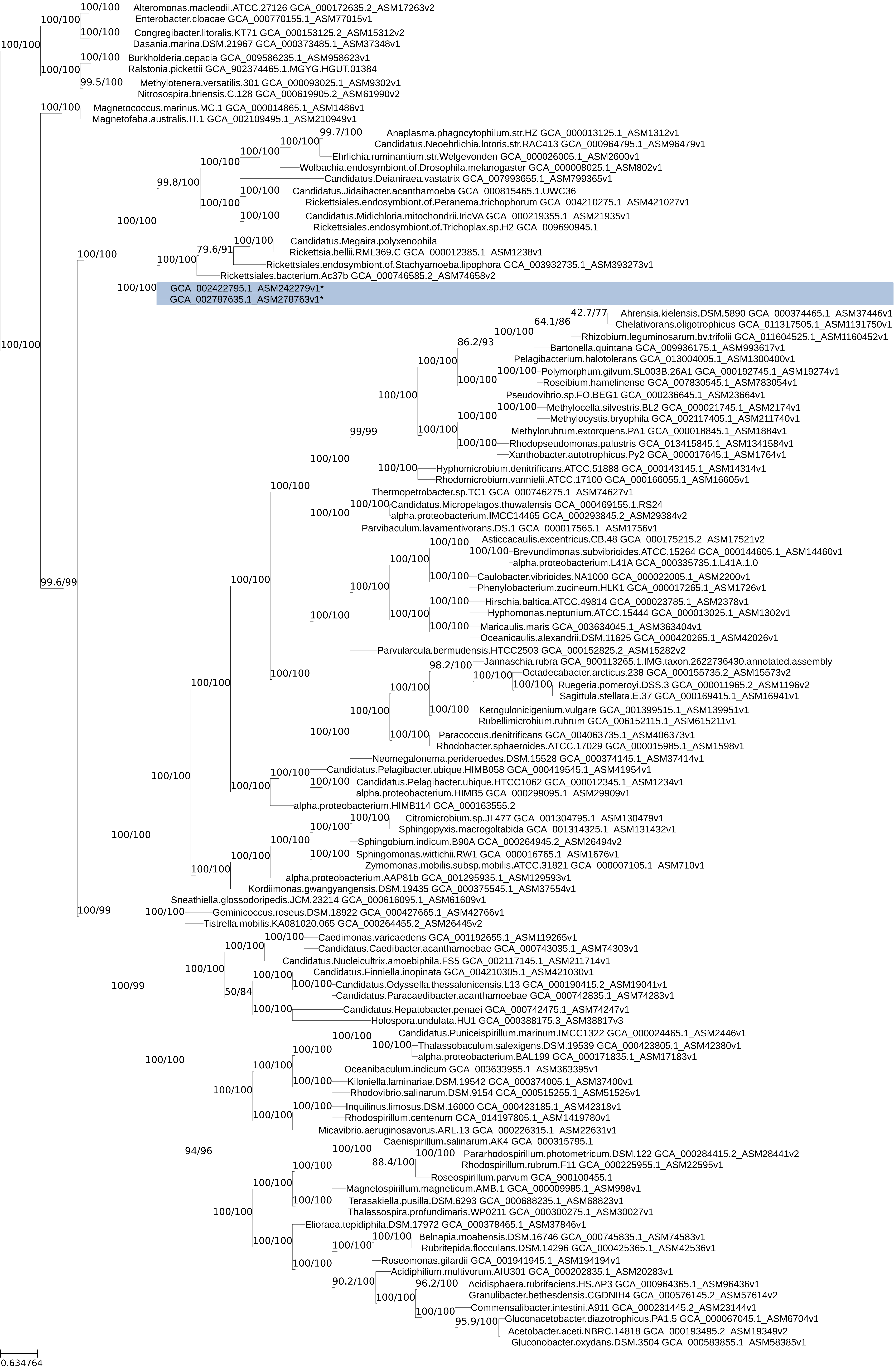

100/100
Alteromonas.macleodii.ATCC.27126 GCA_000172635.2_ASM17263v2
100/100
Enterobacter.cloacae GCA_000770155.1_ASM77015v1
100/100
Congregibacter.litoralis.KT71 GCA_000153125.2_ASM15312v2
100/100
Dasania.marina.DSM.21967 GCA_000373485.1_ASM37348v1
100/100
Burkholderia.cepacia GCA_009586235.1_ASM958623v1
100/100
Ralstonia.pickettii GCA_902374465.1.MGYG.HGUT.01384
99.5/100
Methylotenera.versatilis.301 GCA_000093025.1_ASM9302v1
Nitrosospira.briensis.C.128 GCA_000619905.2_ASM61990v2
100/100
Magnetococcus.marinus.MC.1 GCA_000014865.1_ASM1486v1
Magnetofaba.australis.IT.1 GCA_002109495.1_ASM210949v1
99.7/100
Anaplasma.phagocytophilum.str.HZ GCA_000013125.1_ASM1312v1
100/100
Candidatus.Neoehrlichia.lotoris.str.RAC413 GCA_000964795.1_ASM96479v1
100/100
Ehrlichia.ruminantium.str.Welgevonden GCA_000026005.1_ASM2600v1
100/100
Wolbachia.endosymbiont.of.Drosophila.melanogaster GCA_000008025.1_ASM802v1
99.8/100
Candidatus.Deianiraea.vastatrix GCA_007993655.1_ASM799365v1
100/100
Candidatus.Jidaibacter.acanthamoeba GCA_000815465.1.UWC36
100/100
Rickettsiales.endosymbiont.of.Peranema.trichophorum GCA_004210275.1_ASM421027v1
100/100
Candidatus.Midichloria.mitochondrii.IricVA GCA_000219355.1_ASM21935v1
100/100
Rickettsiales.endosymbiont.of.Trichoplax.sp.H2 GCA_009690945.1
100/100
Candidatus.Megaira.polyxenophila
100/100
79.6/91
Rickettsia.bellii.RML369.C GCA_000012385.1_ASM1238v1
100/100
Rickettsiales.endosymbiont.of.Stachyamoeba.lipophora GCA_003932735.1_ASM393273v1
Rickettsiales.bacterium.Ac37b GCA_000746585.2_ASM74658v2
100/100
GCA_002422795.1_ASM242279v1*
GCA_002787635.1_ASM278763v1*
42.7/77
Ahrensia.kielensis.DSM.5890 GCA_000374465.1_ASM37446v1
64.1/86
Chelativorans.oligotrophicus GCA_011317505.1_ASM1131750v1
100/100
Rhizobium.leguminosarum.bv.trifolii GCA_011604525.1_ASM1160452v1
86.2/93
Bartonella.quintana GCA_009936175.1_ASM993617v1
Pelagibacterium.halotolerans GCA_013004005.1_ASM1300400v1
100/100
100/100
Polymorphum.gilvum.SL003B.26A1 GCA_000192745.1_ASM19274v1
100/100
Roseibium.hamelinense GCA_007830545.1_ASM783054v1
Pseudovibrio.sp.FO.BEG1 GCA_000236645.1_ASM23664v1
100/100
100/100
Methylocella.silvestris.BL2 GCA_000021745.1_ASM2174v1
100/100
Methylocystis.bryophila GCA_002117405.1_ASM211740v1
100/100
Methylorubrum.extorquens.PA1 GCA_000018845.1_ASM1884v1
99/99
100/100
Rhodopseudomonas.palustris GCA_013415845.1_ASM1341584v1
Xanthobacter.autotrophicus.Py2 GCA_000017645.1_ASM1764v1
100/100
100/100
Hyphomicrobium.denitrificans.ATCC.51888 GCA_000143145.1_ASM14314v1
Rhodomicrobium.vannielii.ATCC.17100 GCA_000166055.1_ASM16605v1
100/100
Thermopetrobacter.sp.TC1 GCA_000746275.1_ASM74627v1
100/100
Candidatus.Micropelagos.thuwalensis GCA_000469155.1.RS24
100/100
alpha.proteobacterium.IMCC14465 GCA_000293845.2_ASM29384v2
Parvibaculum.lavamentivorans.DS.1 GCA_000017565.1_ASM1756v1
100/100
Asticcacaulis.excentricus.CB.48 GCA_000175215.2_ASM17521v2
100/100
100/100
Brevundimonas.subvibrioides.ATCC.15264 GCA_000144605.1_ASM14460v1
alpha.proteobacterium.L41A GCA_000335735.1.L41A.1.0
99.6/99
100/100
100/100
Caulobacter.vibrioides.NA1000 GCA_000022005.1_ASM2200v1
100/100
Phenylobacterium.zucineum.HLK1 GCA_000017265.1_ASM1726v1
100/100
Hirschia.baltica.ATCC.49814 GCA_000023785.1_ASM2378v1
100/100
Hyphomonas.neptunium.ATCC.15444 GCA_000013025.1_ASM1302v1
100/100
100/100
Maricaulis.maris GCA_003634045.1_ASM363404v1
Oceanicaulis.alexandrii.DSM.11625 GCA_000420265.1_ASM42026v1
Parvularcula.bermudensis.HTCC2503 GCA_000152825.2_ASM15282v2
98.2/100
Jannaschia.rubra GCA_900113265.1.IMG.taxon.2622736430.annotated.assembly
100/100
Octadecabacter.arcticus.238 GCA_000155735.2_ASM15573v2
100/100
100/100
Ruegeria.pomeroyi.DSS.3 GCA_000011965.2_ASM1196v2
100/100
Sagittula.stellata.E.37 GCA_000169415.1_ASM16941v1
100/100
100/100
Ketogulonicigenium.vulgare GCA_001399515.1_ASM139951v1
Rubellimicrobium.rubrum GCA_006152115.1_ASM615211v1
100/100
100/100
Paracoccus.denitrificans GCA_004063735.1_ASM406373v1
Rhodobacter.sphaeroides.ATCC.17029 GCA_000015985.1_ASM1598v1
Neomegalonema.perideroedes.DSM.15528 GCA_000374145.1_ASM37414v1
100/100
Candidatus.Pelagibacter.ubique.HIMB058 GCA_000419545.1_ASM41954v1
100/100
100/100
100/100
Candidatus.Pelagibacter.ubique.HTCC1062 GCA_000012345.1_ASM1234v1
alpha.proteobacterium.HIMB5 GCA_000299095.1_ASM29909v1
alpha.proteobacterium.HIMB114 GCA_000163555.2
100/100
Citromicrobium.sp.JL477 GCA_001304795.1_ASM130479v1
100/100
Sphingopyxis.macrogoltabida GCA_001314325.1_ASM131432v1
100/100
Sphingobium.indicum.B90A GCA_000264945.2_ASM26494v2
100/100
100/100
Sphingomonas.wittichii.RW1 GCA_000016765.1_ASM1676v1
100/100
Zymomonas.mobilis.subsp.mobilis.ATCC.31821 GCA_000007105.1_ASM710v1
alpha.proteobacterium.AAP81b GCA_001295935.1_ASM129593v1
Kordiimonas.gwangyangensis.DSM.19435 GCA_000375545.1_ASM37554v1
Sneathiella.glossodoripedis.JCM.23214 GCA_000616095.1_ASM61609v1
100/99
100/100
Geminicoccus.roseus.DSM.18922 GCA_000427665.1_ASM42766v1
Tistrella.mobilis.KA081020.065 GCA_000264455.2_ASM26445v2
100/100
Caedimonas.varicaedens GCA_001192655.1_ASM119265v1
100/100
Candidatus.Caedibacter.acanthamoebae GCA_000743035.1_ASM74303v1
100/100
Candidatus.Nucleicultrix.amoebiphila.FS5 GCA_002117145.1_ASM211714v1
100/100
Candidatus.Finniella.inopinata GCA_004210305.1_ASM421030v1
100/99
100/100
Candidatus.Odyssella.thessalonicensis.L13 GCA_000190415.2_ASM19041v1
50/84
Candidatus.Paracaedibacter.acanthamoebae GCA_000742835.1_ASM74283v1
100/100
Candidatus.Hepatobacter.penaei GCA_000742475.1_ASM74247v1
Holospora.undulata.HU1 GCA_000388175.3_ASM38817v3
100/100
Candidatus.Puniceispirillum.marinum.IMCC1322 GCA_000024465.1_ASM2446v1
100/100
100/100
Thalassobaculum.salexigens.DSM.19539 GCA_000423805.1_ASM42380v1
alpha.proteobacterium.BAL199 GCA_000171835.1_ASM17183v1
100/100
100/100
Oceanibaculum.indicum GCA_003633955.1_ASM363395v1
100/100
Kiloniella.laminariae.DSM.19542 GCA_000374005.1_ASM37400v1
100/100
Rhodovibrio.salinarum.DSM.9154 GCA_000515255.1_ASM51525v1
100/100
Inquilinus.limosus.DSM.16000 GCA_000423185.1_ASM42318v1
100/100
Rhodospirillum.centenum GCA_014197805.1_ASM1419780v1
Micavibrio.aeruginosavorus.ARL.13 GCA_000226315.1_ASM22631v1
94/96
Caenispirillum.salinarum.AK4 GCA_000315795.1
100/100
100/100
Pararhodospirillum.photometricum.DSM.122 GCA_000284415.2_ASM28441v2
88.4/100
Rhodospirillum.rubrum.F11 GCA_000225955.1_ASM22595v1
100/100
Roseospirillum.parvum GCA_900100455.1
100/100
Magnetospirillum.magneticum.AMB.1 GCA_000009985.1_ASM998v1
100/100
Terasakiella.pusilla.DSM.6293 GCA_000688235.1_ASM68823v1
100/100
Thalassospira.profundimaris.WP0211 GCA_000300275.1_ASM30027v1
Elioraea.tepidiphila.DSM.17972 GCA_000378465.1_ASM37846v1
100/100
Belnapia.moabensis.DSM.16746 GCA_000745835.1_ASM74583v1
100/100
100/100
Rubritepida.flocculans.DSM.14296 GCA_000425365.1_ASM42536v1
Roseomonas.gilardii GCA_001941945.1_ASM194194v1
100/100
Acidiphilium.multivorum.AIU301 GCA_000202835.1_ASM20283v1
90.2/100
96.2/100
Acidisphaera.rubrifaciens.HS.AP3 GCA_000964365.1_ASM96436v1
100/100
Granulibacter.bethesdensis.CGDNIH4 GCA_000576145.2_ASM57614v2
100/100
Commensalibacter.intestini.A911 GCA_000231445.2_ASM23144v1
95.9/100
Gluconacetobacter.diazotrophicus.PA1.5 GCA_000067045.1_ASM6704v1
Acetobacter.aceti.NBRC.14818 GCA_000193495.2_ASM19349v2
Gluconobacter.oxydans.DSM.3504 GCA_000583855.1_ASM58385v1
0.634764

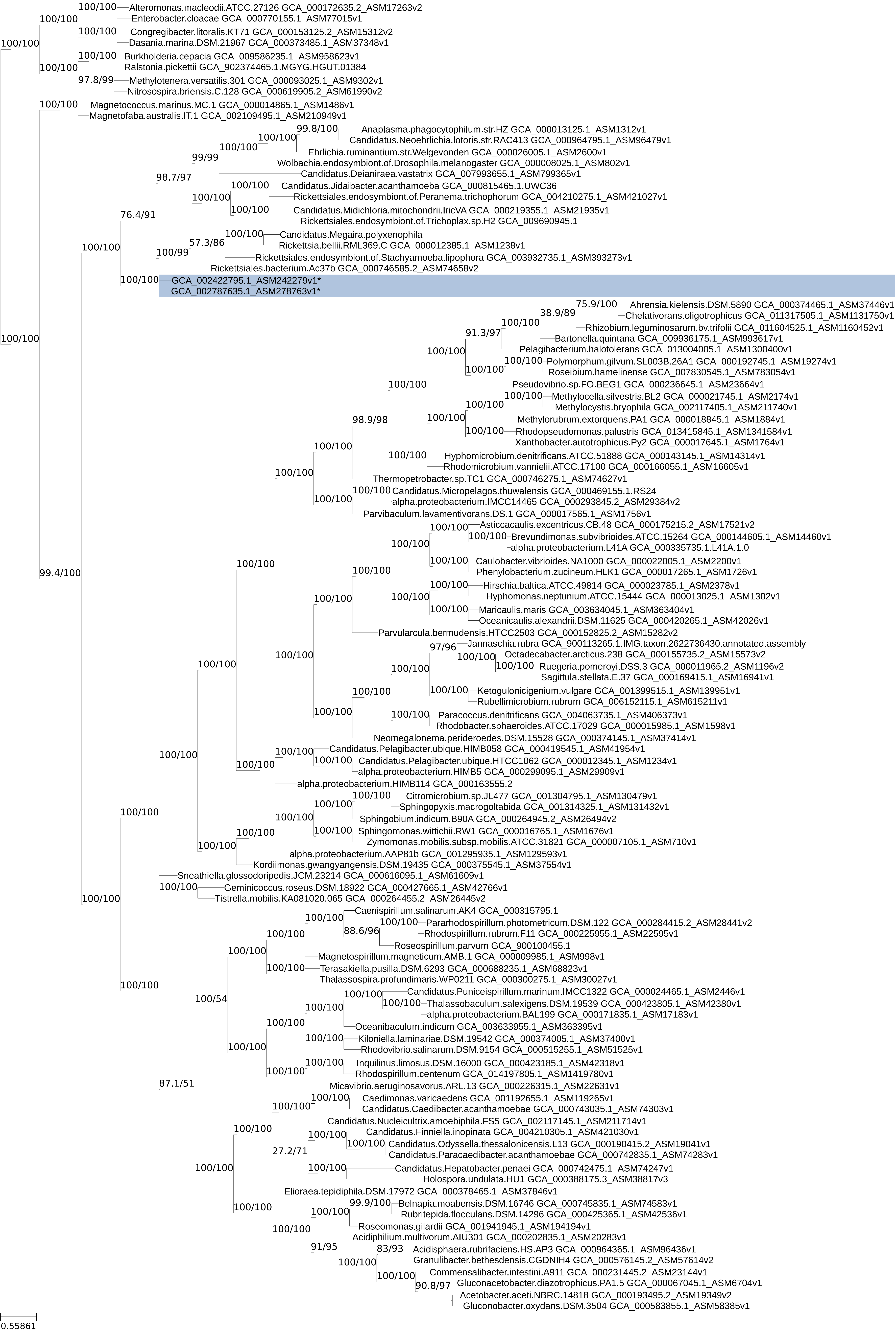

100/100
Alteromonas.macleodii.ATCC.27126 GCA_000172635.2_ASM17263v2
Enterobacter.cloacae GCA_000770155.1_ASM77015v1
100/100
100/100
Congregibacter.litoralis.KT71 GCA_000153125.2_ASM15312v2
Dasania.marina.DSM.21967 GCA_000373485.1_ASM37348v1
100/100
100/100
Burkholderia.cepacia GCA_009586235.1_ASM958623v1
Ralstonia.pickettii GCA_902374465.1.MGYG.HGUT.01384
100/100
97.8/99
Methylotenera.versatilis.301 GCA_000093025.1_ASM9302v1
Nitrosospira.briensis.C.128 GCA_000619905.2_ASM61990v2
100/100
Magnetococcus.marinus.MC.1 GCA_000014865.1_ASM1486v1
Magnetofaba.australis.IT.1 GCA_002109495.1_ASM210949v1
99.8/100
Anaplasma.phagocytophilum.str.HZ GCA_000013125.1_ASM1312v1
100/100
Candidatus.Neoehrlichia.lotoris.str.RAC413 GCA_000964795.1_ASM96479v1
100/100
Ehrlichia.ruminantium.str.Welgevonden GCA_000026005.1_ASM2600v1
99/99
Wolbachia.endosymbiont.of.Drosophila.melanogaster GCA_000008025.1_ASM802v1
Candidatus.Deianiraea.vastatrix GCA_007993655.1_ASM799365v1
98.7/97
100/100
Candidatus.Jidaibacter.acanthamoeba GCA_000815465.1.UWC36
100/100
Rickettsiales.endosymbiont.of.Peranema.trichophorum GCA_004210275.1_ASM421027v1
100/100
Candidatus.Midichloria.mitochondrii.IricVA GCA_000219355.1_ASM21935v1
Rickettsiales.endosymbiont.of.Trichoplax.sp.H2 GCA_009690945.1
76.4/91
100/100
Candidatus.Megaira.polyxenophila
57.3/86
Rickettsia.bellii.RML369.C GCA_000012385.1_ASM1238v1
100/100
Rickettsiales.endosymbiont.of.Stachyamoeba.lipophora GCA_003932735.1_ASM393273v1
100/99
Rickettsiales.bacterium.Ac37b GCA_007746585.2_ASM74658v2
100/100
GCA_002422795.1_ASM242279v1*
GCA_002787635.1_ASM278763v1*
75.9/100
Ahrensia.kielensis.DSM.5890 GCA_000374465.1_ASM37446v1
38.9/89
Chelativorans.oligotrophicus GCA_011317505.1_ASM1131750v1
100/100
Rhizobium.leguminosarum.bv.trifolii GCA_011604525.1_ASM1160452v1
91.3/97
Bartonella.quintana GCA_009936175.1_ASM993617v1
100/100
Pelagibacterium.halotolerans GCA_013004005.1_ASM1300400v1
100/100
Polymorphum.gilvum.SL003B.26A1 GCA_000192745.1_ASM19274v1
100/100
Roseibium.hamelinense GCA_007830545.1_ASM783054v1
Pseudovibrio.sp.FO.BEG1 GCA_000236645.1_ASM23664v1
100/100
100/100
Methylocella.silvestris.BL2 GCA_000021745.1_ASM2174v1
100/100
Methylocystis.bryophila GCA_002117405.1_ASM211740v1
100/100
Methylorubrum.extorquens.PA1 GCA_000018845.1_ASM1884v1
98.9/98
100/100
Rhodopseudomonas.palustris GCA_013415845.1_ASM1341584v1
Xanthobacter.autotrophicus.Py2 GCA_000017645.1_ASM1764v1
100/100
100/100
Hyphomicrobium.denitrificans.ATCC.51888 GCA_000143145.1_ASM14314v1
Rhodomicrobium.vannielii.ATCC.17100 GCA_000166055.1_ASM16605v1
100/100
Thermopetrobacter.sp.TC1 GCA_000746275.1_ASM74627v1
100/100
Candidatus.Micropelagos.thuwalensis GCA_000469155.1.RS24
100/100
alpha.proteobacterium.IMCC14465 GCA_000293845.2_ASM29384v2
Parvibaculum.lavamentivorans.DS.1 GCA_000017565.1_ASM1756v1
100/100
Asticcacaulis.excentricus.CB.48 GCA_000175215.2_ASM17521v2
100/100
100/100
Brevundimonas.subvibrioides.ATCC.15264 GCA_000144605.1_ASM14460v1
alpha.proteobacterium.L41A GCA_000335735.1.L41A.1.0
100/100
100/100
Caulobacter.vibrioides.NA1000 GCA_000022005.1_ASM2200v1
Phenylobacterium.zucineum.HLK1 GCA_000017265.1_ASM1726v1
100/100
100/100
Hirschia.baltica.ATCC.49814 GCA_000023785.1_ASM2378v1
100/100
Hyphomonas.neptunium.ATCC.15444 GCA_000013025.1_ASM1302v1
100/100
Maricaulis.maris GCA_003634045.1_ASM363404v1
100/100
Oceanicaulis.alexandrii.DSM.11625 GCA_000420265.1_ASM42026v1
Parvularcula.bermudensis.HTCC2503 GCA_000152825.2_ASM15282v2
100/100
97/96
Jannaschia.rubra GCA_900113265.1.IMG.taxon.2622736430.annotated.assembly
100/100
Octadecabacter.arcticus.238 GCA_000155735.2_ASM15573v2
100/100
Ruegeria.pomeroyi.DSS.3 GCA_000011965.2_ASM1196v2
Sagittula.stellata.E.37 GCA_000169415.1_ASM16941v1
100/100
100/100
Ketogulonicigenium.vulgare GCA_001399515.1_ASM139951v1
Rubellimicrobium.rubrum GCA_006152115.1_ASM615211v1
100/100
100/100
Paracoccus.denitrificans GCA_004063735.1_ASM406373v1
Rhodobacter.sphaeroides.ATCC.17029 GCA_000015985.1_ASM1598v1
Neomegalonema.perideroedes.DSM.15528 GCA_000374145.1_ASM37414v1
100/100
100/100
Candidatus.Pelagibacter.ubique.HIMB058 GCA_000419545.1_ASM41954v1
100/100
Candidatus.Pelagibacter.ubique.HTCC1062 GCA_000012345.1_ASM1234v1
100/100
alpha.proteobacterium.HIMB5 GCA_000299095.1_ASM29909v1
alpha.proteobacterium.HIMB114 GCA_000163555.2
100/100
Citromicrobium.sp.JL477 GCA_001304795.1_ASM130479v1
100/100
Sphingopyxis.macrogoltabida GCA_001314325.1_ASM131432v1
100/100
Sphingobium.indicum.B90A GCA_000264945.2_ASM26494v2
100/100
100/100
Sphingomonas.wittichii.RW1 GCA_000016765.1_ASM1676v1
100/100
Zymomonas.mobilis.subsp.mobilis.ATCC.31821 GCA_000007105.1_ASM710v1
100/100
alpha.proteobacterium.AAP81b GCA_001295935.1_ASM129593v1
Kordiimonas.gwangyangensis.DSM.19435 GCA_000375545.1_ASM37554v1
Sneathiella.glossodoripedis.JCM.23214 GCA_000616095.1_ASM61609v1
100/100
Geminicoccus.roseus.DSM.18922 GCA_000427665.1_ASM42766v1
Tistrella.mobilis.KA081020.065 GCA_000264455.2_ASM26445v2
100/100
Caenispirillum.salinarum.AK4 GCA_000315795.1
100/100
100/100
Pararhodospirillum.photometricum.DSM.122 GCA_000284415.2_ASM28441v2
88.6/96
Rhodospirillum.rubrum.F11 GCA_000225955.1_ASM22595v1
100/100
Roseospirillum.parvum GCA_900100455.1
100/100
Magnetospirillum.magneticum.AMB.1 GCA_000009985.1_ASM998v1
100/100
Terasakiella.pusilla.DSM.6293 GCA_000688235.1_ASM68823v1
Thalassospira.profundimaris.WP0211 GCA_000300275.1_ASM30027v1
100/100
100/54
Candidatus.Puniceispirillum.marinum.IMCC1322 GCA_000024465.1_ASM2446v1
100/100
100/100
Thalassobaculum.salexigens.DSM.19539 GCA_000423805.1_ASM42380v1
alpha.proteobacterium.BAL199 GCA_000171835.1_ASM17183v1
100/100
Oceanibaculum.indicum GCA_003633955.1_ASM363395v1
100/100
Kiloniella.laminariae.DSM.19542 GCA_000374005.1_ASM37400v1
100/100
Rhodovibrio.salinarum.DSM.9154 GCA_000515255.1_ASM51525v1
100/100
Inquilinus.limosus.DSM.16000 GCA_000423185.1_ASM42318v1
100/100
Rhodospirillum.centenum GCA_014197805.1_ASM1419780v1
87.1/51
Micavibrio.aeruginosavorus.ARL.13 GCA_000226315.1_ASM22631v1
100/100
Caedimonas.varicaedens GCA_001192655.1_ASM119265v1
100/100
Candidatus.Caedibacter.acanthamoebae GCA_000743035.1_ASM74303v1
Candidatus.Nucleicultrix.amoebiphila.FS5 GCA_002117145.1_ASM211714v1
100/100
100/100
Candidatus.Finniella.inopinata GCA_004210305.1_ASM421030v1
100/100
Candidatus.Odyssella.thessalonicensis.L13 GCA_000190415.2_ASM19041v1
27.2/71
Candidatus.Paracaedibacter.acanthamoebae GCA_000742835.1_ASM74283v1
100/100
100/100
Candidatus.Hepatobacter.penaei GCA_000742475.1_ASM74247v1
Holospora.undulata.HU1 GCA_000388175.3_ASM38817v3
Elioraea.tepidiphila.DSM.17972 GCA_000378465.1_ASM37846v1
99.9/100
100/100
Belnapia.moabensis.DSM.16746 GCA_000745835.1_ASM74583v1
100/100
Rubritepida.flocculans.DSM.14296 GCA_000425365.1_ASM42536v1
100/100
Roseomonas.gilardii GCA_001941945.1_ASM194194v1
Acidiphilium.multivorum.AIU301 GCA_000202835.1_ASM20283v1
91/95
83/93
Acidisphaera.rubrifaciens.HS.AP3 GCA_000964365.1_ASM96436v1
100/100
Granulibacter.bethesdensis.CGDNIH4 GCA_000576145.2_ASM57614v2
100/100
Commensalibacter.intestini.A911 GCA_000231445.2_ASM23144v1
90.8/97
Gluconacetobacter.diazotrophicus.PA1.5 GCA_000067045.1_ASM6704v1
Acetobacter.aceti.NBRC.14818 GCA_000193495.2_ASM19349v2
Gluconobacter.oxydans.DSM.3504 GCA_000583855.1_ASM58385v1
100/100
100/100
99.4/100
0.55861

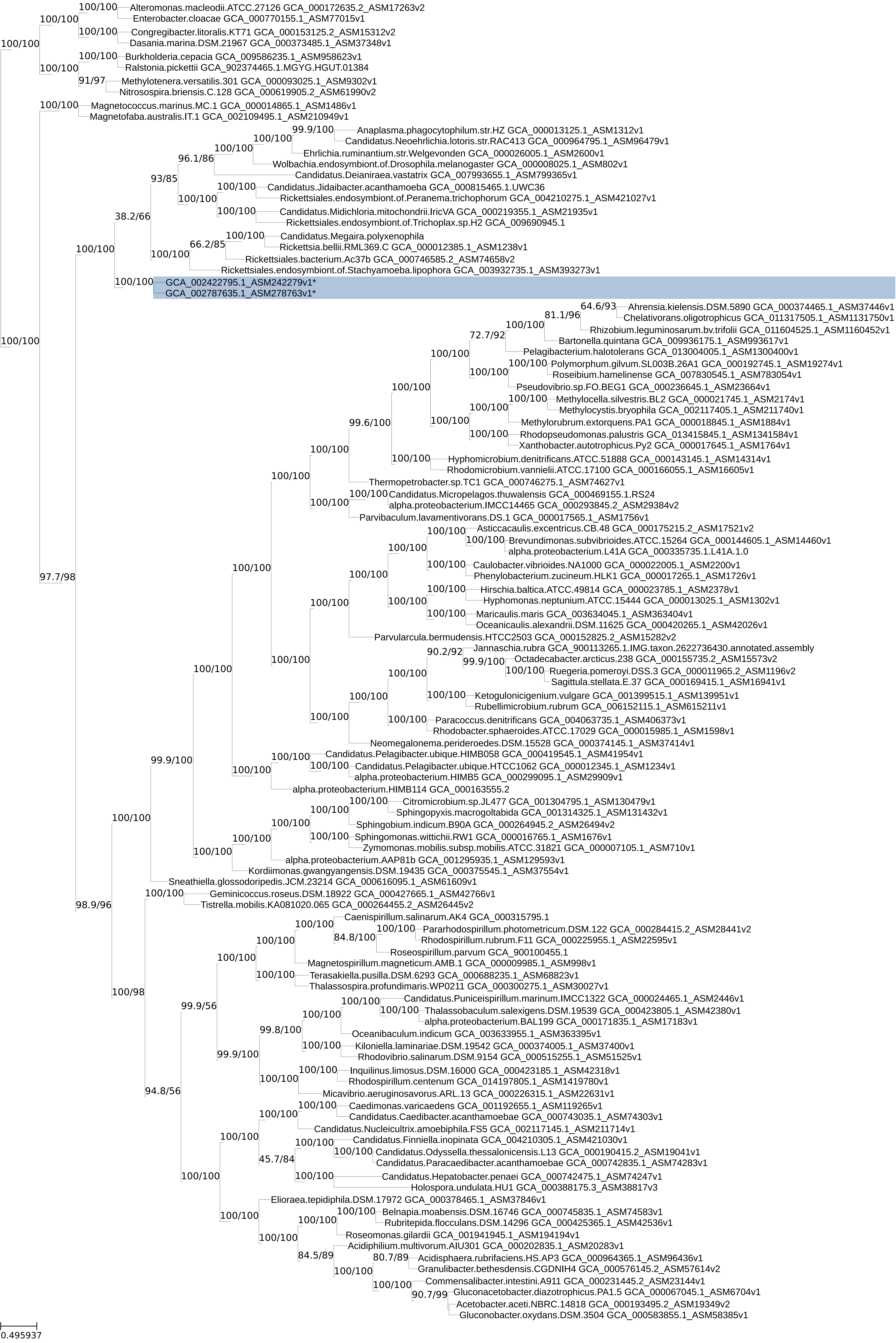

Alteromonas.macleodii.ATCC.27126 GCA_000172635.2_ASM17263v2
Enterobacter.cloacae GCA_000770155.1_ASM77015v1
Congregibacter.litoralis.KT71 GCA_000153125.2_ASM15312v2
Dasania.marina.DSM.21967 GCA_000373485.1_ASM37348v1
Burkholderia.cepacia GCA_009586235.1_ASM958623v1
Ralstonia.pickettii GCA_902374465.1.MGYG.HGUT.01384
Methylotenera.versatilis.301 GCA_000093025.1_ASM9302v1
Nitrosospira.briensis.C.128 GCA_000619905.2_ASM61990v2
Magnetococcus.marinus.MC.1 GCA_000014865.1_ASM1486v1
Magnetofaba.australis.IT.1 GCA_002109495.1_ASM210949v1
Anaplasma.phagocytophilum.str.HZ GCA_000013125.1_ASM1312v1
Candidatus.Neoehrlichia.lotoris.str.RAC413 GCA_000964795.1_ASM96479v1
Ehrlichia.ruminantium.str.Welgevonden GCA_000026005.1_ASM2600v1
Wolbachia.endosymbiont.of.Drosophila.melanogaster GCA_000008025.1_ASM802v1
Candidatus.Deianiraea.vastatrix GCA_007993655.1_ASM799365v1
Candidatus.Jidaibacter.acanthamoeba GCA_000815465.1.UWC36
Rickettsiales.endosymbiont.of.Peranema.trichophorum GCA_004210275.1_ASM421027v1
Candidatus.Midichloria.mitochondrii.IricVA GCA_000219355.1_ASM21935v1
Rickettsiales.endosymbiont.of.Trichoplax.sp.H2 GCA_009690945.1
Candidatus.Megaira.polyxenophila
Rickettsia.bellii.RML369.C GCA_000012385.1_ASM1238v1
Rickettsiales.bacterium.Ac37b GCA_000746585.2_ASM74658v2
Rickettsiales.endosymbiont.of.Stachyamoeba.lipophora GCA_003932735.1_ASM393273v1
GCA_002422795.1_ASM242279v1*
GCA_002787635.1_ASM278763v1*
Ahrensia.kielensis.DSM.5890 GCA_000374465.1_ASM37446v1
Chelativorans.oligotrophicus GCA_011317505.1_ASM1131750v1
Rhizobium.leguminosarum.bv.trifolii GCA_011604525.1_ASM1160452v1
Bartonella.quintana GCA_009936175.1_ASM993617v1
Pelagibacterium.halotolerans GCA_013004005.1_ASM1300400v1
Polymorphum.gilvum.SL003B.26A1 GCA_000192745.1_ASM19274v1
Roseibium.hamelinense GCA_007830545.1_ASM783054v1
Pseudovibrio.sp.FO.BEG1 GCA_000236645.1_ASM23664v1
Methylocella.silvestris.BL2 GCA_000021745.1_ASM2174v1
Methylocystis.bryophila GCA_002117405.1_ASM211740v1
Methylorubrum.extorquens.PA1 GCA_000018845.1_ASM1884v1
Rhodopseudomonas.palustris GCA_013415845.1_ASM1341584v1
Xanthobacter.autotrophicus.Py2 GCA_000017645.1_ASM1764v1
Hyphomicrobium.denitrificans.ATCC.51888 GCA_000143145.1_ASM14314v1
Rhodomicrobium.vannielii.ATCC.17100 GCA_000166055.1_ASM16605v1
Thermopetrobacter.sp.TC1 GCA_007746275.1_ASM74627v1
Candidatus.Micropelagos.thuwalensis GCA_000469155.1.RS24
alpha.proteobacterium.IMCC14465 GCA_000293845.2_ASM29384v2
Parvibaculum.lavamentivorans.DS.1 GCA_000017565.1_ASM1756v1
Asticcacaulis.excentricus.CB.48 GCA_000175215.2_ASM17521v2
Brevundimonas.subvibrioides.ATCC.15264 GCA_000144605.1_ASM14460v1
alpha.proteobacterium.L41A GCA_000335735.1.L41A.1.0
Caulobacter.vibrioides.NA1000 GCA_000022005.1_ASM2200v1
Phenylobacterium.zucineum.HLK1 GCA_000017265.1_ASM1726v1
Hirschia.baltica.ATCC.49814 GCA_000023785.1_ASM2378v1
Hyphomonas.neptunium.ATCC.15444 GCA_000013025.1_ASM1302v1
Maricaulis.maris GCA_003634045.1_ASM363404v1
Oceanicaulis.alexandrii.DSM.11625 GCA_000420265.1_ASM42026v1
Parvularcula.bermudensis.HTCC2503 GCA_000152825.2_ASM15282v2
Jannaschia.rubra GCA_900113265.1.IMG.taxon.2622736430.annotated.assembly
Octadecabacter.arcticus.238 GCA_000155735.2_ASM15573v2
Ruegeria.pomeroyi.DSS.3 GCA_000011965.2_ASM1196v2
Sagittula.stellata.E.37 GCA_000169415.1_ASM16941v1
Ketogulonicigenium.vulgare GCA_001399515.1_ASM139951v1
Rubellimicrobium.rubrum GCA_006152115.1_ASM615211v1
Paracoccus.denitrificans GCA_004063735.1_ASM406373v1
Rhodobacter.sphaeroides.ATCC.17029 GCA_000015985.1_ASM1598v1
Neomegalonema.perideroedes.DSM.15528 GCA_000374145.1_ASM37414v1
Candidatus.Pelagibacter.ubique.HIMB058 GCA_000419545.1_ASM41954v1
Candidatus.Pelagibacter.ubique.HTCC1062 GCA_000012345.1_ASM1234v1
alpha.proteobacterium.HIMB5 GCA_000299095.1_ASM29909v1
alpha.proteobacterium.HIMB114 GCA_000163555.2
Citromicrobium.sp.JL477 GCA_001304795.1_ASM130479v1
Sphingopyxis.macrogoltabida GCA_001314325.1_ASM131432v1
Sphingobium.indicum.B90A GCA_000264945.2_ASM26494v2
Sphingomonas.wittichii.RW1 GCA_000016765.1_ASM1676v1
Zymomonas.mobilis.subsp.mobilis.ATCC.31821 GCA_000007105.1_ASM710v1
alpha.proteobacterium.AAP81b GCA_001295935.1_ASM129593v1
Kordiimonas.gwangyangensis.DSM.19435 GCA_000375545.1_ASM37554v1
Sneathiella.glossodoripedis.JCM.23214 GCA_000616095.1_ASM61609v1
Geminicoccus.roseus.DSM.18922 GCA_000427665.1_ASM42766v1
Tistrella.mobilis.KA081020.065 GCA_000264455.2_ASM26445v2
Caenispirillum.salinarum.AK4 GCA_000315795.1
Pararhodospirillum.photometricum.DSM.122 GCA_000284415.2_ASM28441v2
Rhodospirillum.rubrum.F11 GCA_000225955.1_ASM22595v1
Roseospirillum.parvum GCA_900100455.1
Magnetospirillum.magneticum.AMB.1 GCA_000009985.1_ASM998v1
Terasakiella.pusilla.DSM.6293 GCA_000688235.1_ASM68823v1
Thalassospira.profundimaris.WP0211 GCA_000300275.1_ASM30027v1
Candidatus.Puniceispirillum.marinum.IMCC1322 GCA_000024465.1_ASM2446v1
Thalassobaculum.salexigens.DSM.19539 GCA_000423805.1_ASM42380v1
alpha.proteobacterium.BAL199 GCA_000171835.1_ASM17183v1
Oceanibaculum.indicum GCA_003633955.1_ASM363395v1
Kiloniella.laminariae.DSM.19542 GCA_000374005.1_ASM37400v1
Rhodovibrio.salinarum.DSM.9154 GCA_000515255.1_ASM51525v1
Inquilinus.limosus.DSM.16000 GCA_000423185.1_ASM42318v1
Rhodospirillum.centenum GCA_014197805.1_ASM1419780v1
Micavibrio.aeruginosavorus.ARL.13 GCA_000226315.1_ASM22631v1
Caedimonas.varicaedens GCA_001192655.1_ASM119265v1
Candidatus.Caedibacter.acanthamoebae GCA_000743035.1_ASM74303v1
Candidatus.Nucleicultrix.amoebiphila.FS5 GCA_002117145.1_ASM211714v1
Candidatus.Finniella.inopinata GCA_004210305.1_ASM421030v1
Candidatus.Odyssella.thessalonicensis.L13 GCA_000190415.2_ASM19041v1
Candidatus.Paracaedibacter.acanthamoebae GCA_000742835.1_ASM74283v1
Candidatus.Hepatobacter.penaei GCA_000742475.1_ASM74247v1
Holospora.undulata.HU1 GCA_000388175.3_ASM38817v3
Elioraea.tepidiphila.DSM.17972 GCA_000378465.1_ASM37846v1
Belnapia.moabensis.DSM.16746 GCA_000745835.1_ASM74583v1
Rubritepida.flocculans.DSM.14296 GCA_000425365.1_ASM42536v1
Roseomonas.gilardii GCA_001941945.1_ASM194194v1
Acidiphilium.multivorum.AIU301 GCA_000202835.1_ASM20283v1
Acidisphaera.rubrifaciens.HS.AP3 GCA_000964365.1_ASM96436v1
Granulibacter.bethesdensis.CGDNIH4 GCA_000576145.2_ASM57614v2
Commensalibacter.intestini.A911 GCA_000231445.2_ASM23144v1
Gluconacetobacter.diazotrophicus.PA1.5 GCA_000067045.1_ASM6704v1
Acetobacter.aceti.NBRC.14818 GCA_000193495.2_ASM19349v2
Gluconobacter.oxydans.DSM.3504 GCA_000583855.1_ASM58385v1
0.495937

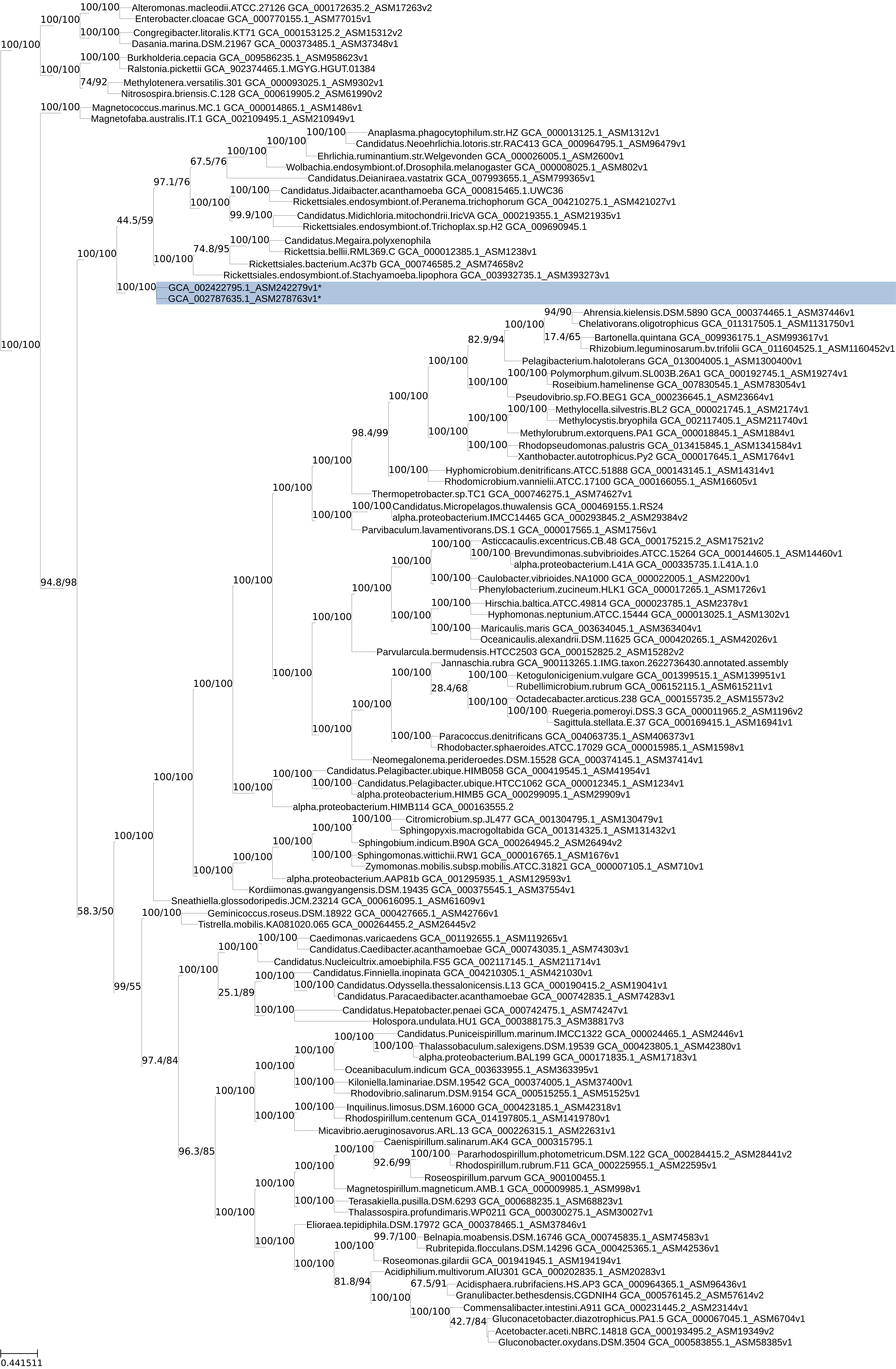

Alteromonas.macleodii.ATCC.27126 GCA_000172635.2_ASM17263v2
Enterobacter.cloacae GCA_000770155.1_ASM77015v1
Congregibacter.litoralis.KT71 GCA_000153125.2_ASM15312v2
Dasania.marina.DSM.21967 GCA_000373485.1_ASM37348v1
Burkholderia.cepacia GCA_009586235.1_ASM958623v1
Ralstonia.pickettii GCA_902374465.1.MGYG.HGUT.01384
Methylotenera.versatilis.301 GCA_000093025.1_ASM9302v1
Nitrosospira.briensis.C.128 GCA_000619905.2_ASM61990v2
Magnetococcus.marinus.MC.1 GCA_000014865.1_ASM1486v1
Magnetofaba.australis.IT.1 GCA_002109495.1_ASM210949v1
Anaplasma.phagocytophilum.str.HZ GCA_000013125.1_ASM1312v1
Candidatus.Neoehrlichia.lotoris.str.RAC413 GCA_000964795.1_ASM96479v1
Ehrlichia.ruminantium.str.Welgevonden GCA_000026005.1_ASM2600v1
Wolbachia.endosymbiont.of.Drosophila.melanogaster GCA_000008025.1_ASM802v1
Candidatus.Deianiraea.vastatrix GCA_007993655.1_ASM799365v1
Candidatus.Jidaibacter.acanthamoeba GCA_000815465.1.UWC36
Rickettsiales.endosymbiont.of.Peranema.trichophorum GCA_004210275.1_ASM421027v1
Candidatus.Midichloria.mitochondrii.IricVA GCA_000219355.1_ASM21935v1
Rickettsiales.endosymbiont.of.Trichoplax.sp.H2 GCA_009690945.1
Candidatus.Megaira.polyxenophila
Rickettsia.bellii.RML369.C GCA_000012385.1_ASM1238v1
Rickettsiales.bacterium.Ac37b GCA_000746585.2_ASM74658v2
Rickettsiales.endosymbiont.of.Stachyamoeba.lipophora GCA_003932735.1_ASM393273v1
GCA_002422795.1_ASM242279v1*
GCA_002787635.1_ASM278763v1*
Ahrensia.kielensis.DSM.5890 GCA_000374465.1_ASM37446v1
Chelativorans.oligotrophicus GCA_011317505.1_ASM1131750v1
Bartonella.quintana GCA_009936175.1_ASM993617v1
Rhizobium.leguminosarum.bv.trifolii GCA_011604525.1_ASM1160452v1
Pelagibacterium.halotolerans GCA_013004005.1_ASM1300400v1
Polymorphum.gilvum.SL003B.26A1 GCA_000192745.1_ASM19274v1
Roseibium.hamelinense GCA_007830545.1_ASM783054v1
Pseudovibrio.sp.FO.BEG1 GCA_000236645.1_ASM23664v1
Methylocella.silvestris.BL2 GCA_000021745.1_ASM2174v1
Methylocystis.bryophila GCA_002117405.1_ASM211740v1
Methylorubrum.extorquens.PA1 GCA_000018845.1_ASM1884v1
Rhodopseudomonas.palustris GCA_013415845.1_ASM1341584v1
Xanthobacter.autotrophicus.Py2 GCA_000017645.1_ASM1764v1
Hyphomicrobium.denitrificans.ATCC.51888 GCA_000143145.1_ASM14314v1
Rhodomicrobium.vannielii.ATCC.17100 GCA_000166055.1_ASM16605v1
Thermopetrobacter.sp.TC1 GCA_000746275.1_ASM74627v1
Candidatus.Micropelagos.thuwalensis GCA_000469155.1.RS24
alpha.proteobacterium.IMCC14465 GCA_000293845.2_ASM29384v2
Parvibaculum.lavamentivorans.DS.1 GCA_000017565.1_ASM1756v1
Asticcacaulis.excentricus.CB.48 GCA_000175215.2_ASM17521v2
Brevundimonas.subvibrioides.ATCC.15264 GCA_000144605.1_ASM14460v1
alpha.proteobacterium.L41A GCA_000335735.1.L41A.1.0
Caulobacter.vibrioides.NA1000 GCA_000022005.1_ASM2200v1
Phenylobacterium.zucineum.HLK1 GCA_000017265.1_ASM1726v1
Hirschia.baltica.ATCC.49814 GCA_000023785.1_ASM2378v1
Hyphomonas.neptunium.ATCC.15444 GCA_000013025.1_ASM1302v1
Maricaulis.maris GCA_003634045.1_ASM363404v1
Oceanicaulis.alexandrii.DSM.11625 GCA_000420265.1_ASM42026v1
Parvularcula.bermudensis.HTCC2503 GCA_000152825.2_ASM15282v2
Jannaschia.rubra GCA_900113265.1.IMG.taxon.2622736430.annotated.assembly
Ketogulonicigenium.vulgare GCA_001399515.1_ASM139951v1
Rubellimicrobium.rubrum GCA_006152115.1_ASM615211v1
Octadecabacter.arcticus.238 GCA_000155735.2_ASM15573v2
Ruegeria.pomeroyi.DSS.3 GCA_000011965.2_ASM1196v2
Sagittula.stellata.E.37 GCA_000169415.1_ASM16941v1
Paracoccus.denitrificans GCA_004063735.1_ASM406373v1
Rhodobacter.sphaeroides.ATCC.17029 GCA_000015985.1_ASM1598v1
Neomegalonema.perideroedes.DSM.15528 GCA_000374145.1_ASM37414v1
Candidatus.Pelagibacter.ubique.HIMB058 GCA_000419545.1_ASM41954v1
Candidatus.Pelagibacter.ubique.HTCC1062 GCA_000012345.1_ASM1234v1
alpha.proteobacterium.HIMB5 GCA_000299095.1_ASM29909v1
alpha.proteobacterium.HIMB114 GCA_000163555.2
Citromicrobium.sp.JL477 GCA_001304795.1_ASM130479v1
Sphingopyxis.macrogoltabida GCA_001314325.1_ASM131432v1
Sphingobium.indicum.B90A GCA_000264945.2_ASM26494v2
Sphingomonas.wittichii.RW1 GCA_000016765.1_ASM1676v1
Zymomonas.mobilis.subsp.mobilis.ATCC.31821 GCA_000007105.1_ASM710v1
alpha.proteobacterium.AAP81b GCA_001295935.1_ASM129593v1
Kordiimonas.gwangyangensis.DSM.19435 GCA_000375545.1_ASM37554v1
Sneathiella.glossodoripedis.JCM.23214 GCA_000616095.1_ASM61609v1
Geminicoccus.roseus.DSM.18922 GCA_000427665.1_ASM42766v1
Tistrella.mobilis.KA081020.065 GCA_000264455.2_ASM26445v2
Caedimonas.varicaedens GCA_001192655.1_ASM119265v1
Candidatus.Caedibacter.acanthamoebae GCA_000743035.1_ASM74303v1
Candidatus.Nucleicultrix.amoebiphila.FS5 GCA_002117145.1_ASM211714v1
Candidatus.Finniella.inopinata GCA_004210305.1_ASM421030v1
Candidatus.Odyssella.thessalonicensis.L13 GCA_000190415.2_ASM19041v1
Candidatus.Paracaedibacter.acanthamoebae GCA_000742835.1_ASM74283v1
Candidatus.Hepatobacter.penaei GCA_000742475.1_ASM74247v1
Holospora.undulata.HU1 GCA_000388175.3_ASM38817v3
Candidatus.Puniceispirillum.marinum.IMCC1322 GCA_000024465.1_ASM2446v1
Thalassobaculum.salexigens.DSM.19539 GCA_000423805.1_ASM42380v1
alpha.proteobacterium.BAL199 GCA_000171835.1_ASM17183v1
Oceanibaculum.indicum GCA_003633955.1_ASM363395v1
Kiloniella.laminariae.DSM.19542 GCA_000374005.1_ASM37400v1
Rhodovibrio.salinarum.DSM.9154 GCA_000515255.1_ASM51525v1
Inquilinus.limosus.DSM.16000 GCA_000423185.1_ASM42318v1
Rhodospirillum.centenum GCA_014197805.1_ASM1419780v1
Micavibrio.aeruginosavorus.ARL.13 GCA_000226315.1_ASM22631v1
Caenispirillum.salinarum.AK4 GCA_000315795.1
Pararhodospirillum.photometricum.DSM.122 GCA_000284415.2_ASM28441v2
Rhodospirillum.rubrum.F11 GCA_000225955.1_ASM22595v1
Roseospirillum.parvum GCA_900100455.1
Magnetospirillum.magneticum.AMB.1 GCA_000009985.1_ASM998v1
Terasakiella.pusilla.DSM.6293 GCA_000688235.1_ASM68823v1
Thalassospira.profundimaris.WP0211 GCA_000300275.1_ASM30027v1
Elioraea.tepidiphila.DSM.17972 GCA_000378465.1_ASM37846v1
Belnapia.moabensis.DSM.16746 GCA_000745835.1_ASM74583v1
Rubritepida.flocculans.DSM.14296 GCA_000425365.1_ASM42536v1
Roseomonas.gilardii GCA_001941945.1_ASM194194v1
Acidiphilium.multivorum.AIU301 GCA_000202835.1_ASM20283v1
Acidisphaera.rubrifaciens.HS.AP3 GCA_000964365.1_ASM96436v1
Granulibacter.bethesdensis.CGDNIH4 GCA_000576145.2_ASM57614v2
Commensalibacter.intestini.A911 GCA_000231445.2_ASM23144v1
Gluconacetobacter.diazotrophicus.PA1.5 GCA_000067045.1_ASM6704v1
Acetobacter.aceti.NBRC.14818 GCA_000193495.2_ASM19349v2
Gluconobacter.oxydans.DSM.3504 GCA_000583855.1_ASM58385v1
0.441511

### Group G

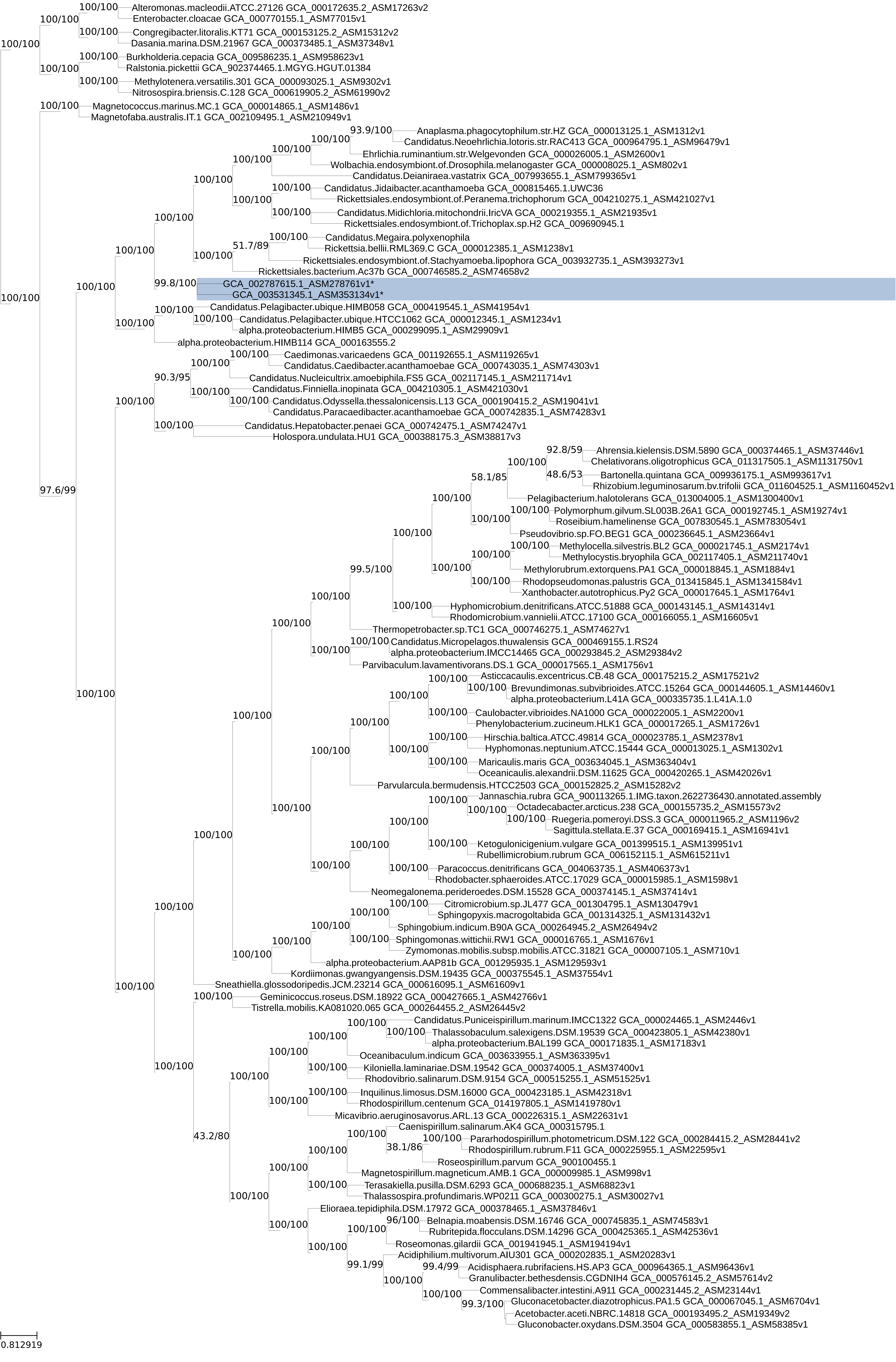
100/100
Alteromonas.macleodii.ATCC.27126 GCA_000172635.2_ASM17263v2
100/100
Enterobacter.cloacae GCA_000770155.1_ASM77015v1
100/100
100/100
Congregibacter.litoralis.KT71 GCA_000153125.2_ASM15312v2
Dasania.marina.DSM.21967 GCA_000373485.1_ASM37348v1
100/100
Burkholderia.cepacia GCA_009586235.1_ASM958623v1
100/100
Ralstonia.pickettii GCA_902374465.1.MGYG.HGUT.01384
100/100
Methylotenera.versatilis.301 GCA_000093025.1_ASM9302v1
Nitrosospira.briensis.C.128 GCA_000619905.2_ASM61990v2
100/100
Magnetococcus.marinus.MC.1 GCA_000014865.1_ASM1486v1
Magnetofaba.australis.IT.1 GCA_002109495.1_ASM210949v1
93.9/100
Anaplasma.phagocytophilum.str.HZ GCA_000013125.1_ASM1312v1
100/100
Candidatus.Neoehrlichia.lotoris.str.RAC413 GCA_000964795.1_ASM96479v1
100/100
Ehrlichia.ruminantium.str.Welgevonden GCA_000026005.1_ASM2600v1
100/100
Wolbachia.endosymbiont.of.Drosophila.melanogaster GCA_000008025.1_ASM802v1
Candidatus.Deianiraea.vastatrix GCA_007993655.1_ASM799365v1
100/100
100/100
Candidatus.Jidaibacter.acanthamoeba GCA_000815465.1.UWC36
100/100
Rickettsiales.endosymbiont.of.Peranema.trichophorum GCA_004210275.1_ASM421027v1
100/100
Candidatus.Midichloria.mitochondrii.IricVA GCA_000219355.1_ASM21935v1
100/100
Rickettsiales.endosymbiont.of.Trichoplax.sp.H2 GCA_009690945.1
100/100
Candidatus.Megaira.polyxenophila
51.7/89
100/100
Rickettsia.bellii.RML369.C GCA_000012385.1_ASM1238v1
100/100
Rickettsiales.endosymbiont.of.Stachyamoeba.lipophora GCA_003932735.1_ASM393273v1
Rickettsiales.bacterium.Ac37b GCA_000746585.2_ASM74658v2
99.8/100
GCA_002787615.1_ASM278761v1*
100/100
GCA_003531345.1_ASM353134v1*
100/100
100/100
Candidatus.Pelagibacter.ubique.HIMB058 GCA_000419545.1_ASM41954v1
100/100
Candidatus.Pelagibacter.ubique.HTCC1062 GCA_000012345.1_ASM1234v1
100/100
alpha.proteobacterium.HIMB5 GCA_000299095.1_ASM29909v1
alpha.proteobacterium.HIMB114 GCA_000163555.2
100/100
Caedimonas.varicaedens GCA_001192655.1_ASM119265v1
100/100
Candidatus.Caedibacter.acanthamoebae GCA_000743035.1_ASM74303v1
90.3/95
Candidatus.Nucleicultrix.amoebiphila.FS5 GCA_002117145.1_ASM211714v1
100/100
Candidatus.Finniella.inopinata GCA_004210305.1_ASM421030v1
100/100
100/100
Candidatus.Odyssella.thessalonicensis.L13 GCA_000190415.2_ASM19041v1
Candidatus.Paracaedibacter.acanthamoebae GCA_000742835.1_ASM74283v1
100/100
Candidatus.Hepatobacter.penaei GCA_000742475.1_ASM74247v1
Holospora.undulata.HU1 GCA_000388175.3_ASM38817v3
92.8/59
Ahrensia.kielensis.DSM.5890 GCA_000374465.1_ASM37446v1
100/100
Chelativorans.oligotrophicus GCA_011317505.1_ASM1131750v1
58.1/85
48.6/53
Bartonella.quintana GCA_009936175.1_ASM993617v1
Rhizobium.leguminosarum.bv.trifolii GCA_011604525.1_ASM1160452v1
97.6/99
Pelagibacterium.halotolerans GCA_013004005.1_ASM1300400v1
100/100
100/100
Polymorphum.gilvum.SL003B.26A1 GCA_000192745.1_ASM19274v1
100/100
Roseibium.hamelinense GCA_007830545.1_ASM783054v1
Pseudovibrio.sp.FO.BEG1 GCA_000236645.1_ASM23664v1
100/100
100/100
Methylocella.silvestris.BL2 GCA_000021745.1_ASM2174v1
100/100
Methylocystis.bryophila GCA_002117405.1_ASM211740v1
99.5/100
100/100
Methylorubrum.extorquens.PA1 GCA_000018845.1_ASM1884v1
100/100
Rhodopseudomonas.palustris GCA_013415845.1_ASM1341584v1
Xanthobacter.autotrophicus.Py2 GCA_000017645.1_ASM1764v1
100/100
100/100
Hyphomicrobium.denitrificans.ATCC.51888 GCA_000143145.1_ASM14314v1
Rhodomicrobium.vannielii.ATCC.17100 GCA_000166055.1_ASM16605v1
100/100
Thermopetrobacter.sp.TC1 GCA_000746275.1_ASM74627v1
100/100
Candidatus.Micropelagos.thuwalensis GCA_000469155.1.RS24
100/100
alpha.proteobacterium.IMCC14465 GCA_000293845.2_ASM29384v2
Parvibaculum.lavamentivorans.DS.1 GCA_000017565.1_ASM1756v1
100/100
100/100
Asticcacaulis.excentricus.CB.48 GCA_000175215.2_ASM17521v2
100/100
Brevundimonas.subvibrioides.ATCC.15264 GCA_000144605.1_ASM14460v1
100/100
alpha.proteobacterium.L41A GCA_000335735.1.L41A.1.0
100/100
Caulobacter.vibrioides.NA1000 GCA_000022005.1_ASM2200v1
100/100
Phenylobacterium.zucineum.HLK1 GCA_000017265.1_ASM1726v1
100/100
Hirschia.baltica.ATCC.49814 GCA_000023785.1_ASM2378v1
100/100
100/100
Hyphomonas.neptunium.ATCC.15444 GCA_000013025.1_ASM1302v1
100/100
Maricaulis.maris GCA_003634045.1_ASM363404v1
Oceanicaulis.alexandrii.DSM.11625 GCA_000420265.1_ASM42026v1
Parvularcula.bermudensis.HTCC2503 GCA_000152825.2_ASM15282v2
100/100
Jannaschia.rubra GCA_900113265.1.IMG.taxon.2622736430.annotated.assembly
100/100
100/100
Octadecabacter.arcticus.238 GCA_000155735.2_ASM15573v2
100/100
Ruegeria.pomeroyi.DSS.3 GCA_000011965.2_ASM1196v2
100/100
Sagittula.stellata.E.37 GCA_000169415.1_ASM16941v1
100/100
100/100
Ketogulonicigenium.vulgare GCA_001399515.1_ASM139951v1
100/100
Rubellimicrobium.rubrum GCA_006152115.1_ASM615211v1
100/100
100/100
Paracoccus.denitrificans GCA_004063735.1_ASM406373v1
Rhodobacter.sphaeroides.ATCC.17029 GCA_000015985.1_ASM1598v1
Neomegalonema.perideroedes.DSM.15528 GCA_000374145.1_ASM37414v1
100/100
100/100
Citromicrobium.sp.JL477 GCA_001304795.1_ASM130479v1
Sphingopyxis.macrogoltabida GCA_001314325.1_ASM131432v1
100/100
100/100
Sphingobium.indicum.B90A GCA_000264945.2_ASM26494v2
100/100
Sphingomonas.wittichii.RW1 GCA_000016765.1_ASM1676v1
100/100
Zymomonas.mobilis.subsp.mobilis.ATCC.31821 GCA_000007105.1_ASM710v1
100/100
alpha.proteobacterium.AAP81b GCA_001295935.1_ASM129593v1
Kordiimonas.gwangyangensis.DSM.19435 GCA_000375545.1_ASM37554v1
100/100
Sneathiella.glossodoripedis.JCM.23214 GCA_000616095.1_ASM61609v1
100/100
Geminicoccus.roseus.DSM.18922 GCA_000427665.1_ASM42766v1
Tistrella.mobilis.KA081020.065 GCA_000264455.2_ASM26445v2
100/100
Candidatus.Puniceispirillum.marinum.IMCC1322 GCA_000024465.1_ASM2446v1
100/100
Thalassobaculum.salexigens.DSM.19539 GCA_000423805.1_ASM42380v1
100/100
alpha.proteobacterium.BAL199 GCA_000171835.1_ASM17183v1
100/100
Oceanibaculum.indicum GCA_003633955.1_ASM363395v1
100/100
Kiloniella.laminariae.DSM.19542 GCA_000374005.1_ASM37400v1
100/100
Rhodovibrio.salinarum.DSM.9154 GCA_000515255.1_ASM51525v1
100/100
100/100
Inquilinus.limosus.DSM.16000 GCA_000423185.1_ASM42318v1
100/100
Rhodospirillum.centenum GCA_014197805.1_ASM1419780v1
Micavibrio.aeruginosavorus.ARL.13 GCA_000226315.1_ASM22631v1
Caenispirillum.salinarum.AK4 GCA_000315795.1
100/100
100/100
Pararhodospirillum.photometricum.DSM.122 GCA_000284415.2_ASM28441v2
43.2/80
38.1/86
Rhodospirillum.rubrum.F11 GCA_000225955.1_ASM22595v1
100/100
Roseospirillum.parvum GCA_900100455.1
100/100
Magnetospirillum.magneticum.AMB.1 GCA_000009985.1_ASM998v1
100/100
Terasakiella.pusilla.DSM.6293 GCA_000688235.1_ASM68823v1
100/100
Thalassospira.profundimaris.WP0211 GCA_000300275.1_ASM30027v1
Elioraea.tepidiphila.DSM.17972 GCA_000378465.1_ASM37846v1
96/100
Belnapia.moabensis.DSM.16746 GCA_000745835.1_ASM74583v1
100/100
Rubritepida.flocculans.DSM.14296 GCA_000425365.1_ASM42536v1
100/100
Roseomonas.gilardii GCA_001941945.1_ASM194194v1
100/100
Acidiphilium.multivorum.AIU301 GCA_000202835.1_ASM20283v1
99.1/99
99.4/99
Acidisphaera.rubrifaciens.HS.AP3 GCA_000964365.1_ASM96436v1
100/100
Granulibacter.bethesdensis.CGDNIH4 GCA_000576145.2_ASM57614v2
100/100
Commensalibacter.intestini.A911 GCA_000231445.2_ASM23144v1
99.3/100
Gluconacetobacter.diazotrophicus.PA1.5 GCA_000067045.1_ASM6704v1
Acetobacter.aceti.NBRC.14818 GCA_000193495.2_ASM19349v2
Gluconobacter.oxydans.DSM.3504 GCA_000583855.1_ASM58385v1
0.812919

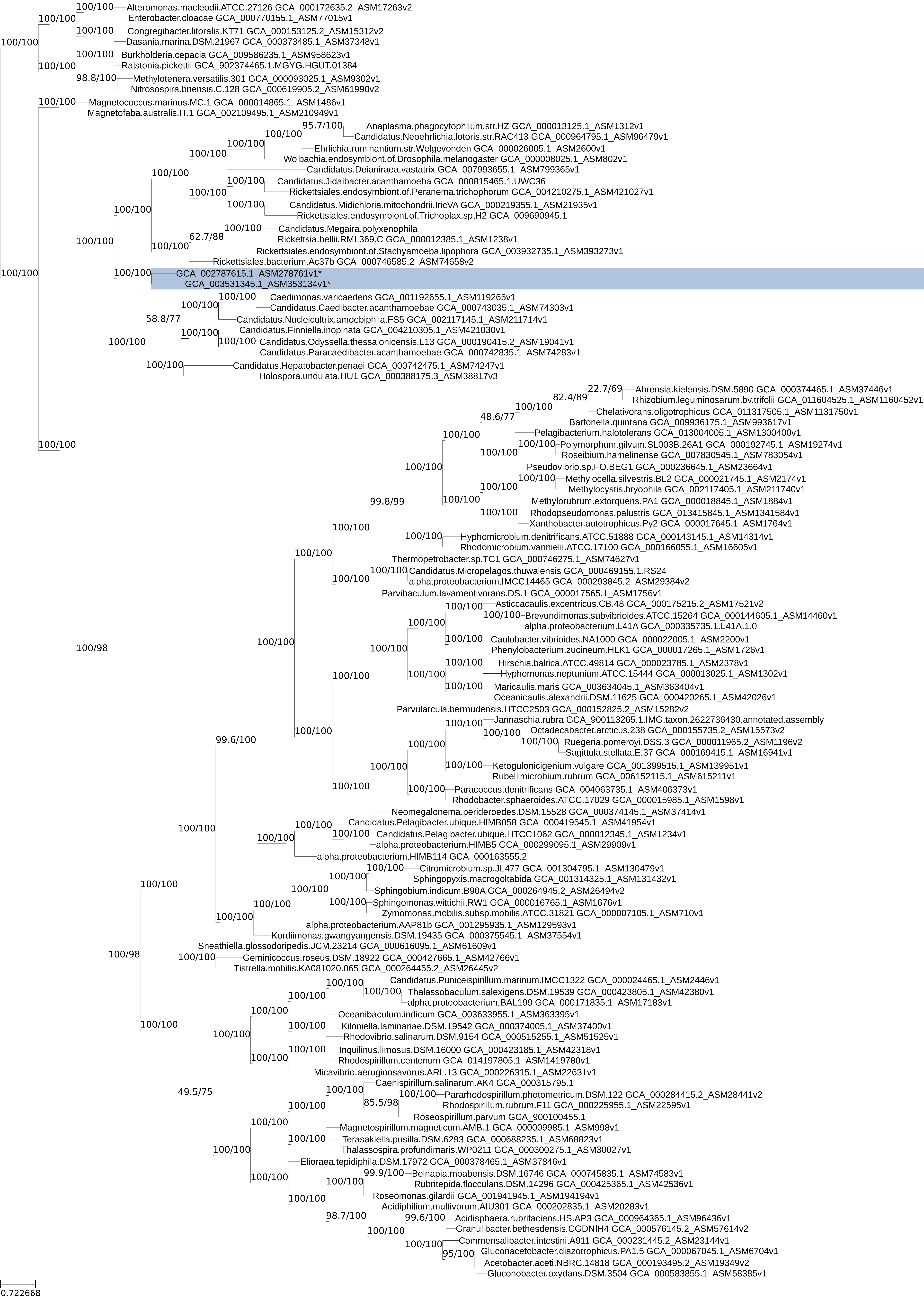

Alteromonas.macleodii.ATCC.27126 GCA_000172635.2_ASM17263v2
Enterobacter.cloacae GCA_000770155.1_ASM77015v1
Congregibacter.litoralis.KT71 GCA_000153125.2_ASM15312v2
Dasania.marina.DSM.21967 GCA_000373485.1_ASM37348v1
Burkholderia.cepacia GCA_009586235.1_ASM958623v1
Ralstonia.pickettii GCA_902374465.1.MGYG.HGUT.01384
Methylotenera.versatilis.301 GCA_000093025.1_ASM9302v1
Nitrosospira.briensis.C.128 GCA_000619905.2_ASM61990v2
Magnetococcus.marinus.MC.1 GCA_000014865.1_ASM1486v1
Magnetofaba.australis.IT.1 GCA_002109495.1_ASM210949v1
Anaplasma.phagocytophilum.str.HZ GCA_000013125.1_ASM1312v1
Candidatus.Neoehrlichia.lotoris.str.RAC413 GCA_000964795.1_ASM96479v1
Ehrlichia.ruminantium.str.Welgevonden GCA_000026005.1_ASM2600v1
Wolbachia.endosymbiont.of.Drosophila.melanogaster GCA_000008025.1_ASM802v1
Candidatus.Deianiraea.vastatrix GCA_007993655.1_ASM799365v1
Candidatus.Jidaibacter.acanthamoeba GCA_000815465.1.UWC36
Rickettsiales.endosymbiont.of.Peranema.trichophorum GCA_004210275.1_ASM421027v1
Candidatus.Midichloria.mitochondrii.IricVA GCA_000219355.1_ASM21935v1
Rickettsiales.endosymbiont.of.Trichoplax.sp.H2 GCA_009690945.1
Candidatus.Megaira.polyxenophila
Rickettsia.bellii.RML369.C GCA_000012385.1_ASM1238v1
Rickettsiales.endosymbiont.of.Stachyamoeba.lipophora GCA_003932735.1_ASM393273v1
Rickettsiales.bacterium.Ac37b GCA_000746585.2_ASM74658v2
GCA_002787615.1_ASM278761v1*
GCA_003531345.1_ASM353134v1*
Caedimonas.varicaedens GCA_001192655.1_ASM119265v1
Candidatus.Caedibacter.acanthamoebae GCA_000743035.1_ASM74303v1
Candidatus.Nucleicultrix.amoebiphila.FS5 GCA_002117145.1_ASM211714v1
Candidatus.Finniella.inopinata GCA_004210305.1_ASM421030v1
Candidatus.Odyssella.thessalonicensis.L13 GCA_000190415.2_ASM19041v1
Candidatus.Paracaedibacter.acanthamoebae GCA_000742835.1_ASM74283v1
Candidatus.Hepatobacter.penaei GCA_000742475.1_ASM74247v1
Holospora.undulata.HU1 GCA_000388175.3_ASM38817v3
Ahrensia.kielensis.DSM.5890 GCA_000374465.1_ASM37446v1
Rhizobium.leguminosarum.bv.trifolii GCA_011604525.1_ASM1160452v1
Chelativorans.oligotrophicus GCA_011317505.1_ASM1131750v1
Bartonella.quintana GCA_009936175.1_ASM993617v1
Pelagibacterium.halotolerans GCA_013004005.1_ASM1300400v1
Polymorphum.gilvum.SL003B.26A1 GCA_000192745.1_ASM19274v1
Roseibium.hamelinense GCA_007830545.1_ASM783054v1
Pseudovibrio.sp.FO.BEG1 GCA_000236645.1_ASM23664v1
Methylocella.silvestris.BL2 GCA_000021745.1_ASM2174v1
Methylocystis.bryophila GCA_002117405.1_ASM211740v1
Methylorubrum.extorquens.PA1 GCA_000018845.1_ASM1884v1
Rhodopseudomonas.palustris GCA_013415845.1_ASM1341584v1
Xanthobacter.autotrophicus.Py2 GCA_000017645.1_ASM1764v1
Hyphomicrobium.denitrificans.ATCC.51888 GCA_000143145.1_ASM14314v1
Rhodomicrobium.vannielii.ATCC.17100 GCA_000166055.1_ASM16605v1
Thermopetrobacter.sp.TC1 GCA_000746275.1_ASM74627v1
Candidatus.Micropelagos.thuwalensis GCA_000469155.1.RS24
alpha.proteobacterium.IMCC14465 GCA_000293845.2_ASM29384v2
Parvibaculum.lavamentivorans.DS.1 GCA_000017565.1_ASM1756v1
Asticcacaulis.excentricus.CB.48 GCA_000175215.2_ASM17521v2
Brevundimonas.subvibrioides.ATCC.15264 GCA_000144605.1_ASM14460v1
alpha.proteobacterium.L41A GCA_000335735.1.L41A.1.0
Caulobacter.vibrioides.NA1000 GCA_000022005.1_ASM2200v1
Phenylobacterium.zucineum.HLK1 GCA_000017265.1_ASM1726v1
Hirschia.baltica.ATCC.49814 GCA_000023785.1_ASM2378v1
Hyphomonas.neptunium.ATCC.15444 GCA_000013025.1_ASM1302v1
Maricaulis.maris GCA_003634045.1_ASM363404v1
Oceanicaulis.alexandrii.DSM.11625 GCA_000420265.1_ASM42026v1
Parvularcula.bermudensis.HTCC2503 GCA_000152825.2_ASM15282v2
Jannaschia.rubra GCA_900113265.1.IMG.taxon.2622736430.annotated.assembly
Octadecabacter.arcticus.238 GCA_000155735.2_ASM15573v2
Ruegeria.pomeroyi.DSS.3 GCA_000011965.2_ASM1196v2
Sagittula.stellata.E.37 GCA_000169415.1_ASM16941v1
Ketogulonicigenium.vulgare GCA_001399515.1_ASM139951v1
Rubellimicrobium.rubrum GCA_006152115.1_ASM615211v1
Paracoccus.denitrificans GCA_004063735.1_ASM406373v1
Rhodobacter.sphaeroides.ATCC.17029 GCA_000015985.1_ASM1598v1
Neomegalonema.perideroedes.DSM.15528 GCA_000374145.1_ASM37414v1
Candidatus.Pelagibacter.ubique.HIMB058 GCA_000419545.1_ASM41954v1
Candidatus.Pelagibacter.ubique.HTCC1062 GCA_000012345.1_ASM1234v1
alpha.proteobacterium.HIMB5 GCA_000299095.1_ASM29909v1
alpha.proteobacterium.HIMB114 GCA_000163555.2
Citromicrobium.sp.JL477 GCA_001304795.1_ASM130479v1
Sphingopyxis.macrogoltabida GCA_001314325.1_ASM131432v1
Sphingobium.indicum.B90A GCA_000264945.2_ASM26494v2
Sphingomonas.wittichii.RW1 GCA_000016765.1_ASM1676v1
Zymomonas.mobilis.subsp.mobilis.ATCC.31821 GCA_000007105.1_ASM710v1
alpha.proteobacterium.AAP81b GCA_001295935.1_ASM129593v1
Kordiimonas.gwangyangensis.DSM.19435 GCA_000375545.1_ASM37554v1
Sneathiella.glossodoripedis.JCM.23214 GCA_000616095.1_ASM61609v1
Geminicoccus.roseus.DSM.18922 GCA_000427665.1_ASM42766v1
Tistrella.mobilis.KA081020.065 GCA_000264455.2_ASM26445v2
Candidatus.Puniceispirillum.marinum.IMCC1322 GCA_000024465.1_ASM2446v1
Thalassobaculum.salexigens.DSM.19539 GCA_000423805.1_ASM42380v1
alpha.proteobacterium.BAL199 GCA_000171835.1_ASM17183v1
Oceanibaculum.indicum GCA_003633955.1_ASM363395v1
Kiloniella.laminariae.DSM.19542 GCA_000374005.1_ASM37400v1
Rhodovibrio.salinarum.DSM.9154 GCA_000515255.1_ASM51525v1
Inquilinus.limosus.DSM.16000 GCA_000423185.1_ASM42318v1
Rhodospirillum.centenum GCA_014197805.1_ASM1419780v1
Micavibrio.aeruginosavorus.ARL.13 GCA_000226315.1_ASM22631v1
Caenispirillum.salinarum.AK4 GCA_000315795.1
Pararhodospirillum.photometricum.DSM.122 GCA_000284415.2_ASM28441v2
Rhodospirillum.rubrum.F11 GCA_000225955.1_ASM22595v1
Roseospirillum.parvum GCA_900100455.1
Magnetospirillum.magneticum.AMB.1 GCA_000009985.1_ASM998v1
Terasakiella.pusilla.DSM.6293 GCA_000688235.1_ASM68823v1
Thalassospira.profundimaris.WP0211 GCA_000300275.1_ASM30027v1
Elioraea.tepidiphila.DSM.17972 GCA_000378465.1_ASM37846v1
Belnapia.moabensis.DSM.16746 GCA_000745835.1_ASM74583v1
Rubritepida.flocculans.DSM.14296 GCA_000425365.1_ASM42536v1
Roseomonas.gilardii GCA_001941945.1_ASM194194v1
Acidiphilium.multivorum.AIU301 GCA_000202835.1_ASM20283v1
Acidisphaera.rubrifaciens.HS.AP3 GCA_000964365.1_ASM96436v1
Granulibacter.bethesdensis.CGDNIH4 GCA_000576145.2_ASM57614v2
Commensalibacter.intestini.A911 GCA_000231445.2_ASM23144v1
Gluconacetobacter.diazotrophicus.PA1.5 GCA_000067045.1_ASM6704v1
Acetobacter.aceti.NBRC.14818 GCA_000193495.2_ASM19349v2
Gluconobacter.oxydans.DSM.3504 GCA_000583855.1_ASM58385v1
0.722668

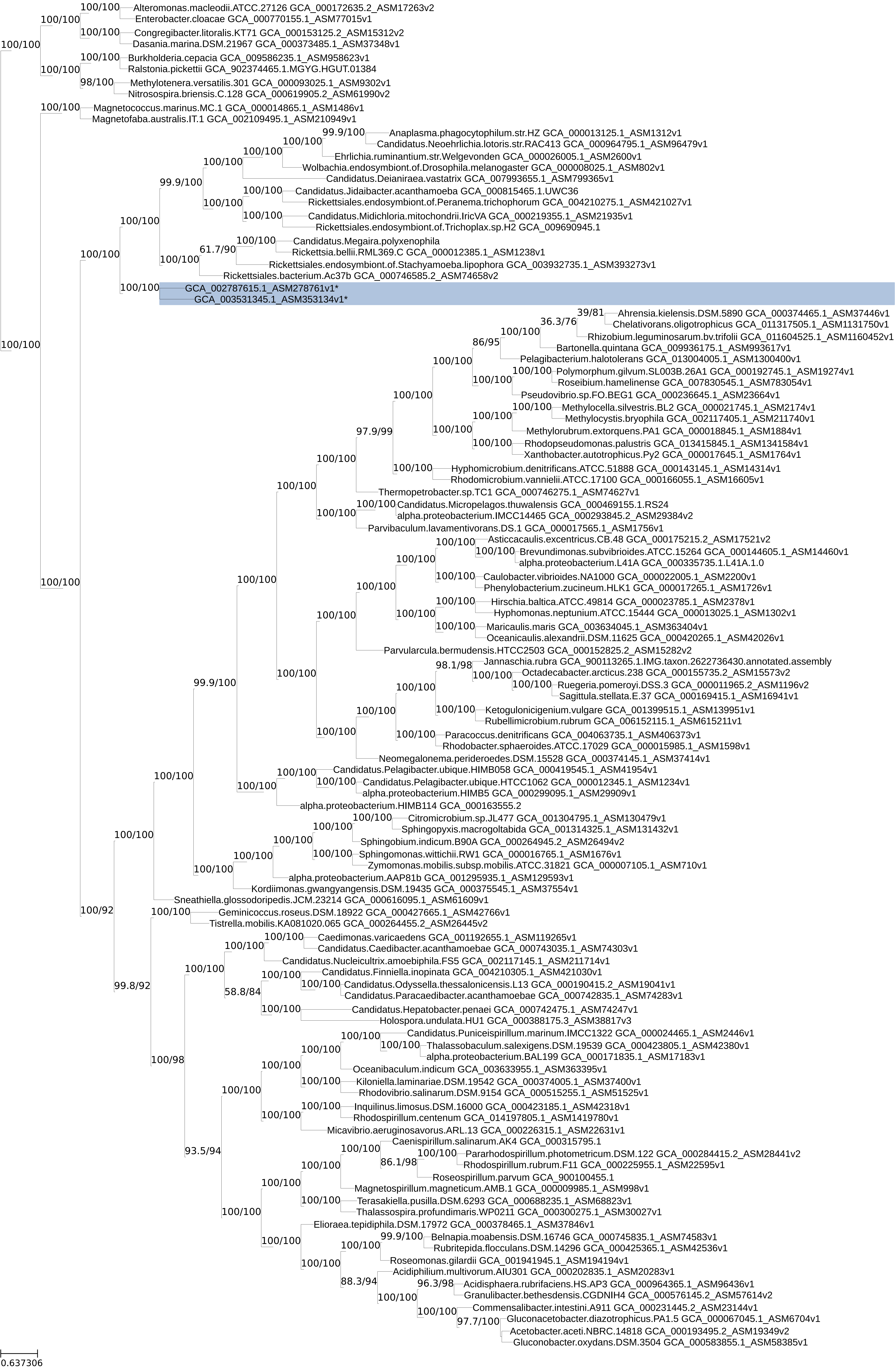
Alteromonas.macleodii.ATCC.27126 GCA_000172635.2_ASM17263v2
Enterobacter.cloacae GCA_000770155.1_ASM77015v1
Congregibacter.litoralis.KT71 GCA_000153125.2_ASM15312v2
Dasania.marina.DSM.21967 GCA_000373485.1_ASM37348v1
Burkholderia.cepacia GCA_009586235.1_ASM958623v1
Ralstonia.pickettii GCA_902374465.1.MGYG.HGUT.01384
Methylotenera.versatilis.301 GCA_000093025.1_ASM9302v1
Nitrosospira.briensis.C.128 GCA_000619905.2_ASM61990v2
Magnetococcus.marinus.MC.1 GCA_000014865.1_ASM1486v1
Magnetofaba.australis.IT.1 GCA_002109495.1_ASM210949v1
Anaplasma.phagocytophilum.str.HZ GCA_000013125.1_ASM1312v1
Candidatus.Neoehrlichia.lotoris.str.RAC413 GCA_000964795.1_ASM96479v1
Ehrlichia.ruminantium.str.Welgevonden GCA_000026005.1_ASM2600v1
Wolbachia.endosymbiont.of.Drosophila.melanogaster GCA_000008025.1_ASM802v1
Candidatus.Deianiraea.vastatrix GCA_007993655.1_ASM799365v1
Candidatus.Jidaibacter.acanthamoeba GCA_000815465.1.UWC36
Rickettsiales.endosymbiont.of.Peranema.trichophorum GCA_004210275.1_ASM421027v1
Candidatus.Midichloria.mitochondrii.IricVA GCA_000219355.1_ASM21935v1
Rickettsiales.endosymbiont.of.Trichoplax.sp.H2 GCA_009690945.1
Candidatus.Megaira.polyxenophila
Rickettsia.bellii.RML369.C GCA_000012385.1_ASM1238v1
Rickettsiales.endosymbiont.of.Stachyamoeba.lipophora GCA_003932735.1_ASM393273v1
Rickettsiales.bacterium.Ac37b GCA_000746585.2_ASM74658v2
GCA_002787615.1_ASM278761v1*
GCA_003531345.1_ASM353134v1*
Ahrensia.kielensis.DSM.5890 GCA_000374465.1_ASM37446v1
Chelativorans.oligotrophicus GCA_011317505.1_ASM1131750v1
Rhizobium.leguminosarum.bv.trifolii GCA_011604525.1_ASM1160452v1
Bartonella.quintana GCA_009936175.1_ASM993617v1
Pelagibacterium.halotolerans GCA_013004005.1_ASM1300400v1
Polymorphum.gilvum.SL003B.26A1 GCA_000192745.1_ASM19274v1
Roseibium.hamelinense GCA_007830545.1_ASM783054v1
Pseudovibrio.sp.FO.BEG1 GCA_000236645.1_ASM23664v1
Methylocella.silvestris.BL2 GCA_000021745.1_ASM2174v1
Methylocystis.bryophila GCA_002117405.1_ASM211740v1
Methylorubrum.extorquens.PA1 GCA_000018845.1_ASM1884v1
Rhodopseudomonas.palustris GCA_013415845.1_ASM1341584v1
Xanthobacter.autotrophicus.Py2 GCA_000017645.1_ASM1764v1
Hyphomicrobium.denitrificans.ATCC.51888 GCA_000143145.1_ASM14314v1
Rhodomicrobium.vannielii.ATCC.17100 GCA_000166055.1_ASM16605v1
Thermopetrobacter.sp.TC1 GCA_000746275.1_ASM74627v1
Candidatus.Micropelagos.thuwalensis GCA_000469155.1.RS24
alpha.proteobacterium.IMCC14465 GCA_000293845.2_ASM29384v2
Parvibaculum.lavamentivorans.DS.1 GCA_000017565.1_ASM1756v1
Asticcacaulis.excentricus.CB.48 GCA_000175215.2_ASM17521v2
Brevundimonas.subvibrioides.ATCC.15264 GCA_000144605.1_ASM14460v1
alpha.proteobacterium.L41A GCA_000335735.1.L41A.1.0
Caulobacter.vibrioides.NA1000 GCA_000022005.1_ASM2200v1
Phenylobacterium.zucineum.HLK1 GCA_000017265.1_ASM1726v1
Hirschia.baltica.ATCC.49814 GCA_000023785.1_ASM2378v1
Hyphomonas.neptunium.ATCC.15444 GCA_000013025.1_ASM1302v1
Maricaulis.maris GCA_003634045.1_ASM363404v1
Oceanicaulis.alexandrii.DSM.11625 GCA_000420265.1_ASM42026v1
Parvularcula.bermudensis.HTCC2503 GCA_000152825.2_ASM15282v2
Jannaschia.rubra GCA_900113265.1.IMG.taxon.2622736430.annotated.assembly
Octadecabacter.arcticus.238 GCA_000155735.2_ASM15573v2
Ruegeria.pomeroyi.DSS.3 GCA_000011965.2_ASM1196v2
Sagittula.stellata.E.37 GCA_000169415.1_ASM16941v1
Ketogulonicigenium.vulgare GCA_001399515.1_ASM139951v1
Rubellimicrobium.rubrum GCA_006152115.1_ASM615211v1
Paracoccus.denitrificans GCA_004063735.1_ASM406373v1
Rhodobacter.sphaeroides.ATCC.17029 GCA_000015985.1_ASM1598v1
Neomegalonema.perideroedes.DSM.15528 GCA_000374145.1_ASM37414v1
Candidatus.Pelagibacter.ubique.HIMB058 GCA_000419545.1_ASM41954v1
Candidatus.Pelagibacter.ubique.HTCC1062 GCA_000012345.1_ASM1234v1
alpha.proteobacterium.HIMB5 GCA_000299095.1_ASM29909v1
alpha.proteobacterium.HIMB114 GCA_000163555.2
Citromicrobium.sp.JL477 GCA_001304795.1_ASM130479v1
Sphingopyxis.macrogoltabida GCA_001314325.1_ASM131432v1
Sphingobium.indicum.B90A GCA_000264945.2_ASM26494v2
Sphingomonas.wittichii.RW1 GCA_000016765.1_ASM1676v1
Zymomonas.mobilis.subsp.mobilis.ATCC.31821 GCA_000007105.1_ASM710v1
alpha.proteobacterium.AAP81b GCA_001295935.1_ASM129593v1
Kordiimonas.gwangyangensis.DSM.19435 GCA_000375545.1_ASM37554v1
Sneathiella.glossodoripedis.JCM.23214 GCA_000616095.1_ASM61609v1
Geminicoccus.roseus.DSM.18922 GCA_000427665.1_ASM42766v1
Tistrella.mobilis.KA081020.065 GCA_000264455.2_ASM26445v2
Caedimonas.varicaedens GCA_001192655.1_ASM119265v1
Candidatus.Caedibacter.acanthamoebae GCA_000743035.1_ASM74303v1
Candidatus.Nucleicultrix.amoebiphila.FS5 GCA_002117145.1_ASM211714v1
Candidatus.Finniella.inopinata GCA_004210305.1_ASM421030v1
Candidatus.Odyssella.thessalonicensis.L13 GCA_000190415.2_ASM19041v1
Candidatus.Paracaedibacter.acanthamoebae GCA_000742835.1_ASM74283v1
Candidatus.Hepatobacter.penaei GCA_000742475.1_ASM74247v1
Holospora.undulata.HU1 GCA_000388175.3_ASM38817v3
Candidatus.Puniceispirillum.marinum.IMCC1322 GCA_000024465.1_ASM2446v1
Thalassobaculum.salexigens.DSM.19539 GCA_000423805.1_ASM42380v1
alpha.proteobacterium.BAL199 GCA_000171835.1_ASM17183v1
Oceanibaculum.indicum GCA_003633955.1_ASM363395v1
Kiloniella.laminariae.DSM.19542 GCA_000374005.1_ASM37400v1
Rhodovibrio.salinarum.DSM.9154 GCA_000515255.1_ASM51525v1
Inquilinus.limosus.DSM.16000 GCA_000423185.1_ASM42318v1
Rhodospirillum.centenum GCA_014197805.1_ASM1419780v1
Micavibrio.aeruginosavorus.ARL.13 GCA_000226315.1_ASM22631v1
Caenispirillum.salinarum.AK4 GCA_000315795.1
Pararhodospirillum.photometricum.DSM.122 GCA_000284415.2_ASM28441v2
Rhodospirillum.rubrum.F11 GCA_000225955.1_ASM22595v1
Roseospirillum.parvum GCA_900100455.1
Magnetospirillum.magneticum.AMB.1 GCA_000009985.1_ASM998v1
Terasakiella.pusilla.DSM.6293 GCA_000688235.1_ASM68823v1
Thalassospira.profundimaris.WP0211 GCA_000300275.1_ASM30027v1
Elioraea.tepidiphila.DSM.17972 GCA_000378465.1_ASM37846v1
Belnapia.moabensis.DSM.16746 GCA_000745835.1_ASM74583v1
Rubritepida.flocculans.DSM.14296 GCA_000425365.1_ASM42536v1
Roseomonas.gilardii GCA_001941945.1_ASM194194v1
Acidiphilium.multivorum.AIU301 GCA_000202835.1_ASM20283v1
Acidisphaera.rubrifaciens.HS.AP3 GCA_000964365.1_ASM96436v1
Granulibacter.bethesdensis.CGDNIH4 GCA_000576145.2_ASM57614v2
Commensalibacter.intestini.A911 GCA_000231445.2_ASM23144v1
Gluconacetobacter.diazotrophicus.PA1.5 GCA_000067045.1_ASM6704v1
Acetobacter.aceti.NBRC.14818 GCA_000193495.2_ASM19349v2
Gluconobacter.oxydans.DSM.3504 GCA_000583855.1_ASM58385v1
0.637306

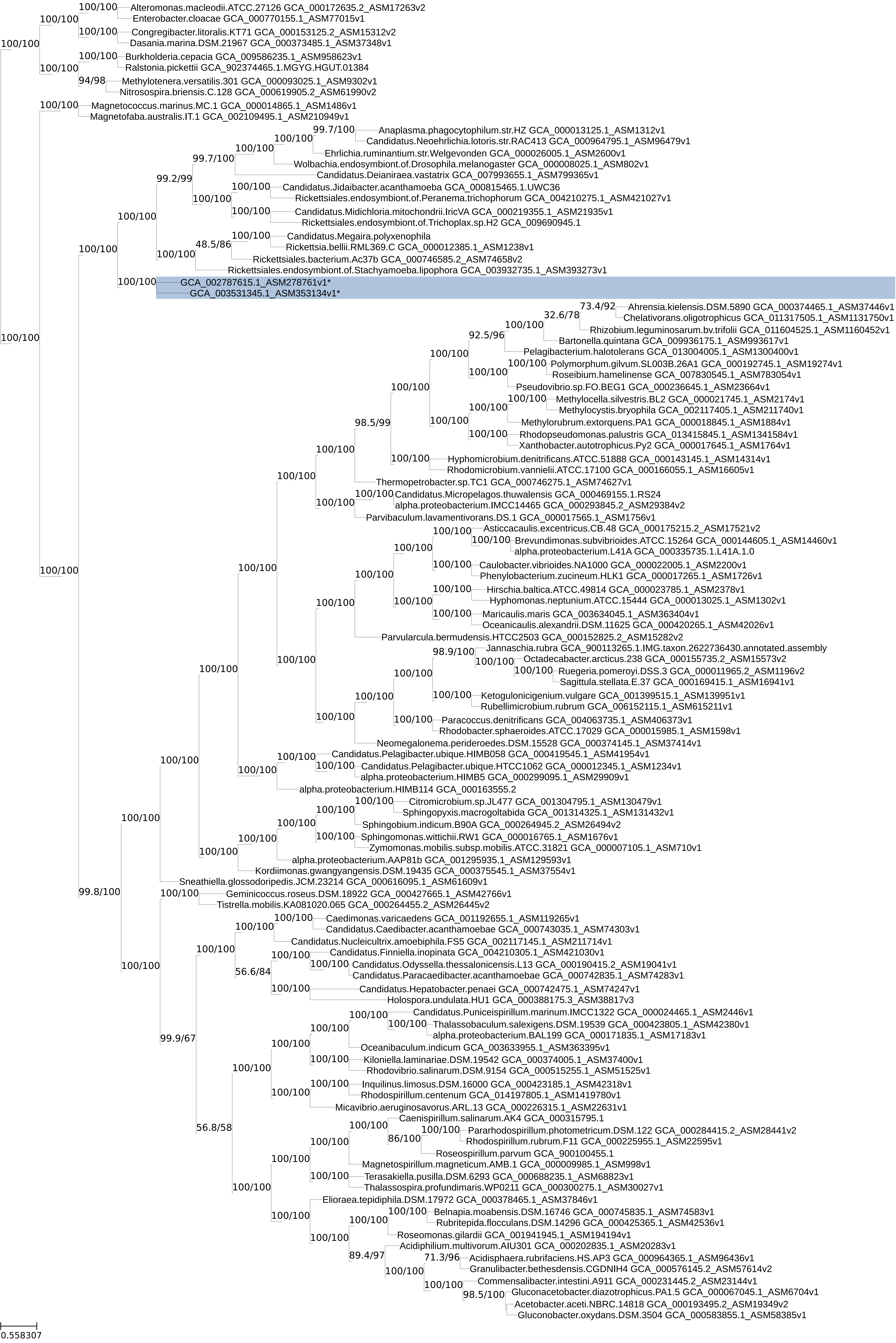
Alteromonas.macleodii.ATCC.27126 GCA_000172635.2_ASM17263v2
Enterobacter.cloacae GCA_000770155.1_ASM77015v1
Congregibacter.litoralis.KT71 GCA_000153125.2_ASM15312v2
Dasania.marina.DSM.21967 GCA_000373485.1_ASM37348v1
Burkholderia.cepacia GCA_009586235.1_ASM958623v1
Ralstonia.pickettii GCA_902374465.1.MGYG.HGUT.01384
Methylotenera.versatilis.301 GCA_000093025.1_ASM9302v1
Nitrosospira.briensis.C.128 GCA_000619905.2_ASM61990v2
Magnetococcus.marinus.MC.1 GCA_000014865.1_ASM1486v1
Magnetofaba.australis.IT.1 GCA_002109495.1_ASM210949v1
Anaplasma.phagocytophilum.str.HZ GCA_000013125.1_ASM1312v1
Candidatus.Neoehrlichia.lotoris.str.RAC413 GCA_000964795.1_ASM96479v1
Ehrlichia.ruminantium.str.Welgevonden GCA_000026005.1_ASM2600v1
Wolbachia.endosymbiont.of.Drosophila.melanogaster GCA_000008025.1_ASM802v1
Candidatus.Deianiraea.vastatrix GCA_007993655.1_ASM799365v1
Candidatus.Jidaibacter.acanthamoeba GCA_000815465.1.UWC36
Rickettsiales.endosymbiont.of.Peranema.trichophorum GCA_004210275.1_ASM421027v1
Candidatus.Midichloria.mitochondrii.IricVA GCA_000219355.1_ASM21935v1
Rickettsiales.endosymbiont.of.Trichoplax.sp.H2 GCA_009690945.1
Candidatus.Megaira.polyxenophila
Rickettsia.bellii.RML369.C GCA_000012385.1_ASM1238v1
Rickettsiales.bacterium.Ac37b GCA_000746585.2_ASM74658v2
Rickettsiales.endosymbiont.of.Stachyamoeba.lipophora GCA_003932735.1_ASM393273v1
GCA_002787615.1_ASM278761v1*
GCA_003531345.1_ASM353134v1*
Ahrensia.kielensis.DSM.5890 GCA_000374465.1_ASM37446v1
Chelativorans.oligotrophicus GCA_011317505.1_ASM1131750v1
Rhizobium.leguminosarum.bv.trifolii GCA_011604525.1_ASM1160452v1
Bartonella.quintana GCA_009936175.1_ASM993617v1
Pelagibacterium.halotolerans GCA_013004005.1_ASM1300400v1
Polymorphum.gilvum.SL003B.26A1 GCA_000192745.1_ASM19274v1
Roseibium.hamelinense GCA_007830545.1_ASM783054v1
Pseudovibrio.sp.FO.BEG1 GCA_000236645.1_ASM23664v1
Methylocella.silvestris.BL2 GCA_000021745.1_ASM2174v1
Methylocystis.bryophila GCA_002117405.1_ASM211740v1
Methylorubrum.extorquens.PA1 GCA_000018845.1_ASM1884v1
Rhodopseudomonas.palustris GCA_013415845.1_ASM1341584v1
Xanthobacter.autotrophicus.Py2 GCA_000017645.1_ASM1764v1
Hyphomicrobium.denitrificans.ATCC.51888 GCA_000143145.1_ASM14314v1
Rhodomicrobium.vannielii.ATCC.17100 GCA_000166055.1_ASM16605v1
Thermopetrobacter.sp.TC1 GCA_000746275.1_ASM74627v1
Candidatus.Micropelagos.thuwalensis GCA_000469155.1.RS24
alpha.proteobacterium.IMCC14465 GCA_000293845.2_ASM29384v2
Parvibaculum.lavamentivorans.DS.1 GCA_000017565.1_ASM1756v1
Asticcacaulis.excentricus.CB.48 GCA_000175215.2_ASM17521v2
Brevundimonas.subvibrioides.ATCC.15264 GCA_000144605.1_ASM14460v1
alpha.proteobacterium.L41A GCA_000335735.1.L41A.1.0
Caulobacter.vibrioides.NA1000 GCA_000022005.1_ASM2200v1
Phenylobacterium.zucineum.HLK1 GCA_000017265.1_ASM1726v1
Hirschia.baltica.ATCC.49814 GCA_000023785.1_ASM2378v1
Hyphomonas.neptunium.ATCC.15444 GCA_000013025.1_ASM1302v1
Maricaulis.maris GCA_003634045.1_ASM363404v1
Oceanicaulis.alexandrii.DSM.11625 GCA_000420265.1_ASM42026v1
Parvularcula.bermudensis.HTCC2503 GCA_000152825.2_ASM15282v2
Jannaschia.rubra GCA_900113265.1.IMG.taxon.2622736430.annotated.assembly
Octadecabacter.arcticus.238 GCA_000155735.2_ASM15573v2
Ruegeria.pomeroyi.DSS.3 GCA_000011965.2_ASM1196v2
Sagittula.stellata.E.37 GCA_000169415.1_ASM16941v1
Ketogulonicigenium.vulgare GCA_001399515.1_ASM139951v1
Rubellimicrobium.rubrum GCA_006152115.1_ASM615211v1
Paracoccus.denitrificans GCA_004063735.1_ASM406373v1
Rhodobacter.sphaeroides.ATCC.17029 GCA_000015985.1_ASM1598v1
Neomegalonema.perideroedes.DSM.15528 GCA_000374145.1_ASM37414v1
Candidatus.Pelagibacter.ubique.HIMB058 GCA_000419545.1_ASM41954v1
Candidatus.Pelagibacter.ubique.HTCC1062 GCA_000012345.1_ASM1234v1
alpha.proteobacterium.HIMB5 GCA_000299095.1_ASM29909v1
alpha.proteobacterium.HIMB114 GCA_000163555.2
Citromicrobium.sp.JL477 GCA_001304795.1_ASM130479v1
Sphingopyxis.macrogoltabida GCA_001314325.1_ASM131432v1
Sphingobium.indicum.B90A GCA_000264945.2_ASM26494v2
Sphingomonas.wittichii.RW1 GCA_000016765.1_ASM1676v1
Zymomonas.mobilis.subsp.mobilis.ATCC.31821 GCA_000007105.1_ASM710v1
alpha.proteobacterium.AAP81b GCA_001295935.1_ASM129593v1
Kordiimonas.gwangyangensis.DSM.19435 GCA_000375545.1_ASM37554v1
Sneathiella.glossodoripedis.JCM.23214 GCA_000616095.1_ASM61609v1
Geminicoccus.roseus.DSM.18922 GCA_000427665.1_ASM42766v1
Tistrella.mobilis.KA081020.065 GCA_000264455.2_ASM26445v2
Caedimonas.varicaedens GCA_001192655.1_ASM119265v1
Candidatus.Caedibacter.acanthamoebae GCA_000743035.1_ASM74303v1
Candidatus.Nucleicultrix.amoebiphila.FS5 GCA_002117145.1_ASM211714v1
Candidatus.Finniella.inopinata GCA_004210305.1_ASM421030v1
Candidatus.Odyssella.thessalonicensis.L13 GCA_000190415.2_ASM19041v1
Candidatus.Paracaedibacter.acanthamoebae GCA_000742835.1_ASM74283v1
Candidatus.Hepatobacter.penaei GCA_000742475.1_ASM74247v1
Holospora.undulata.HU1 GCA_000388175.3_ASM38817v3
Candidatus.Puniceispirillum.marinum.IMCC1322 GCA_000024465.1_ASM2446v1
Thalassobaculum.salexigens.DSM.19539 GCA_000423805.1_ASM42380v1
alpha.proteobacterium.BAL199 GCA_000171835.1_ASM17183v1
Oceanibaculum.indicum GCA_003633955.1_ASM363395v1
Kiloniella.laminariae.DSM.19542 GCA_000374005.1_ASM37400v1
Rhodovibrio.salinarum.DSM.9154 GCA_000515255.1_ASM51525v1
Inquilinus.limosus.DSM.16000 GCA_000423185.1_ASM42318v1
Rhodospirillum.centenum GCA_014197805.1_ASM1419780v1
Micavibrio.aeruginosavorus.ARL.13 GCA_000226315.1_ASM22631v1
Caenispirillum.salinarum.AK4 GCA_000315795.1
Pararhodospirillum.photometricum.DSM.122 GCA_000284415.2_ASM28441v2
Rhodospirillum.rubrum.F11 GCA_000225955.1_ASM22595v1
Roseospirillum.parvum GCA_900100455.1
Magnetospirillum.magneticum.AMB.1 GCA_000009985.1_ASM998v1
Terasakiella.pusilla.DSM.6293 GCA_000688235.1_ASM68823v1
Thalassospira.profundimaris.WP0211 GCA_000300275.1_ASM30027v1
Elioraea.tepidiphila.DSM.17972 GCA_000378465.1_ASM37846v1
Belnapia.moabensis.DSM.16746 GCA_000745835.1_ASM74583v1
Rubritepida.flocculans.DSM.14296 GCA_000425365.1_ASM42536v1
Roseomonas.gilardii GCA_001941945.1_ASM194194v1
Acidiphilium.multivorum.AIU301 GCA_000202835.1_ASM20283v1
Acidisphaera.rubrifaciens.HS.AP3 GCA_000964365.1_ASM96436v1
Granulibacter.bethesdensis.CGDNIH4 GCA_000576145.2_ASM57614v2
Commensalibacter.intestini.A911 GCA_000231445.2_ASM23144v1
Gluconacetobacter.diazotrophicus.PA1.5 GCA_000067045.1_ASM6704v1
Acetobacter.aceti.NBRC.14818 GCA_000193495.2_ASM19349v2
Gluconobacter.oxydans.DSM.3504 GCA_000583855.1_ASM58385v1
0.558307

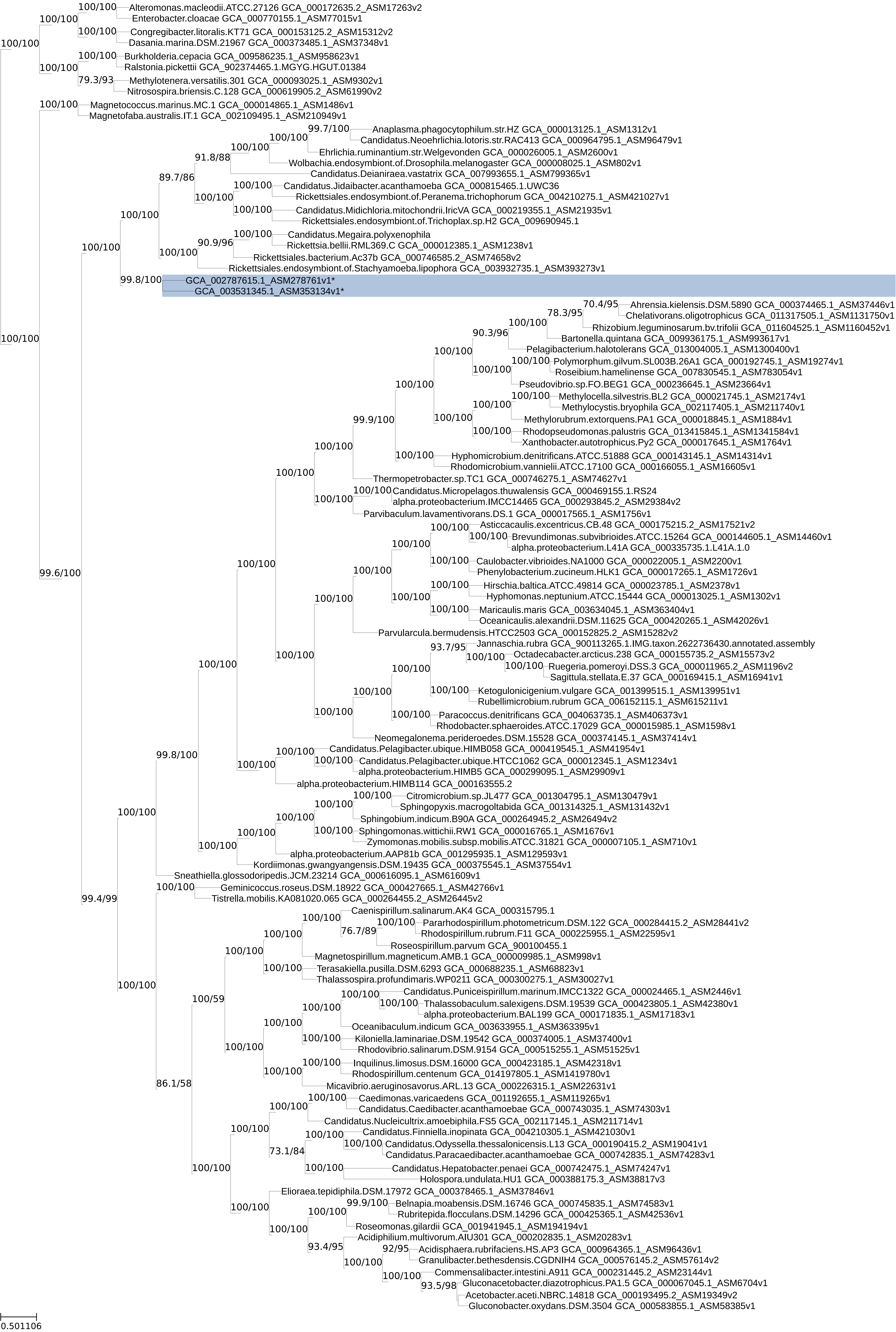
Alteromonas.macleodii.ATCC.27126 GCA_000172635.2_ASM17263v2
Enterobacter.cloacae GCA_000770155.1_ASM77015v1
Congregibacter.litoralis.KT71 GCA_000153125.2_ASM15312v2
Dasania.marina.DSM.21967 GCA_000373485.1_ASM37348v1
Burkholderia.cepacia GCA_009586235.1_ASM958623v1
Ralstonia.pickettii GCA_902374465.1.MGYG.HGUT.01384
Methylotenera.versatilis.301 GCA_000093025.1_ASM9302v1
Nitrosospira.briensis.C.128 GCA_000619905.2_ASM61990v2
Magnetococcus.marinus.MC.1 GCA_000014865.1_ASM1486v1
Magnetofaba.australis.IT.1 GCA_002109495.1_ASM210949v1
Anaplasma.phagocytophilum.str.HZ GCA_000013125.1_ASM1312v1
Candidatus.Neoehrlichia.lotoris.str.RAC413 GCA_000964795.1_ASM96479v1
Ehrlichia.ruminantium.str.Welgevonden GCA_000026005.1_ASM2600v1
Wolbachia.endosymbiont.of.Drosophila.melanogaster GCA_000008025.1_ASM802v1
Candidatus.Deianiraea.vastatrix GCA_007993655.1_ASM799365v1
Candidatus.Jidaibacter.acanthamoeba GCA_000815465.1.UWC36
Rickettsiales.endosymbiont.of.Peranema.trichophorum GCA_004210275.1_ASM421027v1
Candidatus.Midichloria.mitochondrii.IricVA GCA_000219355.1_ASM21935v1
Rickettsiales.endosymbiont.of.Trichoplax.sp.H2 GCA_009690945.1
Candidatus.Megaira.polyxenophila
Rickettsia.bellii.RML369.C GCA_000012385.1_ASM1238v1
Rickettsiales.bacterium.Ac37b GCA_000746585.2_ASM74658v2
Rickettsiales.endosymbiont.of.Stachyamoeba.lipophora GCA_003932735.1_ASM393273v1
GCA_002787615.1_ASM278761v1*
GCA_003531345.1_ASM353134v1*
Ahrensia.kielensis.DSM.5890 GCA_000374465.1_ASM37446v1
Chelativorans.oligotrophicus GCA_011317505.1_ASM1131750v1
Rhizobium.leguminosarum.bv.trifolii GCA_011604525.1_ASM1160452v1
Bartonella.quintana GCA_009936175.1_ASM993617v1
Pelagibacterium.halotolerans GCA_013004005.1_ASM1300400v1
Polymorphum.gilvum.SL003B.26A1 GCA_000192745.1_ASM19274v1
Roseibium.hamelinense GCA_007830545.1_ASM783054v1
Pseudovibrio.sp.FO.BEG1 GCA_000236645.1_ASM23664v1
Methylocella.silvestris.BL2 GCA_000021745.1_ASM2174v1
Methylocystis.bryophila GCA_002117405.1_ASM211740v1
Methylorubrum.extorquens.PA1 GCA_000018845.1_ASM1884v1
Rhodopseudomonas.palustris GCA_013415845.1_ASM1341584v1
Xanthobacter.autotrophicus.Py2 GCA_000017645.1_ASM1764v1
Hyphomicrobium.denitrificans.ATCC.51888 GCA_000143145.1_ASM14314v1
Rhodomicrobium.vannielii.ATCC.17100 GCA_000166055.1_ASM16605v1
Thermopetrobacter.sp.TC1 GCA_000746275.1_ASM74627v1
Candidatus.Micropelagos.thuwalensis GCA_000469155.1.RS24
alpha.proteobacterium.IMCC14465 GCA_000293845.2_ASM29384v2
Parvibaculum.lavamentivorans.DS.1 GCA_000017565.1_ASM1756v1
Asticcacaulis.excentricus.CB.48 GCA_000175215.2_ASM17521v2
Brevundimonas.subvibrioides.ATCC.15264 GCA_000144605.1_ASM14460v1
alpha.proteobacterium.L41A GCA_000335735.1.L41A.1.0
Caulobacter.vibrioides.NA1000 GCA_000022005.1_ASM2200v1
Phenylobacterium.zucineum.HLK1 GCA_000017265.1_ASM1726v1
Hirschia.baltica.ATCC.49814 GCA_000023785.1_ASM2378v1
Hyphomonas.neptunium.ATCC.15444 GCA_000013025.1_ASM1302v1
Maricaulis.maris GCA_003634045.1_ASM363404v1
Oceanicaulis.alexandrii.DSM.11625 GCA_000420265.1_ASM42026v1
Parvularcula.bermudensis.HTCC2503 GCA_000152825.2_ASM15282v2
Jannaschia.rubra GCA_900113265.1.IMG.taxon.2622736430.annotated.assembly
Octadecabacter.arcticus.238 GCA_000155735.2_ASM15573v2
Ruegeria.pomeroyi.DSS.3 GCA_000011965.2_ASM1196v2
Sagittula.stellata.E.37 GCA_000169415.1_ASM16941v1
Ketogulonicigenium.vulgare GCA_001399515.1_ASM139951v1
Rubellimicrobium.rubrum GCA_006152115.1_ASM615211v1
Paracoccus.denitrificans GCA_004063735.1_ASM406373v1
Rhodobacter.sphaeroides.ATCC.17029 GCA_000015985.1_ASM1598v1
Neomegalonema.perideroedes.DSM.15528 GCA_000374145.1_ASM37414v1
Candidatus.Pelagibacter.ubique.HIMB058 GCA_000419545.1_ASM41954v1
Candidatus.Pelagibacter.ubique.HTCC1062 GCA_000012345.1_ASM1234v1
alpha.proteobacterium.HIMB5 GCA_000299095.1_ASM29909v1
alpha.proteobacterium.HIMB114 GCA_000163555.2
Citromicrobium.sp.JL477 GCA_001304795.1_ASM130479v1
Sphingopyxis.macrogoltabida GCA_001314325.1_ASM131432v1
Sphingobium.indicum.B90A GCA_000264945.2_ASM26494v2
Sphingomonas.wittichii.RW1 GCA_000016765.1_ASM1676v1
Zymomonas.mobilis.subsp.mobilis.ATCC.31821 GCA_000007105.1_ASM710v1
alpha.proteobacterium.AAP81b GCA_001295935.1_ASM129593v1
Kordiimonas.gwangyangensis.DSM.19435 GCA_000375545.1_ASM37554v1
Sneathiella.glossodoripedis.JCM.23214 GCA_000616095.1_ASM61609v1
Geminicoccus.roseus.DSM.18922 GCA_000427665.1_ASM42766v1
Tistrella.mobilis.KA081020.065 GCA_000264455.2_ASM26445v2
Caenispirillum.salinarum.AK4 GCA_000315795.1
Pararhodospirillum.photometricum.DSM.122 GCA_000284415.2_ASM28441v2
Rhodospirillum.rubrum.F11 GCA_000225955.1_ASM22595v1
Roseospirillum.parvum GCA_900100455.1
Magnetospirillum.magneticum.AMB.1 GCA_000009985.1_ASM998v1
Terasakiella.pusilla.DSM.6293 GCA_000688235.1_ASM68823v1
Thalassospira.profundimaris.WP0211 GCA_000300275.1_ASM30027v1
Candidatus.Puniceispirillum.marinum.IMCC1322 GCA_000024465.1_ASM2446v1
Thalassobaculum.salexigens.DSM.19539 GCA_000423805.1_ASM42380v1
alpha.proteobacterium.BAL199 GCA_000171835.1_ASM17183v1
Oceanibaculum.indicum GCA_003633955.1_ASM363395v1
Kiloniella.laminariae.DSM.19542 GCA_000374005.1_ASM37400v1
Rhodovibrio.salinarum.DSM.9154 GCA_000515255.1_ASM51525v1
Inquilinus.limosus.DSM.16000 GCA_000423185.1_ASM42318v1
Rhodospirillum.centenum GCA_014197805.1_ASM1419780v1
Micavibrio.aeruginosavorus.ARL.13 GCA_000226315.1_ASM22631v1
Caedimonas.varicaedens GCA_001192655.1_ASM119265v1
Candidatus.Caedibacter.acanthamoebae GCA_000743035.1_ASM74303v1
Candidatus.Nucleicultrix.amoebiphila.FS5 GCA_002117145.1_ASM211714v1
Candidatus.Finniella.inopinata GCA_004210305.1_ASM421030v1
Candidatus.Odyssella.thessalonicensis.L13 GCA_000190415.2_ASM19041v1
Candidatus.Paracaedibacter.acanthamoebae GCA_000742835.1_ASM74283v1
Candidatus.Hepatobacter.penaei GCA_000742475.1_ASM74247v1
Holospora.undulata.HU1 GCA_000388175.3_ASM38817v3
Elioraea.tepidiphila.DSM.17972 GCA_000378465.1_ASM37846v1
Belnapia.moabensis.DSM.16746 GCA_000745835.1_ASM74583v1
Rubritepida.flocculans.DSM.14296 GCA_000425365.1_ASM42536v1
Roseomonas.gilardii GCA_001941945.1_ASM194194v1
Acidiphilium.multivorum.AIU301 GCA_000202835.1_ASM20283v1
Acidisphaera.rubrifaciens.HS.AP3 GCA_000964365.1_ASM96436v1
Granulibacter.bethesdensis.CGDNIH4 GCA_000576145.2_ASM57614v2
Commensalibacter.intestini.A911 GCA_000231445.2_ASM23144v1
Gluconacetobacter.diazotrophicus.PA1.5 GCA_000067045.1_ASM6704v1
Acetobacter.aceti.NBRC.14818 GCA_000193495.2_ASM19349v2
Gluconobacter.oxydans.DSM.3504 GCA_000583855.1_ASM58385v1
0.501106

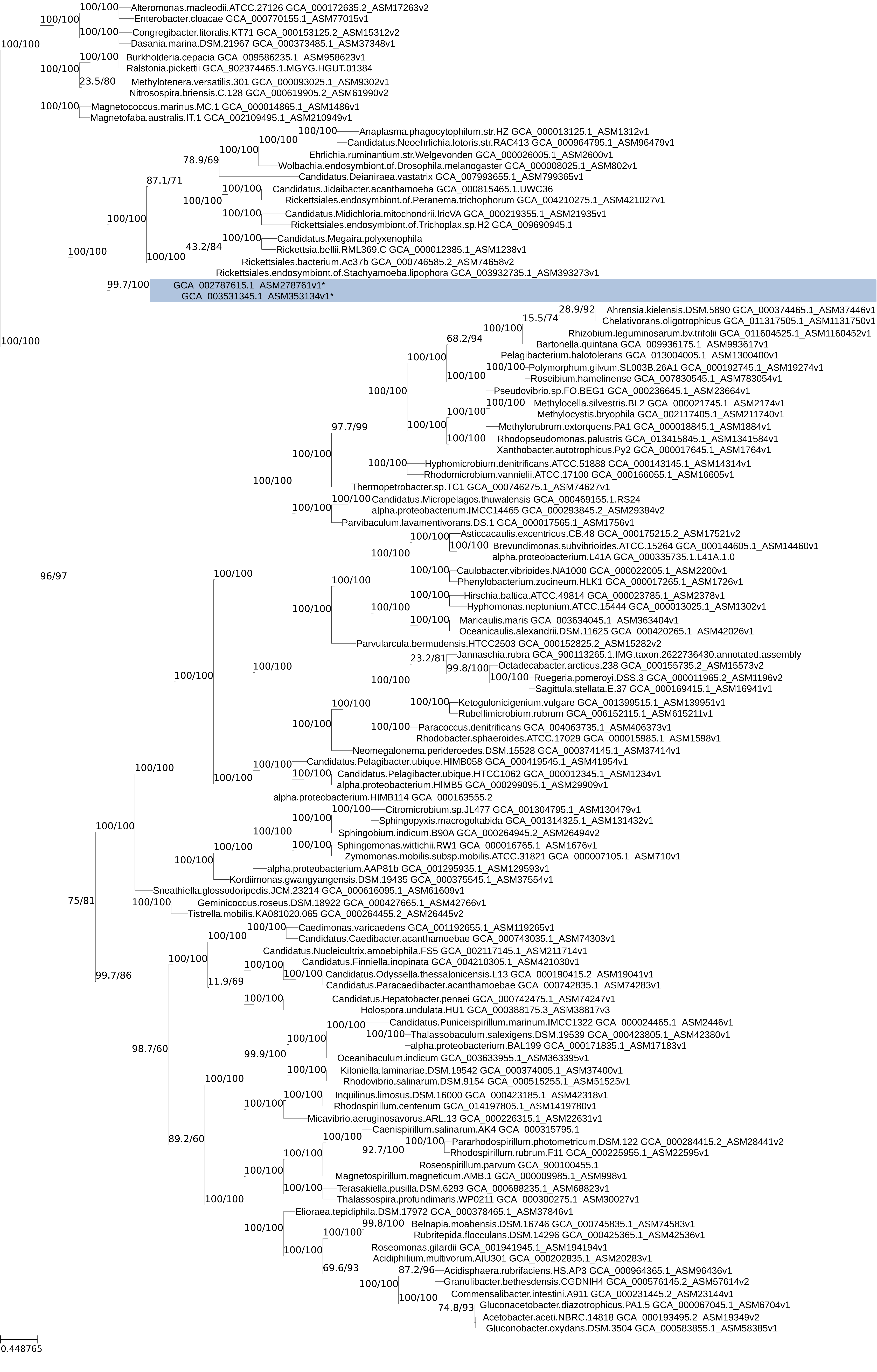
100/100
Alteromonas.macleodii.ATCC.27126 GCA_000172635.2_ASM17263v2
100/100
Enterobacter.cloacae GCA_000770155.1_ASM77015v1
100/100
Congregibacter.litoralis.KT71 GCA_000153125.2_ASM15312v2
100/100
Dasania.marina.DSM.21967 GCA_000373485.1_ASM37348v1
100/100
Burkholderia.cepacia GCA_009586235.1_ASM958623v1
100/100
Ralstonia.pickettii GCA_902374465.1.MGYG.HGUT.01384
23.5/80
Methylotenera.versatilis.301 GCA_000093025.1_ASM9302v1
Nitrosospira.briensis.C.128 GCA_000619905.2_ASM61990v2
100/100
Magnetococcus.marinus.MC.1 GCA_000014865.1_ASM1486v1
Magnetofaba.australis.IT.1 GCA_002109495.1_ASM210949v1
100/100
Anaplasma.phagocytophilum.str.HZ GCA_000013125.1_ASM1312v1
100/100
Candidatus.Neoehrlichia.lotoris.str.RAC413 GCA_000964795.1_ASM96479v1
100/100
Ehrlichia.ruminantium.str.Welgevonden GCA_000026005.1_ASM2600v1
78.9/69
Wolbachia.endosymbiont.of.Drosophila.melanogaster GCA_000008025.1_ASM802v1
Candidatus.Deianiraea.vastatrix GCA_007993655.1_ASM799365v1
87.1/71
100/100
Candidatus.Jidaibacter.acanthamoeba GCA_000815465.1.UWC36
100/100
Rickettsiales.endosymbiont.of.Peranema.trichophorum GCA_004210275.1_ASM421027v1
100/100
100/100
Candidatus.Midichloria.mitochondrii.IricVA GCA_000219355.1_ASM21935v1
Rickettsiales.endosymbiont.of.Trichoplax.sp.H2 GCA_009690945.1
100/100
Candidatus.Megaira.polyxenophila
43.2/84
Rickettsia.bellii.RML369.C GCA_000012385.1_ASM1238v1
100/100
Rickettsiales.bacterium.Ac37b GCA_000746585.2_ASM74658v2
Rickettsiales.endosymbiont.of.Stachyamoeba.lipophora GCA_003932735.1_ASM393273v1
99.7/100
GCA_002787615.1_ASM278761v1*
GCA_003531345.1_ASM353134v1*
28.9/92
Ahrensia.kielensis.DSM.5890 GCA_000374465.1_ASM37446v1
15.5/74
Chelativorans.oligotrophicus GCA_011317505.1_ASM1131750v1
100/100
Rhizobium.leguminosarum.bv.trifolii GCA_011604525.1_ASM1160452v1
68.2/94
Bartonella.quintana GCA_009936175.1_ASM993617v1
Pelagibacterium.halotolerans GCA_013004005.1_ASM1300400v1
100/100
100/100
Polymorphum.gilvum.SL003B.26A1 GCA_000192745.1_ASM19274v1
100/100
Roseibium.hamelinense GCA_007830545.1_ASM783054v1
Pseudovibrio.sp.FO.BEG1 GCA_000236645.1_ASM23664v1
100/100
100/100
Methylocella.silvestris.BL2 GCA_000021745.1_ASM2174v1
100/100
Methylocystis.bryophila GCA_002117405.1_ASM211740v1
100/100
Methylorubrum.extorquens.PA1 GCA_000018845.1_ASM1884v1
100/100
Rhodopseudomonas.palustris GCA_013415845.1_ASM1341584v1
97.7/99
Xanthobacter.autotrophicus.Py2 GCA_000017645.1_ASM1764v1
100/100
Hyphomicrobium.denitrificans.ATCC.51888 GCA_000143145.1_ASM14314v1
100/100
Rhodomicrobium.vannielii.ATCC.17100 GCA_000166055.1_ASM16605v1
100/100
Thermopetrobacter.sp.TC1 GCA_000746275.1_ASM74627v1
100/100
Candidatus.Micropelagos.thuwalensis GCA_000469155.1.RS24
100/100
alpha.proteobacterium.IMCC14465 GCA_000293845.2_ASM29384v2
Parvibaculum.lavamentivorans.DS.1 GCA_000017565.1_ASM1756v1
100/100
Asticcacaulis.excentricus.CB.48 GCA_000175215.2_ASM17521v2
100/100
Brevundimonas.subvibrioides.ATCC.15264 GCA_000144605.1_ASM14460v1
100/100
alpha.proteobacterium.L41A GCA_000335735.1.L41A.1.0
100/100
Caulobacter.vibrioides.NA1000 GCA_000022005.1_ASM2200v1
100/100
Phenylobacterium.zucineum.HLK1 GCA_000017265.1_ASM1726v1
100/100
Hirschia.baltica.ATCC.49814 GCA_000023785.1_ASM2378v1
100/100
Hyphomonas.neptunium.ATCC.15444 GCA_000013025.1_ASM1302v1
100/100
Maricaulis.maris GCA_003634045.1_ASM363404v1
Oceanicaulis.alexandrii.DSM.11625 GCA_000420265.1_ASM42026v1
Parvularcula.bermudensis.HTCC2503 GCA_000152825.2_ASM15282v2
23.2/81
Jannaschia.rubra GCA_900113265.1.IMG.taxon.2622736430.annotated.assembly
100/100
99.8/100
Octadecabacter.arcticus.238 GCA_000155735.2_ASM15573v2
100/100
100/100
Ruegeria.pomeroyi.DSS.3 GCA_000011965.2_ASM1196v2
Sagittula.stellata.E.37 GCA_000169415.1_ASM16941v1
100/100
100/100
Ketogulonicigenium.vulgare GCA_001399515.1_ASM139951v1
Rubellimicrobium.rubrum GCA_006152115.1_ASM615211v1
100/100
100/100
Paracoccus.denitrificans GCA_004063735.1_ASM406373v1
Rhodobacter.sphaeroides.ATCC.17029 GCA_000015985.1_ASM1598v1
Neomegalonema.perideroedes.DSM.15528 GCA_000374145.1_ASM37414v1
100/100
100/100
Candidatus.Pelagibacter.ubique.HIMB058 GCA_000419545.1_ASM41954v1
100/100
Candidatus.Pelagibacter.ubique.HTCC1062 GCA_000012345.1_ASM1234v1
100/100
alpha.proteobacterium.HIMB5 GCA_000299095.1_ASM29909v1
alpha.proteobacterium.HIMB114 GCA_000163555.2
100/100
100/100
Citromicrobium.sp.JL477 GCA_001304795.1_ASM130479v1
100/100
Sphingopyxis.macrogoltabida GCA_001314325.1_ASM131432v1
100/100
Sphingobium.indicum.B90A GCA_000264945.2_ASM26494v2
100/100
100/100
Sphingomonas.wittichii.RW1 GCA_000016765.1_ASM1676v1
100/100
Zymomonas.mobilis.subsp.mobilis.ATCC.31821 GCA_000007105.1_ASM710v1
alpha.proteobacterium.AAP81b GCA_001295935.1_ASM129593v1
Kordiimonas.gwangyangensis.DSM.19435 GCA_000375545.1_ASM37554v1
Sneathiella.glossodoripedis.JCM.23214 GCA_000616095.1_ASM61609v1
100/100
Geminicoccus.roseus.DSM.18922 GCA_000427665.1_ASM42766v1
75/81
Tistrella.mobilis.KA081020.065 GCA_000264455.2_ASM26445v2
100/100
Caedimonas.varicaedens GCA_001192655.1_ASM119265v1
100/100
Candidatus.Caedibacter.acanthamoebae GCA_000743035.1_ASM74303v1
Candidatus.Nucleicultrix.amoebiphila.FS5 GCA_002117145.1_ASM211714v1
100/100
100/100
Candidatus.Finniella.inopinata GCA_004210305.1_ASM421030v1
100/100
Candidatus.Odyssella.thessalonicensis.L13 GCA_000190415.2_ASM19041v1
99.7/86
11.9/69
Candidatus.Paracaedibacter.acanthamoebae GCA_000742835.1_ASM74283v1
100/100
Candidatus.Hepatobacter.penaei GCA_000742475.1_ASM74247v1
Holospora.undulata.HU1 GCA_000388175.3_ASM38817v3
100/100
Candidatus.Puniceispirillum.marinum.IMCC1322 GCA_000024465.1_ASM2446v1
100/100
100/100
Thalassobaculum.salexigens.DSM.19539 GCA_000423805.1_ASM42380v1
alpha.proteobacterium.BAL199 GCA_000171835.1_ASM17183v1
98.7/60
99.9/100
Oceanibaculum.indicum GCA_003633955.1_ASM363395v1
100/100
Kiloniella.laminariae.DSM.19542 GCA_000374005.1_ASM37400v1
100/100
Rhodovibrio.salinarum.DSM.9154 GCA_000515255.1_ASM51525v1
100/100
Inquilinus.limosus.DSM.16000 GCA_000423185.1_ASM42318v1
100/100
Rhodospirillum.centenum GCA_014197805.1_ASM1419780v1
Micavibrio.aeruginosavorus.ARL.13 GCA_000226315.1_ASM22631v1
Caenispirillum.salinarum.AK4 GCA_000315795.1
100/100
100/100
Pararhodospirillum.photometricum.DSM.122 GCA_000284415.2_ASM28441v2
89.2/60
92.7/100
Rhodospirillum.rubrum.F11 GCA_000225955.1_ASM22595v1
100/100
Roseospirillum.parvum GCA_900100455.1
100/100
Magnetospirillum.magneticum.AMB.1 GCA_000009985.1_ASM998v1
100/100
Terasakiella.pusilla.DSM.6293 GCA_000688235.1_ASM68823v1
100/100
Thalassospira.profundimaris.WP0211 GCA_000300275.1_ASM30027v1
Elioraea.tepidiphila.DSM.17972 GCA_000378465.1_ASM37846v1
99.8/100
Belnapia.moabensis.DSM.16746 GCA_000745835.1_ASM74583v1
100/100
100/100
Rubritepida.flocculans.DSM.14296 GCA_000425365.1_ASM42536v1
Roseomonas.gilardii GCA_001941945.1_ASM194194v1
100/100
Acidiphilium.multivorum.AIU301 GCA_000202835.1_ASM20283v1
87.2/96
Acidisphaera.rubrifaciens.HS.AP3 GCA_000964365.1_ASM96436v1
69.6/93
Granulibacter.bethesdensis.CGDNIH4 GCA_000576145.2_ASM57614v2
100/100
Commensalibacter.intestini.A911 GCA_000231445.2_ASM23144v1
100/100
Gluconacetobacter.diazotrophicus.PA1.5 GCA_000067045.1_ASM6704v1
74.8/93
Acetobacter.aceti.NBRC.14818 GCA_000193495.2_ASM19349v2
Gluconobacter.oxydans.DSM.3504 GCA_000583855.1_ASM58385v1
100/100
100/100
96/97
100/100
0.448765

### Group H

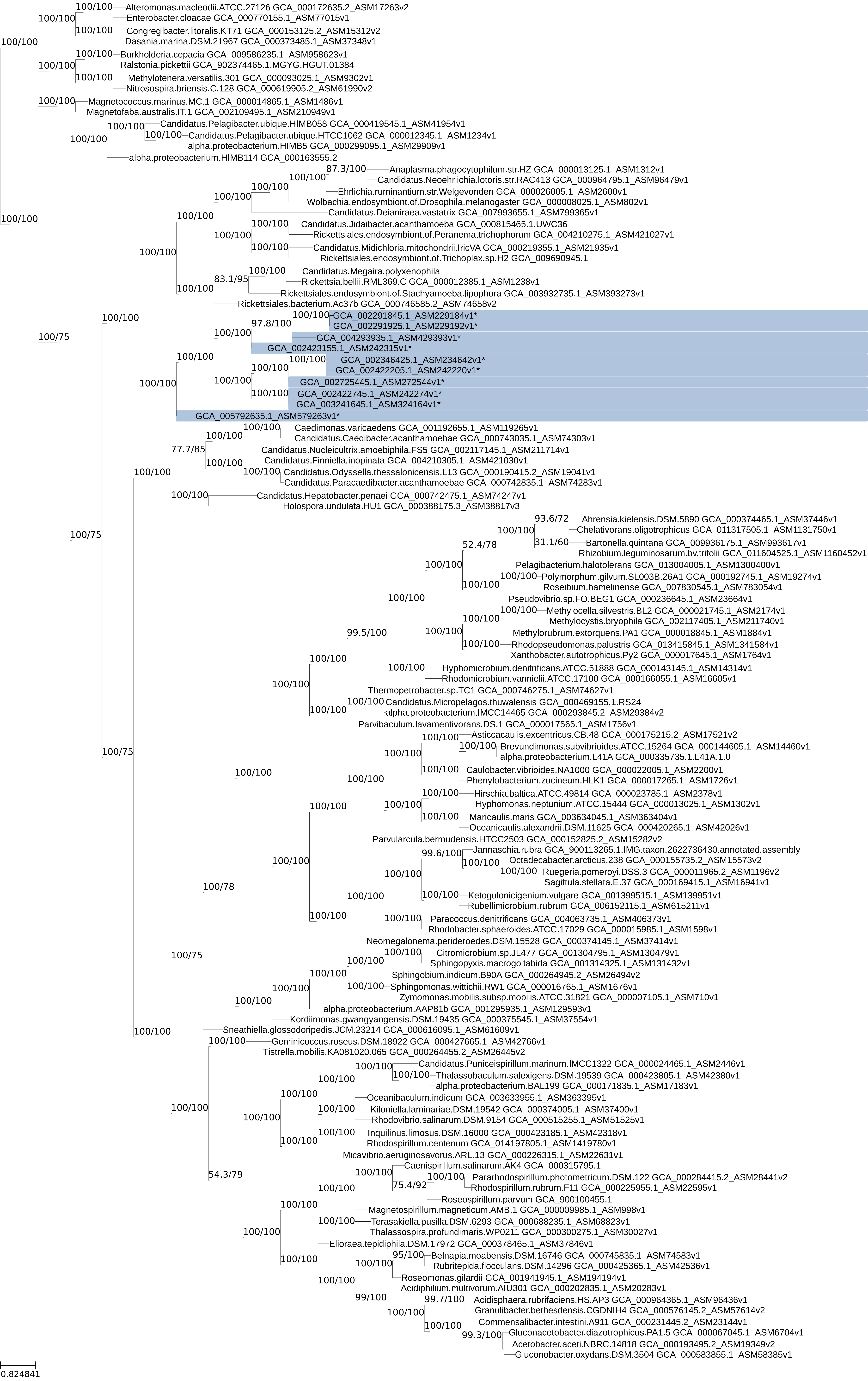

100/100
Alteromonas.macleodii.ATCC.27126 GCA_000172635.2_ASM17263v2
100/100
Enterobacter.cloacae GCA_000770155.1_ASM77015v1
100/100
100/100
Congregibacter.litoralis.KT71 GCA_000153125.2_ASM15312v2
Dasania.marina.DSM.21967 GCA_000373485.1_ASM37348v1
100/100
Burkholderia.cepacia GCA_009586235.1_ASM958623v1
100/100
Ralstonia.pickettii GCA_902374465.1.MGYG.HGUT.01384
100/100
Methylotenera.versatilis.301 GCA_000093025.1_ASM9302v1
Nitrosospira.briensis.C.128 GCA_000619905.2_ASM61990v2
100/100
Magnetococcus.marinus.MC.1 GCA_000014865.1_ASM1486v1
Magnetofaba.australis.IT.1 GCA_002109495.1_ASM210949v1
100/100
Candidatus.Pelagibacter.ubique.HIMB058 GCA_000419545.1_ASM41954v1
100/100
100/100
Candidatus.Pelagibacter.ubique.HTCC1062 GCA_000012345.1_ASM1234v1
alpha.proteobacterium.HIMB5 GCA_000299095.1_ASM29909v1
alpha.proteobacterium.HIMB114 GCA_000163555.2
87.3/100
Anaplasma.phagocytophilum.str.HZ GCA_000013125.1_ASM1312v1
100/100
Candidatus.Neoehrlichia.lotoris.str.RAC413 GCA_000964795.1_ASM96479v1
100/100
Ehrlichia.ruminantium.str.Welgevonden GCA_000026005.1_ASM2600v1
100/100
Wolbachia.endosymbiont.of.Drosophila.melanogaster GCA_000008025.1_ASM802v1
100/100
Candidatus.Deianiraea.vastatrix GCA_007993655.1_ASM799365v1
100/100
Candidatus.Jidaibacter.acanthamoeba GCA_000815465.1.UWC36
100/100
Rickettsiales.endosymbiont.of.Peranema.trichophorum GCA_004210275.1_ASM421027v1
100/100
100/100
Candidatus.Midichloria.mitochondrii.IricVA GCA_000219355.1_ASM21935v1
Rickettsiales.endosymbiont.of.Trichoplax.sp.H2 GCA_009690945.1
100/100
Candidatus.Megaira.polyxenophila
83.1/95
Rickettsia.bellii.RML369.C GCA_000012385.1_ASM1238v1
100/100
Rickettsiales.endosymbiont.of.Stachyamoeba.lipophora GCA_003932735.1_ASM393273v1
Rickettsiales.bacterium.Ac37b GCA_000746585.2_ASM74658v2
100/100
GCA_002291845.1_ASM229184v1*
97.8/100
GCA_002291925.1_ASM229192v1*
100/100
GCA_004293935.1_ASM429393v1*
100/75
GCA_002423155.1_ASM242315v1*
100/100
100/100
GCA_002346425.1_ASM234642v1*
100/100
GCA_002422205.1_ASM242220v1*
100/100
GCA_002725445.1_ASM272544v1*
100/100
100/100
GCA_002422745.1_ASM242274v1*
GCA_003241645.1_ASM324164v1*
GCA_005792635.1_ASM579263v1*
100/100
Caedimonas.varicaedens GCA_001192655.1_ASM119265v1
100/100
Candidatus.Caedibacter.acanthamoebae GCA_000743035.1_ASM74303v1
77.7/85
Candidatus.Nucleicultrix.amoebiphila.FS5 GCA_002117145.1_ASM211714v1
100/100
Candidatus.Finniella.inopinata GCA_004210305.1_ASM421030v1
100/100
100/100
Candidatus.Odyssella.thessalonicensis.L13 GCA_000190415.2_ASM19041v1
Candidatus.Paracaedibacter.acanthamoebae GCA_000742835.1_ASM74283v1
100/100
Candidatus.Hepatobacter.penaei GCA_000742475.1_ASM74247v1
Holospora.undulata.HU1 GCA_000388175.3_ASM38817v3
93.6/72
Ahrensia.kielensis.DSM.5890 GCA_000374465.1_ASM37446v1
100/100
Chelativorans.oligotrophicus GCA_011317505.1_ASM1131750v1
100/75
52.4/78
31.1/60
Bartonella.quintana GCA_009936175.1_ASM993617v1
Rhizobium.leguminosarum.bv.trifolii GCA_011604525.1_ASM1160452v1
100/100
Pelagibacterium.halotolerans GCA_013004005.1_ASM1300400v1
100/100
Polymorphum.gilvum.SL003B.26A1 GCA_000192745.1_ASM19274v1
100/100
Roseibium.hamelinense GCA_007830545.1_ASM783054v1
100/100
Pseudovibrio.sp.FO.BEG1 GCA_000236645.1_ASM23664v1
100/100
Methylocella.silvestris.BL2 GCA_000021745.1_ASM2174v1
100/100
Methylocystis.bryophila GCA_002117405.1_ASM211740v1
99.5/100
100/100
Methylorubrum.extorquens.PA1 GCA_000018845.1_ASM1884v1
100/100
Rhodopseudomonas.palustris GCA_013415845.1_ASM1341584v1
Xanthobacter.autotrophicus.Py2 GCA_000017645.1_ASM1764v1
100/100
100/100
Hyphomicrobium.denitrificans.ATCC.51888 GCA_000143145.1_ASM14314v1
Rhodomicrobium.vannielii.ATCC.17100 GCA_000166055.1_ASM16605v1
100/100
Thermopetrobacter.sp.TC1 GCA_000746275.1_ASM74627v1
100/100
Candidatus.Micropelagos.thuwalensis GCA_000469155.1.RS24
100/100
alpha.proteobacterium.IMCC14465 GCA_000293845.2_ASM29384v2
Parvibaculum.lavamentivorans.DS.1 GCA_000017565.1_ASM1756v1
100/100
Asticcacaulis.excentricus.CB.48 GCA_000175215.2_ASM17521v2
100/100
Brevundimonas.subvibrioides.ATCC.15264 GCA_000144605.1_ASM14460v1
100/100
alpha.proteobacterium.L41A GCA_000335735.1.L41A.1.0
100/100
Caulobacter.vibrioides.NA1000 GCA_000022005.1_ASM2200v1
100/100
100/100
Phenylobacterium.zucineum.HLK1 GCA_000017265.1_ASM1726v1
100/100
Hirschia.baltica.ATCC.49814 GCA_000023785.1_ASM2378v1
100/100
Hyphomonas.neptunium.ATCC.15444 GCA_000013025.1_ASM1302v1
100/100
Maricaulis.maris GCA_003634045.1_ASM363404v1
Oceanicaulis.alexandrii.DSM.11625 GCA_000420265.1_ASM42026v1
Parvularcula.bermudensis.HTCC2503 GCA_000152825.2_ASM15282v2
99.6/100
Jannaschia.rubra GCA_900113265.1.IMG.taxon.2622736430.annotated.assembly
100/100
100/100
Octadecabacter.arcticus.238 GCA_000155735.2_ASM15573v2
100/100
Ruegeria.pomeroyi.DSS.3 GCA_000011965.2_ASM1196v2
Sagittula.stellata.E.37 GCA_000169415.1_ASM16941v1
100/78
100/100
100/100
Ketogulonicigenium.vulgare GCA_001399515.1_ASM139951v1
Rubellimicrobium.rubrum GCA_006152115.1_ASM615211v1
100/100
100/100
Paracoccus.denitrificans GCA_004063735.1_ASM406373v1
Rhodobacter.sphaeroides.ATCC.17029 GCA_000015985.1_ASM1598v1
Neomegalonema.perideroedes.DSM.15528 GCA_000374145.1_ASM37414v1
100/75
100/100
Citromicrobium.sp.JL477 GCA_001304795.1_ASM130479v1
100/100
Sphingopyxis.macrogoltabida GCA_001314325.1_ASM131432v1
100/100
Sphingobium.indicum.B90A GCA_000264945.2_ASM26494v2
100/100
100/100
Sphingomonas.wittichii.RW1 GCA_000016765.1_ASM1676v1
Zymomonas.mobilis.subsp.mobilis.ATCC.31821 GCA_000007105.1_ASM710v1
100/100
alpha.proteobacterium.AAP81b GCA_001295935.1_ASM129593v1
Kordiimonas.gwangyangensis.DSM.19435 GCA_000375545.1_ASM37554v1
100/100
Sneathiella.glossodoripedis.JCM.23214 GCA_000616095.1_ASM61609v1
100/100
Geminicoccus.roseus.DSM.18922 GCA_000427665.1_ASM42766v1
Tistrella.mobilis.KA081020.065 GCA_000264455.2_ASM26445v2
100/100
Candidatus.Puniceispirillum.marinum.IMCC1322 GCA_000024465.1_ASM2446v1
100/100
100/100
Thalassobaculum.salexigens.DSM.19539 GCA_000423805.1_ASM42380v1
alpha.proteobacterium.BAL199 GCA_000171835.1_ASM17183v1
100/100
Oceanibaculum.indicum GCA_003633955.1_ASM363395v1
100/100
Kiloniella.laminariae.DSM.19542 GCA_000374005.1_ASM37400v1
100/100
Rhodovibrio.salinarum.DSM.9154 GCA_000515255.1_ASM51525v1
100/100
100/100
Inquilinus.limosus.DSM.16000 GCA_000423185.1_ASM42318v1
100/100
Rhodospirillum.centenum GCA_014197805.1_ASM1419780v1
Micavibrio.aeruginosavorus.ARL.13 GCA_000226315.1_ASM22631v1
100/100
Caenispirillum.salinarum.AK4 GCA_000315795.1
54.3/79
100/100
100/100
Pararhodospirillum.photometricum.DSM.122 GCA_000284415.2_ASM28441v2
75.4/92
Rhodospirillum.rubrum.F11 GCA_000225955.1_ASM22595v1
Roseospirillum.parvum GCA_900100455.1
100/100
Magnetospirillum.magneticum.AMB.1 GCA_000009985.1_ASM998v1
100/100
Terasakiella.pusilla.DSM.6293 GCA_000688235.1_ASM68823v1
100/100
Thalassospira.profundimaris.WP0211 GCA_000300275.1_ASM30027v1
Elioraea.tepidiphila.DSM.17972 GCA_000378465.1_ASM37846v1
95/100
Belnapia.moabensis.DSM.16746 GCA_000745835.1_ASM74583v1
100/100
100/100
Rubritepida.flocculans.DSM.14296 GCA_000425365.1_ASM42536v1
100/100
Roseomonas.gilardii GCA_001941945.1_ASM194194v1
Acidiphilium.multivorum.AIU301 GCA_000202835.1_ASM20283v1
99/100
99.7/100
Acidisphaera.rubrifaciens.HS.AP3 GCA_000964365.1_ASM96436v1
100/100
Granulibacter.bethesdensis.CGDNIH4 GCA_000576145.2_ASM57614v2
Commensalibacter.intestini.A911 GCA_000231445.2_ASM23144v1
100/100
Gluconacetobacter.diazotrophicus.PA1.5 GCA_000067045.1_ASM6704v1
99.3/100
Acetobacter.aceti.NBRC.14818 GCA_000193495.2_ASM19349v2
Gluconobacter.oxydans.DSM.3504 GCA_000583855.1_ASM58385v1
0.824841

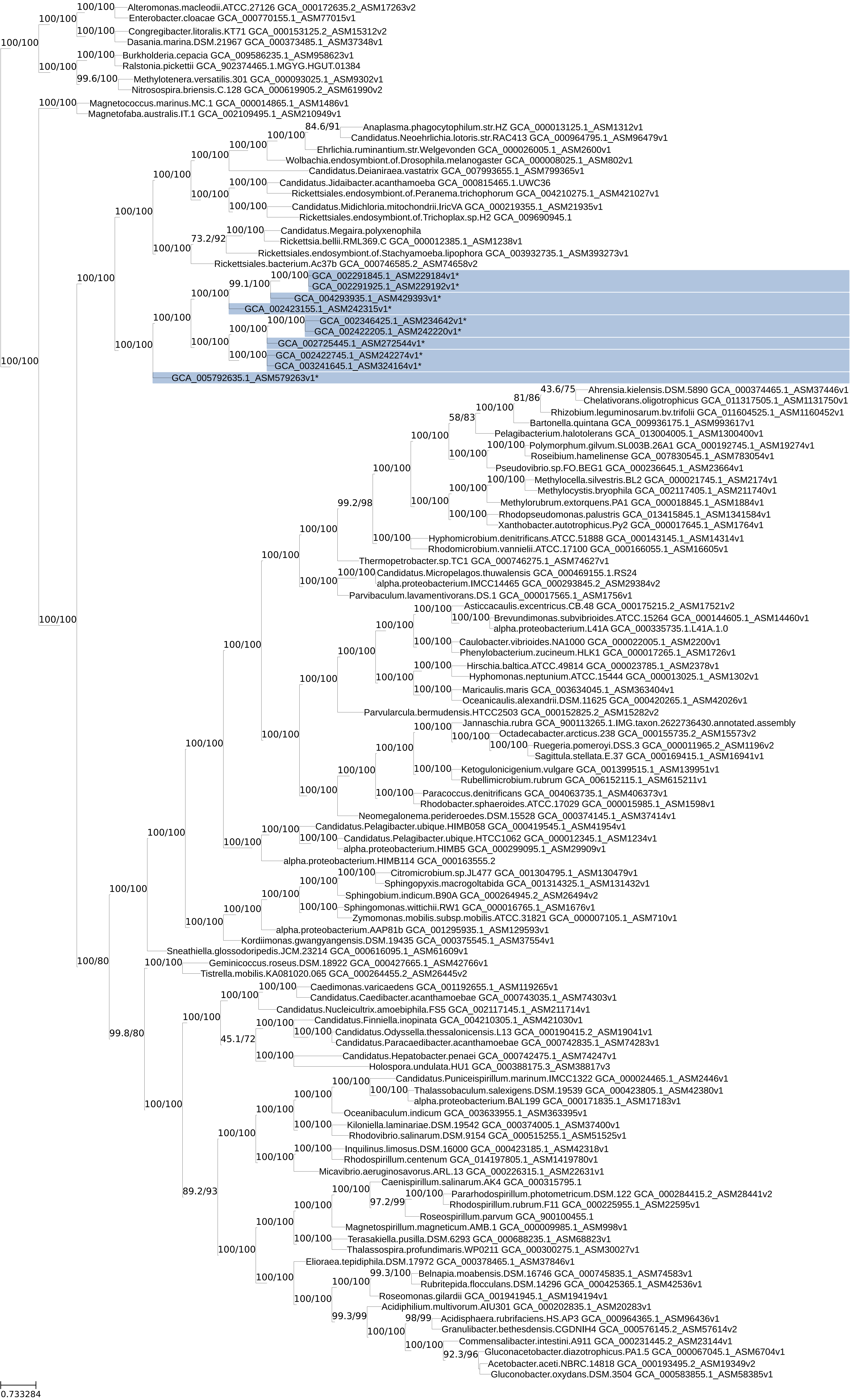
Alteromonas.macleodii.ATCC.27126 GCA_000172635.2_ASM17263v2
Enterobacter.cloacae GCA_000770155.1_ASM77015v1
Congregibacter.litoralis.KT71 GCA_000153125.2_ASM15312v2
Dasania.marina.DSM.21967 GCA_000373485.1_ASM37348v1
Burkholderia.cepacia GCA_009586235.1_ASM958623v1
Ralstonia.pickettii GCA_902374465.1.MGYG.HGUT.01384
Methylotenera.versatilis.301 GCA_000093025.1_ASM9302v1
Nitrosospira.briensis.C.128 GCA_000619905.2_ASM61990v2
Magnetococcus.marinus.MC.1 GCA_000014865.1_ASM1486v1
Magnetofaba.australis.IT.1 GCA_002109495.1_ASM210949v1
Anaplasma.phagocytophilum.str.HZ GCA_000013125.1_ASM1312v1
Candidatus.Neoehrlichia.lotoris.str.RAC413 GCA_000964795.1_ASM96479v1
Ehrlichia.ruminantium.str.Welgevonden GCA_000026005.1_ASM2600v1
Wolbachia.endosymbiont.of.Drosophila.melanogaster GCA_000008025.1_ASM802v1
Candidatus.Deianiraea.vastatrix GCA_007993655.1_ASM799365v1
Candidatus.Jidaibacter.acanthamoeba GCA_000815465.1.UWC36
Rickettsiales.endosymbiont.of.Peranema.trichophorum GCA_004210275.1_ASM421027v1
Candidatus.Midichloria.mitochondrii.IricVA GCA_000219355.1_ASM21935v1
Rickettsiales.endosymbiont.of.Trichoplax.sp.H2 GCA_009690945.1
Candidatus.Megaira.polyxenophila
Rickettsia.bellii.RML369.C GCA_000012385.1_ASM1238v1
Rickettsiales.endosymbiont.of.Stachyamoeba.lipophora GCA_003932735.1_ASM393273v1
Rickettsiales.bacterium.Ac37b GCA_000746585.2_ASM74658v2
GCA_002291845.1_ASM229184v1*
GCA_002291925.1_ASM229192v1*
GCA_004293935.1_ASM429393v1*
GCA_002423155.1_ASM242315v1*
GCA_002346425.1_ASM234642v1*
GCA_002422205.1_ASM242220v1*
GCA_002725445.1_ASM272544v1*
GCA_002422745.1_ASM242274v1*
GCA_003241645.1_ASM324164v1*
GCA_005792635.1_ASM579263v1*
Ahrensia.kielensis.DSM.5890 GCA_000374465.1_ASM37446v1
Chelativorans.oligotrophicus GCA_011317505.1_ASM1131750v1
Rhizobium.leguminosarum.bv.trifolii GCA_011604525.1_ASM1160452v1
Bartonella.quintana GCA_009936175.1_ASM993617v1
Pelagibacterium.halotolerans GCA_013004005.1_ASM1300400v1
Polymorphum.gilvum.SL003B.26A1 GCA_000192745.1_ASM19274v1
Roseibium.hamelinense GCA_007830545.1_ASM783054v1
Pseudovibrio.sp.FO.BEG1 GCA_000236645.1_ASM23664v1
Methylocella.silvestris.BL2 GCA_000021745.1_ASM2174v1
Methylocystis.bryophila GCA_002117405.1_ASM211740v1
Methylorubrum.extorquens.PA1 GCA_000018845.1_ASM1884v1
Rhodopseudomonas.palustris GCA_013415845.1_ASM1341584v1
Xanthobacter.autotrophicus.Py2 GCA_000017645.1_ASM1764v1
Hyphomicrobium.denitrificans.ATCC.51888 GCA_000143145.1_ASM14314v1
Rhodomicrobium.vannielii.ATCC.17100 GCA_000166055.1_ASM16605v1
Thermopetrobacter.sp.TC1 GCA_000746275.1_ASM74627v1
Candidatus.Micropelagos.thuwalensis GCA_000469155.1.RS24
alpha.proteobacterium.IMCC14465 GCA_000293845.2_ASM29384v2
Parvibaculum.lavamentivorans.DS.1 GCA_000017565.1_ASM1756v1
Asticcacaulis.excentricus.CB.48 GCA_000175215.2_ASM17521v2
Brevundimonas.subvibrioides.ATCC.15264 GCA_000144605.1_ASM14460v1
alpha.proteobacterium.L41A GCA_000335735.1.L41A.1.0
Caulobacter.vibrioides.NA1000 GCA_000022005.1_ASM2200v1
Phenylobacterium.zucineum.HLK1 GCA_000017265.1_ASM1726v1
Hirschia.baltica.ATCC.49814 GCA_000023785.1_ASM2378v1
Hyphomonas.neptunium.ATCC.15444 GCA_000013025.1_ASM1302v1
Maricaulis.maris GCA_003634045.1_ASM363404v1
Oceanicaulis.alexandrii.DSM.11625 GCA_000420265.1_ASM42026v1
Parvularcula.bermudensis.HTCC2503 GCA_000152825.2_ASM15282v2
Jannaschia.rubra GCA_900113265.1.IMG.taxon.2622736430.annotated.assembly
Octadecabacter.arcticus.238 GCA_000155735.2_ASM15573v2
Ruegeria.pomeroyi.DSS.3 GCA_000011965.2_ASM1196v2
Sagittula.stellata.E.37 GCA_000169415.1_ASM16941v1
Ketogulonicigenium.vulgare GCA_001399515.1_ASM139951v1
Rubellimicrobium.rubrum GCA_006152115.1_ASM615211v1
Paracoccus.denitrificans GCA_004063735.1_ASM406373v1
Rhodobacter.sphaeroides.ATCC.17029 GCA_000015985.1_ASM1598v1
Neomegalonema.perideroedes.DSM.15528 GCA_000374145.1_ASM37414v1
Candidatus.Pelagibacter.ubique.HIMB058 GCA_000419545.1_ASM41954v1
Candidatus.Pelagibacter.ubique.HTCC1062 GCA_000012345.1_ASM1234v1
alpha.proteobacterium.HIMB5 GCA_000299095.1_ASM29909v1
alpha.proteobacterium.HIMB114 GCA_000163555.2
Citromicrobium.sp.JL477 GCA_001304795.1_ASM130479v1
Sphingopyxis.macrogoltabida GCA_001314325.1_ASM131432v1
Sphingobium.indicum.B90A GCA_000264945.2_ASM26494v2
Sphingomonas.wittichii.RW1 GCA_000016765.1_ASM1676v1
Zymomonas.mobilis.subsp.mobilis.ATCC.31821 GCA_000007105.1_ASM710v1
alpha.proteobacterium.AAP81b GCA_001295935.1_ASM129593v1
Kordiimonas.gwangyangensis.DSM.19435 GCA_000375545.1_ASM37554v1
Sneathiella.glossodoripedis.JCM.23214 GCA_000616095.1_ASM61609v1
Geminicoccus.roseus.DSM.18922 GCA_000427665.1_ASM42766v1
Tistrella.mobilis.KA081020.065 GCA_000264455.2_ASM26445v2
Caedimonas.varicaedens GCA_001192655.1_ASM119265v1
Candidatus.Caedibacter.acanthamoebae GCA_000743035.1_ASM74303v1
Candidatus.Nucleicultrix.amoebiphila.FS5 GCA_002117145.1_ASM211714v1
Candidatus.Finniella.inopinata GCA_004210305.1_ASM421030v1
Candidatus.Odyssella.thessalonicensis.L13 GCA_000190415.2_ASM19041v1
Candidatus.Paracaedibacter.acanthamoebae GCA_000742835.1_ASM74283v1
Candidatus.Hepatobacter.penaei GCA_000742475.1_ASM74247v1
Holospora.undulata.HU1 GCA_000388175.3_ASM38817v3
Candidatus.Puniceispirillum.marinum.IMCC1322 GCA_000024465.1_ASM2446v1
Thalassobaculum.salexigens.DSM.19539 GCA_000423805.1_ASM42380v1
alpha.proteobacterium.BAL199 GCA_000171835.1_ASM17183v1
Oceanibaculum.indicum GCA_003633955.1_ASM363395v1
Kiloniella.laminariae.DSM.19542 GCA_000374005.1_ASM37400v1
Rhodovibrio.salinarum.DSM.9154 GCA_000515255.1_ASM51525v1
Inquilinus.limosus.DSM.16000 GCA_000423185.1_ASM42318v1
Rhodospirillum.centenum GCA_014197805.1_ASM1419780v1
Micavibrio.aeruginosavorus.ARL.13 GCA_000226315.1_ASM22631v1
Caenispirillum.salinarum.AK4 GCA_000315795.1
Pararhodospirillum.photometricum.DSM.122 GCA_000284415.2_ASM28441v2
Rhodospirillum.rubrum.F11 GCA_000225955.1_ASM22595v1
Roseospirillum.parvum GCA_900100455.1
Magnetospirillum.magneticum.AMB.1 GCA_000009985.1_ASM998v1
Terasakiella.pusilla.DSM.6293 GCA_000688235.1_ASM68823v1
Thalassospira.profundimaris.WP0211 GCA_000300275.1_ASM30027v1
Elioraea.tepidiphila.DSM.17972 GCA_000378465.1_ASM37846v1
Belnapia.moabensis.DSM.16746 GCA_000745835.1_ASM74583v1
Rubritepida.flocculans.DSM.14296 GCA_000425365.1_ASM42536v1
Roseomonas.gilardii GCA_001941945.1_ASM194194v1
Acidiphilium.multivorum.AIU301 GCA_000202835.1_ASM20283v1
Acidisphaera.rubrifaciens.HS.AP3 GCA_000964365.1_ASM96436v1
Granulibacter.bethesdensis.CGDNIH4 GCA_000576145.2_ASM57614v2
Commensalibacter.intestini.A911 GCA_000231445.2_ASM23144v1
Gluconacetobacter.diazotrophicus.PA1.5 GCA_000067045.1_ASM6704v1
Acetobacter.aceti.NBRC.14818 GCA_000193495.2_ASM19349v2
Gluconobacter.oxydans.DSM.3504 GCA_000583855.1_ASM58385v1
0.733284

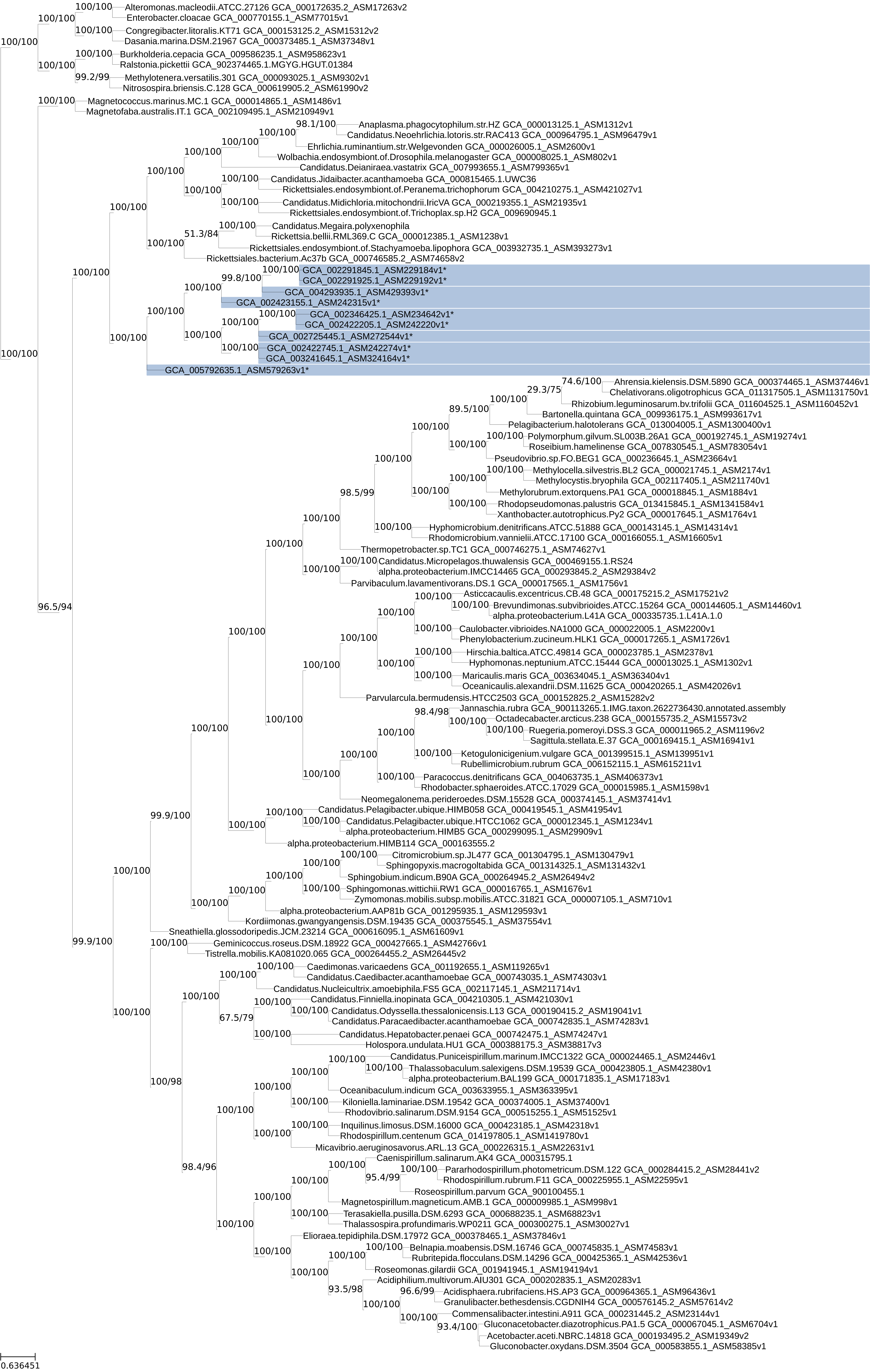

Alteromonas.macleodii.ATCC.27126 GCA_000172635.2_ASM17263v2
Enterobacter.cloacae GCA_000770155.1_ASM77015v1
Congregibacter.litoralis.KT71 GCA_000153125.2_ASM15312v2
Dasania.marina.DSM.21967 GCA_000373485.1_ASM37348v1
Burkholderia.cepacia GCA_009586235.1_ASM958623v1
Ralstonia.pickettii GCA_902374465.1.MGYG.HGUT.01384
Methylotenera.versatilis.301 GCA_000093025.1_ASM9302v1
Nitrosospira.briensis.C.128 GCA_000619905.2_ASM61990v2
Magnetococcus.marinus.MC.1 GCA_000014865.1_ASM1486v1
Magnetofaba.australis.IT.1 GCA_002109495.1_ASM210949v1
Anaplasma.phagocytophilum.str.HZ GCA_000013125.1_ASM1312v1
Candidatus.Neoehrlichia.lotoris.str.RAC413 GCA_000964795.1_ASM96479v1
Ehrlichia.ruminantium.str.Welgevonden GCA_000026005.1_ASM2600v1
Wolbachia.endosymbiont.of.Drosophila.melanogaster GCA_000008025.1_ASM802v1
Candidatus.Deianiraea.vastatrix GCA_007993655.1_ASM799365v1
Candidatus.Jidaibacter.acanthamoeba GCA_000815465.1.UWC36
Rickettsiales.endosymbiont.of.Peranema.trichophorum GCA_004210275.1_ASM421027v1
Candidatus.Midichloria.mitochondrii.IricVA GCA_000219355.1_ASM21935v1
Rickettsiales.endosymbiont.of.Trichoplax.sp.H2 GCA_009690945.1
Candidatus.Megaira.polyxenophila
Rickettsia.bellii.RML369.C GCA_000012385.1_ASM1238v1
Rickettsiales.endosymbiont.of.Stachyamoeba.lipophora GCA_003932735.1_ASM393273v1
Rickettsiales.bacterium.Ac37b GCA_000746585.2_ASM74658v2
GCA_002291845.1_ASM229184v1*
GCA_002291925.1_ASM229192v1*
GCA_004293935.1_ASM429393v1*
GCA_002423155.1_ASM242315v1*
GCA_002346425.1_ASM234642v1*
GCA_002422205.1_ASM242220v1*
GCA_002725445.1_ASM272544v1*
GCA_002422745.1_ASM242274v1*
GCA_003241645.1_ASM324164v1*
GCA_005792635.1_ASM579263v1*
Ahrensia.kielensis.DSM.5890 GCA_000374465.1_ASM37446v1
Chelativorans.oligotrophicus GCA_011317505.1_ASM1131750v1
Rhizobium.leguminosarum.bv.trifolii GCA_011604525.1_ASM1160452v1
Bartonella.quintana GCA_009936175.1_ASM993617v1
Pelagibacterium.halotolerans GCA_013004005.1_ASM1300400v1
Polymorphum.gilvum.SL003B.26A1 GCA_000192745.1_ASM19274v1
Roseibium.hamelinense GCA_007830545.1_ASM783054v1
Pseudovibrio.sp.FO.BEG1 GCA_000236645.1_ASM23664v1
Methylocella.silvestris.BL2 GCA_000021745.1_ASM2174v1
Methylocystis.bryophila GCA_002117405.1_ASM211740v1
Methylorubrum.extorquens.PA1 GCA_000018845.1_ASM1884v1
Rhodopseudomonas.palustris GCA_013415845.1_ASM1341584v1
Xanthobacter.autotrophicus.Py2 GCA_000017645.1_ASM1764v1
Hyphomicrobium.denitrificans.ATCC.51888 GCA_000143145.1_ASM14314v1
Rhodomicrobium.vannielii.ATCC.17100 GCA_000166055.1_ASM16605v1
Thermopetrobacter.sp.TC1 GCA_000746275.1_ASM74627v1
Candidatus.Micropelagos.thuwalensis GCA_000469155.1.RS24
alpha.proteobacterium.IMCC14465 GCA_000293845.2_ASM29384v2
Parvibaculum.lavamentivorans.DS.1 GCA_000017565.1_ASM1756v1
Asticcacaulis.excentricus.CB.48 GCA_000175215.2_ASM17521v2
Brevundimonas.subvibrioides.ATCC.15264 GCA_000144605.1_ASM14460v1
alpha.proteobacterium.L41A GCA_000335735.1.L41A.1.0
Caulobacter.vibrioides.NA1000 GCA_000022005.1_ASM2200v1
Phenylobacterium.zucineum.HLK1 GCA_000017265.1_ASM1726v1
Hirschia.baltica.ATCC.49814 GCA_000023785.1_ASM2378v1
Hyphomonas.neptunium.ATCC.15444 GCA_000013025.1_ASM1302v1
Maricaulis.maris GCA_003634045.1_ASM363404v1
Oceanicaulis.alexandrii.DSM.11625 GCA_000420265.1_ASM42026v1
Parvularcula.bermudensis.HTCC2503 GCA_000152825.2_ASM15282v2
Jannaschia.rubra GCA_900113265.1.IMG.taxon.2622736430.annotated.assembly
Octadecabacter.arcticus.238 GCA_000155735.2_ASM15573v2
Ruegeria.pomeroyi.DSS.3 GCA_000011965.2_ASM1196v2
Sagittula.stellata.E.37 GCA_000169415.1_ASM16941v1
Ketogulonicigenium.vulgare GCA_001399515.1_ASM139951v1
Rubellimicrobium.rubrum GCA_006152115.1_ASM615211v1
Paracoccus.denitrificans GCA_004063735.1_ASM406373v1
Rhodobacter.sphaeroides.ATCC.17029 GCA_000015985.1_ASM1598v1
Neomegalonema.perideroedes.DSM.15528 GCA_000374145.1_ASM37414v1
Candidatus.Pelagibacter.ubique.HIMB058 GCA_000419545.1_ASM41954v1
Candidatus.Pelagibacter.ubique.HTCC1062 GCA_000012345.1_ASM1234v1
alpha.proteobacterium.HIMB5 GCA_000299095.1_ASM29909v1
alpha.proteobacterium.HIMB114 GCA_000163555.2
Citromicrobium.sp.JL477 GCA_001304795.1_ASM130479v1
Sphingopyxis.macrogoltabida GCA_001314325.1_ASM131432v1
Sphingobium.indicum.B90A GCA_000264945.2_ASM26494v2
Sphingomonas.wittichii.RW1 GCA_000016765.1_ASM1676v1
Zymomonas.mobilis.subsp.mobilis.ATCC.31821 GCA_000007105.1_ASM710v1
alpha.proteobacterium.AAP81b GCA_001295935.1_ASM129593v1
Kordiimonas.gwangyangensis.DSM.19435 GCA_000375545.1_ASM37554v1
Sneathiella.glossodoripedis.JCM.23214 GCA_000616095.1_ASM61609v1
Geminicoccus.roseus.DSM.18922 GCA_000427665.1_ASM42766v1
Tistrella.mobilis.KA081020.065 GCA_000264455.2_ASM26445v2
Caedimonas.varicaedens GCA_001192655.1_ASM119265v1
Candidatus.Caedibacter.acanthamoebae GCA_000743035.1_ASM74303v1
Candidatus.Nucleicultrix.amoebiphila.FS5 GCA_002117145.1_ASM211714v1
Candidatus.Finniella.inopinata GCA_004210305.1_ASM421030v1
Candidatus.Odyssella.thessalonicensis.L13 GCA_000190415.2_ASM19041v1
Candidatus.Paracaedibacter.acanthamoebae GCA_000742835.1_ASM74283v1
Candidatus.Hepatobacter.penaei GCA_000742475.1_ASM74247v1
Holospora.undulata.HU1 GCA_000388175.3_ASM38817v3
Candidatus.Puniceispirillum.marinum.IMCC1322 GCA_000024465.1_ASM2446v1
Thalassobaculum.salexigens.DSM.19539 GCA_000423805.1_ASM42380v1
alpha.proteobacterium.BAL199 GCA_000171835.1_ASM17183v1
Oceanibaculum.indicum GCA_003633955.1_ASM363395v1
Kiloniella.laminariae.DSM.19542 GCA_000374005.1_ASM37400v1
Rhodovibrio.salinarum.DSM.9154 GCA_000515255.1_ASM51525v1
Inquilinus.limosus.DSM.16000 GCA_000423185.1_ASM42318v1
Rhodospirillum.centenum GCA_014197805.1_ASM1419780v1
Micavibrio.aeruginosavorus.ARL.13 GCA_000226315.1_ASM22631v1
Caenispirillum.salinarum.AK4 GCA_000315795.1
Pararhodospirillum.photometricum.DSM.122 GCA_000284415.2_ASM28441v2
Rhodospirillum.rubrum.F11 GCA_000225955.1_ASM22595v1
Roseospirillum.parvum GCA_900100455.1
Magnetospirillum.magneticum.AMB.1 GCA_000009985.1_ASM998v1
Terasakiella.pusilla.DSM.6293 GCA_000688235.1_ASM68823v1
Thalassospira.profundimaris.WP0211 GCA_000300275.1_ASM30027v1
Elioraea.tepidiphila.DSM.17972 GCA_000378465.1_ASM37846v1
Belnapia.moabensis.DSM.16746 GCA_000745835.1_ASM74583v1
Rubritepida.flocculans.DSM.14296 GCA_000425365.1_ASM42536v1
Roseomonas.gilardii GCA_001941945.1_ASM194194v1
Acidiphilium.multivorum.AIU301 GCA_000202835.1_ASM20283v1
Acidisphaera.rubrifaciens.HS.AP3 GCA_000964365.1_ASM96436v1
Granulibacter.bethesdensis.CGDNIH4 GCA_000576145.2_ASM57614v2
Commensalibacter.intestini.A911 GCA_000231445.2_ASM23144v1
Gluconacetobacter.diazotrophicus.PA1.5 GCA_000067045.1_ASM6704v1
Acetobacter.aceti.NBRC.14818 GCA_000193495.2_ASM19349v2
Gluconobacter.oxydans.DSM.3504 GCA_000583855.1_ASM58385v1
0.636451

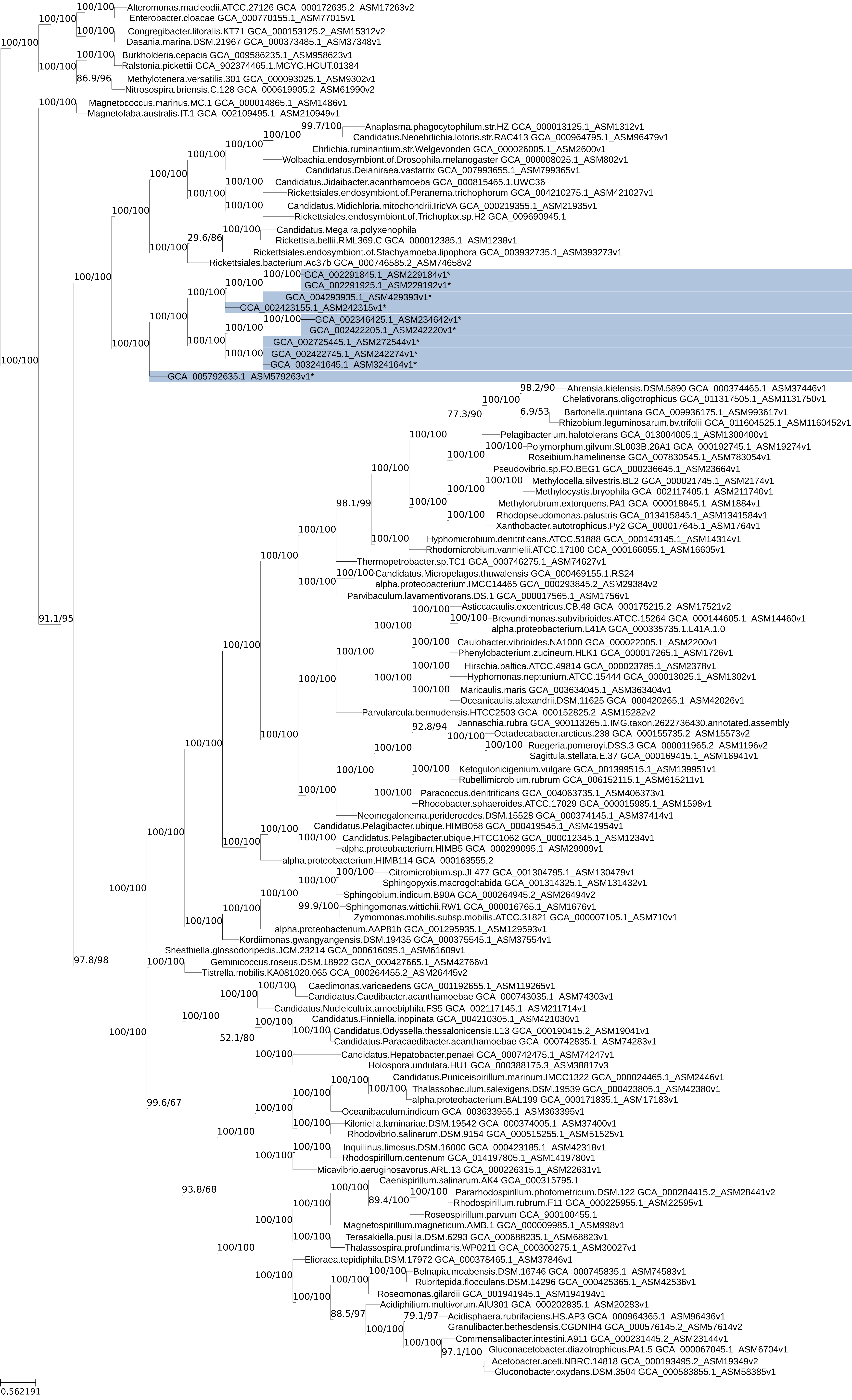

100/100
Alteromonas.macleodii.ATCC.27126 GCA_000172635.2_ASM17263v2
100/100
Enterobacter.cloacae GCA_000770155.1_ASM77015v1
100/100
100/100
Congregibacter.litoralis.KT71 GCA_000153125.2_ASM15312v2
Dasania.marina.DSM.21967 GCA_000373485.1_ASM37348v1
100/100
Burkholderia.cepacia GCA_009586235.1_ASM958623v1
100/100
Ralstonia.pickettii GCA_902374465.1.MGYG.HGUT.01384
86.9/96
Methylotenera.versatilis.301 GCA_000093025.1_ASM9302v1
Nitrosospira.briensis.C.128 GCA_000619905.2_ASM61990v2
100/100
Magnetococcus.marinus.MC.1 GCA_000014865.1_ASM1486v1
Magnetofaba.australis.IT.1 GCA_002109495.1_ASM210949v1
99.7/100
Anaplasma.phagocytophilum.str.HZ GCA_000013125.1_ASM1312v1
100/100
Candidatus.Neoehrlichia.lotoris.str.RAC413 GCA_000964795.1_ASM96479v1
100/100
Ehrlichia.ruminantium.str.Welgevonden GCA_000026005.1_ASM2600v1
100/100
Wolbachia.endosymbiont.of.Drosophila.melanogaster GCA_000008025.1_ASM802v1
Candidatus.Deianiraea.vastatrix GCA_007993655.1_ASM799365v1
100/100
100/100
Candidatus.Jidaibacter.acanthamoeba GCA_000815465.1.UWC36
100/100
Rickettsiales.endosymbiont.of.Peranema.trichophorum GCA_004210275.1_ASM421027v1
100/100
100/100
Candidatus.Midichloria.mitochondrii.IricVA GCA_000219355.1_ASM21935v1
Rickettsiales.endosymbiont.of.Trichoplax.sp.H2 GCA_009690945.1
100/100
Candidatus.Megaira.polyxenophila
29.6/86
Rickettsia.bellii.RML369.C GCA_000012385.1_ASM1238v1
100/100
Rickettsiales.endosymbiont.of.Stachyamoeba.lipophora GCA_003932735.1_ASM393273v1
Rickettsiales.bacterium.Ac37b GCA_000746585.2_ASM74658v2
100/100
100/100
GCA_002291845.1_ASM229184v1*
100/100
GCA_002291925.1_ASM229192v1*
100/100
GCA_004293935.1_ASM429393v1*
GCA_002423155.1_ASM242315v1*
100/100
100/100
GCA_002346425.1_ASM234642v1*
100/100
GCA_002422205.1_ASM242220v1*
100/100
100/100
GCA_002725445.1_ASM272544v1*
100/100
GCA_002422745.1_ASM242274v1*
GCA_003241645.1_ASM324164v1*
GCA_005792635.1_ASM579263v1*
98.2/90
Ahrensia.kielensis.DSM.5890 GCA_000374465.1_ASM37446v1
100/100
Chelativorans.oligotrophicus GCA_011317505.1_ASM1131750v1
77.3/90
6.9/53
Bartonella.quintana GCA_009936175.1_ASM993617v1
Rhizobium.leguminosarum.bv.trifolii GCA_011604525.1_ASM1160452v1
100/100
Pelagibacterium.halotolerans GCA_013004005.1_ASM1300400v1
100/100
Polymorphum.gilvum.SL003B.26A1 GCA_000192745.1_ASM19274v1
100/100
Roseibium.hamelinense GCA_007830545.1_ASM783054v1
100/100
Pseudovibrio.sp.FO.BEG1 GCA_000236645.1_ASM23664v1
100/100
Methylocella.silvestris.BL2 GCA_000021745.1_ASM2174v1
100/100
Methylocystis.bryophila GCA_002117405.1_ASM211740v1
98.1/99
100/100
Methylorubrum.extorquens.PA1 GCA_000018845.1_ASM1884v1
100/100
Rhodopseudomonas.palustris GCA_013415845.1_ASM1341584v1
100/100
Xanthobacter.autotrophicus.Py2 GCA_000017645.1_ASM1764v1
100/100
Hyphomicrobium.denitrificans.ATCC.51888 GCA_000143145.1_ASM14314v1
100/100
Rhodomicrobium.vannielii.ATCC.17100 GCA_000166055.1_ASM16605v1
Thermopetrobacter.sp.TC1 GCA_000746275.1_ASM74627v1
100/100
Candidatus.Micropelagos.thuwalensis GCA_000469155.1.RS24
100/100
alpha.proteobacterium.IMCC14465 GCA_000293845.2_ASM29384v2
Parvibaculum.lavamentivorans.DS.1 GCA_000017565.1_ASM1756v1
100/100
Asticcacaulis.excentricus.CB.48 GCA_000175215.2_ASM17521v2
100/100
100/100
Brevundimonas.subvibrioides.ATCC.15264 GCA_000144605.1_ASM14460v1
alpha.proteobacterium.L41A GCA_000335735.1.L41A.1.0
100/100
100/100
Caulobacter.vibrioides.NA1000 GCA_000022005.1_ASM2200v1
Phenylobacterium.zucineum.HLK1 GCA_000017265.1_ASM1726v1
100/100
100/100
Hirschia.baltica.ATCC.49814 GCA_000023785.1_ASM2378v1
100/100
Hyphomonas.neptunium.ATCC.15444 GCA_000013025.1_ASM1302v1
100/100
Maricaulis.maris GCA_003634045.1_ASM363404v1
Oceanicaulis.alexandrii.DSM.11625 GCA_000420265.1_ASM42026v1
Parvularcula.bermudensis.HTCC2503 GCA_000152825.2_ASM15282v2
92.8/94
Jannaschia.rubra GCA_900113265.1.IMG.taxon.2622736430.annotated.assembly
100/100
100/100
Octadecabacter.arcticus.238 GCA_000155735.2_ASM15573v2
100/100
Ruegeria.pomeroyi.DSS.3 GCA_000011965.2_ASM1196v2
100/100
Sagittula.stellata.E.37 GCA_000169415.1_ASM16941v1
100/100
Ketogulonicigenium.vulgare GCA_001399515.1_ASM139951v1
100/100
Rubellimicrobium.rubrum GCA_006152115.1_ASM615211v1
100/100
Paracoccus.denitrificans GCA_004063735.1_ASM406373v1
Rhodobacter.sphaeroides.ATCC.17029 GCA_000015985.1_ASM1598v1
Neomegalonema.perideroedes.DSM.15528 GCA_000374145.1_ASM37414v1
100/100
Candidatus.Pelagibacter.ubique.HIMB058 GCA_000419545.1_ASM41954v1
100/100
100/100
Candidatus.Pelagibacter.ubique.HTCC1062 GCA_000012345.1_ASM1234v1
alpha.proteobacterium.HIMB5 GCA_000299095.1_ASM29909v1
alpha.proteobacterium.HIMB114 GCA_000163555.2
100/100
Citromicrobium.sp.JL477 GCA_001304795.1_ASM130479v1
100/100
Sphingopyxis.macrogoltabida GCA_001314325.1_ASM131432v1
100/100
Sphingobium.indicum.B90A GCA_000264945.2_ASM26494v2
100/100
Sphingomonas.wittichii.RW1 GCA_000016765.1_ASM1676v1
99.9/100
Zymomonas.mobilis.subsp.mobilis.ATCC.31821 GCA_000007105.1_ASM710v1
100/100
alpha.proteobacterium.AAP81b GCA_001295935.1_ASM129593v1
Kordiimonas.gwangyangensis.DSM.19435 GCA_000375545.1_ASM37554v1
100/100
Sneathiella.glossodoripedis.JCM.23214 GCA_000616095.1_ASM61609v1
100/100
Geminicoccus.roseus.DSM.18922 GCA_000427665.1_ASM42766v1
Tistrella.mobilis.KA081020.065 GCA_000264455.2_ASM26445v2
100/100
Caedimonas.varicaedens GCA_001192655.1_ASM119265v1
100/100
Candidatus.Caedibacter.acanthamoebae GCA_000743035.1_ASM74303v1
100/100
Candidatus.Nucleicultrix.amoebiphila.FS5 GCA_002117145.1_ASM211714v1
100/100
Candidatus.Finniella.inopinata GCA_004210305.1_ASM421030v1
100/100
100/100
Candidatus.Odyssella.thessalonicensis.L13 GCA_000190415.2_ASM19041v1
52.1/80
Candidatus.Paracaedibacter.acanthamoebae GCA_000742835.1_ASM74283v1
100/100
Candidatus.Hepatobacter.penaei GCA_000742475.1_ASM74247v1
Holospora.undulata.HU1 GCA_000388175.3_ASM38817v3
100/100
Candidatus.Puniceispirillum.marinum.IMCC1322 GCA_000024465.1_ASM2446v1
100/100
100/100
Thalassobaculum.salexigens.DSM.19539 GCA_000423805.1_ASM42380v1
alpha.proteobacterium.BAL199 GCA_000171835.1_ASM17183v1
99.6/67
100/100
Oceanibaculum.indicum GCA_003633955.1_ASM363395v1
100/100
Kiloniella.laminariae.DSM.19542 GCA_000374005.1_ASM37400v1
100/100
Rhodovibrio.salinarum.DSM.9154 GCA_000515255.1_ASM51525v1
100/100
Inquilinus.limosus.DSM.16000 GCA_000423185.1_ASM42318v1
100/100
Rhodospirillum.centenum GCA_014197805.1_ASM1419780v1
Micavibrio.aeruginosavorus.ARL.13 GCA_000226315.1_ASM22631v1
100/100
Caenispirillum.salinarum.AK4 GCA_000315795.1
100/100
Pararhodospirillum.photometricum.DSM.122 GCA_000284415.2_ASM28441v2
93.8/68
89.4/100
Rhodospirillum.rubrum.F11 GCA_000225955.1_ASM22595v1
100/100
Roseospirillum.parvum GCA_900100455.1
100/100
Magnetospirillum.magneticum.AMB.1 GCA_000009985.1_ASM998v1
100/100
Terasakiella.pusilla.DSM.6293 GCA_000688235.1_ASM68823v1
100/100
Thalassospira.profundimaris.WP0211 GCA_000300275.1_ASM30027v1
Elioraea.tepidiphila.DSM.17972 GCA_000378465.1_ASM37846v1
100/100
Belnapia.moabensis.DSM.16746 GCA_000745835.1_ASM74583v1
100/100
100/100
Rubritepida.flocculans.DSM.14296 GCA_000425365.1_ASM42536v1
100/100
Roseomonas.gilardii GCA_001941945.1_ASM194194v1
Acidiphilium.multivorum.AIU301 GCA_000202835.1_ASM20283v1
79.1/97
Acidisphaera.rubrifaciens.HS.AP3 GCA_000964365.1_ASM96436v1
88.5/97
100/100
Granulibacter.bethesdensis.CGDNIH4 GCA_000576145.2_ASM57614v2
Commensalibacter.intestini.A911 GCA_000231445.2_ASM23144v1
100/100
97.1/100
Gluconacetobacter.diazotrophicus.PA1.5 GCA_000067045.1_ASM6704v1
Acetobacter.aceti.NBRC.14818 GCA_000193495.2_ASM19349v2
Gluconobacter.oxydans.DSM.3504 GCA_000583855.1_ASM58385v1
100/100
100/100
100/100
97.8/98
91.1/95
100/100
100/100
0.562191

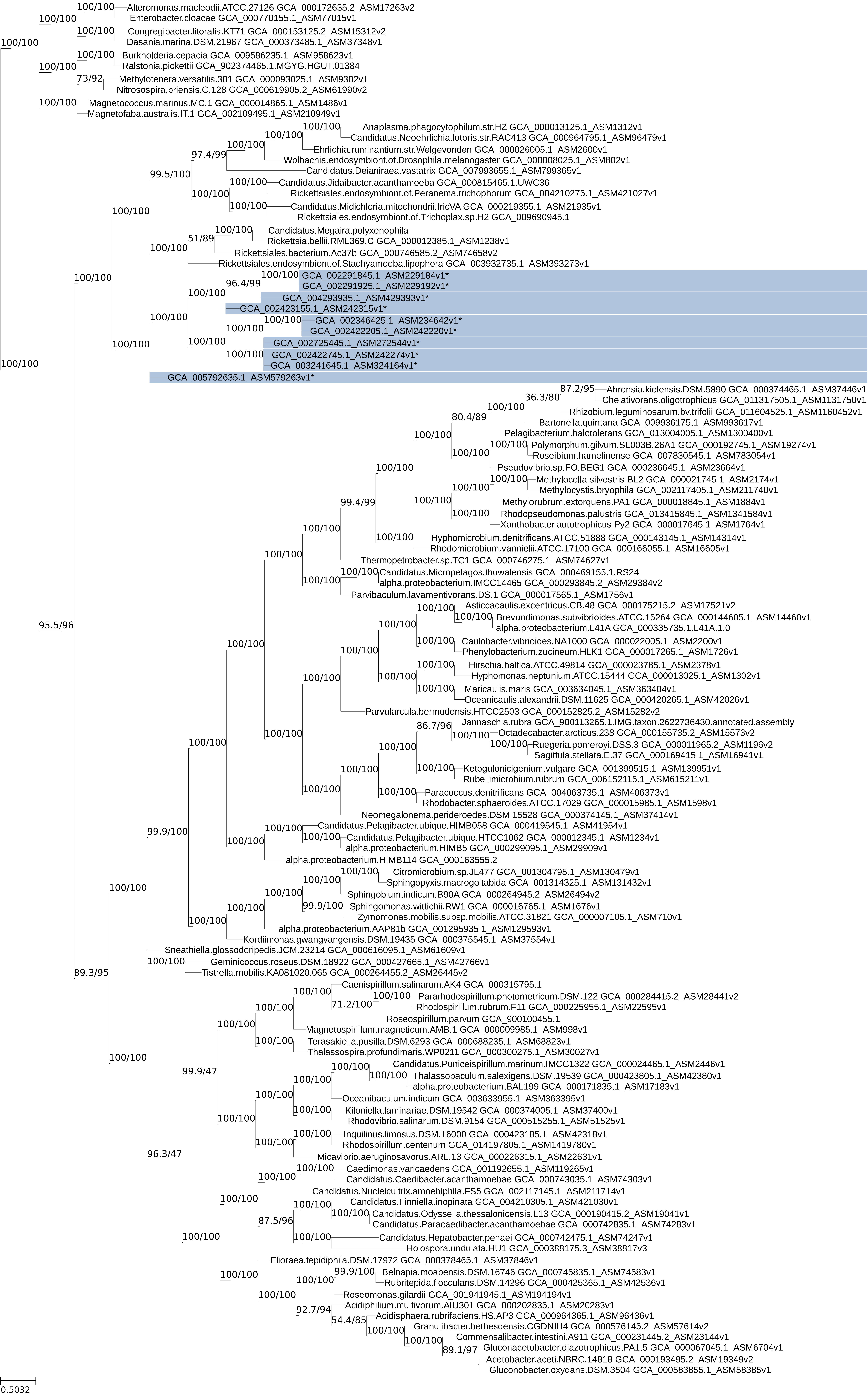

Alteromonas.macleodii.ATCC.27126 GCA_000172635.2_ASM17263v2
Enterobacter.cloacae GCA_000770155.1_ASM77015v1
Congregibacter.litoralis.KT71 GCA_000153125.2_ASM15312v2
Dasania.marina.DSM.21967 GCA_000373485.1_ASM37348v1
Burkholderia.cepacia GCA_009586235.1_ASM958623v1
Ralstonia.pickettii GCA_902374465.1.MGYG.HGUT.01384
Methylotenera.versatilis.301 GCA_000093025.1_ASM9302v1
Nitrosospira.briensis.C.128 GCA_000619905.2_ASM61990v2
Magnetococcus.marinus.MC.1 GCA_000014865.1_ASM1486v1
Magnetofaba.australis.IT.1 GCA_002109495.1_ASM210949v1
Anaplasma.phagocytophilum.str.HZ GCA_000013125.1_ASM1312v1
Candidatus.Neoehrlichia.lotoris.str.RAC413 GCA_000964795.1_ASM96479v1
Ehrlichia.ruminantium.str.Welgevonden GCA_000026005.1_ASM2600v1
Wolbachia.endosymbiont.of.Drosophila.melanogaster GCA_000008025.1_ASM802v1
Candidatus.Deianiraea.vastatrix GCA_007993655.1_ASM799365v1
Candidatus.Jidaibacter.acanthamoeba GCA_000815465.1.UWC36
Rickettsiales.endosymbiont.of.Peranema.trichophorum GCA_004210275.1_ASM421027v1
Candidatus.Midichloria.mitochondrii.IricVA GCA_000219355.1_ASM21935v1
Rickettsiales.endosymbiont.of.Trichoplax.sp.H2 GCA_009690945.1
Candidatus.Megaira.polyxenophila
Rickettsia.bellii.RML369.C GCA_000012385.1_ASM1238v1
Rickettsiales.bacterium.Ac37b GCA_000746585.2_ASM74658v2
Rickettsiales.endosymbiont.of.Stachyamoeba.lipophora GCA_003932735.1_ASM393273v1
GCA_002291845.1_ASM229184v1*
GCA_002291925.1_ASM229192v1*
GCA_004293935.1_ASM429393v1*
GCA_002423155.1_ASM242315v1*
GCA_002346425.1_ASM234642v1*
GCA_002422205.1_ASM242220v1*
GCA_002725445.1_ASM272544v1*
GCA_002422745.1_ASM242274v1*
GCA_003241645.1_ASM324164v1*
GCA_005792635.1_ASM579263v1*
Ahrensia.kielensis.DSM.5890 GCA_000374465.1_ASM37446v1
Chelativorans.oligotrophicus GCA_011317505.1_ASM1131750v1
Rhizobium.leguminosarum.bv.trifolii GCA_011604525.1_ASM1160452v1
Bartonella.quintana GCA_009936175.1_ASM993617v1
Pelagibacterium.halotolerans GCA_013004005.1_ASM1300400v1
Polymorphum.gilvum.SL003B.26A1 GCA_000192745.1_ASM19274v1
Roseibium.hamelinense GCA_007830545.1_ASM783054v1
Pseudovibrio.sp.FO.BEG1 GCA_000236645.1_ASM23664v1
Methylocella.silvestris.BL2 GCA_000021745.1_ASM2174v1
Methylocystis.bryophila GCA_002117405.1_ASM211740v1
Methylorubrum.extorquens.PA1 GCA_000018845.1_ASM1884v1
Rhodopseudomonas.palustris GCA_013415845.1_ASM1341584v1
Xanthobacter.autotrophicus.Py2 GCA_000017645.1_ASM1764v1
Hyphomicrobium.denitrificans.ATCC.51888 GCA_000143145.1_ASM14314v1
Rhodomicrobium.vannielii.ATCC.17100 GCA_000166055.1_ASM16605v1
Thermopetrobacter.sp.TC1 GCA_000746275.1_ASM74627v1
Candidatus.Micropelagos.thuwalensis GCA_000469155.1.RS24
alpha.proteobacterium.IMCC14465 GCA_000293845.2_ASM29384v2
Parvibaculum.lavamentivorans.DS.1 GCA_000017565.1_ASM1756v1
Asticcacaulis.excentricus.CB.48 GCA_000175215.2_ASM17521v2
Brevundimonas.subvibrioides.ATCC.15264 GCA_000144605.1_ASM14460v1
alpha.proteobacterium.L41A GCA_000335735.1.L41A.1.0
Caulobacter.vibrioides.NA1000 GCA_000022005.1_ASM2200v1
Phenylobacterium.zucineum.HLK1 GCA_000017265.1_ASM1726v1
Hirschia.baltica.ATCC.49814 GCA_000023785.1_ASM2378v1
Hyphomonas.neptunium.ATCC.15444 GCA_000013025.1_ASM1302v1
Maricaulis.maris GCA_003634045.1_ASM363404v1
Oceanicaulis.alexandrii.DSM.11625 GCA_000420265.1_ASM42026v1
Parvularcula.bermudensis.HTCC2503 GCA_000152825.2_ASM15282v2
Jannaschia.rubra GCA_900113265.1.IMG.taxon.2622736430.annotated.assembly
Octadecabacter.arcticus.238 GCA_000155735.2_ASM15573v2
Ruegeria.pomeroyi.DSS.3 GCA_000011965.2_ASM1196v2
Sagittula.stellata.E.37 GCA_000169415.1_ASM16941v1
Ketogulonicigenium.vulgare GCA_001399515.1_ASM139951v1
Rubellimicrobium.rubrum GCA_006152115.1_ASM615211v1
Paracoccus.denitrificans GCA_004063735.1_ASM406373v1
Rhodobacter.sphaeroides.ATCC.17029 GCA_000015985.1_ASM1598v1
Neomegalonema.perideroedes.DSM.15528 GCA_000374145.1_ASM37414v1
Candidatus.Pelagibacter.ubique.HIMB058 GCA_000419545.1_ASM41954v1
Candidatus.Pelagibacter.ubique.HTCC1062 GCA_000012345.1_ASM1234v1
alpha.proteobacterium.HIMB5 GCA_000299095.1_ASM29909v1
alpha.proteobacterium.HIMB114 GCA_000163555.2
Citromicrobium.sp.JL477 GCA_001304795.1_ASM130479v1
Sphingopyxis.macrogoltabida GCA_001314325.1_ASM131432v1
Sphingobium.indicum.B90A GCA_000264945.2_ASM26494v2
Sphingomonas.wittichii.RW1 GCA_000016765.1_ASM1676v1
Zymomonas.mobilis.subsp.mobilis.ATCC.31821 GCA_000007105.1_ASM710v1
alpha.proteobacterium.AAP81b GCA_001295935.1_ASM129593v1
Kordiimonas.gwangyangensis.DSM.19435 GCA_000375545.1_ASM37554v1
Sneathiella.glossodoripedis.JCM.23214 GCA_000616095.1_ASM61609v1
Geminicoccus.roseus.DSM.18922 GCA_000427665.1_ASM42766v1
Tistrella.mobilis.KA081020.065 GCA_000264455.2_ASM26445v2
Caenispirillum.salinarum.AK4 GCA_000315795.1
Pararhodospirillum.photometricum.DSM.122 GCA_000284415.2_ASM28441v2
Rhodospirillum.rubrum.F11 GCA_000225955.1_ASM22595v1
Roseospirillum.parvum GCA_900100455.1
Magnetospirillum.magneticum.AMB.1 GCA_000009985.1_ASM998v1
Terasakiella.pusilla.DSM.6293 GCA_000688235.1_ASM68823v1
Thalassospira.profundimaris.WP0211 GCA_000300275.1_ASM30027v1
Candidatus.Puniceispirillum.marinum.IMCC1322 GCA_000024465.1_ASM2446v1
Thalassobaculum.salexigens.DSM.19539 GCA_000423805.1_ASM42380v1
alpha.proteobacterium.BAL199 GCA_000171835.1_ASM17183v1
Oceanibaculum.indicum GCA_003633955.1_ASM363395v1
Kiloniella.laminariae.DSM.19542 GCA_000374005.1_ASM37400v1
Rhodovibrio.salinarum.DSM.9154 GCA_000515255.1_ASM51525v1
Inquilinus.limosus.DSM.16000 GCA_000423185.1_ASM42318v1
Rhodospirillum.centenum GCA_014197805.1_ASM1419780v1
Micavibrio.aeruginosavorus.ARL.13 GCA_000226315.1_ASM22631v1
Caedimonas.varicaedens GCA_001192655.1_ASM119265v1
Candidatus.Caedibacter.acanthamoebae GCA_000743035.1_ASM74303v1
Candidatus.Nucleicultrix.amoebiphila.FS5 GCA_002117145.1_ASM211714v1
Candidatus.Finniella.inopinata GCA_004210305.1_ASM421030v1
Candidatus.Odyssella.thessalonicensis.L13 GCA_000190415.2_ASM19041v2
Candidatus.Paracaedibacter.acanthamoebae GCA_000742835.1_ASM74283v1
Candidatus.Hepatobacter.penaei GCA_000742475.1_ASM74247v1
Holospora.undulata.HU1 GCA_000388175.3_ASM38817v3
Elioraea.tepidiphila.DSM.17972 GCA_000378465.1_ASM37846v1
Belnapia.moabensis.DSM.16746 GCA_000745835.1_ASM74583v1
Rubritepida.flocculans.DSM.14296 GCA_000425365.1_ASM42536v1
Roseomonas.gilardii GCA_001941945.1_ASM194194v1
Acidiphilium.multivorum.AIU301 GCA_000202835.1_ASM20283v1
Acidisphaera.rubrifaciens.HS.AP3 GCA_000964365.1_ASM96436v1
Granulibacter.bethesdensis.CGDNIH4 GCA_000576145.2_ASM57614v2
Commensalibacter.intestini.A911 GCA_000231445.2_ASM23144v1
Gluconacetobacter.diazotrophicus.PA1.5 GCA_000067045.1_ASM6704v1
Acetobacter.aceti.NBRC.14818 GCA_000193495.2_ASM19349v2
Gluconobacter.oxydans.DSM.3504 GCA_000583855.1_ASM58385v1
0.5032

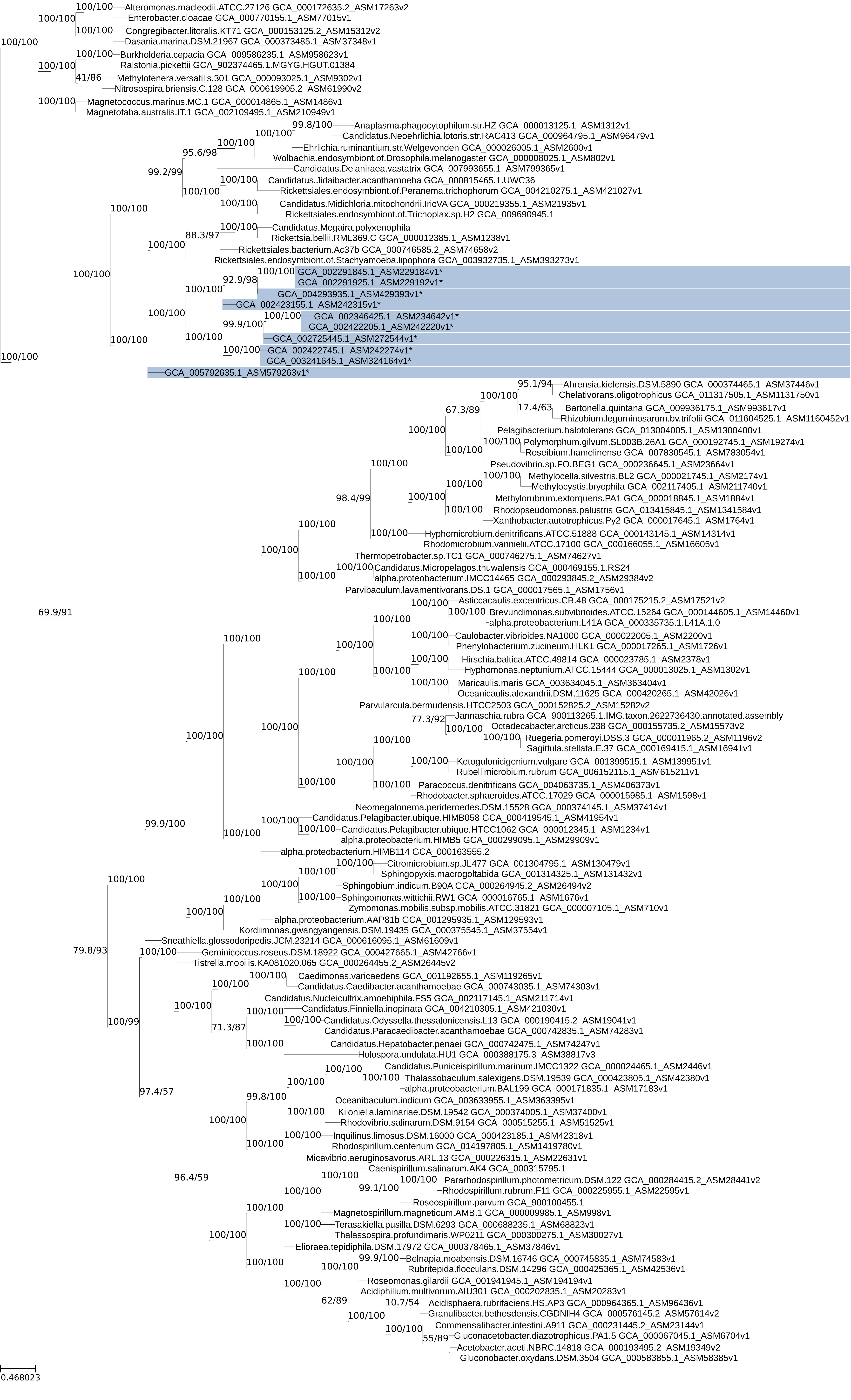
100/100
Alteromonas.macleodii.ATCC.27126 GCA_000172635.2_ASM17263v2
100/100
Enterobacter.cloacae GCA_000770155.1_ASM77015v1
100/100
Congregibacter.litoralis.KT71 GCA_000153125.2_ASM15312v2
100/100
Dasania.marina.DSM.21967 GCA_000373485.1_ASM37348v1
100/100
Burkholderia.cepacia GCA_009586235.1_ASM958623v1
100/100
Ralstonia.pickettii GCA_902374465.1.MGYG.HGUT.01384
41/86
Methylotenera.versatilis.301 GCA_000093025.1_ASM9302v1
Nitrosospira.briensis.C.128 GCA_000619905.2_ASM61990v2
100/100
Magnetococcus.marinus.MC.1 GCA_000014865.1_ASM1486v1
Magnetofaba.australis.IT.1 GCA_002109495.1_ASM210949v1
99.8/100
Anaplasma.phagocytophilum.str.HZ GCA_000013125.1_ASM1312v1
100/100
Candidatus.Neoehrlichia.lotoris.str.RAC413 GCA_000964795.1_ASM96479v1
100/100
Ehrlichia.ruminantium.str.Welgevonden GCA_000026005.1_ASM2600v1
95.6/98
Wolbachia.endosymbiont.of.Drosophila.melanogaster GCA_000008025.1_ASM802v1
99.2/99
Candidatus.Deianiraea.vastatrix GCA_007993655.1_ASM799365v1
100/100
Candidatus.Jidaibacter.acanthamoeba GCA_000815465.1.UWC36
100/100
Rickettsiales.endosymbiont.of.Peranema.trichophorum GCA_004210275.1_ASM421027v1
100/100
Candidatus.Midichloria.mitochondrii.IricVA GCA_000219355.1_ASM21935v1
100/100
Rickettsiales.endosymbiont.of.Trichoplax.sp.H2 GCA_009690945.1
100/100
Candidatus.Megaira.polyxenophila
88.3/97
Rickettsia.bellii.RML369.C GCA_000012385.1_ASM1238v1
100/100
Rickettsiales.bacterium.Ac37b GCA_000746585.2_ASM74658v2
Rickettsiales.endosymbiont.of.Stachyamoeba.lipophora GCA_003932735.1_ASM393273v1
100/100
GCA_002291845.1_ASM229184v1*
100/100
92.9/98
GCA_002291925.1_ASM229192v1*
GCA_004293935.1_ASM429393v1*
100/100
GCA_002423155.1_ASM242315v1*
100/100
GCA_002346425.1_ASM234642v1*
100/100
99.9/100
GCA_002422205.1_ASM242220v1*
100/100
GCA_002725445.1_ASM272544v1*
100/100
100/100
GCA_002422745.1_ASM242274v1*
100/100
GCA_003241645.1_ASM324164v1*
GCA_005792635.1_ASM579263v1*
95.1/94
Ahrensia.kielensis.DSM.5890 GCA_000374465.1_ASM37446v1
100/100
Chelativorans.oligotrophicus GCA_011317505.1_ASM1131750v1
67.3/89
17.4/63
Bartonella.quintana GCA_009936175.1_ASM993617v1
Rhizobium.leguminosarum.bv.trifolii GCA_011604525.1_ASM1160452v1
100/100
Pelagibacterium.halotolerans GCA_013004005.1_ASM1300400v1
100/100
Polymorphum.gilvum.SL003B.26A1 GCA_000192745.1_ASM19274v1
100/100
Roseibium.hamelinense GCA_007830545.1_ASM783054v1
100/100
Pseudovibrio.sp.FO.BEG1 GCA_000236645.1_ASM23664v1
100/100
Methylocella.silvestris.BL2 GCA_000021745.1_ASM2174v1
100/100
Methylocystis.bryophila GCA_002117405.1_ASM211740v1
98.4/99
100/100
Methylorubrum.extorquens.PA1 GCA_000018845.1_ASM1884v1
100/100
Rhodopseudomonas.palustris GCA_013415845.1_ASM1341584v1
Xanthobacter.autotrophicus.Py2 GCA_000017645.1_ASM1764v1
100/100
100/100
Hyphomicrobium.denitrificans.ATCC.51888 GCA_000143145.1_ASM14314v1
Rhodomicrobium.vannielii.ATCC.17100 GCA_000166055.1_ASM16605v1
100/100
Thermopetrobacter.sp.TC1 GCA_000746275.1_ASM74627v1
100/100
Candidatus.Micropelagos.thuwalensis GCA_000469155.1.RS24
100/100
alpha.proteobacterium.IMCC14465 GCA_000293845.2_ASM29384v2
Parvibaculum.lavamentivorans.DS.1 GCA_000017565.1_ASM1756v1
100/100
Asticcacaulis.excentricus.CB.48 GCA_000175215.2_ASM17521v2
100/100
Brevundimonas.subvibrioides.ATCC.15264 GCA_000144605.1_ASM14460v1
69.9/91
100/100
alpha.proteobacterium.L41A GCA_000335735.1.L41A.1.0
100/100
Caulobacter.vibrioides.NA1000 GCA_000022005.1_ASM2200v1
100/100
Phenylobacterium.zucineum.HLK1 GCA_000017265.1_ASM1726v1
100/100
Hirschia.baltica.ATCC.49814 GCA_000023785.1_ASM2378v1
100/100
100/100
Hyphomonas.neptunium.ATCC.15444 GCA_000013025.1_ASM1302v1
100/100
Maricaulis.maris GCA_003634045.1_ASM363404v1
Oceanicaulis.alexandrii.DSM.11625 GCA_000420265.1_ASM42026v1
Parvularcula.bermudensis.HTCC2503 GCA_000152825.2_ASM15282v2
77.3/92
Jannaschia.rubra GCA_900113265.1.IMG.taxon.2622736430.annotated.assembly
100/100
100/100
Octadecabacter.arcticus.238 GCA_000155735.2_ASM15573v2
100/100
100/100
Ruegeria.pomeroyi.DSS.3 GCA_000011965.2_ASM1196v2
Sagittula.stellata.E.37 GCA_000169415.1_ASM16941v1
100/100
100/100
Ketogulonicigenium.vulgare GCA_001399515.1_ASM139951v1
Rubellimicrobium.rubrum GCA_006152115.1_ASM615211v1
100/100
100/100
Paracoccus.denitrificans GCA_004063735.1_ASM406373v1
Rhodobacter.sphaeroides.ATCC.17029 GCA_000015985.1_ASM1598v1
Neomegalonema.perideroedes.DSM.15528 GCA_003741445.1_ASM37414v1
100/100
Candidatus.Pelagibacter.ubique.HIMB058 GCA_000419545.1_ASM41954v1
99.9/100
100/100
100/100
Candidatus.Pelagibacter.ubique.HTCC1062 GCA_000012345.1_ASM1234v1
alpha.proteobacterium.HIMB5 GCA_000299095.1_ASM29909v1
alpha.proteobacterium.HIMB114 GCA_000163555.2
100/100
Citromicrobium.sp.JL477 GCA_001304795.1_ASM130479v1
100/100
100/100
Sphingopyxis.macrogoltabida GCA_001314325.1_ASM131432v1
100/100
Sphingobium.indicum.B90A GCA_000264945.2_ASM26494v2
100/100
100/100
Sphingomonas.wittichii.RW1 GCA_000016765.1_ASM1676v1
100/100
Zymomonas.mobilis.subsp.mobilis.ATCC.31821 GCA_000007105.1_ASM710v1
alpha.proteobacterium.AAP81b GCA_001295935.1_ASM129593v1
Kordiimonas.gwangyangensis.DSM.19435 GCA_000375545.1_ASM37554v1
Sneathiella.glossodoripedis.JCM.23214 GCA_000616095.1_ASM61609v1
79.8/93
100/100
Geminicoccus.roseus.DSM.18922 GCA_000427665.1_ASM42766v1
Tistrella.mobilis.KA081020.065 GCA_000264455.2_ASM26445v2
100/100
Caedimonas.varicaedens GCA_001192655.1_ASM119265v1
100/100
Candidatus.Caedibacter.acanthamoebae GCA_000743035.1_ASM74303v1
100/100
Candidatus.Nucleicultrix.amoebiphila.FS5 GCA_002117145.1_ASM211714v1
100/100
Candidatus.Finniella.inopinata GCA_004210305.1_ASM421030v1
100/99
100/100
Candidatus.Odyssella.thessalonicensis.L13 GCA_000190415.2_ASM19041v1
71.3/87
Candidatus.Paracaedibacter.acanthamoebae GCA_000742835.1_ASM74283v1
100/100
Candidatus.Hepatobacter.penaei GCA_000742475.1_ASM74247v1
Holospora.undulata.HU1 GCA_000388175.3_ASM38817v3
100/100
Candidatus.Puniceispirillum.marinum.IMCC1322 GCA_000024465.1_ASM2446v1
100/100
100/100
Thalassobaculum.salexigens.DSM.19539 GCA_000423805.1_ASM42380v1
97.4/57
alpha.proteobacterium.BAL199 GCA_000171835.1_ASM17183v1
99.8/100
Oceanibaculum.indicum GCA_003633955.1_ASM363395v1
100/100
Kiloniella.laminariae.DSM.19542 GCA_000374005.1_ASM37400v1
100/100
Rhodovibrio.salinarum.DSM.9154 GCA_000515255.1_ASM51525v1
100/100
Inquilinus.limosus.DSM.16000 GCA_000423185.1_ASM42318v1
100/100
Rhodospirillum.centenum GCA_014197805.1_ASM1419780v1
Micavibrio.aeruginosavorus.ARL.13 GCA_000226315.1_ASM22631v1
100/100
Caenispirillum.salinarum.AK4 GCA_000315795.1
96.4/59
100/100
Pararhodospirillum.photometricum.DSM.122 GCA_000284415.2_ASM28441v2
99.1/100
Rhodospirillum.rubrum.F11 GCA_000225955.1_ASM22595v1
100/100
Roseospirillum.parvum GCA_900100455.1
100/100
Magnetospirillum.magneticum.AMB.1 GCA_000009985.1_ASM998v1
100/100
Terasakiella.pusilla.DSM.6293 GCA_000688235.1_ASM68823v1
100/100
Thalassospira.profundimaris.WP0211 GCA_000300275.1_ASM30027v1
Elioraea.tepidiphila.DSM.17972 GCA_000378465.1_ASM37846v1
99.9/100
Belnapia.moabensis.DSM.16746 GCA_000745835.1_ASM74583v1
100/100
100/100
Rubritepida.flocculans.DSM.14296 GCA_000425365.1_ASM42536v1
100/100
Roseomonas.gilardii GCA_001941945.1_ASM194194v1
Acidiphilium.multivorum.AIU301 GCA_000202835.1_ASM20283v1
62/89
10.7/54
Acidisphaera.rubrifaciens.HS.AP3 GCA_000964365.1_ASM96436v1
100/100
Granulibacter.bethesdensis.CGDNIH4 GCA_000576145.2_ASM57614v2
100/100
Commensalibacter.intestini.A911 GCA_000231445.2_ASM23144v1
Gluconacetobacter.diazotrophicus.PA1.5 GCA_000067045.1_ASM6704v1
55/89
Acetobacter.aceti.NBRC.14818 GCA_000193495.2_ASM19349v2
Gluconobacter.oxydans.DSM.3504 GCA_000583855.1_ASM58385v1
0.468023

### Group I

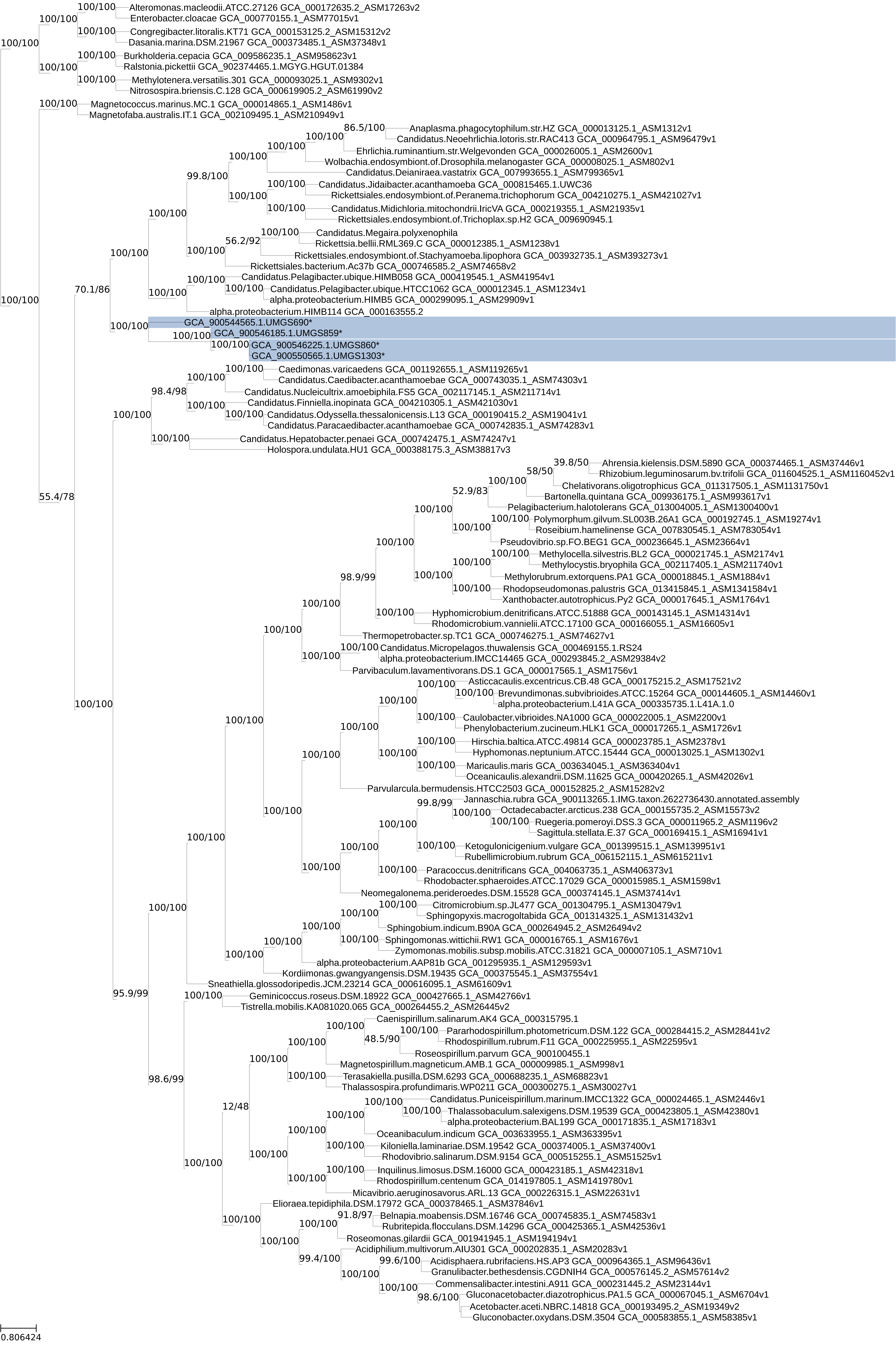
Alteromonas.macleodii.ATCC.27126 GCA_000172635.2_ASM17263v2
Enterobacter.cloacae GCA_000770155.1_ASM77015v1
Congregibacter.litoralis.KT71 GCA_000153125.2_ASM15312v2
Dasania.marina.DSM.21967 GCA_000373485.1_ASM37348v1
Burkholderia.cepacia GCA_009586235.1_ASM958623v1
Ralstonia.pickettii GCA_902374465.1.MGYG.HGUT.01384
Methylotenera.versatilis.301 GCA_000093025.1_ASM9302v1
Nitrosospira.briensis.C.128 GCA_000619905.2_ASM61990v2
Magnetococcus.marinus.MC.1 GCA_000014865.1_ASM1486v1
Magnetofaba.australis.IT.1 GCA_002109495.1_ASM210949v1
Anaplasma.phagocytophilum.str.HZ GCA_000013125.1_ASM1312v1
Candidatus.Neoehrlichia.lotoris.str.RAC413 GCA_000964795.1_ASM96479v1
Ehrlichia.ruminantium.str.Welgevonden GCA_000026005.1_ASM2600v1
Wolbachia.endosymbiont.of.Drosophila.melanogaster GCA_000008025.1_ASM802v1
Candidatus.Deianiraea.vastatrix GCA_007993655.1_ASM799365v1
Candidatus.Jidaibacter.acanthamoeba GCA_000815465.1.UWC36
Rickettsiales.endosymbiont.of.Peranema.trichophorum GCA_004210275.1_ASM421027v1
Candidatus.Midichloria.mitochondrii.IricVA GCA_000219355.1_ASM21935v1
Rickettsiales.endosymbiont.of.Trichoplax.sp.H2 GCA_009690945.1
Candidatus.Megaira.polyxenophila
Rickettsia.bellii.RML369.C GCA_000012385.1_ASM1238v1
Rickettsiales.endosymbiont.of.Stachyamoeba.lipophora GCA_003932735.1_ASM393273v1
Rickettsiales.bacterium.Ac37b GCA_000746585.2_ASM74658v2
Candidatus.Pelagibacter.ubique.HIMB058 GCA_000419545.1_ASM41954v1
Candidatus.Pelagibacter.ubique.HTCC1062 GCA_000012345.1_ASM1234v1
alpha.proteobacterium.HIMB5 GCA_000299095.1_ASM29909v1
alpha.proteobacterium.HIMB114 GCA_000163555.2
GCA_900544565.1.UMGS690*
GCA_900546185.1.UMGS859*
GCA_900546225.1.UMGS860*
GCA_900550565.1.UMGS1303*
Caedimonas.varicaedens GCA_001192655.1_ASM119265v1
Candidatus.Caedibacter.acanthamoebae GCA_000743035.1_ASM74303v1
Candidatus.Nucleicultrix.amoebiphila.FS5 GCA_002117145.1_ASM211714v1
Candidatus.Finniella.inopinata GCA_004210305.1_ASM421030v1
Candidatus.Odyssella.thessalonicensis.L13 GCA_000190415.2_ASM19041v1
Candidatus.Paracaedibacter.acanthamoebae GCA_000742835.1_ASM74283v1
Candidatus.Hepatobacter.penaei GCA_000742475.1_ASM74247v1
Holospora.undulata.HU1 GCA_000388175.3_ASM38817v3
Ahrensia.kielensis.DSM.5890 GCA_000374465.1_ASM37446v1
Rhizobium.leguminosarum.bv.trifolii GCA_011604525.1_ASM1160452v1
Chelativorans.oligotrophicus GCA_011317505.1_ASM1131750v1
Bartonella.quintana GCA_009936175.1_ASM993617v1
Pelagibacterium.halotolerans GCA_013004005.1_ASM1300400v1
Polymorphum.gilvum.SL003B.26A1 GCA_000192745.1_ASM19274v1
Roseibium.hamelinense GCA_007830545.1_ASM783054v1
Pseudovibrio.sp.FO.BEG1 GCA_000236645.1_ASM23664v1
Methylocella.silvestris.BL2 GCA_000021745.1_ASM2174v1
Methylocystis.bryophila GCA_002117405.1_ASM211740v1
Methylorubrum.extorquens.PA1 GCA_000018845.1_ASM1884v1
Rhodopseudomonas.palustris GCA_013415845.1_ASM1341584v1
Xanthobacter.autotrophicus.Py2 GCA_000017645.1_ASM1764v1
Hyphomicrobium.denitrificans.ATCC.51888 GCA_000143145.1_ASM14314v1
Rhodomicrobium.vannielii.ATCC.17100 GCA_000166055.1_ASM16605v1
Thermopetrobacter.sp.TC1 GCA_000746275.1_ASM74627v1
Candidatus.Micropelagos.thuwalensis GCA_000469155.1.RS24
alpha.proteobacterium.IMCC14465 GCA_000293845.2_ASM29384v2
Parvibaculum.lavamentivorans.DS.1 GCA_000017565.1_ASM1756v1
Asticcacaulis.excentricus.CB.48 GCA_000175215.2_ASM17521v2
Brevundimonas.subvibrioides.ATCC.15264 GCA_000144605.1_ASM14460v1
alpha.proteobacterium.L41A GCA_000335735.1.L41A.1.0
Caulobacter.vibrioides.NA1000 GCA_000022005.1_ASM2200v1
Phenylobacterium.zucineum.HLK1 GCA_000017265.1_ASM1726v1
Hirschia.baltica.ATCC.49814 GCA_000023785.1_ASM2378v1
Hyphomonas.neptunium.ATCC.15444 GCA_000013025.1_ASM1302v1
Maricaulis.maris GCA_003634045.1_ASM363404v1
Oceanicaulis.alexandrii.DSM.11625 GCA_000420265.1_ASM42026v1
Parvularcula.bermudensis.HTCC2503 GCA_000152825.2_ASM15282v2
Jannaschia.rubra GCA_900113265.1.IMG.taxon.2622736430.annotated.assembly
Octadecabacter.arcticus.238 GCA_000155735.2_ASM15573v2
Ruegeria.pomeroyi.DSS.3 GCA_000011965.2_ASM1196v2
Sagittula.stellata.E.37 GCA_000169415.1_ASM16941v1
Ketogulonicigenium.vulgare GCA_001399515.1_ASM139951v1
Rubellimicrobium.rubrum GCA_006152115.1_ASM615211v1
Paracoccus.denitrificans GCA_004063735.1_ASM406373v1
Rhodobacter.sphaeroides.ATCC.17029 GCA_000015985.1_ASM1598v1
Neomegalonema.perideroedes.DSM.15528 GCA_000374145.1_ASM37414v1
Citromicrobium.sp.JL477 GCA_001304795.1_ASM130479v1
Sphingopyxis.macrogoltabida GCA_001314325.1_ASM131432v1
Sphingobium.indicum.B90A GCA_000264945.2_ASM26494v2
Sphingomonas.wittichii.RW1 GCA_000016765.1_ASM1676v1
Zymomonas.mobilis.subsp.mobilis.ATCC.31821 GCA_000007105.1_ASM710v1
alpha.proteobacterium.AAP81b GCA_001295935.1_ASM129593v1
Kordiimonas.gwangyangensis.DSM.19435 GCA_000375545.1_ASM37554v1
Sneathiella.glossodoripedis.JCM.23214 GCA_000616095.1_ASM61609v1
Geminicoccus.roseus.DSM.18922 GCA_000427665.1_ASM42766v1
Tistrella.mobilis.KA081020.065 GCA_000264455.2_ASM26445v2
Caenispirillum.salinarum.AK4 GCA_000315795.1
Pararhodospirillum.photometricum.DSM.122 GCA_000284415.2_ASM28441v2
Rhodospirillum.rubrum.F11 GCA_000225955.1_ASM22595v1
Roseospirillum.parvum GCA_900100455.1
Magnetospirillum.magneticum.AMB.1 GCA_000009985.1_ASM998v1
Terasakiella.pusilla.DSM.6293 GCA_000688235.1_ASM68823v1
Thalassospira.profundimaris.WP0211 GCA_000300275.1_ASM30027v1
Candidatus.Puniceispirillum.marinum.IMCC1322 GCA_000024465.1_ASM2446v1
Thalassobaculum.salexigens.DSM.19539 GCA_000423805.1_ASM42380v1
alpha.proteobacterium.BAL199 GCA_000171835.1_ASM17183v1
Oceanibaculum.indicum GCA_003633955.1_ASM363395v1
Kiloniella.laminariae.DSM.19542 GCA_000374005.1_ASM37400v1
Rhodovibrio.salinarum.DSM.9154 GCA_000515255.1_ASM51525v1
Inquilinus.limosus.DSM.16000 GCA_000423185.1_ASM42318v1
Rhodospirillum.centenum GCA_014197805.1_ASM1419780v1
Micavibrio.aeruginosavorus.ARL.13 GCA_000226315.1_ASM22631v1
Elioraea.tepidiphila.DSM.17972 GCA_000378465.1_ASM37846v1
Belnapia.moabensis.DSM.16746 GCA_000745835.1_ASM74583v1
Rubritepida.flocculans.DSM.14296 GCA_000425365.1_ASM42536v1
Roseomonas.gilardii GCA_001941945.1_ASM194194v1
Acidiphilium.multivorum.AIU301 GCA_000202835.1_ASM20283v1
Acidisphaera.rubrifaciens.HS.AP3 GCA_000964365.1_ASM96436v1
Granulibacter.bethesdensis.CGDNIH4 GCA_000576145.2_ASM57614v2
Commensalibacter.intestini.A911 GCA_000231445.2_ASM23144v1
Gluconacetobacter.diazotrophicus.PA1.5 GCA_000067045.1_ASM6704v1
Acetobacter.aceti.NBRC.14818 GCA_000193495.2_ASM19349v2
Gluconobacter.oxydans.DSM.3504 GCA_000583855.1_ASM58385v1
0.806424

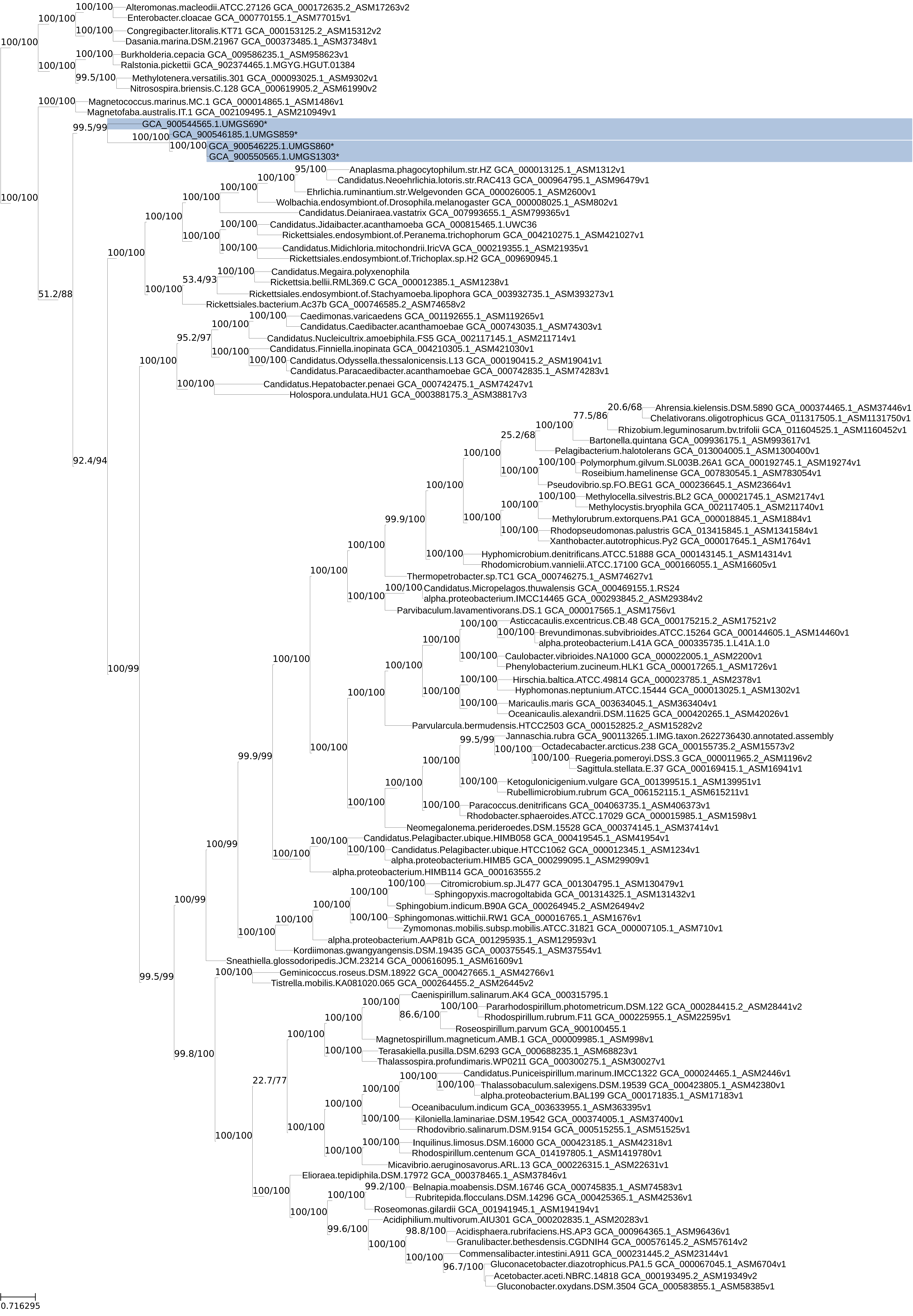

100/100
Alteromonas.macleodii.ATCC.27126 GCA_000172635.2_ASM17263v2
100/100
Enterobacter.cloacae GCA_000770155.1_ASM77015v1
100/100
Congregibacter.litoralis.KT71 GCA_000153125.2_ASM15312v2
100/100
Dasania.marina.DSM.21967 GCA_000373485.1_ASM37348v1
100/100
Burkholderia.cepacia GCA_009586235.1_ASM958623v1
100/100
Ralstonia.pickettii GCA_902374465.1.MGYG.HGUT.01384
99.5/100
Methylotenera.versatilis.301 GCA_000093025.1_ASM9302v1
Nitrosospira.briensis.C.128 GCA_000619905.2_ASM61990v2
100/100
Magnetococcus.marinus.MC.1 GCA_000014865.1_ASM1486v1
Magnetofaba.australis.IT.1 GCA_002109495.1_ASM210949v1
99.5/99
GCA_900544565.1.UMGS690*
100/100
GCA_900546185.1.UMGS859*
100/100
GCA_900546225.1.UMGS860*
GCA_900550565.1.UMGS1303*
95/100
Anaplasma.phagocytophilum.str.HZ GCA_000013125.1_ASM1312v1
100/100
Candidatus.Neoehrlichia.lotoris.str.RAC413 GCA_000964795.1_ASM96479v1
100/100
Ehrlichia.ruminantium.str.Welgevonden GCA_000026005.1_ASM2600v1
100/100
Wolbachia.endosymbiont.of.Drosophila.melanogaster GCA_000008025.1_ASM802v1
100/100
Candidatus.Deianiraea.vastatrix GCA_007993655.1_ASM799365v1
100/100
Candidatus.Jidaibacter.acanthamoeba GCA_000815465.1.UWC36
100/100
Rickettsiales.endosymbiont.of.Peranema.trichophorum GCA_004210275.1_ASM421027v1
100/100
Candidatus.Midichloria.mitochondrii.IricVA GCA_000219355.1_ASM21935v1
100/100
Rickettsiales.endosymbiont.of.Trichoplax.sp.H2 GCA_009690945.1
100/100
Candidatus.Megaira.polyxenophila
53.4/93
Rickettsia.bellii.RML369.C GCA_000012385.1_ASM1238v1
51.2/88
Rickettsiales.endosymbiont.of.Stachyamoeba.lipophora GCA_003932735.1_ASM393273v1
100/100
Rickettsiales.bacterium.Ac37b GCA_000746585.2_ASM74658v2
100/100
Caedimonas.varicaedens GCA_001192655.1_ASM119265v1
100/100
Candidatus.Caedibacter.acanthamoebae GCA_000743035.1_ASM74303v1
95.2/97
Candidatus.Nucleicultrix.amoebiphila.FS5 GCA_002117145.1_ASM211714v1
100/100
Candidatus.Finniella.inopinata GCA_004210305.1_ASM421030v1
100/100
Candidatus.Odyssella.thessalonicensis.L13 GCA_000190415.2_ASM19041v1
100/100
Candidatus.Paracaedibacter.acanthamoebae GCA_000742835.1_ASM74283v1
100/100
Candidatus.Hepatobacter.penaei GCA_000742475.1_ASM74247v1
Holospora.undulata.HU1 GCA_000388175.3_ASM38817v3
20.6/68
Ahrensia.kielensis.DSM.5890 GCA_000374465.1_ASM37446v1
77.5/86
Chelativorans.oligotrophicus GCA_011317505.1_ASM1131750v1
100/100
Rhizobium.leguminosarum.bv.trifolii GCA_011604525.1_ASM1160452v1
25.2/68
Bartonella.quintana GCA_009936175.1_ASM993617v1
Pelagibacterium.halotolerans GCA_013004005.1_ASM1300400v1
100/100
100/100
Polymorphum.gilvum.SL003B.26A1 GCA_000192745.1_ASM19274v1
92.4/94
100/100
Roseibium.hamelinense GCA_007830545.1_ASM783054v1
Pseudovibrio.sp.FO.BEG1 GCA_000236645.1_ASM23664v1
100/100
100/100
Methylocella.silvestris.BL2 GCA_000021745.1_ASM2174v1
100/100
Methylocystis.bryophila GCA_002117405.1_ASM211740v1
99.9/100
100/100
Methylorubrum.extorquens.PA1 GCA_000018845.1_ASM1884v1
100/100
Rhodopseudomonas.palustris GCA_013415845.1_ASM1341584v1
Xanthobacter.autotrophicus.Py2 GCA_000017645.1_ASM1764v1
100/100
100/100
Hyphomicrobium.denitrificans.ATCC.51888 GCA_000143145.1_ASM14314v1
Rhodomicrobium.vannielii.ATCC.17100 GCA_000166055.1_ASM16605v1
100/100
Thermopetrobacter.sp.TC1 GCA_000746275.1_ASM74627v1
100/100
Candidatus.Micropelagos.thuwalensis GCA_000469155.1.RS24
100/100
alpha.proteobacterium.IMCC14465 GCA_000293845.2_ASM29384v2
Parvibaculum.lavamentivorans.DS.1 GCA_000017565.1_ASM1756v1
100/100
Asticcacaulis.excentricus.CB.48 GCA_000175215.2_ASM17521v2
100/100
100/100
Brevundimonas.subvibrioides.ATCC.15264 GCA_000144605.1_ASM14460v1
alpha.proteobacterium.L41A GCA_000335735.1.L41A.1.0
100/100
Caulobacter.vibrioides.NA1000 GCA_000022005.1_ASM2200v1
100/100
100/100
Phenylobacterium.zucineum.HLK1 GCA_000017265.1_ASM1726v1
100/99
100/100
Hirschia.baltica.ATCC.49814 GCA_000023785.1_ASM2378v1
100/100
Hyphomonas.neptunium.ATCC.15444 GCA_000013025.1_ASM1302v1
100/100
100/100
Maricaulis.maris GCA_003634045.1_ASM363404v1
Oceanicaulis.alexandrii.DSM.11625 GCA_000420265.1_ASM42026v1
Parvularcula.bermudensis.HTCC2503 GCA_000152825.2_ASM15282v2
99.5/99
Jannaschia.rubra GCA_900113265.1.IMG.taxon.2622736430.annotated.assembly
100/100
100/100
Octadecabacter.arcticus.238 GCA_000155735.2_ASM15573v2
99.9/99
100/100
Ruegeria.pomeroyi.DSS.3 GCA_000011965.2_ASM1196v2
Sagittula.stellata.E.37 GCA_000169415.1_ASM16941v1
100/100
100/100
Ketogulonicigenium.vulgare GCA_001399515.1_ASM139951v1
Rubellimicrobium.rubrum GCA_006152115.1_ASM615211v1
100/100
100/100
Paracoccus.denitrificans GCA_004063735.1_ASM406373v1
Rhodobacter.sphaeroides.ATCC.17029 GCA_000015985.1_ASM1598v1
Neomegalonema.perideroedes.DSM.15528 GCA_000374145.1_ASM37414v1
100/99
100/100
Candidatus.Pelagibacter.ubique.HIMB058 GCA_000419545.1_ASM41954v1
100/100
100/100
Candidatus.Pelagibacter.ubique.HTCC1062 GCA_000012345.1_ASM1234v1
alpha.proteobacterium.HIMB5 GCA_000299095.1_ASM29909v1
alpha.proteobacterium.HIMB114 GCA_000163555.2
100/100
Citromicrobium.sp.JL477 GCA_001304795.1_ASM130479v1
100/100
Sphingopyxis.macrogoltabida GCA_001314325.1_ASM131432v1
100/99
100/100
Sphingobium.indicum.B90A GCA_000264945.2_ASM26494v2
100/100
100/100
Sphingomonas.wittichii.RW1 GCA_000016765.1_ASM1676v1
100/100
Zymomonas.mobilis.subsp.mobilis.ATCC.31821 GCA_000007105.1_ASM710v1
alpha.proteobacterium.AAP81b GCA_001295935.1_ASM129593v1
Kordiimonas.gwangyangensis.DSM.19435 GCA_000375545.1_ASM37554v1
Sneathiella.glossodoripedis.JCM.23214 GCA_000616095.1_ASM61609v1
100/100
Geminicoccus.roseus.DSM.18922 GCA_000427665.1_ASM42766v1
99.5/99
Tistrella.mobilis.KA081020.065 GCA_000264455.2_ASM26445v2
Caenispirillum.salinarum.AK4 GCA_000315795.1
100/100
100/100
Pararhodospirillum.photometricum.DSM.122 GCA_000284415.2_ASM28441v2
100/100
86.6/100
Rhodospirillum.rubrum.F11 GCA_000225955.1_ASM22595v1
Roseospirillum.parvum GCA_900100455.1
100/100
Magnetospirillum.magneticum.AMB.1 GCA_000009985.1_ASM998v1
99.8/100
100/100
Terasakiella.pusilla.DSM.6293 GCA_000688235.1_ASM68823v1
Thalassospira.profundimaris.WP0211 GCA_000300275.1_ASM30027v1
22.7/77
Candidatus.Puniceispirillum.marinum.IMCC1322 GCA_000024465.1_ASM2446v1
100/100
100/100
Thalassobaculum.salexigens.DSM.19539 GCA_000423805.1_ASM42380v1
100/100
alpha.proteobacterium.BAL199 GCA_000171835.1_ASM17183v1
100/100
Oceanibaculum.indicum GCA_003633955.1_ASM363395v1
100/100
Kiloniella.laminariae.DSM.19542 GCA_000374005.1_ASM37400v1
100/100
Rhodovibrio.salinarum.DSM.9154 GCA_000515255.1_ASM51525v1
100/100
100/100
Inquilinus.limosus.DSM.16000 GCA_000423185.1_ASM42318v1
100/100
Rhodospirillum.centenum GCA_014197805.1_ASM1419780v1
Micavibrio.aeruginosavorus.ARL.13 GCA_000226315.1_ASM22631v1
Elioraea.tepidiphila.DSM.17972 GCA_000378465.1_ASM37846v1
99.2/100
100/100
Belnapia.moabensis.DSM.16746 GCA_000745835.1_ASM74583v1
100/100
Rubritepida.flocculans.DSM.14296 GCA_000425365.1_ASM42536v1
100/100
Roseomonas.gilardii GCA_001941945.1_ASM194194v1
Acidiphilium.multivorum.AIU301 GCA_000202835.1_ASM20283v1
99.6/100
98.8/100
Acidisphaera.rubrifaciens.HS.AP3 GCA_000964365.1_ASM96436v1
100/100
Granulibacter.bethesdensis.CGDNIH4 GCA_000576145.2_ASM57614v2
Commensalibacter.intestini.A911 GCA_000231445.2_ASM23144v1
100/100
Gluconacetobacter.diazotrophicus.PA1.5 GCA_000067045.1_ASM6704v1
96.7/100
Acetobacter.aceti.NBRC.14818 GCA_000193495.2_ASM19349v2
Gluconobacter.oxydans.DSM.3504 GCA_000583855.1_ASM58385v1
100/100
100/100
0.716295

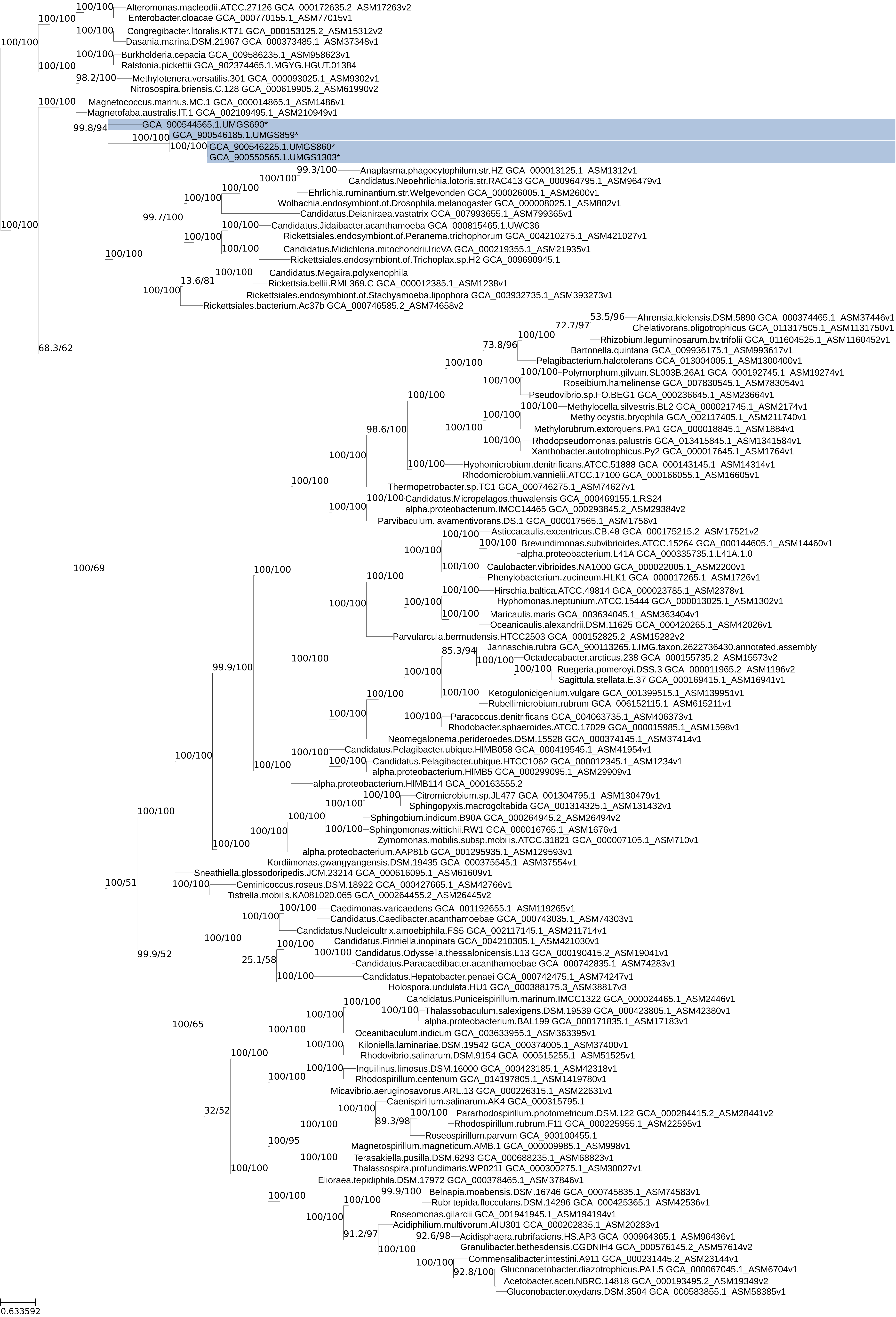
100/100
Alteromonas.macleodii.ATCC.27126 GCA_000172635.2_ASM17263v2
100/100
Enterobacter.cloacae GCA_000770155.1_ASM77015v1
100/100
100/100
Congregibacter.litoralis.KT71 GCA_000153125.2_ASM15312v2
Dasania.marina.DSM.21967 GCA_000373485.1_ASM37348v1
100/100
Burkholderia.cepacia GCA_009586235.1_ASM958623v1
100/100
Ralstonia.pickettii GCA_902374465.1.MGYG.HGUT.01384
98.2/100
Methylotenera.versatilis.301 GCA_000093025.1_ASM9302v1
Nitrosospira.briensis.C.128 GCA_000619905.2_ASM61990v2
100/100
Magnetococcus.marinus.MC.1 GCA_000014865.1_ASM1486v1
Magnetofaba.australis.IT.1 GCA_002109495.1_ASM210949v1
99.8/94
GCA_900544565.1.UMGS690*
100/100
GCA_900546185.1.UMGS859*
100/100
GCA_900546225.1.UMGS860*
GCA_900550565.1.UMGS1303*
99.3/100
Anaplasma.phagocytophilum.str.HZ GCA_000013125.1_ASM1312v1
100/100
Candidatus.Neoehrlichia.lotoris.str.RAC413 GCA_000964795.1_ASM96479v1
100/100
Ehrlichia.ruminantium.str.Welgevonden GCA_000026005.1_ASM2600v1
100/100
Wolbachia.endosymbiont.of.Drosophila.melanogaster GCA_000008025.1_ASM802v1
100/100
Candidatus.Deianiraea.vastatrix GCA_007993655.1_ASM799365v1
99.7/100
100/100
Candidatus.Jidaibacter.acanthamoeba GCA_000815465.1.UWC36
100/100
Rickettsiales.endosymbiont.of.Peranema.trichophorum GCA_004210275.1_ASM421027v1
100/100
100/100
Candidatus.Midichloria.mitochondrii.IricVA GCA_000219355.1_ASM21935v1
Rickettsiales.endosymbiont.of.Trichoplax.sp.H2 GCA_009690945.1
100/100
Candidatus.Megaira.polyxenophila
13.6/81
Rickettsia.bellii.RML369.C GCA_000012385.1_ASM1238v1
100/100
Rickettsiales.endosymbiont.of.Stachyamoeba.lipophora GCA_003932735.1_ASM393273v1
Rickettsiales.bacterium.Ac37b GCA_000746585.2_ASM74658v2
53.5/96
Ahrensia.kielensis.DSM.5890 GCA_000374465.1_ASM37446v1
72.7/97
Chelativorans.oligotrophicus GCA_011317505.1_ASM1131750v1
100/100
Rhizobium.leguminosarum.bv.trifolii GCA_011604525.1_ASM1160452v1
68.3/62
73.8/96
Bartonella.quintana GCA_009936175.1_ASM993617v1
Pelagibacterium.halotolerans GCA_013004005.1_ASM1300400v1
100/100
100/100
Polymorphum.gilvum.SL003B.26A1 GCA_000192745.1_ASM19274v1
100/100
Roseibium.hamelinense GCA_007830545.1_ASM783054v1
100/100
Pseudovibrio.sp.FO.BEG1 GCA_000236645.1_ASM23664v1
100/100
Methylocella.silvestris.BL2 GCA_000021745.1_ASM2174v1
100/100
Methylocystis.bryophila GCA_002117405.1_ASM211740v1
100/100
Methylorubrum.extorquens.PA1 GCA_000018845.1_ASM1884v1
98.6/100
100/100
Rhodopseudomonas.palustris GCA_013415845.1_ASM1341584v1
Xanthobacter.autotrophicus.Py2 GCA_000017645.1_ASM1764v1
100/100
100/100
Hyphomicrobium.denitrificans.ATCC.51888 GCA_000143145.1_ASM14314v1
Rhodomicrobium.vannielii.ATCC.17100 GCA_000166055.1_ASM16605v1
100/100
Thermopetrobacter.sp.TC1 GCA_000746275.1_ASM74627v1
100/100
Candidatus.Micropelagos.thuwalensis GCA_000469155.1.RS24
100/100
alpha.proteobacterium.IMCC14465 GCA_000293845.2_ASM29384v2
Parvibaculum.lavamentivorans.DS.1 GCA_000017565.1_ASM1756v1
100/100
Asticcacaulis.excentricus.CB.48 GCA_000175215.2_ASM17521v2
100/100
100/100
Brevundimonas.subvibrioides.ATCC.15264 GCA_000144605.1_ASM14460v1
alpha.proteobacterium.L41A GCA_000335735.1.L41A.1.0
100/69
100/100
100/100
Caulobacter.vibrioides.NA1000 GCA_000022005.1_ASM2200v1
Phenylobacterium.zucineum.HLK1 GCA_000017265.1_ASM1726v1
100/100
100/100
Hirschia.baltica.ATCC.49814 GCA_000023785.1_ASM2378v1
100/100
Hyphomonas.neptunium.ATCC.15444 GCA_000013025.1_ASM1302v1
100/100
100/100
Maricaulis.maris GCA_003634045.1_ASM363404v1
Oceanicaulis.alexandrii.DSM.11625 GCA_000420265.1_ASM42026v1
Parvularcula.bermudensis.HTCC2503 GCA_000152825.2_ASM15282v2
85.3/94
Jannaschia.rubra GCA_900113265.1.IMG.taxon.2622736430.annotated.assembly
100/100
Octadecabacter.arcticus.238 GCA_000155735.2_ASM15573v2
100/100
100/100
Ruegeria.pomeroyi.DSS.3 GCA_000011965.2_ASM1196v2
99.9/100
Sagittula.stellata.E.37 GCA_000169415.1_ASM16941v1
100/100
100/100
Ketogulonicigenium.vulgare GCA_001399515.1_ASM139951v1
Rubellimicrobium.rubrum GCA_006152115.1_ASM615211v1
100/100
100/100
Paracoccus.denitrificans GCA_004063735.1_ASM406373v1
Rhodobacter.sphaeroides.ATCC.17029 GCA_000015985.1_ASM1598v1
Neomegalonema.perideroedes.DSM.15528 GCA_000374145.1_ASM37414v1
100/100
Candidatus.Pelagibacter.ubique.HIMB058 GCA_000419545.1_ASM41954v1
100/100
100/100
Candidatus.Pelagibacter.ubique.HTCC1062 GCA_000012345.1_ASM1234v1
100/100
alpha.proteobacterium.HIMB5 GCA_000299095.1_ASM29909v1
alpha.proteobacterium.HIMB114 GCA_000163555.2
100/100
Citromicrobium.sp.JL477 GCA_001304795.1_ASM130479v1
100/100
Sphingopyxis.macrogoltabida GCA_001314325.1_ASM131432v1
100/100
100/100
Sphingobium.indicum.B90A GCA_000264945.2_ASM26494v2
100/100
100/100
Sphingomonas.wittichii.RW1 GCA_000016765.1_ASM1676v1
Zymomonas.mobilis.subsp.mobilis.ATCC.31821 GCA_000007105.1_ASM710v1
100/100
alpha.proteobacterium.AAP81b GCA_001295935.1_ASM129593v1
Kordiimonas.gwangyangensis.DSM.19435 GCA_000375545.1_ASM37554v1
Sneathiella.glossodoripedis.JCM.23214 GCA_000616095.1_ASM61609v1
100/51
100/100
Geminicoccus.roseus.DSM.18922 GCA_000427665.1_ASM42766v1
Tistrella.mobilis.KA081020.065 GCA_000264455.2_ASM26445v2
100/100
Caedimonas.varicaedens GCA_001192655.1_ASM119265v1
100/100
Candidatus.Caedibacter.acanthamoebae GCA_000743035.1_ASM74303v1
100/100
Candidatus.Nucleicultrix.amoebiphila.FS5 GCA_002117145.1_ASM211714v1
100/100
Candidatus.Finniella.inopinata GCA_004210305.1_ASM421030v1
99.9/52
100/100
Candidatus.Odyssella.thessalonicensis.L13 GCA_000190415.2_ASM19041v1
25.1/58
Candidatus.Paracaedibacter.acanthamoebae GCA_000742835.1_ASM74283v1
100/100
Candidatus.Hepatobacter.penaei GCA_000742475.1_ASM74247v1
Holospora.undulata.HU1 GCA_000388175.3_ASM38817v3
100/100
Candidatus.Puniceispirillum.marinum.IMCC1322 GCA_000024465.1_ASM2446v1
100/100
100/100
Thalassobaculum.salexigens.DSM.19539 GCA_000423805.1_ASM42380v1
alpha.proteobacterium.BAL199 GCA_000171835.1_ASM17183v1
100/65
100/100
Oceanibaculum.indicum GCA_003633955.1_ASM363395v1
100/100
Kiloniella.laminariae.DSM.19542 GCA_000374005.1_ASM37400v1
100/100
Rhodovibrio.salinarum.DSM.9154 GCA_000515255.1_ASM51525v1
100/100
Inquilinus.limosus.DSM.16000 GCA_000423185.1_ASM42318v1
100/100
Rhodospirillum.centenum GCA_014197805.1_ASM1419780v1
Micavibrio.aeruginosavorus.ARL.13 GCA_000226315.1_ASM22631v1
100/100
Caenispirillum.salinarum.AK4 GCA_000315795.1
32/52
100/100
Pararhodospirillum.photometricum.DSM.122 GCA_000284415.2_ASM28441v2
100/100
89.3/98
Rhodospirillum.rubrum.F11 GCA_000225955.1_ASM22595v1
Roseospirillum.parvum GCA_900100455.1
100/95
Magnetospirillum.magneticum.AMB.1 GCA_000009985.1_ASM998v1
100/100
Terasakiella.pusilla.DSM.6293 GCA_000688235.1_ASM68823v1
100/100
Thalassospira.profundimaris.WP0211 GCA_000300275.1_ASM30027v1
Elioraea.tepidiphila.DSM.17972 GCA_000378465.1_ASM37846v1
99.9/100
Belnapia.moabensis.DSM.16746 GCA_000745835.1_ASM74583v1
100/100
100/100
Rubritepida.flocculans.DSM.14296 GCA_000425365.1_ASM42536v1
Roseomonas.gilardii GCA_001941945.1_ASM194194v1
100/100
Acidiphilium.multivorum.AIU301 GCA_000202835.1_ASM20283v1
91.2/97
92.6/98
Acidisphaera.rubrifaciens.HS.AP3 GCA_000964365.1_ASM96436v1
Granulibacter.bethesdensis.CGDNIH4 GCA_000576145.2_ASM57614v2
100/100
Commensalibacter.intestini.A911 GCA_000231445.2_ASM23144v1
100/100
Gluconacetobacter.diazotrophicus.PA1.5 GCA_000067045.1_ASM6704v1
92.8/100
Acetobacter.aceti.NBRC.14818 GCA_000193495.2_ASM19349v2
Gluconobacter.oxydans.DSM.3504 GCA_000583855.1_ASM58385v1
0.633592

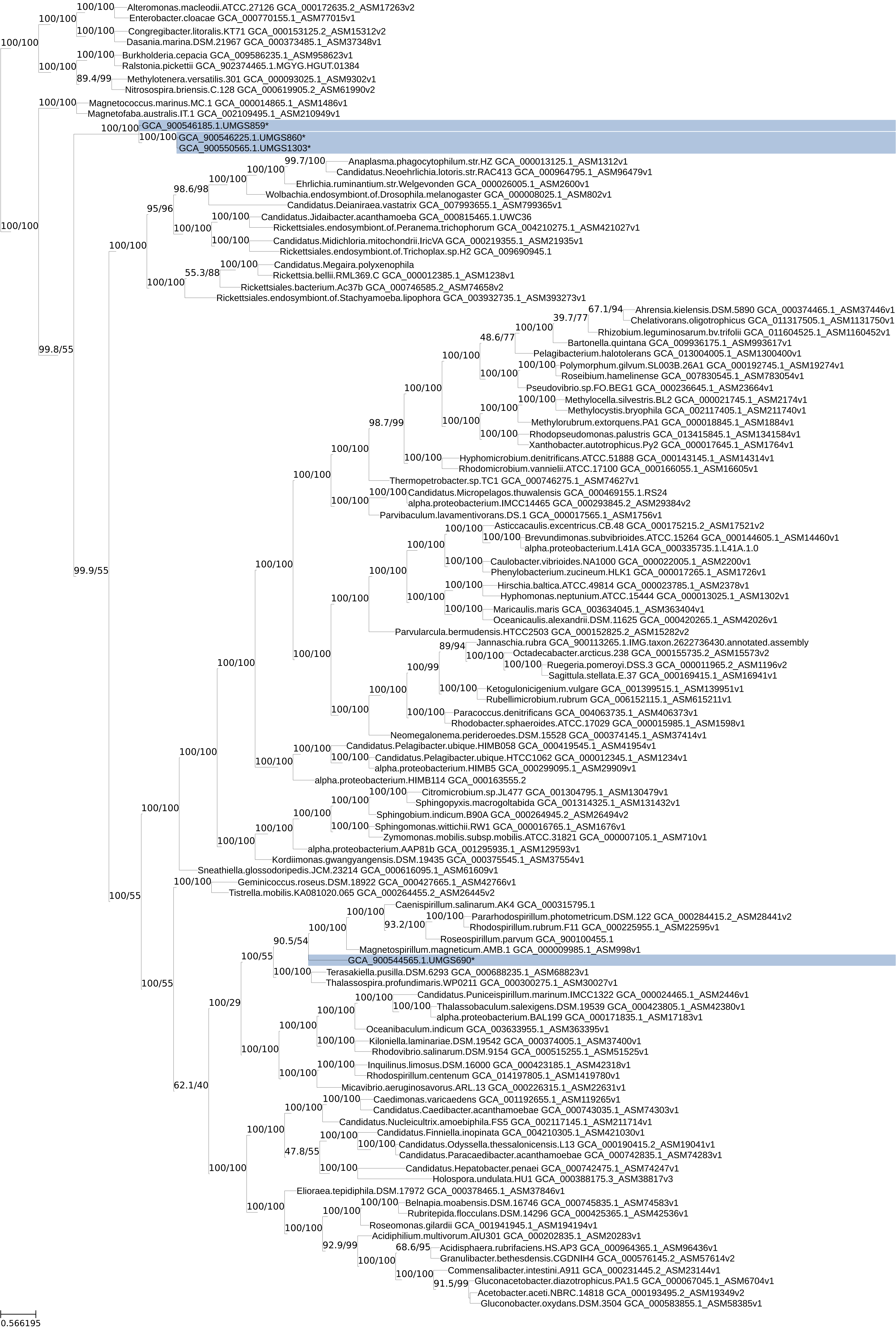

100/100
100/100
Alteromonas.macleodii.ATCC.27126 GCA_000172635.2_ASM17263v2
Enterobacter.cloacae GCA_000770155.1_ASM77015v1
100/100
100/100
Congregibacter.litoralis.KT71 GCA_000153125.2_ASM15312v2
Dasania.marina.DSM.21967 GCA_000373485.1_ASM37348v1
100/100
Burkholderia.cepacia GCA_009586235.1_ASM958623v1
100/100
Ralstonia.pickettii GCA_902374465.1.MGYG.HGUT.01384
89.4/99
Methylotenera.versatilis.301 GCA_000093025.1_ASM9302v1
Nitrosospira.briensis.C.128 GCA_000619905.2_ASM61990v2
100/100
Magnetococcus.marinus.MC.1 GCA_000014865.1_ASM1486v1
Magnetofaba.australis.IT.1 GCA_002109495.1_ASM210949v1
100/100
GCA_900546185.1.UMGS859*
100/100
GCA_900546225.1.UMGS860*
GCA_900550565.1.UMGS1303*
99.7/100
Anaplasma.phagocytophilum.str.HZ GCA_000013125.1_ASM1312v1
100/100
Candidatus.Neoehrlichia.lotoris.str.RAC413 GCA_000964795.1_ASM96479v1
100/100
Ehrlichia.ruminantium.str.Welgevonden GCA_000026005.1_ASM2600v1
98.6/98
Wolbachia.endosymbiont.of.Drosophila.melanogaster GCA_000008025.1_ASM802v1
Candidatus.Deianiraea.vastatrix GCA_007993655.1_ASM799365v1
95/96
100/100
Candidatus.Jidaibacter.acanthamoeba GCA_000815465.1.UWC36
100/100
Rickettsiales.endosymbiont.of.Peranema.trichophorum GCA_004210275.1_ASM421027v1
100/100
100/100
Candidatus.Midichloria.mitochondrii.IricVA GCA_000219355.1_ASM21935v1
100/100
Rickettsiales.endosymbiont.of.Trichoplax.sp.H2 GCA_009690945.1
100/100
Candidatus.Megaira.polyxenophila
55.3/88
Rickettsia.bellii.RML369.C GCA_000012385.1_ASM1238v1
100/100
Rickettsiales.bacterium.Ac37b GCA_000746585.2_ASM74658v2
Rickettsiales.endosymbiont.of.Stachyamoeba.lipophora GCA_003932735.1_ASM393273v1
67.1/94
Ahrensia.kielensis.DSM.5890 GCA_000374465.1_ASM37446v1
39.7/77
Chelativorans.oligotrophicus GCA_011317505.1_ASM1131750v1
100/100
Rhizobium.leguminosarum.bv.trifolii GCA_011604525.1_ASM1160452v1
48.6/77
Bartonella.quintana GCA_009936175.1_ASM993617v1
99.8/55
Pelagibacterium.halotolerans GCA_013004005.1_ASM1300400v1
100/100
100/100
Polymorphum.gilvum.SL003B.26A1 GCA_000192745.1_ASM19274v1
100/100
Roseibium.hamelinense GCA_007830545.1_ASM783054v1
Pseudovibrio.sp.FO.BEG1 GCA_000236645.1_ASM23664v1
100/100
100/100
Methylocella.silvestris.BL2 GCA_000021745.1_ASM2174v1
100/100
Methylocystis.bryophila GCA_002117405.1_ASM211740v1
100/100
Methylorubrum.extorquens.PA1 GCA_000018845.1_ASM1884v1
98.7/99
100/100
Rhodopseudomonas.palustris GCA_013415845.1_ASM1341584v1
Xanthobacter.autotrophicus.Py2 GCA_000017645.1_ASM1764v1
100/100
100/100
Hyphomicrobium.denitrificans.ATCC.51888 GCA_000143145.1_ASM14314v1
100/100
Rhodomicrobium.vannielii.ATCC.17100 GCA_000166055.1_ASM16605v1
Thermopetrobacter.sp.TC1 GCA_000746275.1_ASM74627v1
100/100
Candidatus.Micropelagos.thuwalensis GCA_000469155.1.RS24
100/100
alpha.proteobacterium.IMCC14465 GCA_000293845.2_ASM29384v2
Parvibaculum.lavamentivorans.DS.1 GCA_000017565.1_ASM1756v1
100/100
Asticcacaulis.excentricus.CB.48 GCA_000175215.2_ASM17521v2
100/100
Brevundimonas.subvibrioides.ATCC.15264 GCA_000144605.1_ASM14460v1
100/100
alpha.proteobacterium.L41A GCA_000335735.1_L41A.1.0
99.9/55
100/100
100/100
Caulobacter.vibrioides.NA1000 GCA_000022005.1_ASM2200v1
Phenylobacterium.zucineum.HLK1 GCA_000017265.1_ASM1726v1
100/100
Hirschia.baltica.ATCC.49814 GCA_000023785.1_ASM2378v1
100/100
100/100
Hyphomonas.neptunium.ATCC.15444 GCA_000013025.1_ASM1302v1
100/100
Maricaulis.maris GCA_003634045.1_ASM363404v1
Oceanicaulis.alexandrii.DSM.11625 GCA_000420265.1_ASM42026v1
Parvularcula.bermudensis.HTCC2503 GCA_000152825.2_ASM15282v2
Jannaschia.rubra GCA_900113265.1.IMG.taxon.2622736430.annotated.assembly
100/100
89/94
Octadecabacter.arcticus.238 GCA_000155735.2_ASM15573v2
100/100
100/100
Ruegeria.pomeroyi.DSS.3 GCA_000011965.2_ASM1196v2
100/100
100/99
Sagittula.stellata.E.37 GCA_000169415.1_ASM16941v1
100/100
Ketogulonicigenium.vulgare GCA_001399515.1_ASM139951v1
100/100
Rubellimicrobium.rubrum GCA_006152115.1_ASM615211v1
100/100
100/100
Paracoccus.denitrificans GCA_004063735.1_ASM406373v1
Rhodobacter.sphaeroides.ATCC.17029 GCA_000015985.1_ASM1598v1
Neomegalonema.perideroedes.DSM.15528 GCA_000374145.1_ASM37414v1
100/100
100/100
Candidatus.Pelagibacter.ubique.HIMB058 GCA_000419545.1_ASM41954v1
100/100
Candidatus.Pelagibacter.ubique.HTCC1062 GCA_000012345.1_ASM1234v1
100/100
alpha.proteobacterium.HIMB5 GCA_000299095.1_ASM29909v1
alpha.proteobacterium.HIMB114 GCA_000163555.2
100/100
Citromicrobium.sp.JL477 GCA_001304795.1_ASM130479v1
100/100
Sphingopyxis.macrogoltabida GCA_001314325.1_ASM131432v1
100/100
Sphingobium.indicum.B90A GCA_000264945.2_ASM26494v2
100/100
100/100
Sphingomonas.wittichii.RW1 GCA_000016765.1_ASM1676v1
100/100
Zymomonas.mobilis.subsp.mobilis.ATCC.31821 GCA_000007105.1_ASM710v1
alpha.proteobacterium.AAP81b GCA_001295935.1_ASM129593v1
Kordiimonas.gwangyangensis.DSM.19435 GCA_000375545.1_ASM37554v1
Sneathiella.glossodoripedis.JCM.23214 GCA_000616095.1_ASM61609v1
100/100
Geminicoccus.roseus.DSM.18922 GCA_000427665.1_ASM42766v1
Tistrella.mobilis.KA081020.065 GCA_000264455.2_ASM26445v2
100/55
Caenispirillum.salinarum.AK4 GCA_000315795.1
100/100
100/100
Pararhodospirillum.photometricum.DSM.122 GCA_000284415.2_ASM28441v2
93.2/100
Rhodospirillum.rubrum.F11 GCA_000225955.1_ASM22595v1
100/100
Roseospirillum.parvum GCA_900100455.1
90.5/54
Magnetospirillum.magneticum.AMB.1 GCA_000009985.1_ASM998v1
100/55
GCA_900544565.1.UMGS690*
100/100
Terasakiella.pusilla.DSM.6293 GCA_000688235.1_ASM68823v1
100/55
Thalassospira.profundimaris.WP0211 GCA_000300275.1_ASM30027v1
100/100
Candidatus.Puniceispirillum.marinum.IMCC1322 GCA_000024465.1_ASM2446v1
100/29
100/100
Thalassobaculum.salexigens.DSM.19539 GCA_000423805.1_ASM42380v1
100/100
alpha.proteobacterium.BAL199 GCA_000171835.1_ASM17183v1
100/100
Oceanibaculum.indicum GCA_003633955.1_ASM363395v1
100/100
100/100
Kiloniella.laminariae.DSM.19542 GCA_000374005.1_ASM37400v1
Rhodovibrio.salinarum.DSM.9154 GCA_000515255.1_ASM51525v1
100/100
Inquilinus.limosus.DSM.16000 GCA_000423185.1_ASM42318v1
100/100
Rhodospirillum.centenum GCA_014197805.1_ASM1419780v1
62.1/40
Micavibrio.aeruginosavorus.ARL.13 GCA_000226315.1_ASM22631v1
100/100
Caedimonas.varicaedens GCA_001192655.1_ASM119265v1
100/100
Candidatus.Caedibacter.acanthamoebae GCA_000743035.1_ASM74303v1
100/100
Candidatus.Nucleicultrix.amoebiphila.FS5 GCA_002117145.1_ASM211714v1
100/100
Candidatus.Finniella.inopinata GCA_004210305.1_ASM421030v1
47.8/55
100/100
Candidatus.Odyssella.thessalonicensis.L13 GCA_000190415.2_ASM19041v1
Candidatus.Paracaedibacter.acanthamoebae GCA_000742835.1_ASM74283v1
100/100
100/100
Candidatus.Hepatobacter.penaei GCA_000742475.1_ASM74247v1
Holospora.undulata.HU1 GCA_000388175.3_ASM38817v3
Elioraea.tepidiphila.DSM.17972 GCA_000378465.1_ASM37846v1
100/100
Belnapia.moabensis.DSM.16746 GCA_000745835.1_ASM74583v1
100/100
100/100
Rubritepida.flocculans.DSM.14296 GCA_000425365.1_ASM42536v1
100/100
Roseomonas.gilardii GCA_001941945.1_ASM194194v1
Acidiphilium.multivorum.AIU301 GCA_000202835.1_ASM20283v1
92.9/99
68.6/95
Acidisphaera.rubrifaciens.HS.AP3 GCA_000964365.1_ASM96436v1
100/100
Granulibacter.bethesdensis.CGDNIH4 GCA_000576145.2_ASM57614v2
100/100
Commensalibacter.intestini.A911 GCA_000231445.2_ASM23144v1
91.5/99
Gluconacetobacter.diazotrophicus.PA1.5 GCA_000067045.1_ASM6704v1
Acetobacter.aceti.NBRC.14818 GCA_000193495.2_ASM19349v2
Gluconobacter.oxydans.DSM.3504 GCA_000583855.1_ASM58385v1
0.566195

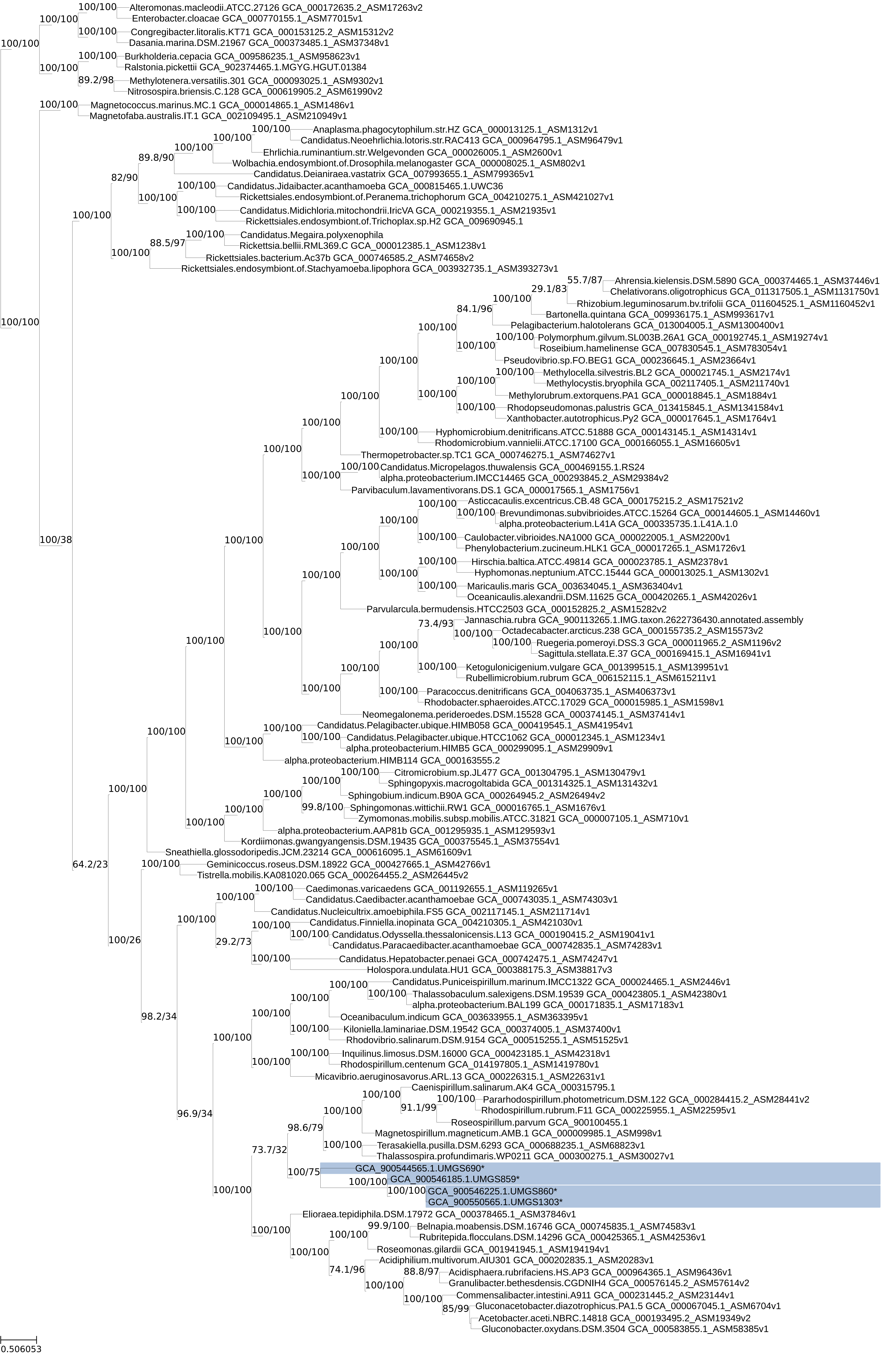
100/100
Alteromonas.macleodii.ATCC.27126 GCA_000172635.2_ASM17263v2
100/100
Enterobacter.cloacae GCA_000770155.1_ASM77015v1
100/100
100/100
Congregibacter.litoralis.KT71 GCA_000153125.2_ASM15312v2
Dasania.marina.DSM.21967 GCA_000373485.1_ASM37348v1
100/100
Burkholderia.cepacia GCA_009586235.1_ASM958623v1
100/100
Ralstonia.pickettii GCA_902374465.1.MGYG.HGUT.01384
89.2/98
Methylotenera.versatilis.301 GCA_000093025.1_ASM9302v1
Nitrosospira.briensis.C.128 GCA_000619905.2_ASM61990v2
100/100
Magnetococcus.marinus.MC.1 GCA_000014865.1_ASM1486v1
Magnetofaba.australis.IT.1 GCA_002109495.1_ASM210949v1
100/100
Anaplasma.phagocytophilum.str.HZ GCA_000013125.1_ASM1312v1
100/100
Candidatus.Neoehrlichia.lotoris.str.RAC413 GCA_000964795.1_ASM96479v1
100/100
Ehrlichia.ruminantium.str.Welgevonden GCA_000026005.1_ASM2600v1
89.8/90
Wolbachia.endosymbiont.of.Drosophila.melanogaster GCA_000008025.1_ASM802v1
Candidatus.Deianiraea.vastatrix GCA_007993655.1_ASM799365v1
82/90
100/100
Candidatus.Jidaibacter.acanthamoeba GCA_000815465.1.UWC36
100/100
Rickettsiales.endosymbiont.of.Peranema.trichophorum GCA_004210275.1_ASM421027v1
100/100
Candidatus.Midichloria.mitochondrii.IricVA GCA_000219355.1_ASM21935v1
100/100
Rickettsiales.endosymbiont.of.Trichoplax.sp.H2 GCA_009690945.1
100/100
Candidatus.Megaira.polyxenophila
88.5/97
Rickettsia.bellii.RML369.C GCA_000012385.1_ASM1238v1
100/100
Rickettsiales.bacterium.Ac37b GCA_000746585.2_ASM74658v2
Rickettsiales.endosymbiont.of.Stachyamoeba.lipophora GCA_003932735.1_ASM393273v1
55.7/87
Ahrensia.kielensis.DSM.5890 GCA_000374465.1_ASM37446v1
29.1/83
Chelativorans.oligotrophicus GCA_011317505.1_ASM1131750v1
100/100
Rhizobium.leguminosarum.bv.trifolii GCA_011604525.1_ASM1160452v1
100/100
84.1/96
Bartonella.quintana GCA_009936175.1_ASM993617v1
Pelagibacterium.halotolerans GCA_013004005.1_ASM1300400v1
100/100
100/100
Polymorphum.gilvum.SL003B.26A1 GCA_000192745.1_ASM19274v1
100/100
Roseibium.hamelinense GCA_007830545.1_ASM783054v1
Pseudovibrio.sp.FO.BEG1 GCA_000236645.1_ASM23664v1
100/100
100/100
Methylocella.silvestris.BL2 GCA_000021745.1_ASM2174v1
100/100
Methylocystis.bryophila GCA_002117405.1_ASM211740v1
100/100
Methylorubrum.extorquens.PA1 GCA_000018845.1_ASM1884v1
100/100
Rhodopseudomonas.palustris GCA_013415845.1_ASM1341584v1
100/100
Xanthobacter.autotrophicus.Py2 GCA_000017645.1_ASM1764v1
100/100
Hyphomicrobium.denitrificans.ATCC.51888 GCA_000143145.1_ASM14314v1
Rhodomicrobium.vannielii.ATCC.17100 GCA_000166055.1_ASM16605v1
100/100
Thermopetrobacter.sp.TC1 GCA_000746275.1_ASM74627v1
100/100
Candidatus.Micropelagos.thuwalensis GCA_000469155.1.RS24
100/100
alpha.proteobacterium.IMCC14465 GCA_000293845.2_ASM29384v2
Parvibaculum.lavamentivorans.DS.1 GCA_000017565.1_ASM1756v1
100/100
Asticcacaulis.excentricus.CB.48 GCA_000175215.2_ASM17521v2
100/100
Brevundimonas.subvibrioides.ATCC.15264 GCA_000144605.1_ASM14460v1
alpha.proteobacterium.L41A GCA_000335735.1.L41A.1.0
100/100
100/100
Caulobacter.vibrioides.NA1000 GCA_000022005.1_ASM2200v1
100/100
Phenylobacterium.zucineum.HLK1 GCA_000017265.1_ASM1726v1
100/100
Hirschia.baltica.ATCC.49814 GCA_000023785.1_ASM2378v1
100/100
Hyphomonas.neptunium.ATCC.15444 GCA_000013025.1_ASM1302v1
100/100
Maricaulis.maris GCA_003634045.1_ASM363404v1
100/100
Oceanicaulis.alexandrii.DSM.11625 GCA_000420265.1_ASM42026v1
Parvularcula.bermudensis.HTCC2503 GCA_000152825.2_ASM15282v2
73.4/93
Jannaschia.rubra GCA_900113265.1.IMG.taxon.2622736430.annotated.assembly
100/100
Octadecabacter.arcticus.238 GCA_000155735.2_ASM15573v2
100/100
Ruegeria.pomeroyi.DSS.3 GCA_000011965.2_ASM1196v2
100/100
Sagittula.stellata.E.37 GCA_000169415.1_ASM16941v1
100/100
100/100
Ketogulonicigenium.vulgare GCA_001399515.1_ASM139951v1
100/100
Rubellimicrobium.rubrum GCA_006152115.1_ASM615211v1
100/100
Paracoccus.denitrificans GCA_004063735.1_ASM406373v1
100/100
Rhodobacter.sphaeroides.ATCC.17029 GCA_000015985.1_ASM1598v1
Neomegalonema.perideroedes.DSM.15528 GCA_000374145.1_ASM37414v1
100/100
Candidatus.Pelagibacter.ubique.HIMB058 GCA_000419545.1_ASM41954v1
100/100
Candidatus.Pelagibacter.ubique.HTCC1062 GCA_000012345.1_ASM1234v1
100/100
alpha.proteobacterium.HIMB5 GCA_000299095.1_ASM29909v1
alpha.proteobacterium.HIMB114 GCA_000163555.2
100/100
Citromicrobium.sp.JL477 GCA_001304795.1_ASM130479v1
100/100
Sphingopyxis.macrogoltabida GCA_001314325.1_ASM131432v1
100/100
Sphingobium.indicum.B90A GCA_000264945.2_ASM26494v2
100/100
99.8/100
Sphingomonas.wittichii.RW1 GCA_000016765.1_ASM1676v1
100/100
Zymomonas.mobilis.subsp.mobilis.ATCC.31821 GCA_000007105.1_ASM710v1
100/100
alpha.proteobacterium.AAP81b GCA_001295935.1_ASM129593v1
Kordiimonas.gwangyangensis.DSM.19435 GCA_000375545.1_ASM37554v1
Sneathiella.glossodoripedis.JCM.23214 GCA_000616095.1_ASM61609v1
100/100
Geminicoccus.roseus.DSM.18922 GCA_000427665.1_ASM42766v1
64.2/23
Tistrella.mobilis.KA081020.065 GCA_000264455.2_ASM26445v2
100/100
Caedimonas.varicaedens GCA_001192655.1_ASM119265v1
100/100
Candidatus.Caedibacter.acanthamoebae GCA_000743035.1_ASM74303v1
Candidatus.Nucleicultrix.amoebiphila.FS5 GCA_002117145.1_ASM211714v1
100/100
100/100
Candidatus.Finniella.inopinata GCA_004210305.1_ASM421030v1
100/100
Candidatus.Odyssella.thessalonicensis.L13 GCA_000190415.2_ASM19041v1
100/26
29.2/73
Candidatus.Paracaedibacter.acanthamoebae GCA_000742835.1_ASM74283v1
100/100
Candidatus.Hepatobacter.penaei GCA_000742475.1_ASM74247v1
Holospora.undulata.HU1 GCA_000388175.3_ASM38817v3
100/100
Candidatus.Puniceispirillum.marinum.IMCC1322 GCA_000024465.1_ASM2446v1
100/100
Thalassobaculum.salexigens.DSM.19539 GCA_000423805.1_ASM42380v1
100/100
alpha.proteobacterium.BAL199 GCA_000171835.1_ASM17183v1
98.2/34
100/100
Oceanibaculum.indicum GCA_003633955.1_ASM363395v1
100/100
Kiloniella.laminariae.DSM.19542 GCA_000374005.1_ASM37400v1
100/100
Rhodovibrio.salinarum.DSM.9154 GCA_000515255.1_ASM51525v1
100/100
Inquilinus.limosus.DSM.16000 GCA_000423185.1_ASM42318v1
100/100
Rhodospirillum.centenum GCA_014197805.1_ASM1419780v1
Micavibrio.aeruginosavorus.ARL.13 GCA_000226315.1_ASM22631v1
Caenispirillum.salinarum.AK4 GCA_000315795.1
100/100
100/100
Pararhodospirillum.photometricum.DSM.122 GCA_000284415.2_ASM28441v2
91.1/99
Rhodospirillum.rubrum.F11 GCA_000225955.1_ASM22595v1
96.9/34
100/100
Roseospirillum.parvum GCA_900100455.1
98.6/79
Magnetospirillum.magneticum.AMB.1 GCA_000009985.1_ASM998v1
100/100
Terasakiella.pusilla.DSM.6293 GCA_000688235.1_ASM68823v1
73.7/32
Thalassospira.profundimaris.WP0211 GCA_000300275.1_ASM30027v1
GCA_900544565.1.UMGS690*
100/75
GCA_900546185.1.UMGS859*
100/100
100/100
GCA_900546225.1.UMGS860*
100/100
GCA_900550565.1.UMGS1303*
Elioraea.tepidiphila.DSM.17972 GCA_000378465.1_ASM37846v1
99.9/100
Belnapia.moabensis.DSM.16746 GCA_000745835.1_ASM74583v1
100/100
100/100
Rubritepida.flocculans.DSM.14296 GCA_000425365.1_ASM42536v1
Roseomonas.gilardii GCA_001941945.1_ASM194194v1
100/100
Acidiphilium.multivorum.AIU301 GCA_000202835.1_ASM20283v1
74.1/96
88.8/97
Acidisphaera.rubrifaciens.HS.AP3 GCA_000964365.1_ASM96436v1
Granulibacter.bethesdensis.CGDNIH4 GCA_000576145.2_ASM57614v2
100/100
Commensalibacter.intestini.A911 GCA_000231445.2_ASM23144v1
100/100
85/99
Gluconacetobacter.diazotrophicus.PA1.5 GCA_000067045.1_ASM6704v1
Acetobacter.aceti.NBRC.14818 GCA_000193495.2_ASM19349v2
Gluconobacter.oxydans.DSM.3504 GCA_000583855.1_ASM58385v1
100/100
100/38
0.506053

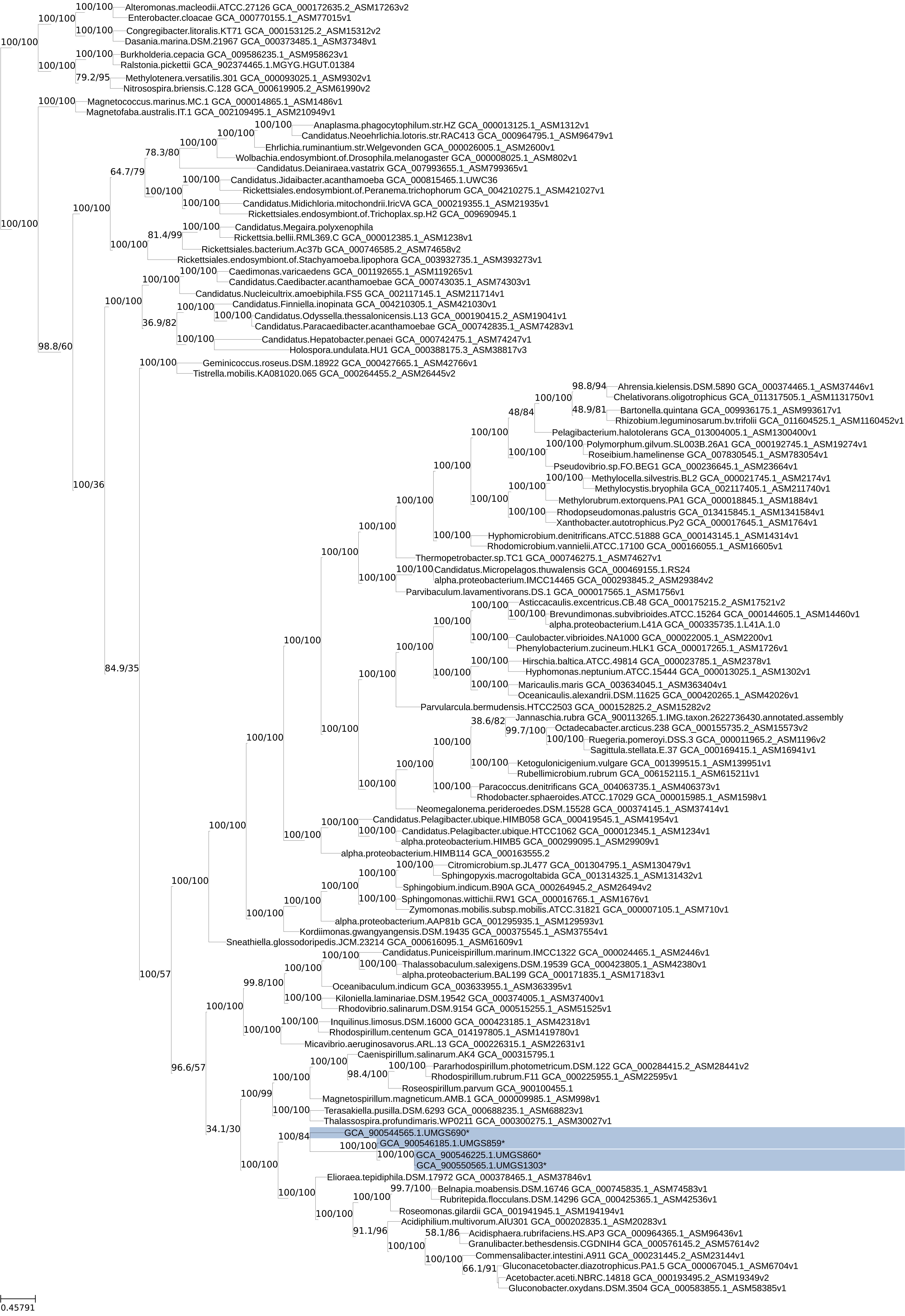

Alteromonas.macleodii.ATCC.27126 GCA_000172635.2_ASM17263v2
Enterobacter.cloacae GCA_000770155.1_ASM77015v1
Congregibacter.litoralis.KT71 GCA_000153125.2_ASM15312v2
Dasania.marina.DSM.21967 GCA_000373485.1_ASM37348v1
Burkholderia.cepacia GCA_009586235.1_ASM958623v1
Ralstonia.pickettii GCA_902374465.1.MGYG.HGUT.01384
Methylotenera.versatilis.301 GCA_000093025.1_ASM9302v1
Nitrosospira.briensis.C.128 GCA_000619905.2_ASM61990v2
Magnetococcus.marinus.MC.1 GCA_000014865.1_ASM1486v1
Magnetofaba.australis.IT.1 GCA_002109495.1_ASM210949v1
Anaplasma.phagocytophilum.str.HZ GCA_000013125.1_ASM1312v1
Candidatus.Neoehrlichia.lotoris.str.RAC413 GCA_000964795.1_ASM96479v1
Ehrlichia.ruminantium.str.Welgevonden GCA_000026005.1_ASM2600v1
Wolbachia.endosymbiont.of.Drosophila.melanogaster GCA_000008025.1_ASM802v1
Candidatus.Deianiraea.vastatrix GCA_007993655.1_ASM799365v1
Candidatus.Jidaibacter.acanthamoeba GCA_000815465.1.UWC36
Rickettsiales.endosymbiont.of.Peranema.trichophorum GCA_004210275.1_ASM421027v1
Candidatus.Midichloria.mitochondrii.IricVA GCA_000219355.1_ASM21935v1
Rickettsiales.endosymbiont.of.Trichoplax.sp.H2 GCA_009690945.1
Candidatus.Megaira.polyxenophila
Rickettsia.bellii.RML369.C GCA_000012385.1_ASM1238v1
Rickettsiales.bacterium.Ac37b GCA_000746585.2_ASM74658v2
Rickettsiales.endosymbiont.of.Stachyamoeba.lipophora GCA_003932735.1_ASM393273v1
Caedimonas.varicaedens GCA_001192655.1_ASM119265v1
Candidatus.Caedibacter.acanthamoebae GCA_000743035.1_ASM74303v1
Candidatus.Nucleicultrix.amoebiphila.FS5 GCA_002117145.1_ASM211714v1
Candidatus.Finniella.inopinata GCA_004210305.1_ASM421030v1
Candidatus.Odyssella.thessalonicensis.L13 GCA_000190415.2_ASM19041v1
Candidatus.Paracaedibacter.acanthamoebae GCA_000742835.1_ASM74283v1
Candidatus.Hepatobacter.penaei GCA_000742475.1_ASM74247v1
Holospora.undulata.HU1 GCA_000388175.3_ASM38817v3
Geminicoccus.roseus.DSM.18922 GCA_000427665.1_ASM42766v1
Tistrella.mobilis.KA081020.065 GCA_000264455.2_ASM26445v2
Ahrensia.kielensis.DSM.5890 GCA_000374465.1_ASM37446v1
Chelativorans.oligotrophicus GCA_011317505.1_ASM1131750v1
Bartonella.quintana GCA_009936175.1_ASM993617v1
Rhizobium.leguminosarum.bv.trifolii GCA_011604525.1_ASM1160452v1
Pelagibacterium.halotolerans GCA_013004005.1_ASM1300400v1
Polymorphum.gilvum.SL003B.26A1 GCA_000192745.1_ASM19274v1
Roseibium.hamelinense GCA_007830545.1_ASM783054v1
Pseudovibrio.sp.FO.BEG1 GCA_000236645.1_ASM23664v1
Methylocella.silvestris.BL2 GCA_000021745.1_ASM2174v1
Methylocystis.bryophila GCA_002117405.1_ASM211740v1
Methylorubrum.extorquens.PA1 GCA_000018845.1_ASM1884v1
Rhodopseudomonas.palustris GCA_013415845.1_ASM1341584v1
Xanthobacter.autotrophicus.Py2 GCA_000017645.1_ASM1764v1
Hyphomicrobium.denitrificans.ATCC.51888 GCA_000143145.1_ASM14314v1
Rhodomicrobium.vannielii.ATCC.17100 GCA_000166055.1_ASM16605v1
Thermopetrobacter.sp.TC1 GCA_000746275.1_ASM74627v1
Candidatus.Micropelagos.thuwalensis GCA_000469155.1.RS24
alpha.proteobacterium.IMCC14465 GCA_000293845.2_ASM29384v2
Parvibaculum.lavamentivorans.DS.1 GCA_000017565.1_ASM1756v1
Asticcacaulis.excentricus.CB.48 GCA_000175215.2_ASM17521v2
Brevundimonas.subvibrioides.ATCC.15264 GCA_000144605.1_ASM14460v1
alpha.proteobacterium.L41A GCA_000335735.1.L41A.1.0
Caulobacter.vibrioides.NA1000 GCA_000022005.1_ASM2200v1
Phenylobacterium.zucineum.HLK1 GCA_000017265.1_ASM1726v1
Hirschia.baltica.ATCC.49814 GCA_000023785.1_ASM2378v1
Hyphomonas.neptunium.ATCC.15444 GCA_000013025.1_ASM1302v1
Maricaulis.maris GCA_003634045.1_ASM363404v1
Oceanicaulis.alexandrii.DSM.11625 GCA_000420265.1_ASM42026v1
Parvularcula.bermudensis.HTCC2503 GCA_000152825.2_ASM15282v2
Jannaschia.rubra GCA_900113265.1.IMG.taxon.2622736430.annotated.assembly
Octadecabacter.arcticus.238 GCA_000155735.2_ASM15573v2
Ruegeria.pomeroyi.DSS.3 GCA_000011965.2_ASM1196v2
Sagittula.stellata.E.37 GCA_000169415.1_ASM16941v1
Ketogulonicigenium.vulgare GCA_001399515.1_ASM139951v1
Rubellimicrobium.rubrum GCA_006152115.1_ASM615211v1
Paracoccus.denitrificans GCA_004063735.1_ASM406373v1
Rhodobacter.sphaeroides.ATCC.17029 GCA_000015985.1_ASM1598v1
Neomegalonema.perideroedes.DSM.15528 GCA_000374145.1_ASM37414v1
Candidatus.Pelagibacter.ubique.HIMB058 GCA_000419545.1_ASM41954v1
Candidatus.Pelagibacter.ubique.HTCC1062 GCA_000012345.1_ASM1234v1
alpha.proteobacterium.HIMB5 GCA_000299095.1_ASM29909v1
alpha.proteobacterium.HIMB114 GCA_000163555.2
Citromicrobium.sp.JL477 GCA_001304795.1_ASM130479v1
Sphingopyxis.macrogoltabida GCA_001314325.1_ASM131432v1
Sphingobium.indicum.B90A GCA_000264945.2_ASM26494v2
Sphingomonas.wittichii.RW1 GCA_000016765.1_ASM1676v1
Zymomonas.mobilis.subsp.mobilis.ATCC.31821 GCA_000007105.1_ASM710v1
alpha.proteobacterium.AAP81b GCA_001295935.1_ASM129593v1
Kordiimonas.gwangyangensis.DSM.19435 GCA_000375545.1_ASM37554v1
Sneathiella.glossodoripedis.JCM.23214 GCA_000616095.1_ASM61609v1
Candidatus.Puniceispirillum.marinum.IMCC1322 GCA_000024465.1_ASM2446v1
Thalassobaculum.salexigens.DSM.19539 GCA_000423805.1_ASM42380v1
alpha.proteobacterium.BAL199 GCA_000171835.1_ASM17183v1
Oceanibaculum.indicum GCA_003633955.1_ASM363395v1
Kiloniella.laminariae.DSM.19542 GCA_000374005.1_ASM37400v1
Rhodovibrio.salinarum.DSM.9154 GCA_000515255.1_ASM51525v1
Inquilinus.limosus.DSM.16000 GCA_000423185.1_ASM42318v1
Rhodospirillum.centenum GCA_014197805.1_ASM1419780v1
Micavibrio.aeruginosavorus.ARL.13 GCA_000226315.1_ASM22631v1
Caenispirillum.salinarum.AK4 GCA_000315795.1
Pararhodospirillum.photometricum.DSM.122 GCA_000284415.2_ASM28441v2
Rhodospirillum.rubrum.F11 GCA_000225955.1_ASM22595v1
Roseospirillum.parvum GCA_900100455.1
Magnetospirillum.magneticum.AMB.1 GCA_000009985.1_ASM998v1
Terasakiella.pusilla.DSM.6293 GCA_000688235.1_ASM68823v1
Thalassospira.profundimaris.WP0211 GCA_000300275.1_ASM30027v1
GCA_900544565.1.UMGS690*
GCA_900546185.1.UMGS859*
GCA_900546225.1.UMGS860*
GCA_900550565.1.UMGS1303*
Elioraea.tepidiphila.DSM.17972 GCA_000378465.1_ASM37846v1
Belnapia.moabensis.DSM.16746 GCA_000745835.1_ASM74583v1
Rubritepida.flocculans.DSM.14296 GCA_000425365.1_ASM42536v1
Roseomonas.gilardii GCA_001941945.1_ASM194194v1
Acidiphilium.multivorum.AIU301 GCA_000202835.1_ASM20283v1
Acidisphaera.rubrifaciens.HS.AP3 GCA_000964365.1_ASM96436v1
Granulibacter.bethesdensis.CGDNIH4 GCA_000576145.2_ASM57614v2
Commensalibacter.intestini.A911 GCA_000231445.2_ASM23144v1
Gluconacetobacter.diazotrophicus.PA1.5 GCA_000067045.1_ASM6704v1
Acetobacter.aceti.NBRC.14818 GCA_000193495.2_ASM19349v2
Gluconobacter.oxydans.DSM.3504 GCA_000583855.1_ASM58385v1
0.45791

### Group J

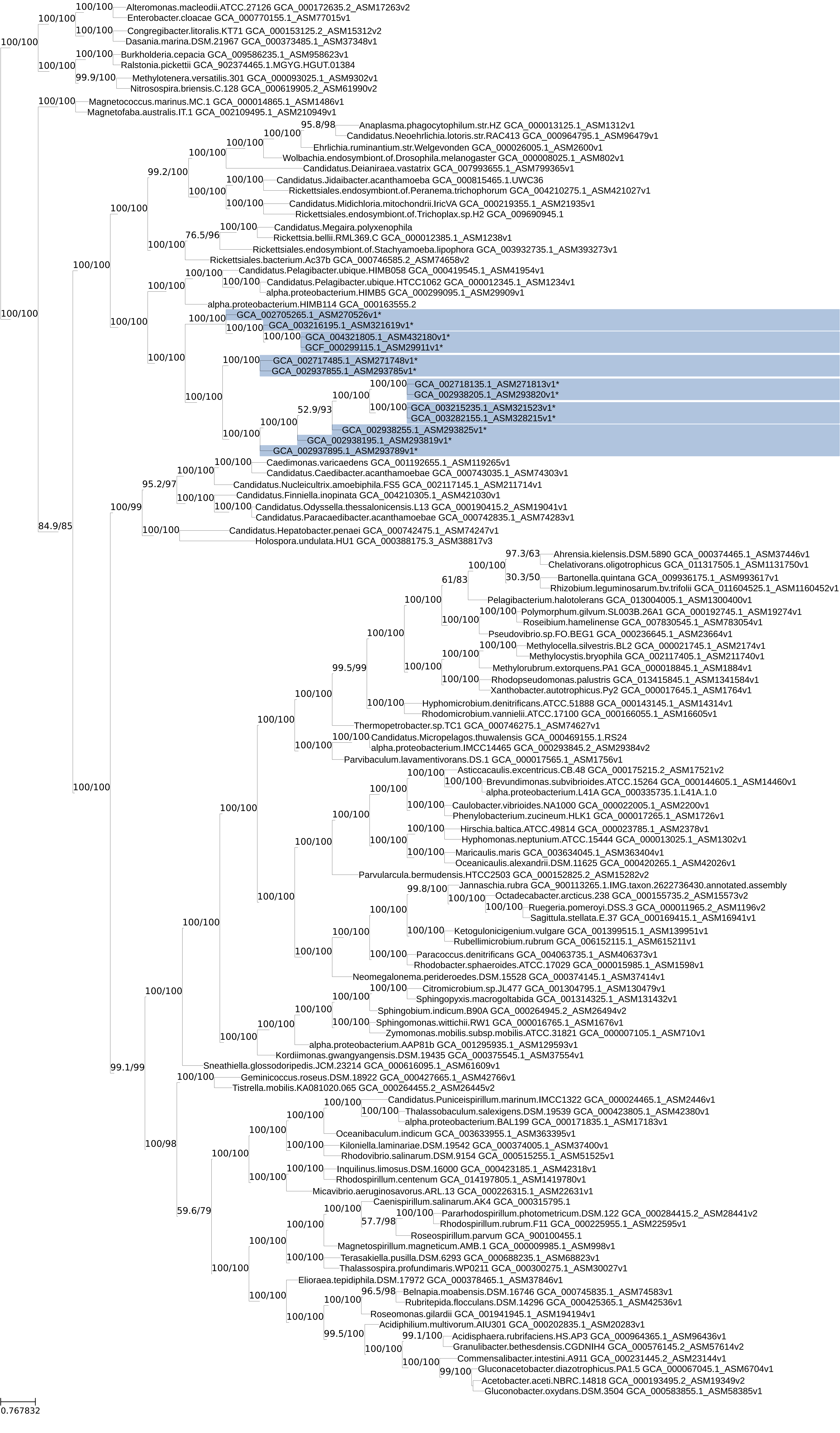

100/100
Alteromonas.macleodii.ATCC.27126 GCA_000172635.2_ASM17263v2
100/100
Enterobacter.cloacae GCA_000770155.1_ASM77015v1
100/100
Congregibacter.litoralis.KT71 GCA_000153125.2_ASM15312v2
100/100
Dasania.marina.DSM.21967 GCA_000373485.1_ASM37348v1
100/100
Burkholderia.cepacia GCA_009586235.1_ASM958623v1
100/100
Ralstonia.pickettii GCA_902374465.1.MGYG.HGUT.01384
99.9/100
Methylotenera.versatilis.301 GCA_000093025.1_ASM9302v1
Nitrosospira.briensis.C.128 GCA_000619905.2_ASM61990v2
100/100
Magnetococcus.marinus.MC.1 GCA_000014865.1_ASM1486v1
Magnetofaba.australis.IT.1 GCA_002109495.1_ASM210949v1
95.8/98
Anaplasma.phagocytophilum.str.HZ GCA_000013125.1_ASM1312v1
100/100
Candidatus.Neoehrlichia.lotoris.str.RAC413 GCA_000964795.1_ASM96479v1
100/100
Ehrlichia.ruminantium.str.Welgevonden GCA_000026005.1_ASM2600v1
100/100
Wolbachia.endosymbiont.of.Drosophila.melanogaster GCA_000008025.1_ASM802v1
99.2/100
Candidatus.Deianiraea.vastatrix GCA_007993655.1_ASM799365v1
100/100
Candidatus.Jidaibacter.acanthamoeba GCA_000815465.1.UWC36
100/100
Rickettsiales.endosymbiont.of.Peranema.trichophorum GCA_004210275.1_ASM421027v1
100/100
Candidatus.Midichloria.mitochondrii.IricVA GCA_000219355.1_ASM21935v1
100/100
Rickettsiales.endosymbiont.of.Trichoplax.sp.H2 GCA_009690945.1
100/100
Candidatus.Megaira.polyxenophila
76.5/96
Rickettsia.bellii.RML369.C GCA_000012385.1_ASM1238v1
100/100
Rickettsiales.endosymbiont.of.Stachyamoeba.lipophora GCA_003932735.1_ASM393273v1
100/100
Rickettsiales.bacterium.Ac37b GCA_000746585.2_ASM74658v2
100/100
Candidatus.Pelagibacter.ubique.HIMB058 GCA_000419545.1_ASM41954v1
100/100
Candidatus.Pelagibacter.ubique.HTCC1062 GCA_000012345.1_ASM1234v1
100/100
alpha.proteobacterium.HIMB5 GCA_000299095.1_ASM29909v1
alpha.proteobacterium.HIMB114 GCA_000163555.2
100/100
GCA_002705265.1_ASM270526v1*
100/100
GCA_003216195.1_ASM321619v1*
100/100
GCA_004321805.1_ASM432180v1*
GCF_000299115.1_ASM29911v1*
100/100
100/100
GCA_002717485.1_ASM271748v1*
GCA_002937855.1_ASM293785v1*
100/100
GCA_002718135.1_ASM271813v1*
100/100
GCA_002938205.1_ASM293820v1*
100/100
GCA_003215235.1_ASM321523v1*
52.9/93
GCA_003282155.1_ASM328215v1*
100/100
GCA_002938255.1_ASM293825v1*
100/100
GCA_002938195.1_ASM293819v1*
GCA_002937895.1_ASM293789v1*
100/100
Caedimonas.varicaedens GCA_001192655.1_ASM119265v1
100/100
Candidatus.Caedibacter.acanthamoebae GCA_000743035.1_ASM74303v1
95.2/97
Candidatus.Nucleicultrix.amoebiphila.FS5 GCA_002117145.1_ASM211714v1
Candidatus.Finniella.inopinata GCA_004210305.1_ASM421030v1
100/100
100/100
Candidatus.Odyssella.thessalonicensis.L13 GCA_000190415.2_ASM19041v1
100/99
Candidatus.Paracaedibacter.acanthamoebae GCA_000742835.1_ASM74283v1
84.9/85
100/100
Candidatus.Hepatobacter.penaei GCA_000742475.1_ASM74247v1
Holospora.undulata.HU1 GCA_000388175.3_ASM38817v3
97.3/63
Ahrensia.kielensis.DSM.5890 GCA_000374465.1_ASM37446v1
100/100
Chelativorans.oligotrophicus GCA_011317505.1_ASM1131750v1
30.3/50
Bartonella.quintana GCA_009936175.1_ASM993617v1
61/83
Rhizobium.leguminosarum.bv.trifolii GCA_011604525.1_ASM1160452v1
100/100
Pelagibacterium.halotolerans GCA_013004005.1_ASM1300400v1
100/100
Polymorphum.gilvum.SL003B.26A1 GCA_000192745.1_ASM19274v1
100/100
Roseibium.hamelinense GCA_007830545.1_ASM783054v1
100/100
Pseudovibrio.sp.FO.BEG1 GCA_000236645.1_ASM23664v1
100/100
Methylocella.silvestris.BL2 GCA_000021745.1_ASM2174v1
100/100
Methylocystis.bryophila GCA_002117405.1_ASM211740v1
99.5/99
100/100
Methylorubrum.extorquens.PA1 GCA_000018845.1_ASM1884v1
100/100
Rhodopseudomonas.palustris GCA_013415845.1_ASM1341584v1
Xanthobacter.autotrophicus.Py2 GCA_000017645.1_ASM1764v1
100/100
Hyphomicrobium.denitrificans.ATCC.51888 GCA_000143145.1_ASM14314v1
100/100
Rhodomicrobium.vannielii.ATCC.17100 GCA_000166055.1_ASM16605v1
Thermopetrobacter.sp.TC1 GCA_000746275.1_ASM74627v1
100/100
Candidatus.Micropelagos.thuwalensis GCA_000469155.1.RS24
100/100
alpha.proteobacterium.IMCC14465 GCA_000293845.2_ASM29384v2
Parvibaculum.lavamentivorans.DS.1 GCA_000017565.1_ASM1756v1
100/100
Asticcacaulis.excentricus.CB.48 GCA_000175215.2_ASM17521v2
100/100
100/100
Brevundimonas.subvibrioides.ATCC.15264 GCA_000144605.1_ASM14460v1
alpha.proteobacterium.L41A GCA_000335735.1.L41A.1.0
100/100
100/100
Caulobacter.vibrioides.NA1000 GCA_000022005.1_ASM2200v1
Phenylobacterium.zucineum.HLK1 GCA_000017265.1_ASM1726v1
100/100
100/100
Hirschia.baltica.ATCC.49814 GCA_000023785.1_ASM2378v1
100/100
Hyphomonas.neptunium.ATCC.15444 GCA_000013025.1_ASM1302v1
100/100
Maricaulis.maris GCA_003634045.1_ASM363404v1
Oceanicaulis.alexandrii.DSM.11625 GCA_000420265.1_ASM42026v1
Parvularcula.bermudensis.HTCC2503 GCA_000152825.2_ASM15282v2
99.8/100
Jannaschia.rubra GCA_900113265.1.IMG.taxon.2622736430.annotated.assembly
100/100
Octadecabacter.arcticus.238 GCA_000155735.2_ASM15573v2
100/100
100/100
Ruegeria.pomeroyi.DSS.3 GCA_000011965.2_ASM1196v2
Sagittula.stellata.E.37 GCA_000169415.1_ASM16941v1
100/100
100/100
Ketogulonicigenium.vulgare GCA_001399515.1_ASM139951v1
100/100
Rubellimicrobium.rubrum GCA_006152115.1_ASM615211v1
100/100
Paracoccus.denitrificans GCA_004063735.1_ASM406373v1
Rhodobacter.sphaeroides.ATCC.17029 GCA_000015985.1_ASM1598v1
Neomegalonema.perideroedes.DSM.15528 GCA_000374145.1_ASM37414v1
100/100
100/100
Citromicrobium.sp.JL477 GCA_001304795.1_ASM130479v1
Sphingopyxis.macrogoltabida GCA_001314325.1_ASM131432v1
100/100
Sphingobium.indicum.B90A GCA_000264945.2_ASM26494v2
100/100
100/100
Sphingomonas.wittichii.RW1 GCA_000016765.1_ASM1676v1
100/100
Zymomonas.mobilis.subsp.mobilis.ATCC.31821 GCA_000007105.1_ASM710v1
alpha.proteobacterium.AAP81b GCA_001295935.1_ASM129593v1
Kordiimonas.gwangyangensis.DSM.19435 GCA_000375545.1_ASM37554v1
99.1/99
Sneathiella.glossodoripedis.JCM.23214 GCA_000616095.1_ASM61609v1
100/100
Geminicoccus.roseus.DSM.18922 GCA_000427665.1_ASM42766v1
Tistrella.mobilis.KA081020.065 GCA_000264455.2_ASM26445v2
100/100
Candidatus.Puniceispirillum.marinum.IMCC1322 GCA_000024465.1_ASM2446v1
100/100
100/100
Thalassobaculum.salexigens.DSM.19539 GCA_000423805.1_ASM42380v1
alpha.proteobacterium.BAL199 GCA_000171835.1_ASM17183v1
100/100
100/98
Oceanibaculum.indicum GCA_003633955.1_ASM363395v1
100/100
Kiloniella.laminariae.DSM.19542 GCA_000374005.1_ASM37400v1
100/100
Rhodovibrio.salinarum.DSM.9154 GCA_000515255.1_ASM51525v1
100/100
Inquilinus.limosus.DSM.16000 GCA_000423185.1_ASM42318v1
100/100
Rhodospirillum.centenum GCA_014197805.1_ASM1419780v1
Micavibrio.aeruginosavorus.ARL.13 GCA_000226315.1_ASM22631v1
100/100
Caenispirillum.salinarum.AK4 GCA_000315795.1
59.6/79
100/100
Pararhodospirillum.photometricum.DSM.122 GCA_000284415.2_ASM28441v2
100/100
57.7/98
Rhodospirillum.rubrum.F11 GCA_000225955.1_ASM22595v1
Roseospirillum.parvum GCA_900100455.1
100/100
Magnetospirillum.magneticum.AMB.1 GCA_000009985.1_ASM998v1
100/100
Terasakiella.pusilla.DSM.6293 GCA_000688235.1_ASM68823v1
100/100
Thalassospira.profundimaris.WP0211 GCA_000300275.1_ASM30027v1
Elioraea.tepidiphila.DSM.17972 GCA_000378465.1_ASM37846v1
96.5/98
Belnapia.moabensis.DSM.16746 GCA_000745835.1_ASM74583v1
100/100
100/100
Rubritepida.flocculans.DSM.14296 GCA_000425365.1_ASM42536v1
100/100
Roseomonas.gilardii GCA_001941945.1_ASM194194v1
Acidiphilium.multivorum.AIU301 GCA_000202835.1_ASM20283v1
99.5/100
99.1/100
Acidisphaera.rubrifaciens.HS.AP3 GCA_000964365.1_ASM96436v1
100/100
Granulibacter.bethesdensis.CGDNIH4 GCA_000576145.2_ASM57614v2
100/100
Commensalibacter.intestini.A911 GCA_000231445.2_ASM23144v1
99/100
Gluconacetobacter.diazotrophicus.PA1.5 GCA_000067045.1_ASM6704v1
Acetobacter.aceti.NBRC.14818 GCA_000193495.2_ASM19349v2
Gluconobacter.oxydans.DSM.3504 GCA_000583855.1_ASM58385v1
0.767832

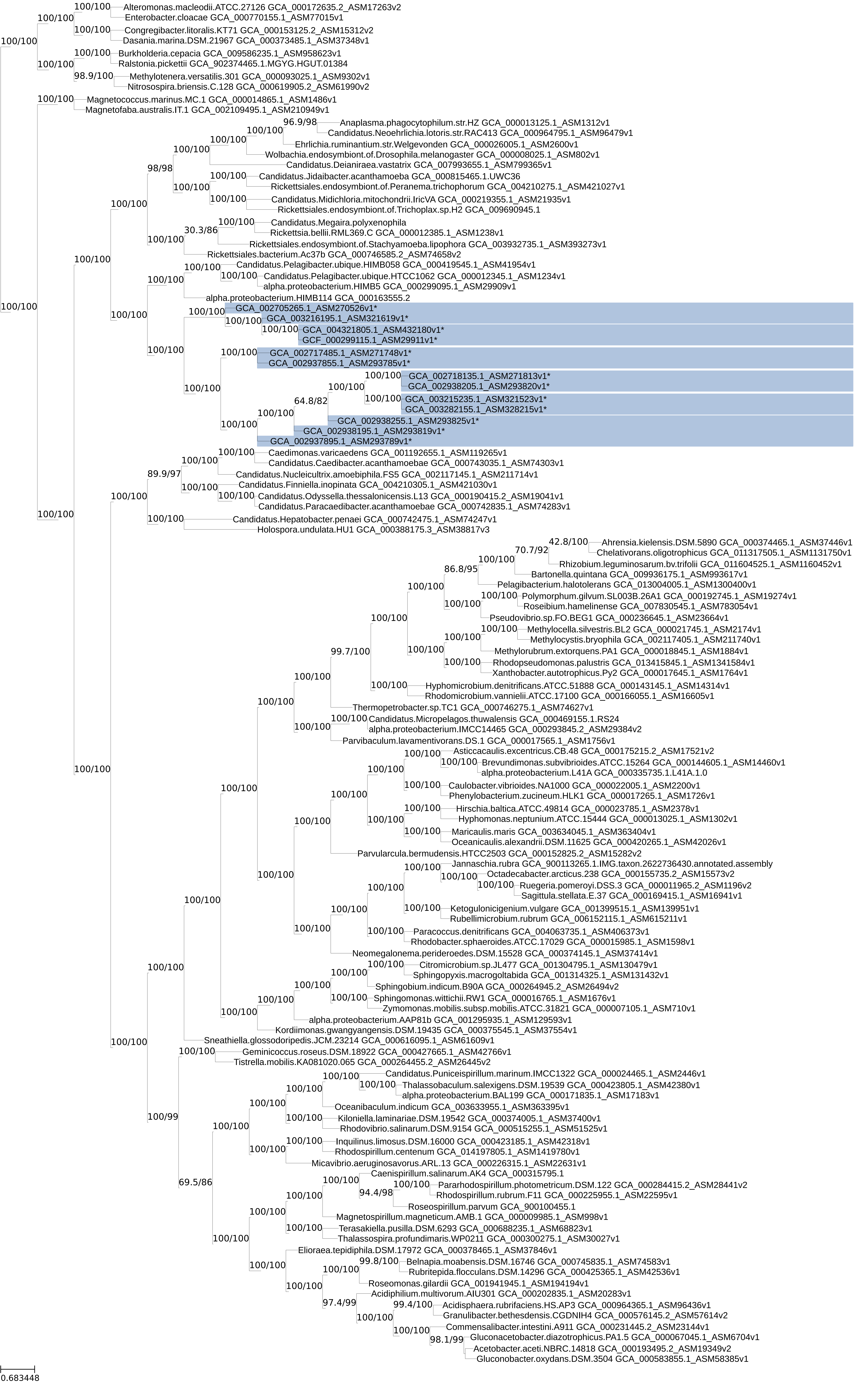

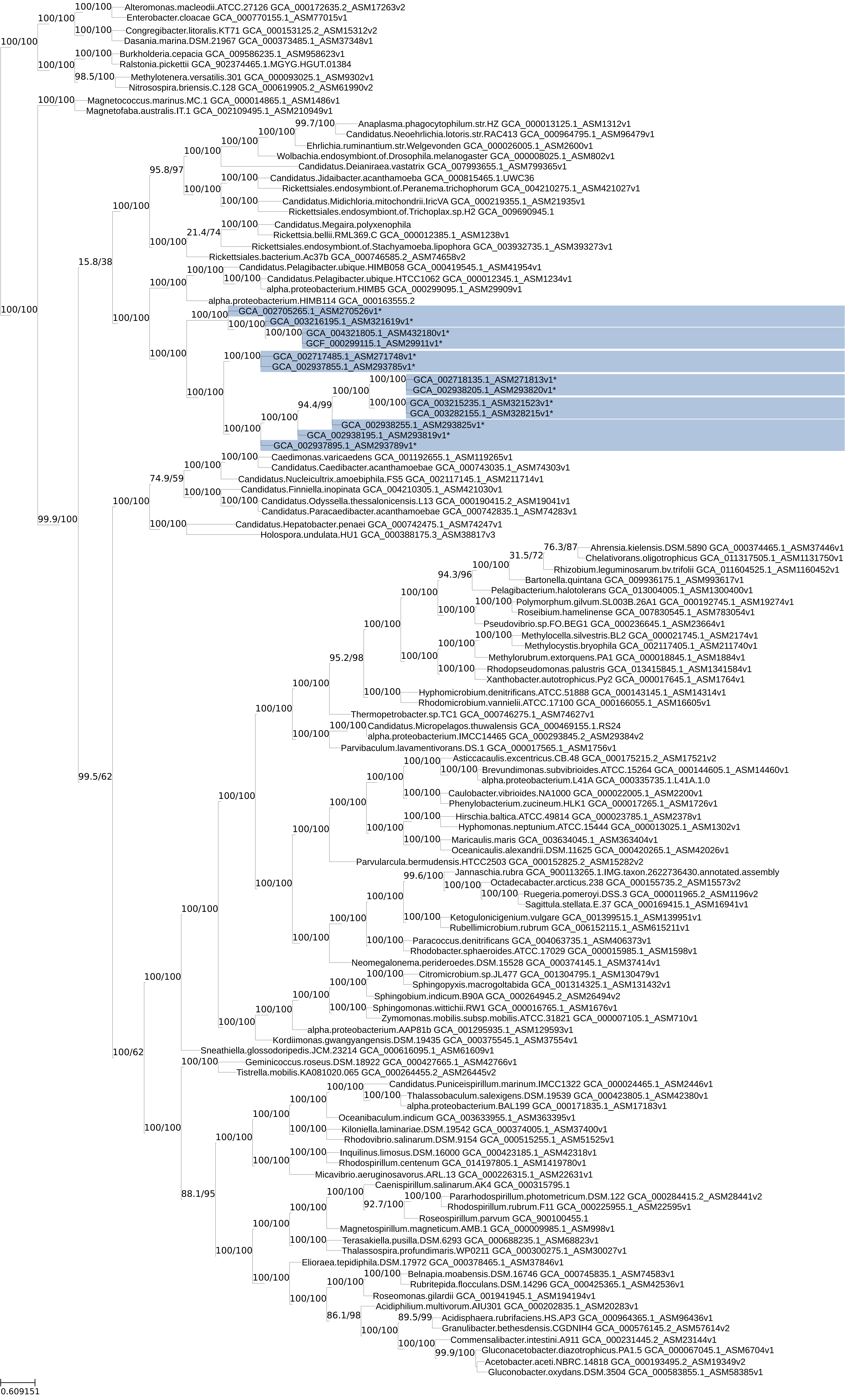

100/100
Alteromonas.macleodii.ATCC.27126 GCA_000172635.2_ASM17263v2
100/100
Enterobacter.cloacae GCA_000770155.1_ASM77015v1
100/100
Congregibacter.litoralis.KT71 GCA_000153125.2_ASM15312v2
100/100
Dasania.marina.DSM.21967 GCA_000373485.1_ASM37348v1
100/100
Burkholderia.cepacia GCA_009586235.1_ASM958623v1
100/100
Ralstonia.pickettii GCA_902374465.1.MGYG.HGUT.01384
98.5/100
Methylotenera.versatilis.301 GCA_000093025.1_ASM9302v1
Nitrosospira.briensis.C.128 GCA_000619905.2_ASM61990v2
100/100
Magnetococcus.marinus.MC.1 GCA_000014865.1_ASM1486v1
Magnetofaba.australis.IT.1 GCA_002109495.1_ASM210949v1
99.7/100
Anaplasma.phagocytophilum.str.HZ GCA_000013125.1_ASM1312v1
100/100
Candidatus.Neoehrlichia.lotoris.str.RAC413 GCA_000964795.1_ASM96479v1
100/100
Ehrlichia.ruminantium.str.Welgevonden GCA_000026005.1_ASM2600v1
100/100
Wolbachia.endosymbiont.of.Drosophila.melanogaster GCA_000008025.1_ASM802v1
95.8/97
Candidatus.Deianiraea.vastatrix GCA_007993655.1_ASM799365v1
100/100
Candidatus.Jidaibacter.acanthamoeba GCA_000815465.1.UWC36
100/100
Rickettsiales.endosymbiont.of.Peranema.trichophorum GCA_004210275.1_ASM421027v1
100/100
Candidatus.Midichloria.mitochondrii.IricVA GCA_000219355.1_ASM21935v1
100/100
Rickettsiales.endosymbiont.of.Trichoplax.sp.H2 GCA_009690945.1
100/100
Candidatus.Megaira.polyxenophila
21.4/74
Rickettsia.bellii.RML369.C GCA_000012385.1_ASM1238v1
100/100
Rickettsiales.endosymbiont.of.Stachyamoeba.lipophora GCA_003932735.1_ASM393273v1
Rickettsiales.bacterium.Ac37b GCA_000746585.2_ASM74658v2
15.8/38
100/100
Candidatus.Pelagibacter.ubique.HIMB058 GCA_000419545.1_ASM41954v1
100/100
100/100
Candidatus.Pelagibacter.ubique.HTCC1062 GCA_000012345.1_ASM1234v1
alpha.proteobacterium.HIMB5 GCA_000299095.1_ASM29909v1
alpha.proteobacterium.HIMB114 GCA_000163555.2
100/100
100/100
GCA_002705265.1_ASM270526v1*
100/100
GCA_003216195.1_ASM321619v1*
100/100
GCA_004321805.1_ASM432180v1*
GCF_000299115.1_ASM29911v1*
100/100
100/100
GCA_002717485.1_ASM271748v1*
GCA_002937855.1_ASM293785v1*
100/100
100/100
GCA_002718135.1_ASM271813v1*
GCA_002938205.1_ASM293820v1*
100/100
GCA_003215235.1_ASM321523v1*
GCA_003282155.1_ASM328215v1*
94.4/99
100/100
GCA_002938255.1_ASM293825v1*
100/100
GCA_002938195.1_ASM293819v1*
GCA_002937895.1_ASM293789v1*
100/100
Caedimonas.varicaedens GCA_001192655.1_ASM119265v1
100/100
Candidatus.Caedibacter.acanthamoebae GCA_000743035.1_ASM74303v1
74.9/59
Candidatus.Nucleicultrix.amoebiphila.FS5 GCA_002117145.1_ASM211714v1
100/100
Candidatus.Finniella.inopinata GCA_004210305.1_ASM421030v1
100/100
Candidatus.Odyssella.thessalonicensis.L13 GCA_000190415.2_ASM19041v1
100/100
Candidatus.Paracaedibacter.acanthamoebae GCA_000742835.1_ASM74283v1
99.9/100
100/100
Candidatus.Hepatobacter.penaei GCA_000742475.1_ASM74247v1
Holospora.undulata.HU1 GCA_000388175.3_ASM38817v3
76.3/87
Ahrensia.kielensis.DSM.5890 GCA_000374465.1_ASM37446v1
31.5/72
Chelativorans.oligotrophicus GCA_011317505.1_ASM1131750v1
100/100
Rhizobium.leguminosarum.bv.trifolii GCA_011604525.1_ASM1160452v1
Bartonella.quintana GCA_009936175.1_ASM993617v1
94.3/96
Pelagibacterium.halotolerans GCA_013004005.1_ASM1300400v1
100/100
100/100
Polymorphum.gilvum.SL003B.26A1 GCA_000192745.1_ASM19274v1
100/100
Roseibium.hamelinense GCA_007830545.1_ASM783054v1
Pseudovibrio.sp.FO.BEG1 GCA_000236645.1_ASM23664v1
100/100
100/100
Methylocella.silvestris.BL2 GCA_000021745.1_ASM2174v1
100/100
Methylocystis.bryophila GCA_002117405.1_ASM211740v1
100/100
Methylorubrum.extorquens.PA1 GCA_000018845.1_ASM1884v1
95.2/98
100/100
Rhodopseudomonas.palustris GCA_013415845.1_ASM1341584v1
Xanthobacter.autotrophicus.Py2 GCA_000017645.1_ASM1764v1
100/100
100/100
Hyphomicrobium.denitrificans.ATCC.51888 GCA_000143145.1_ASM14314v1
Rhodomicrobium.vannielii.ATCC.17100 GCA_000166055.1_ASM16605v1
100/100
Thermopetrobacter.sp.TC1 GCA_000746275.1_ASM74627v1
100/100
Candidatus.Micropelagos.thuwalensis GCA_000469155.1.RS24
100/100
alpha.proteobacterium.IMCC14465 GCA_000293845.2_ASM29384v2
Parvibaculum.lavamentivorans.DS.1 GCA_000017565.1_ASM1756v1
100/100
Asticcacaulis.excentricus.CB.48 GCA_000175215.2_ASM17521v2
100/100
100/100
Brevundimonas.subvibrioides.ATCC.15264 GCA_000144605.1_ASM14460v1
alpha.proteobacterium.L41A GCA_000335735.1.L41A.1.0
100/100
100/100
Caulobacter.vibrioides.NA1000 GCA_000022005.1_ASM2200v1
Phenylobacterium.zucineum.HLK1 GCA_000017265.1_ASM1726v1
100/100
100/100
Hirschia.baltica.ATCC.49814 GCA_000023785.1_ASM2378v1
100/100
Hyphomonas.neptunium.ATCC.15444 GCA_000013025.1_ASM1302v1
100/100
Maricaulis.maris GCA_003634045.1_ASM363404v1
Oceanicaulis.alexandrii.DSM.11625 GCA_000420265.1_ASM42026v1
Parvularcula.bermudensis.HTCC2503 GCA_000152825.2_ASM15282v2
99.6/100
Jannaschia.rubra GCA_900113265.1.IMG.taxon.2622736430.annotated.assembly
100/100
100/100
Octadecabacter.arcticus.238 GCA_000155735.2_ASM15573v2
100/100
Ruegeria.pomeroyi.DSS.3 GCA_000011965.2_ASM1196v2
100/100
Sagittula.stellata.E.37 GCA_000169415.1_ASM16941v1
100/100
Ketogulonicigenium.vulgare GCA_001399515.1_ASM139951v1
100/100
Rubellimicrobium.rubrum GCA_006152115.1_ASM615211v1
100/100
100/100
Paracoccus.denitrificans GCA_004063735.1_ASM406373v1
Rhodobacter.sphaeroides.ATCC.17029 GCA_000015985.1_ASM1598v1
Neomegalonema.perideroedes.DSM.15528 GCA_000374145.1_ASM37414v1
100/100
Citromicrobium.sp.JL477 GCA_001304795.1_ASM130479v1
100/100
100/100
Sphingopyxis.macrogoltabida GCA_001314325.1_ASM131432v1
Sphingobium.indicum.B90A GCA_000264945.2_ASM26494v2
100/100
Sphingomonas.wittichii.RW1 GCA_000016765.1_ASM1676v1
100/100
Zymomonas.mobilis.subsp.mobilis.ATCC.31821 GCA_000007105.1_ASM710v1
100/100
100/100
alpha.proteobacterium.AAP81b GCA_001295935.1_ASM129593v1
Kordiimonas.gwangyangensis.DSM.19435 GCA_000375545.1_ASM37554v1
99.5/62
100/100
Sneathiella.glossodoripedis.JCM.23214 GCA_000616095.1_ASM61609v1
100/62
100/100
Geminicoccus.roseus.DSM.18922 GCA_000427665.1_ASM42766v1
Tistrella.mobilis.KA081020.065 GCA_000264455.2_ASM26445v2
100/100
Candidatus.Puniceispirillum.marinum.IMCC1322 GCA_000024465.1_ASM2446v1
100/100
100/100
Thalassobaculum.salexigens.DSM.19539 GCA_000423805.1_ASM42380v1
alpha.proteobacterium.BAL199 GCA_000171835.1_ASM17183v1
100/100
100/100
Oceanibaculum.indicum GCA_003633955.1_ASM363395v1
100/100
Kiloniella.laminariae.DSM.19542 GCA_000374005.1_ASM37400v1
100/100
Rhodovibrio.salinarum.DSM.9154 GCA_000515255.1_ASM51525v1
100/100
100/100
Inquilinus.limosus.DSM.16000 GCA_000423185.1_ASM42318v1
Rhodospirillum.centenum GCA_014197805.1_ASM1419780v1
Micavibrio.aeruginosavorus.ARL.13 GCA_000226315.1_ASM22631v1
88.1/95
100/100
Caenispirillum.salinarum.AK4 GCA_000315795.1
100/100
Pararhodospirillum.photometricum.DSM.122 GCA_000284415.2_ASM28441v2
92.7/100
Rhodospirillum.rubrum.F11 GCA_000225955.1_ASM22595v1
100/100
Roseospirillum.parvum GCA_900100455.1
100/100
Magnetospirillum.magneticum.AMB.1 GCA_000009985.1_ASM998v1
100/100
Terasakiella.pusilla.DSM.6293 GCA_000688235.1_ASM68823v1
100/100
Thalassospira.profundimaris.WP0211 GCA_000300275.1_ASM30027v1
Elioraea.tepidiphila.DSM.17972 GCA_000378465.1_ASM37846v1
100/100
100/100
Belnapia.moabensis.DSM.16746 GCA_000745835.1_ASM74583v1
100/100
Rubritepida.flocculans.DSM.14296 GCA_000425365.1_ASM42536v1
100/100
Roseomonas.gilardii GCA_001941945.1_ASM194194v1
Acidiphilium.multivorum.AIU301 GCA_000202835.1_ASM20283v1
86.1/98
89.5/99
Acidisphaera.rubrifaciens.HS.AP3 GCA_000964365.1_ASM96436v1
100/100
Granulibacter.bethesdensis.CGDNIH4 GCA_000576145.2_ASM57614v2
100/100
Commensalibacter.intestini.A911 GCA_000231445.2_ASM23144v1
99.9/100
Gluconacetobacter.diazotrophicus.PA1.5 GCA_000067045.1_ASM6704v1
Acetobacter.aceti.NBRC.14818 GCA_000193495.2_ASM19349v2
Gluconobacter.oxydans.DSM.3504 GCA_000583855.1_ASM58385v1
0.609151

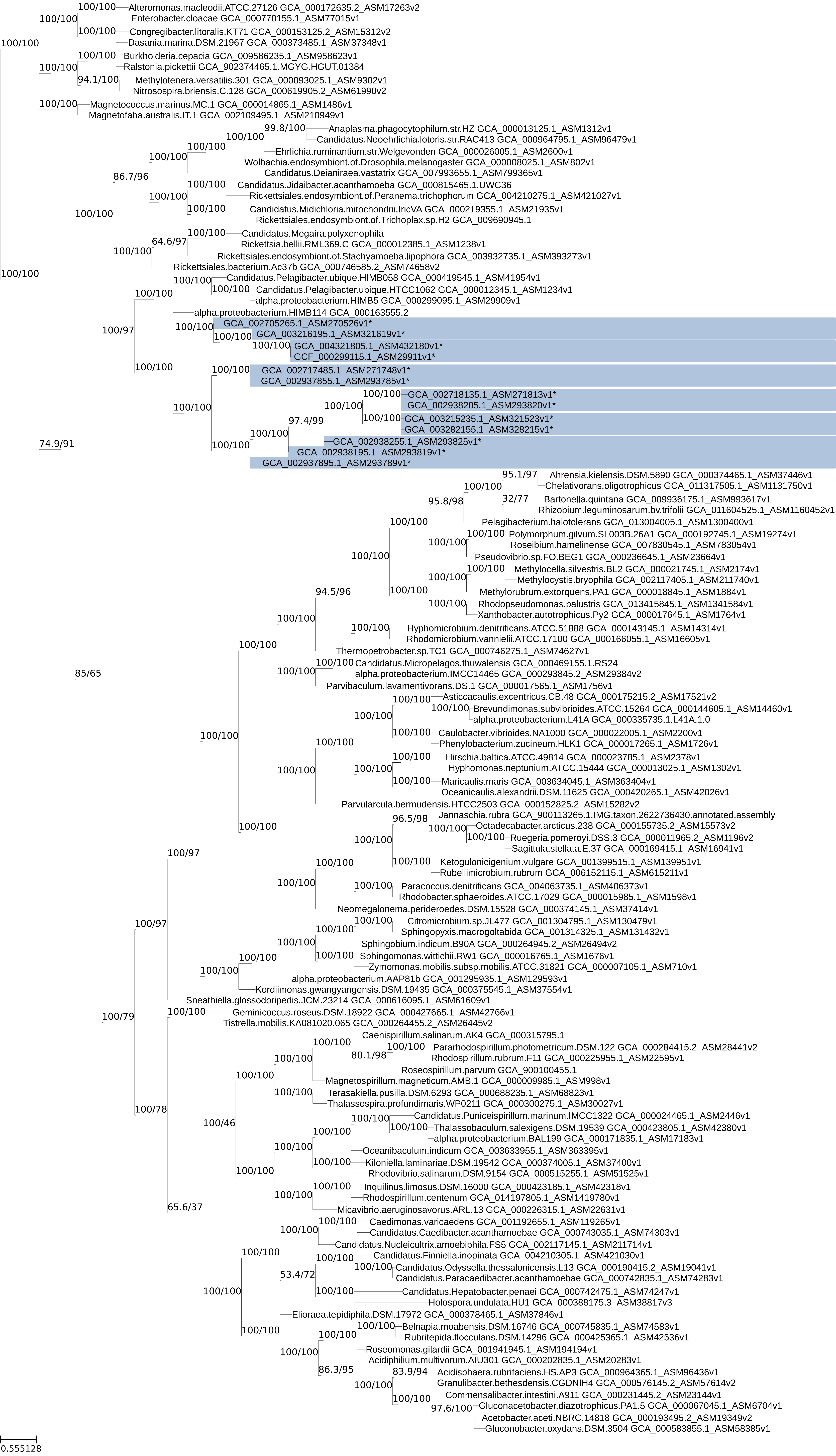

100/100
Alteromonas.macleodii.ATCC.27126 GCA_000172635.2_ASM17263v2
100/100
Enterobacter.cloacae GCA_000770155.1_ASM77015v1
100/100
100/100
Congregibacter.litoralis.KT71 GCA_000153125.2_ASM15312v2
Dasania.marina.DSM.21967 GCA_000373485.1_ASM37348v1
100/100
Burkholderia.cepacia GCA_009586235.1_ASM958623v1
100/100
Ralstonia.pickettii GCA_902374465.1_MGYG.HGUT.01384
94.1/100
Methylotenera.versatilis.301 GCA_000093025.1_ASM9302v1
Nitrosospira.briensis.C.128 GCA_000619905.2_ASM61990v2
100/100
Magnetococcus.marinus.MC.1 GCA_000014865.1_ASM1486v1
Magnetofaba.australis.IT.1 GCA_002109495.1_ASM210949v1
99.8/100
Anaplasma.phagocytophilum.str.HZ GCA_000013125.1_ASM1312v1
100/100
Candidatus.Neoehrlichia.lotoris.str.RAC413 GCA_000964795.1_ASM96479v1
100/100
Ehrlichia.ruminantium.str.Welgevonden GCA_000026005.1_ASM2600v1
100/100
Wolbachia.endosymbiont.of.Drosophila.melanogaster GCA_000008025.1_ASM802v1
86.7/96
Candidatus.Deianiraea.vastatrix GCA_007993655.1_ASM799365v1
100/100
Candidatus.Jidaibacter.acanthamoeba GCA_000815465.1.UWC36
100/100
Rickettsiales.endosymbiont.of.Peranema.trichophorum GCA_004210275.1_ASM421027v1
100/100
Candidatus.Midichloria.mitochondrii.IricVA GCA_000219355.1_ASM21935v1
100/100
Rickettsiales.endosymbiont.of.Trichoplax.sp.H2 GCA_009690945.1
100/100
Candidatus.Megaira.polyxenophila
64.6/97
Rickettsia.bellii.RML369.C GCA_000012385.1_ASM1238v1
100/100
Rickettsiales.endosymbiont.of.Stachyamoeba.lipophora GCA_003932735.1_ASM393273v1
100/100
Rickettsiales.bacterium.Ac37b GCA_000746585.2_ASM74658v2
Candidatus.Pelagibacter.ubique.HIMB058 GCA_000419545.1_ASM41954v1
100/100
100/100
Candidatus.Pelagibacter.ubique.HTCC1062 GCA_000012345.1_ASM1234v1
100/100
alpha.proteobacterium.HIMB5 GCA_000299095.1_ASM29909v1
alpha.proteobacterium.HIMB114 GCA_000163555.2
100/97
100/100
GCA_002705265.1_ASM270526v1*
100/100
GCA_003216195.1_ASM321619v1*
100/100
GCA_004321805.1_ASM432180v1*
100/100
GCF_000299115.1_ASM29911v1*
100/100
GCA_002717485.1_ASM271748v1*
GCA_002937855.1_ASM293785v1*
100/100
GCA_002718135.1_ASM271813v1*
100/100
GCA_002938205.1_ASM293820v1*
100/100
GCA_003215235.1_ASM321523v1*
97.4/99
GCA_003282155.1_ASM328215v1*
100/100
74.9/91
100/100
GCA_002938255.1_ASM293825v1*
GCA_002938195.1_ASM293819v1*
GCA_002937895.1_ASM293789v1*
95.1/97
Ahrensia.kielensis.DSM.5890 GCA_000374465.1_ASM37446v1
100/100
Chelativorans.oligotrophicus GCA_011317505.1_ASM1131750v1
32/77
Bartonella.quintana GCA_009936175.1_ASM993617v1
95.8/98
Rhizobium.leguminosarum.bv.trifolii GCA_011604525.1_ASM1160452v1
100/100
Pelagibacterium.halotolerans GCA_013004005.1_ASM1300400v1
100/100
Polymorphum.gilvum.SL003B.26A1 GCA_000192745.1_ASM19274v1
100/100
Roseibium.hamelinense GCA_007830545.1_ASM783054v1
100/100
Pseudovibrio.sp.FO.BEG1 GCA_000236645.1_ASM23664v1
100/100
Methylocella.silvestris.BL2 GCA_000021745.1_ASM2174v1
100/100
Methylocystis.bryophila GCA_002117405.1_ASM211740v1
100/100
Methylorubrum.extorquens.PA1 GCA_000018845.1_ASM1884v1
94.5/96
100/100
Rhodopseudomonas.palustris GCA_013415845.1_ASM1341584v1
Xanthobacter.autotrophicus.Py2 GCA_000017645.1_ASM1764v1
100/100
100/100
Hyphomicrobium.denitrificans.ATCC.51888 GCA_000143145.1_ASM14314v1
Rhodomicrobium.vannielii.ATCC.17100 GCA_000166055.1_ASM16605v1
100/100
Thermopetrobacter.sp.TC1 GCA_000746275.1_ASM74627v1
100/100
Candidatus.Micropelagos.thuwalensis GCA_000469155.1.RS24
100/100
alpha.proteobacterium.IMCC14465 GCA_000293845.2_ASM29384v2
85/65
Parvibaculum.lavamentivorans.DS.1 GCA_000017565.1_ASM1756v1
100/100
Asticcacaulis.excentricus.CB.48 GCA_000175215.2_ASM17521v2
100/100
Brevundimonas.subvibrioides.ATCC.15264 GCA_000144605.1_ASM14460v1
100/100
alpha.proteobacterium.L41A GCA_000335735.1.L41A.1.0
100/100
Caulobacter.vibrioides.NA1000 GCA_000022005.1_ASM2200v1
100/100
Phenylobacterium.zucineum.HLK1 GCA_000017265.1_ASM1726v1
100/100
100/100
Hirschia.baltica.ATCC.49814 GCA_000023785.1_ASM2378v1
100/100
Hyphomonas.neptunium.ATCC.15444 GCA_000013025.1_ASM1302v1
100/100
Maricaulis.maris GCA_003634045.1_ASM363404v1
Oceanicaulis.alexandrii.DSM.11625 GCA_000420265.1_ASM42026v1
Parvularcula.bermudensis.HTCC2503 GCA_000152825.2_ASM15282v2
96.5/98
Jannaschia.rubra GCA_900113265.1.IMG.taxon.2622736430.annotated.assembly
100/100
100/100
Octadecabacter.arcticus.238 GCA_000155735.2_ASM15573v2
100/100
Ruegeria.pomeroyi.DSS.3 GCA_000011965.2_ASM1196v2
Sagittula.stellata.E.37 GCA_000169415.1_ASM16941v1
100/97
100/100
100/100
Ketogulonicigenium.vulgare GCA_001399515.1_ASM139951v1
Rubellimicrobium.rubrum GCA_006152115.1_ASM615211v1
100/100
100/100
Paracoccus.denitrificans GCA_004063735.1_ASM406373v1
Rhodobacter.sphaeroides.ATCC.17029 GCA_000015985.1_ASM1598v1
100/97
Neomegalonema.perideroedes.DSM.15528 GCA_000374145.1_ASM37414v1
100/100
Citromicrobium.sp.JL477 GCA_001304795.1_ASM130479v1
100/100
Sphingopyxis.macrogoltabida GCA_001314325.1_ASM131432v1
100/100
Sphingobium.indicum.B90A GCA_000264945.2_ASM26494v2
100/100
100/100
Sphingomonas.wittichii.RW1 GCA_000016765.1_ASM1676v1
100/100
Zymomonas.mobilis.subsp.mobilis.ATCC.31821 GCA_000007105.1_ASM710v1
alpha.proteobacterium.AAP81b GCA_001295935.1_ASM129593v1
Kordiimonas.gwangyangensis.DSM.19435 GCA_000375545.1_ASM37554v1
Sneathiella.glossodoripedis.JCM.23214 GCA_000616095.1_ASM61609v1
100/100
Geminicoccus.roseus.DSM.18922 GCA_000427665.1_ASM42766v1
100/79
Tistrella.mobilis.KA081020.065 GCA_000264455.2_ASM26445v2
Caenispirillum.salinarum.AK4 GCA_000315795.1
100/100
100/100
Pararhodospirillum.photometricum.DSM.122 GCA_000284415.2_ASM28441v2
100/100
80.1/98
Rhodospirillum.rubrum.F11 GCA_000225955.1_ASM22595v1
100/100
Roseospirillum.parvum GCA_900100455.1
Magnetospirillum.magneticum.AMB.1 GCA_000009985.1_ASM998v1
100/100
Terasakiella.pusilla.DSM.6293 GCA_000688235.1_ASM68823v1
100/78
Thalassospira.profundimaris.WP0211 GCA_000300275.1_ASM30027v1
100/100
Candidatus.Puniceispirillum.marinum.IMCC1322 GCA_000024465.1_ASM2446v1
100/46
100/100
Thalassobaculum.salexigens.DSM.19539 GCA_000423805.1_ASM42380v1
100/100
alpha.proteobacterium.BAL199 GCA_000171835.1_ASM17183v1
100/100
Oceanibaculum.indicum GCA_003633955.1_ASM363395v1
100/100
Kiloniella.laminariae.DSM.19542 GCA_000374005.1_ASM37400v1
100/100
Rhodovibrio.salinarum.DSM.9154 GCA_000515255.1_ASM51525v1
100/100
Inquilinus.limosus.DSM.16000 GCA_000423185.1_ASM42318v1
100/100
Rhodospirillum.centenum GCA_014197805.1_ASM1419780v1
65.6/37
Micavibrio.aeruginosavorus.ARL.13 GCA_000226315.1_ASM22631v1
100/100
Caedimonas.varicaedens GCA_001192655.1_ASM119265v1
100/100
Candidatus.Caedibacter.acanthamoebae GCA_000743035.1_ASM74303v1
100/100
Candidatus.Nucleicultrix.amoebiphila.FS5 GCA_002117145.1_ASM211714v1
100/100
Candidatus.Finniella.inopinata GCA_004210305.1_ASM421030v1
100/100
Candidatus.Odyssella.thessalonicensis.L13 GCA_000190415.2_ASM19041v1
53.4/72
Candidatus.Paracaedibacter.acanthamoebae GCA_000742835.1_ASM74283v1
100/100
100/100
Candidatus.Hepatobacter.penaei GCA_000742475.1_ASM74247v1
Holospora.undulata.HU1 GCA_000388175.3_ASM38817v3
Elioraea.tepidiphila.DSM.17972 GCA_000378465.1_ASM37846v1
100/100
100/100
Belnapia.moabensis.DSM.16746 GCA_000745835.1_ASM74583v1
100/100
Rubritepida.flocculans.DSM.14296 GCA_000425365.1_ASM42536v1
100/100
Roseomonas.gilardii GCA_001941945.1_ASM194194v1
Acidiphilium.multivorum.AIU301 GCA_000202835.1_ASM20283v1
86.3/95
83.9/94
Acidisphaera.rubrifaciens.HS.AP3 GCA_000964365.1_ASM96436v1
100/100
Granulibacter.bethesdensis.CGDNIH4 GCA_000576145.2_ASM57614v2
100/100
Commensalibacter.intestini.A911 GCA_000231445.2_ASM23144v1
97.6/100
Gluconacetobacter.diazotrophicus.PA1.5 GCA_000067045.1_ASM6704v1
Acetobacter.aceti.NBRC.14818 GCA_000193495.2_ASM19349v2
Gluconobacter.oxydans.DSM.3504 GCA_000583855.1_ASM58385v1
0.555128

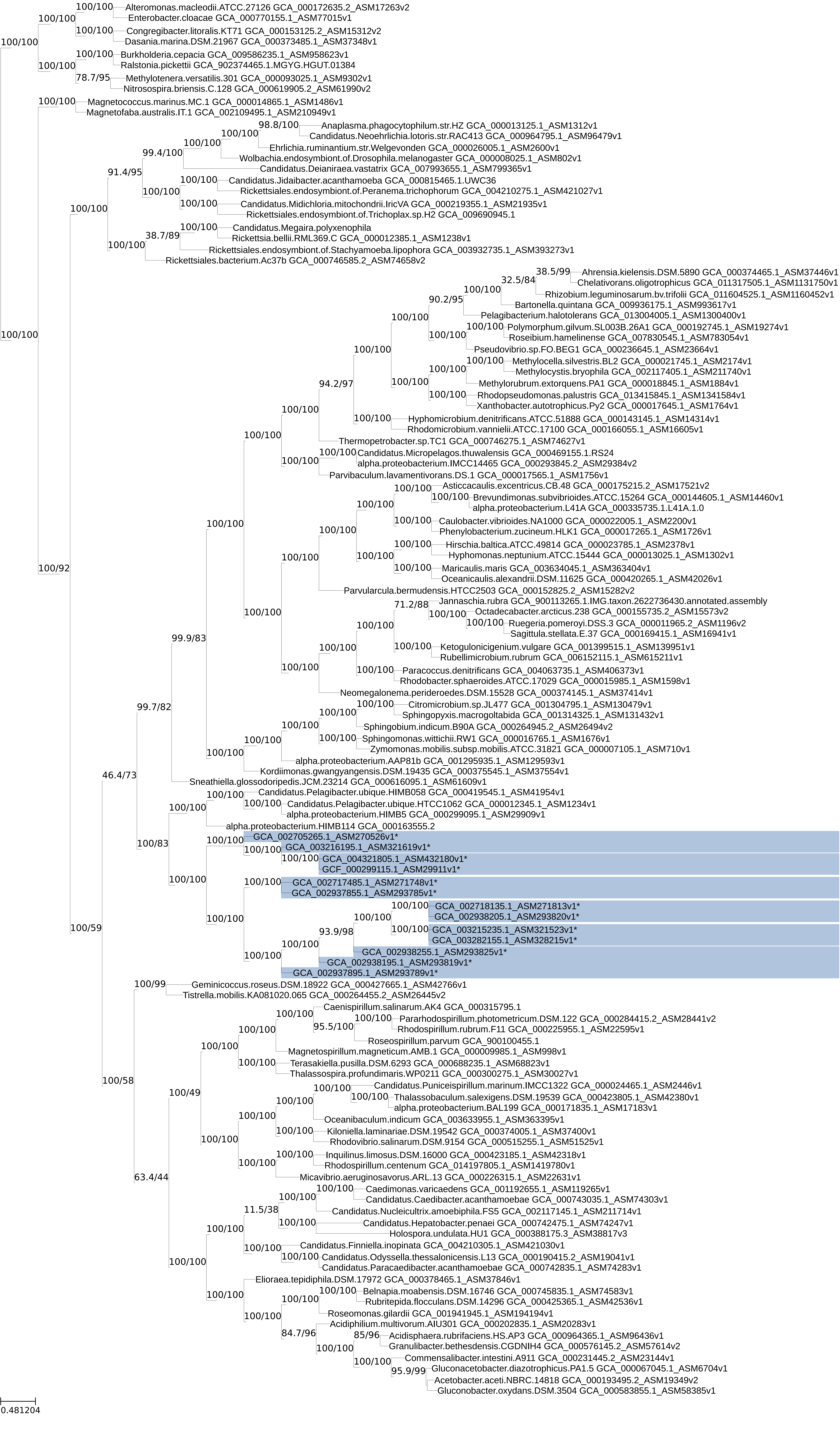

Alteromonas.macleodii.ATCC.27126 GCA_000172635.2_ASM17263v2
Enterobacter.cloacae GCA_000770155.1_ASM77015v1
Congregibacter.litoralis.KT71 GCA_000153125.2_ASM15312v2
Dasania.marina.DSM.21967 GCA_000373485.1_ASM37348v1
Burkholderia.cepacia GCA_009586235.1_ASM958623v1
Ralstonia.pickettii GCA_902374465.1.MGYG.HGUT.01384
Methylotenera.versatilis.301 GCA_000093025.1_ASM9302v1
Nitrosospira.briensis.C.128 GCA_000619905.2_ASM61990v2
Magnetococcus.marinus.MC.1 GCA_000014865.1_ASM1486v1
Magnetofaba.australis.IT.1 GCA_002109495.1_ASM210949v1
Anaplasma.phagocytophilum.str.HZ GCA_000013125.1_ASM1312v1
Candidatus.Neoehrlichia.lotoris.str.RAC413 GCA_000964795.1_ASM96479v1
Ehrlichia.ruminantium.str.Welgevonden GCA_000026005.1_ASM2600v1
Wolbachia.endosymbiont.of.Drosophila.melanogaster GCA_000008025.1_ASM802v1
Candidatus.Deianiraea.vastatrix GCA_007993655.1_ASM799365v1
Candidatus.Jidaibacter.acanthamoeba GCA_000815465.1.UWC36
Rickettsiales.endosymbiont.of.Peranema.trichophorum GCA_004210275.1_ASM421027v1
Candidatus.Midichloria.mitochondrii.IricVA GCA_000219355.1_ASM21935v1
Rickettsiales.endosymbiont.of.Trichoplax.sp.H2 GCA_009690945.1
Candidatus.Megaira.polyxenophila
Rickettsia.bellii.RML369.C GCA_000012385.1_ASM1238v1
Rickettsiales.endosymbiont.of.Stachyamoeba.lipophora GCA_003932735.1_ASM393273v1
Rickettsiales.bacterium.Ac37b GCA_000746585.2_ASM74658v2
Ahrensia.kielensis.DSM.5890 GCA_000374465.1_ASM37446v1
Chelativorans.oligotrophicus GCA_011317505.1_ASM1131750v1
Rhizobium.leguminosarum.bv.trifolii GCA_011604525.1_ASM1160452v1
Bartonella.quintana GCA_009936175.1_ASM993617v1
Pelagibacterium.halotolerans GCA_013004005.1_ASM1300400v1
Polymorphum.gilvum.SL003B.26A1 GCA_000192745.1_ASM19274v1
Roseibium.hamelinense GCA_007830545.1_ASM783054v1
Pseudovibrio.sp.FO.BEG1 GCA_000236645.1_ASM23664v1
Methylocella.silvestris.BL2 GCA_000021745.1_ASM2174v1
Methylocystis.bryophila GCA_002117405.1_ASM211740v1
Methylorubrum.extorquens.PA1 GCA_000018845.1_ASM1884v1
Rhodopseudomonas.palustris GCA_013415845.1_ASM1341584v1
Xanthobacter.autotrophicus.Py2 GCA_000017645.1_ASM1764v1
Hyphomicrobium.denitrificans.ATCC.51888 GCA_000143145.1_ASM14314v1
Rhodomicrobium.vannielii.ATCC.17100 GCA_000166055.1_ASM16605v1
Thermopetrobacter.sp.TC1 GCA_000746275.1_ASM74627v1
Candidatus.Micropelagos.thuwalensis GCA_000469155.1.RS24
alpha.proteobacterium.IMCC14465 GCA_000293845.2_ASM29384v2
Parvibaculum.lavamentivorans.DS.1 GCA_000017565.1_ASM1756v1
Asticcacaulis.excentricus.CB.48 GCA_000175215.2_ASM17521v2
Brevundimonas.subvibrioides.ATCC.15264 GCA_000144605.1_ASM14460v1
alpha.proteobacterium.L41A GCA_000335735.1.L41A.1.0
Caulobacter.vibrioides.NA1000 GCA_000022005.1_ASM2200v1
Phenylobacterium.zucineum.HLK1 GCA_000017265.1_ASM1726v1
Hirschia.baltica.ATCC.49814 GCA_000023785.1_ASM2378v1
Hyphomonas.neptunium.ATCC.15444 GCA_000013025.1_ASM1302v1
Maricaulis.maris GCA_003634045.1_ASM363404v1
Oceanicaulis.alexandrii.DSM.11625 GCA_000420265.1_ASM42026v1
Parvularcula.bermudensis.HTCC2503 GCA_000152825.2_ASM15282v2
Jannaschia.rubra GCA_900113265.1.IMG.taxon.2622736430.annotated.assembly
Octadecabacter.arcticus.238 GCA_000155735.2_ASM15573v2
Ruegeria.pomeroyi.DSS.3 GCA_000011965.2_ASM1196v2
Sagittula.stellata.E.37 GCA_000169415.1_ASM16941v1
Ketogulonicigenium.vulgare GCA_001399515.1_ASM139951v1
Rubellimicrobium.rubrum GCA_006152115.1_ASM615211v1
Paracoccus.denitrificans GCA_004063735.1_ASM406373v1
Rhodobacter.sphaeroides.ATCC.17029 GCA_000015985.1_ASM1598v1
Neomegalonema.perideroedes.DSM.15528 GCA_000374145.1_ASM37414v1
Citromicrobium.sp.JL477 GCA_001304795.1_ASM130479v1
Sphingopyxis.macrogoltabida GCA_001314325.1_ASM131432v1
Sphingobium.indicum.B90A GCA_000264945.2_ASM26494v2
Sphingomonas.wittichii.RW1 GCA_000016765.1_ASM1676v1
Zymomonas.mobilis.subsp.mobilis.ATCC.31821 GCA_000007105.1_ASM710v1
alpha.proteobacterium.AAP81b GCA_001295935.1_ASM129593v1
Kordiimonas.gwangyangensis.DSM.19435 GCA_000375545.1_ASM37554v1
Sneathiella.glossodoripedis.JCM.23214 GCA_000616095.1_ASM61609v1
Candidatus.Pelagibacter.ubique.HIMB058 GCA_000419545.1_ASM41954v1
Candidatus.Pelagibacter.ubique.HTCC1062 GCA_000012345.1_ASM1234v1
alpha.proteobacterium.HIMB5 GCA_000299095.1_ASM29909v1
alpha.proteobacterium.HIMB114 GCA_000163555.2
GCA_002705265.1_ASM270526v1*
GCA_003216195.1_ASM321619v1*
GCA_004321805.1_ASM432180v1*
GCF_000299115.1_ASM29911v1*
GCA_002717485.1_ASM271748v1*
GCA_002937855.1_ASM293785v1*
GCA_002718135.1_ASM271813v1*
GCA_002938205.1_ASM293820v1*
GCA_003215235.1_ASM321523v1*
GCA_003282155.1_ASM328215v1*
GCA_002938255.1_ASM293825v1*
GCA_002938195.1_ASM293819v1*
GCA_002937895.1_ASM293789v1*
Geminicoccus.roseus.DSM.18922 GCA_000427665.1_ASM42766v1
Tistrella.mobilis.KA081020.065 GCA_000264455.2_ASM26445v2
Caenispirillum.salinarum.AK4 GCA_000315795.1
Pararhodospirillum.photometricum.DSM.122 GCA_000284415.2_ASM28441v2
Rhodospirillum.rubrum.F11 GCA_000225955.1_ASM22595v1
Roseospirillum.parvum GCA_900100455.1
Magnetospirillum.magneticum.AMB.1 GCA_000009985.1_ASM998v1
Terasakiella.pusilla.DSM.6293 GCA_000688235.1_ASM68823v1
Thalassospira.profundimaris.WP0211 GCA_000300275.1_ASM30027v1
Candidatus.Puniceispirillum.marinum.IMCC1322 GCA_000024465.1_ASM2446v1
Thalassobaculum.salexigens.DSM.19539 GCA_000423805.1_ASM42380v1
alpha.proteobacterium.BAL199 GCA_000171835.1_ASM17183v1
Oceanibaculum.indicum GCA_003633955.1_ASM363395v1
Kiloniella.laminariae.DSM.19542 GCA_000374005.1_ASM37400v1
Rhodovibrio.salinarum.DSM.9154 GCA_000515255.1_ASM51525v1
Inquilinus.limosus.DSM.16000 GCA_000423185.1_ASM42318v1
Rhodospirillum.centenum GCA_014197805.1_ASM1419780v1
Micavibrio.aeruginosavorus.ARL.13 GCA_000226315.1_ASM22631v1
Caedimonas.varicaedens GCA_001192655.1_ASM119265v1
Candidatus.Caedibacter.acanthamoebae GCA_000743035.1_ASM74303v1
Candidatus.Nucleicultrix.amoebiphila.FS5 GCA_002117145.1_ASM211714v1
Candidatus.Hepatobacter.penaei GCA_000742475.1_ASM74247v1
Holospora.undulata.HU1 GCA_000388175.3_ASM38817v3
Candidatus.Finniella.inopinata GCA_004210305.1_ASM421030v1
Candidatus.Odyssella.thessalonicensis.L13 GCA_000190415.2_ASM19041v2
Candidatus.Paracaedibacter.acanthamoebae GCA_000742835.1_ASM74283v1
Elioraea.tepidiphila.DSM.17972 GCA_000378465.1_ASM37846v1
Belnapia.moabensis.DSM.16746 GCA_000745835.1_ASM74583v1
Rubritepida.flocculans.DSM.14296 GCA_000425365.1_ASM42536v1
Roseomonas.gilardii GCA_001941945.1_ASM194194v1
Acidiphilium.multivorum.AIU301 GCA_000202835.1_ASM20283v1
Acidisphaera.rubrifaciens.HS.AP3 GCA_000964365.1_ASM96436v1
Granulibacter.bethesdensis.CGDNIH4 GCA_000576145.2_ASM57614v2
Commensalibacter.intestini.A911 GCA_000231445.2_ASM23144v1
Gluconacetobacter.diazotrophicus.PA1.5 GCA_000067045.1_ASM6704v1
Acetobacter.aceti.NBRC.14818 GCA_000193495.2_ASM19349v2
Gluconobacter.oxydans.DSM.3504 GCA_000583855.1_ASM58385v1
0.481204

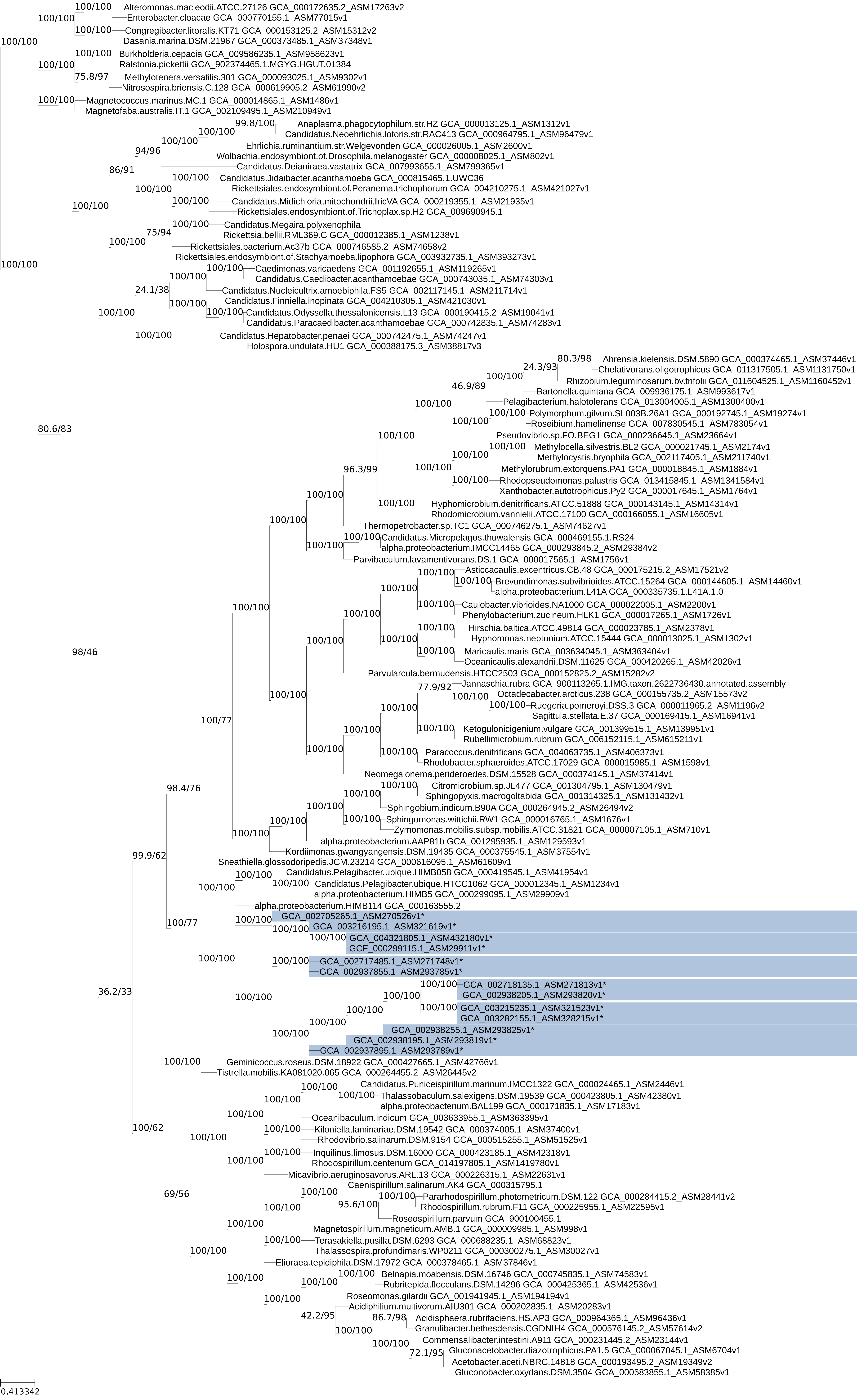

100/100
Alteromonas.macleodii.ATCC.27126 GCA_000172635.2_ASM17263v2
100/100
Enterobacter.cloacae GCA_000770155.1_ASM77015v1
100/100
100/100
Congregibacter.litoralis.KT71 GCA_000153125.2_ASM15312v2
Dasania.marina.DSM.21967 GCA_000373485.1_ASM37348v1
100/100
Burkholderia.cepacia GCA_009586235.1_ASM958623v1
100/100
Ralstonia.pickettii GCA_902374465.1.MGYG.HGUT.01384
75.8/97
Methylotenera.versatilis.301 GCA_000093025.1_ASM9302v1
Nitrosospira.briensis.C.128 GCA_000619905.2_ASM61990v2
100/100
Magnetococcus.marinus.MC.1 GCA_000014865.1_ASM1486v1
Magnetofaba.australis.IT.1 GCA_002109495.1_ASM210949v1
99.8/100
Anaplasma.phagocytophilum.str.HZ GCA_000013125.1_ASM1312v1
100/100
Candidatus.Neoehrlichia.lotoris.str.RAC413 GCA_000964795.1_ASM96479v1
100/100
Ehrlichia.ruminantium.str.Welgevonden GCA_000026005.1_ASM2600v1
94/96
Wolbachia.endosymbiont.of.Drosophila.melanogaster GCA_000008025.1_ASM802v1
Candidatus.Deianiraea.vastatrix GCA_007993655.1_ASM799365v1
86/91
100/100
Candidatus.Jidaibacter.acanthamoeba GCA_000815465.1.UWC36
100/100
Rickettsiales.endosymbiont.of.Peranema.trichophorum GCA_004210275.1_ASM421027v1
100/100
Candidatus.Midichloria.mitochondrii.IricVA GCA_000219355.1_ASM21935v1
100/100
Rickettsiales.endosymbiont.of.Trichoplax.sp.H2 GCA_009690945.1
100/100
Candidatus.Megaira.polyxenophila
75/94
Rickettsia.bellii.RML369.C GCA_000012385.1_ASM1238v1
100/100
Rickettsiales.bacterium.Ac37b GCA_000746585.2_ASM74658v2
Rickettsiales.endosymbiont.of.Stachyamoeba.lipophora GCA_003932735.1_ASM393273v1
100/100
100/100
Caedimonas.varicaedens GCA_001192655.1_ASM119265v1
100/100
Candidatus.Caedibacter.acanthamoebae GCA_000743035.1_ASM74303v1
24.1/38
Candidatus.Nucleicultrix.amoebiphila.FS5 GCA_002117145.1_ASM211714v1
100/100
Candidatus.Finniella.inopinata GCA_004210305.1_ASM421030v1
100/100
100/100
Candidatus.Odyssella.thessalonicensis.L13 GCA_000190415.2_ASM19041v1
Candidatus.Paracaedibacter.acanthamoebae GCA_000742835.1_ASM74283v1
100/100
Candidatus.Hepatobacter.penaei GCA_000742475.1_ASM74247v1
Holospora.undulata.HU1 GCA_000388175.3_ASM38817v3
80.3/98
Ahrensia.kielensis.DSM.5890 GCA_000374465.1_ASM37446v1
24.3/93
Chelativorans.oligotrophicus GCA_011317505.1_ASM1131750v1
100/100
Rhizobium.leguminosarum.bv.trifolii GCA_011604525.1_ASM1160452v1
46.9/89
Bartonella.quintana GCA_009936175.1_ASM993617v1
Pelagibacterium.halotolerans GCA_013004005.1_ASM1300400v1
100/100
100/100
Polymorphum.gilvum.SL003B.26A1 GCA_000192745.1_ASM19274v1
100/100
Roseibium.hamelinense GCA_007830545.1_ASM783054v1
80.6/83
Pseudovibrio.sp.FO.BEG1 GCA_000236645.1_ASM23664v1
100/100
100/100
Methylocella.silvestris.BL2 GCA_000021745.1_ASM2174v1
100/100
Methylocystis.bryophila GCA_002117405.1_ASM211740v1
96.3/99
100/100
Methylorubrum.extorquens.PA1 GCA_000018845.1_ASM1884v1
100/100
Rhodopseudomonas.palustris GCA_013415845.1_ASM1341584v1
100/100
Xanthobacter.autotrophicus.Py2 GCA_000017645.1_ASM1764v1
100/100
Hyphomicrobium.denitrificans.ATCC.51888 GCA_000143145.1_ASM14314v1
Rhodomicrobium.vannielii.ATCC.17100 GCA_000166055.1_ASM16605v1
100/100
Thermopetrobacter.sp.TC1 GCA_000746275.1_ASM74627v1
100/100
Candidatus.Micropelagos.thuwalensis GCA_000469155.1.RS24
100/100
alpha.proteobacterium.IMCC14465 GCA_000293845.2_ASM29384v2
Parvibaculum.lavamentivorans.DS.1 GCA_000017565.1_ASM1756v1
100/100
Asticcacaulis.excentricus.CB.48 GCA_000175215.2_ASM17521v2
100/100
100/100
Brevundimonas.subvibrioides.ATCC.15264 GCA_000144605.1_ASM14460v1
alpha.proteobacterium.L41A GCA_000335735.1.L41A.1.0
100/100
100/100
Caulobacter.vibrioides.NA1000 GCA_000022005.1_ASM2200v1
100/100
Phenylobacterium.zucineum.HLK1 GCA_000017265.1_ASM1726v1
100/100
Hirschia.baltica.ATCC.49814 GCA_000023785.1_ASM2378v1
100/100
100/100
Hyphomonas.neptunium.ATCC.15444 GCA_000013025.1_ASM1302v1
98/46
100/100
Maricaulis.maris GCA_003634045.1_ASM363404v1
Oceanicaulis.alexandrii.DSM.11625 GCA_000420265.1_ASM42026v1
Parvularcula.bermudensis.HTCC2503 GCA_000152825.2_ASM15282v2
77.9/92
Jannaschia.rubra GCA_900113265.1.IMG.taxon.2622736430.annotated.assembly
100/100
100/100
Octadecabacter.arcticus.238 GCA_000155735.2_ASM15573v2
100/100
Ruegeria.pomeroyi.DSS.3 GCA_000011965.2_ASM1196v2
100/100
Sagittula.stellata.E.37 GCA_000169415.1_ASM16941v1
100/77
100/100
Ketogulonicigenium.vulgare GCA_001399515.1_ASM139951v1
100/100
Rubellimicrobium.rubrum GCA_006152115.1_ASM615211v1
100/100
100/100
Paracoccus.denitrificans GCA_004063735.1_ASM406373v1
Rhodobacter.sphaeroides.ATCC.17029 GCA_000015985.1_ASM1598v1
Neomegalonema.perideroedes.DSM.15528 GCA_000374145.1_ASM37414v1
98.4/76
100/100
Citromicrobium.sp.JL477 GCA_001304795.1_ASM130479v1
100/100
Sphingopyxis.macrogoltabida GCA_001314325.1_ASM131432v1
100/100
Sphingobium.indicum.B90A GCA_000264945.2_ASM26494v2
100/100
100/100
Sphingomonas.wittichii.RW1 GCA_000016765.1_ASM1676v1
Zymomonas.mobilis.subsp.mobilis.ATCC.31821 GCA_000007105.1_ASM710v1
100/100
alpha.proteobacterium.AAP81b GCA_001295935.1_ASM129593v1
Kordiimonas.gwangyangensis.DSM.19435 GCA_000375545.1_ASM37554v1
99.9/62
Sneathiella.glossodoripedis.JCM.23214 GCA_000616095.1_ASM61609v1
100/100
Candidatus.Pelagibacter.ubique.HIMB058 GCA_000419545.1_ASM41954v1
100/100
100/100
Candidatus.Pelagibacter.ubique.HTCC1062 GCA_000012345.1_ASM1234v1
alpha.proteobacterium.HIMB5 GCA_000299095.1_ASM29909v1
alpha.proteobacterium.HIMB114 GCA_000163555.2
100/100
GCA_002705265.1_ASM270526v1*
100/100
GCA_003216195.1_ASM321619v1*
100/77
100/100
GCA_004321805.1_ASM432180v1*
GCF_000299115.1_ASM29911v1*
100/100
100/100
GCA_002717485.1_ASM271748v1*
GCA_002937855.1_ASM293785v1*
100/100
GCA_002718135.1_ASM271813v1*
100/100
GCA_002938205.1_ASM293820v1*
36.2/33
100/100
100/100
GCA_003215235.1_ASM321523v1*
100/100
GCA_003282155.1_ASM328215v1*
100/100
GCA_002938255.1_ASM293825v1*
100/100
GCA_002938195.1_ASM293819v1*
GCA_002937895.1_ASM293789v1*
100/100
Geminicoccus.roseus.DSM.18922 GCA_000427665.1_ASM42766v1
Tistrella.mobilis.KA081020.065 GCA_000264455.2_ASM26445v2
100/100
Candidatus.Puniceispirillum.marinum.IMCC1322 GCA_000024465.1_ASM2446v1
100/100
100/100
Thalassobaculum.salexigens.DSM.19539 GCA_000423805.1_ASM42380v1
alpha.proteobacterium.BAL199 GCA_000171835.1_ASM17183v1
100/100
Oceanibaculum.indicum GCA_003633955.1_ASM363395v1
100/62
100/100
Kiloniella.laminariae.DSM.19542 GCA_000374005.1_ASM37400v1
100/100
Rhodovibrio.salinarum.DSM.9154 GCA_000515255.1_ASM51525v1
100/100
Inquilinus.limosus.DSM.16000 GCA_000423185.1_ASM42318v1
100/100
Rhodospirillum.centenum GCA_014197805.1_ASM1419780v1
Micavibrio.aeruginosavorus.ARL.13 GCA_000226315.1_ASM22631v1
Caenispirillum.salinarum.AK4 GCA_000315795.1
100/100
100/100
Pararhodospirillum.photometricum.DSM.122 GCA_000284415.2_ASM28441v2
95.6/100
Rhodospirillum.rubrum.F11 GCA_000225955.1_ASM22595v1
69/56
100/100
Roseospirillum.parvum GCA_900100455.1
100/100
Magnetospirillum.magneticum.AMB.1 GCA_000009985.1_ASM998v1
100/100
Terasakiella.pusilla.DSM.6293 GCA_000688235.1_ASM68823v1
100/100
Thalassospira.profundimaris.WP0211 GCA_000300275.1_ASM30027v1
Elioraea.tepidiphila.DSM.17972 GCA_000378465.1_ASM37846v1
100/100
Belnapia.moabensis.DSM.16746 GCA_000745835.1_ASM74583v1
100/100
100/100
Rubritepida.flocculans.DSM.14296 GCA_000425365.1_ASM42536v1
100/100
Roseomonas.gilardii GCA_001941945.1_ASM194194v1
Acidiphilium.multivorum.AIU301 GCA_000202835.1_ASM20283v1
42.2/95
86.7/98
Acidisphaera.rubrifaciens.HS.AP3 GCA_000964365.1_ASM96436v1
100/100
Granulibacter.bethesdensis.CGDNIH4 GCA_000576145.2_ASM57614v2
Commensalibacter.intestini.A911 GCA_000231445.2_ASM23144v1
100/100
Gluconacetobacter.diazotrophicus.PA1.5 GCA_000067045.1_ASM6704v1
72.1/95
Acetobacter.aceti.NBRC.14818 GCA_000193495.2_ASM19349v2
Gluconobacter.oxydans.DSM.3504 GCA_000583855.1_ASM58385v1
0.413342

### Group K

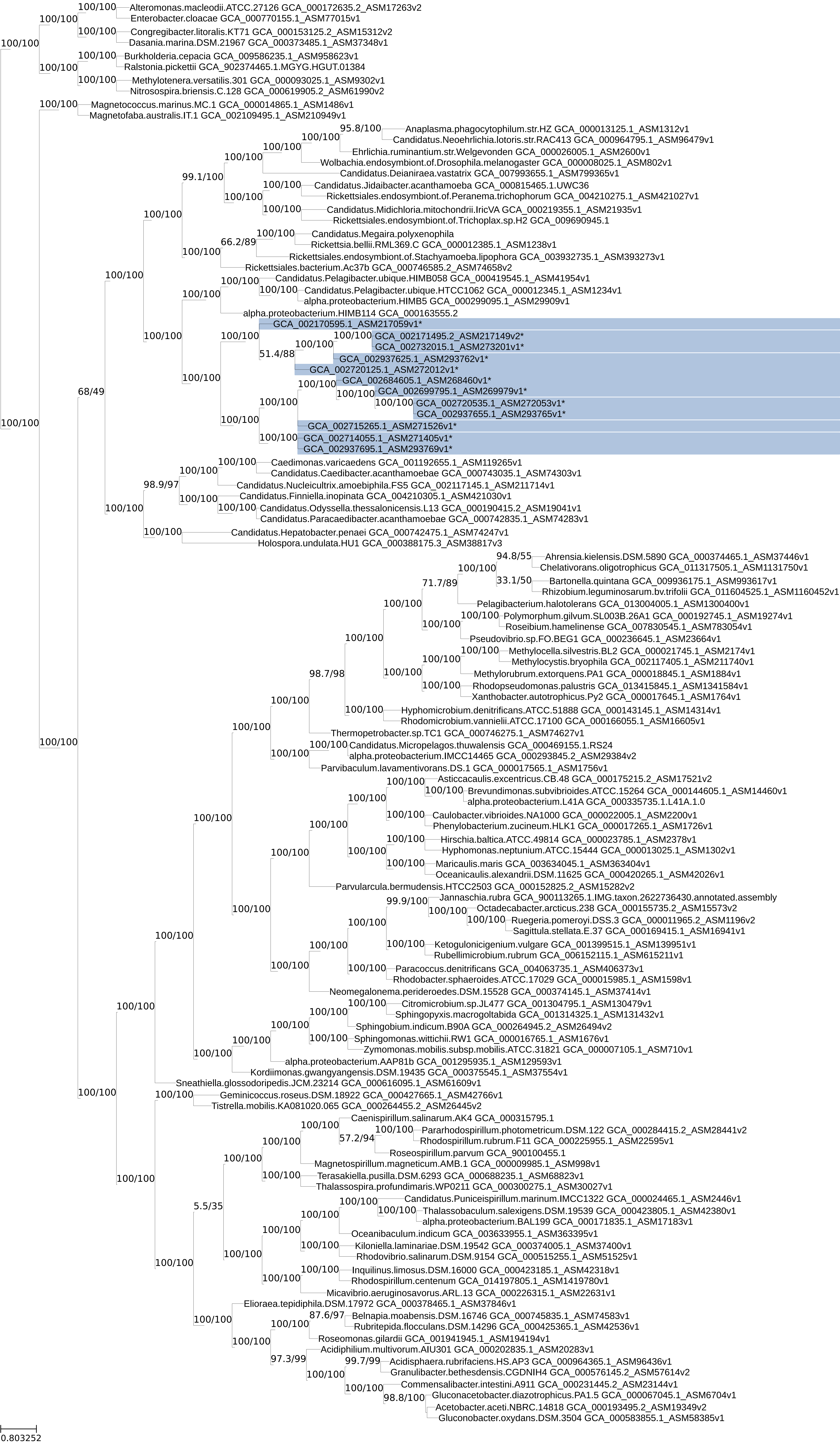

Alteromonas.macleodii.ATCC.27126 GCA_000172635.2_ASM17263v2
Enterobacter.cloacae GCA_000770155.1_ASM77015v1
Congregibacter.litoralis.KT71 GCA_000153125.2_ASM15312v2
Dasania.marina.DSM.21967 GCA_000373485.1_ASM37348v1
Burkholderia.cepacia GCA_009586235.1_ASM958623v1
Ralstonia.pickettii GCA_902374465.1.MGYG.HGUT.01384
Methylotenera.versatilis.301 GCA_000093025.1_ASM9302v1
Nitrosospira.briensis.C.128 GCA_000619905.2_ASM61990v2
Magnetococcus.marinus.MC.1 GCA_000014865.1_ASM1486v1
Magnetofaba.australis.IT.1 GCA_002109495.1_ASM210949v1
Anaplasma.phagocytophilum.str.HZ GCA_000013125.1_ASM1312v1
Candidatus.Neoehrlichia.lotoris.str.RAC413 GCA_000964795.1_ASM96479v1
Ehrlichia.ruminantium.str.Welgevonden GCA_000026005.1_ASM2600v1
Wolbachia.endosymbiont.of.Drosophila.melanogaster GCA_000008025.1_ASM802v1
Candidatus.Deianiraea.vastatrix GCA_007993655.1_ASM799365v1
Candidatus.Jidaibacter.acanthamoeba GCA_000815465.1.UWC36
Rickettsiales.endosymbiont.of.Peranema.trichophorum GCA_004210275.1_ASM421027v1
Candidatus.Midichloria.mitochondrii.IricVA GCA_000219355.1_ASM21935v1
Rickettsiales.endosymbiont.of.Trichoplax.sp.H2 GCA_009690945.1
Candidatus.Megaira.polyxenophila
Rickettsia.bellii.RML369.C GCA_000012385.1_ASM1238v1
Rickettsiales.endosymbiont.of.Stachyamoeba.lipophora GCA_003932735.1_ASM393273v1
Rickettsiales.bacterium.Ac37b GCA_000746585.2_ASM74658v2
Candidatus.Pelagibacter.ubique.HIMB058 GCA_000419545.1_ASM41954v1
Candidatus.Pelagibacter.ubique.HTCC1062 GCA_000012345.1_ASM1234v1
alpha.proteobacterium.HIMB5 GCA_000299095.1_ASM29909v1
alpha.proteobacterium.HIMB114 GCA_000163555.2
GCA_002170595.1_ASM217059v1*
GCA_002171495.2_ASM217149v2*
GCA_002732015.1_ASM273201v1*
GCA_002937625.1_ASM293762v1*
GCA_002720125.1_ASM272012v1*
GCA_002684605.1_ASM268460v1*
GCA_002699795.1_ASM269979v1*
GCA_002720535.1_ASM272053v1*
GCA_002937655.1_ASM293765v1*
GCA_002715265.1_ASM271526v1*
GCA_002714055.1_ASM271405v1*
GCA_002937695.1_ASM293769v1*
Caedimonas.varicaedens GCA_001192655.1_ASM119265v1
Candidatus.Caedibacter.acanthamoebae GCA_000743035.1_ASM74303v1
Candidatus.Nucleicultrix.amoebiphila.FS5 GCA_002117145.1_ASM211714v1
Candidatus.Finniella.inopinata GCA_004210305.1_ASM421030v1
Candidatus.Odyssella.thessalonicensis.L13 GCA_000190415.2_ASM19041v1
Candidatus.Paracaedibacter.acanthamoebae GCA_000742835.1_ASM74283v1
Candidatus.Hepatobacter.penaei GCA_000742475.1_ASM74247v1
Holospora.undulata.HU1 GCA_000388175.3_ASM38817v3
Ahrensia.kielensis.DSM.5890 GCA_000374465.1_ASM37446v1
Chelativorans.oligotrophicus GCA_011317505.1_ASM1131750v1
Bartonella.quintana GCA_009936175.1_ASM993617v1
Rhizobium.leguminosarum.bv.trifolii GCA_011604525.1_ASM1160452v1
Pelagibacterium.halotolerans GCA_013004005.1_ASM1300400v1
Polymorphum.gilvum.SL003B.26A1 GCA_000192745.1_ASM19274v1
Roseibium.hamelinense GCA_007830545.1_ASM783054v1
Pseudovibrio.sp.FO.BEG1 GCA_000236645.1_ASM23664v1
Methylocella.silvestris.BL2 GCA_000021745.1_ASM2174v1
Methylocystis.bryophila GCA_002117405.1_ASM211740v1
Methylorubrum.extorquens.PA1 GCA_000018845.1_ASM1884v1
Rhodopseudomonas.palustris GCA_013415845.1_ASM1341584v1
Xanthobacter.autotrophicus.Py2 GCA_000017645.1_ASM1764v1
Hyphomicrobium.denitrificans.ATCC.51888 GCA_000143145.1_ASM14314v1
Rhodomicrobium.vannielii.ATCC.17100 GCA_000166055.1_ASM16605v1
Thermopetrobacter.sp.TC1 GCA_000746275.1_ASM74627v1
Candidatus.Micropelagos.thuwalensis GCA_000469155.1.RS24
alpha.proteobacterium.IMCC14465 GCA_000293845.2_ASM29384v2
Parvibaculum.lavamentivorans.DS.1 GCA_000017565.1_ASM1756v1
Asticcacaulis.excentricus.CB.48 GCA_000175215.2_ASM17521v2
Brevundimonas.subvibrioides.ATCC.15264 GCA_000144605.1_ASM14460v1
alpha.proteobacterium.L41A GCA_000335735.1.L41A.1.0
Caulobacter.vibrioides.NA1000 GCA_000022005.1_ASM2200v1
Phenylobacterium.zucineum.HLK1 GCA_000017265.1_ASM1726v1
Hirschia.baltica.ATCC.49814 GCA_000023785.1_ASM2378v1
Hyphomonas.neptunium.ATCC.15444 GCA_000013025.1_ASM1302v1
Maricaulis.maris GCA_003634045.1_ASM363404v1
Oceanicaulis.alexandrii.DSM.11625 GCA_000420265.1_ASM42026v1
Parvularcula.bermudensis.HTCC2503 GCA_000152825.2_ASM15282v2
Jannaschia.rubra GCA_900113265.1.IMG.taxon.2622736430.annotated.assembly
Octadecabacter.arcticus.238 GCA_000155735.2_ASM15573v2
Ruegeria.pomeroyi.DSS.3 GCA_000011965.2_ASM1196v2
Sagittula.stellata.E.37 GCA_000169415.1_ASM16941v1
Ketogulonicigenium.vulgare GCA_001399515.1_ASM139951v1
Rubellimicrobium.rubrum GCA_006152115.1_ASM615211v1
Paracoccus.denitrificans GCA_004063735.1_ASM406373v1
Rhodobacter.sphaeroides.ATCC.17029 GCA_000015985.1_ASM1598v1
Neomegalonema.perideroedes.DSM.15528 GCA_000374145.1_ASM37414v1
Citromicrobium.sp.JL477 GCA_001304795.1_ASM130479v1
Sphingopyxis.macrogoltabida GCA_001314325.1_ASM131432v1
Sphingobium.indicum.B90A GCA_000264945.2_ASM26494v2
Sphingomonas.wittichii.RW1 GCA_000016765.1_ASM1676v1
Zymomonas.mobilis.subsp.mobilis.ATCC.31821 GCA_000007105.1_ASM710v1
alpha.proteobacterium.AAP81b GCA_001295935.1_ASM129593v1
Kordiimonas.gwangyangensis.DSM.19435 GCA_000375545.1_ASM37554v1
Sneathiella.glossodoripedis.JCM.23214 GCA_000616095.1_ASM61609v1
Geminicoccus.roseus.DSM.18922 GCA_000427665.1_ASM42766v1
Tistrella.mobilis.KA081020.065 GCA_000264455.2_ASM26445v2
Caenispirillum.salinarum.AK4 GCA_000315795.1
Pararhodospirillum.photometricum.DSM.122 GCA_000284415.2_ASM28441v2
Rhodospirillum.rubrum.F11 GCA_000225955.1_ASM22595v1
Roseospirillum.parvum GCA_900100455.1
Magnetospirillum.magneticum.AMB.1 GCA_000009985.1_ASM998v1
Terasakiella.pusilla.DSM.6293 GCA_000688235.1_ASM68823v1
Thalassospira.profundimaris.WP0211 GCA_000300275.1_ASM30027v1
Candidatus.Puniceispirillum.marinum.IMCC1322 GCA_000024465.1_ASM2446v1
Thalassobaculum.salexigens.DSM.19539 GCA_000423805.1_ASM42380v1
alpha.proteobacterium.BAL199 GCA_000171835.1_ASM17183v1
Oceanibaculum.indicum GCA_003633955.1_ASM363395v1
Kiloniella.laminariae.DSM.19542 GCA_000374005.1_ASM37400v1
Rhodovibrio.salinarum.DSM.9154 GCA_000515255.1_ASM51525v1
Inquilinus.limosus.DSM.16000 GCA_000423185.1_ASM42318v1
Rhodospirillum.centenum GCA_014197805.1_ASM1419780v1
Micavibrio.aeruginosavorus.ARL.13 GCA_000226315.1_ASM22631v1
Elioraea.tepidiphila.DSM.17972 GCA_000378465.1_ASM37846v1
Belnapia.moabensis.DSM.16746 GCA_000745835.1_ASM74583v1
Rubritepida.flocculans.DSM.14296 GCA_000425365.1_ASM42536v1
Roseomonas.gilardii GCA_001941945.1_ASM194194v1
Acidiphilium.multivorum.AIU301 GCA_000202835.1_ASM20283v1
Acidisphaera.rubrifaciens.HS.AP3 GCA_000964365.1_ASM96436v1
Granulibacter.bethesdensis.CGDNIH4 GCA_000576145.2_ASM57614v2
Commensalibacter.intestini.A911 GCA_000231445.2_ASM23144v1
Gluconacetobacter.diazotrophicus.PA1.5 GCA_000067045.1_ASM6704v1
Acetobacter.aceti.NBRC.14818 GCA_000193495.2_ASM19349v2
Gluconobacter.oxydans.DSM.3504 GCA_000583855.1_ASM58385v1
0.803252

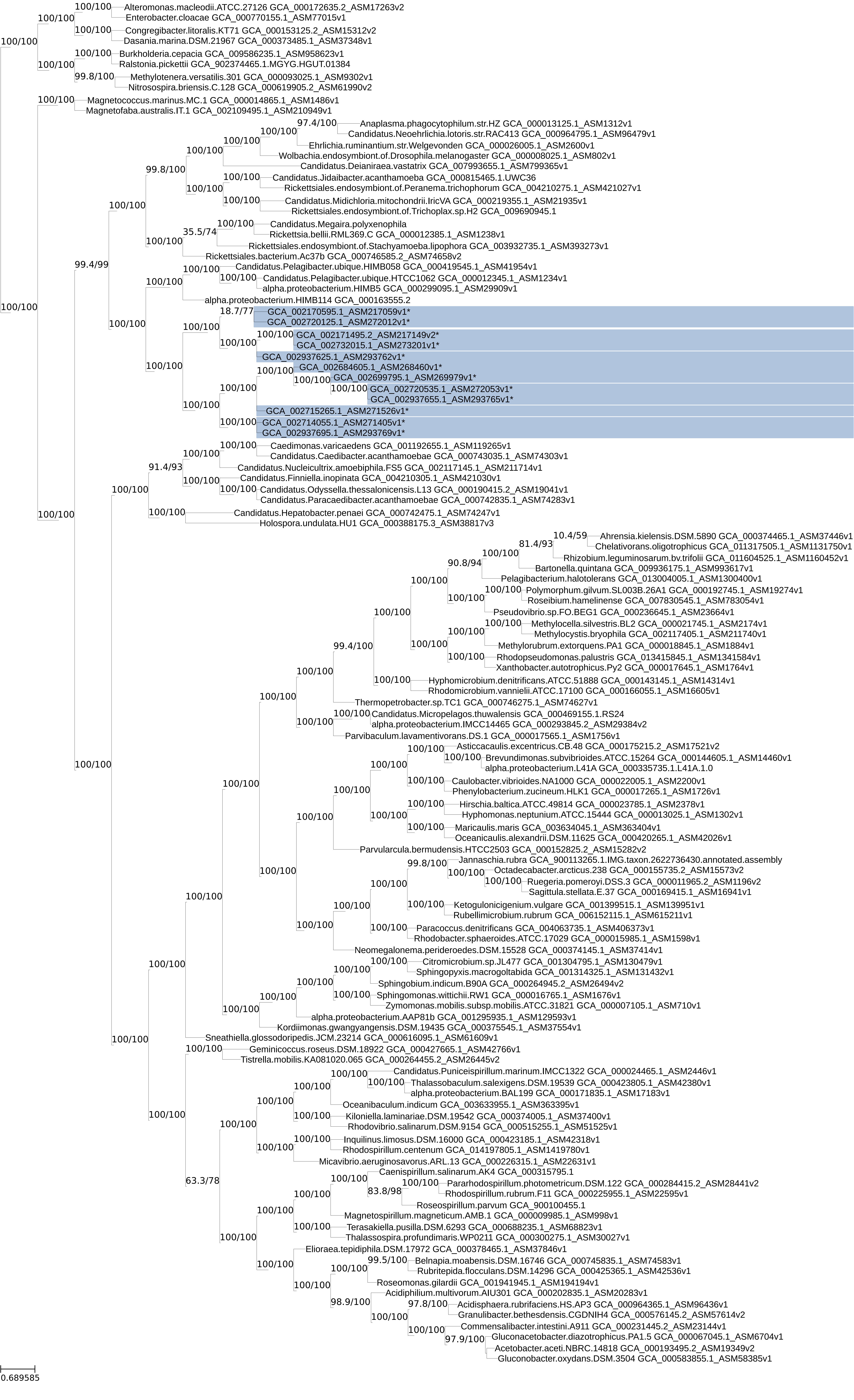

Alteromonas.macleodii.ATCC.27126 GCA_000172635.2_ASM17263v2
Enterobacter.cloacae GCA_000770155.1_ASM77015v1
Congregibacter.litoralis.KT71 GCA_000153125.2_ASM15312v2
Dasania.marina.DSM.21967 GCA_000373485.1_ASM37348v1
Burkholderia.cepacia GCA_009586235.1_ASM958623v1
Ralstonia.pickettii GCA_902374465.1.MGYG.HGUT.01384
Methylotenera.versatilis.301 GCA_000093025.1_ASM9302v1
Nitrosospira.briensis.C.128 GCA_000619905.2_ASM61990v2
Magnetococcus.marinus.MC.1 GCA_000014865.1_ASM1486v1
Magnetofaba.australis.IT.1 GCA_002109495.1_ASM210949v1
Anaplasma.phagocytophilum.str.HZ GCA_000013125.1_ASM1312v1
Candidatus.Neoehrlichia.lotoris.str.RAC413 GCA_000964795.1_ASM96479v1
Ehrlichia.ruminantium.str.Welgevonden GCA_000026005.1_ASM2600v1
Wolbachia.endosymbiont.of.Drosophila.melanogaster GCA_000008025.1_ASM802v1
Candidatus.Deianiraea.vastatrix GCA_007993655.1_ASM799365v1
Candidatus.Jidaibacter.acanthamoeba GCA_000815465.1.UWC36
Rickettsiales.endosymbiont.of.Peranema.trichophorum GCA_004210275.1_ASM421027v1
Candidatus.Midichloria.mitochondrii.IricVA GCA_000219355.1_ASM21935v1
Rickettsiales.endosymbiont.of.Trichoplax.sp.H2 GCA_009690945.1
Candidatus.Megaira.polyxenophila
Rickettsia.bellii.RML369.C GCA_000012385.1_ASM1238v1
Rickettsiales.endosymbiont.of.Stachyamoeba.lipophora GCA_003932735.1_ASM393273v1
Rickettsiales.bacterium.Ac37b GCA_000746585.2_ASM74658v2
Candidatus.Pelagibacter.ubique.HIMB058 GCA_000419545.1_ASM41954v1
Candidatus.Pelagibacter.ubique.HTCC1062 GCA_000012345.1_ASM1234v1
alpha.proteobacterium.HIMB5 GCA_000299095.1_ASM29909v1
alpha.proteobacterium.HIMB114 GCA_000163555.2
GCA_002170595.1_ASM217059v1*
GCA_002720125.1_ASM272012v1*
GCA_002171495.2_ASM217149v2*
GCA_002732015.1_ASM273201v1*
GCA_002937625.1_ASM293762v1*
GCA_002684605.1_ASM268460v1*
GCA_002699795.1_ASM269979v1*
GCA_002720535.1_ASM272053v1*
GCA_002937655.1_ASM293765v1*
GCA_002715265.1_ASM271526v1*
GCA_002714055.1_ASM271405v1*
GCA_002937695.1_ASM293769v1*
Caedimonas.varicaedens GCA_001192655.1_ASM119265v1
Candidatus.Caedibacter.acanthamoebae GCA_000743035.1_ASM74303v1
Candidatus.Nucleicultrix.amoebiphila.FS5 GCA_002117145.1_ASM211714v1
Candidatus.Finniella.inopinata GCA_004210305.1_ASM421030v1
Candidatus.Odyssella.thessalonicensis.L13 GCA_000190415.2_ASM19041v1
Candidatus.Paracaedibacter.acanthamoebae GCA_000742835.1_ASM74283v1
Candidatus.Hepatobacter.penaei GCA_000742475.1_ASM74247v1
Holospora.undulata.HU1 GCA_000388175.3_ASM38817v3
Ahrensia.kielensis.DSM.5890 GCA_000374465.1_ASM37446v1
Chelativorans.oligotrophicus GCA_011317505.1_ASM1131750v1
Rhizobium.leguminosarum.bv.trifolii GCA_011604525.1_ASM1160452v1
Bartonella.quintana GCA_009936175.1_ASM993617v1
Pelagibacterium.halotolerans GCA_013004005.1_ASM1300400v1
Polymorphum.gilvum.SL003B.26A1 GCA_000192745.1_ASM19274v1
Roseibium.hamelinense GCA_007830545.1_ASM783054v1
Pseudovibrio.sp.FO.BEG1 GCA_000236645.1_ASM23664v1
Methylocella.silvestris.BL2 GCA_000021745.1_ASM2174v1
Methylocystis.bryophila GCA_002117405.1_ASM211740v1
Methylorubrum.extorquens.PA1 GCA_000018845.1_ASM1884v1
Rhodopseudomonas.palustris GCA_013415845.1_ASM1341584v1
Xanthobacter.autotrophicus.Py2 GCA_000017645.1_ASM1764v1
Hyphomicrobium.denitrificans.ATCC.51888 GCA_000143145.1_ASM14314v1
Rhodomicrobium.vannielii.ATCC.17100 GCA_000166055.1_ASM16605v1
Thermopetrobacter.sp.TC1 GCA_000746275.1_ASM74627v1
Candidatus.Micropelagos.thuwalensis GCA_000469155.1.RS24
alpha.proteobacterium.IMCC14465 GCA_000293845.2_ASM29384v2
Parvibaculum.lavamentivorans.DS.1 GCA_000017565.1_ASM1756v1
Asticcacaulis.excentricus.CB.48 GCA_000175215.2_ASM17521v2
Brevundimonas.subvibrioides.ATCC.15264 GCA_000144605.1_ASM14460v1
alpha.proteobacterium.L41A GCA_000335735.1.L41A.1.0
Caulobacter.vibrioides.NA1000 GCA_000022005.1_ASM2200v1
Phenylobacterium.zucineum.HLK1 GCA_000017265.1_ASM1726v1
Hirschia.baltica.ATCC.49814 GCA_000023785.1_ASM2378v1
Hyphomonas.neptunium.ATCC.15444 GCA_000013025.1_ASM1302v1
Maricaulis.maris GCA_003634045.1_ASM363404v1
Oceanicaulis.alexandrii.DSM.11625 GCA_000420265.1_ASM42026v1
Parvularcula.bermudensis.HTCC2503 GCA_000152825.2_ASM15282v2
Jannaschia.rubra GCA_900113265.1.IMG.taxon.2622736430.annotated.assembly
Octadecabacter.arcticus.238 GCA_000155735.2_ASM15573v2
Ruegeria.pomeroyi.DSS.3 GCA_000011965.2_ASM1196v2
Sagittula.stellata.E.37 GCA_000169415.1_ASM16941v1
Ketogulonicigenium.vulgare GCA_001399515.1_ASM139951v1
Rubellimicrobium.rubrum GCA_006152115.1_ASM615211v1
Paracoccus.denitrificans GCA_004063735.1_ASM406373v1
Rhodobacter.sphaeroides.ATCC.17029 GCA_000015985.1_ASM1598v1
Neomegalonema.perideroedes.DSM.15528 GCA_000374145.1_ASM37414v1
Citromicrobium.sp.JL477 GCA_001304795.1_ASM130479v1
Sphingopyxis.macrogoltabida GCA_001314325.1_ASM131432v1
Sphingobium.indicum.B90A GCA_000264945.2_ASM26494v2
Sphingomonas.wittichii.RW1 GCA_000016765.1_ASM1676v1
Zymomonas.mobilis.subsp.mobilis.ATCC.31821 GCA_000007105.1_ASM710v1
alpha.proteobacterium.AAP81b GCA_001295935.1_ASM129593v1
Kordiimonas.gwangyangensis.DSM.19435 GCA_000375545.1_ASM37554v1
Sneathiella.glossodoripedis.JCM.23214 GCA_000616095.1_ASM61609v1
Geminicoccus.roseus.DSM.18922 GCA_000427665.1_ASM42766v1
Tistrella.mobilis.KA081020.065 GCA_000264455.2_ASM26445v2
Candidatus.Puniceispirillum.marinum.IMCC1322 GCA_000024465.1_ASM2446v1
Thalassobaculum.salexigens.DSM.19539 GCA_000423805.1_ASM42380v1
alpha.proteobacterium.BAL199 GCA_000171835.1_ASM17183v1
Oceanibaculum.indicum GCA_003633955.1_ASM363395v1
Kiloniella.laminariae.DSM.19542 GCA_000374005.1_ASM37400v1
Rhodovibrio.salinarum.DSM.9154 GCA_000515255.1_ASM51525v1
Inquilinus.limosus.DSM.16000 GCA_000423185.1_ASM42318v1
Rhodospirillum.centenum GCA_014197805.1_ASM1419780v1
Micavibrio.aeruginosavorus.ARL.13 GCA_000226315.1_ASM22631v1
Caenispirillum.salinarum.AK4 GCA_000315795.1
Pararhodospirillum.photometricum.DSM.122 GCA_000284415.2_ASM28441v2
Rhodospirillum.rubrum.F11 GCA_000225955.1_ASM22595v1
Roseospirillum.parvum GCA_900100455.1
Magnetospirillum.magneticum.AMB.1 GCA_000009985.1_ASM998v1
Terasakiella.pusilla.DSM.6293 GCA_000688235.1_ASM68823v1
Thalassospira.profundimaris.WP0211 GCA_000300275.1_ASM30027v1
Elioraea.tepidiphila.DSM.17972 GCA_000378465.1_ASM37846v1
Belnapia.moabensis.DSM.16746 GCA_000745835.1_ASM74583v1
Rubritepida.flocculans.DSM.14296 GCA_000425365.1_ASM42536v1
Roseomonas.gilardii GCA_001941945.1_ASM194194v1
Acidiphilium.multivorum.AIU301 GCA_000202835.1_ASM20283v1
Acidisphaera.rubrifaciens.HS.AP3 GCA_000964365.1_ASM96436v1
Granulibacter.bethesdensis.CGDNIH4 GCA_000576145.2_ASM57614v2
Commensalibacter.intestini.A911 GCA_000231445.2_ASM23144v1
Gluconacetobacter.diazotrophicus.PA1.5 GCA_000067045.1_ASM6704v1
Acetobacter.aceti.NBRC.14818 GCA_000193495.2_ASM19349v2
Gluconobacter.oxydans.DSM.3504 GCA_000583855.1_ASM58385v1
0.689585

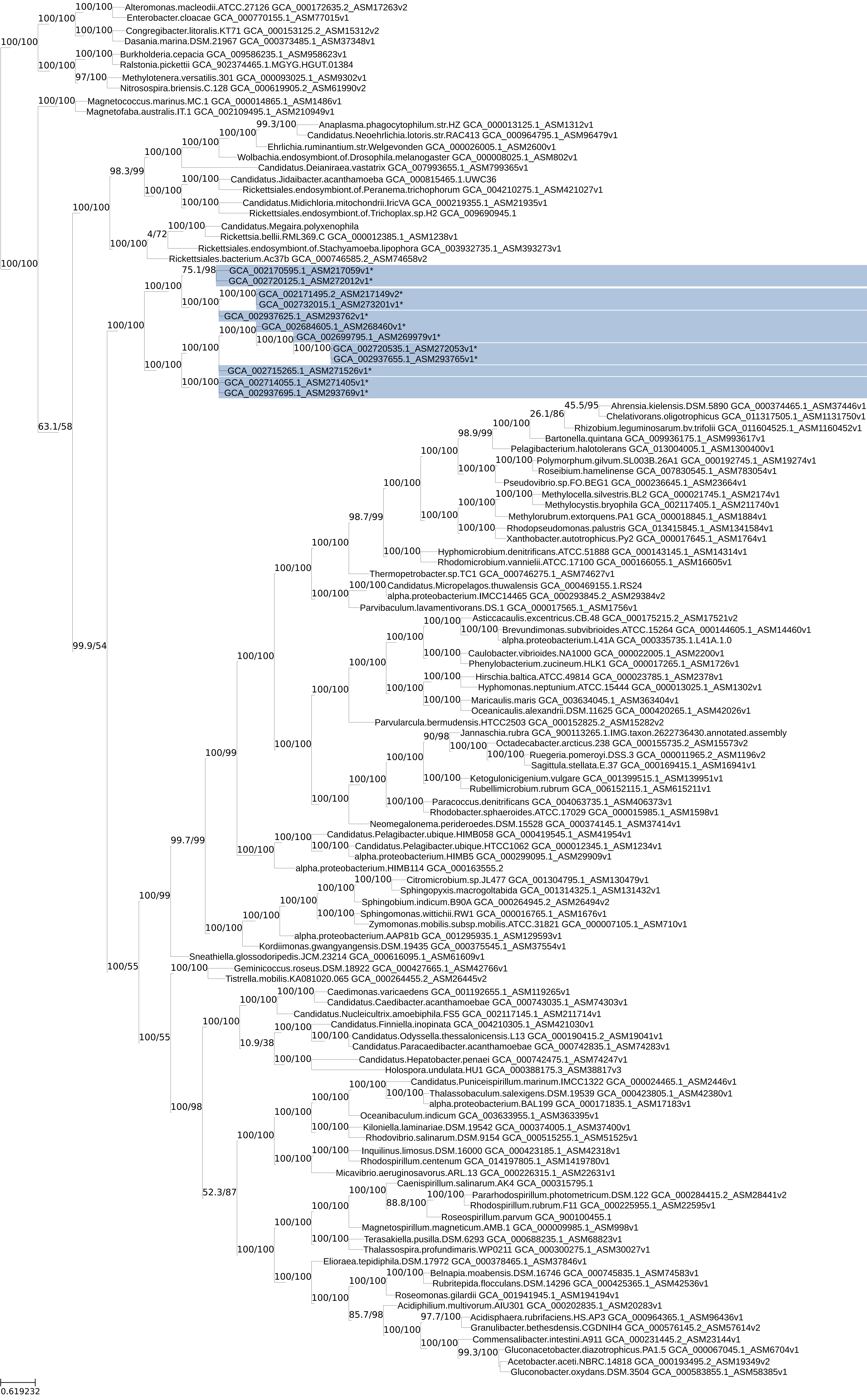

100/100
Alteromonas.macleodii.ATCC.27126 GCA_000172635.2_ASM17263v2
100/100
Enterobacter.cloacae GCA_000770155.1_ASM77015v1
100/100
100/100
Congregibacter.litoralis.KT71 GCA_000153125.2_ASM15312v2
Dasania.marina.DSM.21967 GCA_000373485.1_ASM37348v1
100/100
Burkholderia.cepacia GCA_009586235.1_ASM958623v1
100/100
Ralstonia.pickettii GCA_902374465.1.MGYG.HGUT.01384
97/100
Methylotenera.versatilis.301 GCA_000093025.1_ASM9302v1
Nitrosospira.briensis.C.128 GCA_000619905.2_ASM61990v2
100/100
Magnetococcus.marinus.MC.1 GCA_000014865.1_ASM1486v1
Magnetofaba.australis.IT.1 GCA_002109495.1_ASM210949v1
99.3/100
Anaplasma.phagocytophilum.str.HZ GCA_000013125.1_ASM1312v1
100/100
Candidatus.Neoehrlichia.lotoris.str.RAC413 GCA_000964795.1_ASM96479v1
100/100
Ehrlichia.ruminantium.str.Welgevonden GCA_000026005.1_ASM2600v1
100/100
Wolbachia.endosymbiont.of.Drosophila.melanogaster GCA_000008025.1_ASM802v1
98.3/99
Candidatus.Deianiraea.vastatrix GCA_007993655.1_ASM799365v1
100/100
Candidatus.Jidaibacter.acanthamoeba GCA_000815465.1.UWC36
100/100
Rickettsiales.endosymbiont.of.Peranema.trichophorum GCA_004210275.1_ASM421027v1
100/100
100/100
Candidatus.Midichloria.mitochondrii.IricVA GCA_000219355.1_ASM21935v1
Rickettsiales.endosymbiont.of.Trichoplax.sp.H2 GCA_009690945.1
100/100
Candidatus.Megaira.polyxenophila
4/72
Rickettsia.bellii.RML369.C GCA_000012385.1_ASM1238v1
100/100
Rickettsiales.endosymbiont.of.Stachyamoeba.lipophora GCA_003932735.1_ASM393273v1
100/100
Rickettsiales.bacterium.Ac37b GCA_000746585.2_ASM74658v2
75.1/98
GCA_002170595.1_ASM217059v1*
GCA_002720125.1_ASM272012v1*
100/100
100/100
GCA_002171495.2_ASM217149v2*
100/100
GCA_002732015.1_ASM273201v1*
GCA_002937625.1_ASM293762v1*
100/100
100/100
GCA_002684605.1_ASM268460v1*
GCA_002699795.1_ASM269979v1*
100/100
100/100
GCA_002720535.1_ASM272053v1*
100/100
GCA_002937655.1_ASM293765v1*
100/100
GCA_002715265.1_ASM271526v1*
100/100
GCA_002714055.1_ASM271405v1*
GCA_002937695.1_ASM293769v1*
45.5/95
Ahrensia.kielensis.DSM.5890 GCA_000374465.1_ASM37446v1
26.1/86
Chelativorans.oligotrophicus GCA_011317505.1_ASM1131750v1
100/100
Rhizobium.leguminosarum.bv.trifolii GCA_011604525.1_ASM1160452v1
63.1/58
Bartonella.quintana GCA_009936175.1_ASM993617v1
98.9/99
Pelagibacterium.halotolerans GCA_013004005.1_ASM1300400v1
100/100
100/100
Polymorphum.gilvum.SL003B.26A1 GCA_000192745.1_ASM19274v1
100/100
Roseibium.hamelinense GCA_007830545.1_ASM783054v1
Pseudovibrio.sp.FO.BEG1 GCA_000236645.1_ASM23664v1
100/100
100/100
Methylocella.silvestris.BL2 GCA_000021745.1_ASM2174v1
100/100
Methylocystis.bryophila GCA_002117405.1_ASM211740v1
100/100
Methylorubrum.extorquens.PA1 GCA_000018845.1_ASM1884v1
98.7/99
100/100
Rhodopseudomonas.palustris GCA_013415845.1_ASM1341584v1
Xanthobacter.autotrophicus.Py2 GCA_000017645.1_ASM1764v1
100/100
100/100
Hyphomicrobium.denitrificans.ATCC.51888 GCA_000143145.1_ASM14314v1
Rhodomicrobium.vannielii.ATCC.17100 GCA_000166055.1_ASM16605v1
100/100
Thermopetrobacter.sp.TC1 GCA_000746275.1_ASM74627v1
100/100
Candidatus.Micropelagos.thuwalensis GCA_000469155.1.RS24
100/100
alpha.proteobacterium.IMCC14465 GCA_000293845.2_ASM29384v2
Parvibaculum.lavamentivorans.DS.1 GCA_000017565.1_ASM1756v1
100/100
Asticcacaulis.excentricus.CB.48 GCA_000175215.2_ASM17521v2
100/100
100/100
Brevundimonas.subvibrioides.ATCC.15264 GCA_000144605.1_ASM14460v1
alpha.proteobacterium.L41A GCA_000335735.1.L41A.1.0
99.9/54
100/100
100/100
Caulobacter.vibrioides.NA1000 GCA_000022005.1_ASM2200v1
Phenylobacterium.zucineum.HLK1 GCA_000017265.1_ASM1726v1
100/100
100/100
Hirschia.baltica.ATCC.49814 GCA_000023785.1_ASM2378v1
100/100
Hyphomonas.neptunium.ATCC.15444 GCA_000013025.1_ASM1302v1
100/100
Maricaulis.maris GCA_003634045.1_ASM363404v1
Oceanicaulis.alexandrii.DSM.11625 GCA_000420265.1_ASM42026v1
Parvularcula.bermudensis.HTCC2503 GCA_000152825.2_ASM15282v2
90/98
Jannaschia.rubra GCA_900113265.1.IMG.taxon.2622736430.annotated.assembly
100/100
Octadecabacter.arcticus.238 GCA_000155735.2_ASM15573v2
100/100
100/100
Ruegeria.pomeroyi.DSS.3 GCA_000011965.2_ASM1196v2
100/99
100/100
Sagittula.stellata.E.37 GCA_000169415.1_ASM16941v1
100/100
Ketogulonicigenium.vulgare GCA_001399515.1_ASM139951v1
Rubellimicrobium.rubrum GCA_006152115.1_ASM615211v1
100/100
100/100
Paracoccus.denitrificans GCA_004063735.1_ASM406373v1
Rhodobacter.sphaeroides.ATCC.17029 GCA_000015985.1_ASM1598v1
Neomegalonema.perideroedes.DSM.15528 GCA_000374145.1_ASM37414v1
100/100
Candidatus.Pelagibacter.ubique.HIMB058 GCA_000419545.1_ASM41954v1
99.7/99
100/100
Candidatus.Pelagibacter.ubique.HTCC1062 GCA_000012345.1_ASM1234v1
100/100
alpha.proteobacterium.HIMB5 GCA_000299095.1_ASM29909v1
alpha.proteobacterium.HIMB114 GCA_000163555.2
100/100
Citromicrobium.sp.JL477 GCA_001304795.1_ASM130479v1
100/100
Sphingopyxis.macrogoltabida GCA_001314325.1_ASM131432v1
100/99
100/100
Sphingobium.indicum.B90A GCA_000264945.2_ASM26494v2
100/100
Sphingomonas.wittichii.RW1 GCA_000016765.1_ASM1676v1
100/100
Zymomonas.mobilis.subsp.mobilis.ATCC.31821 GCA_000007105.1_ASM710v1
100/100
alpha.proteobacterium.AAP81b GCA_001295935.1_ASM129593v1
Kordiimonas.gwangyangensis.DSM.19435 GCA_000375545.1_ASM37554v1
Sneathiella.glossodoripedis.JCM.23214 GCA_000616095.1_ASM61609v1
100/55
100/100
Geminicoccus.roseus.DSM.18922 GCA_000427665.1_ASM42766v1
Tistrella.mobilis.KA081020.065 GCA_000264455.2_ASM26445v2
100/100
Caedimonas.varicaedens GCA_001192655.1_ASM119265v1
100/100
Candidatus.Caedibacter.acanthamoebae GCA_000743035.1_ASM74303v1
100/100
Candidatus.Nucleicultrix.amoebiphila.FS5 GCA_002117145.1_ASM211714v1
100/100
Candidatus.Finniella.inopinata GCA_004210305.1_ASM421030v1
100/55
100/100
Candidatus.Odyssella.thessalonicensis.L13 GCA_000190415.2_ASM19041v1
10.9/38
Candidatus.Paracaedibacter.acanthamoebae GCA_000742835.1_ASM74283v1
100/100
Candidatus.Hepatobacter.penaei GCA_000742475.1_ASM74247v1
Holospora.undulata.HU1 GCA_000388175.3_ASM38817v3
100/100
Candidatus.Puniceispirillum.marinum.IMCC1322 GCA_000024465.1_ASM2446v1
100/100
100/100
Thalassobaculum.salexigens.DSM.19539 GCA_000423805.1_ASM42380v1
alpha.proteobacterium.BAL199 GCA_000171835.1_ASM17183v1
100/98
100/100
Oceanibaculum.indicum GCA_003633955.1_ASM363395v1
100/100
Kiloniella.laminariae.DSM.19542 GCA_000374005.1_ASM37400v1
100/100
Rhodovibrio.salinarum.DSM.9154 GCA_000515255.1_ASM51525v1
100/100
Inquilinus.limosus.DSM.16000 GCA_000423185.1_ASM42318v1
100/100
Rhodospirillum.centenum GCA_014197805.1_ASM1419780v1
Micavibrio.aeruginosavorus.ARL.13 GCA_000226315.1_ASM22631v1
52.3/87
100/100
Caenispirillum.salinarum.AK4 GCA_000315795.1
100/100
Pararhodospirillum.photometricum.DSM.122 GCA_000284415.2_ASM28441v2
88.8/100
Rhodospirillum.rubrum.F11 GCA_000225955.1_ASM22595v1
100/100
Roseospirillum.parvum GCA_900100455.1
100/100
Magnetospirillum.magneticum.AMB.1 GCA_000009985.1_ASM998v1
100/100
Terasakiella.pusilla.DSM.6293 GCA_000688235.1_ASM68823v1
100/100
Thalassospira.profundimaris.WP0211 GCA_000300275.1_ASM30027v1
Elioraea.tepidiphila.DSM.17972 GCA_000378465.1_ASM37846v1
100/100
Belnapia.moabensis.DSM.16746 GCA_000745835.1_ASM74583v1
100/100
100/100
Rubritepida.flocculans.DSM.14296 GCA_000425365.1_ASM42536v1
100/100
Roseomonas.gilardii GCA_001941945.1_ASM194194v1
Acidiphilium.multivorum.AIU301 GCA_000202835.1_ASM20283v1
85.7/98
97.7/100
Acidisphaera.rubrifaciens.HS.AP3 GCA_000964365.1_ASM96436v1
100/100
Granulibacter.bethesdensis.CGDNIH4 GCA_000576145.2_ASM57614v2
100/100
Commensalibacter.intestini.A911 GCA_000231445.2_ASM23144v1
99.3/100
Gluconacetobacter.diazotrophicus.PA1.5 GCA_000067045.1_ASM6704v1
Acetobacter.aceti.NBRC.14818 GCA_000193495.2_ASM19349v2
Gluconobacter.oxydans.DSM.3504 GCA_000583855.1_ASM58385v1
100/100
100/100
0.619232

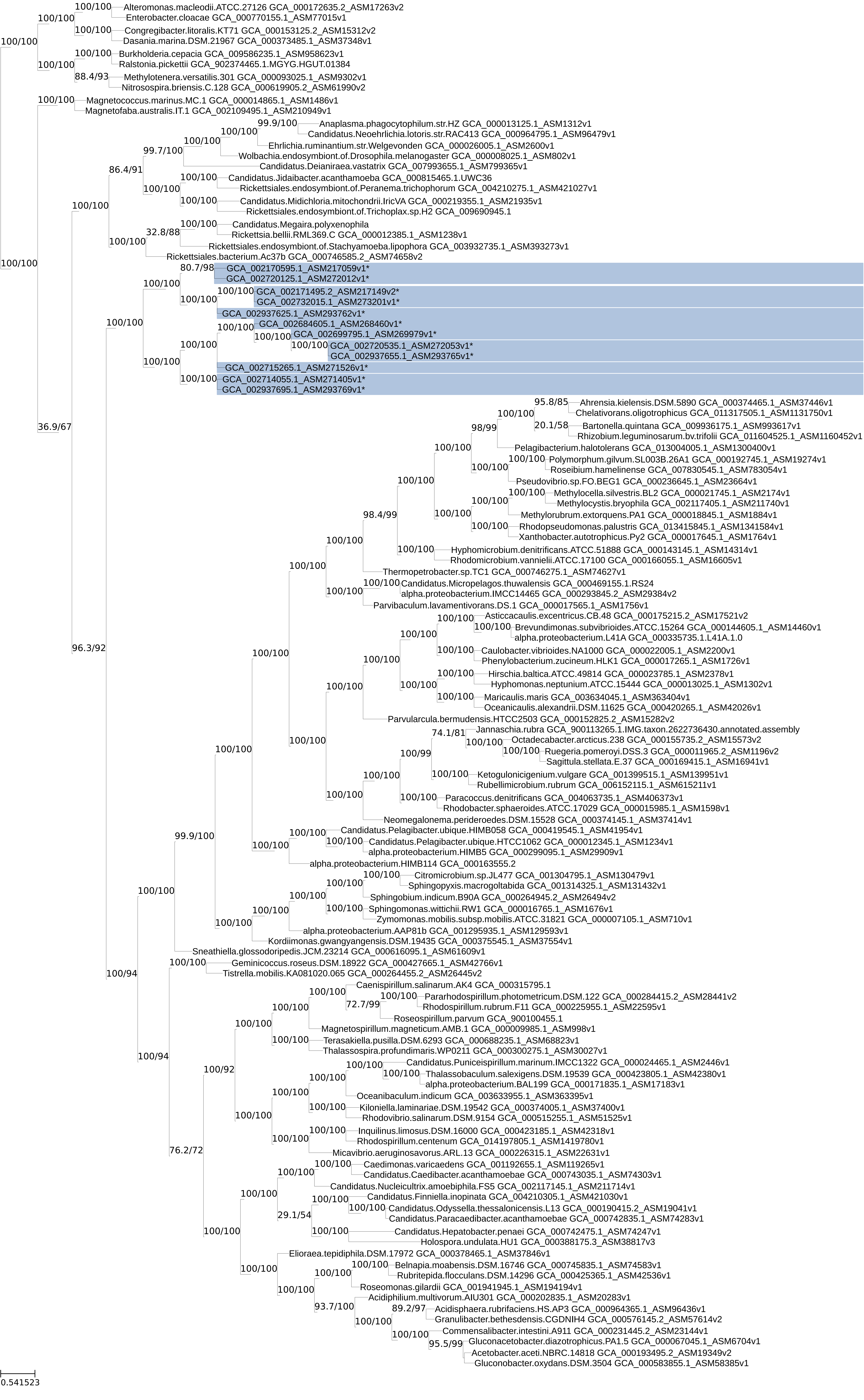
100/100
Alteromonas.macleodii.ATCC.27126 GCA_000172635.2_ASM17263v2
Enterobacter.cloacae GCA_000770155.1_ASM77015v1
100/100
100/100
Congregibacter.litoralis.KT71 GCA_000153125.2_ASM15312v2
Dasania.marina.DSM.21967 GCA_000373485.1_ASM37348v1
100/100
100/100
Burkholderia.cepacia GCA_009586235.1_ASM958623v1
Ralstonia.pickettii GCA_902374465.1.MGYG.HGUT.01384
100/100
88.4/93
Methylotenera.versatilis.301 GCA_000093025.1_ASM9302v1
Nitrosospira.briensis.C.128 GCA_000619905.2_ASM61990v2
100/100
Magnetococcus.marinus.MC.1 GCA_000014865.1_ASM1486v1
Magnetofaba.australis.IT.1 GCA_002109495.1_ASM210949v1
99.9/100
Anaplasma.phagocytophilum.str.HZ GCA_000013125.1_ASM1312v1
100/100
Candidatus.Neoehrlichia.lotoris.str.RAC413 GCA_000964795.1_ASM96479v1
100/100
Ehrlichia.ruminantium.str.Welgevonden GCA_000026005.1_ASM2600v1
99.7/100
Wolbachia.endosymbiont.of.Drosophila.melanogaster GCA_000008025.1_ASM802v1
86.4/91
Candidatus.Deianiraea.vastatrix GCA_007993655.1_ASM799365v1
100/100
Candidatus.Jidaibacter.acanthamoeba GCA_000815465.1.UWC36
100/100
Rickettsiales.endosymbiont.of.Peranema.trichophorum GCA_004210275.1_ASM421027v1
100/100
100/100
Candidatus.Midichloria.mitochondrii.IricVA GCA_000219355.1_ASM21935v1
Rickettsiales.endosymbiont.of.Trichoplax.sp.H2 GCA_009690945.1
100/100
Candidatus.Megaira.polyxenophila
32.8/88
Rickettsia.bellii.RML369.C GCA_000012385.1_ASM1238v1
100/100
Rickettsiales.endosymbiont.of.Stachyamoeba.lipophora GCA_003932735.1_ASM393273v1
Rickettsiales.bacterium.Ac37b GCA_000746585.2_ASM74658v2
100/100
80.7/98
GCA_002170595.1_ASM217059v1*
GCA_002720125.1_ASM272012v1*
100/100
100/100
GCA_002171495.2_ASM217149v2*
100/100
GCA_002732015.1_ASM273201v1*
GCA_002937625.1_ASM293762v1*
100/100
GCA_002684605.1_ASM268460v1*
100/100
GCA_002699795.1_ASM269979v1*
100/100
100/100
GCA_002720535.1_ASM272053v1*
GCA_002937655.1_ASM293765v1*
100/100
GCA_002715265.1_ASM271526v1*
100/100
GCA_002714055.1_ASM271405v1*
GCA_002937695.1_ASM293769v1*
95.8/85
Ahrensia.kielensis.DSM.5890 GCA_000374465.1_ASM37446v1
100/100
Chelativorans.oligotrophicus GCA_011317505.1_ASM1131750v1
36.9/67
98/99
20.1/58
Bartonella.quintana GCA_009936175.1_ASM993617v1
Rhizobium.leguminosarum.bv.trifolii GCA_011604525.1_ASM1160452v1
Pelagibacterium.halotolerans GCA_013004005.1_ASM1300400v1
100/100
100/100
Polymorphum.gilvum.SL003B.26A1 GCA_000192745.1_ASM19274v1
100/100
Roseibium.hamelinense GCA_007830545.1_ASM783054v1
Pseudovibrio.sp.FO.BEG1 GCA_000236645.1_ASM23664v1
100/100
100/100
Methylocella.silvestris.BL2 GCA_000021745.1_ASM2174v1
100/100
Methylocystis.bryophila GCA_002117405.1_ASM211740v1
100/100
Methylorubrum.extorquens.PA1 GCA_000018845.1_ASM1884v1
98.4/99
100/100
Rhodopseudomonas.palustris GCA_013415845.1_ASM1341584v1
Xanthobacter.autotrophicus.Py2 GCA_000017645.1_ASM1764v1
100/100
100/100
Hyphomicrobium.denitrificans.ATCC.51888 GCA_000143145.1_ASM14314v1
Rhodomicrobium.vannielii.ATCC.17100 GCA_000166055.1_ASM16605v1
100/100
Thermopetrobacter.sp.TC1 GCA_000746275.1_ASM74627v1
100/100
Candidatus.Micropelagos.thuwalensis GCA_000469155.1.RS24
100/100
alpha.proteobacterium.IMCC14465 GCA_000293845.2_ASM29384v2
Parvibaculum.lavamentivorans.DS.1 GCA_000017565.1_ASM1756v1
100/100
Asticcacaulis.excentricus.CB.48 GCA_000175215.2_ASM17521v2
100/100
Brevundimonas.subvibrioides.ATCC.15264 GCA_000144605.1_ASM14460v1
100/100
alpha.proteobacterium.L41A GCA_000335735.1.L41A.1.0
100/100
Caulobacter.vibrioides.NA1000 GCA_000022005.1_ASM2200v1
96.3/92
100/100
Phenylobacterium.zucineum.HLK1 GCA_000017265.1_ASM1726v1
100/100
Hirschia.baltica.ATCC.49814 GCA_000023785.1_ASM2378v1
100/100
Hyphomonas.neptunium.ATCC.15444 GCA_000013025.1_ASM1302v1
100/100
100/100
Maricaulis.maris GCA_003634045.1_ASM363404v1
Oceanicaulis.alexandrii.DSM.11625 GCA_000420265.1_ASM42026v1
Parvularcula.bermudensis.HTCC2503 GCA_000152825.2_ASM15282v2
74.1/81
Jannaschia.rubra GCA_900113265.1.IMG.taxon.2622736430.annotated.assembly
100/100
Octadecabacter.arcticus.238 GCA_000155735.2_ASM15573v2
100/100
Ruegeria.pomeroyi.DSS.3 GCA_000011965.2_ASM1196v2
100/99
Sagittula.stellata.E.37 GCA_000169415.1_ASM16941v1
100/100
100/100
Ketogulonicigenium.vulgare GCA_001399515.1_ASM139951v1
100/100
Rubellimicrobium.rubrum GCA_006152115.1_ASM615211v1
100/100
Paracoccus.denitrificans GCA_004063735.1_ASM406373v1
Rhodobacter.sphaeroides.ATCC.17029 GCA_000015985.1_ASM1598v1
Neomegalonema.perideroedes.DSM.15528 GCA_000374145.1_ASM37414v1
100/100
Candidatus.Pelagibacter.ubique.HIMB058 GCA_000419545.1_ASM41954v1
99.9/100
100/100
100/100
Candidatus.Pelagibacter.ubique.HTCC1062 GCA_000012345.1_ASM1234v1
alpha.proteobacterium.HIMB5 GCA_000299095.1_ASM29909v1
alpha.proteobacterium.HIMB114 GCA_000163555.2
100/100
Citromicrobium.sp.JL477 GCA_001304795.1_ASM130479v1
100/100
Sphingopyxis.macrogoltabida GCA_001314325.1_ASM131432v1
100/100
100/100
Sphingobium.indicum.B90A GCA_000264945.2_ASM26494v2
100/100
Sphingomonas.wittichii.RW1 GCA_000016765.1_ASM1676v1
100/100
Zymomonas.mobilis.subsp.mobilis.ATCC.31821 GCA_000007105.1_ASM710v1
100/100
alpha.proteobacterium.AAP81b GCA_001295935.1_ASM129593v1
Kordiimonas.gwangyangensis.DSM.19435 GCA_000375545.1_ASM37554v1
Sneathiella.glossodoripedis.JCM.23214 GCA_000616095.1_ASM61609v1
100/100
Geminicoccus.roseus.DSM.18922 GCA_000427665.1_ASM42766v1
100/94
Tistrella.mobilis.KA081020.065 GCA_000264455.2_ASM26445v2
Caenispirillum.salinarum.AK4 GCA_000315795.1
100/100
100/100
Pararhodospirillum.photometricum.DSM.122 GCA_000284415.2_ASM28441v2
72.7/99
Rhodospirillum.rubrum.F11 GCA_000225955.1_ASM22595v1
100/100
Roseospirillum.parvum GCA_900100455.1
100/100
Magnetospirillum.magneticum.AMB.1 GCA_000009985.1_ASM998v1
100/100
Terasakiella.pusilla.DSM.6293 GCA_000688235.1_ASM68823v1
100/94
Thalassospira.profundimaris.WP0211 GCA_000300275.1_ASM30027v1
100/100
Candidatus.Puniceispirillum.marinum.IMCC1322 GCA_000024465.1_ASM2446v1
100/92
100/100
Thalassobaculum.salexigens.DSM.19539 GCA_000423805.1_ASM42380v1
100/100
alpha.proteobacterium.BAL199 GCA_000171835.1_ASM17183v1
100/100
Oceanibaculum.indicum GCA_003633955.1_ASM363395v1
100/100
Kiloniella.laminariae.DSM.19542 GCA_000374005.1_ASM37400v1
100/100
Rhodovibrio.salinarum.DSM.9154 GCA_000515255.1_ASM51525v1
100/100
Inquilinus.limosus.DSM.16000 GCA_000423185.1_ASM42318v1
100/100
Rhodospirillum.centenum GCA_014197805.1_ASM1419780v1
76.2/72
Micavibrio.aeruginosavorus.ARL.13 GCA_000226315.1_ASM22631v1
100/100
Caedimonas.varicaedens GCA_001192655.1_ASM119265v1
100/100
Candidatus.Caedibacter.acanthamoebae GCA_000743035.1_ASM74303v1
Candidatus.Nucleicultrix.amoebiphila.FS5 GCA_002117145.1_ASM211714v1
100/100
100/100
Candidatus.Finniella.inopinata GCA_004210305.1_ASM421030v1
100/100
29.1/54
Candidatus.Odyssella.thessalonicensis.L13 GCA_000190415.2_ASM19041v1
Candidatus.Paracaedibacter.acanthamoebae GCA_000742835.1_ASM74283v1
100/100
100/100
Candidatus.Hepatobacter.penaei GCA_000742475.1_ASM74247v1
Holospora.undulata.HU1 GCA_000388175.3_ASM38817v3
Elioraea.tepidiphila.DSM.17972 GCA_000378465.1_ASM37846v1
100/100
Belnapia.moabensis.DSM.16746 GCA_000745835.1_ASM74583v1
100/100
100/100
Rubritepida.flocculans.DSM.14296 GCA_000425365.1_ASM42536v1
100/100
Roseomonas.gilardii GCA_001941945.1_ASM194194v1
Acidiphilium.multivorum.AIU301 GCA_000202835.1_ASM20283v1
93.7/100
89.2/97
Acidisphaera.rubrifaciens.HS.AP3 GCA_000964365.1_ASM96436v1
100/100
Granulibacter.bethesdensis.CGDNIH4 GCA_000576145.2_ASM57614v2
100/100
Commensalibacter.intestini.A911 GCA_000231445.2_ASM23144v1
95.5/99
Gluconacetobacter.diazotrophicus.PA1.5 GCA_000067045.1_ASM6704v1
Acetobacter.aceti.NBRC.14818 GCA_000193495.2_ASM19349v2
Gluconobacter.oxydans.DSM.3504 GCA_000583855.1_ASM58385v1
0.541523

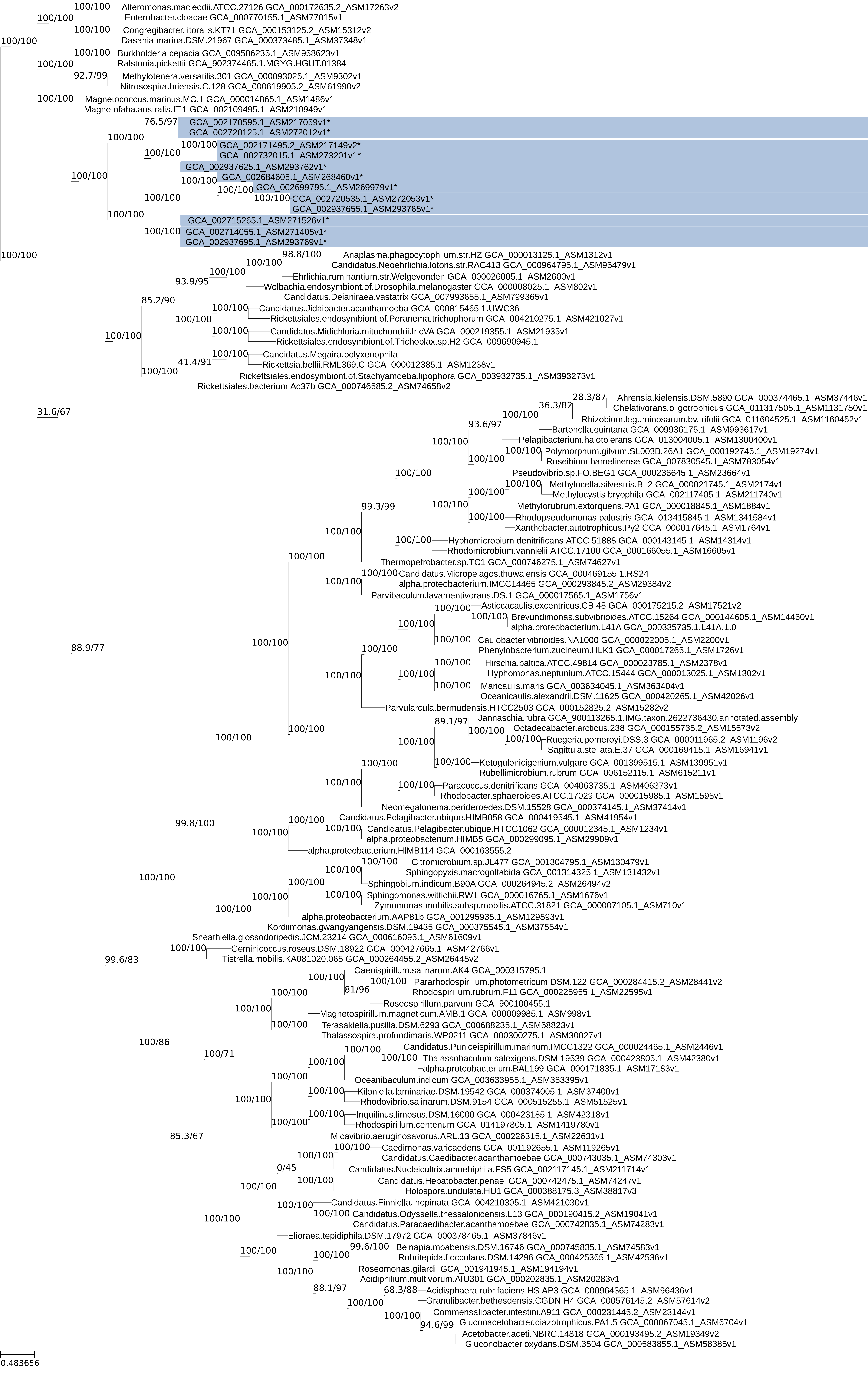

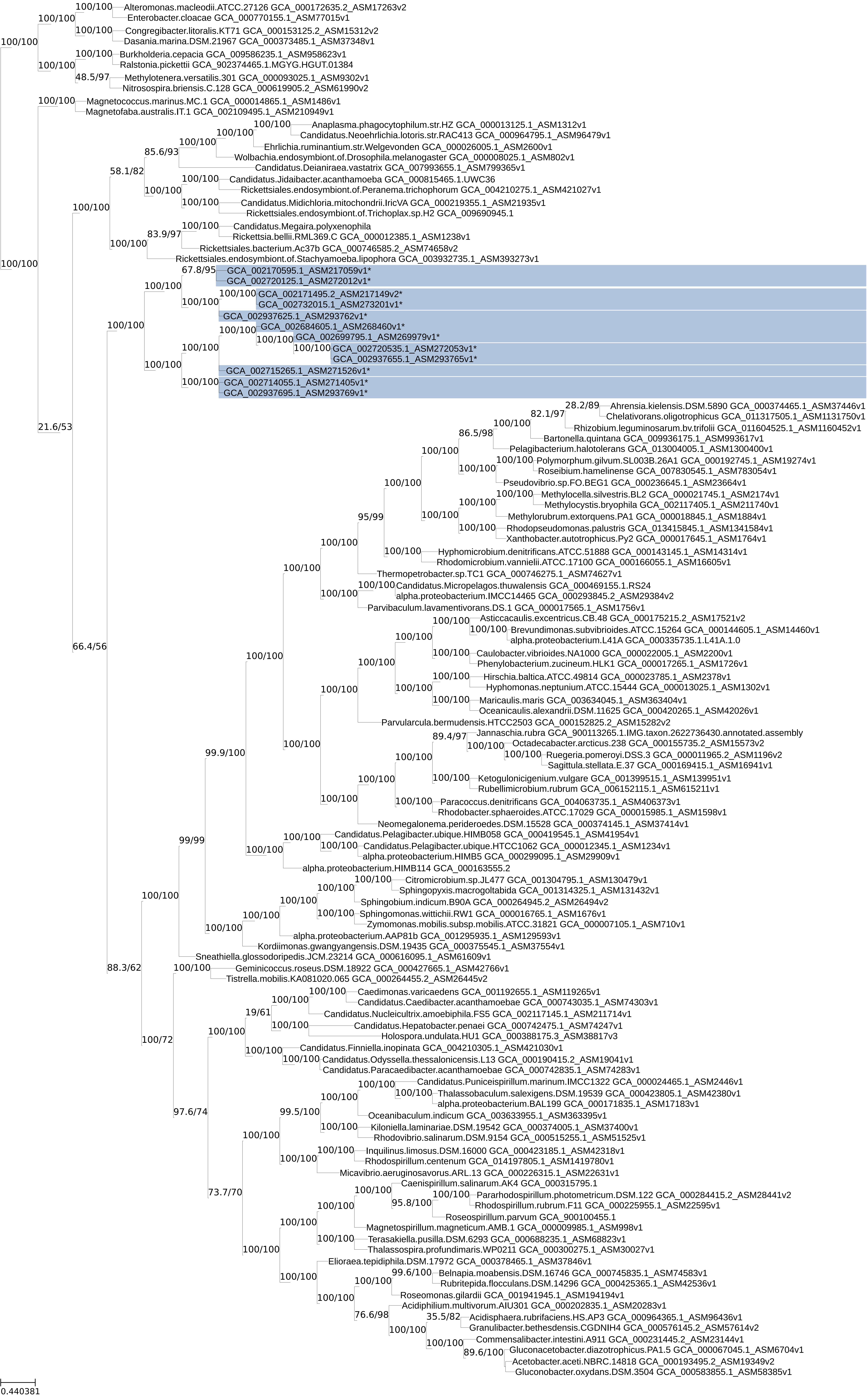
100/100
Alteromonas.macleodii.ATCC.27126 GCA_000172635.2_ASM17263v2
100/100
Enterobacter.cloacae GCA_000770155.1_ASM77015v1
100/100
100/100
Congregibacter.litoralis.KT71 GCA_000153125.2_ASM15312v2
Dasania.marina.DSM.21967 GCA_000373485.1_ASM37348v1
100/100
Burkholderia.cepacia GCA_009586235.1_ASM958623v1
100/100
Ralstonia.pickettii GCA_902374465.1.MGYG.HGUT.01384
48.5/97
Methylotenera.versatilis.301 GCA_000093025.1_ASM9302v1
Nitrosospira.briensis.C.128 GCA_000619905.2_ASM61990v2
100/100
Magnetococcus.marinus.MC.1 GCA_000014865.1_ASM1486v1
Magnetofaba.australis.IT.1 GCA_002109495.1_ASM210949v1
100/100
Anaplasma.phagocytophilum.str.HZ GCA_000013125.1_ASM1312v1
100/100
Candidatus.Neoehrlichia.lotoris.str.RAC413 GCA_000964795.1_ASM96479v1
100/100
Ehrlichia.ruminantium.str.Welgevonden GCA_000026005.1_ASM2600v1
85.6/93
Wolbachia.endosymbiont.of.Drosophila.melanogaster GCA_000008025.1_ASM802v1
Candidatus.Deianiraea.vastatrix GCA_007993655.1_ASM799365v1
58.1/82
100/100
Candidatus.Jidaibacter.acanthamoeba GCA_000815465.1.UWC36
100/100
Rickettsiales.endosymbiont.of.Peranema.trichophorum GCA_004210275.1_ASM421027v1
100/100
100/100
Candidatus.Midichloria.mitochondrii.IricVA GCA_000219355.1_ASM21935v1
Rickettsiales.endosymbiont.of.Trichoplax.sp.H2 GCA_009690945.1
100/100
Candidatus.Megaira.polyxenophila
83.9/97
Rickettsia.bellii.RML369.C GCA_000012385.1_ASM1238v1
100/100
Rickettsiales.bacterium.Ac37b GCA_000746585.2_ASM74658v2
Rickettsiales.endosymbiont.of.Stachyamoeba.lipophora GCA_003932735.1_ASM393273v1
100/100
67.8/95
GCA_002170595.1_ASM217059v1*
GCA_002720125.1_ASM272012v1*
100/100
100/100
GCA_002171495.2_ASM217149v2*
100/100
GCA_002732015.1_ASM273201v1*
GCA_002937625.1_ASM293762v1*
100/100
100/100
GCA_002684605.1_ASM268460v1*
100/100
GCA_002699795.1_ASM269979v1*
100/100
100/100
GCA_002720535.1_ASM272053v1*
100/100
GCA_002937655.1_ASM293765v1*
GCA_002715265.1_ASM271526v1*
100/100
GCA_002714055.1_ASM271405v1*
GCA_002937695.1_ASM293769v1*
28.2/89
Ahrensia.kielensis.DSM.5890 GCA_000374465.1_ASM37446v1
82.1/97
Chelativorans.oligotrophicus GCA_011317505.1_ASM1131750v1
21.6/53
100/100
Rhizobium.leguminosarum.bv.trifolii GCA_011604525.1_ASM1160452v1
86.5/98
Bartonella.quintana GCA_009936175.1_ASM993617v1
Pelagibacterium.halotolerans GCA_013004005.1_ASM1300400v1
100/100
100/100
Polymorphum.gilvum.SL003B.26A1 GCA_000192745.1_ASM19274v1
100/100
Roseibium.hamelinense GCA_007830545.1_ASM783054v1
100/100
Pseudovibrio.sp.FO.BEG1 GCA_000236645.1_ASM23664v1
100/100
Methylocella.silvestris.BL2 GCA_000021745.1_ASM2174v1
100/100
Methylocystis.bryophila GCA_002117405.1_ASM211740v1
100/100
Methylorubrum.extorquens.PA1 GCA_000018845.1_ASM1884v1
95/99
100/100
Rhodopseudomonas.palustris GCA_013415845.1_ASM1341584v1
Xanthobacter.autotrophicus.Py2 GCA_000017645.1_ASM1764v1
100/100
100/100
Hyphomicrobium.denitrificans.ATCC.51888 GCA_000143145.1_ASM14314v1
Rhodomicrobium.vannielii.ATCC.17100 GCA_000166055.1_ASM16605v1
100/100
Thermopetrobacter.sp.TC1 GCA_000746275.1_ASM74627v1
100/100
Candidatus.Micropelagos.thuwalensis GCA_000469155.1.RS24
100/100
alpha.proteobacterium.IMCC14465 GCA_000293845.2_ASM29384v2
Parvibaculum.lavamentivorans.DS.1 GCA_000017565.1_ASM1756v1
100/100
Asticcacaulis.excentricus.CB.48 GCA_000175215.2_ASM17521v2
100/100
100/100
Brevundimonas.subvibrioides.ATCC.15264 GCA_000144605.1_ASM14460v1
alpha.proteobacterium.L41A GCA_000335735.1.L41A.1.0
66.4/56
100/100
100/100
Caulobacter.vibrioides.NA1000 GCA_000022005.1_ASM2200v1
100/100
Phenylobacterium.zucineum.HLK1 GCA_000017265.1_ASM1726v1
100/100
Hirschia.baltica.ATCC.49814 GCA_000023785.1_ASM2378v1
100/100
100/100
Hyphomonas.neptunium.ATCC.15444 GCA_000013025.1_ASM1302v1
100/100
Maricaulis.maris GCA_003634045.1_ASM363404v1
Oceanicaulis.alexandrii.DSM.11625 GCA_000420265.1_ASM42026v1
Parvularcula.bermudensis.HTCC2503 GCA_000152825.2_ASM15282v2
89.4/97
Jannaschia.rubra GCA_900113265.1.IMG.taxon.2622736430.annotated.assembly
100/100
Octadecabacter.arcticus.238 GCA_000155735.2_ASM15573v2
100/100
Ruegeria.pomeroyi.DSS.3 GCA_000011965.2_ASM1196v2
99.9/100
100/100
Sagittula.stellata.E.37 GCA_000169415.1_ASM16941v1
100/100
Ketogulonicigenium.vulgare GCA_001399515.1_ASM139951v1
100/100
Rubellimicrobium.rubrum GCA_006152115.1_ASM615211v1
100/100
100/100
Paracoccus.denitrificans GCA_004063735.1_ASM406373v1
Rhodobacter.sphaeroides.ATCC.17029 GCA_000015985.1_ASM1598v1
Neomegalonema.perideroedes.DSM.15528 GCA_000374145.1_ASM37414v1
100/100
Candidatus.Pelagibacter.ubique.HIMB058 GCA_000419545.1_ASM41954v1
99/99
100/100
100/100
Candidatus.Pelagibacter.ubique.HTCC1062 GCA_000012345.1_ASM1234v1
alpha.proteobacterium.HIMB5 GCA_000299095.1_ASM29909v1
alpha.proteobacterium.HIMB114 GCA_000163555.2
100/100
Citromicrobium.sp.JL477 GCA_001304795.1_ASM130479v1
100/100
Sphingopyxis.macrogoltabida GCA_001314325.1_ASM131432v1
100/100
100/100
Sphingobium.indicum.B90A GCA_000264945.2_ASM26494v2
100/100
Sphingomonas.wittichii.RW1 GCA_000016765.1_ASM1676v1
100/100
Zymomonas.mobilis.subsp.mobilis.ATCC.31821 GCA_000007105.1_ASM710v1
100/100
alpha.proteobacterium.AAP81b GCA_001295935.1_ASM129593v1
Kordiimonas.gwangyangensis.DSM.19435 GCA_000375545.1_ASM37554v1
Sneathiella.glossodoripedis.JCM.23214 GCA_000616095.1_ASM61609v1
88.3/62
100/100
Geminicoccus.roseus.DSM.18922 GCA_000427665.1_ASM42766v1
Tistrella.mobilis.KA081020.065 GCA_000264455.2_ASM26445v2
100/100
Caedimonas.varicaedens GCA_001192655.1_ASM119265v1
100/100
Candidatus.Caedibacter.acanthamoebae GCA_000743035.1_ASM74303v1
19/61
Candidatus.Nucleicultrix.amoebiphila.FS5 GCA_002117145.1_ASM211714v1
100/100
Candidatus.Hepatobacter.penaei GCA_000742475.1_ASM74247v1
100/100
100/72
Holospora.undulata.HU1 GCA_000388175.3_ASM38817v3
Candidatus.Finniella.inopinata GCA_004210305.1_ASM421030v1
100/100
100/100
Candidatus.Odyssella.thessalonicensis.L13 GCA_000190415.2_ASM19041v1
Candidatus.Paracaedibacter.acanthamoebae GCA_000742835.1_ASM74283v1
100/100
Candidatus.Puniceispirillum.marinum.IMCC1322 GCA_000024465.1_ASM2446v1
100/100
100/100
Thalassobaculum.salexigens.DSM.19539 GCA_000423805.1_ASM42380v1
alpha.proteobacterium.BAL199 GCA_000171835.1_ASM17183v1
99.5/100
97.6/74
Oceanibaculum.indicum GCA_003633955.1_ASM363395v1
100/100
Kiloniella.laminariae.DSM.19542 GCA_000374005.1_ASM37400v1
Rhodovibrio.salinarum.DSM.9154 GCA_000515255.1_ASM51525v1
100/100
100/100
Inquilinus.limosus.DSM.16000 GCA_000423185.1_ASM42318v1
100/100
Rhodospirillum.centenum GCA_014197805.1_ASM1419780v1
Micavibrio.aeruginosavorus.ARL.13 GCA_000226315.1_ASM22631v1
100/100
Caenispirillum.salinarum.AK4 GCA_000315795.1
100/100
Pararhodospirillum.photometricum.DSM.122 GCA_000284415.2_ASM28441v2
73.7/70
95.8/100
Rhodospirillum.rubrum.F11 GCA_000225955.1_ASM22595v1
100/100
Roseospirillum.parvum GCA_900100455.1
100/100
Magnetospirillum.magneticum.AMB.1 GCA_000009985.1_ASM998v1
100/100
Terasakiella.pusilla.DSM.6293 GCA_000688235.1_ASM68823v1
100/100
Thalassospira.profundimaris.WP0211 GCA_000300275.1_ASM30027v1
Elioraea.tepidiphila.DSM.17972 GCA_000378465.1_ASM37846v1
99.6/100
Belnapia.moabensis.DSM.16746 GCA_000745835.1_ASM74583v1
100/100
100/100
Rubritepida.flocculans.DSM.14296 GCA_000425365.1_ASM42536v1
100/100
Roseomonas.gilardii GCA_001941945.1_ASM194194v1
Acidiphilium.multivorum.AIU301 GCA_000202835.1_ASM20283v1
35.5/82
76.6/98
Acidisphaera.rubrifaciens.HS.AP3 GCA_000964365.1_ASM96436v1
Granulibacter.bethesdensis.CGDNIH4 GCA_000576145.2_ASM57614v2
100/100
Commensalibacter.intestini.A911 GCA_000231445.2_ASM23144v1
100/100
Gluconacetobacter.diazotrophicus.PA1.5 GCA_000067045.1_ASM6704v1
89.6/100
Acetobacter.aceti.NBRC.14818 GCA_000193495.2_ASM19349v2
Gluconobacter.oxydans.DSM.3504 GCA_000583855.1_ASM58385v1
0.440381

### Group L

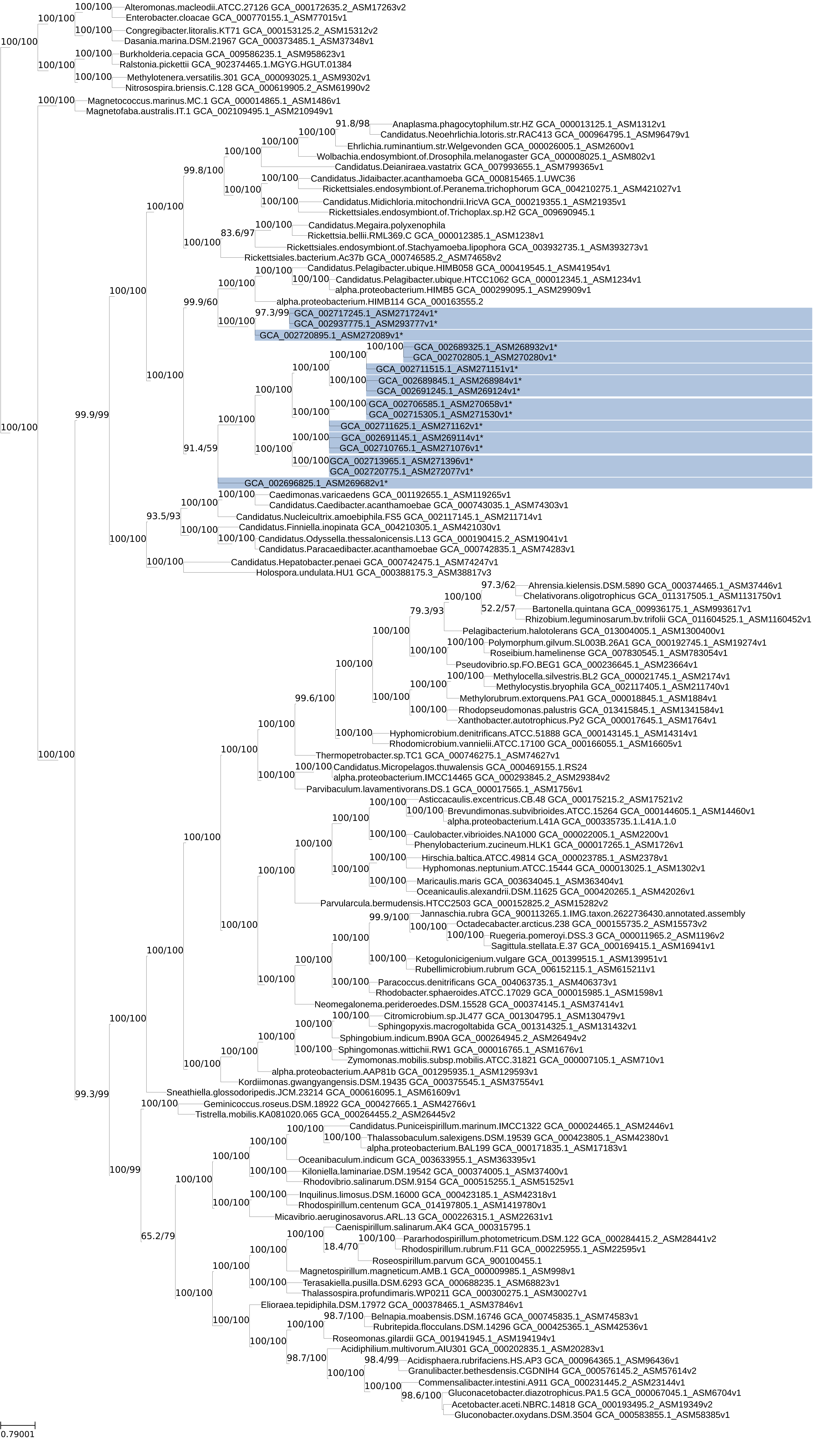

Alteromonas.macleodii.ATCC.27126 GCA_000172635.2_ASM17263v2
Enterobacter.cloacae GCA_000770155.1_ASM77015v1
Congregibacter.litoralis.KT71 GCA_000153125.2_ASM15312v2
Dasania.marina.DSM.21967 GCA_000373485.1_ASM37348v1
Burkholderia.cepacia GCA_009586235.1_ASM958623v1
Ralstonia.pickettii GCA_902374465.1.MGYG.HGUT.01384
Methylotenera.versatilis.301 GCA_000093025.1_ASM9302v1
Nitrosospira.briensis.C.128 GCA_000619905.2_ASM61990v2
Magnetococcus.marinus.MC.1 GCA_000014865.1_ASM1486v1
Magnetofaba.australis.IT.1 GCA_002109495.1_ASM210949v1
Anaplasma.phagocytophilum.str.HZ GCA_000013125.1_ASM1312v1
Candidatus.Neoehrlichia.lotoris.str.RAC413 GCA_000964795.1_ASM96479v1
Ehrlichia.ruminantium.str.Welgevonden GCA_000026005.1_ASM2600v1
Wolbachia.endosymbiont.of.Drosophila.melanogaster GCA_000008025.1_ASM802v1
Candidatus.Deianiraea.vastatrix GCA_007993655.1_ASM799365v1
Candidatus.Jidaibacter.acanthamoeba GCA_000815465.1.UWC36
Rickettsiales.endosymbiont.of.Peranema.trichophorum GCA_004210275.1_ASM421027v1
Candidatus.Midichloria.mitochondrii.IricVA GCA_000219355.1_ASM21935v1
Rickettsiales.endosymbiont.of.Trichoplax.sp.H2 GCA_009690945.1
Candidatus.Megaira.polyxenophila
Rickettsia.bellii.RML369.C GCA_000012385.1_ASM1238v1
Rickettsiales.endosymbiont.of.Stachyamoeba.lipophora GCA_003932735.1_ASM393273v1
Rickettsiales.bacterium.Ac37b GCA_000746585.2_ASM74658v2
Candidatus.Pelagibacter.ubique.HIMB058 GCA_000419545.1_ASM41954v1
Candidatus.Pelagibacter.ubique.HTCC1062 GCA_000012345.1_ASM1234v1
alpha.proteobacterium.HIMB5 GCA_000299095.1_ASM29909v1
alpha.proteobacterium.HIMB114 GCA_000163555.2
GCA_002717245.1_ASM271724v1*
GCA_002937775.1_ASM293777v1*
GCA_002720895.1_ASM272089v1*
GCA_002689325.1_ASM268932v1*
GCA_002702805.1_ASM270280v1*
GCA_002711515.1_ASM271151v1*
GCA_002689845.1_ASM268984v1*
GCA_002691245.1_ASM269124v1*
GCA_002706585.1_ASM270658v1*
GCA_002715305.1_ASM271530v1*
GCA_002711625.1_ASM271162v1*
GCA_002691145.1_ASM269114v1*
GCA_002710765.1_ASM271076v1*
GCA_002713965.1_ASM271396v1*
GCA_002720775.1_ASM272077v1*
GCA_002696825.1_ASM269682v1*
Caedimonas.varicaedens GCA_001192655.1_ASM119265v1
Candidatus.Caedibacter.acanthamoebae GCA_000743035.1_ASM74303v1
Candidatus.Nucleicultrix.amoebiphila.FS5 GCA_002117145.1_ASM211714v1
Candidatus.Finniella.inopinata GCA_004210305.1_ASM421030v1
Candidatus.Odyssella.thessalonicensis.L13 GCA_000190415.2_ASM19041v1
Candidatus.Paracaedibacter.acanthamoebae GCA_000742835.1_ASM74283v1
Candidatus.Hepatobacter.penaei GCA_000742475.1_ASM74247v1
Holospora.undulata.HU1 GCA_000388175.3_ASM38817v3
Ahrensia.kielensis.DSM.5890 GCA_000374465.1_ASM37446v1
Chelativorans.oligotrophicus GCA_011317505.1_ASM1131750v1
Bartonella.quintana GCA_009936175.1_ASM993617v1
Rhizobium.leguminosarum.bv.trifolii GCA_011604525.1_ASM1160452v1
Pelagibacterium.halotolerans GCA_013004005.1_ASM1300400v1
Polymorphum.gilvum.SL003B.26A1 GCA_000192745.1_ASM19274v1
Roseibium.hamelinense GCA_007830545.1_ASM783054v1
Pseudovibrio.sp.FO.BEG1 GCA_000236645.1_ASM23664v1
Methylocella.silvestris.BL2 GCA_000021745.1_ASM2174v1
Methylocystis.bryophila GCA_002117405.1_ASM211740v1
Methylorubrum.extorquens.PA1 GCA_000018845.1_ASM1884v1
Rhodopseudomonas.palustris GCA_013415845.1_ASM1341584v1
Xanthobacter.autotrophicus.Py2 GCA_000017645.1_ASM1764v1
Hyphomicrobium.denitrificans.ATCC.51888 GCA_000143145.1_ASM14314v1
Rhodomicrobium.vannielii.ATCC.17100 GCA_000166055.1_ASM16605v1
Thermopetrobacter.sp.TC1 GCA_000746275.1_ASM74627v1
Candidatus.Micropelagos.thuwalensis GCA_000469155.1.RS24
alpha.proteobacterium.IMCC14465 GCA_000293845.2_ASM29384v2
Parvibaculum.lavamentivorans.DS.1 GCA_000017565.1_ASM1756v1
Asticcacaulis.excentricus.CB.48 GCA_000175215.2_ASM17521v2
Brevundimonas.subvibrioides.ATCC.15264 GCA_000144605.1_ASM14460v1
alpha.proteobacterium.L41A GCA_000335735.1.L41A.1.0
Caulobacter.vibrioides.NA1000 GCA_000022005.1_ASM2200v1
Phenylobacterium.zucineum.HLK1 GCA_000017265.1_ASM1726v1
Hirschia.baltica.ATCC.49814 GCA_000023785.1_ASM2378v1
Hyphomonas.neptunium.ATCC.15444 GCA_000013025.1_ASM1302v1
Maricaulis.maris GCA_003634045.1_ASM363404v1
Oceanicaulis.alexandrii.DSM.11625 GCA_000420265.1_ASM42026v1
Parvularcula.bermudensis.HTCC2503 GCA_000152825.2_ASM15282v2
Jannaschia.rubra GCA_900113265.1.IMG.taxon.2622736430.annotated.assembly
Octadecabacter.arcticus.238 GCA_000155735.2_ASM15573v2
Ruegeria.pomeroyi.DSS.3 GCA_000011965.2_ASM1196v2
Sagittula.stellata.E.37 GCA_000169415.1_ASM16941v1
Ketogulonicigenium.vulgare GCA_001399515.1_ASM139951v1
Rubellimicrobium.rubrum GCA_006152115.1_ASM615211v1
Paracoccus.denitrificans GCA_004063735.1_ASM406373v1
Rhodobacter.sphaeroides.ATCC.17029 GCA_000015985.1_ASM1598v1
Neomegalonema.perideroedes.DSM.15528 GCA_000374145.1_ASM37414v1
Citromicrobium.sp.JL477 GCA_001304795.1_ASM130479v1
Sphingopyxis.macrogoltabida GCA_001314325.1_ASM131432v1
Sphingobium.indicum.B90A GCA_000264945.2_ASM26494v2
Sphingomonas.wittichii.RW1 GCA_000016765.1_ASM1676v1
Zymomonas.mobilis.subsp.mobilis.ATCC.31821 GCA_000007105.1_ASM710v1
alpha.proteobacterium.AAP81b GCA_001295935.1_ASM129593v1
Kordiimonas.gwangyangensis.DSM.19435 GCA_000375545.1_ASM37554v1
Sneathiella.glossodoripedis.JCM.23214 GCA_000616095.1_ASM61609v1
Geminicoccus.roseus.DSM.18922 GCA_000427665.1_ASM42766v1
Tistrella.mobilis.KA081020.065 GCA_000264455.2_ASM26445v2
Candidatus.Puniceispirillum.marinum.IMCC1322 GCA_000024465.1_ASM2446v1
Thalassobaculum.salexigens.DSM.19539 GCA_000423805.1_ASM42380v1
alpha.proteobacterium.BAL199 GCA_000171835.1_ASM17183v1
Oceanibaculum.indicum GCA_003633955.1_ASM363395v1
Kiloniella.laminariae.DSM.19542 GCA_000374005.1_ASM37400v1
Rhodovibrio.salinarum.DSM.9154 GCA_000515255.1_ASM51525v1
Inquilinus.limosus.DSM.16000 GCA_000423185.1_ASM42318v1
Rhodospirillum.centenum GCA_014197805.1_ASM1419780v1
Micavibrio.aeruginosavorus.ARL.13 GCA_000226315.1_ASM22631v1
Caenispirillum.salinarum.AK4 GCA_000315795.1
Pararhodospirillum.photometricum.DSM.122 GCA_000284415.2_ASM28441v2
Rhodospirillum.rubrum.F11 GCA_000225955.1_ASM22595v1
Roseospirillum.parvum GCA_900100455.1
Magnetospirillum.magneticum.AMB.1 GCA_000009985.1_ASM998v1
Terasakiella.pusilla.DSM.6293 GCA_000688235.1_ASM68823v1
Thalassospira.profundimaris.WP0211 GCA_000300275.1_ASM30027v1
Elioraea.tepidiphila.DSM.17972 GCA_000378465.1_ASM37846v1
Belnapia.moabensis.DSM.16746 GCA_000745835.1_ASM74583v1
Rubritepida.flocculans.DSM.14296 GCA_000425365.1_ASM42536v1
Roseomonas.gilardii GCA_001941945.1_ASM194194v1
Acidiphilium.multivorum.AIU301 GCA_000202835.1_ASM20283v1
Acidisphaera.rubrifaciens.HS.AP3 GCA_000964365.1_ASM96436v1
Granulibacter.bethesdensis.CGDNIH4 GCA_000576145.2_ASM57614v2
Commensalibacter.intestini.A911 GCA_000231445.2_ASM23144v1
Gluconacetobacter.diazotrophicus.PA1.5 GCA_000067045.1_ASM6704v1
Acetobacter.aceti.NBRC.14818 GCA_000193495.2_ASM19349v2
Gluconobacter.oxydans.DSM.3504 GCA_000583855.1_ASM58385v1
0.79001

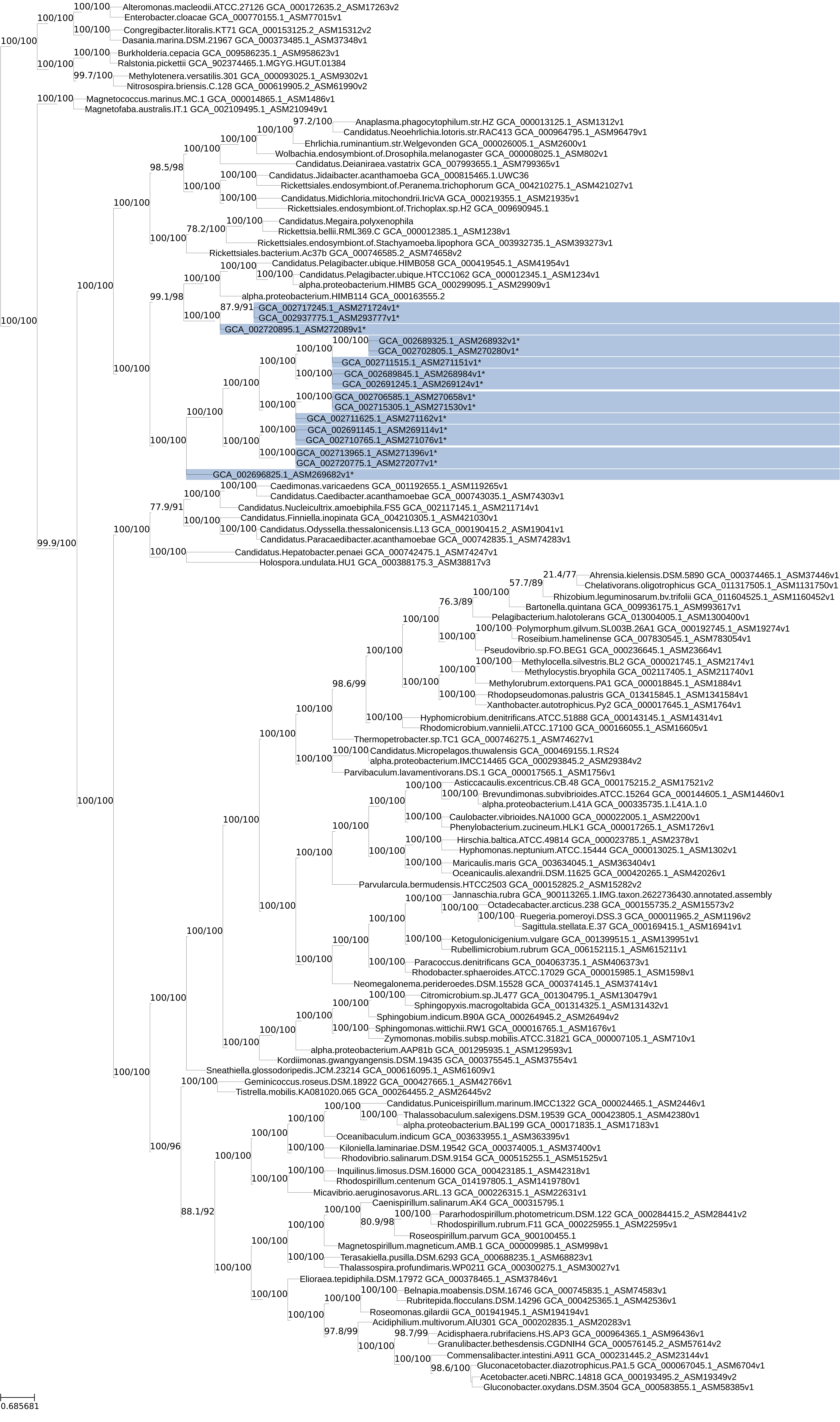

Alteromonas.macleodii.ATCC.27126 GCA_000172635.2_ASM17263v2
Enterobacter.cloacae GCA_000770155.1_ASM77015v1
Congregibacter.litoralis.KT71 GCA_000153125.2_ASM15312v2
Dasania.marina.DSM.21967 GCA_000373485.1_ASM37348v1
Burkholderia.cepacia GCA_009586235.1_ASM958623v1
Ralstonia.pickettii GCA_902374465.1.MGYG.HGUT.01384
Methylotenera.versatilis.301 GCA_000093025.1_ASM9302v1
Nitrosospira.briensis.C.128 GCA_000619905.2_ASM61990v2
Magnetococcus.marinus.MC.1 GCA_000014865.1_ASM1486v1
Magnetofaba.australis.IT.1 GCA_002109495.1_ASM210949v1
Anaplasma.phagocytophilum.str.HZ GCA_000013125.1_ASM1312v1
Candidatus.Neoehrlichia.lotoris.str.RAC413 GCA_000964795.1_ASM96479v1
Ehrlichia.ruminantium.str.Welgevonden GCA_000026005.1_ASM2600v1
Wolbachia.endosymbiont.of.Drosophila.melanogaster GCA_000008025.1_ASM802v1
Candidatus.Deianiraea.vastatrix GCA_007993655.1_ASM799365v1
Candidatus.Jidaibacter.acanthamoeba GCA_000815465.1.UWC36
Rickettsiales.endosymbiont.of.Peranema.trichophorum GCA_004210275.1_ASM421027v1
Candidatus.Midichloria.mitochondrii.IricVA GCA_000219355.1_ASM21935v1
Rickettsiales.endosymbiont.of.Trichoplax.sp.H2 GCA_009690945.1
Candidatus.Megaira.polyxenophila
Rickettsia.bellii.RML369.C GCA_000012385.1_ASM1238v1
Rickettsiales.endosymbiont.of.Stachyamoeba.lipophora GCA_003932735.1_ASM393273v1
Rickettsiales.bacterium.Ac37b GCA_000746585.2_ASM74658v2
Candidatus.Pelagibacter.ubique.HIMB058 GCA_000419545.1_ASM41954v1
Candidatus.Pelagibacter.ubique.HTCC1062 GCA_000012345.1_ASM1234v1
alpha.proteobacterium.HIMB5 GCA_000299095.1_ASM29909v1
alpha.proteobacterium.HIMB114 GCA_000163555.2
GCA_002717245.1_ASM271724v1*
GCA_002937775.1_ASM293777v1*
GCA_002720895.1_ASM272089v1*
GCA_002689325.1_ASM268932v1*
GCA_002702805.1_ASM270280v1*
GCA_002711515.1_ASM271151v1*
GCA_002689845.1_ASM268984v1*
GCA_002691245.1_ASM269124v1*
GCA_002706585.1_ASM270658v1*
GCA_002715305.1_ASM271530v1*
GCA_002711625.1_ASM271162v1*
GCA_002691145.1_ASM269114v1*
GCA_002710765.1_ASM271076v1*
GCA_002713965.1_ASM271396v1*
GCA_002720775.1_ASM272077v1*
GCA_002696825.1_ASM269682v1*
Caedimonas.varicaedens GCA_001192655.1_ASM119265v1
Candidatus.Caedibacter.acanthamoebae GCA_000743035.1_ASM74303v1
Candidatus.Nucleicultrix.amoebiphila.FS5 GCA_002117145.1_ASM211714v1
Candidatus.Finniella.inopinata GCA_004210305.1_ASM421030v1
Candidatus.Odyssella.thessalonicensis.L13 GCA_000190415.2_ASM19041v1
Candidatus.Paracaedibacter.acanthamoebae GCA_000742835.1_ASM74283v1
Candidatus.Hepatobacter.penaei GCA_000742475.1_ASM74247v1
Holospora.undulata.HU1 GCA_000388175.3_ASM38817v3
Ahrensia.kielensis.DSM.5890 GCA_000374465.1_ASM37446v1
Chelativorans.oligotrophicus GCA_011317505.1_ASM1131750v1
Rhizobium.leguminosarum.bv.trifolii GCA_011604525.1_ASM1160452v1
Bartonella.quintana GCA_009936175.1_ASM993617v1
Pelagibacterium.halotolerans GCA_013004005.1_ASM1300400v1
Polymorphum.gilvum.SL003B.26A1 GCA_000192745.1_ASM19274v1
Roseibium.hamelinense GCA_007830545.1_ASM783054v1
Pseudovibrio.sp.FO.BEG1 GCA_000236645.1_ASM23664v1
Methylocella.silvestris.BL2 GCA_000021745.1_ASM2174v1
Methylocystis.bryophila GCA_002117405.1_ASM211740v1
Methylorubrum.extorquens.PA1 GCA_000018845.1_ASM1884v1
Rhodopseudomonas.palustris GCA_013415845.1_ASM1341584v1
Xanthobacter.autotrophicus.Py2 GCA_000017645.1_ASM1764v1
Hyphomicrobium.denitrificans.ATCC.51888 GCA_000143145.1_ASM14314v1
Rhodomicrobium.vannielii.ATCC.17100 GCA_000166055.1_ASM16605v1
Thermopetrobacter.sp.TC1 GCA_000746275.1_ASM74627v1
Candidatus.Micropelagos.thuwalensis GCA_000469155.1.RS24
alpha.proteobacterium.IMCC14465 GCA_000293845.2_ASM29384v2
Parvibaculum.lavamentivorans.DS.1 GCA_000017565.1_ASM1756v1
Asticcacaulis.excentricus.CB.48 GCA_000175215.2_ASM17521v2
Brevundimonas.subvibrioides.ATCC.15264 GCA_000144605.1_ASM14460v1
alpha.proteobacterium.L41A GCA_000335735.1.L41A.1.0
Caulobacter.vibrioides.NA1000 GCA_000022005.1_ASM2200v1
Phenylobacterium.zucineum.HLK1 GCA_000017265.1_ASM1726v1
Hirschia.baltica.ATCC.49814 GCA_000023785.1_ASM2378v1
Hyphomonas.neptunium.ATCC.15444 GCA_000013025.1_ASM1302v1
Maricaulis.maris GCA_003634045.1_ASM363404v1
Oceanicaulis.alexandrii.DSM.11625 GCA_000420265.1_ASM42026v1
Parvularcula.bermudensis.HTCC2503 GCA_000152825.2_ASM15282v2
Jannaschia.rubra GCA_900113265.1.IMG.taxon.2622736430.annotated.assembly
Octadecabacter.arcticus.238 GCA_000155735.2_ASM15573v2
Ruegeria.pomeroyi.DSS.3 GCA_000011965.2_ASM1196v2
Sagittula.stellata.E.37 GCA_000169415.1_ASM16941v1
Ketogulonicigenium.vulgare GCA_001399515.1_ASM139951v1
Rubellimicrobium.rubrum GCA_006152115.1_ASM615211v1
Paracoccus.denitrificans GCA_004063735.1_ASM406373v1
Rhodobacter.sphaeroides.ATCC.17029 GCA_000015985.1_ASM1598v1
Neomegalonema.perideroedes.DSM.15528 GCA_000374145.1_ASM37414v1
Citromicrobium.sp.JL477 GCA_001304795.1_ASM130479v1
Sphingopyxis.macrogoltabida GCA_001314325.1_ASM131432v1
Sphingobium.indicum.B90A GCA_000264945.2_ASM26494v2
Sphingomonas.wittichii.RW1 GCA_000016765.1_ASM1676v1
Zymomonas.mobilis.subsp.mobilis.ATCC.31821 GCA_000007105.1_ASM710v1
alpha.proteobacterium.AAP81b GCA_001295935.1_ASM129593v1
Kordiimonas.gwangyangensis.DSM.19435 GCA_000375545.1_ASM37554v1
Sneathiella.glossodoripedis.JCM.23214 GCA_000616095.1_ASM61609v1
Geminicoccus.roseus.DSM.18922 GCA_000427665.1_ASM42766v1
Tistrella.mobilis.KA081020.065 GCA_000264455.2_ASM26445v2
Candidatus.Puniceispirillum.marinum.IMCC1322 GCA_000024465.1_ASM2446v1
Thalassobaculum.salexigens.DSM.19539 GCA_000423805.1_ASM42380v1
alpha.proteobacterium.BAL199 GCA_000171835.1_ASM17183v1
Oceanibaculum.indicum GCA_003633955.1_ASM363395v1
Kiloniella.laminariae.DSM.19542 GCA_000374005.1_ASM37400v1
Rhodovibrio.salinarum.DSM.9154 GCA_000515255.1_ASM51525v1
Inquilinus.limosus.DSM.16000 GCA_000423185.1_ASM42318v1
Rhodospirillum.centenum GCA_014197805.1_ASM1419780v1
Micavibrio.aeruginosavorus.ARL.13 GCA_000226315.1_ASM22631v1
Caenispirillum.salinarum.AK4 GCA_000315795.1
Pararhodospirillum.photometricum.DSM.122 GCA_000284415.2_ASM28441v2
Rhodospirillum.rubrum.F11 GCA_000225955.1_ASM22595v1
Roseospirillum.parvum GCA_900100455.1
Magnetospirillum.magneticum.AMB.1 GCA_000009985.1_ASM998v1
Terasakiella.pusilla.DSM.6293 GCA_000688235.1_ASM68823v1
Thalassospira.profundimaris.WP0211 GCA_000300275.1_ASM30027v1
Elioraea.tepidiphila.DSM.17972 GCA_000378465.1_ASM37846v1
Belnapia.moabensis.DSM.16746 GCA_000745835.1_ASM74583v1
Rubritepida.flocculans.DSM.14296 GCA_000425365.1_ASM42536v1
Roseomonas.gilardii GCA_001941945.1_ASM194194v1
Acidiphilium.multivorum.AIU301 GCA_000202835.1_ASM20283v1
Acidisphaera.rubrifaciens.HS.AP3 GCA_000964365.1_ASM96436v1
Granulibacter.bethesdensis.CGDNIH4 GCA_000576145.2_ASM57614v2
Commensalibacter.intestini.A911 GCA_000231445.2_ASM23144v1
Gluconacetobacter.diazotrophicus.PA1.5 GCA_000067045.1_ASM6704v1
Acetobacter.aceti.NBRC.14818 GCA_000193495.2_ASM19349v2
Gluconobacter.oxydans.DSM.3504 GCA_000583855.1_ASM58385v1
0.685681

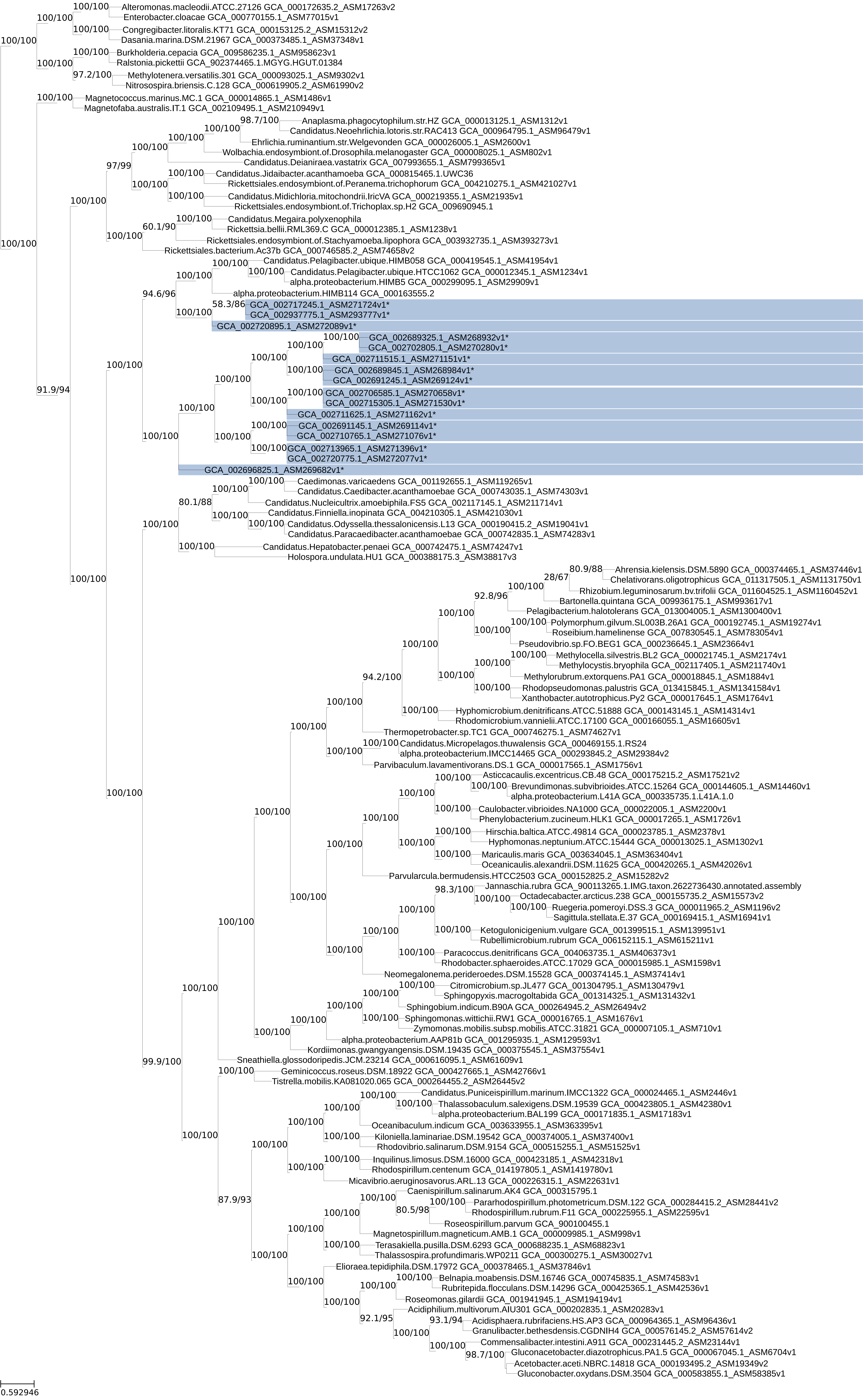

Alteromonas.macleodii.ATCC.27126 GCA_000172635.2_ASM17263v2
Enterobacter.cloacae GCA_000770155.1_ASM77015v1
Congregibacter.litoralis.KT71 GCA_000153125.2_ASM15312v2
Dasania.marina.DSM.21967 GCA_000373485.1_ASM37348v1
Burkholderia.cepacia GCA_009586235.1_ASM958623v1
Ralstonia.pickettii GCA_902374465.1.MGYG.HGUT.01384
Methylotenera.versatilis.301 GCA_000093025.1_ASM9302v1
Nitrosospira.briensis.C.128 GCA_000619905.2_ASM61990v2
Magnetococcus.marinus.MC.1 GCA_000014865.1_ASM1486v1
Magnetofaba.australis.IT.1 GCA_002109495.1_ASM210949v1
Anaplasma.phagocytophilum.str.HZ GCA_000013125.1_ASM1312v1
Candidatus.Neoehrlichia.lotoris.str.RAC413 GCA_000964795.1_ASM96479v1
Ehrlichia.ruminantium.str.Welgevonden GCA_000026005.1_ASM2600v1
Wolbachia.endosymbiont.of.Drosophila.melanogaster GCA_000008025.1_ASM802v1
Candidatus.Deianiraea.vastatrix GCA_007993655.1_ASM799365v1
Candidatus.Jidaibacter.acanthamoeba GCA_000815465.1.UWC36
Rickettsiales.endosymbiont.of.Peranema.trichophorum GCA_004210275.1_ASM421027v1
Candidatus.Midichloria.mitochondrii.IricVA GCA_000219355.1_ASM21935v1
Rickettsiales.endosymbiont.of.Trichoplax.sp.H2 GCA_009690945.1
Candidatus.Megaira.polyxenophila
Rickettsia.bellii.RML369.C GCA_000012385.1_ASM1238v1
Rickettsiales.endosymbiont.of.Stachyamoeba.lipophora GCA_003932735.1_ASM393273v1
Rickettsiales.bacterium.Ac37b GCA_000746585.2_ASM74658v2
Candidatus.Pelagibacter.ubique.HIMB058 GCA_000419545.1_ASM41954v1
Candidatus.Pelagibacter.ubique.HTCC1062 GCA_000012345.1_ASM1234v1
alpha.proteobacterium.HIMB5 GCA_000299095.1_ASM29909v1
alpha.proteobacterium.HIMB114 GCA_000163555.2
GCA_002717245.1_ASM271724v1*
GCA_002937775.1_ASM293777v1*
GCA_002720895.1_ASM272089v1*
GCA_002689325.1_ASM268932v1*
GCA_002702805.1_ASM270280v1*
GCA_002711515.1_ASM271151v1*
GCA_002689845.1_ASM268984v1*
GCA_002691245.1_ASM269124v1*
GCA_002706585.1_ASM270658v1*
GCA_002715305.1_ASM271530v1*
GCA_002711625.1_ASM271162v1*
GCA_002691145.1_ASM269114v1*
GCA_002710765.1_ASM271076v1*
GCA_002713965.1_ASM271396v1*
GCA_002720775.1_ASM272077v1*
GCA_002696825.1_ASM269682v1*
Caedimonas.varicaedens GCA_001192655.1_ASM119265v1
Candidatus.Caedibacter.acanthamoebae GCA_000743035.1_ASM74303v1
Candidatus.Nucleicultrix.amoebiphila.FS5 GCA_002117145.1_ASM211714v1
Candidatus.Finniella.inopinata GCA_004210305.1_ASM421030v1
Candidatus.Odyssella.thessalonicensis.L13 GCA_000190415.2_ASM19041v1
Candidatus.Paracaedibacter.acanthamoebae GCA_000742835.1_ASM74283v1
Candidatus.Hepatobacter.penaei GCA_000742475.1_ASM74247v1
Holospora.undulata.HU1 GCA_000388175.3_ASM38817v3
Ahrensia.kielensis.DSM.5890 GCA_000374465.1_ASM37446v1
Chelativorans.oligotrophicus GCA_011317505.1_ASM1131750v1
Rhizobium.leguminosarum.bv.trifolii GCA_011604525.1_ASM1160452v1
Bartonella.quintana GCA_009936175.1_ASM993617v1
Pelagibacterium.halotolerans GCA_013004005.1_ASM1300400v1
Polymorphum.gilvum.SL003B.26A1 GCA_000192745.1_ASM19274v1
Roseibium.hamelinense GCA_007830545.1_ASM783054v1
Pseudovibrio.sp.FO.BEG1 GCA_000236645.1_ASM23664v1
Methylocella.silvestris.BL2 GCA_000021745.1_ASM2174v1
Methylocystis.bryophila GCA_002117405.1_ASM211740v1
Methylorubrum.extorquens.PA1 GCA_000018845.1_ASM1884v1
Rhodopseudomonas.palustris GCA_013415845.1_ASM1341584v1
Xanthobacter.autotrophicus.Py2 GCA_000017645.1_ASM1764v1
Hyphomicrobium.denitrificans.ATCC.51888 GCA_000143145.1_ASM14314v1
Rhodomicrobium.vannielii.ATCC.17100 GCA_000166055.1_ASM16605v1
Thermopetrobacter.sp.TC1 GCA_000746275.1_ASM74627v1
Candidatus.Micropelagos.thuwalensis GCA_000469155.1.RS24
alpha.proteobacterium.IMCC14465 GCA_000293845.2_ASM29384v2
Parvibaculum.lavamentivorans.DS.1 GCA_000017565.1_ASM1756v1
Asticcacaulis.excentricus.CB.48 GCA_000175215.2_ASM17521v2
Brevundimonas.subvibrioides.ATCC.15264 GCA_000144605.1_ASM14460v1
alpha.proteobacterium.L41A GCA_000335735.1.L41A.1.0
Caulobacter.vibrioides.NA1000 GCA_000022005.1_ASM2200v1
Phenylobacterium.zucineum.HLK1 GCA_000017265.1_ASM1726v1
Hirschia.baltica.ATCC.49814 GCA_000023785.1_ASM2378v1
Hyphomonas.neptunium.ATCC.15444 GCA_000013025.1_ASM1302v1
Maricaulis.maris GCA_003634045.1_ASM363404v1
Oceanicaulis.alexandrii.DSM.11625 GCA_000420265.1_ASM42026v1
Parvularcula.bermudensis.HTCC2503 GCA_000152825.2_ASM15282v2
Jannaschia.rubra GCA_900113265.1.IMG.taxon.2622736430.annotated.assembly
Octadecabacter.arcticus.238 GCA_000155735.2_ASM15573v2
Ruegeria.pomeroyi.DSS.3 GCA_000011965.2_ASM1196v2
Sagittula.stellata.E.37 GCA_000169415.1_ASM16941v1
Ketogulonicigenium.vulgare GCA_001399515.1_ASM139951v1
Rubellimicrobium.rubrum GCA_006152115.1_ASM615211v1
Paracoccus.denitrificans GCA_004063735.1_ASM406373v1
Rhodobacter.sphaeroides.ATCC.17029 GCA_000015985.1_ASM1598v1
Neomegalonema.perideroedes.DSM.15528 GCA_000374145.1_ASM37414v1
Citromicrobium.sp.JL477 GCA_001304795.1_ASM130479v1
Sphingopyxis.macrogoltabida GCA_001314325.1_ASM131432v1
Sphingobium.indicum.B90A GCA_000264945.2_ASM26494v2
Sphingomonas.wittichii.RW1 GCA_000016765.1_ASM1676v1
Zymomonas.mobilis.subsp.mobilis.ATCC.31821 GCA_000007105.1_ASM710v1
alpha.proteobacterium.AAP81b GCA_001295935.1_ASM129593v1
Kordiimonas.gwangyangensis.DSM.19435 GCA_000375545.1_ASM37554v1
Sneathiella.glossodoripedis.JCM.23214 GCA_000616095.1_ASM61609v1
Geminicoccus.roseus.DSM.18922 GCA_000427665.1_ASM42766v1
Tistrella.mobilis.KA081020.065 GCA_000264455.2_ASM26445v2
Candidatus.Puniceispirillum.marinum.IMCC1322 GCA_000024465.1_ASM2446v1
Thalassobaculum.salexigens.DSM.19539 GCA_000423805.1_ASM42380v1
alpha.proteobacterium.BAL199 GCA_000171835.1_ASM17183v1
Oceanibaculum.indicum GCA_003633955.1_ASM363395v1
Kiloniella.laminariae.DSM.19542 GCA_000374005.1_ASM37400v1
Rhodovibrio.salinarum.DSM.9154 GCA_000515255.1_ASM51525v1
Inquilinus.limosus.DSM.16000 GCA_000423185.1_ASM42318v1
Rhodospirillum.centenum GCA_014197805.1_ASM1419780v1
Micavibrio.aeruginosavorus.ARL.13 GCA_000226315.1_ASM22631v1
Caenispirillum.salinarum.AK4 GCA_000315795.1
Pararhodospirillum.photometricum.DSM.122 GCA_000284415.2_ASM28441v2
Rhodospirillum.rubrum.F11 GCA_000225955.1_ASM22595v1
Roseospirillum.parvum GCA_900100455.1
Magnetospirillum.magneticum.AMB.1 GCA_000009985.1_ASM998v1
Terasakiella.pusilla.DSM.6293 GCA_000688235.1_ASM68823v1
Thalassospira.profundimaris.WP0211 GCA_000300275.1_ASM30027v1
Elioraea.tepidiphila.DSM.17972 GCA_000378465.1_ASM37846v1
Belnapia.moabensis.DSM.16746 GCA_000745835.1_ASM74583v1
Rubritepida.flocculans.DSM.14296 GCA_000425365.1_ASM42536v1
Roseomonas.gilardii GCA_001941945.1_ASM194194v1
Acidiphilium.multivorum.AIU301 GCA_000202835.1_ASM20283v1
Acidisphaera.rubrifaciens.HS.AP3 GCA_000964365.1_ASM96436v1
Granulibacter.bethesdensis.CGDNIH4 GCA_000576145.2_ASM57614v2
Commensalibacter.intestini.A911 GCA_000231445.2_ASM23144v1
Gluconacetobacter.diazotrophicus.PA1.5 GCA_000067045.1_ASM6704v1
Acetobacter.aceti.NBRC.14818 GCA_000193495.2_ASM19349v2
Gluconobacter.oxydans.DSM.3504 GCA_000583855.1_ASM58385v1
0.592946

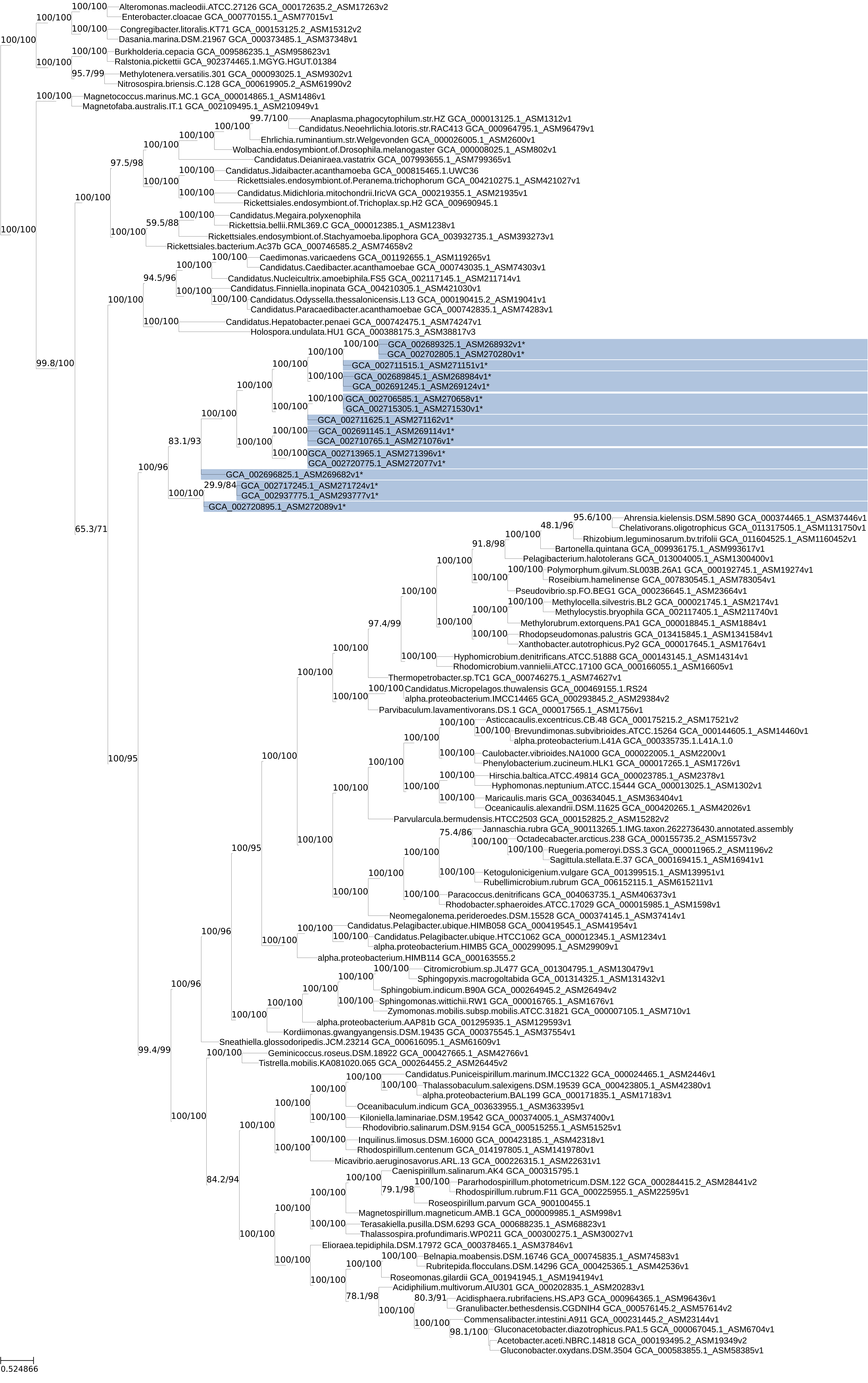

100/100
100/100
100/100
Alteromonas.macleodii.ATCC.27126 GCA_000172635.2_ASM17263v2
Enterobacter.cloacae GCA_000770155.1_ASM77015v1
100/100
Congregibacter.litoralis.KT71 GCA_000153125.2_ASM15312v2
Dasania.marina.DSM.21967 GCA_000373485.1_ASM37348v1
100/100
100/100
Burkholderia.cepacia GCA_009586235.1_ASM958623v1
Ralstonia.pickettii GCA_902374465.1.MGYG.HGUT.01384
95.7/99
Methylotenera.versatilis.301 GCA_000093025.1_ASM9302v1
Nitrosospira.briensis.C.128 GCA_000619905.2_ASM61990v2
100/100
100/100
Magnetococcus.marinus.MC.1 GCA_000014865.1_ASM1486v1
Magnetofaba.australis.IT.1 GCA_002109495.1_ASM210949v1
99.7/100
Anaplasma.phagocytophilum.str.HZ GCA_000013125.1_ASM1312v1
Candidatus.Neoehrlichia.lotoris.str.RAC413 GCA_000964795.1_ASM96479v1
100/100
Ehrlichia.ruminantium.str.Welgevonden GCA_000026005.1_ASM2600v1
100/100
Wolbachia.endosymbiont.of.Drosophila.melanogaster GCA_000008025.1_ASM802v1
100/100
Candidatus.Deianiraea.vastatrix GCA_007993655.1_ASM799365v1
97.5/98
100/100
Candidatus.Jidaibacter.acanthamoeba GCA_000815465.1.UWC36
Rickettsiales.endosymbiont.of.Peranema.trichophorum GCA_004210275.1_ASM421027v1
100/100
100/100
Candidatus.Midichloria.mitochondrii.IricVA GCA_000219355.1_ASM21935v1
Rickettsiales.endosymbiont.of.Trichoplax.sp.H2 GCA_009690945.1
100/100
100/100
Candidatus.Megaira.polyxenophila
59.5/88
Rickettsia.bellii.RML369.C GCA_000012385.1_ASM1238v1
100/100
Rickettsiales.endosymbiont.of.Stachyamoeba.lipophora GCA_003932735.1_ASM393273v1
Rickettsiales.bacterium.Ac37b GCA_000746585.2_ASM74658v2
100/100
Caedimonas.varicaedens GCA_001192655.1_ASM119265v1
100/100
Candidatus.Caedibacter.acanthamoebae GCA_000743035.1_ASM74303v1
Candidatus.Nucleicultrix.amoebiphila.FS5 GCA_002117145.1_ASM211714v1
94.5/96
Candidatus.Finniella.inopinata GCA_004210305.1_ASM421030v1
100/100
100/100
Candidatus.Odyssella.thessalonicensis.L13 GCA_000190415.2_ASM19041v1
100/100
Candidatus.Paracaedibacter.acanthamoebae GCA_000742835.1_ASM74283v1
100/100
Candidatus.Hepatobacter.penaei GCA_000742475.1_ASM74247v1
Holospora.undulata.HU1 GCA_000388175.3_ASM38817v3
99.8/100
100/100
GCA_002689325.1_ASM268932v1*
100/100
GCA_002702805.1_ASM270280v1*
100/100
GCA_002711515.1_ASM271151v1*
100/100
GCA_002689845.1_ASM268984v1*
GCA_002691245.1_ASM269124v1*
100/100
100/100
GCA_002706585.1_ASM270658v1*
100/100
GCA_002715305.1_ASM271530v1*
GCA_002711625.1_ASM271162v1*
100/100
100/100
GCA_002691145.1_ASM269114v1*
GCA_002710765.1_ASM271076v1*
83.1/93
100/100
GCA_002713965.1_ASM271396v1*
GCA_002720775.1_ASM272077v1*
100/96
GCA_002696825.1_ASM269682v1*
29.9/84
GCA_002717245.1_ASM271724v1*
100/100
GCA_002937775.1_ASM293777v1*
GCA_002720895.1_ASM272089v1*
95.6/100
Ahrensia.kielensis.DSM.5890 GCA_000374465.1_ASM37446v1
48.1/96
Chelativorans.oligotrophicus GCA_011317505.1_ASM1131750v1
100/100
Rhizobium.leguminosarum.bv.trifolii GCA_011604525.1_ASM1160452v1
65.3/71
91.8/98
Bartonella.quintana GCA_009936175.1_ASM993617v1
Pelagibacterium.halotolerans GCA_013004005.1_ASM1300400v1
100/100
100/100
Polymorphum.gilvum.SL003B.26A1 GCA_000192745.1_ASM19274v1
100/100
Roseibium.hamelinense GCA_007830545.1_ASM783054v1
Pseudovibrio.sp.FO.BEG1 GCA_000236645.1_ASM23664v1
100/100
100/100
Methylocella.silvestris.BL2 GCA_000021745.1_ASM2174v1
100/100
Methylocystis.bryophila GCA_002117405.1_ASM211740v1
97.4/99
100/100
Methylorubrum.extorquens.PA1 GCA_000018845.1_ASM1884v1
100/100
Rhodopseudomonas.palustris GCA_013415845.1_ASM1341584v1
Xanthobacter.autotrophicus.Py2 GCA_000017645.1_ASM1764v1
100/100
Hyphomicrobium.denitrificans.ATCC.51888 GCA_000143145.1_ASM14314v1
Rhodomicrobium.vannielii.ATCC.17100 GCA_000166055.1_ASM16605v1
100/100
100/100
Thermopetrobacter.sp.TC1 GCA_000746275.1_ASM74627v1
100/100
Candidatus.Micropelagos.thuwalensis GCA_000469155.1.RS24
100/100
alpha.proteobacterium.IMCC14465 GCA_000293845.2_ASM29384v2
Parvibaculum.lavamentivorans.DS.1 GCA_000017565.1_ASM1756v1
100/100
Asticcacaulis.excentricus.CB.48 GCA_000175215.2_ASM17521v2
100/100
100/100
Brevundimonas.subvibrioides.ATCC.15264 GCA_000144605.1_ASM14460v1
alpha.proteobacterium.L41A GCA_000335735.1.L41A.1.0
100/95
100/100
Caulobacter.vibrioides.NA1000 GCA_000022005.1_ASM2200v1
100/100
Phenylobacterium.zucineum.HLK1 GCA_000017265.1_ASM1726v1
100/100
100/100
Hirschia.baltica.ATCC.49814 GCA_000023785.1_ASM2378v1
100/100
Hyphomonas.neptunium.ATCC.15444 GCA_000013025.1_ASM1302v1
100/100
Maricaulis.maris GCA_003634045.1_ASM363404v1
Oceanicaulis.alexandrii.DSM.11625 GCA_000420265.1_ASM42026v1
Parvularcula.bermudensis.HTCC2503 GCA_000152825.2_ASM15282v2
75.4/86
Jannaschia.rubra GCA_900113265.1.IMG.taxon.2622736430.annotated.assembly
100/100
Octadecabacter.arcticus.238 GCA_000155735.2_ASM15573v2
100/95
100/100
100/100
Ruegeria.pomeroyi.DSS.3 GCA_000011965.2_ASM1196v2
Sagittula.stellata.E.37 GCA_000169415.1_ASM16941v1
100/100
100/100
Ketogulonicigenium.vulgare GCA_001399515.1_ASM139951v1
Rubellimicrobium.rubrum GCA_006152115.1_ASM615211v1
100/100
100/100
Paracoccus.denitrificans GCA_004063735.1_ASM406373v1
Rhodobacter.sphaeroides.ATCC.17029 GCA_000015985.1_ASM1598v1
Neomegalonema.perideroedes.DSM.15528 GCA_000374145.1_ASM37414v1
100/96
100/100
Candidatus.Pelagibacter.ubique.HIMB058 GCA_000419545.1_ASM41954v1
100/100
100/100
Candidatus.Pelagibacter.ubique.HTCC1062 GCA_000012345.1_ASM1234v1
alpha.proteobacterium.HIMB5 GCA_000299095.1_ASM29909v1
alpha.proteobacterium.HIMB114 GCA_000163555.2
100/100
Citromicrobium.sp.JL477 GCA_001304795.1_ASM130479v1
100/96
100/100
Sphingopyxis.macrogoltabida GCA_001314325.1_ASM131432v1
100/100
Sphingobium.indicum.B90A GCA_000264945.2_ASM26494v2
100/100
Sphingomonas.wittichii.RW1 GCA_000016765.1_ASM1676v1
100/100
100/100
Zymomonas.mobilis.subsp.mobilis.ATCC.31821 GCA_000007105.1_ASM710v1
alpha.proteobacterium.AAP81b GCA_001295935.1_ASM129593v1
Kordiimonas.gwangyangensis.DSM.19435 GCA_000375545.1_ASM37554v1
Sneathiella.glossodoripedis.JCM.23214 GCA_000616095.1_ASM61609v1
99.4/99
100/100
Geminicoccus.roseus.DSM.18922 GCA_000427665.1_ASM42766v1
Tistrella.mobilis.KA081020.065 GCA_000264455.2_ASM26445v2
100/100
Candidatus.Puniceispirillum.marinum.IMCC1322 GCA_000024465.1_ASM2446v1
100/100
100/100
Thalassobaculum.salexigens.DSM.19539 GCA_000423805.1_ASM42380v1
alpha.proteobacterium.BAL199 GCA_000171835.1_ASM17183v1
100/100
100/100
Oceanibaculum.indicum GCA_003633955.1_ASM363395v1
100/100
Kiloniella.laminariae.DSM.19542 GCA_000374005.1_ASM37400v1
100/100
Rhodovibrio.salinarum.DSM.9154 GCA_000515255.1_ASM51525v1
100/100
100/100
Inquilinus.limosus.DSM.16000 GCA_000423185.1_ASM42318v1
Rhodospirillum.centenum GCA_014197805.1_ASM1419780v1
Micavibrio.aeruginosavorus.ARL.13 GCA_000226315.1_ASM22631v1
100/100
Caenispirillum.salinarum.AK4 GCA_000315795.1
84.2/94
100/100
Pararhodospirillum.photometricum.DSM.122 GCA_000284415.2_ASM28441v2
100/100
79.1/98
Rhodospirillum.rubrum.F11 GCA_000225955.1_ASM22595v1
Roseospirillum.parvum GCA_900100455.1
100/100
Magnetospirillum.magneticum.AMB.1 GCA_000009985.1_ASM998v1
100/100
Terasakiella.pusilla.DSM.6293 GCA_000688235.1_ASM68823v1
100/100
Thalassospira.profundimaris.WP0211 GCA_000300275.1_ASM30027v1
Elioraea.tepidiphila.DSM.17972 GCA_000378465.1_ASM37846v1
100/100
100/100
Belnapia.moabensis.DSM.16746 GCA_000745835.1_ASM74583v1
100/100
Rubritepida.flocculans.DSM.14296 GCA_000425365.1_ASM42536v1
100/100
Roseomonas.gilardii GCA_001941945.1_ASM194194v1
Acidiphilium.multivorum.AIU301 GCA_000202835.1_ASM20283v1
80.3/91
Acidisphaera.rubrifaciens.HS.AP3 GCA_000964365.1_ASM96436v1
78.1/98
100/100
Granulibacter.bethesdensis.CGDNIH4 GCA_000576145.2_ASM57614v2
100/100
Commensalibacter.intestini.A911 GCA_000231445.2_ASM23144v1
98.1/100
Gluconacetobacter.diazotrophicus.PA1.5 GCA_000067045.1_ASM6704v1
Acetobacter.aceti.NBRC.14818 GCA_000193495.2_ASM19349v2
Gluconobacter.oxydans.DSM.3504 GCA_000583855.1_ASM58385v1
0.524866

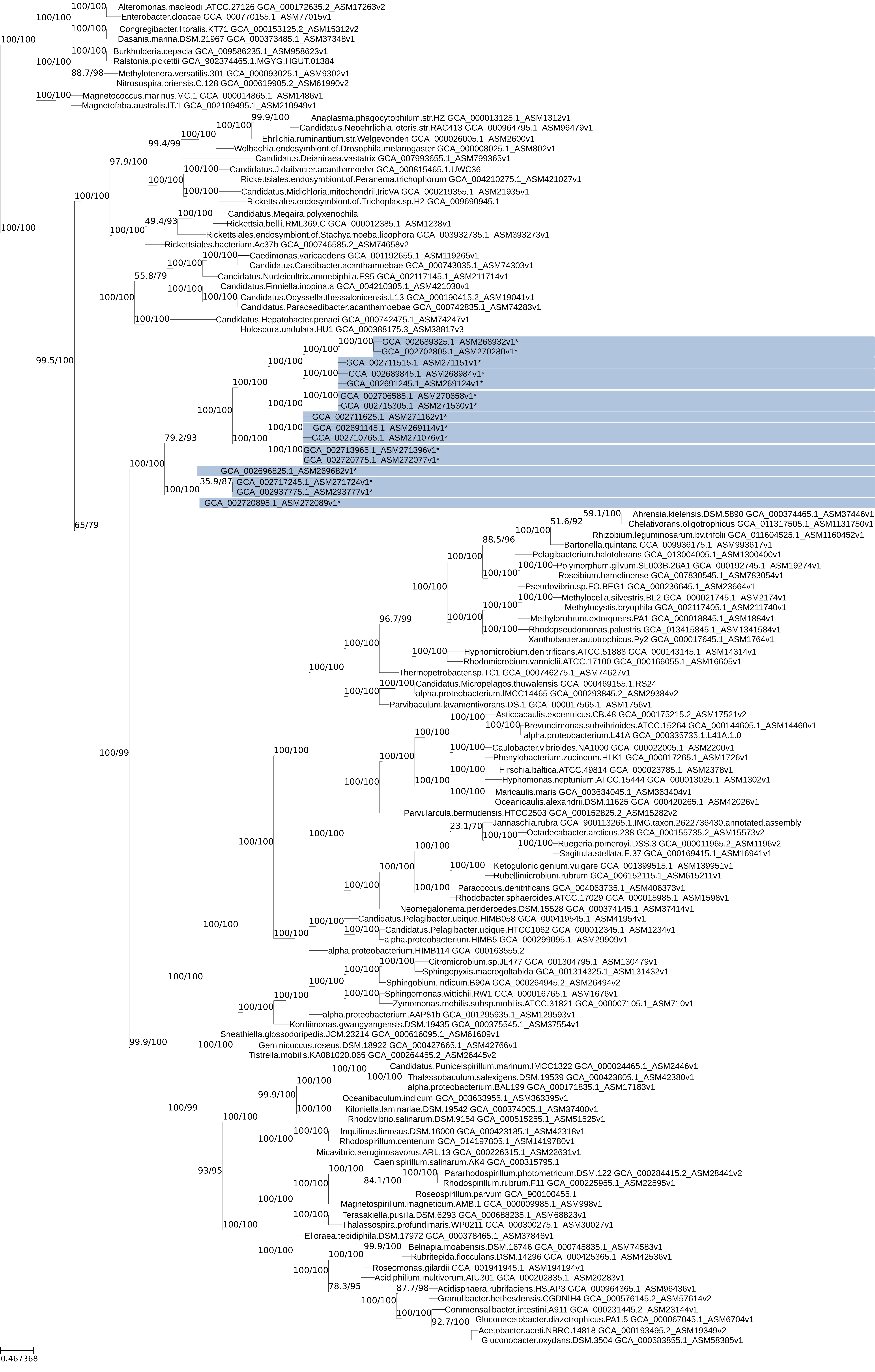

100/100
Alteromonas.macleodii.ATCC.27126 GCA_000172635.2_ASM17263v2
100/100
Enterobacter.cloacae GCA_000770155.1_ASM77015v1
100/100
Congregibacter.litoralis.KT71 GCA_000153125.2_ASM15312v2
100/100
Dasania.marina.DSM.21967 GCA_000373485.1_ASM37348v1
100/100
Burkholderia.cepacia GCA_009586235.1_ASM958623v1
100/100
Ralstonia.pickettii GCA_902374465.1.MGYG.HGUT.01384
88.7/98
Methylotenera.versatilis.301 GCA_000093025.1_ASM9302v1
Nitrosospira.briensis.C.128 GCA_000619905.2_ASM61990v2
100/100
Magnetococcus.marinus.MC.1 GCA_000014865.1_ASM1486v1
Magnetofaba.australis.IT.1 GCA_002109495.1_ASM210949v1
99.9/100
Anaplasma.phagocytophilum.str.HZ GCA_000013125.1_ASM1312v1
100/100
Candidatus.Neoehrlichia.lotoris.str.RAC413 GCA_000964795.1_ASM96479v1
100/100
Ehrlichia.ruminantium.str.Welgevonden GCA_000026005.1_ASM2600v1
99.4/99
Wolbachia.endosymbiont.of.Drosophila.melanogaster GCA_000008025.1_ASM802v1
97.9/100
Candidatus.Deianiraea.vastatrix GCA_007993655.1_ASM799365v1
100/100
Candidatus.Jidaibacter.acanthamoeba GCA_000815465.1.UWC36
100/100
Rickettsiales.endosymbiont.of.Peranema.trichophorum GCA_004210275.1_ASM421027v1
100/100
100/100
Candidatus.Midichloria.mitochondrii.IricVA GCA_000219355.1_ASM21935v1
Rickettsiales.endosymbiont.of.Trichoplax.sp.H2 GCA_009690945.1
100/100
Candidatus.Megaira.polyxenophila
49.4/93
Rickettsia.bellii.RML369.C GCA_000012385.1_ASM1238v1
100/100
Rickettsiales.endosymbiont.of.Stachyamoeba.lipophora GCA_003932735.1_ASM393273v1
Rickettsiales.bacterium.Ac37b GCA_000746585.2_ASM74658v2
100/100
Caedimonas.varicaedens GCA_001192655.1_ASM119265v1
100/100
Candidatus.Caedibacter.acanthamoebae GCA_000743035.1_ASM74303v1
55.8/79
Candidatus.Nucleicultrix.amoebiphila.FS5 GCA_002117145.1_ASM211714v1
100/100
Candidatus.Finniella.inopinata GCA_004210305.1_ASM421030v1
100/100
100/100
Candidatus.Odyssella.thessalonicensis.L13 GCA_000190415.2_ASM19041v1
Candidatus.Paracaedibacter.acanthamoebae GCA_000742835.1_ASM74283v1
100/100
Candidatus.Hepatobacter.penaei GCA_000742475.1_ASM74247v1
Holospora.undulata.HU1 GCA_000388175.3_ASM38817v3
99.5/100
100/100
GCA_002689325.1_ASM268932v1*
100/100
GCA_002702805.1_ASM270280v1*
100/100
GCA_002711515.1_ASM271151v1*
100/100
GCA_002689845.1_ASM268984v1*
100/100
GCA_002691245.1_ASM269124v1*
100/100
GCA_002706585.1_ASM270658v1*
100/100
100/100
GCA_002715305.1_ASM271530v1*
GCA_002711625.1_ASM271162v1*
100/100
GCA_002691145.1_ASM269114v1*
100/100
GCA_002710765.1_ASM271076v1*
79.2/93
100/100
GCA_002713965.1_ASM271396v1*
100/100
GCA_002720775.1_ASM272077v1*
GCA_002696825.1_ASM269682v1*
35.9/87
GCA_002717245.1_ASM271724v1*
100/100
GCA_002937775.1_ASM293777v1*
GCA_002720895.1_ASM272089v1*
59.1/100
Ahrensia.kielensis.DSM.5890 GCA_000374465.1_ASM37446v1
51.6/92
Chelativorans.oligotrophicus GCA_011317505.1_ASM1131750v1
65/79
100/100
Rhizobium.leguminosarum.bv.trifolii GCA_011604525.1_ASM1160452v1
88.5/96
Bartonella.quintana GCA_009936175.1_ASM993617v1
100/100
Pelagibacterium.halotolerans GCA_013004005.1_ASM1300400v1
100/100
Polymorphum.gilvum.SL003B.26A1 GCA_000192745.1_ASM19274v1
100/100
Roseibium.hamelinense GCA_007830545.1_ASM783054v1
100/100
Pseudovibrio.sp.FO.BEG1 GCA_000236645.1_ASM23664v1
100/100
Methylocella.silvestris.BL2 GCA_000021745.1_ASM2174v1
100/100
Methylocystis.bryophila GCA_002117405.1_ASM211740v1
96.7/99
100/100
Methylorubrum.extorquens.PA1 GCA_000018845.1_ASM1884v1
100/100
Rhodopseudomonas.palustris GCA_013415845.1_ASM1341584v1
Xanthobacter.autotrophicus.Py2 GCA_000017645.1_ASM1764v1
100/100
100/100
Hyphomicrobium.denitrificans.ATCC.51888 GCA_000143145.1_ASM14314v1
100/100
Rhodomicrobium.vannielii.ATCC.17100 GCA_000166055.1_ASM16605v1
Thermopetrobacter.sp.TC1 GCA_000746275.1_ASM74627v1
100/100
Candidatus.Micropelagos.thuwalensis GCA_000469155.1.RS24
100/100
alpha.proteobacterium.IMCC14465 GCA_000293845.2_ASM29384v2
Parvibaculum.lavamentivorans.DS.1 GCA_000017565.1_ASM1756v1
100/100
Asticcacaulis.excentricus.CB.48 GCA_000175215.2_ASM17521v2
100/100
Brevundimonas.subvibrioides.ATCC.15264 GCA_000144605.1_ASM14460v1
100/100
alpha.proteobacterium.L41A GCA_000335735.1.L41A.1.0
100/100
Caulobacter.vibrioides.NA1000 GCA_000022005.1_ASM2200v1
100/100
Phenylobacterium.zucineum.HLK1 GCA_000017265.1_ASM1726v1
100/99
100/100
100/100
Hirschia.baltica.ATCC.49814 GCA_000023785.1_ASM2378v1
100/100
Hyphomonas.neptunium.ATCC.15444 GCA_000013025.1_ASM1302v1
100/100
Maricaulis.maris GCA_003634045.1_ASM363404v1
Oceanicaulis.alexandrii.DSM.11625 GCA_000420265.1_ASM42026v1
Parvularcula.bermudensis.HTCC2503 GCA_000152825.2_ASM15282v2
23.1/70
Jannaschia.rubra GCA_900113265.1.IMG.taxon.2622736430.annotated.assembly
100/100
Octadecabacter.arcticus.238 GCA_000155735.2_ASM15573v2
100/100
100/100
Ruegeria.pomeroyi.DSS.3 GCA_000011965.2_ASM1196v2
100/100
Sagittula.stellata.E.37 GCA_000169415.1_ASM16941v1
100/100
100/100
Ketogulonicigenium.vulgare GCA_001399515.1_ASM139951v1
Rubellimicrobium.rubrum GCA_006152115.1_ASM615211v1
100/100
100/100
Paracoccus.denitrificans GCA_004063735.1_ASM406373v1
Rhodobacter.sphaeroides.ATCC.17029 GCA_000015985.1_ASM1598v1
Neomegalonema.perideroedes.DSM.15528 GCA_000374145.1_ASM37414v1
100/100
100/100
Candidatus.Pelagibacter.ubique.HIMB058 GCA_000419545.1_ASM41954v1
100/100
Candidatus.Pelagibacter.ubique.HTCC1062 GCA_000012345.1_ASM1234v1
100/100
alpha.proteobacterium.HIMB5 GCA_000299095.1_ASM29909v1
alpha.proteobacterium.HIMB114 GCA_000163555.2
100/100
Citromicrobium.sp.JL477 GCA_001304795.1_ASM130479v1
100/100
Sphingopyxis.macrogoltabida GCA_001314325.1_ASM131432v1
100/100
Sphingobium.indicum.B90A GCA_000264945.2_ASM26494v2
100/100
Sphingomonas.wittichii.RW1 GCA_000016765.1_ASM1676v1
100/100
Zymomonas.mobilis.subsp.mobilis.ATCC.31821 GCA_000007105.1_ASM710v1
alpha.proteobacterium.AAP81b GCA_001295935.1_ASM129593v1
Kordiimonas.gwangyangensis.DSM.19435 GCA_000375545.1_ASM37554v1
99.9/100
Sneathiella.glossodoripedis.JCM.23214 GCA_000616095.1_ASM61609v1
100/100
Geminicoccus.roseus.DSM.18922 GCA_000427665.1_ASM42766v1
Tistrella.mobilis.KA081020.065 GCA_000264455.2_ASM26445v2
100/100
Candidatus.Puniceispirillum.marinum.IMCC1322 GCA_000024465.1_ASM2446v1
100/100
Thalassobaculum.salexigens.DSM.19539 GCA_000423805.1_ASM42380v1
100/100
alpha.proteobacterium.BAL199 GCA_000171835.1_ASM17183v1
99.9/100
Oceanibaculum.indicum GCA_003633955.1_ASM363395v1
100/99
100/100
Kiloniella.laminariae.DSM.19542 GCA_000374005.1_ASM37400v1
100/100
Rhodovibrio.salinarum.DSM.9154 GCA_000515255.1_ASM51525v1
100/100
Inquilinus.limosus.DSM.16000 GCA_000423185.1_ASM42318v1
100/100
Rhodospirillum.centenum GCA_014197805.1_ASM1419780v1
Micavibrio.aeruginosavorus.ARL.13 GCA_000226315.1_ASM22631v1
100/100
Caenispirillum.salinarum.AK4 GCA_000315795.1
93/95
100/100
Pararhodospirillum.photometricum.DSM.122 GCA_000284415.2_ASM28441v2
84.1/100
Rhodospirillum.rubrum.F11 GCA_000225955.1_ASM22595v1
100/100
Roseospirillum.parvum GCA_900100455.1
100/100
Magnetospirillum.magneticum.AMB.1 GCA_000009985.1_ASM998v1
100/100
Terasakiella.pusilla.DSM.6293 GCA_000688235.1_ASM68823v1
100/100
Thalassospira.profundimaris.WP0211 GCA_000300275.1_ASM30027v1
Elioraea.tepidiphila.DSM.17972 GCA_000378465.1_ASM37846v1
99.9/100
Belnapia.moabensis.DSM.16746 GCA_000745835.1_ASM74583v1
100/100
100/100
Rubritepida.flocculans.DSM.14296 GCA_000425365.1_ASM42536v1
100/100
Roseomonas.gilardii GCA_001941945.1_ASM194194v1
Acidiphilium.multivorum.AIU301 GCA_000202835.1_ASM20283v1
78.3/95
87.7/98
Acidisphaera.rubrifaciens.HS.AP3 GCA_000964365.1_ASM96436v1
100/100
Granulibacter.bethesdensis.CGDNIH4 GCA_000576145.2_ASM57614v2
100/100
Commensalibacter.intestini.A911 GCA_000231445.2_ASM23144v1
92.7/100
Gluconacetobacter.diazotrophicus.PA1.5 GCA_000067045.1_ASM6704v1
Acetobacter.aceti.NBRC.14818 GCA_000193495.2_ASM19349v2
Gluconobacter.oxydans.DSM.3504 GCA_000583855.1_ASM58385v1
0.467368

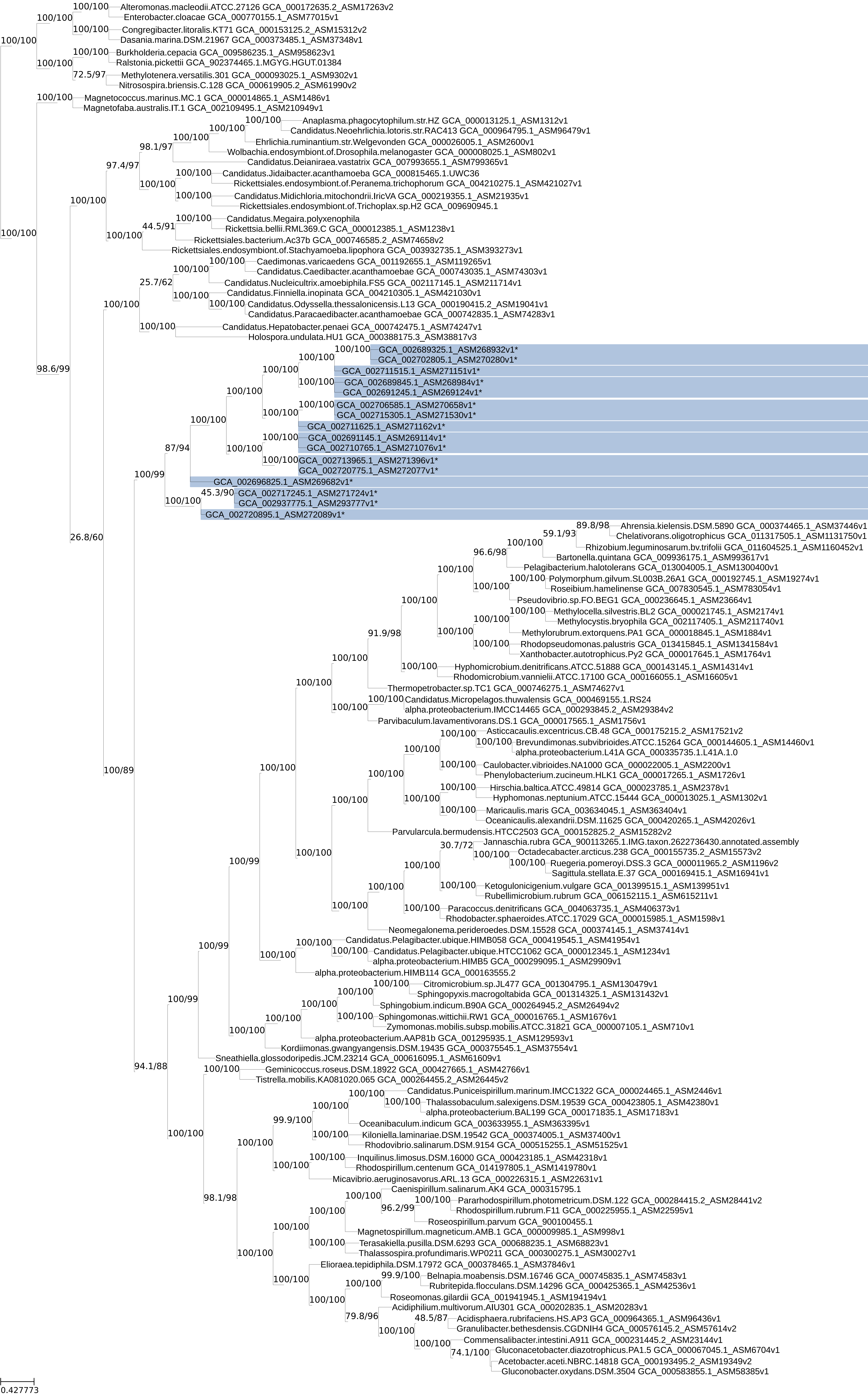
Alteromonas.macleodii.ATCC.27126 GCA_000172635.2_ASM17263v2
Enterobacter.cloacae GCA_000770155.1_ASM77015v1
Congregibacter.litoralis.KT71 GCA_000153125.2_ASM15312v2
Dasania.marina.DSM.21967 GCA_000373485.1_ASM37348v1
Burkholderia.cepacia GCA_009586235.1_ASM958623v1
Ralstonia.pickettii GCA_902374465.1.MGYG.HGUT.01384
Methylotenera.versatilis.301 GCA_000093025.1_ASM9302v1
Nitrosospira.briensis.C.128 GCA_000619905.2_ASM61990v2
Magnetococcus.marinus.MC.1 GCA_000014865.1_ASM1486v1
Magnetofaba.australis.IT.1 GCA_002109495.1_ASM210949v1
Anaplasma.phagocytophilum.str.HZ GCA_000013125.1_ASM1312v1
Candidatus.Neoehrlichia.lotoris.str.RAC413 GCA_000964795.1_ASM96479v1
Ehrlichia.ruminantium.str.Welgevonden GCA_000026005.1_ASM2600v1
Wolbachia.endosymbiont.of.Drosophila.melanogaster GCA_000008025.1_ASM802v1
Candidatus.Deianiraea.vastatrix GCA_007993655.1_ASM799365v1
Candidatus.Jidaibacter.acanthamoeba GCA_000815465.1.UWC36
Rickettsiales.endosymbiont.of.Peranema.trichophorum GCA_004210275.1_ASM421027v1
Candidatus.Midichloria.mitochondrii.IricVA GCA_000219355.1_ASM21935v1
Rickettsiales.endosymbiont.of.Trichoplax.sp.H2 GCA_009690945.1
Candidatus.Megaira.polyxenophila
Rickettsia.bellii.RML369.C GCA_000012385.1_ASM1238v1
Rickettsiales.bacterium.Ac37b GCA_000746585.2_ASM74658v2
Rickettsiales.endosymbiont.of.Stachyamoeba.lipophora GCA_003932735.1_ASM393273v1
Caedimonas.varicaedens GCA_001192655.1_ASM119265v1
Candidatus.Caedibacter.acanthamoebae GCA_000743035.1_ASM74303v1
Candidatus.Nucleicultrix.amoebiphila.FS5 GCA_002117145.1_ASM211714v1
Candidatus.Finniella.inopinata GCA_004210305.1_ASM421030v1
Candidatus.Odyssella.thessalonicensis.L13 GCA_000190415.2_ASM19041v1
Candidatus.Paracaedibacter.acanthamoebae GCA_000742835.1_ASM74283v1
Candidatus.Hepatobacter.penaei GCA_000742475.1_ASM74247v1
Holospora.undulata.HU1 GCA_000388175.3_ASM38817v3
GCA_002689325.1_ASM268932v1*
GCA_002702805.1_ASM270280v1*
GCA_002711515.1_ASM271151v1*
GCA_002689845.1_ASM268984v1*
GCA_002691245.1_ASM269124v1*
GCA_002706585.1_ASM270658v1*
GCA_002715305.1_ASM271530v1*
GCA_002711625.1_ASM271162v1*
GCA_002691145.1_ASM269114v1*
GCA_002710765.1_ASM271076v1*
GCA_002713965.1_ASM271396v1*
GCA_002720775.1_ASM272077v1*
GCA_002696825.1_ASM269682v1*
GCA_002717245.1_ASM271724v1*
GCA_002937775.1_ASM293777v1*
GCA_002720895.1_ASM272089v1*
Ahrensia.kielensis.DSM.5890 GCA_000374465.1_ASM37446v1
Chelativorans.oligotrophicus GCA_011317505.1_ASM1131750v1
Rhizobium.leguminosarum.bv.trifolii GCA_011604525.1_ASM1160452v1
Bartonella.quintana GCA_009936175.1_ASM993617v1
Pelagibacterium.halotolerans GCA_013004005.1_ASM1300400v1
Polymorphum.gilvum.SL003B.26A1 GCA_000192745.1_ASM19274v1
Roseibium.hamelinense GCA_007830545.1_ASM783054v1
Pseudovibrio.sp.FO.BEG1 GCA_000236645.1_ASM23664v1
Methylocella.silvestris.BL2 GCA_000021745.1_ASM2174v1
Methylocystis.bryophila GCA_002117405.1_ASM211740v1
Methylorubrum.extorquens.PA1 GCA_000018845.1_ASM1884v1
Rhodopseudomonas.palustris GCA_013415845.1_ASM1341584v1
Xanthobacter.autotrophicus.Py2 GCA_000017645.1_ASM1764v1
Hyphomicrobium.denitrificans.ATCC.51888 GCA_000143145.1_ASM14314v1
Rhodomicrobium.vannielii.ATCC.17100 GCA_000166055.1_ASM16605v1
Thermopetrobacter.sp.TC1 GCA_000746275.1_ASM74627v1
Candidatus.Micropelagos.thuwalensis GCA_000469155.1.RS24
alpha.proteobacterium.IMCC14465 GCA_000293845.2_ASM29384v2
Parvibaculum.lavamentivorans.DS.1 GCA_000017565.1_ASM1756v1
Asticcacaulis.excentricus.CB.48 GCA_000175215.2_ASM17521v2
Brevundimonas.subvibrioides.ATCC.15264 GCA_000144605.1_ASM14460v1
alpha.proteobacterium.L41A GCA_000335735.1.L41A.1.0
Caulobacter.vibrioides.NA1000 GCA_000022005.1_ASM2200v1
Phenylobacterium.zucineum.HLK1 GCA_000017265.1_ASM1726v1
Hirschia.baltica.ATCC.49814 GCA_000023785.1_ASM2378v1
Hyphomonas.neptunium.ATCC.15444 GCA_000013025.1_ASM1302v1
Maricaulis.maris GCA_003634045.1_ASM363404v1
Oceanicaulis.alexandrii.DSM.11625 GCA_000420265.1_ASM42026v1
Parvularcula.bermudensis.HTCC2503 GCA_000152825.2_ASM15282v2
Jannaschia.rubra GCA_900113265.1.IMG.taxon.2622736430.annotated.assembly
Octadecabacter.arcticus.238 GCA_000155735.2_ASM15573v2
Ruegeria.pomeroyi.DSS.3 GCA_000011965.2_ASM1196v2
Sagittula.stellata.E.37 GCA_000169415.1_ASM16941v1
Ketogulonicigenium.vulgare GCA_001399515.1_ASM139951v1
Rubellimicrobium.rubrum GCA_006152115.1_ASM615211v1
Paracoccus.denitrificans GCA_004063735.1_ASM406373v1
Rhodobacter.sphaeroides.ATCC.17029 GCA_000015985.1_ASM1598v1
Neomegalonema.perideroedes.DSM.15528 GCA_000374145.1_ASM37414v1
Candidatus.Pelagibacter.ubique.HIMB058 GCA_000419545.1_ASM41954v1
Candidatus.Pelagibacter.ubique.HTCC1062 GCA_000012345.1_ASM1234v1
alpha.proteobacterium.HIMB5 GCA_000299095.1_ASM29909v1
alpha.proteobacterium.HIMB114 GCA_000163555.2
Citromicrobium.sp.JL477 GCA_001304795.1_ASM130479v1
Sphingopyxis.macrogoltabida GCA_001314325.1_ASM131432v1
Sphingobium.indicum.B90A GCA_000264945.2_ASM26494v2
Sphingomonas.wittichii.RW1 GCA_000016765.1_ASM1676v1
Zymomonas.mobilis.subsp.mobilis.ATCC.31821 GCA_000007105.1_ASM710v1
alpha.proteobacterium.AAP81b GCA_001295935.1_ASM129593v1
Kordiimonas.gwangyangensis.DSM.19435 GCA_000375545.1_ASM37554v1
Sneathiella.glossodoripedis.JCM.23214 GCA_000616095.1_ASM61609v1
Geminicoccus.roseus.DSM.18922 GCA_000427665.1_ASM42766v1
Tistrella.mobilis.KA081020.065 GCA_000264455.2_ASM26445v2
Candidatus.Puniceispirillum.marinum.IMCC1322 GCA_000024465.1_ASM2446v1
Thalassobaculum.salexigens.DSM.19539 GCA_000423805.1_ASM42380v1
alpha.proteobacterium.BAL199 GCA_000171835.1_ASM17183v1
Oceanibaculum.indicum GCA_003633955.1_ASM363395v1
Kiloniella.laminariae.DSM.19542 GCA_000374005.1_ASM37400v1
Rhodovibrio.salinarum.DSM.9154 GCA_000515255.1_ASM51525v1
Inquilinus.limosus.DSM.16000 GCA_000423185.1_ASM42318v1
Rhodospirillum.centenum GCA_014197805.1_ASM1419780v1
Micavibrio.aeruginosavorus.ARL.13 GCA_000226315.1_ASM22631v1
Caenispirillum.salinarum.AK4 GCA_000315795.1
Pararhodospirillum.photometricum.DSM.122 GCA_000284415.2_ASM28441v2
Rhodospirillum.rubrum.F11 GCA_000225955.1_ASM22595v1
Roseospirillum.parvum GCA_900100455.1
Magnetospirillum.magneticum.AMB.1 GCA_000009985.1_ASM998v1
Terasakiella.pusilla.DSM.6293 GCA_000688235.1_ASM68823v1
Thalassospira.profundimaris.WP0211 GCA_000300275.1_ASM30027v1
Elioraea.tepidiphila.DSM.17972 GCA_000378465.1_ASM37846v1
Belnapia.moabensis.DSM.16746 GCA_000745835.1_ASM74583v1
Rubritepida.flocculans.DSM.14296 GCA_000425365.1_ASM42536v1
Roseomonas.gilardii GCA_001941945.1_ASM194194v1
Acidiphilium.multivorum.AIU301 GCA_000202835.1_ASM20283v1
Acidisphaera.rubrifaciens.HS.AP3 GCA_000964365.1_ASM96436v1
Granulibacter.bethesdensis.CGDNIH4 GCA_000576145.2_ASM57614v2
Commensalibacter.intestini.A911 GCA_000231445.2_ASM23144v1
Gluconacetobacter.diazotrophicus.PA1.5 GCA_000067045.1_ASM6704v1
Acetobacter.aceti.NBRC.14818 GCA_000193495.2_ASM19349v2
Gluconobacter.oxydans.DSM.3504 GCA_000583855.1_ASM58385v1
0.427773

### Group M

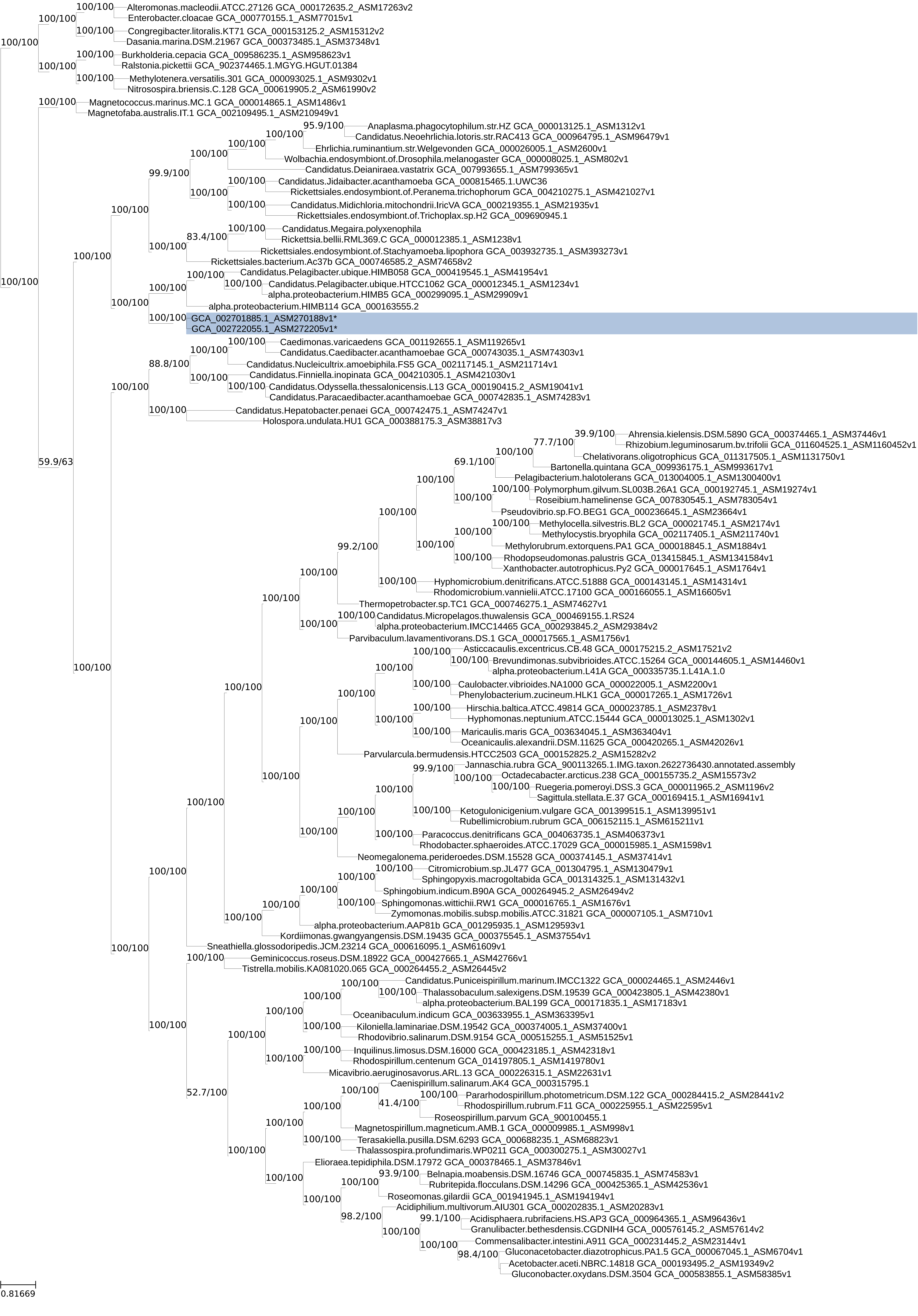
Alteromonas.macleodii.ATCC.27126 GCA_000172635.2_ASM17263v2
Enterobacter.cloacae GCA_000770155.1_ASM77015v1
Congregibacter.litoralis.KT71 GCA_000153125.2_ASM15312v2
Dasania.marina.DSM.21967 GCA_000373485.1_ASM37348v1
Burkholderia.cepacia GCA_009586235.1_ASM958623v1
Ralstonia.pickettii GCA_902374465.1.MGYG.HGUT.01384
Methylotenera.versatilis.301 GCA_000093025.1_ASM9302v1
Nitrosospira.briensis.C.128 GCA_000619905.2_ASM61990v2
Magnetococcus.marinus.MC.1 GCA_000014865.1_ASM1486v1
Magnetofaba.australis.IT.1 GCA_002109495.1_ASM210949v1
Anaplasma.phagocytophilum.str.HZ GCA_000013125.1_ASM1312v1
Candidatus.Neoehrlichia.lotoris.str.RAC413 GCA_000964795.1_ASM96479v1
Ehrlichia.ruminantium.str.Welgevonden GCA_000026005.1_ASM2600v1
Wolbachia.endosymbiont.of.Drosophila.melanogaster GCA_000008025.1_ASM802v1
Candidatus.Deianiraea.vastatrix GCA_007993655.1_ASM799365v1
Candidatus.Jidaibacter.acanthamoeba GCA_000815465.1.UWC36
Rickettsiales.endosymbiont.of.Peranema.trichophorum GCA_004210275.1_ASM421027v1
Candidatus.Midichloria.mitochondrii.IricVA GCA_000219355.1_ASM21935v1
Rickettsiales.endosymbiont.of.Trichoplax.sp.H2 GCA_009690945.1
Candidatus.Megaira.polyxenophila
Rickettsia.bellii.RML369.C GCA_000012385.1_ASM1238v1
Rickettsiales.endosymbiont.of.Stachyamoeba.lipophora GCA_003932735.1_ASM393273v1
Rickettsiales.bacterium.Ac37b GCA_000746585.2_ASM74658v2
Candidatus.Pelagibacter.ubique.HIMB058 GCA_000419545.1_ASM41954v1
Candidatus.Pelagibacter.ubique.HTCC1062 GCA_000012345.1_ASM1234v1
alpha.proteobacterium.HIMB5 GCA_000299095.1_ASM29909v1
alpha.proteobacterium.HIMB114 GCA_000163555.2
GCA_002701885.1_ASM270188v1*
GCA_002722055.1_ASM272205v1*
Caedimonas.varicaedens GCA_001192655.1_ASM119265v1
Candidatus.Caedibacter.acanthamoebae GCA_000743035.1_ASM74303v1
Candidatus.Nucleicultrix.amoebiphila.FS5 GCA_002117145.1_ASM211714v1
Candidatus.Finniella.inopinata GCA_004210305.1_ASM421030v1
Candidatus.Odyssella.thessalonicensis.L13 GCA_000190415.2_ASM19041v1
Candidatus.Paracaedibacter.acanthamoebae GCA_000742835.1_ASM74283v1
Candidatus.Hepatobacter.penaei GCA_000742475.1_ASM74247v1
Holospora.undulata.HU1 GCA_000388175.3_ASM38817v3
Ahrensia.kielensis.DSM.5890 GCA_000374465.1_ASM37446v1
Rhizobium.leguminosarum.bv.trifolii GCA_011604525.1_ASM1160452v1
Chelativorans.oligotrophicus GCA_011317505.1_ASM1131750v1
Bartonella.quintana GCA_009936175.1_ASM993617v1
Pelagibacterium.halotolerans GCA_013004005.1_ASM1300400v1
Polymorphum.gilvum.SL003B.26A1 GCA_000192745.1_ASM19274v1
Roseibium.hamelinense GCA_007830545.1_ASM783054v1
Pseudovibrio.sp.FO.BEG1 GCA_000236645.1_ASM23664v1
Methylocella.silvestris.BL2 GCA_000021745.1_ASM2174v1
Methylocystis.bryophila GCA_002117405.1_ASM211740v1
Methylorubrum.extorquens.PA1 GCA_000018845.1_ASM1884v1
Rhodopseudomonas.palustris GCA_013415845.1_ASM1341584v1
Xanthobacter.autotrophicus.Py2 GCA_000017645.1_ASM1764v1
Hyphomicrobium.denitrificans.ATCC.51888 GCA_000143145.1_ASM14314v1
Rhodomicrobium.vannielii.ATCC.17100 GCA_000166055.1_ASM16605v1
Thermopetrobacter.sp.TC1 GCA_000746275.1_ASM74627v1
Candidatus.Micropelagos.thuwalensis GCA_000469155.1.RS24
alpha.proteobacterium.IMCC14465 GCA_000293845.2_ASM29384v2
Parvibaculum.lavamentivorans.DS.1 GCA_000017565.1_ASM1756v1
Asticcacaulis.excentricus.CB.48 GCA_000175215.2_ASM17521v2
Brevundimonas.subvibrioides.ATCC.15264 GCA_000144605.1_ASM14460v1
alpha.proteobacterium.L41A GCA_000335735.1.L41A.1.0
Caulobacter.vibrioides.NA1000 GCA_000022005.1_ASM2200v1
Phenylobacterium.zucineum.HLK1 GCA_000017265.1_ASM1726v1
Hirschia.baltica.ATCC.49814 GCA_000023785.1_ASM2378v1
Hyphomonas.neptunium.ATCC.15444 GCA_000013025.1_ASM1302v1
Maricaulis.maris GCA_003634045.1_ASM363404v1
Oceanicaulis.alexandrii.DSM.11625 GCA_000420265.1_ASM42026v1
Parvularcula.bermudensis.HTCC2503 GCA_000152825.2_ASM15282v2
Jannaschia.rubra GCA_900113265.1.IMG.taxon.2622736430.annotated.assembly
Octadecabacter.arcticus.238 GCA_000155735.2_ASM15573v2
Ruegeria.pomeroyi.DSS.3 GCA_000011965.2_ASM1196v2
Sagittula.stellata.E.37 GCA_000169415.1_ASM16941v1
Ketogulonicigenium.vulgare GCA_001399515.1_ASM139951v1
Rubellimicrobium.rubrum GCA_006152115.1_ASM615211v1
Paracoccus.denitrificans GCA_004063735.1_ASM406373v1
Rhodobacter.sphaeroides.ATCC.17029 GCA_000015985.1_ASM1598v1
Neomegalonema.perideroedes.DSM.15528 GCA_000374145.1_ASM37414v1
Citromicrobium.sp.JL477 GCA_001304795.1_ASM130479v1
Sphingopyxis.macrogoltabida GCA_001314325.1_ASM131432v1
Sphingobium.indicum.B90A GCA_000264945.2_ASM26494v2
Sphingomonas.wittichii.RW1 GCA_000016765.1_ASM1676v1
Zymomonas.mobilis.subsp.mobilis.ATCC.31821 GCA_000007105.1_ASM710v1
alpha.proteobacterium.AAP81b GCA_001295935.1_ASM129593v1
Kordiimonas.gwangyangensis.DSM.19435 GCA_000375545.1_ASM37554v1
Sneathiella.glossodoripedis.JCM.23214 GCA_000616095.1_ASM61609v1
Geminicoccus.roseus.DSM.18922 GCA_000427665.1_ASM42766v1
Tistrella.mobilis.KA081020.065 GCA_000264455.2_ASM26445v2
Candidatus.Puniceispirillum.marinum.IMCC1322 GCA_000024465.1_ASM2446v1
Thalassobaculum.salexigens.DSM.19539 GCA_000423805.1_ASM42380v1
alpha.proteobacterium.BAL199 GCA_000171835.1_ASM17183v1
Oceanibaculum.indicum GCA_003633955.1_ASM363395v1
Kiloniella.laminariae.DSM.19542 GCA_000374005.1_ASM37400v1
Rhodovibrio.salinarum.DSM.9154 GCA_000515255.1_ASM51525v1
Inquilinus.limosus.DSM.16000 GCA_000423185.1_ASM42318v1
Rhodospirillum.centenum GCA_014197805.1_ASM1419780v1
Micavibrio.aeruginosavorus.ARL.13 GCA_000226315.1_ASM22631v1
Caenispirillum.salinarum.AK4 GCA_000315795.1
Pararhodospirillum.photometricum.DSM.122 GCA_000284415.2_ASM28441v2
Rhodospirillum.rubrum.F11 GCA_000225955.1_ASM22595v1
Roseospirillum.parvum GCA_900100455.1
Magnetospirillum.magneticum.AMB.1 GCA_000009985.1_ASM998v1
Terasakiella.pusilla.DSM.6293 GCA_000688235.1_ASM68823v1
Thalassospira.profundimaris.WP0211 GCA_000300275.1_ASM30027v1
Elioraea.tepidiphila.DSM.17972 GCA_000378465.1_ASM37846v1
Belnapia.moabensis.DSM.16746 GCA_000745835.1_ASM74583v1
Rubritepida.flocculans.DSM.14296 GCA_000425365.1_ASM42536v1
Roseomonas.gilardii GCA_001941945.1_ASM194194v1
Acidiphilium.multivorum.AIU301 GCA_000202835.1_ASM20283v1
Acidisphaera.rubrifaciens.HS.AP3 GCA_000964365.1_ASM96436v1
Granulibacter.bethesdensis.CGDNIH4 GCA_000576145.2_ASM57614v2
Commensalibacter.intestini.A911 GCA_000231445.2_ASM23144v1
Gluconacetobacter.diazotrophicus.PA1.5 GCA_000067045.1_ASM6704v1
Acetobacter.aceti.NBRC.14818 GCA_000193495.2_ASM19349v2
Gluconobacter.oxydans.DSM.3504 GCA_000583855.1_ASM58385v1
0.81669

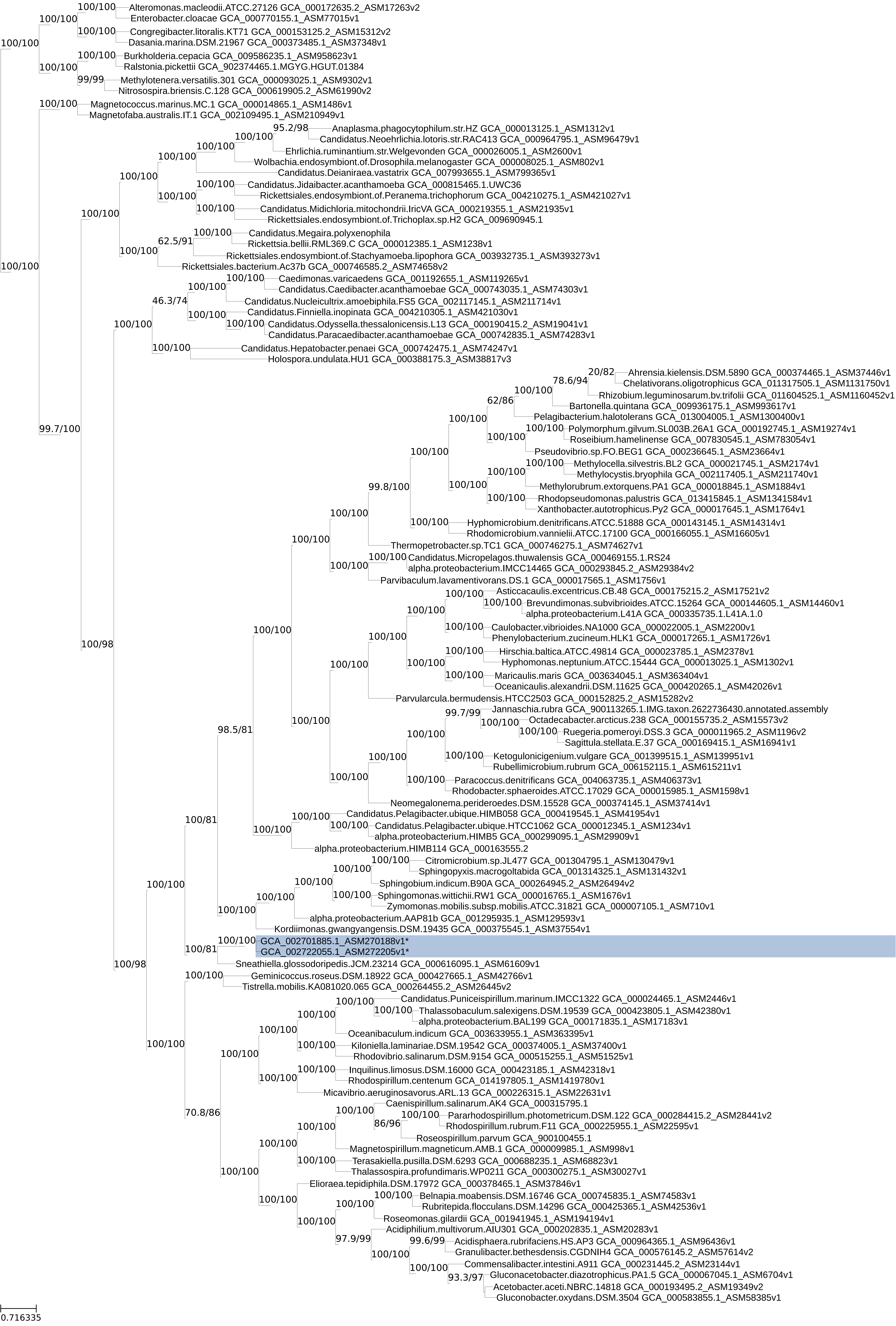
Alteromonas.macleodii.ATCC.27126 GCA_000172635.2_ASM17263v2
Enterobacter.cloacae GCA_000770155.1_ASM77015v1
Congregibacter.litoralis.KT71 GCA_000153125.2_ASM15312v2
Dasania.marina.DSM.21967 GCA_000373485.1_ASM37348v1
Burkholderia.cepacia GCA_009586235.1_ASM958623v1
Ralstonia.pickettii GCA_902374465.1.MGYG.HGUT.01384
Methylotenera.versatilis.301 GCA_000093025.1_ASM9302v1
Nitrosospira.briensis.C.128 GCA_000619905.2_ASM61990v2
Magnetococcus.marinus.MC.1 GCA_000014865.1_ASM1486v1
Magnetofaba.australis.IT.1 GCA_002109495.1_ASM210949v1
Anaplasma.phagocytophilum.str.HZ GCA_000013125.1_ASM1312v1
Candidatus.Neoehrlichia.lotoris.str.RAC413 GCA_000964795.1_ASM96479v1
Ehrlichia.ruminantium.str.Welgevonden GCA_000026005.1_ASM2600v1
Wolbachia.endosymbiont.of.Drosophila.melanogaster GCA_000008025.1_ASM802v1
Candidatus.Deianiraea.vastatrix GCA_007993655.1_ASM799365v1
Candidatus.Jidaibacter.acanthamoeba GCA_000815465.1.UWC36
Rickettsiales.endosymbiont.of.Peranema.trichophorum GCA_004210275.1_ASM421027v1
Candidatus.Midichloria.mitochondrii.IricVA GCA_000219355.1_ASM21935v1
Rickettsiales.endosymbiont.of.Trichoplax.sp.H2 GCA_009690945.1
Candidatus.Megaira.polyxenophila
Rickettsia.bellii.RML369.C GCA_000012385.1_ASM1238v1
Rickettsiales.endosymbiont.of.Stachyamoeba.lipophora GCA_003932735.1_ASM393273v1
Rickettsiales.bacterium.Ac37b GCA_000746585.2_ASM74658v2
Caedimonas.varicaedens GCA_001192655.1_ASM119265v1
Candidatus.Caedibacter.acanthamoebae GCA_000743035.1_ASM74303v1
Candidatus.Nucleicultrix.amoebiphila.FS5 GCA_002117145.1_ASM211714v1
Candidatus.Finniella.inopinata GCA_004210305.1_ASM421030v1
Candidatus.Odyssella.thessalonicensis.L13 GCA_000190415.2_ASM19041v1
Candidatus.Paracaedibacter.acanthamoebae GCA_000742835.1_ASM74283v1
Candidatus.Hepatobacter.penaei GCA_000742475.1_ASM74247v1
Holospora.undulata.HU1 GCA_000388175.3_ASM38817v3
Ahrensia.kielensis.DSM.5890 GCA_000374465.1_ASM37446v1
Chelativorans.oligotrophicus GCA_011317505.1_ASM1131750v1
Rhizobium.leguminosarum.bv.trifolii GCA_011604525.1_ASM1160452v1
Bartonella.quintana GCA_009936175.1_ASM993617v1
Pelagibacterium.halotolerans GCA_013004005.1_ASM1300400v1
Polymorphum.gilvum.SL003B.26A1 GCA_000192745.1_ASM19274v1
Roseibium.hamelinense GCA_007830545.1_ASM783054v1
Pseudovibrio.sp.FO.BEG1 GCA_000236645.1_ASM23664v1
Methylocella.silvestris.BL2 GCA_000021745.1_ASM2174v1
Methylocystis.bryophila GCA_002117405.1_ASM211740v1
Methylorubrum.extorquens.PA1 GCA_000018845.1_ASM1884v1
Rhodopseudomonas.palustris GCA_013415845.1_ASM1341584v1
Xanthobacter.autotrophicus.Py2 GCA_000017645.1_ASM1764v1
Hyphomicrobium.denitrificans.ATCC.51888 GCA_000143145.1_ASM14314v1
Rhodomicrobium.vannielii.ATCC.17100 GCA_000166055.1_ASM16605v1
Thermopetrobacter.sp.TC1 GCA_000746275.1_ASM74627v1
Candidatus.Micropelagos.thuwalensis GCA_000469155.1.RS24
alpha.proteobacterium.IMCC14465 GCA_000293845.2_ASM29384v2
Parvibaculum.lavamentivorans.DS.1 GCA_000017565.1_ASM1756v1
Asticcacaulis.excentricus.CB.48 GCA_000175215.2_ASM17521v2
Brevundimonas.subvibrioides.ATCC.15264 GCA_000144605.1_ASM14460v1
alpha.proteobacterium.L41A GCA_000335735.1.L41A.1.0
Caulobacter.vibrioides.NA1000 GCA_000022005.1_ASM2200v1
Phenylobacterium.zucineum.HLK1 GCA_000017265.1_ASM1726v1
Hirschia.baltica.ATCC.49814 GCA_000023785.1_ASM2378v1
Hyphomonas.neptunium.ATCC.15444 GCA_000013025.1_ASM1302v1
Maricaulis.maris GCA_003634045.1_ASM363404v1
Oceanicaulis.alexandrii.DSM.11625 GCA_000420265.1_ASM42026v1
Parvularcula.bermudensis.HTCC2503 GCA_000152825.2_ASM15282v2
Jannaschia.rubra GCA_900113265.1.IMG.taxon.2622736430.annotated.assembly
Octadecabacter.arcticus.238 GCA_000155735.2_ASM15573v2
Ruegeria.pomeroyi.DSS.3 GCA_000011965.2_ASM1196v2
Sagittula.stellata.E.37 GCA_000169415.1_ASM16941v1
Ketogulonicigenium.vulgare GCA_001399515.1_ASM139951v1
Rubellimicrobium.rubrum GCA_006152115.1_ASM615211v1
Paracoccus.denitrificans GCA_004063735.1_ASM406373v1
Rhodobacter.sphaeroides.ATCC.17029 GCA_000015985.1_ASM1598v1
Neomegalonema.perideroedes.DSM.15528 GCA_000374145.1_ASM37414v1
Candidatus.Pelagibacter.ubique.HIMB058 GCA_000419545.1_ASM41954v1
Candidatus.Pelagibacter.ubique.HTCC1062 GCA_000012345.1_ASM1234v1
alpha.proteobacterium.HIMB5 GCA_000299095.1_ASM29909v1
alpha.proteobacterium.HIMB114 GCA_000163555.2
Citromicrobium.sp.JL477 GCA_001304795.1_ASM130479v1
Sphingopyxis.macrogoltabida GCA_001314325.1_ASM131432v1
Sphingobium.indicum.B90A GCA_000264945.2_ASM26494v2
Sphingomonas.wittichii.RW1 GCA_000016765.1_ASM1676v1
Zymomonas.mobilis.subsp.mobilis.ATCC.31821 GCA_000007105.1_ASM710v1
alpha.proteobacterium.AAP81b GCA_001295935.1_ASM129593v1
Kordiimonas.gwangyangensis.DSM.19435 GCA_000375545.1_ASM37554v1
GCA_002701885.1_ASM270188v1*
GCA_002722055.1_ASM272205v1*
Sneathiella.glossodoripedis.JCM.23214 GCA_000616095.1_ASM61609v1
Geminicoccus.roseus.DSM.18922 GCA_000427665.1_ASM42766v1
Tistrella.mobilis.KA081020.065 GCA_000264455.2_ASM26445v2
Candidatus.Puniceispirillum.marinum.IMCC1322 GCA_000024465.1_ASM2446v1
Thalassobaculum.salexigens.DSM.19539 GCA_000423805.1_ASM42380v1
alpha.proteobacterium.BAL199 GCA_000171835.1_ASM17183v1
Oceanibaculum.indicum GCA_003633955.1_ASM363395v1
Kiloniella.laminariae.DSM.19542 GCA_000374005.1_ASM37400v1
Rhodovibrio.salinarum.DSM.9154 GCA_000515255.1_ASM51525v1
Inquilinus.limosus.DSM.16000 GCA_000423185.1_ASM42318v1
Rhodospirillum.centenum GCA_014197805.1_ASM1419780v1
Micavibrio.aeruginosavorus.ARL.13 GCA_000226315.1_ASM22631v1
Caenispirillum.salinarum.AK4 GCA_000315795.1
Pararhodospirillum.photometricum.DSM.122 GCA_000284415.2_ASM28441v2
Rhodospirillum.rubrum.F11 GCA_000225955.1_ASM22595v1
Roseospirillum.parvum GCA_900100455.1
Magnetospirillum.magneticum.AMB.1 GCA_000009985.1_ASM998v1
Terasakiella.pusilla.DSM.6293 GCA_000688235.1_ASM68823v1
Thalassospira.profundimaris.WP0211 GCA_000300275.1_ASM30027v1
Elioraea.tepidiphila.DSM.17972 GCA_000378465.1_ASM37846v1
Belnapia.moabensis.DSM.16746 GCA_000745835.1_ASM74583v1
Rubritepida.flocculans.DSM.14296 GCA_000425365.1_ASM42536v1
Roseomonas.gilardii GCA_001941945.1_ASM194194v1
Acidiphilium.multivorum.AIU301 GCA_000202835.1_ASM20283v1
Acidisphaera.rubrifaciens.HS.AP3 GCA_000964365.1_ASM96436v1
Granulibacter.bethesdensis.CGDNIH4 GCA_000576145.2_ASM57614v2
Commensalibacter.intestini.A911 GCA_000231445.2_ASM23144v1
Gluconacetobacter.diazotrophicus.PA1.5 GCA_000067045.1_ASM6704v1
Acetobacter.aceti.NBRC.14818 GCA_000193495.2_ASM19349v2
Gluconobacter.oxydans.DSM.3504 GCA_000583855.1_ASM58385v1
0.716335

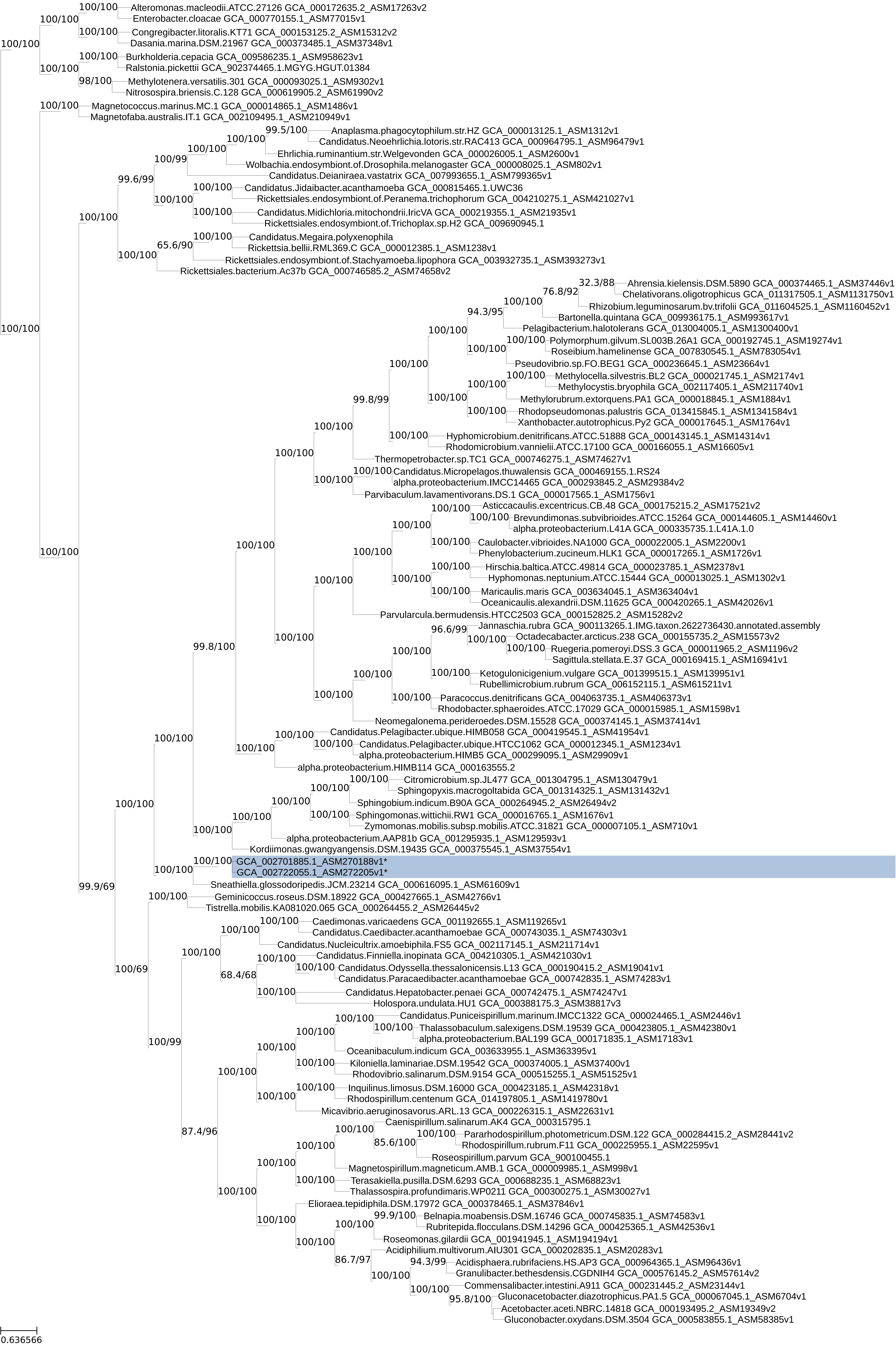

Alteromonas.macleodii.ATCC.27126 GCA_000172635.2_ASM17263v2
Enterobacter.cloacae GCA_000770155.1_ASM77015v1
Congregibacter.litoralis.KT71 GCA_000153125.2_ASM15312v2
Dasania.marina.DSM.21967 GCA_000373485.1_ASM37348v1
Burkholderia.cepacia GCA_009586235.1_ASM958623v1
Ralstonia.pickettii GCA_902374465.1.MGYG.HGUT.01384
Methylotenera.versatilis.301 GCA_000093025.1_ASM9302v1
Nitrosospira.briensis.C.128 GCA_000619905.2_ASM61990v2
Magnetococcus.marinus.MC.1 GCA_000014865.1_ASM1486v1
Magnetofaba.australis.IT.1 GCA_002109495.1_ASM210949v1
Anaplasma.phagocytophilum.str.HZ GCA_000013125.1_ASM1312v1
Candidatus.Neoehrlichia.lotoris.str.RAC413 GCA_000964795.1_ASM96479v1
Ehrlichia.ruminantium.str.Welgevonden GCA_000026005.1_ASM2600v1
Wolbachia.endosymbiont.of.Drosophila.melanogaster GCA_000008025.1_ASM802v1
Candidatus.Deianiraea.vastatrix GCA_007993655.1_ASM799365v1
Candidatus.Jidaibacter.acanthamoeba GCA_000815465.1.UWC36
Rickettsiales.endosymbiont.of.Peranema.trichophorum GCA_004210275.1_ASM421027v1
Candidatus.Midichloria.mitochondrii.IricVA GCA_000219355.1_ASM21935v1
Rickettsiales.endosymbiont.of.Trichoplax.sp.H2 GCA_009690945.1
Candidatus.Megaira.polyxenophila
Rickettsia.bellii.RML369.C GCA_000012385.1_ASM1238v1
Rickettsiales.endosymbiont.of.Stachyamoeba.lipophora GCA_003932735.1_ASM393273v1
Rickettsiales.bacterium.Ac37b GCA_000746585.2_ASM74658v2
Ahrensia.kielensis.DSM.5890 GCA_000374465.1_ASM37446v1
Chelativorans.oligotrophicus GCA_011317505.1_ASM1131750v1
Rhizobium.leguminosarum.bv.trifolii GCA_011604525.1_ASM1160452v1
Bartonella.quintana GCA_009936175.1_ASM993617v1
Pelagibacterium.halotolerans GCA_013004005.1_ASM1300400v1
Polymorphum.gilvum.SL003B.26A1 GCA_000192745.1_ASM19274v1
Roseibium.hamelinense GCA_007830545.1_ASM783054v1
Pseudovibrio.sp.FO.BEG1 GCA_000236645.1_ASM23664v1
Methylocella.silvestris.BL2 GCA_000021745.1_ASM2174v1
Methylocystis.bryophila GCA_002117405.1_ASM211740v1
Methylorubrum.extorquens.PA1 GCA_000018845.1_ASM1884v1
Rhodopseudomonas.palustris GCA_013415845.1_ASM1341584v1
Xanthobacter.autotrophicus.Py2 GCA_000017645.1_ASM1764v1
Hyphomicrobium.denitrificans.ATCC.51888 GCA_000143145.1_ASM14314v1
Rhodomicrobium.vannielii.ATCC.17100 GCA_000166055.1_ASM16605v1
Thermopetrobacter.sp.TC1 GCA_000746275.1_ASM74627v1
Candidatus.Micropelagos.thuwalensis GCA_000469155.1.RS24
alpha.proteobacterium.IMCC14465 GCA_000293845.2_ASM29384v2
Parvibaculum.lavamentivorans.DS.1 GCA_000017565.1_ASM1756v1
Asticcacaulis.excentricus.CB.48 GCA_000175215.2_ASM17521v2
Brevundimonas.subvibrioides.ATCC.15264 GCA_000144605.1_ASM14460v1
alpha.proteobacterium.L41A GCA_000335735.1.L41A.1.0
Caulobacter.vibrioides.NA1000 GCA_000022005.1_ASM2200v1
Phenylobacterium.zucineum.HLK1 GCA_000017265.1_ASM1726v1
Hirschia.baltica.ATCC.49814 GCA_000023785.1_ASM2378v1
Hyphomonas.neptunium.ATCC.15444 GCA_000013025.1_ASM1302v1
Maricaulis.maris GCA_003634045.1_ASM363404v1
Oceanicaulis.alexandrii.DSM.11625 GCA_000420265.1_ASM42026v1
Parvularcula.bermudensis.HTCC2503 GCA_000152825.2_ASM15282v2
Jannaschia.rubra GCA_900113265.1.IMG.taxon.2622736430.annotated.assembly
Octadecabacter.arcticus.238 GCA_000155735.2_ASM15573v2
Ruegeria.pomeroyi.DSS.3 GCA_000011965.2_ASM1196v2
Sagittula.stellata.E.37 GCA_000169415.1_ASM16941v1
Ketogulonicigenium.vulgare GCA_001399515.1_ASM139951v1
Rubellimicrobium.rubrum GCA_006152115.1_ASM615211v1
Paracoccus.denitrificans GCA_004063735.1_ASM406373v1
Rhodobacter.sphaeroides.ATCC.17029 GCA_000015985.1_ASM1598v1
Neomegalonema.perideroedes.DSM.15528 GCA_000374145.1_ASM37414v1
Candidatus.Pelagibacter.ubique.HIMB058 GCA_000419545.1_ASM41954v1
Candidatus.Pelagibacter.ubique.HTCC1062 GCA_000012345.1_ASM1234v1
alpha.proteobacterium.HIMB5 GCA_000299095.1_ASM29909v1
alpha.proteobacterium.HIMB114 GCA_000163555.2
Citromicrobium.sp.JL477 GCA_001304795.1_ASM130479v1
Sphingopyxis.macrogoltabida GCA_001314325.1_ASM131432v1
Sphingobium.indicum.B90A GCA_000264945.2_ASM26494v2
Sphingomonas.wittichii.RW1 GCA_000016765.1_ASM1676v1
Zymomonas.mobilis.subsp.mobilis.ATCC.31821 GCA_000007105.1_ASM710v1
alpha.proteobacterium.AAP81b GCA_001295935.1_ASM129593v1
Kordiimonas.gwangyangensis.DSM.19435 GCA_000375545.1_ASM37554v1
GCA_002701885.1_ASM270188v1*
GCA_002722055.1_ASM272205v1*
Sneathiella.glossodoripedis.JCM.23214 GCA_000616095.1_ASM61609v1
Geminicoccus.roseus.DSM.18922 GCA_000427665.1_ASM42766v1
Tistrella.mobilis.KA081020.065 GCA_000264455.2_ASM26445v2
Caedimonas.varicaedens GCA_001192655.1_ASM119265v1
Candidatus.Caedibacter.acanthamoebae GCA_000743035.1_ASM74303v1
Candidatus.Nucleicultrix.amoebiphila.FS5 GCA_002117145.1_ASM211714v1
Candidatus.Finniella.inopinata GCA_004210305.1_ASM421030v1
Candidatus.Odyssella.thessalonicensis.L13 GCA_000190415.2_ASM19041v1
Candidatus.Paracaedibacter.acanthamoebae GCA_000742835.1_ASM74283v1
Candidatus.Hepatobacter.penaei GCA_000742475.1_ASM74247v1
Holospora.undulata.HU1 GCA_000388175.3_ASM38817v3
Candidatus.Puniceispirillum.marinum.IMCC1322 GCA_000024465.1_ASM2446v1
Thalassobaculum.salexigens.DSM.19539 GCA_000423805.1_ASM42380v1
alpha.proteobacterium.BAL199 GCA_000171835.1_ASM17183v1
Oceanibaculum.indicum GCA_003633955.1_ASM363395v1
Kiloniella.laminariae.DSM.19542 GCA_000374005.1_ASM37400v1
Rhodovibrio.salinarum.DSM.9154 GCA_000515255.1_ASM51525v1
Inquilinus.limosus.DSM.16000 GCA_000423185.1_ASM42318v1
Rhodospirillum.centenum GCA_014197805.1_ASM1419780v1
Micavibrio.aeruginosavorus.ARL.13 GCA_000226315.1_ASM22631v1
Caenispirillum.salinarum.AK4 GCA_000315795.1
Pararhodospirillum.photometricum.DSM.122 GCA_000284415.2_ASM28441v2
Rhodospirillum.rubrum.F11 GCA_000225955.1_ASM22595v1
Roseospirillum.parvum GCA_900100455.1
Magnetospirillum.magneticum.AMB.1 GCA_000009985.1_ASM998v1
Terasakiella.pusilla.DSM.6293 GCA_000688235.1_ASM68823v1
Thalassospira.profundimaris.WP0211 GCA_000300275.1_ASM30027v1
Elioraea.tepidiphila.DSM.17972 GCA_000378465.1_ASM37846v1
Belnapia.moabensis.DSM.16746 GCA_000745835.1_ASM74583v1
Rubritepida.flocculans.DSM.14296 GCA_000425365.1_ASM42536v1
Roseomonas.gilardii GCA_001941945.1_ASM194194v1
Acidiphilium.multivorum.AIU301 GCA_000202835.1_ASM20283v1
Acidisphaera.rubrifaciens.HS.AP3 GCA_000964365.1_ASM96436v1
Granulibacter.bethesdensis.CGDNIH4 GCA_000576145.2_ASM57614v2
Commensalibacter.intestini.A911 GCA_000231445.2_ASM23144v1
Gluconacetobacter.diazotrophicus.PA1.5 GCA_000067045.1_ASM6704v1
Acetobacter.aceti.NBRC.14818 GCA_000193495.2_ASM19349v2
Gluconobacter.oxydans.DSM.3504 GCA_000583855.1_ASM58385v1
0.636566

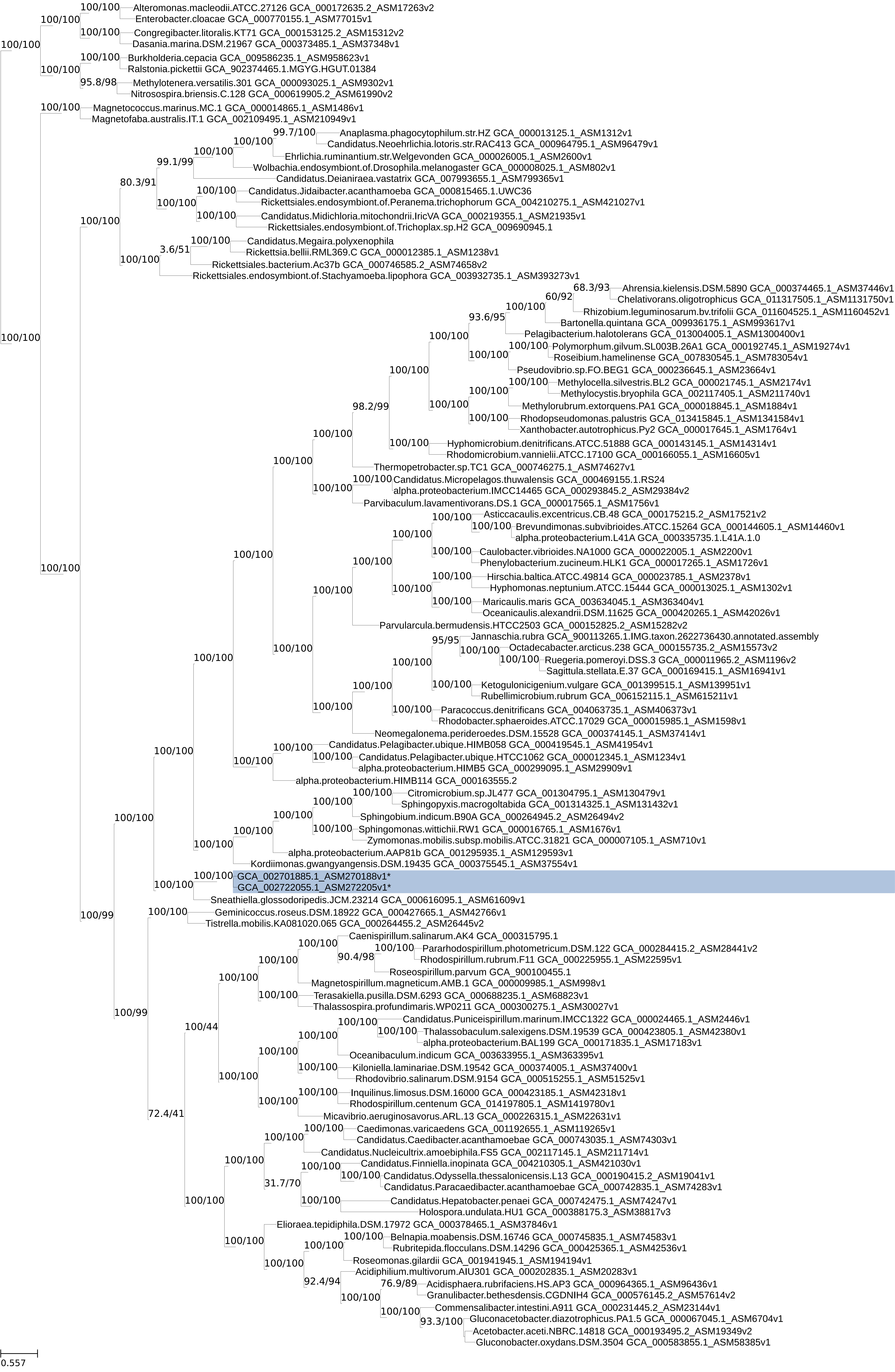
100/100
Alteromonas.macleodii.ATCC.27126 GCA_000172635.2_ASM17263v2
100/100
Enterobacter.cloacae GCA_000770155.1_ASM77015v1
100/100
Congregibacter.litoralis.KT71 GCA_000153125.2_ASM15312v2
100/100
Dasania.marina.DSM.21967 GCA_000373485.1_ASM37348v1
100/100
Burkholderia.cepacia GCA_009586235.1_ASM958623v1
100/100
Ralstonia.pickettii GCA_902374465.1.MGYG.HGUT.01384
95.8/98
Methylotenera.versatilis.301 GCA_000093025.1_ASM9302v1
Nitrosospira.briensis.C.128 GCA_000619905.2_ASM61990v2
100/100
Magnetococcus.marinus.MC.1 GCA_000014865.1_ASM1486v1
Magnetofaba.australis.IT.1 GCA_002109495.1_ASM210949v1
99.7/100
Anaplasma.phagocytophilum.str.HZ GCA_000013125.1_ASM1312v1
100/100
Candidatus.Neoehrlichia.lotoris.str.RAC413 GCA_000964795.1_ASM96479v1
100/100
Ehrlichia.ruminantium.str.Welgevonden GCA_000026005.1_ASM2600v1
99.1/99
Wolbachia.endosymbiont.of.Drosophila.melanogaster GCA_000008025.1_ASM802v1
Candidatus.Deianiraea.vastatrix GCA_007993655.1_ASM799365v1
80.3/91
100/100
Candidatus.Jidaibacter.acanthamoeba GCA_000815465.1.UWC36
100/100
Rickettsiales.endosymbiont.of.Peranema.trichophorum GCA_004210275.1_ASM421027v1
100/100
Candidatus.Midichloria.mitochondrii.IricVA GCA_000219355.1_ASM21935v1
100/100
Rickettsiales.endosymbiont.of.Trichoplax.sp.H2 GCA_009690945.1
100/100
Candidatus.Megaira.polyxenophila
3.6/51
Rickettsia.bellii.RML369.C GCA_000012385.1_ASM1238v1
100/100
Rickettsiales.bacterium.Ac37b GCA_000746585.2_ASM74658v2
Rickettsiales.endosymbiont.of.Stachyamoeba.lipophora GCA_003932735.1_ASM393273v1
68.3/93
Ahrensia.kielensis.DSM.5890 GCA_000374465.1_ASM37446v1
60/92
Chelativorans.oligotrophicus GCA_011317505.1_ASM1131750v1
100/100
Rhizobium.leguminosarum.bv.trifolii GCA_011604525.1_ASM1160452v1
93.6/95
Bartonella.quintana GCA_009936175.1_ASM993617v1
Pelagibacterium.halotolerans GCA_013004005.1_ASM1300400v1
100/100
100/100
Polymorphum.gilvum.SL003B.26A1 GCA_000192745.1_ASM19274v1
100/100
Roseibium.hamelinense GCA_007830545.1_ASM783054v1
100/100
Pseudovibrio.sp.FO.BEG1 GCA_000236645.1_ASM23664v1
100/100
100/100
Methylocella.silvestris.BL2 GCA_000021745.1_ASM2174v1
Methylocystis.bryophila GCA_002117405.1_ASM211740v1
100/100
Methylorubrum.extorquens.PA1 GCA_000018845.1_ASM1884v1
100/100
Rhodopseudomonas.palustris GCA_013415845.1_ASM1341584v1
98.2/99
Xanthobacter.autotrophicus.Py2 GCA_000017645.1_ASM1764v1
100/100
100/100
Hyphomicrobium.denitrificans.ATCC.51888 GCA_000143145.1_ASM14314v1
Rhodomicrobium.vannielii.ATCC.17100 GCA_000166055.1_ASM16605v1
100/100
Thermopetrobacter.sp.TC1 GCA_000746275.1_ASM74627v1
100/100
Candidatus.Micropelagos.thuwalensis GCA_000469155.1.RS24
100/100
alpha.proteobacterium.IMCC14465 GCA_000293845.2_ASM29384v2
Parvibaculum.lavamentivorans.DS.1 GCA_000017565.1_ASM1756v1
100/100
Asticcacaulis.excentricus.CB.48 GCA_000175215.2_ASM17521v2
100/100
Brevundimonas.subvibrioides.ATCC.15264 GCA_000144605.1_ASM14460v1
100/100
alpha.proteobacterium.L41A GCA_000335735.1.L41A.1.0
100/100
100/100
Caulobacter.vibrioides.NA1000 GCA_000022005.1_ASM2200v1
Phenylobacterium.zucineum.HLK1 GCA_000017265.1_ASM1726v1
100/100
100/100
Hirschia.baltica.ATCC.49814 GCA_000023785.1_ASM2378v1
100/100
Hyphomonas.neptunium.ATCC.15444 GCA_000013025.1_ASM1302v1
100/100
Maricaulis.maris GCA_003634045.1_ASM363404v1
Oceanicaulis.alexandrii.DSM.11625 GCA_000420265.1_ASM42026v1
Parvularcula.bermudensis.HTCC2503 GCA_000152825.2_ASM15282v2
95/95
Jannaschia.rubra GCA_900113265.1.IMG.taxon.2622736430.annotated.assembly
100/100
100/100
Octadecabacter.arcticus.238 GCA_000155735.2_ASM15573v2
100/100
Ruegeria.pomeroyi.DSS.3 GCA_000011965.2_ASM1196v2
100/100
Sagittula.stellata.E.37 GCA_000169415.1_ASM16941v1
100/100
100/100
Ketogulonicigenium.vulgare GCA_001399515.1_ASM139951v1
Rubellimicrobium.rubrum GCA_006152115.1_ASM615211v1
100/100
100/100
Paracoccus.denitrificans GCA_004063735.1_ASM406373v1
Rhodobacter.sphaeroides.ATCC.17029 GCA_000015985.1_ASM1598v1
Neomegalonema.perideroedes.DSM.15528 GCA_000374145.1_ASM37414v1
100/100
100/100
Candidatus.Pelagibacter.ubique.HIMB058 GCA_000419545.1_ASM41954v1
100/100
100/100
Candidatus.Pelagibacter.ubique.HTCC1062 GCA_000012345.1_ASM1234v1
alpha.proteobacterium.HIMB5 GCA_000299095.1_ASM29909v1
alpha.proteobacterium.HIMB114 GCA_000163555.2
100/100
Citromicrobium.sp.JL477 GCA_001304795.1_ASM130479v1
100/100
Sphingopyxis.macrogoltabida GCA_001314325.1_ASM131432v1
100/100
Sphingobium.indicum.B90A GCA_000264945.2_ASM26494v2
100/100
Sphingomonas.wittichii.RW1 GCA_000016765.1_ASM1676v1
100/100
Zymomonas.mobilis.subsp.mobilis.ATCC.31821 GCA_000007105.1_ASM710v1
100/100
alpha.proteobacterium.AAP81b GCA_001295935.1_ASM129593v1
100/100
Kordiimonas.gwangyangensis.DSM.19435 GCA_000375545.1_ASM37554v1
100/100
GCA_002701885.1_ASM270188v1*
100/100
GCA_002722055.1_ASM272205v1*
Sneathiella.glossodoripedis.JCM.23214 GCA_000616095.1_ASM61609v1
100/100
Geminicoccus.roseus.DSM.18922 GCA_000427665.1_ASM42766v1
100/99
Tistrella.mobilis.KA081020.065 GCA_000264455.2_ASM26445v2
100/100
Caenispirillum.salinarum.AK4 GCA_000315795.1
100/100
Pararhodospirillum.photometricum.DSM.122 GCA_000284415.2_ASM28441v2
90.4/98
Rhodospirillum.rubrum.F11 GCA_000225955.1_ASM22595v1
100/100
Roseospirillum.parvum GCA_900100455.1
100/100
Magnetospirillum.magneticum.AMB.1 GCA_000009985.1_ASM998v1
100/100
Terasakiella.pusilla.DSM.6293 GCA_000688235.1_ASM68823v1
100/99
Thalassospira.profundimaris.WP0211 GCA_000300275.1_ASM30027v1
100/44
100/100
Candidatus.Puniceispirillum.marinum.IMCC1322 GCA_000024465.1_ASM2446v1
100/100
Thalassobaculum.salexigens.DSM.19539 GCA_000423805.1_ASM42380v1
100/100
alpha.proteobacterium.BAL199 GCA_000171835.1_ASM17183v1
100/100
Oceanibaculum.indicum GCA_003633955.1_ASM363395v1
100/100
Kiloniella.laminariae.DSM.19542 GCA_000374005.1_ASM37400v1
100/100
Rhodovibrio.salinarum.DSM.9154 GCA_000515255.1_ASM51525v1
100/100
Inquilinus.limosus.DSM.16000 GCA_000423185.1_ASM42318v1
100/100
Rhodospirillum.centenum GCA_014197805.1_ASM1419780v1
72.4/41
Micavibrio.aeruginosavorus.ARL.13 GCA_000226315.1_ASM22631v1
100/100
Caedimonas.varicaedens GCA_001192655.1_ASM119265v1
100/100
Candidatus.Caedibacter.acanthamoebae GCA_000743035.1_ASM74303v1
Candidatus.Nucleicultrix.amoebiphila.FS5 GCA_002117145.1_ASM211714v1
100/100
100/100
Candidatus.Finniella.inopinata GCA_004210305.1_ASM421030v1
100/100
Candidatus.Odyssella.thessalonicensis.L13 GCA_000190415.2_ASM19041v1
31.7/70
Candidatus.Paracaedibacter.acanthamoebae GCA_000742835.1_ASM74283v1
100/100
100/100
Candidatus.Hepatobacter.penaei GCA_000742475.1_ASM74247v1
Holospora.undulata.HU1 GCA_000388175.3_ASM38817v3
Elioraea.tepidiphila.DSM.17972 GCA_000378465.1_ASM37846v1
100/100
Belnapia.moabensis.DSM.16746 GCA_000745835.1_ASM74583v1
100/100
Rubritepida.flocculans.DSM.14296 GCA_000425365.1_ASM42536v1
100/100
Roseomonas.gilardii GCA_001941945.1_ASM194194v1
100/100
Acidiphilium.multivorum.AIU301 GCA_000202835.1_ASM20283v1
92.4/94
76.9/89
Acidisphaera.rubrifaciens.HS.AP3 GCA_000964365.1_ASM96436v1
100/100
Granulibacter.bethesdensis.CGDNIH4 GCA_000576145.2_ASM57614v2
100/100
Commensalibacter.intestini.A911 GCA_000231445.2_ASM23144v1
93.3/100
Gluconacetobacter.diazotrophicus.PA1.5 GCA_000067045.1_ASM6704v1
Acetobacter.aceti.NBRC.14818 GCA_000193495.2_ASM19349v2
Gluconobacter.oxydans.DSM.3504 GCA_000583855.1_ASM58385v1
0.557

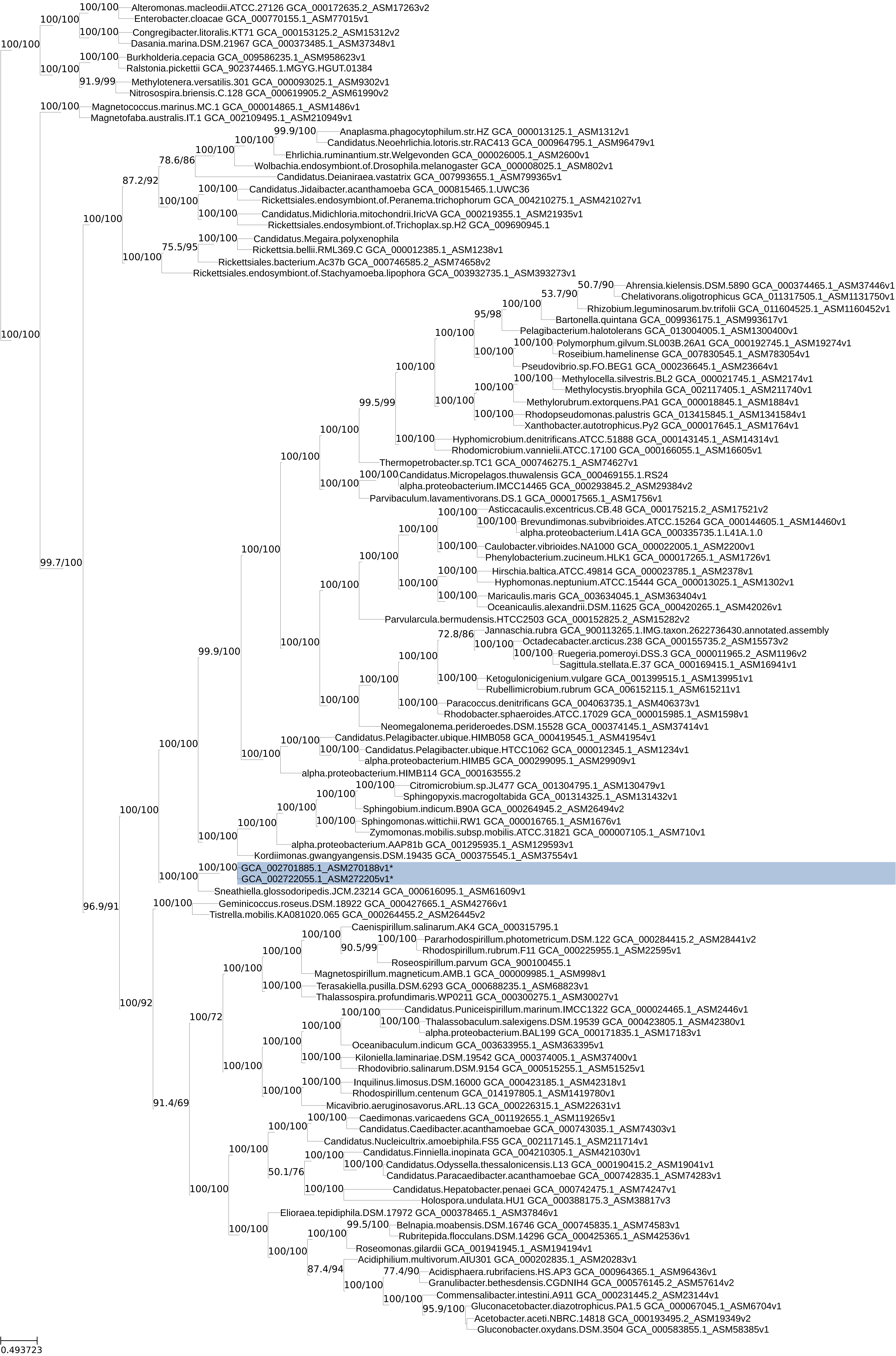
100/100
Alteromonas.macleodii.ATCC.27126 GCA_000172635.2_ASM17263v2
Enterobacter.cloacae GCA_000770155.1_ASM77015v1
100/100
100/100
Congregibacter.litoralis.KT71 GCA_000153125.2_ASM15312v2
Dasania.marina.DSM.21967 GCA_000373485.1_ASM37348v1
100/100
100/100
Burkholderia.cepacia GCA_009586235.1_ASM958623v1
Ralstonia.pickettii GCA_902374465.1.MGYG.HGUT.01384
100/100
91.9/99
Methylotenera.versatilis.301 GCA_000093025.1_ASM9302v1
Nitrosospira.briensis.C.128 GCA_000619905.2_ASM61990v2
100/100
Magnetococcus.marinus.MC.1 GCA_000014865.1_ASM1486v1
Magnetofaba.australis.IT.1 GCA_002109495.1_ASM210949v1
99.9/100
Anaplasma.phagocytophilum.str.HZ GCA_000013125.1_ASM1312v1
100/100
Candidatus.Neoehrlichia.lotoris.str.RAC413 GCA_000964795.1_ASM96479v1
100/100
Ehrlichia.ruminantium.str.Welgevonden GCA_000026005.1_ASM2600v1
78.6/86
Wolbachia.endosymbiont.of.Drosophila.melanogaster GCA_000008025.1_ASM802v1
Candidatus.Deianiraea.vastatrix GCA_007993655.1_ASM799365v1
87.2/92
100/100
Candidatus.Jidaibacter.acanthamoeba GCA_000815465.1.UWC36
100/100
Rickettsiales.endosymbiont.of.Peranema.trichophorum GCA_004210275.1_ASM421027v1
100/100
Candidatus.Midichloria.mitochondrii.IricVA GCA_000219355.1_ASM21935v1
Rickettsiales.endosymbiont.of.Trichoplax.sp.H2 GCA_009690945.1
100/100
100/100
Candidatus.Megaira.polyxenophila
75.5/95
Rickettsia.bellii.RML369.C GCA_000012385.1_ASM1238v1
100/100
Rickettsiales.bacterium.Ac37b GCA_000746585.2_ASM74658v2
Rickettsiales.endosymbiont.of.Stachyamoeba.lipophora GCA_003932735.1_ASM393273v1
50.7/90
Ahrensia.kielensis.DSM.5890 GCA_000374465.1_ASM37446v1
53.7/90
Chelativorans.oligotrophicus GCA_011317505.1_ASM1131750v1
100/100
Rhizobium.leguminosarum.bv.trifolii GCA_011604525.1_ASM1160452v1
95/98
Bartonella.quintana GCA_009936175.1_ASM993617v1
Pelagibacterium.halotolerans GCA_013004005.1_ASM1300400v1
100/100
100/100
Polymorphum.gilvum.SL003B.26A1 GCA_000192745.1_ASM19274v1
100/100
Roseibium.hamelinense GCA_007830545.1_ASM783054v1
Pseudovibrio.sp.FO.BEG1 GCA_000236645.1_ASM23664v1
100/100
100/100
100/100
Methylocella.silvestris.BL2 GCA_000021745.1_ASM2174v1
Methylocystis.bryophila GCA_002117405.1_ASM211740v1
100/100
Methylorubrum.extorquens.PA1 GCA_000018845.1_ASM1884v1
99.5/99
100/100
Rhodopseudomonas.palustris GCA_013415845.1_ASM1341584v1
Xanthobacter.autotrophicus.Py2 GCA_000017645.1_ASM1764v1
100/100
100/100
Hyphomicrobium.denitrificans.ATCC.51888 GCA_000143145.1_ASM14314v1
Rhodomicrobium.vannielii.ATCC.17100 GCA_000166055.1_ASM16605v1
100/100
Thermopetrobacter.sp.TC1 GCA_000746275.1_ASM74627v1
100/100
Candidatus.Micropelagos.thuwalensis GCA_000469155.1.RS24
100/100
alpha.proteobacterium.IMCC14465 GCA_000293845.2_ASM29384v2
Parvibaculum.lavamentivorans.DS.1 GCA_000017565.1_ASM1756v1
100/100
Asticcacaulis.excentricus.CB.48 GCA_000175215.2_ASM17521v2
100/100
100/100
Brevundimonas.subvibrioides.ATCC.15264 GCA_000144605.1_ASM14460v1
alpha.proteobacterium.L41A GCA_000335735.1.L41A.1.0
100/100
100/100
Caulobacter.vibrioides.NA1000 GCA_000022005.1_ASM2200v1
Phenylobacterium.zucineum.HLK1 GCA_000017265.1_ASM1726v1
100/100
100/100
Hirschia.baltica.ATCC.49814 GCA_000023785.1_ASM2378v1
100/100
Hyphomonas.neptunium.ATCC.15444 GCA_000013025.1_ASM1302v1
100/100
Maricaulis.maris GCA_003634045.1_ASM363404v1
Oceanicaulis.alexandrii.DSM.11625 GCA_000420265.1_ASM42026v1
Parvularcula.bermudensis.HTCC2503 GCA_000152825.2_ASM15282v2
72.8/86
Jannaschia.rubra GCA_900113265.1.IMG.taxon.2622736430.annotated.assembly
100/100
Octadecabacter.arcticus.238 GCA_000155735.2_ASM15573v2
100/100
Ruegeria.pomeroyi.DSS.3 GCA_000011965.2_ASM1196v2
99.9/100
100/100
Sagittula.stellata.E.37 GCA_000169415.1_ASM16941v1
100/100
100/100
Ketogulonicigenium.vulgare GCA_001399515.1_ASM139951v1
Rubellimicrobium.rubrum GCA_006152115.1_ASM615211v1
100/100
100/100
Paracoccus.denitrificans GCA_004063735.1_ASM406373v1
Rhodobacter.sphaeroides.ATCC.17029 GCA_000015985.1_ASM1598v1
Neomegalonema.perideroedes.DSM.15528 GCA_000374145.1_ASM37414v1
100/100
100/100
Candidatus.Pelagibacter.ubique.HIMB058 GCA_000419545.1_ASM41954v1
100/100
Candidatus.Pelagibacter.ubique.HTCC1062 GCA_000012345.1_ASM1234v1
100/100
alpha.proteobacterium.HIMB5 GCA_000299095.1_ASM29909v1
alpha.proteobacterium.HIMB114 GCA_000163555.2
100/100
Citromicrobium.sp.JL477 GCA_001304795.1_ASM130479v1
100/100
Sphingopyxis.macrogoltabida GCA_001314325.1_ASM131432v1
100/100
Sphingobium.indicum.B90A GCA_000264945.2_ASM26494v2
100/100
100/100
Sphingomonas.wittichii.RW1 GCA_000016765.1_ASM1676v1
100/100
Zymomonas.mobilis.subsp.mobilis.ATCC.31821 GCA_000007105.1_ASM710v1
alpha.proteobacterium.AAP81b GCA_001295935.1_ASM129593v1
Kordiimonas.gwangyangensis.DSM.19435 GCA_000375545.1_ASM37554v1
100/100
GCA_002701885.1_ASM270188v1*
100/100
GCA_002722055.1_ASM272205v1*
Sneathiella.glossodoripedis.JCM.23214 GCA_000616095.1_ASM61609v1
96.9/91
100/100
Geminicoccus.roseus.DSM.18922 GCA_000427665.1_ASM42766v1
Tistrella.mobilis.KA081020.065 GCA_000264455.2_ASM26445v2
100/100
Caenispirillum.salinarum.AK4 GCA_000315795.1
100/100
Pararhodospirillum.photometricum.DSM.122 GCA_000284415.2_ASM28441v2
100/100
90.5/99
Rhodospirillum.rubrum.F11 GCA_000225955.1_ASM22595v1
Roseospirillum.parvum GCA_900100455.1
100/100
Magnetospirillum.magneticum.AMB.1 GCA_000009985.1_ASM998v1
100/100
Terasakiella.pusilla.DSM.6293 GCA_000688235.1_ASM68823v1
100/92
Thalassospira.profundimaris.WP0211 GCA_000300275.1_ASM30027v1
100/72
100/100
Candidatus.Puniceispirillum.marinum.IMCC1322 GCA_000024465.1_ASM2446v1
100/100
100/100
Thalassobaculum.salexigens.DSM.19539 GCA_000423805.1_ASM42380v1
alpha.proteobacterium.BAL199 GCA_000171835.1_ASM17183v1
100/100
Oceanibaculum.indicum GCA_003633955.1_ASM363395v1
100/100
Kiloniella.laminariae.DSM.19542 GCA_000374005.1_ASM37400v1
100/100
Rhodovibrio.salinarum.DSM.9154 GCA_000515255.1_ASM51525v1
100/100
Inquilinus.limosus.DSM.16000 GCA_000423185.1_ASM42318v1
100/100
Rhodospirillum.centenum GCA_014197805.1_ASM1419780v1
91.4/69
Micavibrio.aeruginosavorus.ARL.13 GCA_000226315.1_ASM22631v1
100/100
Caedimonas.varicaedens GCA_001192655.1_ASM119265v1
100/100
Candidatus.Caedibacter.acanthamoebae GCA_000743035.1_ASM74303v1
Candidatus.Nucleicultrix.amoebiphila.FS5 GCA_002117145.1_ASM211714v1
100/100
100/100
Candidatus.Finniella.inopinata GCA_004210305.1_ASM421030v1
100/100
Candidatus.Odyssella.thessalonicensis.L13 GCA_000190415.2_ASM19041v1
50.1/76
Candidatus.Paracaedibacter.acanthamoebae GCA_000742835.1_ASM74283v1
100/100
Candidatus.Hepatobacter.penaei GCA_000742475.1_ASM74247v1
100/100
Holospora.undulata.HU1 GCA_000388175.3_ASM38817v3
Elioraea.tepidiphila.DSM.17972 GCA_000378465.1_ASM37846v1
99.5/100
Belnapia.moabensis.DSM.16746 GCA_000745835.1_ASM74583v1
100/100
100/100
Rubritepida.flocculans.DSM.14296 GCA_000425365.1_ASM42536v1
Roseomonas.gilardii GCA_001941945.1_ASM194194v1
100/100
Acidiphilium.multivorum.AIU301 GCA_000202835.1_ASM20283v1
77.4/90
Acidisphaera.rubrifaciens.HS.AP3 GCA_000964365.1_ASM96436v1
87.4/94
100/100
Granulibacter.bethesdensis.CGDNIH4 GCA_000576145.2_ASM57614v2
Commensalibacter.intestini.A911 GCA_000231445.2_ASM23144v1
100/100
Gluconacetobacter.diazotrophicus.PA1.5 GCA_000067045.1_ASM6704v1
95.9/100
Acetobacter.aceti.NBRC.14818 GCA_000193495.2_ASM19349v2
Gluconobacter.oxydans.DSM.3504 GCA_000583855.1_ASM58385v1
0.493723

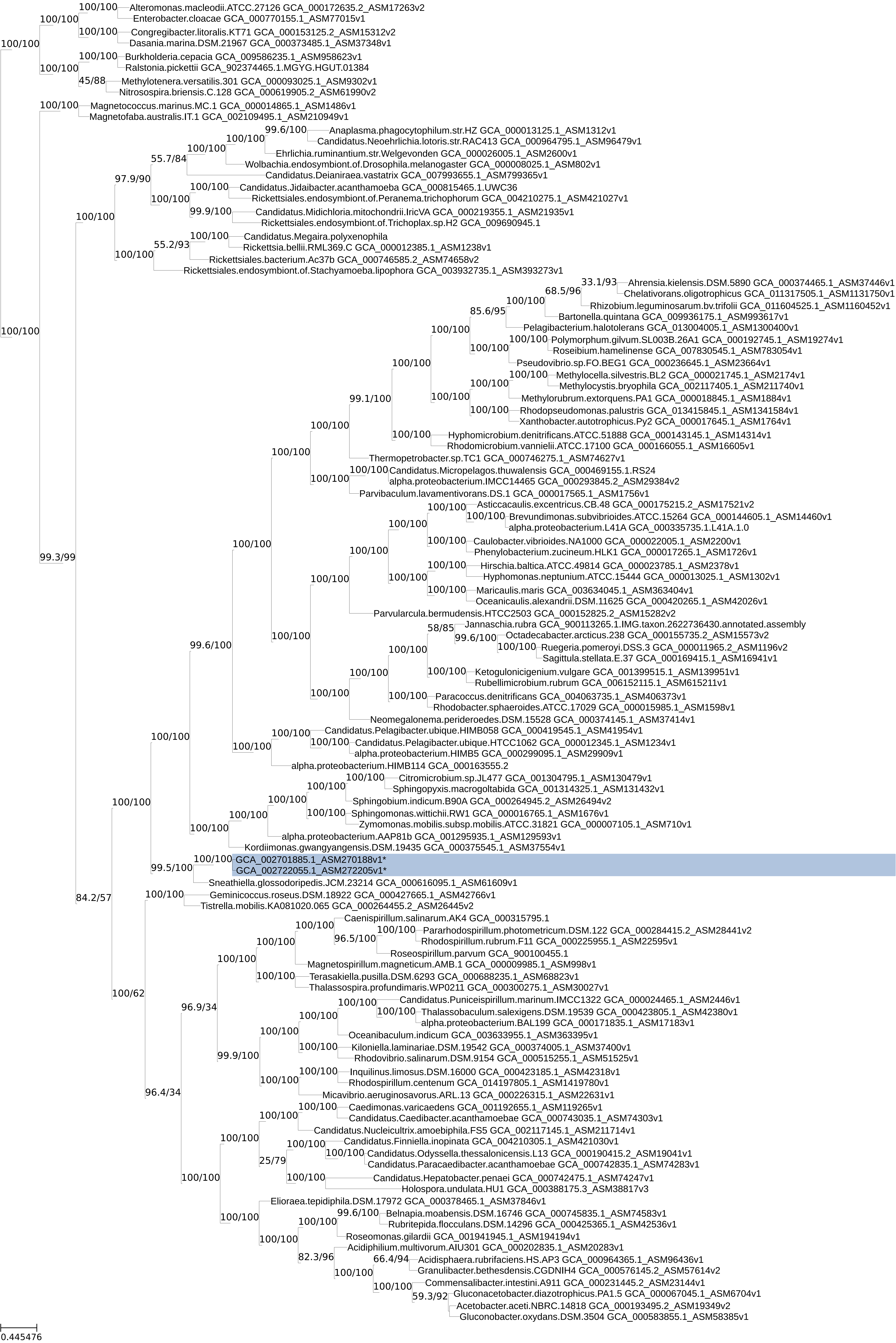
100/100 Alteromonas.macleodii.ATCC.27126 GCA_000172635.2_ASM17263v2
Enterobacter.cloacae GCA_000770155.1_ASM77015v1
100/100
100/100 Congregibacter.litoralis.KT71 GCA_000153125.2_ASM15312v2
Dasania.marina.DSM.21967 GCA_000373485.1_ASM37348v1
100/100
100/100 Burkholderia.cepacia GCA_009586235.1_ASM958623v1
Ralstonia.pickettii GCA_902374465.1.MGYG.HGUT.01384
100/100
45/88 Methylotenera.versatilis.301 GCA_000093025.1_ASM9302v1
Nitrosospira.briensis.C.128 GCA_000619905.2_ASM61990v2
100/100 Magnetococcus.marinus.MC.1 GCA_000014865.1_ASM1486v1
Magnetofaba.australis.IT.1 GCA_002109495.1_ASM210949v1
99.6/100 Anaplasma.phagocytophilum.str.HZ GCA_000013125.1_ASM1312v1
100/100 Candidatus.Neoehrlichia.lotoris.str.RAC413 GCA_000964795.1_ASM96479v1
100/100 Ehrlichia.ruminantium.str.Welgevonden GCA_000026005.1_ASM2600v1
55.7/84 Wolbachia.endosymbiont.of.Drosophila.melanogaster GCA_000008025.1_ASM802v1
Candidatus.Deianiraea.vastatrix GCA_007993655.1_ASM799365v1
97.9/90
100/100 Candidatus.Jidaibacter.acanthamoeba GCA_000815465.1.UWC36
100/100 Rickettsiales.endosymbiont.of.Peranema.trichophorum GCA_004210275.1_ASM421027v1
100/100
99.9/100 Candidatus.Midichloria.mitochondrii.IricVA GCA_000219355.1_ASM21935v1
Rickettsiales.endosymbiont.of.Trichoplax.sp.H2 GCA_009690945.1
100/100 Candidatus.Megaira.polyxenophila
55.2/93 Rickettsia.bellii.RML369.C GCA_000012385.1_ASM1238v1
100/100 Rickettsiales.bacterium.Ac37b GCA_000746585.2_ASM74658v2
Rickettsiales.endosymbiont.of.Stachyamoeba.lipophora GCA_003932735.1_ASM393273v1
33.1/93 Ahrensia.kielensis.DSM.5890 GCA_000374465.1_ASM37446v1
68.5/96 Chelativorans.oligotrophicus GCA_011317505.1_ASM1131750v1
100/100 Rhizobium.leguminosarum.bv.trifolii GCA_011604525.1_ASM1160452v1
100/100
85.6/95 Bartonella.quintana GCA_009936175.1_ASM993617v1
Pelagibacterium.halotolerans GCA_013004005.1_ASM1300400v1
100/100
100/100 Polymorphum.gilvum.SL003B.26A1 GCA_000192745.1_ASM19274v1
100/100 Roseibium.hamelinense GCA_007830545.1_ASM783054v1
Pseudovibrio.sp.FO.BEG1 GCA_000236645.1_ASM23664v1
100/100
100/100 Methylocella.silvestris.BL2 GCA_000021745.1_ASM2174v1
100/100 Methylocystis.bryophila GCA_002117405.1_ASM211740v1
99.1/100
100/100 Methylorubrum.extorquens.PA1 GCA_000018845.1_ASM1884v1
100/100 Rhodopseudomonas.palustris GCA_013415845.1_ASM1341584v1
Xanthobacter.autotrophicus.Py2 GCA_000017645.1_ASM1764v1
100/100
100/100 Hyphomicrobium.denitrificans.ATCC.51888 GCA_000143145.1_ASM14314v1
Rhodomicrobium.vannielii.ATCC.17100 GCA_000166055.1_ASM16605v1
100/100
Thermopetrobacter.sp.TC1 GCA_000746275.1_ASM74627v1
100/100 Candidatus.Micropelagos.thuwalensis GCA_000469155.1.RS24
100/100 alpha.proteobacterium.IMCC14465 GCA_000293845.2_ASM29384v2
Parvibaculum.lavamentivorans.DS.1 GCA_000017565.1_ASM1756v1
100/100 Asticcacaulis.excentricus.CB.48 GCA_000175215.2_ASM17521v2
100/100 Brevundimonas.subvibrioides.ATCC.15264 GCA_000144605.1_ASM14460v1
100/100 alpha.proteobacterium.L41A GCA_000335735.1.L41A.1.0
100/100
100/100 Caulobacter.vibrioides.NA1000 GCA_000022005.1_ASM2200v1
Phenylobacterium.zucineum.HLK1 GCA_000017265.1_ASM1726v1
100/100
100/100 Hirschia.baltica.ATCC.49814 GCA_000023785.1_ASM2378v1
100/100 Hyphomonas.neptunium.ATCC.15444 GCA_000013025.1_ASM1302v1
100/100
100/100 Maricaulis.maris GCA_003634045.1_ASM363404v1
Oceanicaulis.alexandrii.DSM.11625 GCA_000420265.1_ASM42026v1
Parvularcula.bermudensis.HTCC2503 GCA_000152825.2_ASM15282v2
58/85 Jannaschia.rubra GCA_900113265.1.IMG.taxon.2622736430.annotated.assembly
100/100
99.6/100 Octadecabacter.arcticus.238 GCA_000155735.2_ASM15573v2
100/100 Ruegeria.pomeroyi.DSS.3 GCA_000011965.2_ASM1196v2
Sagittula.stellata.E.37 GCA_000169415.1_ASM16941v1
100/100
100/100 Ketogulonicigenium.vulgare GCA_001399515.1_ASM139951v1
Rubellimicrobium.rubrum GCA_006152115.1_ASM615211v1
99.6/100
100/100
100/100 Paracoccus.denitrificans GCA_004063735.1_ASM406373v1
Rhodobacter.sphaeroides.ATCC.17029 GCA_000015985.1_ASM1598v1
Neomegalonema.perideroedes.DSM.15528 GCA_000374145.1_ASM37414v1
100/100 Candidatus.Pelagibacter.ubique.HIMB058 GCA_000419545.1_ASM41954v1
100/100
100/100 Candidatus.Pelagibacter.ubique.HTCC1062 GCA_000012345.1_ASM1234v1
alpha.proteobacterium.HIMB5 GCA_000299095.1_ASM29909v1
alpha.proteobacterium.HIMB114 GCA_000163555.2
100/100 Citromicrobium.sp.JL477 GCA_001304795.1_ASM130479v1
100/100 Sphingopyxis.macrogoltabida GCA_001314325.1_ASM131432v1
100/100 Sphingobium.indicum.B90A GCA_000264945.2_ASM26494v2
100/100 Sphingomonas.wittichii.RW1 GCA_000016765.1_ASM1676v1
100/100 Zymomonas.mobilis.subsp.mobilis.ATCC.31821 GCA_000007105.1_ASM710v1
100/100 alpha.proteobacterium.AAP81b GCA_001295935.1_ASM129593v1
100/100 Kordiimonas.gwangyangensis.DSM.19435 GCA_000375545.1_ASM37554v1
100/100 GCA_002701885.1_ASM270188v1*
99.5/100 GCA_002722055.1_ASM272205v1*
Sneathiella.glossodoripedis.JCM.23214 GCA_000616095.1_ASM61609v1
100/100 Geminicoccus.roseus.DSM.18922 GCA_000427665.1_ASM42766v1
Tistrella.mobilis.KA081020.065 GCA_000264455.2_ASM26445v2
100/100 Caenispirillum.salinarum.AK4 GCA_000315795.1
100/100 Pararhodospirillum.photometricum.DSM.122 GCA_000284415.2_ASM28441v2
96.5/100 Rhodospirillum.rubrum.F11 GCA_000225955.1_ASM22595v1
100/100 Roseospirillum.parvum GCA_900100455.1
100/100 Magnetospirillum.magneticum.AMB.1 GCA_000009985.1_ASM998v1
100/100 Terasakiella.pusilla.DSM.6293 GCA_000688235.1_ASM68823v1
Thalassospira.profundimaris.WP0211 GCA_000300275.1_ASM30027v1
100/100 Candidatus.Puniceispirillum.marinum.IMCC1322 GCA_000024465.1_ASM2446v1
96.9/34
100/100 Thalassobaculum.salexigens.DSM.19539 GCA_000423805.1_ASM42380v1
100/100 alpha.proteobacterium.BAL199 GCA_000171835.1_ASM17183v1
100/100 Oceanibaculum.indicum GCA_003633955.1_ASM363395v1
100/100 Kiloniella.laminariae.DSM.19542 GCA_000374005.1_ASM37400v1
99.9/100 Rhodovibrio.salinarum.DSM.9154 GCA_000515255.1_ASM51525v1
100/100 Inquilinus.limosus.DSM.16000 GCA_000423185.1_ASM42318v1
100/100 Rhodospirillum.centenum GCA_014197805.1_ASM1419780v1
96.4/34 Micavibrio.aeruginosavorus.ARL.13 GCA_000226315.1_ASM22631v1
100/100 Caedimonas.varicaedens GCA_001192655.1_ASM119265v1
100/100 Candidatus.Caedibacter.acanthamoebae GCA_000743035.1_ASM74303v1
Candidatus.Nucleicultrix.amoebiphila.FS5 GCA_002117145.1_ASM211714v1
100/100 Candidatus.Finniella.inopinata GCA_004210305.1_ASM421030v1
100/100
100/100 Candidatus.Odyssella.thessalonicensis.L13 GCA_000190415.2_ASM19041v1
25/79 Candidatus.Paracaedibacter.acanthamoebae GCA_000742835.1_ASM74283v1
100/100
100/100 Candidatus.Hepatobacter.penaei GCA_000742475.1_ASM74247v1
Holospora.undulata.HU1 GCA_000388175.3_ASM38817v3
Elioraea.tepidiphila.DSM.17972 GCA_000378465.1_ASM37846v1
99.6/100 Belnapia.moabensis.DSM.16746 GCA_000745835.1_ASM74583v1
100/100
100/100 Rubritepida.flocculans.DSM.14296 GCA_000425365.1_ASM42536v1
Roseomonas.gilardii GCA_001941945.1_ASM194194v1
100/100 Acidiphilium.multivorum.AIU301 GCA_000202835.1_ASM20283v1
82.3/96
66.4/94 Acidisphaera.rubrifaciens.HS.AP3 GCA_000964365.1_ASM96436v1
100/100 Granulibacter.bethesdensis.CGDNIH4 GCA_000576145.2_ASM57614v2
100/100 Commensalibacter.intestini.A911 GCA_000231445.2_ASM23144v1
59.3/92 Gluconacetobacter.diazotrophicus.PA1.5 GCA_000067045.1_ASM6704v1
Acetobacter.aceti.NBRC.14818 GCA_000193495.2_ASM19349v2
Gluconobacter.oxydans.DSM.3504 GCA_000583855.1_ASM58385v1
100/100
99.3/99
84.2/57
100/62
100/100
0.445476
