## Supplementary material for "Host association and intracellularity evolved multiple times independently in the *Rickettsiales*": figures_and_supplementary_materials: Supplementary_text_1_taxonomic_descriptions.docx

**Description of “*Candidatus* Vederia obscura” gen. nov. sp. nov.**

“Candidatus Ve.de’ri.a ob.scu’ra”. (Ve.de’ri.a: N.L. fem. n., light as a feather, due to the small genome size, from the Dutch veder, and resembling a character’s name in George Lucas’ Star Wars saga; ob.scu’ra. L. fem. adj. obscura, hidden, unseen, as the bacterium was not directly observed). This bacterium was found in association with the ciliate *Plagiopyla frontata* strain IBS-3, and belongs to the family “*Candidatus* Midichloriaceae”. Uncultured so far. Basis of the assignment, genome sequence.

**Description of “*Candidatus* Diomedesia obscura” gen. nov. sp. nov.**

“*Candidatus* Di.o.me.de’si.a ob.scu’ra”. (Di.o.me.de’si.a: N.L. fem. n., dedicated to Diomedes, in reference to mythological Thracian king Diomedes, owner of mares stolen by Heracles in one of his labours; ob.scu’ra. L. fem. adj. obscura, hidden, unseen, as the bacterium was not directly observed). The genome sequence of this bacterium was obtained from an environmental water sample collected from Black Sea, next to Varna, Bulgaria. Uncultured so far. Basis of the assignment, genome sequence (GCA_016780625.1).

**Description of “*Candidatus* Diomedesiaceae” fam. nov.**

“*Candidatus* Diomedesiaceae” (Di.o.me.de.si.a'ce.ae, N.L. fem. n. “*Candidatus* Diomedesia” type

genus of the family; suff. -aceae ending to denote a family; N.L. fem. pl. n. “*Candidatus*

Diomedesiaceae” the family of genus “*Candidatus* Diomedesia”).

The family “*Candidatus* Diomedesiaceae” is defined based on phylogenetic analyses of concatenated orthologous genes of uncultured representatives from various marine environments. The family belongs to the order *Rickettsiales*, and currently contains one genus, “*Candidatus* Diomedesia”.

**Description of “*Candidatus* Jistubacter obscurus” gen. nov. sp. nov.**

“*Candidatus* Jis.tu.bac’ter ob.scu’rus”. (Jis.tu.bac’ter: N.L. Jistu, the Native American mythological trickster rabbit, a light-hearted character prone to humorously inappropriate behaviour; N. L. masc. n. bacter, rod; N. L. masc. n. Jistubacter, Jistu rod [also intended as an ironic compliment to one of the authors]; ob.scu’rus. L. adj. obscurus, hidden, unseen, as the bacterium was not directly observed). The genome sequence of this bacterium was obtained from environmental freshwater sample collected from the Chattahoochee river, USA. Uncultured so far. Basis of the assignment, genome sequence (GCA_010031735.1).

**Description of “Candidatus Jistubacteraceae” fam. nov.**

“*Candidatus* Jistubacteraceae” (Jis.tu.bac.te.ra'ce.ae, N.L. fem. n. “*Candidatus* Jistubacter” type

genus of the family; suff. -aceae ending to denote a family; N.L. fem. pl. n. “*Candidatus*

Jistubacteraceae” the family of genus “*Candidatus* Jistubacter”).

The family “*Candidatus* Jistubacteraceae” is defined based on phylogenetic analyses of concatenated orthologous genes of uncultured representatives from various freshwater and terrestrial environments. The family belongs to the order *Rickettsiales*, and currently contains one genus, “*Candidatus* Jistubacter”.

**Description of “*Candidatus* Arkhamia obscura” gen. nov. sp. nov.**

“*Candidatus* Ar.kha’mi.a ob.scu’ra”. (Ar.kha’mi.a: N.L. fem. n., dedicated to Arkham, fictional town in Massachussets created by Howard Phillips Lovecraft; ob.scu’ra. L. fem. adj. obscura, hidden, unseen, as the bacterium was not directly observed). The genome sequence of this bacterium was found in an environmental soil sample from Massachussets, USA. Basis of the assignment, genome sequence (GCA_013288575.1).

**Description of “Candidatus Arkhamiaceae” fam. nov.**

“*Candidatus* Arkhamiaceae” (Ar.kha.mi.a'ce.ae, N.L. fem. n. “*Candidatus* Arkhamia” type

genus of the family; suff. -aceae ending to denote a family; N.L. fem. pl. n. “*Candidatus*

Arkhamiaceae” the family of genus “*Candidatus* Arkhamia”).

The family “*Candidatus* Arkhamiaceae” is defined based on phylogenetic analyses of concatenated orthologous genes of uncultured representatives from freshwater and terrestrial environments. The family belongs to the order *Rickettsiales*, and currently contains one genus, “*Candidatus* Arkhamia”.
