## Supplementary material for "Host association and intracellularity evolved multiple times independently in the *Rickettsiales*": figures_and_supplementary_materials: Supplementary_text_2_assembly.docx

**Supplementary text 2: genome assembly procedures**

In this work, the novel genome sequences of 9 *Rickettsiales* bacteria were assembled. This was obtained starting from 7 ciliate strains/populations (with *Euplotes woodruffi* NDG2 harbouring 2 different *Rickettsiales* bacteria) and from one sample derived from a previous study (Gloekner et al. 2014) on the foraminiferan *Reticulomyxa filosa*.

**Sample origin and maintenance**

Among, the seven ciliate strains/populations employed in this study, five were obtained and described in previous studies (Catania et al. 2009; Vannini et al. 2010; Tarcz et al. 2012; Boscaro et al. 2013; Szokoli et al. 2016; Lanzoni et al. 2019; Mironov and Sabaneyeva 2020). The origin and laboratory maintenance of the other two strains is reported below.

The population of *Plagiopyla frontata* IBS-3 was obtained from a brackish water sample collected in May 2018 from the Baltic Sea in Inkoo (Finland), following a previously described protocol for the laboratory propagation of anaerobic ciliates (Nitla et al. 2019).

The monoclonal strain *Euplotes woodruffi* NDG2 was isolated from a water sample collected on 15^th^ March 2013 at Naval Dockyard Gate in Visakhapatnam (Andhra Pradesh, India). It was maintained in the laboratory by regular feeding (every 1-2 weeks) with *Dunaliella tertiolecta*, grown in artificial water at 5‰ salinity as previously described (e.g. Vannini et al. 2010).

**Sample preparation and Illumina sequencing**

Four ciliates (*Paramecium biaurelia* US_Bl 11III1, *Paramecium* *nephridiatum* Sr 2-6, *Euplotes woodruffi* NDG2, *Plagiopyla frontata* IBS-3) were subjected to whole-genome amplification (WGA) with the REPLI-g Single Cell Kit (Qiagen), starting from a limited number of cells (see Supplementary table 1), and following the protocol for live cells (as in Castelli et al. 2021). This procedure allows to obtain a sufficient amount of DNA for Illumina sequencing even for samples (*Plagiopyla* *frontata* IBS-3) that are not easily propagated in the laboratory, as well as to minimise the presence of food bacteria contamination thanks to the manual washing procedure, which in any case would not be amenable for larger amounts of host cells. For the sample *Euplotes harpa* BOD18,which was not available as live material, WGA was performed from a DNA extract obtained in a previous study (Vannini et al. 2010), following the dedicated protocol of the Repli-G kit.

For the sample *Paramecium biaurelia* USBL-36I1, 1.2 liters of laboratory culture (ca. 1,500-2,000 *Paramecium* cells/ml) were processed and DNA was extracted for sequencing using a modified CTAB protocol, as previously described (Floriano et al. 2018).

For sample *Paramecium multimicronucleatum* Kr154-4, 400 ml of laboratory culture (ca. 600 *Paramecium* cells/ml) were process and DNA extraction was performed following a phenol-chloroform protocol (Sambrook and Russell 2006).

Each of the seven DNA samples from ciliates was processed through a Nextera XT library, and sequenced by Admera Health (South Plainfield, NJ, USA) on a Illumina HiSeq X machine producing 2x150 bp paired-end reads (Supplementary table 1).

The reads from the *Reticulomyxa filosa* sample (Gloeckner et al. 2014), were kindly provided by Gernot Gloeckner and Marco Groth (University of Koeln, Germany).

**Sample preparation and Oxford Nanopore sequencing**

In order to perform Nanopore sequencing, a large enough amount of DNA extract was obtained from *Euplotes woodruffii* NDG2 as follows. A total of 3 liters of NDG2 culture were obtained (~ 300,000 *Euplotes* cells). The culture was passed through a sieve with 20 µm-sized pores, so that cells were retained by the filter, resulting concentrated in the final 12 ml of unfiltered liquid, which was then centrifuged at 5000 g for 5 min. The pellet was resuspended in 2 ml of distilled water, and centrifuged again. The final pellet was fixed in ethanol and stored until DNA extraction, performed with the NucleoSpin™ Tissue Kit (Macherey-Nagel™). The DNA extract was processed through a SQK-LSK109 ligation-sequencing library, following the manufacturer’s instructions. Afterwards, library was loaded onto a FLO-MIN106 flowcell and sequenced on a Minion device using the MinKNOW software 18.12.9, for 72 total hours.

Basecalling was then performed with the guppy 5.0.11 software by Oxford Nanopore. Then, reads were processed with Porechop 0.2.4 (Wick et al. 2017a) with default options. Quality of the reads was assessed with NanoPlot 1.23.0 (De Coster et al. 2018). The following steps are described in the detailed report on the selective assembly of the sample NDG2 below.

**Selective assembly of the *Rickettsiales* bacterial genomes from shotgun sequencing of their host samples**

In this section a general account of the procedures is presented. Subsequently, a detailed report for each sample is available.

For each sample, after a quality assessment with FastQC (Andrews 2010), the reads were assembled using SPAdes 3.6 (Bankevich et al. 2012) with default settings (except for the *Reticulomyxa filosa* sample, which was assembled with Unicycler 0.4.8 (Wick et al. 2017b)), obtaining a “preliminary assembly”.

Then, a multi-step procedure was applied, in order to select only those contigs belonging to the symbiont of interest and discard those belonging to the host and to additional organisms present in the sample (e.g. residual food, additional associated bacteria), as described previously (e.g. Castelli et al. 2019).

For this purpose, the blobology pipeline was applied (Kumar et al. 2013). Briefly, the contigs of the “preliminary assembly” were classified according to their length, GC% content, sequencing coverage based on reads mapped with Bowtie2 (Langmead and Salzberg 2012), and NCBI taxonomy of the best megablast hit on NCBI nucleotide. Results were visualised by the R package ggplot2 (Wickham 2016). Accordingly, a set of contigs was selected, considering also ribosomal RNA genes identified with barrnap (Seemann 2013), using default options for *Bacteria*. Thus, reads mapping on the selected contigs were reassembled separately with SPAdes. Results were manually revised prior and after reassembly, by examining blastp results on NCBI nr protein database after annotation with Prokka (Seemann 2014), as well as taking advantage of Bandage (Wick et al. 2015) for visualisation of assembly graphs to identify connections between contigs. The “contigs.fasta”, “contigs.fastg”, “scaffolds.fasta”, and “scaffolds.fastg” SPAdes output files were considered, and, unless specified, reported results refer to the “contigs” files. Only for the sample *E. woodruffi* NDG2, for which Nanopore reads were produced, re-assembly was directly performed as hybrid (Illumina+Nanopore) with Unicycler 0.4.8-beta (see details below).

Moreover, an additional round of screening was performed in order to account for potential sequences belonging to *Rickettsiales* symbiont(s) in contigs falling outside the selected sets of contigs, especially considering possible “hybrid” contigs between host and symbiont, which may otherwise have been discarded in case of best hit on host sequences. In detail, all the respectively annotated ORFs (Hyatt et al. 2010) where queried on NCBI nr protein database with DIAMOND (Bunchfink et al. 2015), applying an e-value threshold of 1e-5. Contigs with at least one ORF with a best hit on *Bacteria* were inspected manually (by their blast and DIAMOND hits), and, in case, further contigs selections and/or refinements of the previous ones were made, followed by re-assemblies and manual revision, as described above.

**Genome closing by PCR and Sanger sequencing**

After selecting the *Rickettsiales* genome sequences from SPAdes assemblies, two samples (*P. multimicronucleatum* 12 and *P. biaurelia* US_Bl 11III1) were selected, based on assembly quality, for molecular biology based finishing. PCR primers were designed next to the ends of larger contigs, and PCR reactions were performed in order to test their connections and join them into larger contigs/scaffolds. PCRs were performed with TaKaRa Ex Taq and reagents (Takara Bio, Japan), following manufacturer’s instructions (each primer with final concentration 0.5 μM). Successful PCR results were confirmed by bidirectional Sanger sequencing (after purification with the EuroGOLD Cycle-Pure kit - EuroClone, Italy) performed by GATC Biotech (Germany) with amplification primers and/or internal primers (see sections below for details on each sample).

***Paramecium multimicronucleatum* Kr154-4 hosting “*Ca*. Trichorickettsia mobilis”**

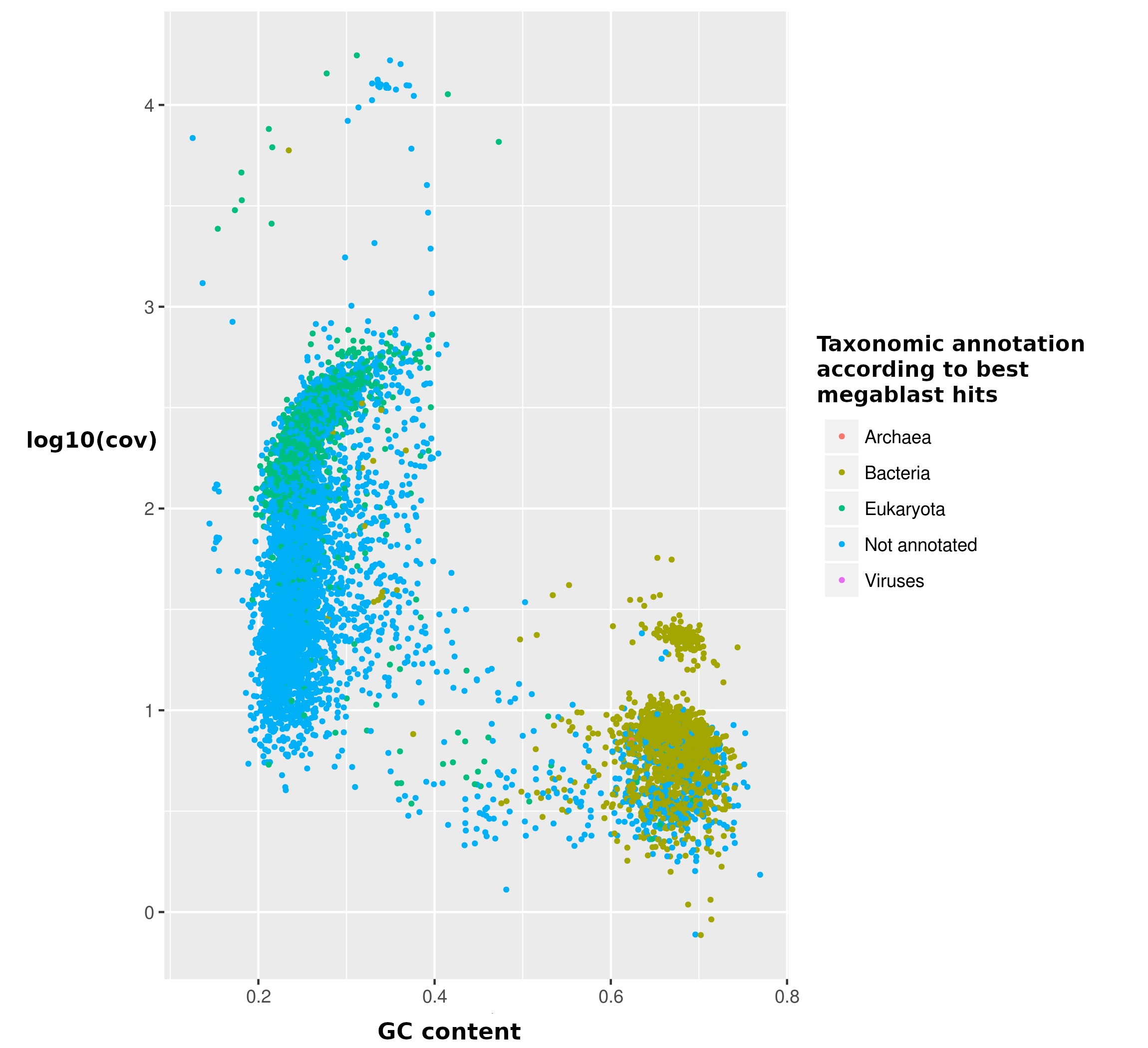

**Figure A** Plot of the preliminary assembly contigs according to their GC content and log_10_ of sequencing coverage, coloured according to the respective best megablast hit. Only contigs with length higher than or equal to 1000 bp are shown for viewers’ clarity.

In total, 24 full-length or partial rRNA genes were identified in the “preliminary assembly” (Supplementary table 4):

- 13 complete or partial rRNA genes had best megablast hits on nuclear or mitochondrial *Paramecium* spp., and were thus assigned to the host. The respective contigs ranged from 18.59 to 17607.88 in sequencing coverage
- 3 rRNA genes (16S, 23S and 5S) were identified as belonging to the symbiont, based on their best megablast hits on “*Ca*. Trichorickettsia” (16S) or *Rickettsia* (23S), or for being in the same contig as the 23S (5S). The respective contigs ranged from 36.57 to 38.55 in sequencing coverage, and ~0.34 GC content
- 8 remaining complete or partial rRNA gene sequences had megablast hits on other organisms. The respective contigs ranged from 3.85 to 356.00 sequencing coverage, having generally a higher GC content than the those assigned to “*Ca*. Trichorickettsia” (>0.44), with just one exception (0.26 GC)

Taking into account the plot and the rRNA gene features, we designed a “selection from preliminary assembly” for “*Ca*. Trichorickettsia”, including all contigs with log_10_ of coverage lower than 2.5 and GC content between 0.25 and 0.4, with the exclusion of those having best megablast hit on members of order Peniculida (to with the *Paramecium* host belongs).

The reads mapping to “selection from preliminary assembly” were extracted and reassembled separately with SPAdes default settings: “final SPAdes assembly” (10,563,744 bp, 11700 contigs, N50=2,955 bp; L50=570). In this assembly, a selection was made of contigs belonging to “*Ca*. Trichorickettsia (14 contigs, 1,468,577 bp; N50=1,094,380 bp; L50=1; Figure B).

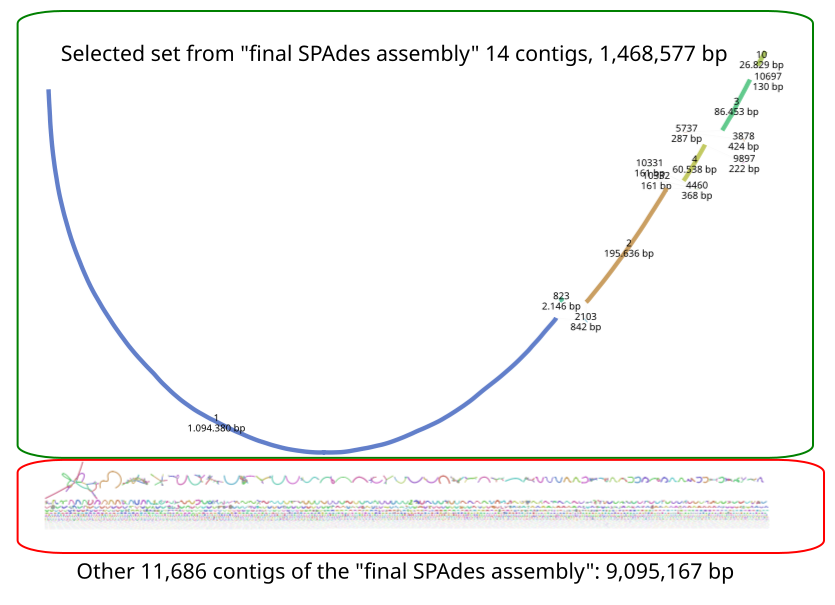

**Figure B** Visual representation of the 11700 contigs of the “final SPAdes assembly” and the relative connections in the assembly graph, obtained with the Bandage software. The green square highlights the selected set of contigs. The red square highlights the remaining contigs of the assembly.

Then, primers were designed next to the ends of the 6 larger contigs in the selected set, and PCR reactions were performed in order to identify/confirm junctions among those contigs. The combinations of primers used for amplification and sequencing, primer sequences, and the reaction protocols used are reported below.

| **Reaction** | **Amplification primer 1** | **Amplification primer 2** | **Sequencing primer 1** | **Sequencing primer 2** | **Protocol** |
| --- | --- | --- | --- | --- | --- |
| 1 | Tricho_10Fa | Tricho_3Ra | Tricho_10Fb | Tricho_3Rb | A |
| 2 | Tricho_3F | Tricho_4R | Tricho_3F | Tricho_4R | A |
| 3 | Tricho_4Fa | Tricho_2Ra | Tricho_4Fb | Tricho_2Rb | A |
| 4 | Tricho_2F | Tricho_823F | Tricho_2F | Tricho_823F | B |
| 5 | Tricho_823Ra | Tricho_1Fa | Tricho_823Rb | Tricho_1Fb | A |
| 6 | Tricho_1Ra | Tricho_10R | Tricho_1Rb | Tricho_10R | A |

| **Primer** | **Sequence (F: forward; R: reverse)** |
| --- | --- |
| Tricho_1Fa | 5'-TTGGCAGAAGCACTAATTGAG-3' |
| Tricho_1Fb | 5'-TAACTGGTAATGATCTCCACG-3' |
| Tricho_1Ra | 5'-AGCGAAGCAGTGACATTGG-3' |
| Tricho_1Rb | 5'-TACAGTGACCACAGTTCGG-3' |
| Tricho_2F | 5'-CCCTCATATCTAGGTTGAGC-3' |
| Tricho_2Ra | 5'-TGATAACATACGAGTAGCAGC-3' |
| Tricho_2Rb | 5'-TTAACGCAGGCACTACTCC-3' |
| Tricho_3F | 5'-TCAACGCTCTTTCCTGTGG-3' |
| Tricho_3Ra | 5'-AACCTGCCGTGAGGTGC-3' |
| Tricho_3Rb | 5'-ATCAGTAGCAGCGCCTAC-3' |
| Tricho_4Fa | 5'-AACAGCAGCATGCACTGAG-3' |
| Tricho_4Fb | 5'-CTACCCTCATAGGGTATCC-3' |
| Tricho_4R | 5'-CGACTCAAGGCTCGTGG-3' |
| Tricho_10Fa | 5'-CCAGAGTCCGTTAGAATAGG-3' |
| Tricho_10Fb | 5'-GCCCCAGGCCTTAACATC-3' |
| Tricho_10R | 5'-GTAATTGGAGCCTTAGAGTTC-3' |
| Tricho_823F | 5'-CTTGATTAGGTCTTGCTTCC-3' |
| Tricho_823Ra | 5'-ACTTATCTCGATAGGCTGAAC-3' |
| Tricho_823Rb | 5'-CTGACCTCCTCTATCAAGC-3' |

Protocol A (standard)

| 94°C | 3 min |  |
| --- | --- | --- |
| 94°C | 30 s | 40 cycles |
| 50°C | 30 s |  |
| 72°C | 4 min |  |
| 72°C | 10 min |  |

Protocol B (touchdown)

| 94°C | 3 min |  |
| --- | --- | --- |
| 94°C | 30 s | 5 cycles |
| 56°C | 30 s |  |
| 72°C | 4 min |  |
| 94°C | 30 s | 10 cycles |
| 53°C | 30 s |  |
| 72°C | 4 min |  |
| 94°C | 30 s | 30 cycles |
| 50°C | 30 s |  |
| 72°C | 4 min |  |
| 72°C | 10 min |  |

Thanks to these PCR experiments, it was possible to close the “*Ca*. Trichorickettsia mobilis” genome into a single circular scaffold (1,470,091 bp).

In addition, the DIAMOND hit-based inspection of contigs of the “preliminary assembly” outside the “selection from preliminary assembly” evidenced the presence of putative bacterial/*Rickettsia*-like sequences in higher coverage contigs, which were assigned to putative “*Ca*. Trichorickettsia” plasmid(s). Thus, we designed an additional “selection for plasmids” from the “preliminary assembly”, including all contigs with log_10_ of coverage higher than 1.5.

The reads mapping to “selection for plasmids” were extracted and reassembled separately with SPAdes (based on different attempts, the option -k 21,33,55,77,99 was chosen): “plasmid re-assembly” (39,539,931 bp, 4182 contigs, N50=82,014 bp; L50=128). In this assembly, at least 3 plasmids were identified. Two plasmids were assembled in single circular contigs, 57,766 bp and 17,134 bp long, respectively. The other plasmidic sequences were assembled into 69 contigs (457,770 bp), and may even constitute one or more additional plasmids: “additional plasmid(s)” (Figure C).

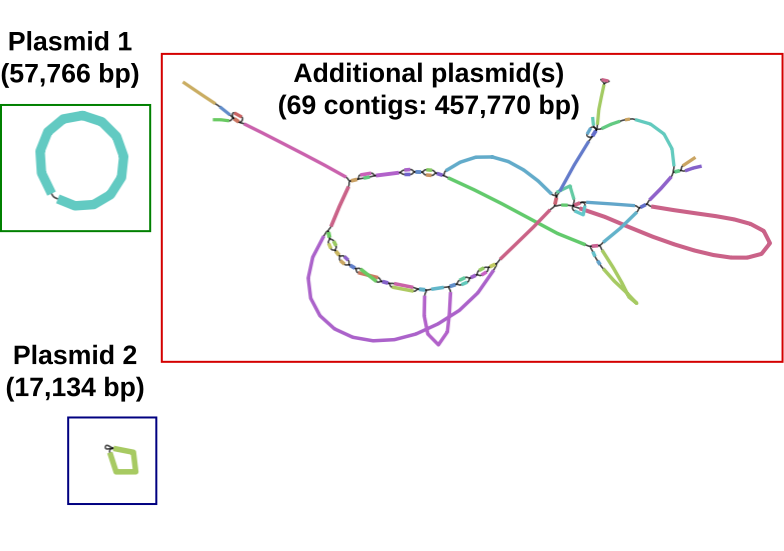

**Figure C** Visual representation of the selected 71 contigs of the “plasmid re-assembly” and the relative connections in the assembly graph, obtained with the Bandage software. The green square highlights the plasmid 1, the blue square highlights plasmid 2, and the red square highlights the “additional plasmid(s)”

Finally, the assembly of “additional plasmid(s)” was refined by removing all contigs <5000bp containing no validated ORF (i.e. with no hits on NCBI nr or with hits only on hypothetical proteins, transposases, phages). Thus, the “final assembly” of the “additional plasmid(s)” included 36 contigs (420,184 bp; N50=28,079 bp; L50=26).

In total, the assembly of “*Ca*. Trichorickettsia”, including chromosome and all plasmids, consisted of 39 contigs (1,965,805 bp).

***Paramecium nephridiatum* Sr 2-6 hosting “*Ca.* Megaira venefica”**

The “preliminary assembly” was constituted by 21,901 contigs (21,515,176 bp; N50 = 2,009 bp; L50 = 2817)

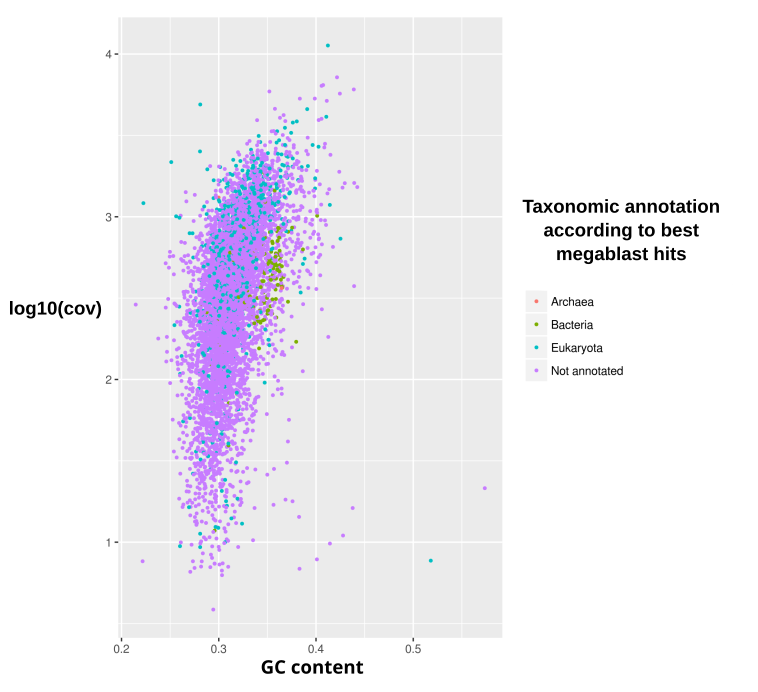

**Figure D** Plot of the preliminary contigs of the sample Sr 2-6 according to their GC content and log_10_ of sequencing coverage, coloured according to the respective best megablast hit. Only contigs with length higher than or equal to 1000 bp are shown for viewers’ clarity.

In total, 36 full-length or partial rRNA genes were identified in the “preliminary assembly” (Supplementary table 4):

- 34 complete rRNA genes had best megablast hits on nuclear *Paramecium* sequences, and were thus assigned to the host. The respective contigs displayed 7.28 and 11285.11 sequencing coverage
- 2 rRNA gene sequences (16S and 5S) were identified as belonging to “*Ca*. Megaira venefica”, based on their respective megablast hits. Namely the 16S rRNA gene had high identity (>99%) with “*Ca*. Megaira spp.”, while 5S rRNA gene had no “informative” hit (low identity hits ~80% with several other proteobacterial sequences), but its full-length contig had best hit on *Rickettsia japonica*. The two respective contigs displayed 595.70 and 569.53 sequencing coverage, and ~0.36 GC content.

As done for the other samples, we selected contigs based on sequencing coverage, GC content, and best megablast hits. Then, we reassembled the reads mapping on these contigs. However, no significant improvement of the assembly of “*Ca*. Megaira venefica” genome was observed (not shown). Therefore, selection of “*Ca*. Megaira venefica” sequences was performed directly on the “preliminary assembly”, counting 628 contigs (2,047,029 bp).

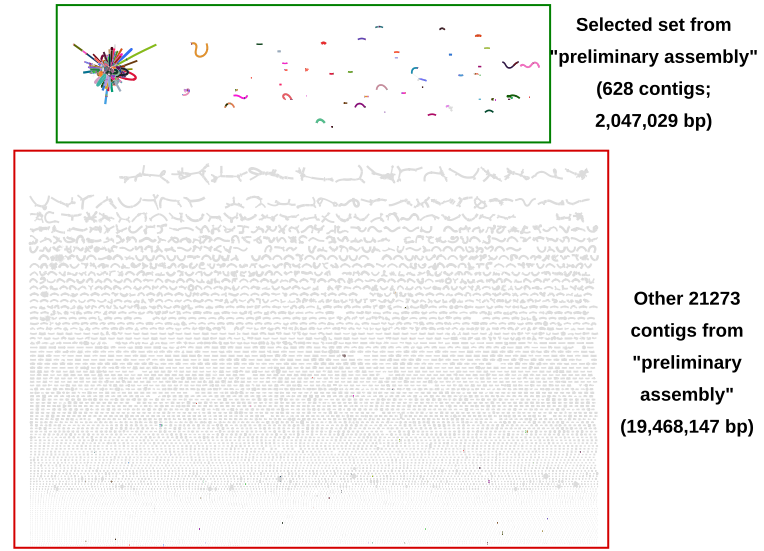

**Figure E** Visual representation of the 21,901 contigs of the “preliminary assembly” and the relative connections in the assembly graph, obtained with the Bandage software. The green square highlights the selected set of contigs, the red square highlights the remaining contigs of the assembly.

The relatively high fragmentation degree and the presence of several small contigs in the selection from “preliminary assembly” were interpreted as due to the presence of many repeated elements in the genome of “*Ca*. Megaira venefica”, considering the high connection degree (Figure E) and the presence of multiple transposase genes. Accordingly, and accounting both for the careful manual revision applied and for the comparative BUSCO results with other *Rickettsiales* (Supplementary table 3), we concluded that this assembly likely contains the whole genome sequence (in terms of gene sequences) of “*Ca*. Megaira venefica”.

Finally, the assembly was refined by removing all contigs <3000bp containing no validated ORF (i.e. with no hits on NCBI nr or with hits only on hypothetical proteins, transposases, phages). Thus, the “final assembly” of “*Ca*. Megaira venefica” counted 281 contigs (1,862,567 bp; N50=10,544 bp; L50=58).

***Euplotes* *woodruffi* NDG2 hosting “*Ca*. Megaira polyxenophila” and “*Ca*. Bandiella woodruffii”**

The “preliminary assembly” was constituted by 26,682 contigs (15,101,154 bp; N50 = 772 bp; L50 = 1633)

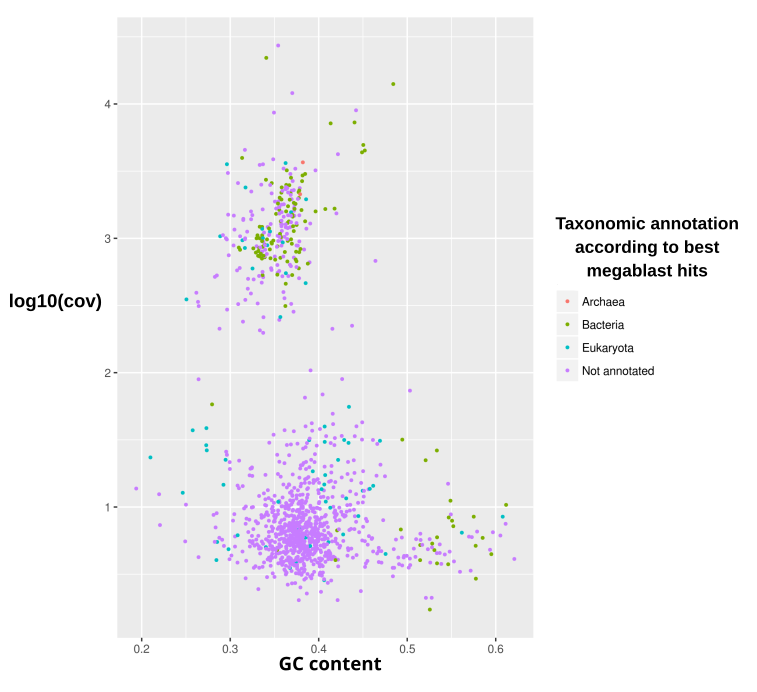

**Figure F** Plot of the preliminary contigs of the sample NDG2 according to their GC content and log_10_ of sequencing coverage, coloured according to the respective best megablast hit. Only contigs with length higher than or equal to 1000 bp are shown for viewers’ clarity.

In total, 17 full-length or partial rRNA genes were identified in the “preliminary assembly” (Supplementary table 4):

- 3 partial rRNA genes had best megablast hits on nuclear *Euplotes* sequences, and were thus assigned to the host. The two respective contigs displayed quite lower sequencing coverage (31.12 and 37.25)
- 3 rRNA genes (5S, 16S, 23S) formed a putative operon on a single contig. Based on megablast hits, they were identified as belonging to *Polynucleobacter necessarius*, a betaproteobacterial obligate symbiont of many *Euplotes spp.*, including *E. woodruffi* (e.g. Senra et al. 2016). This contig displayed 4363.69 sequencing coverage
- 3 rRNA genes (5S, 16S, 23S) were identified as belonging to a “*Ca*. Bandiella woodruffii” symbiont, based on their respective best megablast hits on “*Ca*. Midichloriaceae” bacteria. The precise species assignment was due to the 99.70% identity of the 16S rRNA gene with the type “*Ca*. Bandiella woodruffii” endosymbiont of *E. woodruffi* Sq1 (Senra et al. 2016), though not representing the best hit due to the relatively short length of this sequence. The two respective contigs displayed 648.22 and 1664.01 sequencing coverage
- 3 rRNA gene sequences (16S, 23S and 5S) were identified as belonging to a “*Ca*. Megaira polyxenophila” symbiont, based on their respective megablast hits, including for the 16S rRNA gene hits with high identity (>99%) on “*Ca*. Megaira polyxenophila” sequences (including symbiont of *Carteria cerasiformis* (Kawafune et al. 2012)), and for the 23S rRNA hits on *Rickettsia* *japonica*. The two respective contigs displayed 899.13 and 914.37 sequencing coverage
- 5 remaining complete or partial rRNA gene sequences had megablast hits on other organisms. The respective contigs had comparably much lower coverage (<26.42) than the ones assigned to the two *Rickettsiales* bacteria

Taking into account the plot and the rRNA gene features, we designed a “selection from preliminary assembly” targeting the two *Rickettsiales*, including all contigs with log_10_ of coverage higher than 1.5, and excluding those with best megablast hit on *Polynucleobacter*.

The reads mapping to this “selection from preliminary assembly” were extracted. In parallel, the corresponding Nanopore reads were extracted by blasting total reads on the contigs of the same “selection from preliminary assembly” (default megablast options) and keeping all reads with hits. The hybrid assembly of these two sets of reads “Unicycler hybrid re-assembly” included 195 contigs (3,240,079 bp, N50=503,502 bp; L50=2). In this assembly, two selection were made for sequences belonging respectively to “*Ca*. Megaira polyxenophila” (28 contigs, 1,995,794 bp), and to “*Ca*. Bandiella woodruffii” (163 contigs; 1,205,980 bp; Figure G). Each selection contained multiple putative plasmids (at least 9 for “*Ca*. Megaira”, 2 for “*Ca*. Bandiella”), which were assigned to each organism based on the connection in the Illumina assembly graphs, as well as on the sequence similarity of the annotated genes among them and with published sequences (e.g. published plasmidic sequences in *Rickettsia* for sequences assigned to “*Ca*. Megaira”).

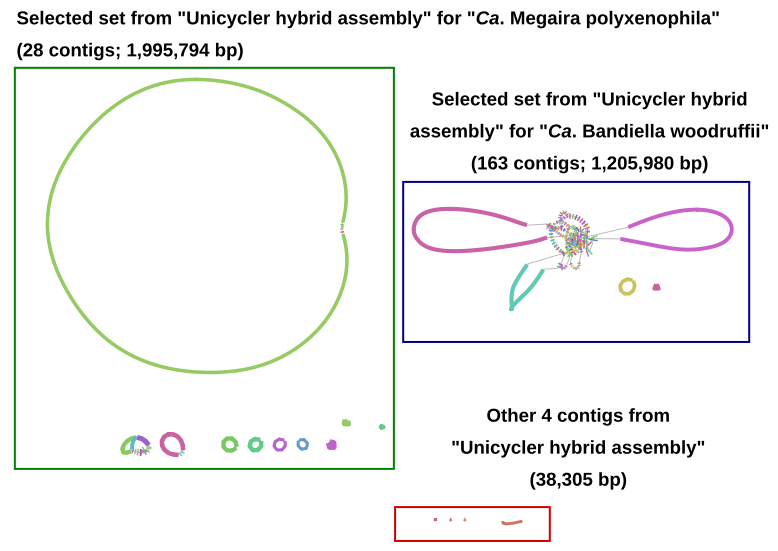

**Figure G** Visual representation of the 195 contigs of the “Unicycler hybrid assembly” and the relative connections in the assembly graph, obtained with the Bandage software. The green square highlights the selected set of contigs for “*Ca*. Megaira polyxenophila”, the blue square highlights the selected set of contigs for “*Ca*. Bandiella woodruffii”, and red square highlights the remaining contigs of the assembly.

For what concerns the putative “*Ca*. Megaira polyxenophila” and “*Ca*. Bandiella woodruffii” chromosomes, a further manual step was performed in order to improve the assembly. Namely, all selected Nanopore reads were blasted (default megablast options) on the contig ends (1000 bp, only for contigs larger than 1000 bp), and those with hits were manually inspected, in order to identify potentially bridging reads between contigs. Accordingly, it was possible to close each genome into a single circular contig (“*Ca*. Megaira polyxenophila”: 1,508,753 bp; “*Ca*. Bandiella woodruffii”: 1,129,747 bp). Moreover, the assembly of plasmids was further refined by removing all contigs ≤1000bp containing no validated ORF (i.e. with no hits on NCBI nr or with hits only on hypothetical proteins or transposases). Thus, the two “final assemblies”, including chromosome and plasmids, counted respectively 14 contigs (1,994,314 bp) for “*Ca*. Megaira polyxenophila”, and 3 contigs (1,225,299 bp) for “*Ca*. Bandiella woodruffii” (see Supplementary table 1 for details on the length of each plasmid).

***Euplotes* *harpa* BOD18 hosting “*Ca*. Cyrtobacter comes”**

The “preliminary assembly” was constituted by 135,782 contigs (99,489,754 bp; N50 = 1,566 bp; L50 = 16270)

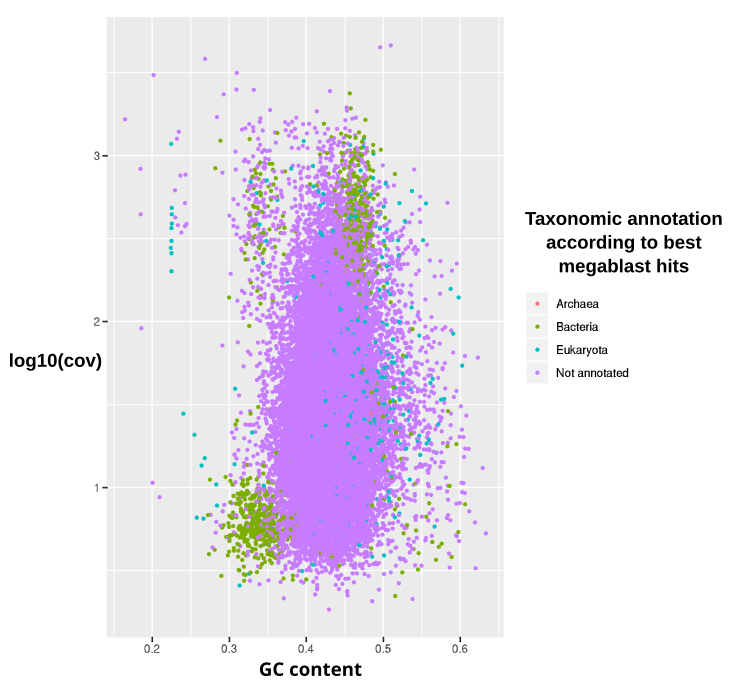

**Figure H** Plot of the preliminary contigs of the sample BOD18 according to their GC content and log_10_ of sequencing coverage, coloured according to the respective best megablast hit. Only contigs with length higher than or equal to 1000 bp are shown for viewers’ clarity.

In total, 15 full-length or partial rRNA genes were identified in the “preliminary assembly” (Supplementary table 4):

- 2 complete rRNA genes had best megablast hits on nuclear *Euplotes* sequences, and were thus assigned to the host. The respective contigs displayed 50.00 and 61.05 sequencing coverage
- one 16 rRNA gene sequence was identified as belonging to “*Ca*. Cyrtobacter comes”, based on its best megablast hit on this organism with 100% identity. The respective contig displayed 668.69 sequencing coverage, and 0.36 GC content
- one complete and one partial rRNA gene sequences were identified based on their megablast hits as belonging to the *Polynucleobacter necessarius* (*Betaproteobacteria*) symbiont previously characterised in the BOD18 strain (Vannini et al. 2010). The respective contigs had relatively comparable coverage (330.76 and 557.69) to the “*Ca*. Cyrtobacter” one, but significantly higher GC content (0.48 and 0.52).
- 10 remaining complete or partial rRNA gene sequences had megablast hits on other organisms (9) or no hit (1). The respective contigs had a clearly lower coverage (<37.57) than the one assigned to “*Ca*. Cyrtobacter”

Taking into account the plot and the rRNA gene features, we designed a “selection from preliminary assembly” targeting “*Ca*. Cyrtobacter”, including all contigs with log_10_ of coverage higher then 1.5.

The reads mapping to this “selection from preliminary assembly” were extracted and reassembled separately with SPAdes (based on different attempts, the option -k 21,33,55,77,99 was chosen). This “SPAdes re-assembly” included 22442 contigs (36,602,319 bp, N50=3,630 bp; L50=2747). In this assembly, a selection was made of sequences belonging to “*Ca*. Cyrtobacter” (689 contigs, 1,524,096 bp; Figure I).

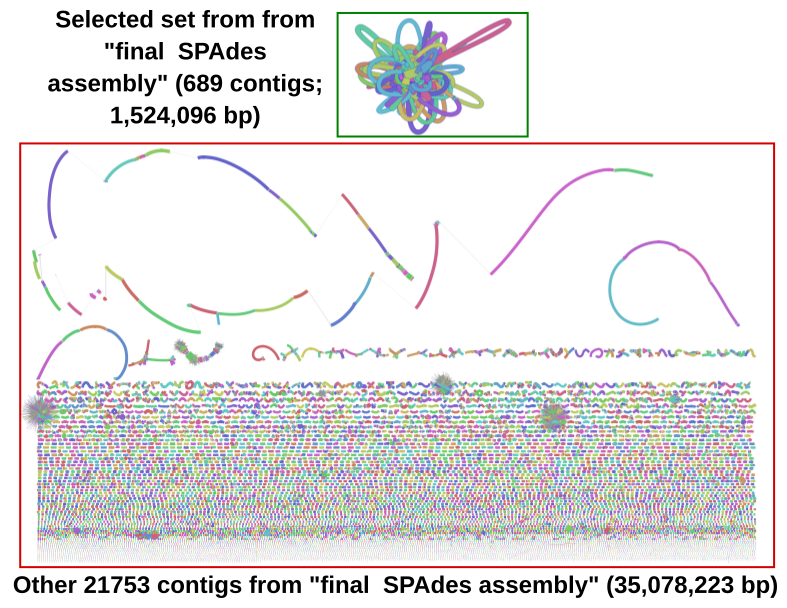

**Figure I** Visual representation of the 22442 contigs of the “final SPAdes assembly” and the relative connections in the assembly graph, obtained with the Bandage software. The green square highlights the selected set of contigs, the red square highlights the remaining contigs of the assembly.

The relatively high fragmentation degree and the presence of several small contigs in “final SPAdes assembly” were interpreted as due to the presence of many repeated elements in the genome of “*Ca*. Cyrtobacter”, considering the high level of connection between the contigs (Figure I) and the presence of multiple transposase genes (data not shown). Accordingly, and accounting both for the careful manual revision applied and for the comparative BUSCO results with other *Rickettsiales* (Supplementary table 3):, we concluded that this assembly likely contains the whole genome sequence (in terms of gene sequences) of “*Ca*. Cyrtobacter comes”.

Finally, the assembly was refined by removing all contigs <2000bp containing no validated ORF (i.e. with no hits on NCBI nr or with hits only on hypothetical proteins or transposases). Thus, the “final assembly” of the “*Ca*. Cyrtobacter comes” counted 141 contigs (1,308,703 bp; N50=14,445 bp; L50=28).

***Paramecium biaurelia* US_Bl 11III1 hosting “*Ca*. Fokinia cryptica”**

The “preliminary assembly” was constituted by 115,057 contigs (136,752,753 bp; N50 = 10,679 bp; L50 = 2209)

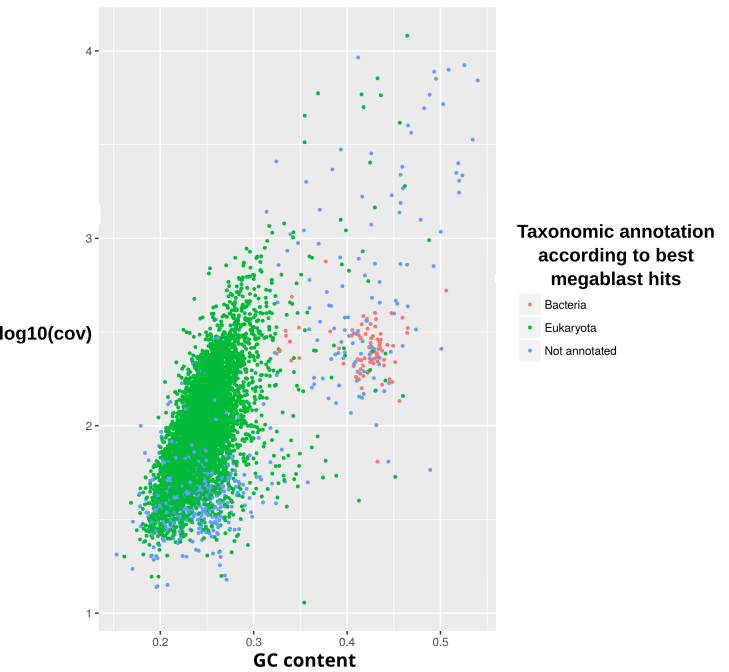

**Figure J** Plot of the preliminary contigs of the sample US_Bl 11III1 according to their GC content and log_10_ of sequencing coverage, coloured according to the respective best megablast hit. Only contigs with length higher than or equal to 1000 bp are shown for viewers’ clarity.

In total, 31 full-length or partial rRNA genes were identified in the “preliminary assembly” (Supplementary table 4):

- 25 complete or partial rRNA genes had best megablast hits on nuclear or mitochondrial *Paramecium* spp., and were thus assigned to the host (in one case, the partial gene sequence had no hit and the assignment was based on the megablast hit of the whole respective contig). The respective contigs ranged from 6.34 to 5944.93 in sequencing coverage
- 3 rRNA gene sequences (16S, 23S, 5S) were identified as belonging to “*Ca*. Fokinia cryptica”, based on their best megablast hits on “*Ca*. Fokinia spp.”. The two respective contigs displayed 281.37 and 301.99 sequencing coverage, and ~0.34 GC content
- 3 rRNA genes (5S, 16S, 23S) formed a putative operon on a single contig. Based on megablast hits, they were identified as belonging to the “*Ca*. Bealeia paramacronuclearis” (*Holosporales*) symbiont previously characterised in the US_Bl 11III1 strain (Szokoli et al. 2016). The contig had comparable coverage (312.91) to the “*Ca*. Fokinia” ones, but significantly higher GC content (0.46).

Taking into account the plot and the rRNA gene features, we designed a “selection from preliminary assembly” for “*Ca.* Fokinia cryptica” , including all contigs with log_10_ of coverage higher than 1.6, and GC content higher than 0.3, with the exception of those having best megablast hit on members of order Peniculida (to with the *Paramecium* host belongs).

The reads mapping to this “selection from preliminary assembly” were extracted and reassembled separately with SPAdes (based on different attempts, the option -k 21,33,55,77,99,127 was chosen). This “SPAdes re-assembly” included 840 contigs (2,952,405 bp, N50=23,802 bp; L50=25). In this assembly, a selection was made of sequences belonging to “*Ca.* Fokinia cryptica” (15 contigs, 812,007 bp; Figure K).

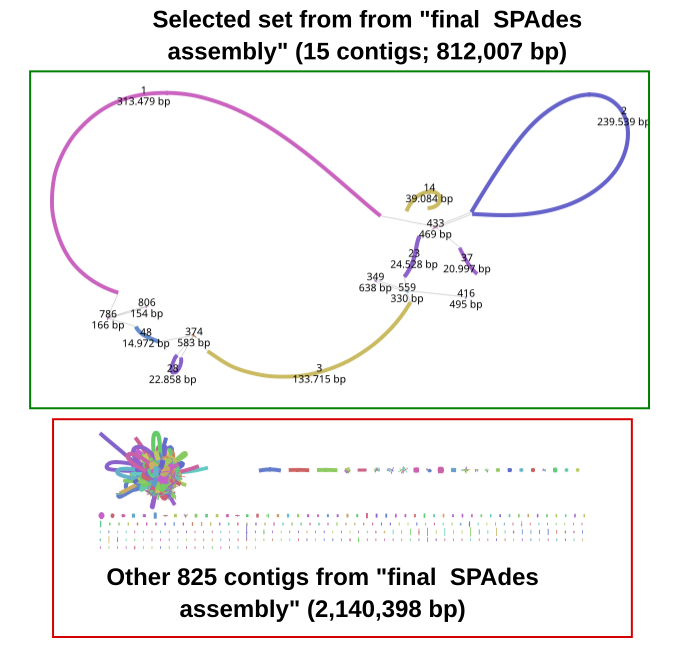

**Figure K** Visual representation of the 7695 contigs of the “final SPAdes assembly” and the relative connections in the assembly graph, obtained with the Bandage software. The green square highlights the selected set of contigs, the red square highlights the remaining contigs of the assembly.

Then, primers were designed next to the ends of the 8 larger contigs in the selected set, and PCR reactions were performed in order to identify/confirm junctions among those contigs. The combinations of primers used for amplification and sequencing, primer sequences, and the reaction protocols used are reported below.

| **Reaction** | **Amplification primer 1** | **Amplification primer 2** | **Sequencing primer 1** | **Sequencing primer 2** | **Protocol** |
| --- | --- | --- | --- | --- | --- |
| 1 | Fcry_1F | Fcry_48R_a | Fcry_1F | Fcry_48R_b | B |
| 2 | Fcry_48F_a | Fcry_28F_a | Fcry_48F_b | Fcry_28F_b | A |
| 3 | Fcry_28R_a | Fcry_3F_a | Fcry_28R_b | Fcry_3F_b | B |
| 4 | Fcry_3R | Fcry_23F | Fcry_3R | Fcry_23F | A |
| 5 | Fcry_23R_a | Fcry_14R_a | Fcry_23R_b | Fcry_14R_b | A |
| 6 | Fcry_37F_a | Fcry_2F_a | Fcry_37F_b | Fcry_2F_b | B |
| 7 | Fcry_2R_a | Fcry_1R | Fcry_2R_b | Fcry_1R | B |

| **Primer** | **Sequence (F: forward; R: reverse)** |
| --- | --- |
| Fcry_1F | 5'-GACACATCAACCGGAACG-3' |
| Fcry_48R_a | 5'-GTTATACATCAGAGAGAAGCAC-3' |
| Fcry_48R_b | 5'-TGATATGTTACGAGAGTGTGC-3' |
| Fcry_48F_a | 5'-TCTTTAGACTACCACAGAGTG-3' |
| Fcry_48F_b | 5'-AGCAGTTTGCTCAGCCTC-3' |
| Fcry_28F_a | 5'-GGTATCTTACGGGTCAAGC-3' |
| Fcry_28F_b | 5'-ATTGTGAAGCCGTAGTTGTAG-3' |
| Fcry_28R_a | 5'-ACATTATGATACGCTCAACGC-3' |
| Fcry_28R_b | 5'-TGGAGCATACGGAGTGAG-3' |
| Fcry_3F_a | 5'-AGATGTGCCAAGGATTCGC-3' |
| Fcry_3F_b | 5'-CCATTTCCGCCTTATTACCC-3' |
| Fcry_3R_b | 5'-TTATGACGAACGATACAACTAG-3' |
| Fcry_23F | 5'-CGATGTACGTTGTTCTCCC-3' |
| Fcry_23R_a | 5'-GAAATGGGAAGGCAGTAGC-3' |
| Fcry_23R_b | 5'-AGAAGTACTAGGAGCCCAC-3' |
| Fcry_14R_a | 5'-CATGAAAGACGTGTAACGGC-3' |
| Fcry_14R_b | 5'-GTTACTATAATGCCGCGAGG-3' |
| Fcry_14F_a | 5'-ATGTTAAGAGCAGCATCGTAC-3' |
| Fcry_14F_b | 5'-AAGAGGAACAAGAGACGAAC-3' |
| Fcry_37R_a | 5'-GTAGATGTATGGTGGCTTGC-3' |
| Fcry_37R_b | 5'-TTGGAGCAGTGAAGGGATG-3' |
| Fcry_37F_a | 5'-AACATCGCTATTGCCGGTG-3' |
| Fcry_37F_b | 5'-GAGTTACCACAATACCGATG-3' |
| Fcry_2F_a | 5'-AGGTGGTGACTCCTGTTG-3' |
| Fcry_2F_b | 5'-CTAAATAGCAGACGCGGTAG-3' |
| Fcry_2R_a | 5'-TCGCATAACACTGCATTACC-3' |
| Fcry_2R_b | 5'-ATCAAGTAGTGAAGCATCAGG-3' |
| Fcry_1R | 5'-CACAACAGAAGGAATCGCG-3' |

Protocol A (standard)

| 94°C | 3 min |  |
| --- | --- | --- |
| 94°C | 30 s | 40 cycles |
| 50°C | 30 s |  |
| 72°C | 4 min |  |
| 72°C | 10 min |  |

Protocol B (touchdown)

| 94°C | 3 min |  |
| --- | --- | --- |
| 94°C | 30 s | 5 cycles |
| 56°C | 30 s |  |
| 72°C | 4 min |  |
| 94°C | 30 s | 10 cycles |
| 53°C | 30 s |  |
| 72°C | 4 min |  |
| 94°C | 30 s | 30 cycles |
| 50°C | 30 s |  |
| 72°C | 4 min |  |
| 72°C | 10 min |  |

Thanks to such PCR experiments, it was possible to close the “*Ca*. Fokinia cryptica” genome into a single scaffold (815,586 bp).

***Paramecium biaurelia* USBL-36I1 hosting *Lyticum linuosum***

The “preliminary assembly” was constituted by 115,057 contigs (136,752,753 bp; N50 = 10,679 bp; L50 = 2209)

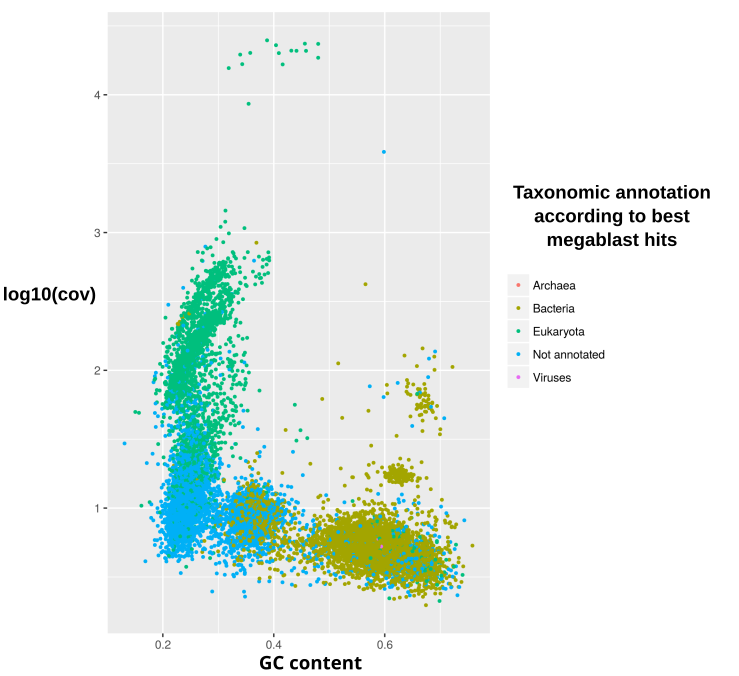

**Figure L** Plot of the preliminary contigs of the sample USBL-36I1 according to their GC content and log_10_ of sequencing coverage, coloured according to the respective best megablast hit. Only contigs with length higher than or equal to 1000 bp are shown for viewers’ clarity.

In total, 52 full-length or partial rRNA genes were identified in the “preliminary assembly” (Supplementary table 4):

- 27 complete or partial rRNA genes had best megablast hits on nuclear or mitochondrial *Paramecium* spp., and were thus assigned to the host. The respective contigs ranged from 71.83 to 20140.69 in sequencing coverage
- 2 rRNA gene sequences (16S, 23S) were identified as belonging to *Lyticum*, based on their best megablast hits on *L. sinuosum* (16S), and on the “*Ca*. Midichloriaceae” endosymbiont of *Acanthamoeba* sp. UWC8 (23S). The respective contigs displayed 258.45 and 843.75 sequencing coverage, and 0.37 and 0.25 GC content
- 23 remaining complete or partial rRNA gene sequences had megablast hits on other organisms (22) or no hit (1). The respective contigs had comparably lower coverage and/or clearly higher GC content (≥0.42) than those with *Lyticum* genes

Taking into account the plot and the rRNA gene features, we designed a “selection from preliminary assembly” for *Lyticum,* including all those contigs with log_10_ of coverage between 2 and 3, and GC content lower than 0.4.

The reads mapping to this “selection from preliminary assembly” were extracted and reassembled separately with SPAdes (based on different attempts, the option -k 21,33,55,77,99 was chosen). This “SPAdes re-assembly” included 7695 scaffolds (59,286,337 bp, N50=41,667 bp; L50=396). In this assembly, a selection was made of sequences belonging to *Lyticum* (30 scaffolds, 1,038,633 bp; Figure M).

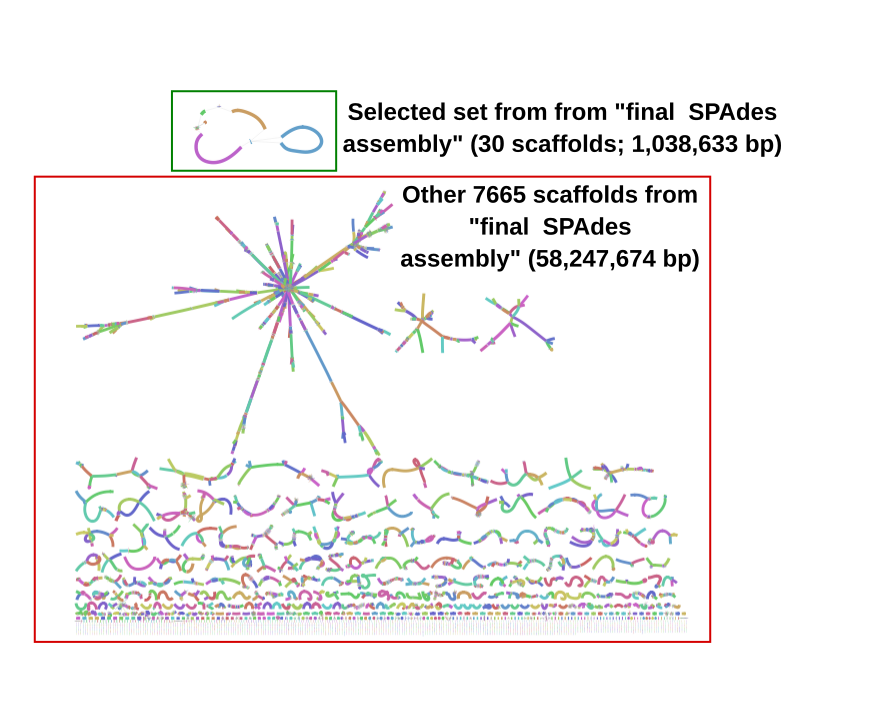

**Figure M** Visual representation of the 7695 scaffolds of the “final SPAdes assembly” and the relative connections in the assembly graph, obtained with the Bandage software. The green square highlights the selected set of scaffolds, the red square highlights the remaining scaffolds of the assembly.

Finally, the assembly was refined by removing all scaffolds <5000 bp containing no validated ORF (i.e. with no hits on NCBI nr or with hits only on hypothetical proteins). Thus, the “final assembly” of *Lyticum* counted 6 scaffolds (1,018,517 bp; N50=336,044 bp; L50=2).

***Plagiopyla* *frontata* IBS-3 hosting the novel “*Ca*. Midichloriaceae” bacterium “*Ca*. Vederia obscura”**

The “preliminary assembly” counted 49,376 contigs (66,833,832 bp; N50 = 33,240
 bp; L50 = 443).

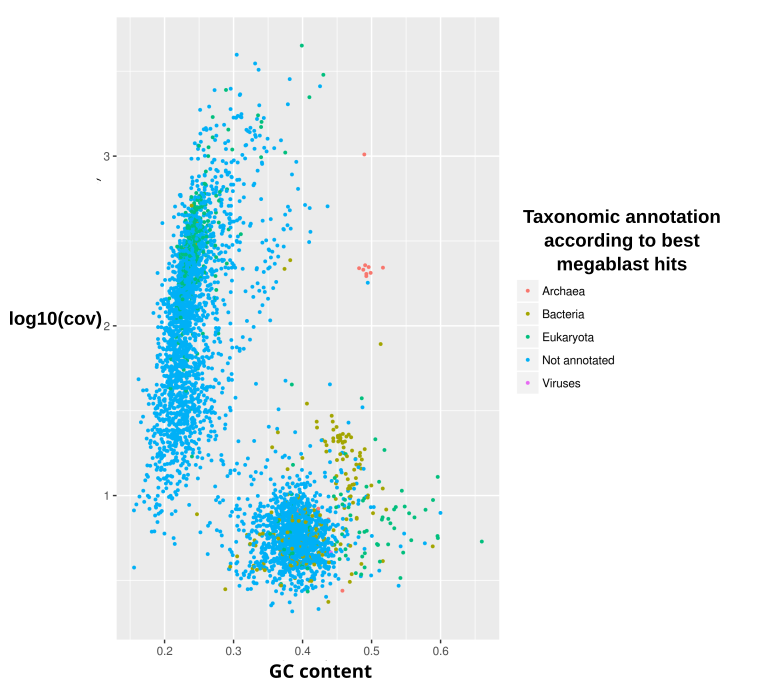

**Figure N** Plot of the preliminary contigs of the sample IBS-3 according to their GC content and log_10_ of sequencing coverage, coloured according to the respective best megablast hit. Only contigs with length higher than or equal to 1000 bp are shown for viewers’ clarity.

In total, 21 full-length or partial rRNA genes were identified in the “preliminary assembly” (Supplementary table 4):

- 1 partial LSU rRNA gene had best megablast hit on the ciliate *Cryptocaryon irritans*. This sequence was assigned to the host, considering the absence of LSU rRNA genes of *Plagiopyla* spp. on NCBI nucleotide. The respective contig had 3014.34 sequencing coverage
- 3 rRNA genes (5S, 16S, 23S) formed a putative operon on a single contig. They had best hits on *Methanocorpospulum* spp., and were thus assigned to a putative archaeal symbiont of *Plagiopyla*. The contig displayed 1022.82 sequencing coverage and 0.49 GC content
- 2 rRNA genes (16S, 23S) were identified as belonging to a novel “*Ca*. Midichloriaceae” bacterium from a new genus, based on their best megablast hits (with <90% identity) on the “*Ca*. Midichloriaceae” endosymbiont of *Acanthamoeba* sp. UWC8. The respective contigs displayed 243.72 and 216.54 sequencing coverage, and 0.38 and 0.37 GC content
- 15 remaining complete or partial rRNA gene sequences had megablast hits on other organisms. The respective contigs had comparably lower coverage (≤78.10) and higher GC (>0.40) than the ones assigned to the “*Ca*. Midichloriaceae” bacterium

Taking into account the plot and the rRNA gene features, we designed a “selection from preliminary assembly” for this “*Ca.* Midichloriaceae” bacterium (from now on, “*Ca*. Vederia obscura”), including all contigs with log_10_ of coverage higher than 1.5, excluding those having best megablast hits on ciliate (Ciliophora) sequences.

The reads mapping to “selection from preliminary assembly” were extracted and reassembled separately with SPAdes (based on different attempts, the option -k 21,33,55,77,99 was chosen). This “SPAdes re-assembly” included 3875 contigs (39,068,512 bp, N50=55,917 bp; L50=188). In this assembly, a selection was made of contigs belonging to *“Ca.* Vederia obscura” (35 contigs, 894,612 bp; Figure O).

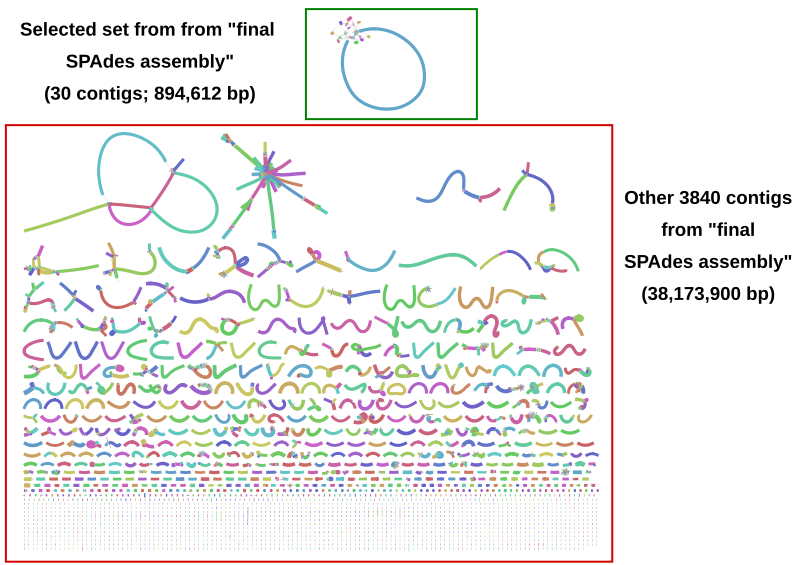

**Figure O** Visual representation of the 3875 contigs of the “final SPAdes assembly” and the relative connections in the assembly graph, obtained with the Bandage software. The green square highlights the selected set of contigs, the red square highlights the remaining contigs of the assembly.

Finally, the assembly was refined by removing all contigs <5000 bp containing no validated ORF (i.e. with no hits on NCBI nr or with hits only on hypothetical proteins. Thus, the “final assembly” of “*Ca*. Vederia obscura” counted 19 contigs (886,222 bp; N50=785,721 bp; L50=1).

***Reticulomyxa filosa* hosting a novel “*Ca*. Midichloriaceae” bacterium**

The “preliminary assembly” counted 25064 contigs (85,838,763 bp; N50 = 5,657
 bp; L50 = 4263).

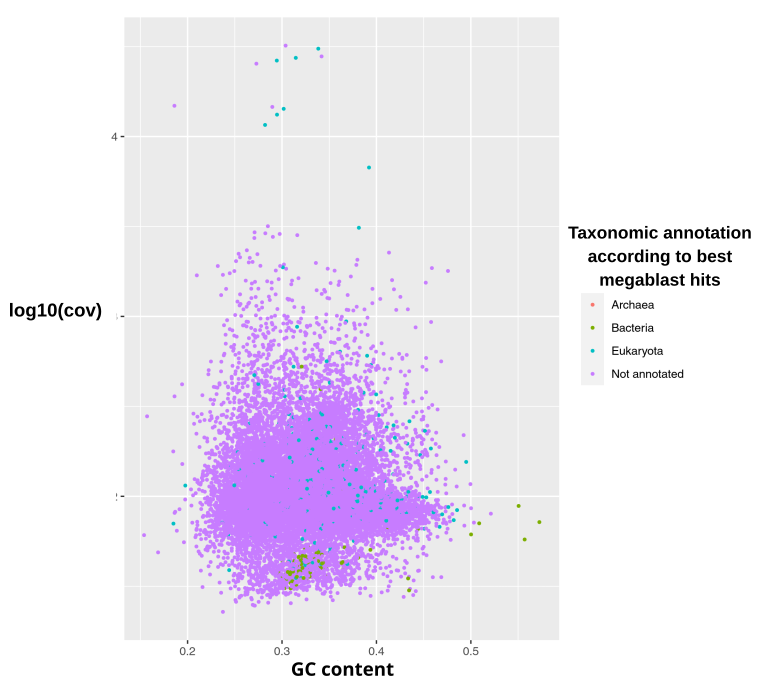

**Figure P** Plot of the preliminary contigs of the sample *Reticulomyxa* *filosa* according to their GC content and log_10_ of sequencing coverage, coloured according to the respective best megablast hit. Only contigs with length higher than or equal to 1000 bp are shown for viewers’ clarity.

In total, 11 full-length or partial rRNA genes were identified in the “preliminary assembly” (Supplementary table 4):

- 3 rRNA genes (16S, 23S, 5S) were identified as belonging to a novel “*Ca*. Midichloriaceae” bacterium from a new genus, based on their best megablast hits (with <94% identity) on the “*Ca*. Midichloriaceae” endosymbiont of *Acanthamoeba* sp. UWC8 and on “*Candidatus* Fokinia solitaria”. The respective contigs displayed 89.09 and 101.35 sequencing coverage, and 0.26 and 0.34 GC content
- 8 remaining complete or partial rRNA gene sequences had megablast hits on other organisms (7) or no hit (1). The respective contigs had comparable coverage (78.23-133.15) and GC (0.23-0.31) with the ones assigned to the “*Ca*. Midichloriaceae” bacterium

Taking into account the plot and the rRNA gene features, we designed a “selection from preliminary assembly” for this “*Ca.* Midichloriaceae” bacterium, including all contigs with log_10_ of coverage lower than 1.5. Moreover, the careful inspection of the DIAMOND hits of contigs with higher coverage revealed the presence of 61 additional contigs putatively belonging to the “*Ca.* Midichloriaceae” bacterium, which were thus individually added to the “selection from preliminary assembly”.

The reads mapping to such “selection from preliminary assembly” were extracted and reassembled separately with SPAdes (based on different attempts, the option -k 21,33,55,63 was chosen). This “SPAdes re-assembly” included 14367 contigs (33,864,241 bp, N50=6,105 bp; L50=1629). In this assembly, a selection was made of contigs belonging to the *“Ca. Midichloriaceae”* bacterium (498 contigs, 1,378,145 bp; Figure Q).

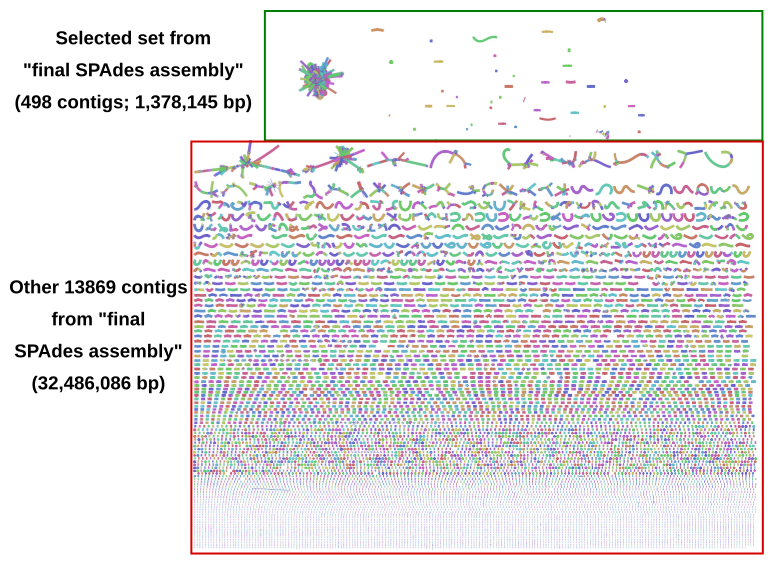

**Figure Q** Visual representation of the 14367 contigs of the “final SPAdes assembly” and the relative connections in the assembly graph, obtained with the Bandage software. The green square highlights the selected set of contigs, the red square highlights the remaining contigs of the assembly.

Finally, the assembly was refined by removing all contigs <1000 bp containing no validated ORF (i.e. with no hits on NCBI nr or with hits only on hypothetical proteins or transposases) Thus, the “final assembly” of the “*Ca*. Midichloriaceae bacterium” counted 242 contigs (1,305,893 bp; N50=7,721 bp; L50=53).

**References**

Andrews S. FastQC: a quality control tool for high throughput sequence data. Available online at: http://www.bioinformatics.babraham.ac.uk/projects/fastqc (2010).

Bankevich A, Nurk S, Antipov D, Gurevich AA, Dvorkin M, Kulikov AS, Lesin VM, Nikolenko SI, Pham S, Prjibelski AD, Pyshkin AV, Sirotkin AV, Vyahhi N, Tesler G, Alekseyev MA, Pevzner PA. SPAdes: A new genome assembly algorithm and its applications to single-cell sequencing. J Comp Biol 19: 455-477 (2012)

Boscaro V, Schrallhammer M, Benken KA, Krenek S, Szokoli F, Berendonk TU, Schweikert M, Verni F, Sabaneyeva EV, Petroni G. Rediscovering the genus *Lyticum*, multiflagellated symbionts of the order *Rickettsiales*. Sci. Rep. 3:3305 (2013)

Buchfink B, Xie C, Huson D. Fast and sensitive protein alignment using DIAMOND, Nature Methods 12: 59–60 (2015)

Castelli M, Sabaneyeva E, Lanzoni O, Lebedeva N, Floriano AM, Gaiarsa S, et al. *Deianiraea*, an extracellular bacterium associated with the ciliate *Paramecium*, suggests an alternative scenario for the evolution of *Rickettsiales*. ISME J 13: 2280-2294 (2019)
Castelli M, Lanzoni O, Nardi, T, Lometto S, Modeo L, Potekhin A, Sassera D, Petroni G. “*Candidatus* Sarmatiella mevalonica” endosymbiont of the ciliate *Paramecium* provides insights on evolutionary plasticity among *Rickettsiales*. Environ Microbiol. 23:1684-1701 (2021)

Catania F, Wurmser F, Potekhin AA, Przybos E, Lynch M. Genetic diversity in the *Paramecium* *aurelia* species complex. Mol Biol Evol. 26: 421-431 (2009)

De Coster W, D'Hert S, Schultz DT, Cruts M, Van Broeckhoven C. NanoPack: visualizing and processing long-read sequencing data. Bioinformatics 34: 2666-2669 (2018)

Floriano AM, Castelli M, Krenek S, Berendonk TU, Bazzocchi C, Petroni G, Sassera D. The genome sequence of “*Candidatus* Fokinia solitaria”: insights on reductive evolution in *Rickettsiales*. Genome Biol Evol. 10:1120–1126 (2018)

Hyatt D, Chen GL, Locascio PF, Land ML, Larimer FW, Hauser LJ. Prodigal: prokaryotic gene recognition and translation initiation site identification. BMC Bioinformatics 11:119 (2010)
Kawafune K, Hongoh Y, Hamaji T, Nozaki H. Molecular identification of rickettsial endosymbionts in the non-phagotrophic volvocalean green algae. PLoS One 7: e31749 (2012)

Kumar S, Jones M, Koutsovoulos G, Clarke M, Blaxter M. Blobology: exploring raw genome data for contaminants, symbionts and parasites using taxon-annotated GC-coverage plots. Front. Genet. 4: 237 (2013)

Langmead B, Salzberg S. Fast gapped-read alignment with Bowtie 2. Nat Methods 9: 357-359 (2012)

Lanzoni O, Sabaneyeva E, Modeo L, Castelli M, Lebedeva N, Verni F, Schrallhammer M, Potekhin A, Petroni G. Diversity and environmental distribution of the cosmopolitan endosymbiont “*Candidatus* Megaira”. Sci Rep 9: 1179 (2019)

Mironov T, Sabaneyeva E. A Robust symbiotic relationship between the ciliate *Paramecium* *multimicronucleatum* and the bacterium “*Ca*. Trichorickettsia mobilis”. Front Microbiol 11: 603335 (2020)

Nitla V, Serra V, Fokin SI, Modeo L, Verni F, Sandeep BV, Kalavati C, Petroni G. Critical revision of the family Plagiopylidae (Ciliophora: Plagiopylea), including the description of two novel species, *Plagiopyla* *ramani* and *Plagiopyla* *narasimhamurtii*, and redescription of *Plagiopyla* *nasuta* Stein, 1860 from India. Zool J Linn Soc 186:1-45 (2019)

Seemann T. barrnap 0.5 : rapid ribosomal RNA prediction. http://www.vicbioinformatics.com/ (2013)

Seemann T. Prokka: Rapid prokaryotic genome annotation. Bioinformatics 30, 2068-2069 (2014)

Senra MVX, Dias RJP, Castelli M, Silva-Neto ID, Verni F, Soares CAG, Petroni G. A house for two-double bacterial infection in *Euplotes woodruffi* Sq1 (Ciliophora, Euplotia) sampled in southeastern Brazil. Microb Ecol. 71: 505–517 (2016)

Simão FA, Waterhouse RM, Ioannidis P, Kriventseva EV, Zdobnov EM. BUSCO: assessing genome assembly and annotation completeness with single-copy orthologs. Bioinformatics 31, 3210–3212 (2015)

Szokoli F, Castelli M, Sabaneyeva E, Schrallhammer M, Krenek S, Doak TG, Berendonk TU, Petroni G*.* Disentangling the taxonomy of *Rickettsiales* and description of two novel symbionts (“*Candidatus* Bealeia paramacronuclearis” and “*Candidatus* Fokinia cryptica”) sharing the cytoplasm of the ciliate protist *Paramecium* *biaurelia*. Appl Environ Microbiol 82: 7236–7247 (2016)
Tarcz S, Potekhin A, Rautian M, Przyboś E. Variation in ribosomal and mitochondrial DNA sequences demonstrates the existence of intraspecific groups in *Paramecium multimicronucleatum* (Ciliophora, Oligohymenophorea). Mol Phylogenet Evol 63: 500-509 (2012)
Vannini C, Ferrantini F, Schleifer KH, Ludwig W, Verni F, Petroni G. “*Candidatus* Anadelfobacter veles” and “*Candidatus* Cyrtobacter comes,” two new *Rickettsiales* species hosted by the protist ciliate *Euplotes harpa (*Ciliophora, Spirotrichea). Appl Environ Microbiol. 76:4047–44054 (2010)

Wick RR, Schultz MB, Zobel J, Holt KE. Bandage: interactive visualization of *de novo* genome assemblies. Bioinformatics 31: 3350-3352 (2015)

Wick RR, Judd LM, Gorrie CL, Holt KE. Completing bacterial genome assemblies with multiplex MinION sequencing. Microb Genom. 3: e000132 (2017a)

Wick RR, Judd LM, Gorrie CL, Holt KE. Unicycler: Resolving bacterial genome assemblies from short and long sequencing reads. PLOS Comp Biol 13: e1005595 (2017b)

Wickham H. ggplot2: elegant graphics for data analysis. Springer-Verlag New York. ISBN 978-3-319-24277-4 (2016)
