## Supplementary material for "Host association and intracellularity evolved multiple times independently in the *Rickettsiales*": figures_and_supplementary_materials: Supplementary_text_3_metabolic_descriptions.docx

**Supplementary text 3: detailed comparative description of additional metabolic and functional features of *Rickettsiales***

A number of relevant traits/functions for the interaction with eukaryotic cells were selected for detailed analyses, described in the main text, in particular biosynthesis and uptake of metabolic precursors (i.e. amino acids and nucleotides), secretion/adhesion/motility apparatuses and putative effector molecules. A general overview of gene content variation and evolution in *Rickettsiales*, with a special focus on general “family-level” trends, as well as on the single newly characterised genomes, is presented below.

Housekeeping functions and DNA repair

Central housekeeping functions (DNA replication. cell division, general transcription, translation, chaperones) are conserved in all families and their representatives. The main DNA repair systems, namely mismatch repair (rare in *Gamibacteraceae* and *Deianiraeaceae*), nucleotide-excision repair, single- and double-strand break homologous repair/recombination, are present and overall common in all families. The most notable peculiarities in the novel genomes include the lack of nucleotide-excision and single-strand break repair systems in *Lyticum* and *Vederia*, and the split into two ORFs of *mutS* in *Bandiella*, possibly non-functional. Moreover, as previously observed in other unrelated *Rickettsiales* symbionts of ciliates(Floriano *et al.*, 2018; Castelli *et al.*, 2019, 2021), only the 5’-3’ exonuclease domain of DNA polymerase I is present in *Fokinia* *cryptica* and *Lyticum*.

Carbohydrate and energy metabolism

The repertoire for carbohydrate and energy metabolism is overall rich in *Rickettsiales*, but quite variable between and within families (especially in crown families). The “canonical” repertoire composed by central reversible core of gluconeogenesis and glycolysis (linking fructose 1-6-bisphosphate and phosphoenolpyruvate) plus some gluconeogenesis-specific enzymes (malate dehydrogenase, fructose 1-6 bisphosphatase, phosphoenolpyruvate synthase), alongside with non-oxidative pentose-phosphate pathway, pyruvate dehydrogenase, Krebs cycle, electron transport chain (including NADH-quinone oxidoreductase, qor type NADPH-quinone oxidoreductase, alternatively followed by ubiquinol-cytochrome c oxidoreductase+cytochrome c oxidase, and/or by the bd-cytochrome ubiquinol terminal oxidase), and ATP synthase are present in all families.

The lineage-specific and possible convergent streamlining of the glycolysis/gluconeogenesis is quite frequent, up to retaining only the initial steps (leading to phosphoenolpyruvate), as in many “classical” *Rickettsiaceae* (namely the monophyletic lineage comprising all characterised organisms, thereby with the exclusion of “basal *Rickettsiaceae*”, as previously defined(Castelli *et al.*, 2021)), including the novel genomes obtained in this study, but also occurring in other families, e.g. among *Midichloriaceae,* *Vederia* has the same subpathway, plus triose-phosphate isomerase. Pentose-phosphate pathway is missing in *Cyrtobacter*, and composed only by ribose-5-phosphate isomerase in *F. cryptica* and *Lyticum*. Moreover, genes belonging to the oxidative branch were found in basal *Rickettsiales*, in particular in *Mitibacteracae*.

Noteworthy is the case of cytochrome c oxidase, of which two distinct alternative forms are available, Cco or cbb3, the latter considered advantageous in anaerobic conditions(Pitcher and Watmough, 2004). Basal *Rickettsiales* present both forms, while in each crown family in general just one alternative is present, namely Cco in *Rickettsiaceae*, *Anaplasmataceae*, *Diomedesiaceae*, and cbb3 in *Midichloriaceae*, *Gamibacteraceae*, and neither in most *Deianiraceaeae* (lacking also ubiquinol-cytochrome c oxidoreductase, thus likely exploiting only their bd alternative oxidase). Such a pattern is clearly not consistent with phylogenetic relationships.

Multiple, probably independent, instances of combined reduction of Krebs cycle and electron transport chain were found. Besides some previously known cases(Floriano *et al.*, 2018; George *et al.*, 2020; Castelli *et al.*, 2021), among the newly sequenced *Midichloriaceae*, *Bandiella* lacks the bd oxidase, conversely *Cyrtobacter* lacks c reductase and oxidase, while both *F. cryptica* and *Lyticum* miss the (almost) entire Krebs cycle and possess only NADH-quinone and bd terminal oxidases. The latter two organisms probably exploit NADH from other sources for ATP synthesis(Floriano *et al.*, 2018), or only in case of *Lyticum*, the succinyl-CoA synthase. *Vederia* and the *Midichloriaceae* symbiont of *Reticulomyxa* are fully devoid of Krebs and electron transport, and possibly take ATP directly from their hosts or exploit alternative paths, such as a peculiar protein homologous to the glycolytic enzyme pyruvate kinase in *Vederia*.

As previously observed(Schön *et al.*, 2022), glyoxylate cycle is present in basal *Rickettsiales*, in particular *Mitibacteraceae*. Additionally, the phosphoenolpyruvate carboxylase, which forms oxaloacetate, thus potentially fuelling both Krebs and glyoxylate cycle, was found in basal *Rickettsiales* and in few crown *Rickettsiales* that are early diverging in the respective families, e.g. *Jidaibacter* and the *Rickettsiaceae* endosymbiont of *Amblyomma* Ac37b, while the nitrate reductase, which may enable the use of this compound as terminal electron acceptor(Moreno-Vivián *et al.*, 1999), was found in *Mitibacteraceae*, *Jistubacteraceae*, and *Gamibacteraceae*.

The main biosynthetic pathways overall mostly follow comparable presence/absence patterns among *Rickettsiales*. They are present in all families, and experienced multiple, probably independent, reductions or even full losses in single organisms/lineages in crown *Rickettsiales* families, as a possible consequence of the ability to obtain intermediates or final products directly from their hosts.

Lipids and membranes

The biosynthesis of lipids and phospholipids is well conserved among *Rickettsiales*, with exceptions including the pathway holes (lack of plsX and plsY) in *Rickettsia**(Driscoll et al., 2017)* and many classical *Rickettsiaceae* (including the newly sequenced genomes), but not in basal *Rickettsiaceae*. Concerning isoprenoid precursor biosynthesis, a previously proposed evolutionary scenario appears confirmed(Castelli *et al.*, 2021), proposing an ancestral and widespread methylerythritol pathway, and a replacement by a horizontally-acquired mevalonate pathway in few representatives of *Deianiraeaceae* and classical *Rickettsiaceae*, which include *Trichorickettsia* and the two symbionts of Apicomplexa(Hunter, Paight and Lane, 2020; Paight, Hunter and Lane, 2022), possibly indicating some link with Alveolata hosts.

Peptidoglycan biosynthesis likely experienced an independent reduction in different *Rickettsiales* sublineages for transpeptidase and transglycosylase activities, in particular *mrcB* gene is absent in *Deianiraeaceae*, *Anaplasmatacae* and many *Rickettsiaceae* (except basal members). Different further steps of streamlining can be observed in lineages such as *Orientia**(Min et al., 2008)* and multiple *Anaplasmataceae*, up to the condition of *Ehrlichia*, lacking multiple genes related to such activities (e.g. *fstI*, *mrdA*, *mrcA*). Lipopolysaccharide (LPS) biosynthesis is globally absent in *Anaplasmataceae*, and in some members of other crown families, such as *Orientia* and *Fokinia solitaria**(Min et al., 2008; Floriano et al., 2018)*, as well as *F.* *cryptica* and *Cyrtobacter*, while *Lyticum* is equipped with most LPS biosynthetic steps, with the exclusion of those producing deoxy-manno-octulosonate.

An interesting pattern is found for the synthesis of N-acetylglucosamine, precursor of both peptidoglycan and LPS in the classical *Rickettsiaceae.* The full pathway from fructose was present in the ancestor, with the last two steps catalysed by distinct polypeptides, due to a split of *glmU* gene, a situation conserved in *Megaira polyxenophila* and *Tisiphia*. Then, most of the pathway was lost independently in *Rickettsia**(Driscoll et al., 2017)* and *Megaira venefica*, both retaining only the N-terminus of *glmU*, catalysing the very last step.

The synthesis of polyhydroxyalkanoate (PHA) granules as storage compounds, present in all four basal families, is quite common in *Rickettsiaceae*, and found also in several *Midichloriaceae* (including all the novel genomes except *Bandiella*) and in few *Anaplasmataceae* from aquatic environments such as *Xenolissoclinum*.

Cofactors/vitamins

Biosynthetic pathways for cofactors/vitamins are quite common in *Rickettsiales*, differently from those for aminoacids and nucleotides (see below), though some lineages present reduction or complete absence of the pathways for one or more cofactors, with some degree of consistence with the lack of enzymes requiring them. Very common in all families are the synthesis of lipoate, ubiquinone (notable the exception of *Vederia*), and NADP from NAD (with *Lyticum* among few exceptions). Other quite frequent pathways are those for folate and heme (the latter absent in *Cyrtobacter*, *F. cryptica*, *Vederia* and endosymbiont of *Reticulomyxa*, consistently with the lack of cytochrome c reductase and oxidase).

The case of biotin is noteworthy, also considering a beneficial role hypothesised for this vitamin in some invertebrate hosts of *Rickettsiales*, and the multiple previous reports of putative horizontal-gene-transfer events involving genes of this biosynthetic pathway (Gillespie *et al.*, 2012; Nikoh *et al.*, 2014; Gerth and Bleidorn, 2016). The presence/absence of this pathway in *Rickettsiales* somehow inversely correlates with the one of the bioY biotin transporter, conversely present in representatives of genus *Wolbachia*, the *Midichloriaceae* endosymbiont of *Reticulomyxa*, as well as a group of closely related *Rickettsiaceae* (*Rickettsia*, *Tisiphia*, *Trichorickettsia*, *Megaira*, and symbiont of *Cardiosporidium cionae*). Some exceptions exist, e.g. *Trichorickettsia* and *Rickettsia* endosymbiont of *Ixodes* *scapularis* have both, while many other classical *Rickettsiaceae* have neither (e.g. *Orientia*, *Sarmatiella*, the latter possibly representing a recent loss of the biosynthesis(Castelli *et al.*, 2021)). Similarly, a single and short gene (*bioA*) is a possible sign of recent loss of the pathway also in *Bandiella*.

NAD synthesis is quite rare in *Rickettsiaceae* (exceptions include *Sarmatiella* and *Megaira* *venefica*), while in *Midichloriacae*, if present, it is represented only by the final two steps (among the novel genomes only in the *Reticulomyxa* endosymbiont). Similarly, FAD synthesis is quite rare among *Rickettsiaceae*, being found only in the basal representatives (such as endosymbionts of *Stachyamoeba* and of *Amblyomma* Ac37b), while in the novel *Midichloriaceae* genomes, except for *F.* *cryptica*, at least the last step is present. For what concerns pantothenate and CoA, the pathway is overall present in all families at variable levels of completeness, starting from the minimal final steps (coaDE), found in most representatives (among the novel genomes, the exception is *Lyticum*, while the richest is *F. cryptica*, from the bifunctional dfp on), up to including also the initial steps from 3-methyl-2-oxobutanoate, common only in the basal families and in *Gamibacteraceae*. Pyridoxal phosphate synthesis is quite rare in crown families, with earlier steps of the pathway being more common in *Rickettsiaceae* and *Midichloriaceae*, while differences can be seen through the order for the final steps, with most *Rickettsiales* (including *Anaplasmataceae*) displaying pdxJ and pdxH, while *Midichloriaceae* display mostly the alternative pdxT and pdxS, as well as the salvage pathway. The latter pathway is the only one present in *F. cryptica*, while *Vederia* is fully devoid of pathways for pyridoxal production/recycling. The presence of biosynthetic pathways for thiamine follows a comparable pattern, with initial steps (thiH) only present in *Mitibacteraceae*, successive steps found also in the other basal families, *Anaplasmataceae* and DDG clade (*Gamibacteraceae* and *Diomedesiaceae* in particular), and the final steps (thiD, thiE and thiL) rather common in all families, though rare in classical *Rickettsiaceae*. This pathway is uncommon in the novel genomes, and only its final step is present solely in *Vederia* and *Bandiella*. Additional cofactor biosynthetic pathways, such as molybdopterin and cobalamin are quite rare in *Rickettsiales* and found mostly/exclusively in basal families*,* namely both cofactors in *Mitibacteraceae*, and the former also in *Gamibacteraceae*.

Additional functions

Multiple *Rickettsiales* bear signs of the presence of different kinds of mobile elements, including plasmids (among which identified plasmidic contigs in published(El Karkouri *et al.*, 2016; George *et al.*, 2020) and in novel genomes, or plasmidic-like genes such as parA and toxin-antitoxins in other assemblies including e.g. *Vederia*, *Cyrtobacter* and *M. venefica*), phages (e.g. *Cyrtobacter*, the two *Megaira*, *Vederia*, and *Trichorickettsia*, the latter consistent with previous observations(Mironov and Sabaneyeva, 2020)), transposons (e.g. *Cyrtobacter*, the two *Megaira*, *Bandiella*), and retrotransposons (e.g *Bandiella* and *M. polyxenophila*).

Another noteworthy feature is the presence of cytoskeletal bactofilin proteins, such as the one previously described in *Deianiraea**(Castelli et al., 2019)*, in members of the three DDG-clade families, as well as in the basal *Rickettsiales*.

Interestingly, some detoxification and inorganic nutrient uptake proteins, previously found only in basal *Rickettsiales**(Schön et al., 2022)*, were also found in some crown lineages. Namely, KatG catalase/peroxidase and arsenic resistance mechanisms (efflux pump and/or reductase) were found in several *Gamibacteraceae* and even in *Diomedesiacae*, whereas ammonium transporters were found in *Diomedesiaceae*, *Gamibacteracaeae*, and a basal *Rickettsiaceae* MAG.

**References**

Castelli, M. *et al.* (2019) ‘Deianiraea, an extracellular bacterium associated with the ciliate Paramecium, suggests an alternative scenario for the evolution of Rickettsiales’, *The ISME journal*, 13(9), pp. 2280–2294.

Castelli, M. *et al.* (2021) ‘“Candidatus Sarmatiella mevalonica” endosymbiont of the ciliate Paramecium provides insights on evolutionary plasticity among Rickettsiales’, *Environmental microbiology*, 23(3), pp. 1684–1701.

Driscoll, T.P. *et al.* (2017) ‘Wholly ! Reconstructed Metabolic Profile of the Quintessential Bacterial Parasite of Eukaryotic Cells’, *mBio*, 8(5). doi:[10.1128/mBio.00859-17](http://dx.doi.org/10.1128/mBio.00859-17).

El Karkouri, K. *et al.* (2016) ‘Origin and Evolution of Rickettsial Plasmids’, *PloS one*, 11(2), p. e0147492.

Floriano, A.M. *et al.* (2018) ‘The Genome Sequence of “Candidatus Fokinia solitaria”: Insights on Reductive Evolution in Rickettsiales’, *Genome biology and evolution*, 10(4), pp. 1120–1126.

George, E.E. *et al.* (2020) ‘Highly Reduced Genomes of Protist Endosymbionts Show Evolutionary Convergence’, *Current biology: CB*, 30(5), pp. 925–933.e3.

Gerth, M. and Bleidorn, C. (2016) ‘Comparative genomics provides a timeframe for Wolbachia evolution and exposes a recent biotin synthesis operon transfer’, *Nature microbiology*, 2, p. 16241.

Gillespie, J.J. *et al.* (2012) ‘A Rickettsia genome overrun by mobile genetic elements provides insight into the acquisition of genes characteristic of an obligate intracellular lifestyle’, *Journal of bacteriology*, 194(2), pp. 376–394.

Hunter, E.S., Paight, C. and Lane, C.E. (2020) ‘Metabolic Contributions of an Alphaproteobacterial Endosymbiont in the Apicomplexan’, *Frontiers in microbiology*, 11, p. 580719.

Min, C.-K. *et al.* (2008) ‘Genome-based construction of the metabolic pathways of Orientia tsutsugamushi and comparative analysis within the Rickettsiales order’, *Comparative and functional genomics*, p. 623145.

Mironov, T. and Sabaneyeva, E. (2020) ‘A Robust Symbiotic Relationship Between the Ciliate and the Bacterium . Trichorickettsia Mobilis’, *Frontiers in microbiology*, 11, p. 603335.

Moreno-Vivián, C. *et al.* (1999) ‘Prokaryotic nitrate reduction: molecular properties and functional distinction among bacterial nitrate reductases’, *Journal of bacteriology*, 181(21), pp. 6573–6584.

Nikoh, N. *et al.* (2014) ‘Evolutionary origin of insect-Wolbachia nutritional mutualism’, *Proceedings of the National Academy of Sciences of the United States of America*, 111(28), pp. 10257–10262.

Paight, C., Hunter, E.S. and Lane, C.E. (2022) ‘Codependence of individuals in the Nephromyces species swarm requires heterospecific bacterial endosymbionts’, *Current biology: CB*, 32(13), pp. 2948–2955.e4.

Pitcher, R.S. and Watmough, N.J. (2004) ‘The bacterial cytochrome cbb3 oxidases’, *Biochimica et biophysica acta*, 1655(1-3), pp. 388–399.

[Schön, M.E. *et al.* (2022) ‘The evolutionary origin of host association in the Rickettsiales’, *Nature microbiology* [Preprint]. doi:](http://paperpile.com/b/w8D7dr/tS63Y)[10.1038/s41564-022-01169-x](http://dx.doi.org/10.1038/s41564-022-01169-x).
