## Supplementary material for "Host association and intracellularity evolved multiple times independently in the *Rickettsiales*": figures_and_supplementary_materials: Supplementary_text_4_phylogenomics.docx

**Supplementary text 3: Phylogenomics methods**

Phylogenomic analyses were aimed to get a comprehensive view on the evolution and diversity of *Rickettsiales*. To do so, we collected a representative dataset of *Rickettsiales*, including at least one representative per genus. We considered all available genomes and MAGs (metagenome-assembled genomes) belonging to “core” described *Rickettsiales* lineages as of February 2021 (i.e. the four families *Rickettsiaceae*, *Anaplasmataceae*, “*Ca*. Midichloriaceae”, “*Ca*. Deianiraeaceae”), and looked also for additional *Rickettsiales-*related organisms, namely any lineage forming a supported monophyletic group with *Rickettsiales* with the exclusion of other (alpha)proteobacterial orders. GenBank assemblies for the phylogenomics were downloaded from NCBI ftp (ftp.ncbi.nlm.nih.gov/genomes/all/GCA) initially in February 2021, and later updated in July 2021 (see below for details).

To reach such objective, the final species dataset for the phylogenomic analyses was built with a progressive refining procedure, described in detail below. The main steps were the following:

1. Collection of the initial dataset including *Rickettsiales* genomes, putative *Rickettsiales* MAGs, other *Alphaproteobacteria* and outgroup organisms (February 2021)
2. Expansion of the set of putative *Rickettsiales* MAGs, and 1^st^ filtering by assembly quality
3. Selection of orthologs for phylogenomics from *Rickettsiales* genomes, *Alphaproteobacteria*, and outgroup
4. Phylogeny-based identification of main MAG clades, and 2^nd^ filtering by phylogenetic relatedness
5. Phylogeny-based tests for the affiliation of each MAG clade to *Rickettsiales*
6. Creation of the final dataset composed by *Rickettsiales* genomes, confirmed *Rickettsiales* MAGs, other *Alphaproteobacteria,* and outgroup organisms, including newly available assemblies (July 2021)
7. Phylogenomic analyses on the final dataset to reconstruct the inner relationships among *Rickettsiales*

Step 1: Collection of the initial dataset including *Rickettsiales* genomes, putative *Rickettsiales* MAGs, other *Alphaproteobacteria* and outgroup organisms (February 2021)

The sequences of other 89 representative non-*Rickettsiales* *Alphaproteobacteria* (including *Magnetococcia* *sensu* Parks et al. 2018) as well as 8 *Gammaproteobacteria* and *Betaproteobacteria* as outgroup were downloaded, taking inspiration from the selection by Muñoz-Gómez and co-authors (2019).

All assemblies assigned by NCBI taxid to *Rickettsiales* (as of February 2021) were downloaded in the latest version, excluding anomalous assemblies. This set was manually edited to remove “*Ca*. Hepatobacter penaei” (GCA_000742475.1), actually belonging to *Holosporales*, and conversely to add the endosymbiont of *Acanthamoeba* UWC8 (Wang and Wu 2014), which, though labelled as member of *Holosporales*, is actually a close relative of the “*Ca*. Midichloriaceae” bacterium “*Ca*. Jidaibacter acanthamoeba” UWC36 (Schulz et al. 2016), thus ending up with 1679 assemblies. Out of those, 1164 were obtained in a large-scale study screening *Wolbachia* sequences from host genome sequencing projects (Scholz et al. 2020). We considered those *Wolbachia* assemblies as phylogenetically redundant for the divergence scale of the present phylogenomic dataset, and thus they were excluded from our analyses. The other assemblies were inspected in detail, and 401 of them, which were found to be derived from genome sequencing studies (in particular, though not exclusively, those labelled as described species or described genera), were treated as “*Rickettsiales* genome assemblies”, while the other 114 were treated as “putative *Rickettsiales* MAGs”. A manual selection was applied to the 401 “*Rickettsiales* genome assemblies”, to obtain a set accounting for at least one representative for each described genus, and in few cases (highly represented/diversified genera, e.g. *Rickettsia*, *Wolbachia*) more than one, resulting in 35 “selected *Rickettsiales* genomes”.

Step 2: Expansion of the set of putative *Rickettsiales* MAGs and 1^st^ filtering by assembly quality

The set of “putative *Rickettsiales* MAGs” was expanded in two ways. On one side, aiming to account for other deep-branching alphaproteobacterial MAGs (which might be potentially related to *Rickettsiales*), those identified in a previous study (Martjin et al. 2018), i.e. “MarineAlpha”, were downloaded. On the other side, in order to collect other potentially overlooked (e.g. due to mislabelling) lineages related to *Rickettsiales* or to “MarineAlpha”, the respective taxonomy of all previously identified assemblies was collected from gtdb r95 (Parks et al. 2022), and all other additional MAGs belonging to the same lineages (as well as closely related lineages forming monophyletic clades in the gtdb reference tree) were downloaded as well and added up to the previously selected ones, ending up with 294 total “extended putative *Rickettsiales* MAGs”. The quality of the extended MAG set was assessed with BUSCO 5.0.0 (Simão et al. 2015) with a search on 219 proteobacterial orthologs (“proteobacteria_odb10”) (Supplementary table 3). Based on a comparison with the “selected *Rickettsiales* genomes”, MAGs were retained only if having ≥50% single-copy orthologs and <5% duplicated orthologs, thereby discarding potentially poor quality/contaminated assemblies, obtaining 219 “BUSCO-filtered putative *Rickettsiales* MAGs”.

Step 3: Selection of orthologs for phylogenomics from *Rickettsiales* genomes, *Alphaproteobacteria*, and outgroup

For all “non-MAG genomes”, whenever available, protein sequences were taken directly from the NCBI annotation included in the assembly. In all other cases, i.e. non-annotated genomes (including newly obtained ones) and MAGs, genome sequences were annotated with Prokka 1.10 (Seemann 2014).

For all assemblies (including *Rickettsiales* genomes downloaded from NCBI and newly sequenced ones, MAGs, *Alphaproteobacteria* and outgroup) the eggNOG orthogroups (Huerta-Cepas et al. 2019) were then predicted on the annotated proteins with eggnog-mapper 2.0.6 (Cantalapiedra et al. 2021) with a taxonomic scope on *Alphaproteobacteria* (options: --tax_scope 28211 --go_evidence non-electronic --target_orthologs all --seed_ortholog_evalue 0.001 --seed_ortholog_score 60 --query_cover 20 --subject_cover 50).

Taking into account all assemblies except MAGs, proteins were grouped based on the assigned orthogroup, and each resulting orthogroup was aligned with MAFFT 7.475 L-INS-i (Katoh and Standley 2013) and trimmed with BMGE 1.12 (Criscuolo and Gribaldo 2010) with BLOSUM30 matrix. Single gene phylogenies were then performed with FastTree 2.1 (Price et al. 2010), and individually examined to remove paralogs and short/poorly aligned sequences. After this “polishing” step, orthogroups were kept only if present in single copy in at least 85% of the genomes of the “*Rickettsiales* genome set for phylogeny” (which, taking together “selected *Rickettsiales* genomes” and newly sequenced genomes, totals 44 organisms), in at least 85% of other *Alphaproteobacteria*, and in at least 50% of outgroup. Thus, by this procedure 179 orthogroups were selected, which were employed for all following phylogenetic analyses (Supplementary table 5).

Step 4: Phylogeny-based identification of main MAG clades, and 2^nd^ filtering of MAGs by phylogenetic relatedness

The genes assigned to the 179 selected orthogroups in the “BUSCO-filtered putative *Rickettsiales* MAGs” were added to those of the previous selection of organisms, each orthogroup was re-aligned and trimmed with MAFFT and BMGE, and single-gene phylogenies were inferred as described above. Each tree was manually inspected, for removing paralogs and short/poorly aligned sequences derived from the MAGs, ending up with the final sequence selection for those MAGs (Supplementary table 5). By this procedures 8 MAG were found to harbour <10 alphaproteobacterial orthogroups. Via online blastp searches on NCBI nr, it was found that (a significant part of) their proteome had much higher identities with non-alphaproteobacterial sequences, and were thus deemed as taxonomically mislabelled and removed from successive analyses, leaving 211 “BUSCO+eggNOG-filtered putative *Rickettsiales* MAGs” (Supplementary table 5).

Then, a first phylogenomic tree was inferred, with the aim to get a general view on the phylogenetic diversity of included MAGs, and on their relationships with *Rickettsiales* and other *Alphaproteobacteria*. Thus, the 179 orthogroup sequences of all the “BUSCO+eggNOG-filtered putative *Rickettsiales* MAGs” were used, together with a “Genome assembly set for MAG classification”, namely the other *Alphaproteobacteria*, outgroup organisms and, to speed up phylogenetic computations, a small and phylogenetically-balanced selection of 13 *Rickettsiales* genome assemblies (“*Ca*. Deianiraea vastatrix” for “*Ca*. Deianiraeaceae” and 4 for each other family). Each orthogroup was *de novo* aligned with MAFFT, and trimmed with BMGE, as described above, and all orthogroups were concatenated using AMAS (Borowiec 2016). Then, in order to avoid artefacts due to compositional heterogeneity in the datasets (in particular potential “erroneous” phylogenetic proximity of MAG lineages to “core” *Rickettsiales* due GC/AT biases in the respective genomes), the approach by Muñoz-Gómez and co-authors (2019) was applied, and the concatenated alignment was trimmed removing 10%, 20%, 30%, 40% or 50% of most heterogeneous sites. For each of these six alignments (the original and the five trimmed ones), maximum likelihood phylogenetic trees were inferred with IQ-TREE 1.6.12 (Nguyen et al. 2015) employing ModelFinder (Kalyaanamoorthy et al, 2017) for model selection, performing 1000 ultrafast bootstraps (Minh et al. 2013) and SH-aLRT with 1000 replicates.

In all the resulting trees, 35 MAGs were found consistently branching within “core” *Rickettsiales* with full support, and thus were directly treated as “confirmed *Rickettsiales* MAGs” (Supplementary figure 11). Other 2 MAGs were found consistently branching with high support within other alphaproteobacterial lineages (respectively, *Holosporales* and *Rhodospirillales*), and thus were considered as non-*Rickettsiales* and directly discarded. The other 174 MAGs were grouped into 13 groups (named A,B,C,D,E,F,G,H,I,J,K,L,M), based on monophyly observed across the obtained trees, with the aim to verify the position of each group with a dedicated phylogenomic dataset (see below, Step 5). Moreover, in order to speed up all the following analyses, phylogenetically-redundant MAGs (both within the 13 groups and within the already “confirmed *Rickettsiales* MAGs”) were removed as follows. Clusters of MAGs were defined based on supported monophyly and reciprocal AAI (Average Amino acid Identity) >0.85 (Rodriguez-R and Konstantinidis 2016; Supplementary figure 12), and for each cluster only the single MAG with best BUSCO scores (in terms of higher single-copy orthologs, or, secondarily, lower duplicated orthologs) (Supplementary table 3) was kept, thus resulting in 21 “confirmed *Rickettsiales* MAGs” and collectively 106 MAGs in the 13 groups.

Step 5: Phylogeny-based tests for the affiliation of each MAG clade to *Rickettsiales*

The MAGs belonging to each of the 13 previously identified groups were tested separately for the potential relatedness to “core” *Rickettsiales*, with a separate dataset. This was done in order to remove phylogenetic “noise” and potential artefacts caused by the contemporary addition of multiple deep-branching and fast-evolving lineages, and to get the “cleanest” phylogenetic position for the members of each group (such an approach has been previously utilized for similar problematic datasets, e.g. Otero-Bravo et al. 2018). Accordingly, for each group, the 179 ortholog sequences of the respective organisms were *de novo* aligned with those of the “Genome assembly set for MAG classification” using MAFFT, trimmed with BMGE, and all ortholgs were concatenated with AMAS, as described above. Each concatenated alignment was trimmed removing 10%, 20%, 30%, 40% or 50% of most heterogeneous sites, as described above. For each resulting alignment, maximum likelihood phylogeny was inferred with IQ-TREE and the LG+C60+F+R6 model, performing 1000 ultrafast bootstraps and SH-aLRT with 1000 replicates (Supplementary figure 13).

All the resulting trees were inspected, allowing the classification of 14 MAGs belonging to groups F, G, and H as additional, deep-branching “confirmed *Rickettsiales* MAGs”, based on strongly supported monophyly with “core” *Rickettsiales* on the original concatenated alignment and on all the five alignments with removal of heterogeneous sites. On the other side, none of the MAGs from the other 10 groups was found as monophyletic with *Rickettsiales* with high support in the respective trees, and were thus deemed as non-*Rickettsiales*, and not considered for the following analyses.

Step 6: Creation of the final dataset composed by *Rickettsiales* genomes, confirmed *Rickettsiales* MAGs, other *Alphaproteobacteria,* and outgroup organisms, including newly available assemblies (July 2021)

Then, the assembly and MAG selection was updated according to the latest NCBI version (July 2021), employing the same selection criteria as described above. Namely, the newly published genome of “*Ca*. Echinorickettsia raffii” (Carrier et al. 2021) was directly added to the “*Rickettsiales* genome set for phylogeny”, while 100 “new putative *Rickettsiales* MAGs” were processed as described above and 95 of them passed the BUSCO-based filtering by assembly quality (Supplementary table 3). In order to test the evolutionary relatedness of these novel MAGs to *Rickettsiales*, similarly to what described above, after assignment of their genes to the 179 orthogroups, the respective sequences were newly aligned together with those of the “Genome assembly set for MAG classification” and of the previously identified deep-branching *Rickettiales* lineages (now treated as ascertained *Rickettsiales*, similarly to the “core” lineages). Aligned genes were trimmed and concatenated, and the concatenated alignment was trimmed to remove more compositionally heterogeneous sites, as described above. For each resulting concatenated alignment, maximum-likelihood phylogeny was inferred with IQ-TREE employing ModelFinder for model selection, as described above (Supplementary figure 14). In all the resulting trees, 77 MAGs were found consistently branching within “core” or “deep-branching” *Rickettsiales* with full support, and thus were directly added to the previously identified “confirmed *Rickettsiales* MAGs”. The other 18 MAGs were found consistently and with high support branching within other alphaproteobacterial lineages (respectively, 1 *Sphingomonadales*, 2 *Rhizobiales*, and 15 *Holosporales*), and thus were considered as non-*Rickettsiales* and discarded. No MAG fell outside previously identified lineages, so no additional phylogeny-based test was required.

In order to remove phylogenetically redundant genomes, among all “confirmed *Rickettsiales* MAGs”, including those derived from the July 2021, updated clusters were again defined based on monophyly and reciprocal AAI >0.85 (Supplementary figure 12) As described above, for each cluster just the MAG with the best BUSCO scores was kept (Supplementary table 3).

Moreover, few MAGs were found to be closely related to some representatives “*Rickettsiales* genome set for phylogeny”. Thus, based on the monophyly and AAI criteria described above, among those the additional “phylogenetically-redundant” MAGs were removed (Supplementary figure 12).

Therefore, the final “*Rickettsiales* selection for phylogeny” comprised 45 genome assemblies and 68 MAGs, and, together with the 89 other representative *Alphaproteobacteria* and 8 outgroup organisms, formed the “final species selection for phylogeny” (210 total organisms).

Step 7: Phylogenomic analyses on the final dataset to reconstruct the inner relationships among *Rickettsiales*

The gene sequences of the 179 orthogroups for the 210 taxa of the final “*Rickettsiales* selection for phylogeny” were aligned, trimmed, and concatenated as described above. The phylogeny on this final dataset was aimed to reconstruct accurately the inner relationships among core and deep-branching *Rickettsiales* organisms, and at the same time, the relationship of *Rickettsiales* with other *Alphaproteobacteria*, building up the basis for the following gene-content analyses. It is quite well-ascertained that compositional heterogeneity leads to repeatable biased phylogenetic reconstructions among *Alphaproteobacteria*, producing artefactual grouping of *Rickettsiales* with other unrelated AT-rich lineages, in particular *Holosporales* and “*Ca*. Pelagibacterales”. Such artefacts can be resolved by the removal of the most compositionally heterogeneous sites (e.g. Viklund et al. 2012; Muñoz-Gómez et al. 2019). However, removal of such sites leads in parallel to the loss of a considerable amount of phylogenetic information (Fan et al. 2020), likely important for the reconstruction of the inner relationships within *Rickettsiales.* Therefore, the following approach was applied. The full alignment was used for phylogenetic reconstruction, with a guide tree indicating the ascertained relationships between *Rickettsiales* and other alphaproteobacterial lineages (-g option in IQ-TREE), so that only trees consistent with the included bipartitions were considered by the software in a partially constrained tree search. In detail, the guide tree included only a “minimal” set of just two constrained bipartitions (as based on the results by Muñoz-Gómez et al. 2019), namely a branch separating *Holosporales*+*Rhodospirillales* from all other organisms in the dataset, and another one separating “*Ca*. Pelagibacterales” + *Rhizobiales* + *Rhodobacterales* + *Caulobacterales* from all other organisms in the dataset, while leaving freely unconstrained tree search for all other possible (bi)partitions of the tree, in particular for what concerns the inner relationships within *Rickettsiales*. Maximum likelihood phylogeny was inferred with IQ-TREE employing the LG+C60+F+R6 model, performing 1000 ultrafast bootstraps and SH-aLRT with 1000 replicates (Supplementary figure 1).

**References**

Borowiec ML. AMAS: a fast tool for alignment manipulation and computing of summary statistics. PeerJ 4:e1660 (2016)

Cantalapiedra CP, Hernández-Plaza A, Letunic I, Bork P, Huerta-Cepas J. eggNOG-mapper v2: Functional annotation, orthology assignments, and domain prediction at the metagenomic scale. Mol Biol Evol: msab293 (2021)

Carrier TJ, Leigh BA, Deaker DJ, Devens HR, Wray GA, Bordenstein SR, Byrne M, Reitzel AM. Microbiome reduction and endosymbiont gain from a switch in sea urchin life history. Proc Natl Acad Sci U S A 118:e2022023118 (2021)

Criscuolo A, Gribaldo S. BMGE (Block Mapping and Gathering with Entropy): a new software for selection of phylogenetic informative regions from multiple sequence alignments. BMC Evol. Biol. 10:210 (2010)

Fan L, Wu D, Goremykin V, Xiao J, Xu Y, Garg S, Zhang C, Martin WF, Zhu R. Phylogenetic analyses with systematic taxon sampling show that mitochondria branch within *Alphaproteobacteria*. Nat Ecol Evol 4:1213–1219 (2020)

Huerta-Cepas J, Szklarczyk D, Heller D, Hernández-Plaza A, Forslund SK, Cook H, Mende DR, Letunic I, Rattei T, Jensen LJ, von Mering C, Bork P. eggNOG 5.0: a hierarchical, functionally and phylogenetically annotated orthology resource based on 5090 organisms and 2502 viruses. Nucleic Acids Res. 47:D309-D314 (2019)
Kalyaanamoorthy S., Minh B., Wong T., von Haeseler A, Jermin LS*.* ModelFinder: fast model selection for accurate phylogenetic estimates. Nat Methods 14:587–589 (2017)
Katoh K, Standley DM. MAFFT multiple sequence alignment software version 7: improvements in performance and usability. Mol Biol Evol. 30:772-80 (2013)

Lartillot N, Philippe H. A Bayesian mixture model for across-site heterogeneities in the amino-acid replacement process. Mol Biol Evol. 21:1095-109 (2004)

Martijn J, Vosseberg J, Guy L, Offre P, Ettema TJG. Deep mitochondrial origin outside the sampled alphaproteobacteria. Nature 557:101–105 (2018)

Minh BQ, Nguyen MA, von Haeseler A. Ultrafast approximation for phylogenetic bootstrap. Mol Biol Evol 30:1188-95 (2013)

Muñoz-Gómez S, Hess S, Burger G, Lang BF, Susko E, Slamovits CH, Roger A. An updated phylogeny of the *Alphaproteobacteria* reveals that the parasitic *Rickettsiales* and *Holosporales* have independent origins. eLife 8:e42535 (2019)

Nguyen LT, Schmidt HA, von Haeseler A, Minh BQ. IQ-TREE: a fast and effective stochastic algorithm for estimating maximum-likelihood phylogenies. Mol Biol Evol. 32:268-274 (2015)

Otero-Bravo A, Goffredi S, Sabree ZL. Cladogenesis and genomic streamlining in extracellular endosymbionts of tropical stink bugs. Genome Biol Evol 10: 680–693 (2018)

Parks DH, Chuvochina M, Waite DW, Rinke C, Skarshewski A, Chaumeil PA, Hugenholtz P. A standardized bacterial taxonomy based on genome phylogeny substantially revises the tree of life. Nat Biotechnol 36:996-1004 (2018)

Parks DH, Chuvochina M, Rinke C, Mussig AJ, Chaumeil PA, Hugenholtz H. GTDB: an ongoing census of bacterial and archaeal diversity through a phylogenetically consistent, rank normalized and complete genome-based taxonomy. Nucl Acids Res 50: D785–D794 (2022)

Price MN, Dehal PS, Arkin AP. FastTree 2--approximately maximum-likelihood trees for large alignments. PLoS One 5:e9490 (2010)

Rodriguez-R LM, Konstantinidis KT. The enveomics collection: a toolbox for specialized analyses of microbial genomes and metagenomes. PeerJ Preprints 4:e1900v1 (2016).

Scholz M, Albanese D, Tuohy K, Donati C, Segata N, Rota-Stabelli O. Large scale genome reconstructions illuminate *Wolbachia* evolution. Nat Commun 11:5235. (2020)

Schulz F, Martijn J, Wascher F, Lagkouvardos I, Kostanjšek R, Ettema TJG, Horn M. A *Rickettsiales* symbiont of amoebae with ancient features. Environ Microbiol. 18:2326–2342 (2016)

Seemann T. Prokka: Rapid prokaryotic genome annotation. Bioinformatics 30: 2068-2069 (2014)

Simão FA, Waterhouse RM, Ioannidis P, Kriventseva EV, Zdobnov EM. BUSCO: assessing genome assembly and annotation completeness with single-copy orthologs. Bioinformatics 31, 3210–3212 (2015)

Viklund J, Ettema TJ, Andersson SG. Independent genome reduction and phylogenetic reclassification of the oceanic SAR11 clade. Mol Biol Evol. 29:599-615 (2012)
Wang Z, Wu M. Complete genome sequence of the endosymbiont of *Acanthamoeba* strain UWC8, an amoeba endosymbiont belonging to the “*Candidatus* Midichloriaceae” family in *Rickettsiales*. Genome Announc. 2:e00791–14 (2014)
