## Supplementary material for "Host association and intracellularity evolved multiple times independently in the *Rickettsiales*": figures_and_supplementary_materials: Supplementary_text_5_gene_content_analyses.docx

**Supplementary text 4: gene content analyses**

**Annotation**

The newly obtained genomes were all annotated with Prokka 1.10 (Seemann 2014), using the --rfam option. Afterwards, annotation of the genomes of ciliate symbionts was manually curated by a detailed inspection of blastp hits on NCBI nr and on *Rickettsiales* proteins as described previously (Castelli et al. 2019).

**Creation of a set of orthogroups for gene content comparisons**

In order to perform gene content comparisons and reconstruct its variation along the inferred species tree, a set of orthogroups was obtained for the 210 total organisms in the final phylogeny, starting from the previously obtained eggNOG orthogroups (see Supplementary text 4).

The eggNOG (Huerta-Cepas et al. 2019) database is organised by taxonomy, namely a dedicated set of orthogroups is identified for each lineage from the domain to the order/family level (domain, phylum, class, order, family), and orthogroups are hierarchically linked. For example 2TSRZ@28211 (DNA sliding clamp in *Alphaproteobacteria –* NCBI taxid 28211) is linked to corresponding orthogroups in higher taxonomic ranks, such as 1MVD9@1224 (same function in *Proteobacteria* – NCBI taxid 1224) and COG0592@2 (same function in *Bacteria* – NCBI taxid 2). Such hierarchical organisation allows to take advantage of lineage-specific annotations, such as labelling of lineage-specific paralogs, and at the same time gives the possibility to compare orthogroups identified in different lineages.

However, we noticed that, with the dataset of this study, this system could prevent the identification of possible orthologs. An extreme example is that of some ribosomal protein genes in “*Candidatus* Aquarickettsia rohweri” (Klinges et al. 2019), namely S10 (RST68169.1), L3 (RST68170.1), L10 (RST68925.1), L14 (RST68179.1), L24 (RST68180.1), which have their best hits in eggNOG orthogroups assigned to lineages as distant as Metazoa (3C1KH@33208, 3BT1X@33208, 3C01C@33208), *Acyclobacillaceae* (*Firmicutes*), and *Sphingomonadales* (2K3XB@204457). This condition is likely due, at least partly, to the high sequence evolution rates in *Rickettsiales*, which may result in extremely high sequence divergence with respect to any other organism (including other *Rickettsiales*), so that hits can be almost equivalently distant from closely related and unrelated lineages (Castelli et al. 2019), even for housekeeping genes.

The specific analysis pipeline of this work, in the following steps (described below), is specifically intended to reconstruct ancestral states of gene presence/absence, including identification of horizontal gene transfer events, employing a phylogeny-based approach, more powerful and suitable for this purpose than an eggNOG assignment. Thus, in this step we aimed to identify the largest groups of (at least potentially) orthologous genes, including such “ambiguous” situations, to be tested directly later. For this purpose, i.e. circumventing the assignment issue described above and at the same time retaining as much lineage specific information as possible from the eggnog database, we designed the following two-step “telescopic” approach, taking advantage of the hierarchical structure of the eggNOG database:

- We started from the eggnog taxonomic path leading to *Rickettsiales* (root; *Bacteria*; *Proteobacteria*; *Alphaproteobacteria*; *Rickettsiales*). When a gene was assigned as best match to an orthogroup belonging to a taxon outside such path, it was moved “upwards” to the respectively proximate higher rank. Such passage was applied recursively until a taxon along the *Rickettsiales* path was met. This allowed also the “connection” of orthologs in the non-*Rickettsiales* organisms of the dataset. For example, if a gene was assigned to the 2JQAW@204441 orthogroup (DNA clamp in *Rhodospirillales* – NCBI taxid 204441), it was labelled as the corresponding alphaproteobacterial orthogroup 2TSRZ@28211
- The second step was intended to “connect” the information between different orthogroups, namely, whenever appropriate, joining taxonomically-linked orthogroups at the respectively highest taxonomic rank. In detail for each orthogroup at the “root” level resulting from the previous step, all genes assigned to orthogroups of lower taxonomic ranks (*Bacteria*; *Proteobacteria*; *Alphaproteobacteria*; *Rickettsiales*) that were linked by the eggnog hierarchy to the given root orthogroup were moved “upwards” to such root orthogroup. For example, if a gene was assigned to the root DNA clamp COG0592@1, any other gene assigned to the linked 2TSRZ@28211 (*Alphaproteobacteria*) or 1MVD9@1224 (*Proteobacteria*) was moved “upwards” to the COG0592@1 orthogroup. This passage was performed progressively for remaining orthogroups of lower ranks. It was thus possible to “merge” genes into larger meaningful orthogroups, while still keeping as much as possible lineage-specific refined annotation.

Applying such “telescopic” approach, a total of 444,226 genes were assigned to 20,041 orthogroups. Only 4009 of such orthogroups were present in at least one member of *Rickettsiales* (“core” or “deep-branching”), and only 2990 of those were present in 4 or more organisms of the total dataset, and were thus considered for following gene content and phylogenetic analyses.

**Reconstruction of ancestral states of gene copy number**

Reconstruction of ancestral states (in terms of gene copy number in each orthogroup) was performed by a tree-reconciliation approach, employing ALE 0.4 (Szöllősi et al. 2013). In detail, each of the 2990 orthogroups was aligned with MAFFT 7.475 L-INS-I (Katoh and Standley 2013) and trimmed with BMGE 1.12 (Criscuolo and Gribaldo 2010) with BLOSUM30 matrix. Then, aligned orthogroups shorter than 30 amino acids after trimming were removed, ending with 2871 orthogroups. For each of those, a sample of gene trees was obtained running 10000 iterations with PhyloBayes 4.1 (Lartillot and Philippe 2004). Then, amalgamated likelihood estimation was performed with ALEobserve, run with 10% burn-in, followed by ALEml_undated (Szöllősi et al. 2015).

In order to get a comprehensive view of general evolutionary trends in terms of copy number variations in *Rickettsiales* and of the underlying events (losses, duplications, origins, transfers) predicted by ALEml_undated, for each node of the species tree predicted events and predicted (or observed at tips) copy numbers were were rounded up or down (using a 0.3 threshold as in Martijn et al. 2020), then summed, considering the whole dataset of 2871 orthogroups, as well as separately for each eggNOG functional category (Supplementary figure 2).

Based on the inspection of such general and orthogroup-specific results of the ALE-reconstructed events, further in-depth and more refined analyses were performed on selected functions, as described in the following sections.

**Phylogenetic analyses on biosynthetic pathways for amino acids and nucleotides**

Reference biosynthetic pathways for amino acids and nucleotides were obtained from the Biocyc database (Karp et al. 2019). In order to get more robust and reliable phylogenetic inferences, we aimed to concatenate together sequences of the proteins involved in the same pathway, thus obtaining more phylogenetically informative datasets. For the same purpose, pathways sharing common reactions (e.g. purines, pyrimidines, branched-chain amino acids, aromatic amino acids) were considered together (Supplementary table 6). Under similar assumptions, single (or two) gene pathways, namely those for glycine, aspartate, asparagine, glutamate, glutamine, and alanine, were not considered phylogenetically informative enough, and were thus not employed in the analyses.

For each gene involved in a chosen pathway we selected an eggNOG COG, blasting the selected Biocyc reference sequence against the protein sequences from all the 210 selected taxa and manually inspecting the results. Then, the composition of each orthogroup was manually refined, by visual inspection of the respective alignment and of a maximum-likelihood tree obtained with IQ-TREE 1.6.12 (Nguyen et al. 2015) employing ModelFinder (Kalyaanamoorthy et al, 2017) for model selection, and performing 1000 ultrafast bootstraps (Minh et al. 2013) and SH-aLRT with 1000 replicates. In particular, short and/or poorly aligned sequences and putative paralogs were manually removed, especially among non-*Rickettsiales* organisms, and, in some cases (e.g. biosynthetic genes for histidine) single domains from alternative fusions of multi-domain proteins were selected and analysed separately. Each manually refined orthogroup was then re-aligned with MAFFT 7.475 L-INS-i and trimmed with BMGE 1.12 with the BLOSUM30 matrix. Then, orthogroups belonging to the same pathway (or same group of related pathways) were concatenated together using AMAS (Borowiec 2016).

In many cases, we noticed that some organisms, including members of *Rickettsiales*, displayed only few genes of a given pathway, and among those, the last enzymatic reaction(s) leading to the final product were frequently missing. All those cases could be indicative of non-specific roles of the identified proteins, thus representing possible “false positives” for the presence of the respective pathway. Alternatively, the apparently missing steps might be “filled” by additional non-specific enzymes, as hypothesised for some *Rickettsiales* (Driscoll et al. 2017). It must also be considered that, if a significant proportion of the total genes of a pathway was missing in a given organism, this would lead to a proportionally significant loss of phylogenetic information (and also to an “unbalanced” availability of sites among the included organisms), which could hamper the accuracy of the phylogenetic inference. Taking into account all the above, for each concatenated alignment we opted for two alternative strategies in parallel:

- “full organism dataset”, keeping all organisms
- “selected organism dataset”: keeping only those organisms displaying at least 50% of the included genes, or, alternatively, at least a significant proportion of selected sub-branches of the pathway (see Supplementary figure 6 for details on each pathway)

Then, in order to account for compositional heterogeneity (see Supplementary text 4), each of the two alignments was separately processed for removing 10%, 20%, 30%, 40% or 50% of the most compositionally heterogeneous sites, following the approach by Muñoz-Gómez and co-authors (2019), thus ending up with 12 alignments for each pathway. On each alignment, phylogeny was inferred with IQ-TREE and the LG+C60+F+R6 model, performing 1000 ultrafast bootstraps and SH-aLRT with 1000 replicates (Supplementary figure 8).

**Identification of amino acid transporters**

Amino acids transporters were identified from the TCDB database (Saier et al. 2021). The transporter families in TCDB were queried by keywords such as “amino acid(s)” and the name of each of the twenty proteinogenic amino acids. The single database entries (i.e. proteins) belonging to each member of the retrieved families were individually examined, in particular information on the kind of substrate (if known) were collected, and those members with ascertained substrate(s) that did not include amino acids were discarded, ending up with a list of trusted and substrate-labelled entries of putative amino acid transporters from the TCDB database. We then blasted all the *Rickettsiales* proteins of our dataset against the trusted entries, and for each genome the number of proteins having a best significant hit (e-value threshold of 1e-5) on each trusted entry were counted. However, we realized this approach led to a number of false positive hits by distant paralogous sequences. While a hypothetical solution could have been to increase blastp threshold, at the same time this could have led to false negatives (i.e. preventing the identification of “good” hits with a putatively similar substrate specificity with respect to the database sequence), considering high sequence evolutionary rates and consequent sequence divergence in *Rickettsiales*. Therefore, to solve the issue, we redid the blast search, but in this case all the *Rickettsiales* proteins of our dataset were blasted on the full TCDB database, and for each genome only the proteins having a best significant hit (e-value threshold of 1e-5) on each trusted entry were selected, and counted (Supplementary figure 7,10).

**Identification and phylogenetic analysis of the tlc nucleotide translocasaes**

Analyses on nucleotide transporters were focused on the tlc nucleotide translocase transporters, common in *Rickettsiales* and in other host-associated lineages (Major et al. 2017). Starting from the eggNOG orthogroup classification, a blastp-based inspection using the protein sequences from the paper by Major and co-authors (2017) as queries led to the identification of the corresponding orthogroup assigned to tlc translocases (COG3202@1), which allowed to count the number of genes present for each organism of our dataset. Then, the phylogenetic dataset of tlc translocases by Major and co-authors (2017) was downloaded. Considering the large size of this dataset, a subset was selected, corresponding to a clade of sequences consisting only of the nucleotide transport protein domain and belonging to multiple bacteria (including *Chlamydiae* and several *Proteobacteria* such as *Rickettsiales* and *Holosporales*), as well as to eukaryotes such as Microsporidia, Stramenopiles, and Viridiplantae. The selected sequences from the published dataset and those identified in our dataset (COG3202@1) were all aligned together with MAFFT 7.475 L-INS-i and trimmed with BMGE 1.12 with the BLOSUM30 matrix, and phylogeny was inferred as in (Major et al. 2017), namely with IQ-TREE with the LG+C60 model, performing 1000 ultrafast bootstraps and SH-aLRT with 1000 replicates (Supplementary figure 9).

**Identification of genes involved in the interaction with host cells**

For getting information on the presence and multiplicity of genes involved in multiple features of the interaction of *Rickettsiales* with host cells, such as secretion systems, putative toxins and other secreted effectors, flagella, adhesion and invasion molecules, the VFDB core reference database was employed (Liu et al. 2022- downloaded on 6^th^ May 2022). The VFDB is organised in order to have a separate focus on single pathogens, which implies a significant redundancy of entries of orthologous genes (e.g. flagellar genes) identified in each pathogen. This organisation would not be suitable for the analyses of our large dataset composed almost exclusively by non-model bacteria. Therefore, orthologs were identified within the database with the following approach. Sequences were divided by organism (i.e. single strain) and by VFC (VF Classes). Then, within each selected VFC (“Adherence”, “Biofilm”, “Effector delivery system”, “Exotoxin”, “Exoenzyme”, “Invasion”, “Motility”) orthogroups were identified with OrthoFinder 2.5.4 (Emms and Kelly 2019), employing blast as sequence search program. The composition of orthogroups produced by OrthoFinder was then manually inspected and curated. Then, all proteins of our dataset of *Rickettsiales* were blasted on the full VFDB core database. Similarly to the previous TCDB blastp search, the whole VFDB set of sequences was employed as database in order to minimise both false negatives and false positive hits. Then, for each *Rickettsiales* genome, proteins were “assigned” to each curated orthogroup as follows: a protein was counted if displaying a significant (evalue 1e-5) best hit on any sequence belonging to the orthogroup, and the total number of proteins having such hits were summed over each orthogroup (Supplementary figure 3,4,5,6).

**References**

Borowiec, M.L. AMAS: a fast tool for alignment manipulation and computing of summary statistics. PeerJ 4:e1660 (2016)

Castelli M, Sabaneyeva E, Lanzoni O, Lebedeva N, Floriano AM, Gaiarsa S, et al. *Deianiraea*, an extracellular bacterium associated with the ciliate *Paramecium*, suggests an alternative scenario for the evolution of *Rickettsiales*. ISME J 13, 2280-2294 (2019).

Criscuolo A, Gribaldo S, BMGE (Block Mapping and Gathering with Entropy): a new software for selection of phylogenetic informative regions from multiple sequence alignments, BMC Evol. Biol. 2010;10:210.

Driscoll TP, Verhoeve VI, Guillotte ML, Lehman SS, Rennoll SA, Beier-Sexton M, Rahman MS, Azad AF, Gillespie JJ. Wholly *Rickettsia*! Reconstructed metabolic profile of the quintessential bacterial parasite of eukaryotic cells. mBio 8: e00859-17 (2017)

Emms DM, Kelly S. OrthoFinder: phylogenetic orthology inference for comparative genomics. Genome Biol. 20:238 (2019)

Huerta-Cepas J, Szklarczyk D, Heller D, Hernández-Plaza A, Forslund SK, Cook H, Mende DR, Letunic I, Rattei T, Jensen LJ, von Mering C, Bork P. eggNOG 5.0: a hierarchical, functionally and phylogenetically annotated orthology resource based on 5090 organisms and 2502 viruses. Nucleic Acids Res. 2019 Jan 8;47(D1):D309-D314
Kalyaanamoorthy, S., Minh, B., Wong, T. *et al.* ModelFinder: fast model selection for accurate phylogenetic estimates. *Nat Methods* **14,** 587–589 (2017).
Karp PD, Billington R, Caspi R, Fulcher CA, Latendresse M, Kothari A, Keseler IM, Krummenacker M, Midford PE, Ong Q, Ong WK, Paley SM, Subhraveti P. The BioCyc collection of microbial genomes and metabolic pathways. Brief Bioinform. 20: 1085-1093 (2019)
Katoh K, Standley DM. MAFFT multiple sequence alignment software version 7: improvements in performance and usability. Mol Biol Evol. 30:772-80 (2013)
Klinges JG, Rosales SM, McMinds R, Shaver EC, Shantz AA, Peters EC, Eitel M, Wörheide G, Sharp KH, Burkepile DE, Silliman BR, Vega Thurber RL. Phylogenetic, genomic, and biogeographic characterization of a novel and ubiquitous marine invertebrate-associated Rickettsiales parasite, Candidatus Aquarickettsia rohweri, gen. nov., sp. nov. ISME J. 13:2938-2953 (2019)

Lartillot N, Philippe H. A Bayesian mixture model for across-site heterogeneities in the amino-acid replacement process. Mol Biol Evol. 21:1095-109 (2004)

Liu B, Zheng D, Zhou S, Chen L, Yang J. VFDB 2022: a general classification scheme for bacterial virulence factors. Nucleic Acids Res. 50: D912-D917 (2022)

Major P, Embley TM, Williams TA. Phylogenetic Diversity of NTT Nucleotide transport proteins in free-living and parasitic bacteria and eukaryotes. Genome Biol Evol. 9:480-487 (2017)

Martijn J, Schön ME, Lind AE, Vosseberg J, Williams TA, Spang A, Ettema TJG. Hikarchaeia demonstrate an intermediate stage in the methanogen-to-halophile transition. Nat Commun. 11:5490 (2020)

Muñoz-Gómez S, Hess S, Burger G, Lang BF, Susko E, Slamovits CH, et al. An updated phylogeny of the *Alphaproteobacteria* reveals that the parasitic *Rickettsiales* and *Holosporales* have independent origins. eLife 2019;8:e42535.

Minh BQ, Nguyen MA, von Haeseler A. Ultrafast approximation for phylogenetic bootstrap. Mol Biol Evol. 30:1188-95 (2013)

Nguyen LT, Schmidt HA, von Haeseler A, Minh BQ. IQ-TREE: a fast and effective stochastic algorithm for estimating maximum-likelihood phylogenies. Mol Biol Evol. 32:268-74. doi: 10.1093/molbev/msu300. (2015)

Saier MH, Reddy VS, Moreno-Hagelsieb G, Hendargo KJ, Zhang Y, Iddamsetty V, Lam KJK, Tian N, Russum S, Wang J, Medrano-Soto A. The Transporter Classification Database (TCDB): 2021 update. Nucleic Acids Res. 49:D461-D467 (2021)

Seemann T. Prokka: Rapid prokaryotic genome annotation. Bioinformatics 30, 2068-2069 (2014)

Szöllõsi GJ, Rosikiewicz W, Boussau B, Tannier E, Daubin V. Efficient exploration of the space of reconciled gene trees. Syst Biol. 62:901-12 (2013)

Szöllősi GJ, Davín AA, Tannier E, Daubin V, Boussau B. Genome-scale phylogenetic analysis finds extensive gene transfer among fungi. Philos Trans R Soc Lond B Biol Sci. 370:20140335 (2015)
