## Supplementary figures and images for "Host association and intracellularity evolved multiple times independently in the *Rickettsiales*"

### Fig_1.pdf

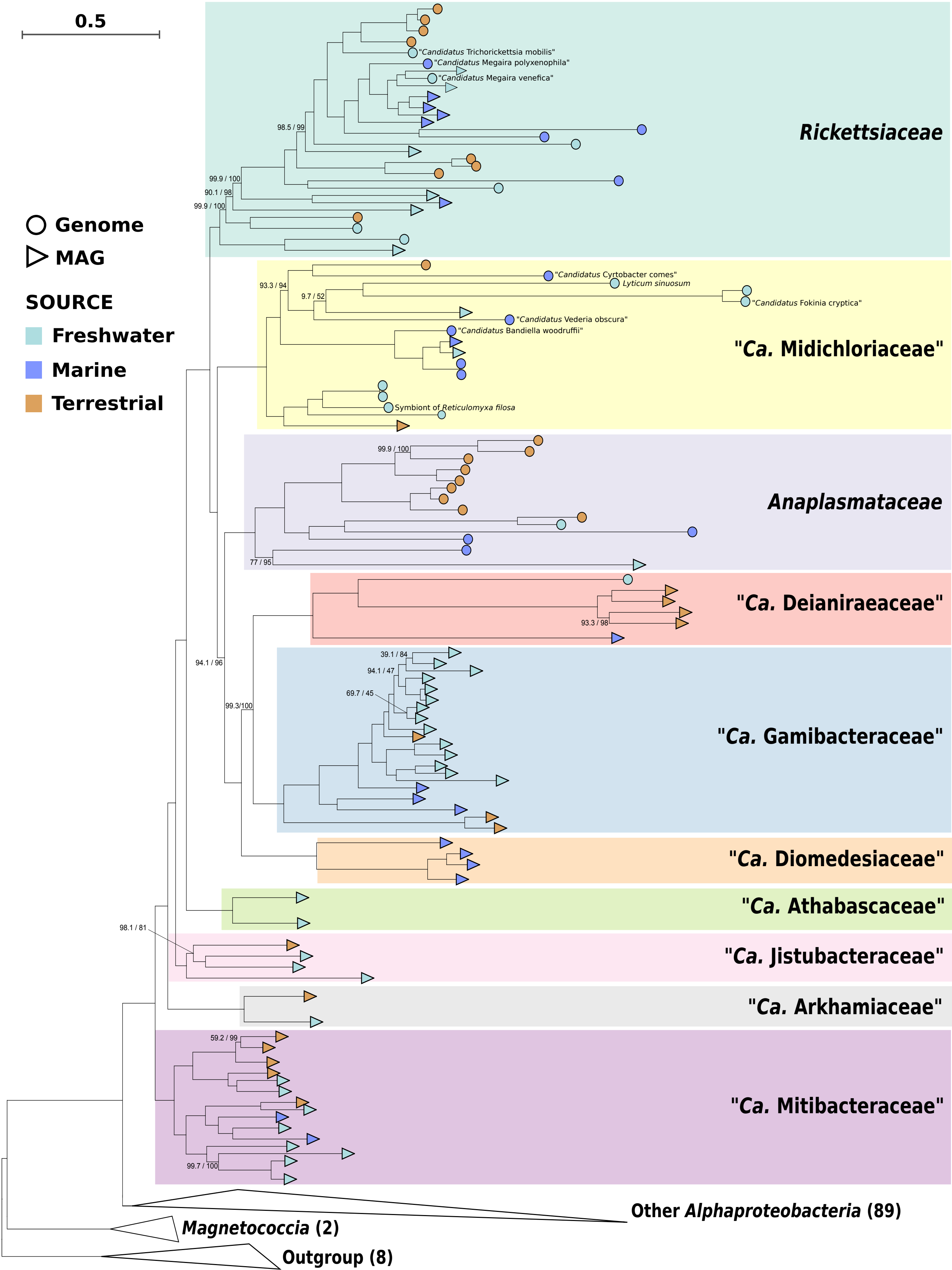

### Fig_3.pdf

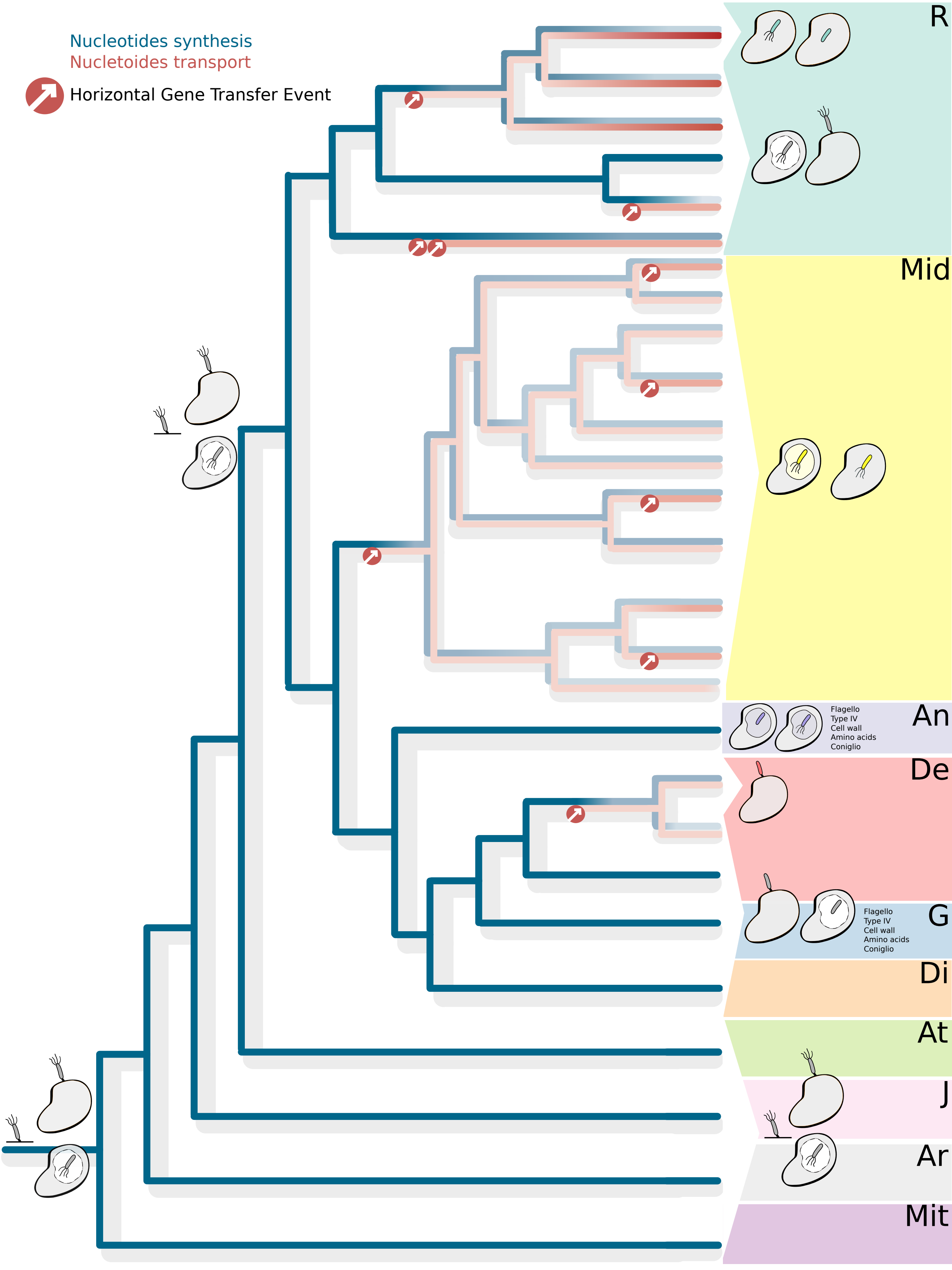

### Supplementary_figure_3_heatmap_secretion.pdf

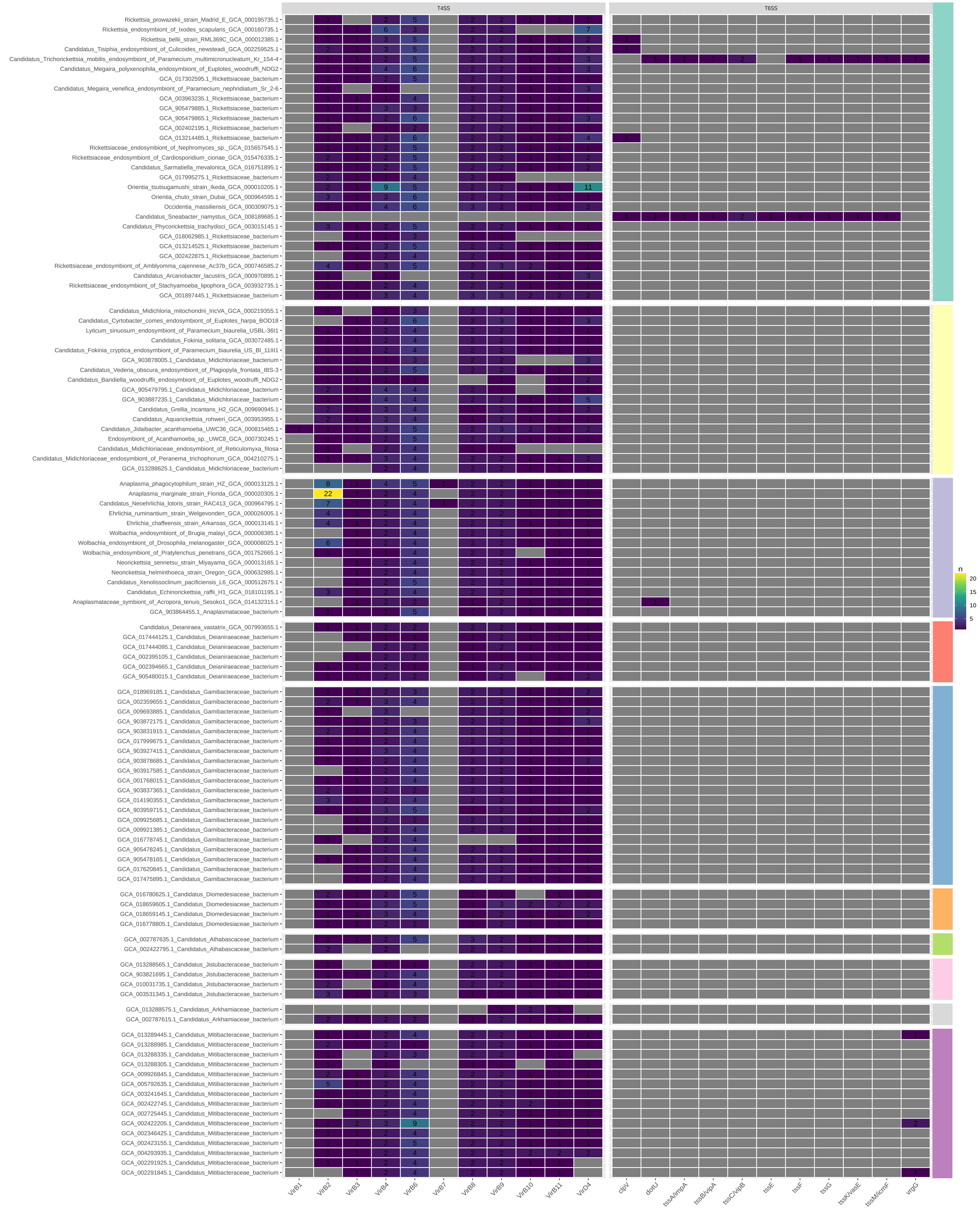

### Supplementary_figure_4_heatmap_effectors.pdf

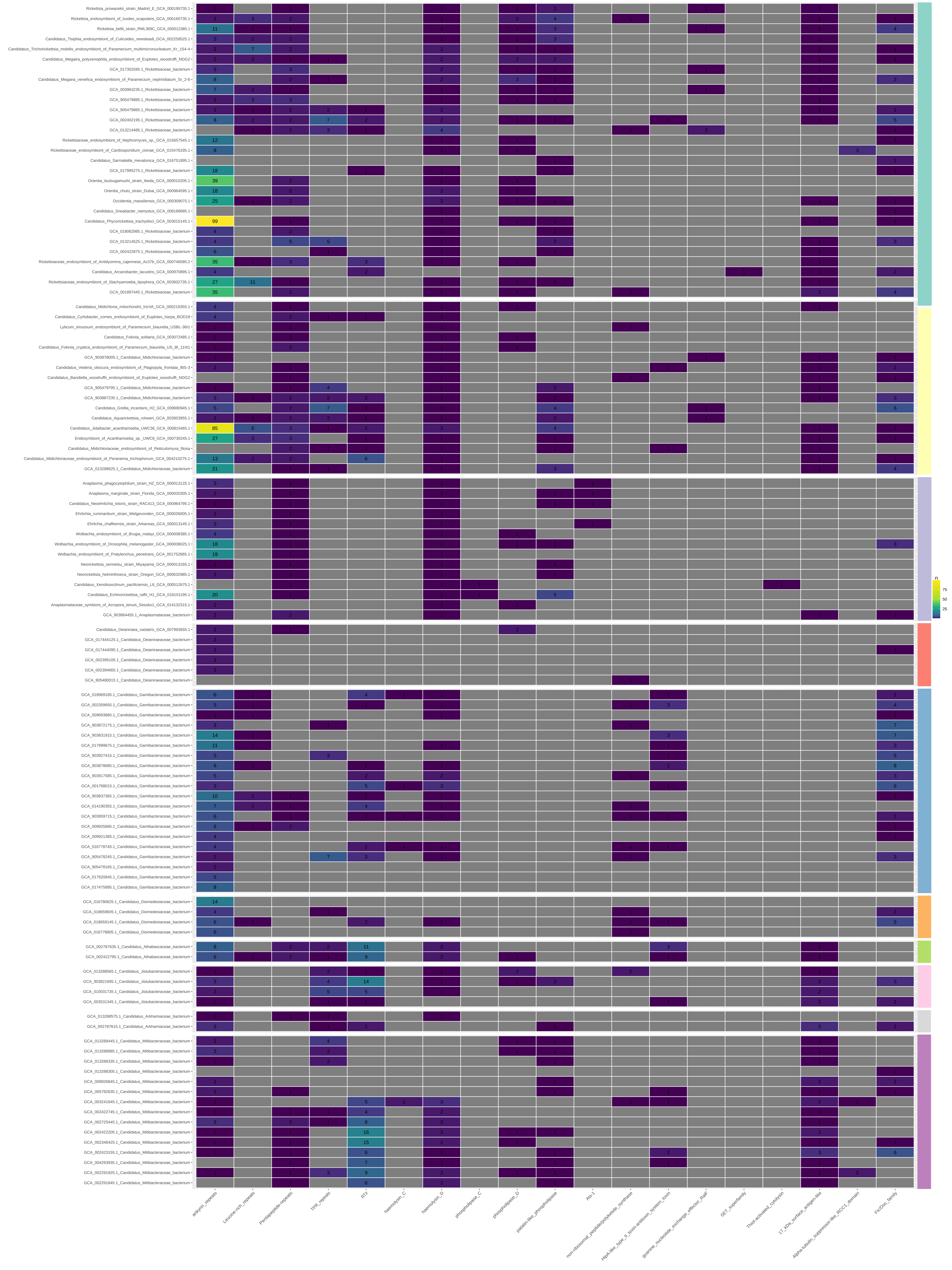

### Supplementary_figure_5_heatmap_flagellum.pdf

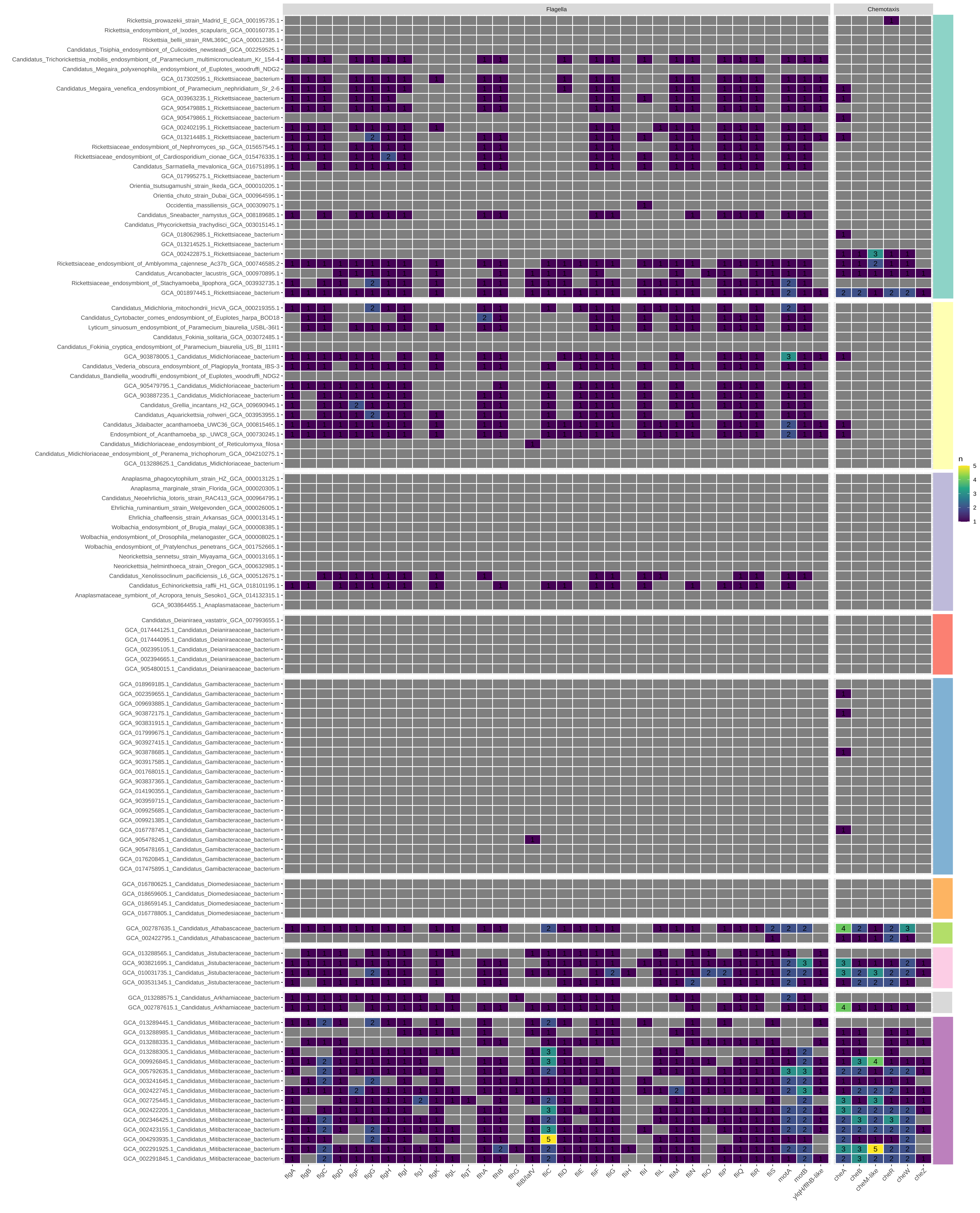

### Supplementary_figure_6_heatmap_adherence_miscellaneous.pdf

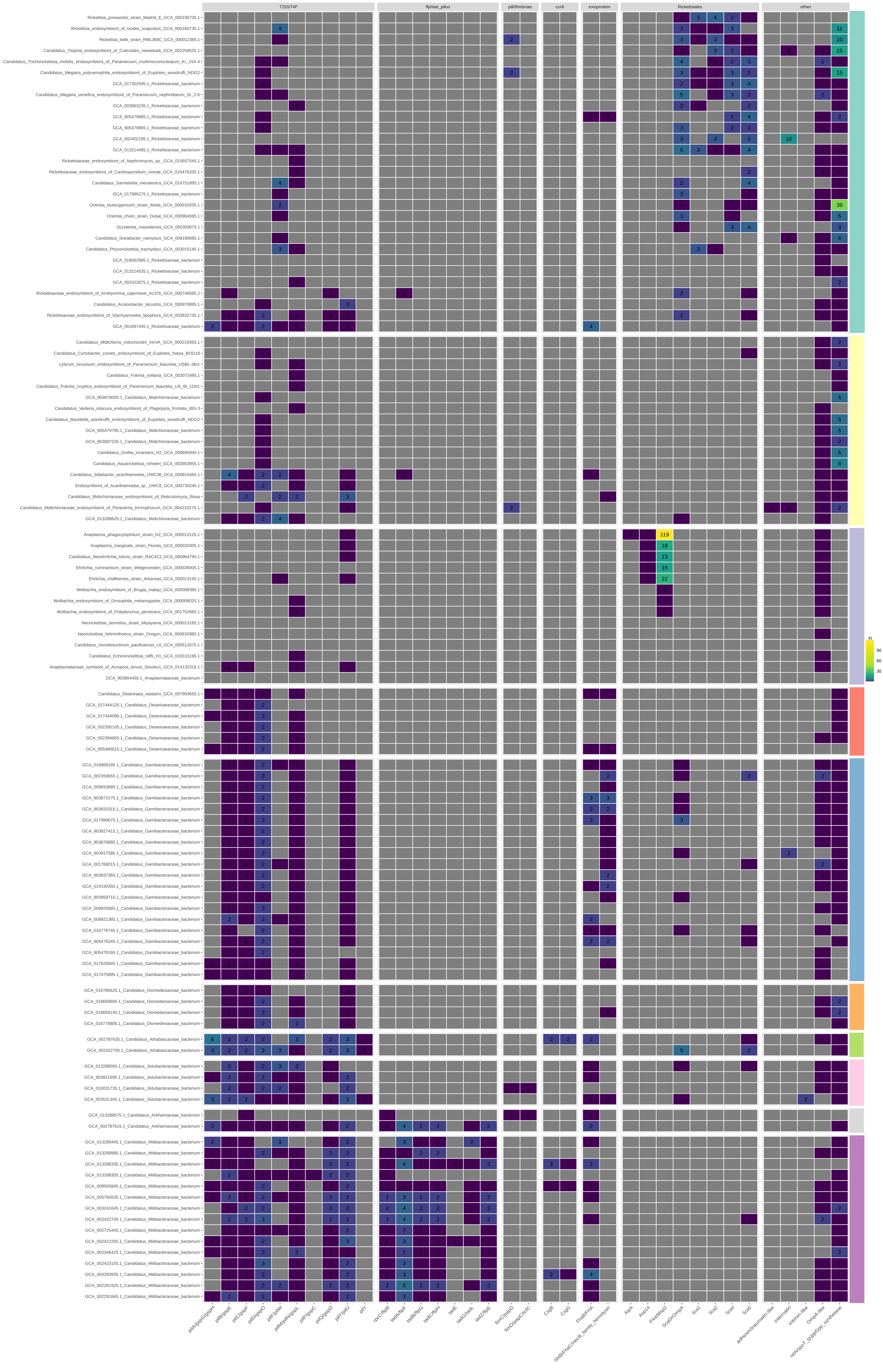

### Supplementary_figure_7_nucleotide_synthesis_transport.pdf

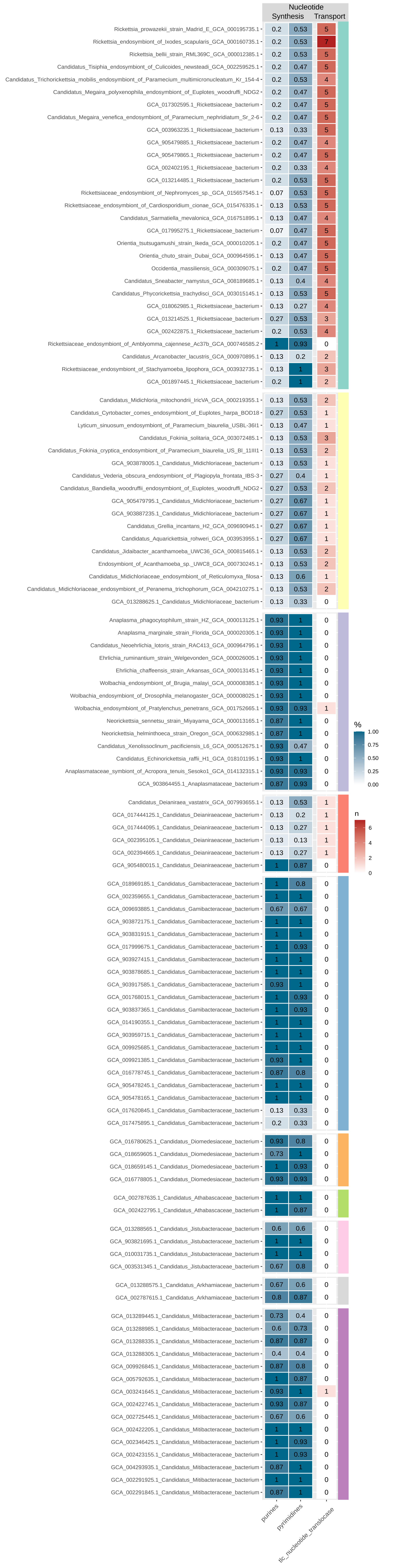

### Supplementary_figure_9_phylogeny_tlc_translocases.pdf

#### ADP/ATP translocase

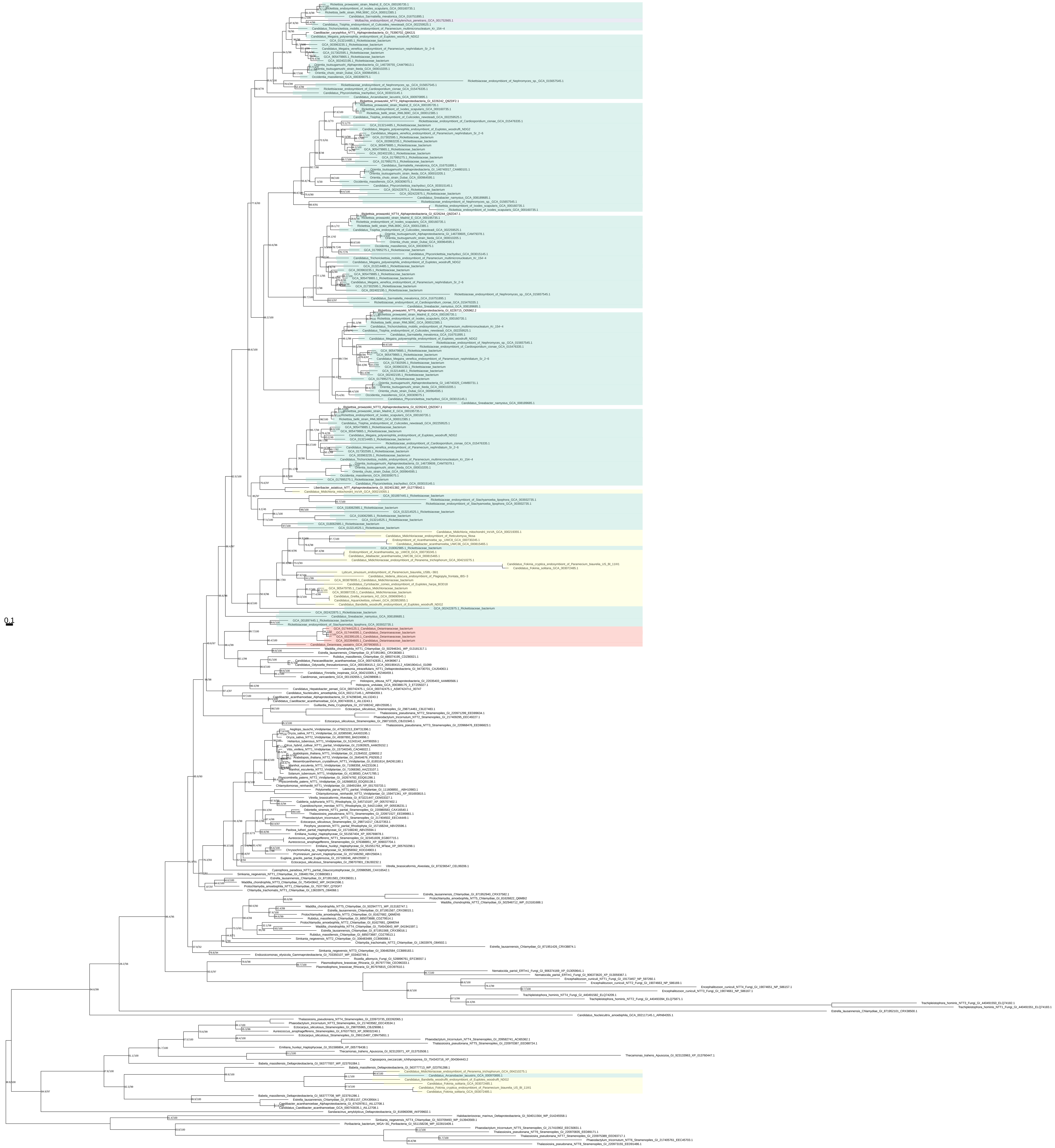

### Supplementary_figure_10_amino_acid_synthesis_transport.pdf

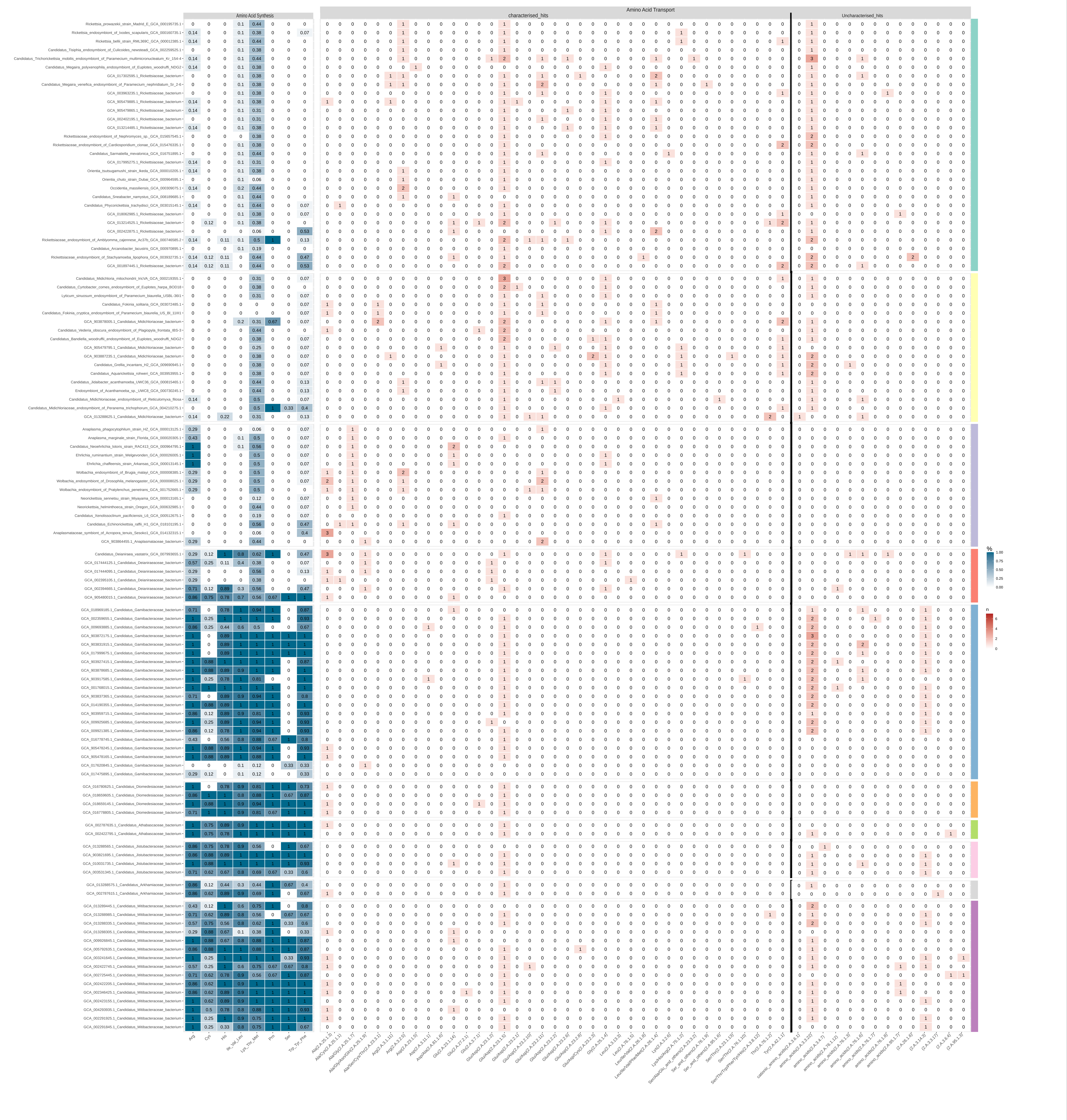

### Supplementary_figure_14_Phylogenomics_step6_updated_dataset_trees.pdf

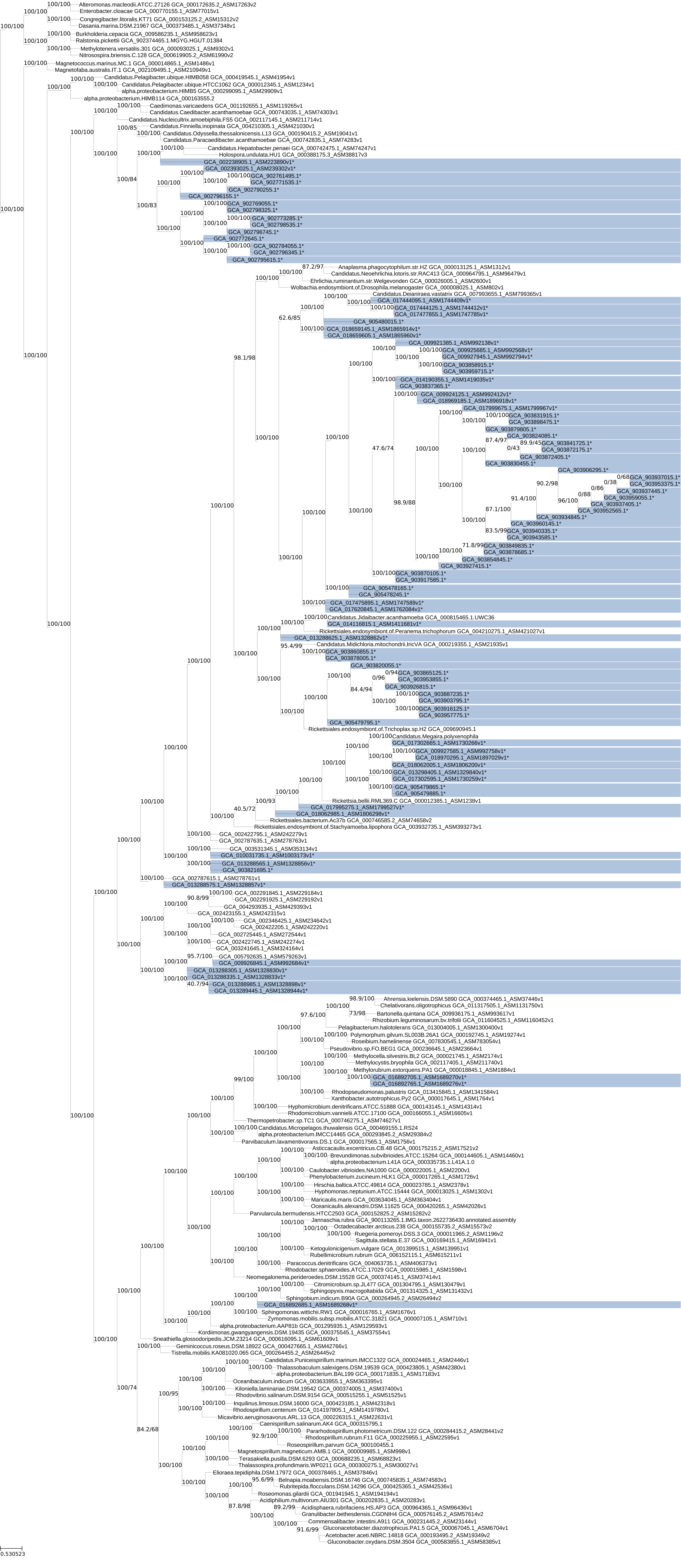

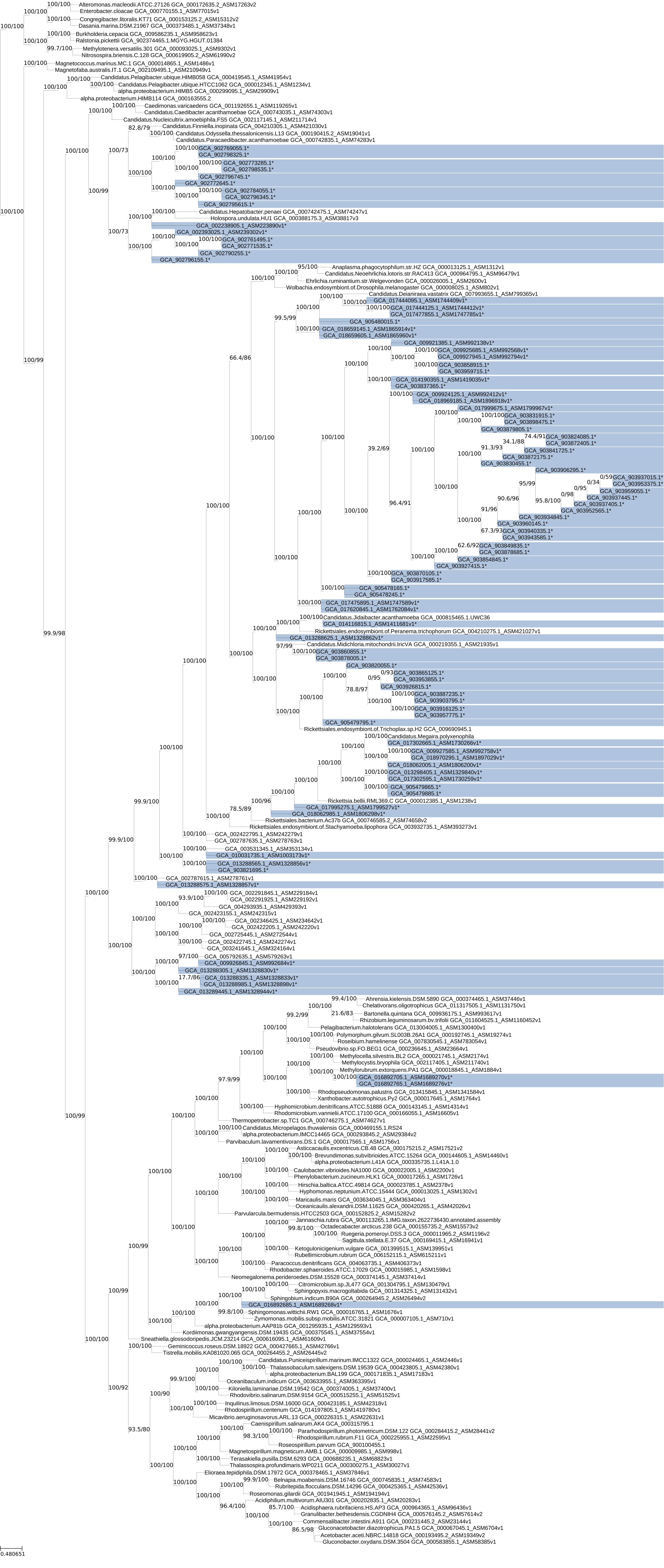

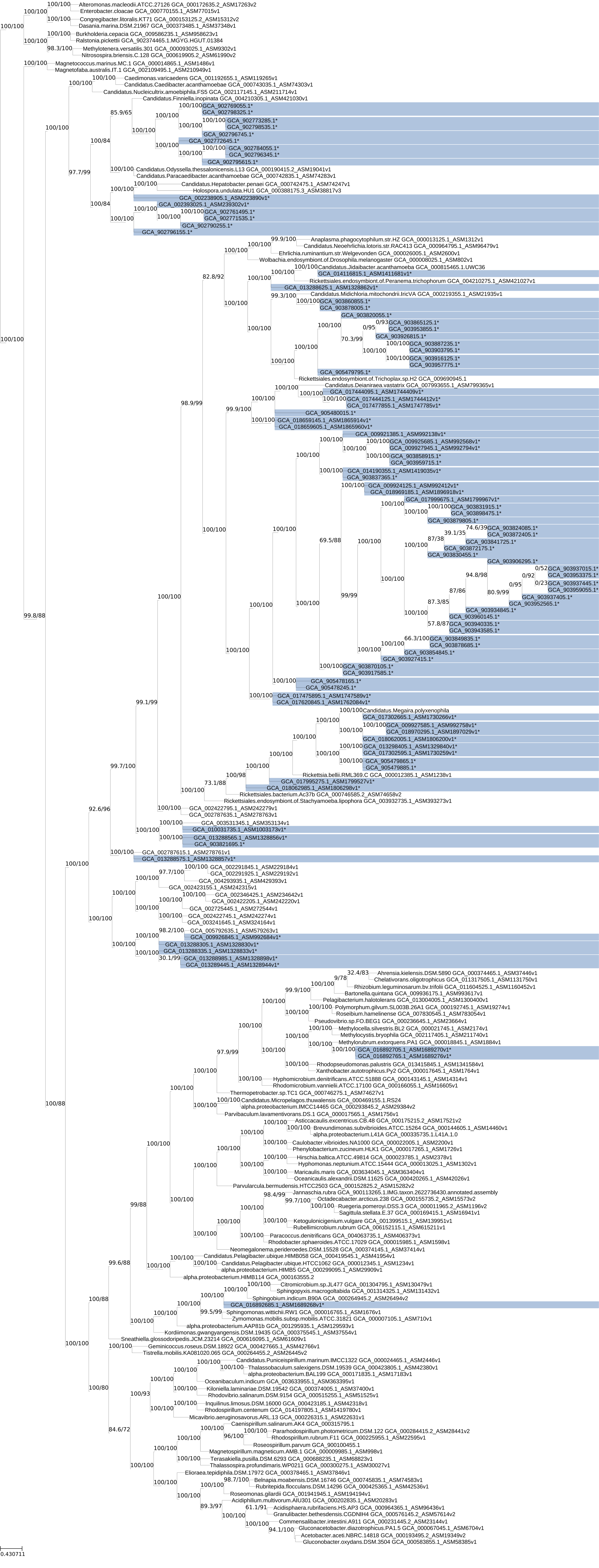

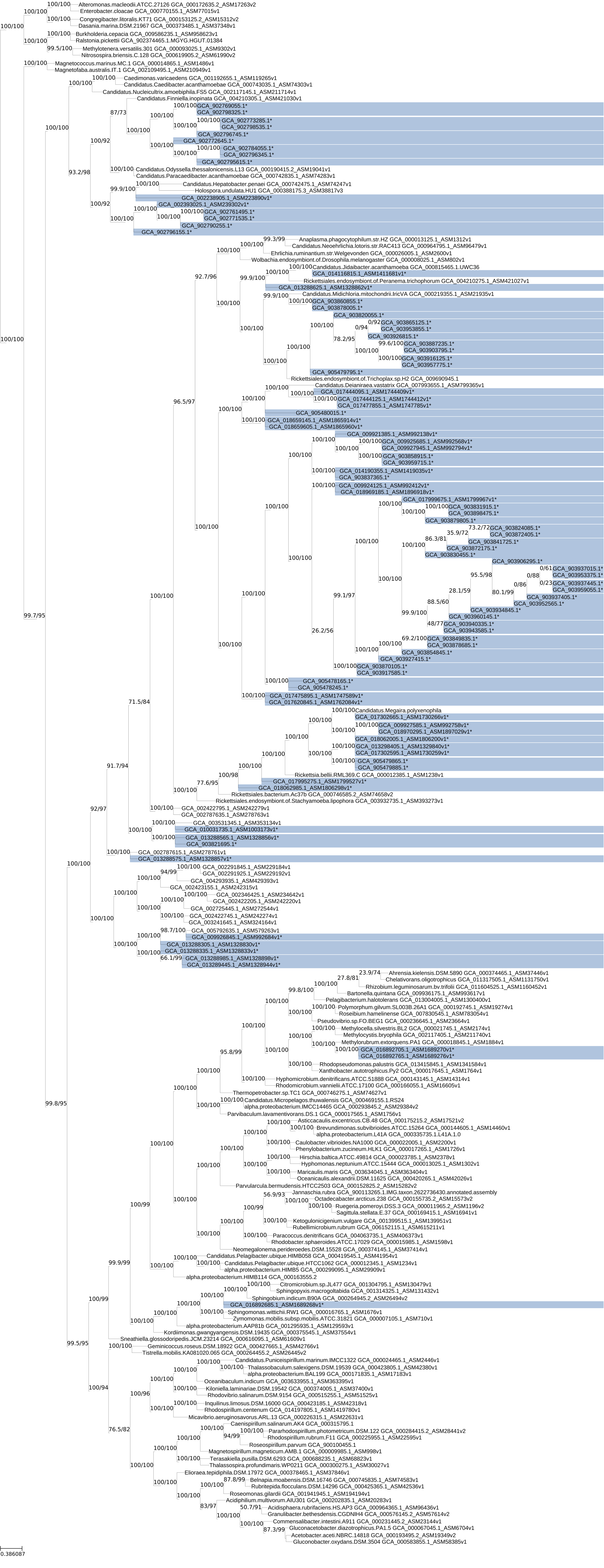

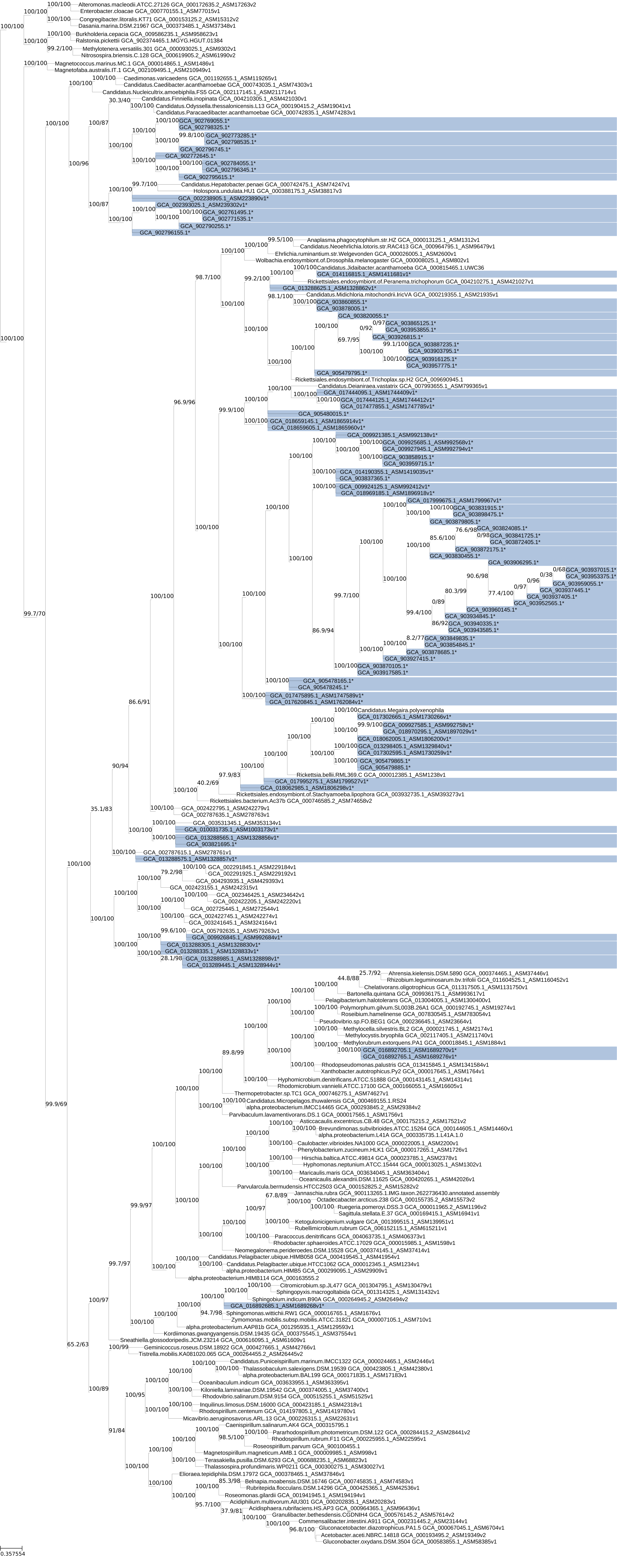

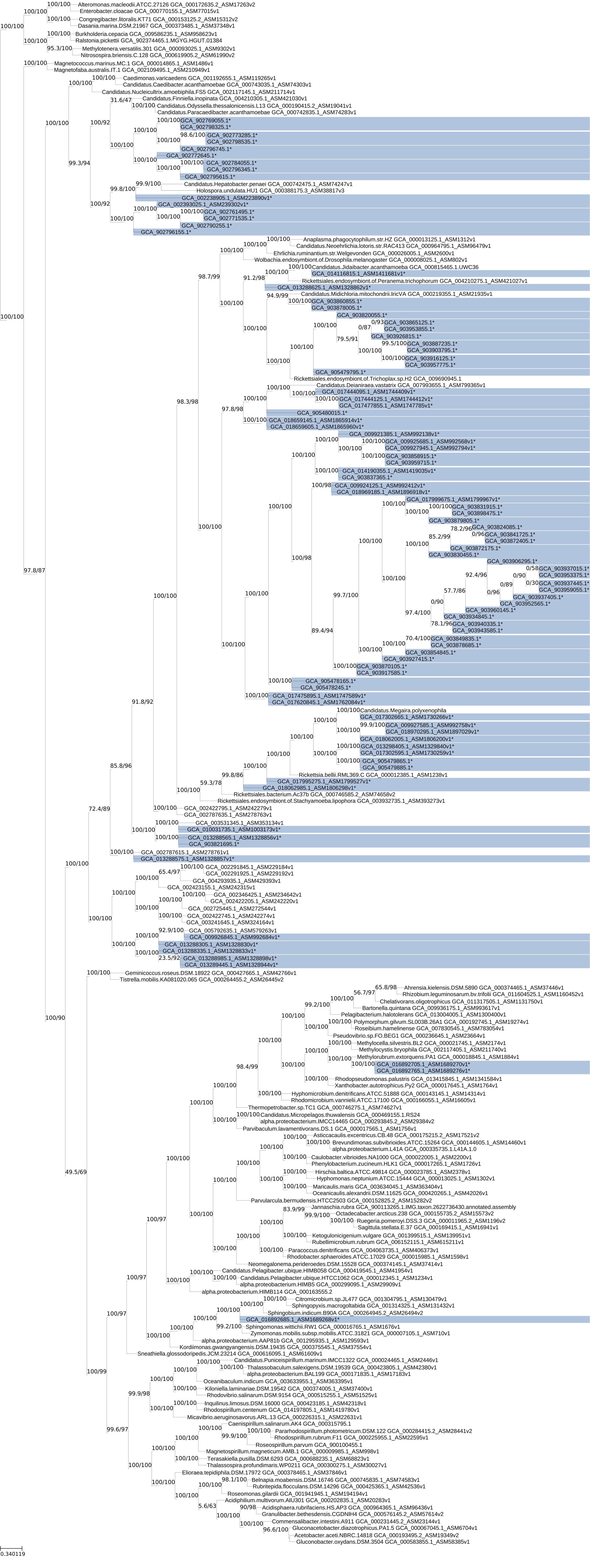
